# Searching for Sulfotyrosines (sY) in a HA(pY)STACK

**DOI:** 10.1101/2024.10.14.618131

**Authors:** Jordan Tzvetkov, Claire E Eyers, Patrick A Eyers, Kerry A Ramsbottom, Sally O Oswald, John A Harris, Zhi Sun, Eric W Deutsch, Andrew R Jones

## Abstract

Protein sulfation can be crucial in regulating protein-protein interactions but remains largely underexplored. Sulfation is near-isobaric to phosphorylation, making it particularly challenging to investigate using mass spectrometry. The degree to which tyrosine sulfation (sY) is misidentified as phosphorylation (pY) is thus an unresolved concern. This study explores the extent of sY misidentification within the human phosphoproteome by distinguishing between sulfation and phosphorylation based on their mass difference.

Using Gaussian mixture models (GMMs), we screened ∼45M peptide-spectrum matches (PSMs) from the PeptideAtlas Human Phosphoproteome build for peptidoforms with mass error shifts indicative of sulfation. This analysis pinpointed 104 candidate sulfated peptidoforms, backed-up by Gene Ontology (GO) terms and custom terms linked to sulfation. False positive filtering by manual annotation resulted in 31 convincing peptidoforms spanning 7 known and 7 novel sY sites. Y47 in Calumenin was particularly intriguing since mass error shifts, acidic motif conservation, and MS^2^ neutral loss patterns characteristic of sulfation provided strong evidence that this site is sulfated rather than phosphorylated.

Overall, although misidentification of sulfation in phosphoproteomics datasets derived from cell and tissue intracellular extracts can occur, it appears relatively rare and should not be considered a substantive confounding factor in high-quality phosphoproteomics datasets.

## Introduction

Protein sulfonation (usually referred to as sulfation) is an understudied post-translational modification (PTM), preferentially occurring on extracellular/secreted proteins, with roles in facilitating extracellular protein-protein interactions and modulating host-pathogen interactions^1–4^. Unlike phosphorylation, covalent addition of sulfate is believed to be irreversible and to occur almost exclusively on Y residues. The cellular mechanism for Y sulfation was originally described in 1983^5^, with enzymatically-deposited sulfation on Y residues occurring on an acidic substrate consensus^6,7^ in the trans-Golgi compartment^8^.In humans, Y sulfation is catalysed by either of two tyrosyl protein-sulfotransferases, termed TPST1 and TPST2. Knockout mice lacking both sulfotransferases possess little detectable sulfotyrosine^9^. Early work employed ^35^S-based labelling and sY antibodies to evaluate this modification, but these techniques are unreliable and do not generate context-dependent information, so have become obsolete in the age of mass spectrometry-based proteomics. As of mid-2024, there are 51 human proteins annotated in UniProtKB^10^ as being sulfated on tyrosine. Some two-thirds of these have been experimentally validated. Our groups have recently developed a mass spectrometry (MS)-based workflow incorporating low collision energy-induced neutral loss for preferential sulfopeptide fragmentation, increasing the number of experimentally identified sulfated proteins from 33 to 54^11^. A number of these were confirmed experimentally using *in vitro* TPST1/2 sulfation assays with purified components^12,13^. One such protein was Golgi-localized Heparan-sulfate 6-O-sulfotransferase 1/2 (H6ST1/2) suggesting potential interplay between protein and carbohydrate sulfation^11^. Intriguingly, previously published data suggests that up to 7% of all Y residues could also potentially be sulfated, which raises the question as to the scale and cellular functions of this vastly underexplored PTM^14,15^.

Even though MS is a powerful analytical tool for the characterisation of covalent protein modifications, there are several challenges in the accurate identification of sites of protein sulfation using MS^16^. First, the sulfoester bond is highly labile, meaning that the modification typically undergoes complete neutral loss during most types of fragmentation commonly employed in tandem MS, and even during electrospray ionisation^11,17^. Second, even optimised methods of sulfopeptide enrichment e.g. using TiO_2_ or immobilised metal-affinity chromatography (similar to the approaches used for phosphopeptide enrichment), or unreliable sulfotyrosine antibodies, are not as efficient or as specific as the strategies typically employed in phosphopeptide-based analytical pipelines^11^. Third, the masses of sulfotyrosine and phosphotyrosine are near isobaric - pY: 79.966331 Da and sY: 79.956815 Da (i.e. 0.0095 Da mass difference). The precursor mass tolerance for modern Orbitrap-based instruments can be accurate to around 1-2 ppm if perfectly calibrated, although in practice it is typical in a sequence database search approach to apply a +/- 5 or 10 ppm mass error window. In a search with ppm tolerance of +/-10 ppm, and for a peptide of mass 2000 Da, the actual window would thus be +/-0.02 Da, and for a 4000 Da peptide, +/-0.04 Da. As such, during MS/MS analysis of a phosphopeptide-enriched sample, a tyrosine-sulfated peptide would be captured within the same precursor window as a tyrosine-phosphorylated peptide. Furthermore, it is not typical to search for sulfation and phosphorylation at the same time during database searching. Combined with the fact that there is almost complete neutral loss of the sulfate group from the amino acid side chain during analysis, and the isobaric nature of the covalent moiety, it is nearly impossible to prove unequivocally that a peptide was indeed sulfated, rather than phosphorylated, in a typical tandem MS acquisition pipeline. We thus hypothesised that large-scale phosphoproteome datasets may contain potential sulfated peptides that have been misidentified as phosphorylated.

There have been several attempts to perform very large-scale meta-analyses of phosphopeptide data for different species^18,19^, as well as collate such data into databases, like PhosphoSitePlus^20^. One of these – the human phosphoproteome build in PeptideAtlas^21^ contains > 400 million spectra in 129 datasets generated from human samples enriched for phosphopeptides, identifying 264K distinct phosphopeptides (https://peptideatlas.org/builds/human/phospho/). In this work, we were intrigued by the possibility of exploring whether we would still robustly identify phosphopeptides using a precursor mass error distribution that better fits the hypothesis of one or more sulfated sites on the peptide, specifically within such a large data collection comprising many different enrichment protocols and data acquisition settings. We describe a method based on experimental mass calibration and model fitting to identify outliers in the precursor mass distribution, which we demonstrate is highly robust for detecting *bona fide* sulfated peptides where they exist in a ‘phosphoproteome’ dataset.

## Methods

An overview of the analytical pipeline is presented in **Fig. 1**.

**Fig. 1.**
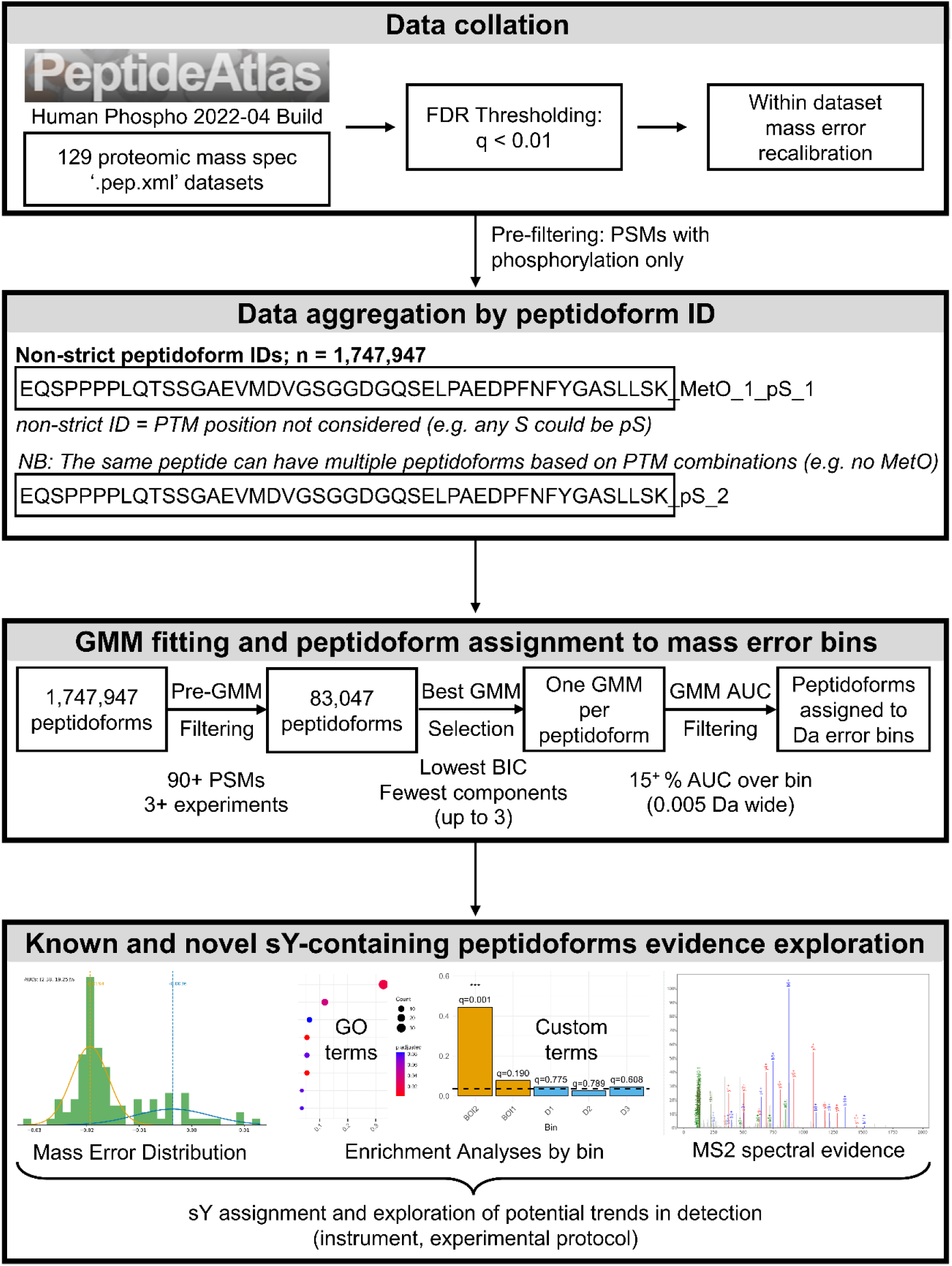
Study design and workflow. **Abbreviations: AUC** - area under the curve; **BIC** - Bayesian information criterion for model selection; **GMM** - Gaussian mixture model; **FDR** - false discovery rate; **PSM** – peptide-spectrum match; **PTM** – post-translational modification

### Data collation and aggregation by peptidoform

Our study employed the largest publicly available human phosphoprotein tandem mass spectrometry build from PeptideAtlas - Human Phosphoproteome 2022-04. The build includes data from 129 MS/MS (MS^2^) datasets spanning 365 experiments, giving a total of 19,852 MS runs. This contributed to the identification of 264,244 peptides derived from 9,810 “canonical” proteins (leading proteins within protein groups). This dataset was selected due to its positive enrichment for phosphopeptides, which we hypothesised may include sulfopeptides misidentified as phosphopeptides. The peptide spectrum matches (PSMs) and scores were extracted and converted from PepXML^22^ to ‘.tsv’ format using custom Python scripts. False discovery rate thresholding was applied based on PeptideProphet^23^ with q < 0.01 followed by recalibration of the thresholded data. Recalibration was performed on a per raw file basis using custom scripts written in Python, by calculating the median mass error (*mme*) in Daltons and subtracting *mme* from the originally assigned mass error for each PSM. Subsequently, filtering was performed to include only peptides with assigned phosphoserine (pS), phosphothreonine (pT), and phosphotyrosine (pY) PTMs. We did not apply any thresholding based on site localisation statistics, since we know that sulfotyrosine cannot be confidently determined, due to complete neutral loss, and thus discoverable sY sites could appear in sites with any PTM localisation score.

Based on the peptide sequence, and type and number of PTMs assigned to it, a peptidoform ID was generated for every peptide-spectrum match (PSM). A peptidoform is a variation of a peptide sequence with a specific set of PTMs. The same peptide sequence can have multiple peptidoforms as illustrated in Fig. 1. In the provided example, the first peptidoform of the same peptide is oxidized on one methionine (MetO_1) and phosphorylated on one serine (pS_1), while the second peptidoform has no oxidation and has two serines phosphorylated (pS_2). Notably, these were ‘non-strict’ peptidoform IDs since precise PTM positional information was not taken into account. However, these IDs do consider the amino acid that has been modified. So, for example, if the same peptide is sometimes assigned as phosphorylated on a serine and other times as phosphorylated on a tyrosine, these would be two different peptidoforms, with pS and pY reflected in the ID. This resulted in 1,747,947 unique phosphorylated peptidoform IDs across the build. For each of these peptidoforms, their associated PSM data was pooled together, and the recalibrated mass errors were taken forward for subsequent analyses.

### Mass error calculation

For each candidate PTM site, we typically have large collections of PSMs emerging from within a single run, across different runs, and from across different data sets. We thus calculated a distribution of mass error values across all PSMs, identifying the same PTM site (and produced histograms for visualisation) for the same peptidoform. Based on accurate mass, if the distribution of mass errors has a peak centred around -0.0096 Da, the modification identified as phosphorylation may be more likely to be a sulfation event. The accurate masses used were as reliable as possible based on recalibration of the original datasets as described above. Despite recalibration, mass accuracy can depend on the instrument settings used (a small fraction of calibrated mass errors may still be incorrect), which warranted the use of robustly detected peptidoforms in this study. Using a method fitting one or more Gaussian mixture models (GMMs) to the mass error distributions, we aimed to profile the extent to which there is potential for misidentification of true sulfation or true phosphorylation in the human PeptideAtlas phosphoproteome build. We then applied further contextual information to determine the likelihood that some of these putative sulfation sites were indeed correct, as follows.

### Gaussian Mixture Model (GMM) fitting

A custom script in Python was built to fit Gaussian mixture models imported from sklearn (the code is publicly available on GitHub: https://github.com/CBFLivUni/Sulfo_Tyrosine). Prior to GMM fitting, data were filtered to include only robustly detected peptidoforms across the build by selecting peptidoforms detected in three or more experiments (n = 413,010 peptidoforms). To ensure sufficient data for fitting high-quality Gaussian models, and minimise the random chance of mass errors fitting the profile for a sulfation candidate, we applied a further filter to include only peptidoforms with 90 or more PSMs, as previously suggested^24^, resulting in 83,047 peptidoforms taken forward for analysis. The threshold was selected to apply an optimal trade-off between specificity and sensitivity. A peptidoform with few PSMs cannot be modelled accurately by the GMM, and it is common for a PSM to have a chance match with a shifted mass error. For a confident detection of sulfation, we wished to observe a large number of (ideally) independent observations of the mass shifted peptidoform.

GMMs with 1, 2, or 3 components were fit on the recalibrated mass error PSM data for each peptidoform. The reason for including up to three components was to explore the following scenarios in a search of sulfation misidentified as phosphorylation:

#### One component

Peptidoforms that are correctly identified as phosphorylated should have a normal distribution centred around a 0 Da shift. The mass error for peptidoforms that are always singly sulfated and misidentified as phosphorylated would be expected to follow a normal distribution with median falling around a -0.0095 Da shift. Similarly, doubly sulfated peptidoforms would exhibit a median shift of approximately -0.019 Da.

#### Two components

Two Gaussians would be an optimal fit for scenarios where a single tyrosine residue of a peptide is sometimes phosphorylated and sometimes sulfated (both 0 Da and -0.0095 Da medians present in singly sulfated peptides or both -0.0095 Da and -0.019 Da medians in doubly sulfated peptides).

#### Three components

Models with three components would cover rare scenarios where two different tyrosine residues within the same peptide could either be sulfated or phosphorylated or any of the previous scenarios mixed with random misidentifications of other PTMs causing a different shift in mass.

The best fitting number of components was individually selected for each peptidoform based on Bayesian Information Criterion (BIC) scores with lower scores indicating a better fit. To avoid overfitting the data, the smallest number of components were prioritised unless the BIC score of a model with a larger number of components was 10 or more points smaller, which is indicative of ‘very strong evidence’ of a better model^25^.

### Peptidoform assignment to mass error bins

Based on the scenarios described above, mass error bins were pre-defined spanning -0.0225 Da to 0.0225 Da. The width of each bin was set to 0.005 Da, and bins were split into three groups as described below:

<u>Bins of interest:</u> three Bins of Interest (BOIs) covering potential single (−0.0096 Da) and double (−0.0192 Da) sulfation events, as follows: BOI1 (−0.0125 : -0.0075); BOI2 (−0.0175 : -0.0125); BOI3 (−0.0225 : -0.0175)

<u>Phosphorylation bins:</u> three bins over the mass error shifts expected for true phosphorylation events (TRUEp), ranging from -0.0075 Da to +0.0075 Da

<u>Decoy bins:</u> three bins for mass shifts opposite to those expected for sulfation (>0.0075 Da, mirroring the BOIs).

For each peptidoform ID, the area under the curve (AUC) spanning each bin’s mass error shift range was computed based on the probability density function of the best-fit GMM for that peptidoform. Peptidoforms were assigned to a bin if the resulting AUC for that bin was >= 15% of the total AUC for the model. This threshold was used to limit false negatives (e.g. sulfated peptidoforms with narrow distribution curves slightly outside the BOIs), while allowing some false positives (e.g. phosphorylated peptidoforms with wide distribution curves that would be filtered out by higher AUC % thresholds). One ‘side effect’ of this threshold is that we would expect BOI2 to give the most comprehensive overview of sulfated peptidoforms by capturing both singly and doubly sulfated peptidoforms with wide enough distributions while being less prone to false positives than BOI1 as it is further away from the 0 Da calibrated mass error expected for phosphorylated peptidoforms. Additionally, using AUC thresholding results in some peptidoforms being assigned to multiple bins. For example, this would be expected when phosphorylation (and assignment to TRUEp bins) and sulfation (and assignment to BOIs) are both possible, or when a sulfated peptidoform with wide enough distribution is assigned to multiple BOIs. The use of BOIs and calibrated mass errors served as a tool for shortlisting potentially sulfated peptidoforms. Combined with the subsequent biological contextualisation and manual curation described below, we were able to identify strong candidate sulfation sites.

### Data exploration for known and potential novel sulfotyrosines sites

All analyses described below were carried out using custom scripts in R version 4.2.0. The associated code is publicly available on GitHub.

#### GO term enrichment analyses

Peptidoforms were filtered based on their protein ID assignment, with proteins absent from the UniProtKB human reference database (20429 entries, downloaded 17/01/2024) excluded from analysis. A total of 2,758 UniProtKB proteins were associated with the 83,047 peptidoforms for which GMMs were fitted. The associated UniProtKB entry IDs for these proteins were converted to Ensembl gene IDs using package biomaRt^26,27^. These were selected as the universe of background genes for overrepresentation analyses. Ensembl gene IDs associated with the peptidoforms within a bin were used as list of foreground genes to test for GO term enrichment using the enrichGO() function from the clusterProfiler package version 4.6.2^28^. A separate enrichment analysis was carried out within each BOI and DECOY bin, testing for molecular function, biological process and cellular compartment GO term enrichment. For each analysis, p-values were adjusted using the Benjamini-Hochberg method, org.Hs.eg.db version 3.16.0 was used as the annotation database, and the q-value cutoff was set to 0.1^29^.

#### Custom terms enrichment analyses

To incorporate biological context that is believed to be strongly-associated with tyrosine sulfation into the analysis, custom term to protein mappings were generated based on the UniProtKB database as of 20^th^ February 2024, as follows:

<u>‘known_sY’:</u> 69 protein entries with known sulfation PTMs on tyrosine residues based on a combination of our previous experimental validation and the following UniProtKB search: “(proteome:UP000005640) AND (ft_mod_res:sulfotyrosine)”. Of these, 12 were present in the background universe (entire dataset for which GMMs were fit).

<u>‘Secreted’</u>: 2,112 proteins for which the word ‘secreted’ appears in the column “Subcellular location” of the UniProtKB dataset. Of these 102 were present in the background universe.

<u>‘Transmembrane’:</u> – 5,221 proteins for which the column “Transmembrane” of the UniProtKB dataset was not empty. Of these 356 were present in the background universe.

<u>‘Golgi’</u>: 1,135 proteins for which the protein name or the column “Subcellular location” of the UniProtKB dataset contained the word “Golgi”. Of these 102 were present in the background universe.

<u>‘unlikely_sY’</u>: 12,876 UniProtKB entries that did not match any of the above criteria. Of these 2,197 were present in the background universe.

Custom term overrepresentation analyses were carried out for each bin in an analogous fashion to the GO terms analyses, using the enricher() function from clusterProfiler. To accommodate for the overall small number of known sY proteins in the background (n=6), the minimum gene set size limit was set to 1. Given the small number of terms, no q-value cutoffs were imposed. Benjamini-Hochberg-adjusted p-values were plotted and reported.

#### Manual annotation of peptidoforms assigned to BOIs

The mass error histograms of Y-containing peptidoforms assigned to any of the BOIs by the GMM AUC pipeline were manually inspected and scored based on how convincing they were using the following criteria: i) ‘Convincing’ if at least one of the Gaussians was centred near the Da error shift expected for a singly (−0.0095 Da) or doubly (−0.019 Da) sulfated peptidoform; ii) ‘Undetermined’ if the Gaussian was widely distributed and had a mean mass error shift between -0.06 and -0.04 Da; iii) ‘Not Convincing’ if the Gaussian was widely distributed and had a mean mass error shift > -0.04 Da. Additionally, a histogram type was assigned to each peptidoform and abbreviated based on the presence of multiple PSMs with a mass shift error supporting one sY site that has been misidentified as phosphorylated (s), two sY sites that have been misidentified as phosphorylated (ss), or no sY sites have been misidentified (p; denotes a correct phosphorylation assignment). Combinations of the above were separated by ’/’ in instances where multiple options were supported in the histogram. For example, ‘ss/s/p’ indicates some PSMs for that peptidoform support two sY sites, some support one, and some suggest no sulfation. All ‘Undetermined’ histograms fell between an ‘s’ and a ‘p’ type and were thus labelled as ‘s/p’. Finally, biological context was summarised using the custom term to protein mappings described in the previous section but known_sY labels were replaced with ‘Sulfated’ and ‘unlikely_sY’ with ‘No prior knowledge’. Peptidoforms supported by biological context with convincing histograms were considered likely to contain a sY site and were taken forward for further MS^2^ validation.

#### Assessment of MS^2^ evidence of sulfation

Using a custom R script, Universal Spectrum Identifiers^30^ (USIs) were generated for each PSM across the build associated with peptidoforms manually annotated as likely to contain a sY. USIs for each PSM are in the following format:

mzspec:<COLLECTIONID>:<MSRUNCOMPONENT>:<INDEXFLAG>:<SCANNUMBER>:<<bold>Spectrum_Interpretation</bold>>/<CHARGE>

Briefly, the script generates ‘original’ and ‘alternative’ USIs. The **Spectrum_Interpretation** portion of an ‘original’ USI is based on the peptide sequence and PTMs assigned in the original search, even if a misidentification of a sulfation event as phosphorylation is suspected. In an ‘alternative’ USI, the suspected misidentification is rectified - where possible a tyrosine phosphorylation is replaced by a tyrosine sulfation (Y[Phospho] -> Y{Sulfo}). The effect of the curly braces in the {Sulfo} notation as prescribed in the ProForma 2.0^31^ peptidoform notation standard means that the spectral interpretation engine expects a full neutral loss of the sulfate group in the fragment ion (MS^2^) spectrum.

If phosphorylation was assigned only to non-tyrosine residues, a random phosphorylation assignment was removed, and a sulfation was placed on a random tyrosine. Two alternative USIs were generated in cases where double sulfation is possible by repeating the phosphorylation replacement process. The ProteomeXchange^32,33^ USI spectrum viewer (https://proteomecentral.proteomexchange.org/usi/) was used to visualise the MS^2^ spectral evidence supporting the two USI alternatives of the same spectrum. The default ion visualisation settings were used, but H_3_PO_4_ and HPO_3_ neutral loss mapping was disabled. As sulfation typically undergoes full neutral loss (−79.95 Da), the HPO_3_ (−79.96 Da from phospho) could confuse the interpretation.

Peak assignment was based on the most intense monoisotopic peak within an m/z tolerance of 0.05 Th. For USIs where a larger portion of the sequence was supported and a larger percentage of the total ion current was accounted by the sulfated variety compared to the phosphorylated variety, sulfation was believed more likely than phosphorylation. Deamidated peptidoforms were not assessed.

#### Purification and TPST-catalyzed sulfation of recombinant human Calumenin

A Genscript cDNA encoding full-length human calumenin (Uniprot: Q6IAW5) was synthesized and cloned into PasI and NotI restriction sites of pGex-6P-1, which encodes an N-terminal GST-tag in frame with calumenin comprising amino acids 1-315. Expression of the GST fusion protein was induced in the *E. coli* strain SoluBL21(DE3) using standard procedures^34^. After *E. coli* extract centrifugation, soluble GST-calumenin was purified to near homogeneity using sequential glutathione-Sepharose affinity and size-exclusion chromatography. Calumenin is a multiple EF-hand containing acidic protein (predicted pI= 4.47) migrating as a tight doublet of around 70kDa (likely monomeric) and as a higher molecular weight multimeric complex of around 250 kDa after SDS-PAGE. To evaluate the kinetics of Tyrosine protein sulfation of the soluble calumenin species present, they were incubated at 30°C in the presence of purified TPST1, TPST2 or a combination of TPST1 and TPST2 with the sulfate donor PAPS (1 mM) and 5 mM MgCl2, using previously described assay conditions^12,35^. Covalently-bound sY was detected using a ZooMAb mouse monoclonal anti-sulfotyrosine (Sigma).

#### Analysis of calumenin by liquid chromatography-tandem mass spectrometry (LC-MS/MS)

Recombinant calumenin pre-incubated with TPST1/2 and PAPS for 16 hours at 20°C was diluted ∼4-fold in 100 mM ammonium bicarbonate (pH 8.0) and subject to reduction with dithiothreitol and alkylation with iodoacetamide as previously described (Ferries et al., 2017). Samples underwent an SP3-based trypsin digestion protocol adapted from Daly et al. 2023^11^, using 100 mM ammonium bicarbonate (pH 8.0) and 0.5 μg of Trypsin gold (Promega)^11^. Dried peptides were solubilized in 20 μl of 3% (v/v) acetonitrile and 0.1% (v/v) TFA in water, sonicated for 10 min, and centrifuged at 13,000x g for 10 min at 4 °C prior to reversed-phase HPLC separation using an Ultimate3000 nano system (Dionex) over a 30 min gradient, as described^36^. All data acquisition was performed using a Thermo QExactive mass spectrometer (Thermo Scientific), with higher-energy C-trap dissociation (HCD) fragmentation set at 30% normalized collision energy for 2+ to 4+ charge states. MS1 spectra were acquired in the Orbitrap (70K resolution at 200 m/z) over a range of 300 to 2000 m/z, AGC target = 1e6, maximum injection time = 250 ms, with an intensity threshold for fragmentation of 1e3. MS2 spectra were acquired in the Orbitrap (17,500 resolution at 200 m/z), maximum injection time = 50 ms, AGC target = 1e5 with a 20 s dynamic exclusion window applied with a 10 ppm tolerance. For sulfation PTM analysis, raw data files (PRIDE ID will be provided during proofs) were converted into mgf format using MSConvert, with peak picking filter set to “2-” and searched with the MASCOT^37^ search engine against the UniProt Human Reviewed database (updated weekly, accessed November 2024) (UniProt 2024) with variable modifications = carbamidomethylation (C), oxidation (M), sulfo (STY), instrument type = electrospray ionization–Fourier-transform ion cyclotron resonance (ESI-FTICR) with internal fragments from 200-2000 *m/z*, MS1 mass tolerance = 10 ppm, MS2 mass tolerance = 0.01 Da. For the best MASCOT scoring peptide spectrum match (PSM) for a sulfation-containing peptide, the mgf file was extracted from the raw file and imported into a custom R script for re-drawing and manual annotation.

#### Analysis of acidity motifs

Motif analysis was carried out using rmotifx^38^. The foreground comprised sequences 7 amino acids up- and down-stream of Y residues detected in peptidoforms with convincing histograms, and the background comprised all sequences surrounding Y residues detected across all peptidoforms for which GMMs were fit. The count of basic, polar, acidic, and neutral residues was computed for the same sets of sequences and fractions were calculated in R.

#### Multiple sequence alignments

A strong candidate Y sulfation site was identified on human Calumenin (UniProt: O43852), a member of the CREC family of ER-localised proteins^39^. This is in agreement with recently published data analysing sY residues found in the human secretome^11^. We next explored the evolutionary conservation of the tyrosine site and nearby amino acids, using an alignment extracted from our previous work^40^ exploring the evolutionary conservation of phosphorylation sites – sourced from 100 UniProt proteomes, aligned with MUSCLE^41^. Example USIs for PSMs giving evidence for sulfation were visualised via the ProteomeXchange USI tool.

#### Exploration of potential factors contributing to sY detection

Custom R scripts were used to explore potential relationships between different datasets, the instrument used to generate the data, and the ability to pinpoint potential sY sites. Briefly, the total count of all PSMs contributed by each instrument and each dataset ID were established across i) the entire phosphobuild, ii) peptidoforms assigned to the BOIs, iii) peptidoforms with manually assigned likely sY sites supported by convincing histograms and biological context, and iv) peptidoforms containing a known sY site. Contingency tables were generated for each category by instrument and by dataset ID (SupplementaryFile1). These contain the PSM count for each instrument-peptidoform and dataset-peptidoform combination respectively. Additionally, mass error shift histograms colour-coded by instrument, experiment tag, and dataset ID were generated to assess potential biases (SupplementaryFile2).

## Results

To explore our hypothesis that sulfopeptides may potentially have been misidentified as phosphorylation events in phosphoproteome datasets, FDR thresholding and subsequent per-raw-file mass error recalibration were performed on the PeptideAtlas human phosphoproteome build. We then aggregated PSM mass errors by peptidoform for phosphorylated peptidoforms (n = 1,747,947) and applied a GMM AUC filtering algorithm to assign peptidoforms into predefined mass error bins.

### GMM AUC filtering highlights potentially sulfated peptidoforms

While there is limited evidence of S and T sulfation in cells, almost all sulfation is found in the context of Y^42^. We thus hypothesised that if true sulfation events were detected by our pipeline, BOIs would be enriched in sY misidentified as pY and would thus be enriched in peptidoforms containing a pY PTM assignment. By association, we suspected that the fraction of peptidoforms that contain a Y may also be greater in the BOIs than in the DECOY and TRUEp bins. Our initial overview of the peptidoform proportions across each bin confirmed this hypothesis, revealing pY- and Y-containing peptidoforms were enriched in the BOIs, especially BOI2 and BOI3 (**Fig. 2**).

**Fig. 2.**
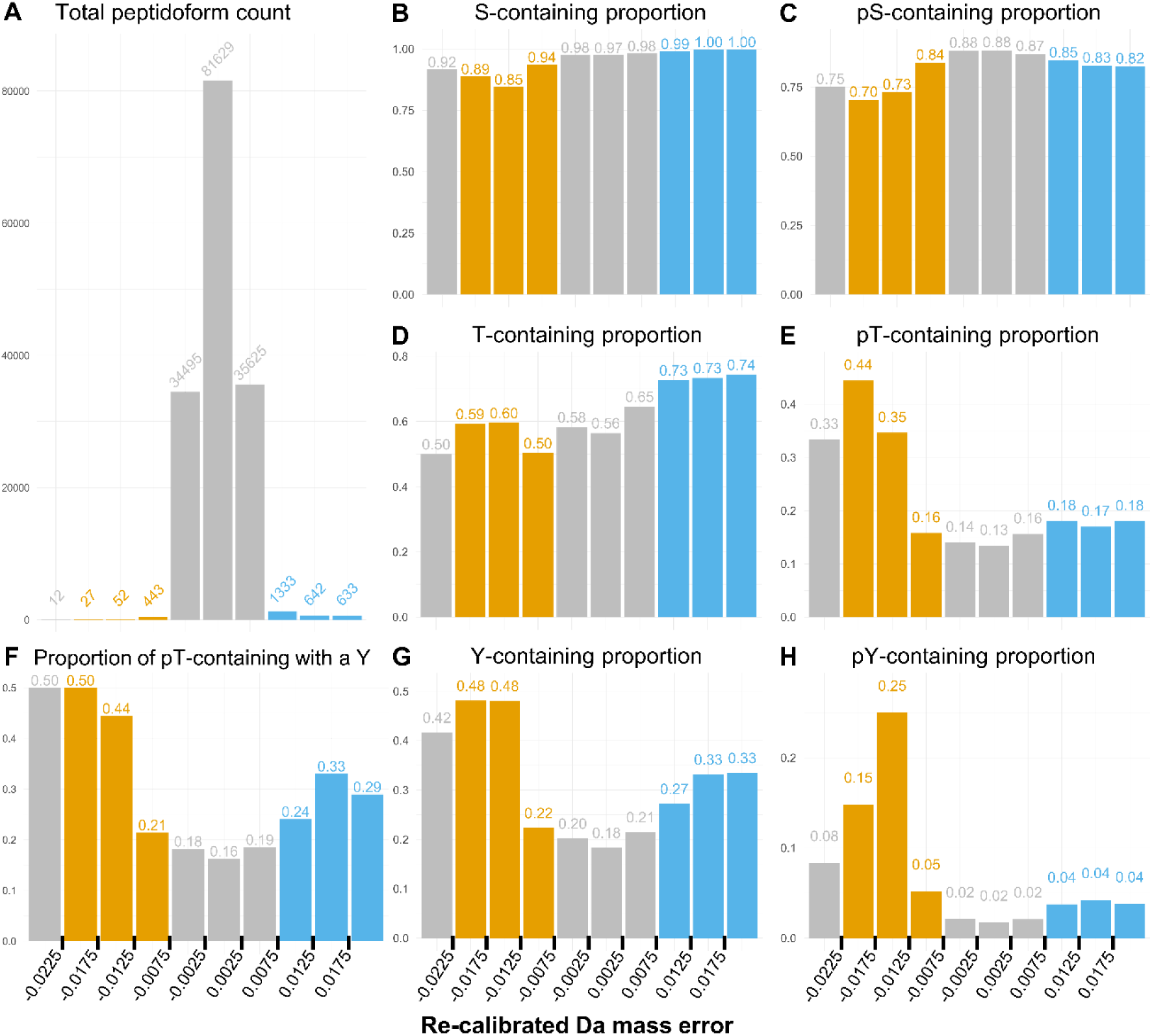
Peptidoform proportions across calibrated mass error bins. Of the total number of peptidoforms assigned to each bin (**A**), the proportion of peptidoforms that contain the specified amino acid (**B, D, G**), or phosphorylation assignment by the search engine (**C, E, H**) as per the plot title are presented. Additionally, the proportion of peptidoforms with a phosphorylation on Threonine that also contain a Tyrosine residue is illustrated (**F**). Orange denotes BOIs, blue denotes DECOY bins, the grey bins in-between are TRUEp bins. The left most grey bin comprises 12 peptidoforms, all of which are a subset of BOI3 to its right and was excluded from further analysis.

Of the 83,047 robustly detected peptidoforms, few were assigned to BOIs (**Fig. 2A**). In terms of absolute number, once data have been correctly calibrated, it is very rare overall to have peptidoforms assigned to BOIs (with mass shift -0.0225 to -0.0075 Da). It appears much more common to have peptidoforms with a positive mass shift, in the DECOY bins. These appear to be mostly caused by false identifications of deamidation on these peptides, which was included as a search parameter (+0.98 Da). A common phenomenon is that the instrument selects the +1 isotopomer for fragmentation, resulting in an experimental mass that is +1.003 Da higher than the actual peptide. In the DECOY1 bin, 13.65 % of peptidoforms are deamidated, rising to 87% and 90% in DECOY2 and 3 respectively. This is a >20-fold increase compared to the ∼ 4% of peptidoforms with deamidation assignment across the dataset (Supplementary Fig. 1). As such, our method can discover false assignments of deamidation.

As anticipated, our pipeline predominantly impacted the fraction of Y-containing peptidoforms but not the fraction of S-containing peptidoforms or T-containing peptidoforms, which served as negative controls in this comparison between bins (**Fig. 2 B, D, G**). Similarly to S-containing, pS-containing proportions were slightly lower in the BOIs, but not strongly impacted (**Fig. 2C**). Surprisingly, peptidoforms in BOI2 and BOI3 were enriched in pT assignments (**Fig. 2E**). We observed that of the pT-containing peptidoforms, a larger proportion contains a Y residue in BOI2 and BOI3 compared to the remaining bins (**Fig. 2F**). Given the small number of peptidoforms in these bins it was difficult to draw conclusions from this observation. Possible explanations include: i) these could be peptidoforms with T sulfation events misidentified as phosphorylation; ii) pT may accompany sY events; iii) the PTM assignment algorithm of the source pipeline may be more prone to incorrectly assigning pT instead of sY in peptides with both a T and a Y; or a combination of all these.

While peptidoforms containing a tyrosine made up roughly 20% of all peptidoforms with a phosphorylation event involved in this analysis, 50% of all peptidoforms in BOI2 and BOI3 contained a tyrosine residue (**Fig. 2G**). The most prominent shift was observed in the proportion of pY-containing peptidoforms, which had increased more than ten-fold in BOI2, reaching 25%, compared to TRUEp bins, where only 2% of all peptidoforms contained a pY assignment (**Fig. 2H**). Although these findings were promising evidence for the pipeline’s ability to distil sY-containing peptidoforms misidentified as pY, the relatively small number of peptidoforms assigned to BOI3 and BOI2 make the observed fractions more prone to random chance. In addition, the fraction of pY-containing peptidoforms in BOI1 was higher than in the TRUEp bins, but not much higher than in the DECOY bins.

We hypothesised that the 15% AUC filtering cutoff for assigning a peptidoform to a bin may have resulted in some false positives with wide mass error distributions, or mass error distributions slightly shifted in the negative direction being assigned to BOI1.

To explore this possibility, we manually assessed the mass error histograms of the 104 Y-containing peptidoforms assigned to the BOIs (SupplementaryFile3). Of these, 49 peptidoforms had convincing histograms, 9 were filtered out as false positive assignments to the BOIs, and 46 were labelled as ‘Undetermined’, indicating a mass error shift towards sulfation, but of less than -0.006 Da magnitude. Example histograms are presented in **Fig. 3**.

**Fig. 3.**
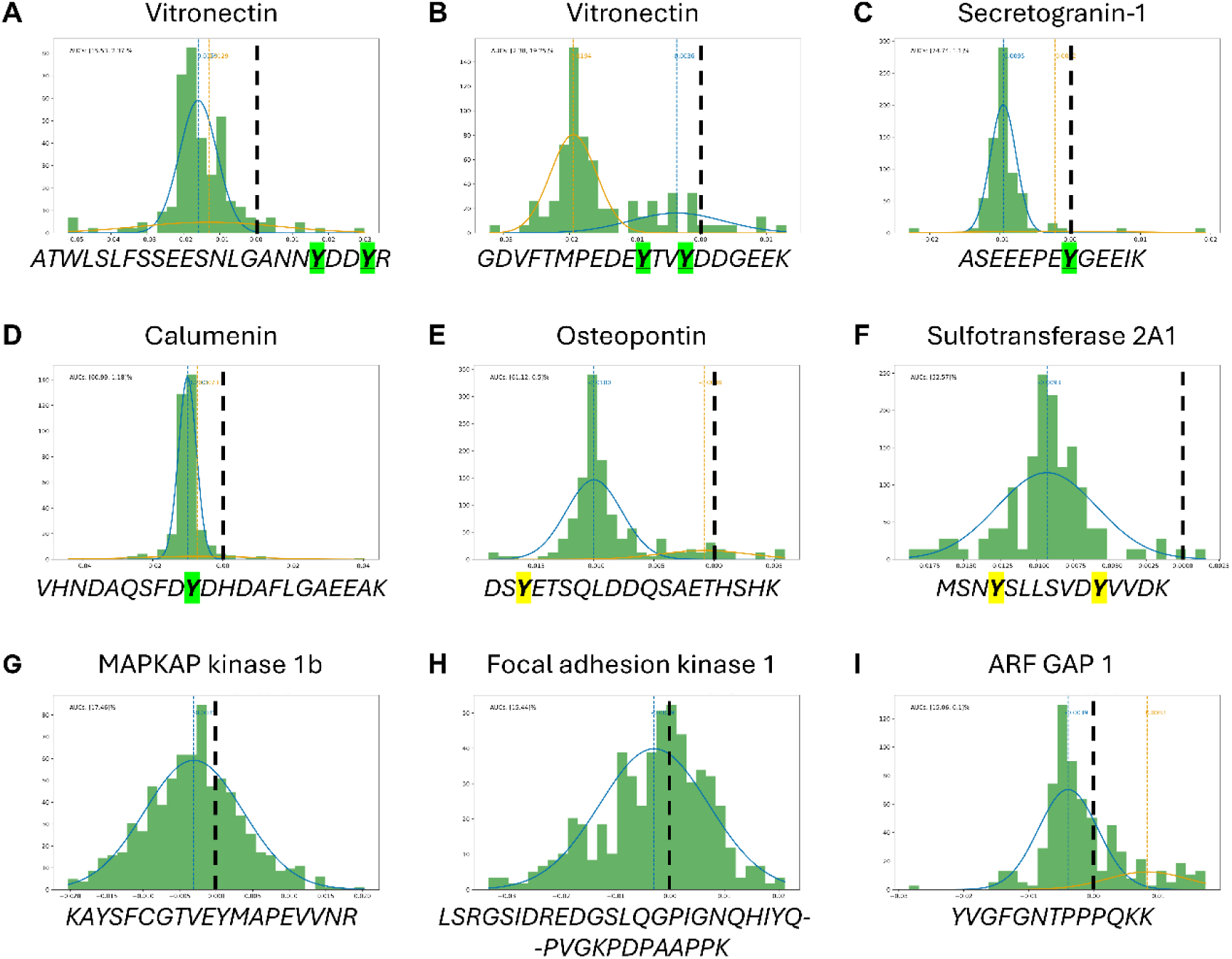
Example histograms of peptidoforms assigned to BOIs. Following GMM filtering, peptidoforms assigned to the BOIs could be split into three groups: peptidoforms with known sY sites (**A-D**); peptidoforms with potential novel sY sites (**E-F**); and hits with unconvincing histograms – either false positives or undetermined (**G-I**). Each plot presented is a histogram of the calibrated mass error of all PSMs for a peptidoform, with the best fit lines of each GMM component. The black dashed line intersects the mass error shift point of 0 Da on the x-axis. The assigned protein ID is printed above the histogram, and the peptide sequence is printed below, with known and potential novel sY sites highlighted. Presented are examples of each group, the full list of peptidoforms assigned to BOIs can be found in SupplementaryFile3.

Notably, 24, or roughly half of the convincing histograms were for peptidoforms covering known sY sites. Of these, 15 peptidoforms covered two peptide sequences of human Vitronectin (Uniprot ID: P04004), both with two known sY sites (**Fig. 3A, B**), and one covered a known singly-sulfated peptide of Secretogranin-1 (**Fig. 3C**). Strikingly, our pipeline identified one Calumenin (Uniprot ID: O43852) and one Golgi integral membrane protein 4 (GIMPc, Uniprot ID: O00461) sY site for which we recently provided experimental validation in separate work^11^. The Calumenin sulfation site at Y47 was supported by six peptidoforms in the BOIs, making it a particularly convincing identification (**Fig. 3D**). GIMPc was supported by two peptidoforms spanning the acidic peptide sequence ELEHNAEET**<u>Y</u>**GENDEN**T**DDKNNDGEEQEVR. The histograms for both demonstrated a similar mass error distribution indicative of a singly sulfated peptide (SupplementaryFile4). Interestingly, in one peptidoform, phosphorylation had originally been assigned to the threonine, while in the other it had been assigned to the tyrosine (SupplementaryFile3). This could be an example of a sY sometimes misidentified as pT, and sometimes misidentified as pY.

Given the presence of different singly phosphorylated peptidoforms of the same peptide, it would be of interest for future studies to incorporate site specificity into the analysis. This could be done based on positional assignment scores (e.g. from PTM Prophet) while retaining more peptidoform granularity by pooling data for peptidoforms with the same positional assignments, rather than the ‘non-strict’ peptidoforms we analyse here. However, this could reduce the number of robustly detected peptidoforms, lowering the coverage of the analysis.

The remaining 25 peptidoforms with convincing histograms constituted examples of peptidoforms with potential novel sY sites. Notably, six of these covered two distinct Osteopontin (Uniprot ID: P10451) peptide sequences (**Fig. 3E, SupplementaryFile3**). This was an exciting finding because Osteopontin is known to be subject to numerous PTMs including an identified sulfation site at Y165 ^43,44^. However, no prior knowledge exists for the sY sites identified by our pipeline. For the remaining potential novel sulfated peptidoforms, one peptidoform was detected per protein, as in the case of human Sulfotransferase 2A1 (Uniprot ID: Q06520) (**Fig. 3F**). Sulfotransferase 2A1 was of particular interest, since our previous research has identified other sulfotransferases with sY residues^11^.

Another example to note were two peptidoforms with convincing histograms spanning the EDSMDMDMSPLRPQN**<u>Y</u>**LFGCELK peptide sequence of the highly modified protein Nucleophosmin (Uniprot ID: P06748). The mass shifts for one of these were indicative of two sulfation events (ss Histogram type, centred at ∼ -0.0195 Da), but the peptide sequence only contains one Y residue (SupplementaryFile3). Since no prior biological knowledge makes Nucleophosmin a likely candidate for tyrosine sulfation, this may be a false positive hit (e.g. a misidentified peptide). However, this example may also be evidence of sulfation in amino acids other than Y, such as T and S, which require further validation in a biological system.

To explore the biological context of our findings without bias to previously reported sY sites, we investigated the presence of amino acid motifs surrounding the Y sites using convincing histograms. We hypothesised that motifs would match the acidic consensus for sulfation by TPST1 and TPST2^45^. Although not present across all motifs, it was evident that acidic residues (D/E) are commonly found (7 out of 15 motifs) within a span of 7 amino acids up- and down-stream of the Y sites of interest, as expected (Supplementary Fig. 2). However, this analysis was limited by the small number of unique peptide sequences in the foreground (n=35). This resulted in motifs supported by only two or three sequences with weak enrichment scores. Nevertheless, the low abundance of these motifs in the background was reassuring, resulting in large fold changes for the presented motifs.

To further explore the Y sulfation acidity hypothesis, which was proposed several decades ago^46^ the total number of basic, acidic, neutral, and polar residues within 7 amino acids of a Y was computed for Y sites of interest and all Y sites found across the 83, 047 peptidoforms for which GMM models were fit. Across the dataset, 14% of amino acids surrounding a Y were acidic (D or E). The acidic composition increased to 19.72% for Y sites covered by peptidoforms with convincing histograms, while the neutral amino acid composition (A, V, L, I, M, F, W, P, or G) remained similar at nearly 38%. The increase in acidity was accompanied by a decrease in basic residues (K, R, or H) from 12.42% to 10.7% and a decrease in polar residues (S, T, Y, N, Q, or C) from 35.67% to 31.83% for the Y sites of interest. Overall, these results suggest that peptidoforms with convincing histograms are more acidic on average, which supports the biological context of tyrosine sulfation by TPSTs.

### Peptidoform assignment to BOIs is supported by biological context

To further explore the biological context of the peptidoforms assigned to the BOIs in an unbiased manner, we carried out GO term enrichment analyses with each BOI and DECOY bin as foreground, and all 83,047 peptidoforms derived from 2,758 proteins as background. Given that Y sulfation is primarily observed in secreted and transmembrane proteins and is catalysed in the Golgi^11,47^, we hypothesised that terms associated with these cellular components would be overrepresented in the BOIs. The DECOY bins served as negative controls.

Significantly enriched terms across the BOIs could be linked to sulfation (**Fig. 4**). More specifically, ‘integrin binding’ was the only GO term significantly enriched in BOI1. This term was close to significant in BOI2 (p adj. < 0.1) alongside ‘sulfur compound binding’, ‘ECM structural constituent’, ‘ER lumen’, and ‘Golgi lumen’. None of these terms were significantly enriched in any of the DECOY bins. Surprisingly, multiple GO terms were significantly or close to significantly enriched (p adj.<0.1) in DECOY1 and DECOY3, but not DECOY2. These were predominantly terms associated with RNA binding, some of which overlapped between the two DECOY bins, for reasons unclear. None of the terms significant in the DECOY bins were directly linked to sulfation.

**Fig. 4.**
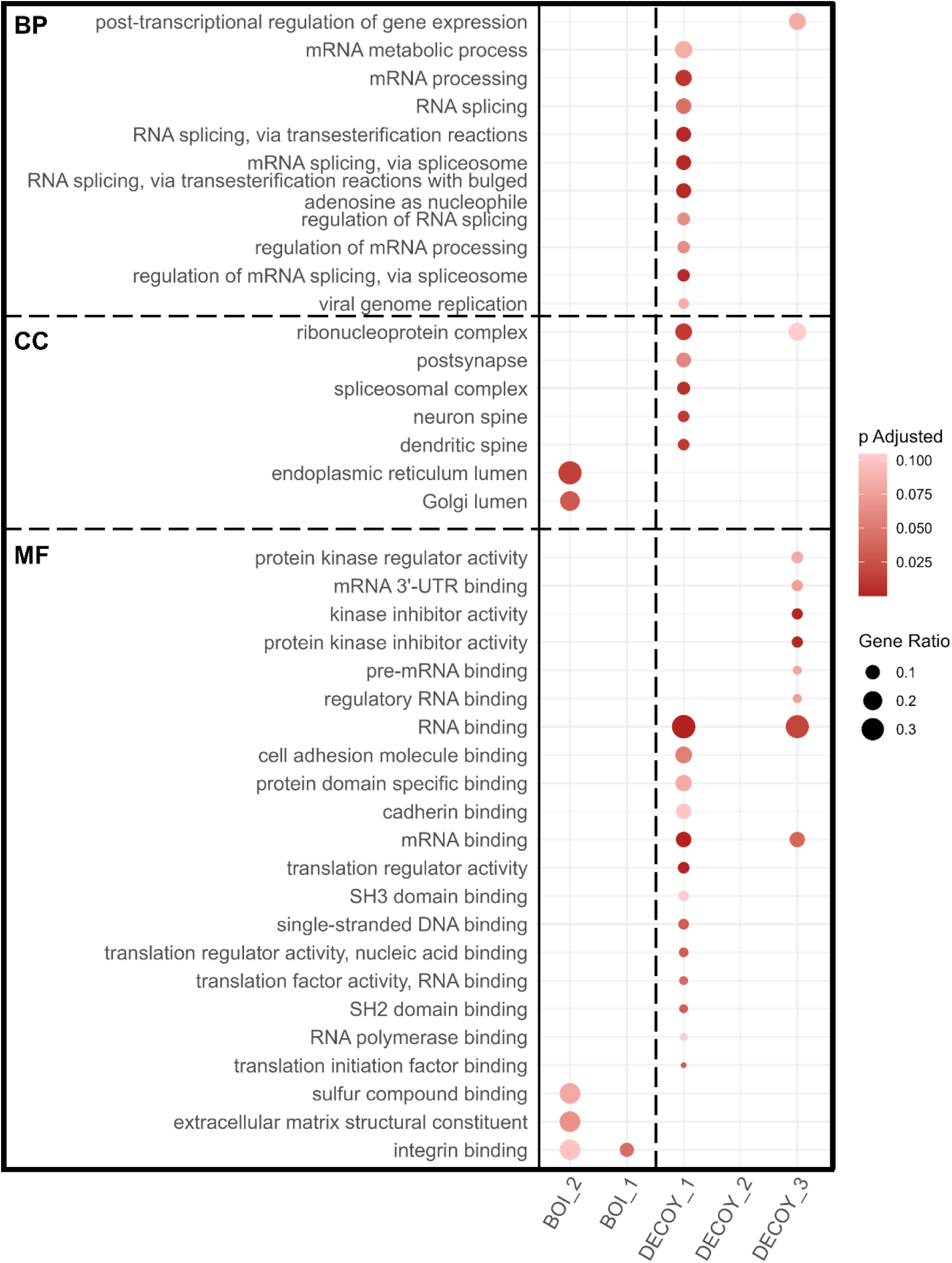
GO Term overrepresentation analysis results by bin. Rows represent labelled GO terms enriched (q<0.01) in one or more bins. BP, CC, and MF terms are separated by a horizontal dashed line and ordered by gene ratio (represented by circle size) and by bin. BOI and DECOY bin results are separated by a vertical dashed line. Colour-coded are the Benjamini-Hochberg-adjusted p-values, with most significant hits coloured darker hues of red. All protein lists used for the analysis can be found in SupplementaryFile5.

One limitation of this analysis in exploring the hypothesised biological context stemmed from GO terms not being strictly associated with sY. Given the initial evidence of some GO terms that could be linked to sulfation, we repeated the overrepresentation analyses for each bin using custom term mappings for the proteins in UniProtKB to explore results in greater depth (**Fig. 5**). The custom terms were: ‘known sY’, ‘cytokines’, ‘Secreted’, ‘Golgi’, ‘Transmembrane’, and all other proteins as ‘unlikely sY’. Term to protein mapping is available in SupplementaryFile5.

**Fig. 5.**
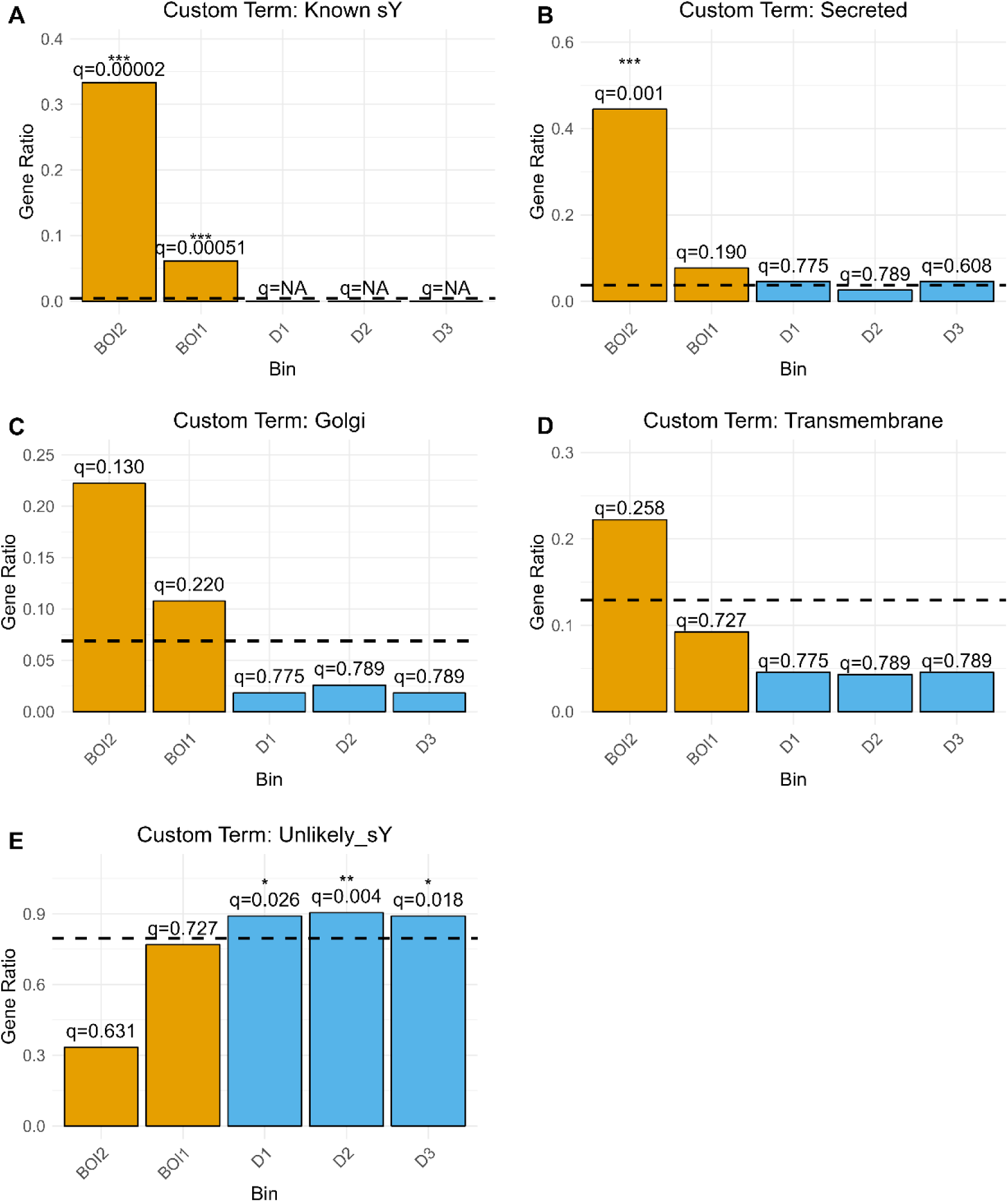
Custom term overrepresentation analysis results by bin. Presented are the ORA results for each custom gene term across BOIs (orange) and DECOY bins (blue). Panels **A-D** outline the results for enrichment of peptidoforms mapped to tyrosine sulfation-related terms, as indicated by each plot title. Panel **E** outlines the results for enrichment of peptidoforms unlikely to be related to tyrosine sulfation. The dashed line in each plot represents the gene ratio for that term detected across the entire dataset. The q-values and asterisk above indicate significance level.

Peptidoforms derived from proteins with known sY sites were significantly enriched in BOI1 and BOI2 (q<0.001), while no such peptidoforms were detected in the DECOY bins (**Fig. 5A**). The term ‘Secreted’ was strongly enriched in BOI2 (q<0.01) but not BOI1 or any of the DECOY bins (**Fig. 5B**). Nevertheless, the ratio of secreted protein peptidoforms was higher in BOI1 compared to their ratio across the entire dataset. Although not significantly, Golgi protein peptidoforms were detected in BOI1 and BOI2 at higher ratios than in the background. In contrast, their detection rate in the DECOY bins was halved or lower compared to the background (**Fig. 5C**). A similar pattern was observed for Transmembrane protein peptidoforms, except their ratio in BOI1 was below the background level (**Fig. 5D**). The exact opposite was observed for peptidoforms derived from proteins unlikely to contain sulfated tyrosine residues – these were significantly enriched in all DECOY bins (q<0.05), while their ratio in BOI1 was similar to that in the background and their ratio in BOI2 was diminished (**Fig. 5E**). Overall, the enrichment analyses of GO terms and custom terms suggest we can find known sY based on mass error shifts, and there is supporting biological evidence that we may be able to find potential novel sY sites. However, the number of proteins associated with peptidoforms that have passed the stringent pre-GMM filtering and are likely to contain sY is small (9 in BOI2, and 65 in BOI1). Notably, these analyses aimed to be unbiased and did not consider the manual annotation of peptidoform mass error histograms.

### A combination of mass error accuracy, biological context, and MS^2^ spectral evidence supports known and novel sY sites

To identify the most promising potentially sulfated peptidoforms, biological context and mass error evidence were integrated, resulting in a final list of 31 candidates derived from seven proteins (SupplementaryFile3). In total, fourteen sY sites were identified, of which seven were novel (**Table 1**). Notably, some singly sulfated peptides (histogram type ‘s’) had two potential sY sites. In such cases both sites were deemed novel sites that will require further experimental disambiguation.

**Table 1.**
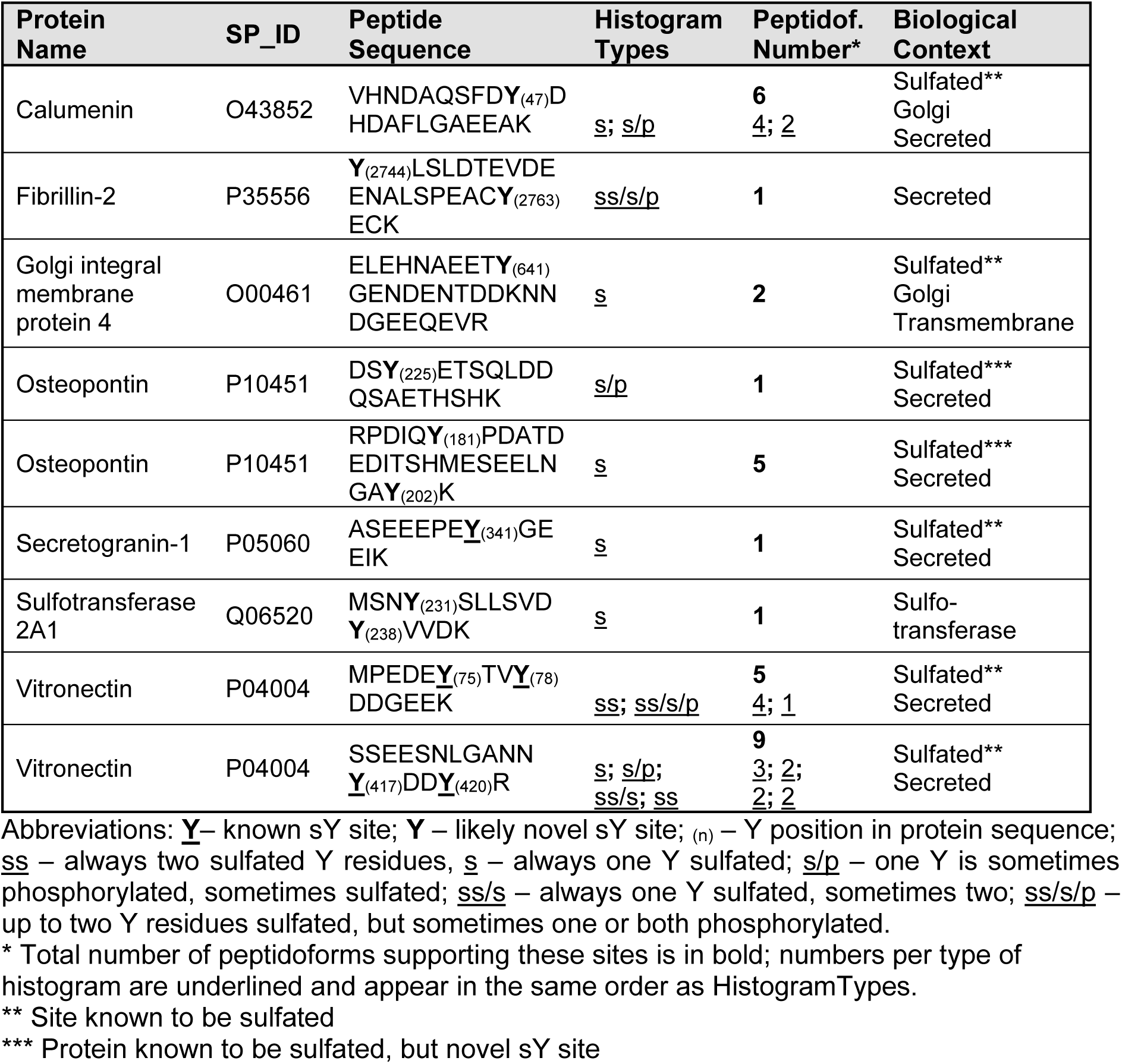
High confidence sY sites identified by our pipeline.

We hypothesised that the strong candidate sulfation sites in Table 1 would be further supported by the MS^2^ spectra across their associated PSMs. To test this hypothesis, we generated USIs for all PSMs across all convincing peptidoforms covering both phosphorylation and sulfation instances (SupplementaryFile6). We expected candidate sulfated peptides to have fuller annotations and explain a larger total ion current percentage than phosphorylation candidates in instances where sulfation had been misidentified.

We focused on the Calumenin Y47 site which was of particular interest, since this site was shown to be sulfated rather than phosphorylated in the secretome of immortalised human embryonic kidney (HEK-293) cells following the development of our sY LC-MS workflow^11^. Fig. 6 displays the spectrum of the lowest q-value PSM for a peptidoform supporting sulfation of Calumenin Y47, with annotations inferred from sulfation with a full neutral loss (A) versus a phosphorylation hypothesis with no assumed neutral loss (B). There is a run of five b2+ ions (b13-b17), only observed under the hypothesis that the peptide contains a fully labile sulfation event on peptide position 10. It is noteworthy that UniProtKB has annotated position 47 as a pY, based on multiple MS/MS datasets. We cannot find any evidence in PeptideAtlas for an MS1 experimental mass observation that supports phosphorylation at this position, and thus believe Y47 to be sulfated but not phosphorylated, as further supported by the current meta-analysis (Supplementary Fig. 3).

**Fig. 6.**
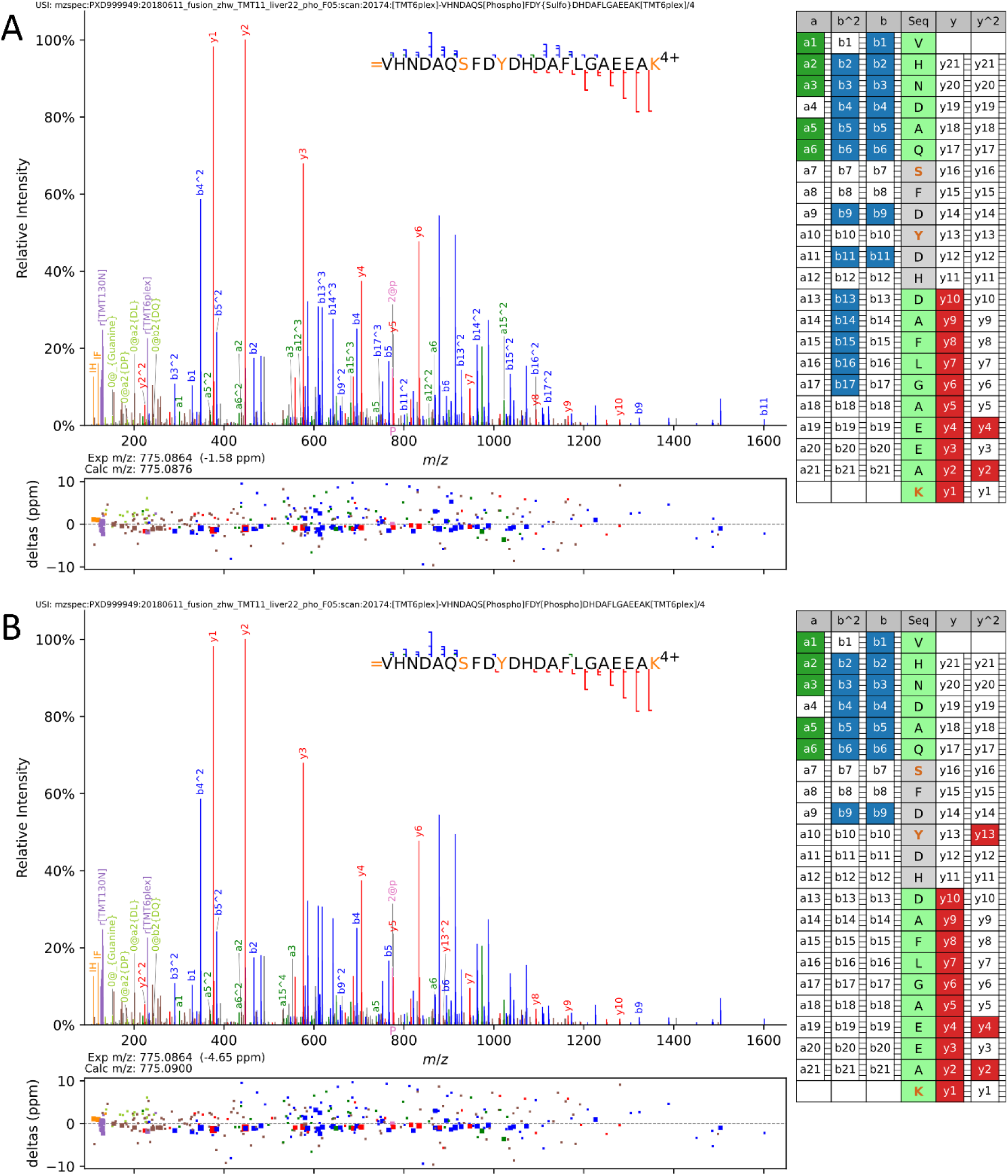
MS^2^ evidence for Calumenin sY47. Visualisation of spectra and annotation evidence for the top scoring peptidoform for VHNDAQSFDYDHDAFLGAEEAK from Calumenin, comparing A) a hypothesis of phosphorylation on position 7 (S44) and fully labile sulfation on position 10 (Y47) against B) phosphorylation on positions 7 and 10. Hypothesis A has a convincing run of b2+ ions (13-17) in series that are only detected with a full loss of the sulfation mass, but not observed with a hypothesis of phosphorylation. Spectra were visualised using Quetzal^48^, available at https://proteomecentral.proteomexchange.org/quetzal with a tolerance of 10 ppm, and labelling of neutral loss ions and internal fragmentation ions disabled in order to reduce clutter from the many other annotations.

To evaluate enzymatic Y sulfation of calumenin, we purified a non-sulfated human GST-full length calumenin fusion protein. Interestingly, even after size-exclusion, calumenin behaves anomalously during SDS-PAGE, presumably due to its acidic nature (pI=4.47). Soluble GST-calumenin migrates as both a monomeric and multimeric complex (Supplementary Fig. 4A). We next subjected calumenin to *in vitro* sulfation^13^ using catalytically-active TPST1 or TPST2 or a combination of the two enzymes and Mg^2+^-PAPS as sulfuryl donor. As shown in Supplementary Fig. 4B, immunoblotting with a sTyr-specific antibody revealed that calumenin becomes sulfated by TPST1, and hypersulfated by a combination of TPST1 and 2, in a time-dependent manner. LC-MS/MS and comprehensive spectral annotation confirmed that Y47 was amongst the sulfated residues present after TPST1/2-treatment (Supplementary Figure 5, Top), and two novel sites, including Y263, which also lies in a DYD motif, were confirmed through manual inspection of the peptide spectra (Supplementary Figure 5, Bottom and Supplementary Table 1). For further biological contextualisation, we explored the conservation of Y47 in non-human species, using a previously-published protein alignment of orthologs^40^. The Y47 site displayed a strong conservation within an acidic consensus sequence across a wide range of vertebrates, although this was less commonly observed in plants, protists, and invertebrates (Supplementary Fig. 6).

We also explored the trends in instrument and dataset contributions to sY detection looking for patterns that may lead to improved sY detection. The majority of datasets with large contributions to convincing sY peptidoforms were from the National Cancer Institute’s Clinical Proteomic Tumor Analysis Consortium (CPTAC)^49^ and other cancer-related datasets (Supplementary Fig. 7). This was in line with the bias of the Human Phosphobuild, in which 92% of PSMs originate from samples categorised as cancer cell lines or tissues. Another bias of the dataset was toward intracellular extracts. Therefore, given that thus far tyrosine sulfation has been found predominantly on secreted and transmembrane proteins, the current study may not accurately represent the rate of misidentification of sY as pY in secretome samples, where sulfation should be a more common event. In terms of instrumentation, the Q Exactive contributed a disproportionately large number of PSMs to the BOIs (close to 70%), likely being a major contributor of false positive peptidoform assignments to this bin (Supplementary Fig. 8). Overall, the contribution pattern across the convincing peptidoforms was similar to the pattern across the dataset, with the Orbitrap Fusion Lumos, Q Exactive, and Q Exactive HF being top contributors. This suggests that when operated at sufficient resolution, modern instruments should enable sY to be distinguished from pY based on accurate precursor mass shifts.

It would be interesting for future studies to explore in more depth the correlation between sulfation and different sample types, tissues, instrument settings and other technical aspects. This would require advancements in our ability to distinguish sulfation from phosphorylation within individual PSMs, perhaps by collating large amounts of MS^2^ data for known sulfated and phosphorylated peptides and using it to train machine learning models to classify PSMs into sulfated or phosphorylated on the basis of fragmentation patterns (e.g. based on differences in the propensity for neutral loss fragments for sulfo-versus phospho-peptides).

### Conclusions

This study provides insights into the potential for detection of sulfotyrosine within phosphoproteomics datasets, and the extent to which physiological sulfation sites may have been misidentified as phosphorylation sites in cell and tissue intracellular extracts. The combined application of GMMs and manual annotation of peptidoform mass error distributions have proven effective in identifying a set of novel sY sites, which will need to be experimentally validated in the future. The enrichment of specific GO terms and custom terms related to sulfation among our candidate peptidoforms and the supporting MS^2^ spectra confirmed the likely biological relevance of these results. Our findings clearly show that sY can be misidentified as pY, as demonstrated in the case of Calumenin Y47, for which we present multiple lines of evidence that this residue is sulfated, rather than phosphorylated. It will be interesting to compare and contrast Y sulfation and phosphorylation on other members of the human CREC-family of low-affinity calcium-binding proteins, which lie in the same part of the secretory pathway where tyrosine sulfation is catalysed. Our analysis demonstrates that although possible, sulfation misidentification as phosphorylation is an extremely rare occurrence (tens of convincing misidentifications in a ‘haystack’ of tens of thousands peptidoforms from cellular proteomics datasets). It would be interesting to compare these rates in secretome samples, where phosphorylation and sulfation are relatively common PTMs^50^. Overall, our data strongly indicate that current phosphoproteomic methods are generally highly reliable for correct attribution of a mass shift to the named phosphorylation-based PTM, especially when appropriate optimised sample preparation, MS techniques and analysis workflows are employed.

## Acknowledgements

We would like to thank the Computational Biology Facility team at the University of Liverpool for their constructive feedback and technical expertise that assisted this project.

## Author contributions

The manuscript was written through contributions of all authors. All authors have given approval to the final version of the manuscript.

## Data Availability

All reanalysed phosphoproteomics datasets are open access. The relevant dataset identifiers and metadata can be found on the Human Phospho PeptideAtlas 2022-24 Build detailed summary page under the Experiment Contribution tab (https://db.systemsbiology.net/sbeams/cgi/PeptideAtlas/buildDetails?atlas_build_id= 537). The newly acquired mass spectrometry proteomics data supporting Calumenin Y47 sulfation have been deposited to the PRIDE Archive (http://www.ebi.ac.uk/pride/archive/) via the PRIDE partner repository with the data set identifier PXDXXXXX (identifier nor received yet, to be provided during proofs).

## SUPPORTING INFORMATION

**The following supporting information is available free of charge at ACS website** http://pubs.acs.org

Supplementary information: Supplementary_Information_Figures_and_Tables.docx; Contains all supplementary figures and tables. Some additional narrative is provided to the Calumenin Y47 sulfation evidence and our brief exploration of factors that may contribute to sY detection (DOCX).

Supplementary File 1: SupplementaryFile1.zip; ZIP folder containing PSM counts contingency tables (CSV)

Supplementary File 2: SupplementaryFile2.zip; ZIP folder containing histograms for peptidoforms of interest colour-coded by dataset/instrument/experiment tag (PDF) Supplementary File 3: SupplementaryFile3.xlsx; Manual annotations of peptidoforms assigned to BOIs (XLSX)

Supplementary File 4: SupplementaryFile4.zip; ZIP folder containing a collection of histograms for each Bin of Interest (PDF); Histograms are displayed for peptidoforms with > 15% AUC falling over that bin (actual AUC % shown on plot)

Supplementary File 5: SupplementaryFile5.zip; ZIP folder containing the foreground proteins in each bin used in the overrepresentation analyses, the background universe (all proteins detected across all bins, including TRUEp), and the term to protein mapping for the custom terms analyses.

Supplementary File 6: SupplementaryFile6.xlsx; USIs generated for convincing peptidoforms (XLSX)

## For Table of Contents Only

### Graphical Abstract

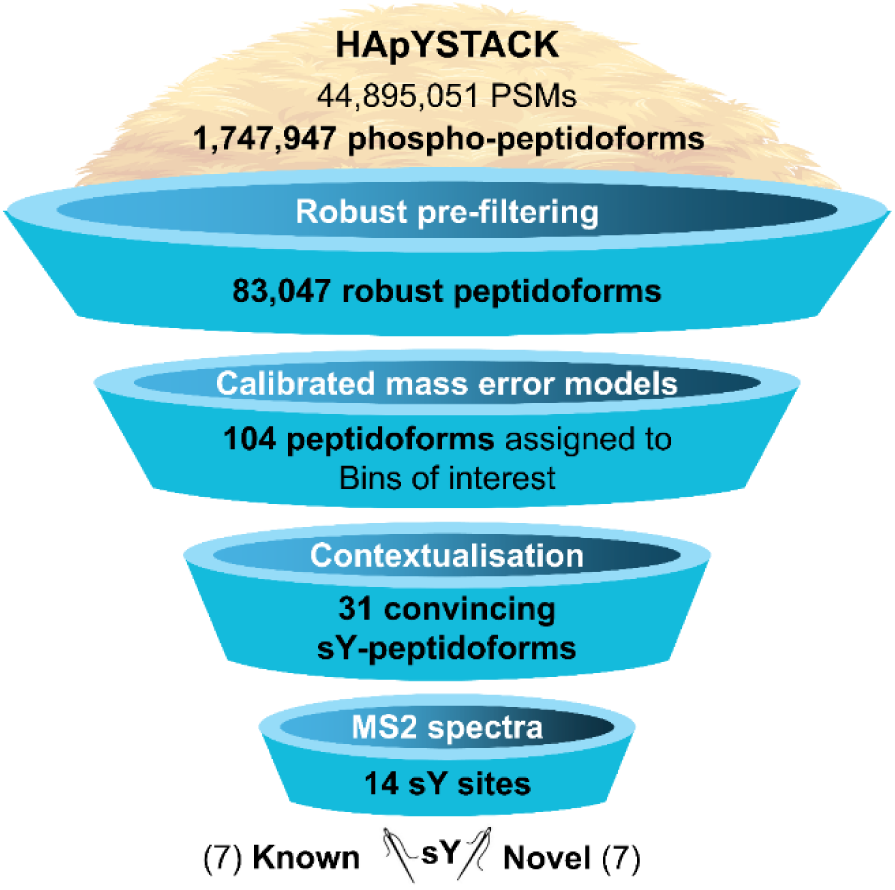

