## Supplementary information, tables and figures for "Searching for Sulfotyrosines (sY) in a HA(pY)STACK"

**Table of Contents**

Supplementary information (This document): Supplementary_Information_Figures_and_Tables.docx; lists all supplementary files and contains all supplementary figures and tables. Some additional narrative is provided to the Calumenin Y47 sulfation evidence and our brief exploration of factors that may contribute to sY detection (DOCX).

Supplementary File 1: SupplementaryFile1.zip; ZIP folder containing PSM counts contingency tables (CSV)

Supplementary File 2: SupplementaryFile2.zip; ZIP folder containing histograms for peptidoforms of interest colour-coded by dataset/instrument/experiment tag (PDF)

Supplementary File 3: SupplementaryFile3.xlsx; Manual annotations of peptidoforms assigned to BOIs (CSV)

Supplementary File 4: SupplementaryFile4.zip; ZIP folder containing a collection of histograms for each Bin of Interest (PDF); Histograms are displayed for peptidoforms with > 15% AUC falling over that bin (actual AUC % shown on plot)

Supplementary File 5: SupplementaryFile5.zip; ZIP folder containing the foreground proteins in each bin used in the overrepresentation analyses, the background universe (all proteins detected across all bins, including TRUEp), and the term to protein mapping for the custom terms analyses.

Supplementary File 6: SupplementaryFile6.xlsx; USIs generated for convincing peptidoforms (CSV)

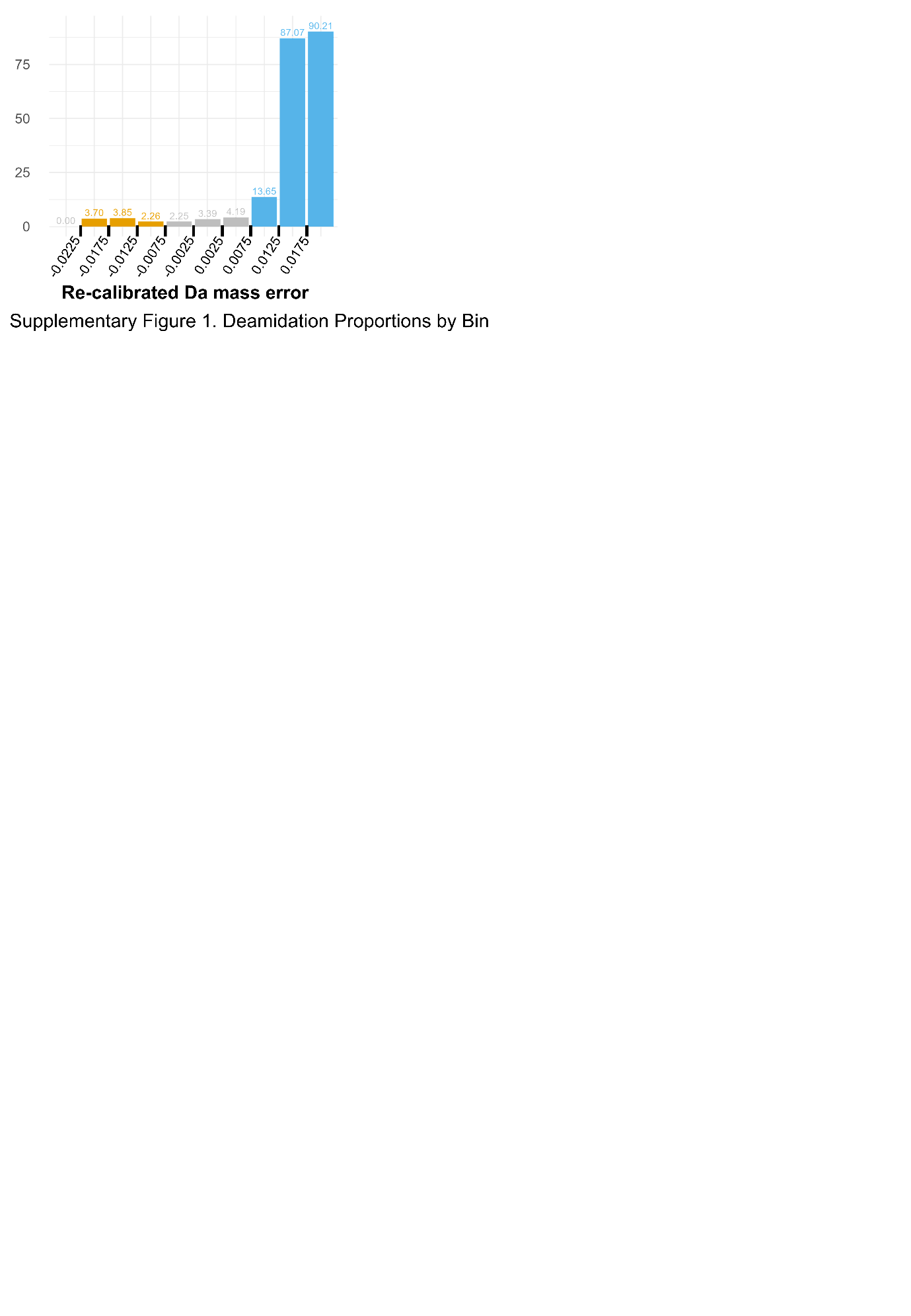

**Supplementary Figure 1.** Deamidation Proportions by Bin

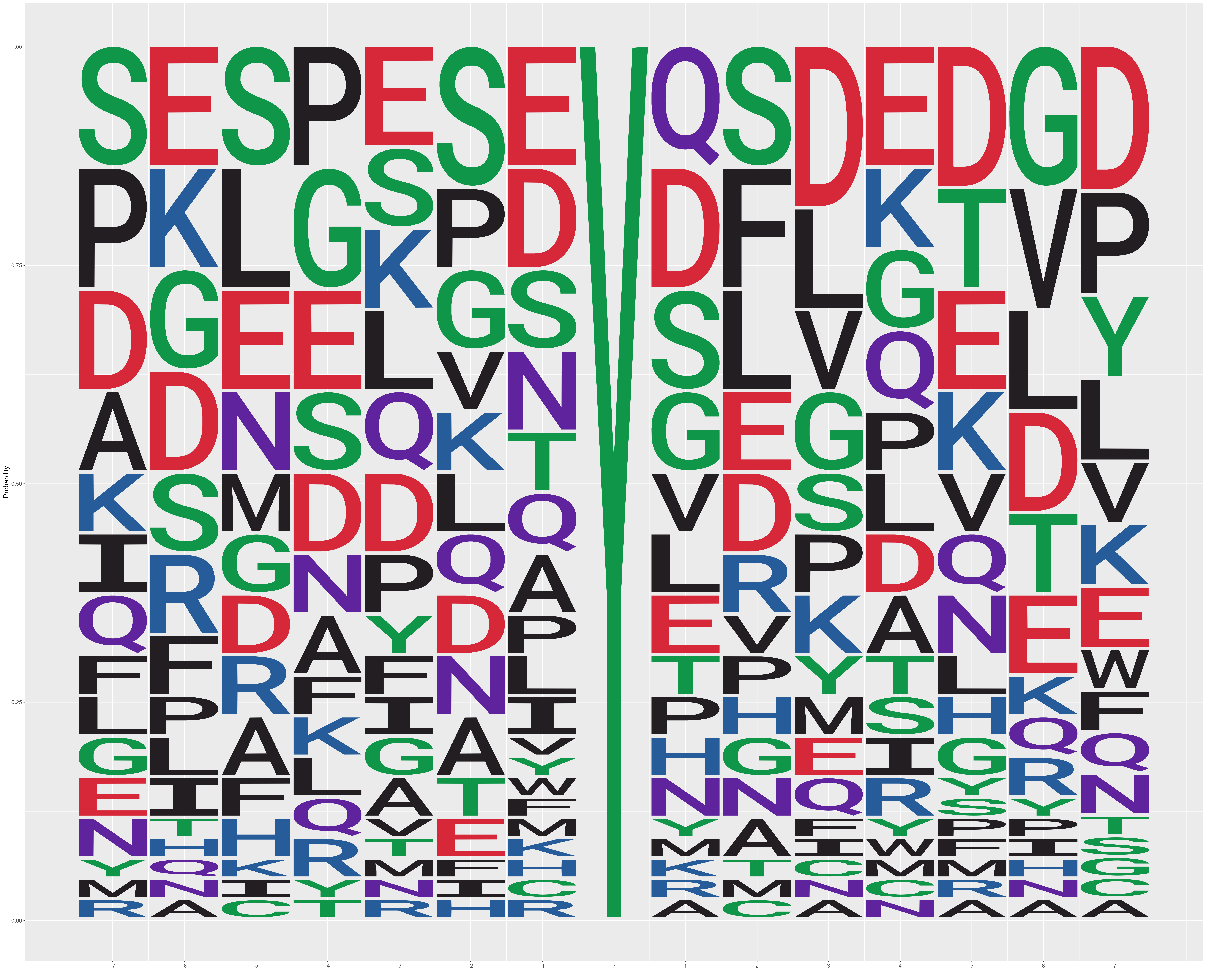

**Supplementary Figure 2.** Motif Enrichment in peptidoforms with convincing histograms

**Calumenin Y47 site is sulfated and not phosphorylated**

Even in the two peptidoforms with histograms scored as type ‘s/p’ that span the Calumenin Y47 site, it is evident that pY47 assignments are likely erroneous due to the ambiguity in the search engine when assigning the phosphorylation PTM position (Supplementary Figure 3). Since both peptidoforms are singly phosphorylated, ambiguity arises between assigning pS44 and pY47. In doubly phosphorylated peptidoforms the positional assignment is disambiguated since these are the only two residues that could be phosphorylated. For such peptidoforms, the mass errors are normally distributed around the sulfo-shifted mass error of ~ -0.01 Da with no evidence of tyrosine phosphorylation. All of these peptidoforms were scored as ‘s’ for sulfated-only during the manual annotation phase as summarised in Table 1.

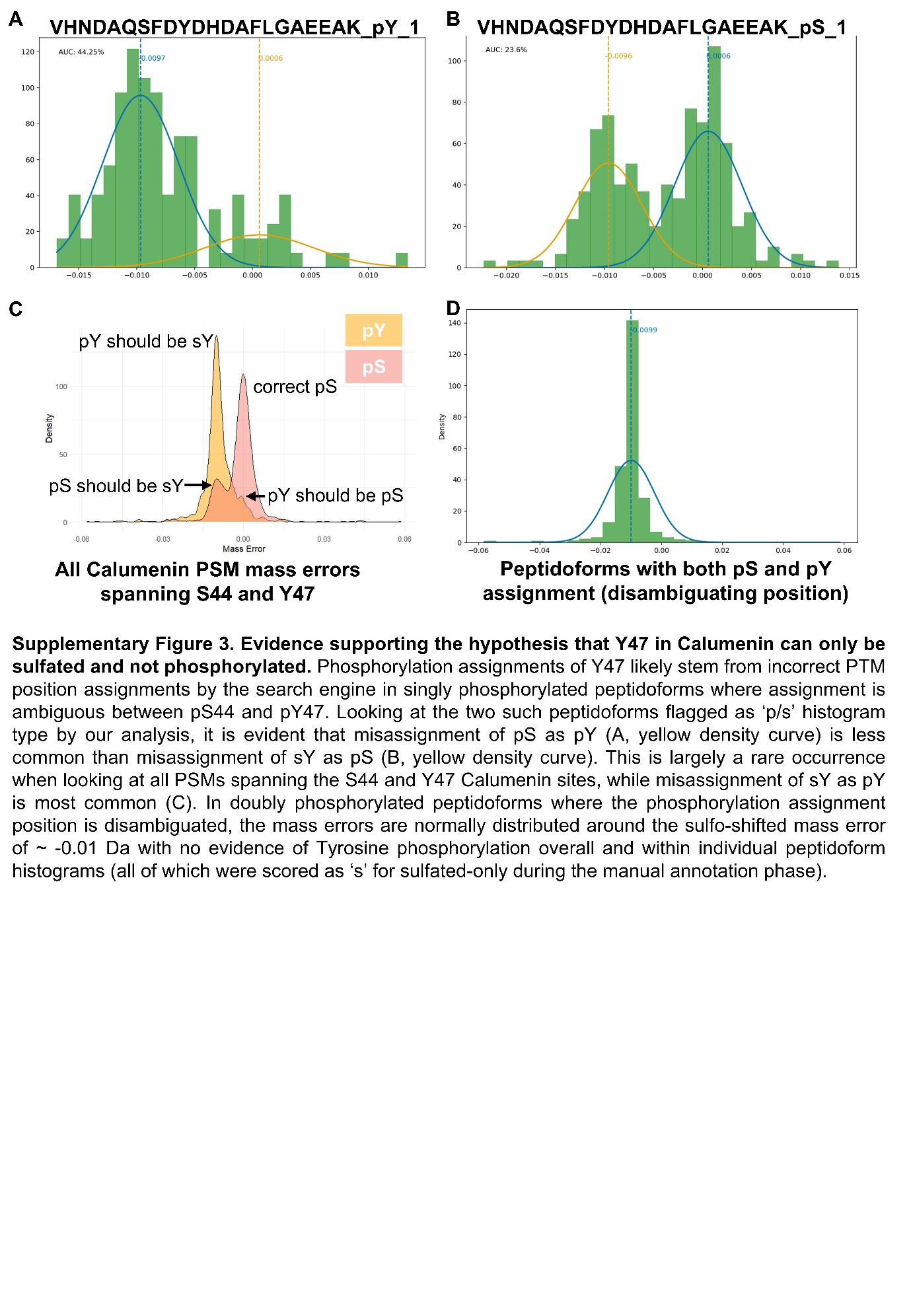

**Supplementary Figure 3. Evidence supporting the hypothesis that Y47 in Calumenin can only be sulfated and not phosphorylated.** Phosphorylation assignments of Y47 likely stem from incorrect PTM position assignments by the search engine in singly phosphorylated peptidoforms where assignment is ambiguous between pS44 and pY47. Looking at the two such peptidoforms flagged as ‘p/s’ histogram type by our analysis, it is evident that misassignment of pS as pY (A, yellow density curve) is less common than misassignment of sY as pS (B, yellow density curve). This is largely a rare occurrence when looking at all PSMs spanning the S44 and Y47 Calumenin sites, while misassignment of sY as pY is most common (C). In doubly phosphorylated peptidoforms where the phosphorylation assignment position is disambiguated, the mass errors are normally distributed around the sulfo-shifted mass error of ~ -0.01 Da with no evidence of Tyrosine phosphorylation overall and within individual peptidoform histograms (all of which were scored as ‘s’ for sulfated-only during the manual annotation phase).

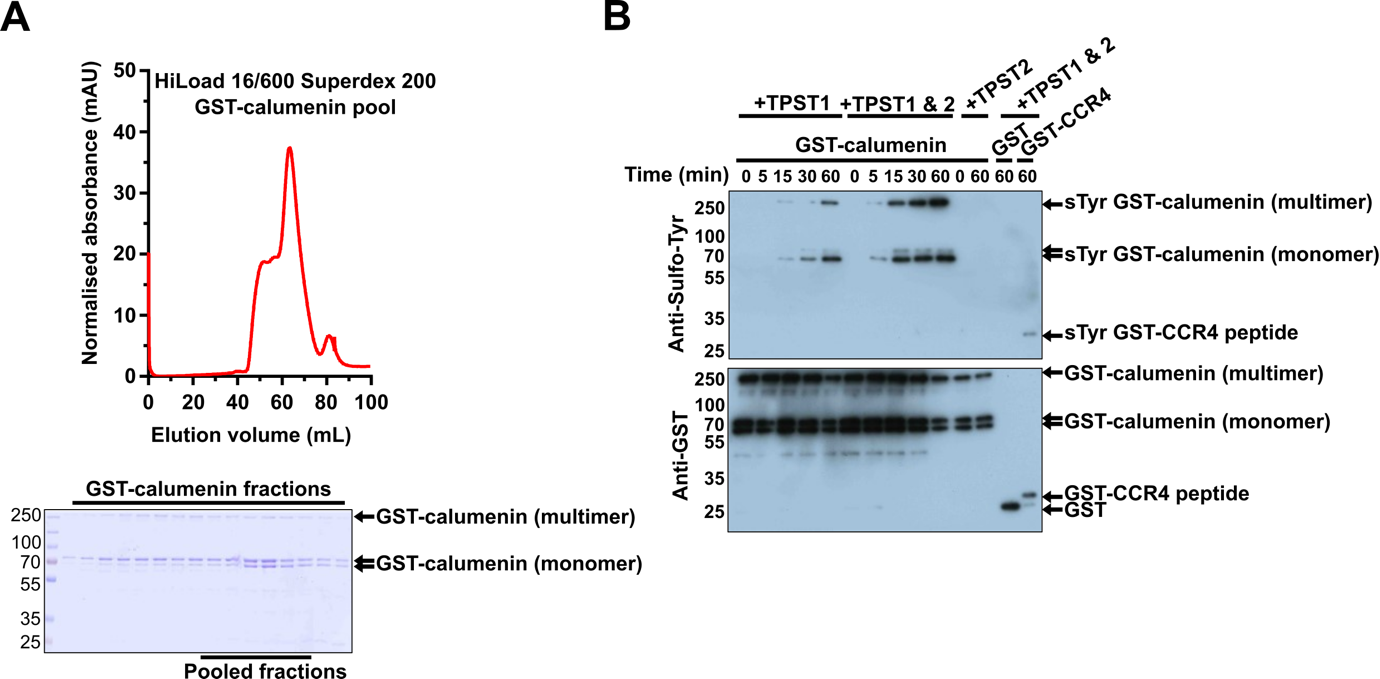

**Supplementary Figure 4. Purification and TPST-catalyzed sulfation of recombinant human Calumenin.**

(A) Purification of GST-calumenin showing UV protein elution profile from Superdex S200 (top) and calumenin analysis after SDS-PAGE and Coomassie blue staining (bottom). The pooled fractions employed for sulfation analysis are indicated, and are composed of both monomeric and multimeric calumenin species. Note that calumenin migrates anomalously by SDS-PAGE, likely due to a very high content of acidic residues (B) Time-course of GST-calumenin sulfation in the presence of the sulfate donor PAPS in the presence of either TPST1, TPST1/2 or TPST2. At the indicated time point, the reaction was terminated and samples were immunoblotted with a sTyr-specific antibody (top panel) or a GST antibody (bottom panel). The migration of each species is indicated on the right. No sulfation of GST was detected by TPST1/2 and a GST-CCR4 peptide served as a positive control.

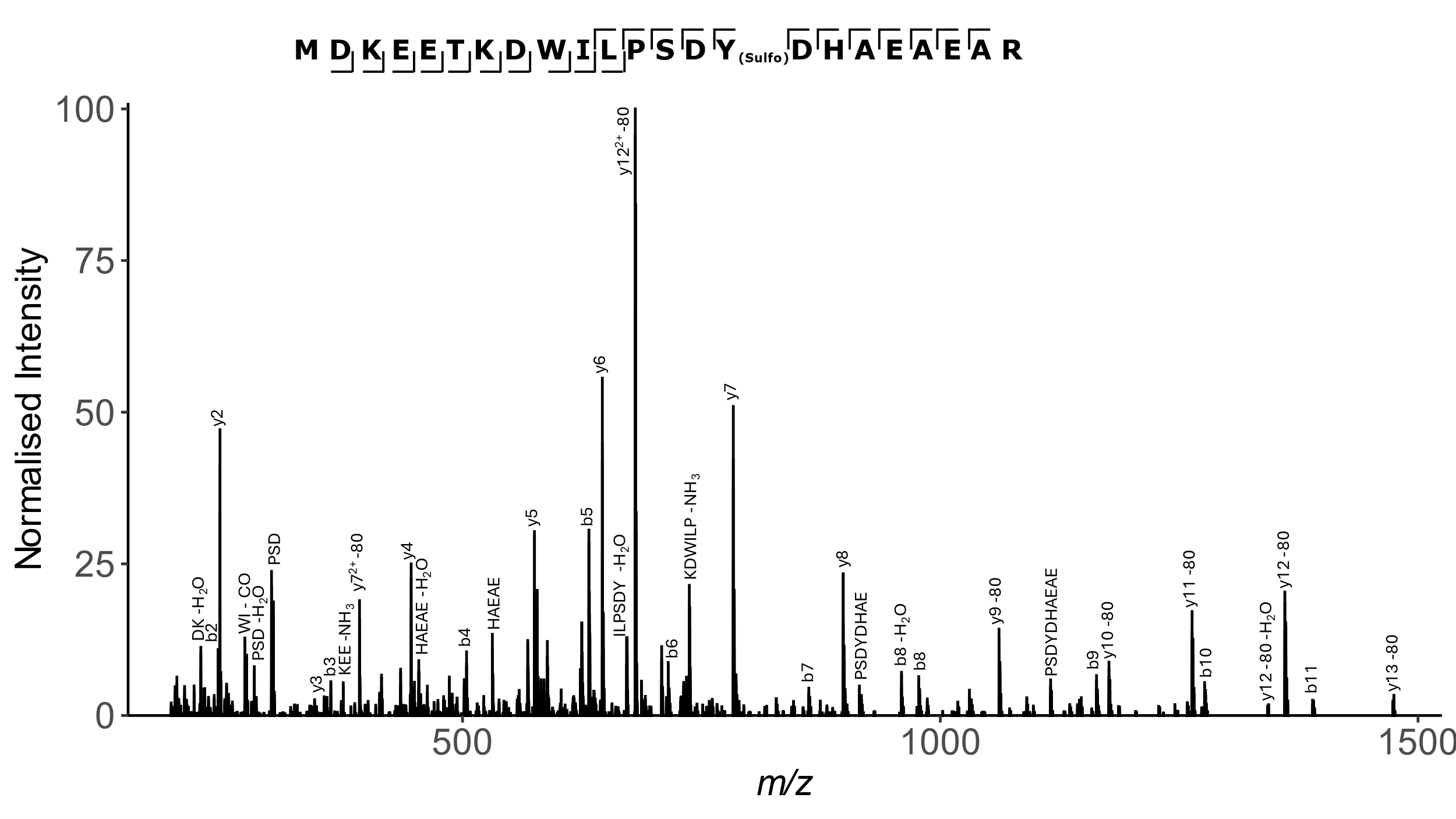

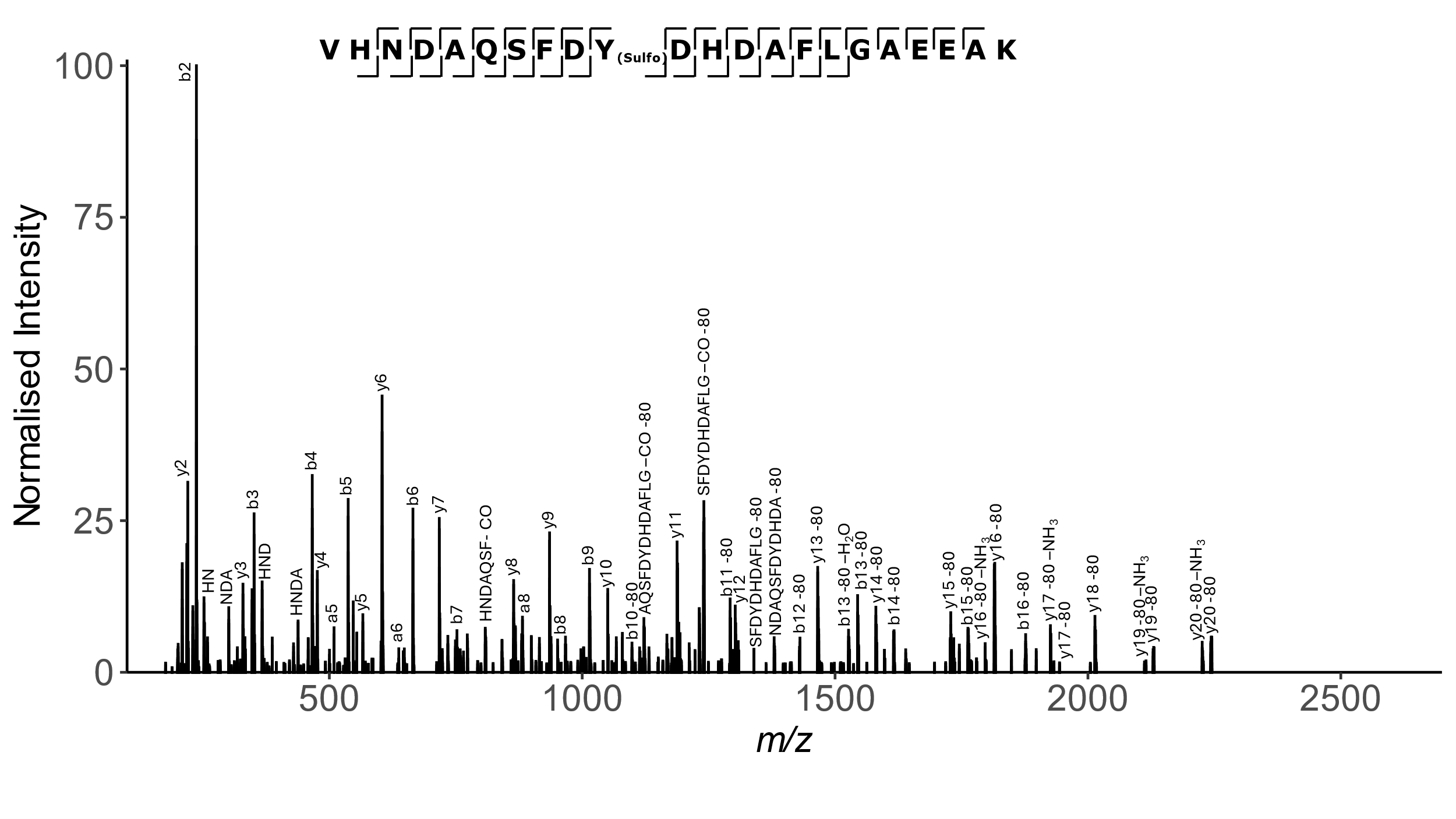
 **Supplementary Figure 5. Analysis of calumenin by Liquid chromatography mass spectrometry (LC-MS/MS).**

Annotated tandem mass spectra of sulfation-containing tryptic peptide ions obtained from calumenin incubated with TPST1/2 and PAPS for 4 hours. Peptide sequences are displayed with the annotated HCD product ions labelled, a/b ions, y ions, and internal ions, including charge state and -80 neutral losses. (Top) MS/MS spectrum of the doubly charged peptide ion of *m/z* 1280.0249, fragmented using HCD, encompassing TPST1/2-sulfated site Y47. (Bottom) MS/MS spectrum of the quadruply charged peptide ion of *m/z* 708.0563, fragmented using HCD, encompassing TPST1/2 sulfated site Y263.

| **Calumenin site of sulfation** | **Tryptic calumenin peptide sequence** | **Covalent Modification** |
| --- | --- | --- |
| Y47 | VHNDAQSFDs**Y^47^**DHDAFLGAEEAK | 1 X Sulfo |
| Y263 | MDKEETKDWILPSDs**Y^263^**DHAEAEAR | 1 X Sulfo |
| Y134 or Y136 or Y149 | NAT**Y^134^**G**Y^136^**VLDDPDPDDGFN**Y^149^**K | 1 X Sulfo |
| Y129 or Y134 or Y136 or Y149 | GHDLNEDGLVSWEE**Y^129^**KNAT**Y^134^**G**Y^136^**VLDDPDPDDGFN**Y^149^**K | 1 X Sulfo |

**Supplementary Table 1**

List of tyrosine-sulfated calumenin peptides identified by LC-MS/MS. sTyr 47 and sTyr263 were identified unequivocally and lie in a classical acidic DYD sulfation-motif. Four other potential sites of modification were also identified within longer tryptic peptides, including a 33 mer containing a site of trypsin mis-cleavage. Although none of these Tyr lie in an acidic -1 or +1 context, they are associated with acidic stretches of amino acids N or C-terminal to the putative sulfation site(s).

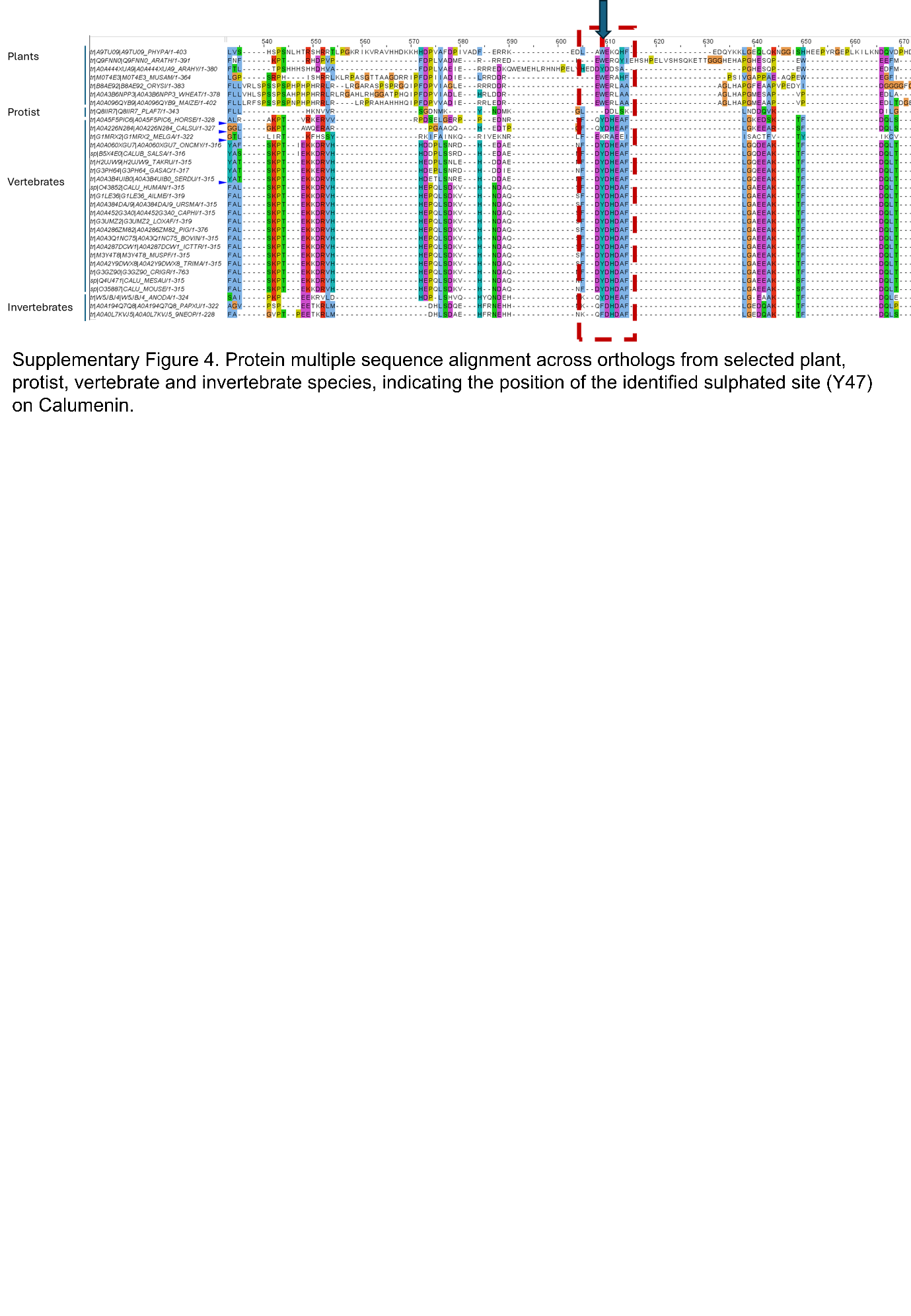

**Supplementary Figure 6**

Protein multiple sequence alignment across orthologs from selected plant, protist, vertebrate and invertebrate species, indicating the position of the identified sulphated site (Y47) on Calumenin.

**Exploration of potential factors that may impact sY detection in MS-based proteomics**

Expanding on our previously published sY LC-MS workflow, which used an Orbitrap Fusion Lumos Tribrid mass spectrometer to interrogate the secretome from immortalised human embryonic kidney (HEK-293) cells, we were interested in whether different datasets or instruments were better at detecting sY.

First, we looked at the number of PSMs contributed to the peptidoforms across BOIs by dataset. The proportion of PSMs contributed by each dataset were similar for all PSMs compared to PSMs shifted in the direction of sulfation with calibrated mass error < -0.006 Da (Supplementary Figure 7). The ProteomeXchange dataset PXD015901 had a large contribution to the PSMs that contributed to all BOI peptidoforms, but a smaller contribution to the convincing peptidoforms. PXD005336 also had a large contribution to the BOI, which remained large in the convincing peptidoforms, but not the known sY peptidoforms. The majority of datasets with large contributions to convincing sY peptidoforms were from the National Cancer Institute's Clinical Proteomic Tumor Analysis Consortium (CPTAC). For instance, across the convincing and known sY peptidoforms, CPTAC-S049 was the largest PSM contributor dataset. This was an interesting observation, since the CPTAC datasets are derived from various cancer studies. Upon further investigation, all datasets with PSM contributions to the convincing sY sites were directly linked to cancer, except PXD005336 which had an indirect link. It investigated the target landscape of clinical kinase inhibitors, which are often altered in cancer cells and have been used as promising cancer therapeutic targets. While these findings appear promising, it is important to recognise that the observed dataset trends are biased due to the overwhelming contribution of cancer datasets to the Human Phosphobuild. Of the total 44.8 million phosphopeptide-spectrum matches (pPSMs) included in the analysis, 92% originate from samples categorised as cancer cell lines or tissues. Therefore, the potential over-representation of cancer-related phosphosites/sulfosites should be taken into account when interpreting these results. Nevertheless, there may be a link between tyrosine sulfation and cancer, as has been recently discussed in a study of pancreatic cancer, where increased expression of SLC35B2 and TPST2 are associated with poor prognosis (Cai et al. 2023). Recent *in vitro* and *in vivo* studies have also shown that inhibiting tyrosine sulfation within this signaling axis can reduce the growth and metastasis of pancreatic cancer (Zhang and Pasca di Magliano 2023).

We also explored the trends in instrumentation (in the form of number of PSM contributions) across the entire dataset, across the BOIs, across peptidoforms with confident sY assignment, and across peptidoforms with known sY sites (Supplementary Figure 8). As expected, Orbitrap based instruments, specifically the Orbitrap Fusion Lumos (launched in 2015) and the Q Exactive (launched in 2011) were the highest contributors across the phosphobuild, each generation close to a third of all PSMs in our analysis. The pattern of instrument contribution across all peptidoforms assigned to the BOIs (including these with undetermined and false positive histogram types) was largely dominated by PSMs derived from the Q Exactive (close to 70% of all PSMs). In contrast, the Q Exactive had a negligible (~5%) contribution to the overall PSMs in the analysis linked to peptidoforms with known sY sites. Across these peptidoforms, the Orbitrap Fusion Lumos and Q Exactive HF were top contributors. The contribution pattern across the convincing peptidoforms was similar to the overall pattern across the dataset. Overall, this suggests that when operated at sufficient resolution, modern instruments should enable distinguishing sY from pY based on accurate precursor mass shifts.

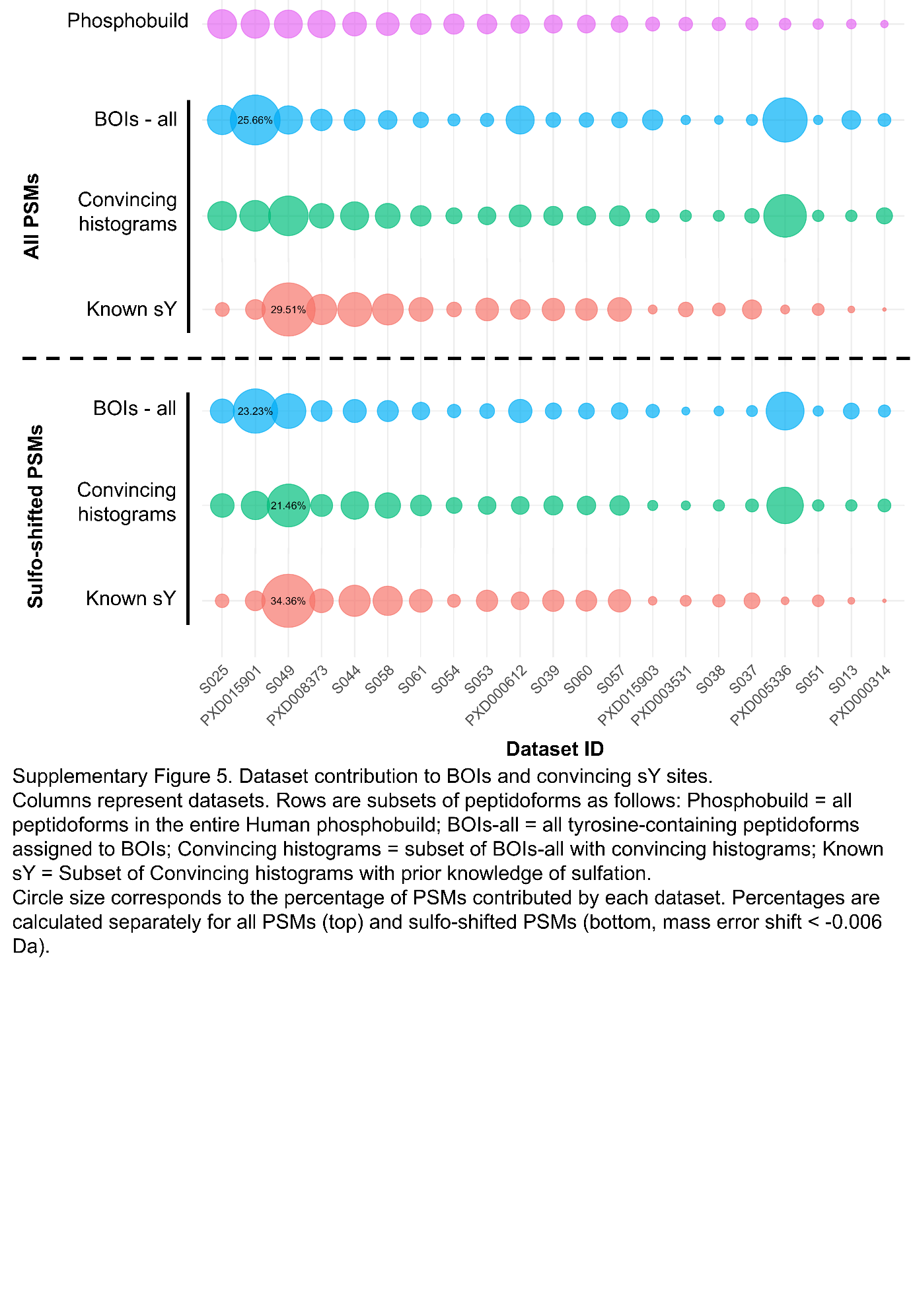

**Supplementary Figure 7.** Dataset contribution to BOIs and convincing sY sites.

Columns represent datasets. Rows are subsets of peptidoforms as follows: Phosphobuild = all peptidoforms in the entire Human phosphobuild; BOIs-all = all tyrosine-containing peptidoforms assigned to BOIs; Convincing histograms = subset of BOIs-all with convincing histograms; Known sY = Subset of Convincing histograms with prior knowledge of sulfation.

Circle size corresponds to the percentage of PSMs contributed by each dataset. Percentages are calculated separately for all PSMs (top) and sulfo-shifted PSMs (bottom, mass error shift < -0.006 Da).

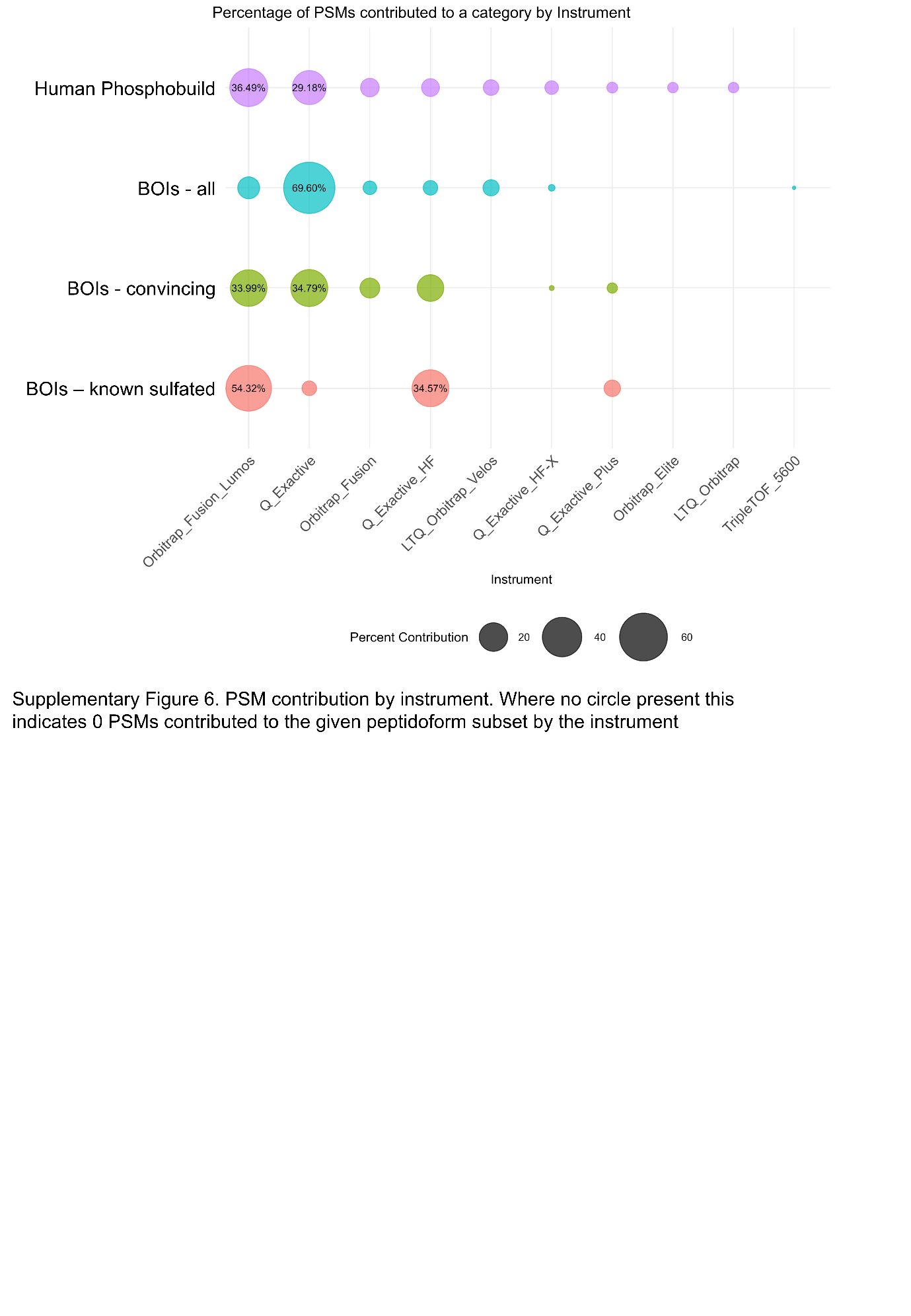

**Supplementary Figure 8.** PSM contribution by instrument. Where no circle present this indicates 0 PSMs contributed to the given peptidoform subset by the instrument

**Supplementary Information References**

Cai, Xinran, Sihan Li, Xuemei Zeng, Meishu Xu, Zehua Wang, Aatur D. Singhi, Daolin Tang, et al. 2023. “Inhibition of the SLC35B2-TPST2 Axis of Tyrosine Sulfation Attenuates the Growth and Metastasis of Pancreatic Ductal Adenocarcinom.” *Cellular and Molecular Gastroenterology and Hepatology*. Elsevier BV.

Zhang, Yaqing, and Marina Pasca di Magliano. 2023. “Tyrosine Sulfation: A New Player and Potential Target in Pancreatic Cancer.” *Cellular and Molecular Gastroenterology and Hepatology* 16 (3): 501–2.
