## Supplementary File 1 for "Searching for Sulfotyrosines (sY) in a HA(pY)STACK": README.txt

Contingency tables used in balloon plots for Supplementary Figures 5 (by dataset) and 6 (by instrument).
The by dataset counts are also split into sulfo-supporting vs all counts.

The first part of the file name indicates PSM counts are for:
BOI = peptidofomrs in BOIs
convincing = peptidoforms with convincing histograms
known_sY = previously annotated as sulfation 
PSMs = counts in the entire build
