## Supplementary File 2 for "Searching for Sulfotyrosines (sY) in a HA(pY)STACK": faceted_plots_by_dataset_ID.pdf

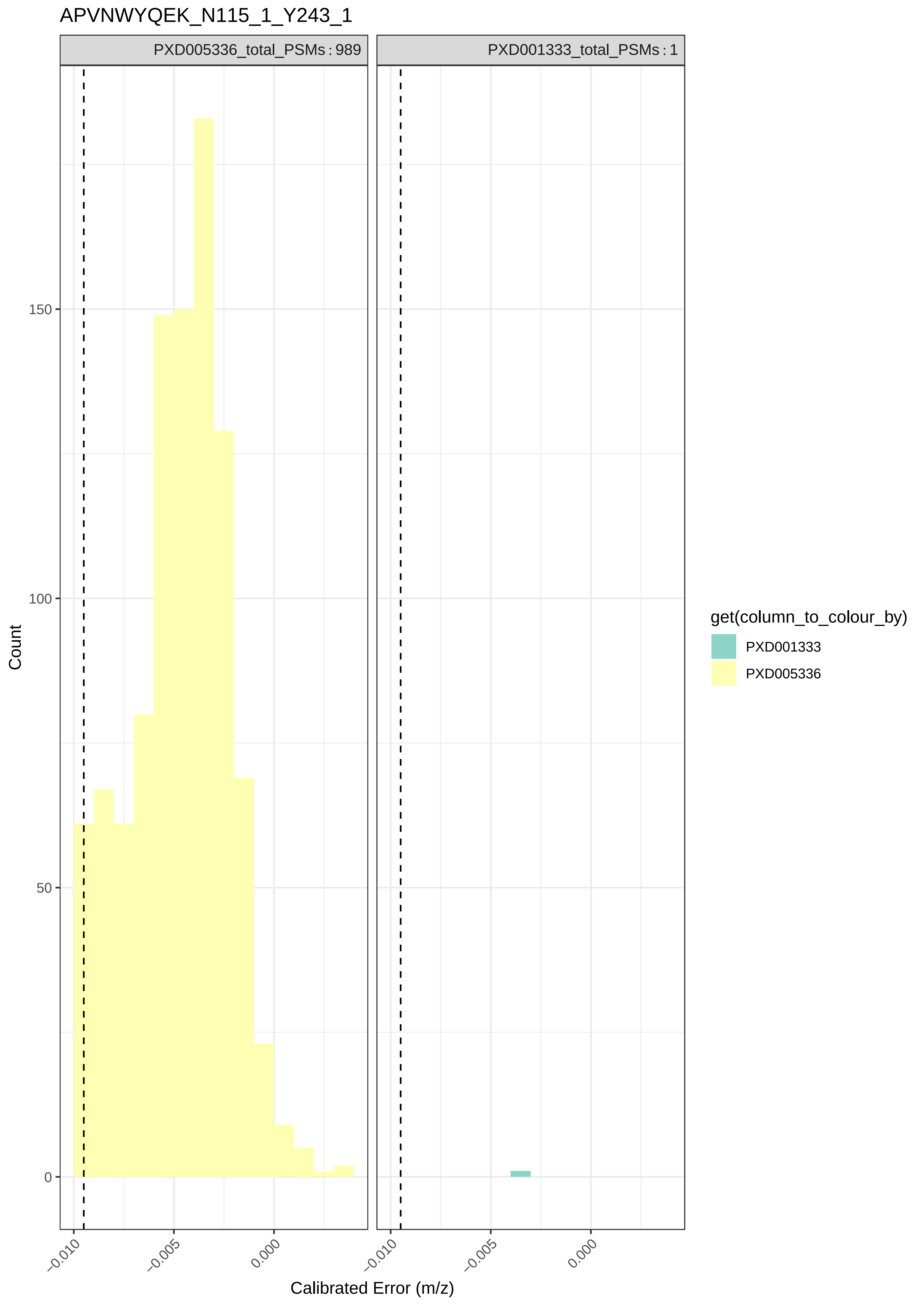

APVNWYQEK_N115_1_Y243_1
PXD005336_total_PSMs : 989
PXD001333_total_PSMs : 1
Count
150
100
50
0
−0.010
−0.005
0.000
Calibrated Error (m/z)
get(column_to_colour_by)
PXD001333
PXD005336

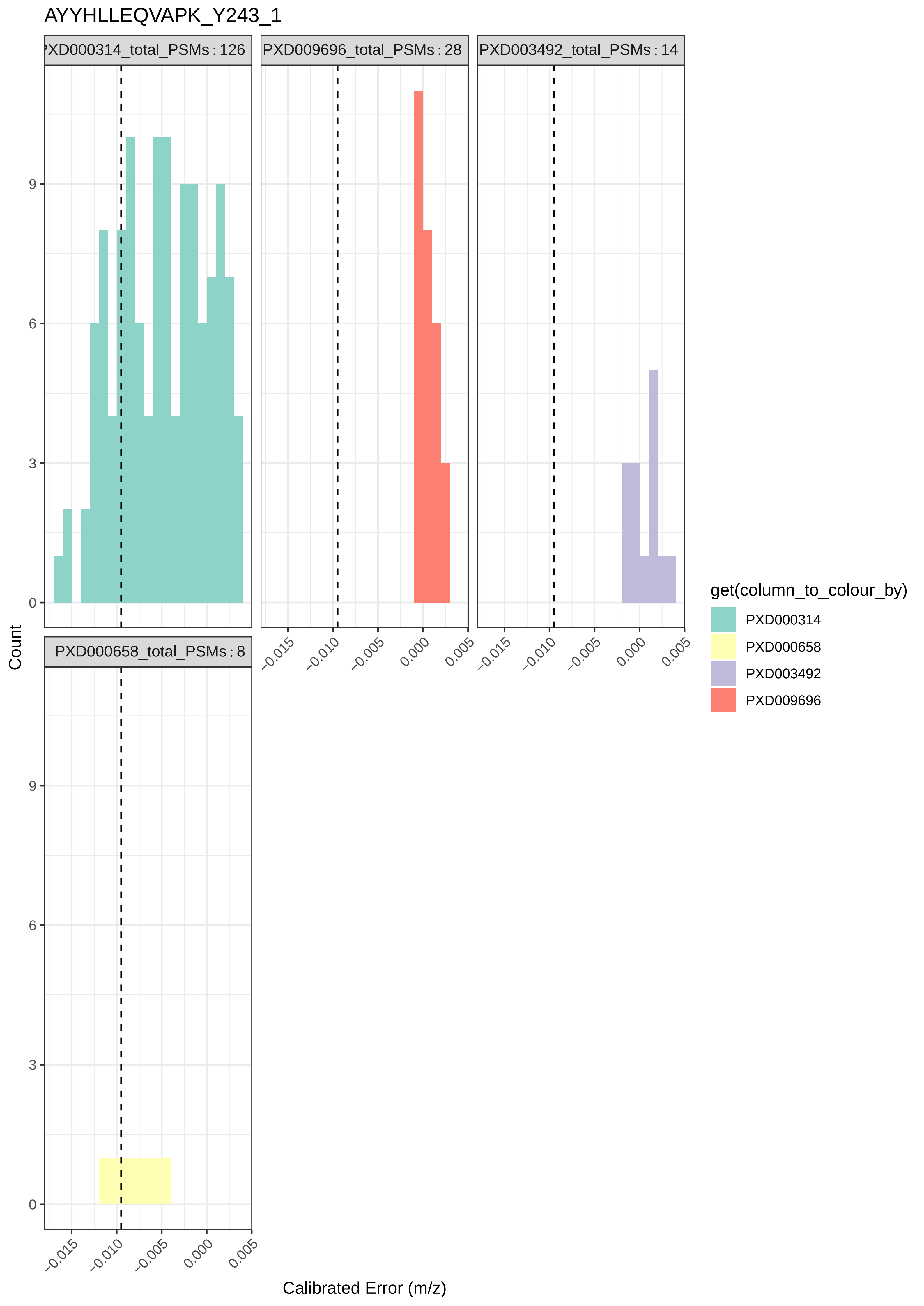

AYYHLLEQVAPK_Y243_1
PXD000314_total_PSMs : 126
PXD009696_total_PSMs : 28
PXD003492_total_PSMs : 14
PXD000658_total_PSMs : 8
Count
Calibrated Error (m/z)
get(column_to_colour_by)
PXD000314
PXD000658
PXD003492
PXD009696

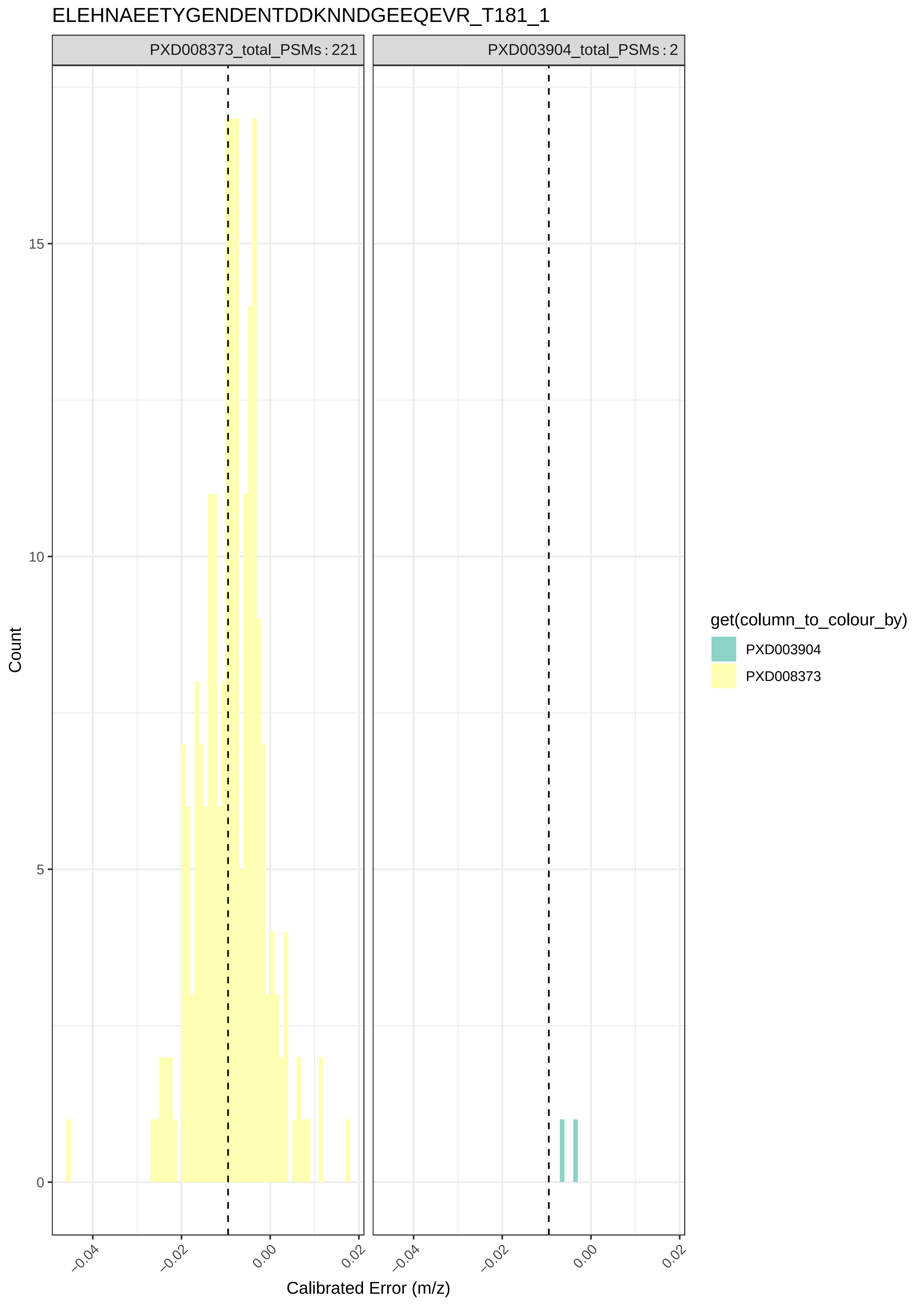
ELEHNAEETYGENDENTDDKNNDGEEQEVR_T181_1
PXD008373_total_PSMs : 221
PXD003904_total_PSMs : 2
Count
15
10
5
0
−0.04
−0.02
0.00
0.02
Calibrated Error (m/z)
get(column_to_colour_by)
PXD003904
PXD008373

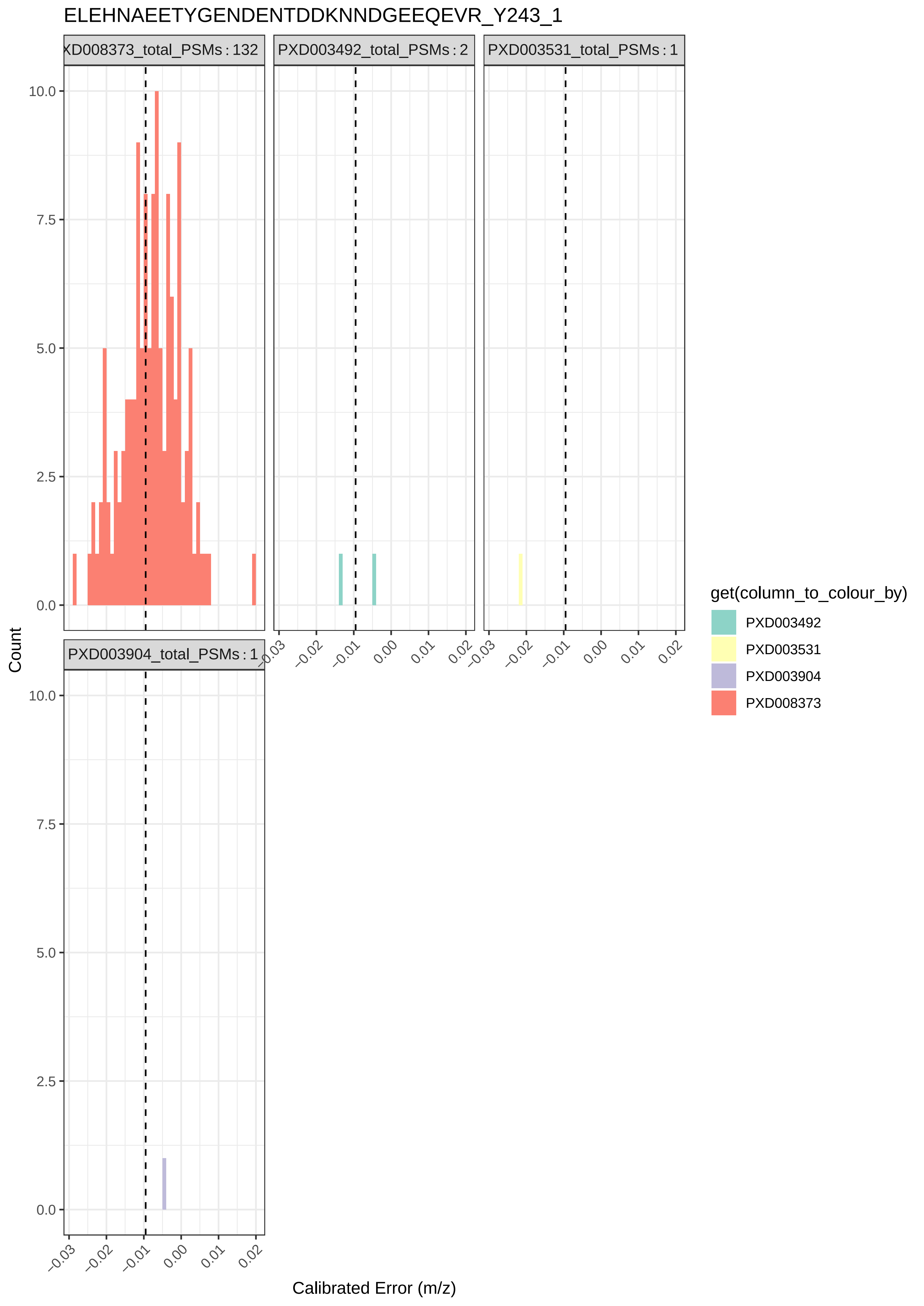

ELEHNAEETYGENDENTDDKNNDGEEQEVR_Y243_1
XD008373_total_PSMs : 132
PXD003492_total_PSMs : 2
PXD003531_total_PSMs : 1
PXD003904_total_PSMs : 1
Count
Calibrated Error (m/z)
0.0
2.5
5.0
7.5
10.0
−0.03
−0.02
−0.01
0.00
0.01
0.02
get(column_to_colour_by)
PXD003492
PXD003531
PXD003904
PXD008373

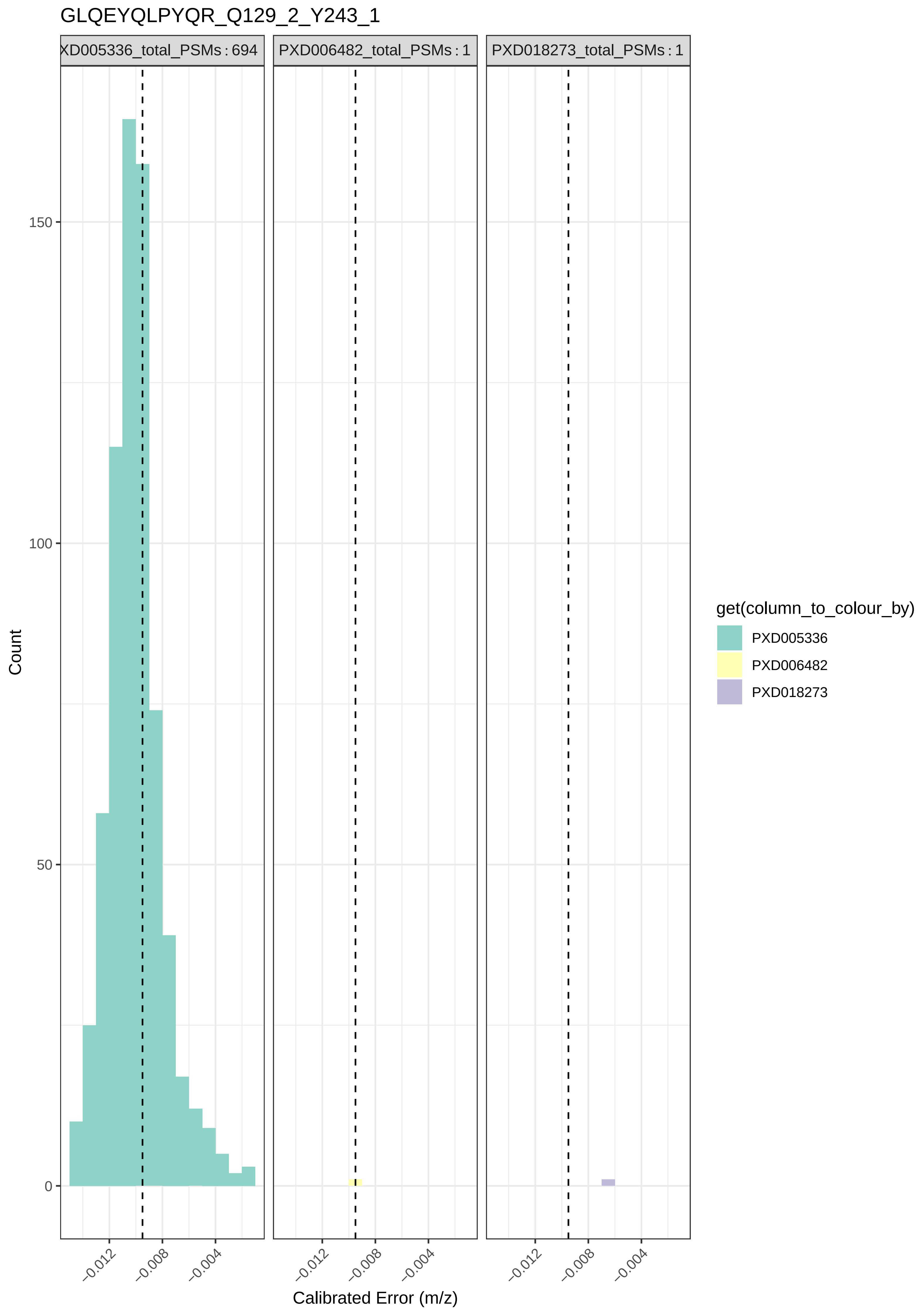

GLQEYQLPYQR_Q129_2_Y243_1
XD005336_total_PSMs : 694
PXD006482_total_PSMs : 1
PXD018273_total_PSMs : 1
Count
150
100
50
0
−0.012
−0.008
−0.004
Calibrated Error (m/z)
get(column_to_colour_by)
PXD005336
PXD006482
PXD018273

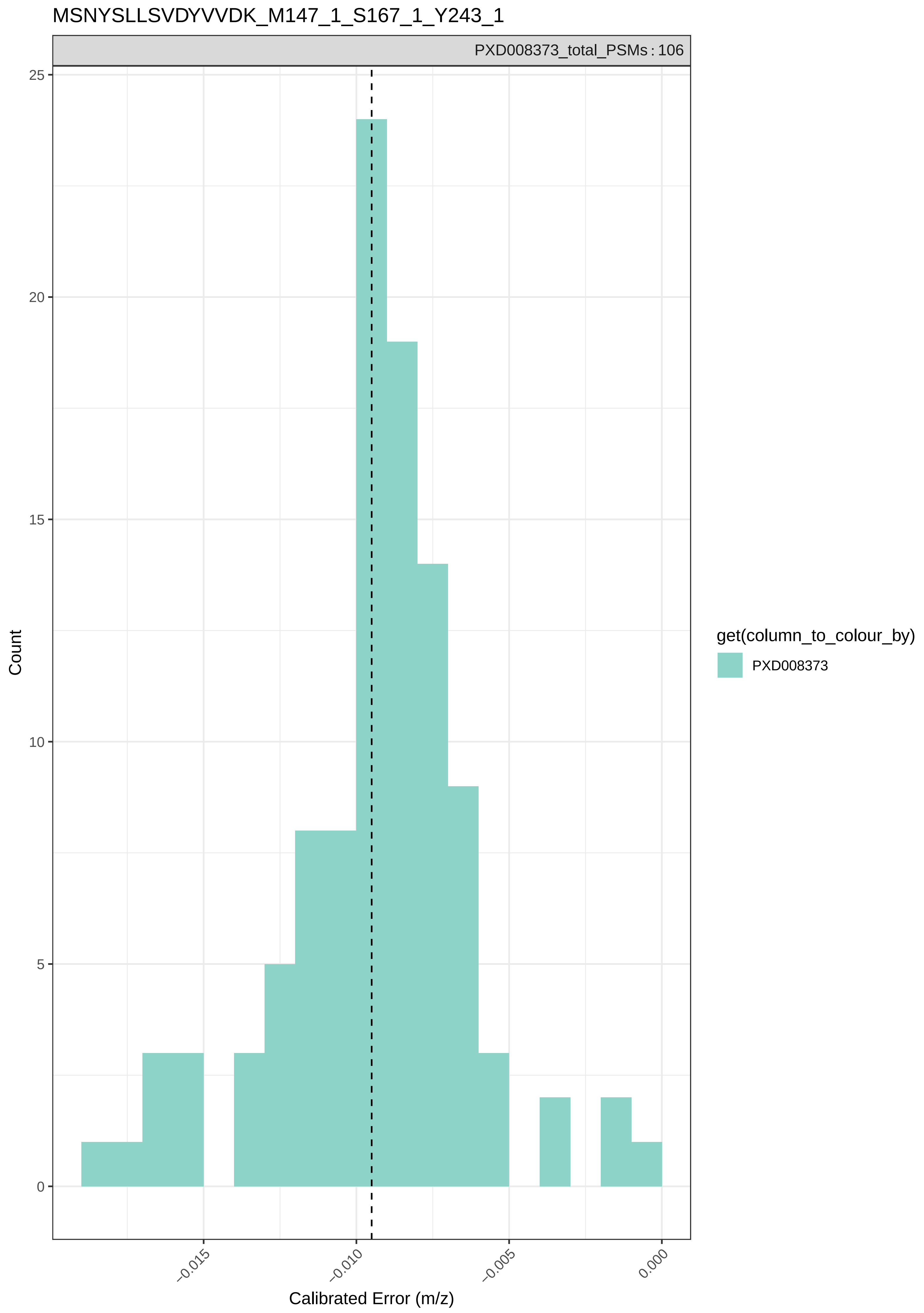

MSNYSLLSVDYVVDK_M147_1_S167_1_Y243_1
PXD008373_total_PSMs : 106
25
20
15
10
5
0
Count
−0.015
−0.010
−0.005
0.000
Calibrated Error (m/z)
get(column_to_colour_by)
PXD008373

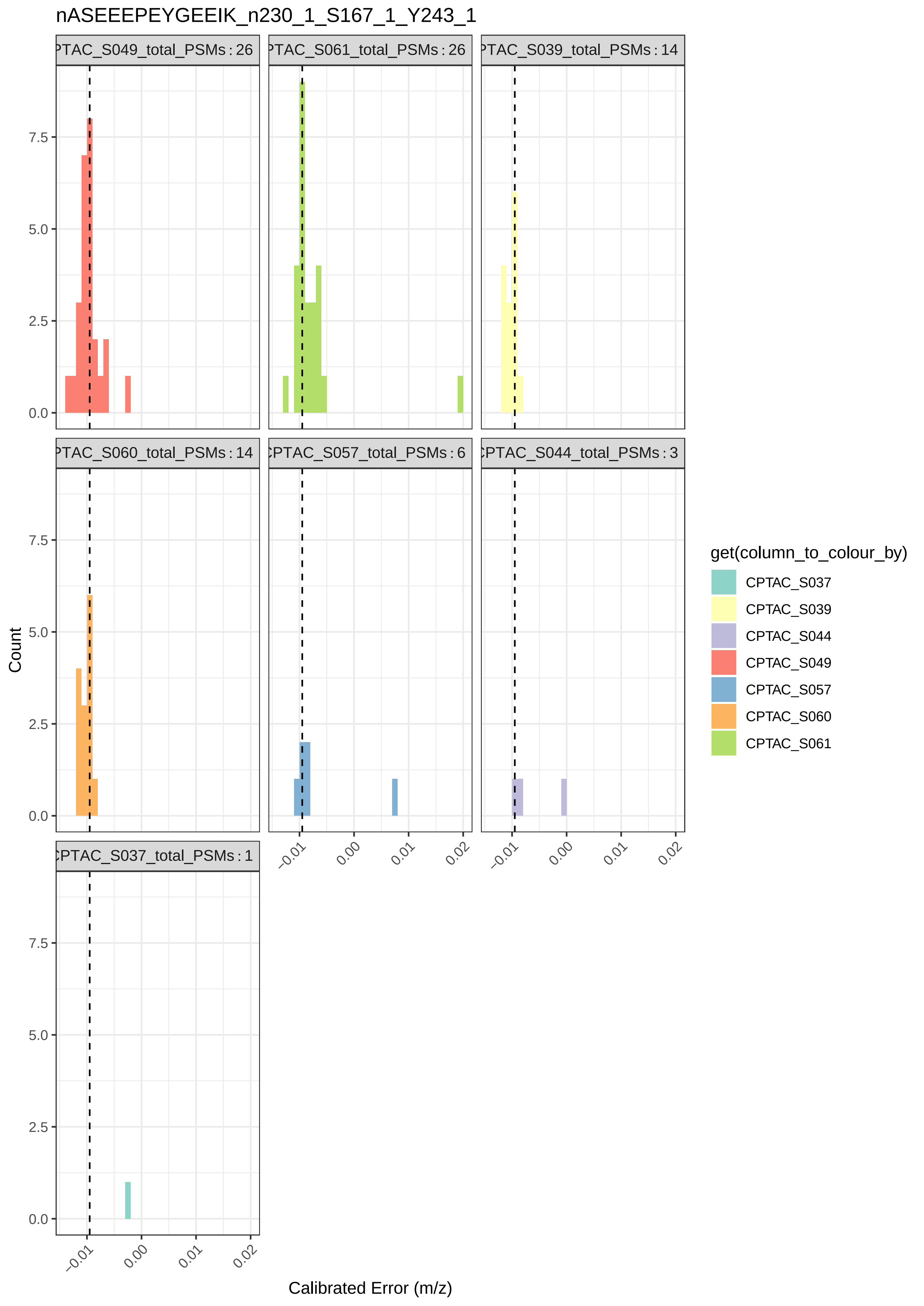

nASEEEPEYGEEIK_n230_1_S167_1_Y243_1
PTAC_S049_total_PSMs : 26
PTAC_S061_total_PSMs : 26
PTAC_S039_total_PSMs : 14
PTAC_S060_total_PSMs : 14
CPTAC_S057_total_PSMs : 6
CPTAC_S044_total_PSMs : 3
CPTAC_S037_total_PSMs : 1
Count
0.0
2.5
5.0
7.5
−0.01
0.00
0.01
0.02
Calibrated Error (m/z)
get(column_to_colour_by)
CPTAC_S037
CPTAC_S039
CPTAC_S044
CPTAC_S049
CPTAC_S057
CPTAC_S060
CPTAC_S061

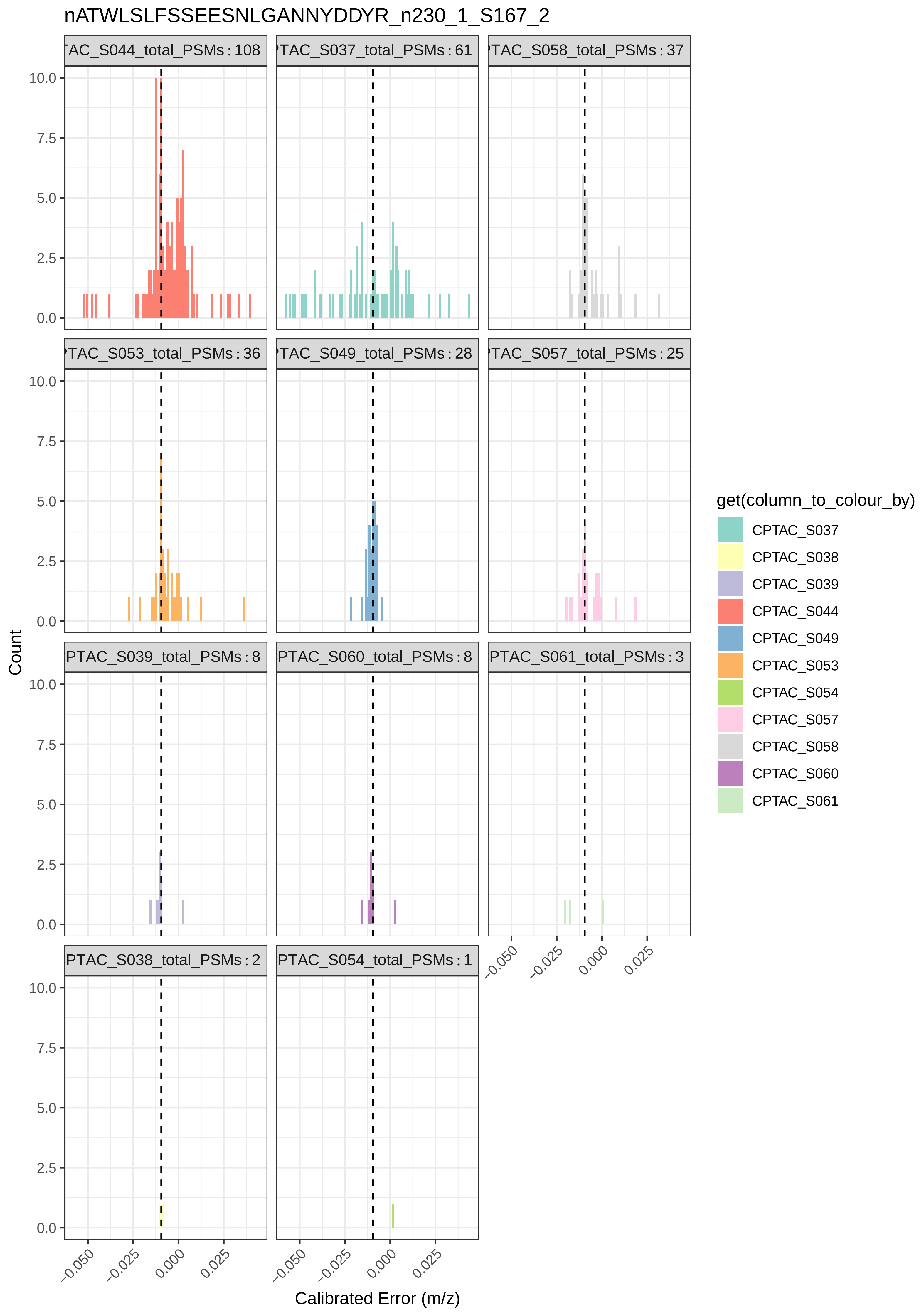

nATWLSLFSSEESNLGANNYDDYR_n230_1_S167_2
TAC_S044_total_PSMs : 108
TAC_S037_total_PSMs : 61
TAC_S058_total_PSMs : 37
TAC_S053_total_PSMs : 36
TAC_S049_total_PSMs : 28
TAC_S057_total_PSMs : 25
PTAC_S039_total_PSMs : 8
PTAC_S060_total_PSMs : 8
PTAC_S061_total_PSMs : 3
PTAC_S038_total_PSMs : 2
PTAC_S054_total_PSMs : 1
Count
10.0
7.5
5.0
2.5
0.0
−0.050
−0.025
0.000
0.025
Calibrated Error (m/z)
get(column_to_colour_by)
CPTAC_S037
CPTAC_S038
CPTAC_S039
CPTAC_S044
CPTAC_S049
CPTAC_S053
CPTAC_S054
CPTAC_S057
CPTAC_S058
CPTAC_S060
CPTAC_S061

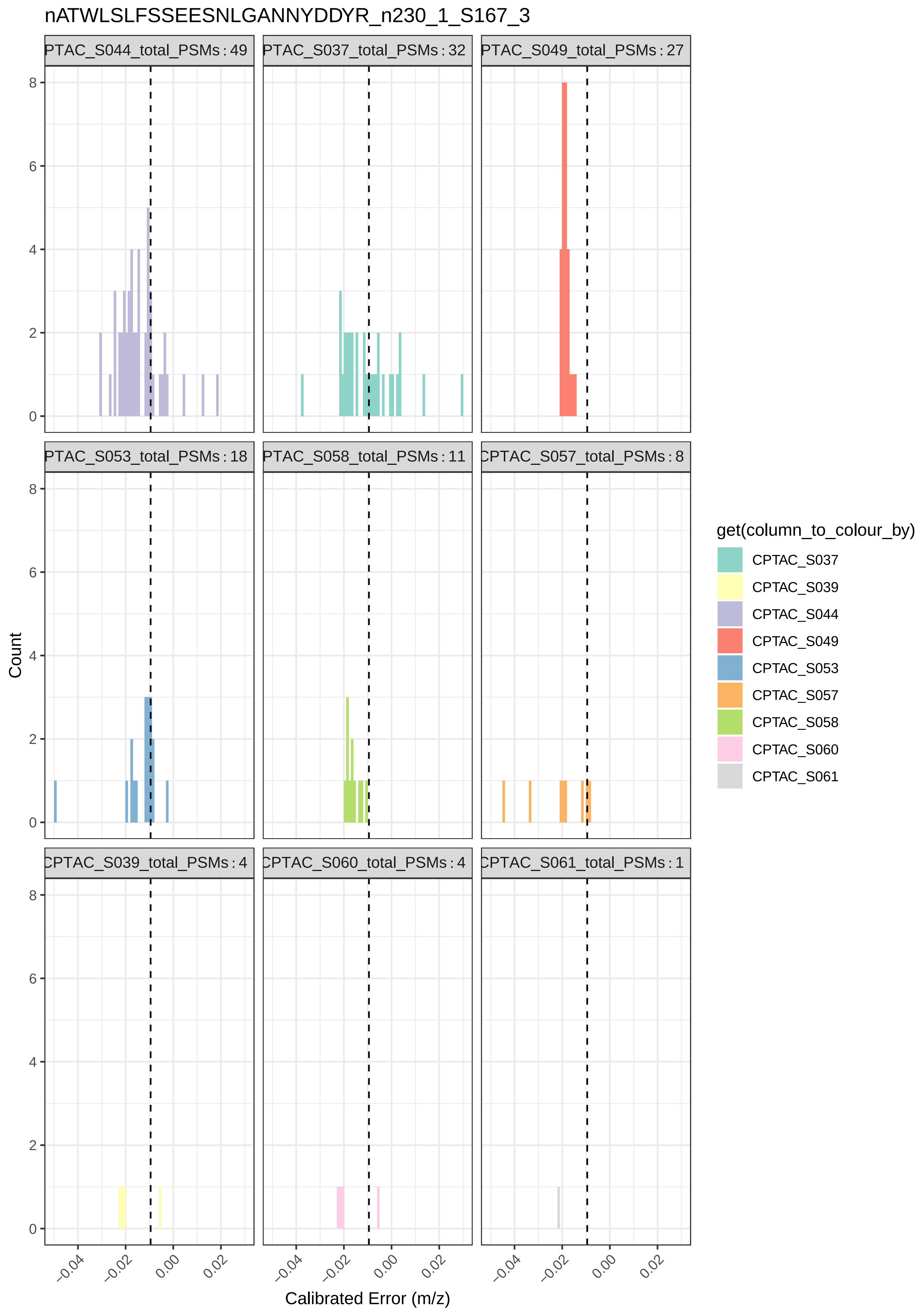

nATWLSLFSSEESNLGANNYDDYR_n230_1_S167_3
PTAC_S044_total_PSMs : 49
PTAC_S037_total_PSMs : 32
PTAC_S049_total_PSMs : 27
PTAC_S053_total_PSMs : 18
PTAC_S058_total_PSMs : 11
CPTAC_S057_total_PSMs : 8
CPTAC_S039_total_PSMs : 4
CPTAC_S060_total_PSMs : 4
CPTAC_S061_total_PSMs : 1
Count
0
2
4
6
8
−0.04
−0.02
0.00
0.02
Calibrated Error (m/z)
get(column_to_colour_by)
CPTAC_S037
CPTAC_S039
CPTAC_S044
CPTAC_S049
CPTAC_S053
CPTAC_S057
CPTAC_S058
CPTAC_S060
CPTAC_S061

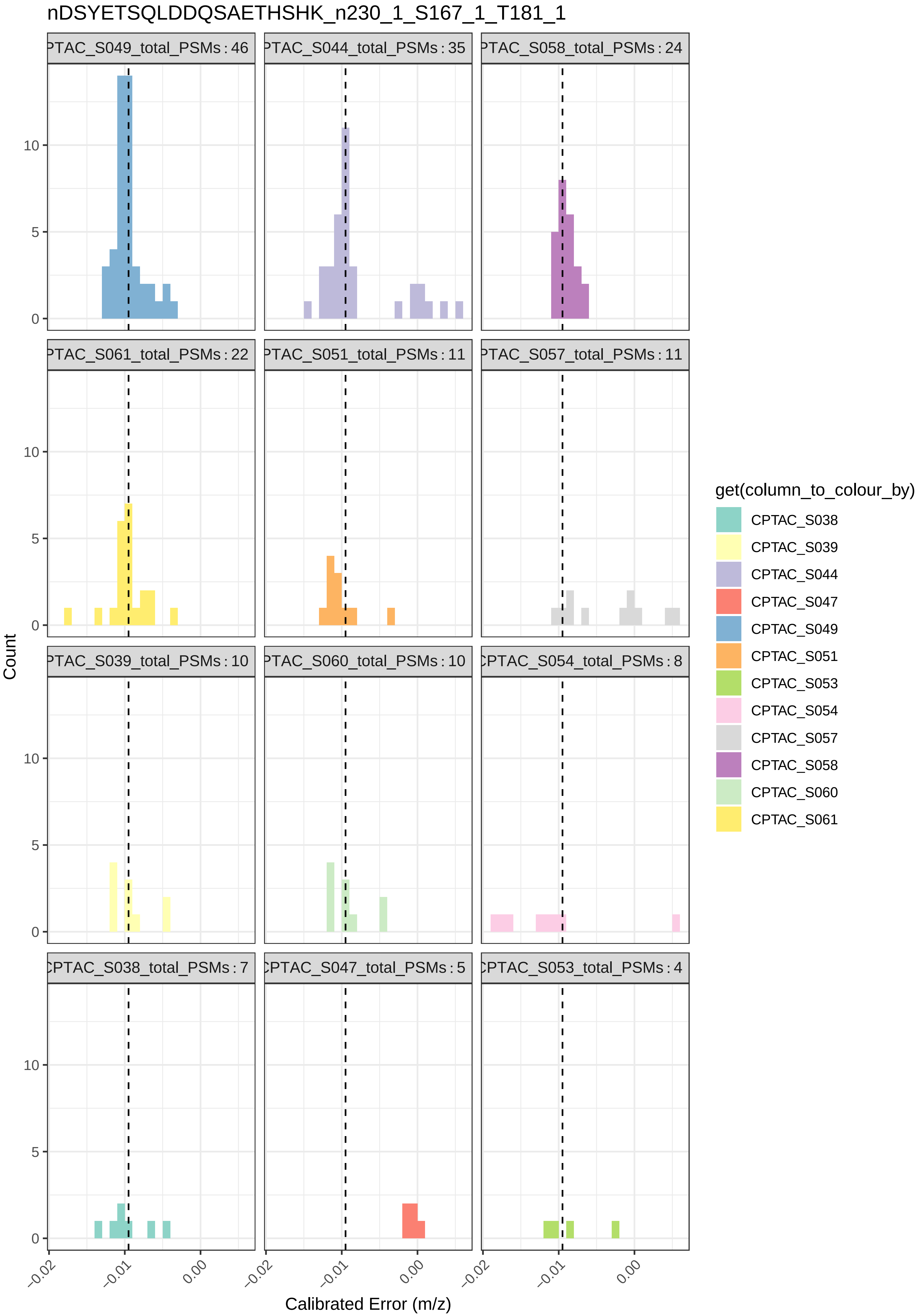
nDSYETSQLDDQSAETHSHK_n230_1_S167_1_T181_1
PTAC_S049_total_PSMs : 46
PTAC_S044_total_PSMs : 35
PTAC_S058_total_PSMs : 24
PTAC_S061_total_PSMs : 22
PTAC_S051_total_PSMs : 11
PTAC_S057_total_PSMs : 11
PTAC_S039_total_PSMs : 10
PTAC_S060_total_PSMs : 10
CPTAC_S054_total_PSMs : 8
CPTAC_S038_total_PSMs : 7
CPTAC_S047_total_PSMs : 5
CPTAC_S053_total_PSMs : 4
Count
0
5
10
−0.02
−0.01
0.00
Calibrated Error (m/z)
get(column_to_colour_by)
CPTAC_S038
CPTAC_S039
CPTAC_S044
CPTAC_S047
CPTAC_S049
CPTAC_S051
CPTAC_S053
CPTAC_S054
CPTAC_S057
CPTAC_S058
CPTAC_S060
CPTAC_S061

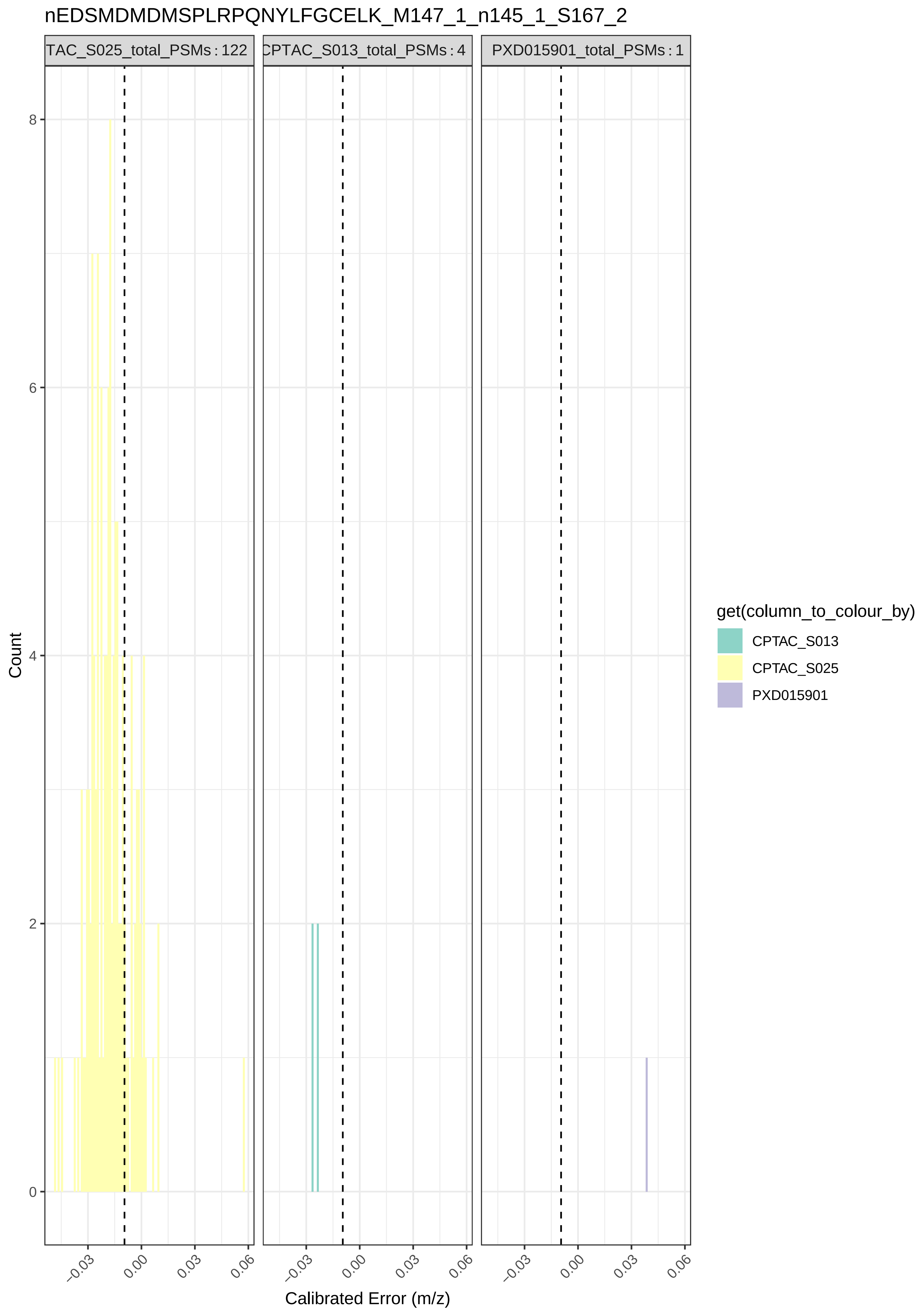

nEDSMDMDMSPLRPQNYLFGCELK_M147_1_n145_1_S167_2
TAC_S025_total_PSMs : 122
CPTAC_S013_total_PSMs : 4
PXD015901_total_PSMs : 1
Count
0
2
4
6
8
−0.03
0.00
0.03
0.06
Calibrated Error (m/z)
get(column_to_colour_by)
CPTAC_S013
CPTAC_S025
PXD015901

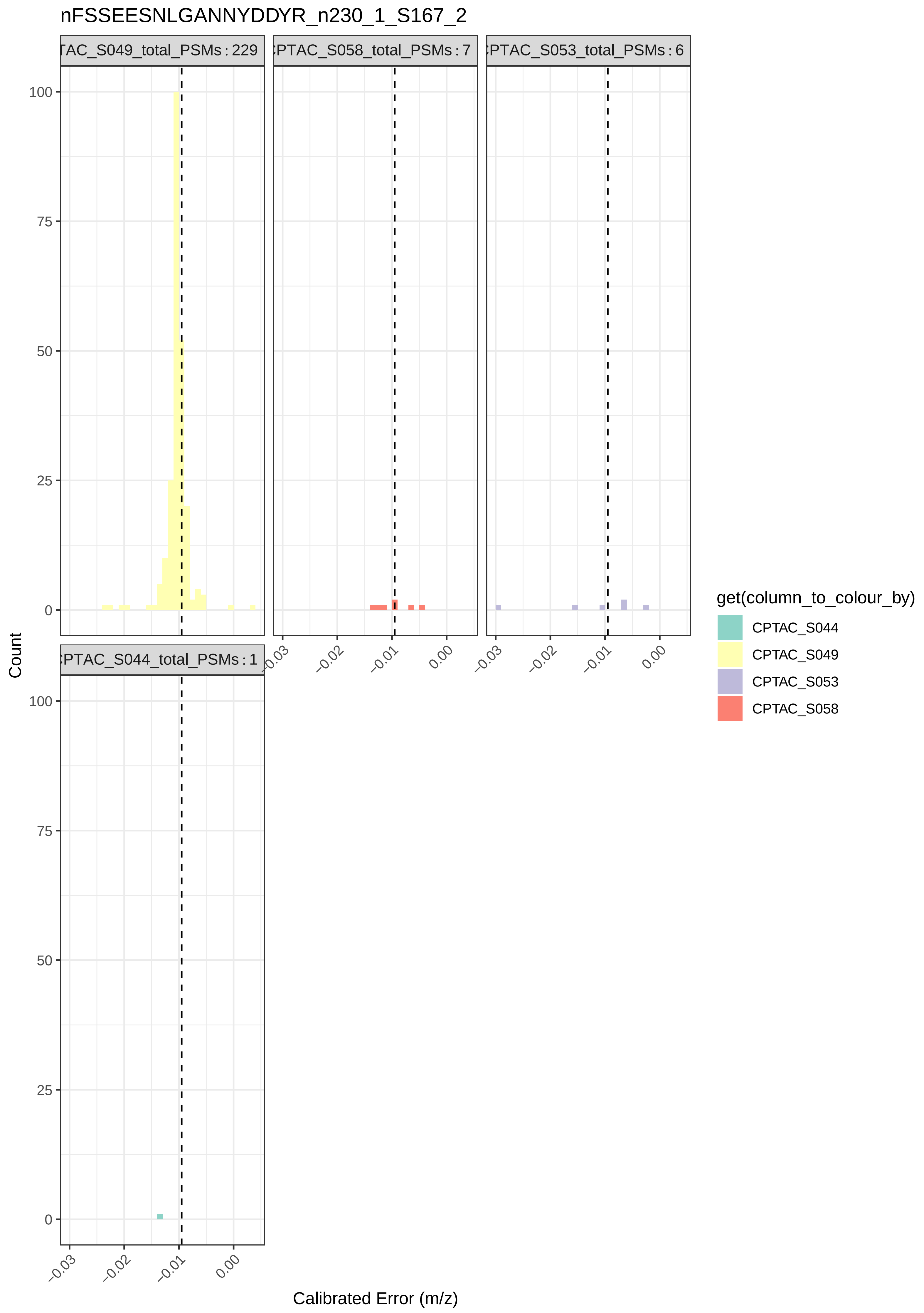
nFSSEESNLGANNYDDYR_n230_1_S167_2
TAC_S049_total_PSMs : 229
PTAC_S058_total_PSMs : 7
PTAC_S053_total_PSMs : 6
PTAC_S044_total_PSMs : 1
100
75
50
25
0
Count
−0.03
−0.02
−0.01
0.00
Calibrated Error (m/z)
get(column_to_colour_by)
CPTAC_S044
CPTAC_S049
CPTAC_S053
CPTAC_S058

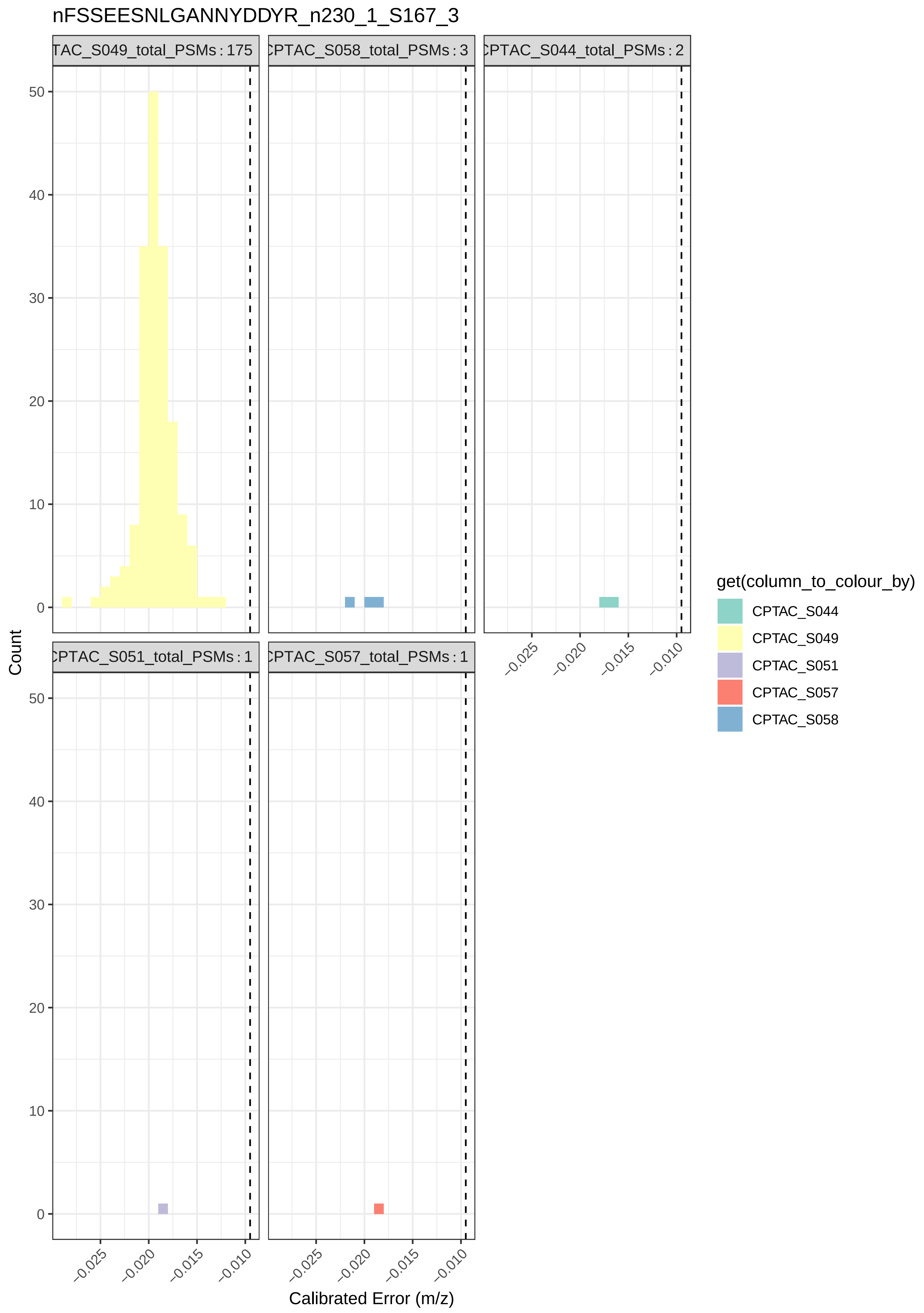

nFSSEESNLGANNYDDYR_n230_1_S167_3
TAC_S049_total_PSMs : 175
CPTAC_S058_total_PSMs : 3
CPTAC_S044_total_PSMs : 2
CPTAC_S051_total_PSMs : 1
CPTAC_S057_total_PSMs : 1
Count
0
10
20
30
40
50
−0.025
−0.020
−0.015
−0.010
Calibrated Error (m/z)
get(column_to_colour_by)
CPTAC_S044
CPTAC_S049
CPTAC_S051
CPTAC_S057
CPTAC_S058

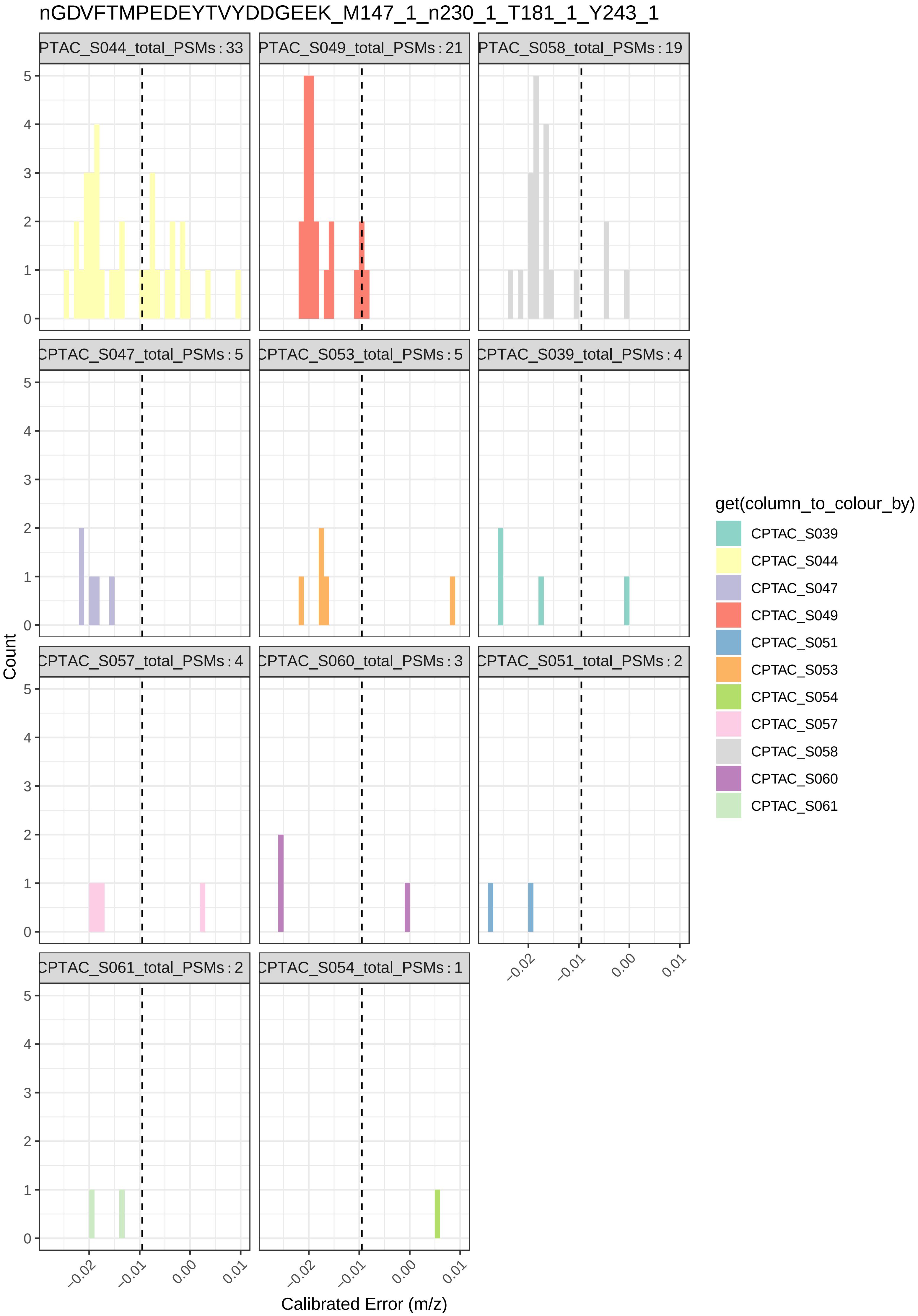

nGDVFTMPEDEYTVYDDGEEK_M147_1_n230_1_T181_1_Y243_1
PTAC_S044_total_PSMs : 33
PTAC_S049_total_PSMs : 21
PTAC_S058_total_PSMs : 19
CPTAC_S047_total_PSMs : 5
CPTAC_S053_total_PSMs : 5
CPTAC_S039_total_PSMs : 4
CPTAC_S057_total_PSMs : 4
CPTAC_S060_total_PSMs : 3
CPTAC_S051_total_PSMs : 2
CPTAC_S061_total_PSMs : 2
CPTAC_S054_total_PSMs : 1
Count
Calibrated Error (m/z)
−0.02
−0.01
0.00
0.01
0
1
2
3
4
5
get(column_to_colour_by)
CPTAC_S039
CPTAC_S044
CPTAC_S047
CPTAC_S049
CPTAC_S051
CPTAC_S053
CPTAC_S054
CPTAC_S057
CPTAC_S058
CPTAC_S060
CPTAC_S061

### nGDVFTMPEDEYTVYDDGEEK_n145_1_T181_1_Y243_1

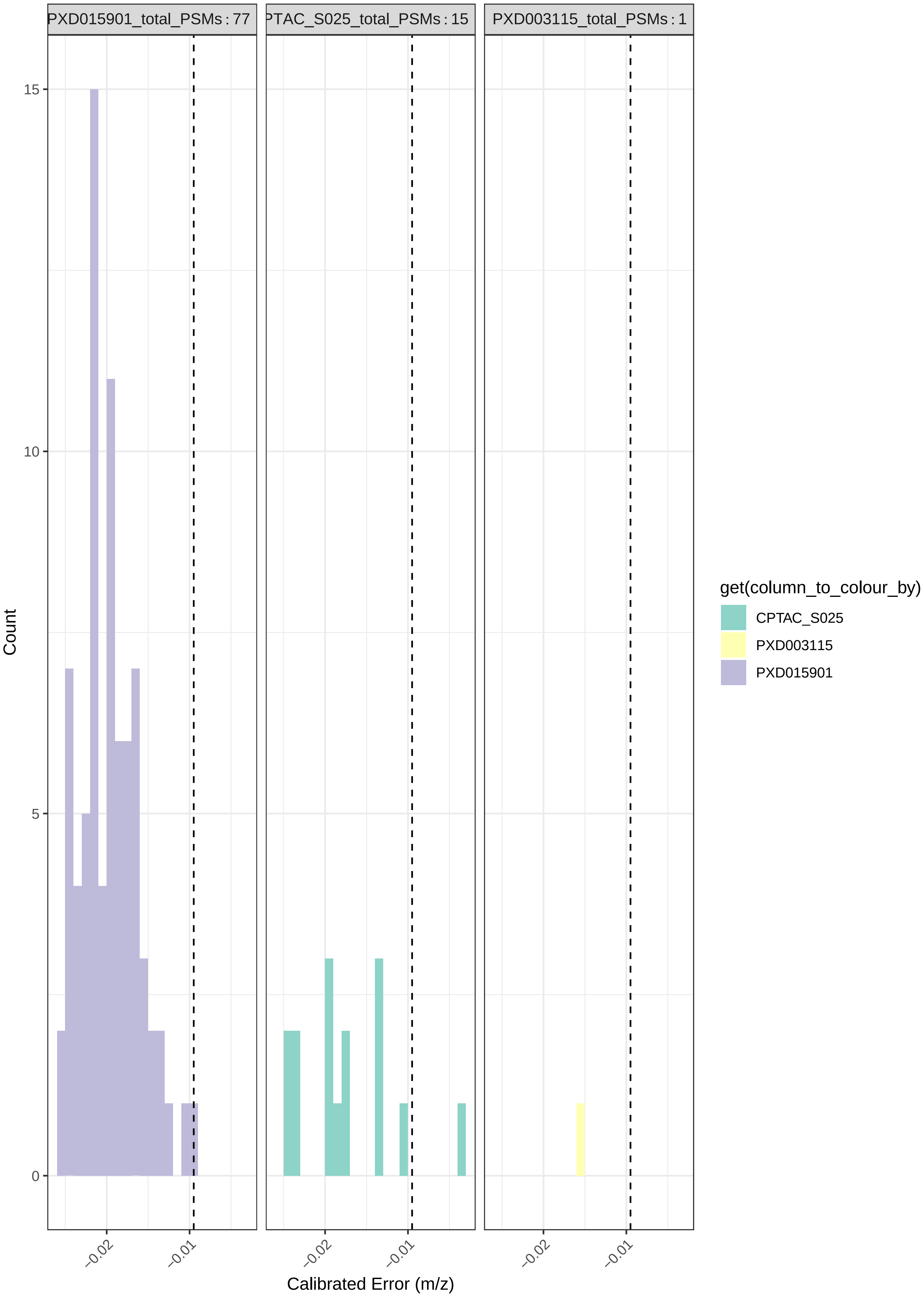

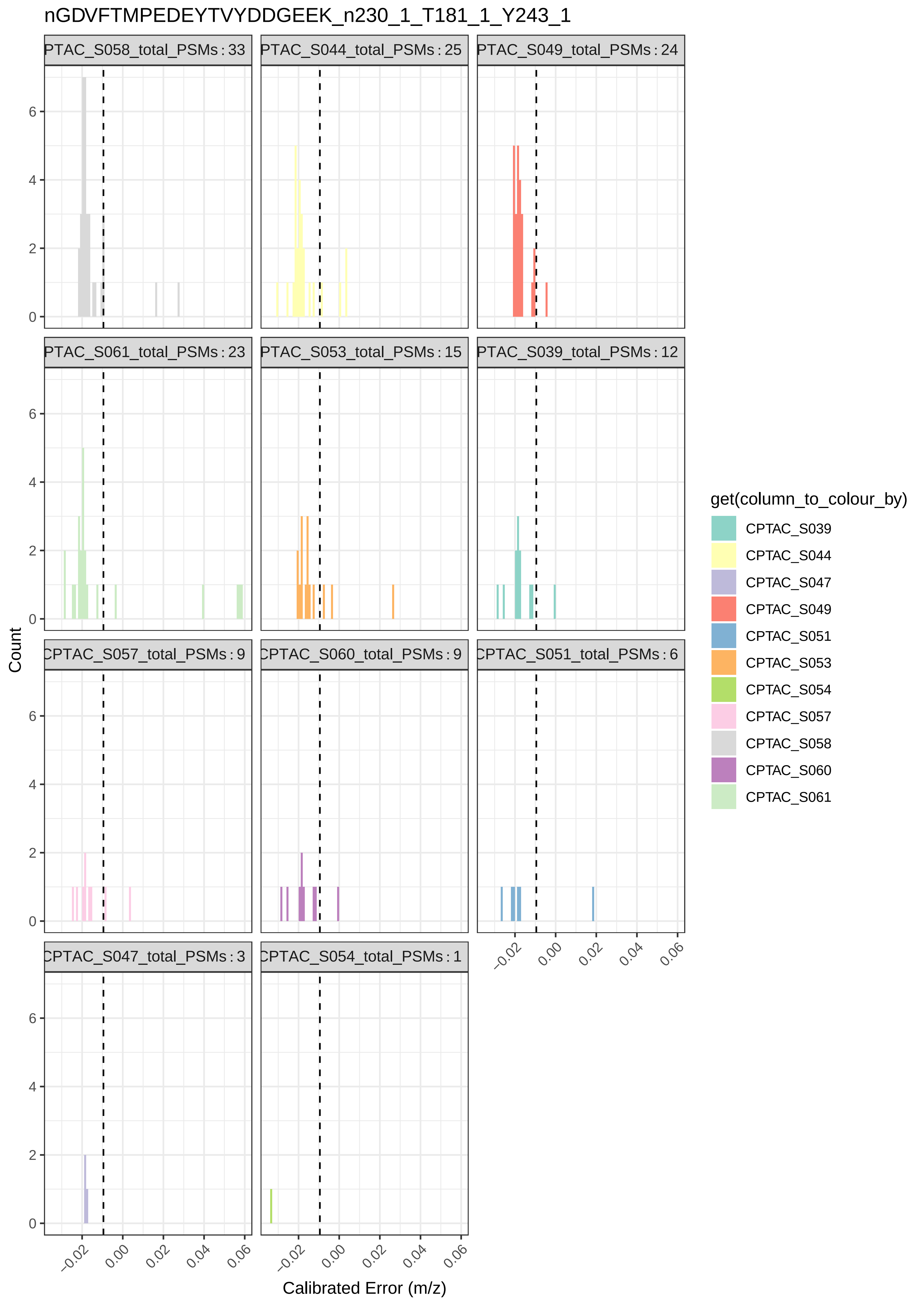

nGDVFTMPEDEYTVYDDGEEK_n230_1_T181_1_Y243_1
PTAC_S058_total_PSMs : 33
PTAC_S044_total_PSMs : 25
PTAC_S049_total_PSMs : 24
PTAC_S061_total_PSMs : 23
PTAC_S053_total_PSMs : 15
PTAC_S039_total_PSMs : 12
CPTAC_S057_total_PSMs : 9
CPTAC_S060_total_PSMs : 9
CPTAC_S051_total_PSMs : 6
CPTAC_S047_total_PSMs : 3
CPTAC_S054_total_PSMs : 1
Count
Calibrated Error (m/z)
get(column_to_colour_by)
CPTAC_S039
CPTAC_S044
CPTAC_S047
CPTAC_S049
CPTAC_S051
CPTAC_S053
CPTAC_S054
CPTAC_S057
CPTAC_S058
CPTAC_S060
CPTAC_S061

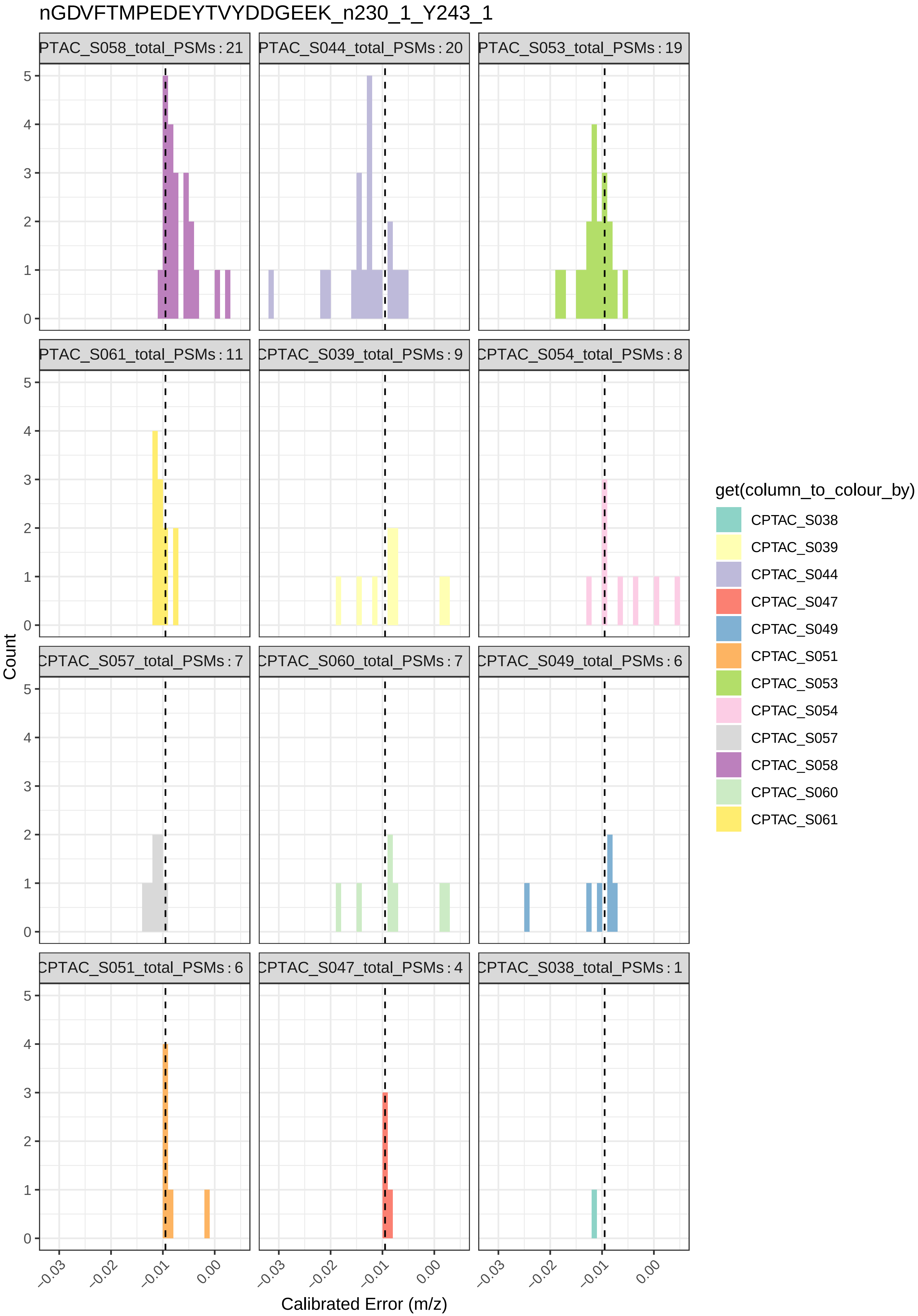

nGDVFTMPEDEYTVYDDGEEK_n230_1_Y243_1
PTAC_S058_total_PSMs : 21
PTAC_S044_total_PSMs : 20
PTAC_S053_total_PSMs : 19
PTAC_S061_total_PSMs : 11
CPTAC_S039_total_PSMs : 9
CPTAC_S054_total_PSMs : 8
CPTAC_S057_total_PSMs : 7
CPTAC_S060_total_PSMs : 7
CPTAC_S049_total_PSMs : 6
CPTAC_S051_total_PSMs : 6
CPTAC_S047_total_PSMs : 4
CPTAC_S038_total_PSMs : 1
Count
Calibrated Error (m/z)
−0.03
−0.02
−0.01
0.00
0
1
2
3
4
5
get(column_to_colour_by)
CPTAC_S038
CPTAC_S039
CPTAC_S044
CPTAC_S047
CPTAC_S049
CPTAC_S051
CPTAC_S053
CPTAC_S054
CPTAC_S057
CPTAC_S058
CPTAC_S060
CPTAC_S061

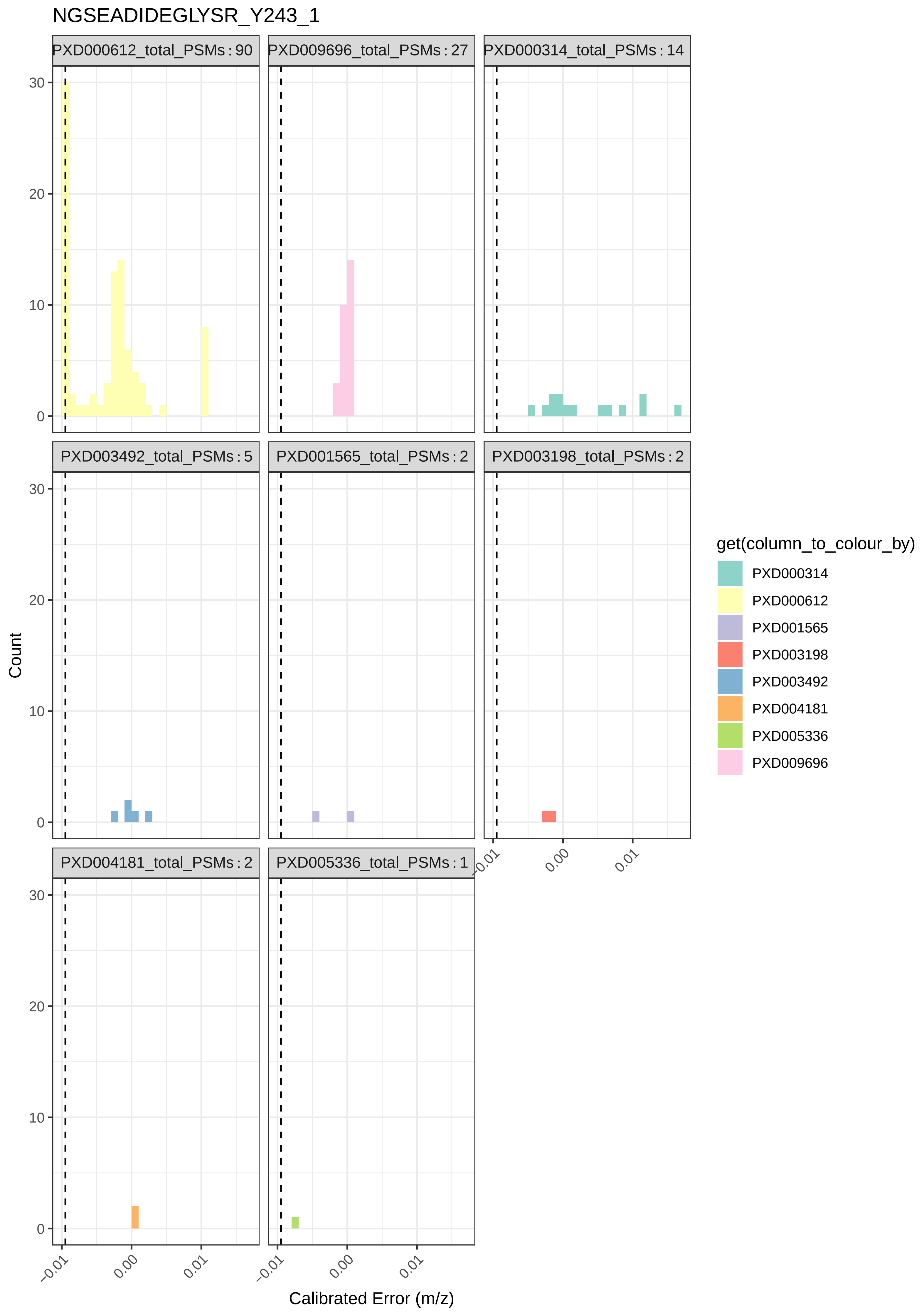

NGSEADIDEGLYSR_Y243_1
PXD000612_total_PSMs : 90
PXD009696_total_PSMs : 27
PXD000314_total_PSMs : 14
PXD003492_total_PSMs : 5
PXD001565_total_PSMs : 2
PXD003198_total_PSMs : 2
PXD004181_total_PSMs : 2
PXD005336_total_PSMs : 1
Count
0
10
20
30
−0.01
0.00
0.01
Calibrated Error (m/z)
get(column_to_colour_by)
PXD000314
PXD000612
PXD001565
PXD003198
PXD003492
PXD004181
PXD005336
PXD009696

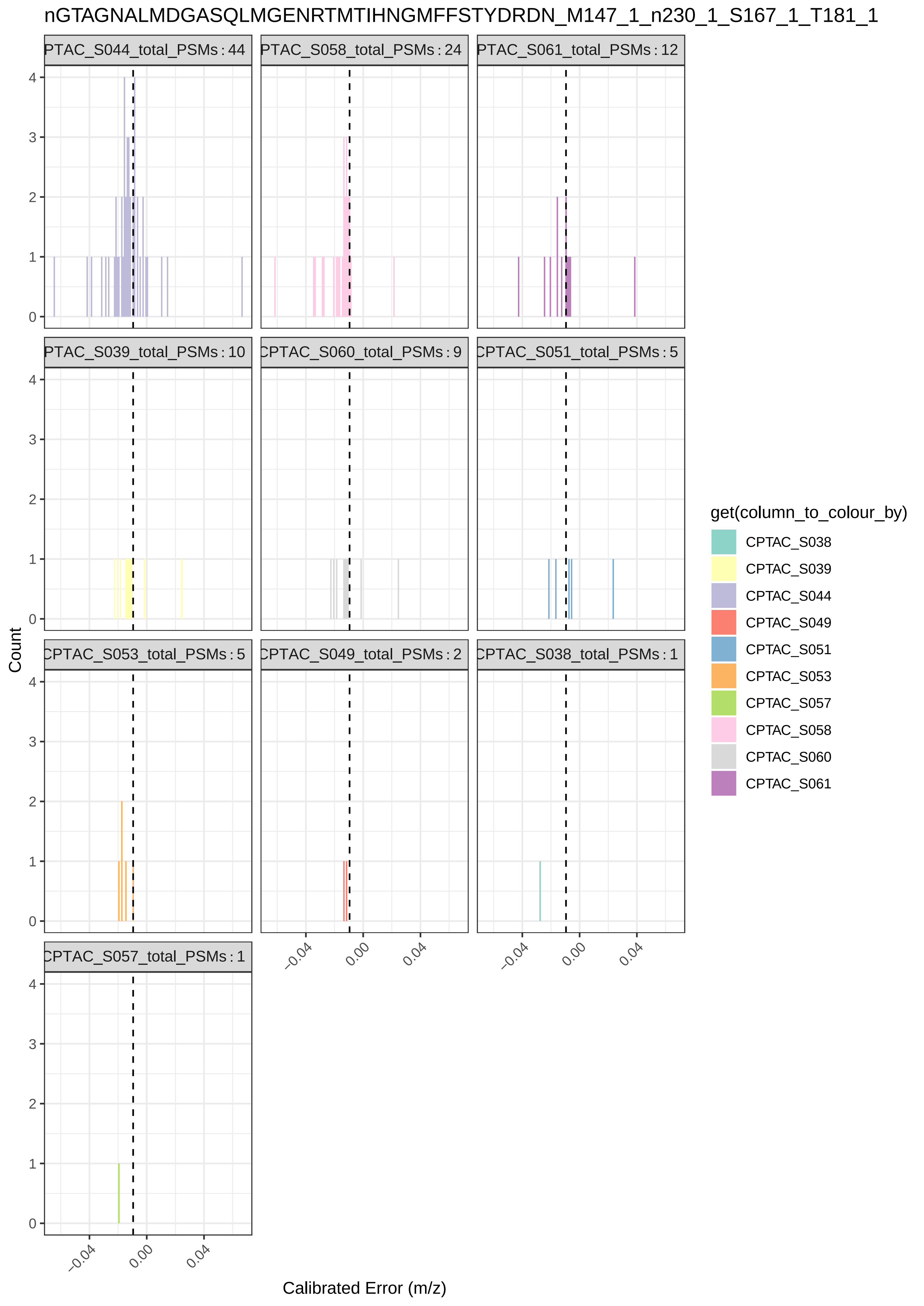

nGTAGNALMDGASQLMGENRTMTIHNGMFFSTYDRDN_M147_1_n230_1_S167_1_T181_1
PTAC_S044_total_PSMs : 44
PTAC_S058_total_PSMs : 24
PTAC_S061_total_PSMs : 12
PTAC_S039_total_PSMs : 10
CPTAC_S060_total_PSMs : 9
CPTAC_S051_total_PSMs : 5
CPTAC_S053_total_PSMs : 5
CPTAC_S049_total_PSMs : 2
CPTAC_S038_total_PSMs : 1
CPTAC_S057_total_PSMs : 1
Count
Calibrated Error (m/z)
−0.04
0.00
0.04
0
1
2
3
4
get(column_to_colour_by)
CPTAC_S038
CPTAC_S039
CPTAC_S044
CPTAC_S049
CPTAC_S051
CPTAC_S053
CPTAC_S057
CPTAC_S058
CPTAC_S060
CPTAC_S061

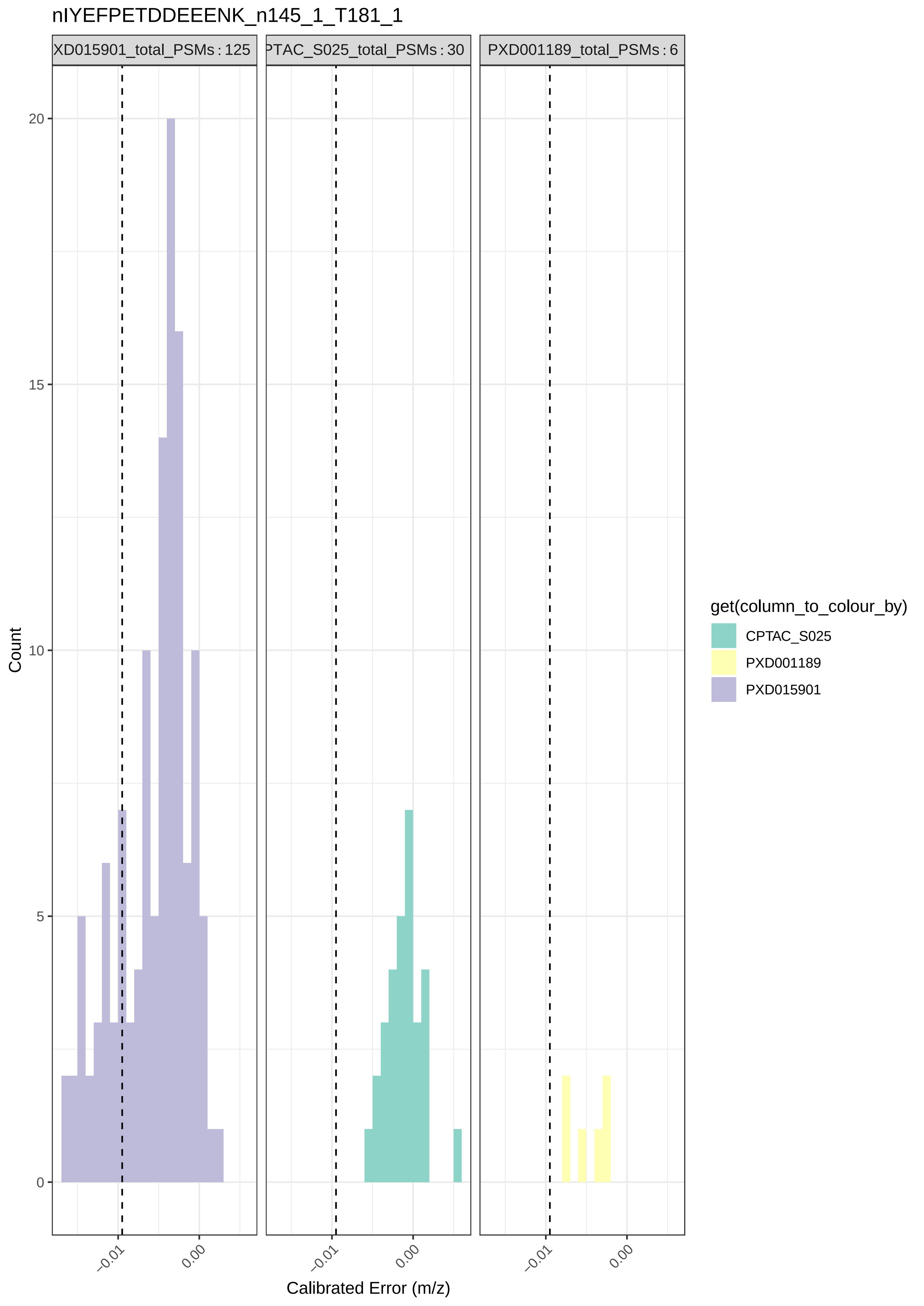

nIYEFPETDDEEENK_n145_1_T181_1
XD015901_total_PSMs : 125
PTAC_S025_total_PSMs : 30
PXD001189_total_PSMs : 6
Count
20
15
10
5
0
−0.01
0.00
Calibrated Error (m/z)
get(column_to_colour_by)
CPTAC_S025
PXD001189
PXD015901

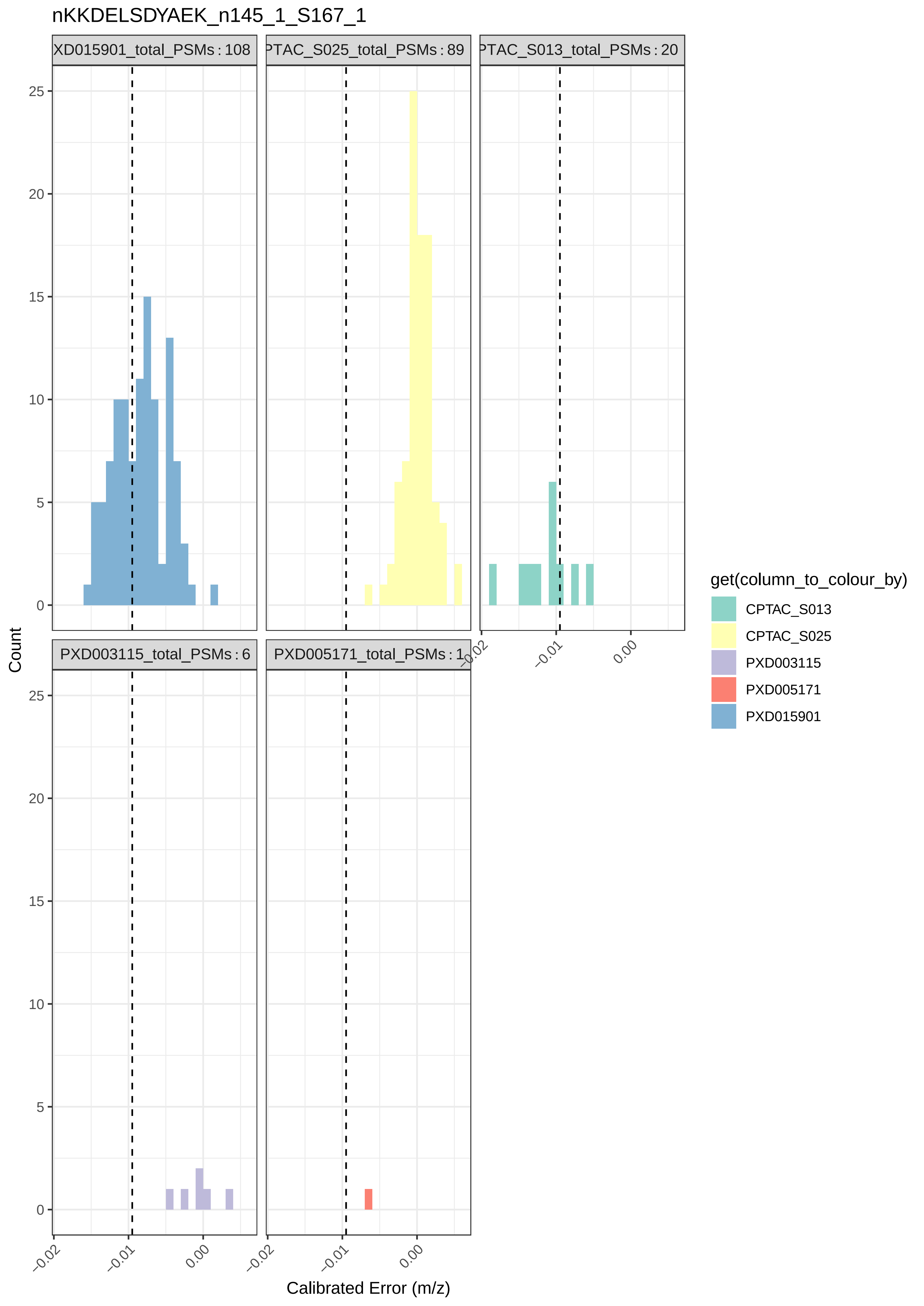

nKKDELSDYAEK_n145_1_S167_1
XD015901_total_PSMs : 108
PTAC_S025_total_PSMs : 89
PTAC_S013_total_PSMs : 20
PXD003115_total_PSMs : 6
PXD005171_total_PSMs : 1
Count
Calibrated Error (m/z)
0
5
10
15
20
25
−0.02
−0.01
0.00
get(column_to_colour_by)
CPTAC_S013
CPTAC_S025
PXD003115
PXD005171
PXD015901

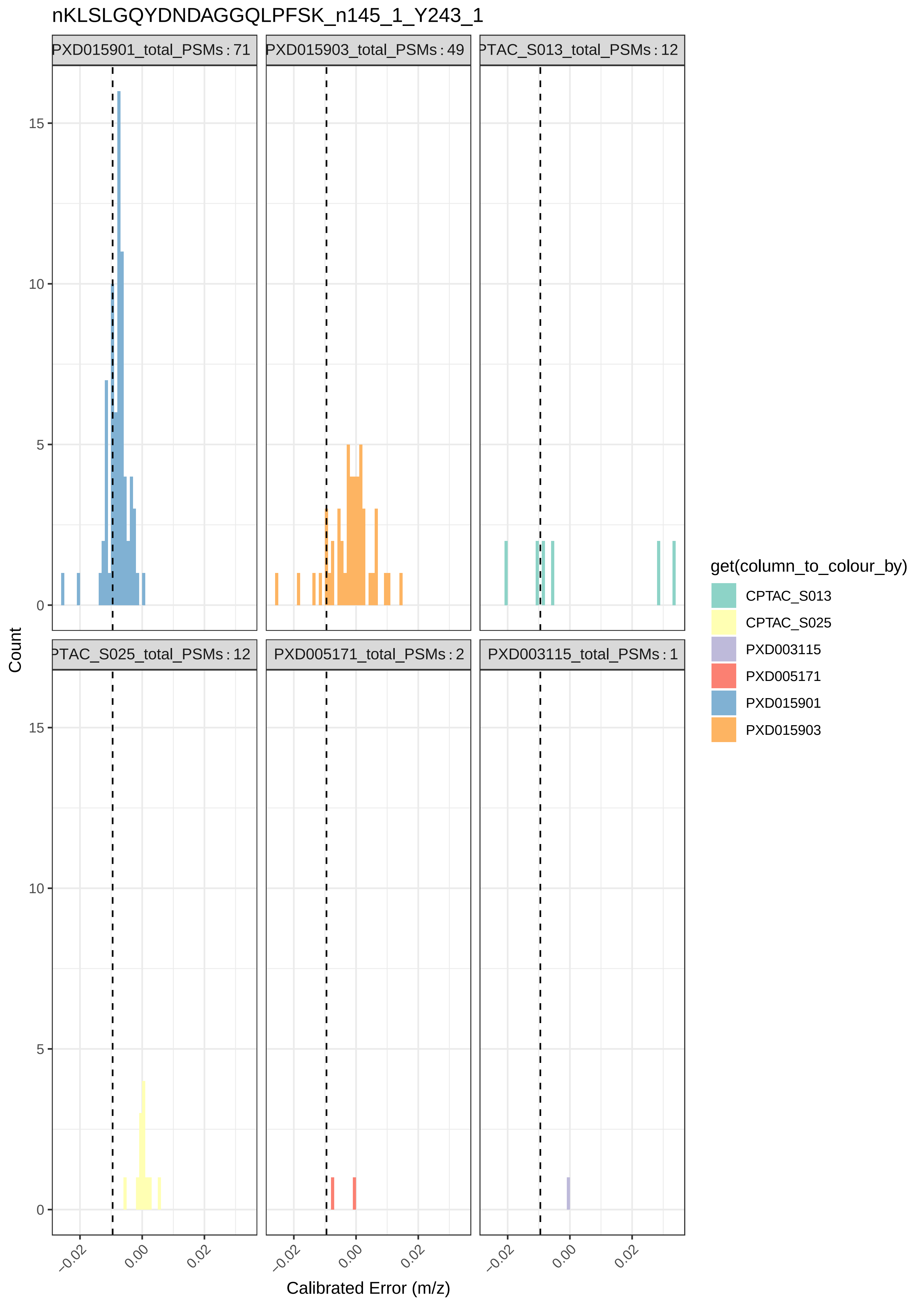

nKLSLGQYDNDAGGQLPFSK_n145_1_Y243_1
PXD015901_total_PSMs : 71
PXD015903_total_PSMs : 49
PTAC_S013_total_PSMs : 12
PTAC_S025_total_PSMs : 12
PXD005171_total_PSMs : 2
PXD003115_total_PSMs : 1
Count
0
5
10
15
−0.02
0.00
0.02
Calibrated Error (m/z)
get(column_to_colour_by)
CPTAC_S013
CPTAC_S025
PXD003115
PXD005171
PXD015901
PXD015903

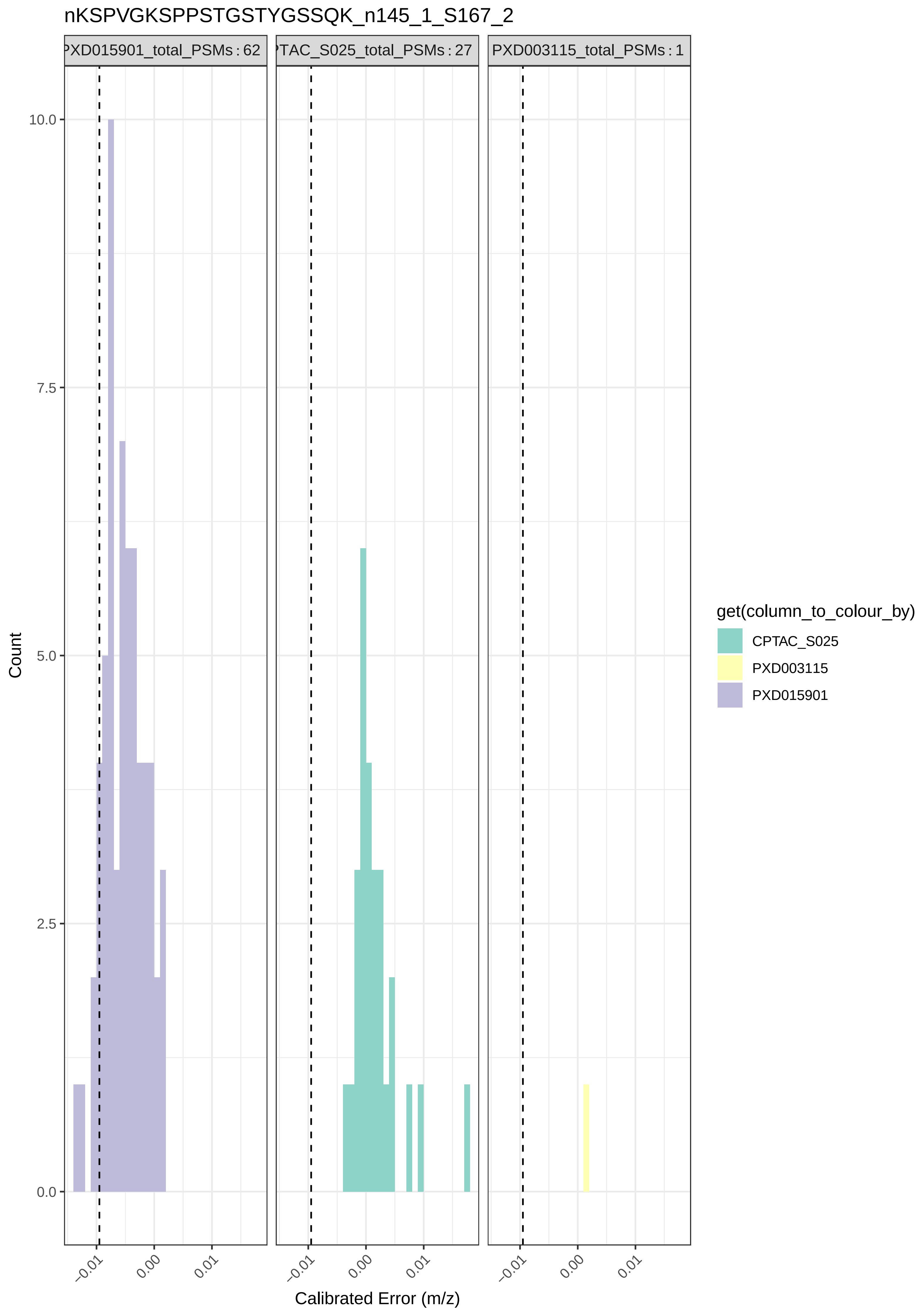

nKSPVGKSPPSTGSTYGSSQK_n145_1_S167_2
PXD015901_total_PSMs : 62
TAC_S025_total_PSMs : 27
PXD003115_total_PSMs : 1
Count
10.0
7.5
5.0
2.5
0.0
−0.01
0.00
0.01
Calibrated Error (m/z)
get(column_to_colour_by)
CPTAC_S025
PXD003115
PXD015901

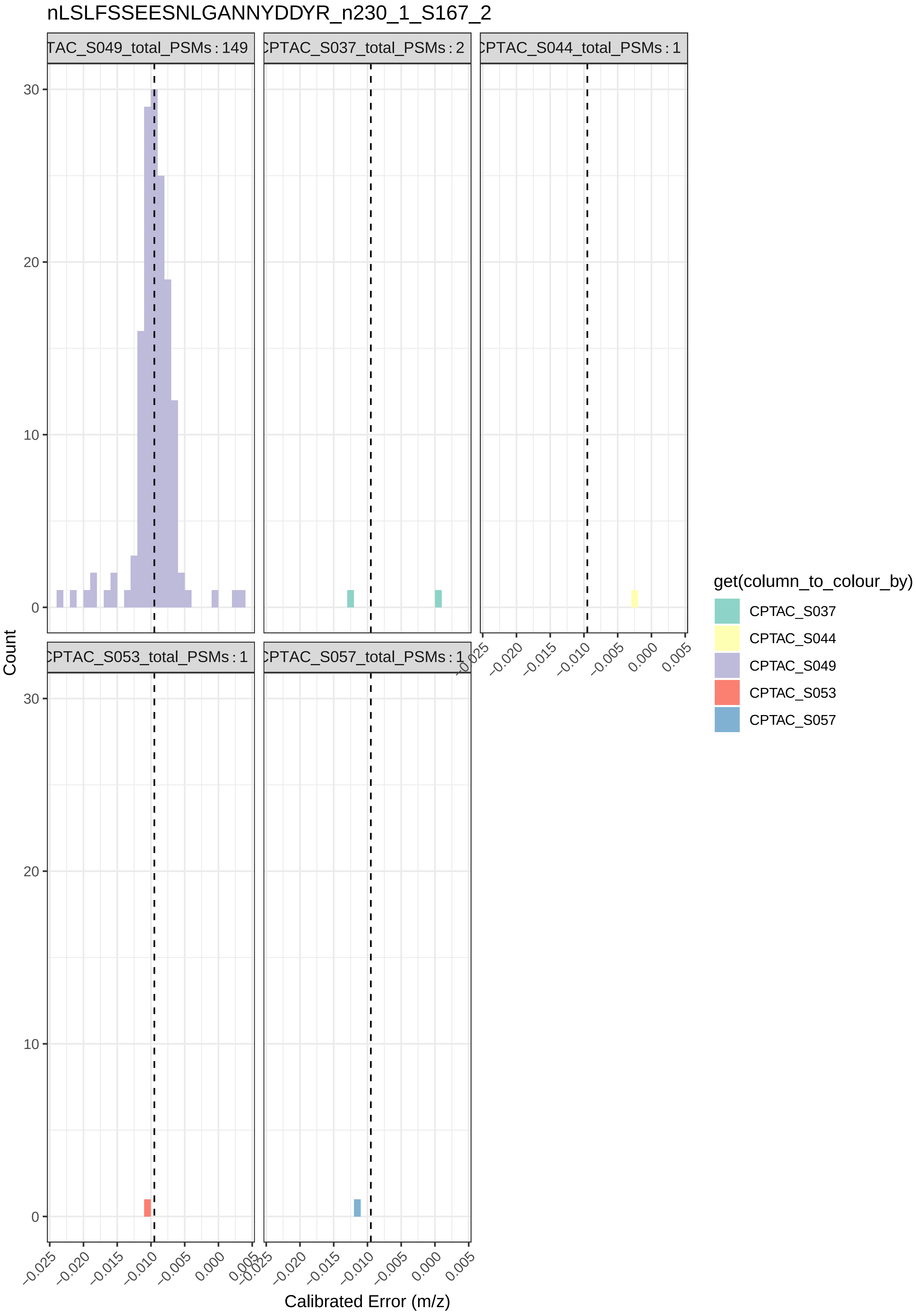
nLSLFSSEESNLGANNYDDYR_n230_1_S167_2
TAC_S049_total_PSMs : 149
CPTAC_S037_total_PSMs : 2
CPTAC_S044_total_PSMs : 1
CPTAC_S053_total_PSMs : 1
CPTAC_S057_total_PSMs : 1
Count
Calibrated Error (m/z)
0
10
20
30
−0.025
−0.020
−0.015
−0.010
−0.005
0.000
0.005
get(column_to_colour_by)
CPTAC_S037
CPTAC_S044
CPTAC_S049
CPTAC_S053
CPTAC_S057

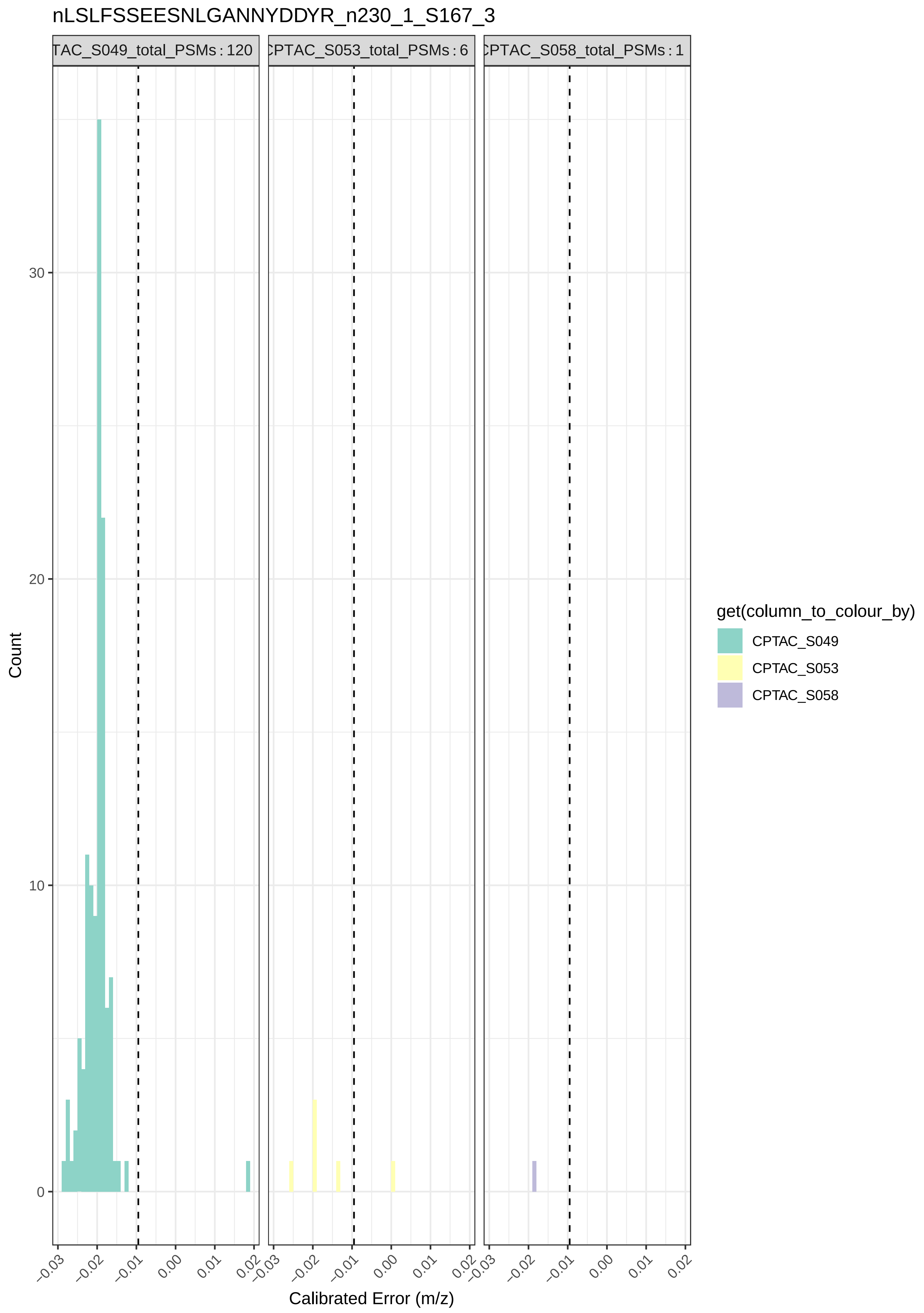
nLSLFSSEESNLGANNYDDYR_n230_1_S167_3
TAC_S049_total_PSMs : 120
CPTAC_S053_total_PSMs : 6
CPTAC_S058_total_PSMs : 1
Count
0
10
20
30
−0.03
−0.02
−0.01
0.00
0.01
0.02
Calibrated Error (m/z)
get(column_to_colour_by)
CPTAC_S049
CPTAC_S053
CPTAC_S058

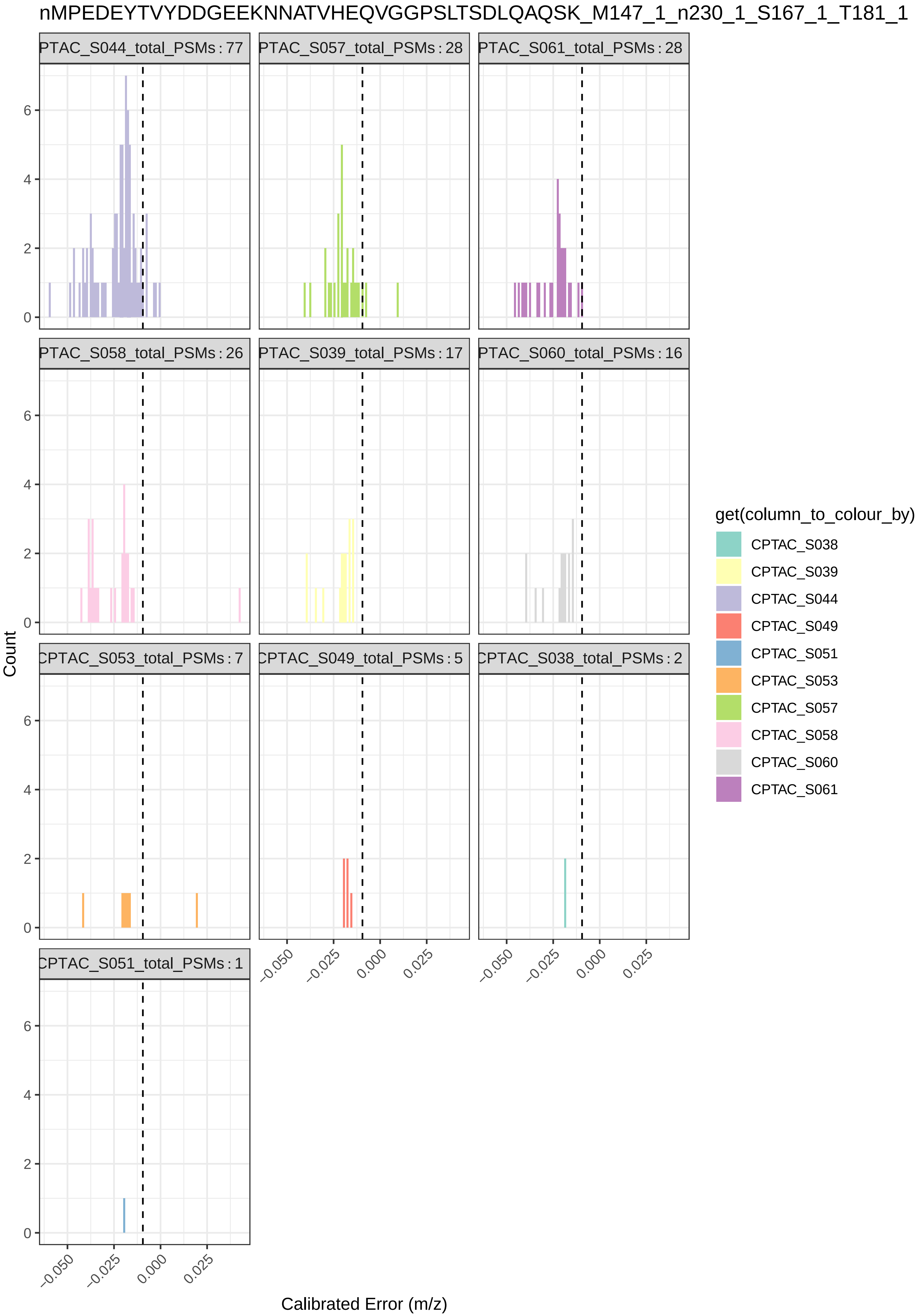

nMPEDEYTVYDDGEEKNNATVHEQVGGPSLTSDLQAQSK_M147_1_n230_1_S167_1_T181_1
PTAC_S044_total_PSMs : 77
PTAC_S057_total_PSMs : 28
PTAC_S061_total_PSMs : 28
PTAC_S058_total_PSMs : 26
PTAC_S039_total_PSMs : 17
PTAC_S060_total_PSMs : 16
CPTAC_S053_total_PSMs : 7
CPTAC_S049_total_PSMs : 5
CPTAC_S038_total_PSMs : 2
CPTAC_S051_total_PSMs : 1
Count
Calibrated Error (m/z)
0
2
4
6
−0.050
−0.025
0.000
0.025
get(column_to_colour_by)
CPTAC_S038
CPTAC_S039
CPTAC_S044
CPTAC_S049
CPTAC_S051
CPTAC_S053
CPTAC_S057
CPTAC_S058
CPTAC_S060
CPTAC_S061

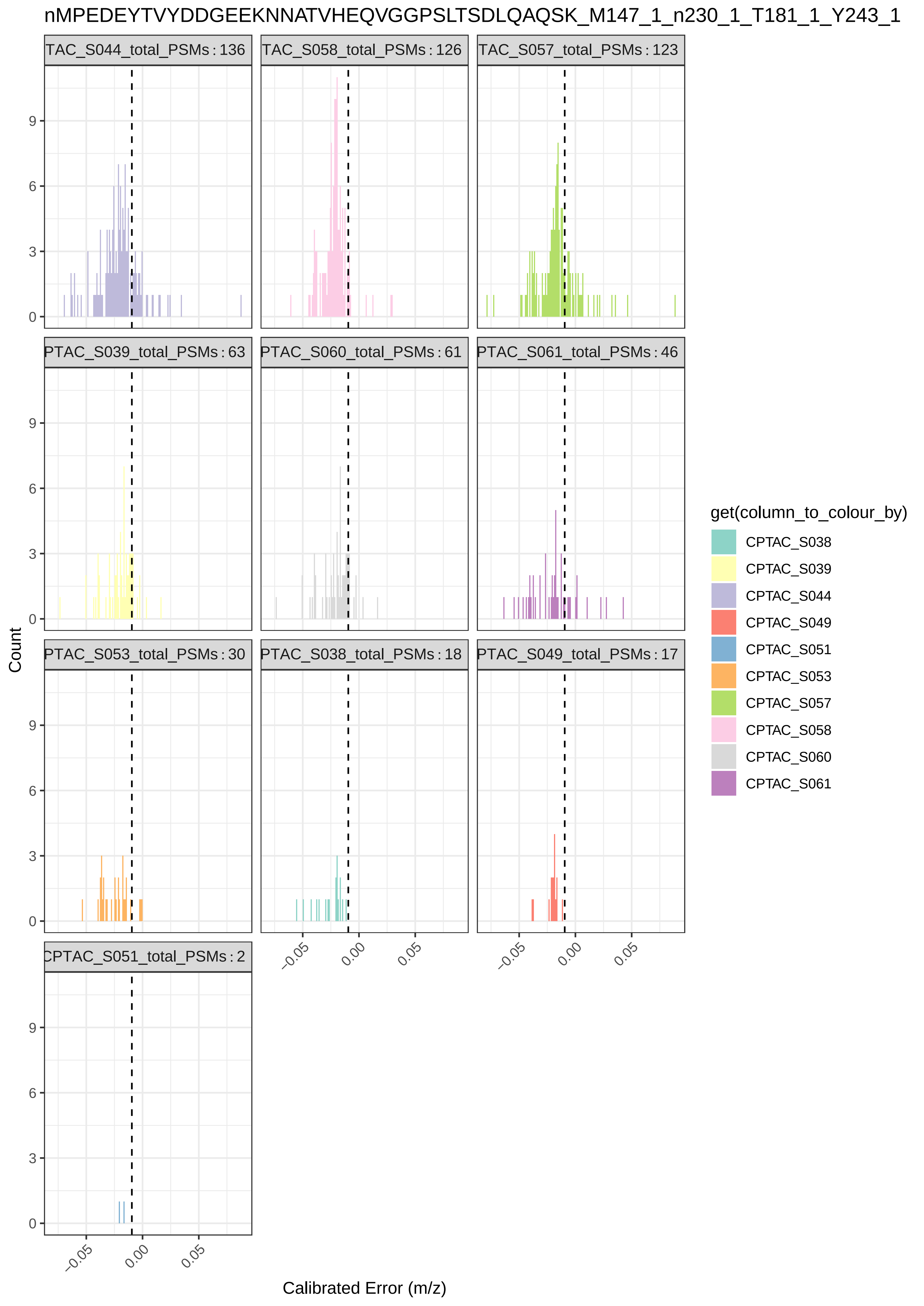

nMPEDEYTVYDDGEEKNNATVHEQVGGPSLTSDLQAQSK_M147_1_n230_1_T181_1_Y243_1
TAC_S044_total_PSMs : 136
TAC_S058_total_PSMs : 126
TAC_S057_total_PSMs : 123
PTAC_S039_total_PSMs : 63
PTAC_S060_total_PSMs : 61
PTAC_S061_total_PSMs : 46
PTAC_S053_total_PSMs : 30
PTAC_S038_total_PSMs : 18
PTAC_S049_total_PSMs : 17
CPTAC_S051_total_PSMs : 2
Count
Calibrated Error (m/z)
get(column_to_colour_by)
CPTAC_S038
CPTAC_S039
CPTAC_S044
CPTAC_S049
CPTAC_S051
CPTAC_S053
CPTAC_S057
CPTAC_S058
CPTAC_S060
CPTAC_S061

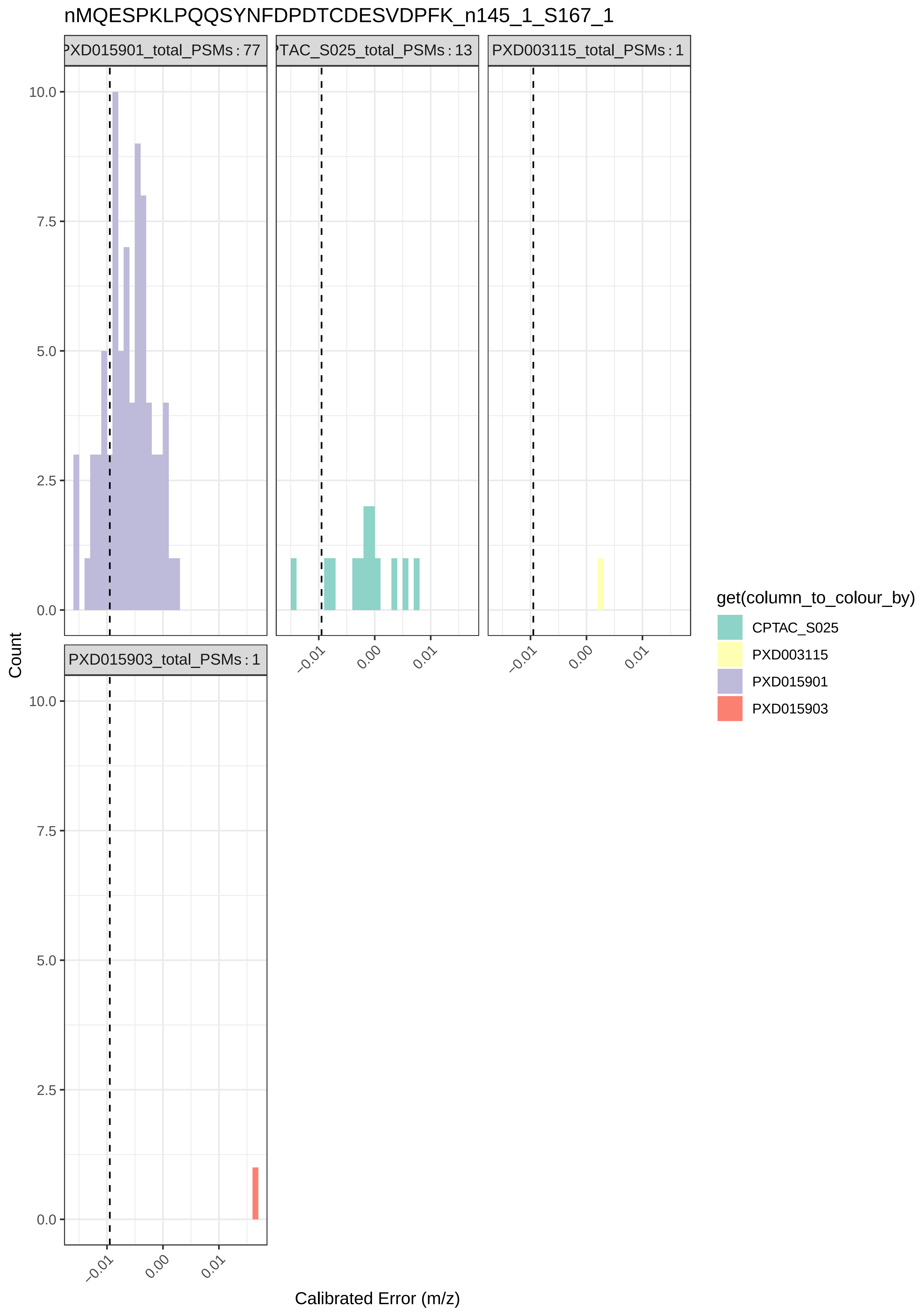

nMQESPKLPQQSYNFDPDTCDESVDPFK_n145_1_S167_1
PXD015901_total_PSMs : 77
TAC_S025_total_PSMs : 13
PXD003115_total_PSMs : 1
PXD015903_total_PSMs : 1
10.0
7.5
5.0
2.5
0.0
−0.01
0.00
0.01
Count
Calibrated Error (m/z)
get(column_to_colour_by)
CPTAC_S025
PXD003115
PXD015901
PXD015903

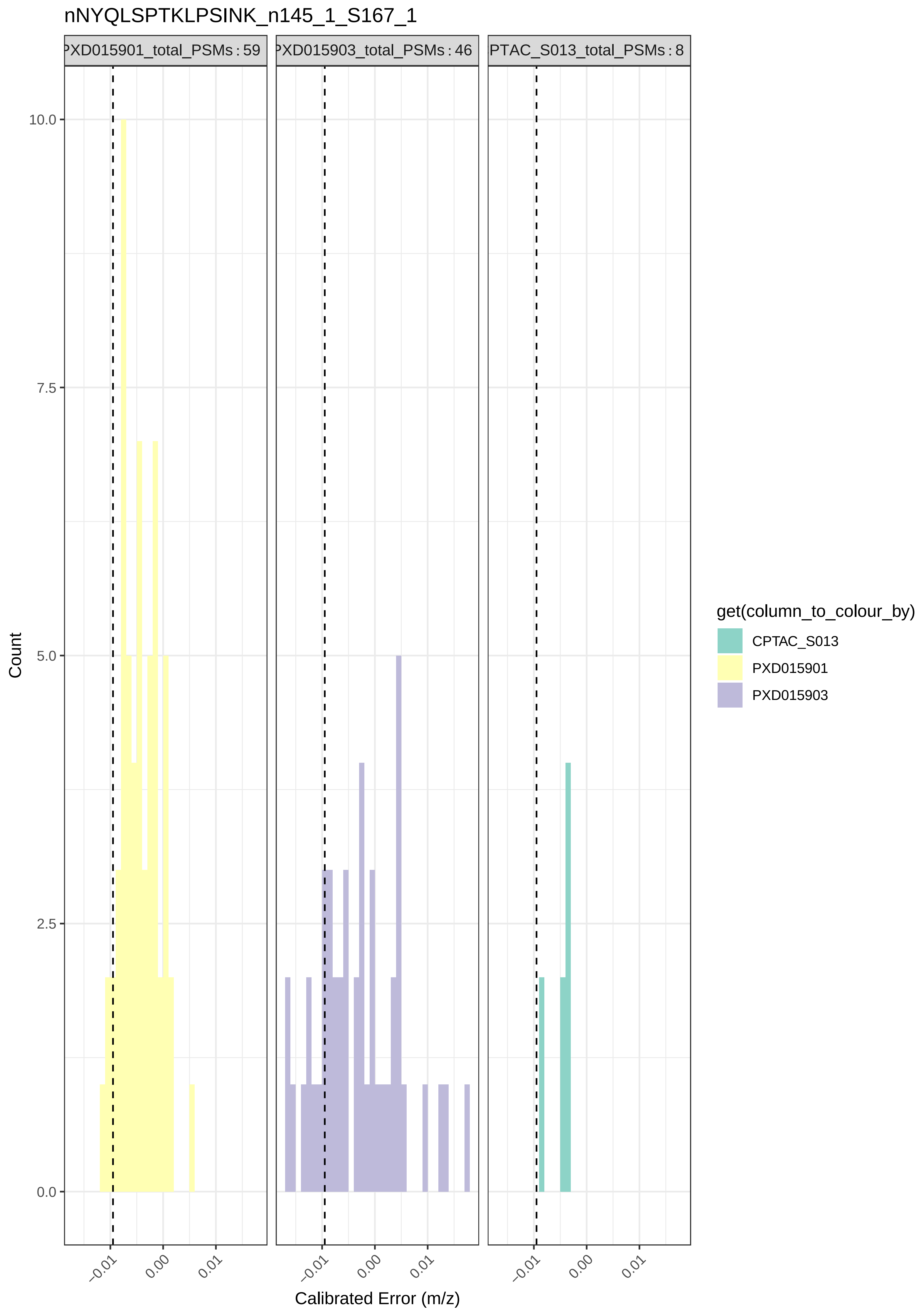

nNYQLSPTKLPSINK_n145_1_S167_1
PXD015901_total_PSMs : 59
PXD015903_total_PSMs : 46
PTAC_S013_total_PSMs : 8
Count
10.0
7.5
5.0
2.5
0.0
−0.01
0.00
0.01
Calibrated Error (m/z)
get(column_to_colour_by)
CPTAC_S013
PXD015901
PXD015903

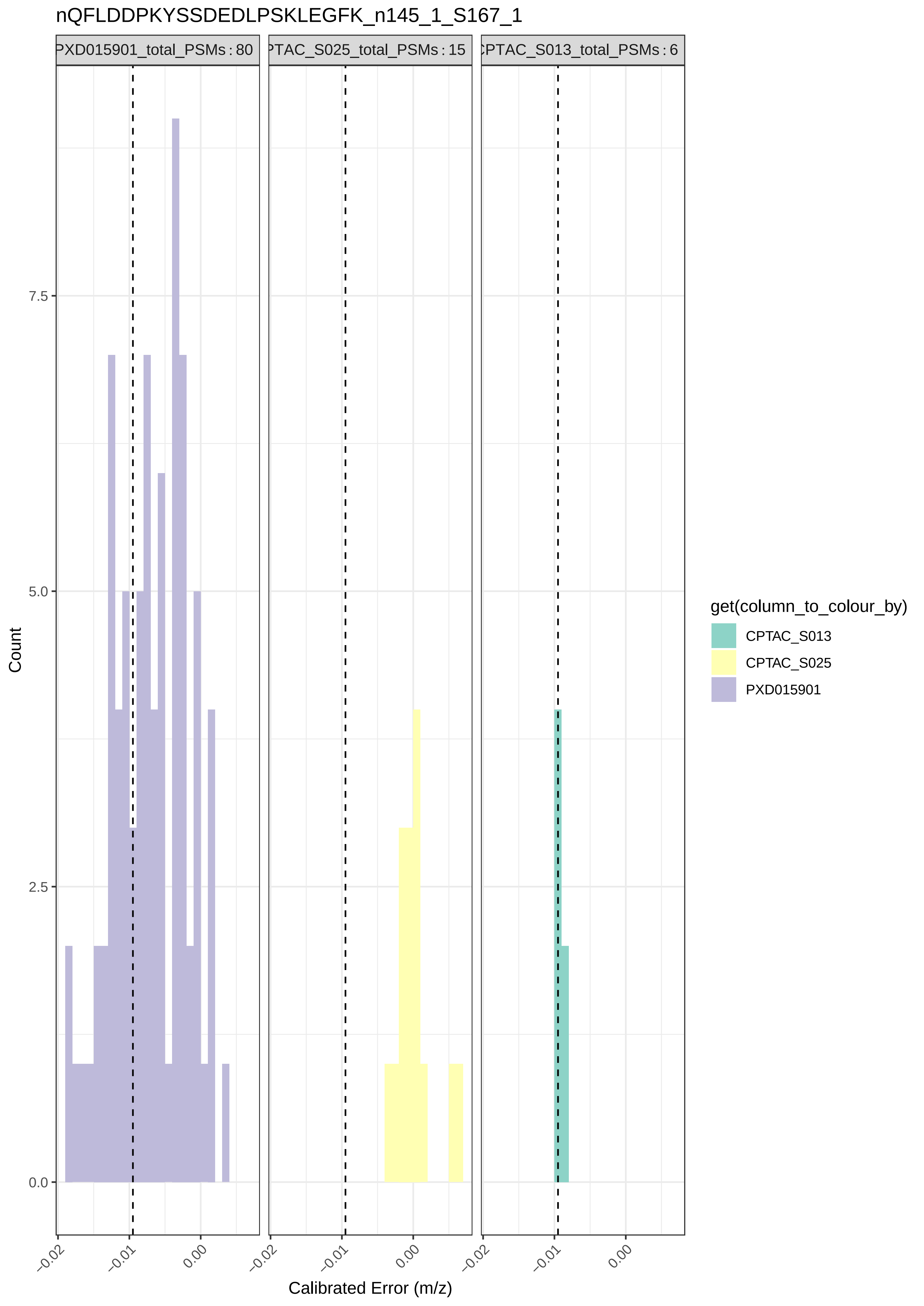
nQFLDDPKYSSDEDLPSKLEGFK_n145_1_S167_1
PXD015901_total_PSMs : 80
PTAC_S025_total_PSMs : 15
CPTAC_S013_total_PSMs : 6
Count
7.5
5.0
2.5
0.0
−0.02
−0.01
0.00
Calibrated Error (m/z)
get(column_to_colour_by)
CPTAC_S013
CPTAC_S025
PXD015901

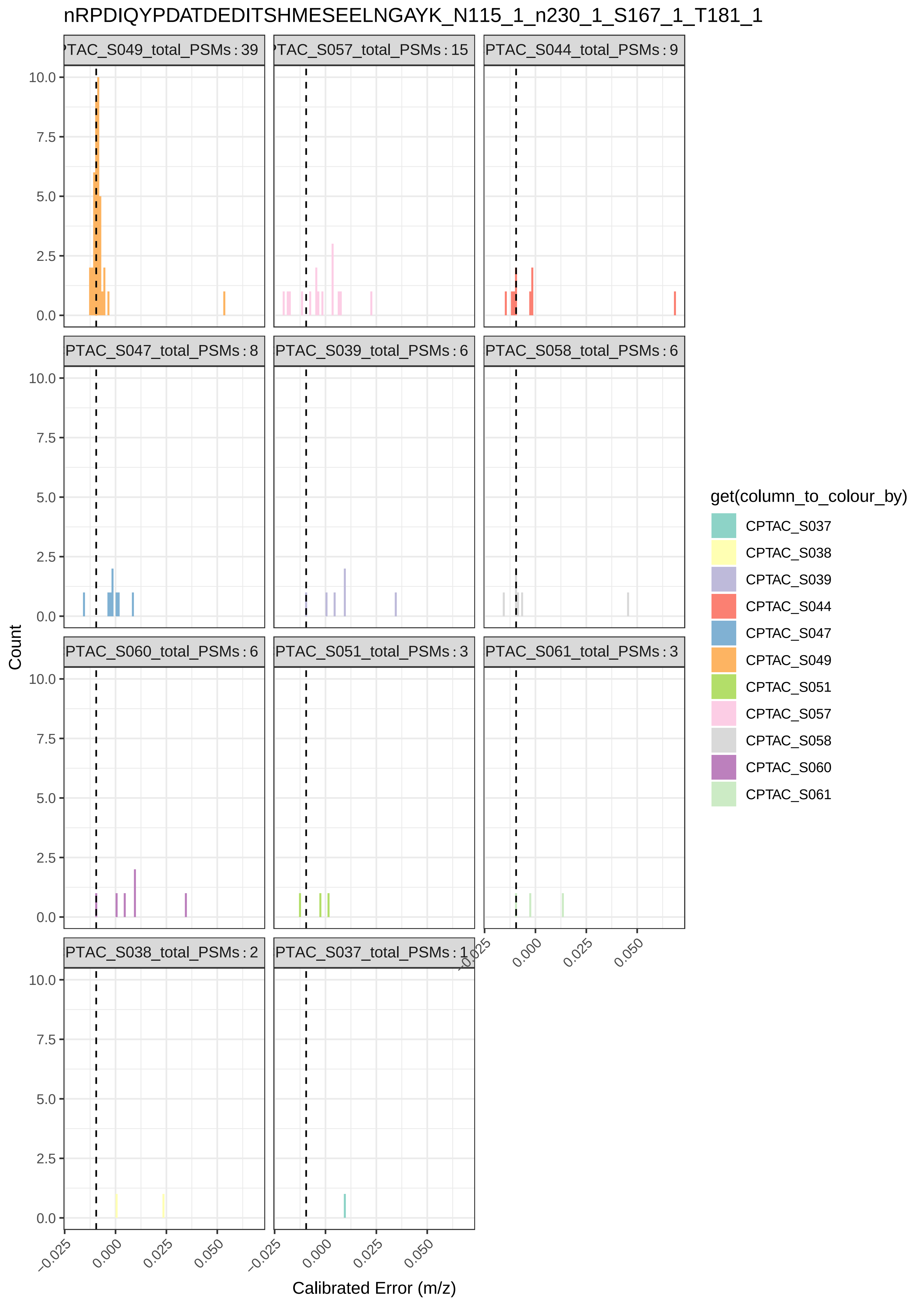
nRPDIQYPDATDEDITSHMESEELNGAYK_N115_1_n230_1_S167_1_T181_1
PTAC_S049_total_PSMs : 39
PTAC_S057_total_PSMs : 15
PTAC_S044_total_PSMs : 9
PTAC_S047_total_PSMs : 8
PTAC_S039_total_PSMs : 6
PTAC_S058_total_PSMs : 6
PTAC_S060_total_PSMs : 6
PTAC_S051_total_PSMs : 3
PTAC_S061_total_PSMs : 3
PTAC_S038_total_PSMs : 2
PTAC_S037_total_PSMs : 1
Count
Calibrated Error (m/z)
get(column_to_colour_by)
CPTAC_S037
CPTAC_S038
CPTAC_S039
CPTAC_S044
CPTAC_S047
CPTAC_S049
CPTAC_S051
CPTAC_S057
CPTAC_S058
CPTAC_S060
CPTAC_S061

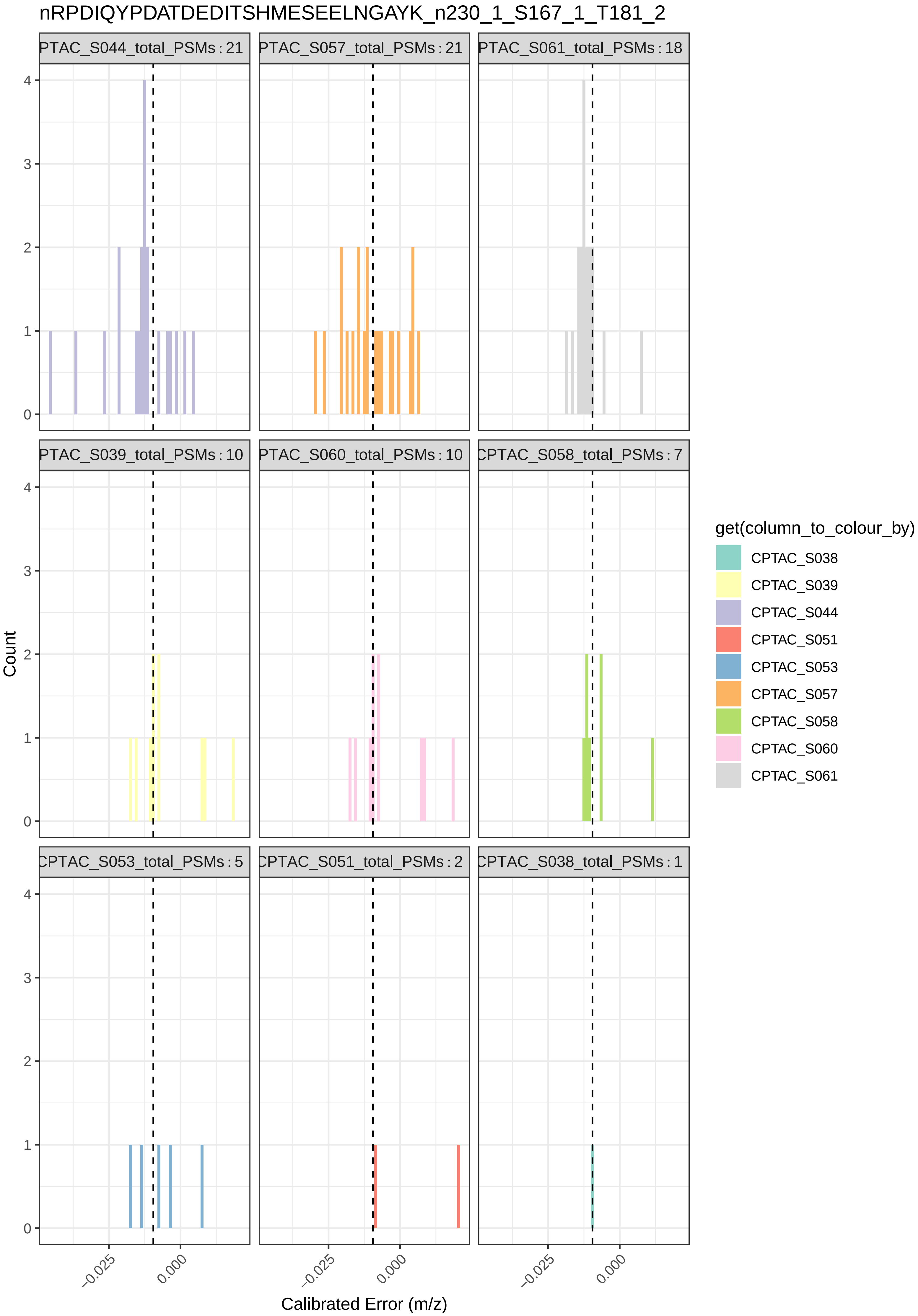

nRPDIQYPDATDEDITSHMESEELNGAYK_n230_1_S167_1_T181_2
PTAC_S044_total_PSMs : 21
PTAC_S057_total_PSMs : 21
PTAC_S061_total_PSMs : 18
PTAC_S039_total_PSMs : 10
PTAC_S060_total_PSMs : 10
CPTAC_S058_total_PSMs : 7
CPTAC_S053_total_PSMs : 5
CPTAC_S051_total_PSMs : 2
CPTAC_S038_total_PSMs : 1
Count
0
1
2
3
4
−0.025
0.000
Calibrated Error (m/z)
get(column_to_colour_by)
CPTAC_S038
CPTAC_S039
CPTAC_S044
CPTAC_S051
CPTAC_S053
CPTAC_S057
CPTAC_S058
CPTAC_S060
CPTAC_S061

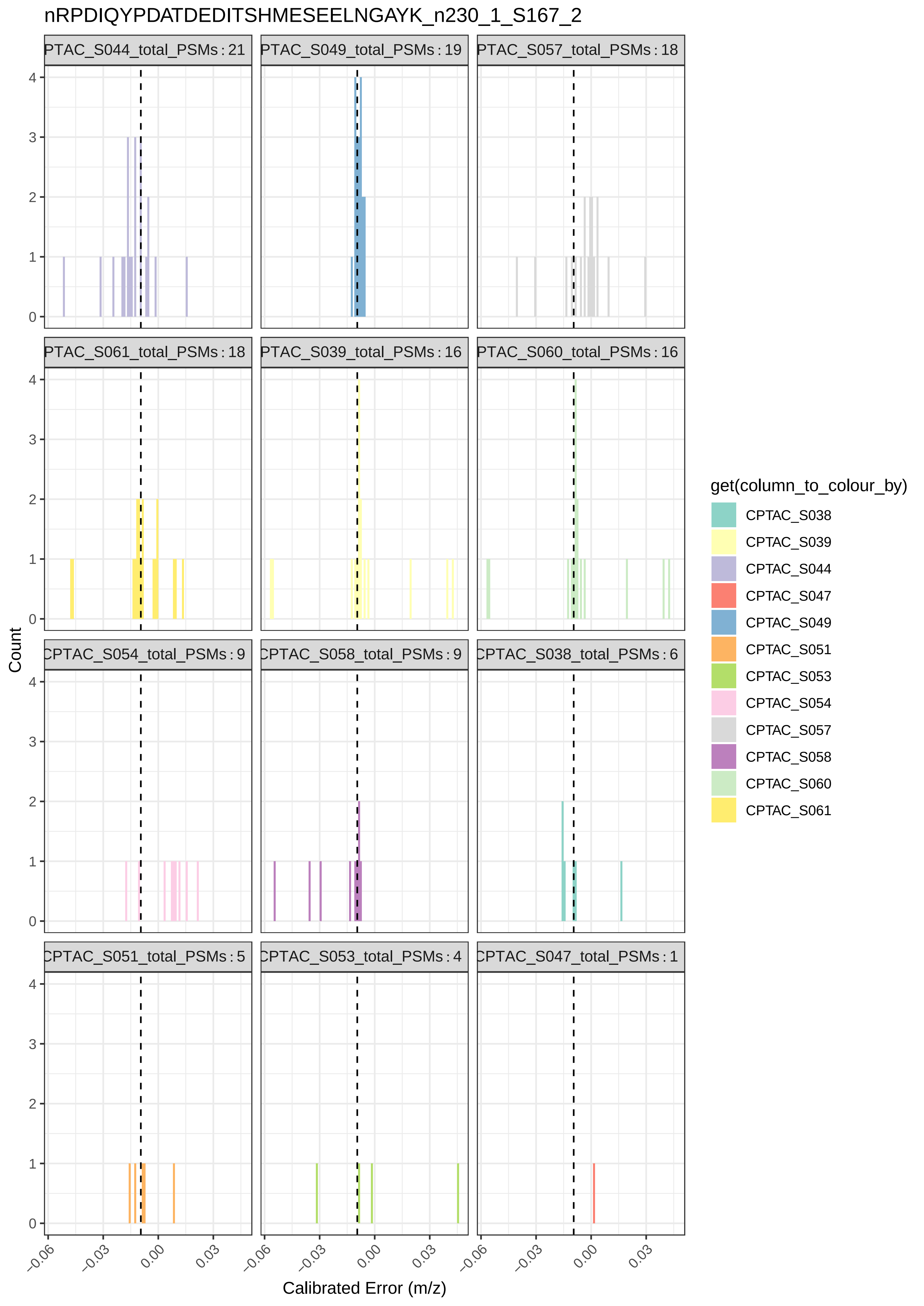

nRPDIQYPDATDEDITSHMESEELNGAYK_n230_1_S167_2
PTAC_S044_total_PSMs : 21
PTAC_S049_total_PSMs : 19
PTAC_S057_total_PSMs : 18
PTAC_S061_total_PSMs : 18
PTAC_S039_total_PSMs : 16
PTAC_S060_total_PSMs : 16
CPTAC_S054_total_PSMs : 9
CPTAC_S058_total_PSMs : 9
CPTAC_S038_total_PSMs : 6
CPTAC_S051_total_PSMs : 5
CPTAC_S053_total_PSMs : 4
CPTAC_S047_total_PSMs : 1
Count
0
1
2
3
4
−0.06
−0.03
0.00
0.03
Calibrated Error (m/z)
get(column_to_colour_by)
CPTAC_S038
CPTAC_S039
CPTAC_S044
CPTAC_S047
CPTAC_S049
CPTAC_S051
CPTAC_S053
CPTAC_S054
CPTAC_S057
CPTAC_S058
CPTAC_S060
CPTAC_S061

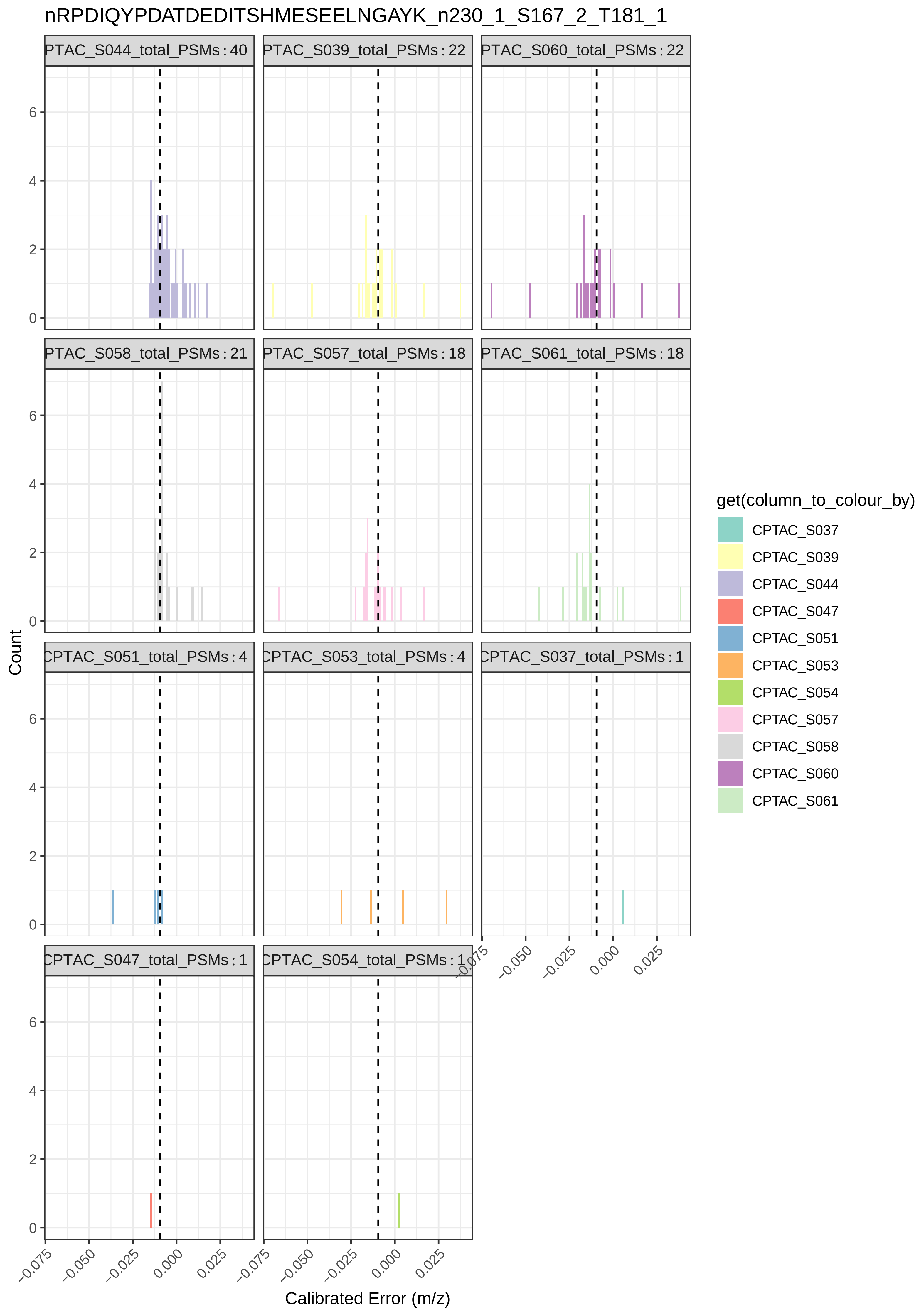

nRPDIQYPDATDEDITSHMESEELNGAYK_n230_1_S167_2_T181_1
PTAC_S044_total_PSMs : 40
PTAC_S039_total_PSMs : 22
PTAC_S060_total_PSMs : 22
PTAC_S058_total_PSMs : 21
PTAC_S057_total_PSMs : 18
PTAC_S061_total_PSMs : 18
CPTAC_S051_total_PSMs : 4
CPTAC_S053_total_PSMs : 4
CPTAC_S037_total_PSMs : 1
CPTAC_S047_total_PSMs : 1
CPTAC_S054_total_PSMs : 1
Count
0
2
4
6
−0.075
−0.050
−0.025
0.000
0.025
Calibrated Error (m/z)
get(column_to_colour_by)
CPTAC_S037
CPTAC_S039
CPTAC_S044
CPTAC_S047
CPTAC_S051
CPTAC_S053
CPTAC_S054
CPTAC_S057
CPTAC_S058
CPTAC_S060
CPTAC_S061

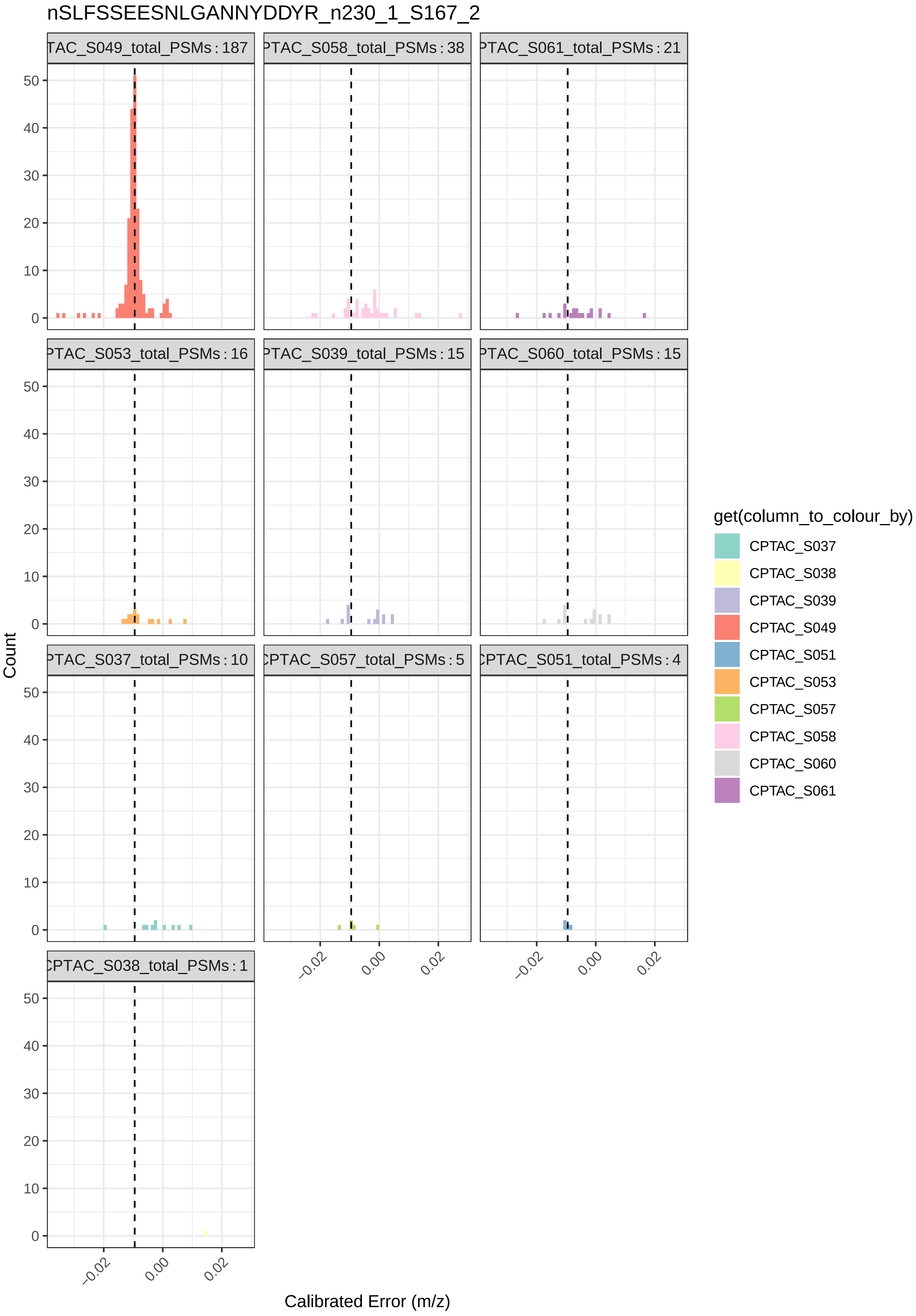

nSLFSSEESNLGANNYDDYR_n230_1_S167_2
TAC_S049_total_PSMs : 187
PTAC_S058_total_PSMs : 38
PTAC_S061_total_PSMs : 21
PTAC_S053_total_PSMs : 16
PTAC_S039_total_PSMs : 15
PTAC_S060_total_PSMs : 15
PTAC_S037_total_PSMs : 10
CPTAC_S057_total_PSMs : 5
CPTAC_S051_total_PSMs : 4
CPTAC_S038_total_PSMs : 1
Count
Calibrated Error (m/z)
get(column_to_colour_by)
CPTAC_S037
CPTAC_S038
CPTAC_S039
CPTAC_S049
CPTAC_S051
CPTAC_S053
CPTAC_S057
CPTAC_S058
CPTAC_S060
CPTAC_S061

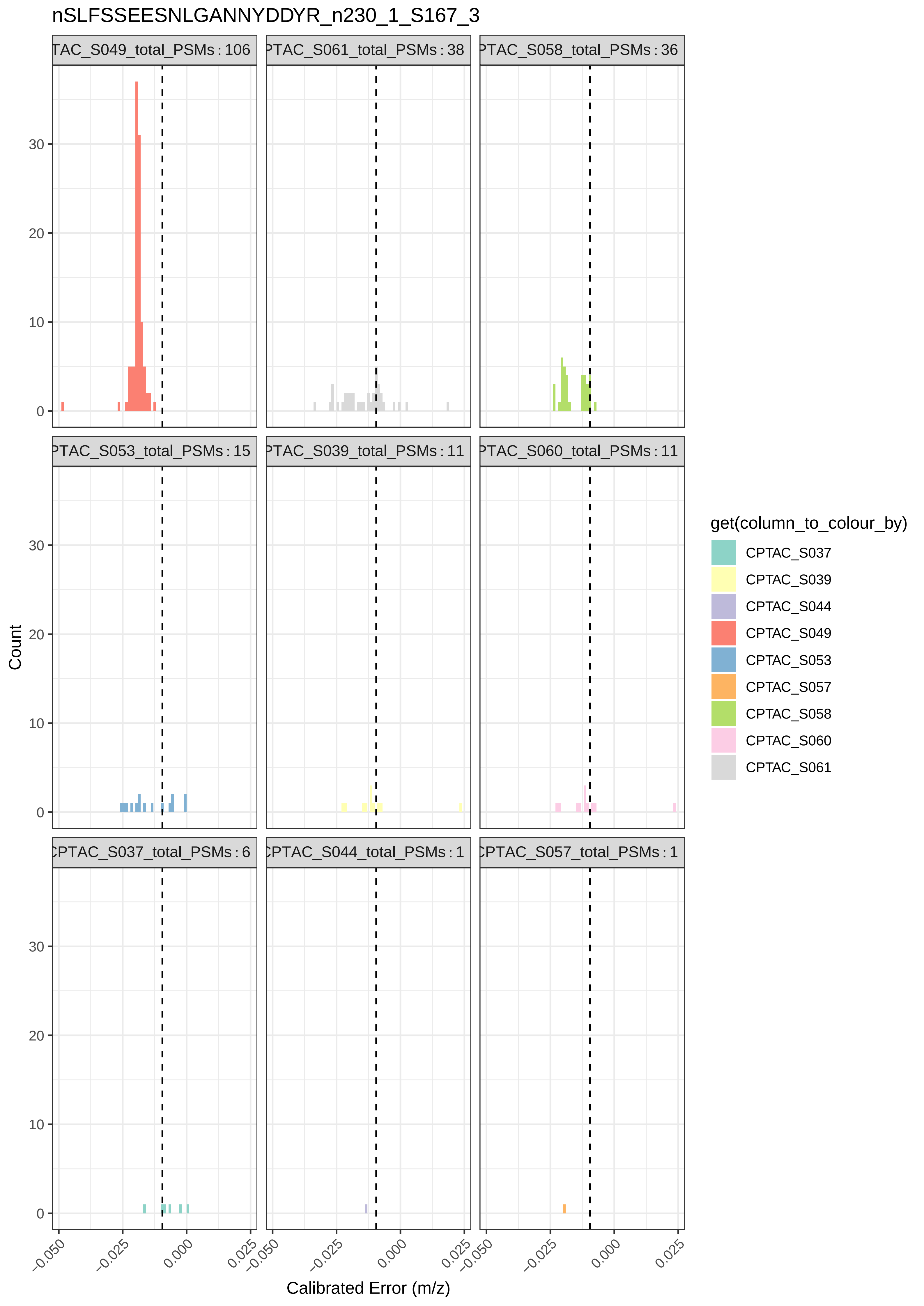
nSLFSSEESNLGANNYDDYR_n230_1_S167_3
TAC_S049_total_PSMs : 106
PTAC_S061_total_PSMs : 38
PTAC_S058_total_PSMs : 36
PTAC_S053_total_PSMs : 15
PTAC_S039_total_PSMs : 11
PTAC_S060_total_PSMs : 11
CPTAC_S037_total_PSMs : 6
CPTAC_S044_total_PSMs : 1
CPTAC_S057_total_PSMs : 1
Count
Calibrated Error (m/z)
0
10
20
30
−0.050
−0.025
0.000
0.025
get(column_to_colour_by)
CPTAC_S037
CPTAC_S039
CPTAC_S044
CPTAC_S049
CPTAC_S053
CPTAC_S057
CPTAC_S058
CPTAC_S060
CPTAC_S061

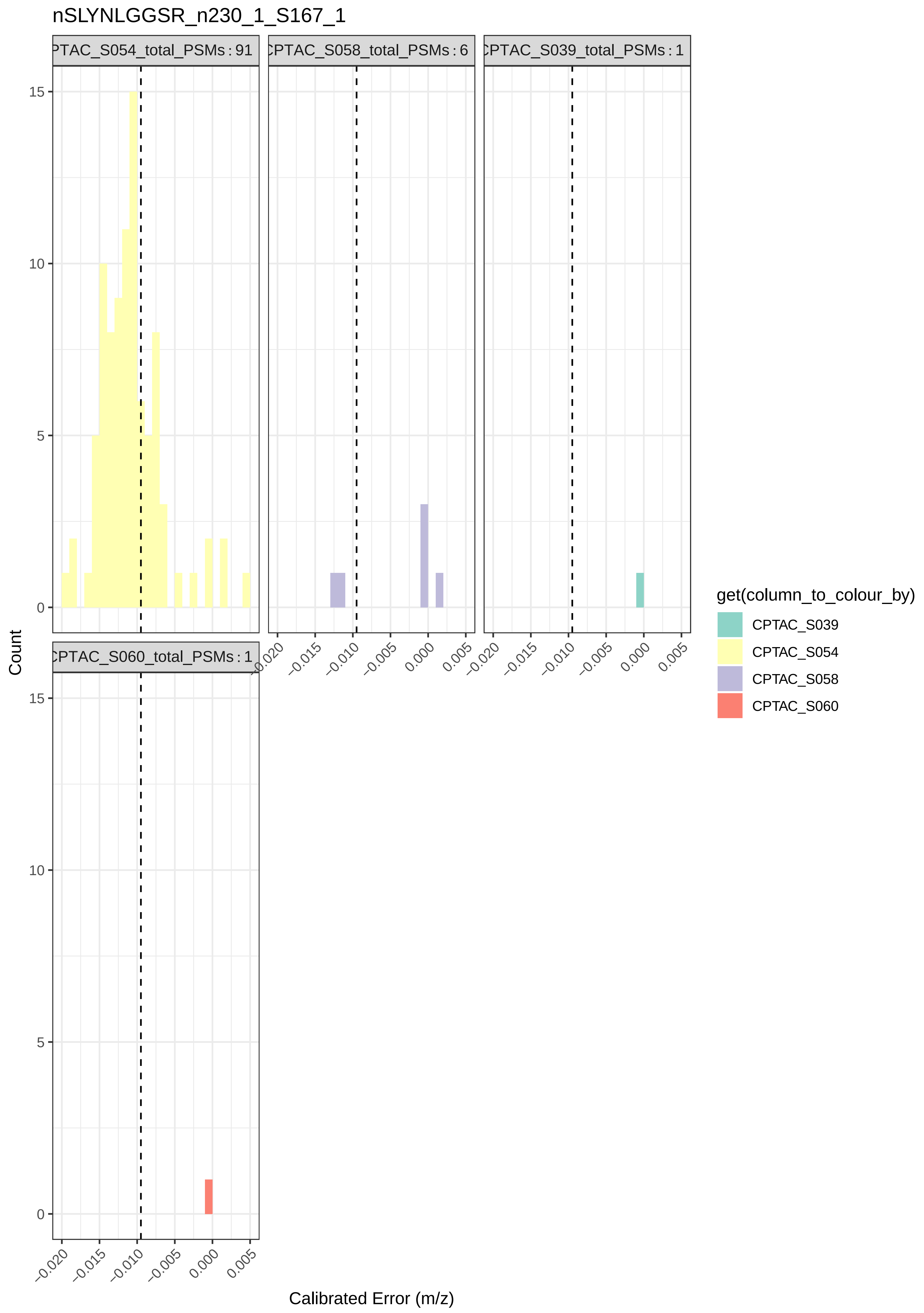

nSLYNLGGSR_n230_1_S167_1
PTAC_S054_total_PSMs : 91
CPTAC_S058_total_PSMs : 6
CPTAC_S039_total_PSMs : 1
CPTAC_S060_total_PSMs : 1
Count
Calibrated Error (m/z)
0
5
10
15
−0.020
−0.015
−0.010
−0.005
0.000
0.005
get(column_to_colour_by)
CPTAC_S039
CPTAC_S054
CPTAC_S058
CPTAC_S060

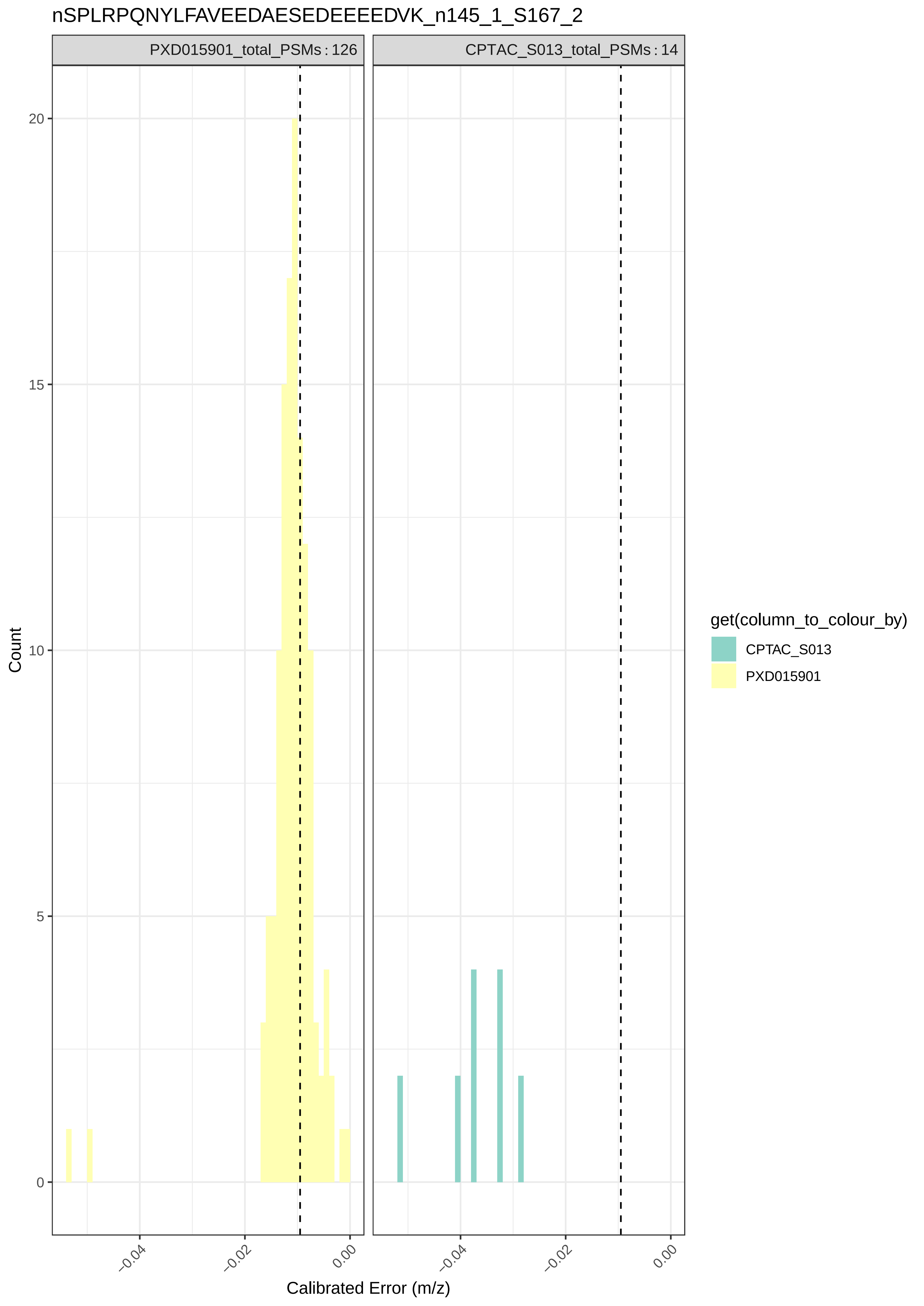
nSPLRPQNYLFAVEEDAESEDEEEEDVK_n145_1_S167_2
PXD015901_total_PSMs : 126
CPTAC_S013_total_PSMs : 14
Count
20
15
10
5
0
−0.04
−0.02
0.00
Calibrated Error (m/z)
get(column_to_colour_by)
CPTAC_S013
PXD015901

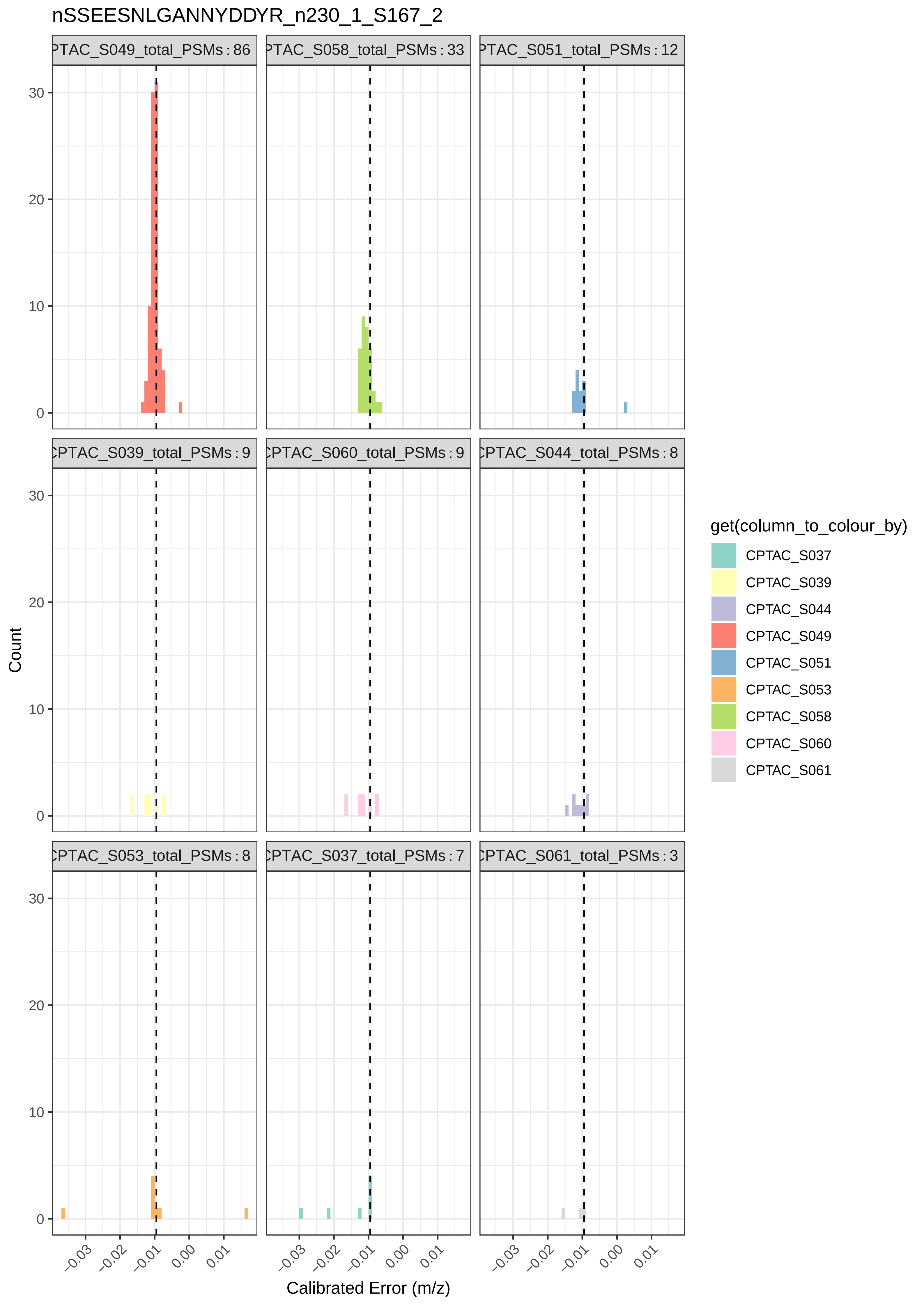

nSSEESNLGANNYDDYR_n230_1_S167_2
PTAC_S049_total_PSMs : 86
PTAC_S058_total_PSMs : 33
PTAC_S051_total_PSMs : 12
CPTAC_S039_total_PSMs : 9
CPTAC_S060_total_PSMs : 9
CPTAC_S044_total_PSMs : 8
CPTAC_S053_total_PSMs : 8
CPTAC_S037_total_PSMs : 7
CPTAC_S061_total_PSMs : 3
Count
0
10
20
30
−0.03
−0.02
−0.01
0.00
0.01
Calibrated Error (m/z)
get(column_to_colour_by)
CPTAC_S037
CPTAC_S039
CPTAC_S044
CPTAC_S049
CPTAC_S051
CPTAC_S053
CPTAC_S058
CPTAC_S060
CPTAC_S061

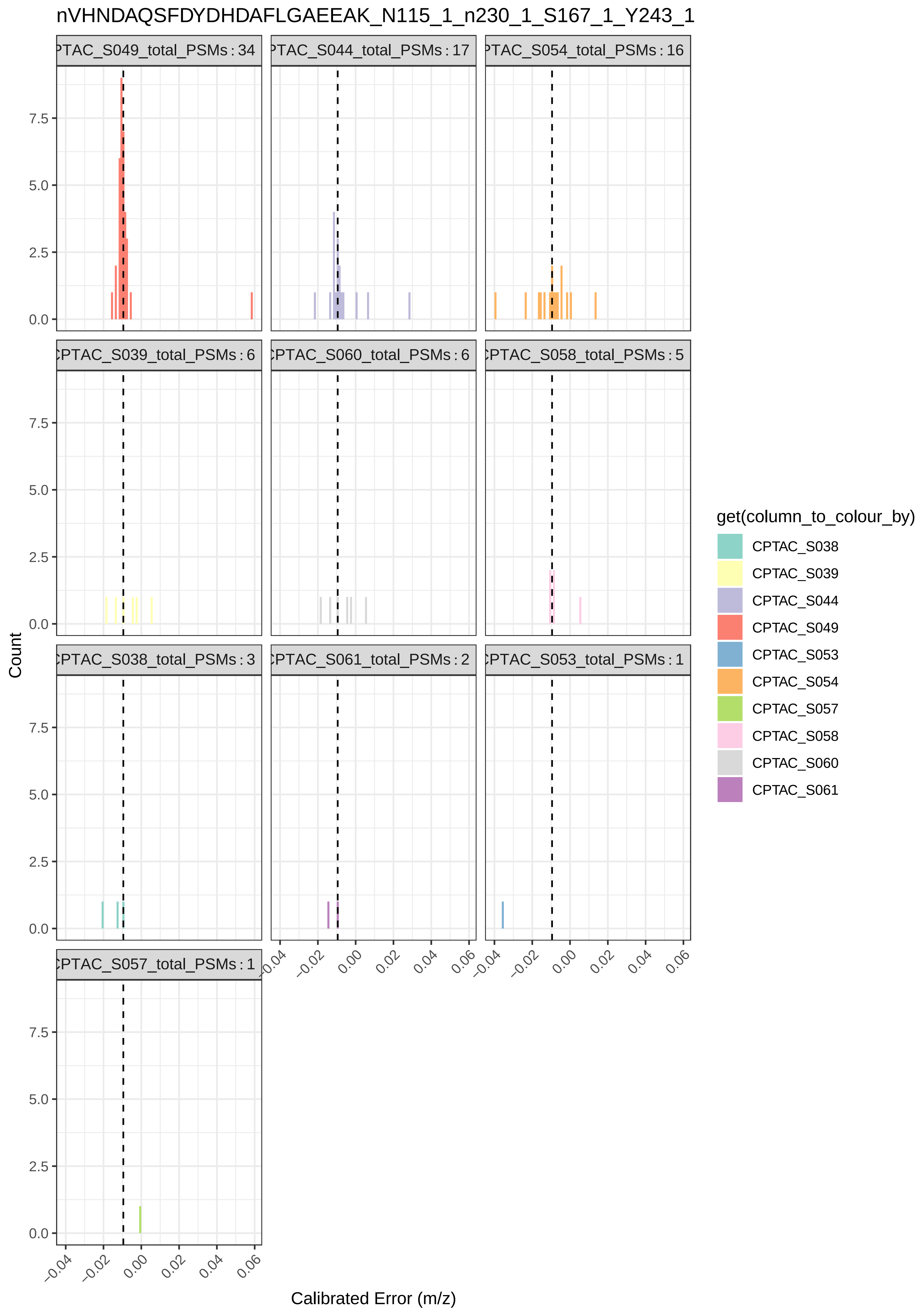

nVHNDAQSFDYDHDAFLGAEEAK_N115_1_n230_1_S167_1_Y243_1
PTAC_S049_total_PSMs : 34
PTAC_S044_total_PSMs : 17
PTAC_S054_total_PSMs : 16
CPTAC_S039_total_PSMs : 6
PTAC_S060_total_PSMs : 6
CPTAC_S058_total_PSMs : 5
CPTAC_S038_total_PSMs : 3
PTAC_S061_total_PSMs : 2
CPTAC_S053_total_PSMs : 1
CPTAC_S057_total_PSMs : 1
Count
0.0
2.5
5.0
7.5
−0.04
−0.02
0.00
0.02
0.04
0.06
Calibrated Error (m/z)
get(column_to_colour_by)
CPTAC_S038
CPTAC_S039
CPTAC_S044
CPTAC_S049
CPTAC_S053
CPTAC_S054
CPTAC_S057
CPTAC_S058
CPTAC_S060
CPTAC_S061

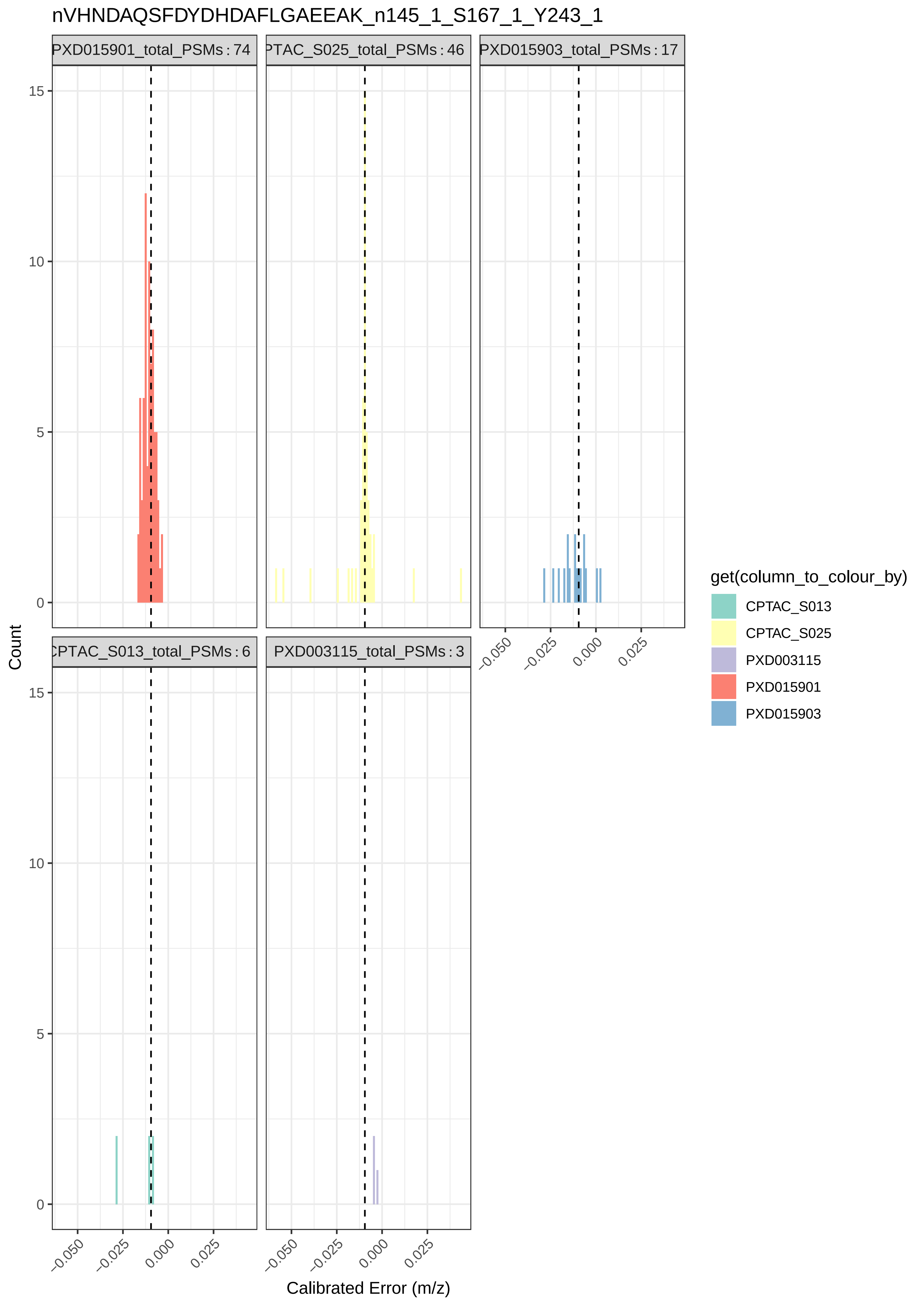

nVHNDAQSFDYDHDAFLGAEEAK_n145_1_S167_1_Y243_1
PXD015901_total_PSMs : 74
PTAC_S025_total_PSMs : 46
PXD015903_total_PSMs : 17
CPTAC_S013_total_PSMs : 6
PXD003115_total_PSMs : 3
Count
0
5
10
15
−0.050
−0.025
0.000
0.025
Calibrated Error (m/z)
get(column_to_colour_by)
CPTAC_S013
CPTAC_S025
PXD003115
PXD015901
PXD015903

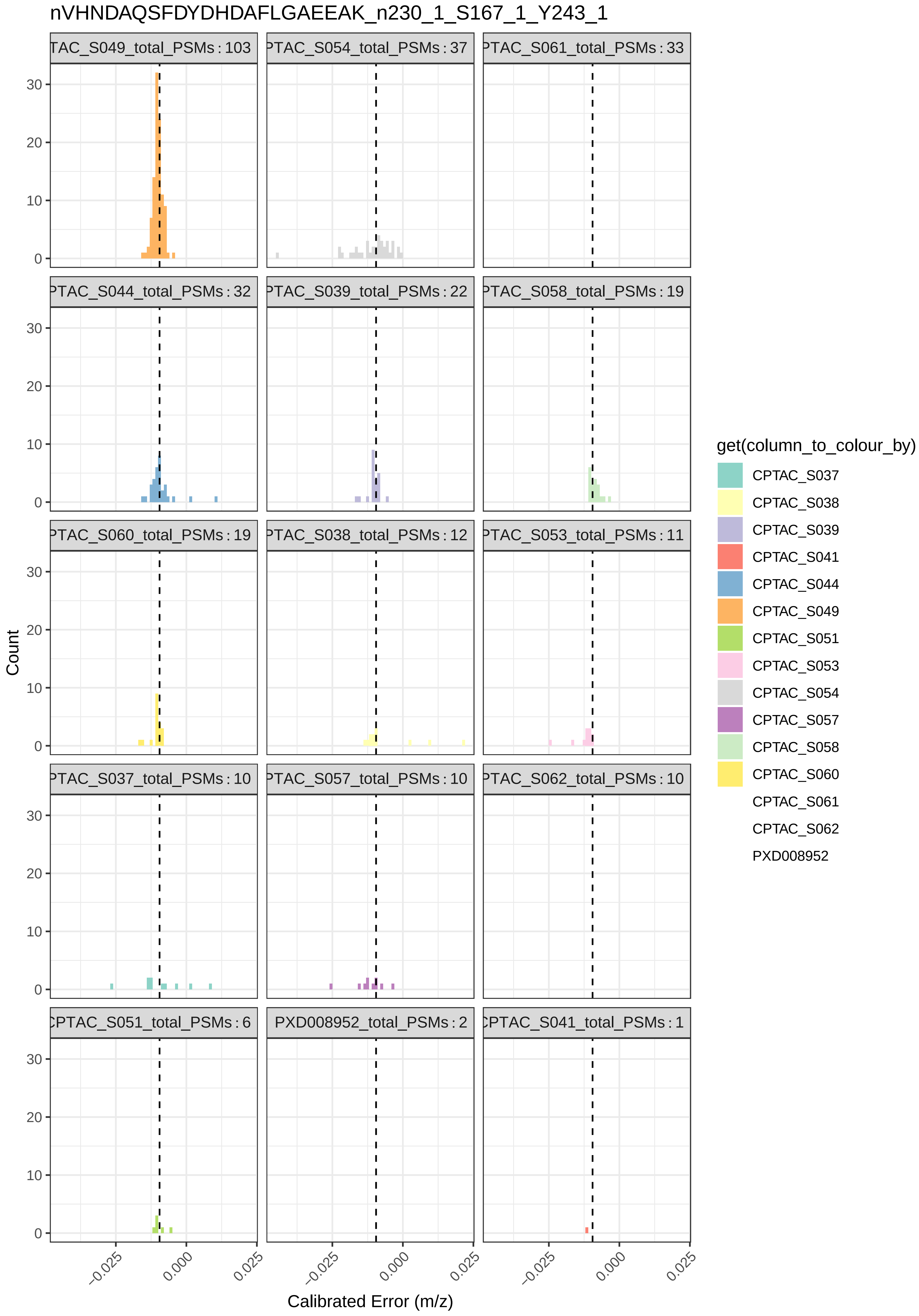

nVHNDAQSFDYDHDAFLGAEEAK_n230_1_S167_1_Y243_1
TAC_S049_total_PSMs : 103
PTAC_S054_total_PSMs : 37
PTAC_S061_total_PSMs : 33
PTAC_S044_total_PSMs : 32
PTAC_S039_total_PSMs : 22
PTAC_S058_total_PSMs : 19
PTAC_S060_total_PSMs : 19
PTAC_S038_total_PSMs : 12
PTAC_S053_total_PSMs : 11
PTAC_S037_total_PSMs : 10
PTAC_S057_total_PSMs : 10
PTAC_S062_total_PSMs : 10
CPTAC_S051_total_PSMs : 6
PXD008952_total_PSMs : 2
CPTAC_S041_total_PSMs : 1
Count
0
10
20
30
-0.025
0.000
0.025
Calibrated Error (m/z)
get(column_to_colour_by)
CPTAC_S037
CPTAC_S038
CPTAC_S039
CPTAC_S041
CPTAC_S044
CPTAC_S049
CPTAC_S051
CPTAC_S053
CPTAC_S054
CPTAC_S057
CPTAC_S058
CPTAC_S060
CPTAC_S061
CPTAC_S062
PXD008952

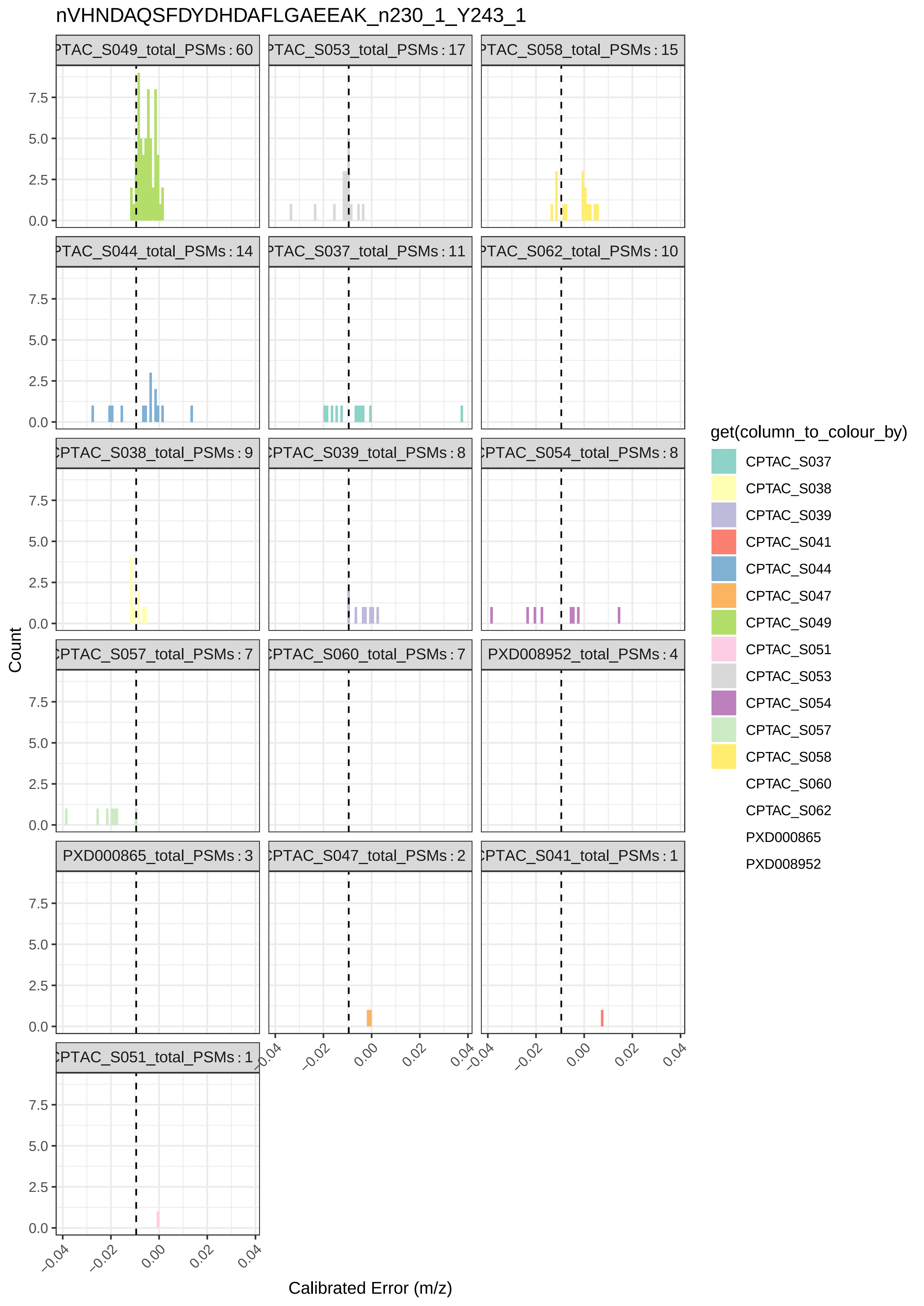
nVHNDAQSFDYDHDAFLGAEEAK_n230_1_Y243_1
PTAC_S049_total_PSMs : 60
PTAC_S053_total_PSMs : 17
PTAC_S058_total_PSMs : 15
PTAC_S044_total_PSMs : 14
PTAC_S037_total_PSMs : 11
PTAC_S062_total_PSMs : 10
CPTAC_S038_total_PSMs : 9
CPTAC_S039_total_PSMs : 8
CPTAC_S054_total_PSMs : 8
CPTAC_S057_total_PSMs : 7
CPTAC_S060_total_PSMs : 7
PXD008952_total_PSMs : 4
PXD000865_total_PSMs : 3
CPTAC_S047_total_PSMs : 2
CPTAC_S041_total_PSMs : 1
CPTAC_S051_total_PSMs : 1
0.0
2.5
5.0
7.5
−0.04
−0.02
0.00
0.02
0.04
Count
Calibrated Error (m/z)
get(column_to_colour_by)
CPTAC_S037
CPTAC_S038
CPTAC_S039
CPTAC_S041
CPTAC_S044
CPTAC_S047
CPTAC_S049
CPTAC_S051
CPTAC_S053
CPTAC_S054
CPTAC_S057
CPTAC_S058
CPTAC_S060
CPTAC_S062
PXD000865
PXD008952

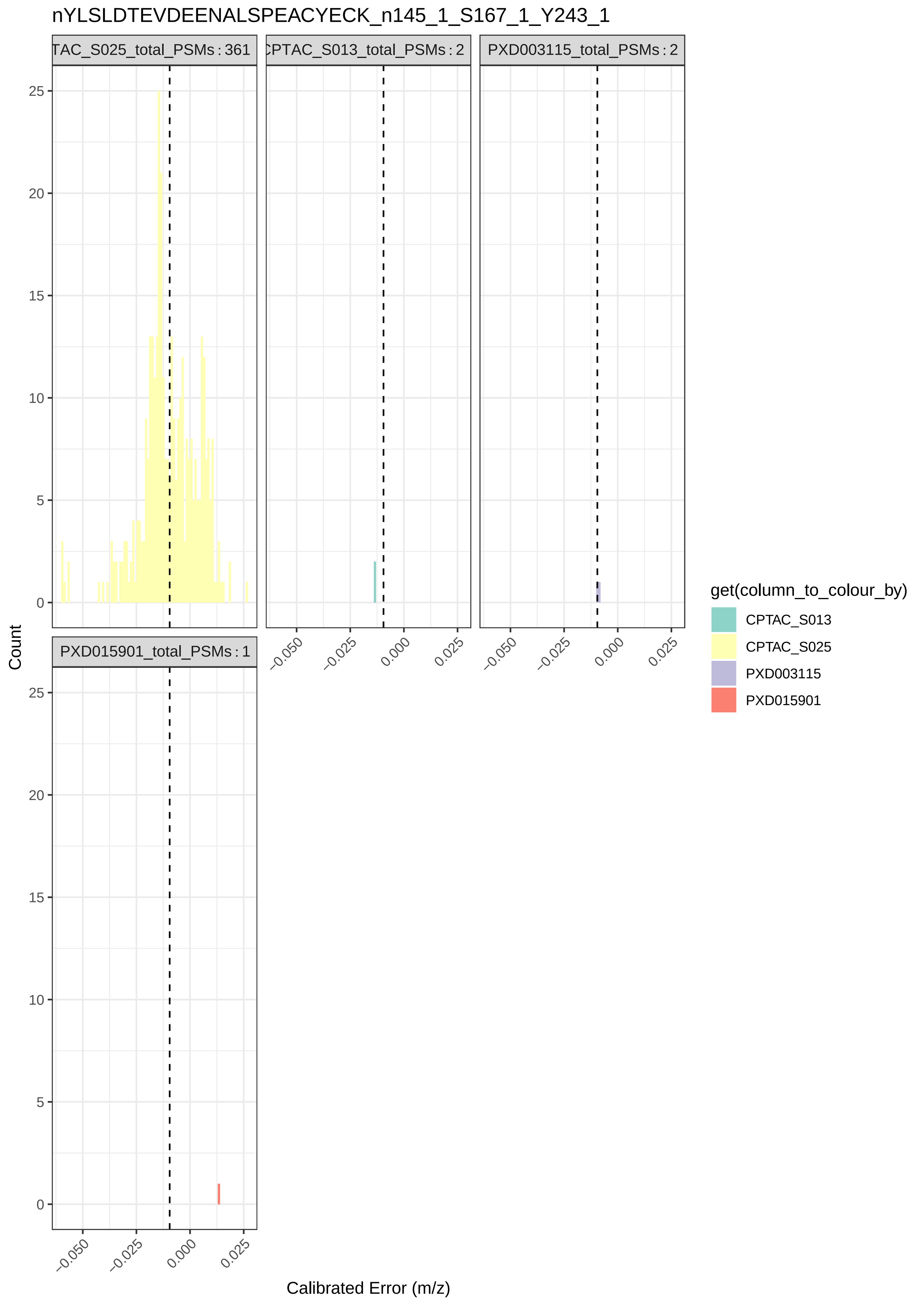
nYLSLDTEVDEENALSPEACYECK_n145_1_S167_1_Y243_1
TAC_S025_total_PSMs : 361
CPTAC_S013_total_PSMs : 2
PXD003115_total_PSMs : 2
PXD015901_total_PSMs : 1
Count
Calibrated Error (m/z)
0
5
10
15
20
25
−0.050
−0.025
0.000
0.025
get(column_to_colour_by)
CPTAC_S013
CPTAC_S025
PXD003115
PXD015901

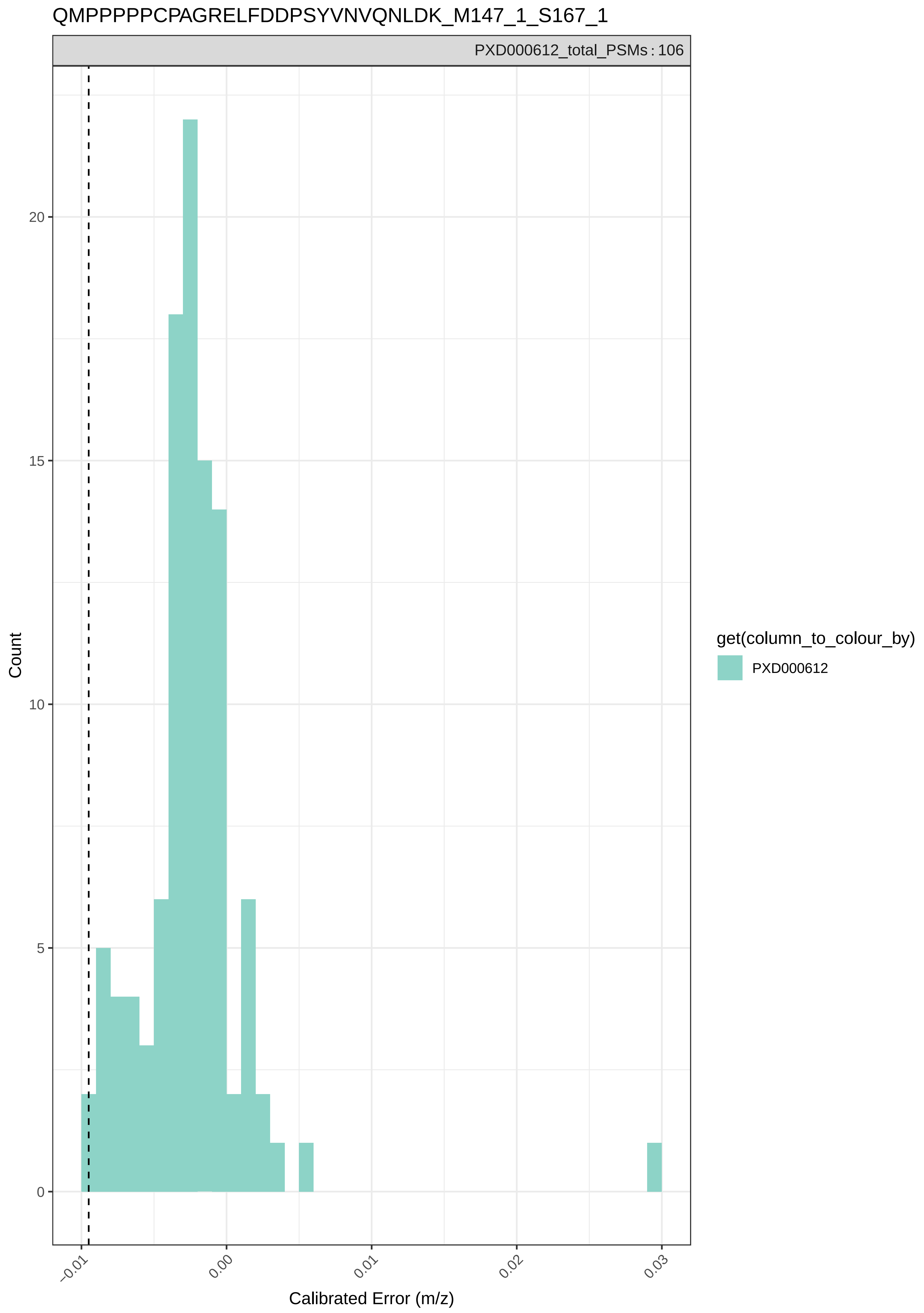

QMPPPPPCPAGRELFDDPSYVNVQNLDK_M147_1_S167_1
PXD000612_total_PSMs : 106
Count
20
15
10
5
0
−0.01
0.00
0.01
0.02
0.03
Calibrated Error (m/z)
get(column_to_colour_by)
PXD000612

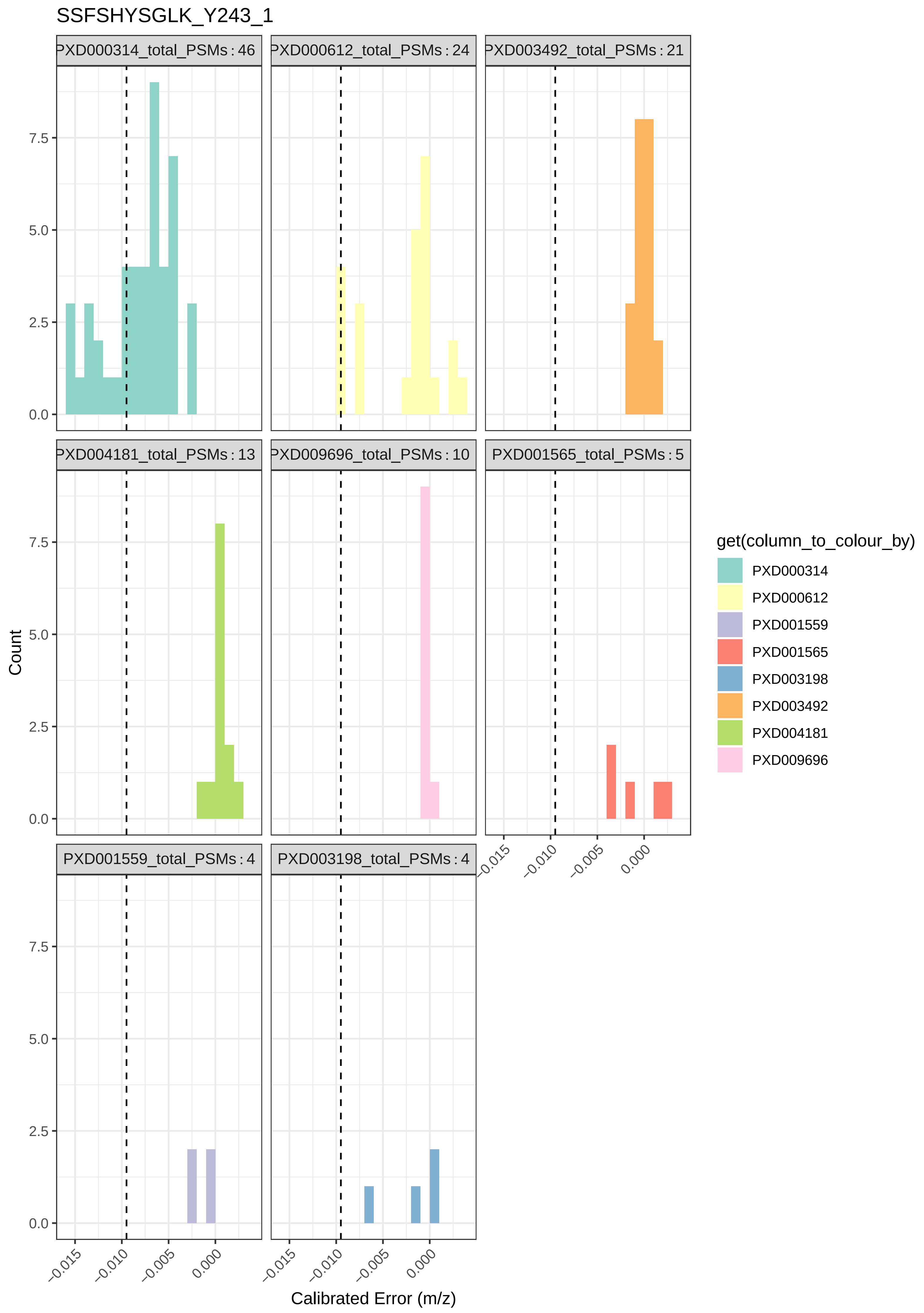
SSFSHYSGLK_Y243_1
PXD000314_total_PSMs : 46
PXD000612_total_PSMs : 24
PXD003492_total_PSMs : 21
PXD004181_total_PSMs : 13
PXD009696_total_PSMs : 10
PXD001565_total_PSMs : 5
PXD001559_total_PSMs : 4
PXD003198_total_PSMs : 4
Count
0.0
2.5
5.0
7.5
−0.015
−0.010
−0.005
0.000
Calibrated Error (m/z)
get(column_to_colour_by)
PXD000314
PXD000612
PXD001559
PXD001565
PXD003198
PXD003492
PXD004181
PXD009696

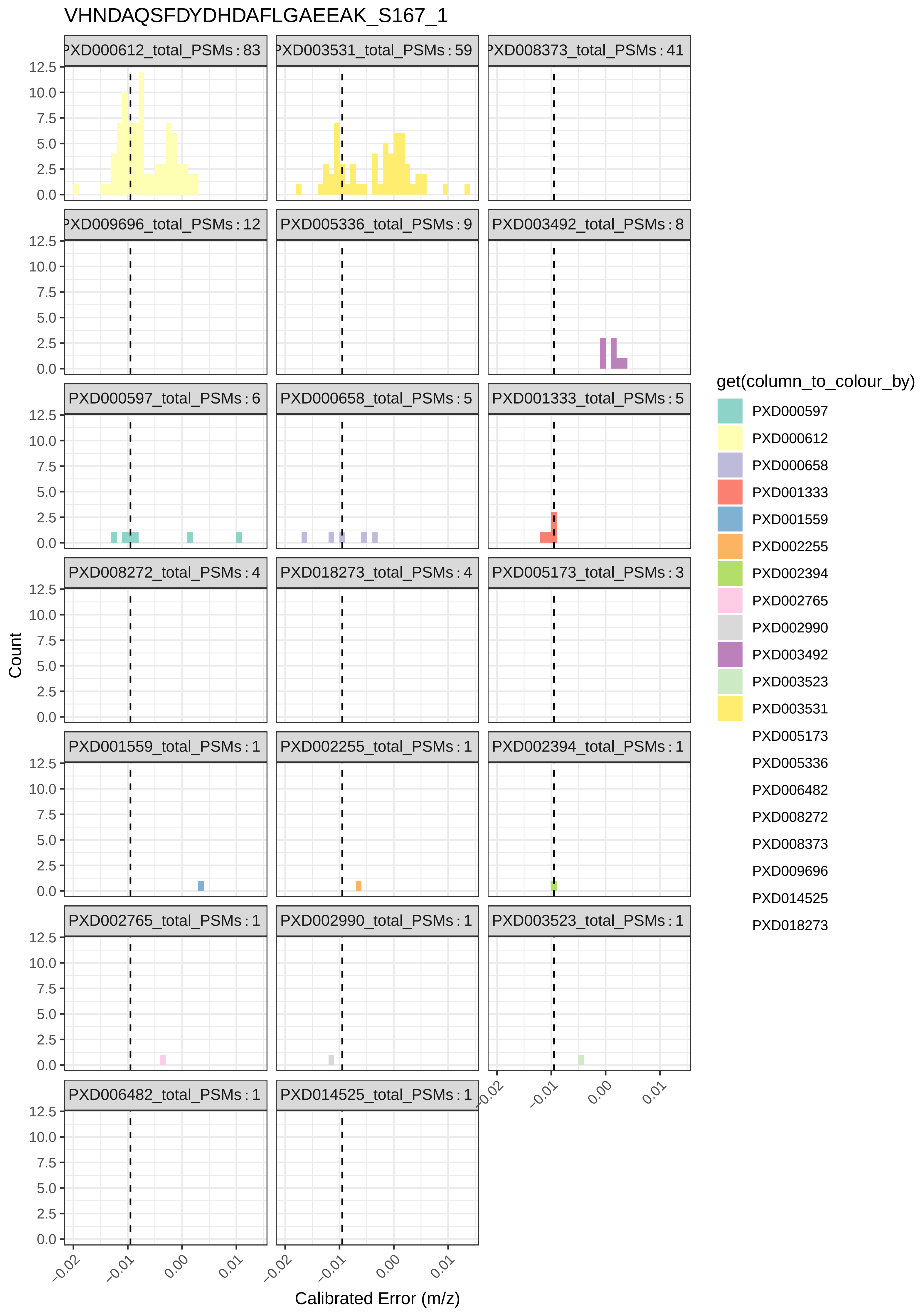
VHNDAQSFDYDHDAFLGAEEAK_S167_1
PXD000612_total_PSMs : 83
PXD003531_total_PSMs : 59
PXD008373_total_PSMs : 41
PXD009696_total_PSMs : 12
PXD005336_total_PSMs : 9
PXD003492_total_PSMs : 8
PXD000597_total_PSMs : 6
PXD000658_total_PSMs : 5
PXD001333_total_PSMs : 5
PXD008272_total_PSMs : 4
PXD018273_total_PSMs : 4
PXD005173_total_PSMs : 3
PXD001559_total_PSMs : 1
PXD002255_total_PSMs : 1
PXD002394_total_PSMs : 1
PXD002765_total_PSMs : 1
PXD002990_total_PSMs : 1
PXD003523_total_PSMs : 1
PXD006482_total_PSMs : 1
PXD014525_total_PSMs : 1
Count
0.0
2.5
5.0
7.5
10.0
12.5
−0.02
−0.01
0.00
0.01
Calibrated Error (m/z)
get(column_to_colour_by)
PXD000597
PXD000612
PXD000658
PXD001333
PXD001559
PXD002255
PXD002394
PXD002765
PXD002990
PXD003492
PXD003523
PXD003531
PXD005173
PXD005336
PXD006482
PXD008272
PXD008373
PXD009696
PXD014525
PXD018273

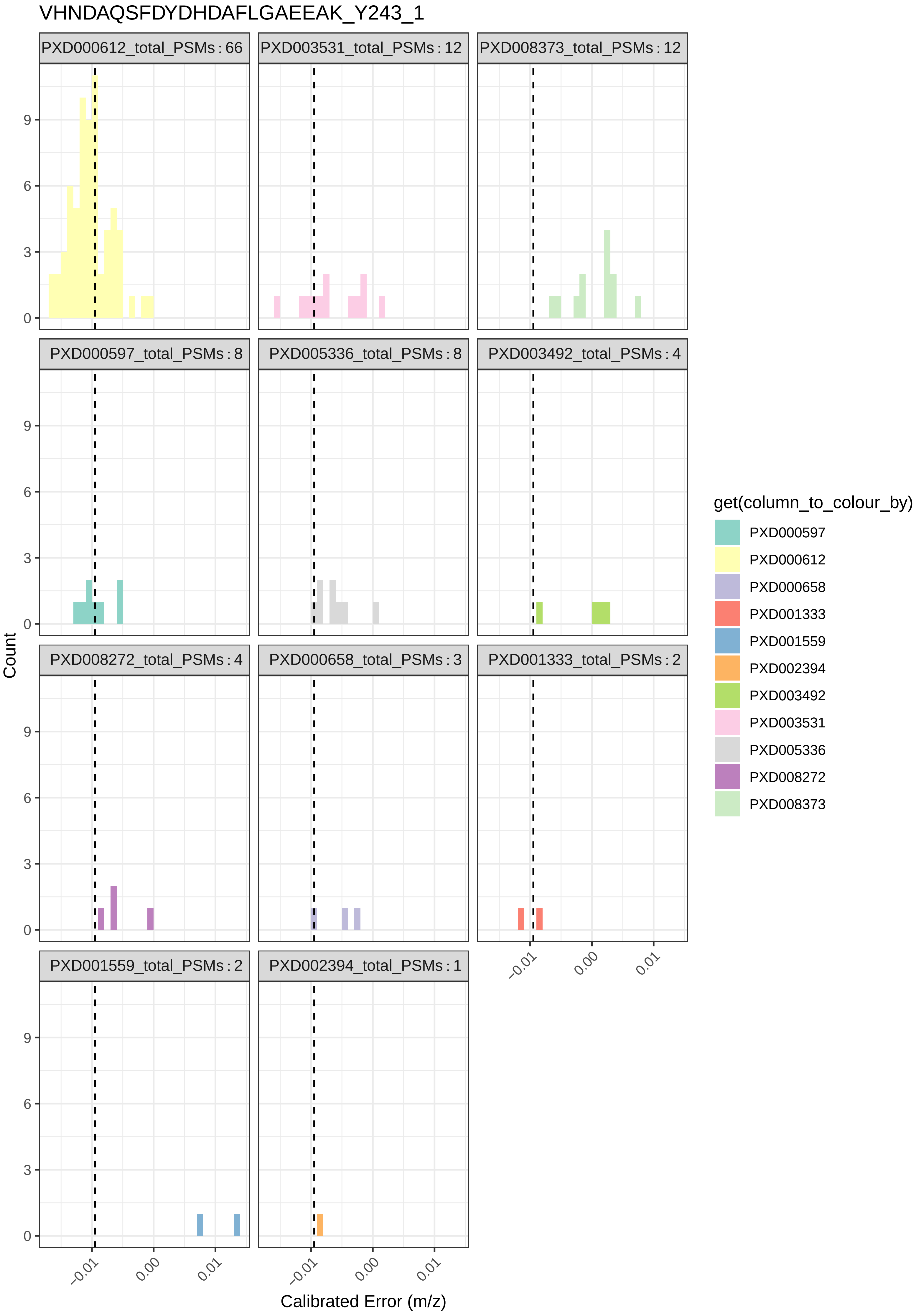
VHNDAQSFDYDHDAFLGAEEAK_Y243_1
PXD000612_total_PSMs : 66
PXD003531_total_PSMs : 12
PXD008373_total_PSMs : 12
PXD000597_total_PSMs : 8
PXD005336_total_PSMs : 8
PXD003492_total_PSMs : 4
PXD008272_total_PSMs : 4
PXD000658_total_PSMs : 3
PXD001333_total_PSMs : 2
PXD001559_total_PSMs : 2
PXD002394_total_PSMs : 1
Count
0
3
6
9
−0.01
0.00
0.01
Calibrated Error (m/z)
get(column_to_colour_by)
PXD000597
PXD000612
PXD000658
PXD001333
PXD001559
PXD002394
PXD003492
PXD003531
PXD005336
PXD008272
PXD008373
