## Supplementary File 2 for "Searching for Sulfotyrosines (sY) in a HA(pY)STACK": faceted_plots_by_experiment_tag.pdf

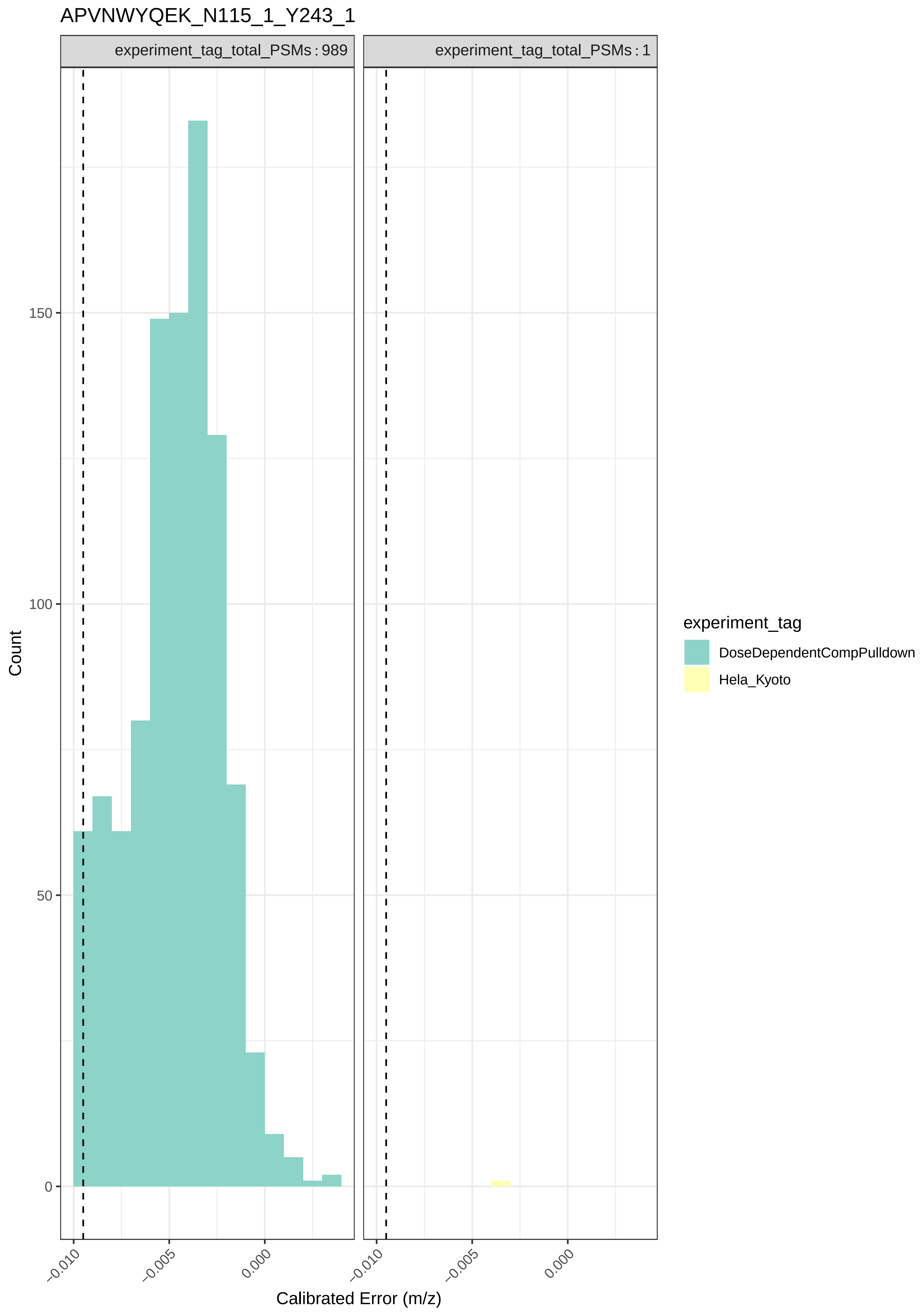

APVNWYQEK_N115_1_Y243_1
experiment_tag_total_PSMs : 989
experiment_tag_total_PSMs : 1
Count
150
100
50
0
−0.010
−0.005
0.000
−0.010
−0.005
0.000
Calibrated Error (m/z)
experiment_tag
DoseDependentCompPulldown
Hela_Kyoto

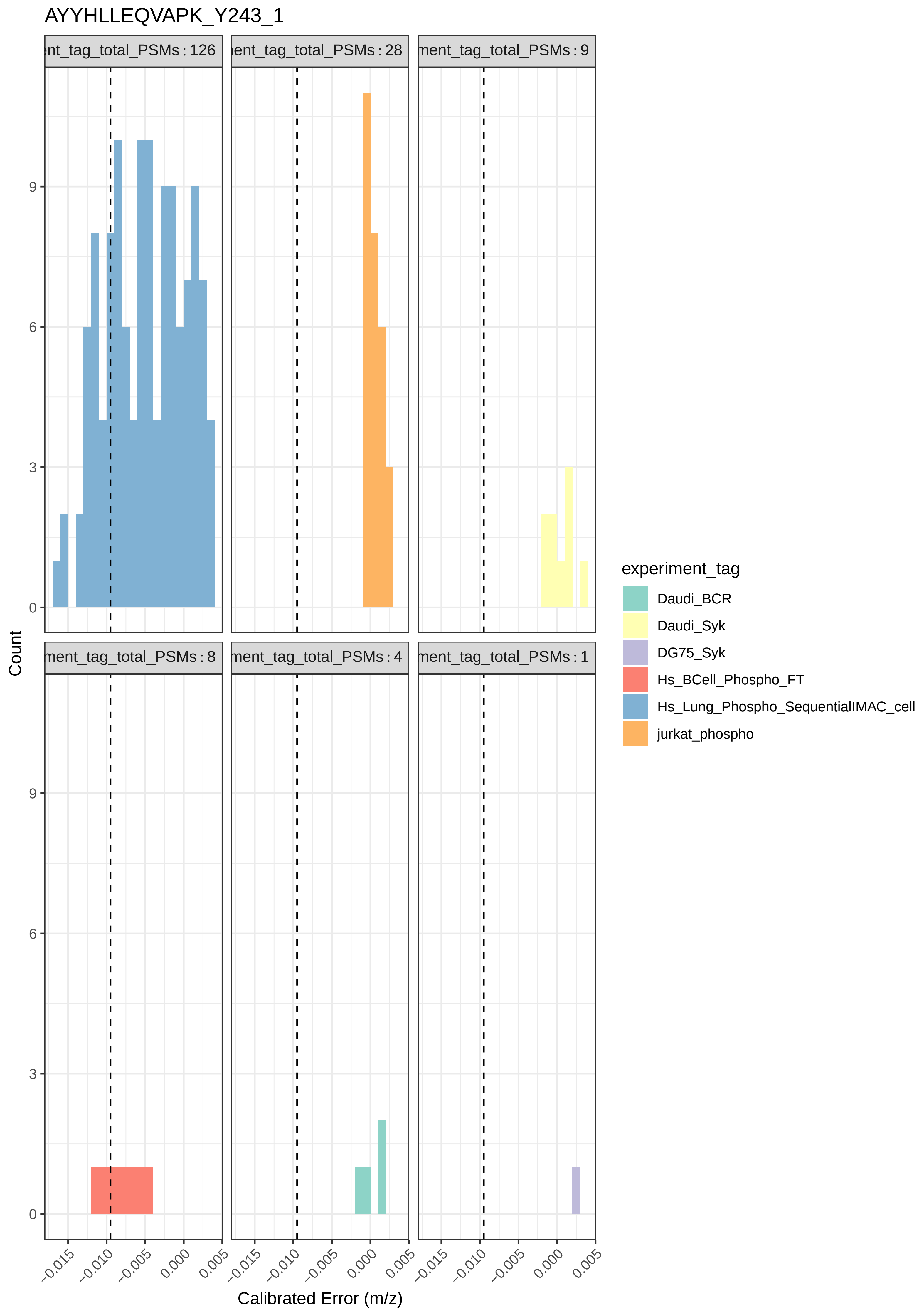

AYYHLLEQVAPK_Y243_1
ent_tag_total_PSMs : 126
ent_tag_total_PSMs : 28
ment_tag_total_PSMs : 9
ment_tag_total_PSMs : 8
ment_tag_total_PSMs : 4
ment_tag_total_PSMs : 1
Count
0
3
6
9
−0.015
−0.010
−0.005
0.000
0.005
Calibrated Error (m/z)
experiment_tag
Daudi_BCR
Daudi_Syk
DG75_Syk
Hs_BCell_Phospho_FT
Hs_Lung_Phospho_SequentialIMAC_cell
jurkat_phospho

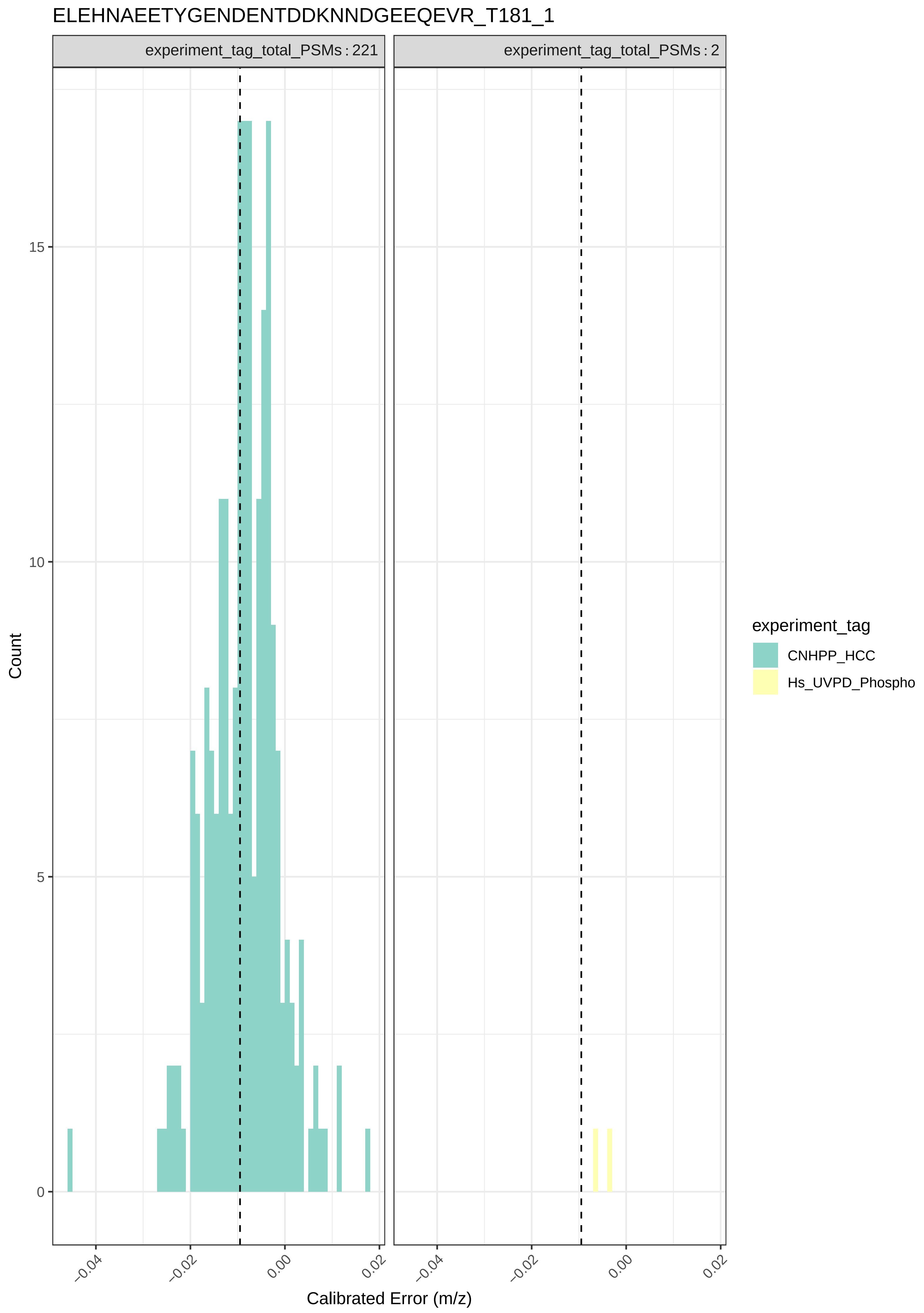

ELEHNAEETYGENDENTDDKNNDGEEQEVR_T181_1
experiment_tag_total_PSMs : 221
experiment_tag_total_PSMs : 2
Count
15
10
5
0
−0.04
−0.02
0.00
0.02
Calibrated Error (m/z)
experiment_tag
CNHPP_HCC
Hs_UVPD_Phospho

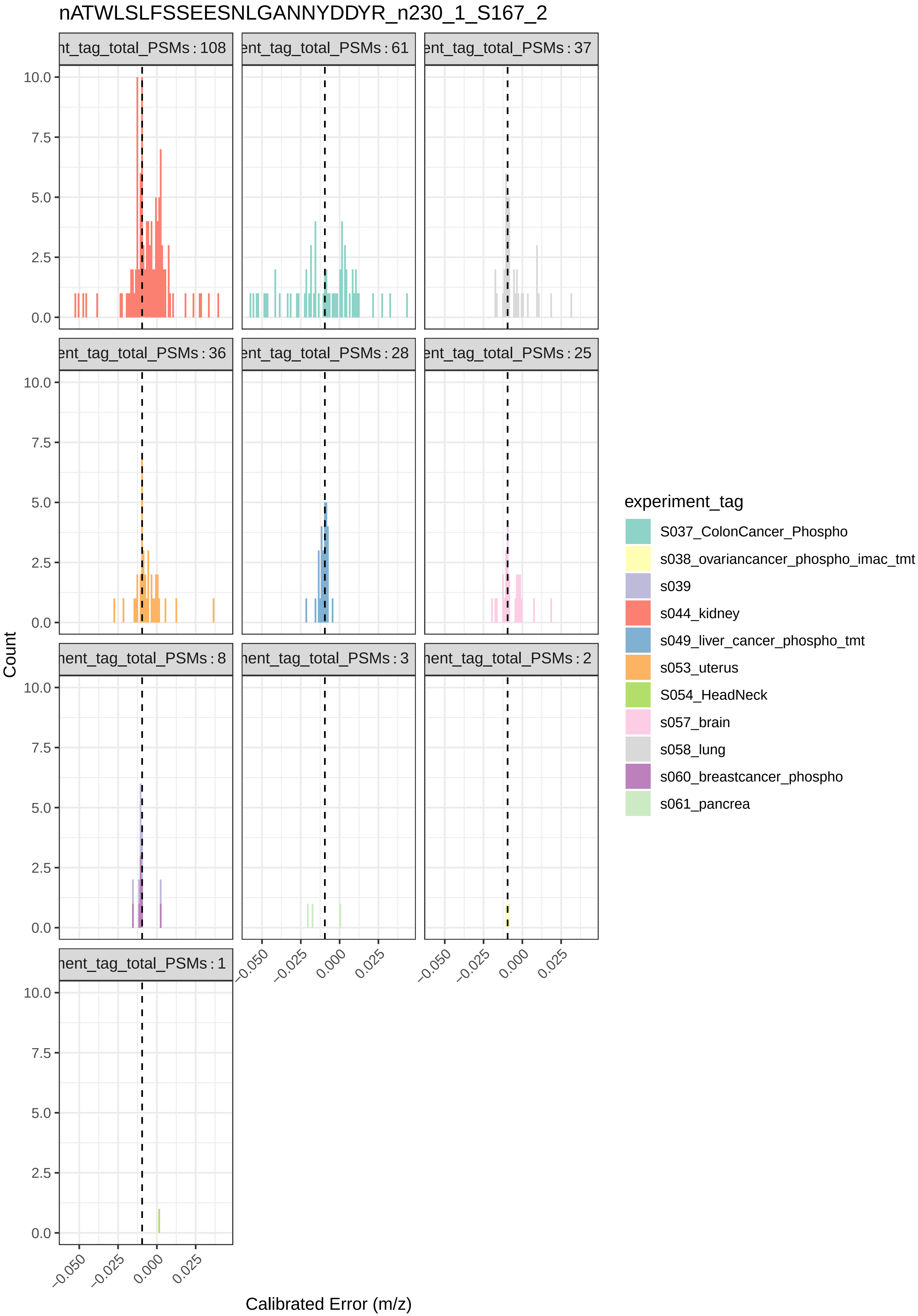
nATWLSLFSSEESNLGANNYDDYR_n230_1_S167_2
nt_tag_total_PSMs : 108
ent_tag_total_PSMs : 61
ent_tag_total_PSMs : 37
ent_tag_total_PSMs : 36
ent_tag_total_PSMs : 28
ent_tag_total_PSMs : 25
nent_tag_total_PSMs : 8
nent_tag_total_PSMs : 3
nent_tag_total_PSMs : 2
nent_tag_total_PSMs : 1
Count
Calibrated Error (m/z)
experiment_tag
S037_ColonCancer_Phospho
s038_ovariancancer_phospho_imac_tmt
s039
s044_kidney
s049_liver_cancer_phospho_tmt
s053_uterus
S054_HeadNeck
s057_brain
s058_lung
s060_breastcancer_phospho
s061_pancrea

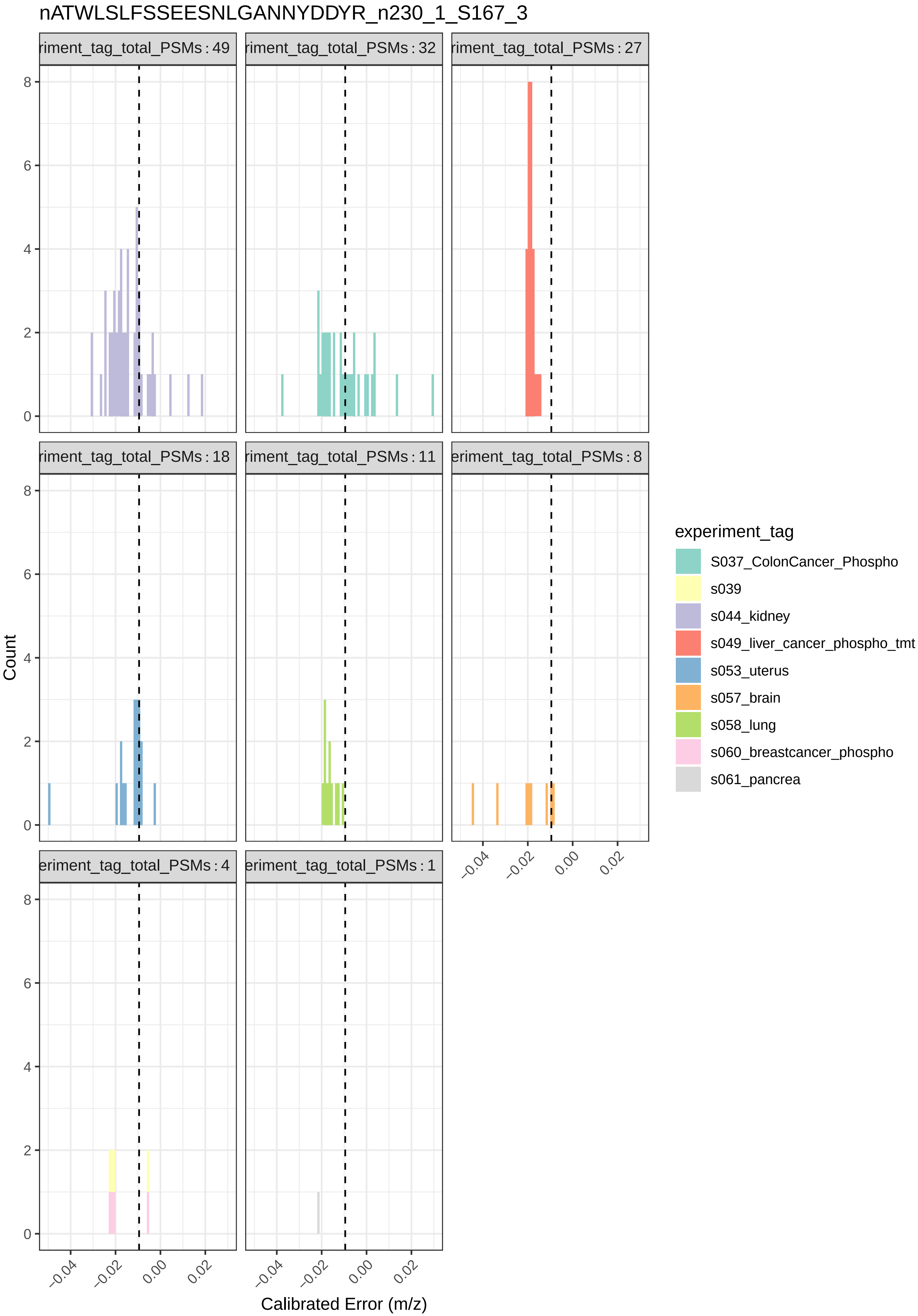
nATWLSLFSSEESNLGANNYDDYR_n230_1_S167_3
riment_tag_total_PSMs : 49
riment_tag_total_PSMs : 32
riment_tag_total_PSMs : 27
riment_tag_total_PSMs : 18
riment_tag_total_PSMs : 11
eriment_tag_total_PSMs : 8
eriment_tag_total_PSMs : 4
eriment_tag_total_PSMs : 1
Count
Calibrated Error (m/z)
experiment_tag
S037_ColonCancer_Phospho
s039
s044_kidney
s049_liver_cancer_phospho_tmt
s053_uterus
s057_brain
s058_lung
s060_breastcancer_phospho
s061_pancrea

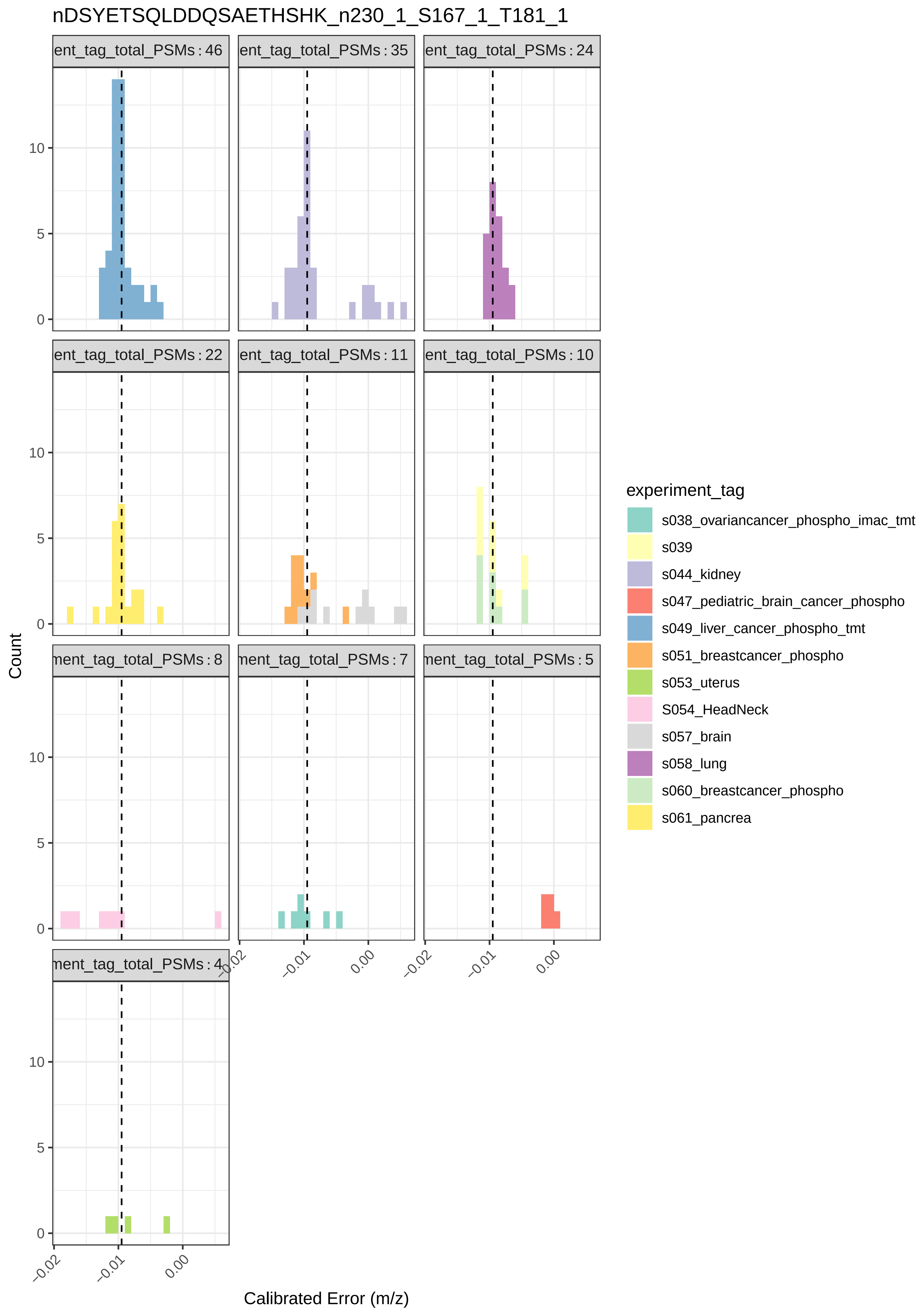

nDSYETSQLDDQSAETHSHK_n230_1_S167_1_T181_1
ent_tag_total_PSMs : 46
ent_tag_total_PSMs : 35
ent_tag_total_PSMs : 24
ent_tag_total_PSMs : 22
ent_tag_total_PSMs : 11
ent_tag_total_PSMs : 10
ment_tag_total_PSMs : 8
ment_tag_total_PSMs : 7
ment_tag_total_PSMs : 5
ment_tag_total_PSMs : 4
Count
0
5
10
−0.02
−0.01
0.00
Calibrated Error (m/z)
experiment_tag
s038_ovariancancer_phospho_imac_tmt
s039
s044_kidney
s047_pediatric_brain_cancer_phospho
s049_liver_cancer_phospho_tmt
s051_breastcancer_phospho
s053_uterus
S054_HeadNeck
s057_brain
s058_lung
s060_breastcancer_phospho
s061_pancrea

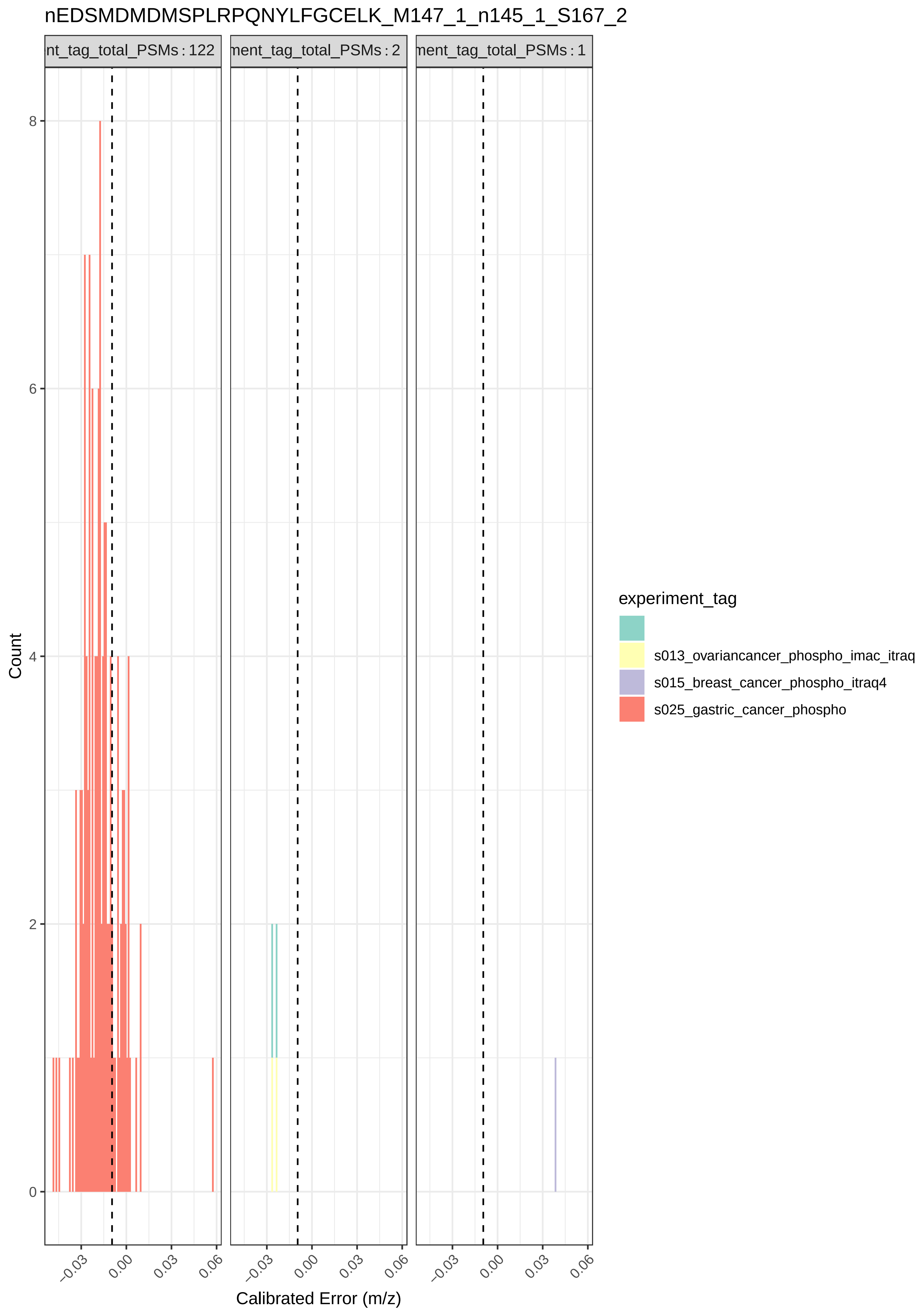
nEDSMDMDMSPLRPQNYLFGCELK_M147_1_n145_1_S167_2
nt_tag_total_PSMs : 122
ment_tag_total_PSMs : 2
ment_tag_total_PSMs : 1
Count
8
6
4
2
0
−0.03
0.00
0.03
0.06
Calibrated Error (m/z)
experiment_tag
s013_ovariancancer_phospho_imac_itraq
s015_breast_cancer_phospho_itraq4
s025_gastric_cancer_phospho

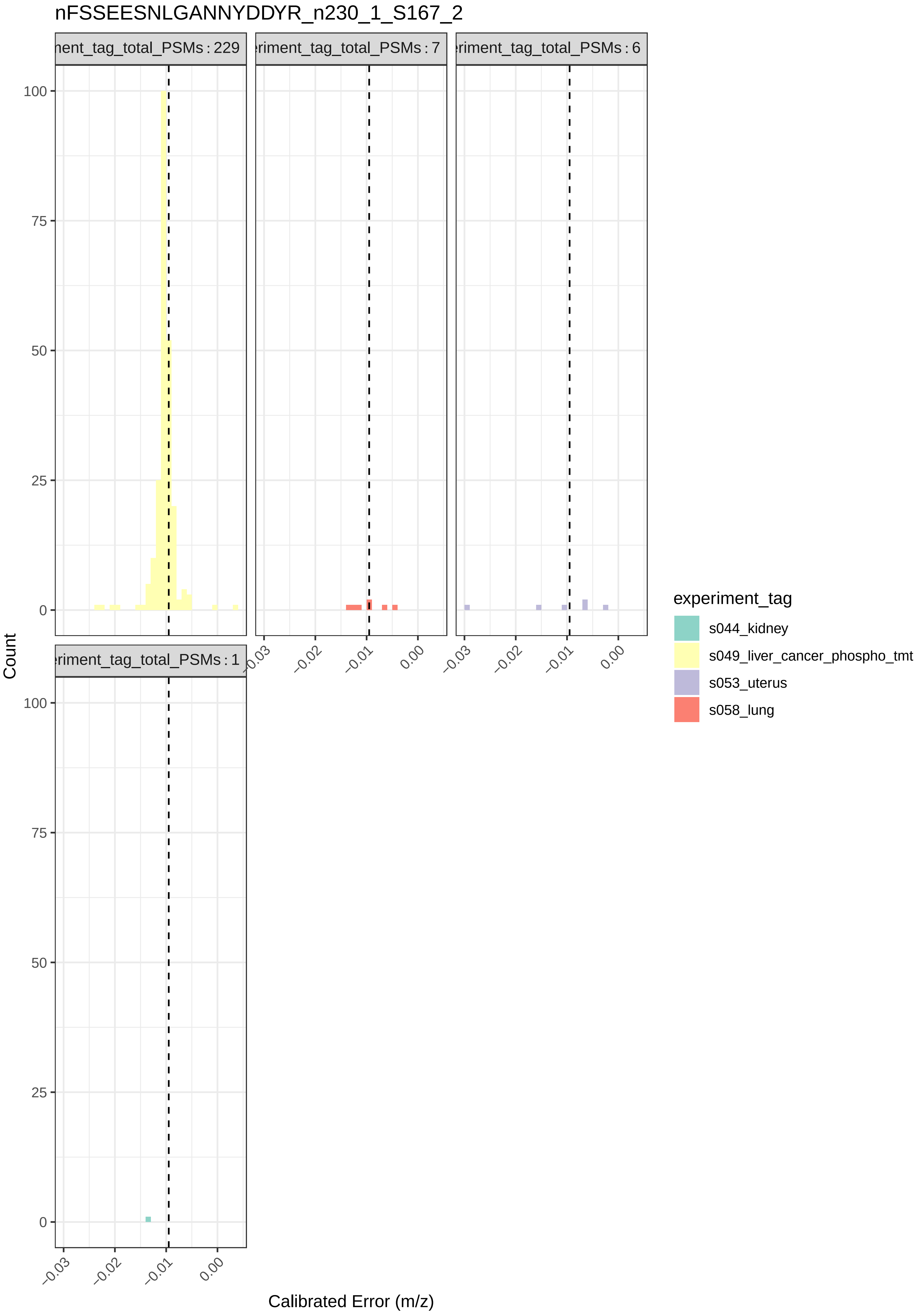

nFSSEESNLGANNYDDYR_n230_1_S167_2
nent_tag_total_PSMs : 229
eriment_tag_total_PSMs : 7
eriment_tag_total_PSMs : 6
eriment_tag_total_PSMs : 1
Count
100
75
50
25
0
−0.03
−0.02
−0.01
0.00
Calibrated Error (m/z)
experiment_tag
s044_kidney
s049_liver_cancer_phospho_tmt
s053_uterus
s058_lung

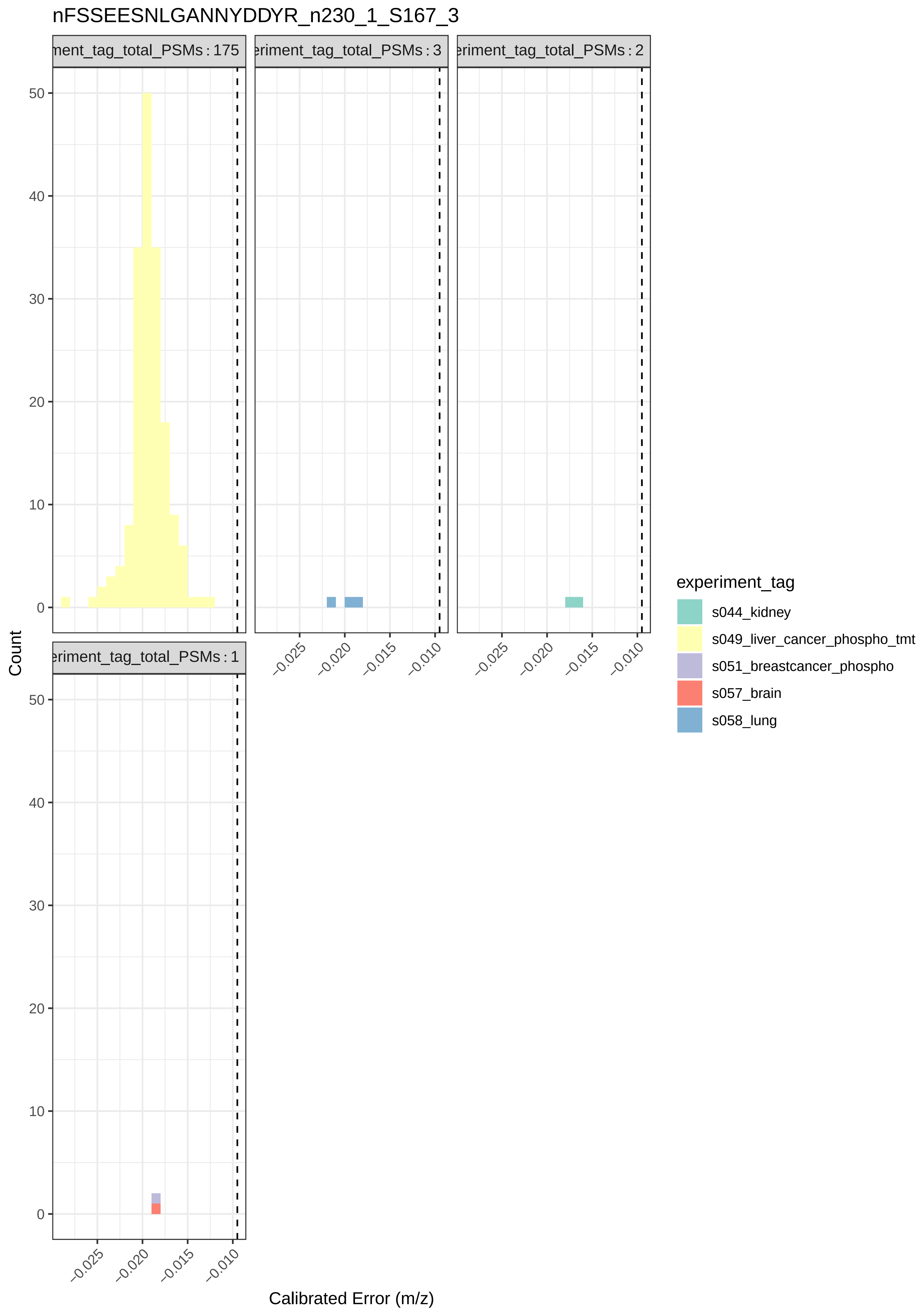

nFSSEESNLGANNYDDYR_n230_1_S167_3
ment_tag_total_PSMs : 175
eriment_tag_total_PSMs : 3
eriment_tag_total_PSMs : 2
eriment_tag_total_PSMs : 1
Count
Calibrated Error (m/z)
0
10
20
30
40
50
−0.025
−0.020
−0.015
−0.010
experiment_tag
s044_kidney
s049_liver_cancer_phospho_tmt
s051_breastcancer_phospho
s057_brain
s058_lung

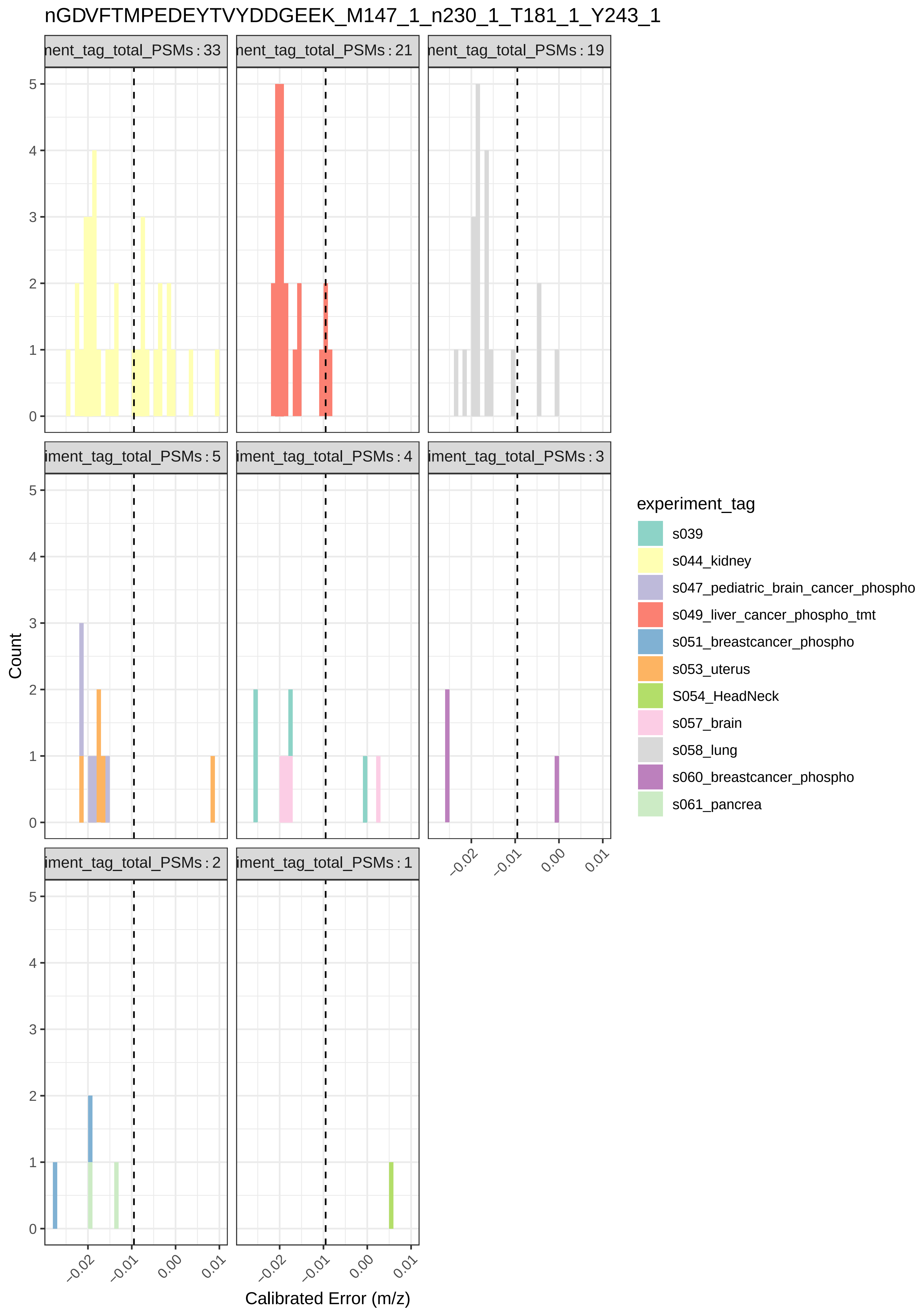

nGDVFTMPEDEYTVYDDGEEK_M147_1_n230_1_T181_1_Y243_1
nent_tag_total_PSMs : 33
nent_tag_total_PSMs : 21
nent_tag_total_PSMs : 19
iment_tag_total_PSMs : 5
iment_tag_total_PSMs : 4
iment_tag_total_PSMs : 3
iment_tag_total_PSMs : 2
iment_tag_total_PSMs : 1
Count
Calibrated Error (m/z)
−0.02
−0.01
0.00
0.01
experiment_tag
s039
s044_kidney
s047_pediatric_brain_cancer_phospho
s049_liver_cancer_phospho_tmt
s051_breastcancer_phospho
s053_uterus
S054_HeadNeck
s057_brain
s058_lung
s060_breastcancer_phospho
s061_pancrea

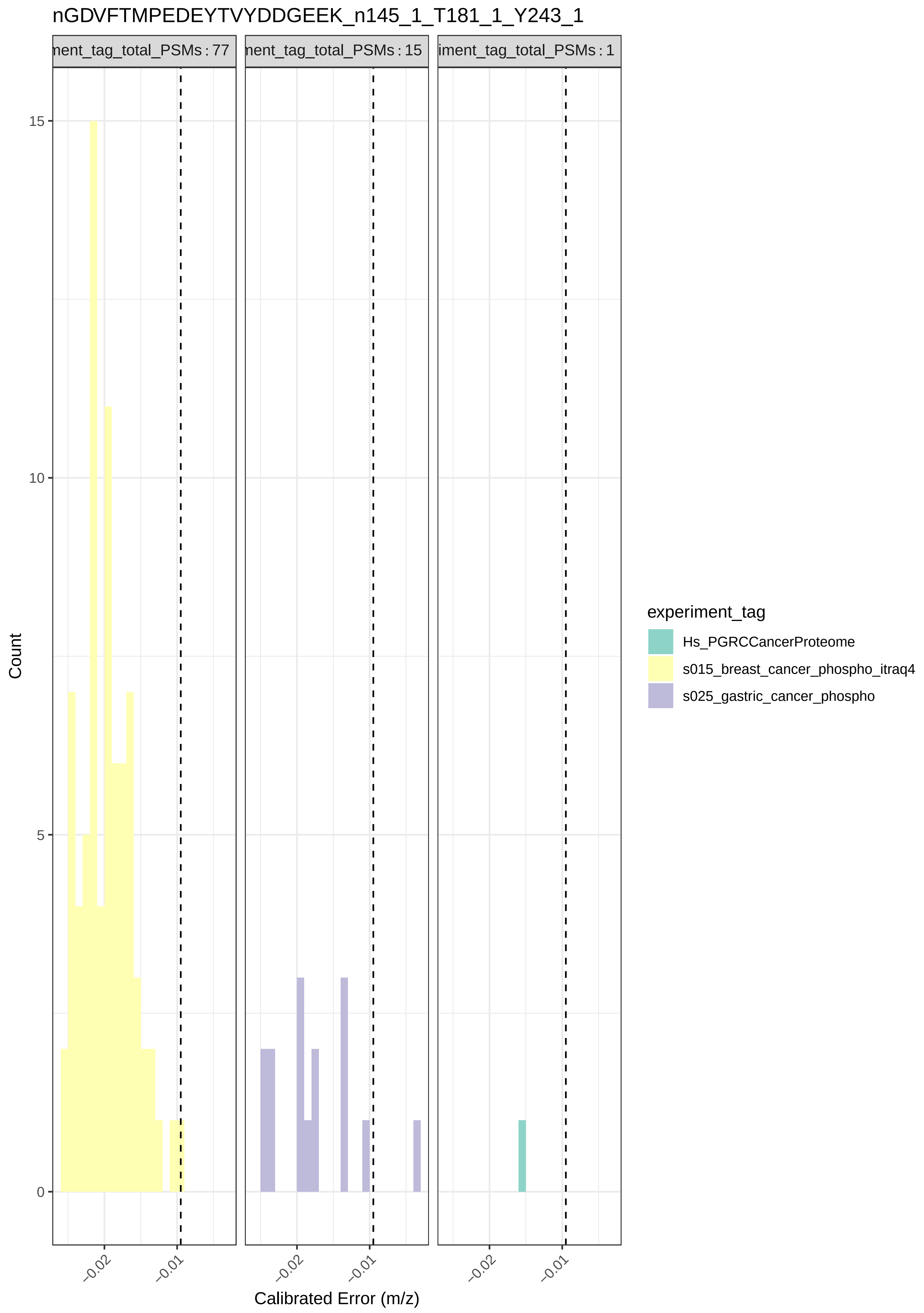

nGDVFTMPEDEYTVYDDGEEK_n145_1_T181_1_Y243_1
nent_tag_total_PSMs : 77
nent_tag_total_PSMs : 15
iment_tag_total_PSMs : 1
Count
15
10
5
0
−0.02
−0.01
Calibrated Error (m/z)
experiment_tag
Hs_PGRCCancerProteome
s015_breast_cancer_phospho_itraq4
s025_gastric_cancer_phospho

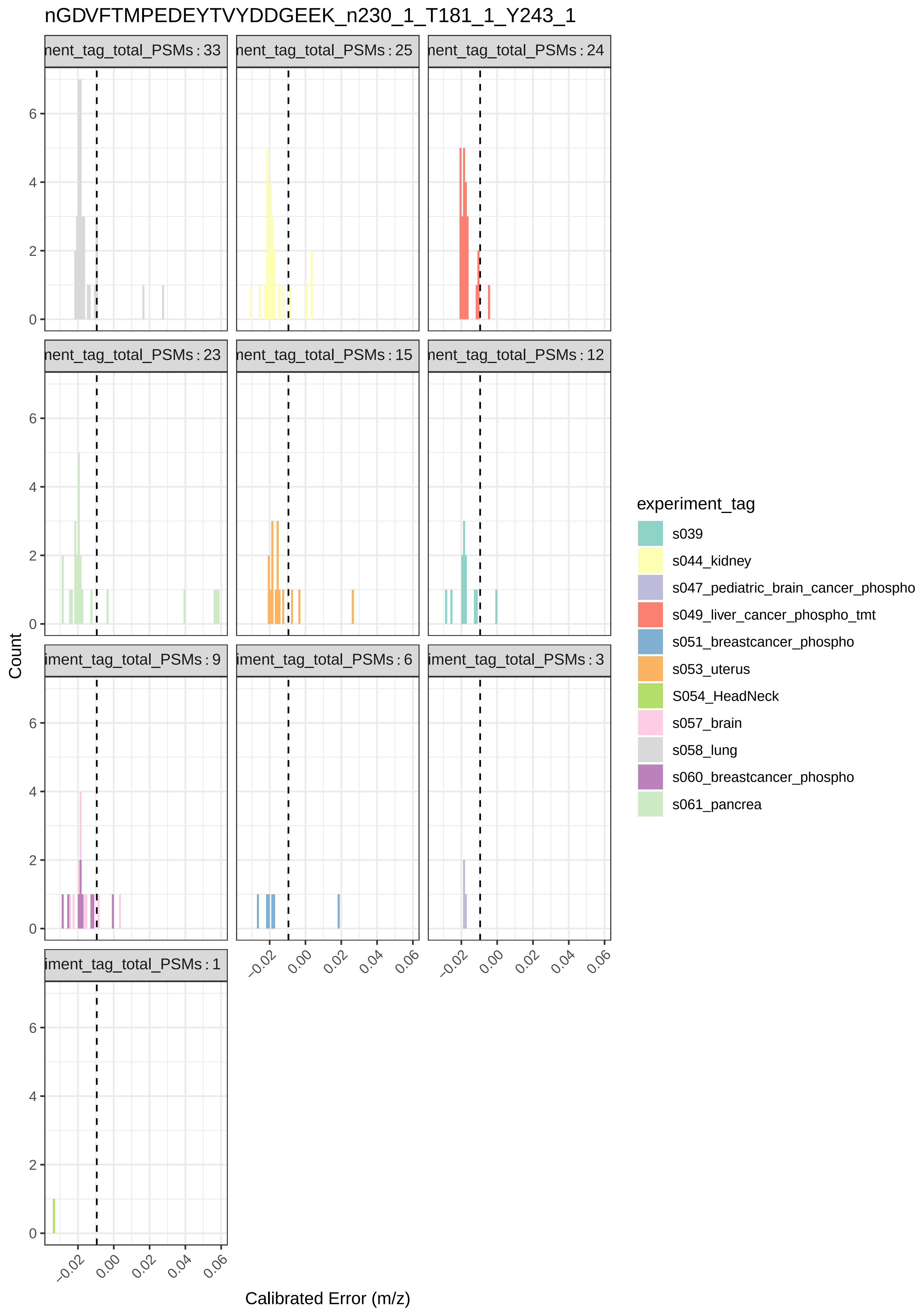
nGDVFTMPEDEYTVYDDGEEK_n230_1_T181_1_Y243_1
nent_tag_total_PSMs : 33
nent_tag_total_PSMs : 25
nent_tag_total_PSMs : 24
nent_tag_total_PSMs : 23
nent_tag_total_PSMs : 15
nent_tag_total_PSMs : 12
iment_tag_total_PSMs : 9
iment_tag_total_PSMs : 6
iment_tag_total_PSMs : 3
iment_tag_total_PSMs : 1
Count
Calibrated Error (m/z)
−0.02
0.00
0.02
0.04
0.06
0
2
4
6
experiment_tag
s039
s044_kidney
s047_pediatric_brain_cancer_phospho
s049_liver_cancer_phospho_tmt
s051_breastcancer_phospho
s053_uterus
S054_HeadNeck
s057_brain
s058_lung
s060_breastcancer_phospho
s061_pancrea

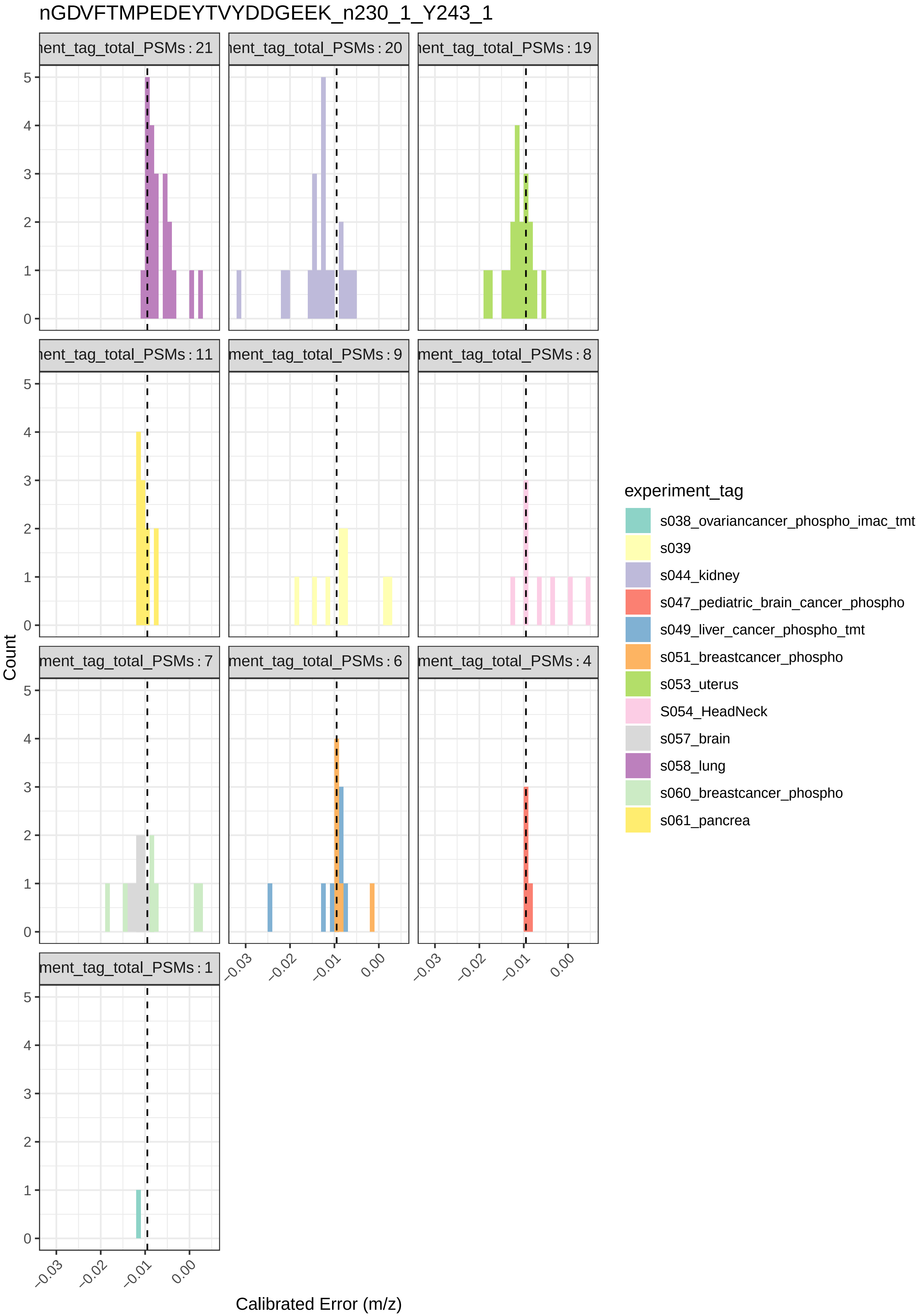

nGDVFTMPEDEYTVYDDGEEK_n230_1_Y243_1
ment_tag_total_PSMs : 21
ment_tag_total_PSMs : 20
ment_tag_total_PSMs : 19
ment_tag_total_PSMs : 11
ment_tag_total_PSMs : 9
ment_tag_total_PSMs : 8
ment_tag_total_PSMs : 7
ment_tag_total_PSMs : 6
ment_tag_total_PSMs : 4
ment_tag_total_PSMs : 1
Count
0
1
2
3
4
5
−0.03
−0.02
−0.01
0.00
Calibrated Error (m/z)
experiment_tag
s038_ovariancancer_phospho_imac_tmt
s039
s044_kidney
s047_pediatric_brain_cancer_phospho
s049_liver_cancer_phospho_tmt
s051_breastcancer_phospho
s053_uterus
S054_HeadNeck
s057_brain
s058_lung
s060_breastcancer_phospho
s061_pancrea

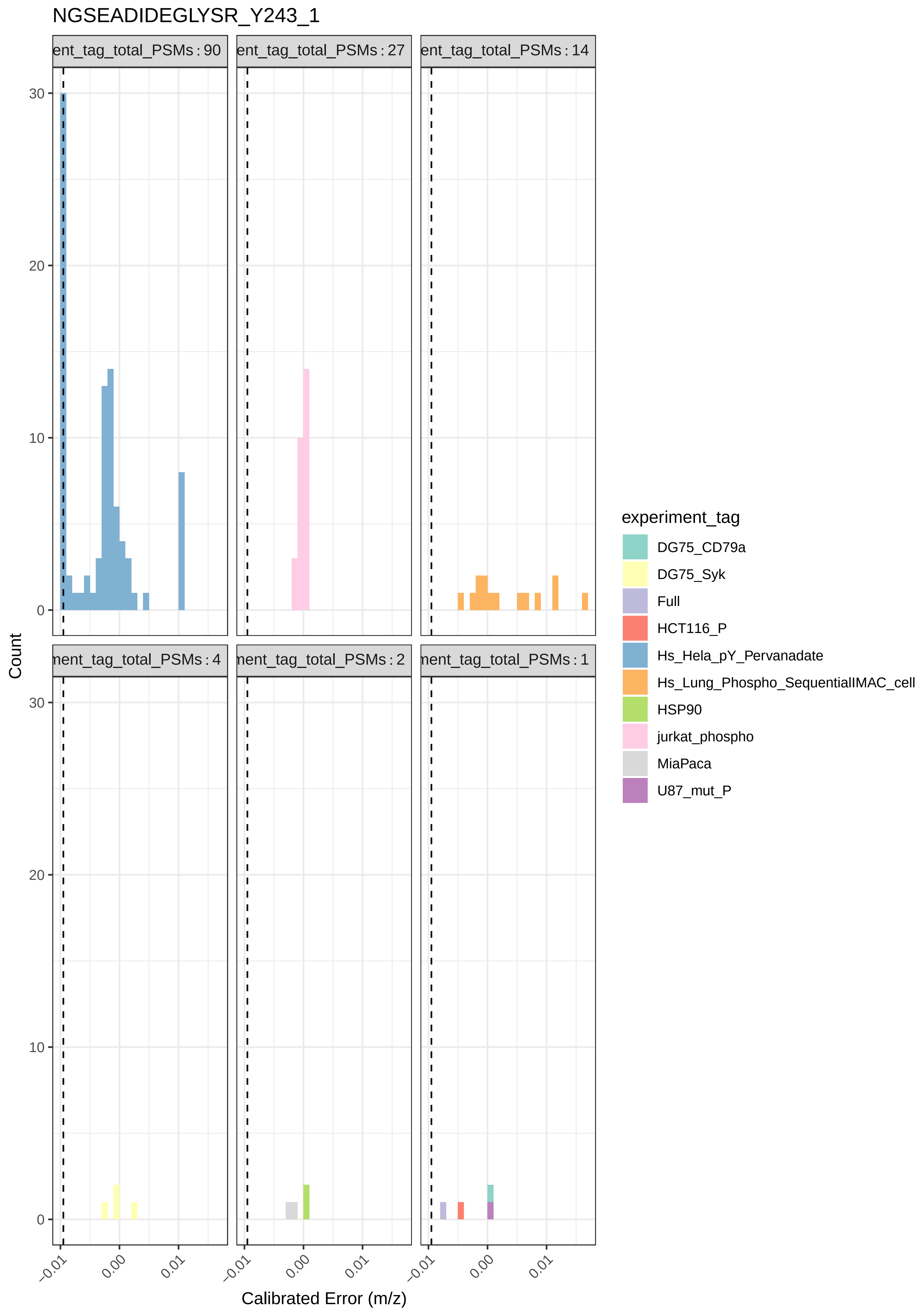
NGSEADIDEGLYSR_Y243_1
ent_tag_total_PSMs : 90
ent_tag_total_PSMs : 27
ent_tag_total_PSMs : 14
ment_tag_total_PSMs : 4
ment_tag_total_PSMs : 2
ment_tag_total_PSMs : 1
Count
0
10
20
30
−0.01
0.00
0.01
Calibrated Error (m/z)
experiment_tag
DG75_CD79a
DG75_Syk
Full
HCT116_P
Hs_Hela_pY_Pervanadate
Hs_Lung_Phospho_SequentialIMAC_cell
HSP90
jurkat_phospho
MiaPaca
U87_mut_P

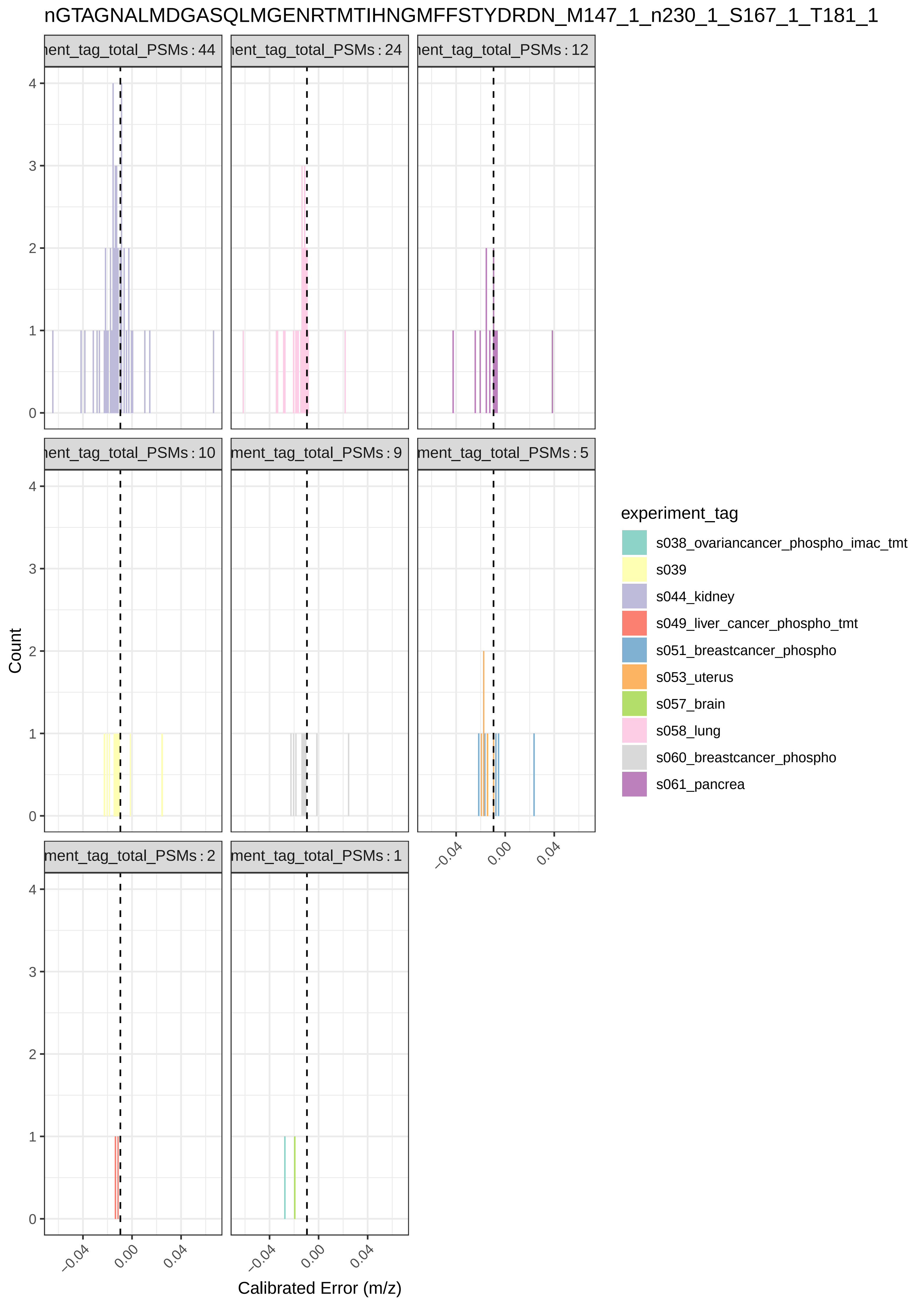

nGTAGNALMDGASQLMGENRTMTIHNGMFFSTYDRDN_M147_1_n230_1_S167_1_T181_1
ment_tag_total_PSMs : 44
ment_tag_total_PSMs : 24
ment_tag_total_PSMs : 12
ment_tag_total_PSMs : 10
ment_tag_total_PSMs : 9
ment_tag_total_PSMs : 5
ment_tag_total_PSMs : 2
ment_tag_total_PSMs : 1
Count
Calibrated Error (m/z)
−0.04
0.00
0.04
experiment_tag
s038_ovariancancer_phospho_imac_tmt
s039
s044_kidney
s049_liver_cancer_phospho_tmt
s051_breastcancer_phospho
s053_uterus
s057_brain
s058_lung
s060_breastcancer_phospho
s061_pancrea

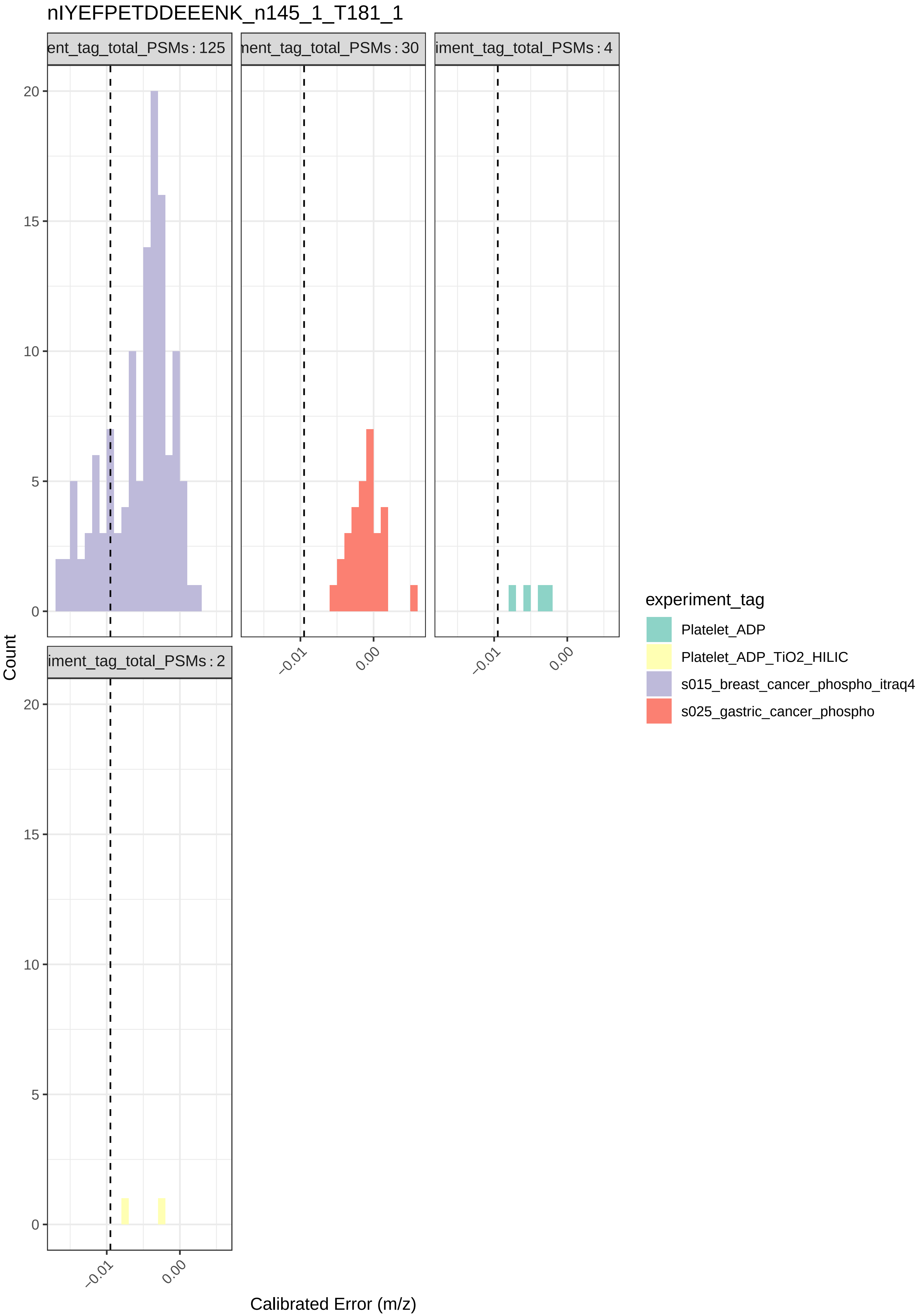

nIYEFPETDDEEENK_n145_1_T181_1
ent_tag_total_PSMs : 125
nent_tag_total_PSMs : 30
iment_tag_total_PSMs : 4
iment_tag_total_PSMs : 2
20
15
10
5
0
Count
−0.01
0.00
Calibrated Error (m/z)
experiment_tag
Platelet_ADP
Platelet_ADP_TiO2_HILIC
s015_breast_cancer_phospho_itraq4
s025_gastric_cancer_phospho

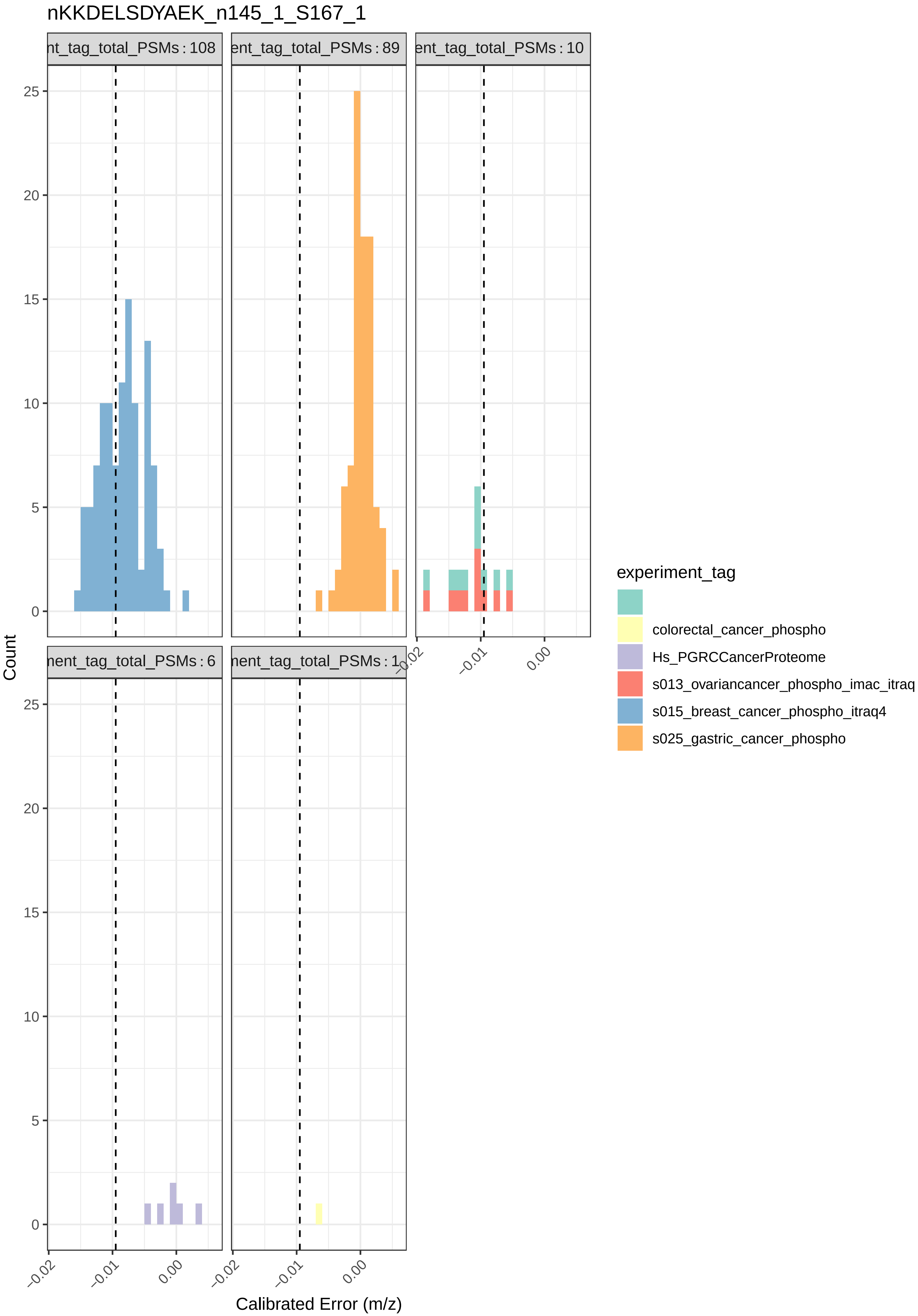

nKKDELSDYAEK_n145_1_S167_1
nt_tag_total_PSMs : 108
ent_tag_total_PSMs : 89
ent_tag_total_PSMs : 10
nent_tag_total_PSMs : 6
nent_tag_total_PSMs : 1
Count
Calibrated Error (m/z)
experiment_tag
colorectal_cancer_phospho
Hs_PGRCCancerProteome
s013_ovariancancer_phospho_imac_itraq
s015_breast_cancer_phospho_itraq4
s025_gastric_cancer_phospho

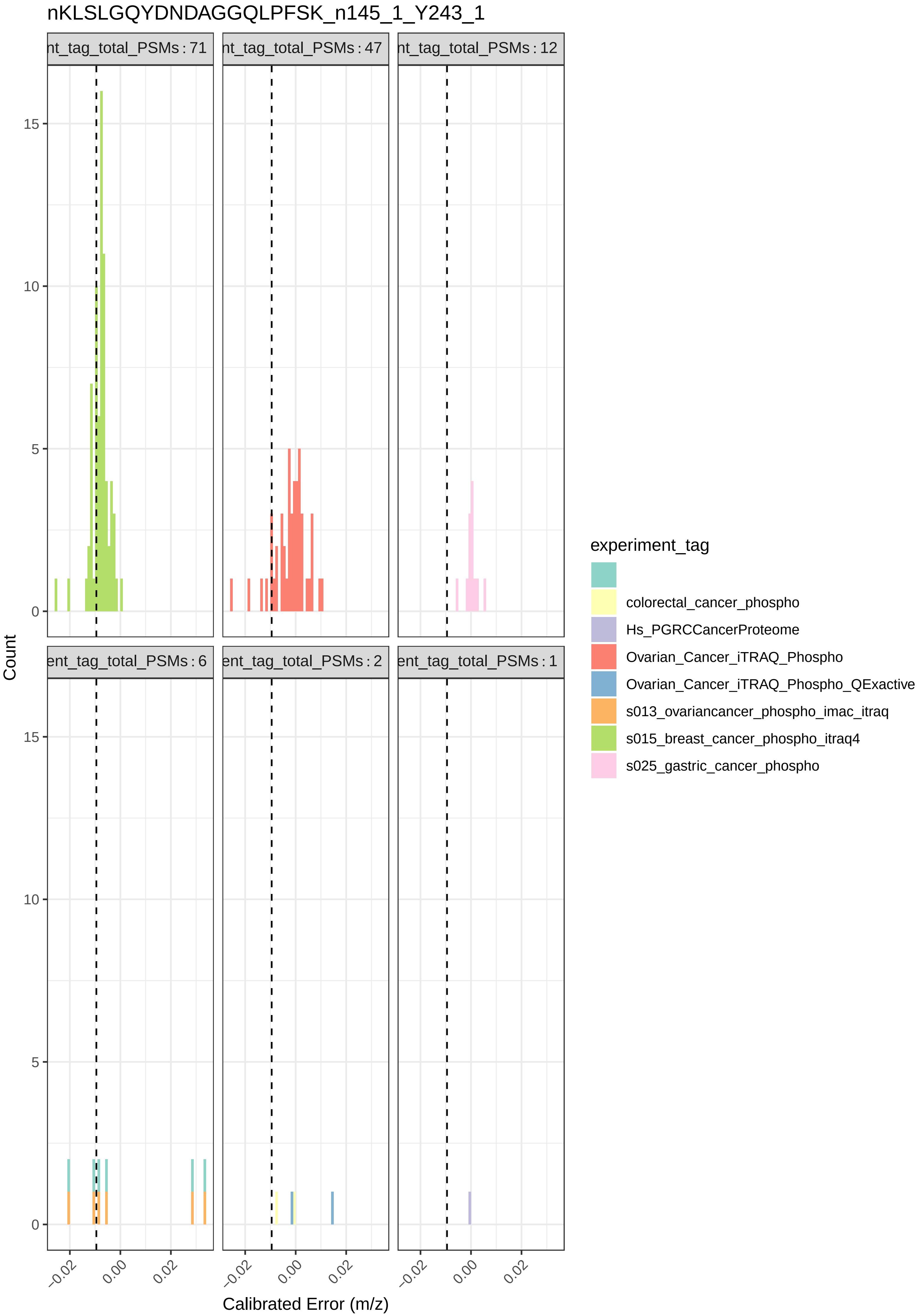
nKLSLGQYDNDAGGQLPFSK_n145_1_Y243_1
nt_tag_total_PSMs : 71
nt_tag_total_PSMs : 47
nt_tag_total_PSMs : 12
ent_tag_total_PSMs : 6
ent_tag_total_PSMs : 2
ent_tag_total_PSMs : 1
Count
Calibrated Error (m/z)
0
5
10
15
-0.02
0.00
0.02
experiment_tag
colorectal_cancer_phospho
Hs_PGRCCancerProteome
Ovarian_Cancer_iTRAQ_Phospho
Ovarian_Cancer_iTRAQ_Phospho_QExactive
s013_ovariancancer_phospho_imac_itraq
s015_breast_cancer_phospho_itraq4
s025_gastric_cancer_phospho

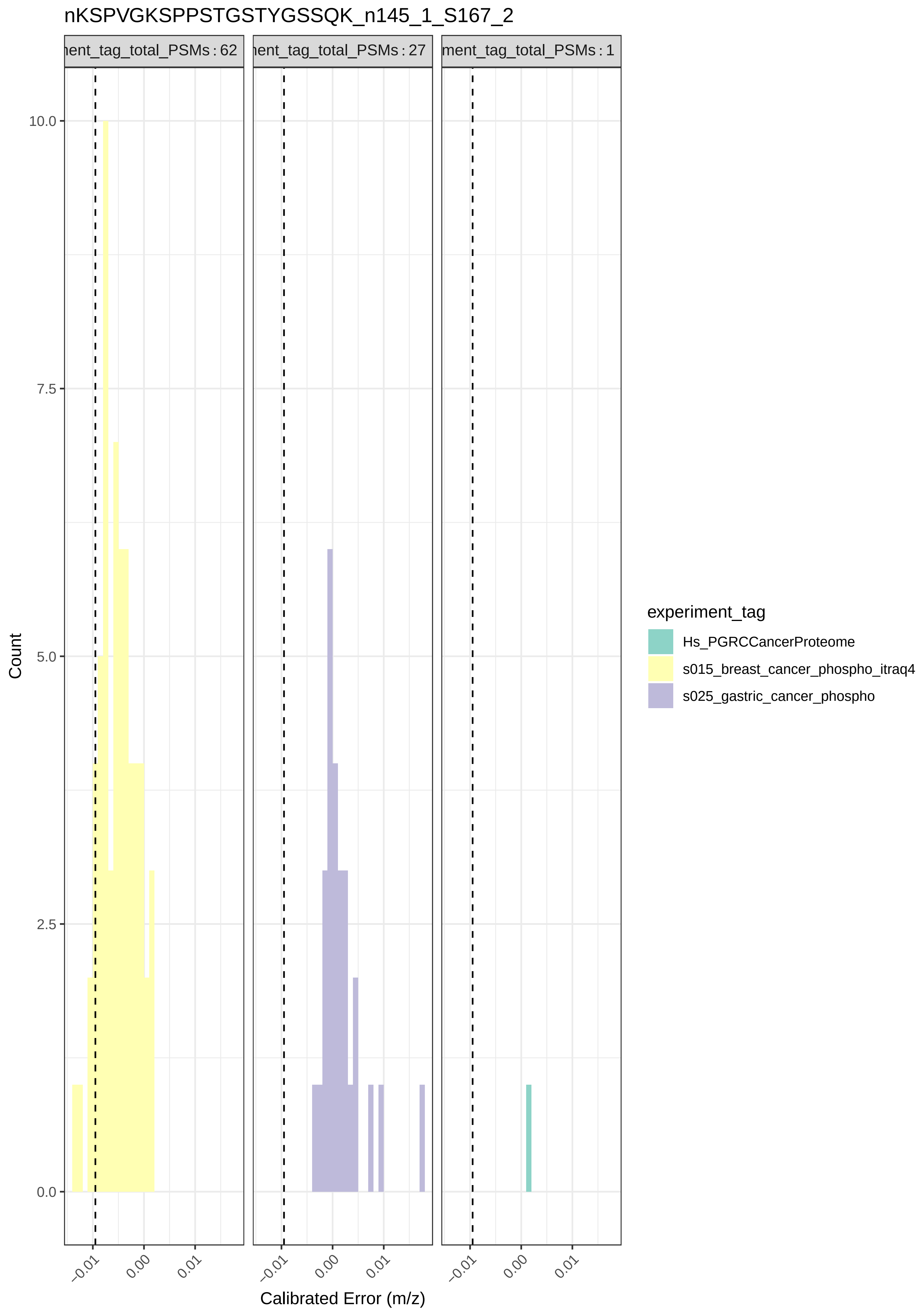

nKSPVGKSPPSTGSTYGSSQK_n145_1_S167_2
ment_tag_total_PSMs : 62
ment_tag_total_PSMs : 27
ment_tag_total_PSMs : 1
Count
10.0
7.5
5.0
2.5
0.0
−0.01
0.00
0.01
Calibrated Error (m/z)
experiment_tag
Hs_PGRCCancerProteome
s015_breast_cancer_phospho_itraq4
s025_gastric_cancer_phospho

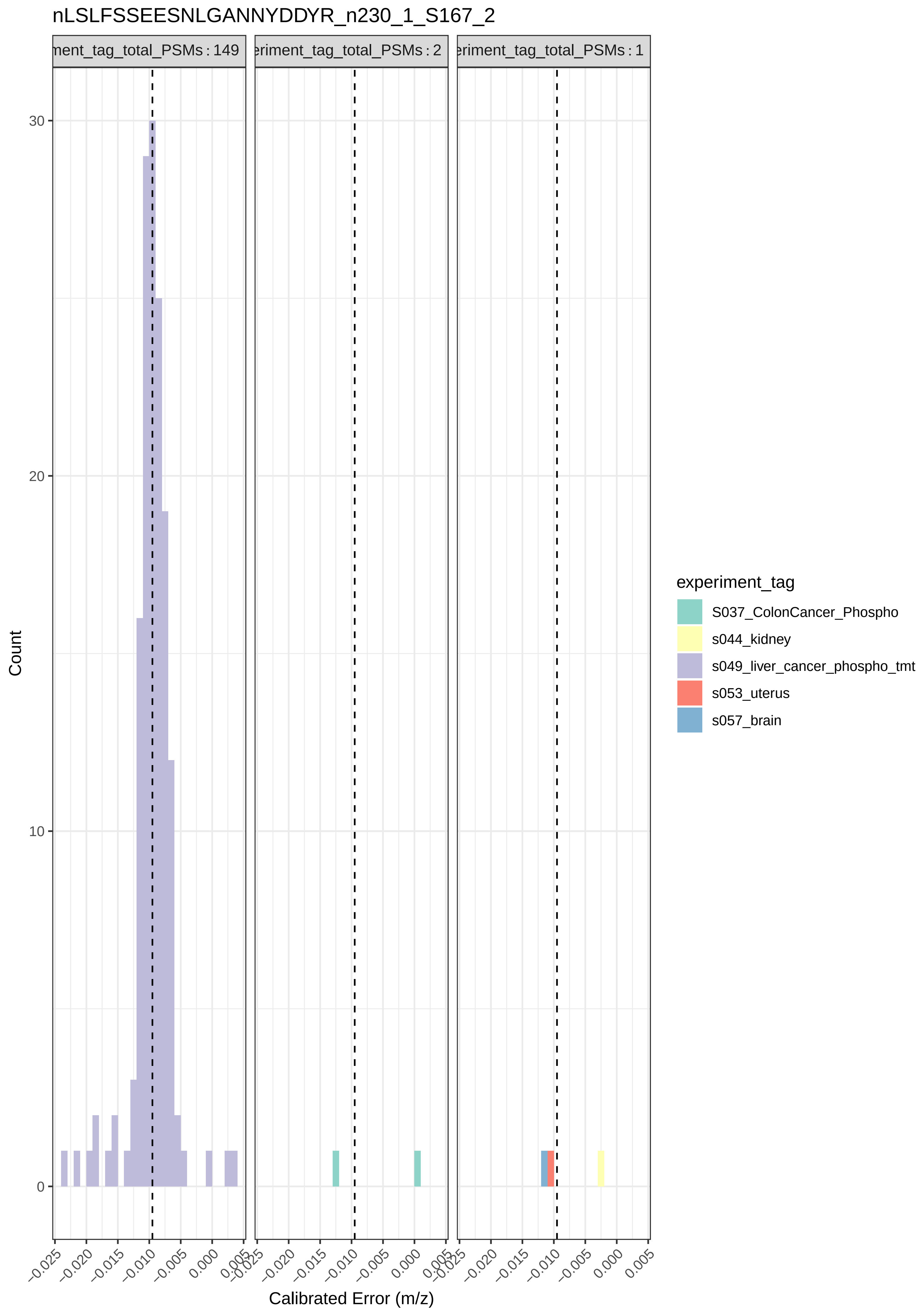

nLSLFSSEESNLGANNYDDYR_n230_1_S167_2
ment_tag_total_PSMs : 149
eriment_tag_total_PSMs : 2
eriment_tag_total_PSMs : 1
Count
30
20
10
0
−0.025
−0.020
−0.015
−0.010
−0.005
0.000
0.005
Calibrated Error (m/z)
experiment_tag
S037_ColonCancer_Phospho
s044_kidney
s049_liver_cancer_phospho_tmt
s053_uterus
s057_brain

nLSLFSSEESNLGANNYDDYR_n230_1_S167_3

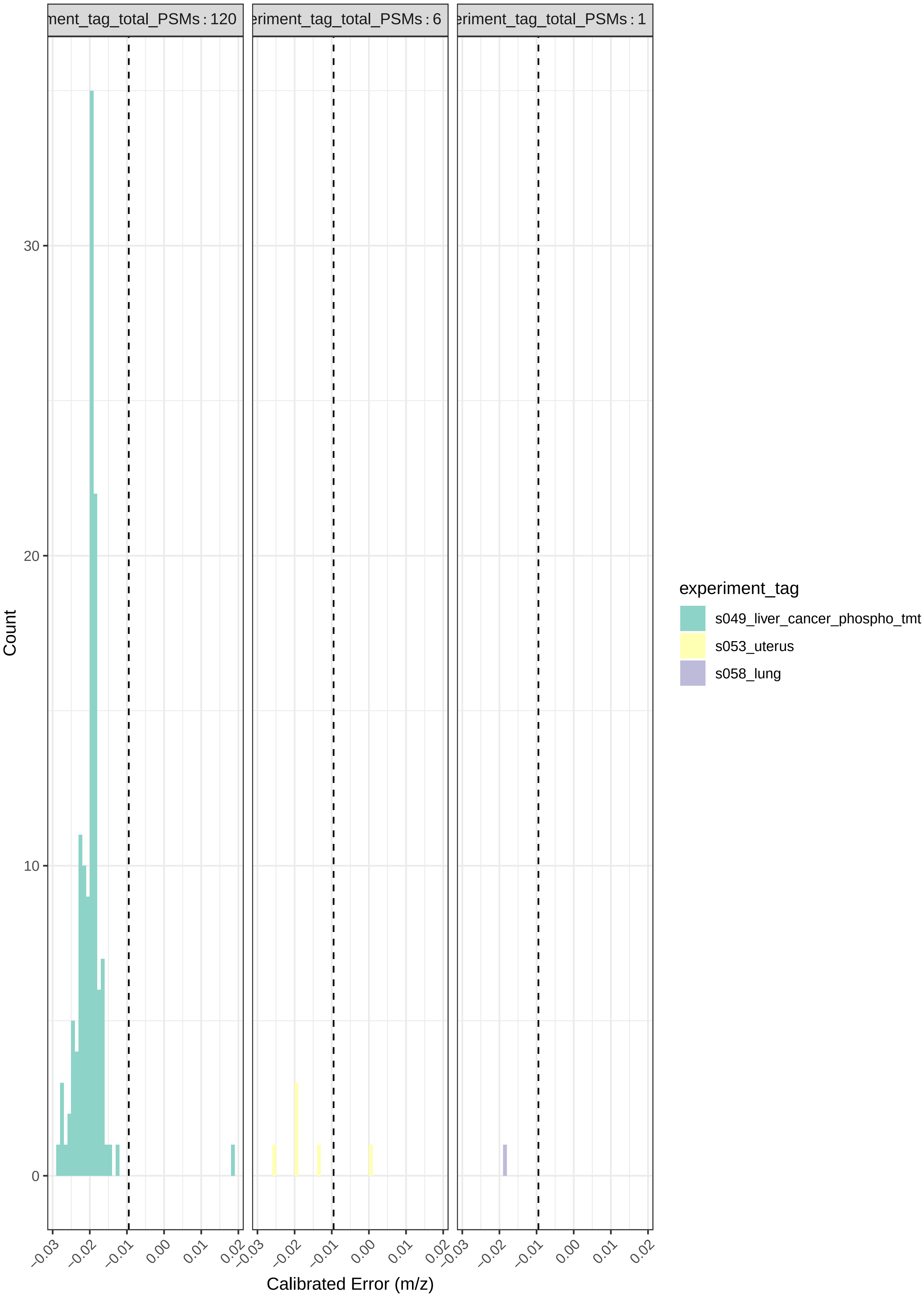

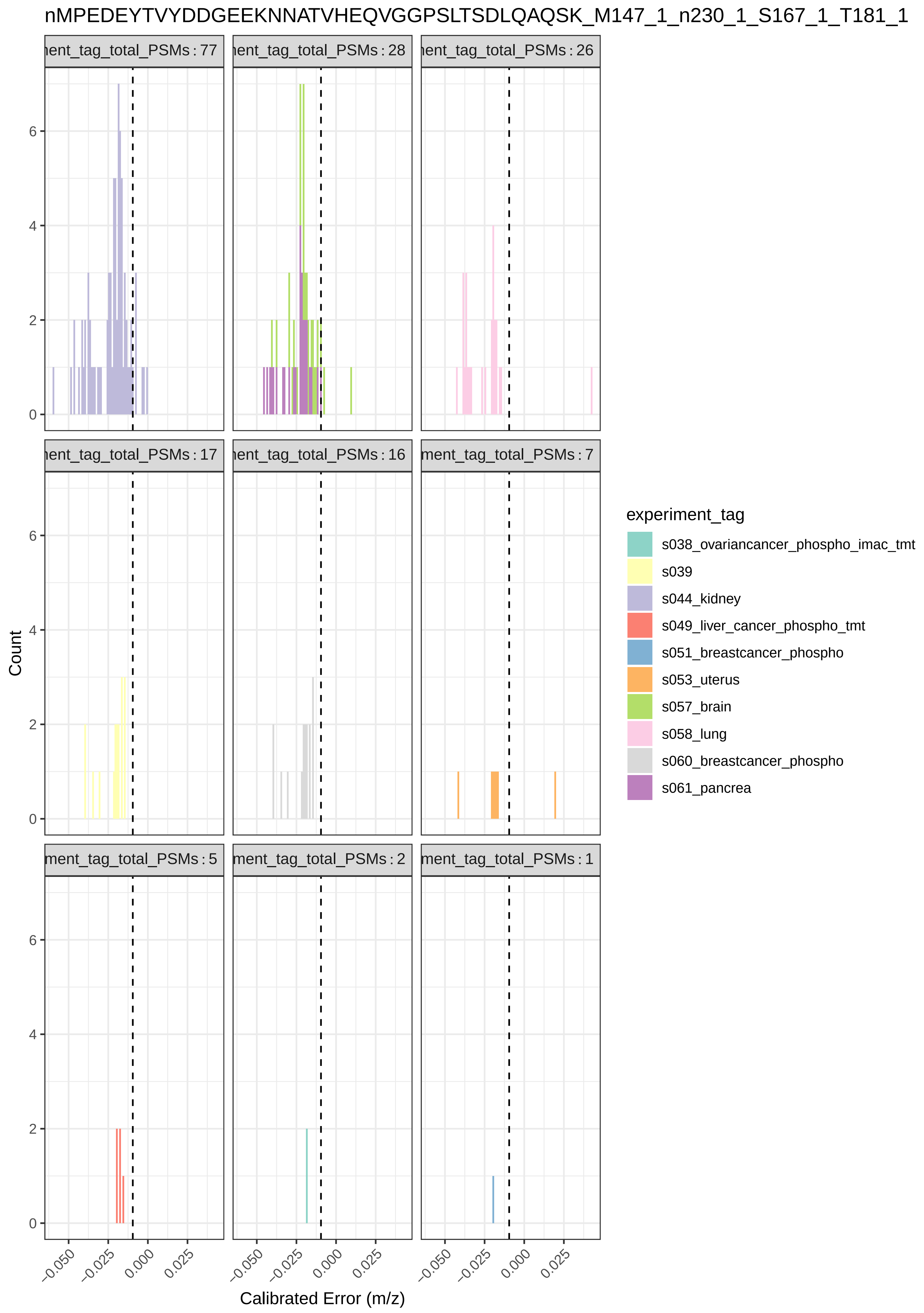

nMPEDEYTVYDDGEEKNNATVHEQVGGPSLTSDLQAQSK_M147_1_n230_1_S167_1_T181_1
ment_tag_total_PSMs : 77
ment_tag_total_PSMs : 28
ment_tag_total_PSMs : 26
ment_tag_total_PSMs : 17
ment_tag_total_PSMs : 16
ment_tag_total_PSMs : 7
ment_tag_total_PSMs : 5
ment_tag_total_PSMs : 2
ment_tag_total_PSMs : 1
Count
0
2
4
6
−0.050
−0.025
0.000
0.025
Calibrated Error (m/z)
experiment_tag
s038_ovariancancer_phospho_imac_tmt
s039
s044_kidney
s049_liver_cancer_phospho_tmt
s051_breastcancer_phospho
s053_uterus
s057_brain
s058_lung
s060_breastcancer_phospho
s061_pancrea

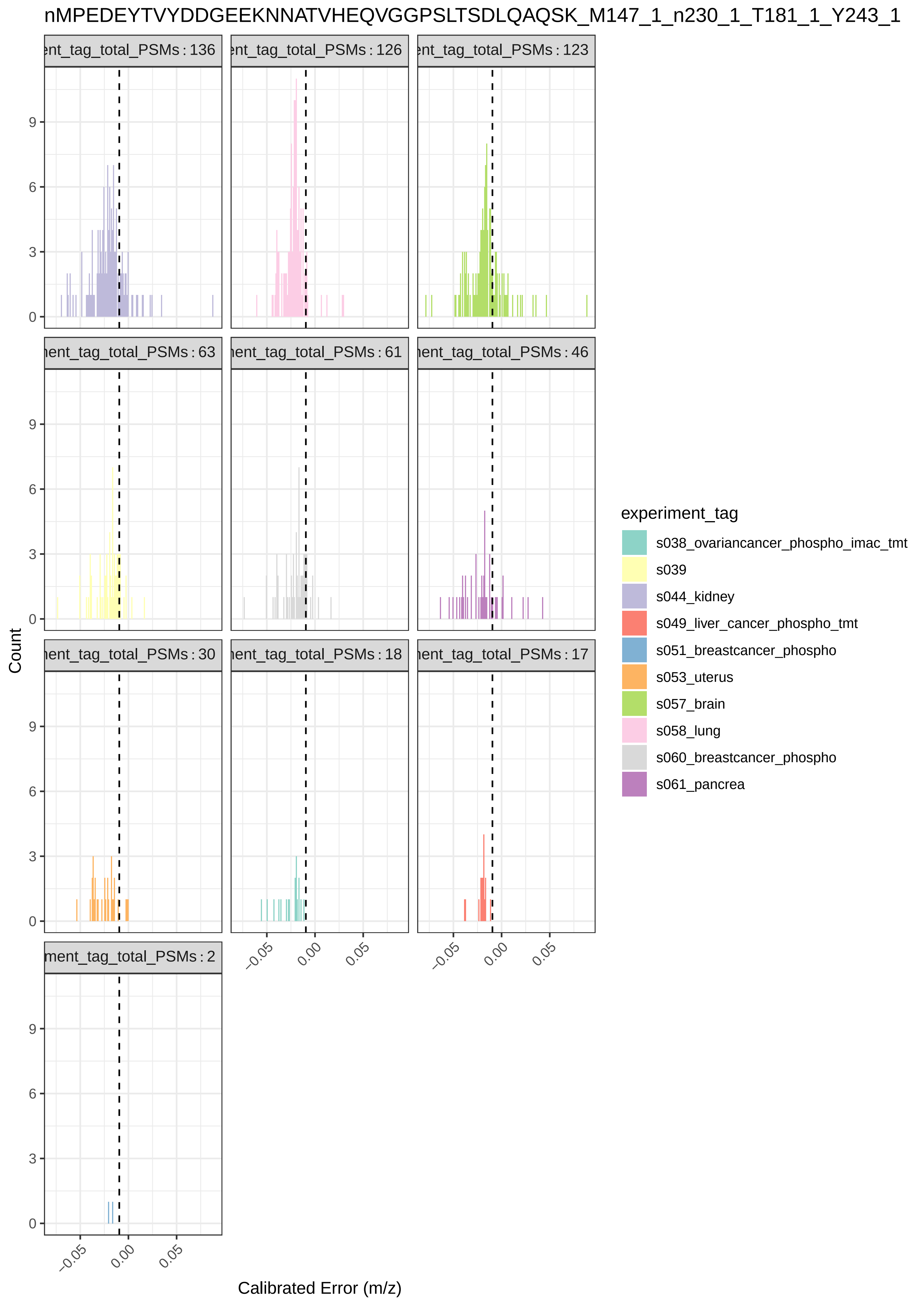
nMPEDEYTVYDDGEEKNNATVHEQVGGPSLTSDLQAQSK_M147_1_n230_1_T181_1_Y243_1
ent_tag_total_PSMs : 136
ent_tag_total_PSMs : 126
ent_tag_total_PSMs : 123
nent_tag_total_PSMs : 63
nent_tag_total_PSMs : 61
nent_tag_total_PSMs : 46
nent_tag_total_PSMs : 30
nent_tag_total_PSMs : 18
nent_tag_total_PSMs : 17
ment_tag_total_PSMs : 2
Count
Calibrated Error (m/z)
0
3
6
9
−0.05
0.00
0.05
experiment_tag
s038_ovariancancer_phospho_imac_tmt
s039
s044_kidney
s049_liver_cancer_phospho_tmt
s051_breastcancer_phospho
s053_uterus
s057_brain
s058_lung
s060_breastcancer_phospho
s061_pancrea

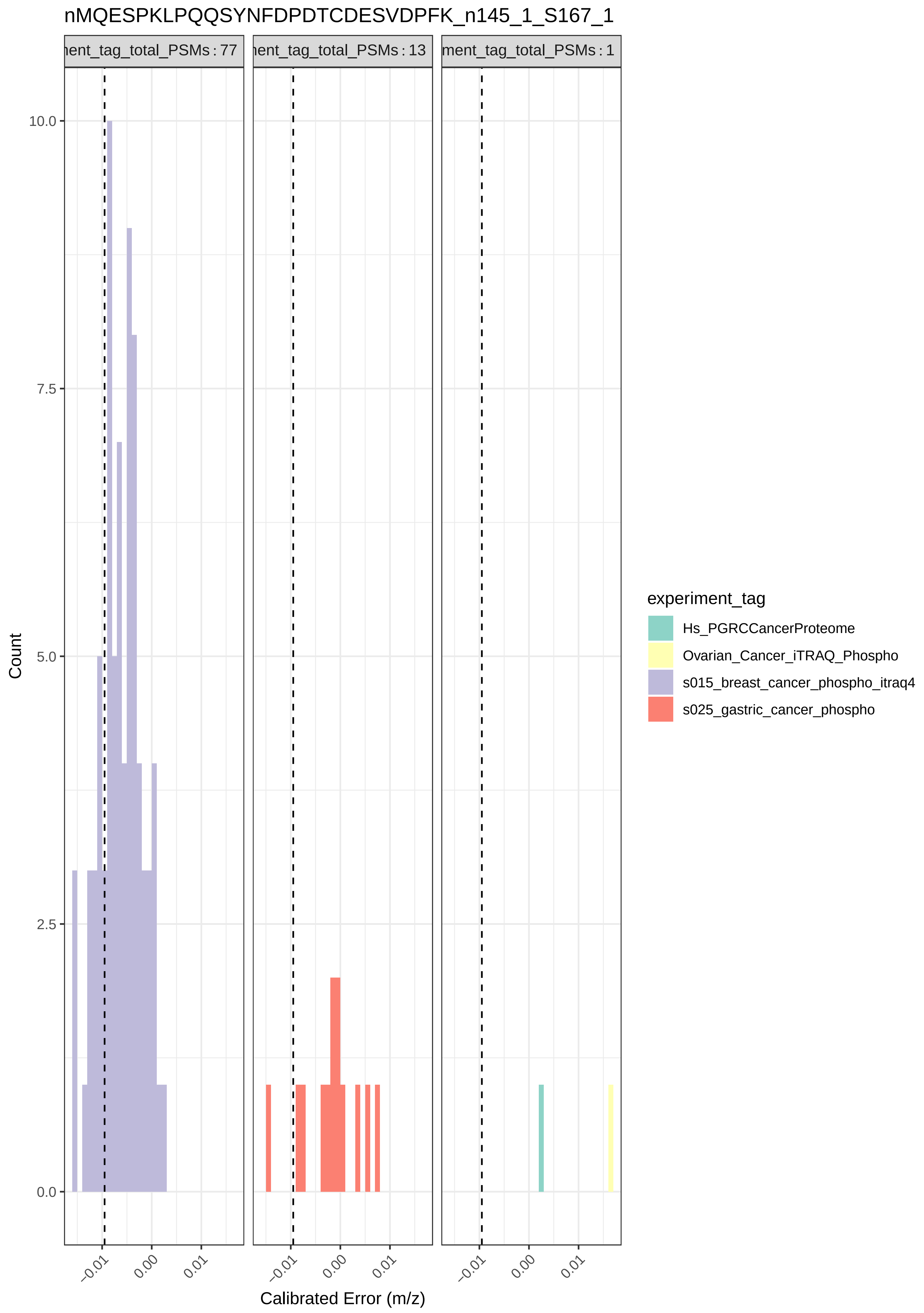
nMQESPKLPQQSYNFDPDTCDESVDPFK_n145_1_S167_1
ent_tag_total_PSMs : 77
ent_tag_total_PSMs : 13
ment_tag_total_PSMs : 1
Count
10.0
7.5
5.0
2.5
0.0
−0.01
0.00
0.01
Calibrated Error (m/z)
experiment_tag
Hs_PGRCCancerProteome
Ovarian_Cancer_iTRAQ_Phospho
s015_breast_cancer_phospho_itraq4
s025_gastric_cancer_phospho

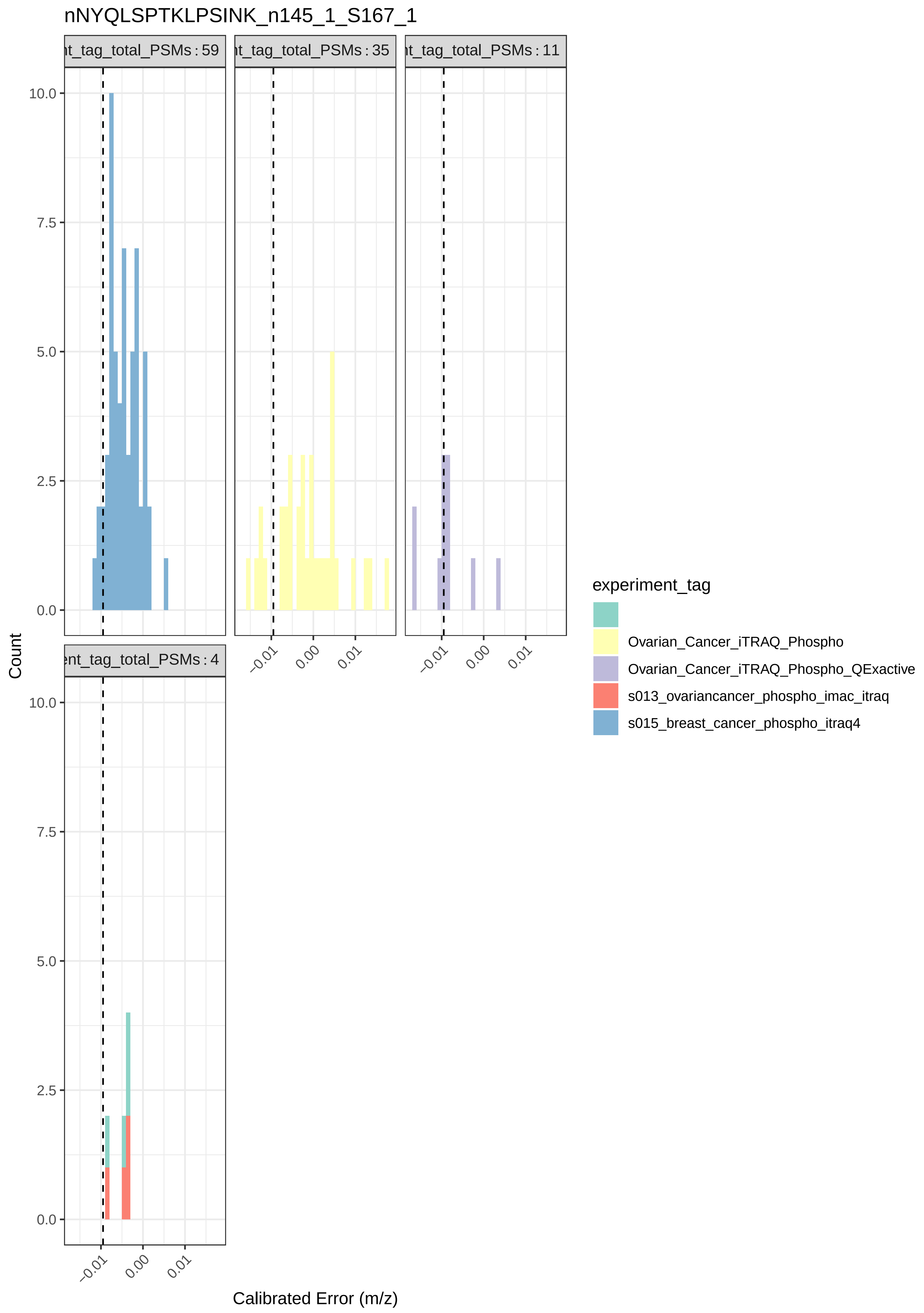
nNYQLSPTKLPSINK_n145_1_S167_1
nt_tag_total_PSMs : 59
nt_tag_total_PSMs : 35
nt_tag_total_PSMs : 11
ent_tag_total_PSMs : 4
10.0
7.5
5.0
2.5
0.0
Count
−0.01
0.00
0.01
Calibrated Error (m/z)
experiment_tag
Ovarian_Cancer_iTRAQ_Phospho
Ovarian_Cancer_iTRAQ_Phospho_QExactive
s013_ovariancancer_phospho_imac_itraq
s015_breast_cancer_phospho_itraq4

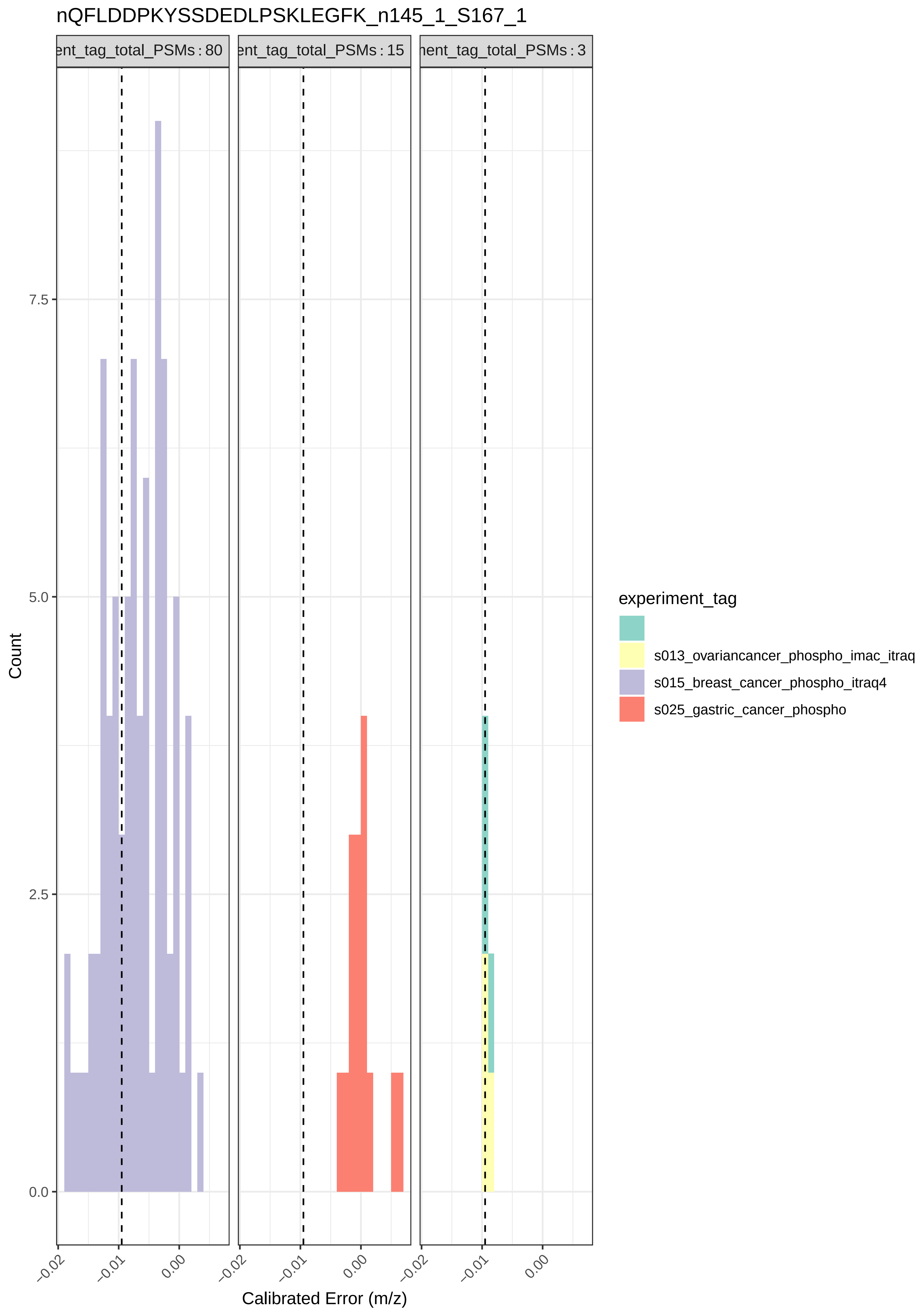

nQFLDDPKYSSDEDLPSKLEGFK_n145_1_S167_1
ent_tag_total_PSMs : 80
ent_tag_total_PSMs : 15
nent_tag_total_PSMs : 3
Count
7.5
5.0
2.5
0.0
−0.02
−0.01
0.00
Calibrated Error (m/z)
experiment_tag
s013_ovariancancer_phospho_imac_itraq
s015_breast_cancer_phospho_itraq4
s025_gastric_cancer_phospho

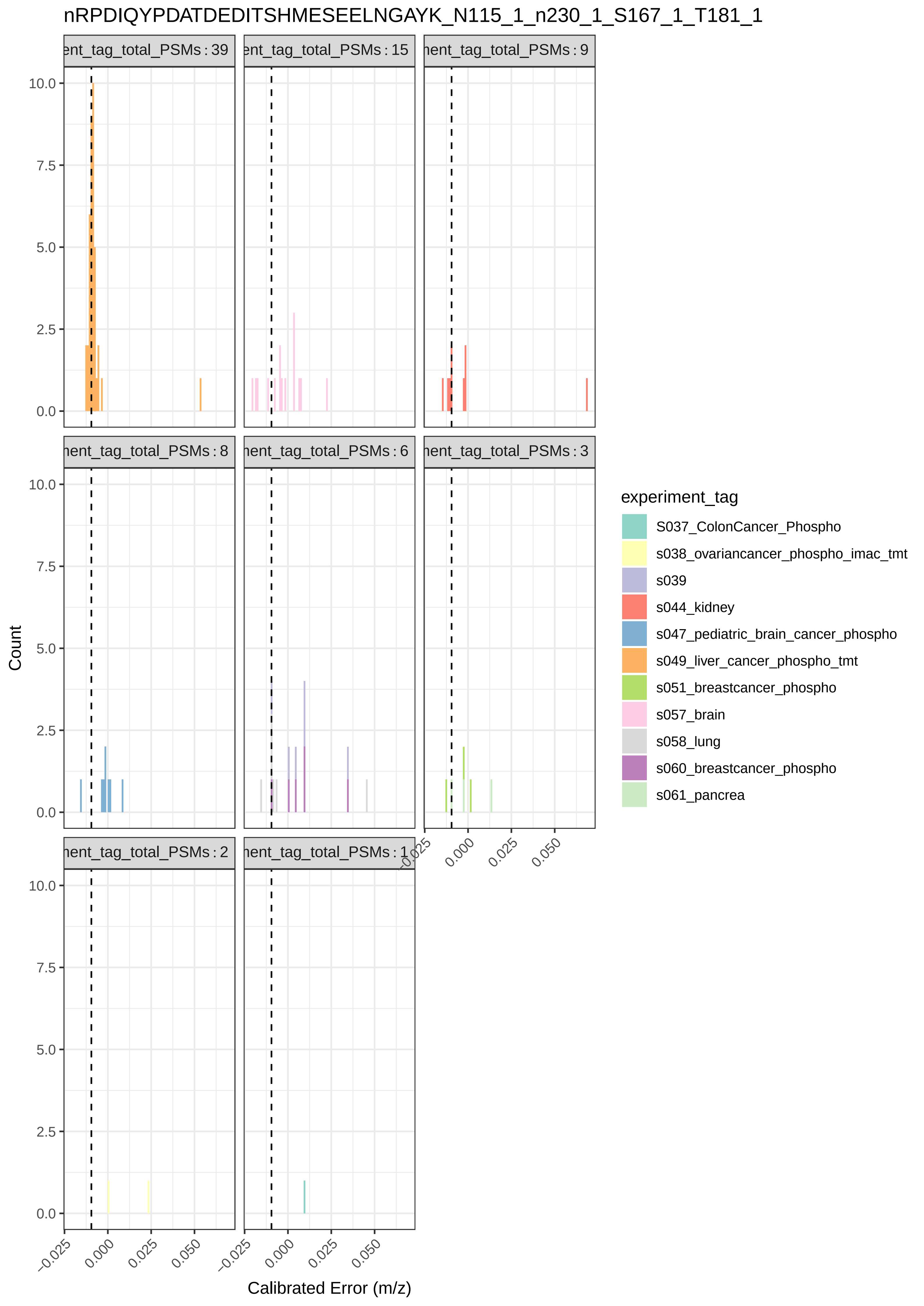
nRPDIQYPDATDEDITSHMESEELNGAYK_N115_1_n230_1_S167_1_T181_1
ent_tag_total_PSMs : 39
ent_tag_total_PSMs : 15
nent_tag_total_PSMs : 9
nent_tag_total_PSMs : 8
nent_tag_total_PSMs : 6
nent_tag_total_PSMs : 3
nent_tag_total_PSMs : 2
nent_tag_total_PSMs : 1
Count
Calibrated Error (m/z)
0.0
2.5
5.0
7.5
10.0
−0.025
0.000
0.025
0.050
experiment_tag
S037_ColonCancer_Phospho
s038_ovariancancer_phospho_imac_tmt
s039
s044_kidney
s047_pediatric_brain_cancer_phospho
s049_liver_cancer_phospho_tmt
s051_breastcancer_phospho
s057_brain
s058_lung
s060_breastcancer_phospho
s061_pancrea

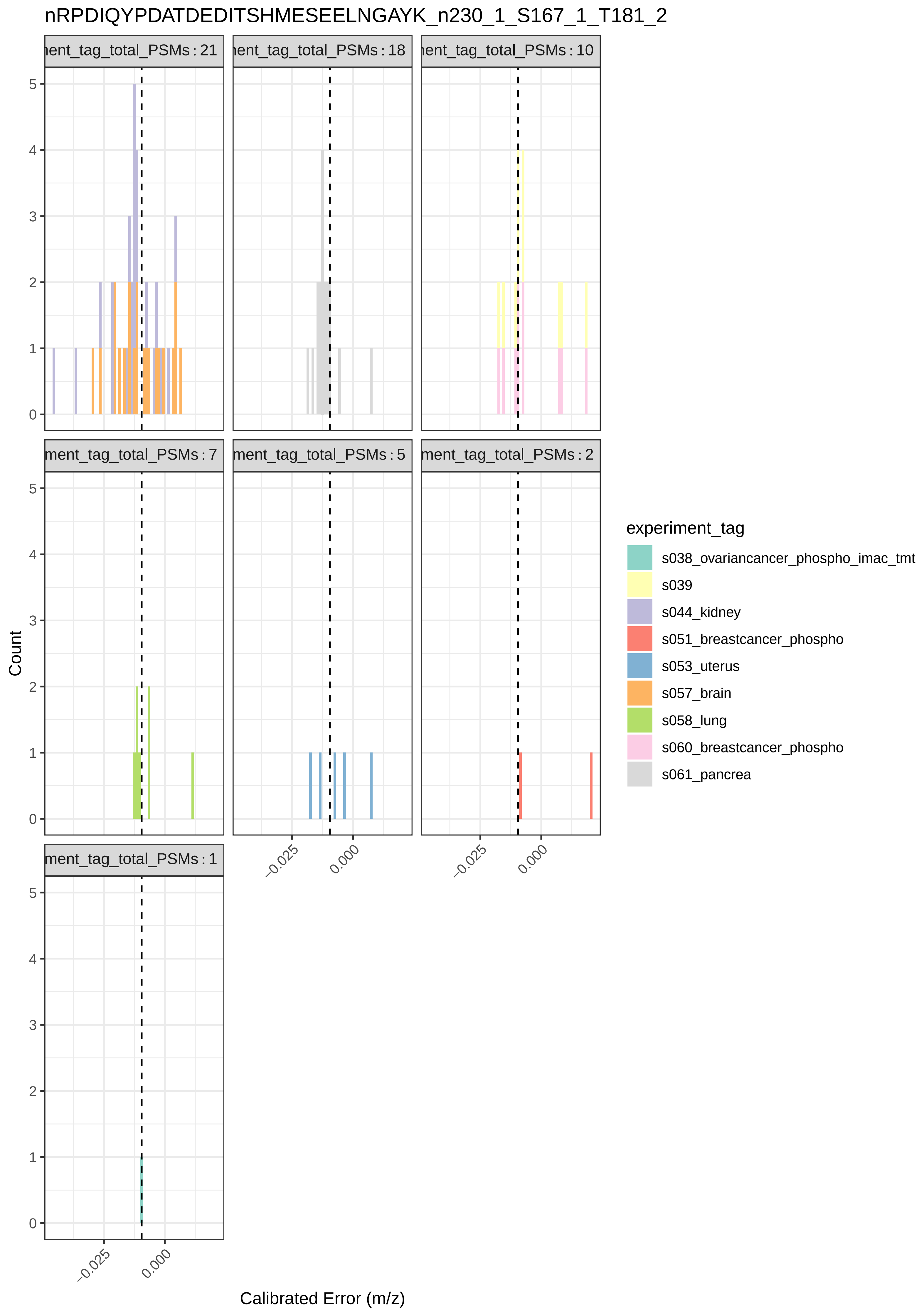

nRPDIQYPDATDEDITSHMESEELNGAYK_n230_1_S167_1_T181_2
ment_tag_total_PSMs : 21
ment_tag_total_PSMs : 18
ment_tag_total_PSMs : 10
ment_tag_total_PSMs : 7
ment_tag_total_PSMs : 5
ment_tag_total_PSMs : 2
ment_tag_total_PSMs : 1
Count
0
1
2
3
4
5
−0.025
0.000
Calibrated Error (m/z)
experiment_tag
s038_ovariancancer_phospho_imac_tmt
s039
s044_kidney
s051_breastcancer_phospho
s053_uterus
s057_brain
s058_lung
s060_breastcancer_phospho
s061_pancrea

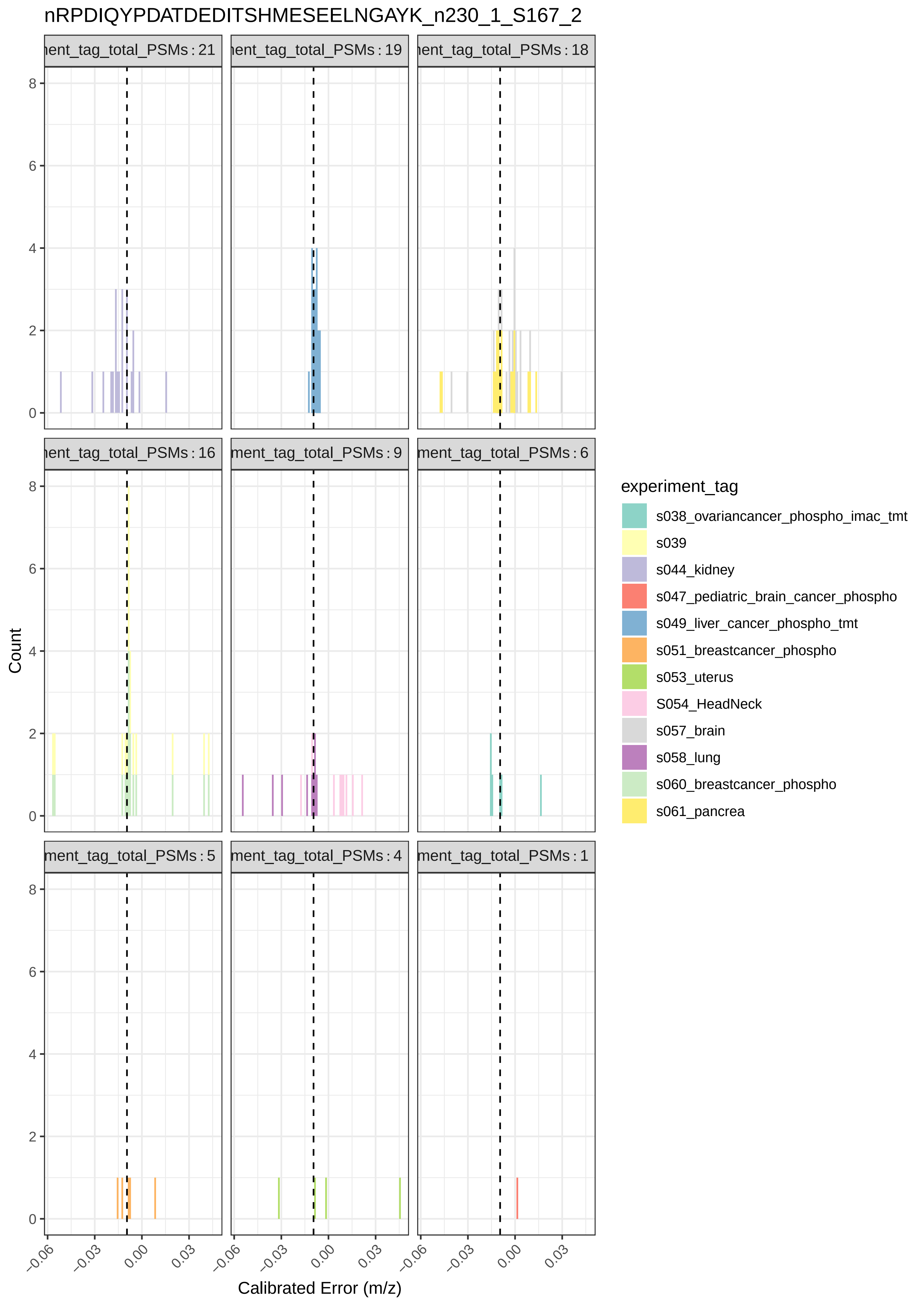

nRPDIQYPDATDEDITSHMESEELNGAYK_n230_1_S167_2
ment_tag_total_PSMs : 21
ment_tag_total_PSMs : 19
ment_tag_total_PSMs : 18
ment_tag_total_PSMs : 16
ment_tag_total_PSMs : 9
ment_tag_total_PSMs : 6
ment_tag_total_PSMs : 5
ment_tag_total_PSMs : 4
ment_tag_total_PSMs : 1
Count
Calibrated Error (m/z)
experiment_tag
s038_ovariancancer_phospho_imac_tmt
s039
s044_kidney
s047_pediatric_brain_cancer_phospho
s049_liver_cancer_phospho_tmt
s051_breastcancer_phospho
s053_uterus
S054_HeadNeck
s057_brain
s058_lung
s060_breastcancer_phospho
s061_pancrea

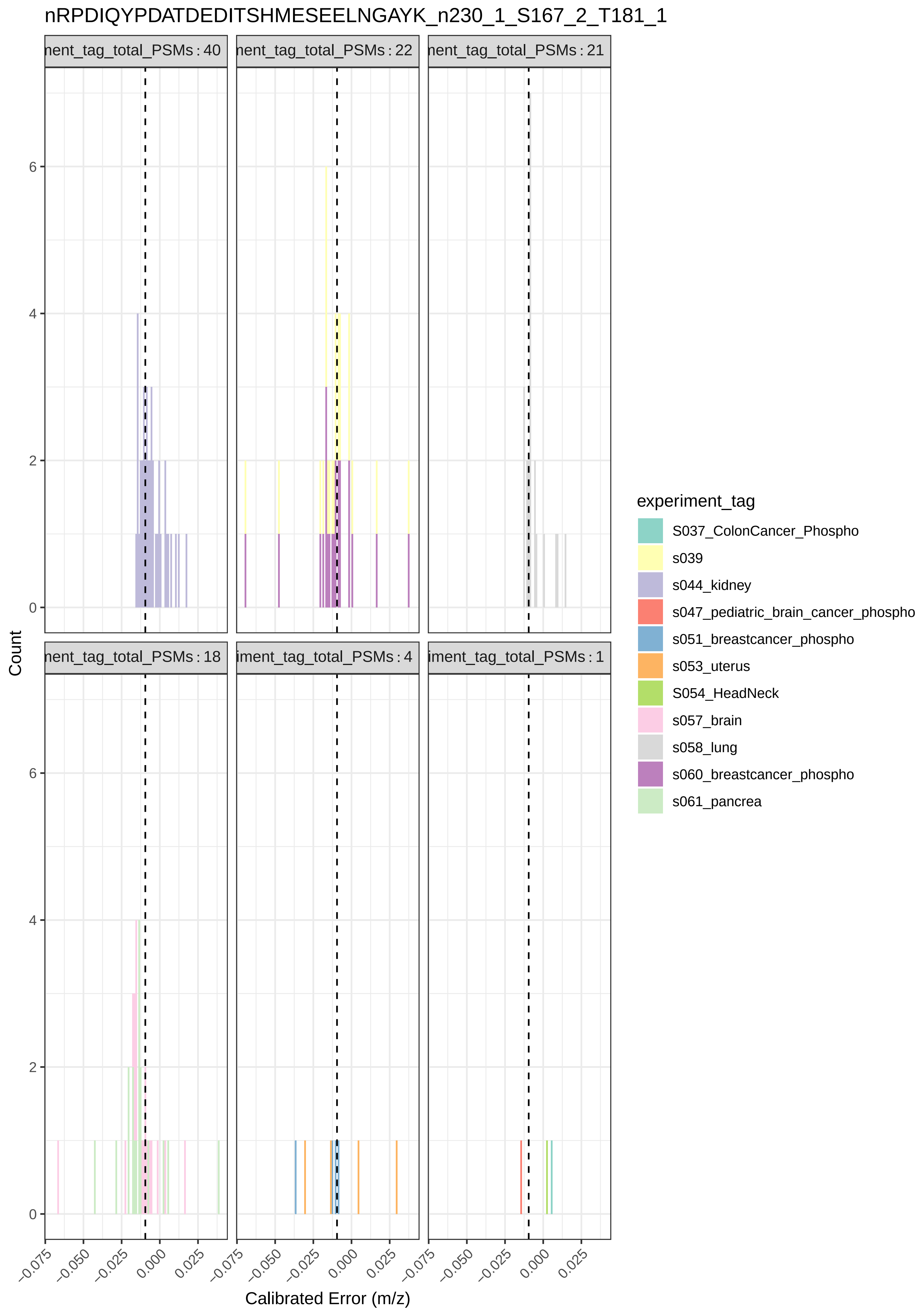
nRPDIQYPDATDEDITSHMESEELNGAYK_n230_1_S167_2_T181_1
nent_tag_total_PSMs : 40
nent_tag_total_PSMs : 22
nent_tag_total_PSMs : 21
nent_tag_total_PSMs : 18
iment_tag_total_PSMs : 4
iment_tag_total_PSMs : 1
Count
0
2
4
6
−0.075
−0.050
−0.025
0.000
0.025
Calibrated Error (m/z)
experiment_tag
S037_ColonCancer_Phospho
s039
s044_kidney
s047_pediatric_brain_cancer_phospho
s051_breastcancer_phospho
s053_uterus
S054_HeadNeck
s057_brain
s058_lung
s060_breastcancer_phospho
s061_pancrea

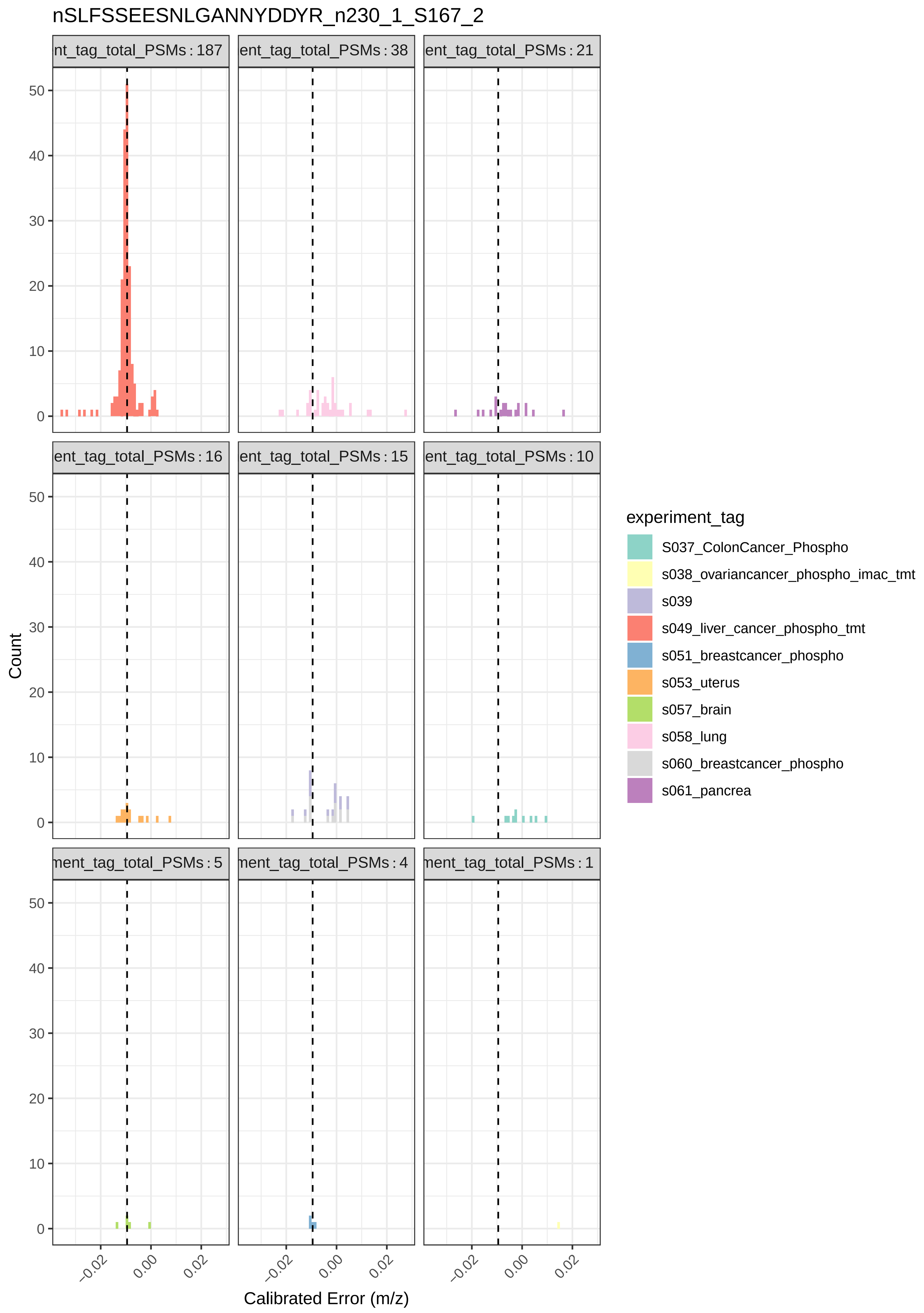
nSLFSSEESNLGANNYDDYR_n230_1_S167_2
nt_tag_total_PSMs : 187
ent_tag_total_PSMs : 38
ent_tag_total_PSMs : 21
ent_tag_total_PSMs : 16
ent_tag_total_PSMs : 15
ent_tag_total_PSMs : 10
ment_tag_total_PSMs : 5
ment_tag_total_PSMs : 4
ment_tag_total_PSMs : 1
Count
Calibrated Error (m/z)
−0.02
0.00
0.02
0
10
20
30
40
50
experiment_tag
S037_ColonCancer_Phospho
s038_ovariancancer_phospho_imac_tmt
s039
s049_liver_cancer_phospho_tmt
s051_breastcancer_phospho
s053_uterus
s057_brain
s058_lung
s060_breastcancer_phospho
s061_pancrea

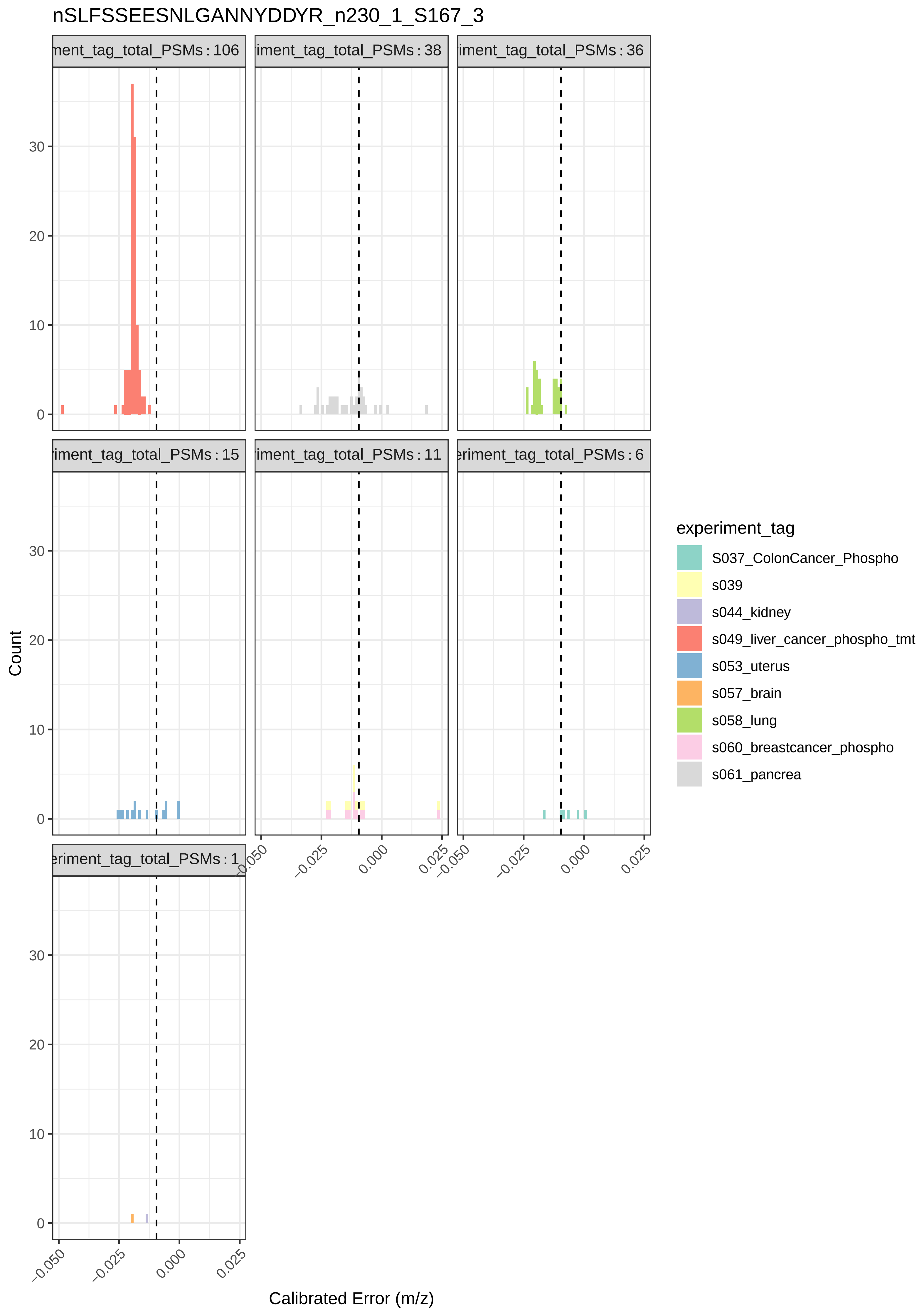
nSLFSSEESNLGANNYDDYR_n230_1_S167_3
ment_tag_total_PSMs : 106
riment_tag_total_PSMs : 38
riment_tag_total_PSMs : 36
riment_tag_total_PSMs : 15
riment_tag_total_PSMs : 11
eriment_tag_total_PSMs : 6
eriment_tag_total_PSMs : 1
Count
Calibrated Error (m/z)
0
10
20
30
−0.050
−0.025
0.000
0.025
experiment_tag
S037_ColonCancer_Phospho
s039
s044_kidney
s049_liver_cancer_phospho_tmt
s053_uterus
s057_brain
s058_lung
s060_breastcancer_phospho
s061_pancrea

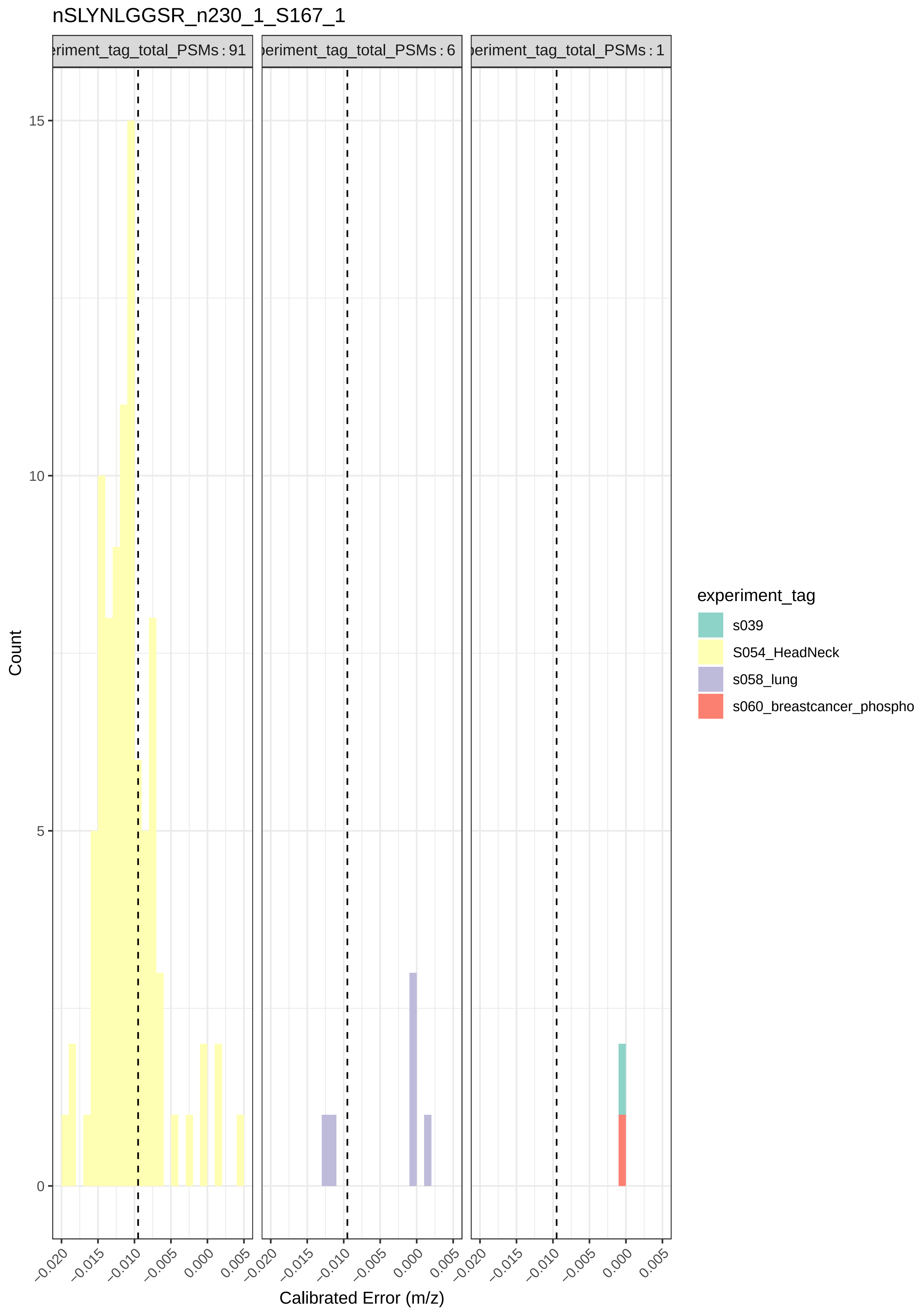

nSLYNLGGSR_n230_1_S167_1
eriment_tag_total_PSMs : 91
periment_tag_total_PSMs : 6
periment_tag_total_PSMs : 1
Count
0
5
10
15
−0.020
−0.015
−0.010
−0.005
0.000
0.005
Calibrated Error (m/z)
experiment_tag
s039
S054_HeadNeck
s058_lung
s060_breastcancer_phospho

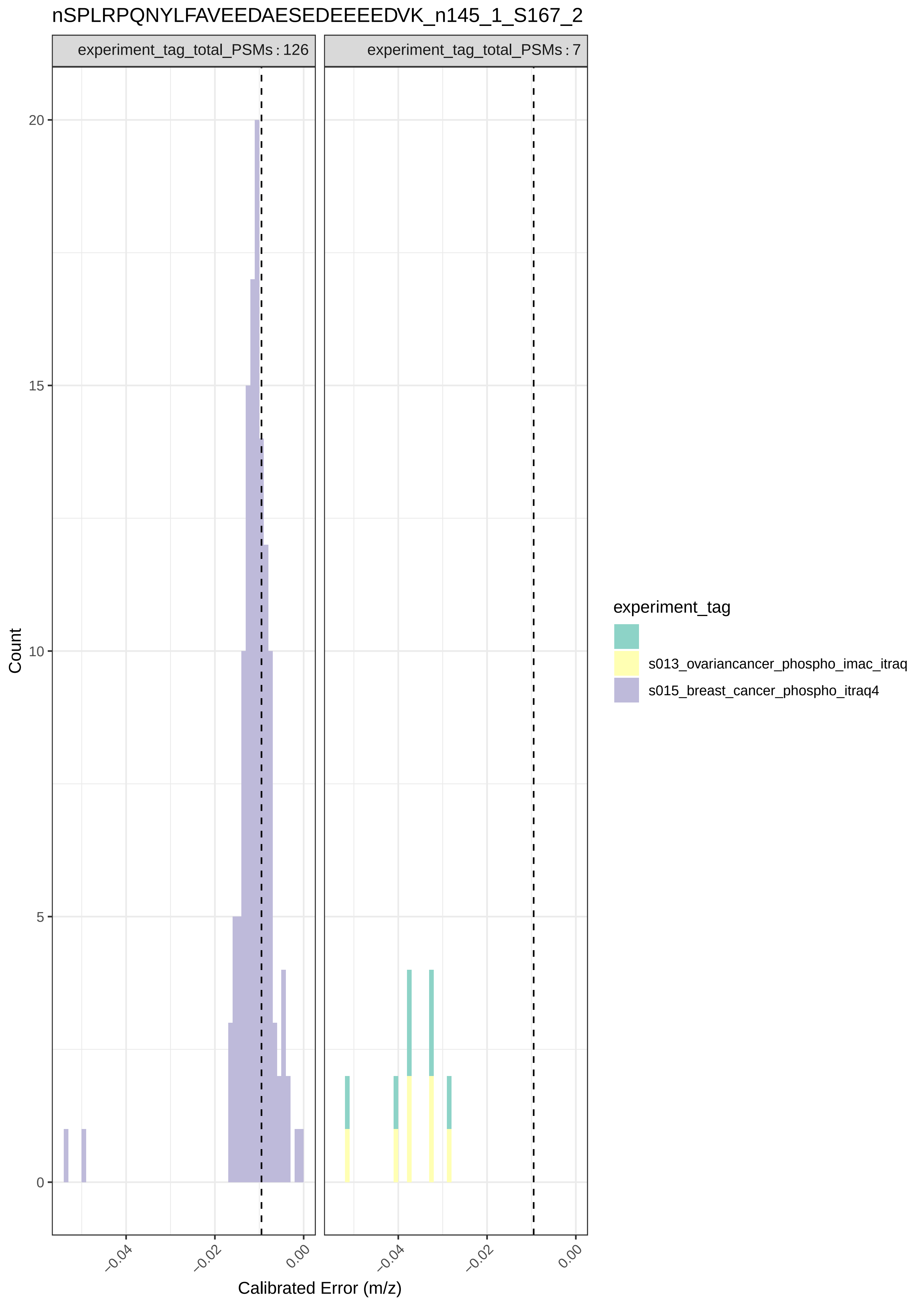
nSPLRPQNYLFAVEEDAESEDEEEEDVK_n145_1_S167_2
experiment_tag_total_PSMs : 126
experiment_tag_total_PSMs : 7
Count
20
15
10
5
0
-0.04
-0.02
0.00
Calibrated Error (m/z)
experiment_tag
s013_ovariancancer_phospho_imac_itraq
s015_breast_cancer_phospho_itraq4

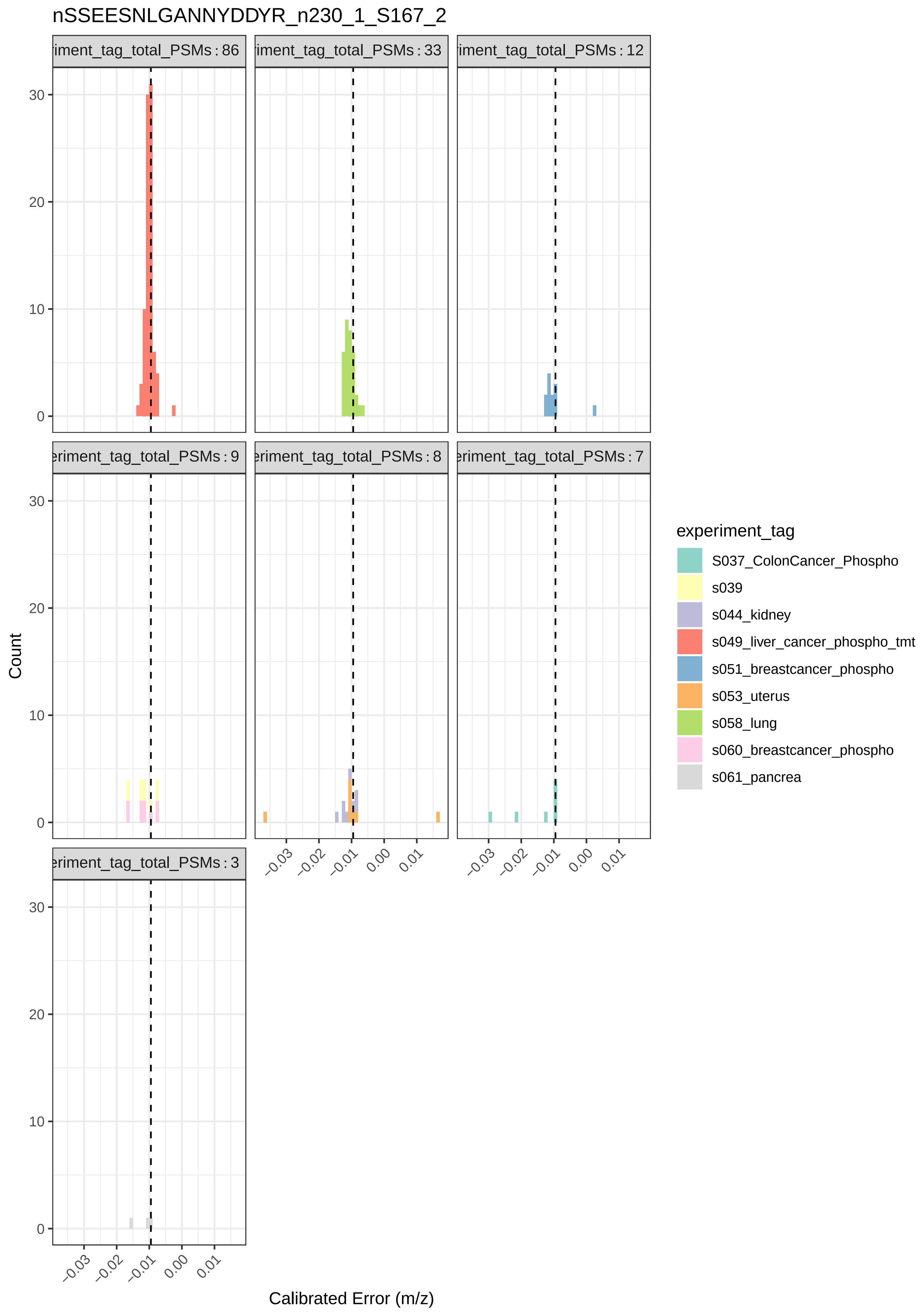

nSSEESNLGANNYDDYR_n230_1_S167_2
iment_tag_total_PSMs : 86
iment_tag_total_PSMs : 33
iment_tag_total_PSMs : 12
eriment_tag_total_PSMs : 9
eriment_tag_total_PSMs : 8
eriment_tag_total_PSMs : 7
eriment_tag_total_PSMs : 3
Count
0
10
20
30
−0.03
−0.02
−0.01
0.00
0.01
Calibrated Error (m/z)
experiment_tag
S037_ColonCancer_Phospho
s039
s044_kidney
s049_liver_cancer_phospho_tmt
s051_breastcancer_phospho
s053_uterus
s058_lung
s060_breastcancer_phospho
s061_pancrea

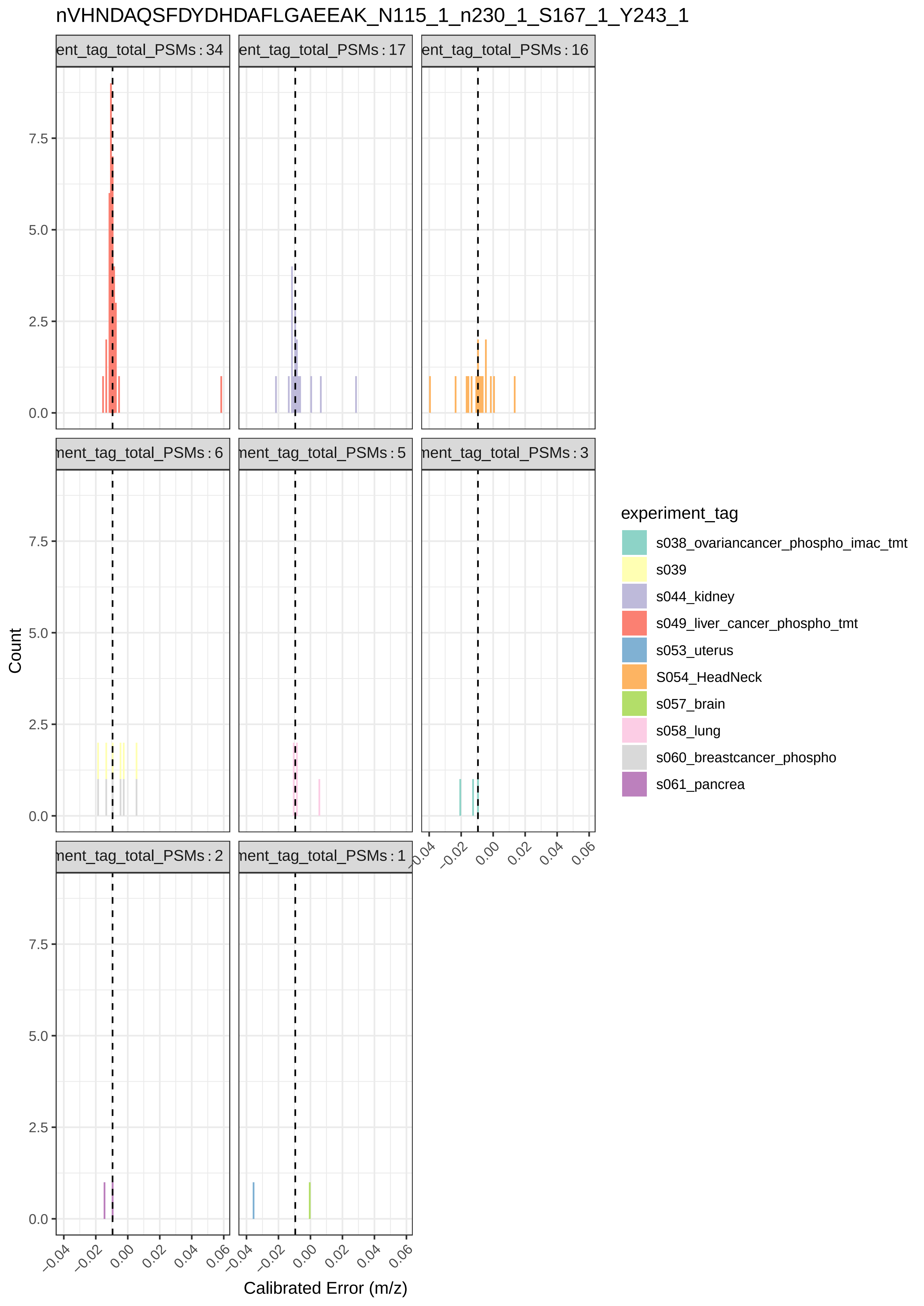
nVHNDAQSFDYDHDAFLGAEEAK_N115_1_n230_1_S167_1_Y243_1
ent_tag_total_PSMs : 34
ent_tag_total_PSMs : 17
ent_tag_total_PSMs : 16
nent_tag_total_PSMs : 6
nent_tag_total_PSMs : 5
nent_tag_total_PSMs : 3
nent_tag_total_PSMs : 2
nent_tag_total_PSMs : 1
Count
0.0
2.5
5.0
7.5
−0.04
−0.02
0.00
0.02
0.04
0.06
Calibrated Error (m/z)
experiment_tag
s038_ovariancancer_phospho_imac_tmt
s039
s044_kidney
s049_liver_cancer_phospho_tmt
s053_uterus
S054_HeadNeck
s057_brain
s058_lung
s060_breastcancer_phospho
s061_pancrea

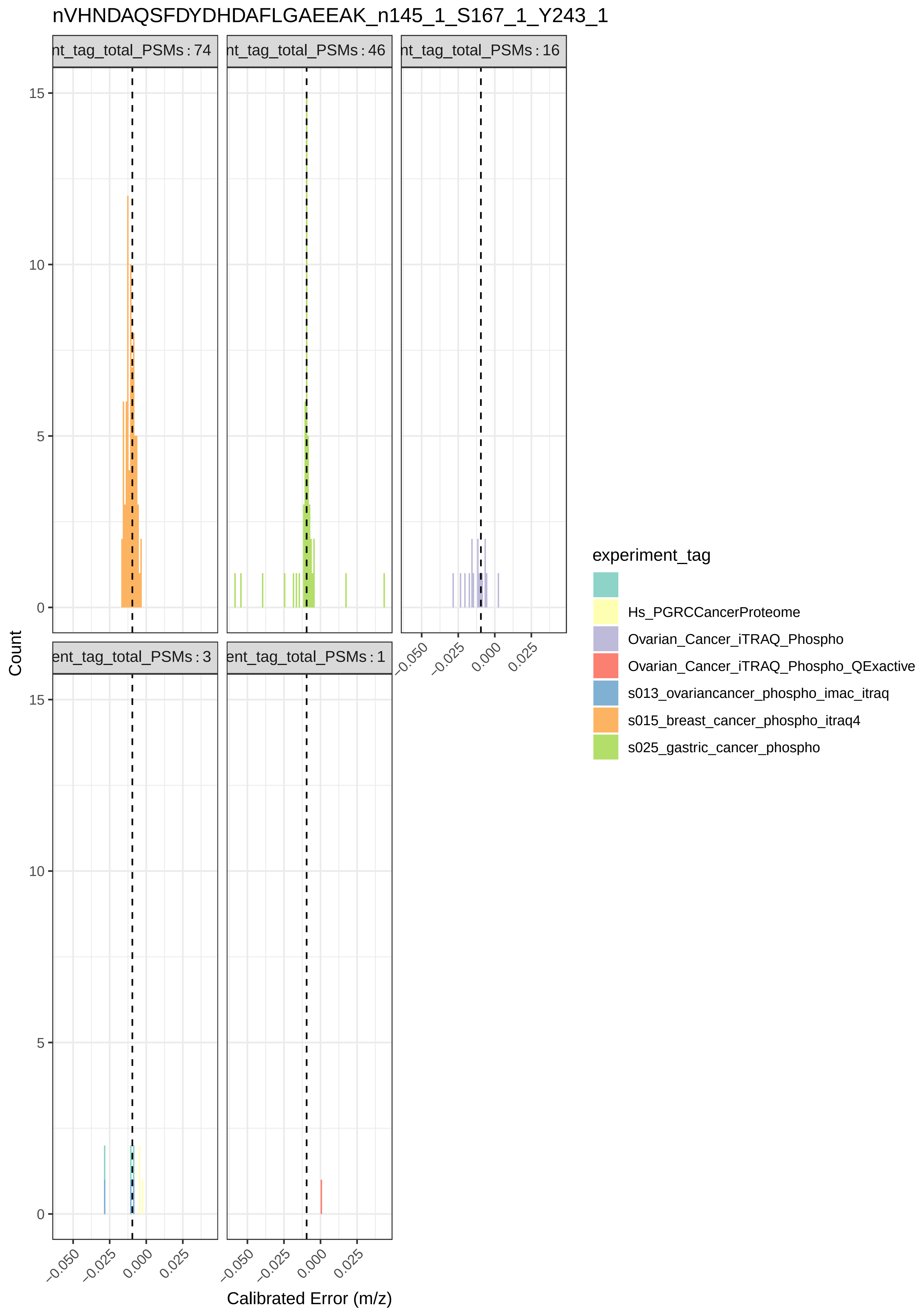
nVHNDAQSFDYDHDAFLGAEEAK_n145_1_S167_1_Y243_1
nt_tag_total_PSMs : 74
nt_tag_total_PSMs : 46
nt_tag_total_PSMs : 16
ent_tag_total_PSMs : 3
ent_tag_total_PSMs : 1
Count
Calibrated Error (m/z)
0
5
10
15
−0.050
−0.025
0.000
0.025
experiment_tag
Hs_PGRCCancerProteome
Ovarian_Cancer_iTRAQ_Phospho
Ovarian_Cancer_iTRAQ_Phospho_QExactive
s013_ovariancancer_phospho_imac_itraq
s015_breast_cancer_phospho_itraq4
s025_gastric_cancer_phospho

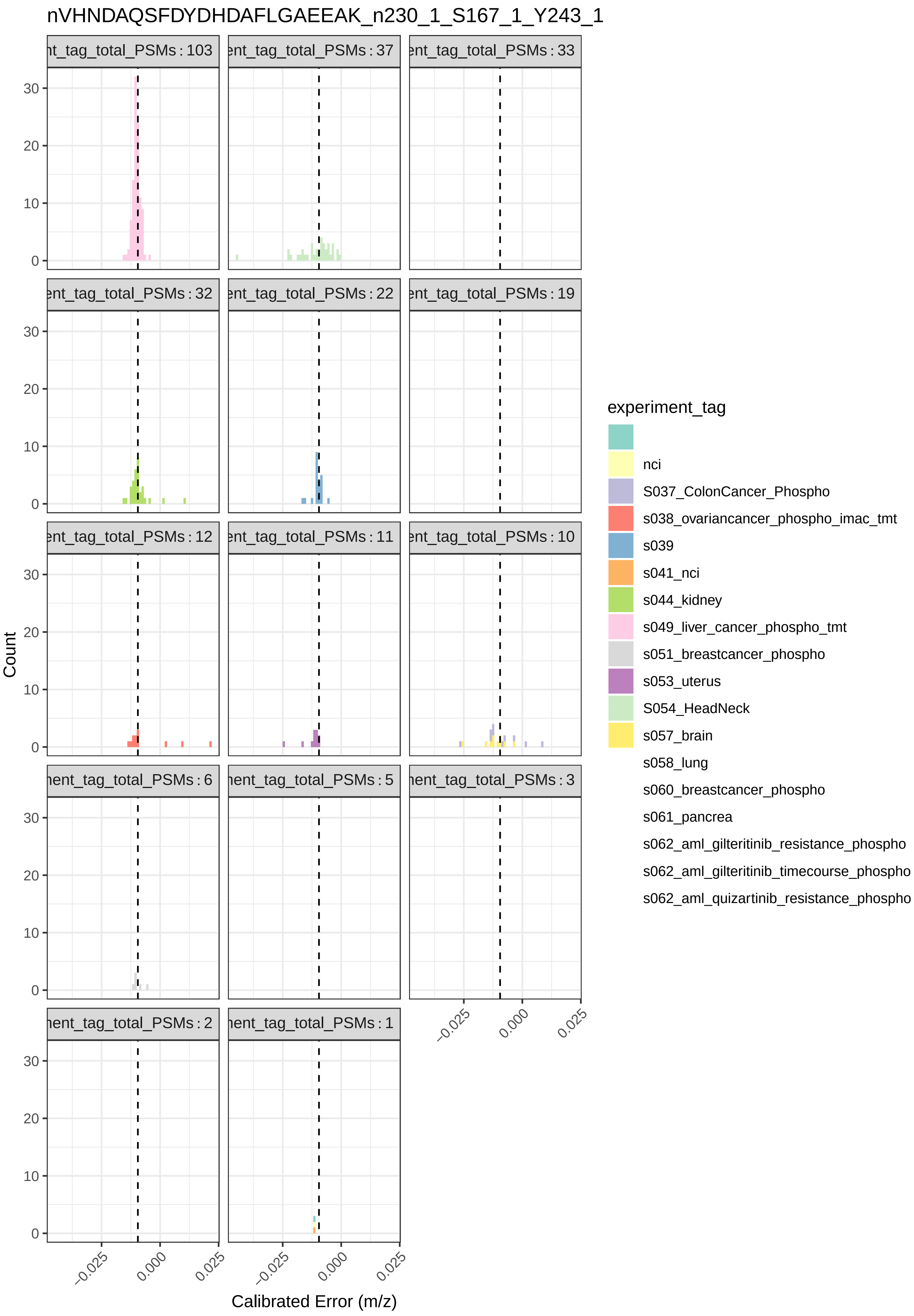
nVHNDAQSFDYDHDAFLGAEEAK_n230_1_S167_1_Y243_1
nt_tag_total_PSMs : 103
ent_tag_total_PSMs : 37
ent_tag_total_PSMs : 33
ent_tag_total_PSMs : 32
ent_tag_total_PSMs : 22
ent_tag_total_PSMs : 19
ent_tag_total_PSMs : 12
ent_tag_total_PSMs : 11
ent_tag_total_PSMs : 10
nent_tag_total_PSMs : 6
nent_tag_total_PSMs : 5
nent_tag_total_PSMs : 3
nent_tag_total_PSMs : 2
nent_tag_total_PSMs : 1
Count
Calibrated Error (m/z)
experiment_tag
nci
S037_ColonCancer_Phospho
s038_ovariancancer_phospho_imac_tmt
s039
s041_nci
s044_kidney
s049_liver_cancer_phospho_tmt
s051_breastcancer_phospho
s053_uterus
S054_HeadNeck
s057_brain
s058_lung
s060_breastcancer_phospho
s061_pancrea
s062_aml_gilteritinib_resistance_phospho
s062_aml_gilteritinib_timecourse_phospho
s062_aml_quizartinib_resistance_phospho

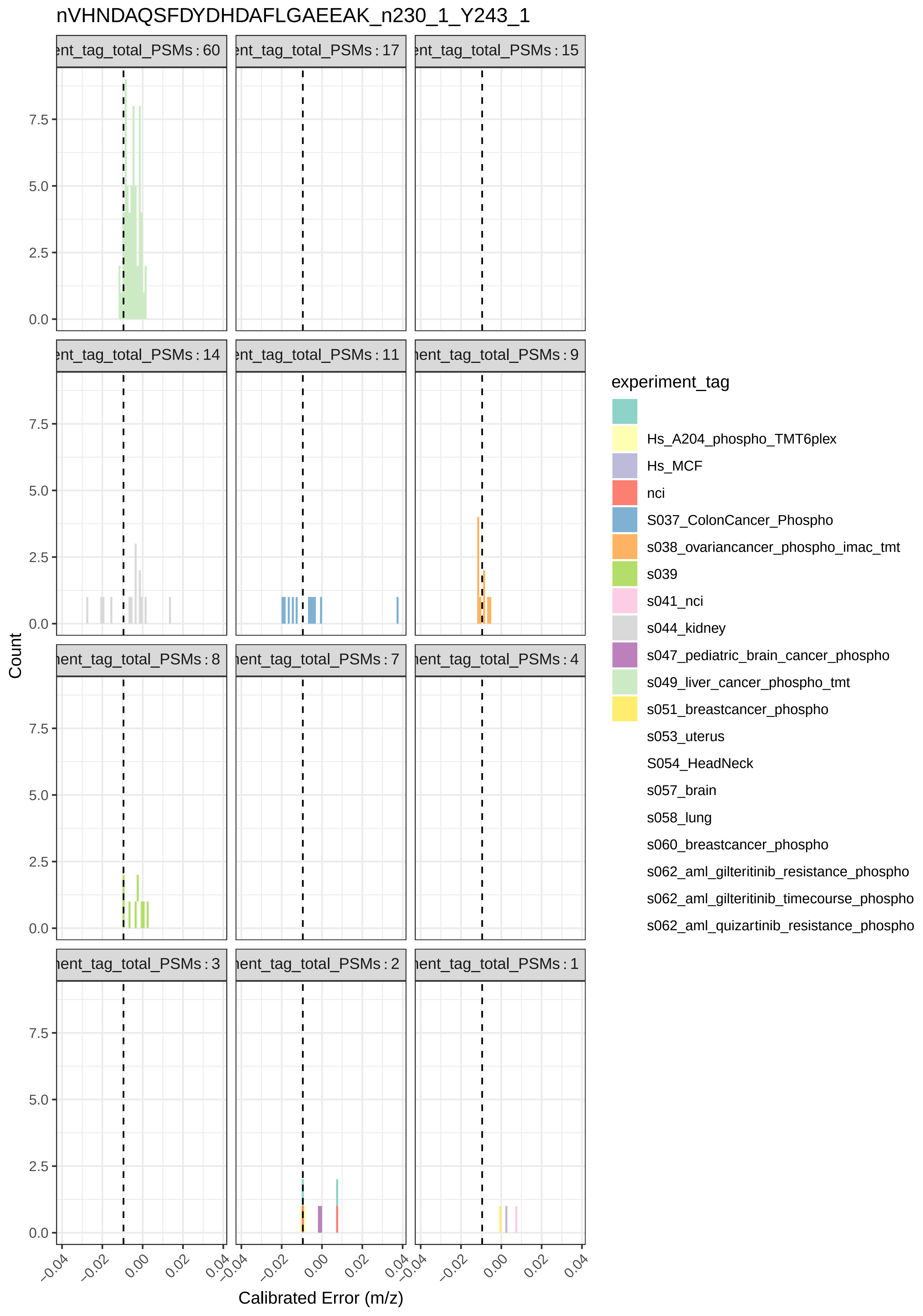
nVHNDAQSFDYDHDAFLGAEEAK_n230_1_Y243_1
ent_tag_total_PSMs : 60
ent_tag_total_PSMs : 17
ent_tag_total_PSMs : 15
ent_tag_total_PSMs : 14
ent_tag_total_PSMs : 11
nent_tag_total_PSMs : 9
nent_tag_total_PSMs : 8
nent_tag_total_PSMs : 7
nent_tag_total_PSMs : 4
nent_tag_total_PSMs : 3
nent_tag_total_PSMs : 2
nent_tag_total_PSMs : 1
Count
Calibrated Error (m/z)
experiment_tag
Hs_A204_phospho_TMT6plex
Hs_MCF
nci
S037_ColonCancer_Phospho
s038_ovariancancer_phospho_imac_tmt
s039
s041_nci
s044_kidney
s047_pediatric_brain_cancer_phospho
s049_liver_cancer_phospho_tmt
s051_breastcancer_phospho
s053_uterus
S054_HeadNeck
s057_brain
s058_lung
s060_breastcancer_phospho
s062_aml_gilteritinib_resistance_phospho
s062_aml_gilteritinib_timecourse_phospho
s062_aml_quizartinib_resistance_phospho

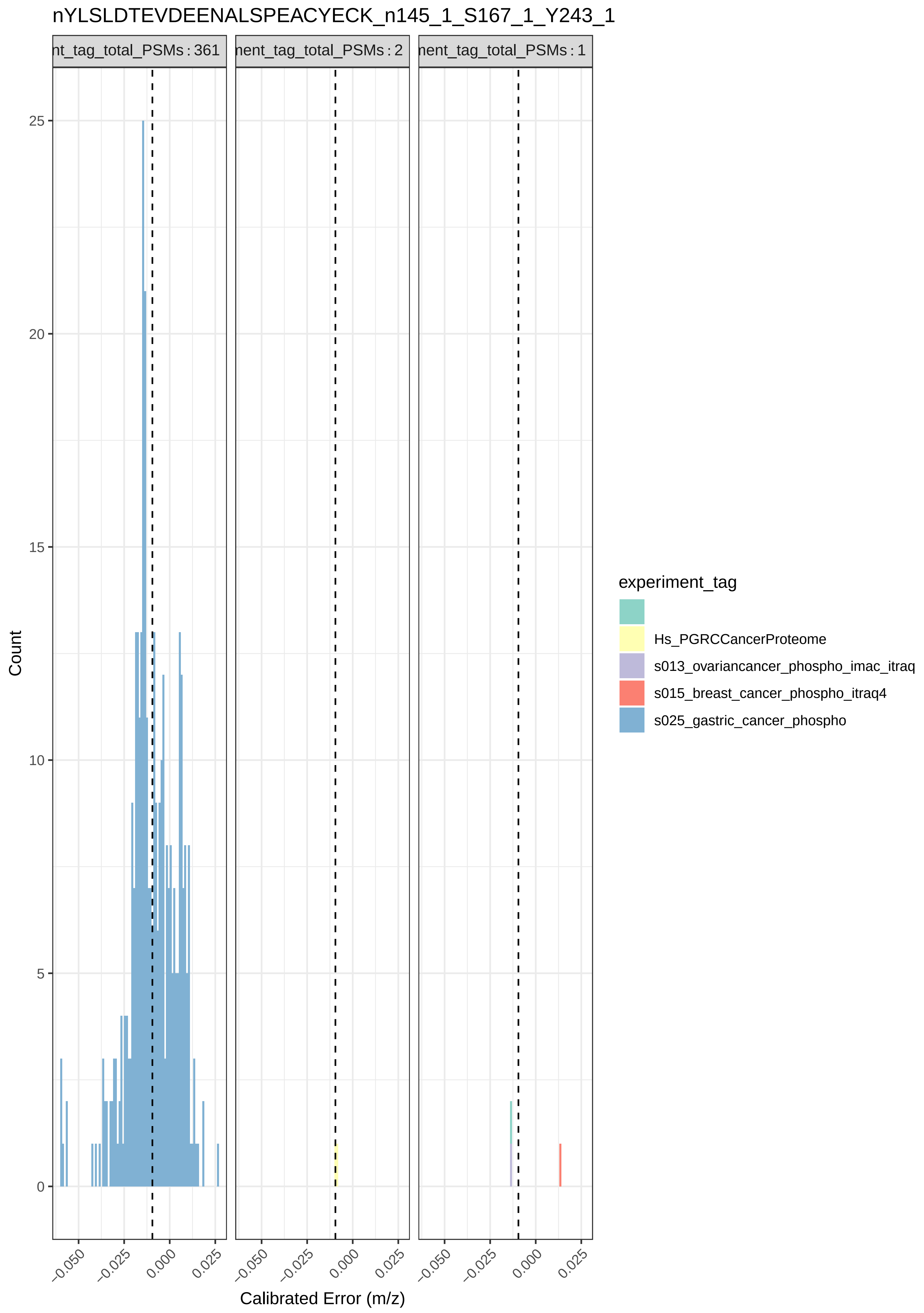

nYLSLDTEVDEENALSPEACYECK_n145_1_S167_1_Y243_1
nt_tag_total_PSMs : 361
nent_tag_total_PSMs : 2
nent_tag_total_PSMs : 1
Count
0
5
10
15
20
25
−0.050
−0.025
0.000
0.025
Calibrated Error (m/z)
experiment_tag
Hs_PGRCCancerProteome
s013_ovariancancer_phospho_imac_itraq
s015_breast_cancer_phospho_itraq4
s025_gastric_cancer_phospho

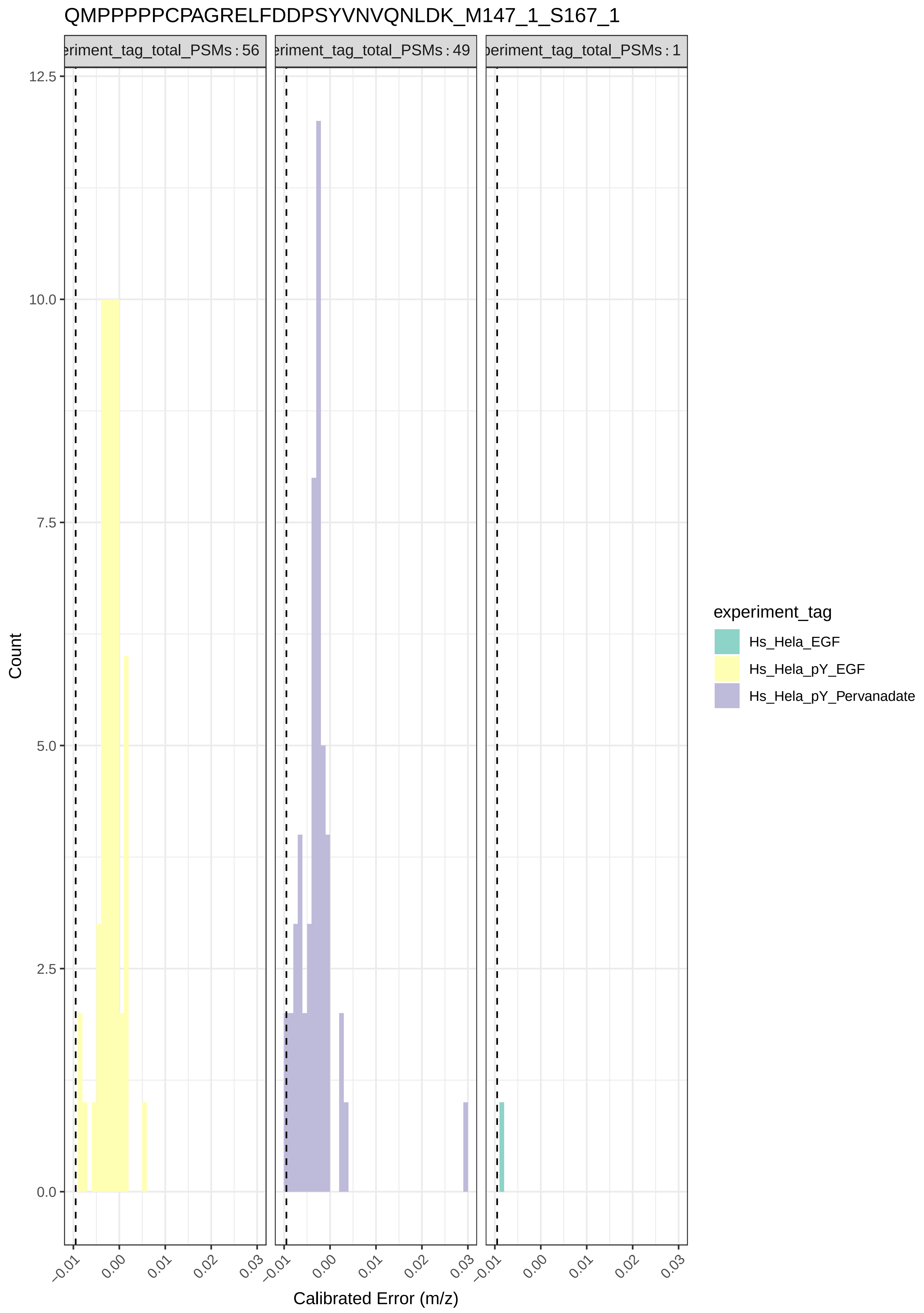

QMPPPPPCPAGRELFDDPSYVNVQNLDK_M147_1_S167_1
eriment_tag_total_PSMs : 56
eriment_tag_total_PSMs : 49
periment_tag_total_PSMs : 1
Count
Calibrated Error (m/z)
experiment_tag
Hs_Hela_EGF
Hs_Hela_pY_EGF
Hs_Hela_pY_Pervanadate

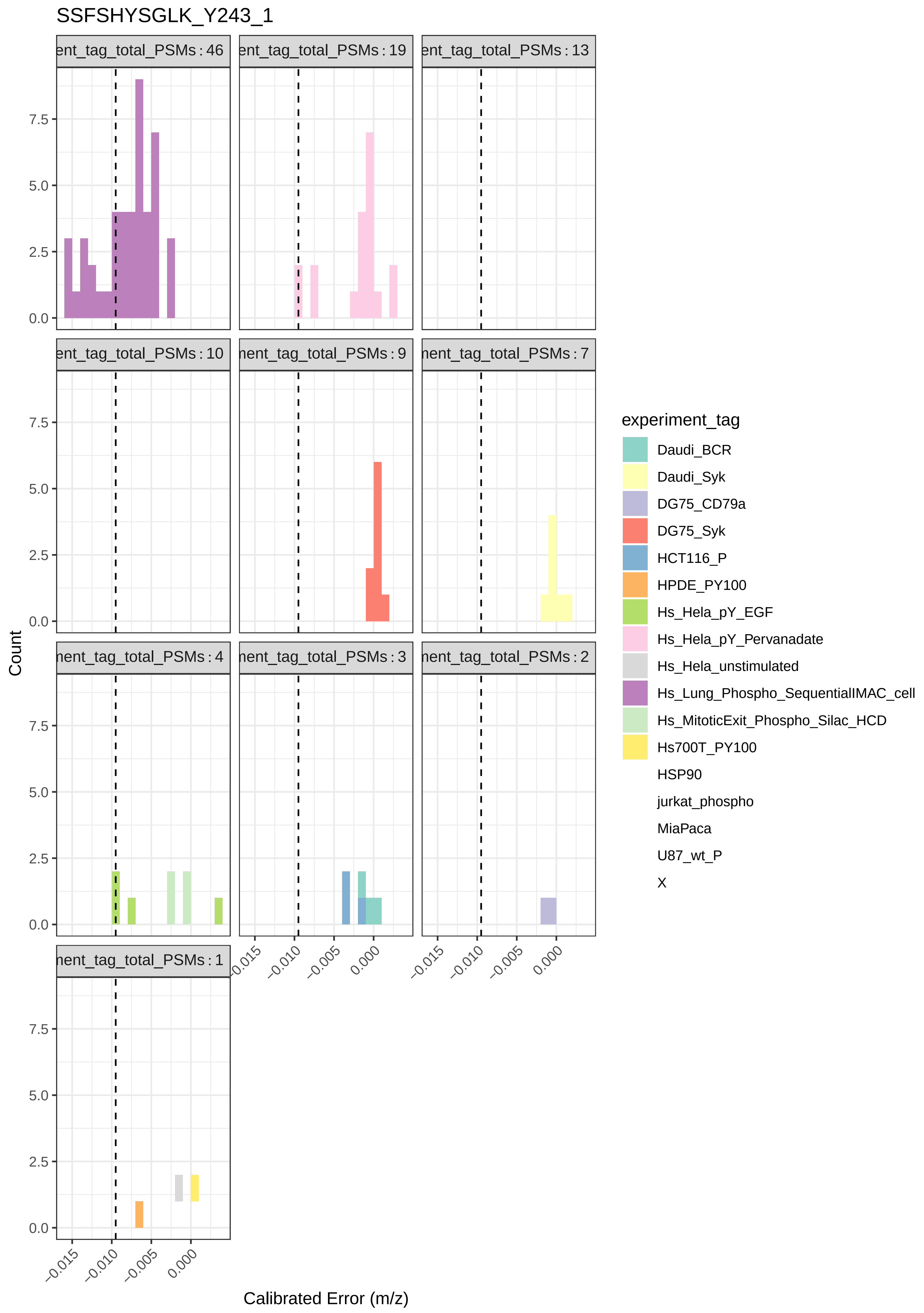
SSFSHYSGLK_Y243_1
ent_tag_total_PSMs : 46
ent_tag_total_PSMs : 19
ent_tag_total_PSMs : 13
ent_tag_total_PSMs : 10
nent_tag_total_PSMs : 9
nent_tag_total_PSMs : 7
nent_tag_total_PSMs : 4
nent_tag_total_PSMs : 3
nent_tag_total_PSMs : 2
nent_tag_total_PSMs : 1
Count
0.0
2.5
5.0
7.5
−0.015
−0.010
−0.005
0.000
Calibrated Error (m/z)
experiment_tag
Daudi_BCR
Daudi_Syk
DG75_CD79a
DG75_Syk
HCT116_P
HPDE_PY100
Hs_Hela_pY_EGF
Hs_Hela_pY_Pervanadate
Hs_Hela_unstimulated
Hs_Lung_Phospho_SequentialIMAC_cell
Hs_MitoticExit_Phospho_Silac_HCD
Hs700T_PY100
HSP90
jurkat_phospho
MiaPaca
U87_wt_P
X

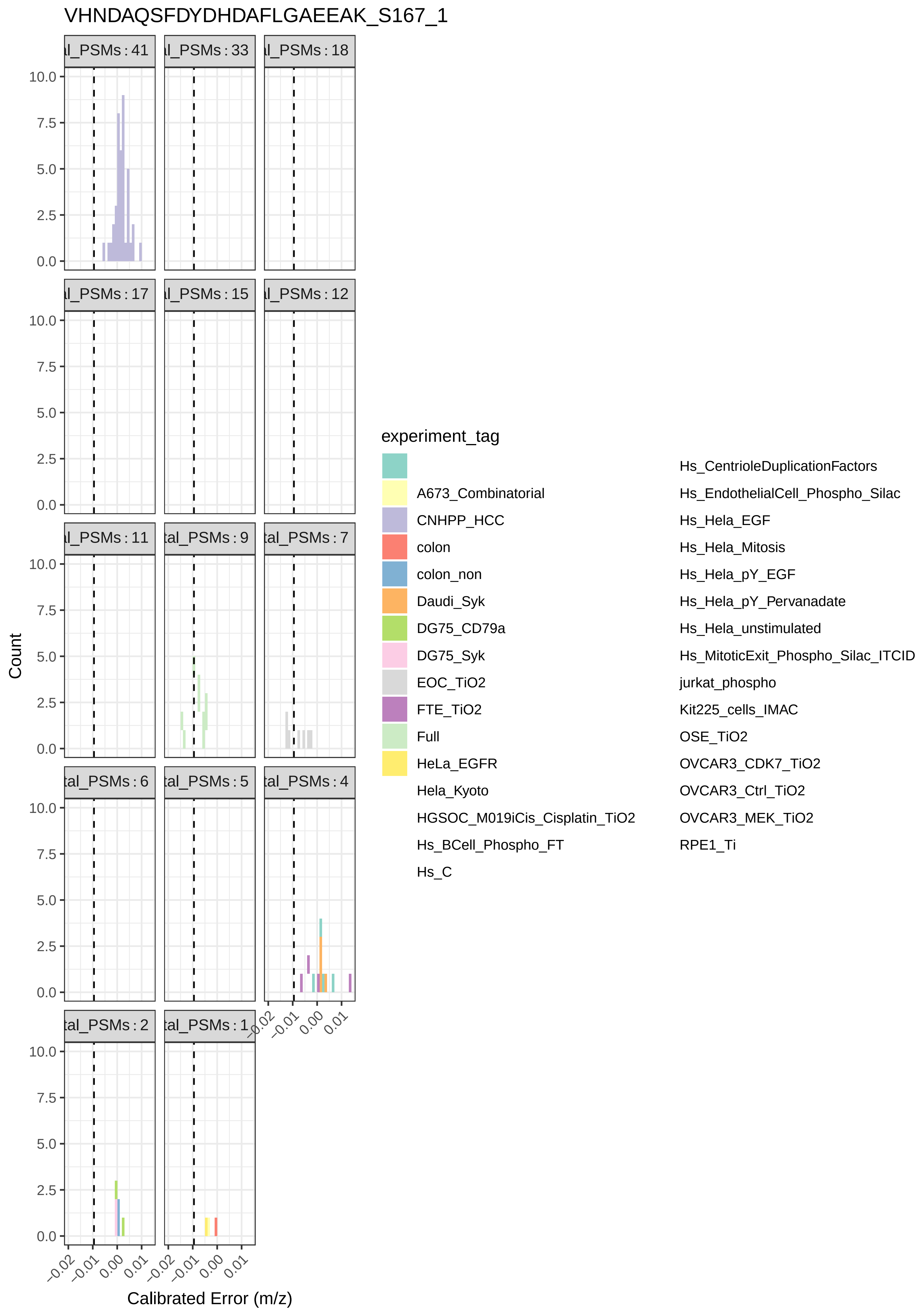
VHNDAQSFDYDHDAFLGAEEAK_S167_1
l_PSMs : 41
l_PSMs : 33
l_PSMs : 18
l_PSMs : 17
l_PSMs : 15
l_PSMs : 12
l_PSMs : 11
tal_PSMs : 9
tal_PSMs : 7
tal_PSMs : 6
tal_PSMs : 5
tal_PSMs : 4
tal_PSMs : 2
tal_PSMs : 1
Count
Calibrated Error (m/z)
experiment_tag
A673_Combinatorial
CNHPP_HCC
colon
colon_non
Daudi_Syk
DG75_CD79a
DG75_Syk
EOC_TiO2
FTE_TiO2
Full
HeLa_EGFR
Hela_Kyoto
HGSOC_M019iCis_Cisplatin_TiO2
Hs_BCell_Phospho_FT
Hs_C
Hs_CentrioleDuplicationFactors
Hs_EndothelialCell_Phospho_Silac
Hs_Hela_EGF
Hs_Hela_Mitosis
Hs_Hela_pY_EGF
Hs_Hela_pY_Pervanadate
Hs_Hela_unstimulated
Hs_MitoticExit_Phospho_Silac_ITCID
jurkat_phospho
Kit225_cells_IMAC
OSE_TiO2
OVCAR3_CDK7_TiO2
OVCAR3_Ctrl_TiO2
OVCAR3_MEK_TiO2
RPE1_Ti

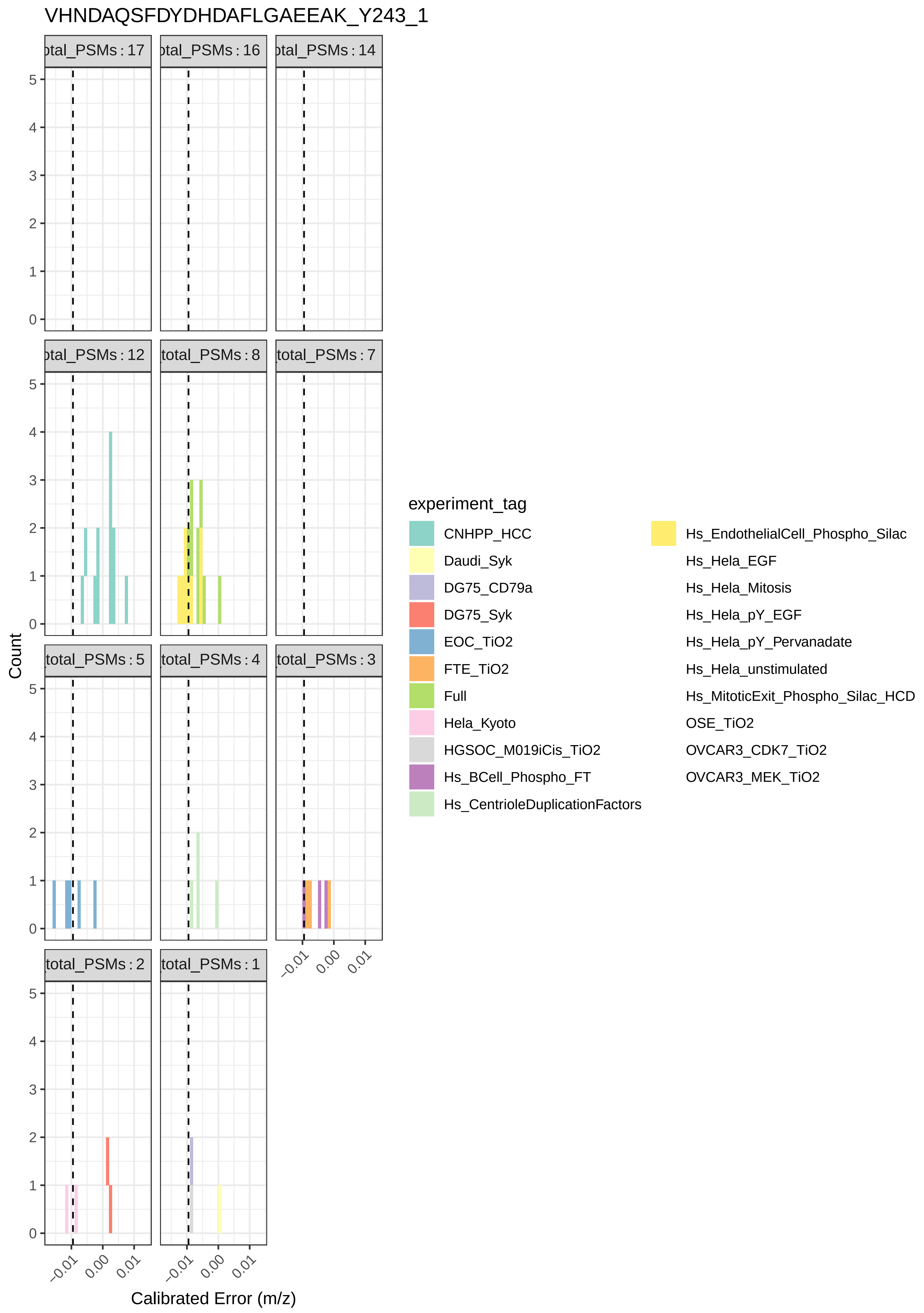

VHNDAQSFDYDHDAFLGAEEAK_Y243_1
otal_PSMs : 17
otal_PSMs : 16
otal_PSMs : 14
otal_PSMs : 12
total_PSMs : 8
total_PSMs : 7
total_PSMs : 5
total_PSMs : 4
total_PSMs : 3
total_PSMs : 2
total_PSMs : 1
Count
Calibrated Error (m/z)
−0.01
0.00
0.01
experiment_tag
CNHPP_HCC
Daudi_Syk
DG75_CD79a
DG75_Syk
EOC_TiO2
FTE_TiO2
Full
Hela_Kyoto
HGSOC_M019iCis_TiO2
Hs_BCell_Phospho_FT
Hs_CentrioleDuplicationFactors
Hs_EndothelialCell_Phospho_Silac
Hs_Hela_EGF
Hs_Hela_Mitosis
Hs_Hela_pY_EGF
Hs_Hela_pY_Pervanadate
Hs_Hela_unstimulated
Hs_MitoticExit_Phospho_Silac_HCD
OSE_TiO2
OVCAR3_CDK7_TiO2
OVCAR3_MEK_TiO2
