## Supplementary File 2 for "Searching for Sulfotyrosines (sY) in a HA(pY)STACK": faceted_plots_by_instrument.pdf

APVNWYQEK_N115_1_Y243_1

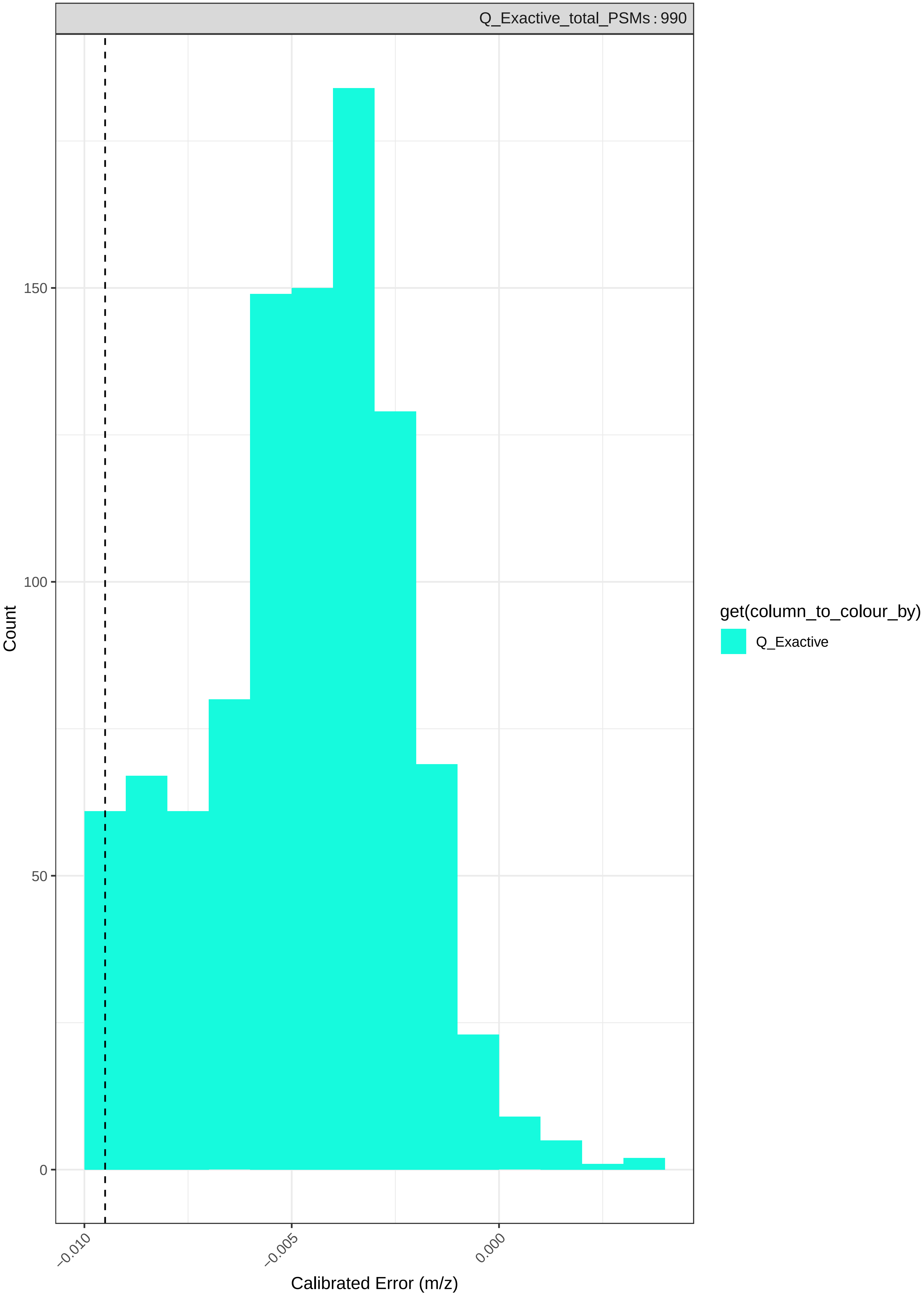

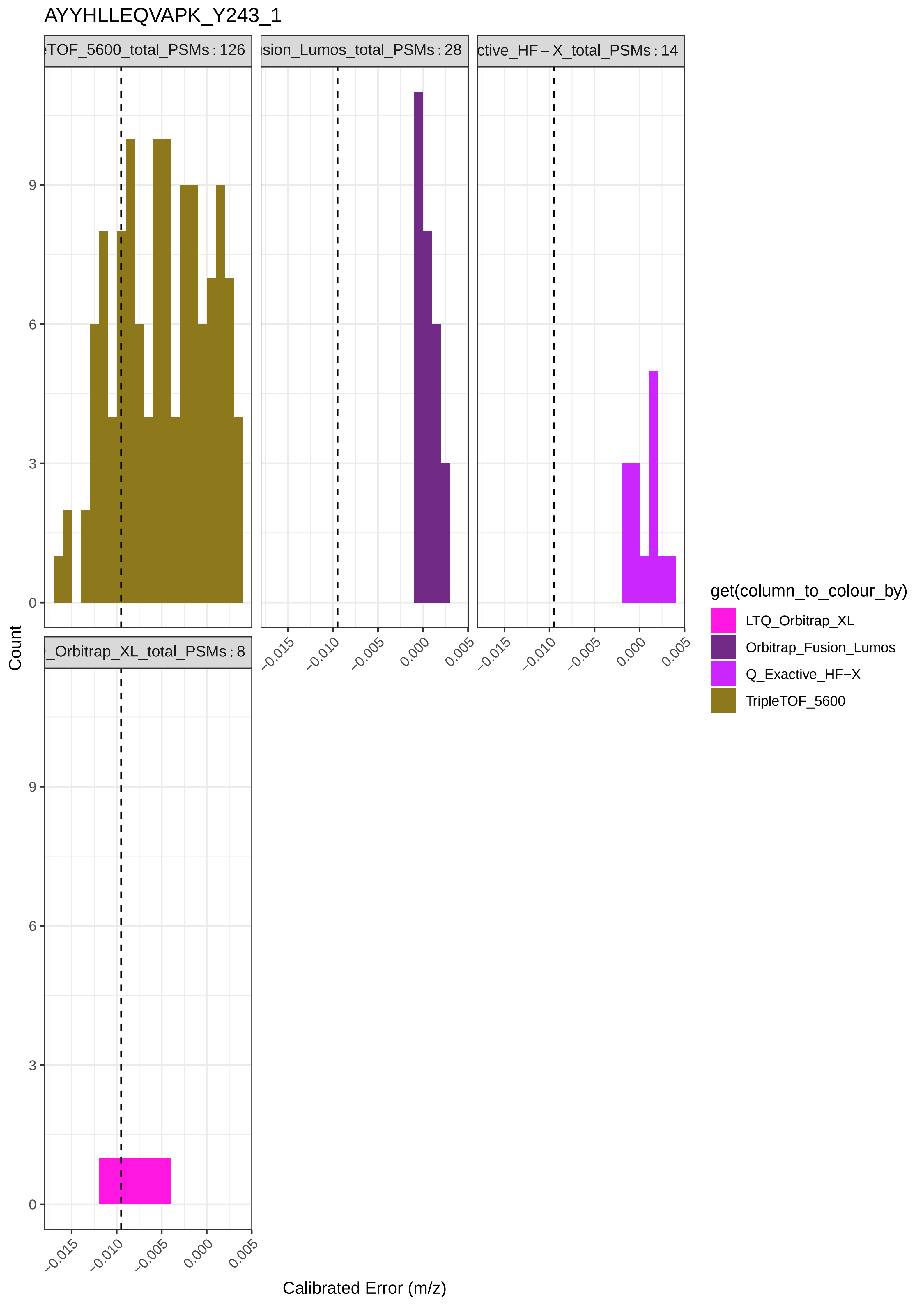

AYYHLLEQVAPK_Y243_1
TOF_5600_total_PSMs : 126
sion_Lumos_total_PSMs : 28
ctive_HF – X_total_PSMs : 14
_Orbitrap_XL_total_PSMs : 8
Count
Calibrated Error (m/z)
0
3
6
9
–0.015
–0.010
–0.005
0.000
0.005
get(column_to_colour_by)
LTQ_Orbitrap_XL
Orbitrap_Fusion_Lumos
Q_Exactive_HF−X
TripleTOF_5600

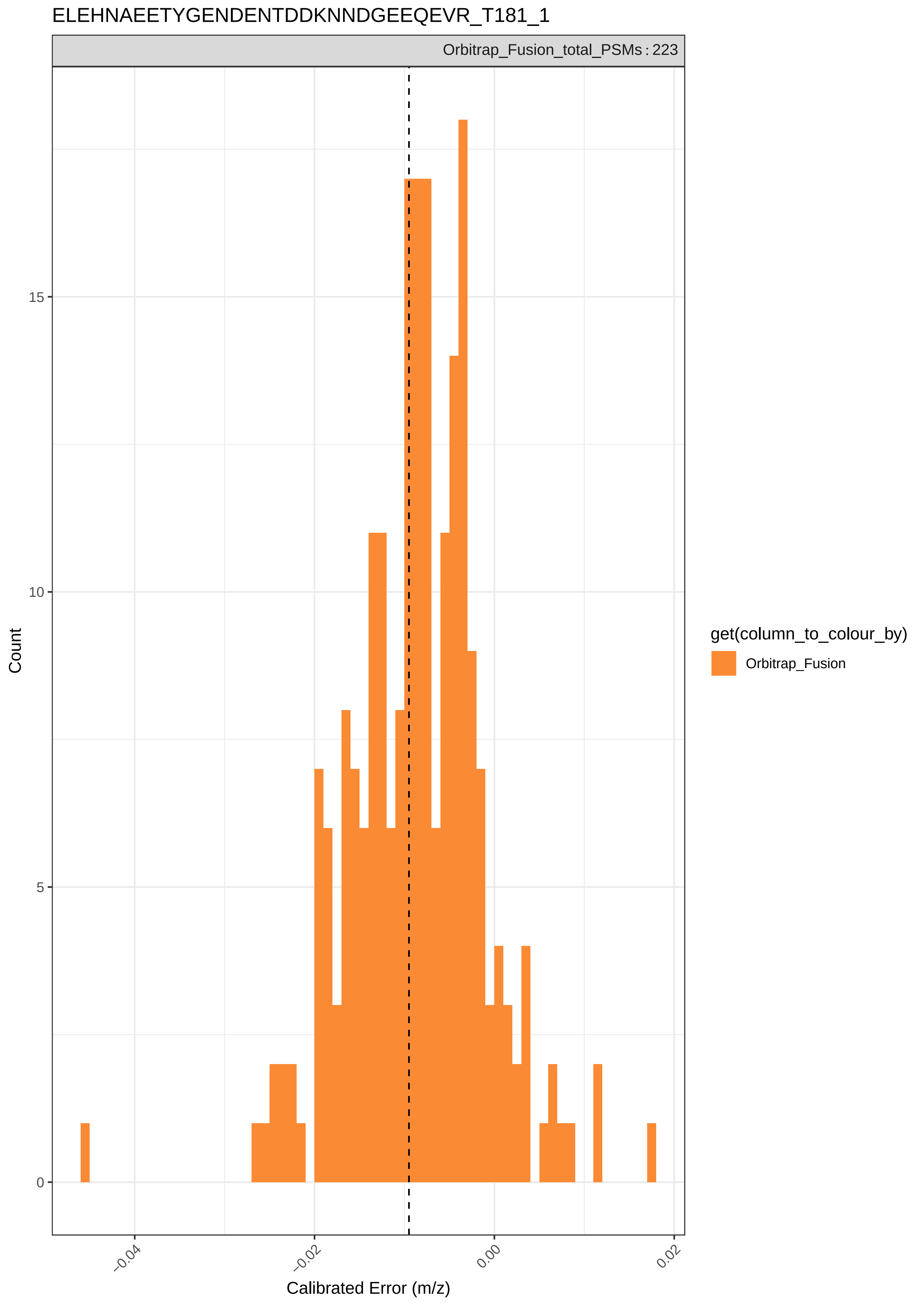

ELEHNAEETYGENDENTDDKNNDGEEQEVR_T181_1
Orbitrap_Fusion_total_PSMs : 223
Count
15
10
5
0
–0.04
–0.02
0.00
0.02
Calibrated Error (m/z)
get(column_to_colour_by)
Orbitrap_Fusion

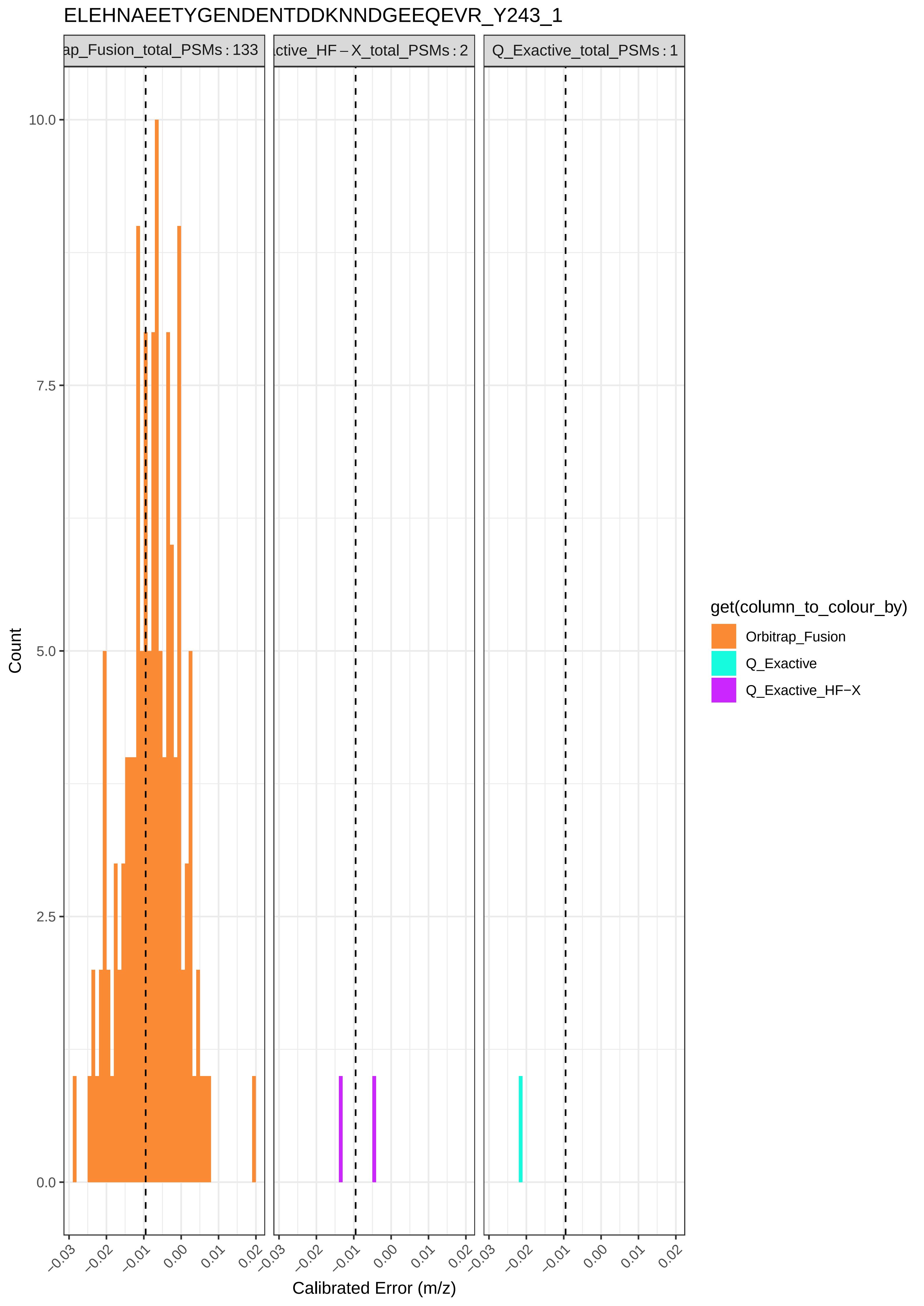

ELEHNAEETYGENDENTDDKNNDGEEQEVR_Y243_1
ap_Fusion_total_PSMs : 133
ctive_HF – X_total_PSMs : 2
Q_Exactive_total_PSMs : 1
10.0
7.5
5.0
2.5
0.0
Count
−0.03
−0.02
−0.01
0.00
0.01
0.02
Calibrated Error (m/z)
get(column_to_colour_by)
Orbitrap_Fusion
Q_Exactive
Q_Exactive_HF−X

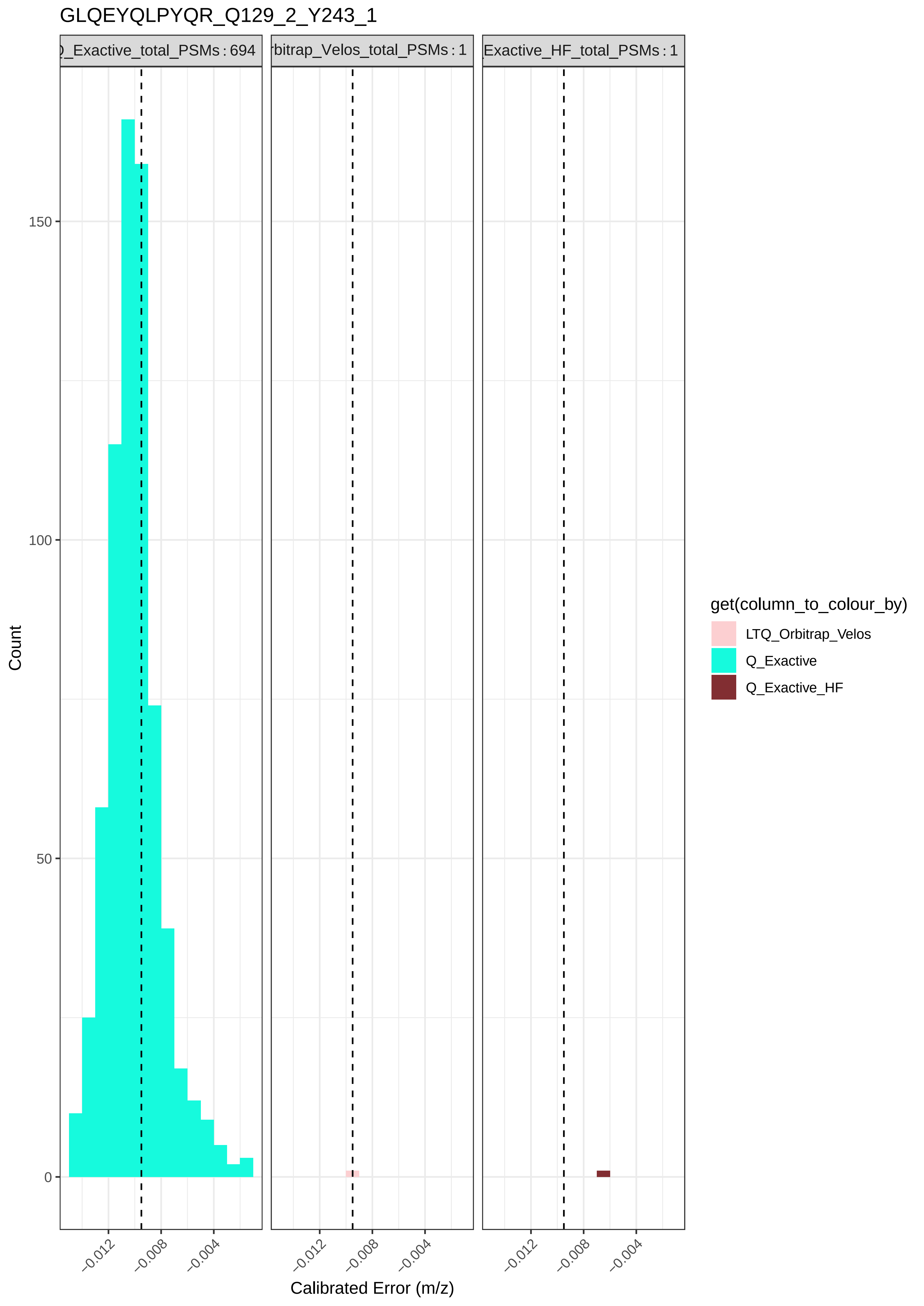

GLQEYQLPYQR_Q129_2_Y243_1
)_Exactive_total_PSMs : 694
rbitrap_Velos_total_PSMs : 1
Exactive_HF_total_PSMs : 1
Count
150
100
50
0
−0.012
−0.008
−0.004
Calibrated Error (m/z)
get(column_to_colour_by)
LTQ_Orbitrap_Velos
Q_Exactive
Q_Exactive_HF

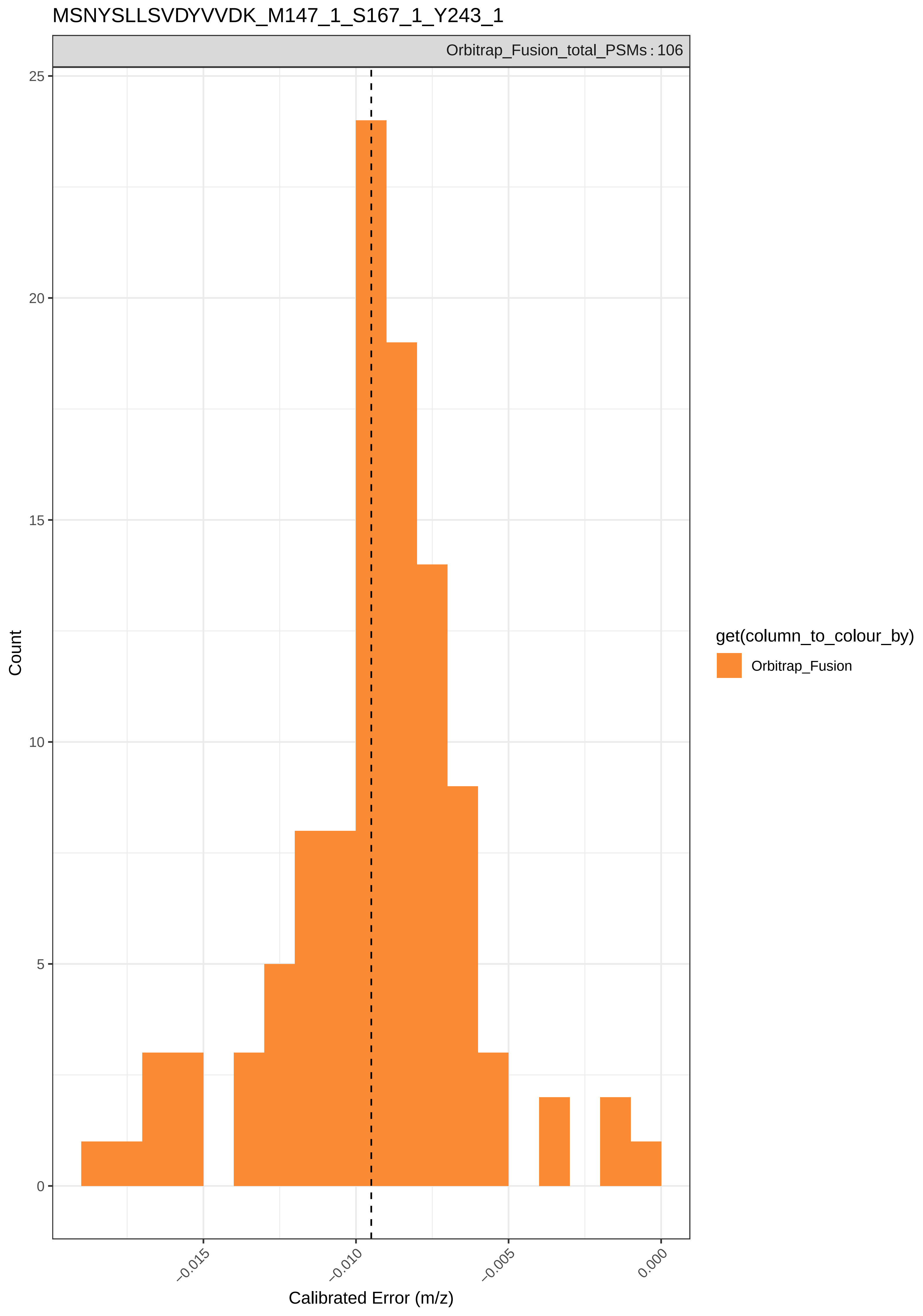

MSNYSLLSVDYVVDK_M147_1_S167_1_Y243_1
Orbitrap_Fusion_total_PSMs : 106
Count
25
20
15
10
5
0
−0.015
−0.010
−0.005
0.000
Calibrated Error (m/z)
get(column_to_colour_by)
Orbitrap_Fusion

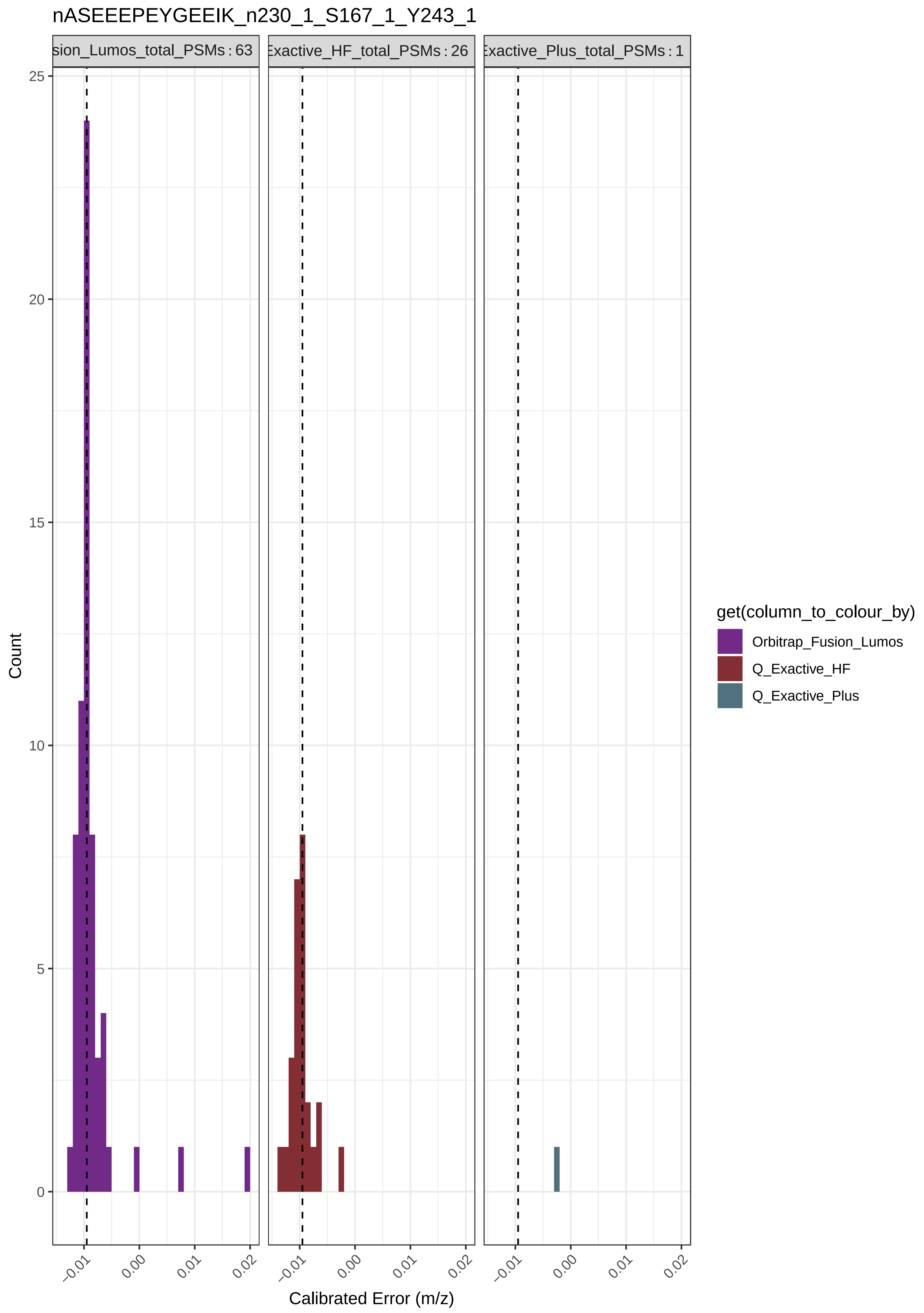

nASEEEPEYGEEIK_n230_1_S167_1_Y243_1
sion_Lumos_total_PSMs : 63
Exactive_HF_total_PSMs : 26
Exactive_Plus_total_PSMs : 1
Count
25
20
15
10
5
0
−0.01
0.00
0.01
0.02
Calibrated Error (m/z)
get(column_to_colour_by)
Orbitrap_Fusion_Lumos
Q_Exactive_HF
Q_Exactive_Plus

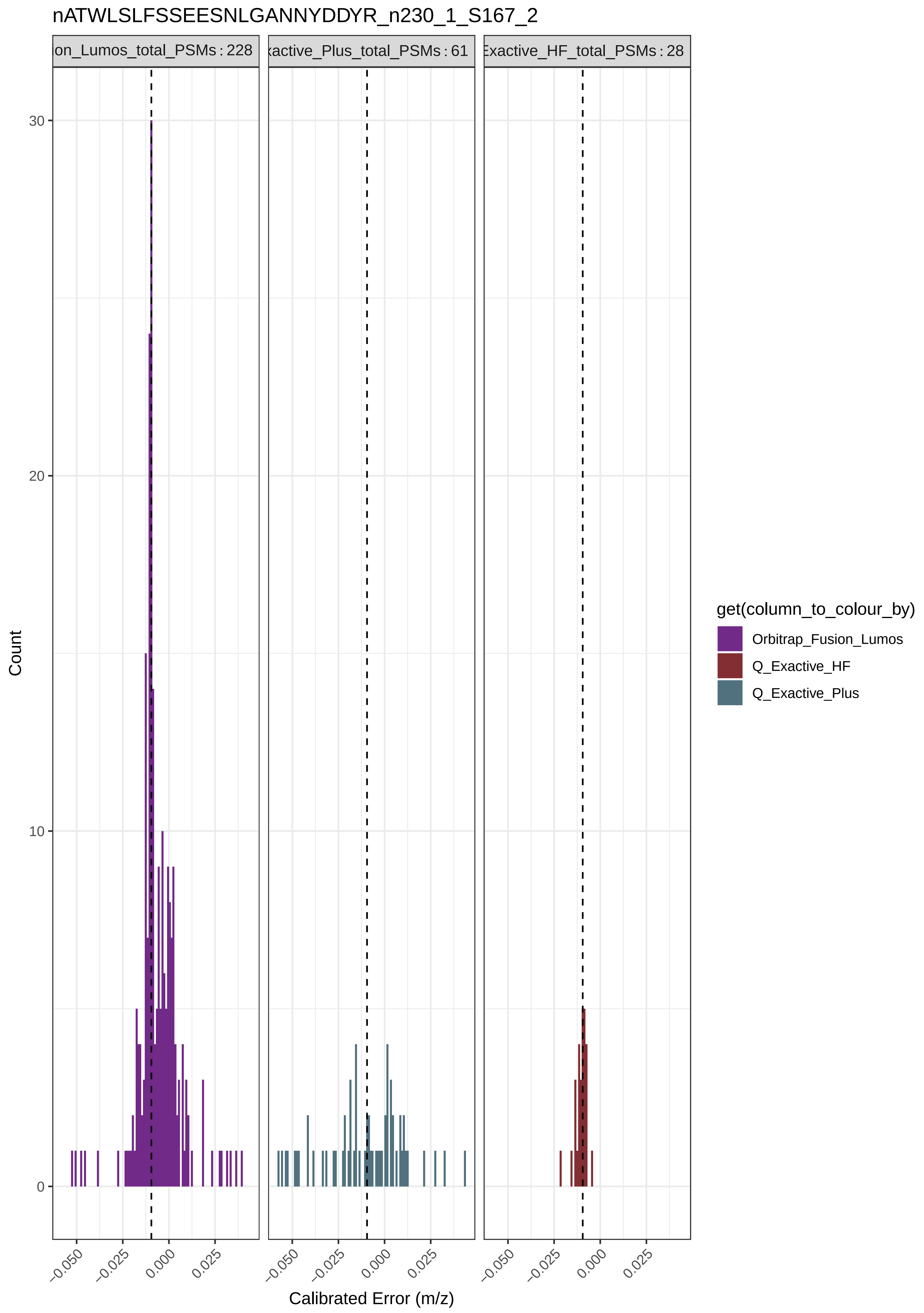
nATWLSLFSSEESNLGANNYDDYR_n230_1_S167_2
ion_Lumos_total_PSMs : 228
xactive_Plus_total_PSMs : 61
Exactive_HF_total_PSMs : 28
Count
0
10
20
30
−0.050
−0.025
0.000
0.025
Calibrated Error (m/z)
get(column_to_colour_by)
Orbitrap_Fusion_Lumos
Q_Exactive_HF
Q_Exactive_Plus

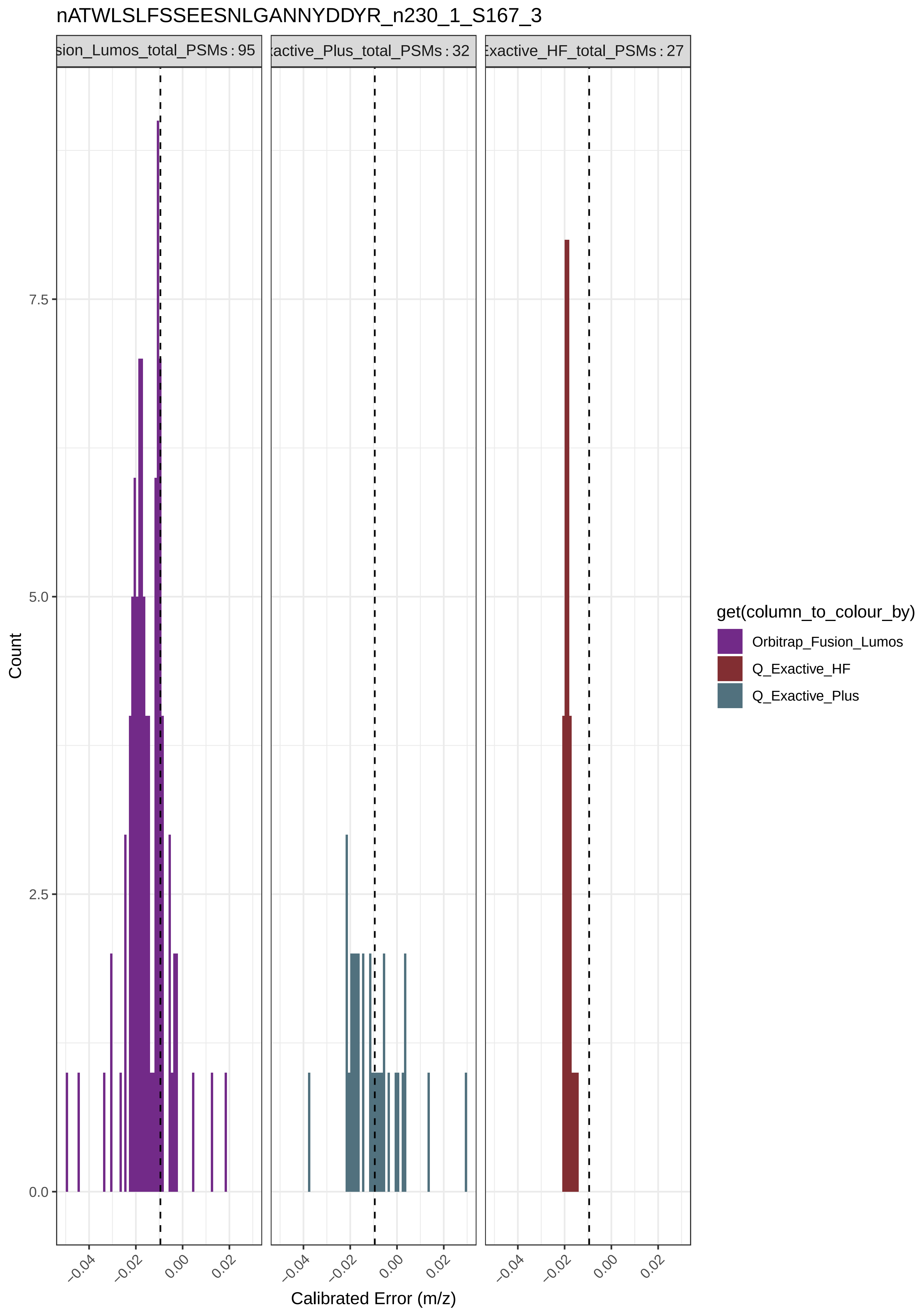
nATWLSLFSSEESNLGANNYDDYR_n230_1_S167_3
sion_Lumos_total_PSMs : 95
:active_Plus_total_PSMs : 32
Exactive_HF_total_PSMs : 27
Count
0.0
2.5
5.0
7.5
−0.04
−0.02
0.00
0.02
Calibrated Error (m/z)
get(column_to_colour_by)
Orbitrap_Fusion_Lumos
Q_Exactive_HF
Q_Exactive_Plus

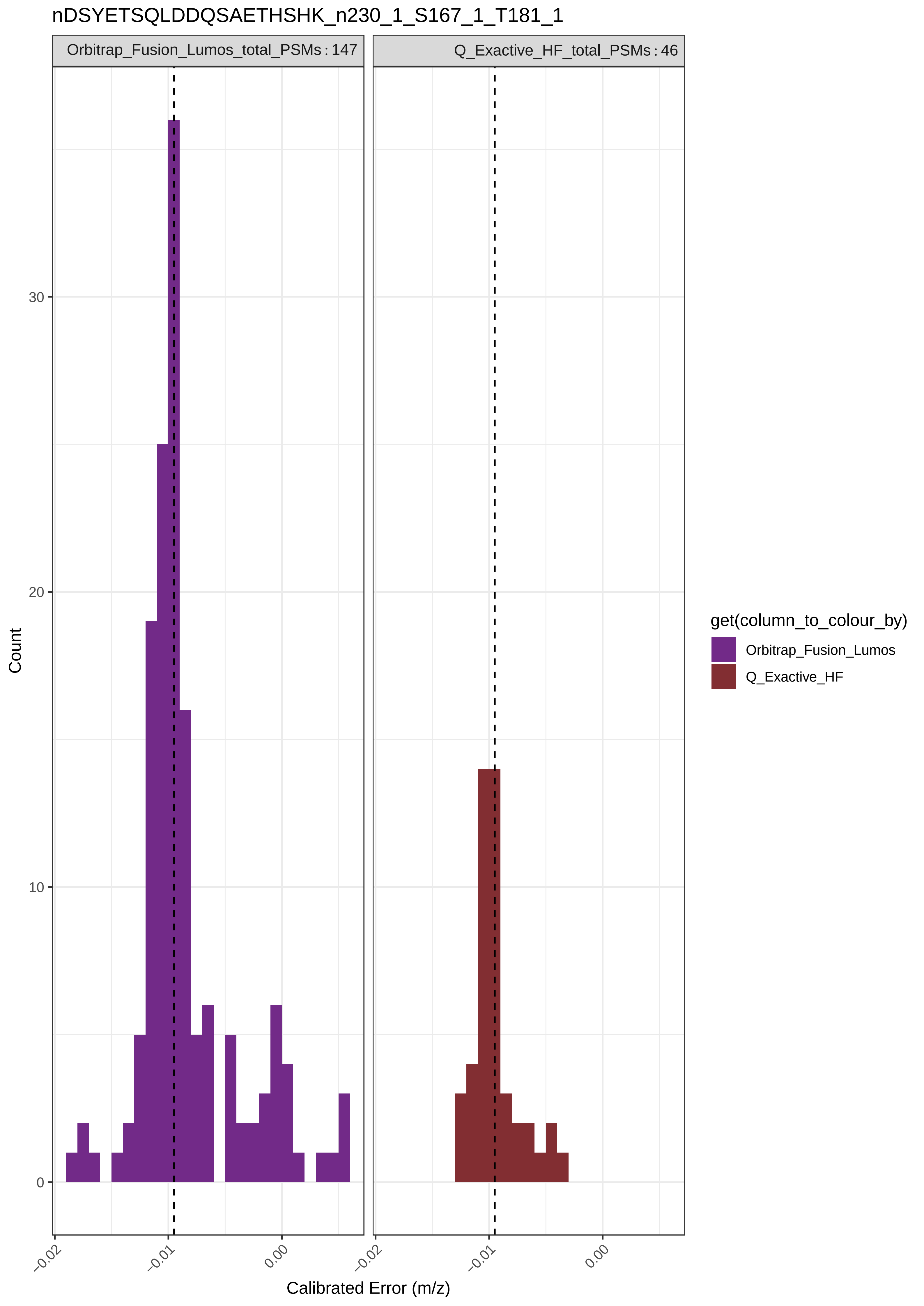

nDSYETSQLDDQSAETHSHK_n230_1_S167_1_T181_1
Orbitrap_Fusion_Lumos_total_PSMs : 147
Q_Exactive_HF_total_PSMs : 46
Count
30
20
10
0
−0.02
−0.01
0.00
−0.02
−0.01
0.00
Calibrated Error (m/z)
get(column_to_colour_by)
Orbitrap_Fusion_Lumos
Q_Exactive_HF

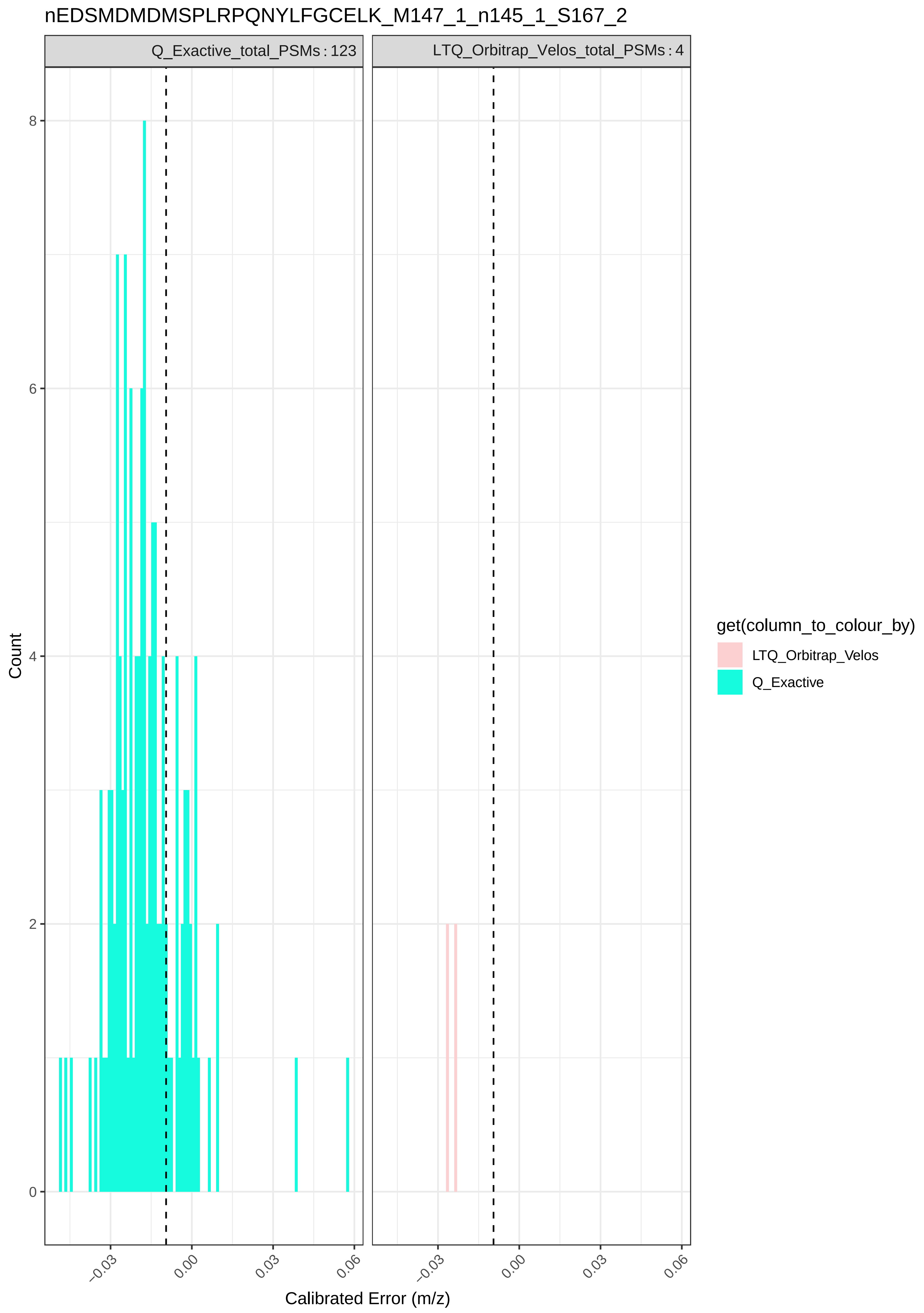
nEDSMDMDMSPLRPQNYLFGCELK_M147_1_n145_1_S167_2
Q_Exactive_total_PSMs : 123
LTQ_Orbitrap_Velos_total_PSMs : 4
Count
0
2
4
6
8
−0.03
0.00
0.03
0.06
Calibrated Error (m/z)
get(column_to_colour_by)
LTQ_Orbitrap_Velos
Q_Exactive

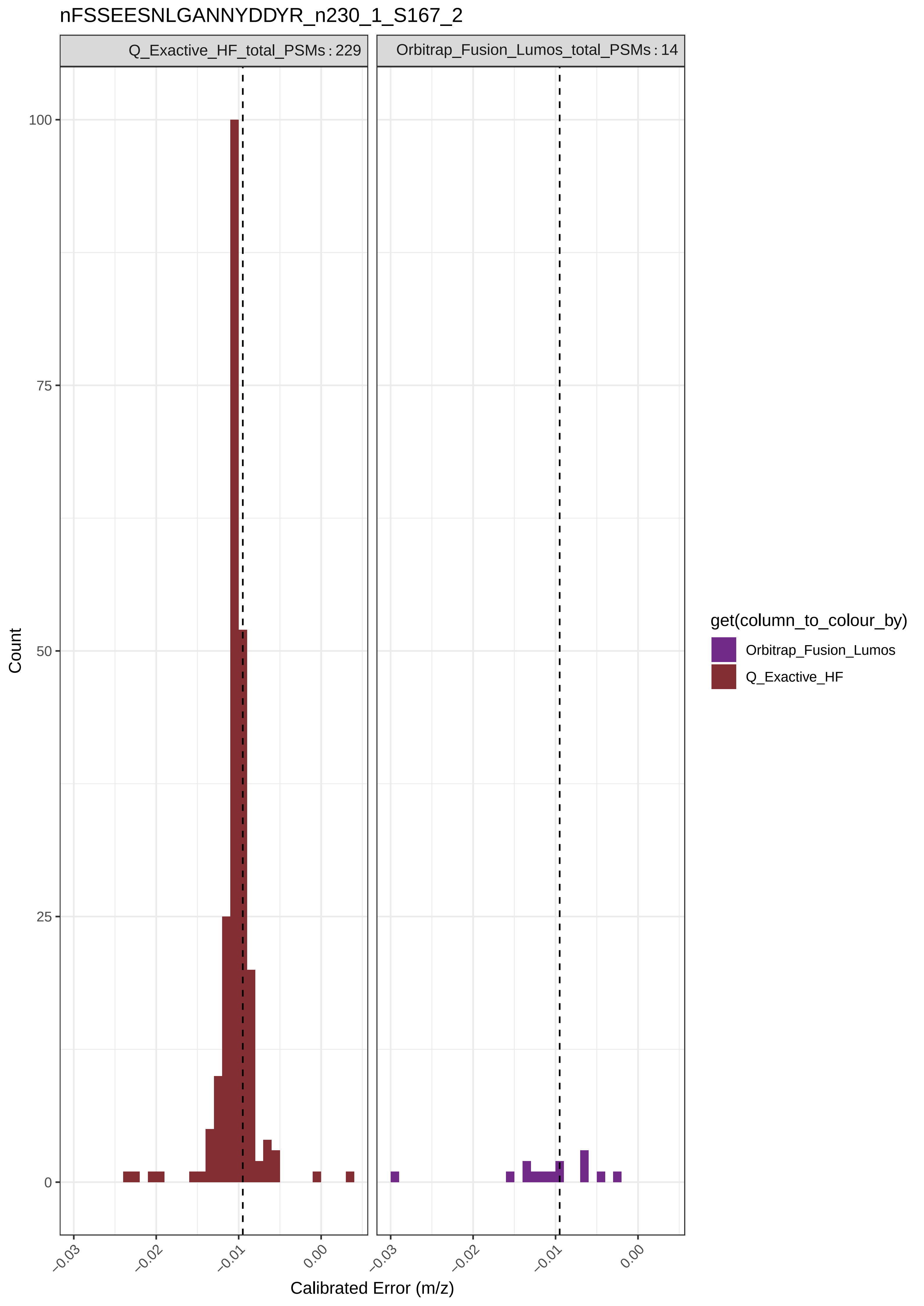
nFSSEESNLGANNYDDYR_n230_1_S167_2
Q_Exactive_HF_total_PSMs : 229
Orbitrap_Fusion_Lumos_total_PSMs : 14
Count
100
75
50
25
0
−0.03
−0.02
−0.01
0.00
Calibrated Error (m/z)
get(column_to_colour_by)
Orbitrap_Fusion_Lumos
Q_Exactive_HF

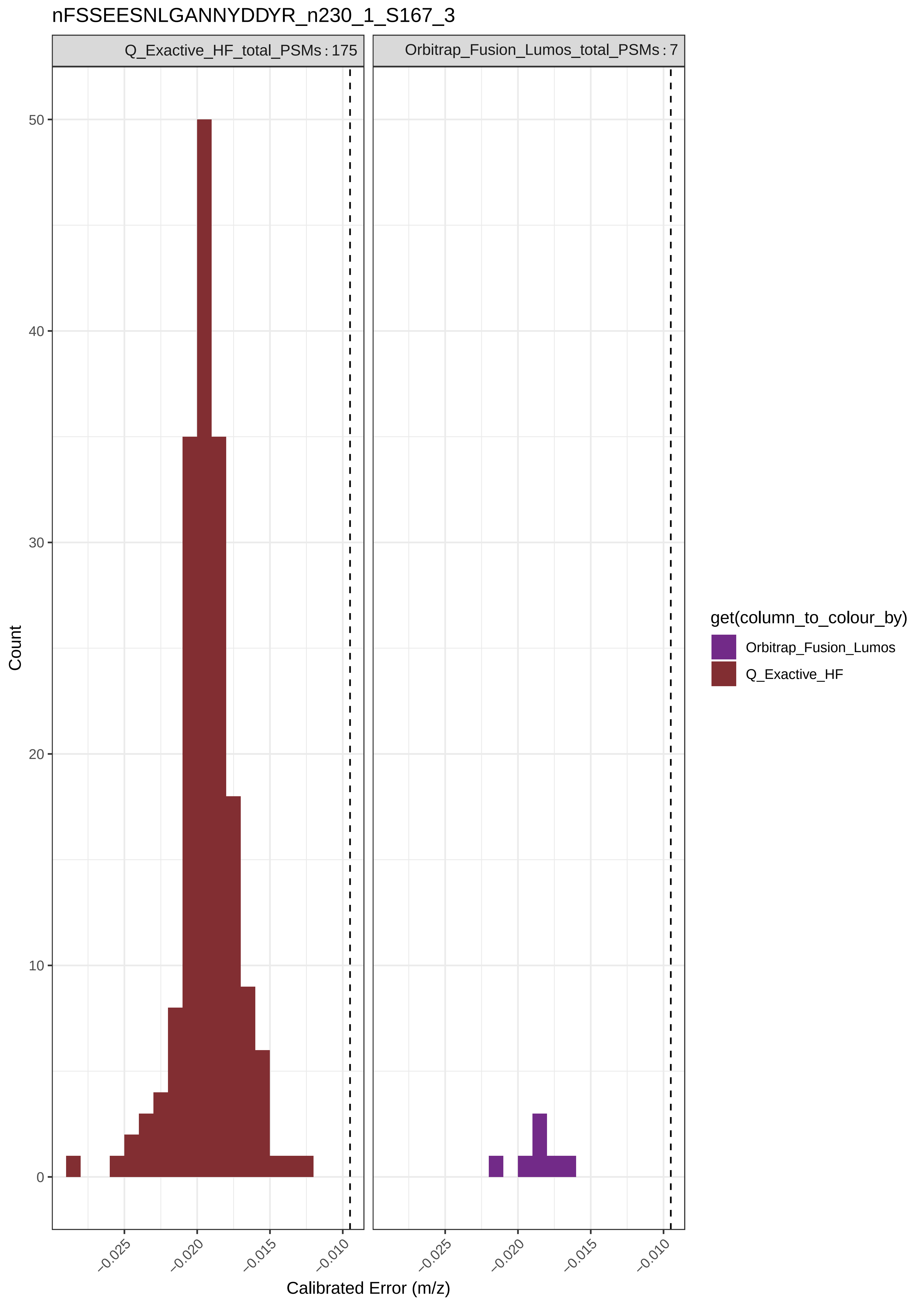
nFSSEESNLGANNYDDYR_n230_1_S167_3
Q_Exactive_HF_total_PSMs : 175
Orbitrap_Fusion_Lumos_total_PSMs : 7
Count
0
10
20
30
40
50
−0.025
−0.020
−0.015
−0.010
Calibrated Error (m/z)
get(column_to_colour_by)
Orbitrap_Fusion_Lumos
Q_Exactive_HF

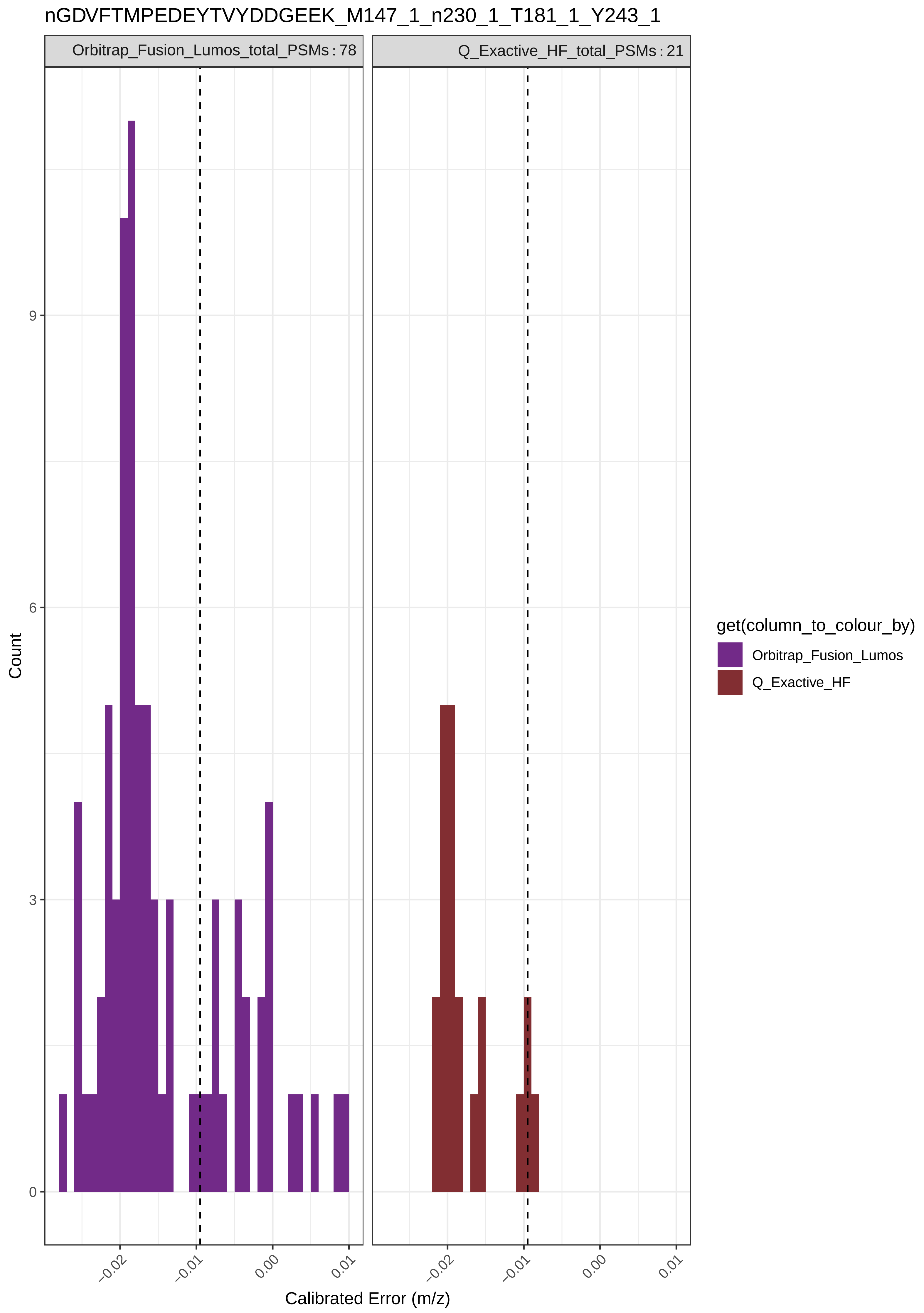

nGDVFTMPEDEYTVYDDGEEK_M147_1_n230_1_T181_1_Y243_1
Orbitrap_Fusion_Lumos_total_PSMs : 78
Q_Exactive_HF_total_PSMs : 21
Count
9
6
3
0
−0.02
−0.01
0.00
0.01
Calibrated Error (m/z)
get(column_to_colour_by)
Orbitrap_Fusion_Lumos
Q_Exactive_HF

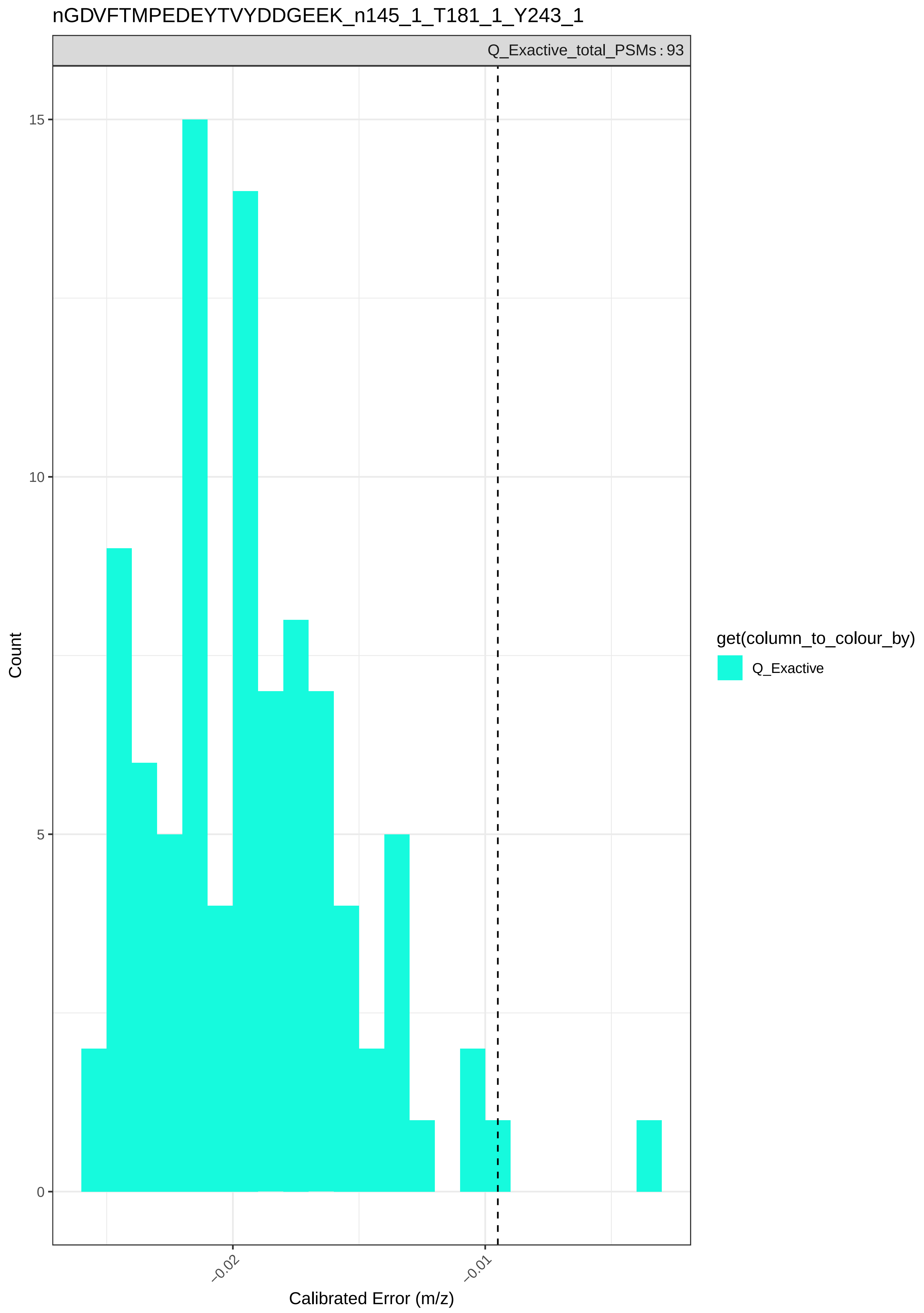

nGDVFTMPEDEYTVYDDGEEK_n145_1_T181_1_Y243_1
Q_Exactive_total_PSMs : 93
15
10
5
0
Count
−0.02
−0.01
Calibrated Error (m/z)
get(column_to_colour_by)
Q_Exactive

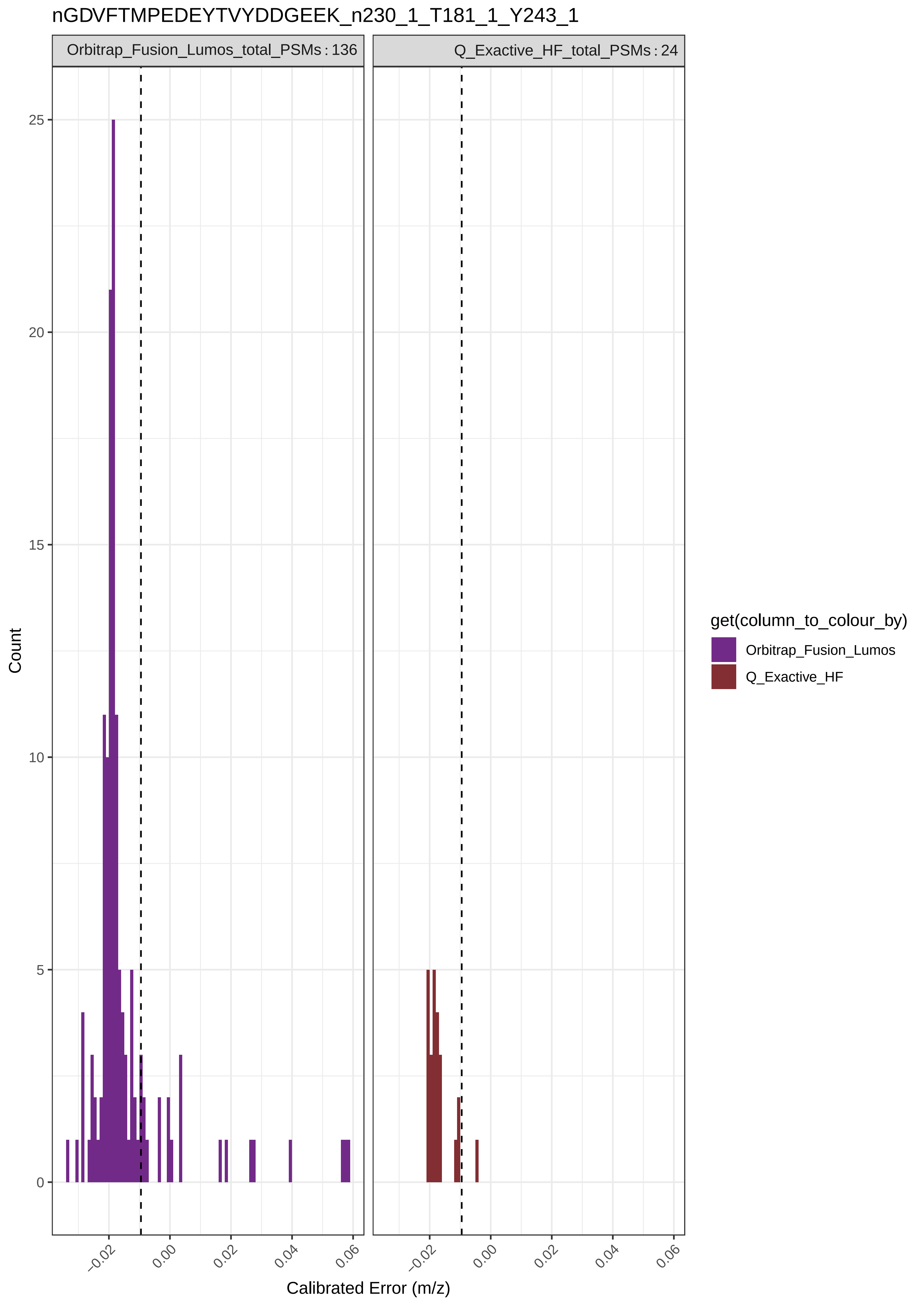

nGDVFTMPEDEYTVYDDGEEK_n230_1_T181_1_Y243_1
Orbitrap_Fusion_Lumos_total_PSMs : 136
Q_Exactive_HF_total_PSMs : 24
Count
25
20
15
10
5
0
−0.02
0.00
0.02
0.04
0.06
Calibrated Error (m/z)
get(column_to_colour_by)
Orbitrap_Fusion_Lumos
Q_Exactive_HF

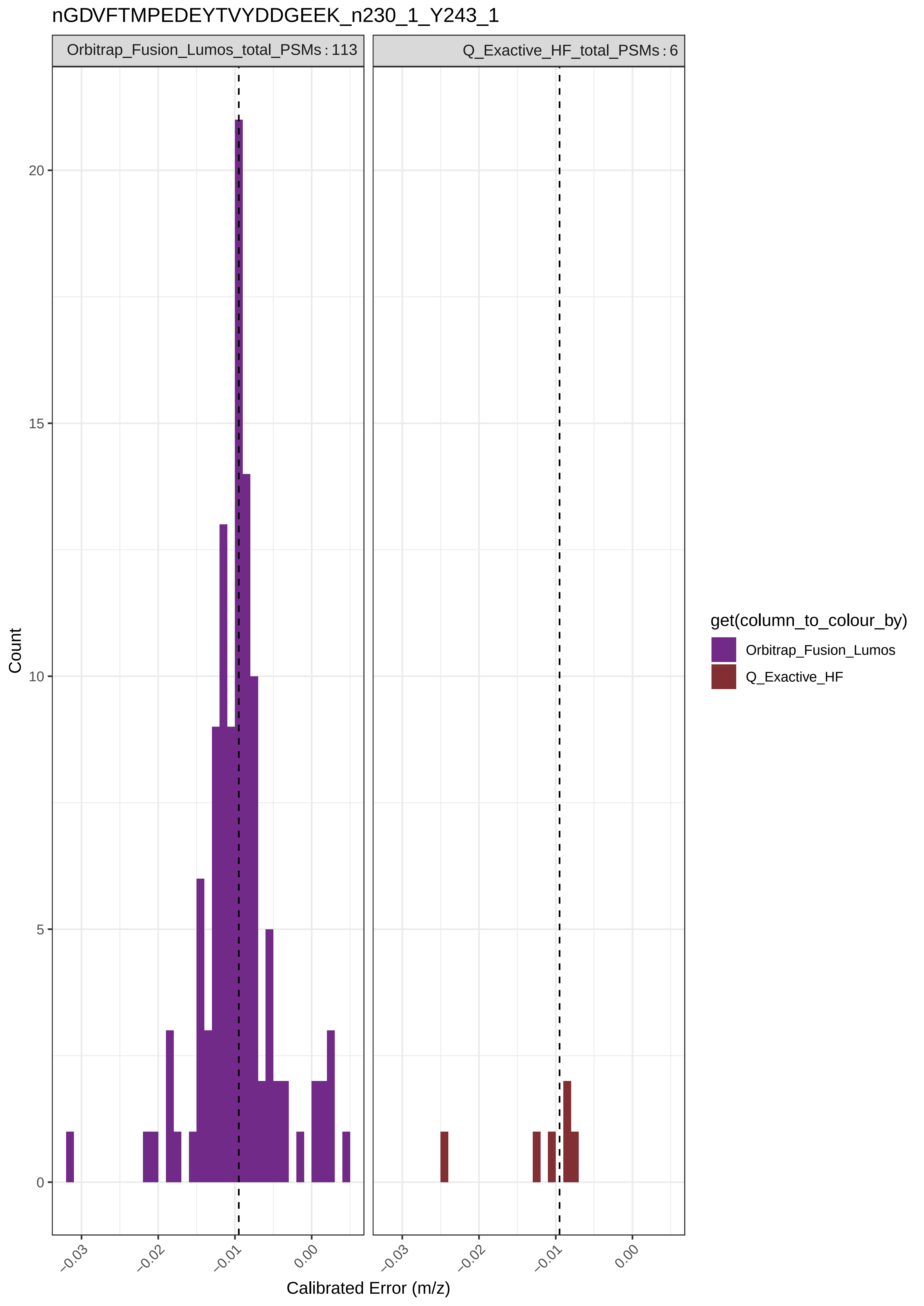
nGDVFTMPEDEYTVYDDGEEK_n230_1_Y243_1
Orbitrap_Fusion_Lumos_total_PSMs : 113
Q_Exactive_HF_total_PSMs : 6
Count
20
15
10
5
0
−0.03
−0.02
−0.01
0.00
Calibrated Error (m/z)
get(column_to_colour_by)
Orbitrap_Fusion_Lumos
Q_Exactive_HF

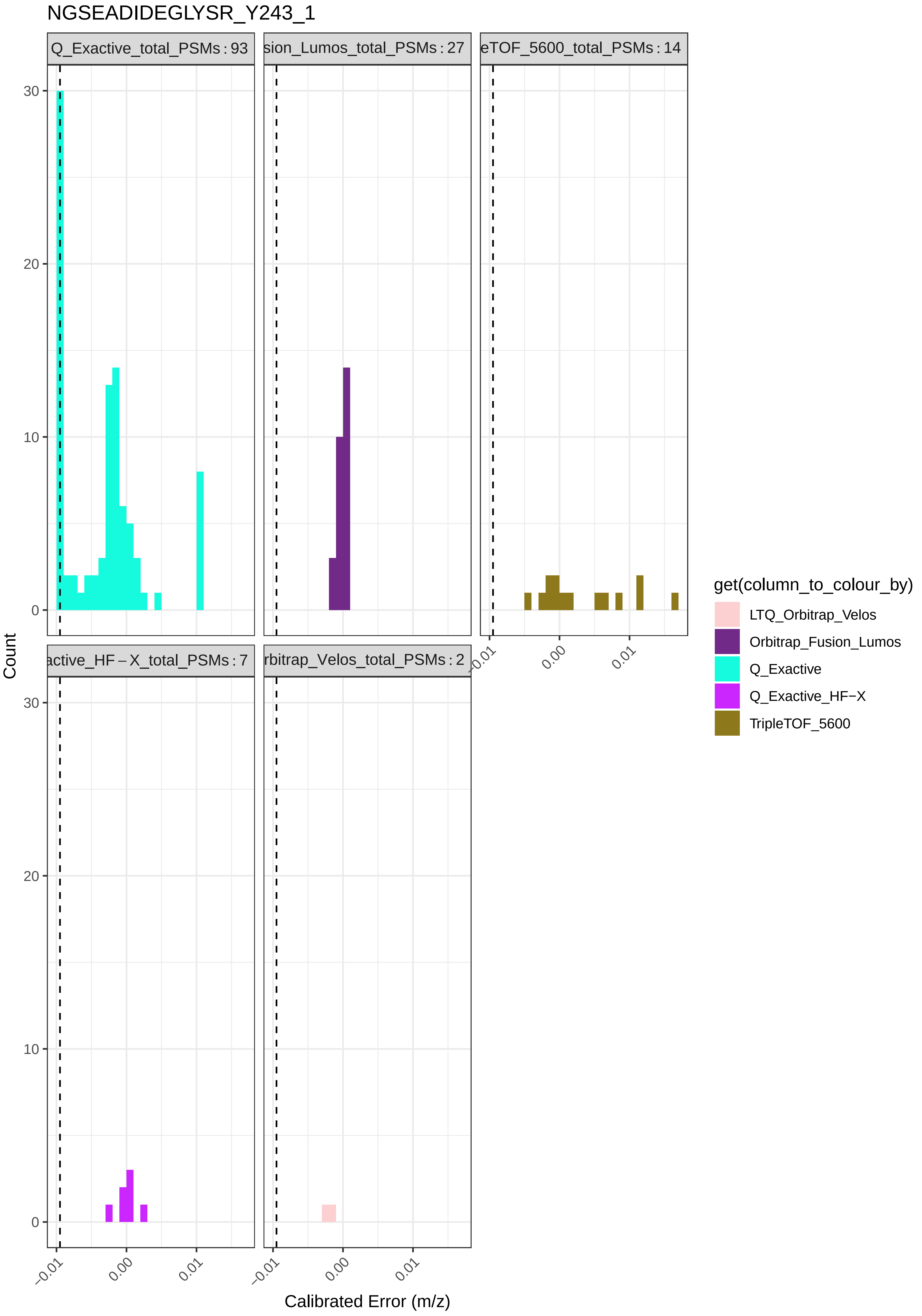
NGSEADIDEGLYSR_Y243_1
Q_Exactive_total_PSMs : 93
sion_Lumos_total_PSMs : 27
eTOF_5600_total_PSMs : 14
active_HF – X_total_PSMs : 7
rbitrap_Velos_total_PSMs : 2
Count
Calibrated Error (m/z)
get(column_to_colour_by)
LTQ_Orbitrap_Velos
Orbitrap_Fusion_Lumos
Q_Exactive
Q_Exactive_HF−X
TripleTOF_5600

nGTAGNALMDGASQLMGENRTMTIHNGMFFSTYDRDN_M147_1_n230_1_S167_1_T181_1

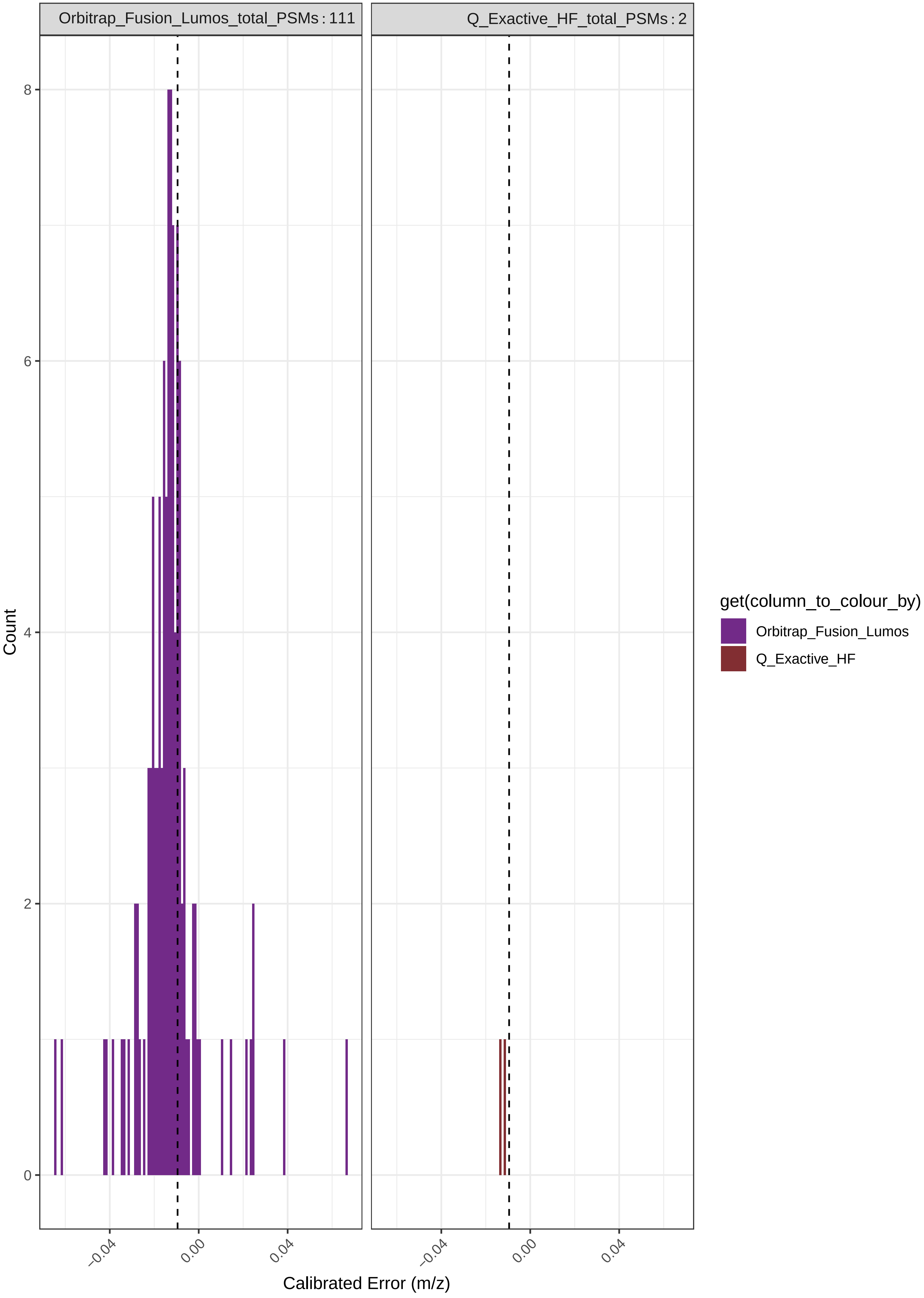

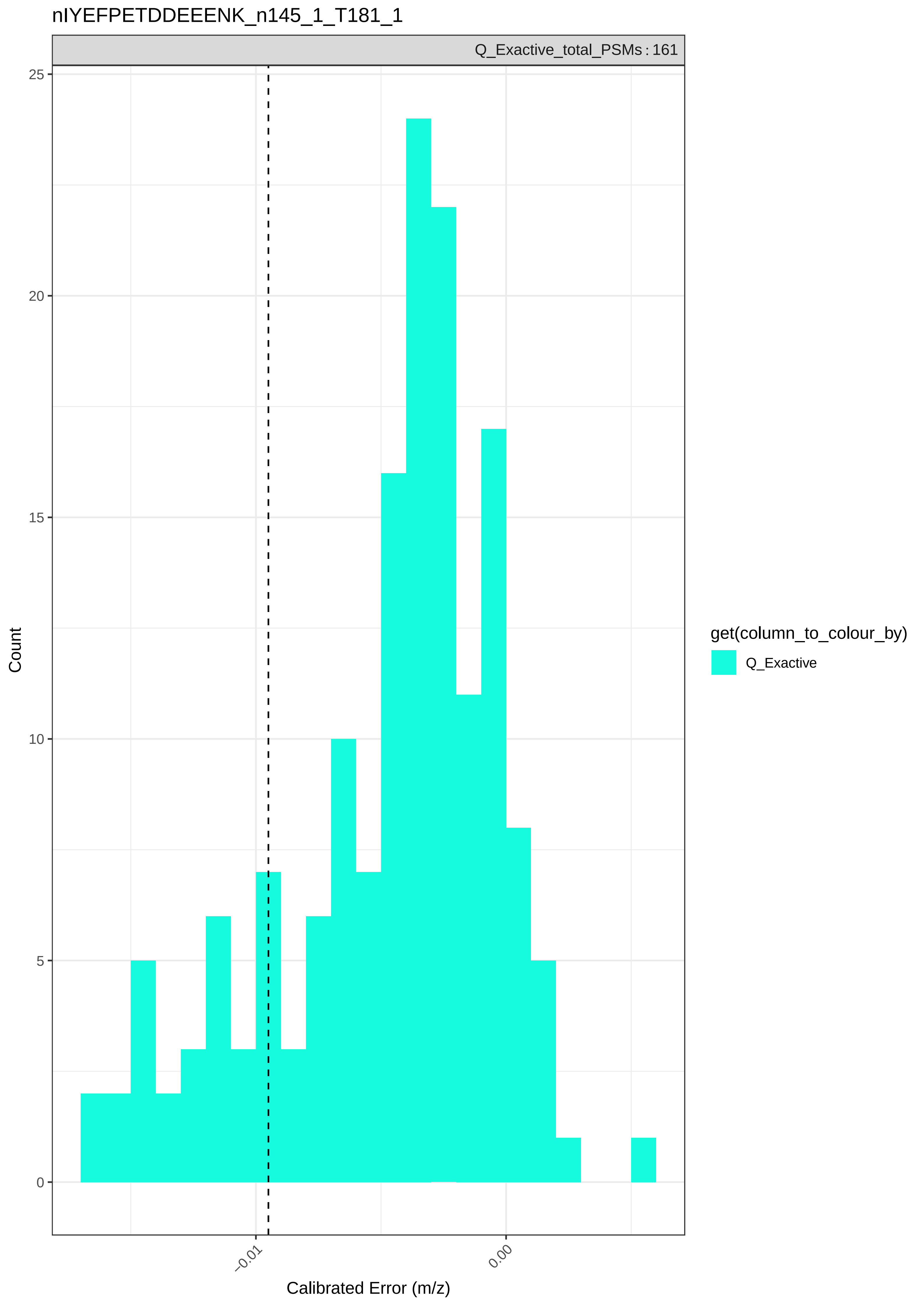
nIYEFPETDDEEENK_n145_1_T181_1
Q_Exactive_total_PSMs : 161
25
20
15
10
5
0
Count
−0.01
0.00
Calibrated Error (m/z)
get(column_to_colour_by)
Q_Exactive

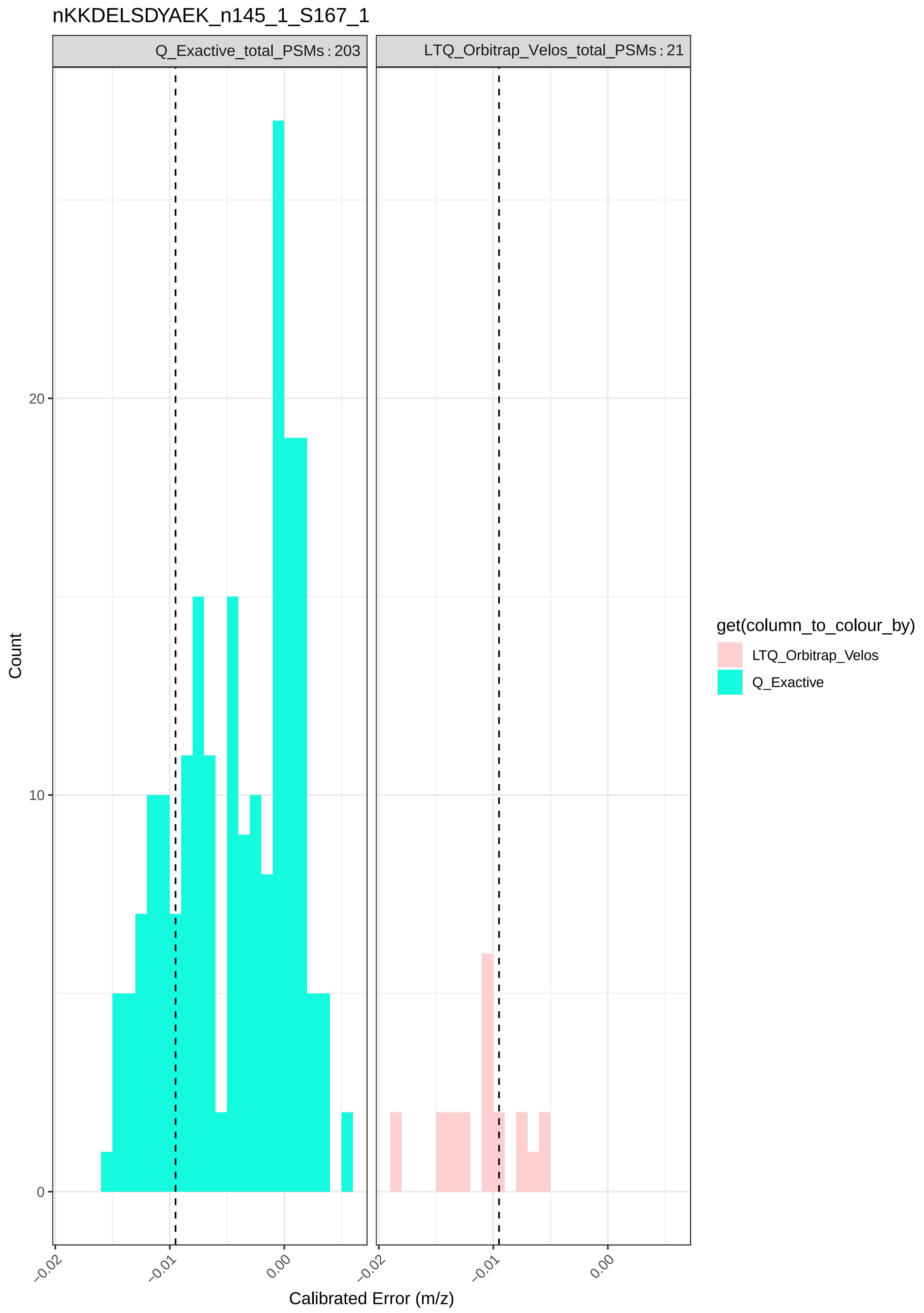

nKKDELSDYAEK_n145_1_S167_1
Q_Exactive_total_PSMs : 203
LTQ_Orbitrap_Velos_total_PSMs : 21
Count
20
10
0
−0.02
−0.01
0.00
Calibrated Error (m/z)
get(column_to_colour_by)
LTQ_Orbitrap_Velos
Q_Exactive

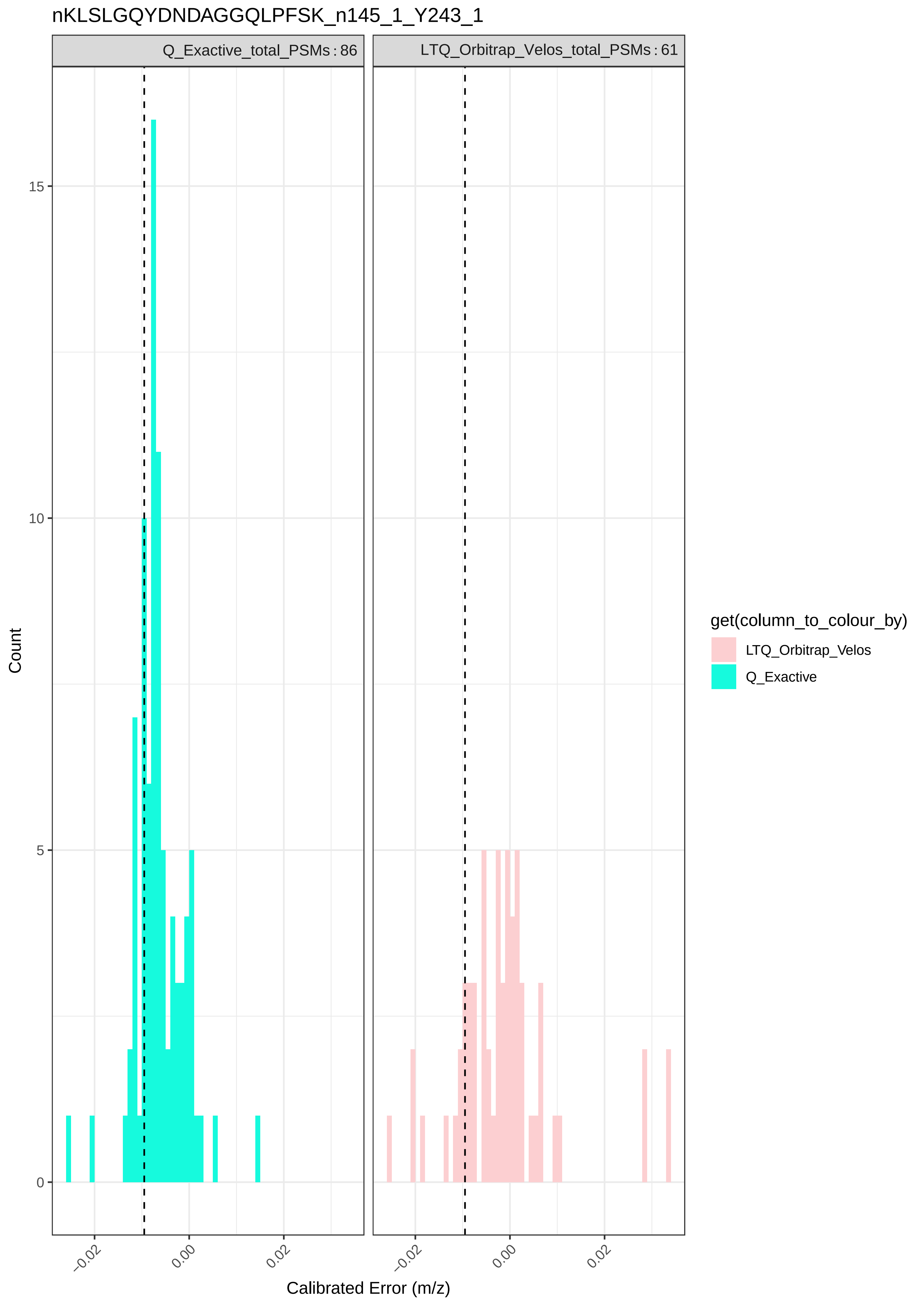
nKLSLGQYDNDAGGQLPFSK_n145_1_Y243_1
Q_Exactive_total_PSMs : 86
LTQ_Orbitrap_Velos_total_PSMs : 61
Count
15
10
5
0
−0.02
0.00
0.02
Calibrated Error (m/z)
get(column_to_colour_by)
LTQ_Orbitrap_Velos
Q_Exactive

nKSPVGKSPPSTGSTYGSSQK_n145_1_S167_2

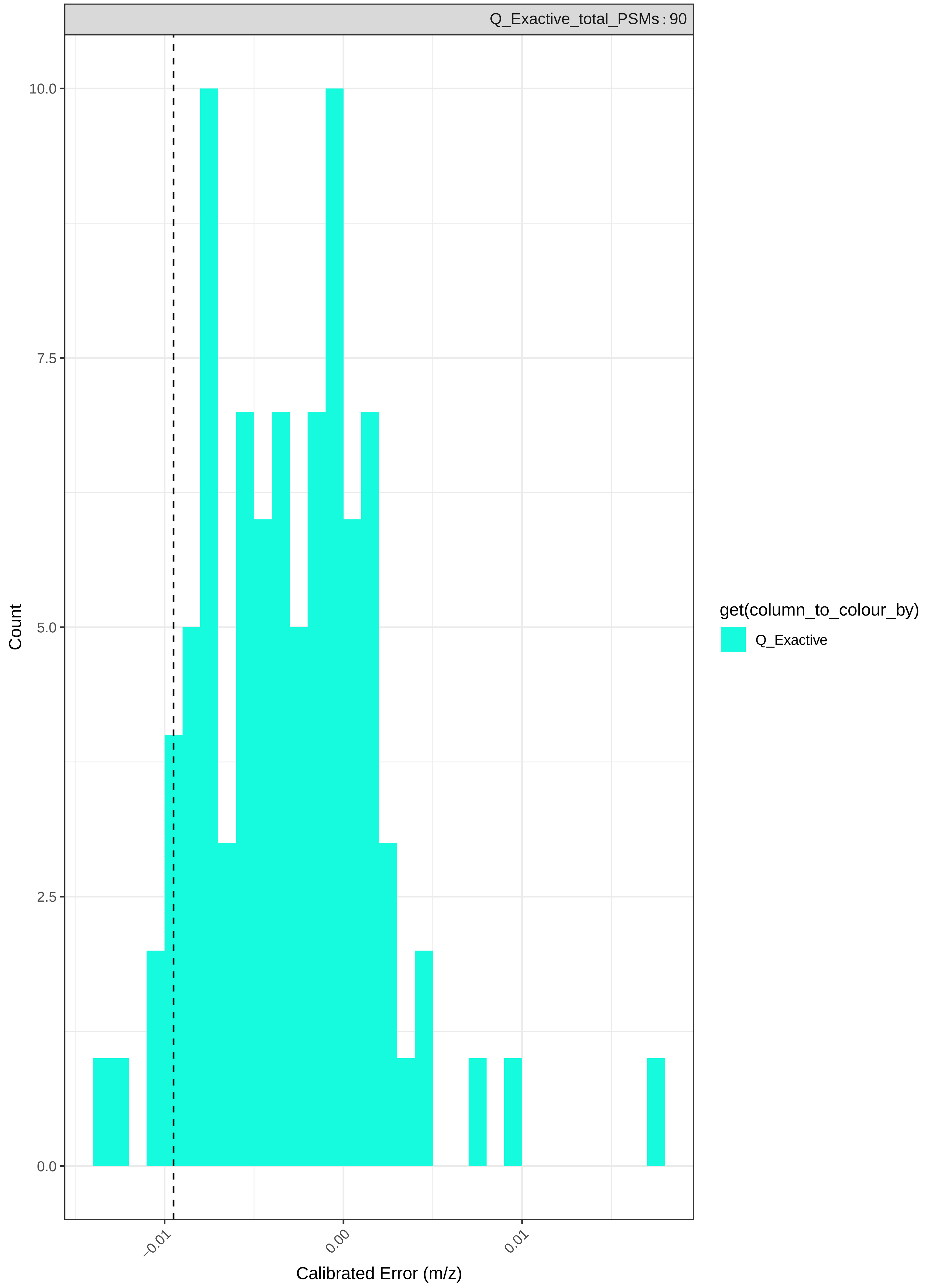

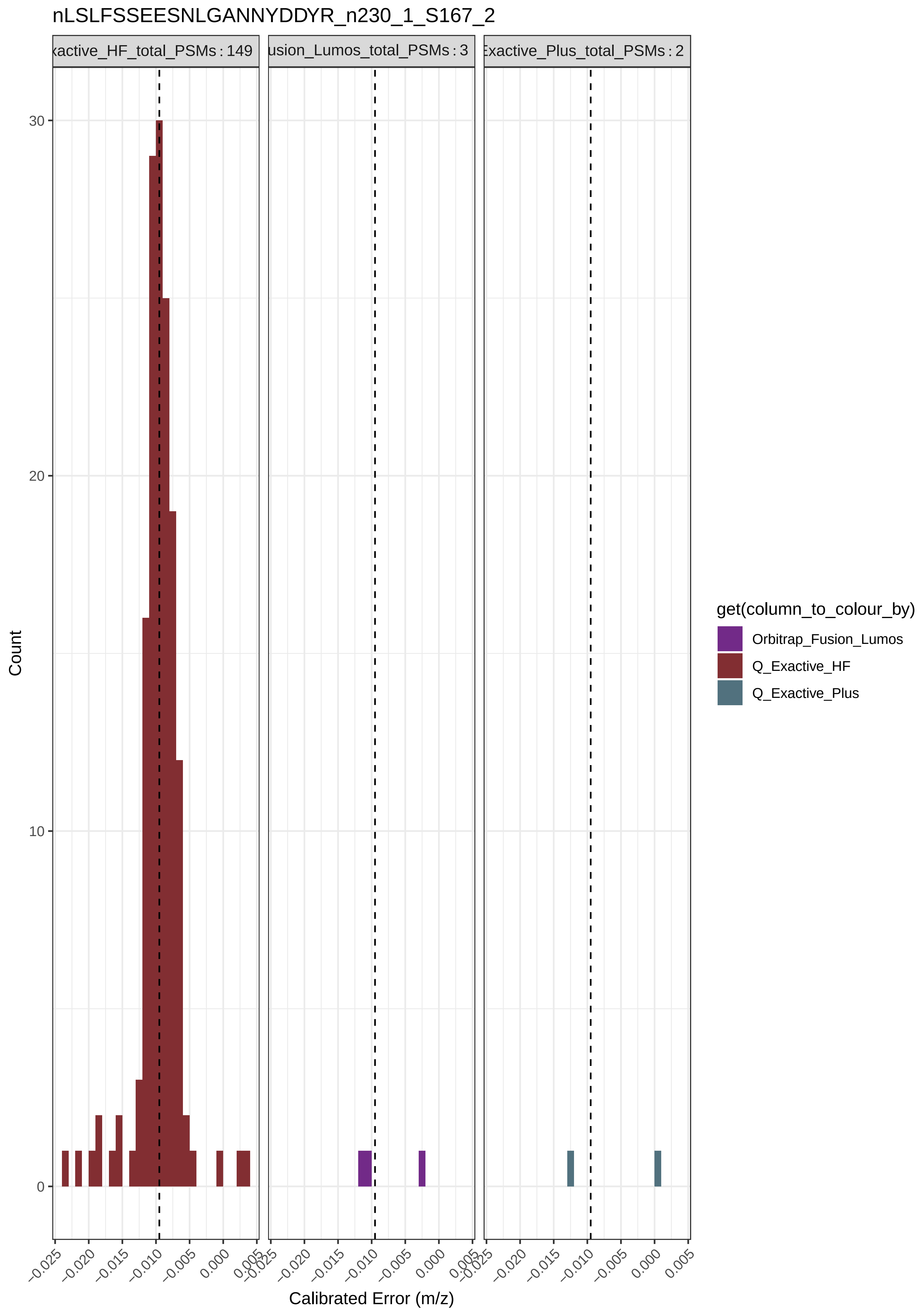
nLSLFSSEESNLGANNYDDYR_n230_1_S167_2
xactive_HF_total_PSMs : 149
usion_Lumos_total_PSMs : 3
Exactive_Plus_total_PSMs : 2
Count
0
10
20
30
−0.025
−0.020
−0.015
−0.010
−0.005
0.000
0.005
Calibrated Error (m/z)
get(column_to_colour_by)
Orbitrap_Fusion_Lumos
Q_Exactive_HF
Q_Exactive_Plus

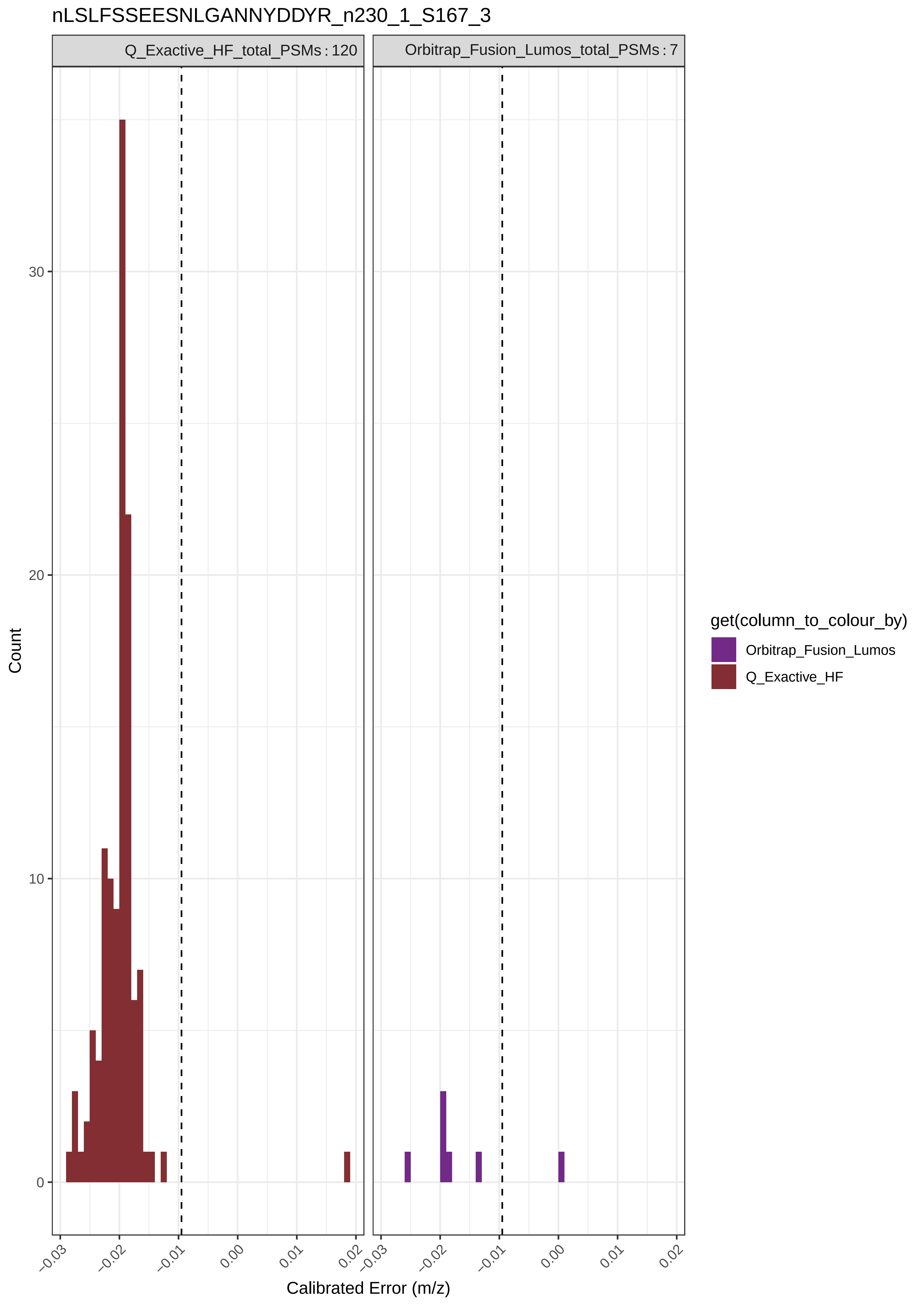
nLSLFSSEESNLGANNYDDYR_n230_1_S167_3
Q_Exactive_HF_total_PSMs : 120
Orbitrap_Fusion_Lumos_total_PSMs : 7
Count
30
20
10
0
−0.03
−0.02
−0.01
0.00
0.01
0.02
Calibrated Error (m/z)
get(column_to_colour_by)
Orbitrap_Fusion_Lumos
Q_Exactive_HF

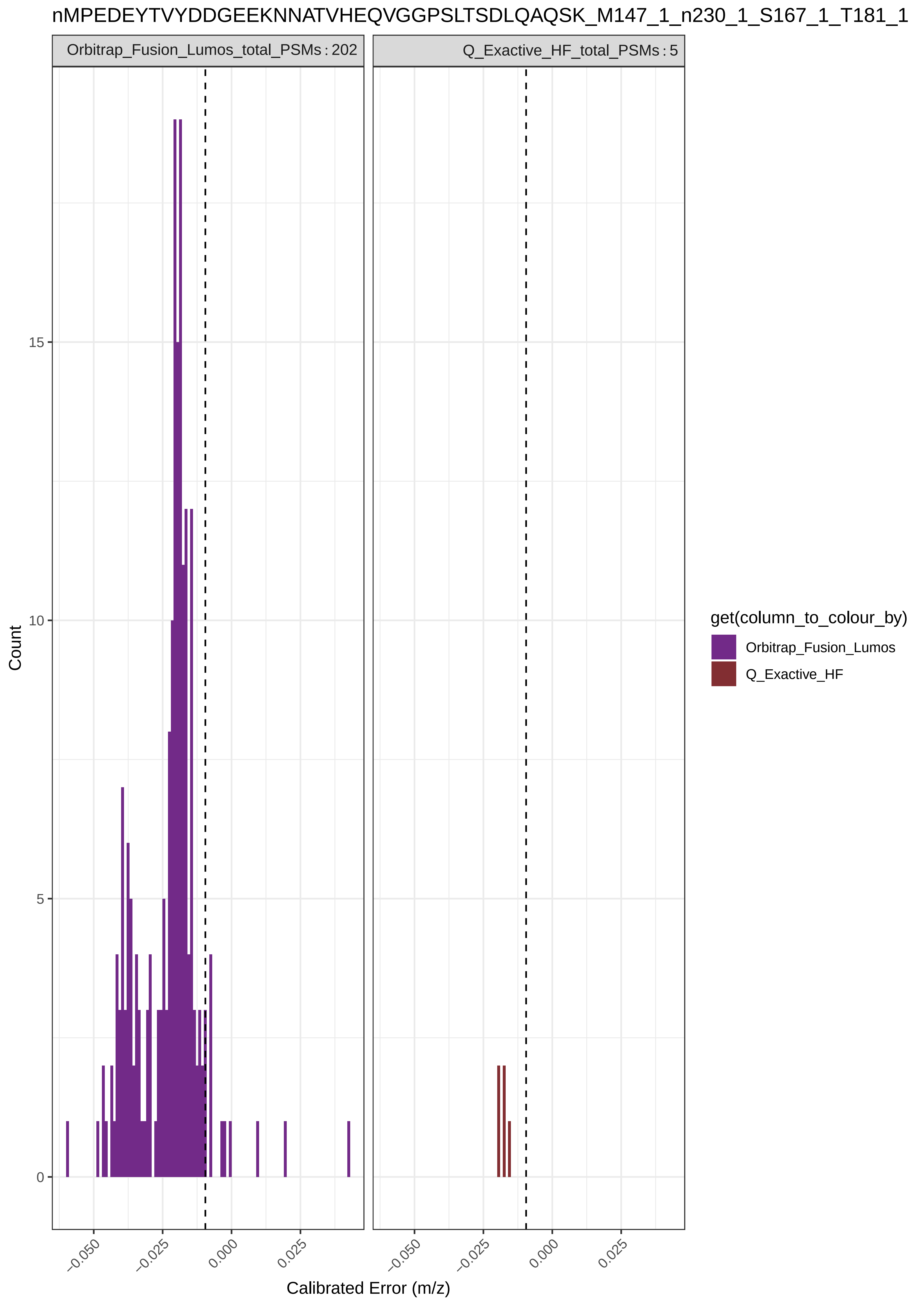
nMPEDEYTVYDDGEEKNNATVHEQVGGPSLTSDLQAQSK_M147_1_n230_1_S167_1_T181_1
Orbitrap_Fusion_Lumos_total_PSMs : 202
Q_Exactive_HF_total_PSMs : 5
Count
0
5
10
15
−0.050
−0.025
0.000
0.025
Calibrated Error (m/z)
get(column_to_colour_by)
Orbitrap_Fusion_Lumos
Q_Exactive_HF

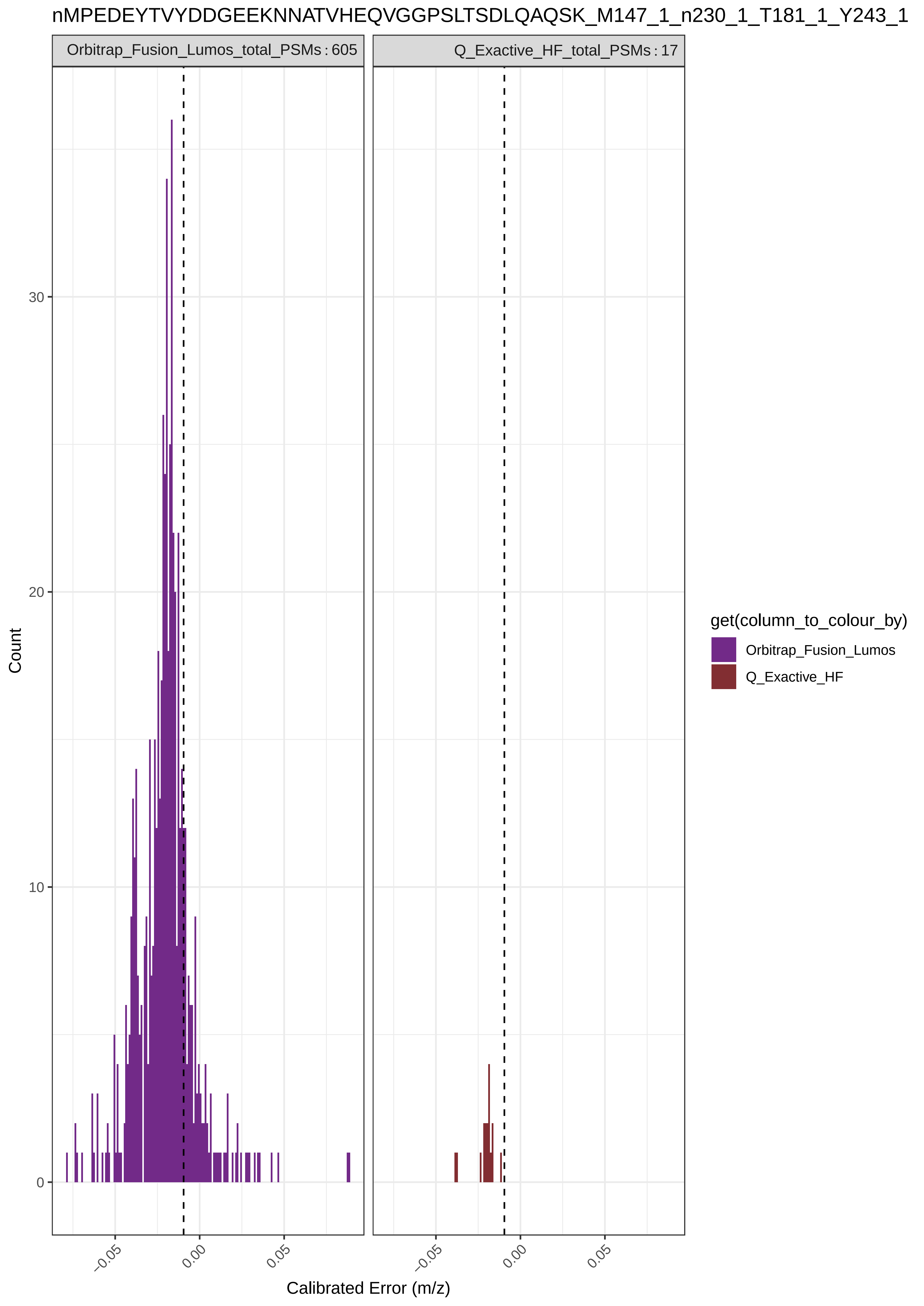

nMPEDEYTVYDDGEEKNNATVHEQVGGPSLTSDLQAQSK_M147_1_n230_1_T181_1_Y243_1
Orbitrap_Fusion_Lumos_total_PSMs : 605
Q_Exactive_HF_total_PSMs : 17
Count
0
10
20
30
−0.05
0.00
0.05
Calibrated Error (m/z)
get(column_to_colour_by)
Orbitrap_Fusion_Lumos
Q_Exactive_HF

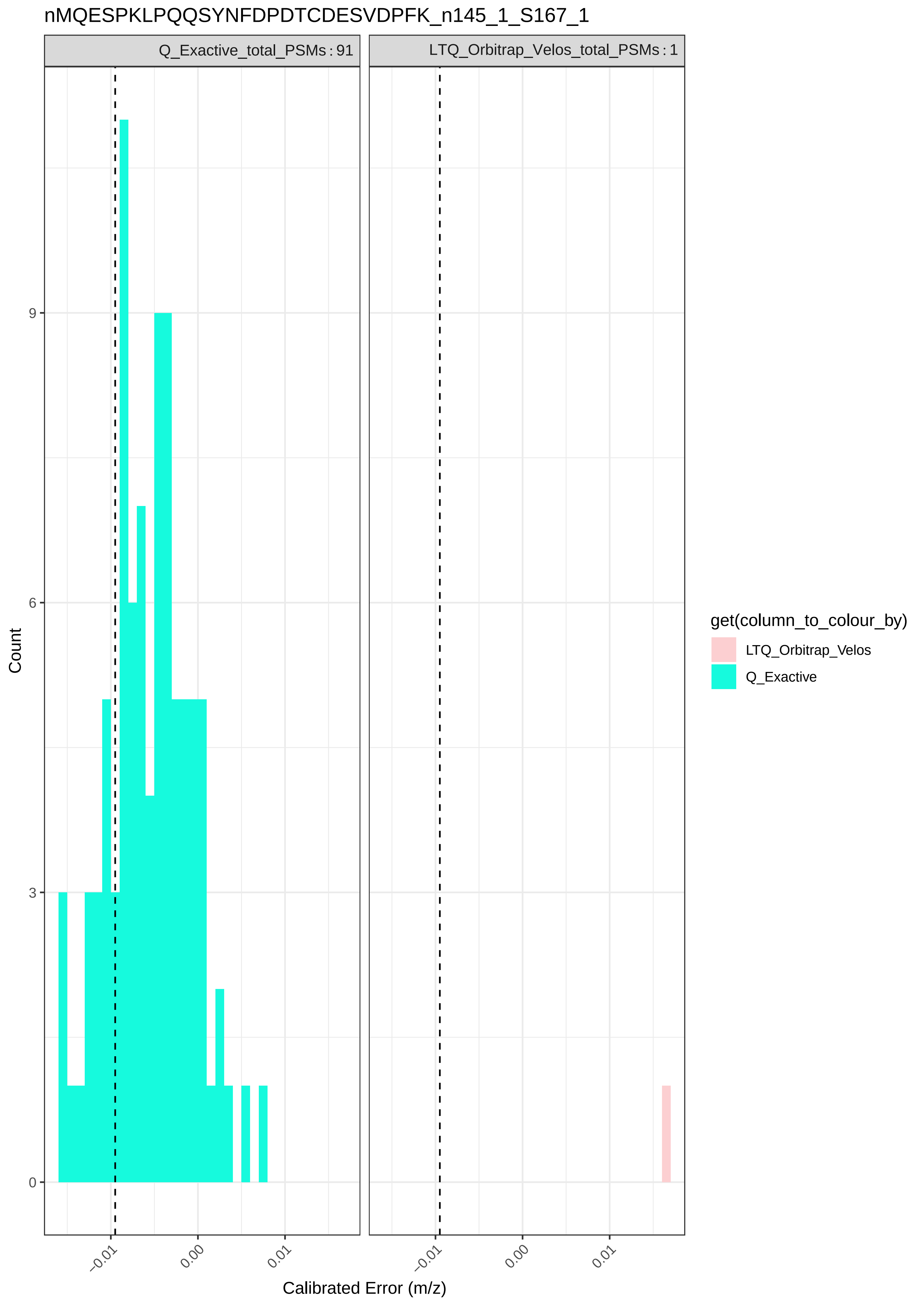

nMQESPKLPQQSYNFDPDTCDESVDPFK_n145_1_S167_1
Q_Exactive_total_PSMs : 91
LTQ_Orbitrap_Velos_total_PSMs : 1
Count
0
3
6
9
−0.01
0.00
0.01
Calibrated Error (m/z)
get(column_to_colour_by)
LTQ_Orbitrap_Velos
Q_Exactive

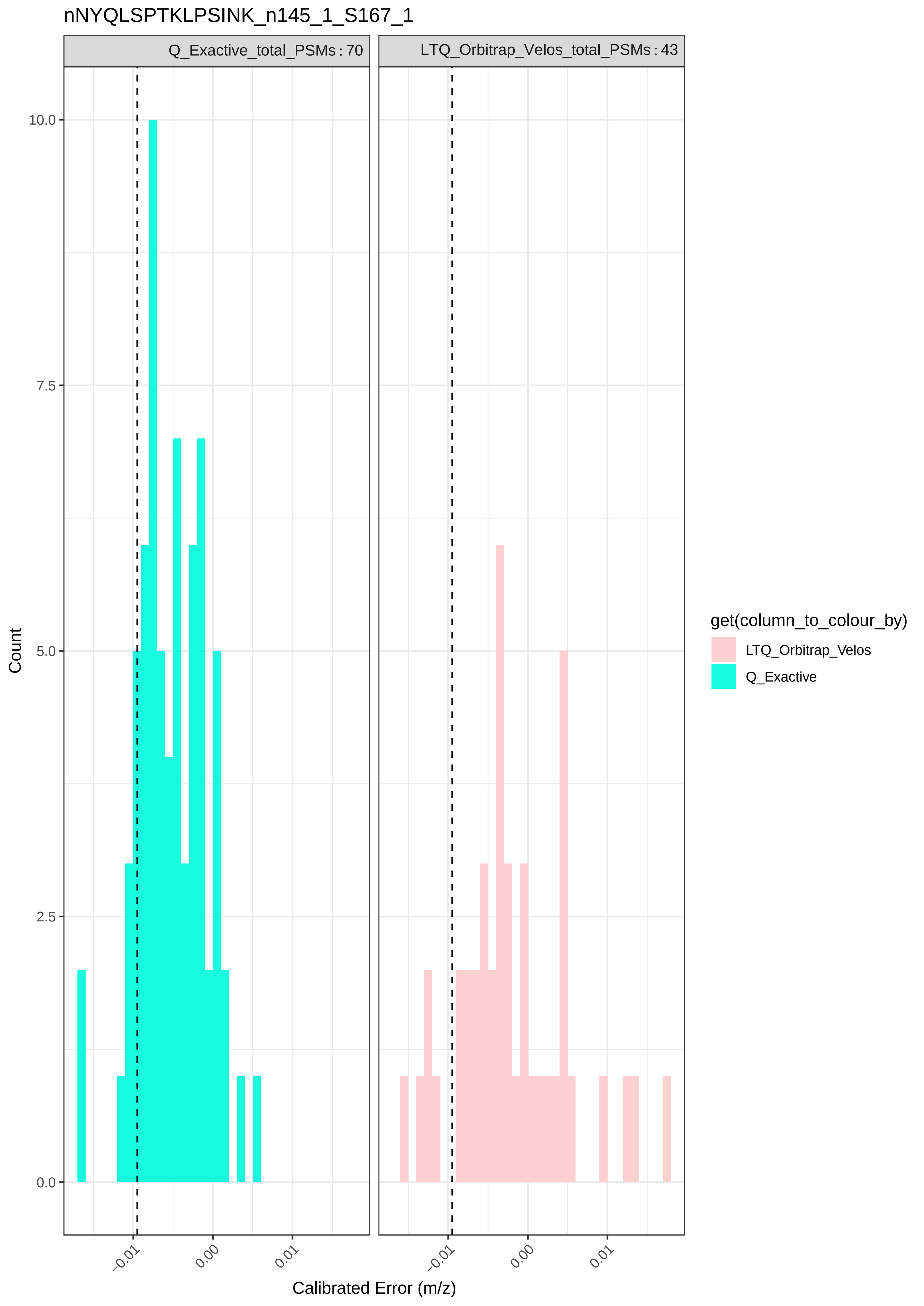

nNYQLSPTKLPSINK_n145_1_S167_1
Q_Exactive_total_PSMs : 70
LTQ_Orbitrap_Velos_total_PSMs : 43
Count
10.0
7.5
5.0
2.5
0.0
−0.01
0.00
0.01
Calibrated Error (m/z)
get(column_to_colour_by)
LTQ_Orbitrap_Velos
Q_Exactive

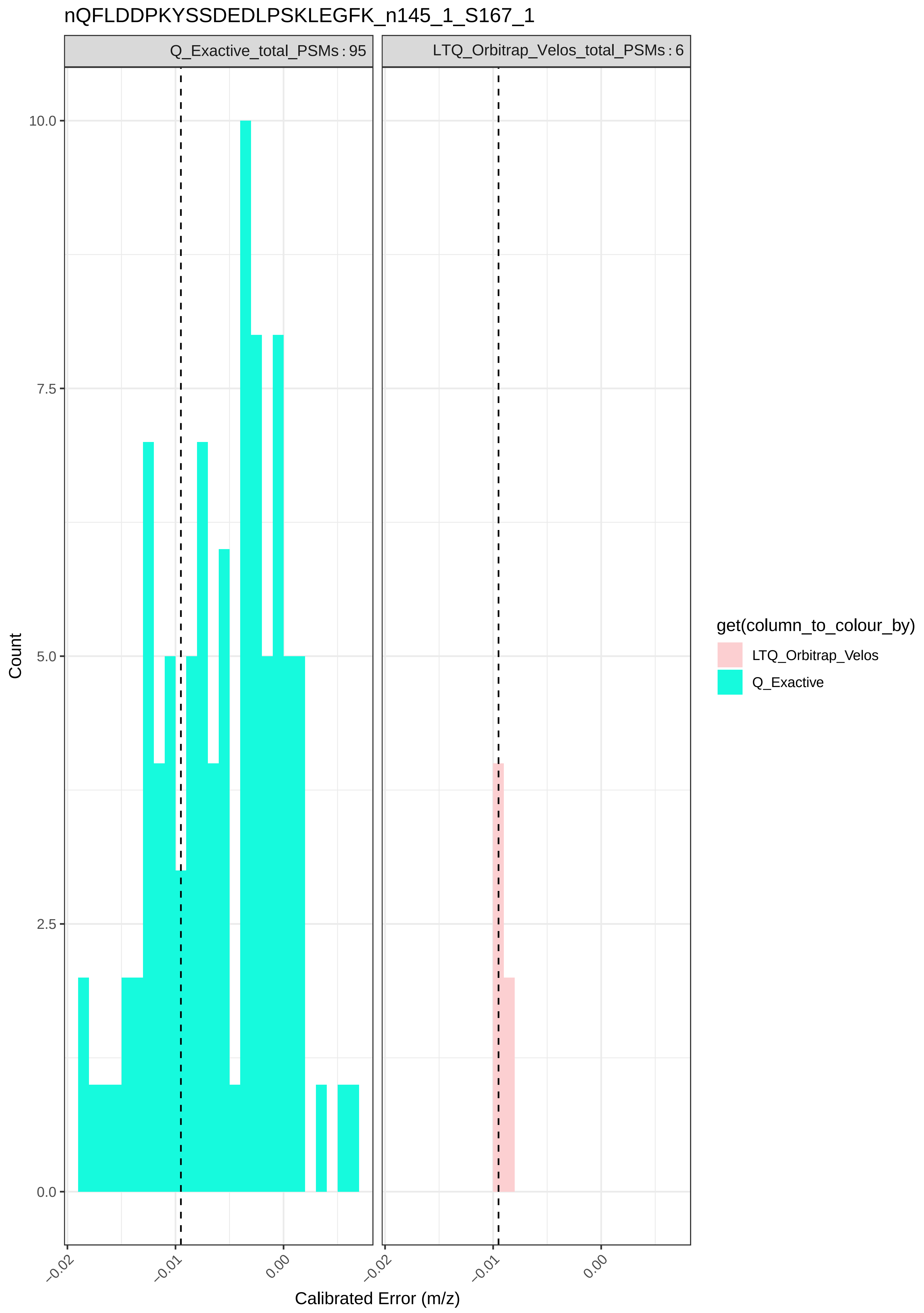
nQFLDDPKYSSDEDLPSKLEGFK_n145_1_S167_1
Q_Exactive_total_PSMs : 95
LTQ_Orbitrap_Velos_total_PSMs : 6
10.0
7.5
5.0
2.5
0.0
Count
−0.02
−0.01
0.00
Calibrated Error (m/z)
get(column_to_colour_by)
LTQ_Orbitrap_Velos
Q_Exactive

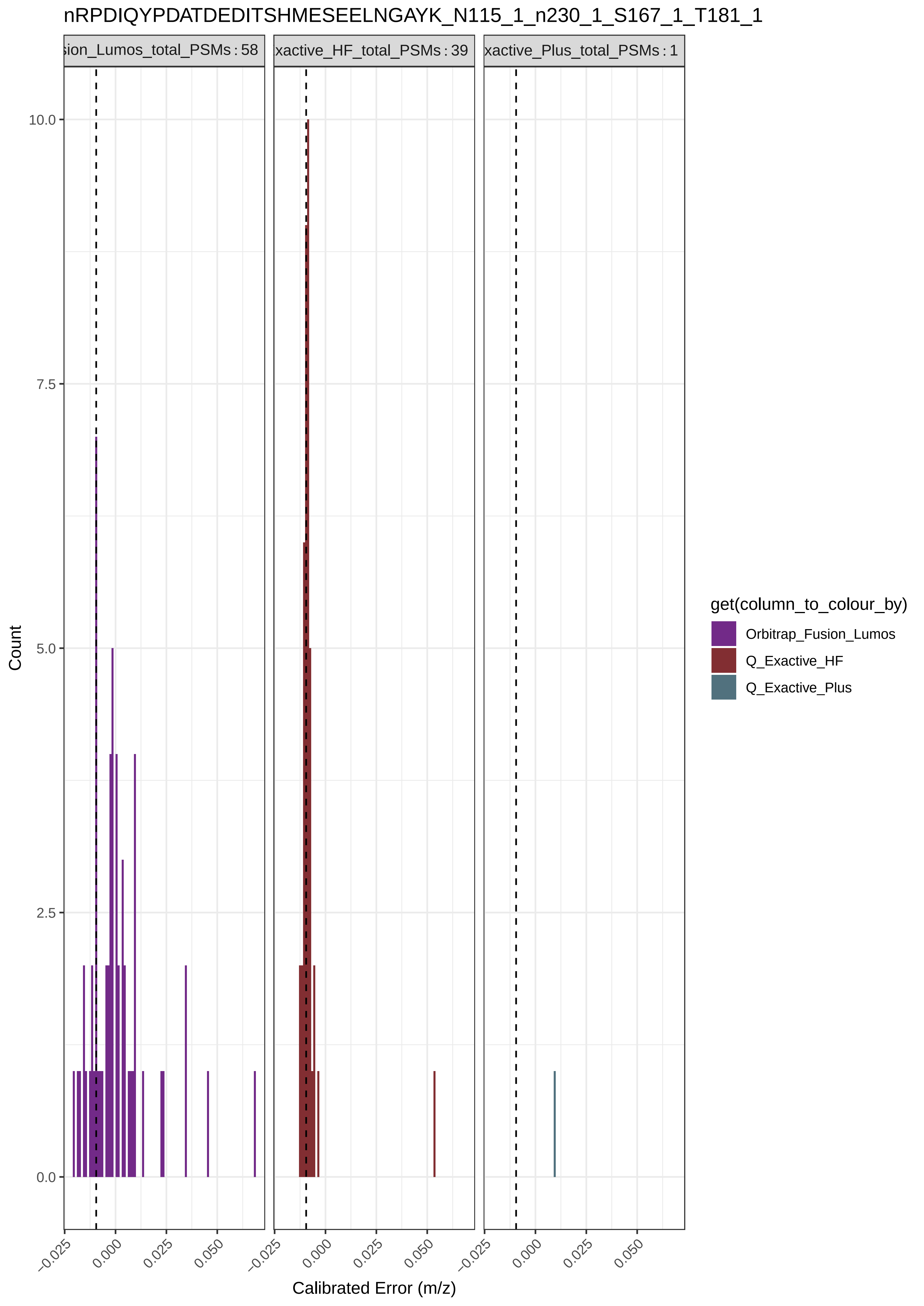
nRPDIQYPDATDEDITSHMESEELNGAYK_N115_1_n230_1_S167_1_T181_1
sion_Lumos_total_PSMs : 58
xactive_HF_total_PSMs : 39
xactive_Plus_total_PSMs : 1
Count
10.0
7.5
5.0
2.5
0.0
−0.025
0.000
0.025
0.050
Calibrated Error (m/z)
get(column_to_colour_by)
Orbitrap_Fusion_Lumos
Q_Exactive_HF
Q_Exactive_Plus

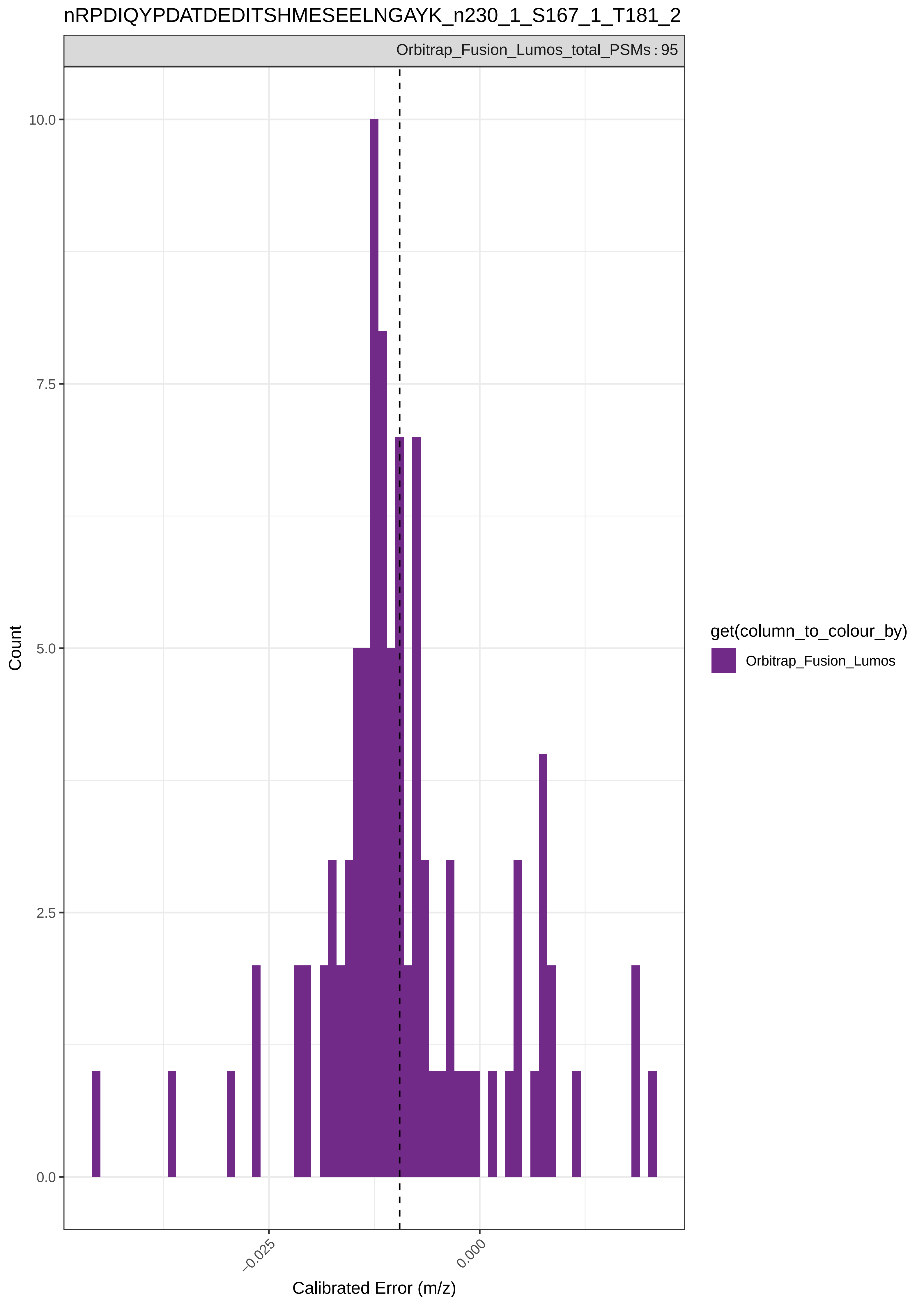
nRPDIQYPDATDEDITSHMESEELNGAYK_n230_1_S167_1_T181_2
Orbitrap_Fusion_Lumos_total_PSMs : 95
10.0
7.5
5.0
2.5
0.0
Count
−0.025
0.000
Calibrated Error (m/z)
get(column_to_colour_by)
Orbitrap_Fusion_Lumos

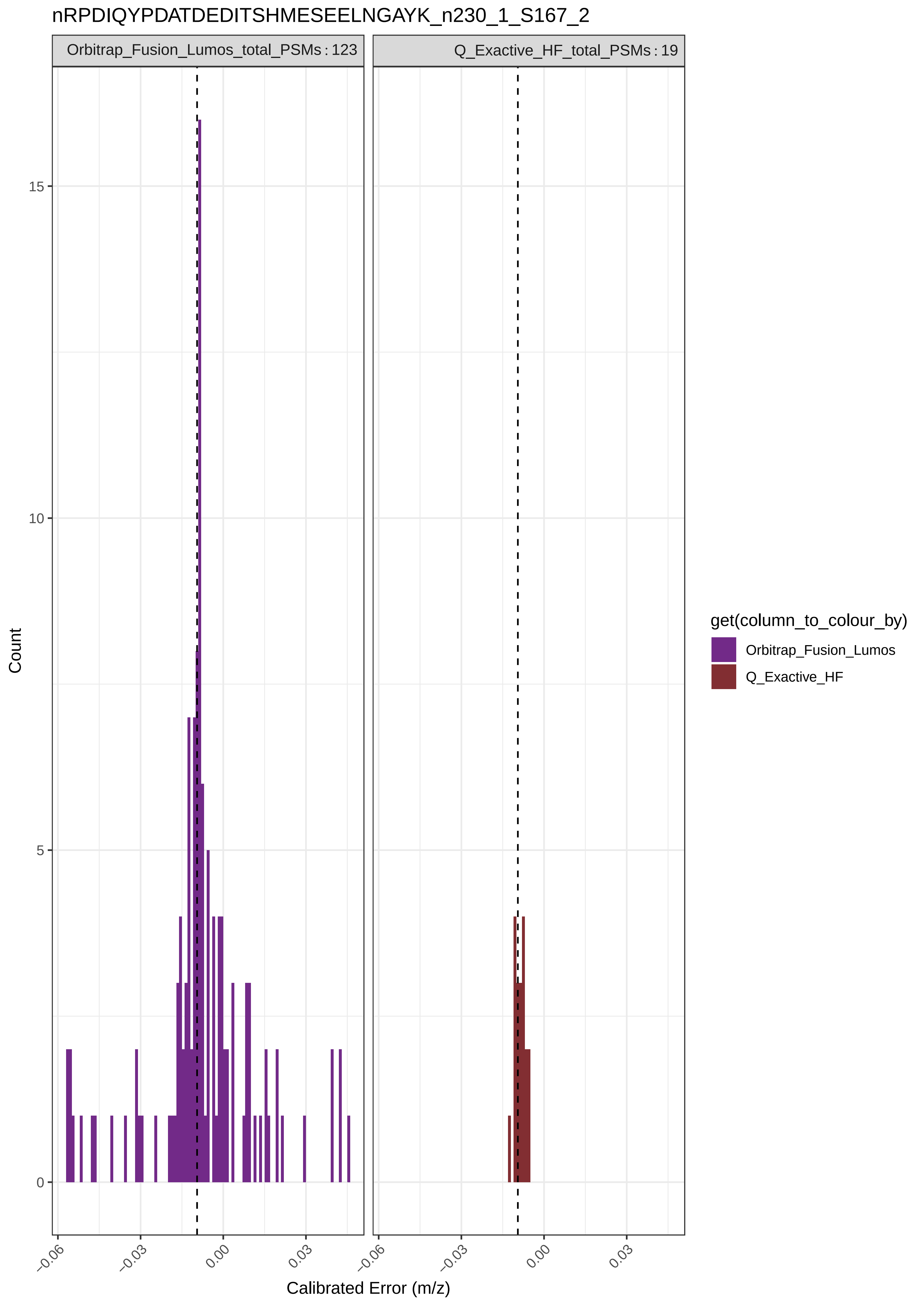
nRPDIQYPDATDEDITSHMESEELNGAYK_n230_1_S167_2
Orbitrap_Fusion_Lumos_total_PSMs : 123
Q_Exactive_HF_total_PSMs : 19
Count
15
10
5
0
−0.06
−0.03
0.00
0.03
Calibrated Error (m/z)
get(column_to_colour_by)
Orbitrap_Fusion_Lumos
Q_Exactive_HF

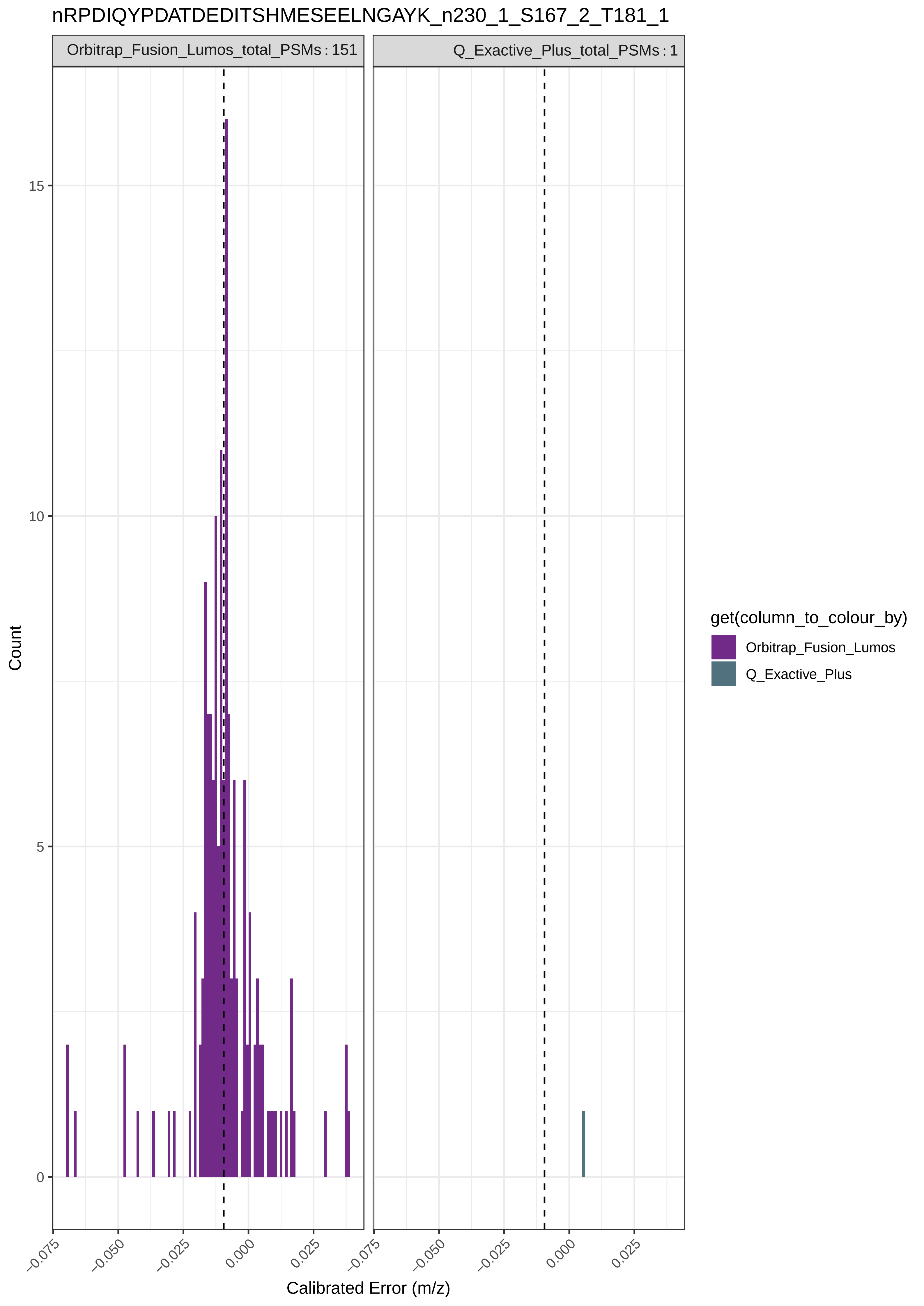
nRPDIQYPDATDEDITSHMESEELNGAYK_n230_1_S167_2_T181_1
Orbitrap_Fusion_Lumos_total_PSMs : 151
Q_Exactive_Plus_total_PSMs : 1
Count
15
10
5
0
−0.075
−0.050
−0.025
0.000
0.025
Calibrated Error (m/z)
get(column_to_colour_by)
Orbitrap_Fusion_Lumos
Q_Exactive_Plus

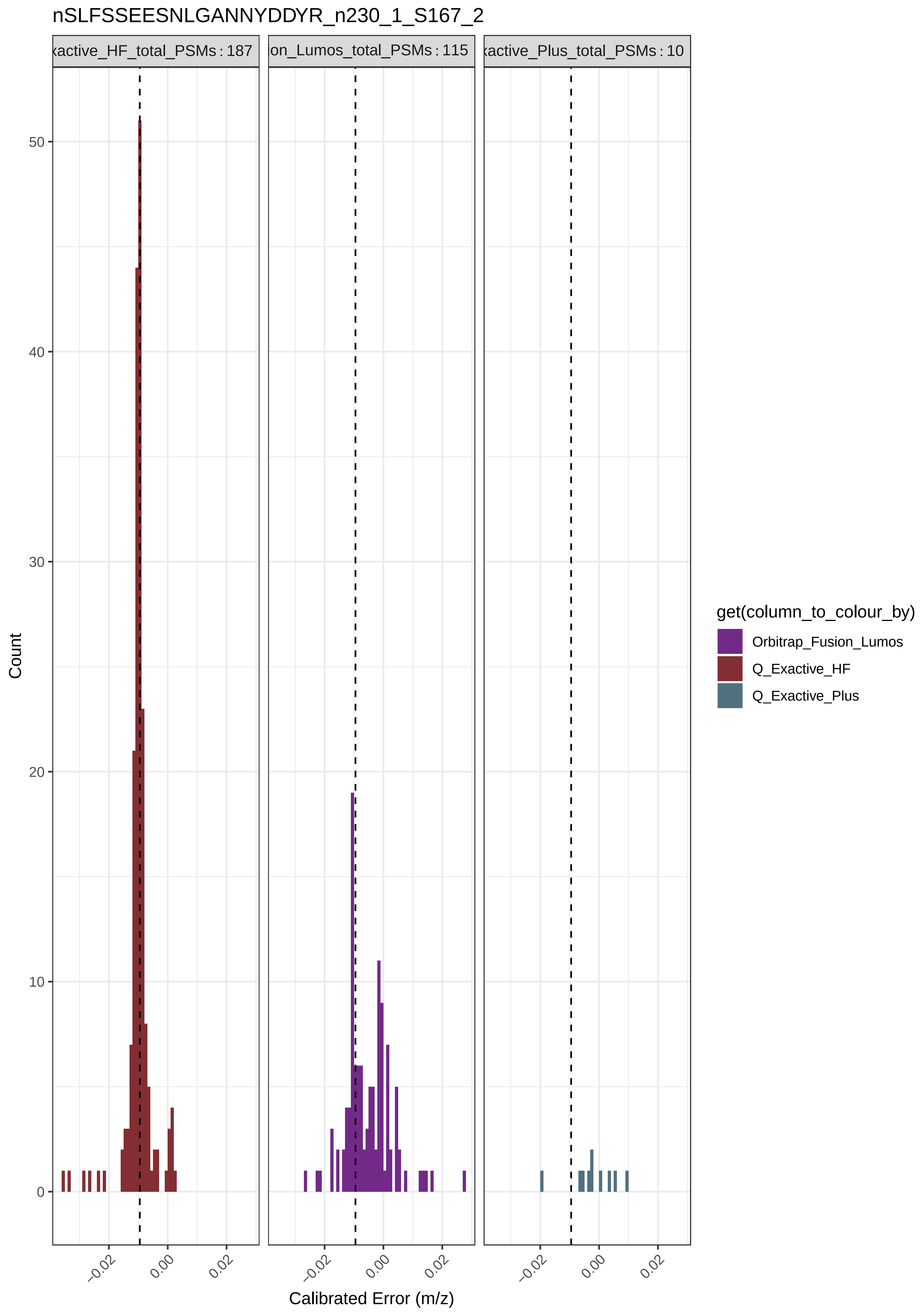
nSLFSSEESNLGANNYDDYR_n230_1_S167_2
xactive_HF_total_PSMs : 187
ion_Lumos_total_PSMs : 115
xactive_Plus_total_PSMs : 10
Count
0
10
20
30
40
50
−0.02
0.00
0.02
Calibrated Error (m/z)
get(column_to_colour_by)
Orbitrap_Fusion_Lumos
Q_Exactive_HF
Q_Exactive_Plus

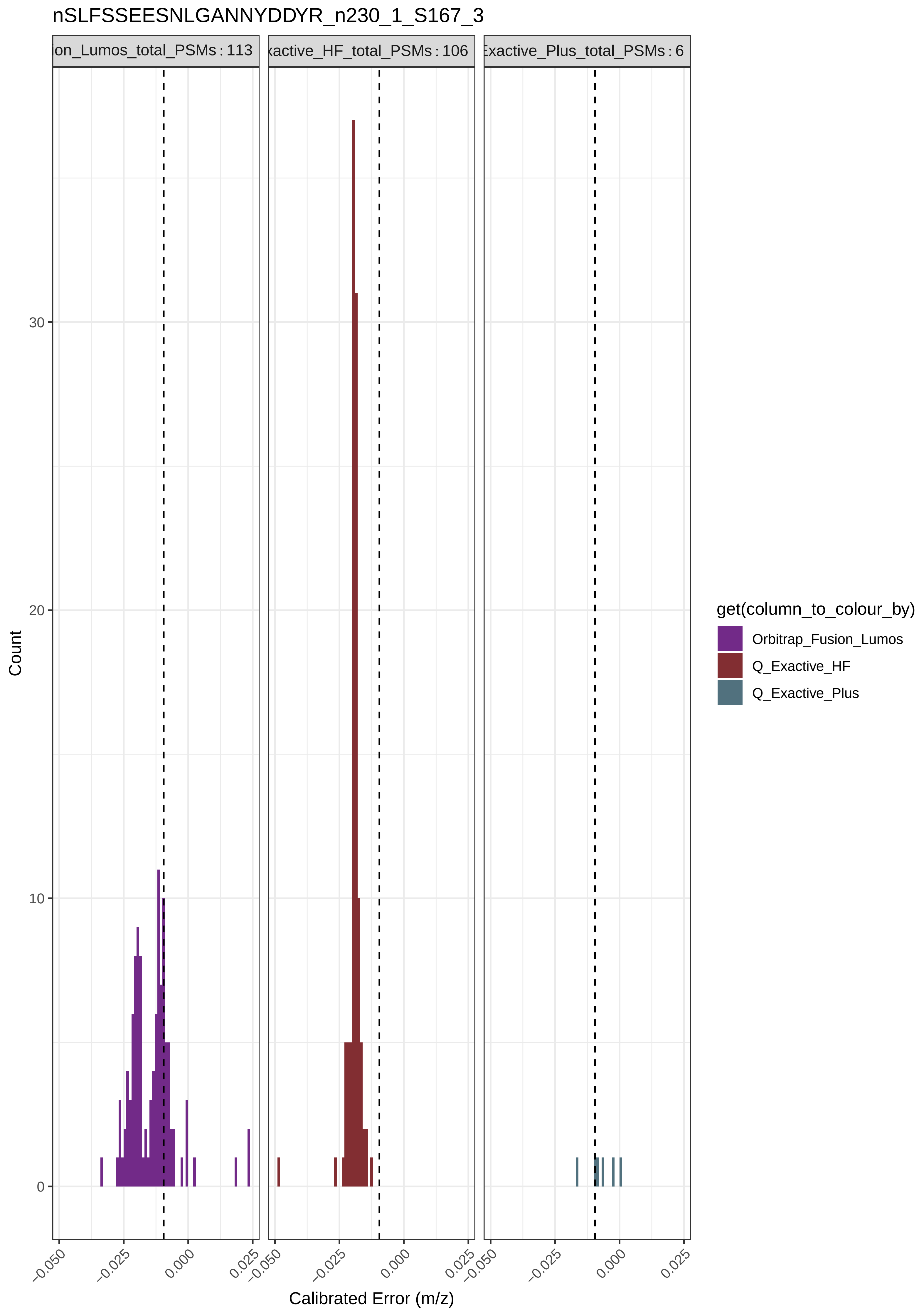

nSLFSSEESNLGANNYDDYR_n230_1_S167_3
ion_Lumos_total_PSMs : 113
xactive_HF_total_PSMs : 106
Exactive_Plus_total_PSMs : 6
Count
30
20
10
0
−0.050
−0.025
0.000
0.025
Calibrated Error (m/z)
get(column_to_colour_by)
Orbitrap_Fusion_Lumos
Q_Exactive_HF
Q_Exactive_Plus

nSLYNLGGSR_n230_1_S167_1

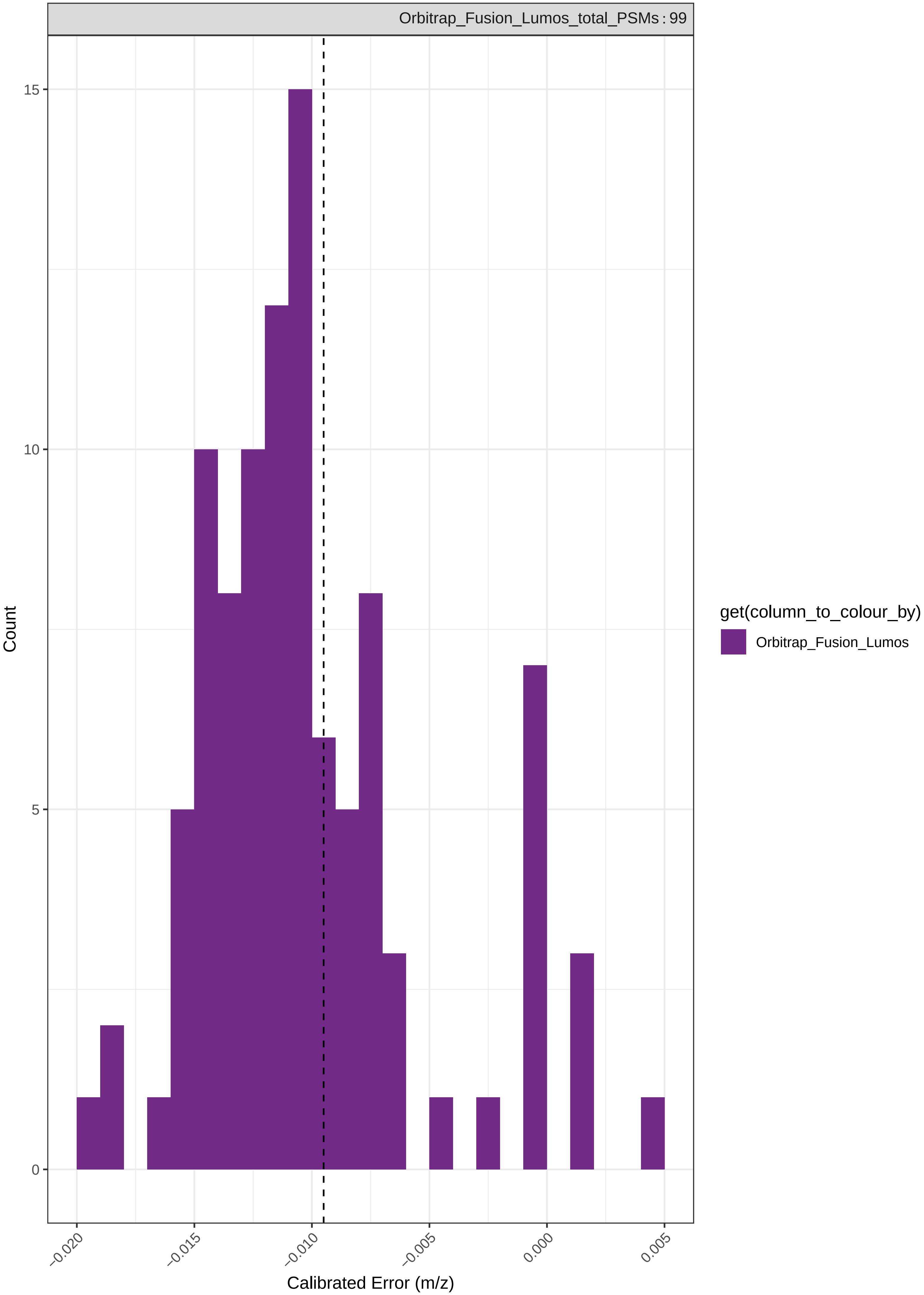

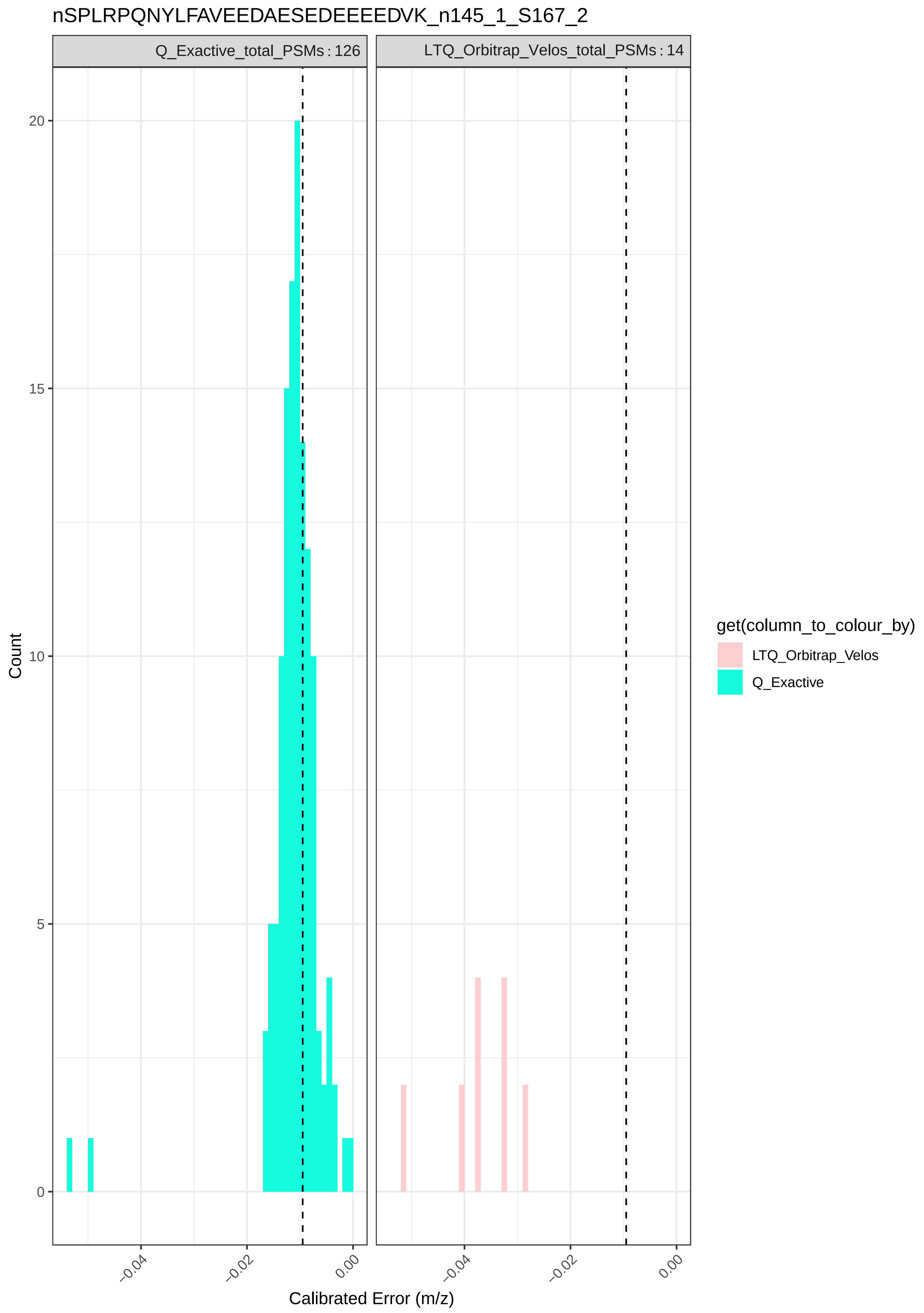

nSPLRPQNYLFAVEEDAESEDEEEEDVK_n145_1_S167_2
Q_Exactive_total_PSMs : 126
LTQ_Orbitrap_Velos_total_PSMs : 14
Count
20
15
10
5
0
−0.04
−0.02
0.00
−0.04
−0.02
0.00
Calibrated Error (m/z)
get(column_to_colour_by)
LTQ_Orbitrap_Velos
Q_Exactive

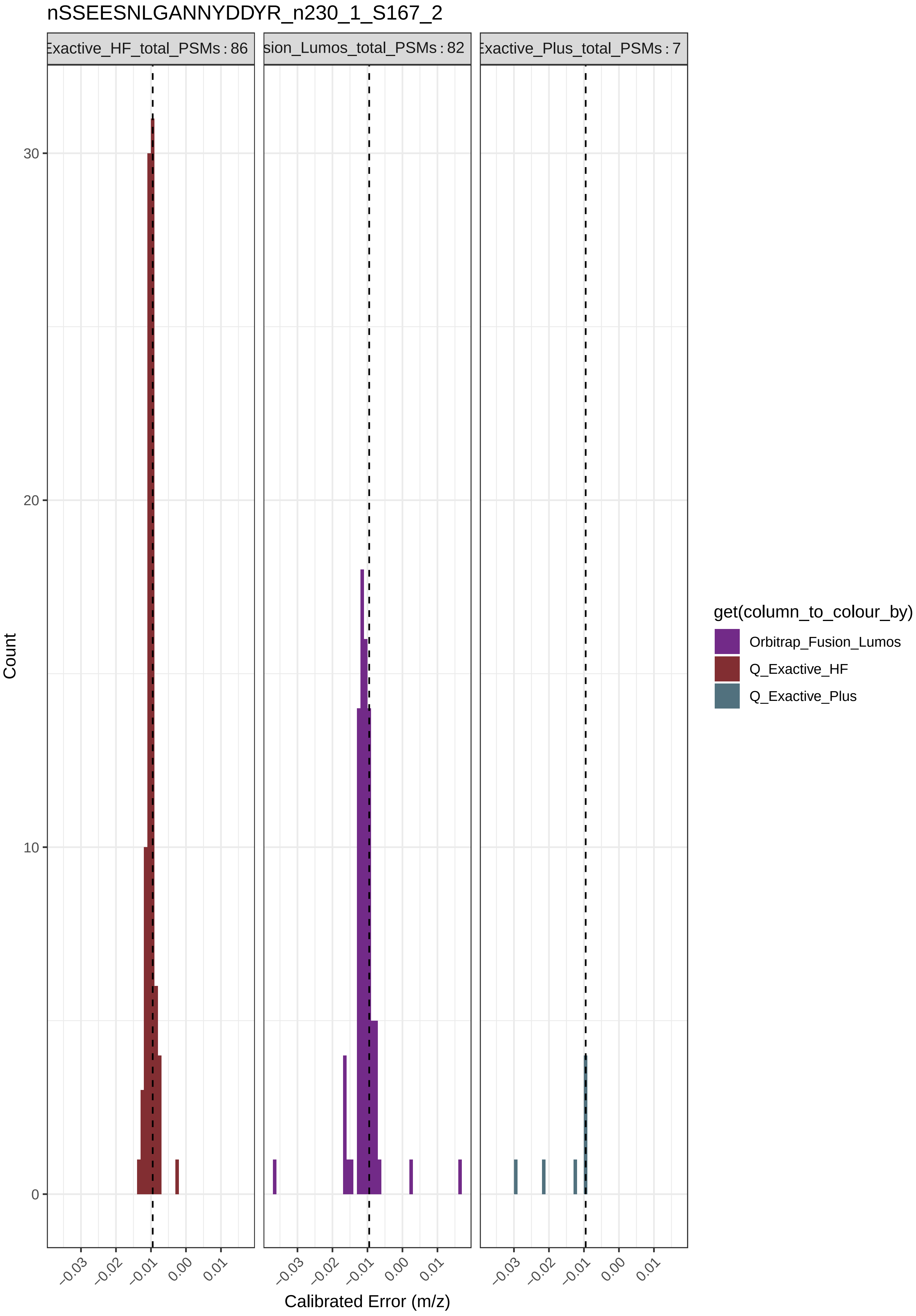

nSSEESNLGANNYDDYR_n230_1_S167_2
Exactive_HF_total_PSMs : 86
sion_Lumos_total_PSMs : 82
Exactive_Plus_total_PSMs : 7
Count
30
20
10
0
−0.03
−0.02
−0.01
0.00
0.01
Calibrated Error (m/z)
get(column_to_colour_by)
Orbitrap_Fusion_Lumos
Q_Exactive_HF
Q_Exactive_Plus

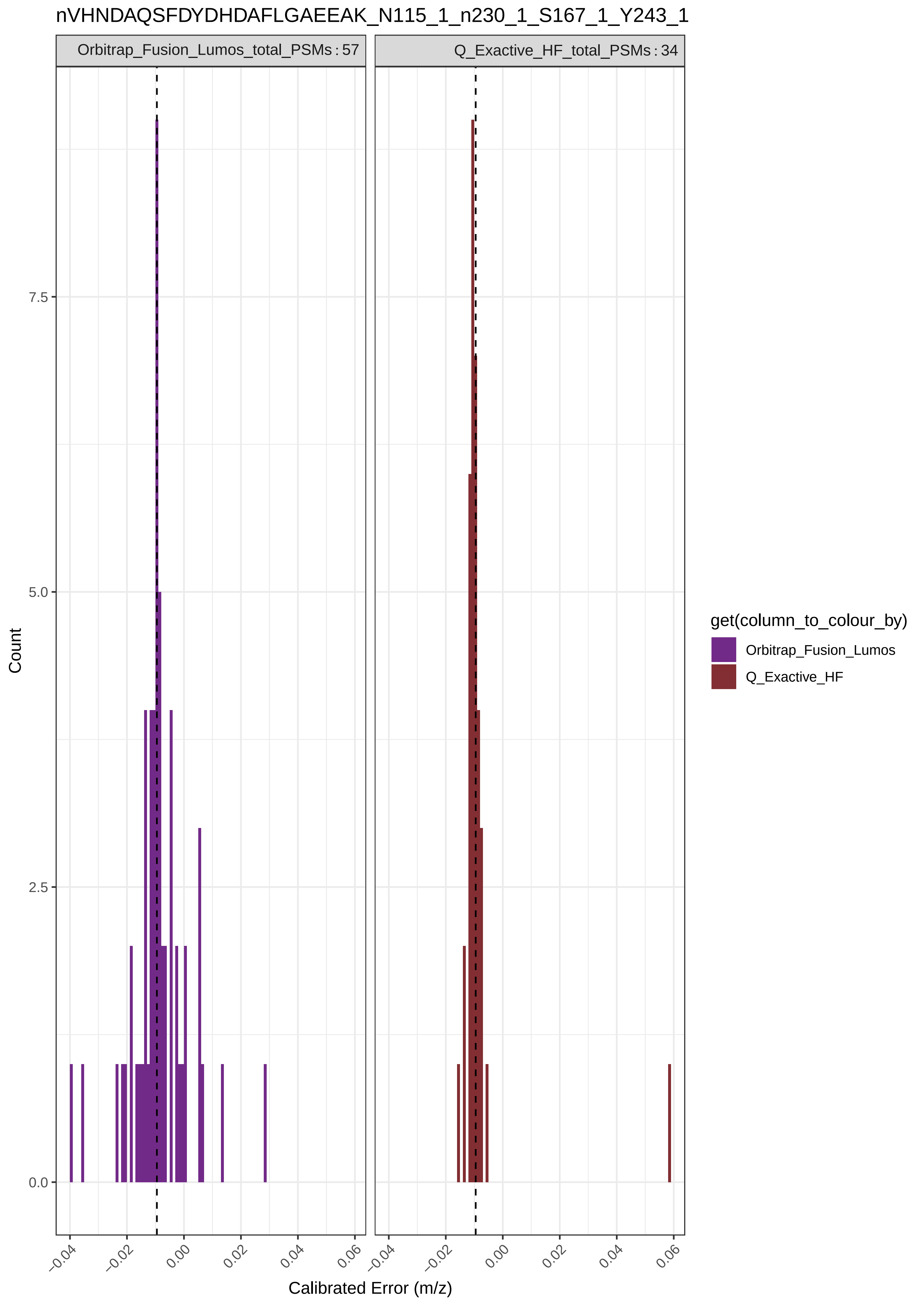
nVHNDAQSFDYDHDAFLGAEEAK_N115_1_n230_1_S167_1_Y243_1
Orbitrap_Fusion_Lumos_total_PSMs : 57
Q_Exactive_HF_total_PSMs : 34
Count
0.0
2.5
5.0
7.5
−0.04
−0.02
0.00
0.02
0.04
0.06
Calibrated Error (m/z)
get(column_to_colour_by)
Orbitrap_Fusion_Lumos
Q_Exactive_HF

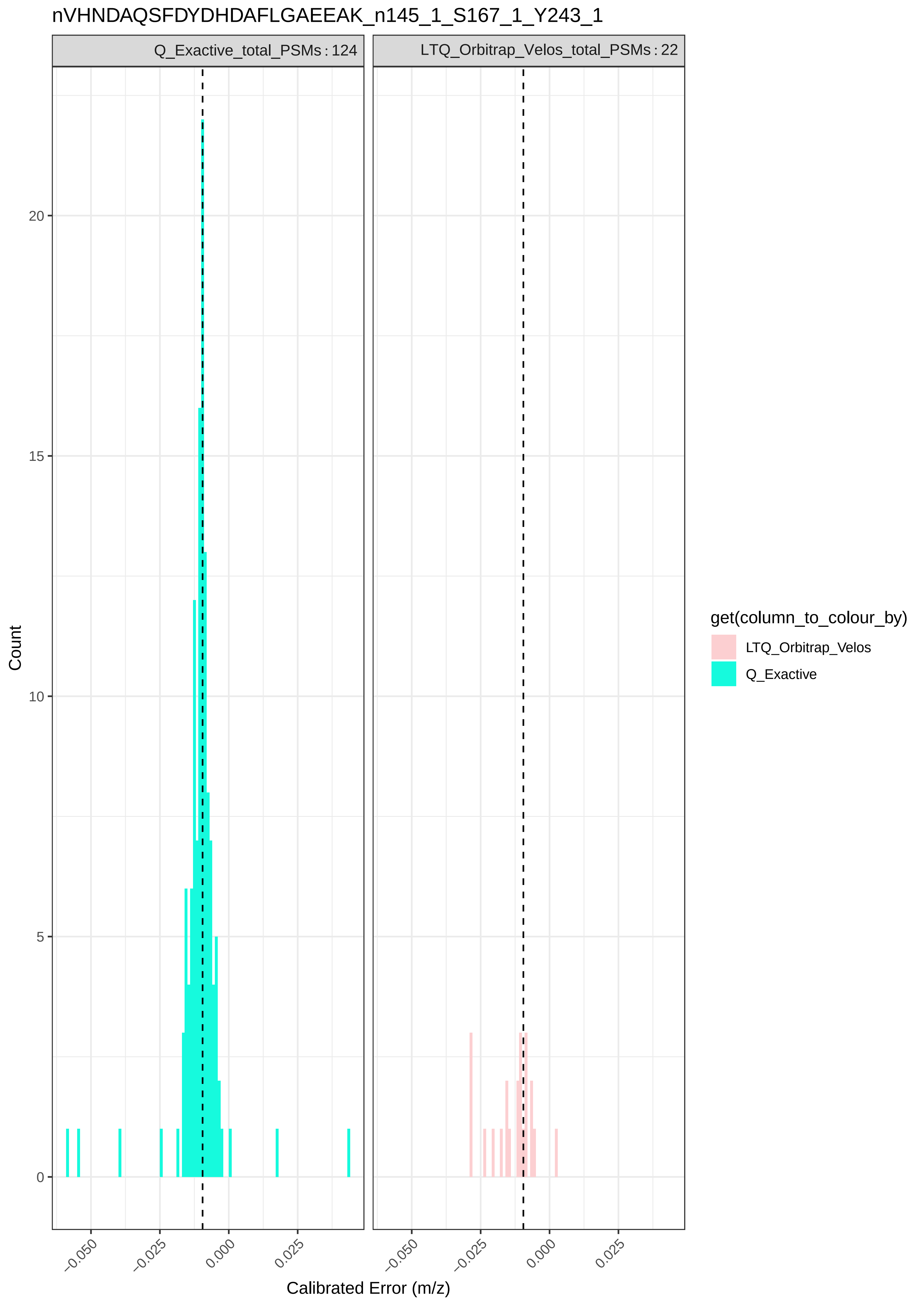
nVHNDAQSFDYDHDAFLGAEEAK_n145_1_S167_1_Y243_1
Q_Exactive_total_PSMs : 124
LTQ_Orbitrap_Velos_total_PSMs : 22
Count
0
5
10
15
20
−0.050
−0.025
0.000
0.025
Calibrated Error (m/z)
get(column_to_colour_by)
LTQ_Orbitrap_Velos
Q_Exactive

### nVHNDAQSFDYDHDAFLGAEEAK_n230_1_S167_1_Y243_1

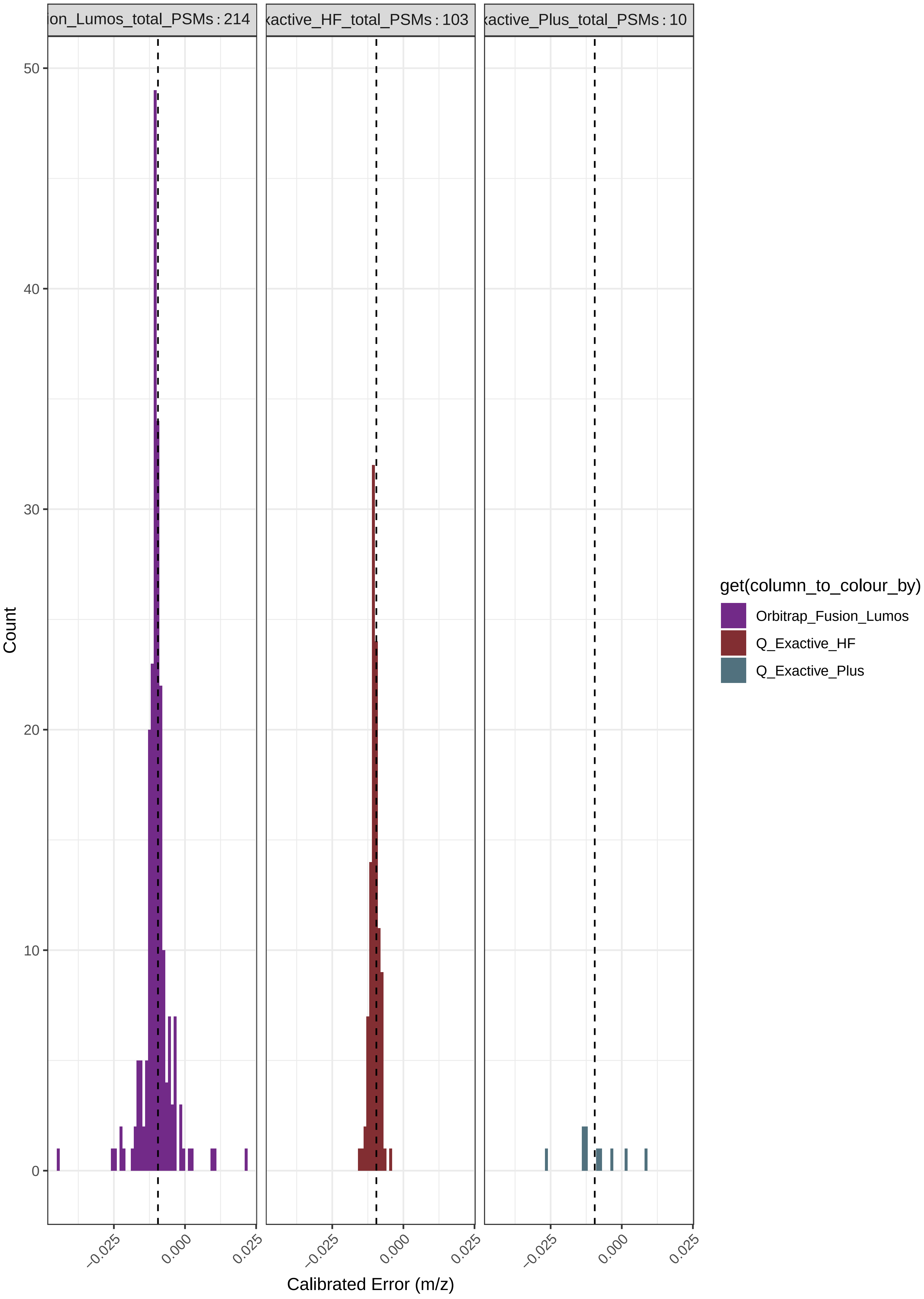

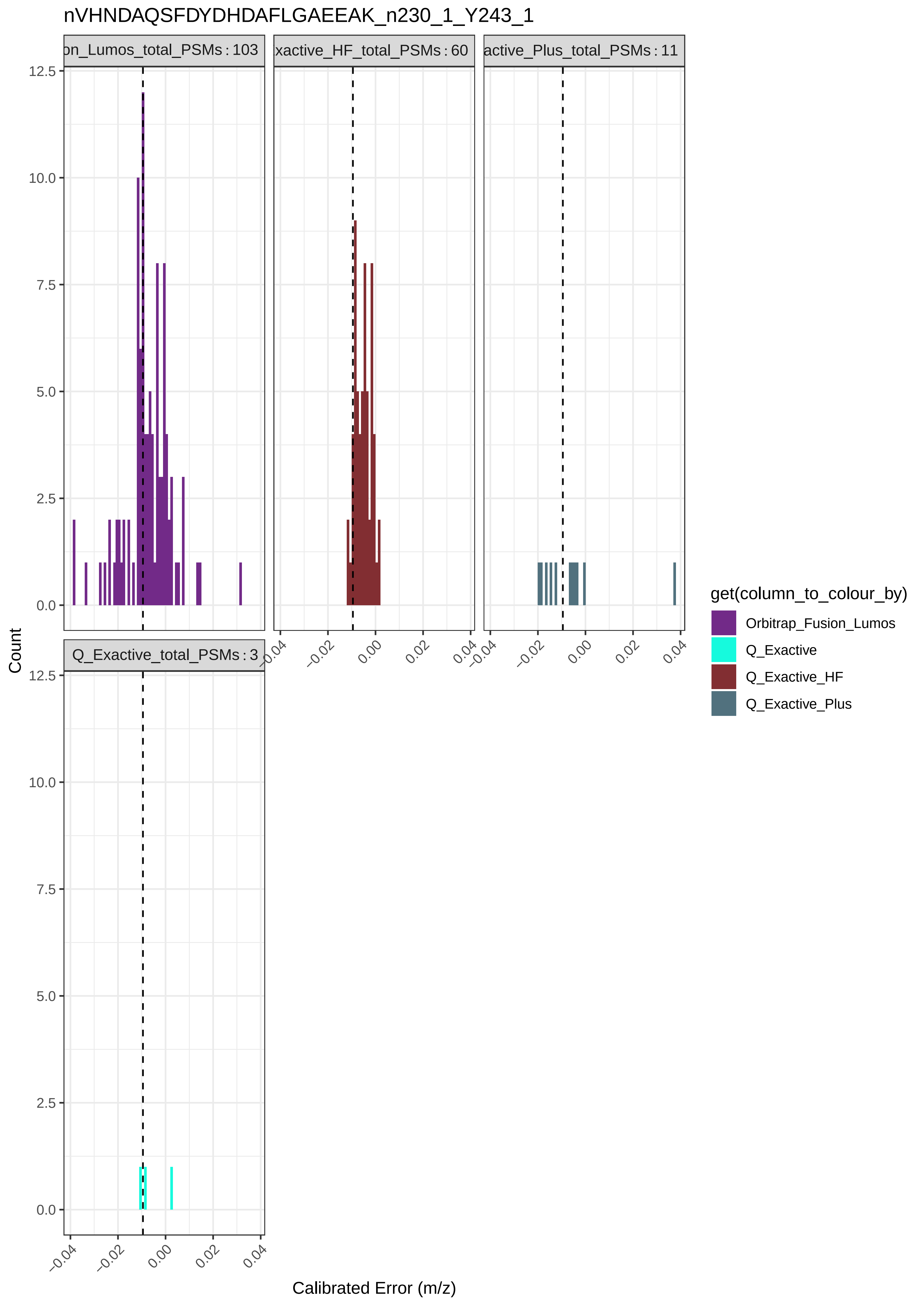

nVHNDAQSFDYDHDAFLGAEEAK_n230_1_Y243_1
on_Lumos_total_PSMs : 103
xactive_HF_total_PSMs : 60
active_Plus_total_PSMs : 11
Q_Exactive_total_PSMs : 3
Count
Calibrated Error (m/z)
0.0
2.5
5.0
7.5
10.0
12.5
−0.04
−0.02
0.00
0.02
0.04
get(column_to_colour_by)
Orbitrap_Fusion_Lumos
Q_Exactive
Q_Exactive_HF
Q_Exactive_Plus

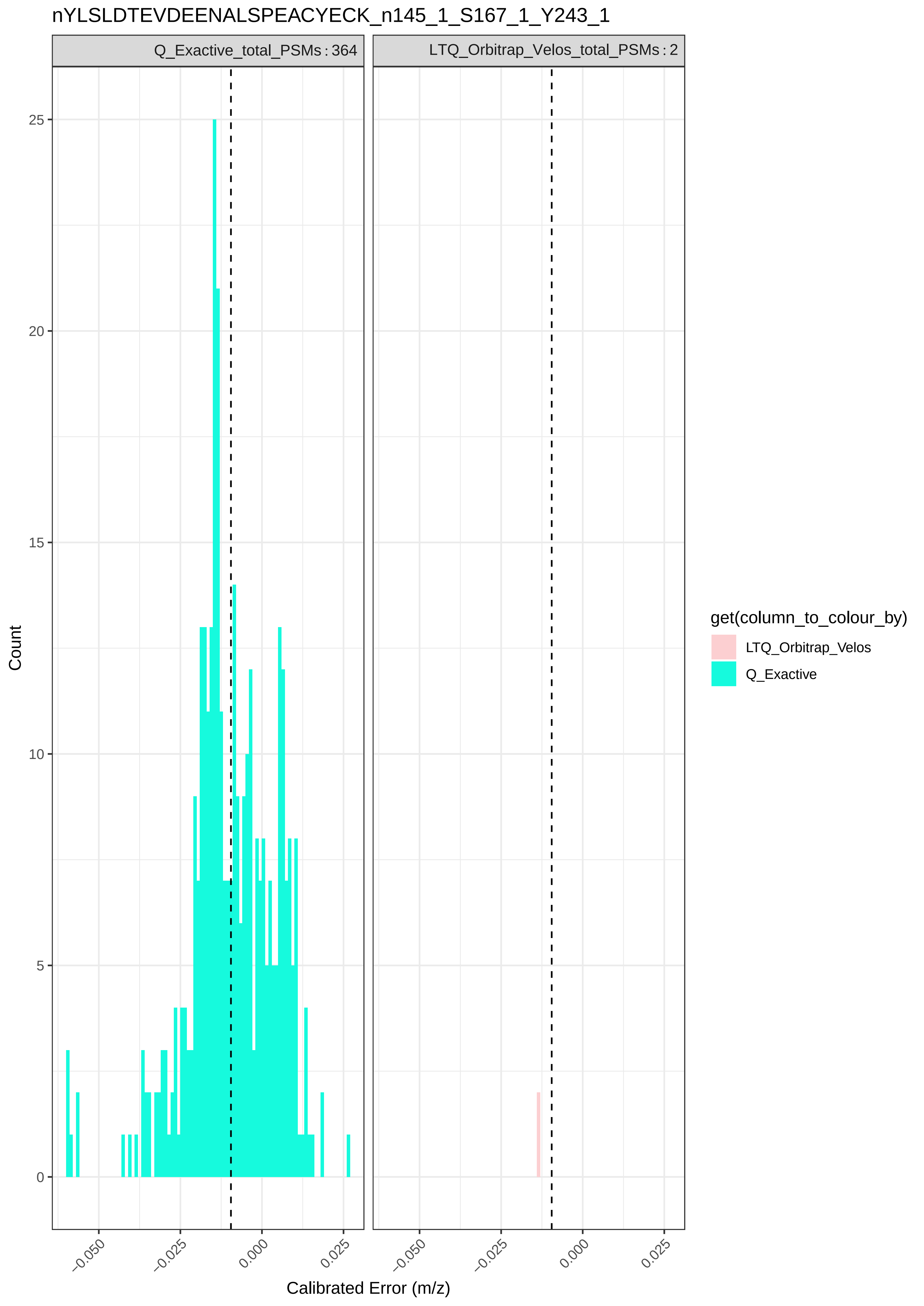

nYLSLDTEVDEENALSPEACYECK_n145_1_S167_1_Y243_1
Q_Exactive_total_PSMs : 364
LTQ_Orbitrap_Velos_total_PSMs : 2
Count
25
20
15
10
5
0
−0.050
−0.025
0.000
0.025
Calibrated Error (m/z)
get(column_to_colour_by)
LTQ_Orbitrap_Velos
Q_Exactive

QMPPPPPCPAGRELFDDPSYVNVQNLDK_M147_1_S167_1

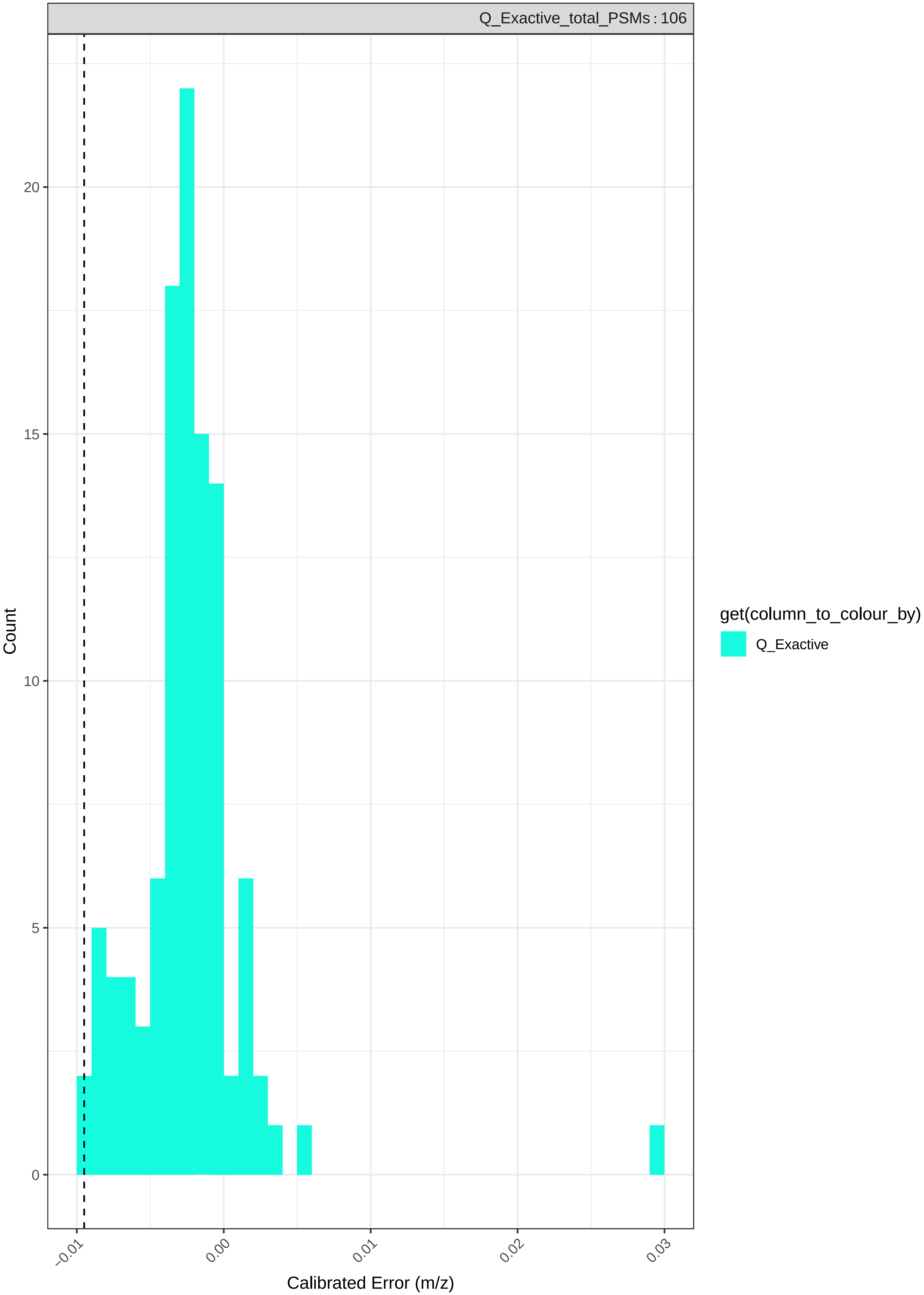

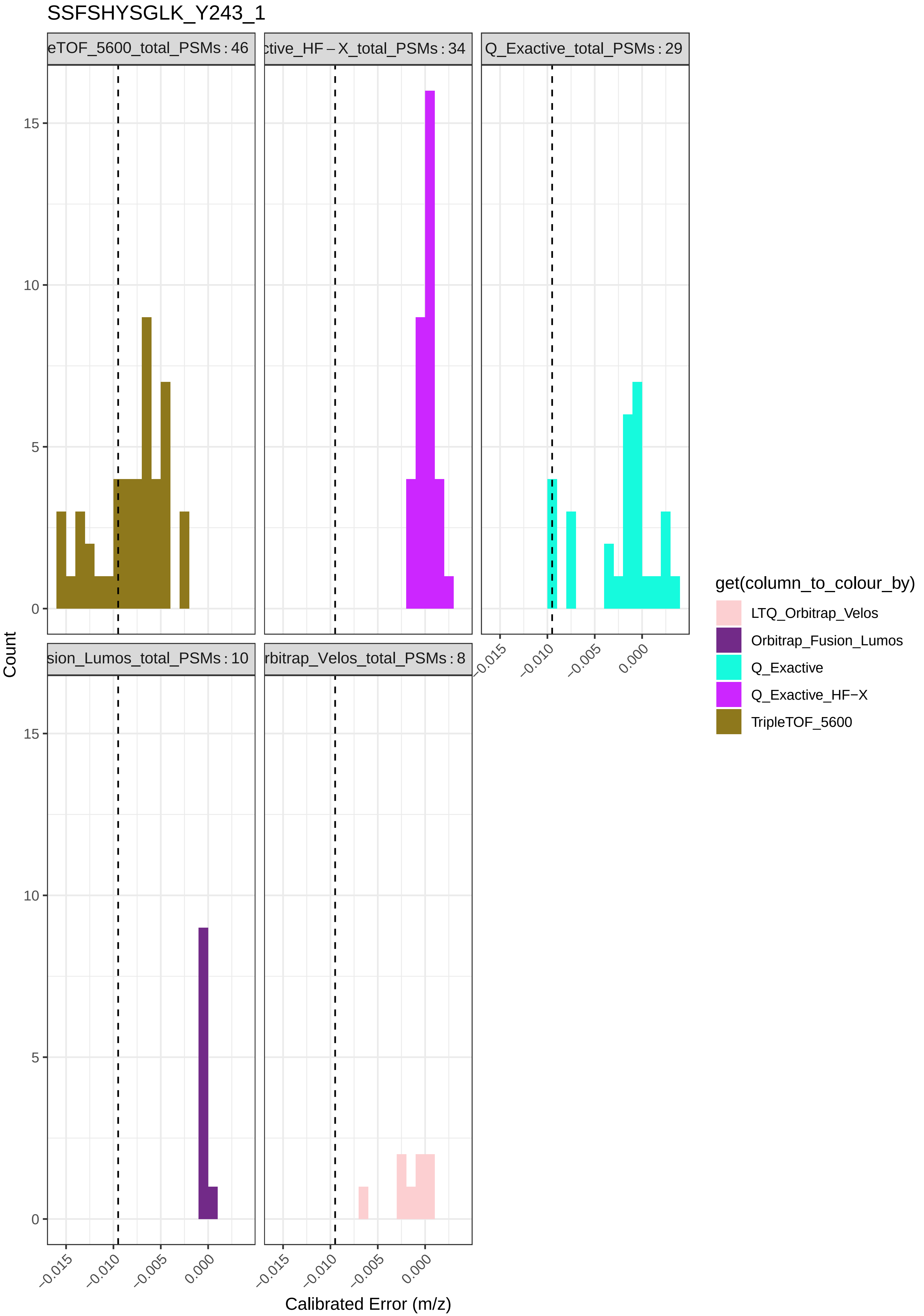
SSFSHYSGLK_Y243_1
eTOF_5600_total_PSMs : 46
ctive_HF – X_total_PSMs : 34
Q_Exactive_total_PSMs : 29
sion_Lumos_total_PSMs : 10
rbitrap_Velos_total_PSMs : 8
Count
0
5
10
15
–0.015
–0.010
–0.005
0.000
Calibrated Error (m/z)
get(column_to_colour_by)
LTQ_Orbitrap_Velos
Orbitrap_Fusion_Lumos
Q_Exactive
Q_Exactive_HF−X
TripleTOF_5600

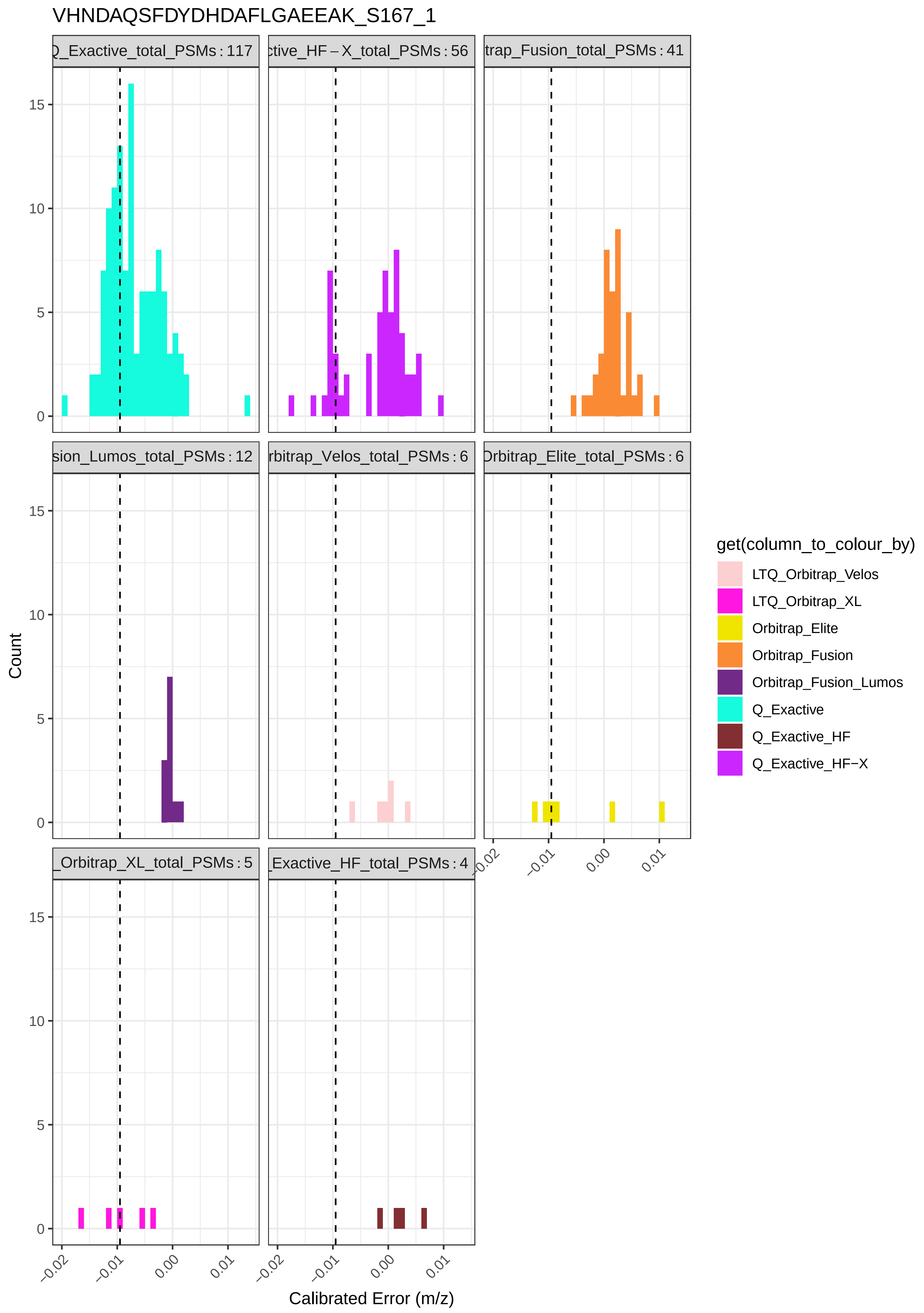

VHNDAQSFDYDHDAFLGAEEAK_S167_1
Q_Exactive_total_PSMs : 117
ctive_HF−X_total_PSMs : 56
trap_Fusion_total_PSMs : 41
sion_Lumos_total_PSMs : 12
rbitrap_Velos_total_PSMs : 6
Orbitrap_Elite_total_PSMs : 6
_Orbitrap_XL_total_PSMs : 5
Exactive_HF_total_PSMs : 4
Count
Calibrated Error (m/z)
get(column_to_colour_by)
LTQ_Orbitrap_Velos
LTQ_Orbitrap_XL
Orbitrap_Elite
Orbitrap_Fusion
Orbitrap_Fusion_Lumos
Q_Exactive
Q_Exactive_HF
Q_Exactive_HF−X

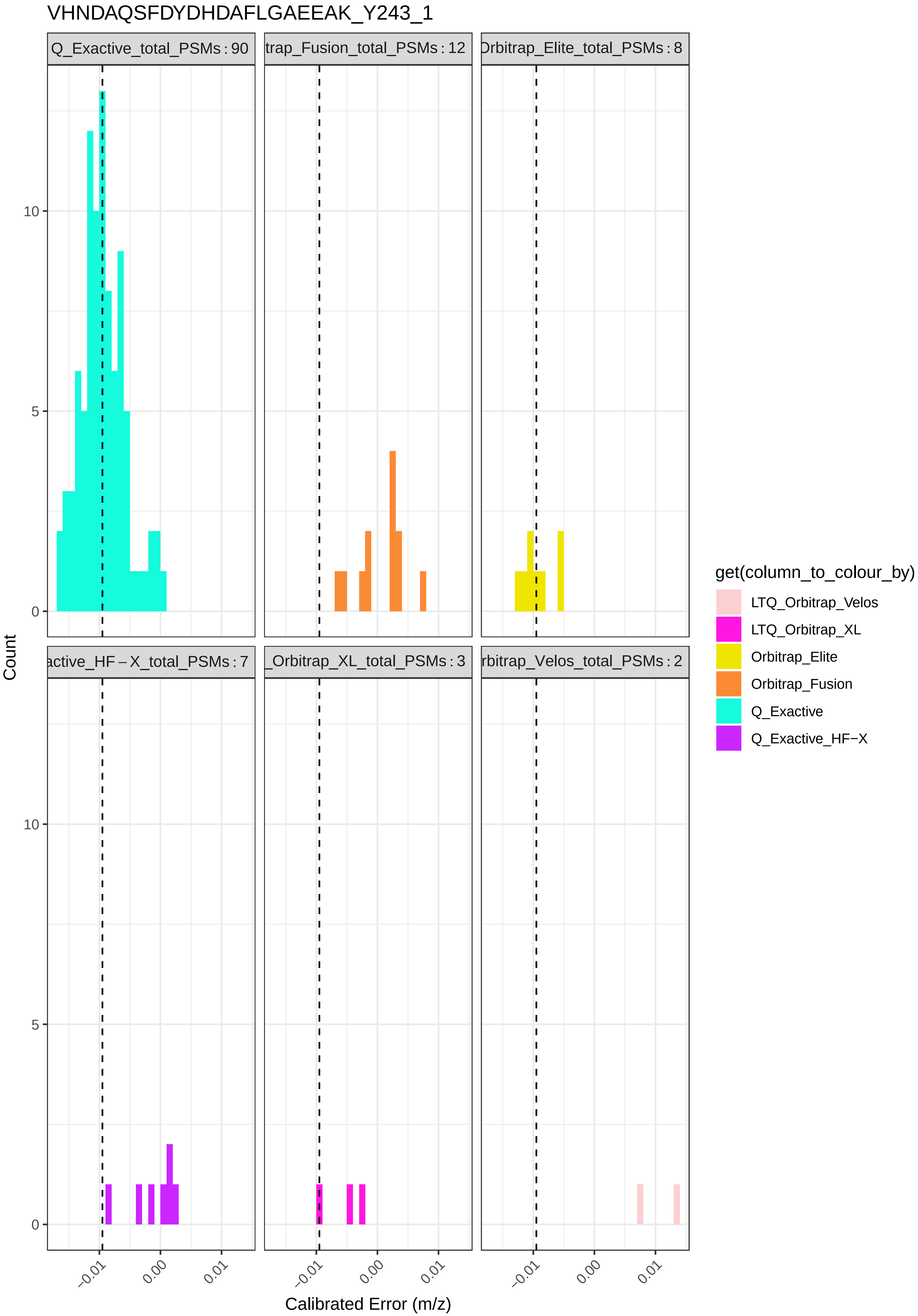
VHNDAQSFDYDHDAFLGAEEAK_Y243_1
Q_Exactive_total_PSMs : 90
trap_Fusion_total_PSMs : 12
Orbitrap_Elite_total_PSMs : 8
active_HF – X_total_PSMs : 7
_Orbitrap_XL_total_PSMs : 3
rbitrap_Velos_total_PSMs : 2
Count
0
5
10
−0.01
0.00
0.01
Calibrated Error (m/z)
get(column_to_colour_by)
LTQ_Orbitrap_Velos
LTQ_Orbitrap_XL
Orbitrap_Elite
Orbitrap_Fusion
Q_Exactive
Q_Exactive_HF−X
