## Supplementary File 4 for "Searching for Sulfotyrosines (sY) in a HA(pY)STACK": histograms_post_auc_filtering_bin_-0.0125_-0.0075.pdf

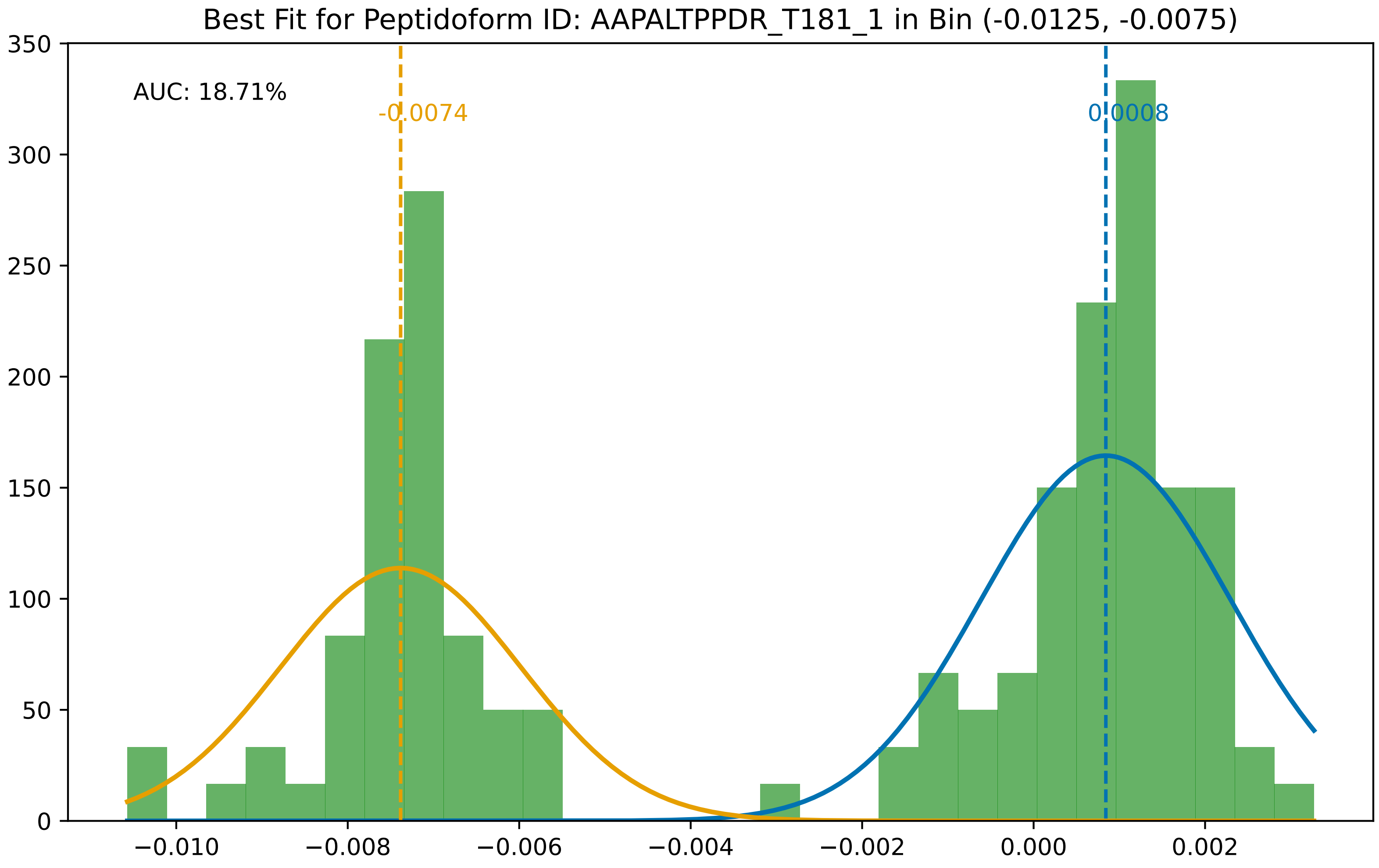
Best Fit for Peptidoform ID: AAPALTPPDR_T181_1 in Bin (-0.0125, -0.0075)
AUC: 18.71%
-0.0074
0.0008
0
50
100
150
200
250
300
350
−0.010
−0.008
−0.006
−0.004
−0.002
0.000
0.002

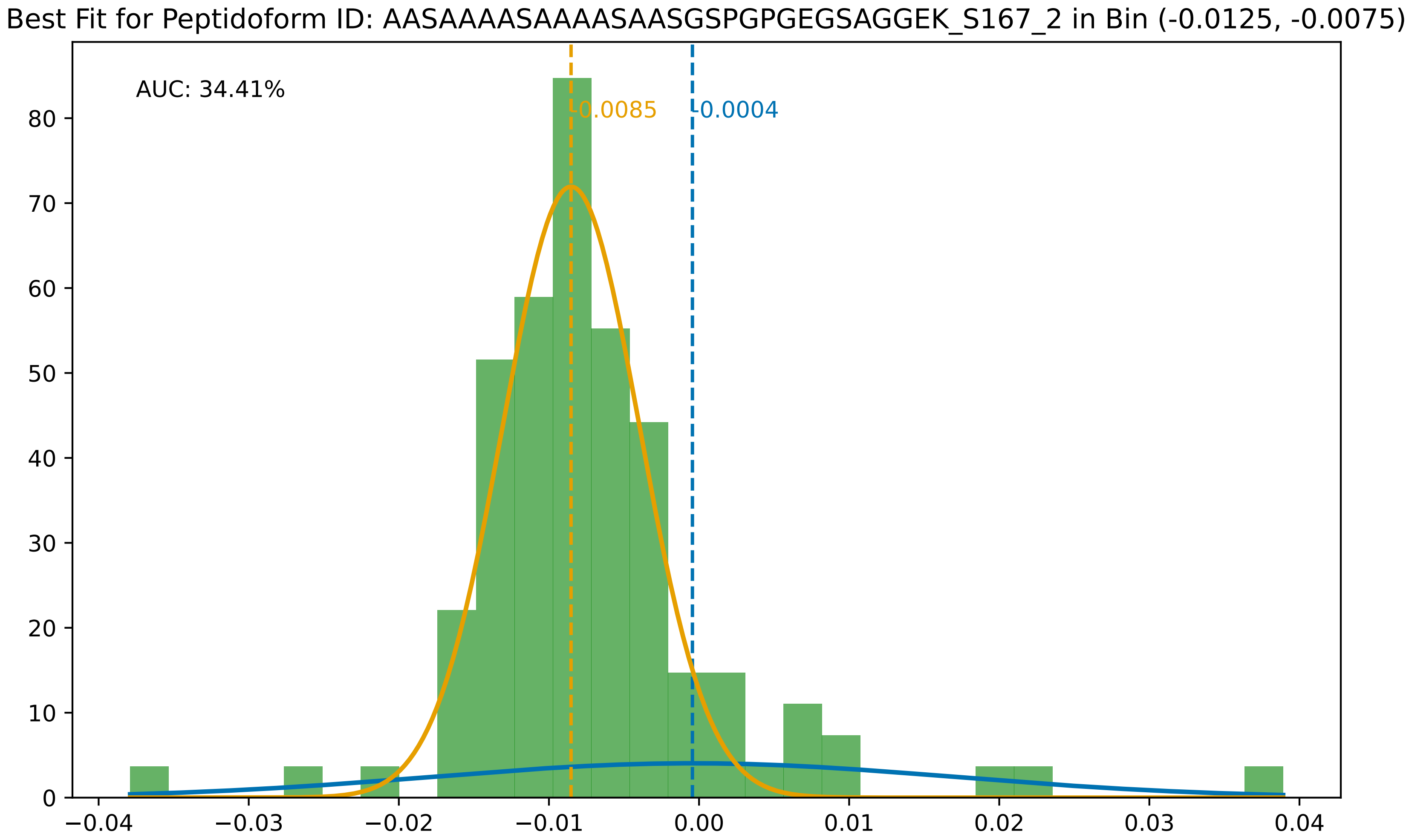
Best Fit for Peptidoform ID: AASAAAASAAAASAASGSPGPGEGSAGGEK_S167_2 in Bin (-0.0125, -0.0075)
AUC: 34.41%
-0.0085
-0.0004
0
10
20
30
40
50
60
70
80
−0.04
−0.03
−0.02
−0.01
0.00
0.01
0.02
0.03
0.04

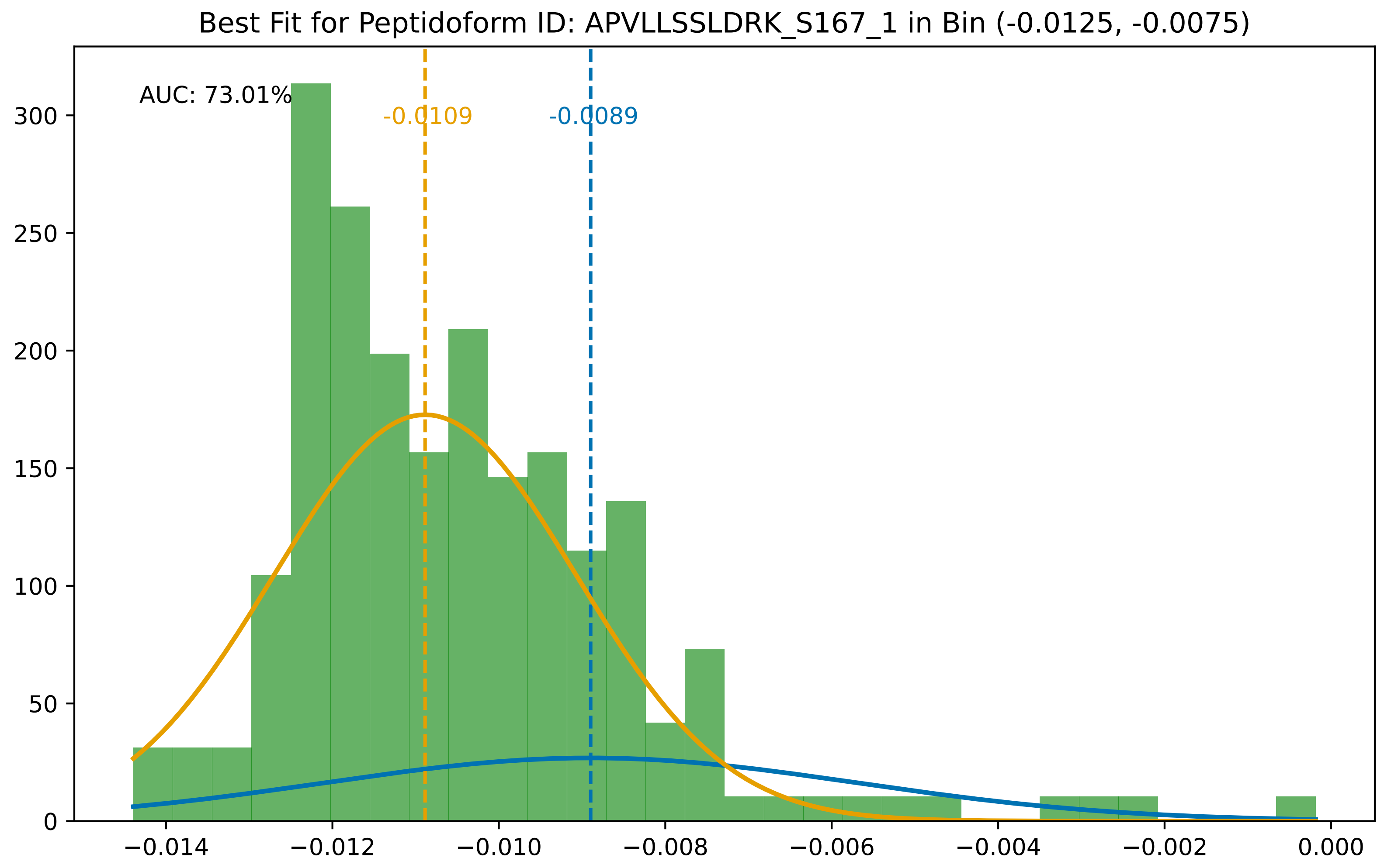

Best Fit for Peptidoform ID: APVLLSSLDRK_S167_1 in Bin (-0.0125, -0.0075)
AUC: 73.01%
-0.0109
-0.0089
300
250
200
150
100
50
0
−0.014
−0.012
−0.010
−0.008
−0.006
−0.004
−0.002
0.000

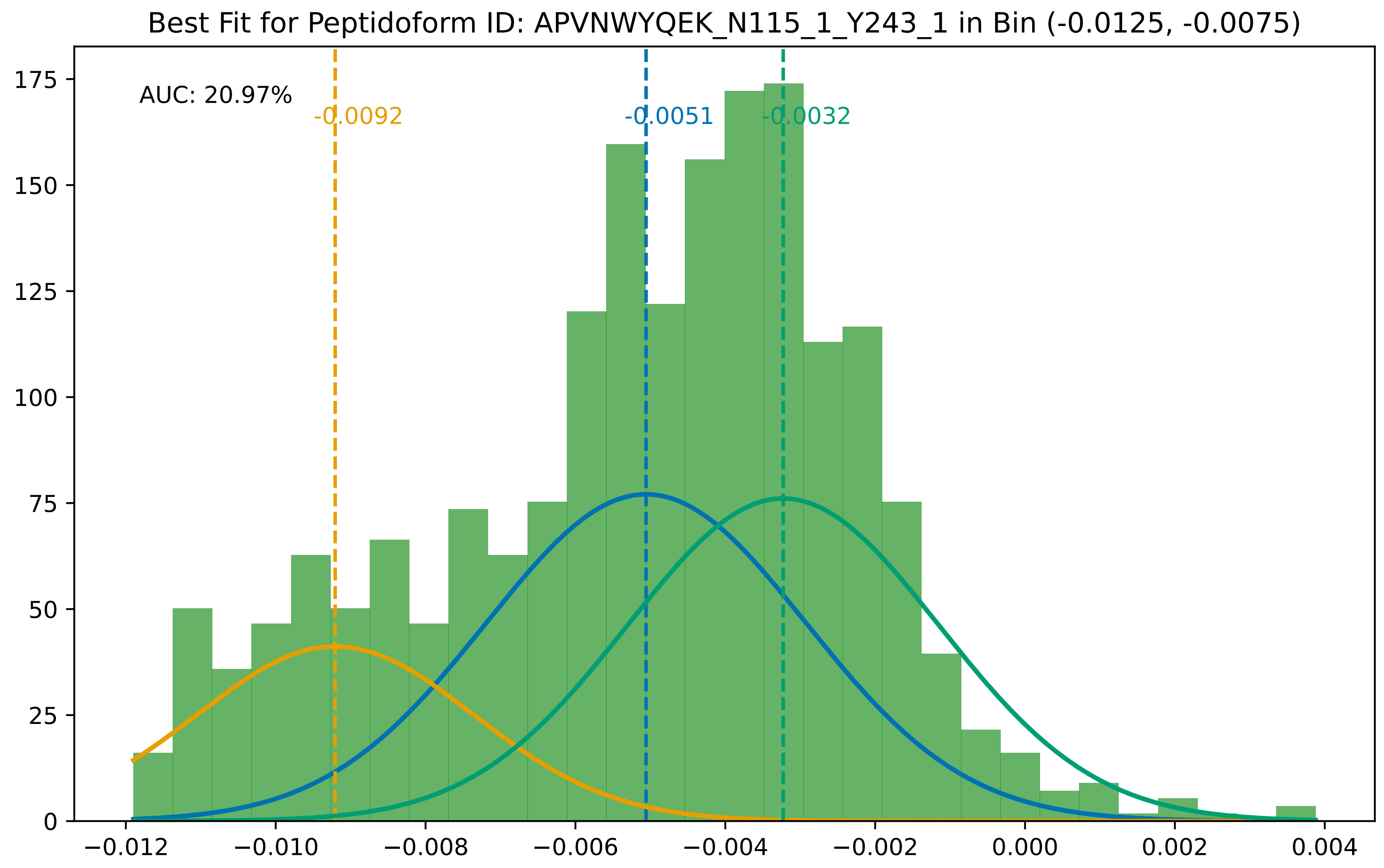
Best Fit for Peptidoform ID: APVNWYQEK_N115_1_Y243_1 in Bin (-0.0125, -0.0075)
AUC: 20.97%
-0.0092
-0.0051
-0.0032
0
25
50
75
100
125
150
175
−0.012
−0.010
−0.008
−0.006
−0.004
−0.002
0.000
0.002
0.004

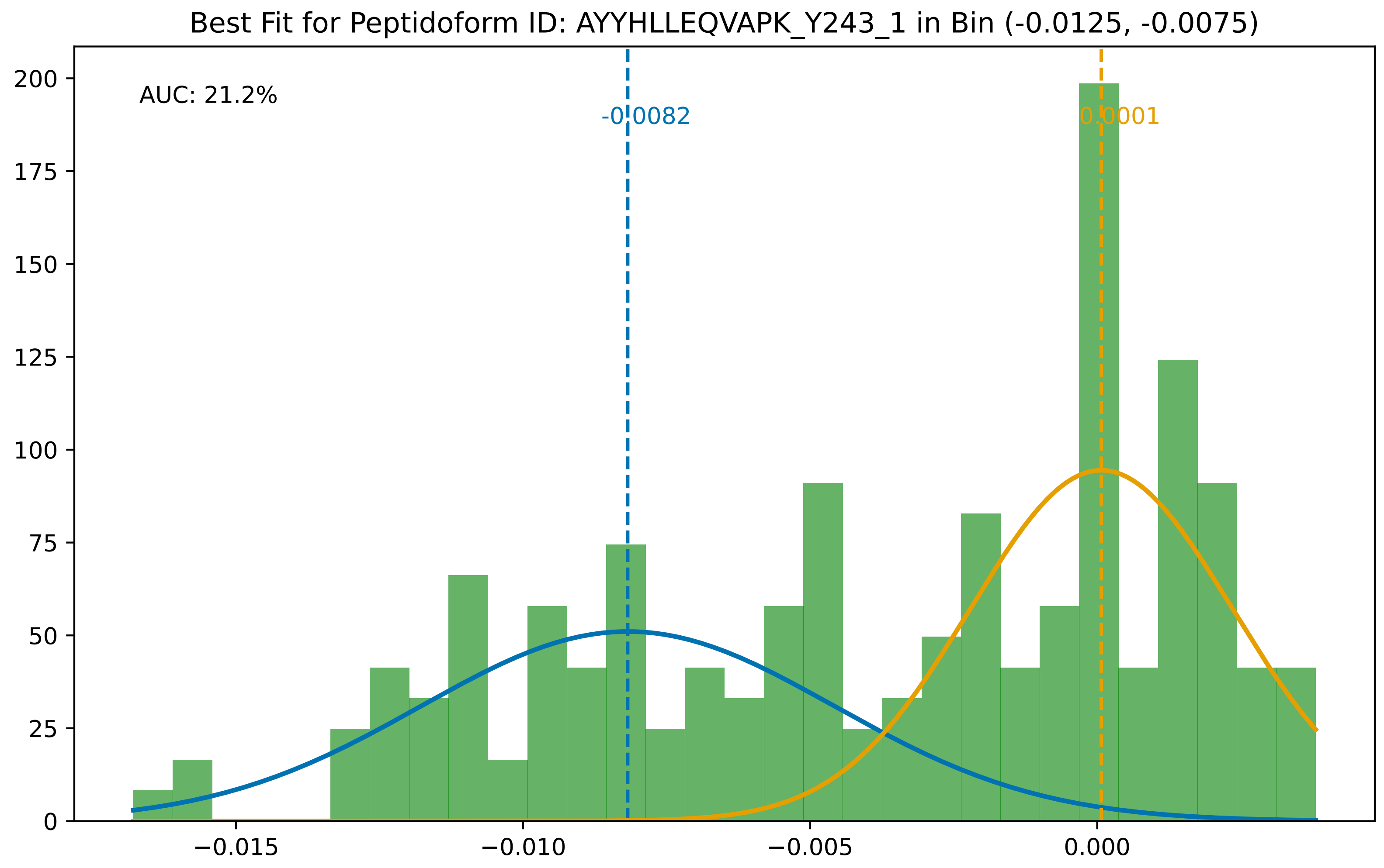

Best Fit for Peptidoform ID: AYYHLLEQVAPK_Y243_1 in Bin (-0.0125, -0.0075)
AUC: 21.2%
-0.0082
0.0001
0
25
50
75
100
125
150
175
200
−0.015
−0.010
−0.005
0.000

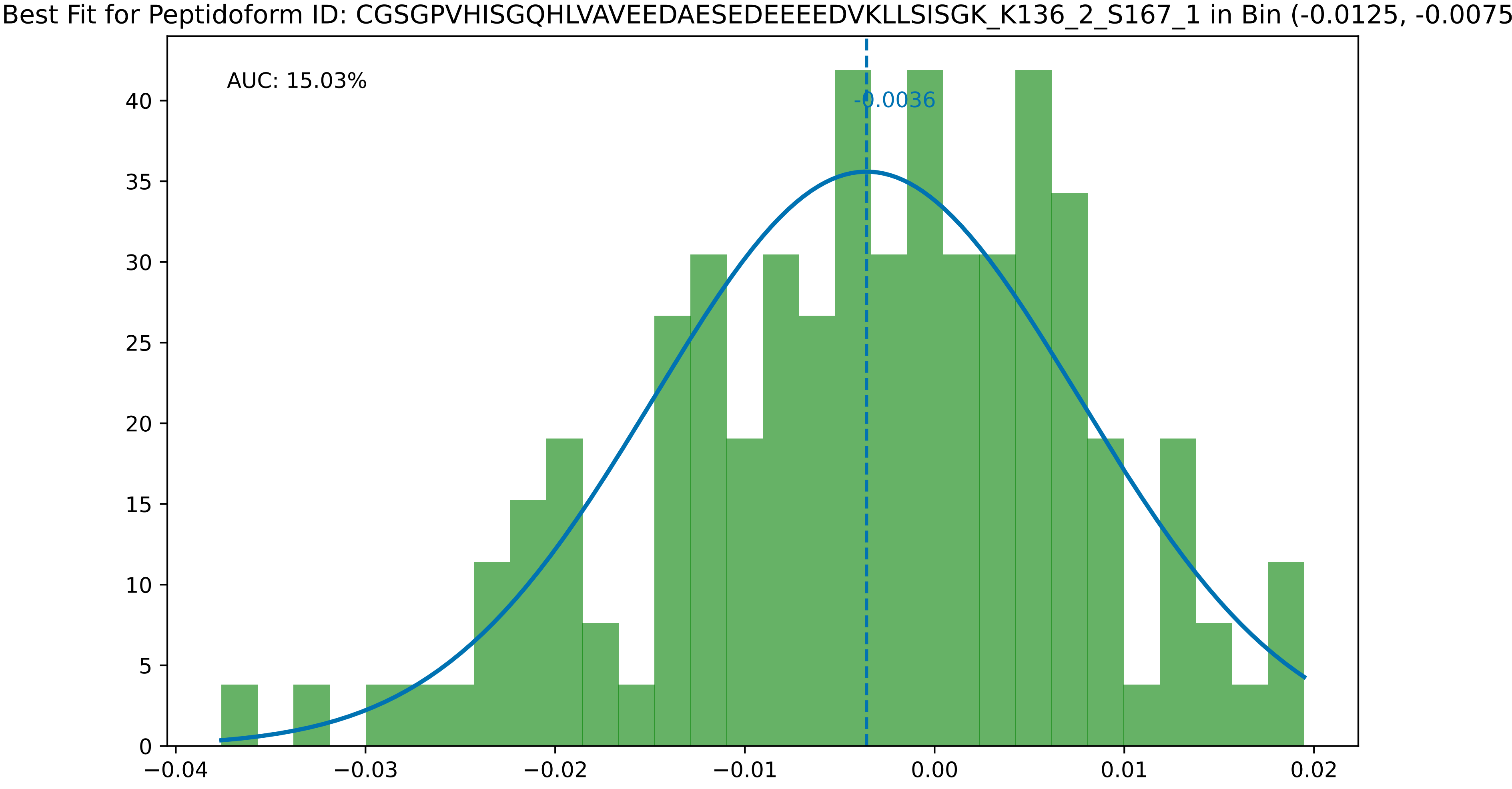
Best Fit for Peptidoform ID: CGSGPVHISGQHLVAVEEDAESEDEEEEDVKLLSISGK_K136_2_S167_1 in Bin (-0.0125, -0.0075
AUC: 15.03%
-0.0036
0
5
10
15
20
25
30
35
40
−0.04
−0.03
−0.02
−0.01
0.00
0.01
0.02

Best Fit for Peptidoform ID: CGSGPVHISGQHLVAVEEDAESEDEEEEDVK_C143_1_K132_1_S167_1 in Bin (-0.0125, -0.0075

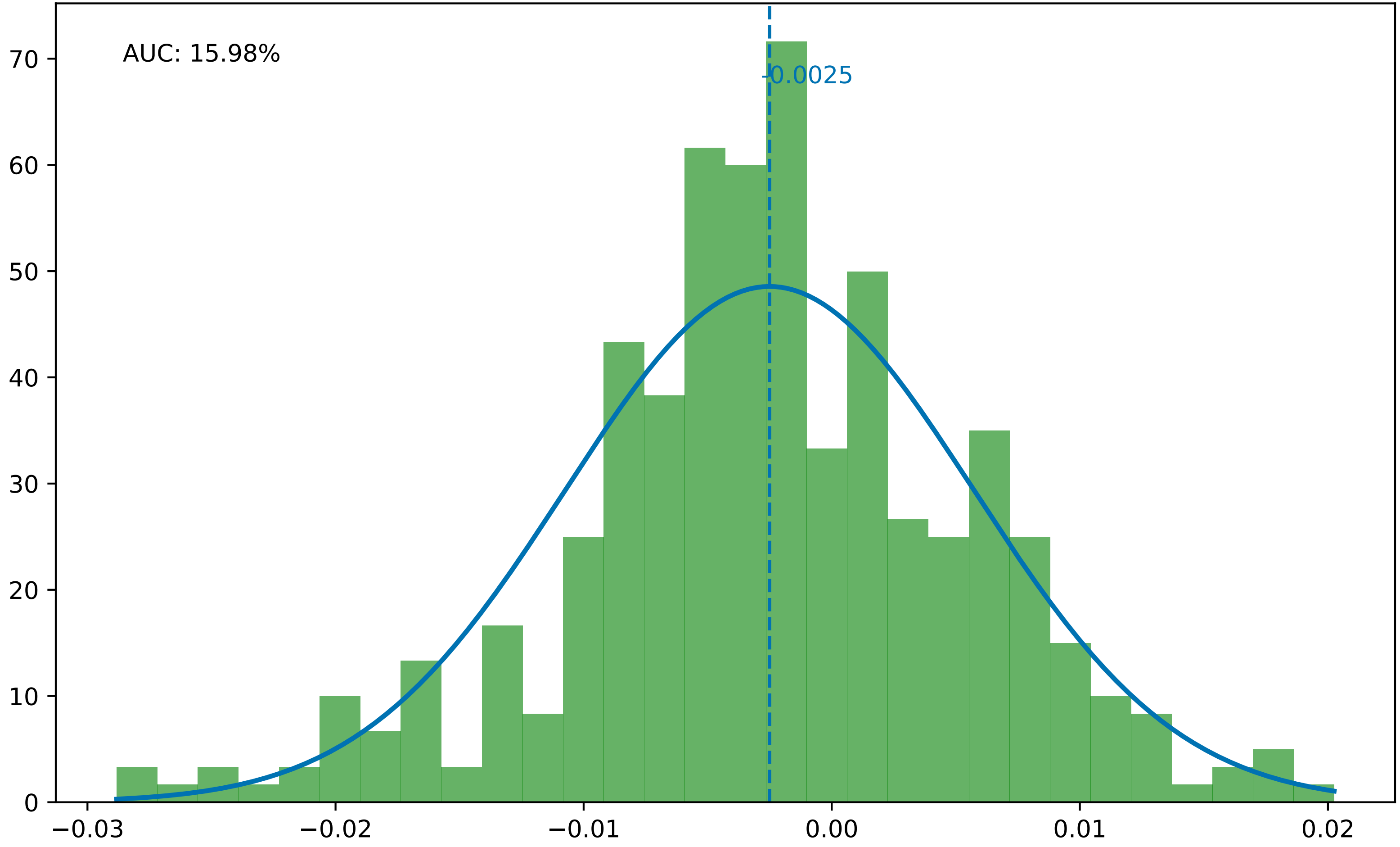

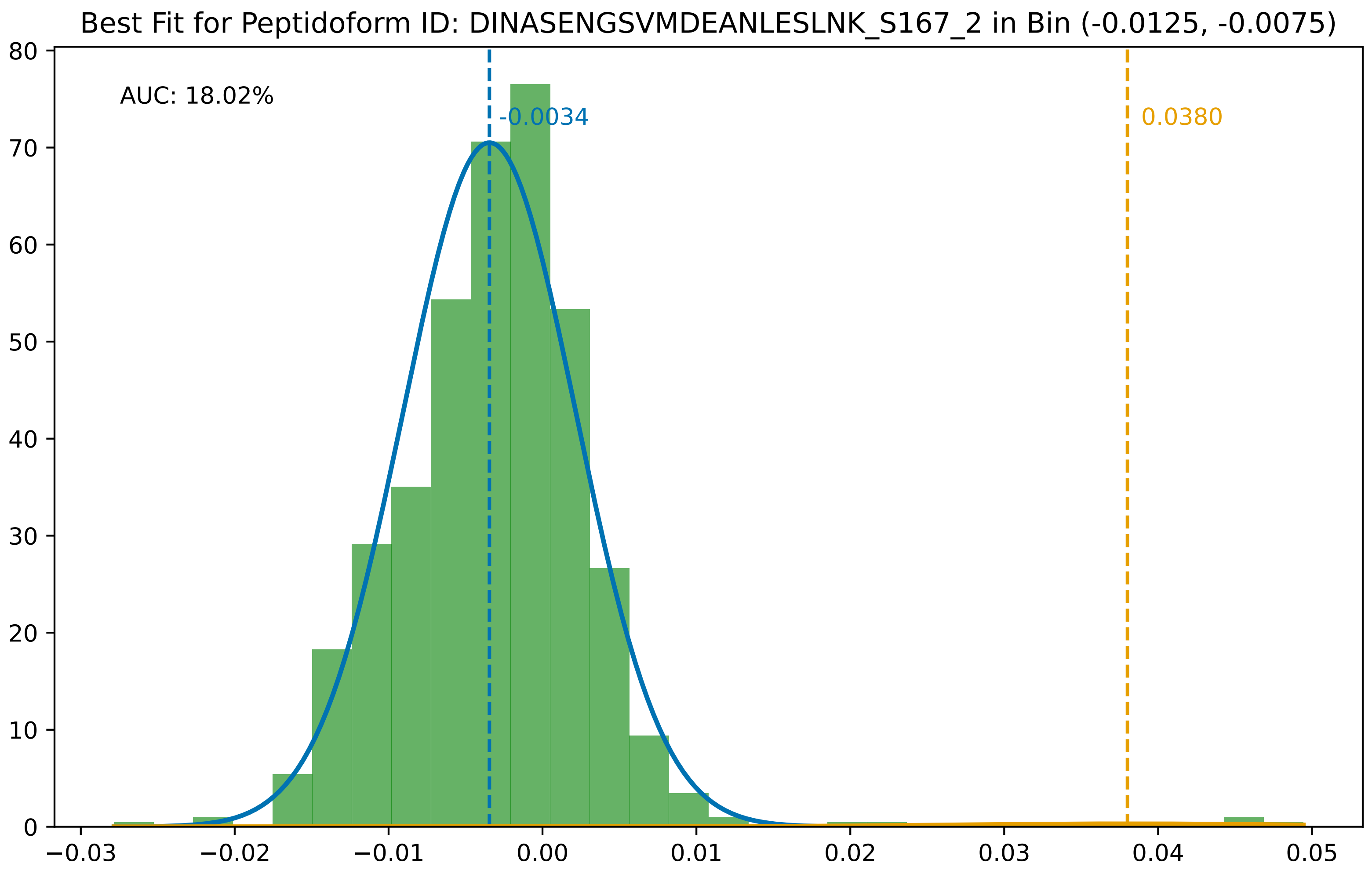

Best Fit for Peptidoform ID: DINASENGSVMDEANLESLNK_S167_2 in Bin (-0.0125, -0.0075)
AUC: 18.02%
-0.0034
0.0380
80
70
60
50
40
30
20
10
0
−0.03
−0.02
−0.01
0.00
0.01
0.02
0.03
0.04
0.05

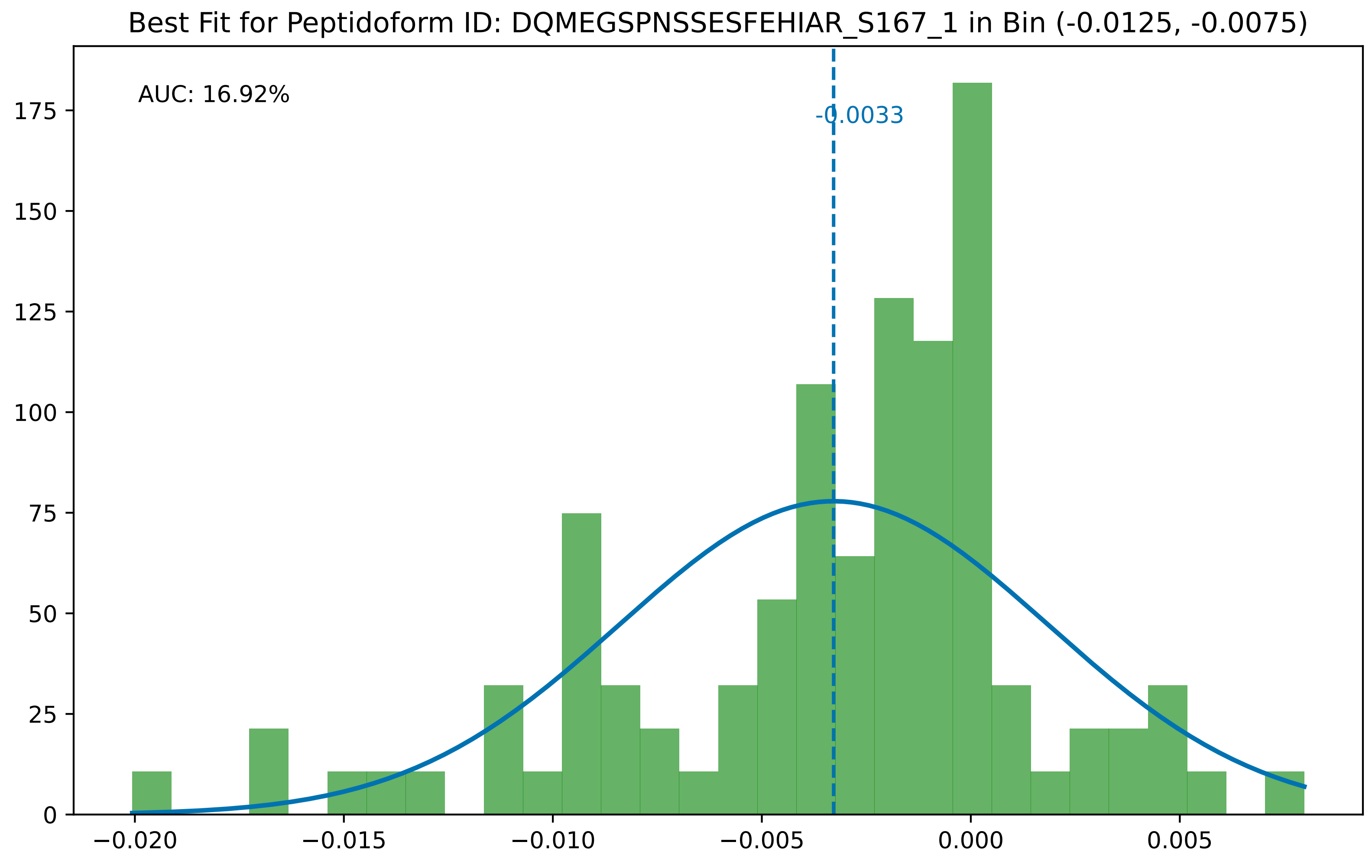

Best Fit for Peptidoform ID: DQMEGSPNSSESFEHIAR_S167_1 in Bin (-0.0125, -0.0075)
AUC: 16.92%
-0.0033
0
25
50
75
100
125
150
175
−0.020
−0.015
−0.010
−0.005
0.000
0.005

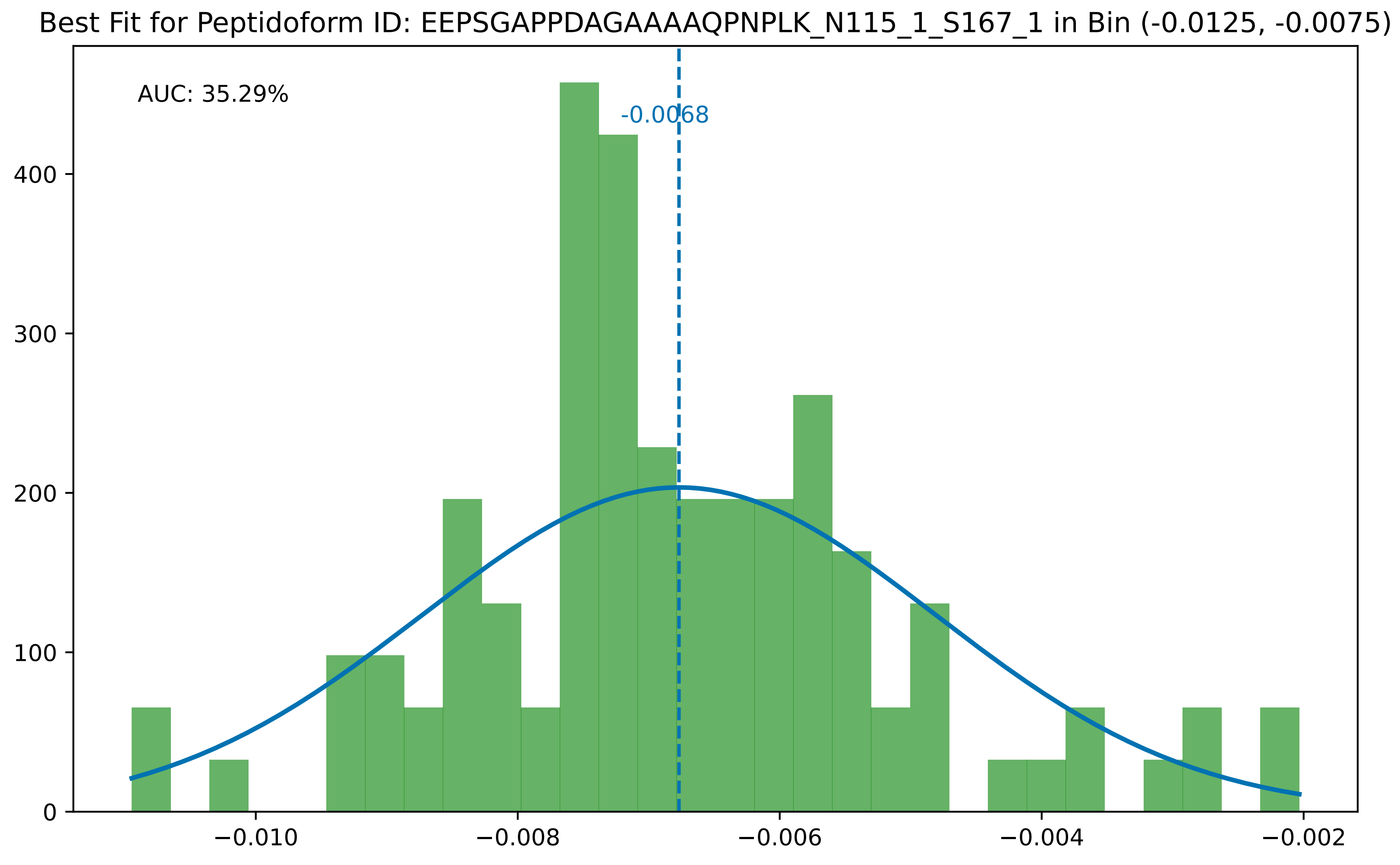
Best Fit for Peptidoform ID: EEPSGAPPDAGAAAAQPNPLK_N115_1_S167_1 in Bin (-0.0125, -0.0075)
AUC: 35.29%
-0.0068
0
100
200
300
400
−0.010
−0.008
−0.006
−0.004
−0.002

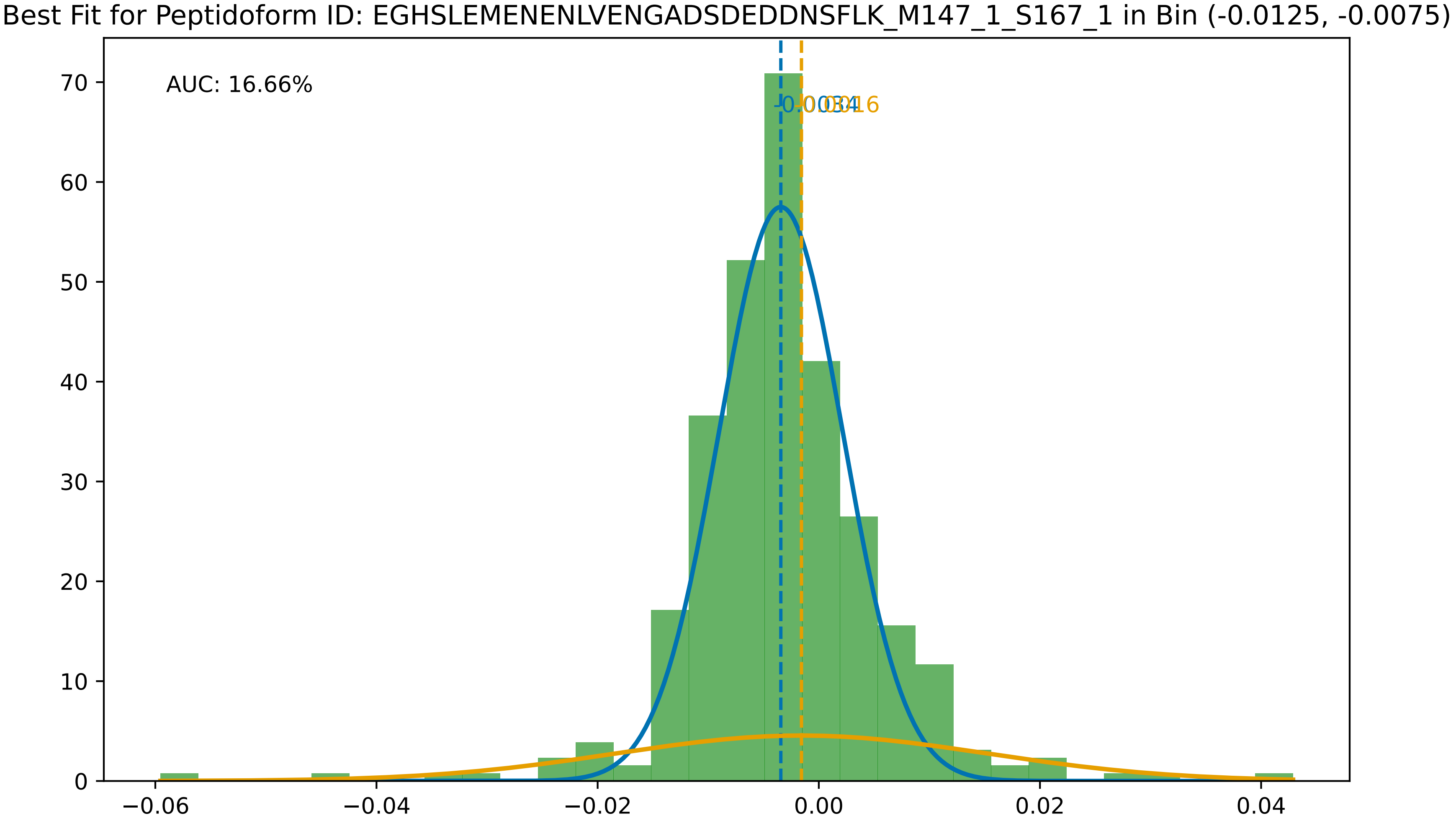
Best Fit for Peptidoform ID: EGHSLEMENENLVENGADSDEDDNSFLK_M147_1_S167_1 in Bin (-0.0125, -0.0075)
AUC: 16.66%
-0.0034
-0.0016
70
60
50
40
30
20
10
0
−0.06
−0.04
−0.02
0.00
0.02
0.04

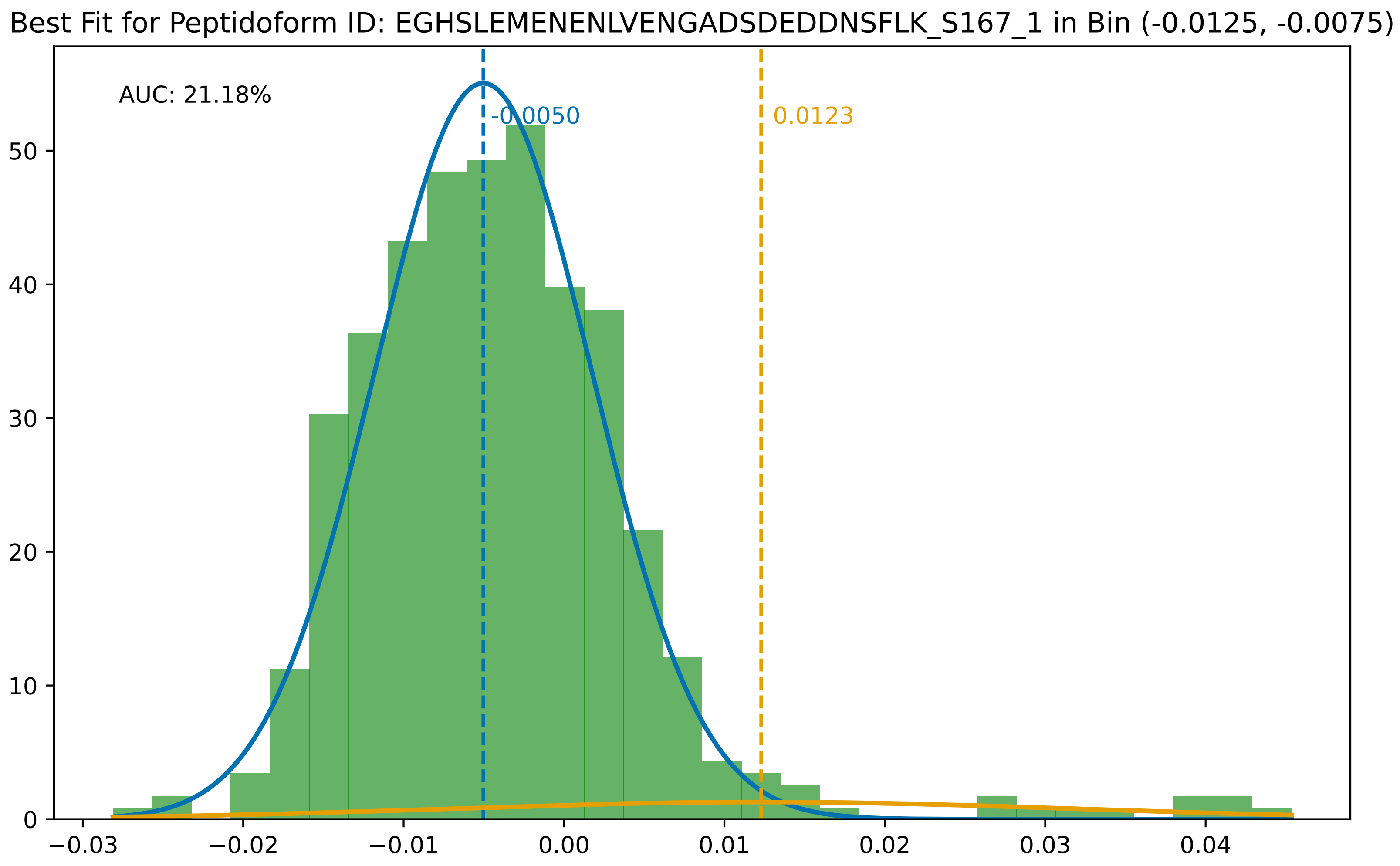

Best Fit for Peptidoform ID: EGHSLEMENENLVENGADSDEDDNSFLK_S167_1 in Bin (-0.0125, -0.0075)
AUC: 21.18%
-0.0050
0.0123
0
10
20
30
40
50
−0.03
−0.02
−0.01
0.00
0.01
0.02
0.03
0.04

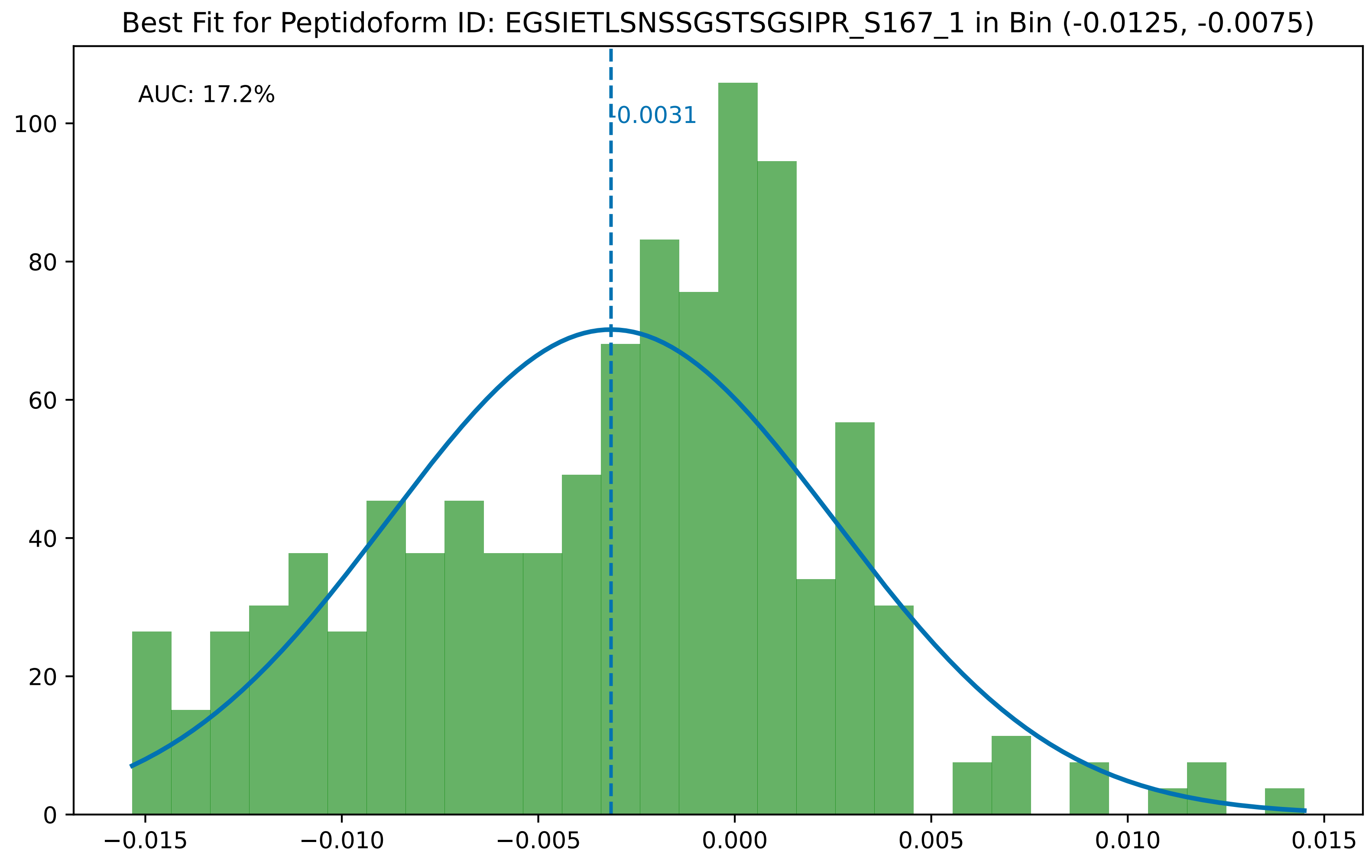

Best Fit for Peptidoform ID: EGSIETLSNSSGSTSGSIPR_S167_1 in Bin (-0.0125, -0.0075)
AUC: 17.2%
-0.0031
0
20
40
60
80
100
−0.015
−0.010
−0.005
0.000
0.005
0.010
0.015

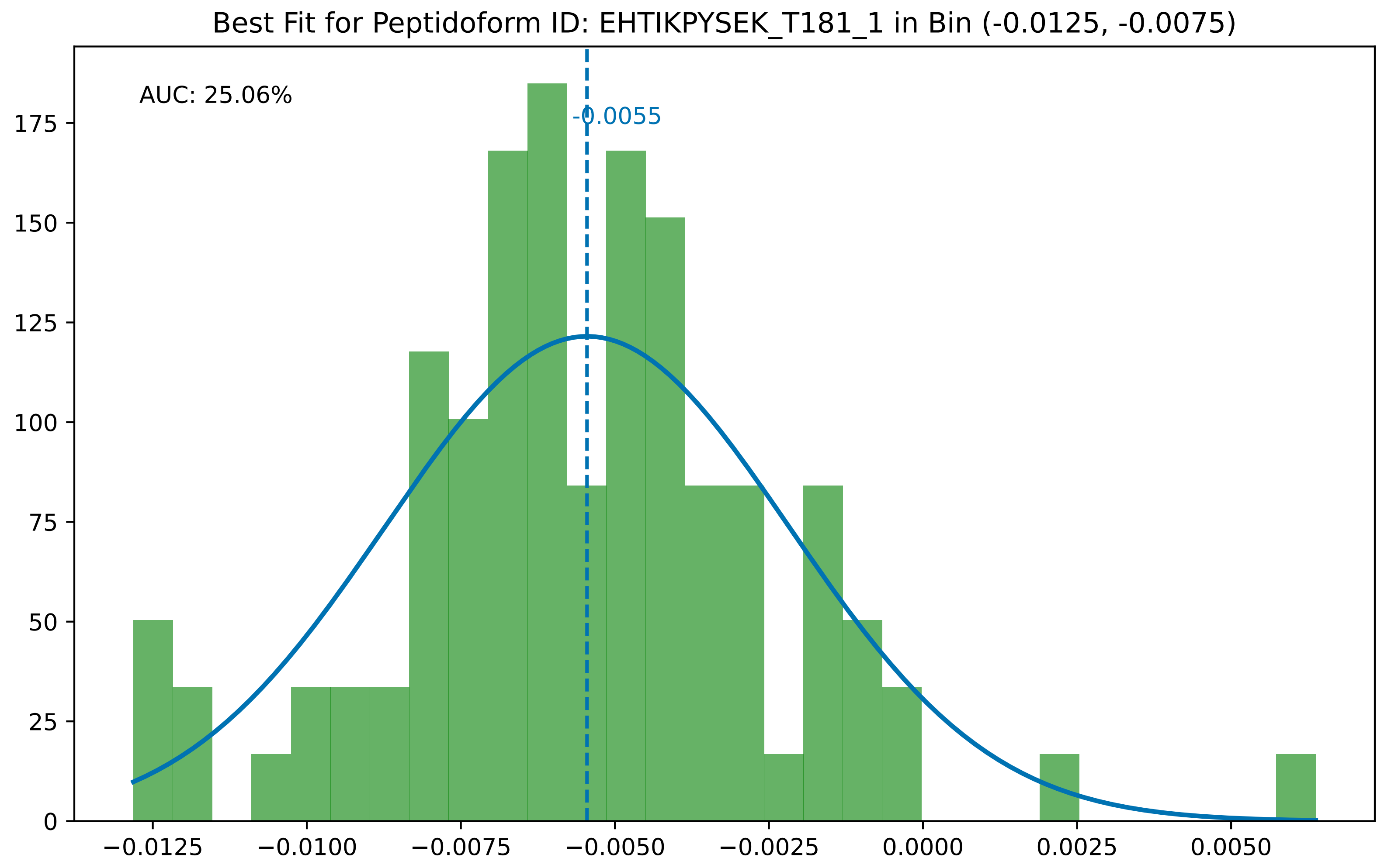

Best Fit for Peptidoform ID: EHTIKPYSEK_T181_1 in Bin (-0.0125, -0.0075)
AUC: 25.06%
-0.0055
175
150
125
100
75
50
25
0
−0.0125
−0.0100
−0.0075
−0.0050
−0.0025
0.0000
0.0025
0.0050

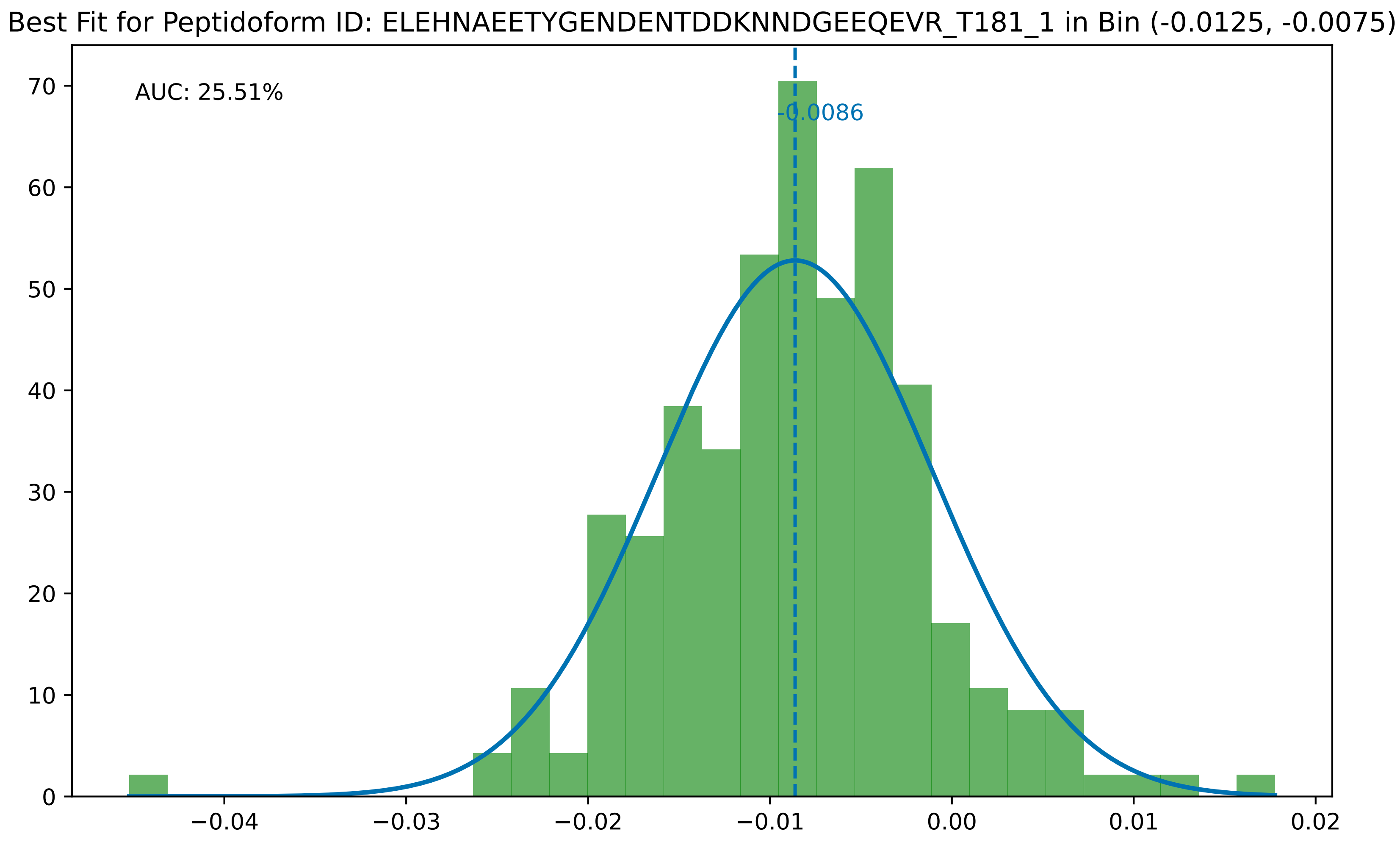
Best Fit for Peptidoform ID: ELEHNAEETYGENDENTDDKNNDGEEQEVR_T181_1 in Bin (-0.0125, -0.0075)
AUC: 25.51%
-0.0086
0
10
20
30
40
50
60
70
−0.04
−0.03
−0.02
−0.01
0.00
0.01
0.02

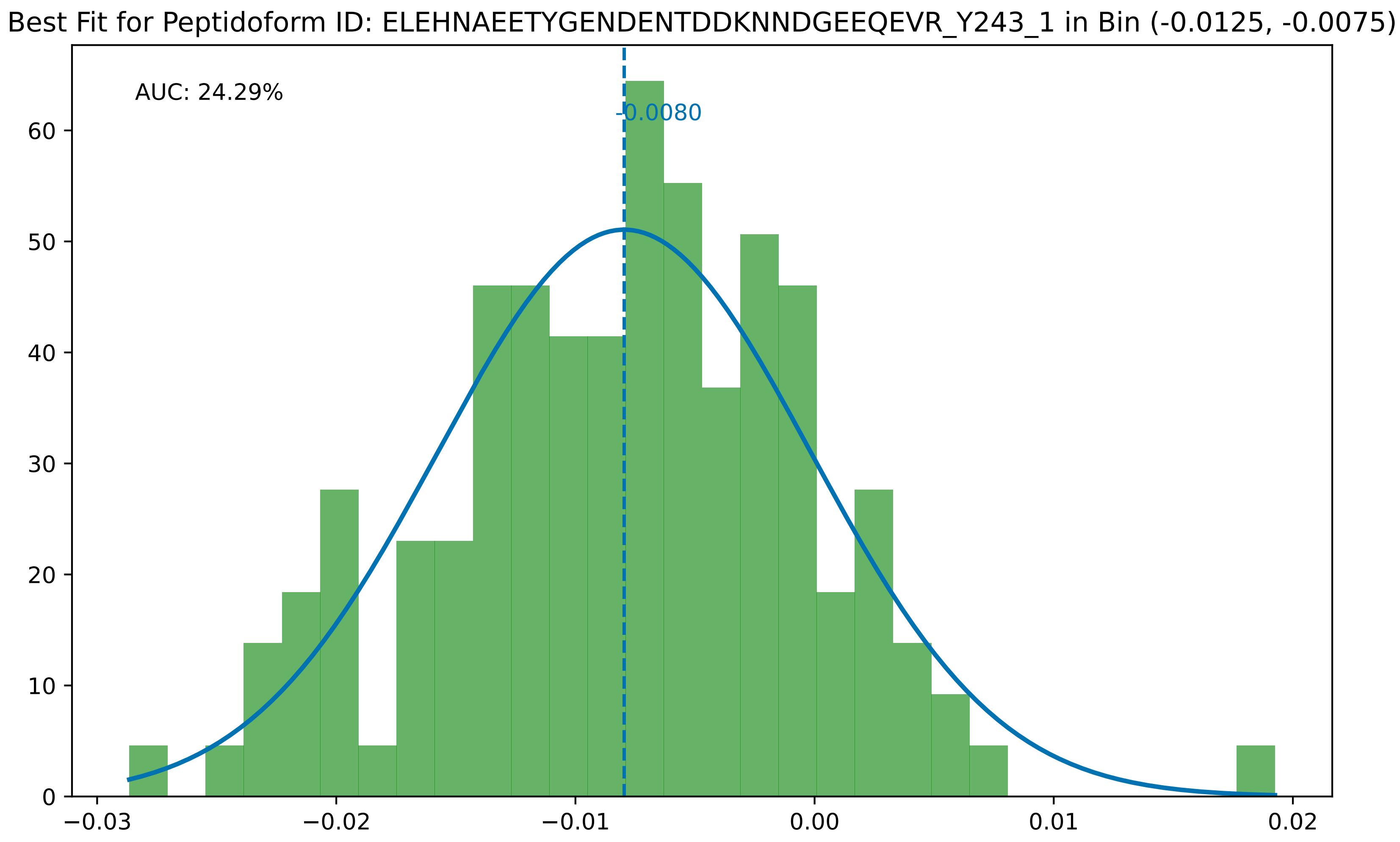

Best Fit for Peptidoform ID: ELEHNAEETYGENDENTDDKNNDGEEQEVR_Y243_1 in Bin (-0.0125, -0.0075)
AUC: 24.29%
-0.0080
0
10
20
30
40
50
60
−0.03
−0.02
−0.01
0.00
0.01
0.02

Best Fit for Peptidoform ID: FEEALQTIFNR_N115_1_Q129_1_T181_1 in Bin (-0.0125, -0.0075)

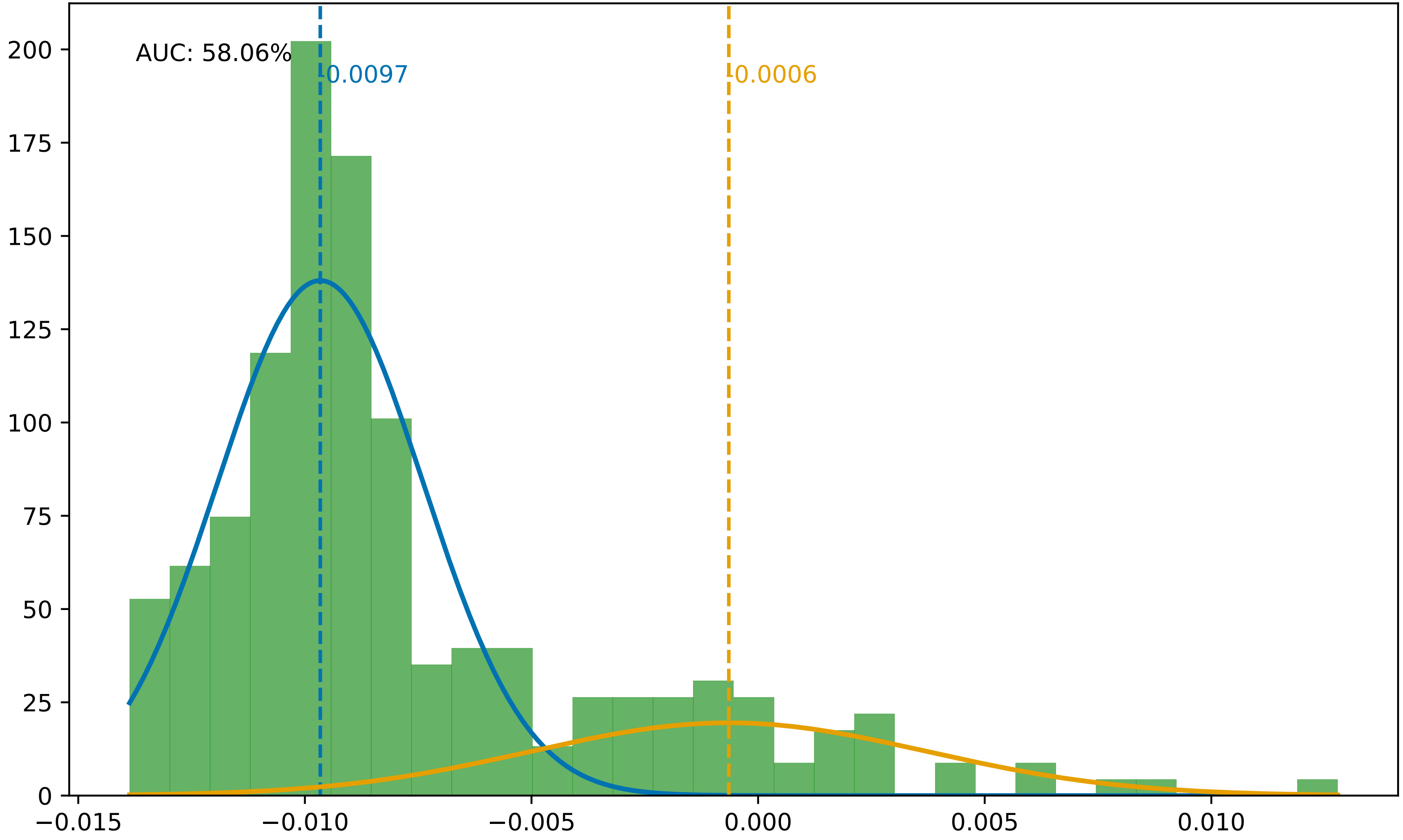

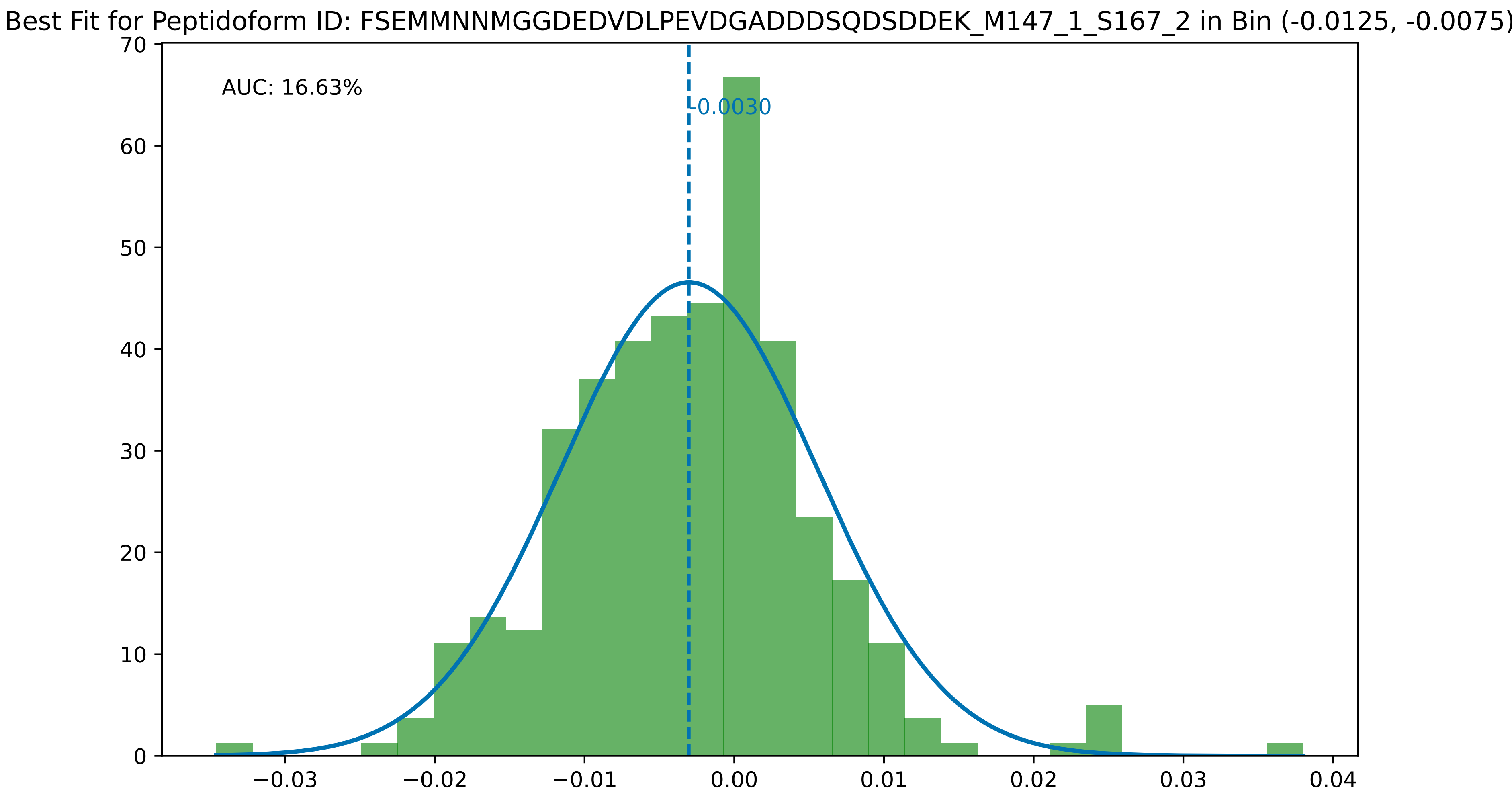
Best Fit for Peptidoform ID: FSEMMNNMGGDEDVDLPEVDGADDDSQDSDDEK_M147_1_S167_2 in Bin (-0.0125, -0.0075)
AUC: 16.63%
-0.0030
0
10
20
30
40
50
60
70
−0.03
−0.02
−0.01
0.00
0.01
0.02
0.03
0.04

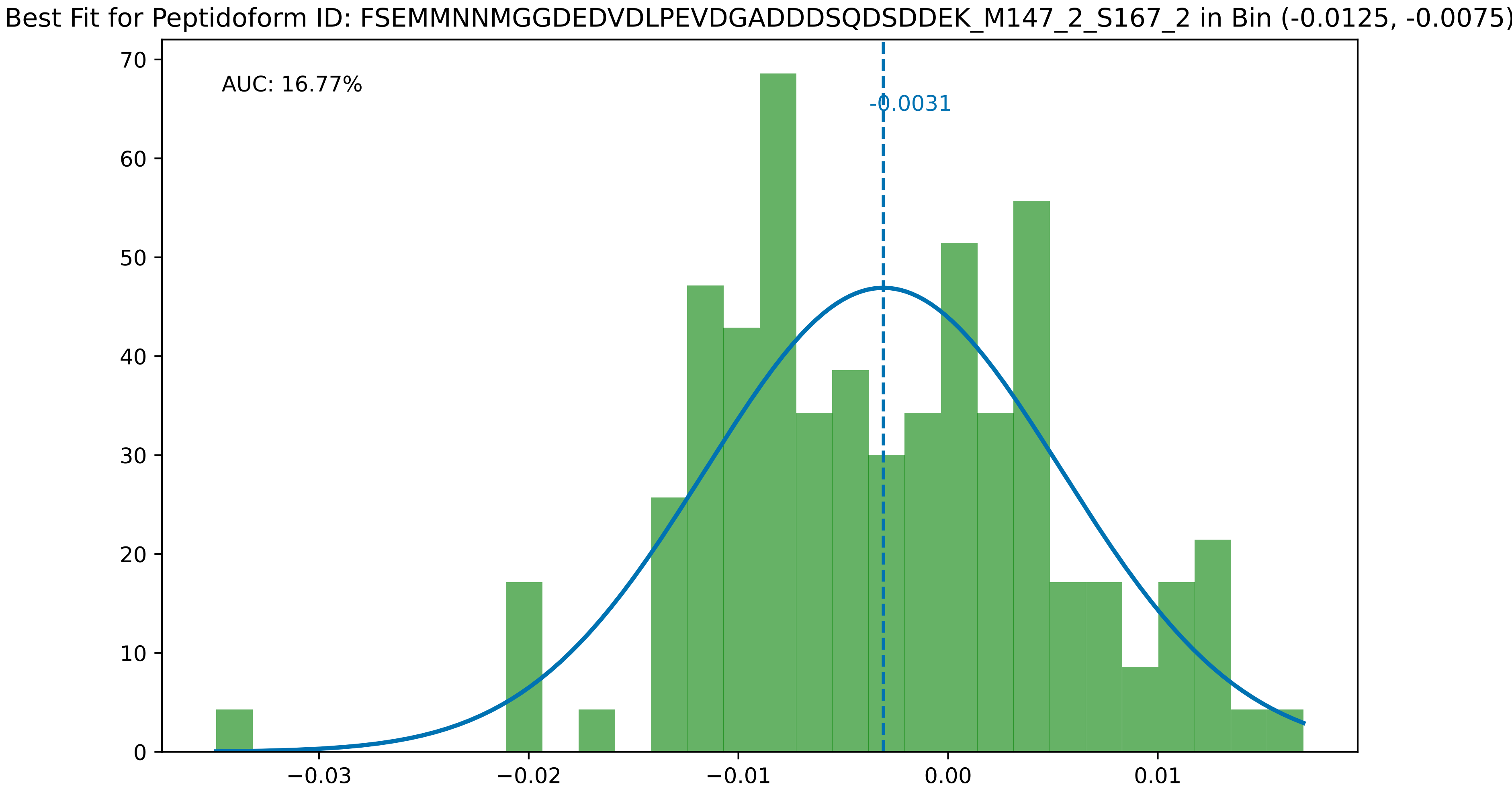
Best Fit for Peptidoform ID: FSEMMNNMGGDEDVDLPEVDGADDDSQDSDDEK_M147_2_S167_2 in Bin (-0.0125, -0.0075)
AUC: 16.77%
-0.0031
70
60
50
40
30
20
10
0
−0.03
−0.02
−0.01
0.00
0.01

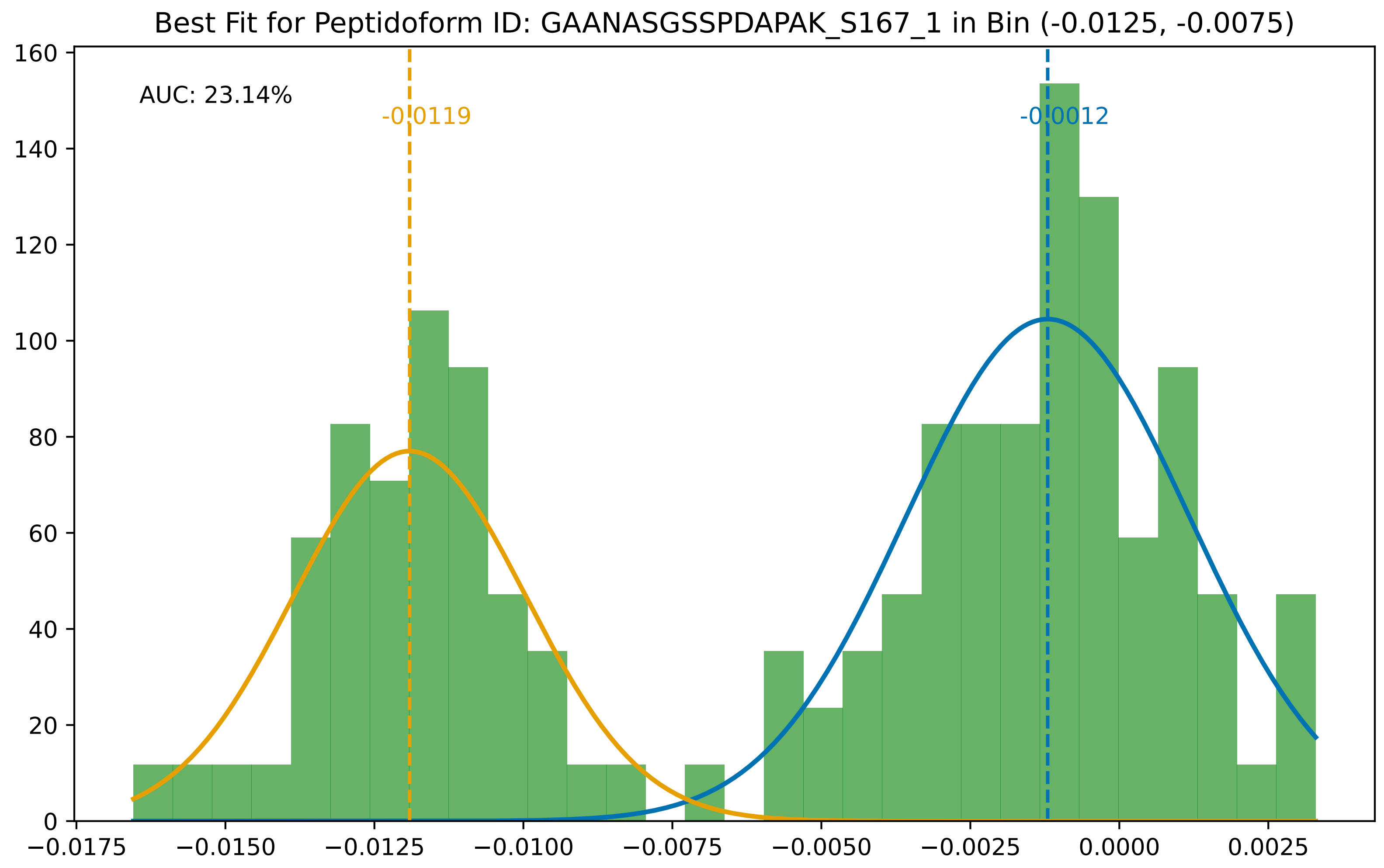

Best Fit for Peptidoform ID: GAANASGSSPDAPAK_S167_1 in Bin (-0.0125, -0.0075)
AUC: 23.14%
-0.0119
-0.0012
0
20
40
60
80
100
120
140
160
−0.0175
−0.0150
−0.0125
−0.0100
−0.0075
−0.0050
−0.0025
0.0000
0.0025

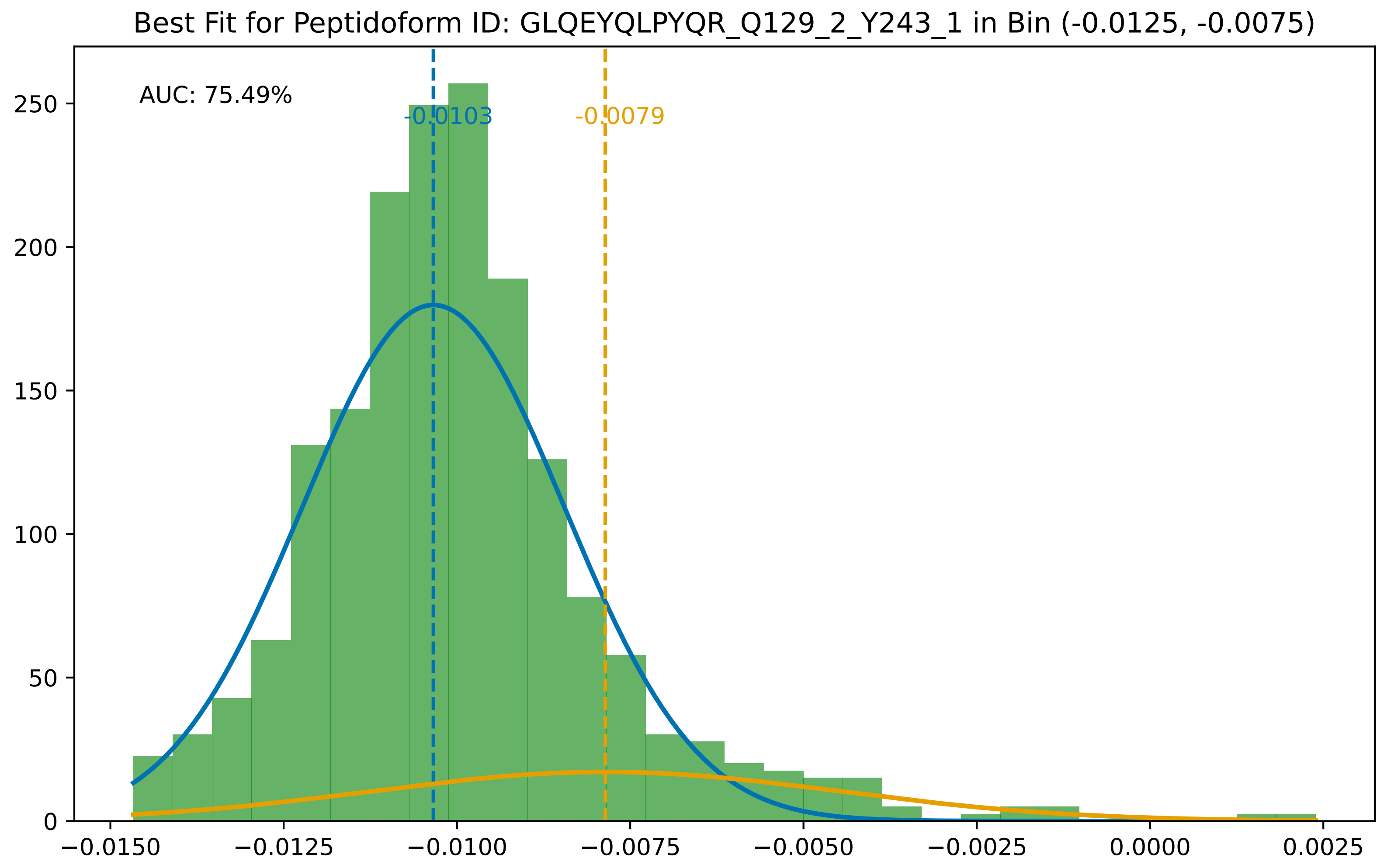
Best Fit for Peptidoform ID: GLQEYQLPYQR_Q129_2_Y243_1 in Bin (-0.0125, -0.0075)
AUC: 75.49%
-0.0103
-0.0079
0
50
100
150
200
250
−0.0150
−0.0125
−0.0100
−0.0075
−0.0050
−0.0025
0.0000
0.0025

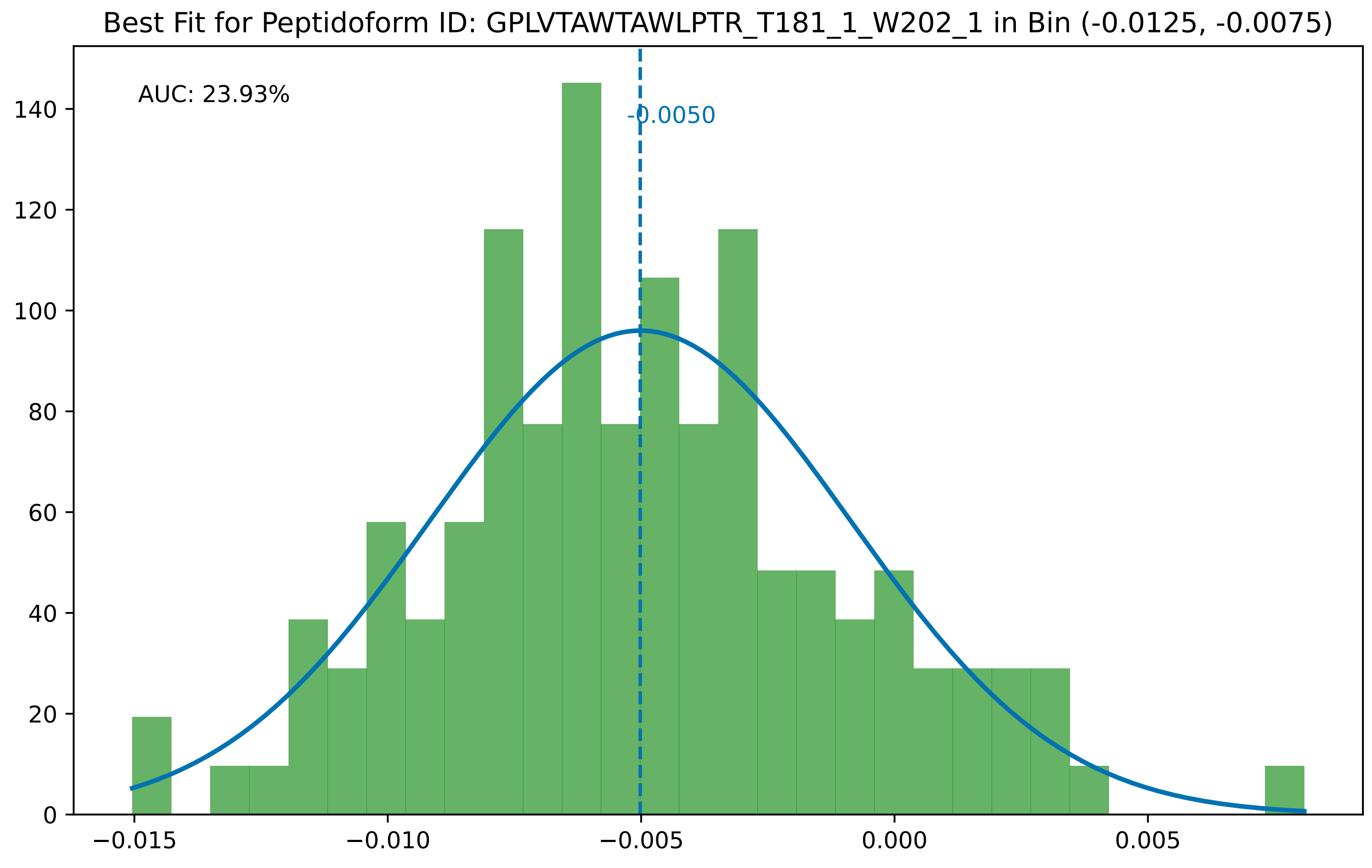

Best Fit for Peptidoform ID: GPLVTAWTAWLPTR_T181_1_W202_1 in Bin (-0.0125, -0.0075)
AUC: 23.93%
-0.0050
0
20
40
60
80
100
120
140
−0.015
−0.010
−0.005
0.000
0.005

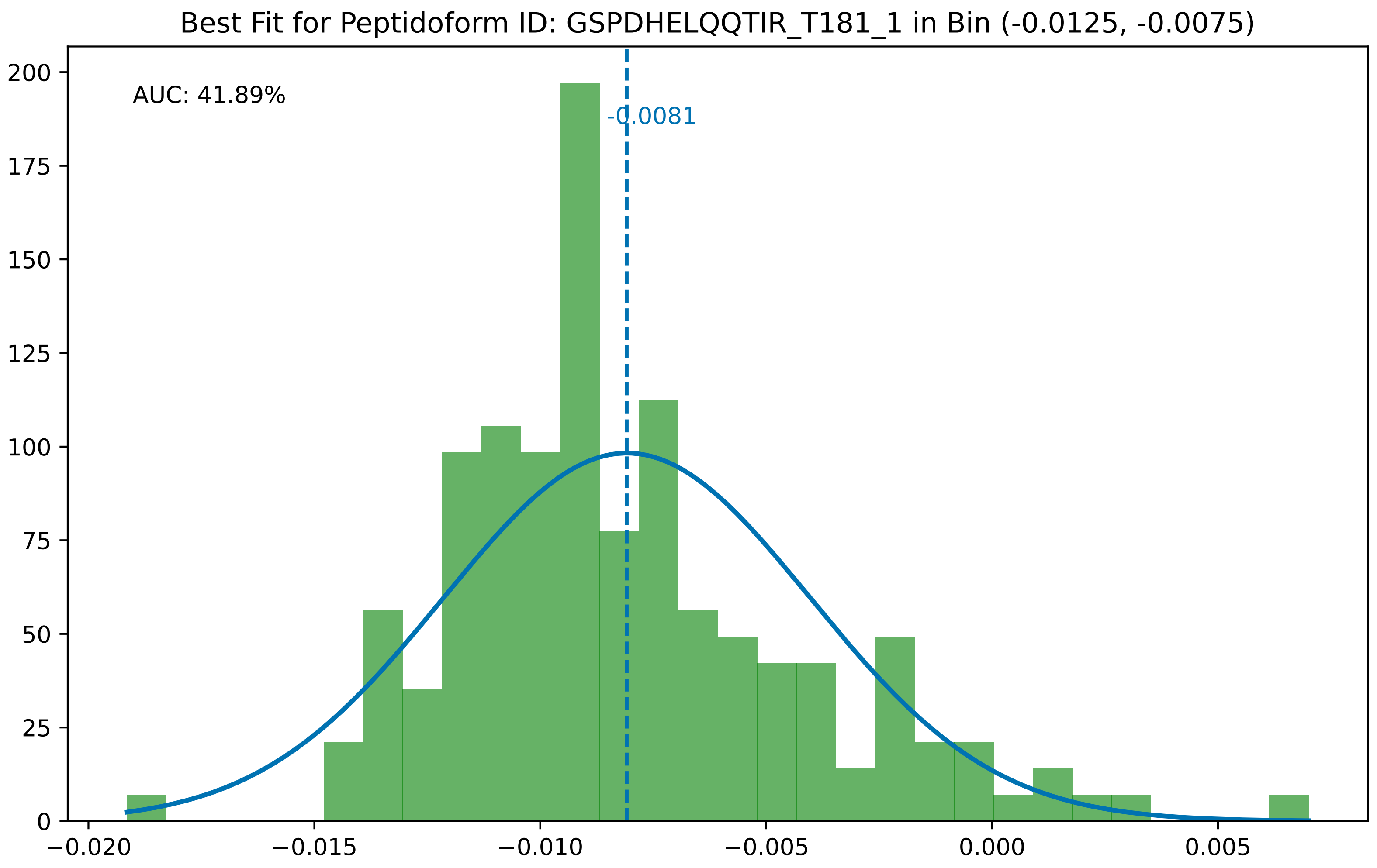
Best Fit for Peptidoform ID: GSPDHELQQTIR_T181_1 in Bin (-0.0125, -0.0075)
AUC: 41.89%
-0.0081
200
175
150
125
100
75
50
25
0
−0.020
−0.015
−0.010
−0.005
0.000
0.005

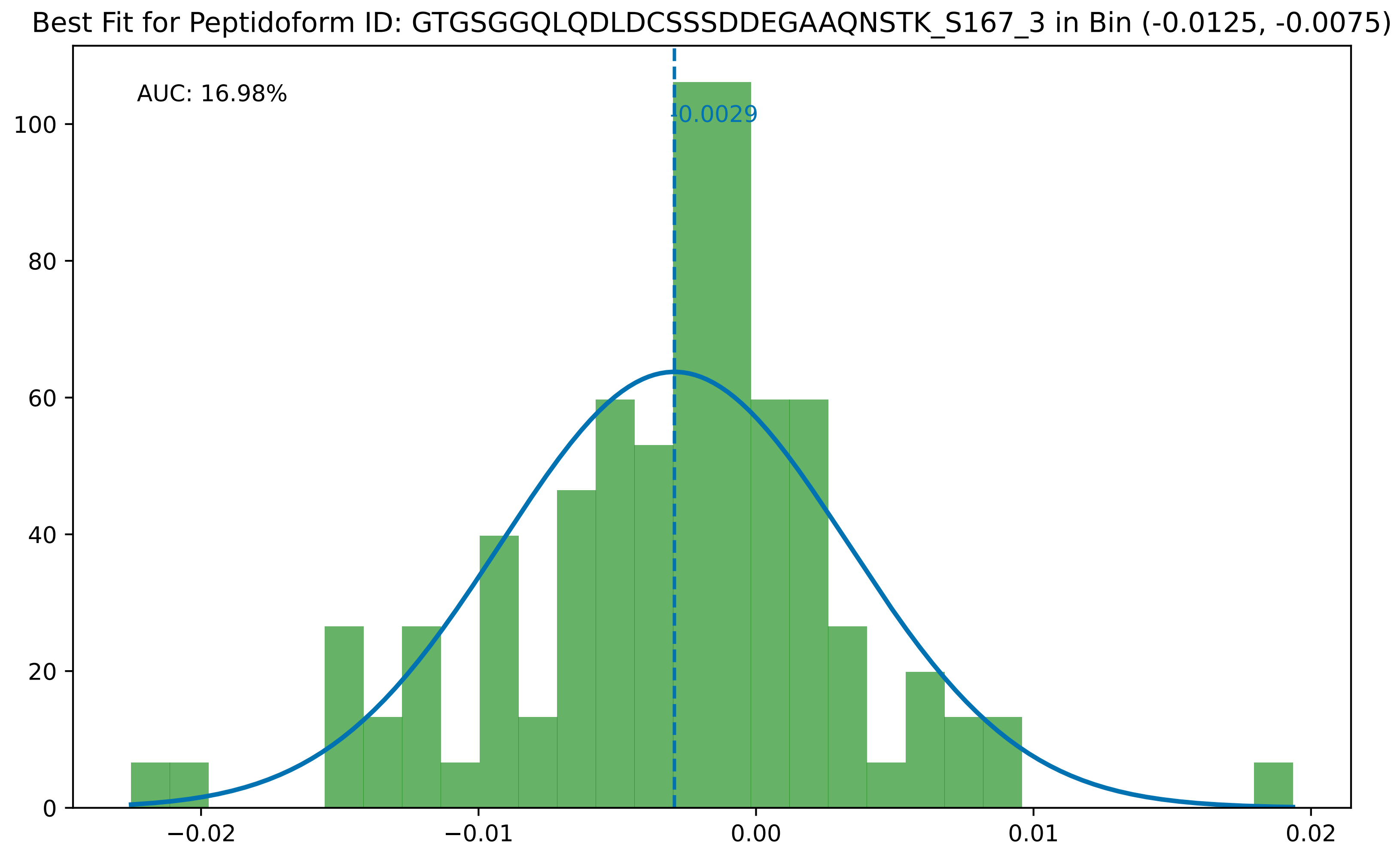
Best Fit for Peptidoform ID: GTGSGGQLQDLDCSSSDDEGAAQNSTK_S167_3 in Bin (-0.0125, -0.0075)
AUC: 16.98%
-0.0029
0
20
40
60
80
100
−0.02
−0.01
0.00
0.01
0.02

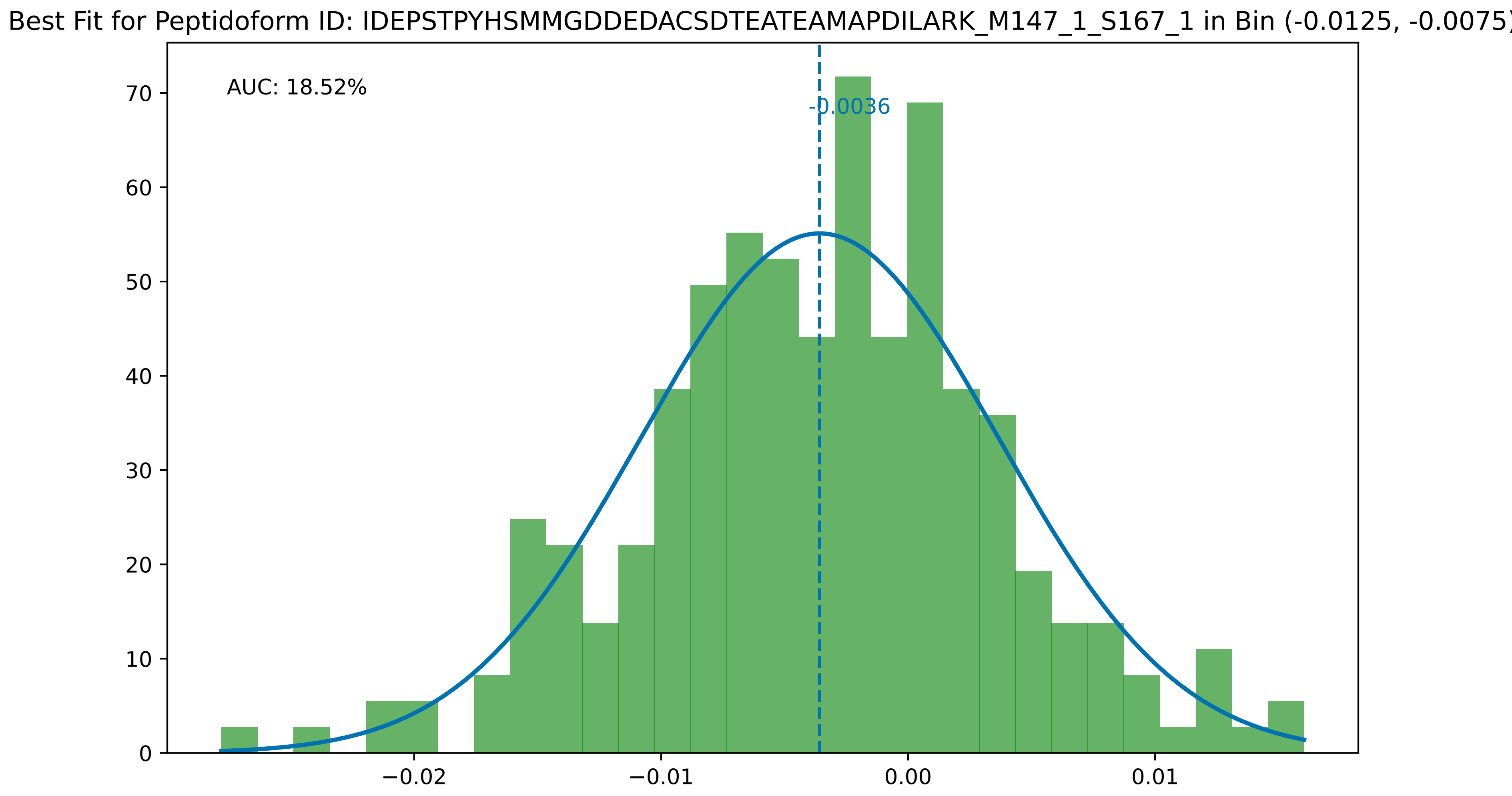

Best Fit for Peptidoform ID: IDEPSTPYHSMMGDDEDACSDTEATEAMAPDILARK_M147_1_S167_1 in Bin (-0.0125, -0.0075)
AUC: 18.52%
-0.0036
0
10
20
30
40
50
60
70
−0.02
−0.01
0.00
0.01

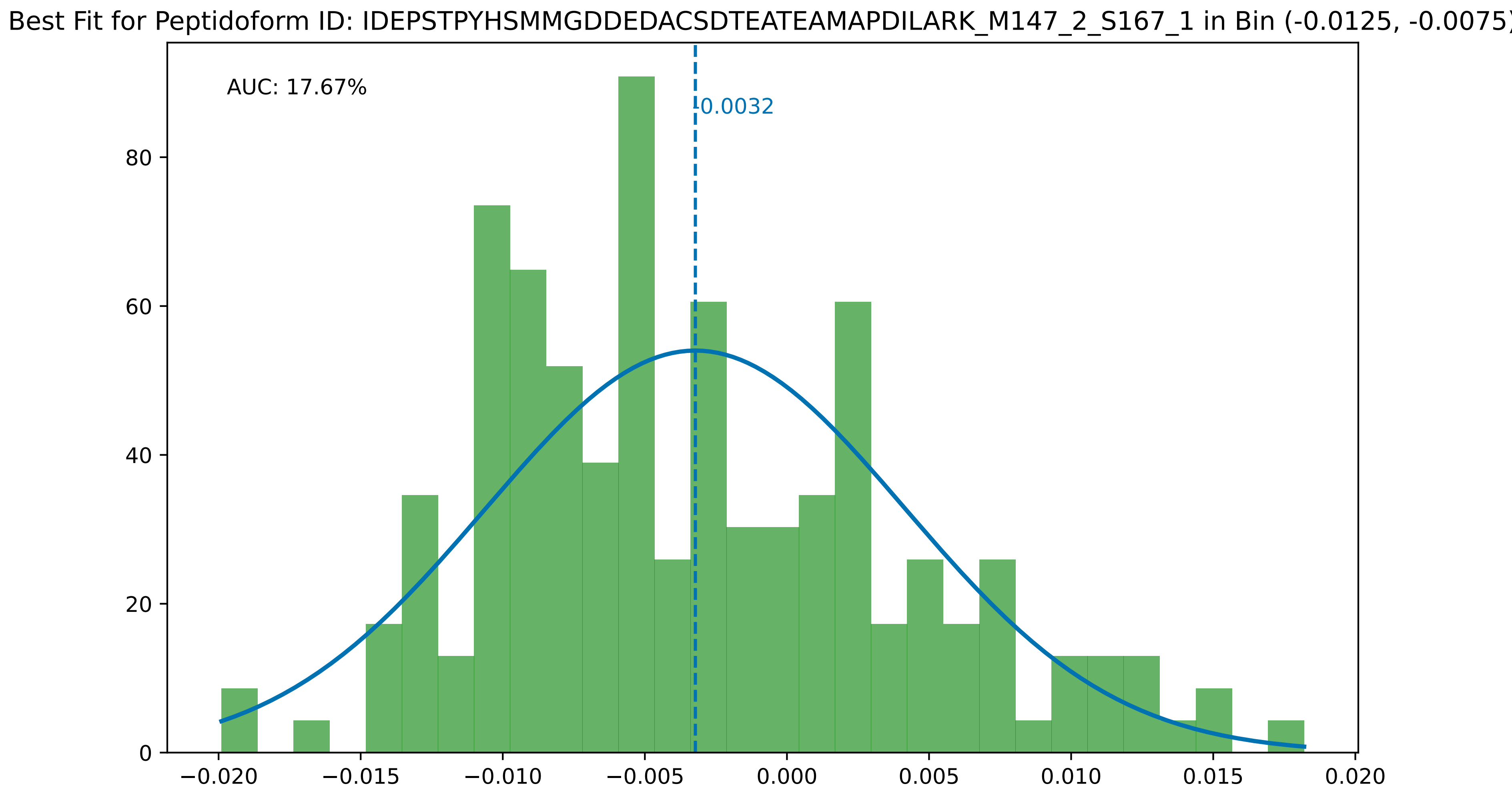

Best Fit for Peptidoform ID: IDEPSTPYHSMMGDDEDACSDTEATEAMAPDILARK_M147_2_S167_1 in Bin (-0.0125, -0.0075)
AUC: 17.67%
-0.0032
0
20
40
60
80
−0.020
−0.015
−0.010
−0.005
0.000
0.005
0.010
0.015
0.020

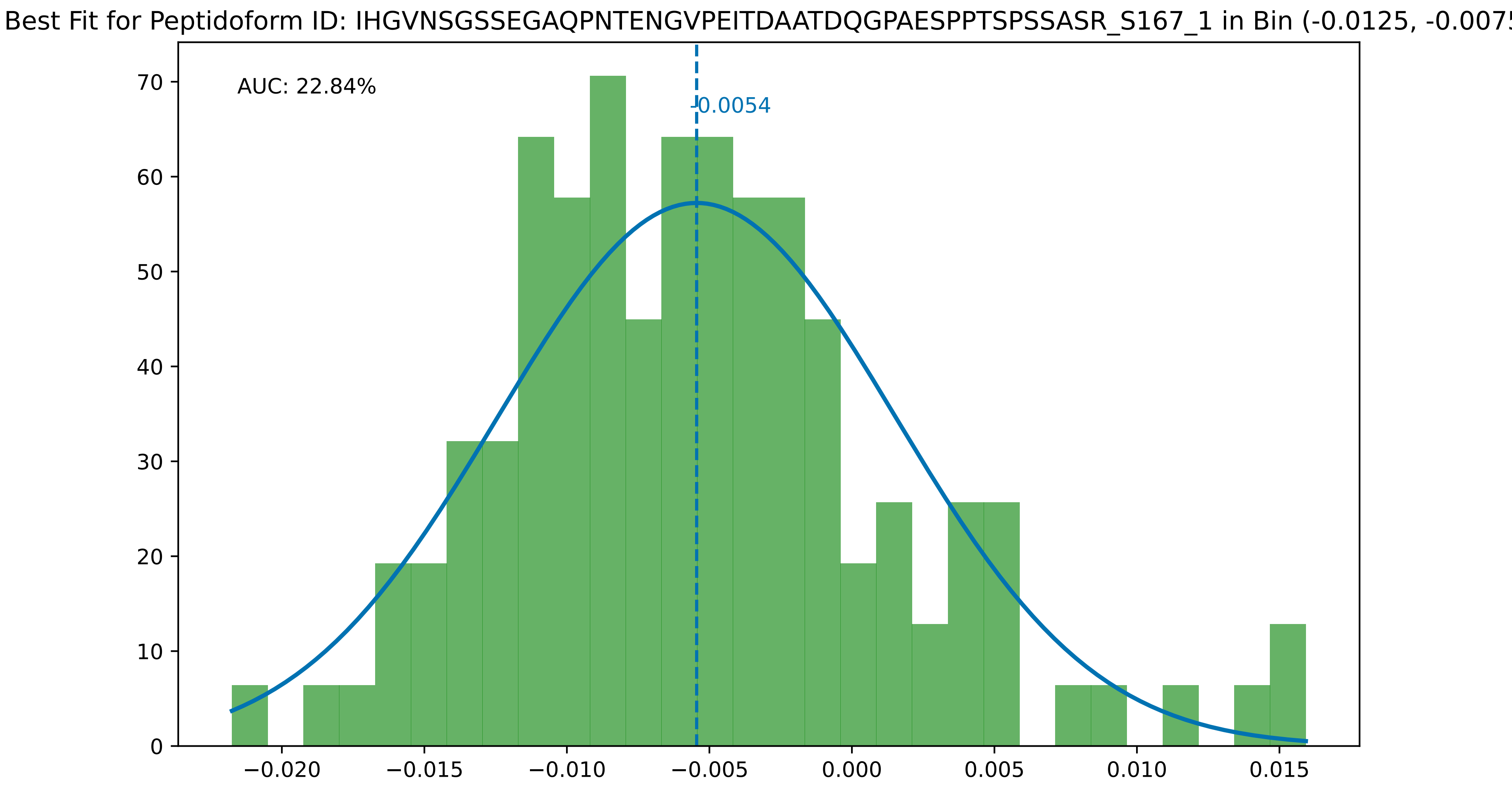

Best Fit for Peptidoform ID: IHGVNSGSSEGAQPNTENGVPEITDAATDQGPAESPPTSPSSASR_S167_1 in Bin (-0.0125, -0.0075
AUC: 22.84%
-0.0054
70
60
50
40
30
20
10
0
−0.020
−0.015
−0.010
−0.005
0.000
0.005
0.010
0.015

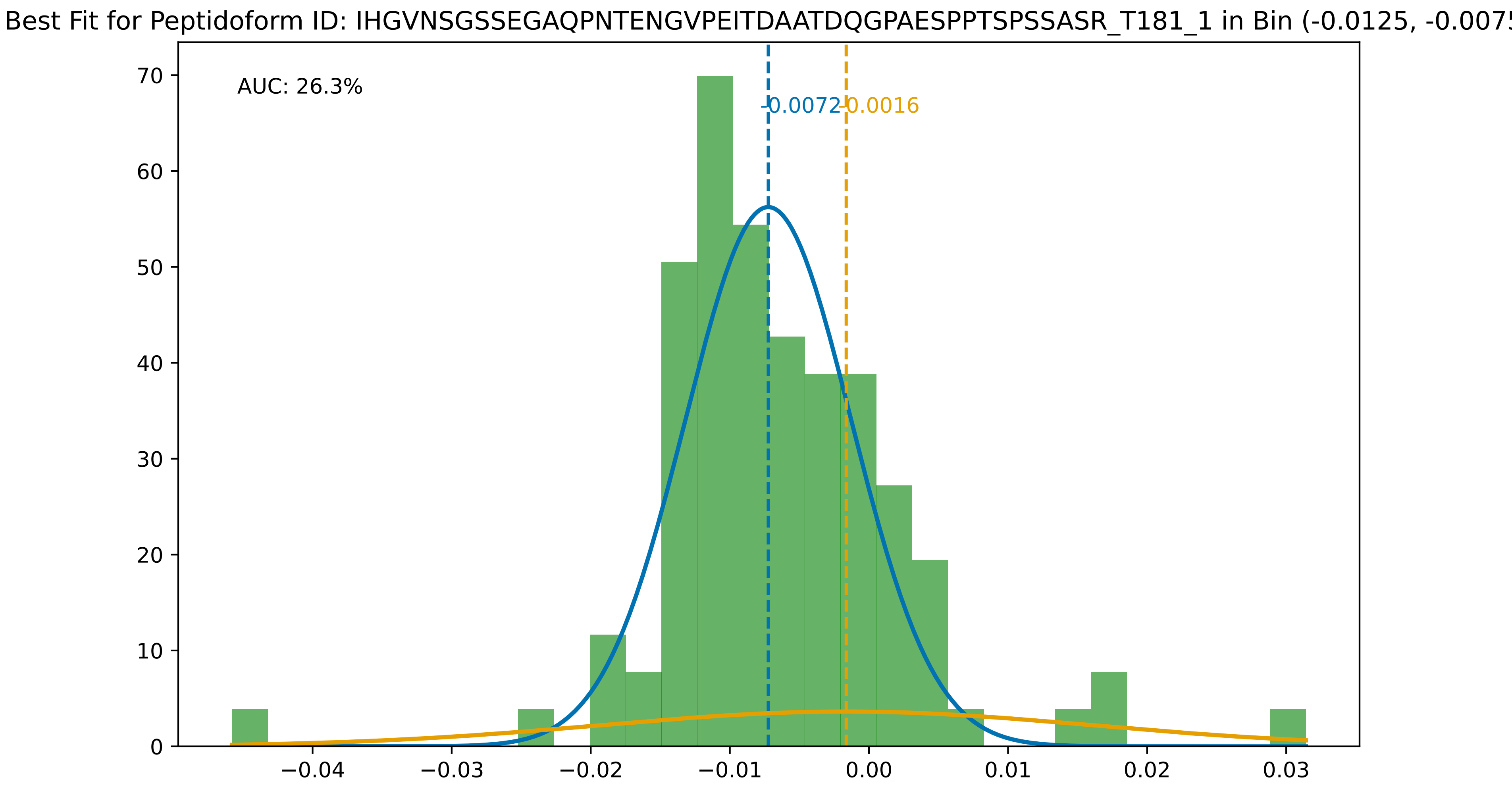

Best Fit for Peptidoform ID: IHGVNSGSSEGAQPNTENGVPEITDAATDQGPAESPPTSPSSASR_T181_1 in Bin (-0.0125, -0.0075
AUC: 26.3%
-0.0072
-0.0016
0
10
20
30
40
50
60
70
−0.04
−0.03
−0.02
−0.01
0.00
0.01
0.02
0.03

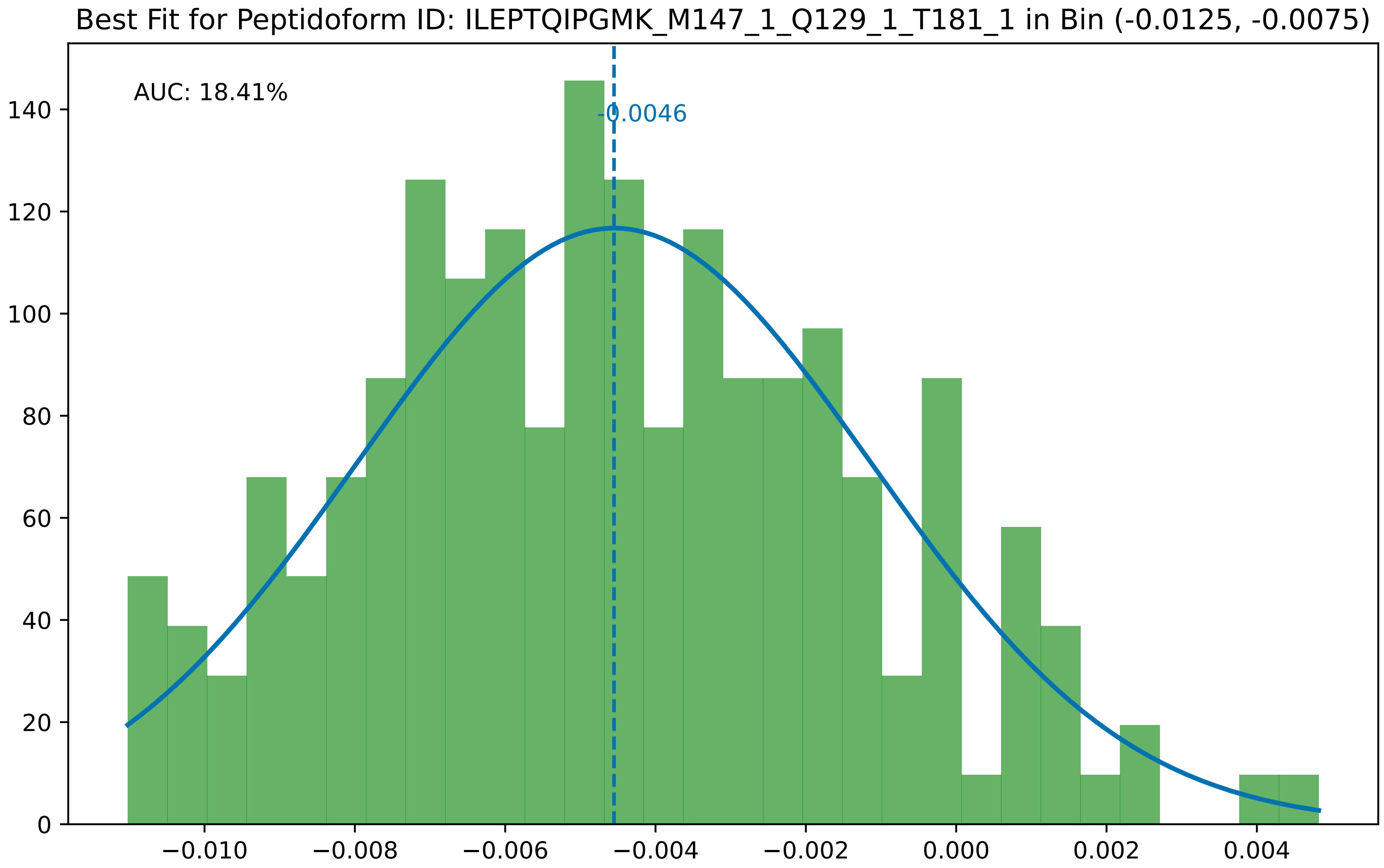

Best Fit for Peptidoform ID: ILEPTQIPGMK_M147_1_Q129_1_T181_1 in Bin (-0.0125, -0.0075)
AUC: 18.41%
-0.0046
0
20
40
60
80
100
120
140
−0.010
−0.008
−0.006
−0.004
−0.002
0.000
0.002
0.004

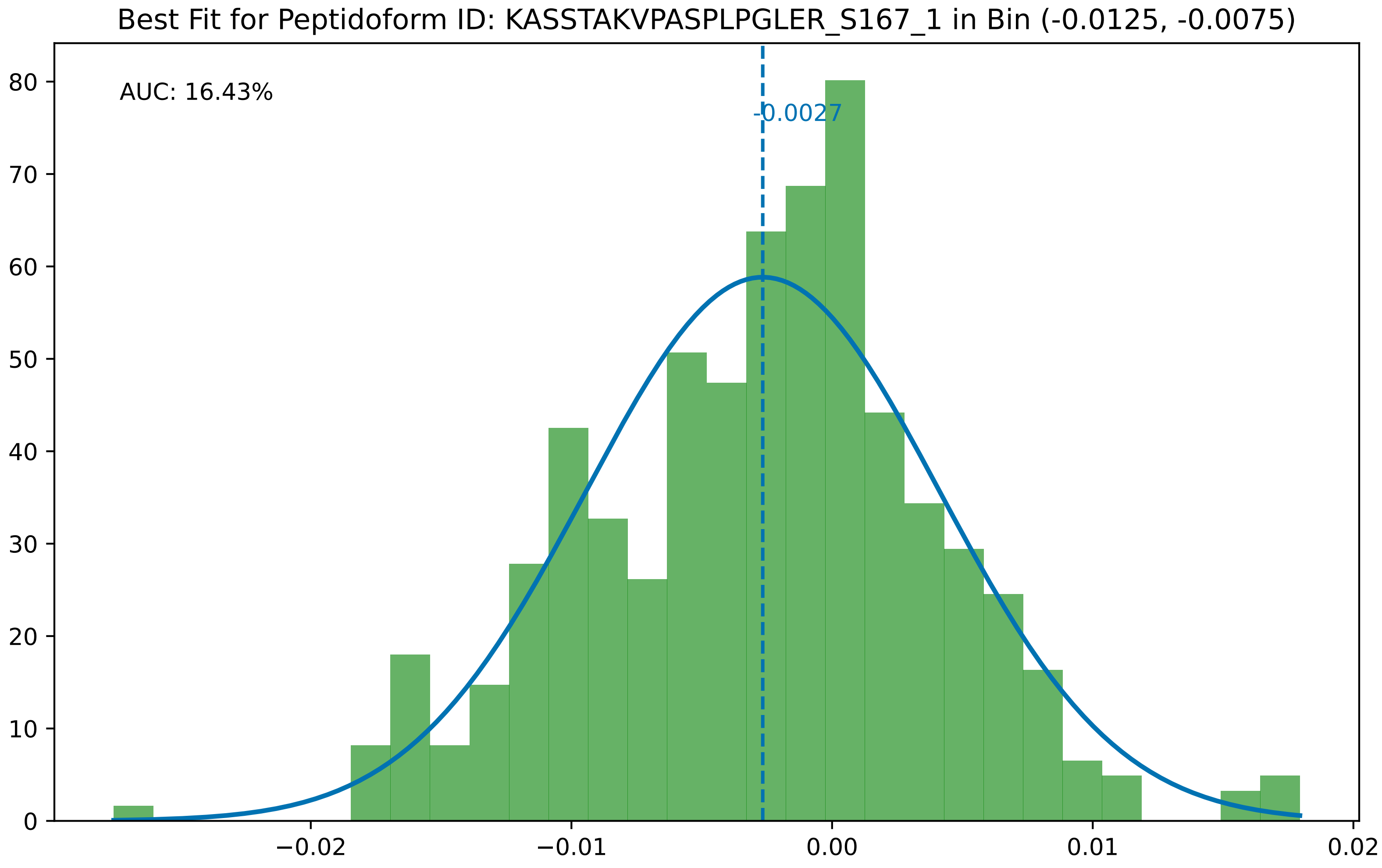
Best Fit for Peptidoform ID: KASSTAKVPASPLPGLER_S167_1 in Bin (-0.0125, -0.0075)
AUC: 16.43%
-0.0027
0
10
20
30
40
50
60
70
80
−0.02
−0.01
0.00
0.01
0.02

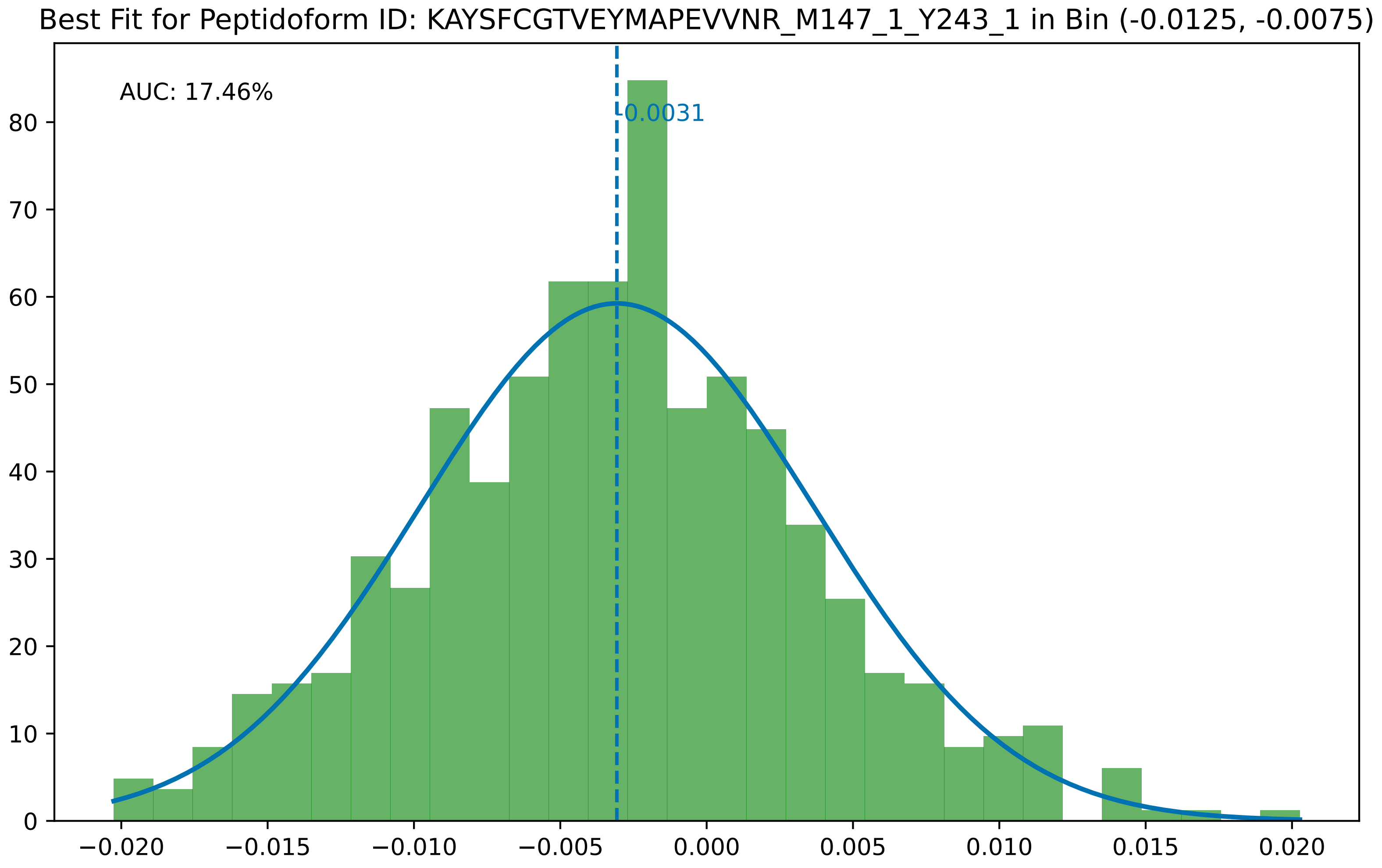
Best Fit for Peptidoform ID: KAYSFCGTVEYMAPEVVNR_M147_1_Y243_1 in Bin (-0.0125, -0.0075)
AUC: 17.46%
-0.0031
0
10
20
30
40
50
60
70
80
−0.020
−0.015
−0.010
−0.005
0.000
0.005
0.010
0.015
0.020

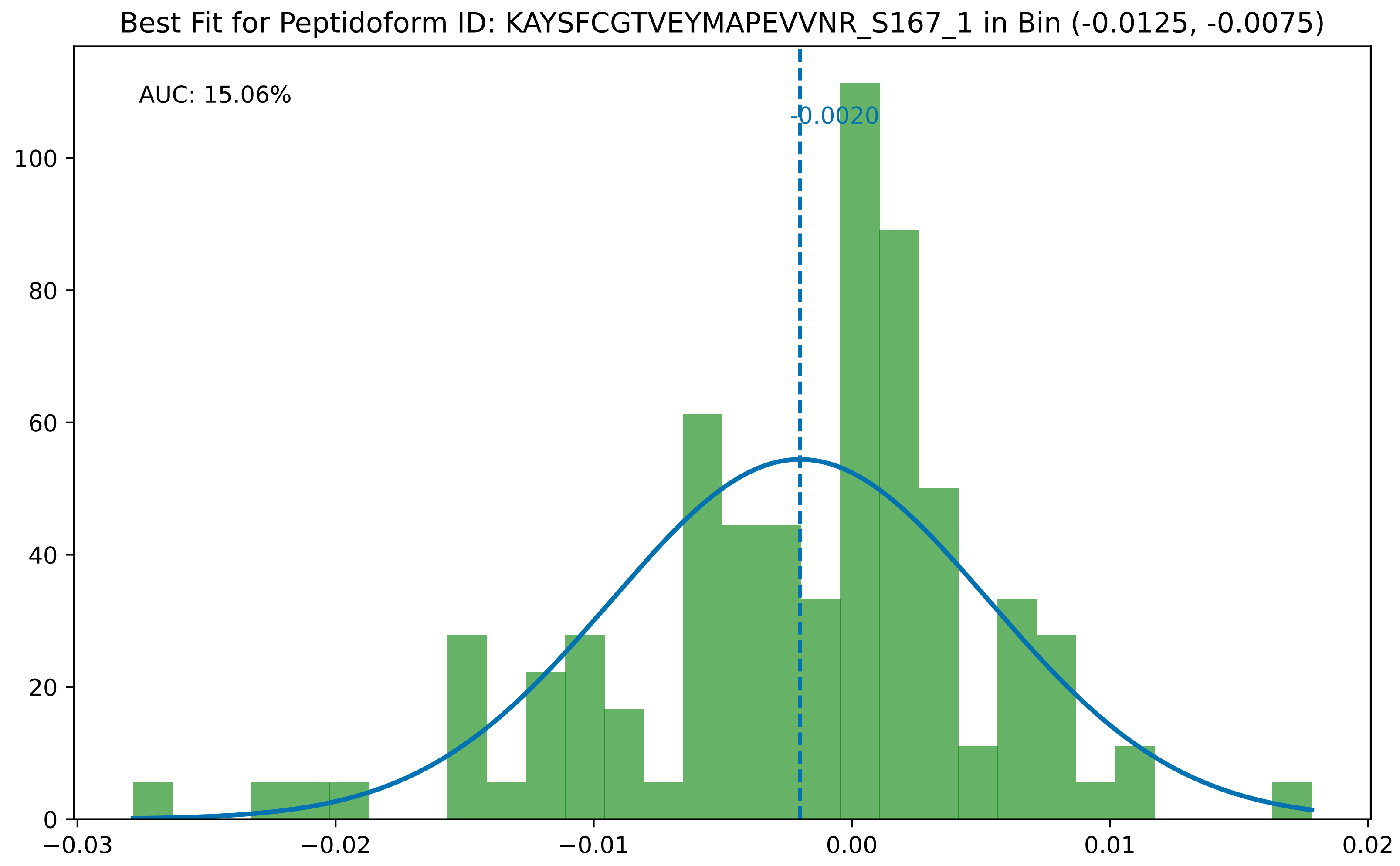
Best Fit for Peptidoform ID: KAYSFCGTVEYMAPEVVNR_S167_1 in Bin (-0.0125, -0.0075)
AUC: 15.06%
-0.0020
0
20
40
60
80
100
−0.03
−0.02
−0.01
0.00
0.01
0.02

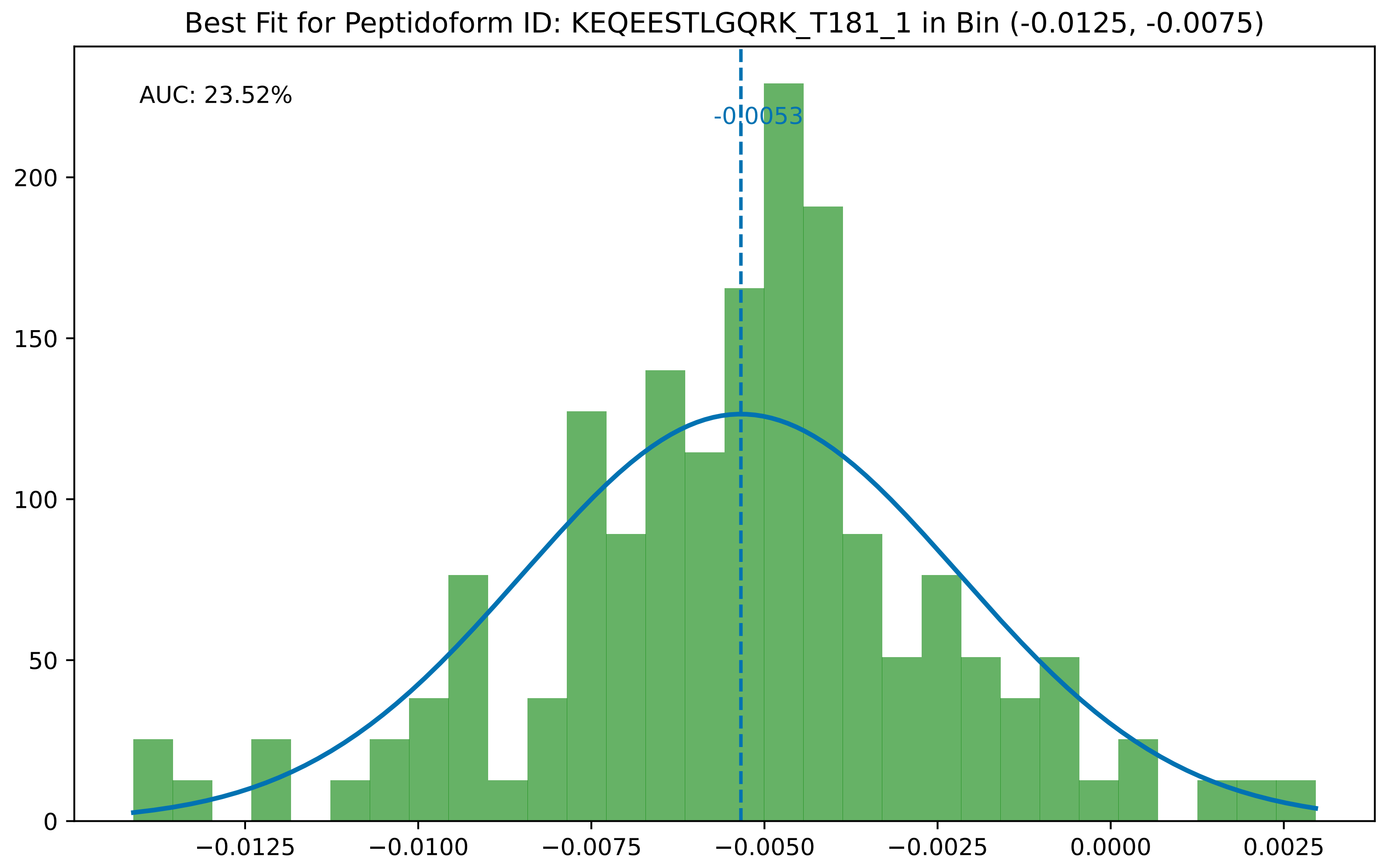
Best Fit for Peptidoform ID: KEQEESTLGQRK_T181_1 in Bin (-0.0125, -0.0075)
AUC: 23.52%
-0.0053
0
50
100
150
200
−0.0125
−0.0100
−0.0075
−0.0050
−0.0025
0.0000
0.0025

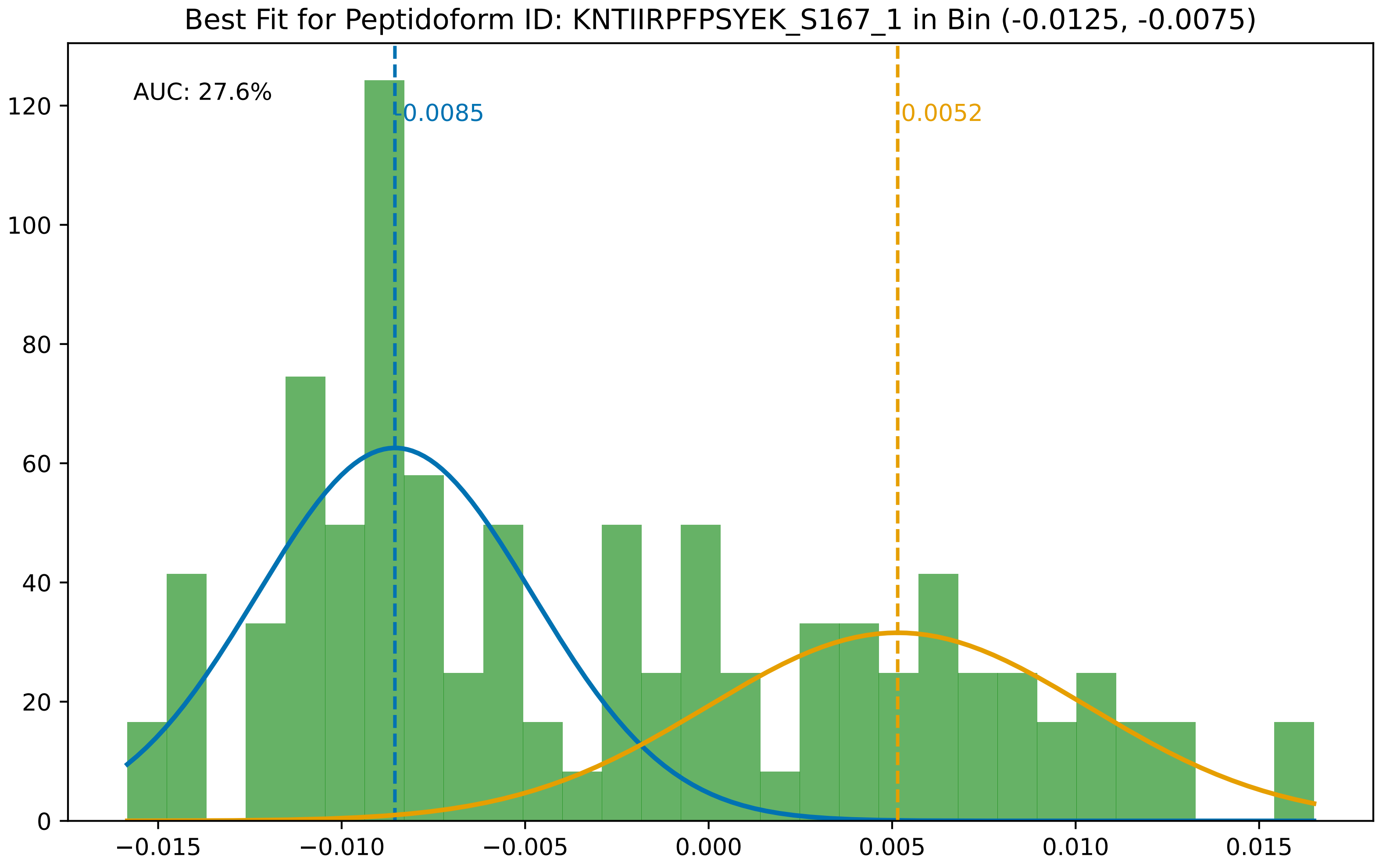

Best Fit for Peptidoform ID: KNTIIRPFPSYEK_S167_1 in Bin (-0.0125, -0.0075)
AUC: 27.6%
-0.0085
0.0052
0
20
40
60
80
100
120
−0.015
−0.010
−0.005
0.000
0.005
0.010
0.015

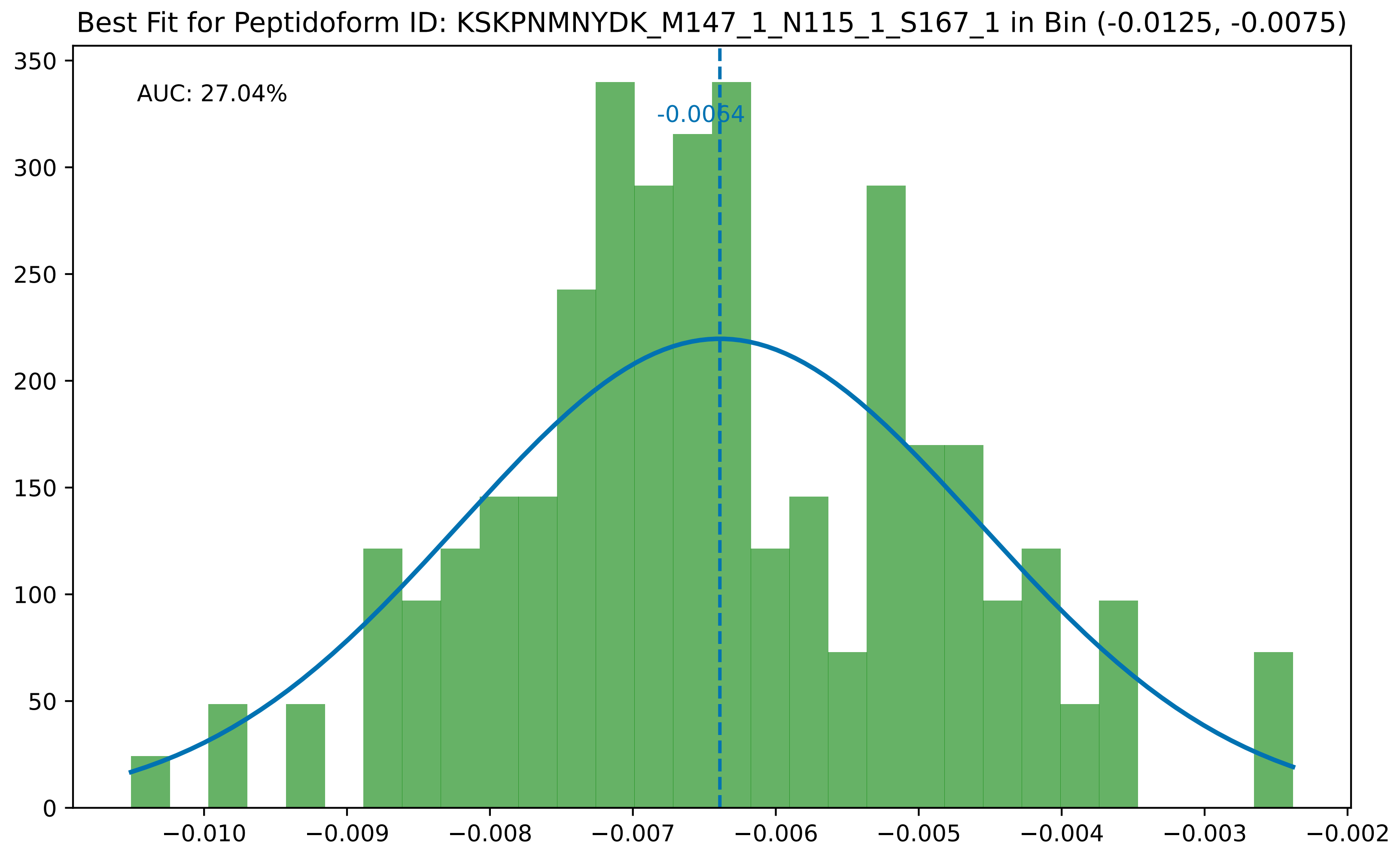
Best Fit for Peptidoform ID: KSKPNMNYDK_M147_1_N115_1_S167_1 in Bin (-0.0125, -0.0075)
AUC: 27.04%
-0.0064
0
50
100
150
200
250
300
350
−0.010
−0.009
−0.008
−0.007
−0.006
−0.005
−0.004
−0.003
−0.002

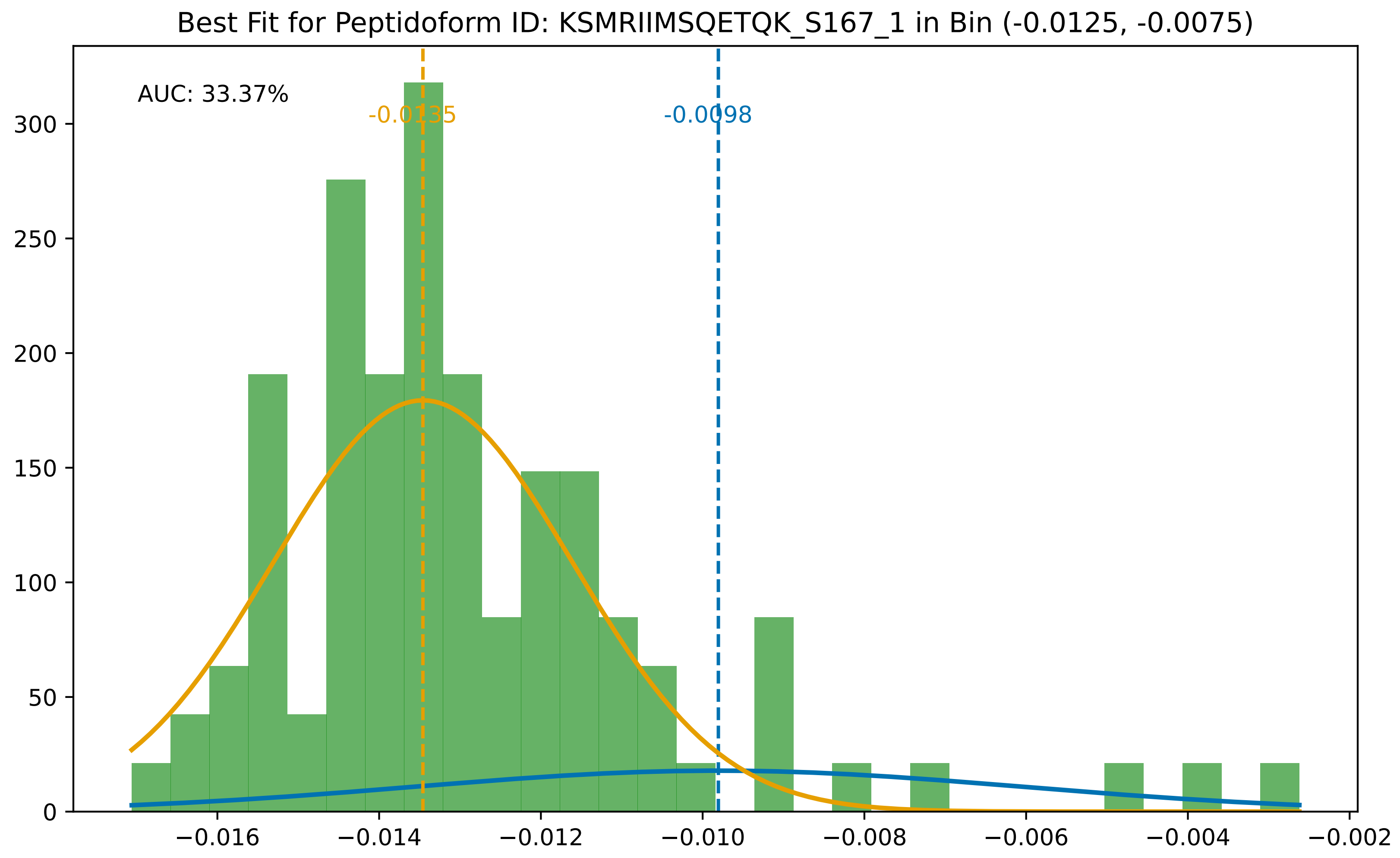

Best Fit for Peptidoform ID: KSMRIIMSQETQK_S167_1 in Bin (-0.0125, -0.0075)
AUC: 33.37%
-0.0135
-0.0098
0
50
100
150
200
250
300
−0.016
−0.014
−0.012
−0.010
−0.008
−0.006
−0.004
−0.002

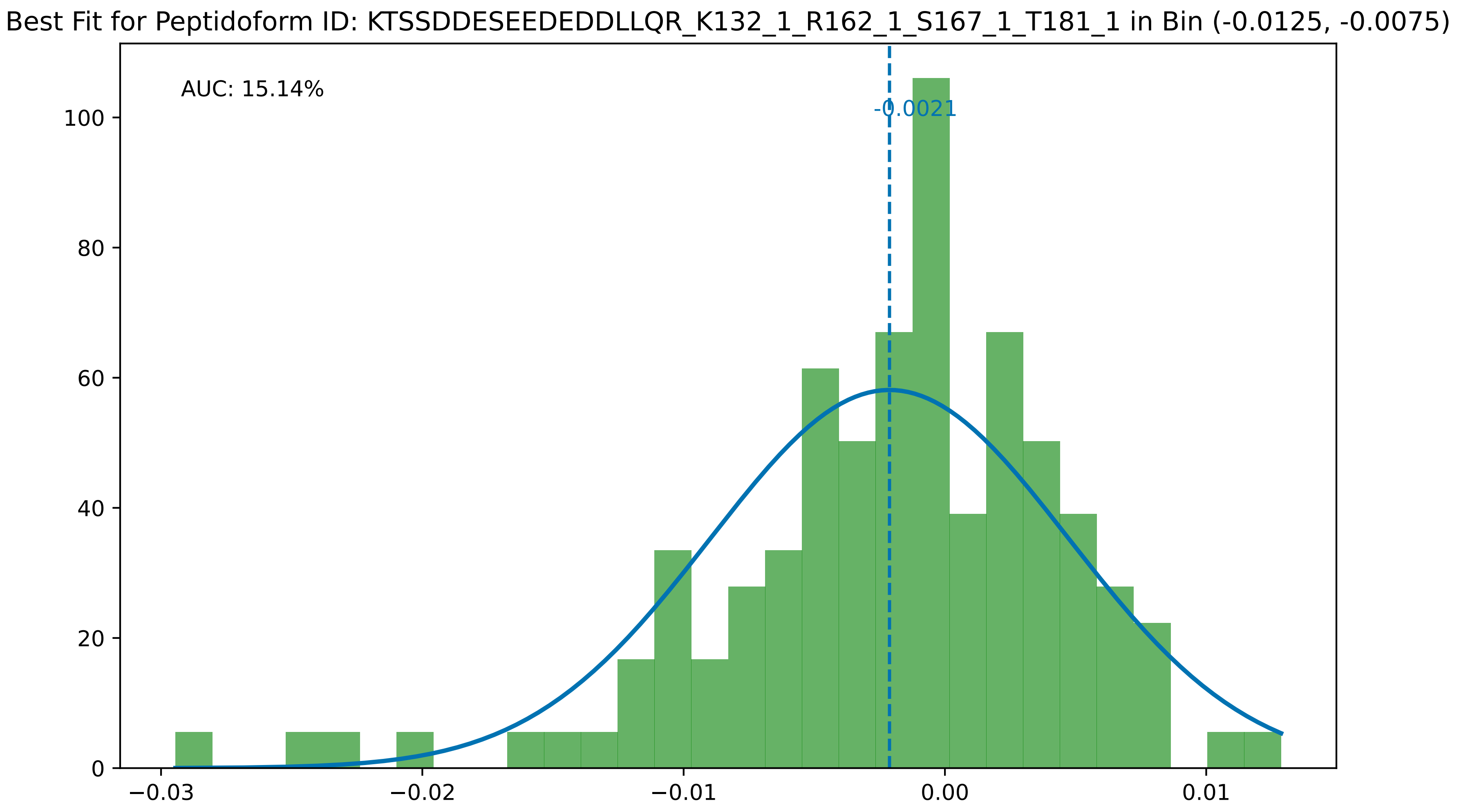

Best Fit for Peptidoform ID: KTSSDDESEEDEDDLLQR_K132_1_R162_1_S167_1_T181_1 in Bin (-0.0125, -0.0075)
AUC: 15.14%
-0.0021
0
20
40
60
80
100
−0.03
−0.02
−0.01
0.00
0.01

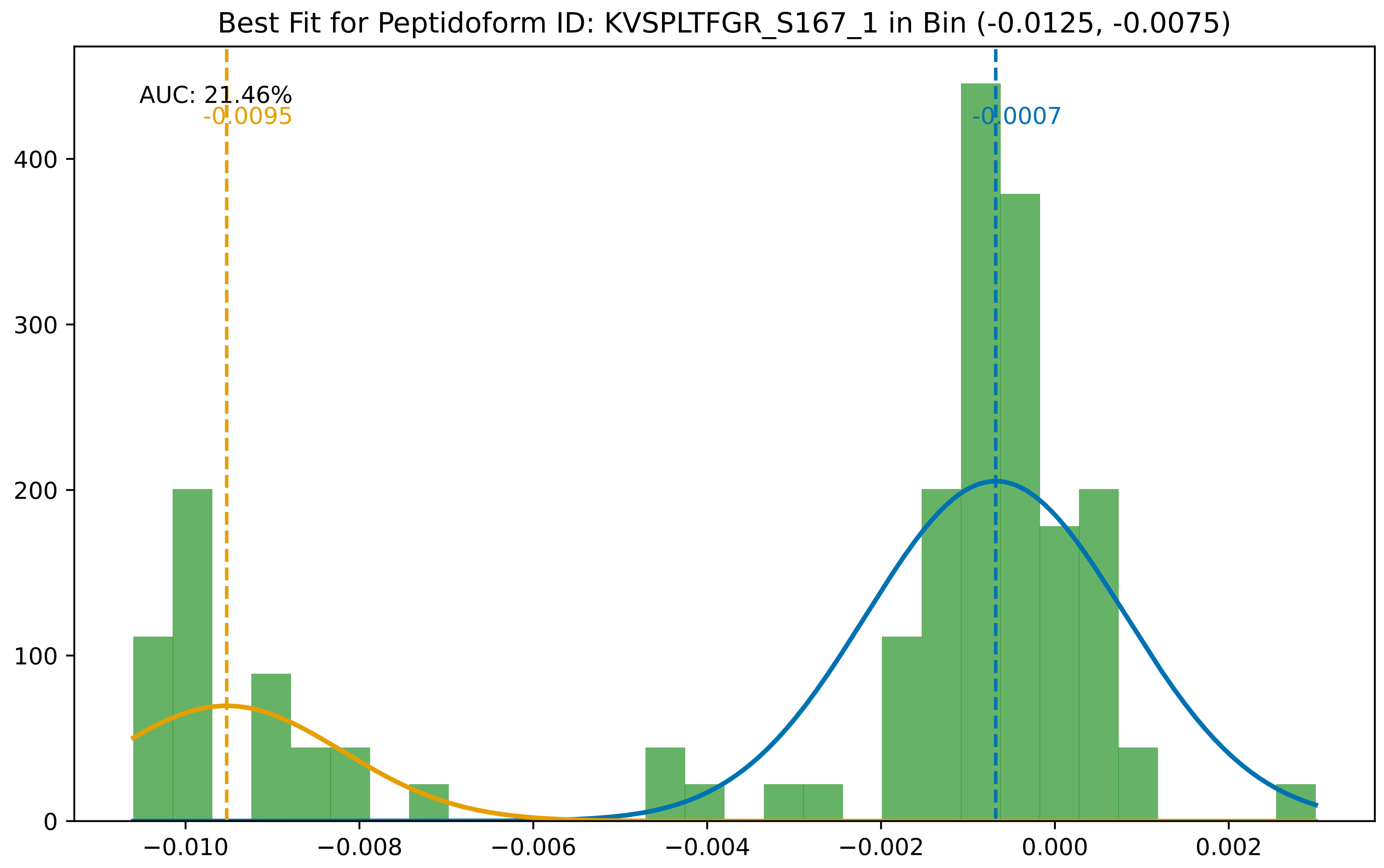

Best Fit for Peptidoform ID: KVSPLTFGR_S167_1 in Bin (-0.0125, -0.0075)
AUC: 21.46%
-0.0095
-0.0007
400
300
200
100
0
−0.010
−0.008
−0.006
−0.004
−0.002
0.000
0.002

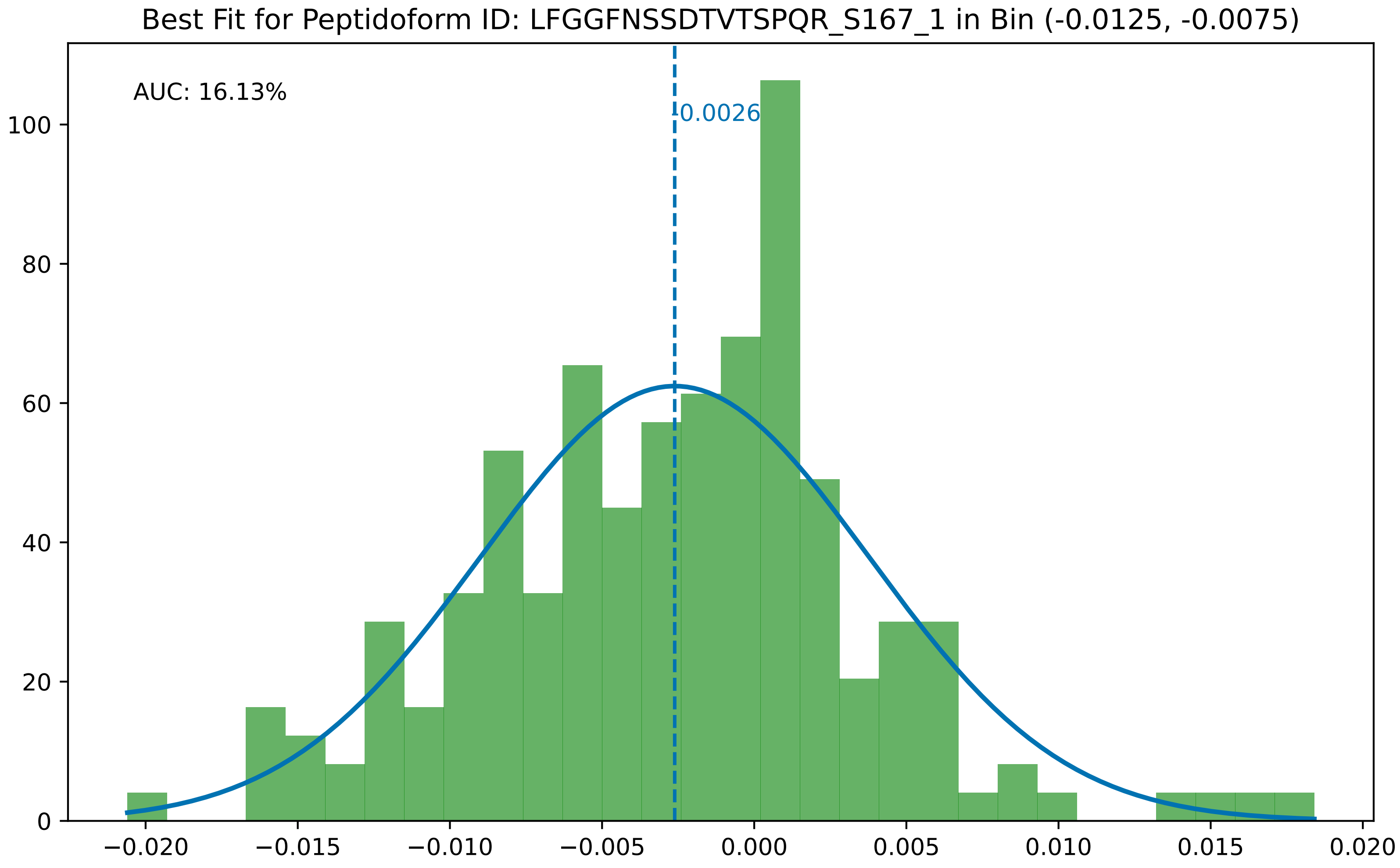
Best Fit for Peptidoform ID: LFGGFNSSDTVTSPQR_S167_1 in Bin (-0.0125, -0.0075)
AUC: 16.13%
-0.0026
0
20
40
60
80
100
−0.020
−0.015
−0.010
−0.005
0.000
0.005
0.010
0.015
0.020

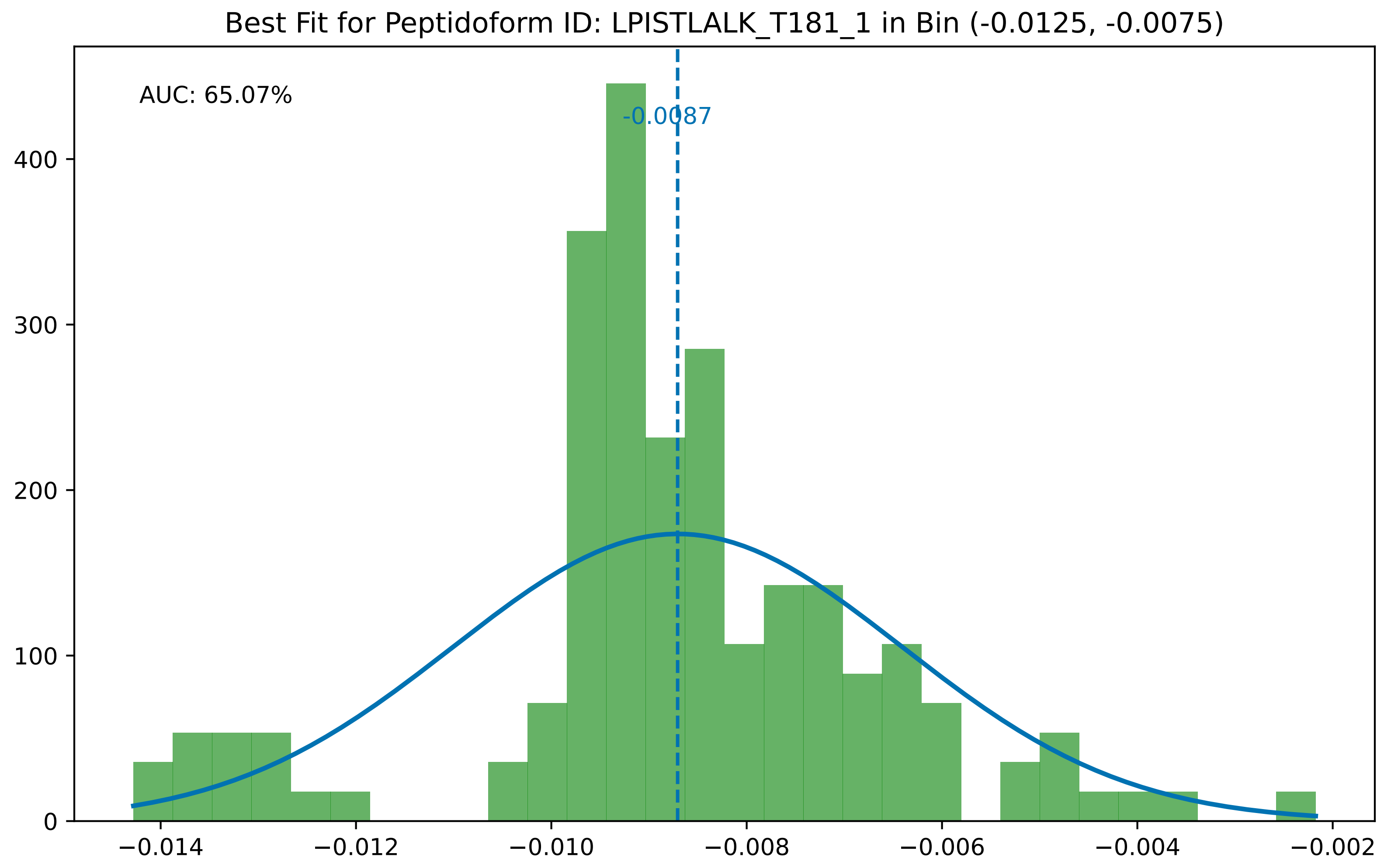

Best Fit for Peptidoform ID: LPISTLALK_T181_1 in Bin (-0.0125, -0.0075)
AUC: 65.07%
-0.0087
0
100
200
300
400
−0.014
−0.012
−0.010
−0.008
−0.006
−0.004
−0.002

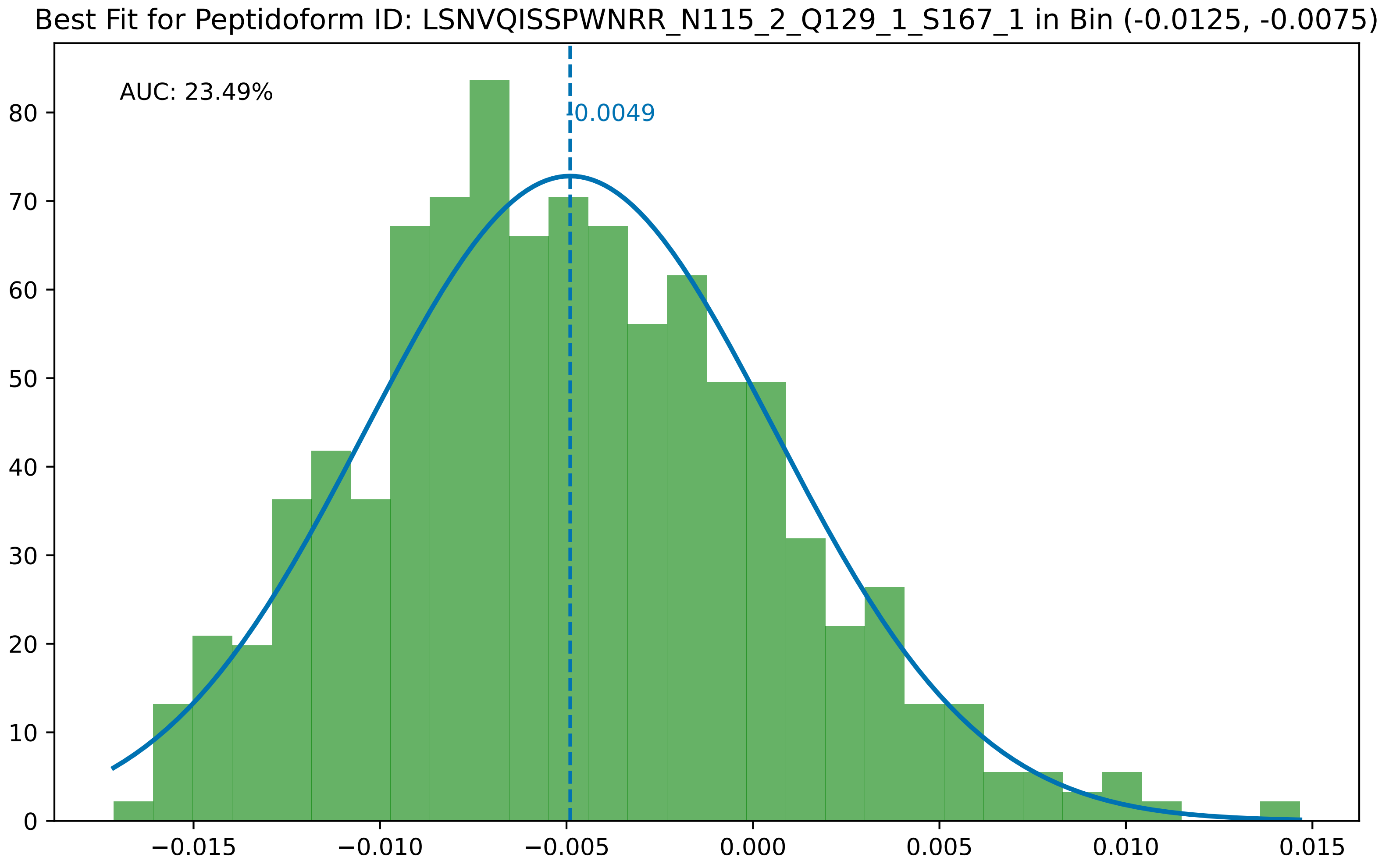

Best Fit for Peptidoform ID: LSNVQISSPWNRR_N115_2_Q129_1_S167_1 in Bin (-0.0125, -0.0075)
AUC: 23.49%
-0.0049
0
10
20
30
40
50
60
70
80
−0.015
−0.010
−0.005
0.000
0.005
0.010
0.015

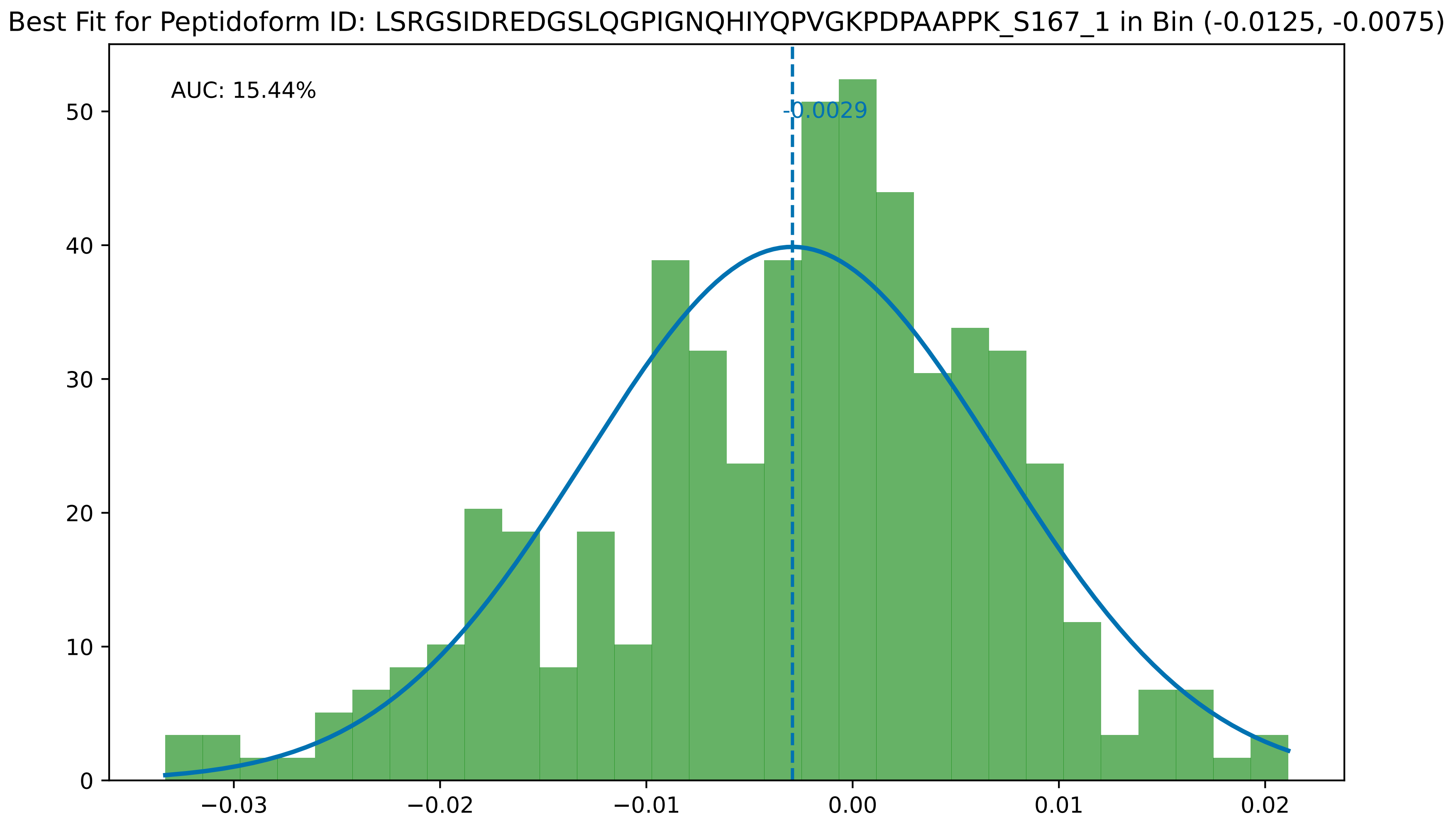

Best Fit for Peptidoform ID: LSRGSIDREDGSLQGPIGNQHIYQPVGKPDPAAPPK_S167_1 in Bin (-0.0125, -0.0075)
AUC: 15.44%
-0.0029
0
10
20
30
40
50
−0.03
−0.02
−0.01
0.00
0.01
0.02

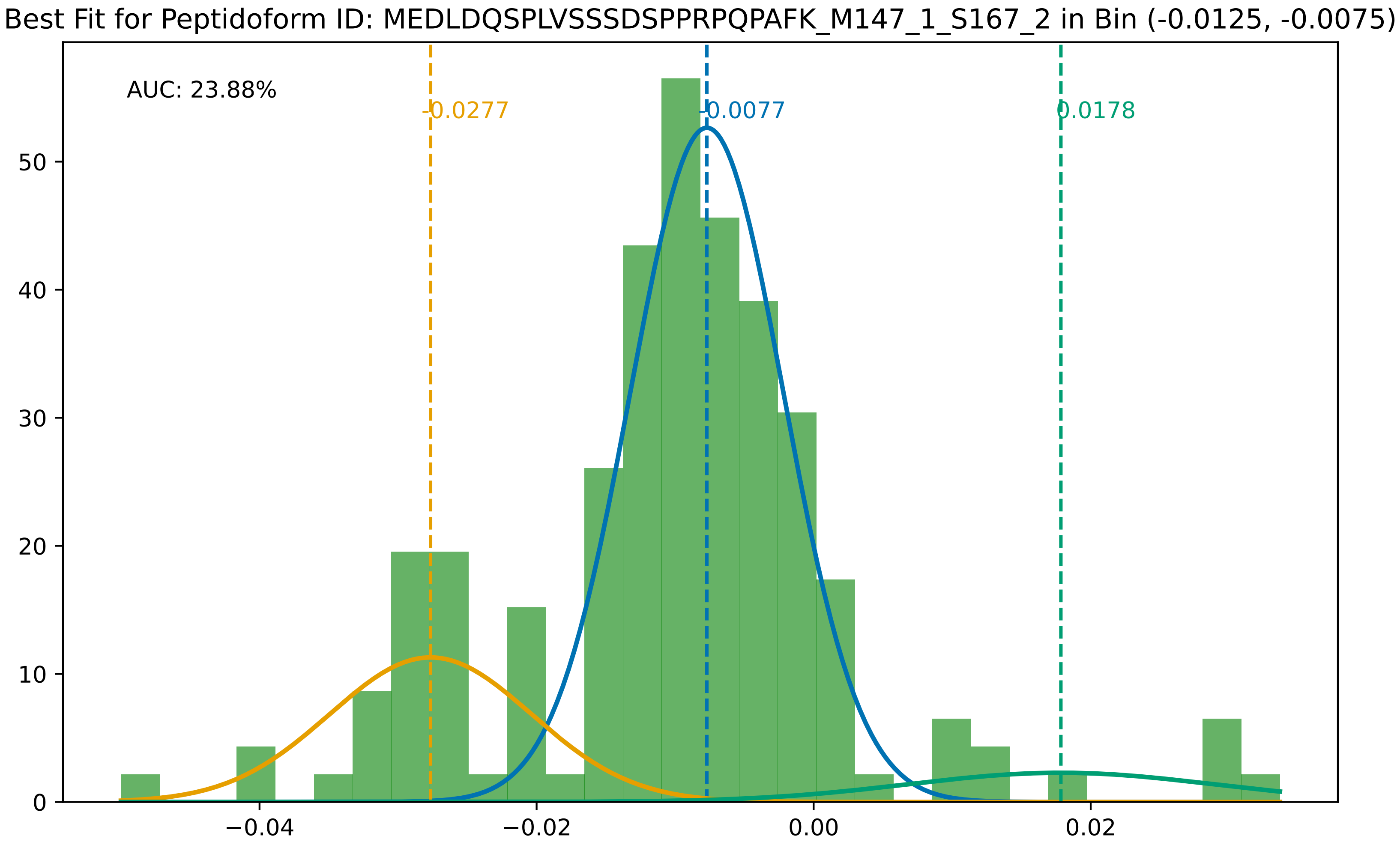

Best Fit for Peptidoform ID: MEDLDQSPLVSSSDSPPRPQPAFK_M147_1_S167_2 in Bin (-0.0125, -0.0075)
AUC: 23.88%
-0.0277
-0.0077
0.0178
0
10
20
30
40
50
−0.04
−0.02
0.00
0.02

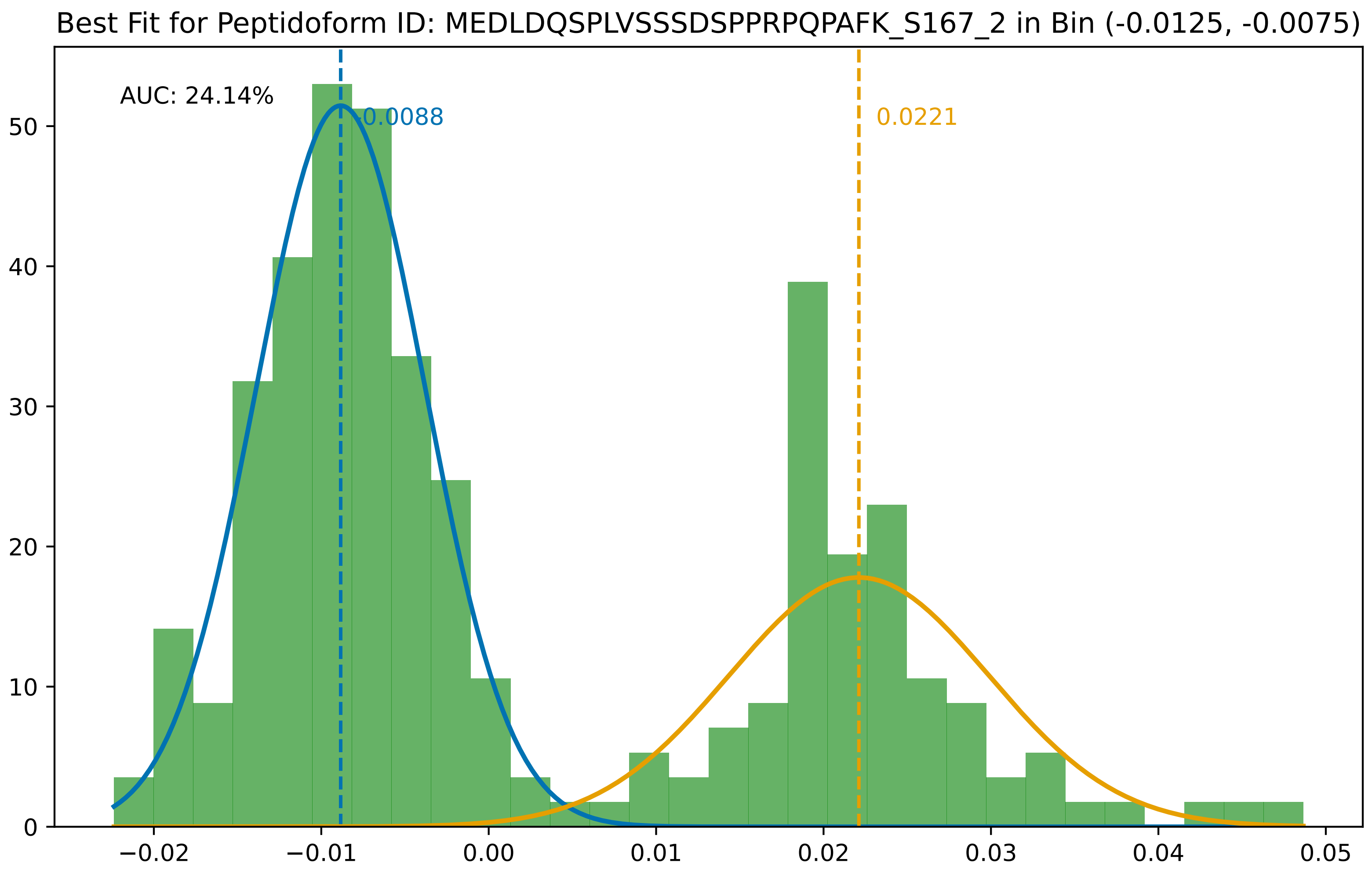
Best Fit for Peptidoform ID: MEDLDQSPLVSSSDSPPRPQPAFK_S167_2 in Bin (-0.0125, -0.0075)
AUC: 24.14%
-0.0088
0.0221
0
10
20
30
40
50
−0.02
−0.01
0.00
0.01
0.02
0.03
0.04
0.05

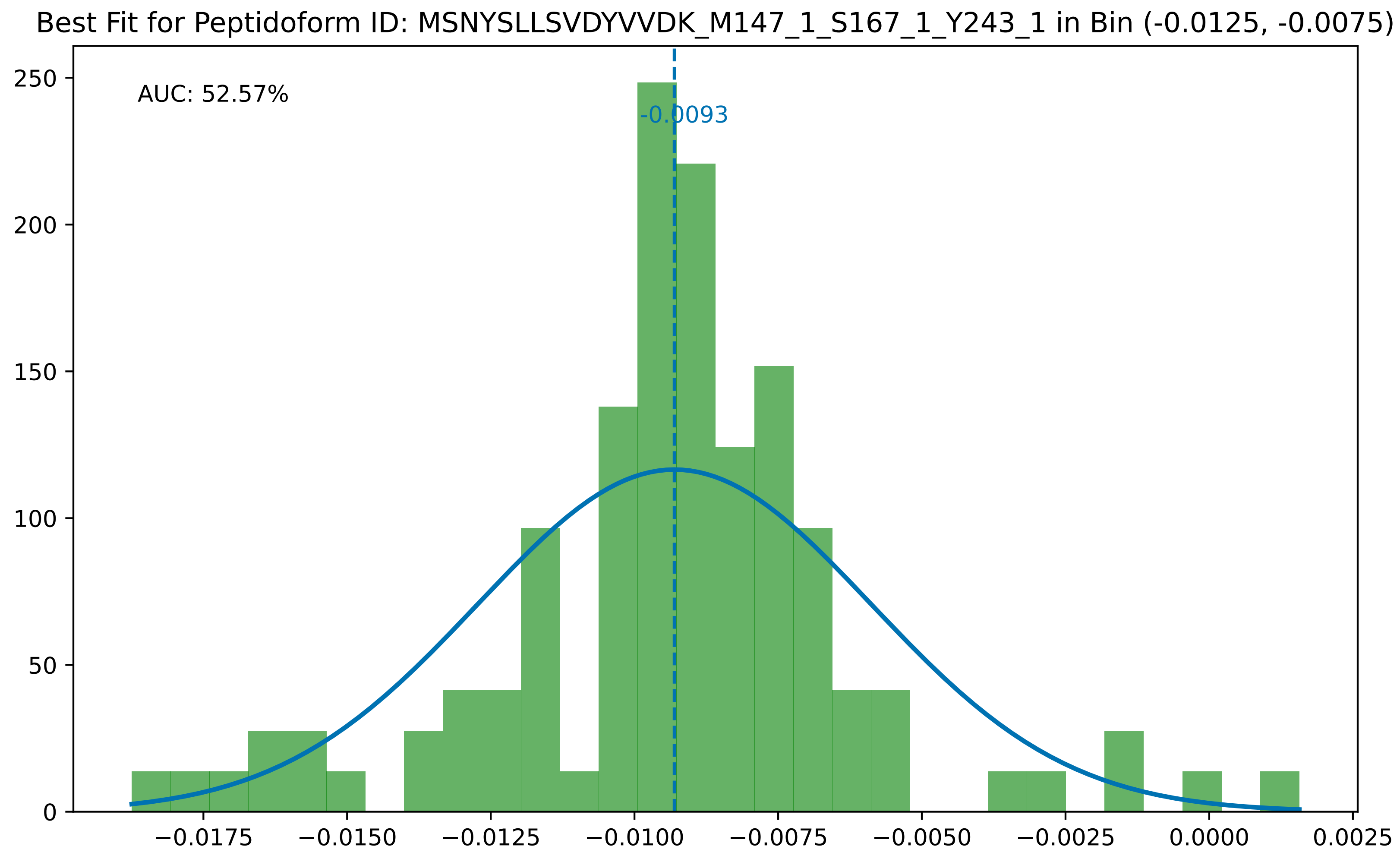

Best Fit for Peptidoform ID: MSNYSLLSVDYVVDK_M147_1_S167_1_Y243_1 in Bin (-0.0125, -0.0075)
AUC: 52.57%
-0.0093
250
200
150
100
50
0
−0.0175
−0.0150
−0.0125
−0.0100
−0.0075
−0.0050
−0.0025
0.0000
0.0025

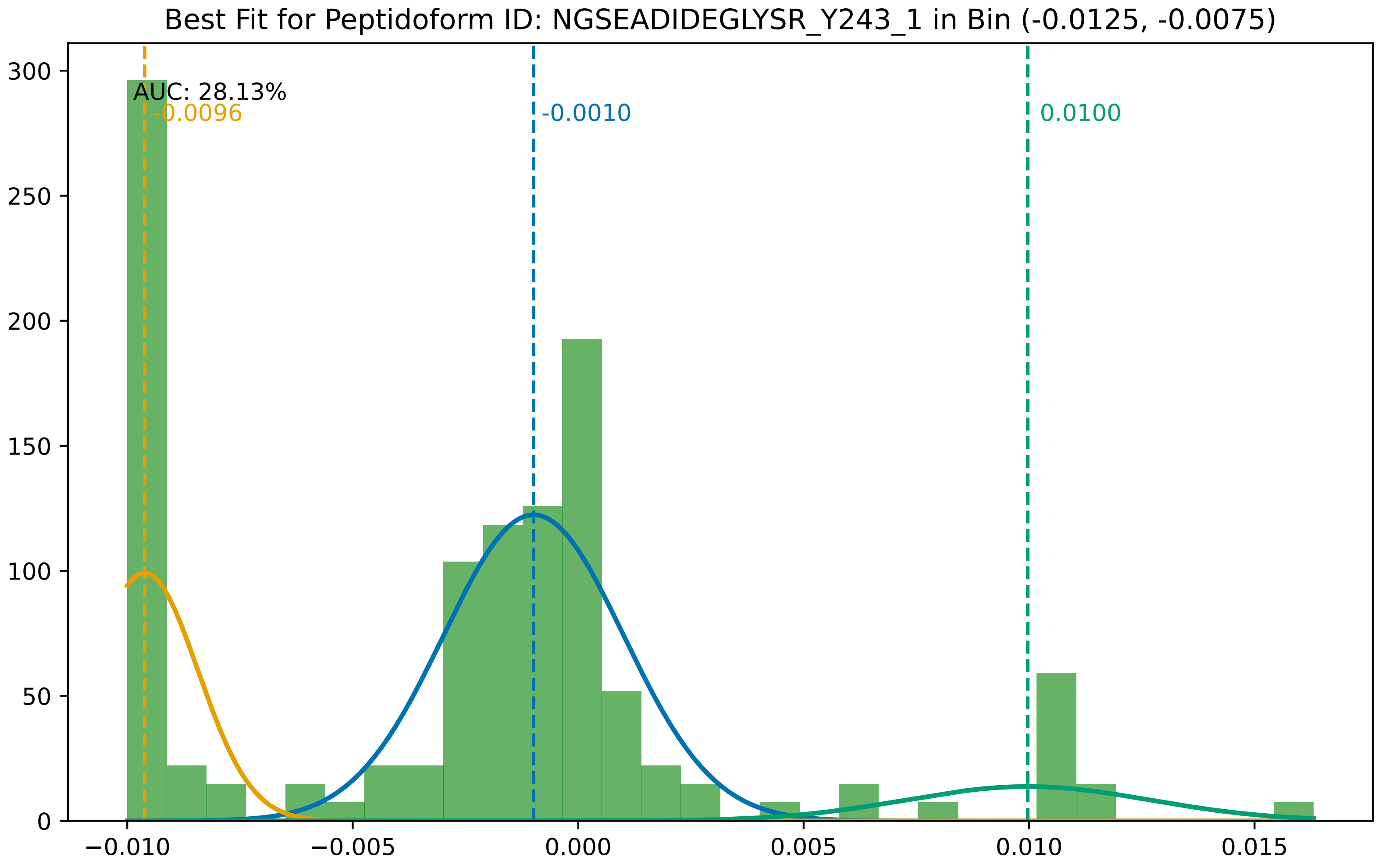

Best Fit for Peptidoform ID: NGSEADIDEGLYSR_Y243_1 in Bin (-0.0125, -0.0075)
AUC: 28.13%
-0.0096
-0.0010
0.0100
300
250
200
150
100
50
0
−0.010
−0.005
0.000
0.005
0.010
0.015

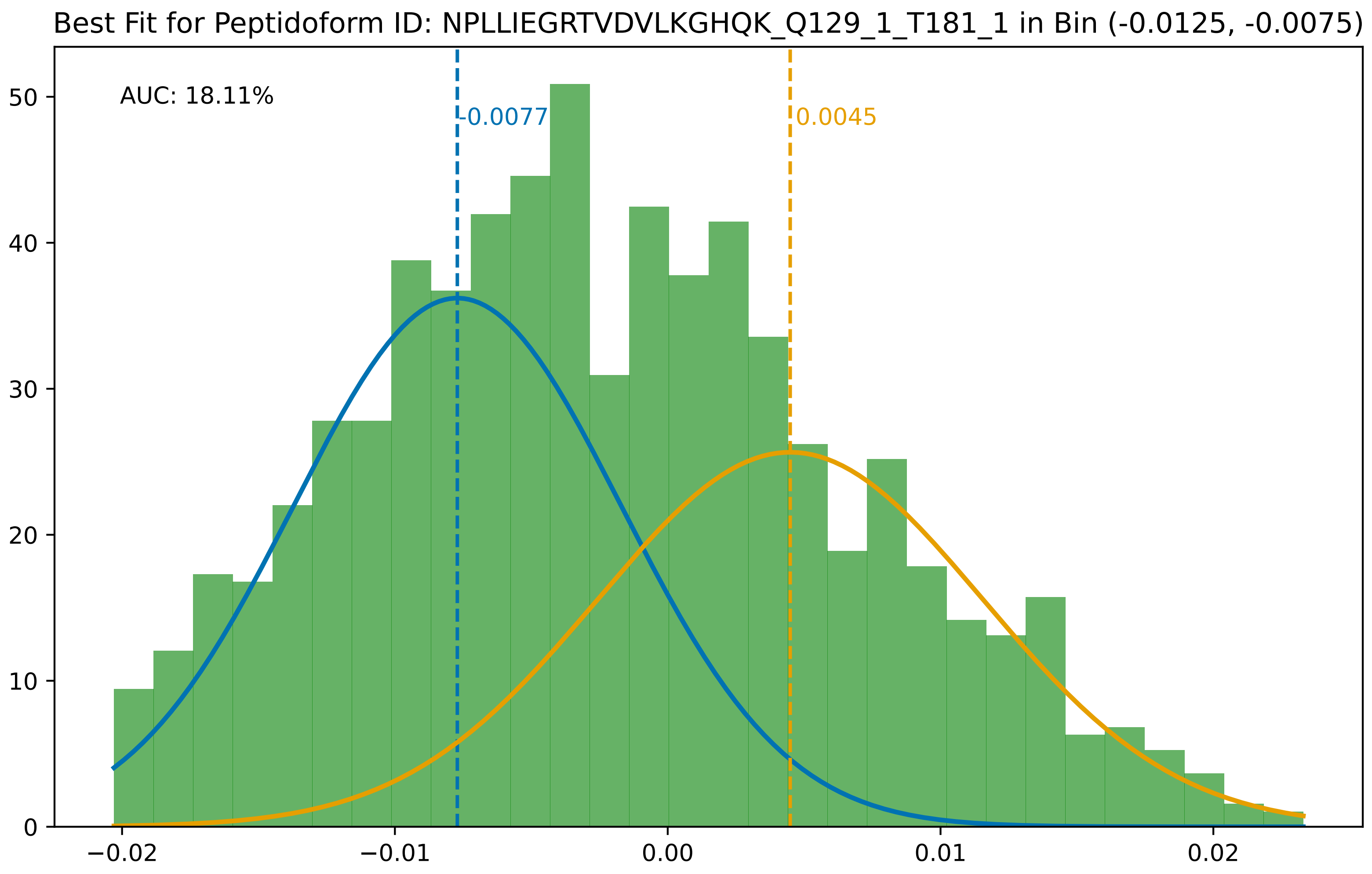

Best Fit for Peptidoform ID: NPLLIEGRTVDVLKGHQK_Q129_1_T181_1 in Bin (-0.0125, -0.0075)
AUC: 18.11%
-0.0077
0.0045
0
10
20
30
40
50
−0.02
−0.01
0.00
0.01
0.02

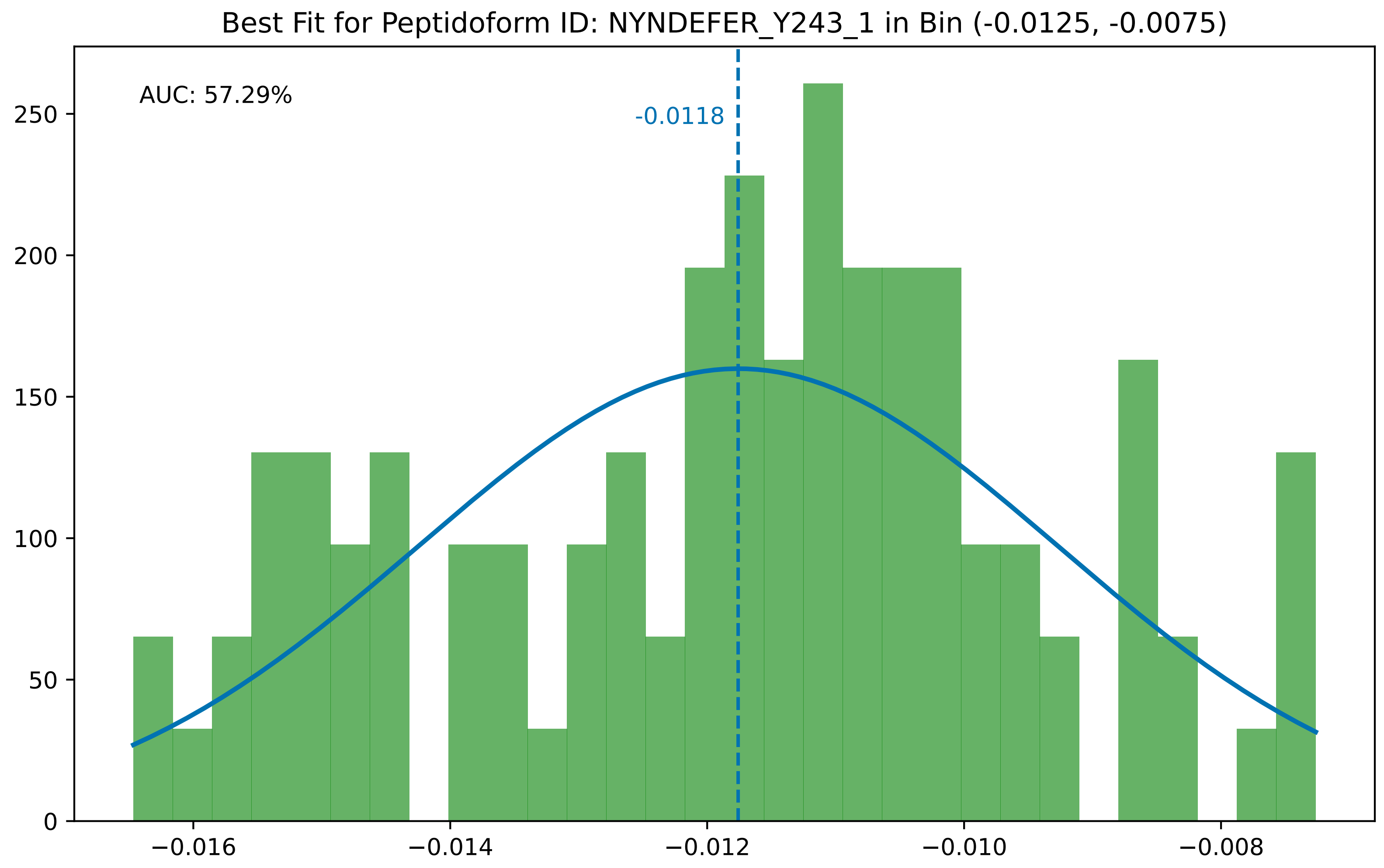
Best Fit for Peptidoform ID: NYNDEFER_Y243_1 in Bin (-0.0125, -0.0075)
AUC: 57.29%
-0.0118
0
50
100
150
200
250
−0.016
−0.014
−0.012
−0.010
−0.008

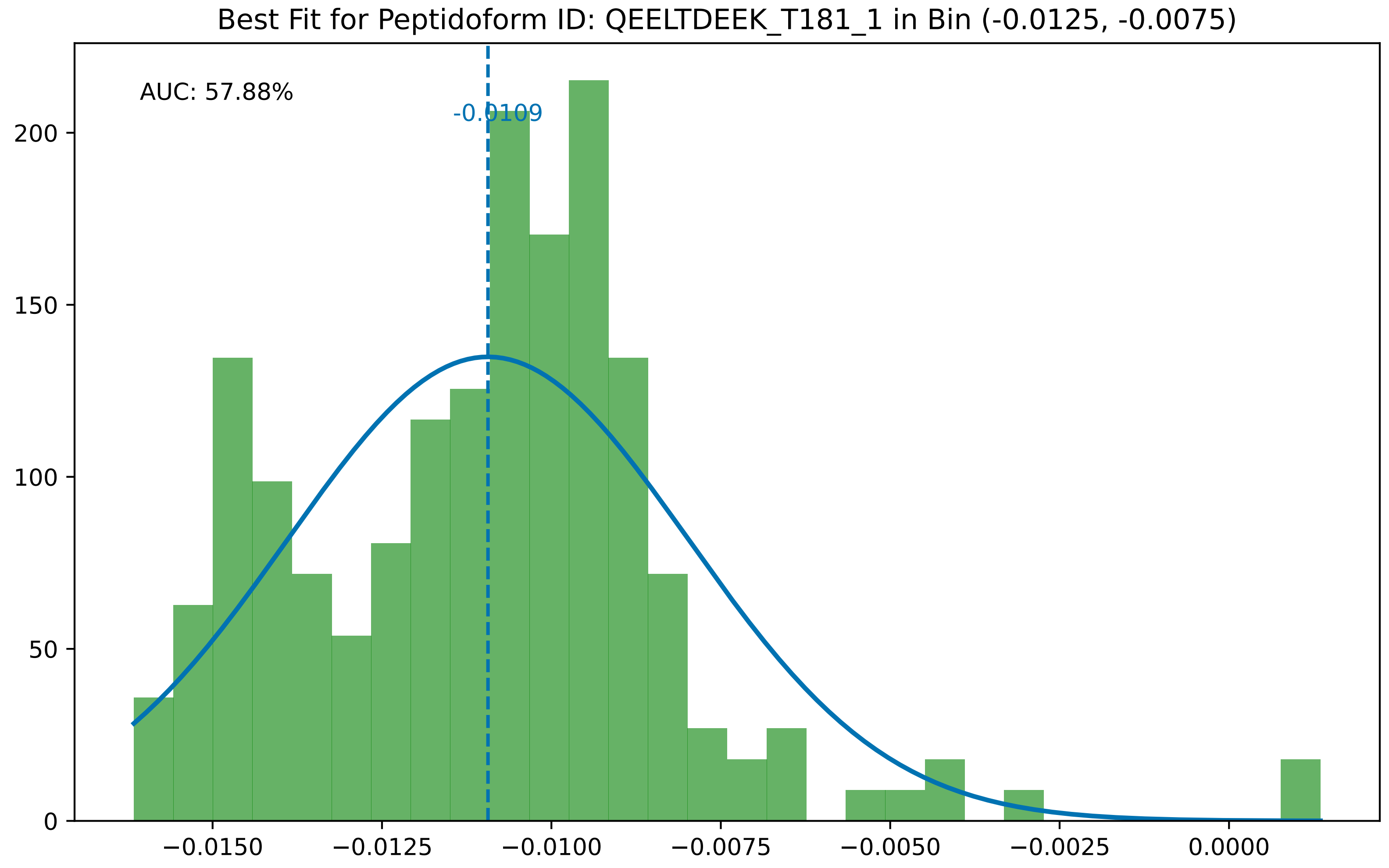
Best Fit for Peptidoform ID: QEELTDEEK_T181_1 in Bin (-0.0125, -0.0075)
AUC: 57.88%
-0.0109
200
150
100
50
0
−0.0150
−0.0125
−0.0100
−0.0075
−0.0050
−0.0025
0.0000

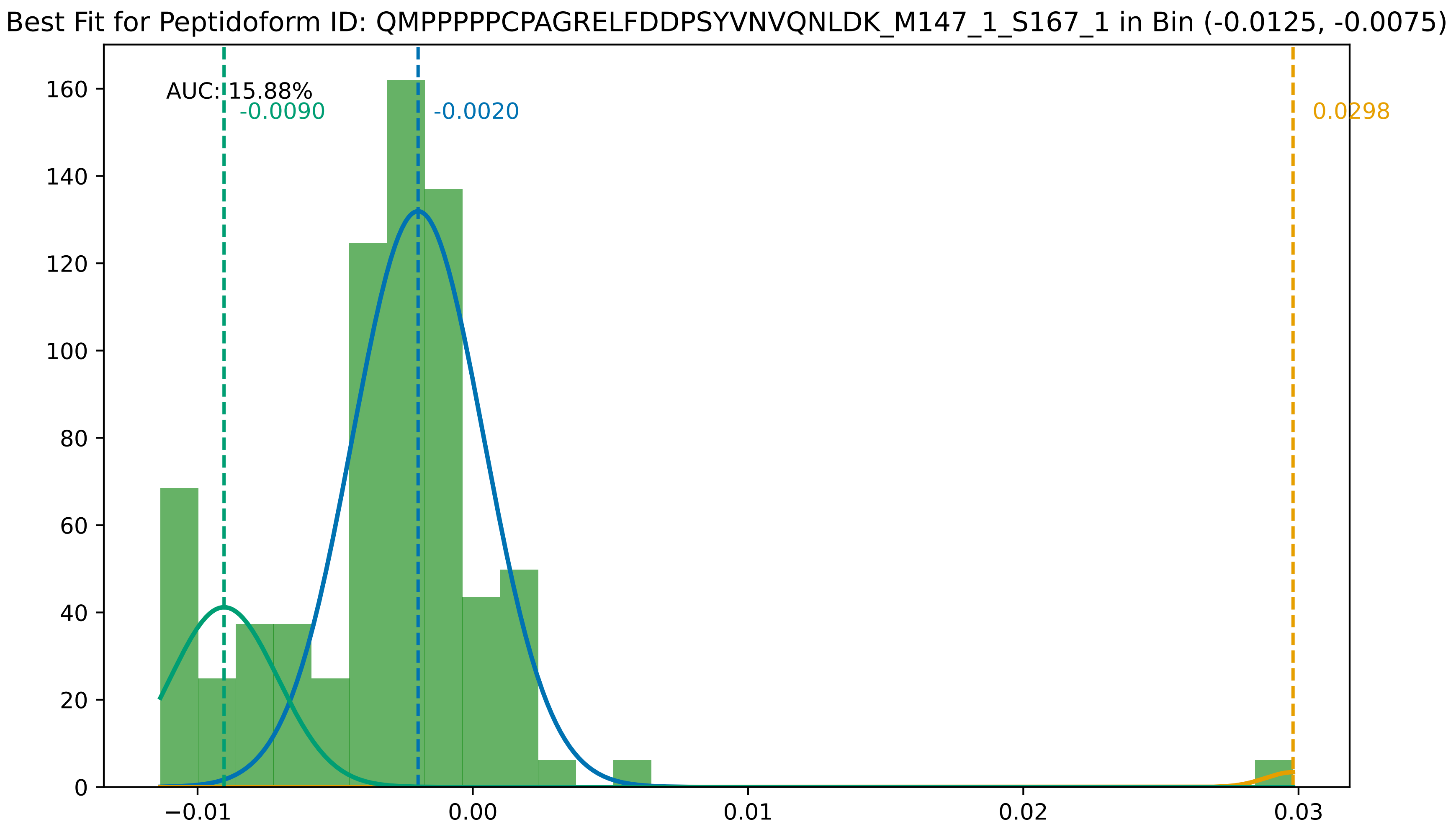
Best Fit for Peptidoform ID: QMPPPPPCPAGRELFDDPSYVNVQNLDK_M147_1_S167_1 in Bin (-0.0125, -0.0075)
AUC: 15.88%
-0.0090
-0.0020
0.0298
0
20
40
60
80
100
120
140
160
−0.01
0.00
0.01
0.02
0.03

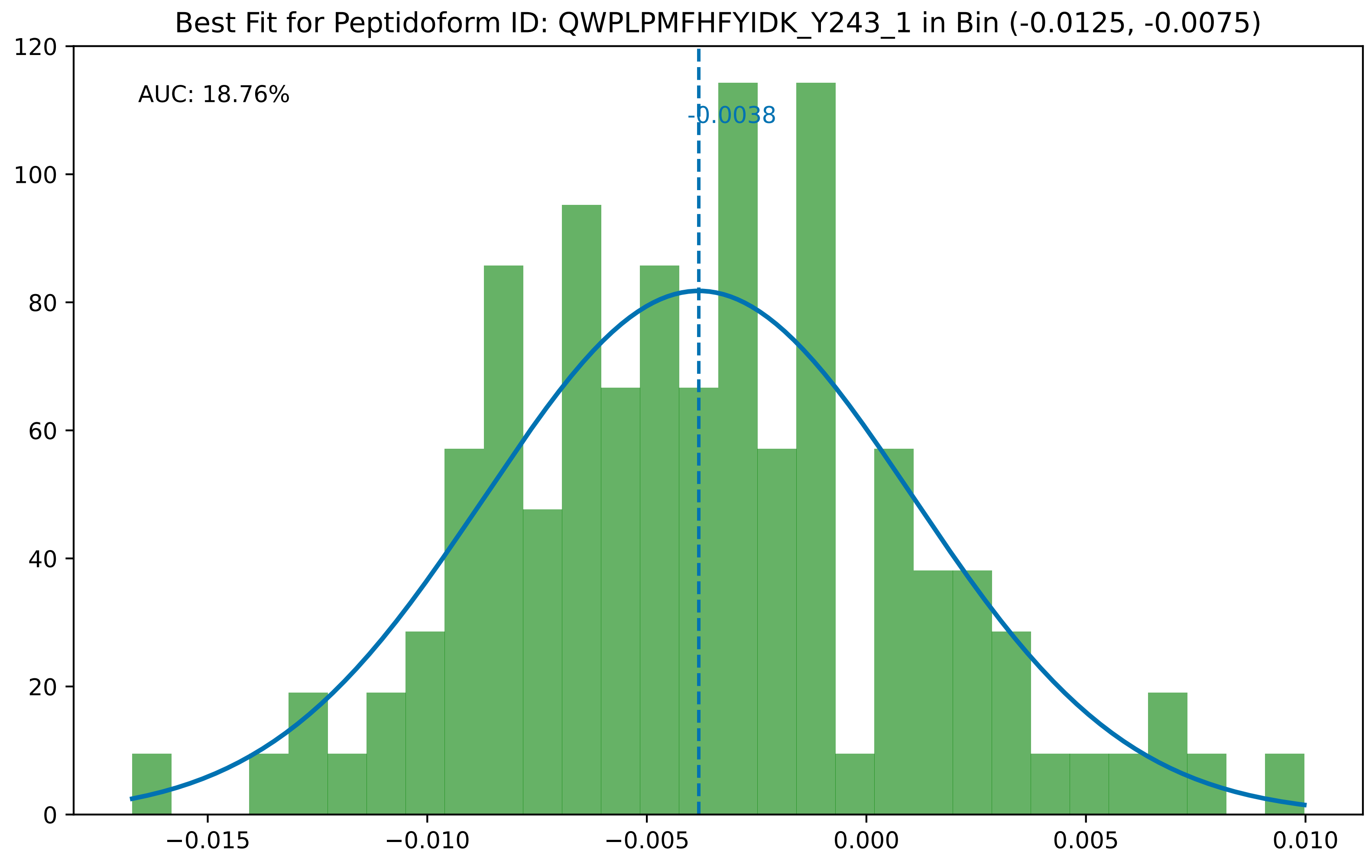
Best Fit for Peptidoform ID: QWPLPMFHFYIDK_Y243_1 in Bin (-0.0125, -0.0075)
AUC: 18.76%
-0.0038
120
100
80
60
40
20
0
−0.015
−0.010
−0.005
0.000
0.005
0.010

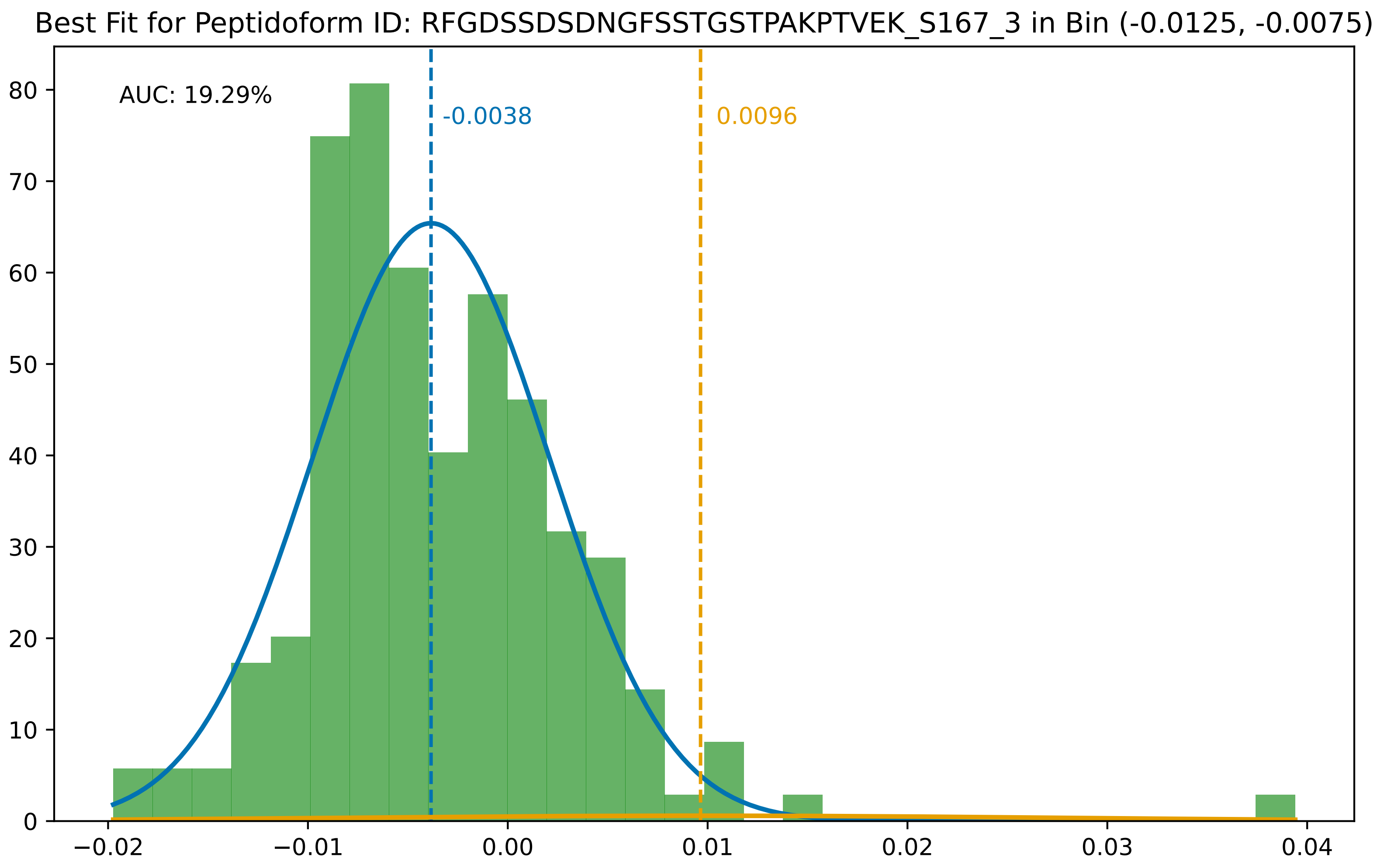

Best Fit for Peptidoform ID: RFGDSSDSDNGFSSTGSTPAKPTVEK_S167_3 in Bin (-0.0125, -0.0075)
AUC: 19.29%
-0.0038
0.0096
0
10
20
30
40
50
60
70
80
−0.02
−0.01
0.00
0.01
0.02
0.03
0.04

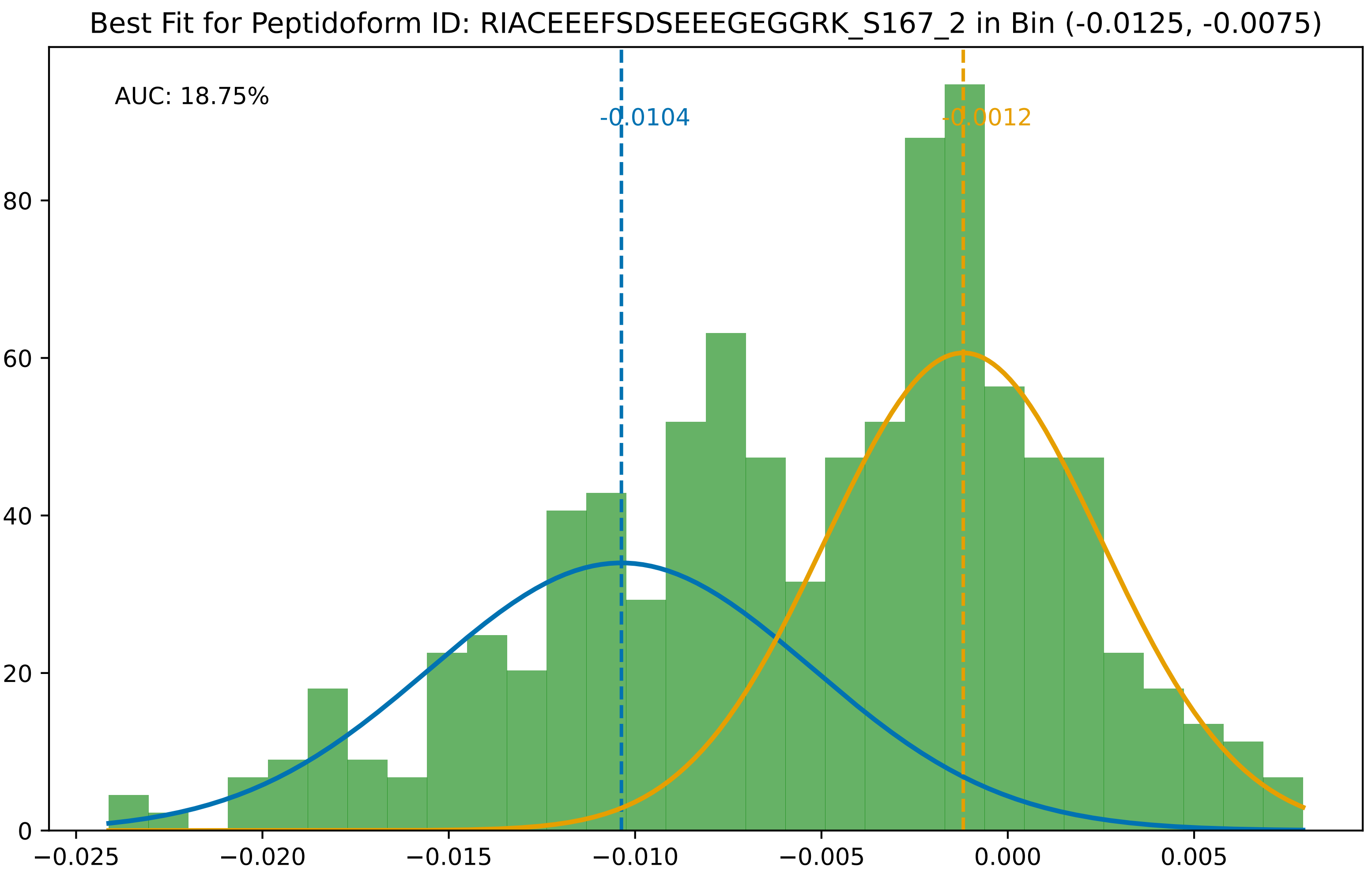

Best Fit for Peptidoform ID: RIACEEEFSDSEEEGEGGRK_S167_2 in Bin (-0.0125, -0.0075)
AUC: 18.75%
-0.0104
-0.0012
0
20
40
60
80
−0.025
−0.020
−0.015
−0.010
−0.005
0.000
0.005

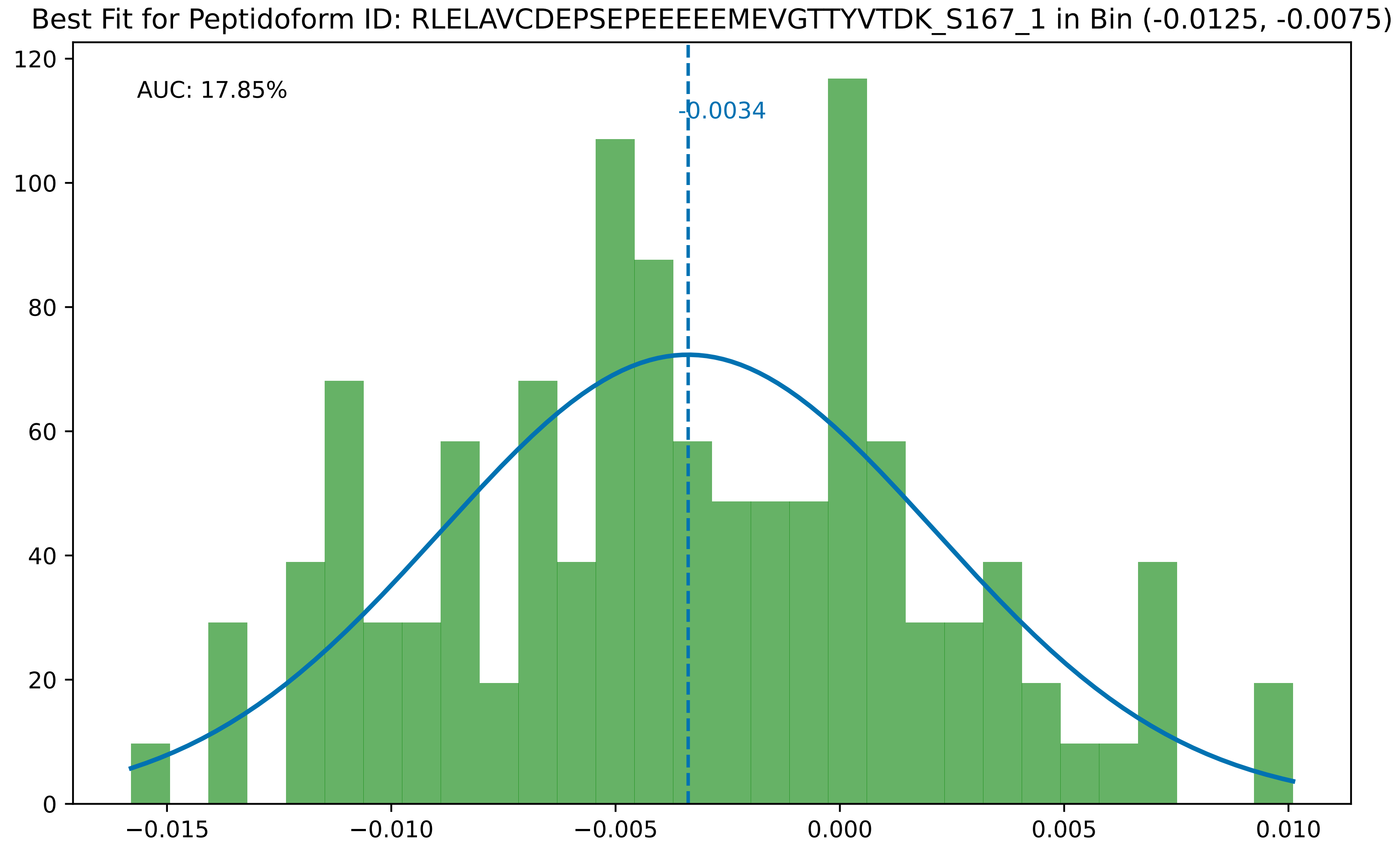

Best Fit for Peptidoform ID: RLELAVCDEPSEPEEEEEMEVGTTYVTDK_S167_1 in Bin (-0.0125, -0.0075)
AUC: 17.85%
-0.0034
0
20
40
60
80
100
120
−0.015
−0.010
−0.005
0.000
0.005
0.010

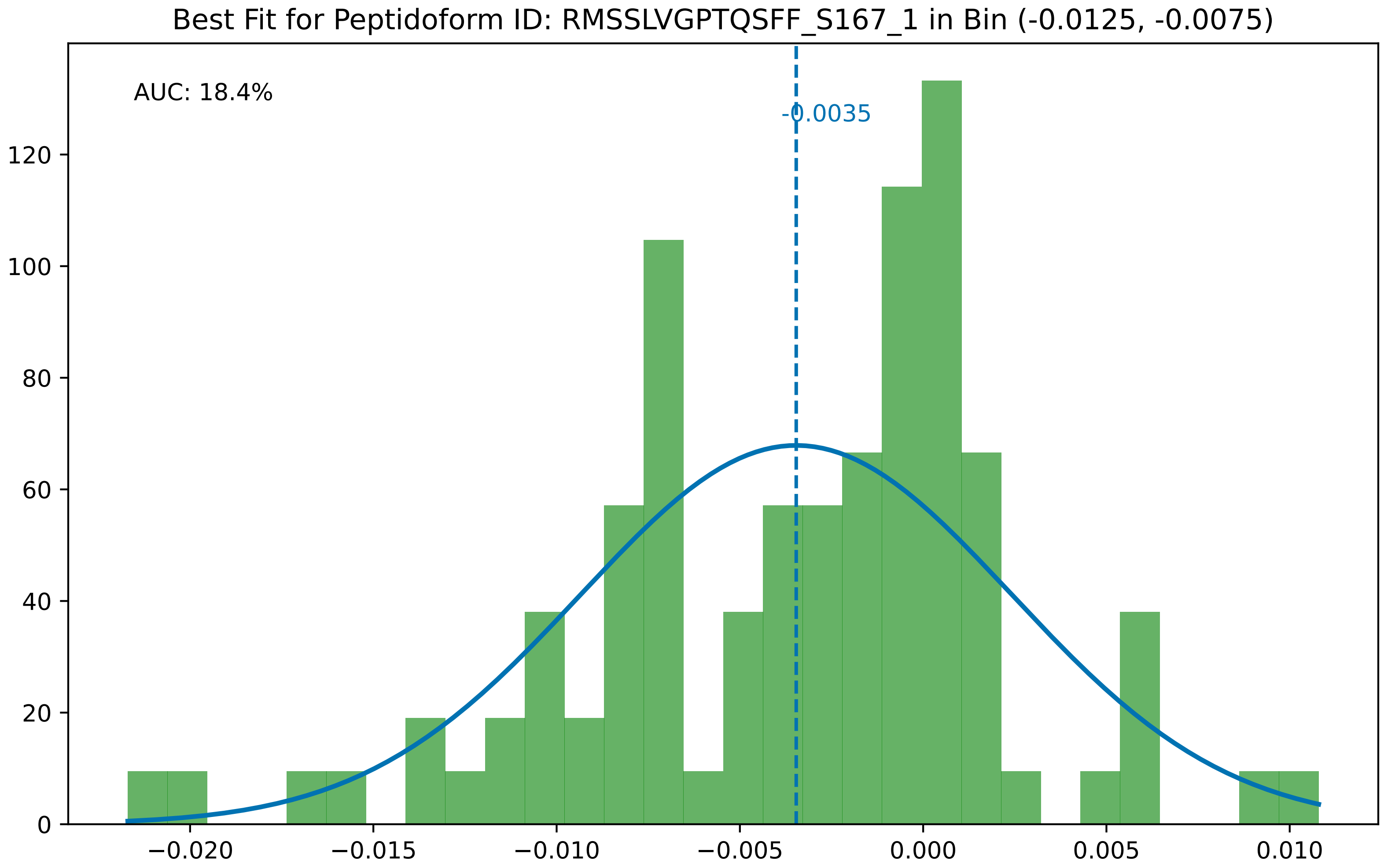
Best Fit for Peptidoform ID: RMSSLVGPTQSFF_S167_1 in Bin (-0.0125, -0.0075)
AUC: 18.4%
-0.0035
0
20
40
60
80
100
120
−0.020
−0.015
−0.010
−0.005
0.000
0.005
0.010

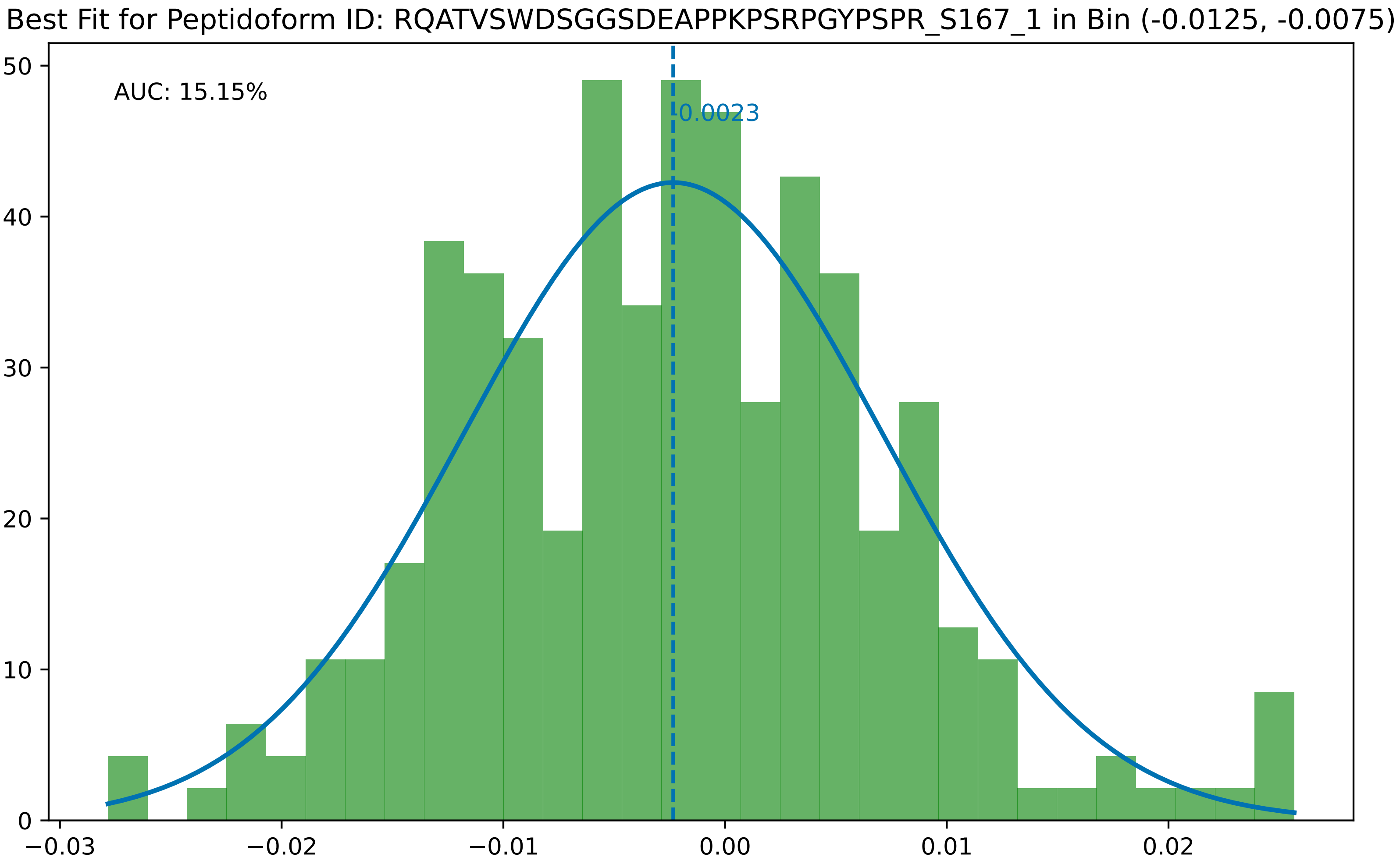

Best Fit for Peptidoform ID: RQATVSWDSGGSDEAPPKPSRPGYPSPR_S167_1 in Bin (-0.0125, -0.0075)
AUC: 15.15%
-0.0023
0
10
20
30
40
50
−0.03
−0.02
−0.01
0.00
0.01
0.02

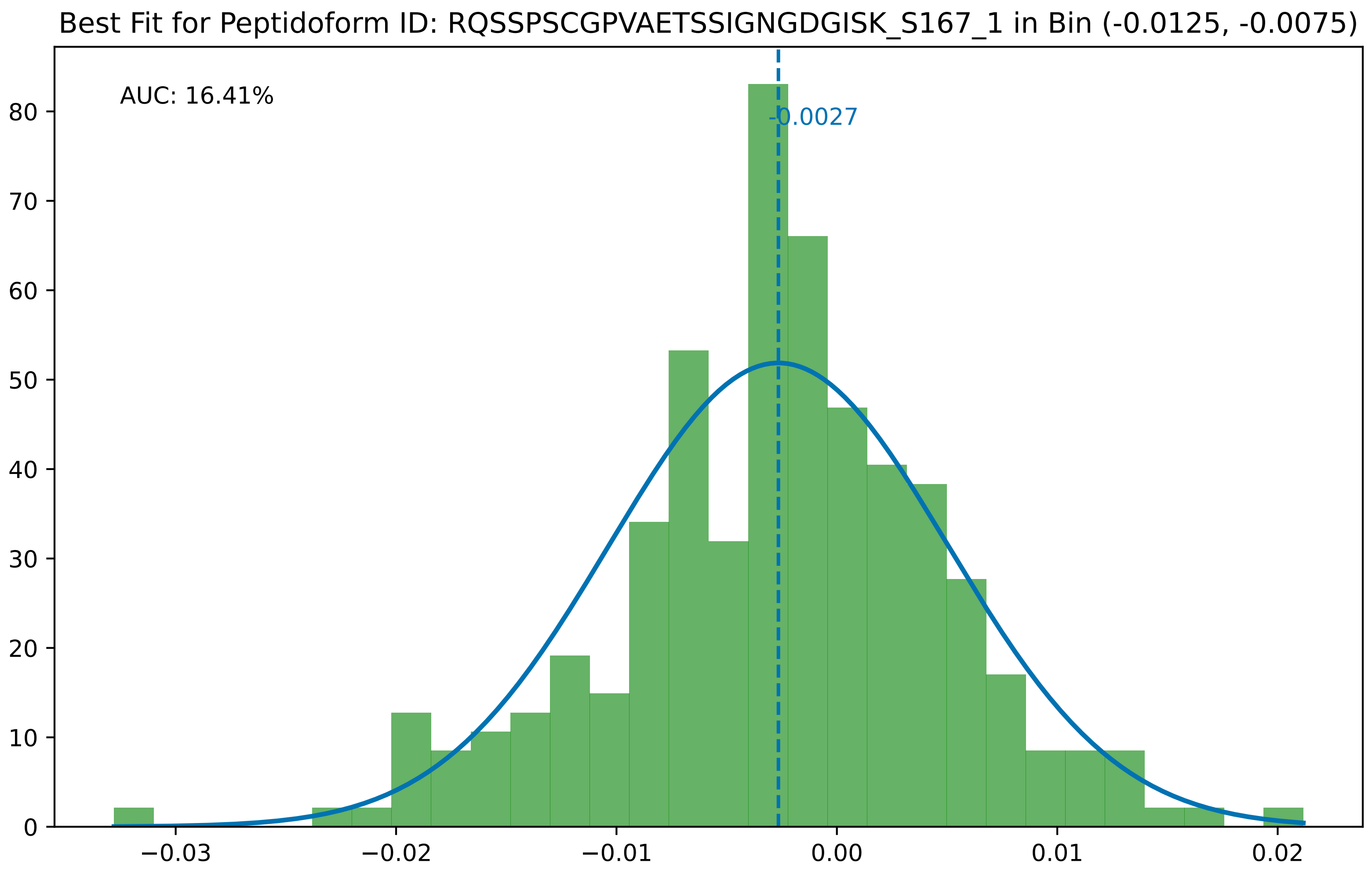

Best Fit for Peptidoform ID: RQSSPSCGPVAETSSIGNGDGISK_S167_1 in Bin (-0.0125, -0.0075)
AUC: 16.41%
-0.0027
0
10
20
30
40
50
60
70
80
−0.03
−0.02
−0.01
0.00
0.01
0.02

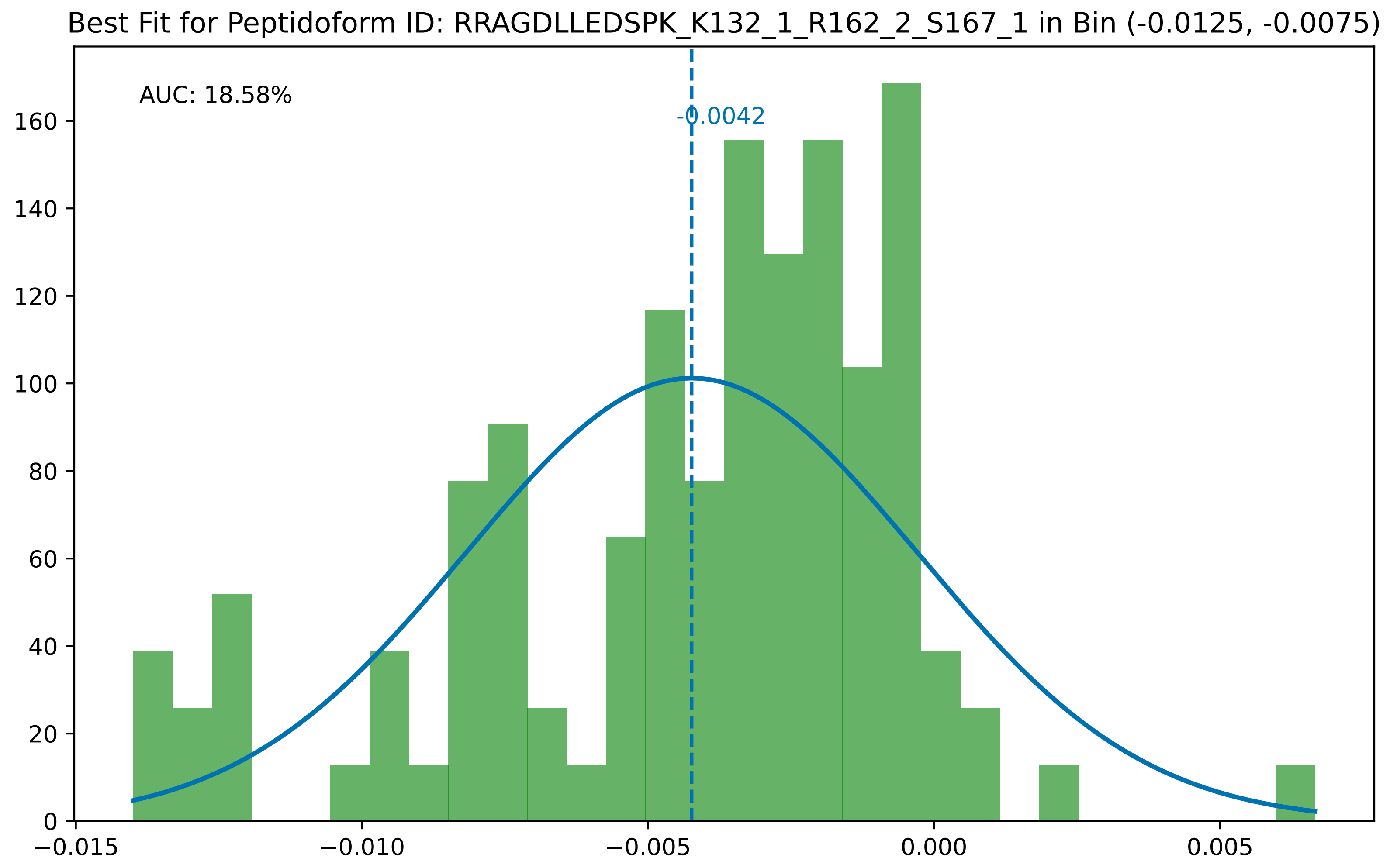
Best Fit for Peptidoform ID: RRAGDLLEDSPK_K132_1_R162_2_S167_1 in Bin (-0.0125, -0.0075)
AUC: 18.58%
-0.0042
0
20
40
60
80
100
120
140
160
−0.015
−0.010
−0.005
0.000
0.005

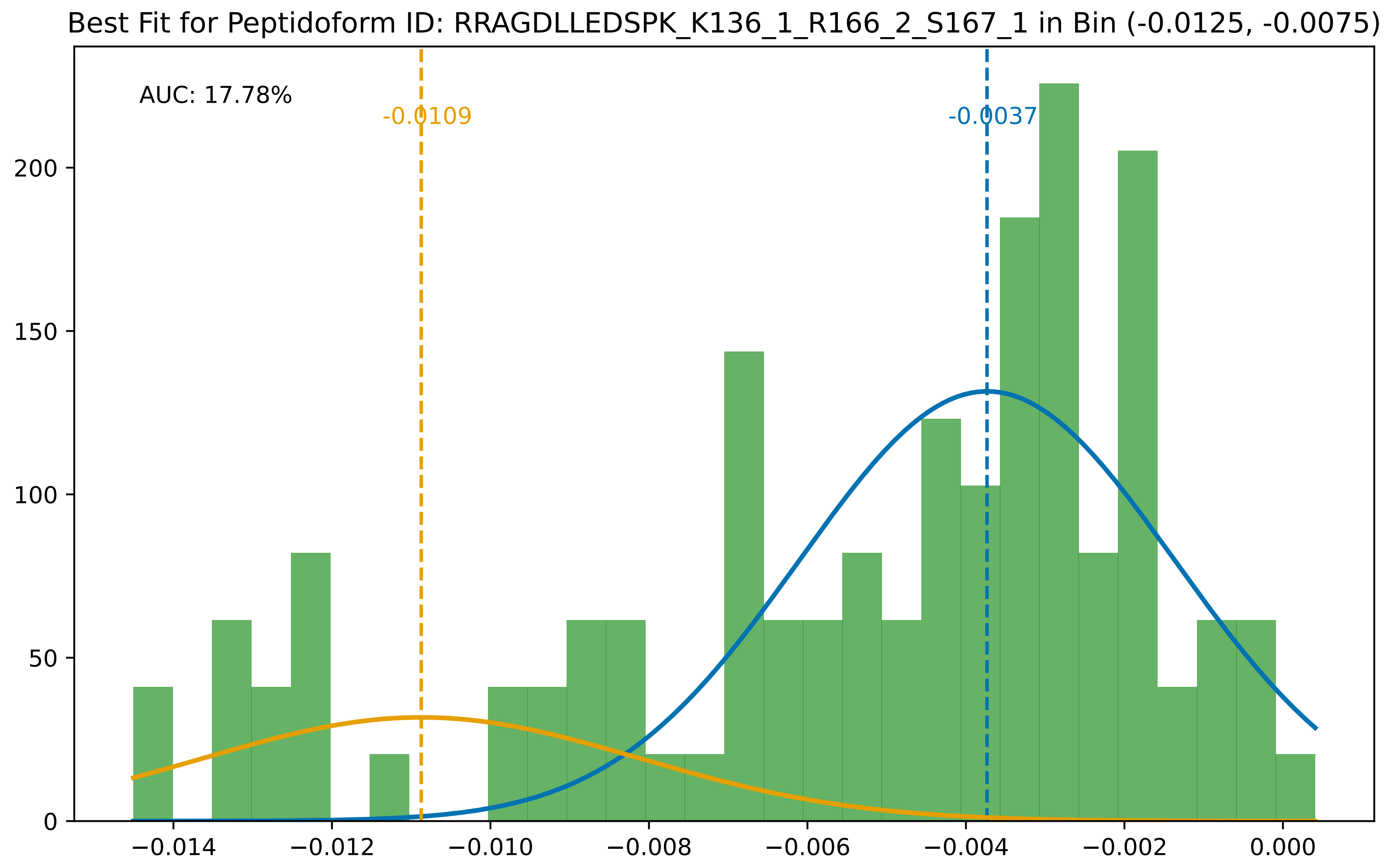
Best Fit for Peptidoform ID: RRAGDLLEDSPK_K136_1_R166_2_S167_1 in Bin (-0.0125, -0.0075)
AUC: 17.78%
-0.0109
-0.0037
0
50
100
150
200
−0.014
−0.012
−0.010
−0.008
−0.006
−0.004
−0.002
0.000

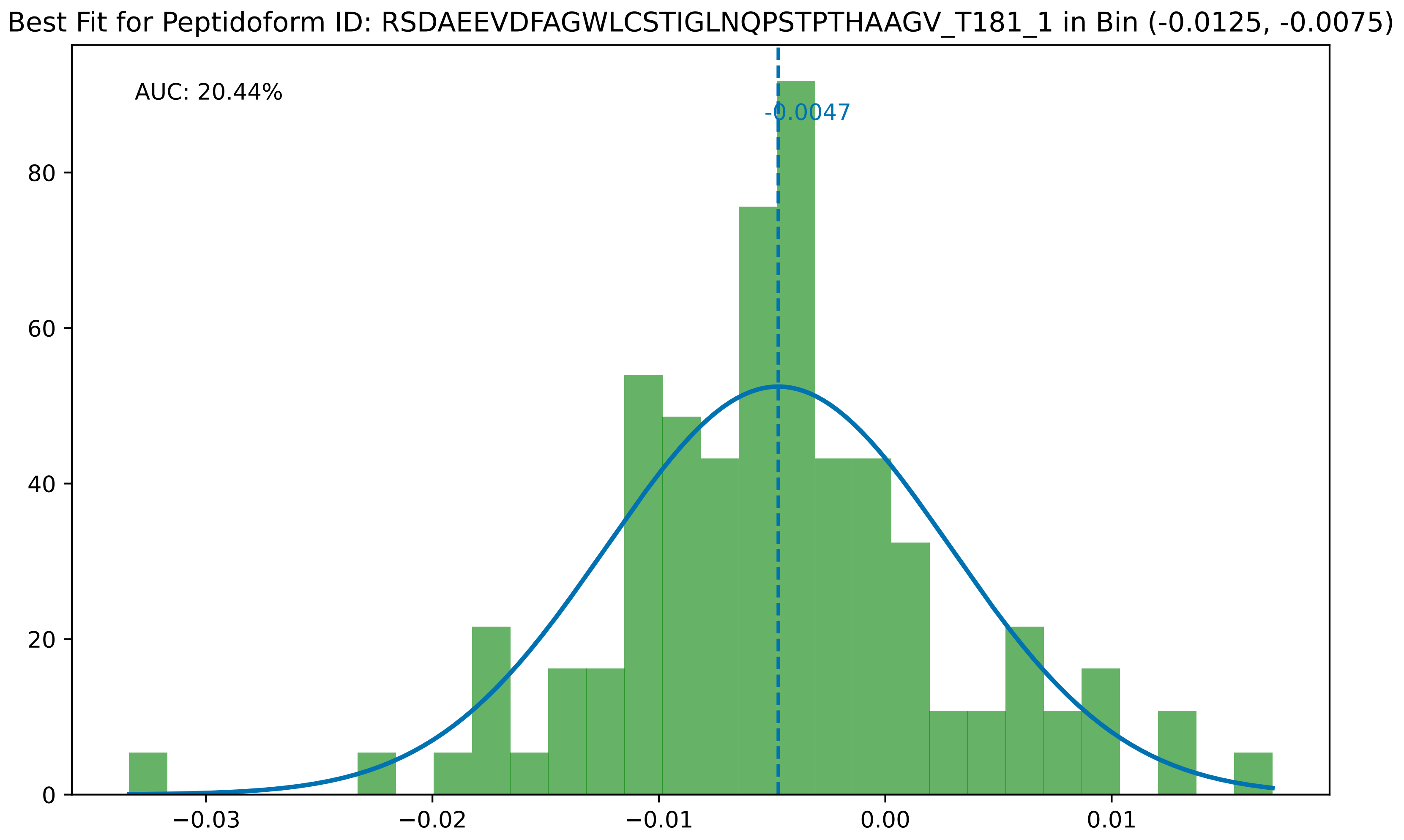
Best Fit for Peptidoform ID: RSDAEEVDFAGWLCSTIGLNQPSTPTHAAGV_T181_1 in Bin (-0.0125, -0.0075)
AUC: 20.44%
-0.0047
0
20
40
60
80
−0.03
−0.02
−0.01
0.00
0.01

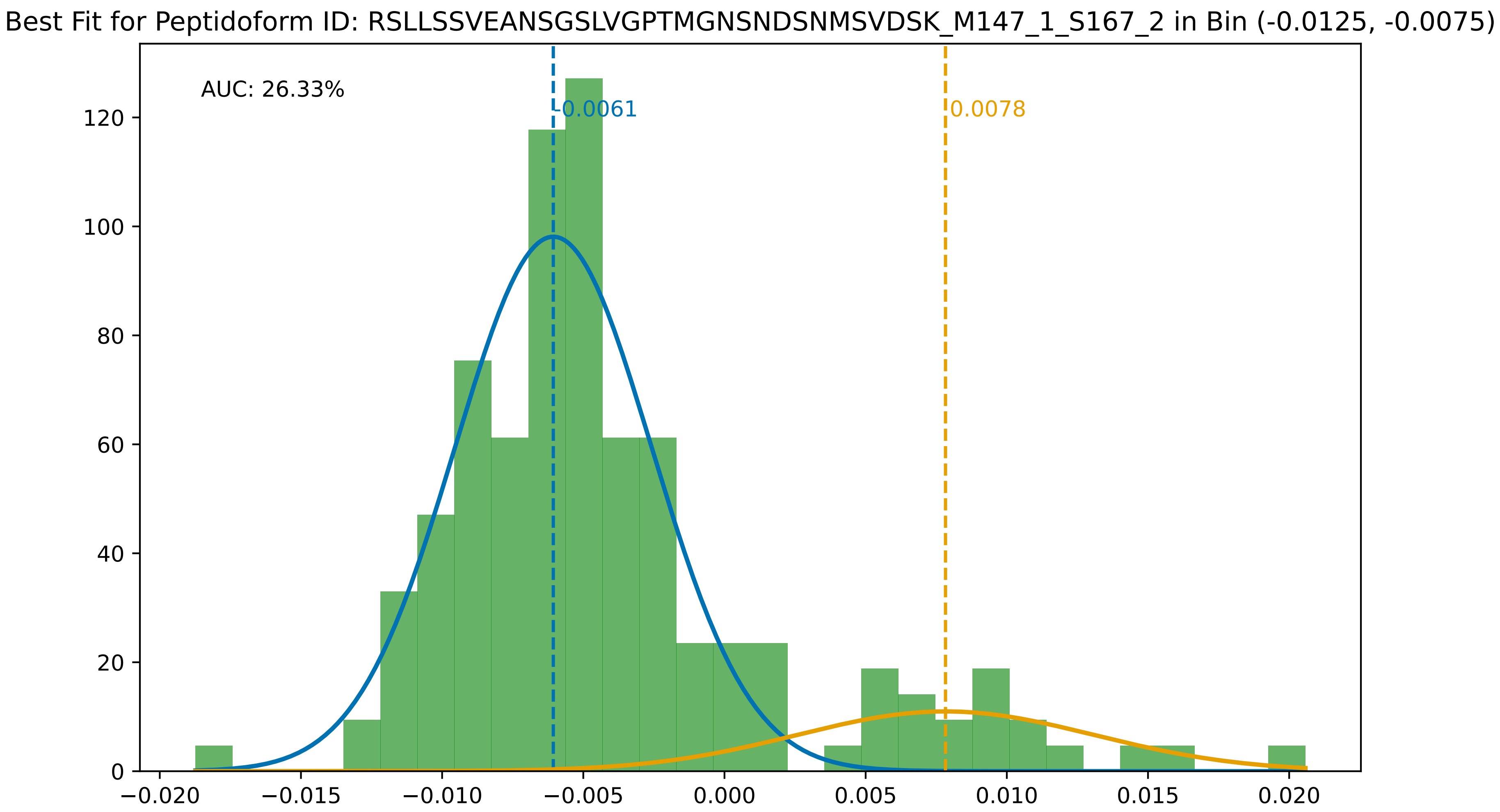
Best Fit for Peptidoform ID: RSLLSSVEANSGSLVGPTMGNSNDSNMSVDSK_M147_1_S167_2 in Bin (-0.0125, -0.0075)
AUC: 26.33%
-0.0061
0.0078
0
20
40
60
80
100
120
−0.020
−0.015
−0.010
−0.005
0.000
0.005
0.010
0.015
0.020

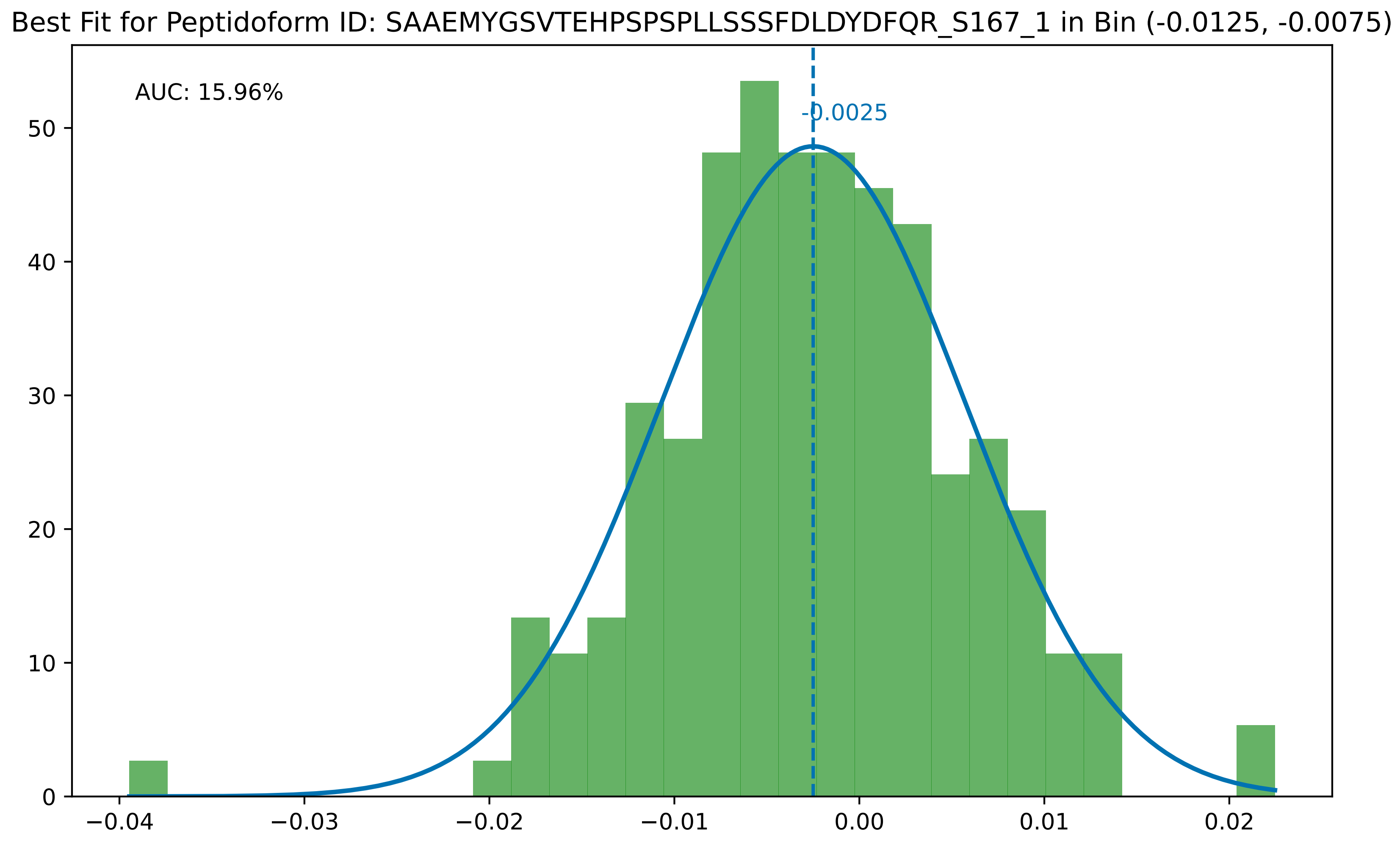

Best Fit for Peptidoform ID: SAAEMYGSVTEHPSPSPLLSSSFDLDYDFQR_S167_1 in Bin (-0.0125, -0.0075)
AUC: 15.96%
-0.0025
0
10
20
30
40
50
−0.04
−0.03
−0.02
−0.01
0.00
0.01
0.02

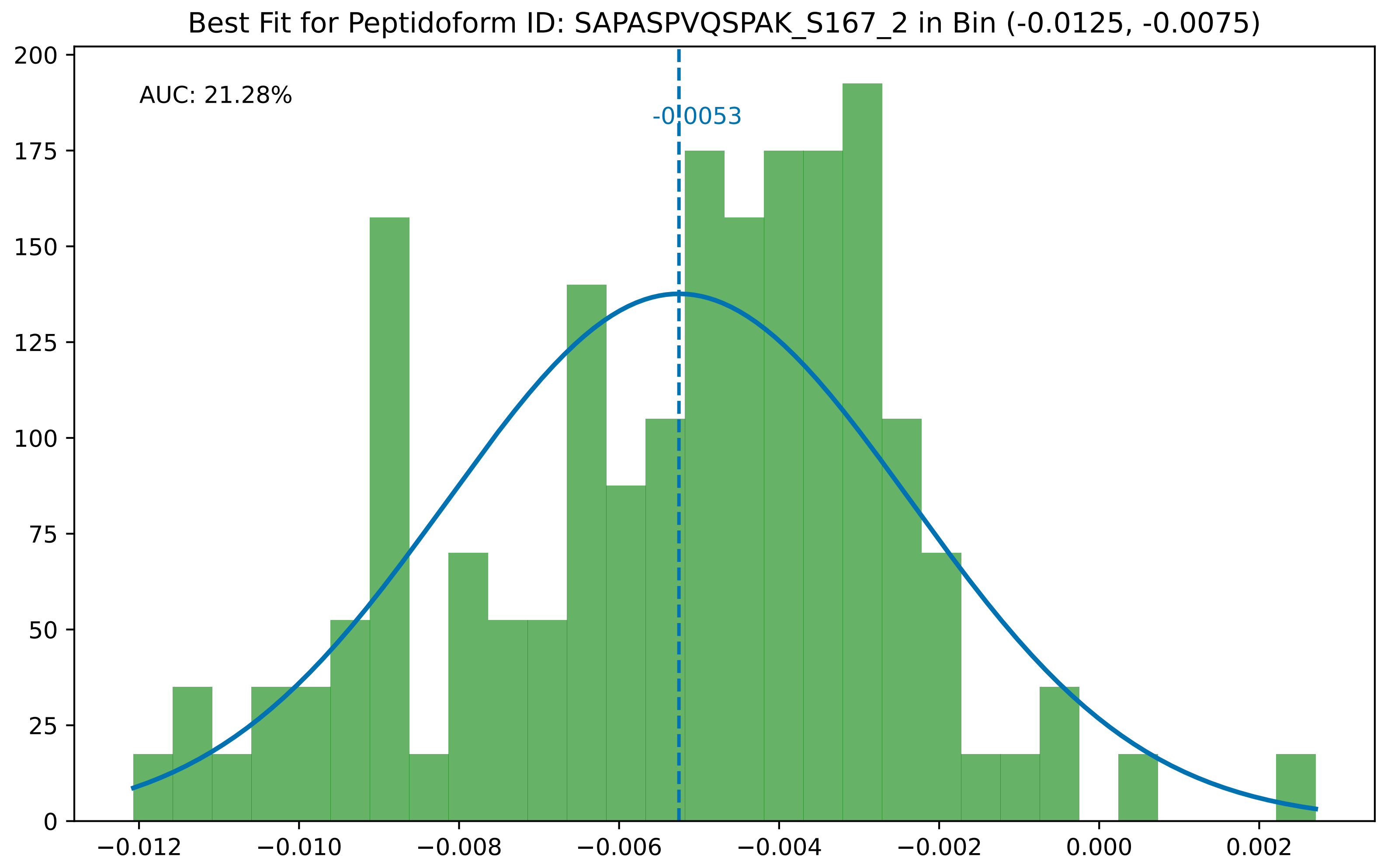
Best Fit for Peptidoform ID: SAPASPVQSPAK_S167_2 in Bin (-0.0125, -0.0075)
AUC: 21.28%
-0.0053
0
25
50
75
100
125
150
175
200
−0.012
−0.010
−0.008
−0.006
−0.004
−0.002
0.000
0.002

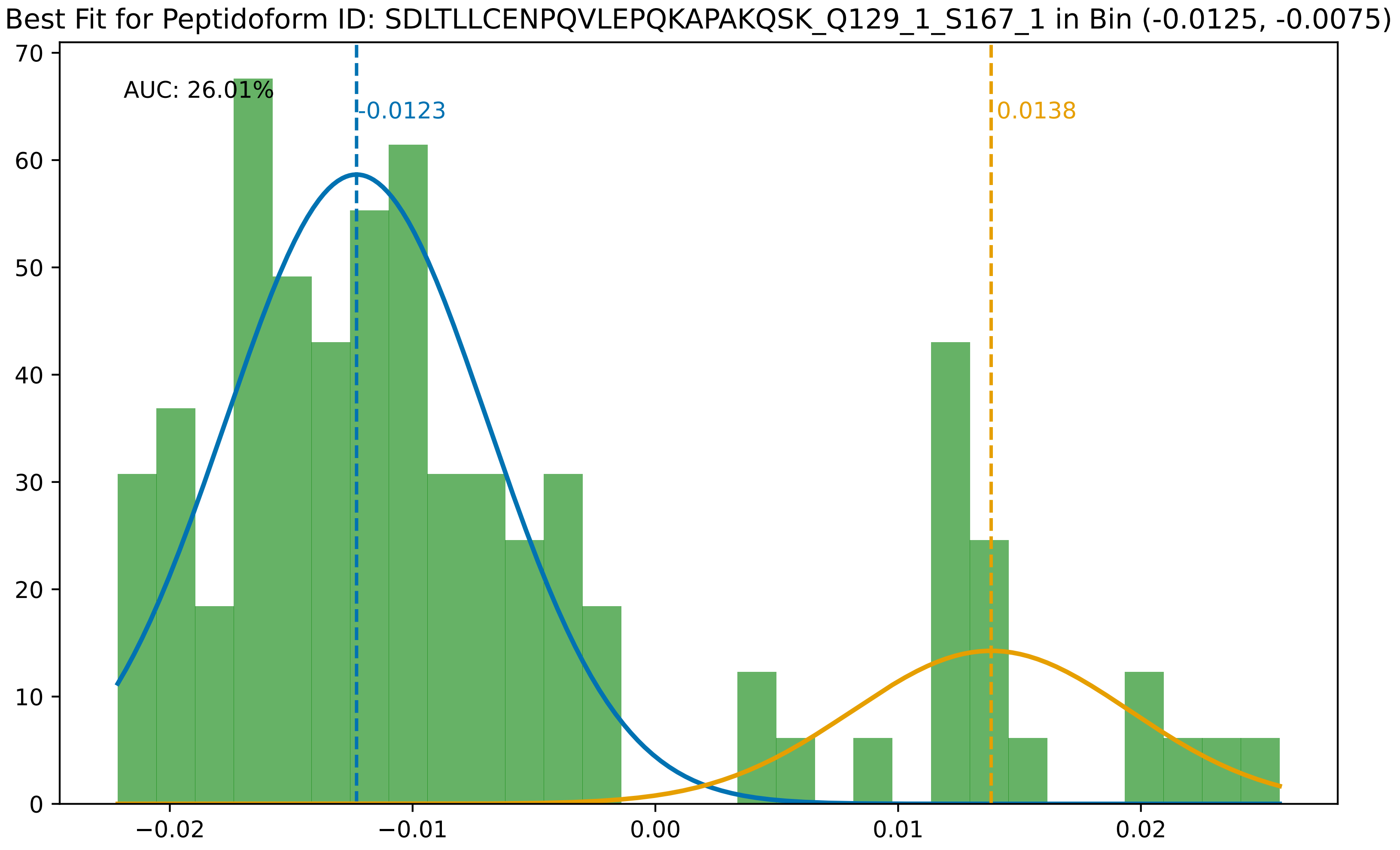

Best Fit for Peptidoform ID: SDLTLLCENPQVLEPQKAPAKQSK_Q129_1_S167_1 in Bin (-0.0125, -0.0075)
AUC: 26.01%
-0.0123
0.0138
0
10
20
30
40
50
60
70
−0.02
−0.01
0.00
0.01
0.02

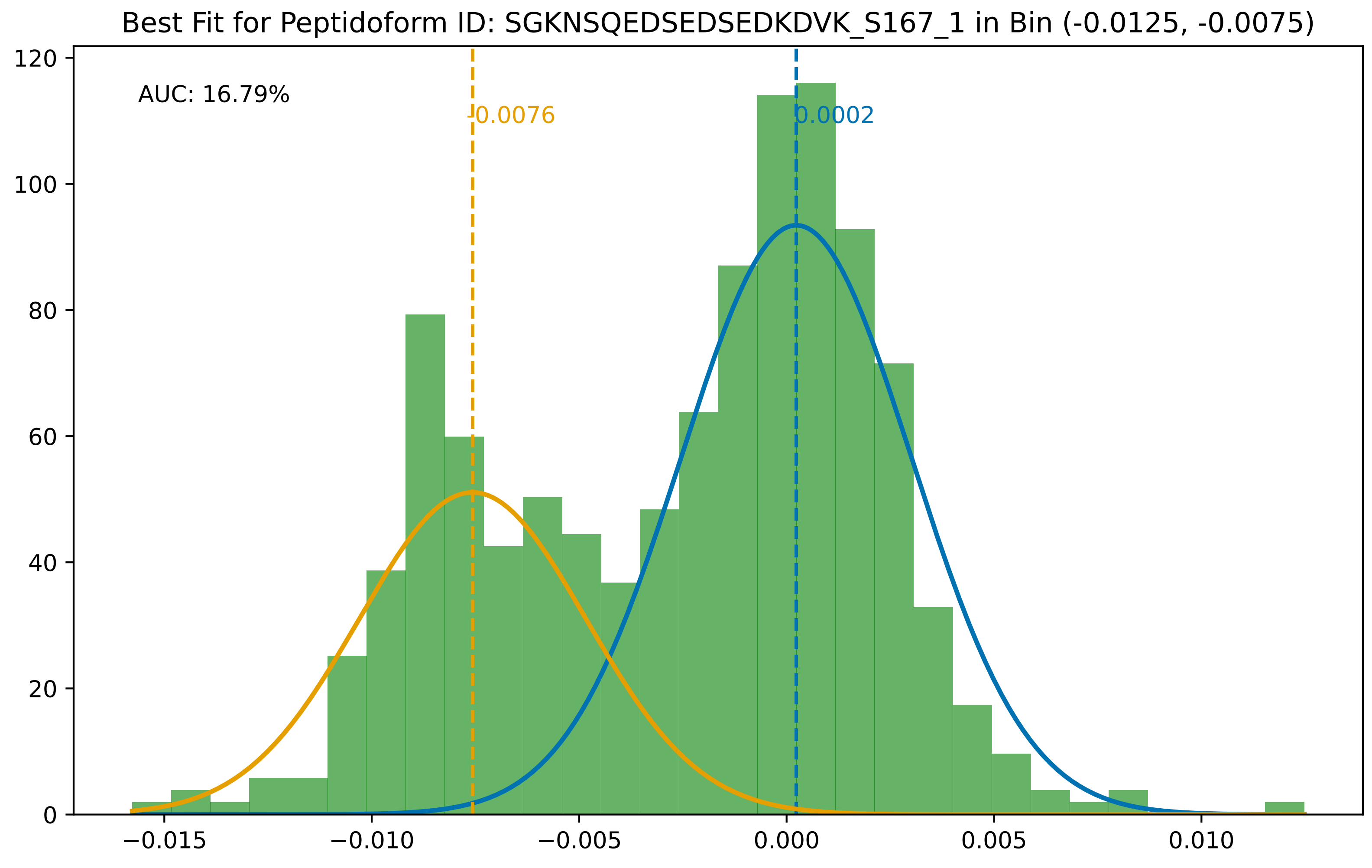

Best Fit for Peptidoform ID: SGKNSQEDSEDSEDKDVK_S167_1 in Bin (-0.0125, -0.0075)
AUC: 16.79%
-0.0076
0.0002
0
20
40
60
80
100
120
−0.015
−0.010
−0.005
0.000
0.005
0.010

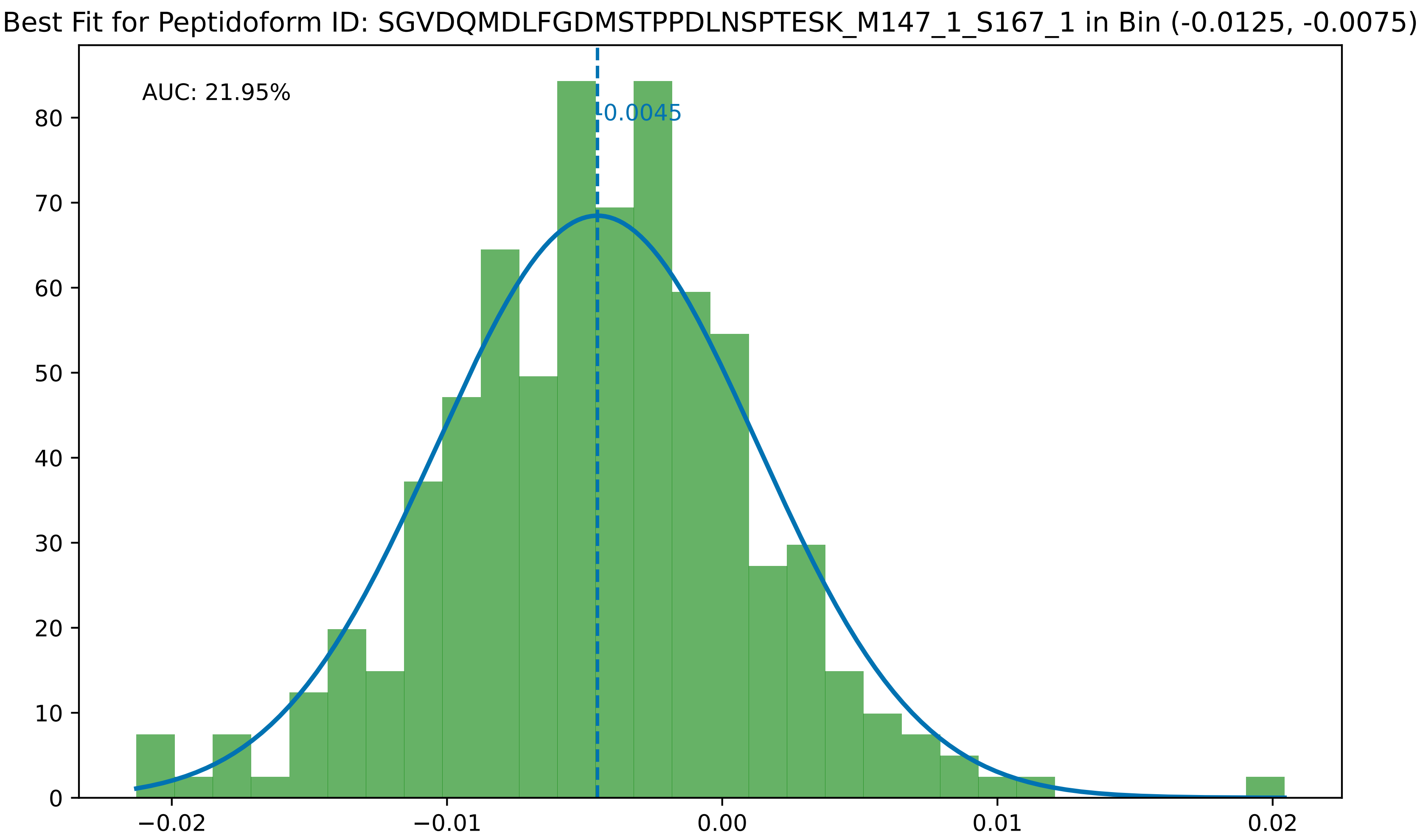

Best Fit for Peptidoform ID: SGVDQMDLFGDMSTPPDLNSPTESK_M147_1_S167_1 in Bin (-0.0125, -0.0075)
AUC: 21.95%
-0.0045
0
10
20
30
40
50
60
70
80
−0.02
−0.01
0.00
0.01
0.02

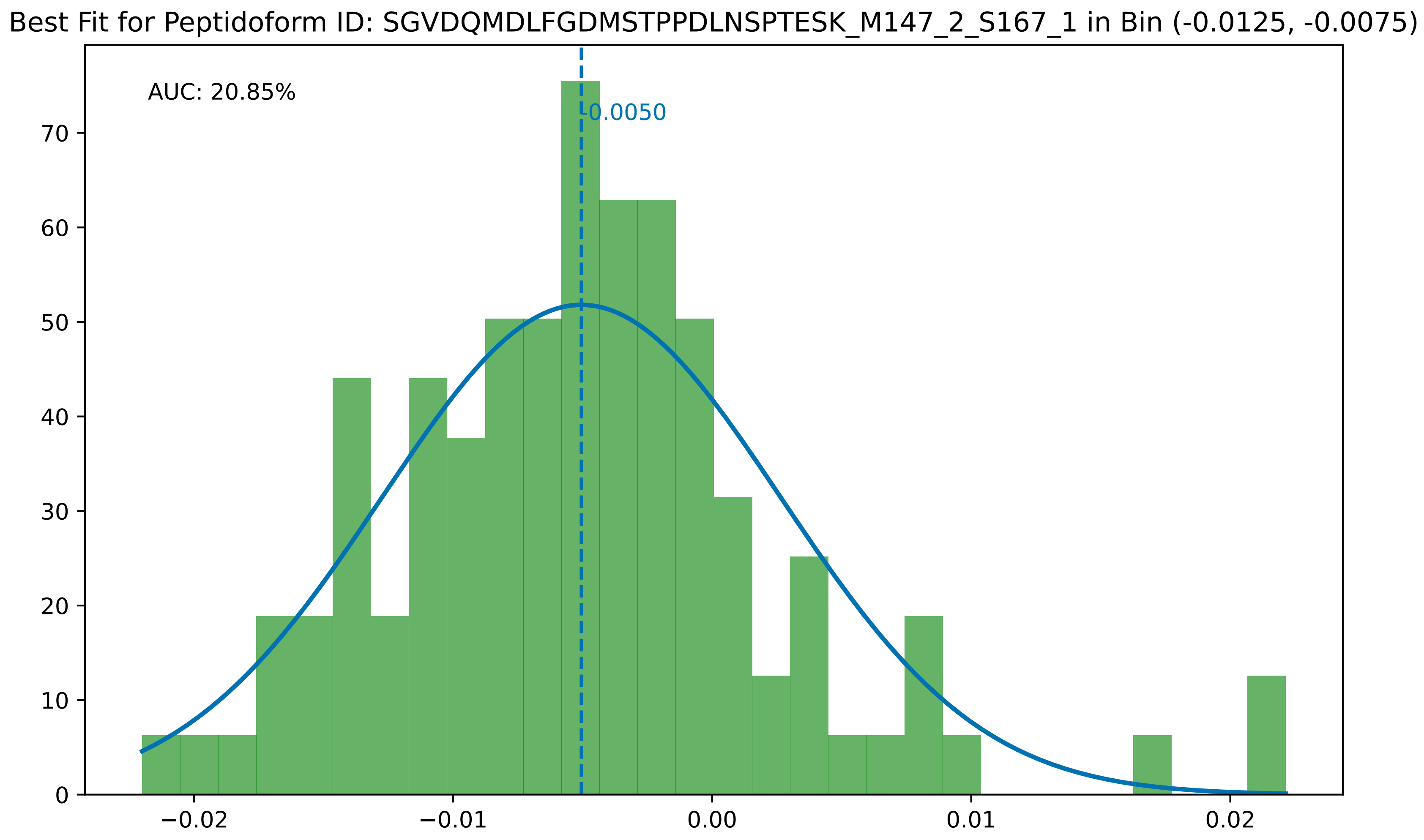

Best Fit for Peptidoform ID: SGVDQMDLFGDMSTPPDLNSPTESK_M147_2_S167_1 in Bin (-0.0125, -0.0075)
AUC: 20.85%
-0.0050
0
10
20
30
40
50
60
70
−0.02
−0.01
0.00
0.01
0.02

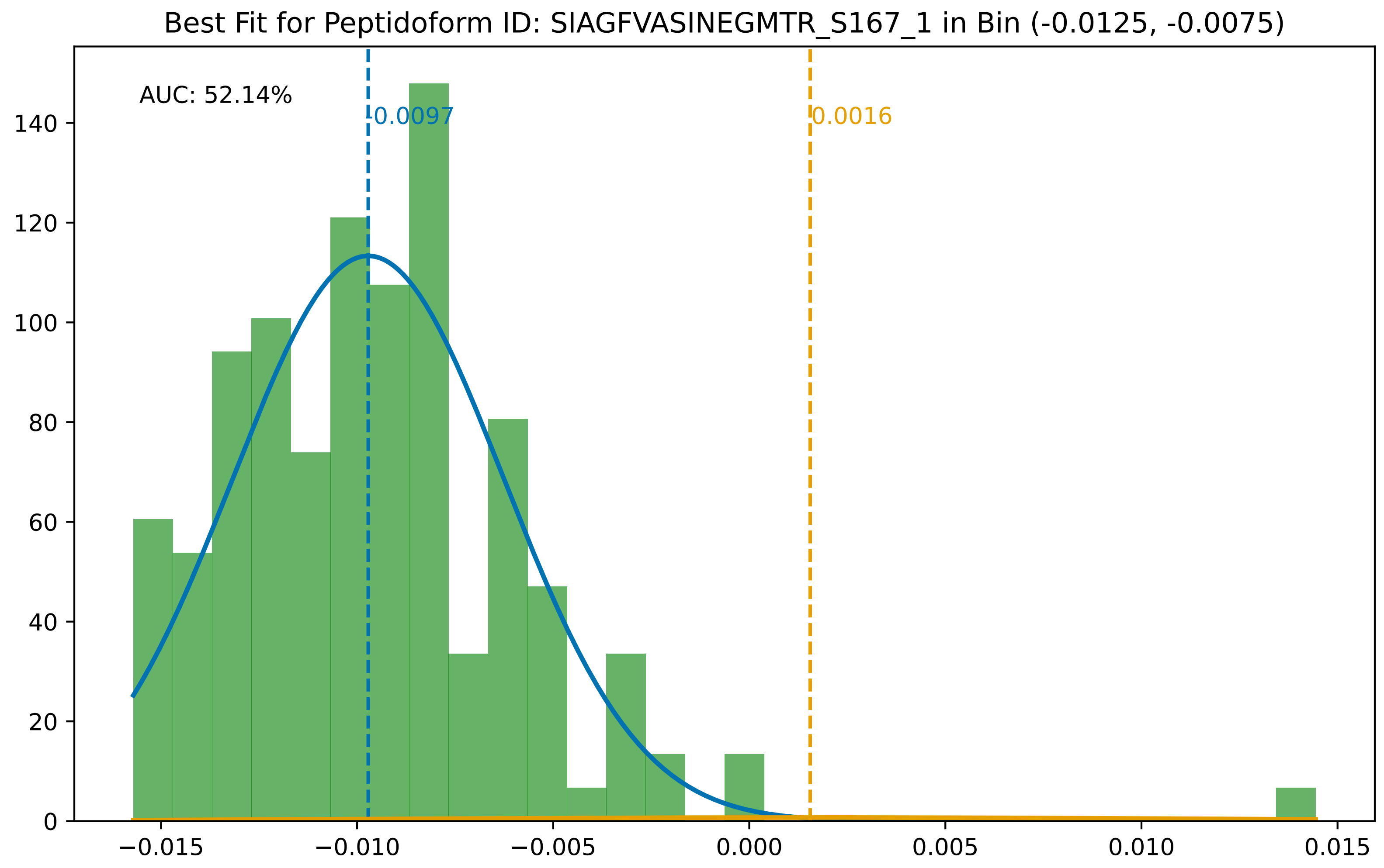

Best Fit for Peptidoform ID: SIAGFVASINEGMTR_S167_1 in Bin (-0.0125, -0.0075)
AUC: 52.14%
-0.0097
0.0016
0
20
40
60
80
100
120
140
−0.015
−0.010
−0.005
0.000
0.005
0.010
0.015

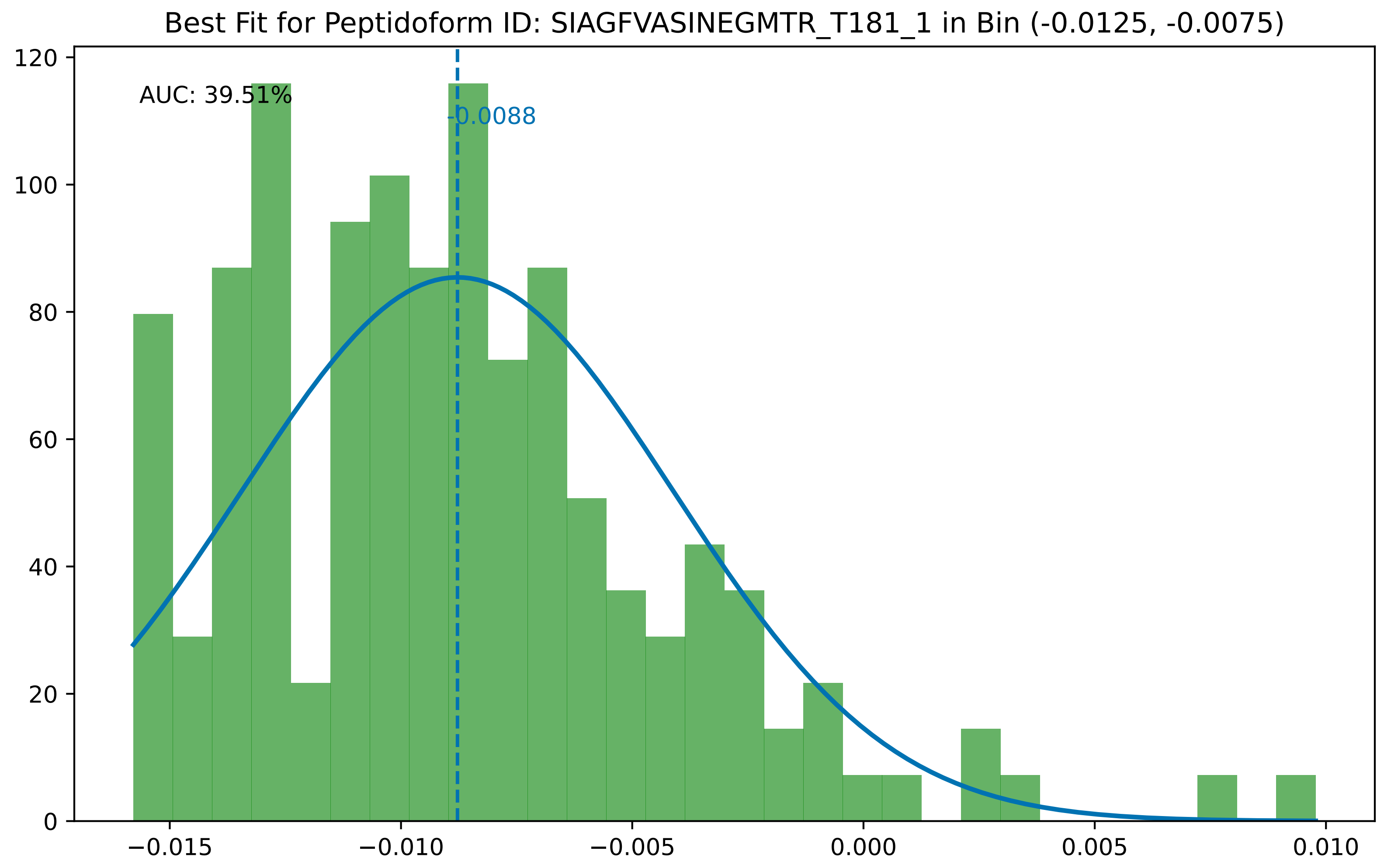
Best Fit for Peptidoform ID: SIAGFVASINEGMTR_T181_1 in Bin (-0.0125, -0.0075)
AUC: 39.51%
-0.0088
0
20
40
60
80
100
120
−0.015
−0.010
−0.005
0.000
0.005
0.010

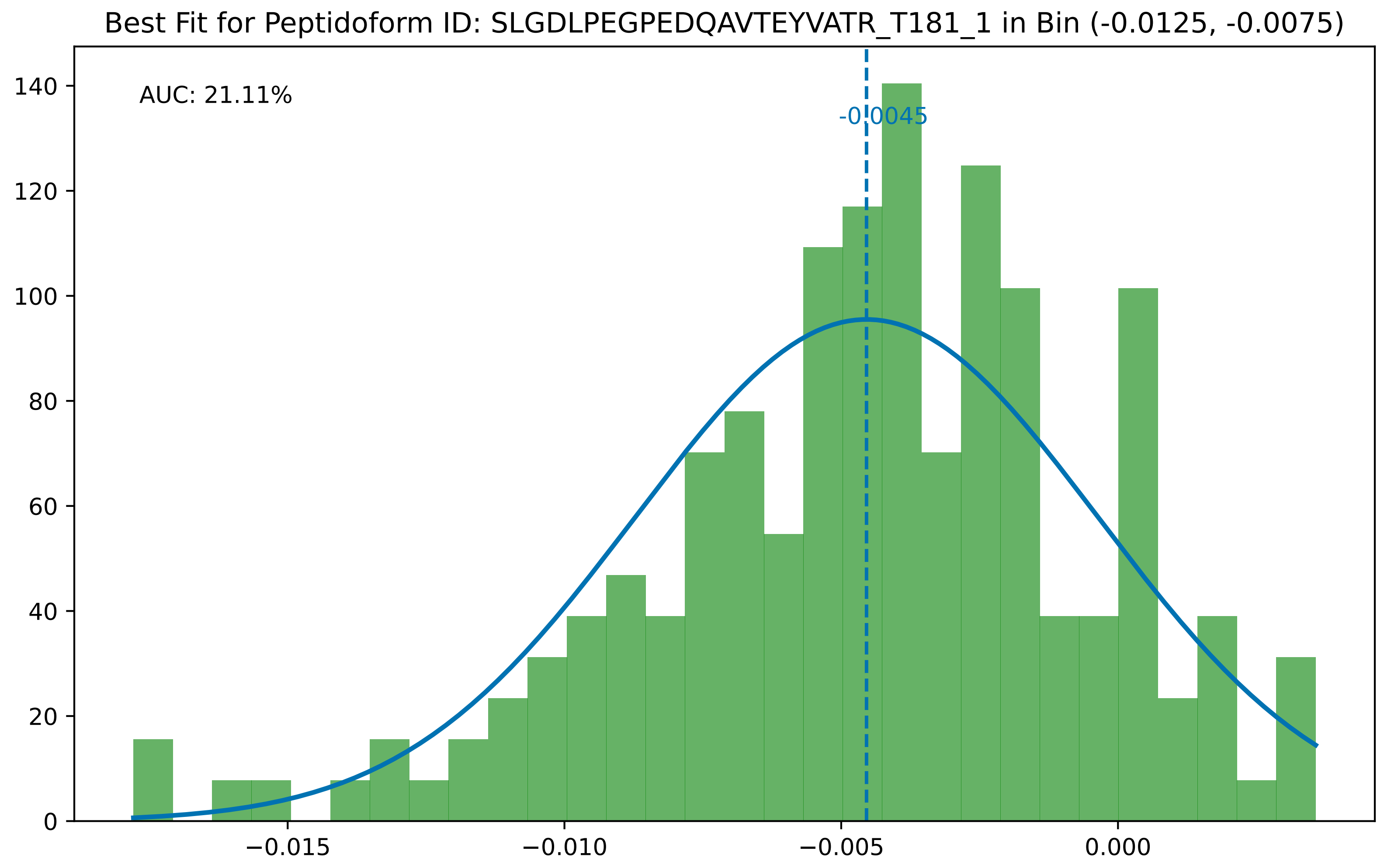
Best Fit for Peptidoform ID: SLGDLPEGPEDQAVTEYVATR_T181_1 in Bin (-0.0125, -0.0075)
AUC: 21.11%
-0.0045
0
20
40
60
80
100
120
140
−0.015
−0.010
−0.005
0.000

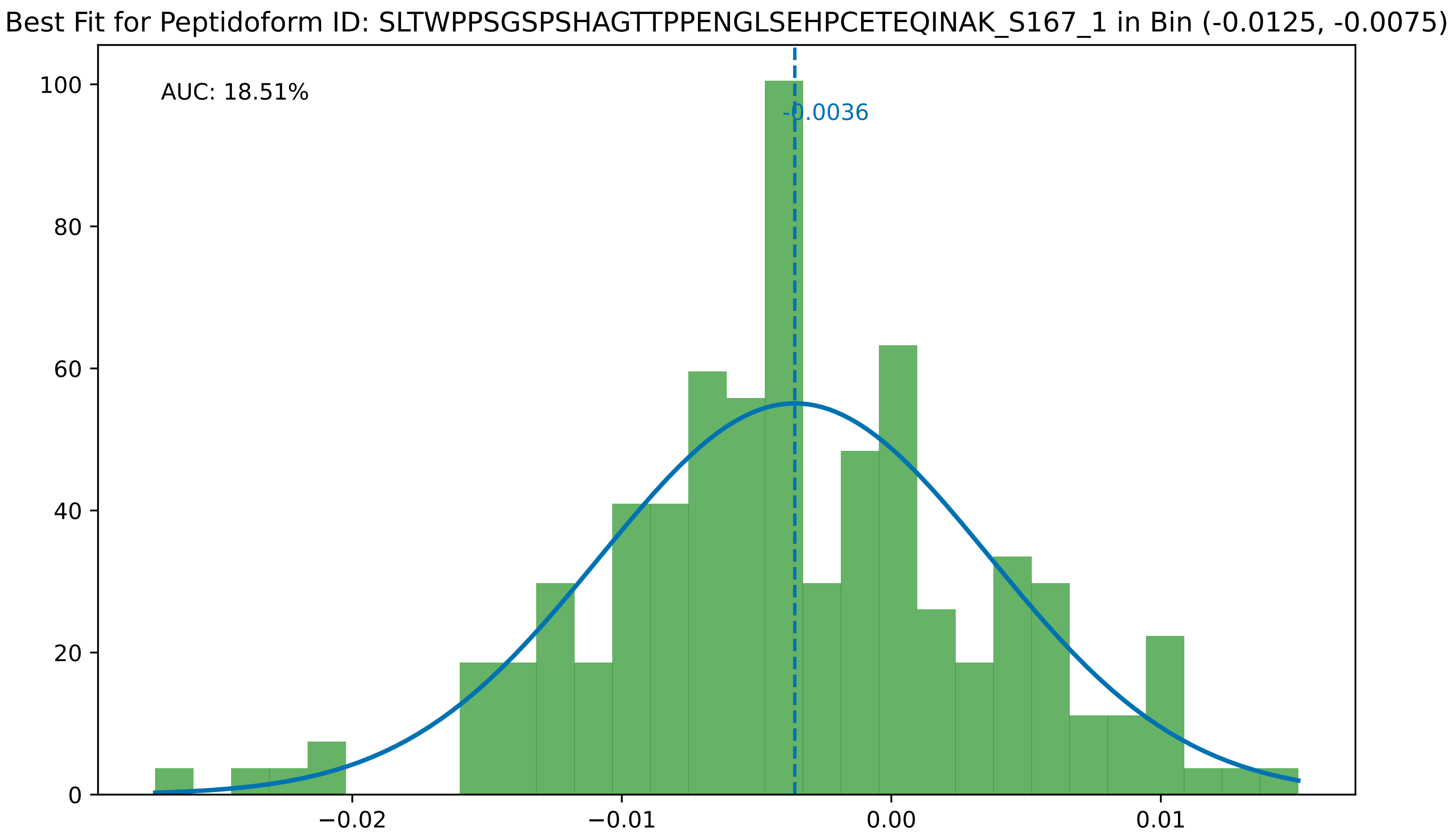

Best Fit for Peptidoform ID: SLTWPPSGSPSHAGTTPPENGLSEHPCETEQINAK_S167_1 in Bin (-0.0125, -0.0075)
AUC: 18.51%
-0.0036
100
80
60
40
20
0
−0.02
−0.01
0.00
0.01

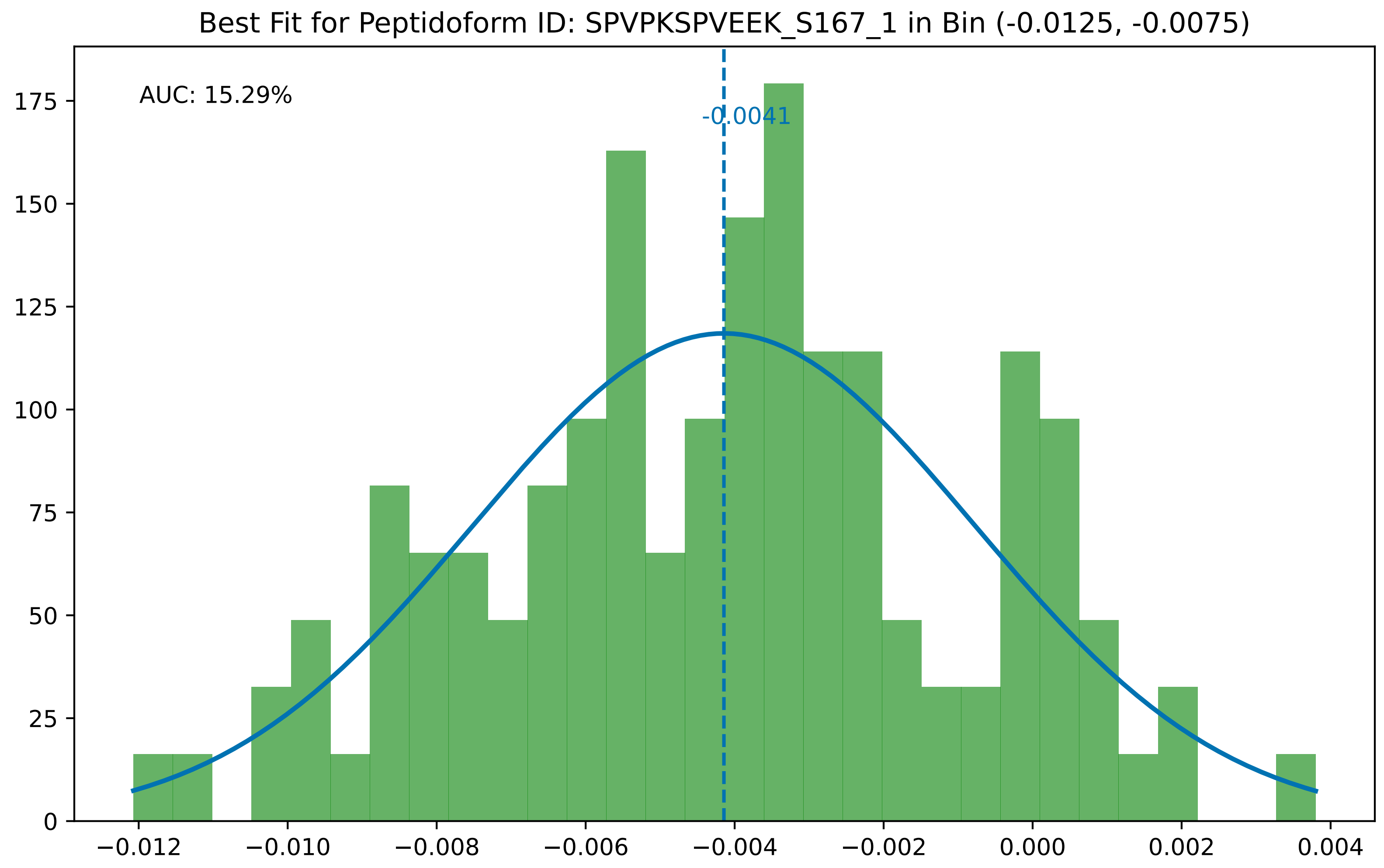

Best Fit for Peptidoform ID: SPVPKSPVEEK_S167_1 in Bin (-0.0125, -0.0075)
AUC: 15.29%
-0.0041
0
25
50
75
100
125
150
175
−0.012
−0.010
−0.008
−0.006
−0.004
−0.002
0.000
0.002
0.004

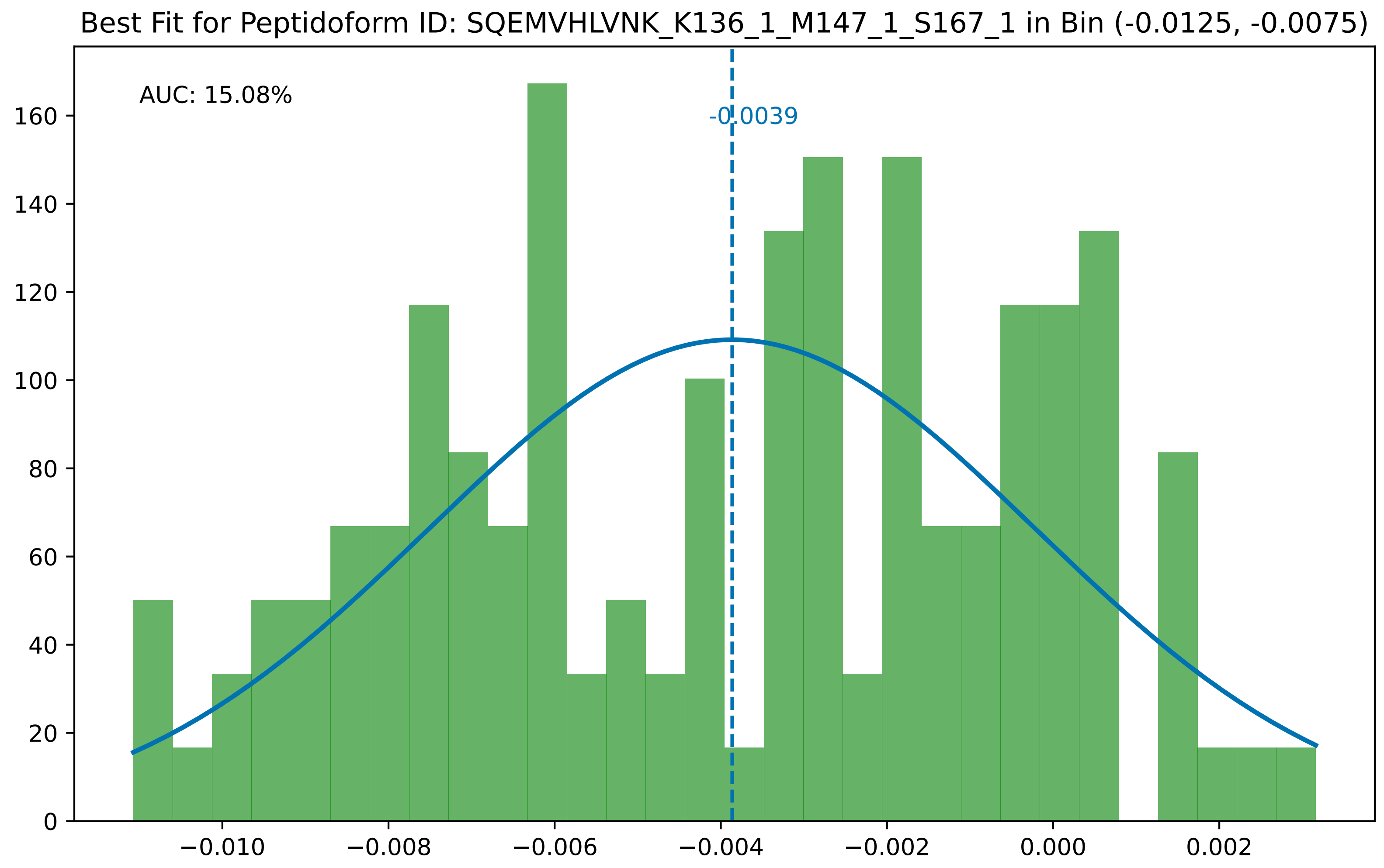

Best Fit for Peptidoform ID: SQEMVHLVNK_K136_1_M147_1_S167_1 in Bin (-0.0125, -0.0075)
AUC: 15.08%
-0.0039
0
20
40
60
80
100
120
140
160
−0.010
−0.008
−0.006
−0.004
−0.002
0.000
0.002

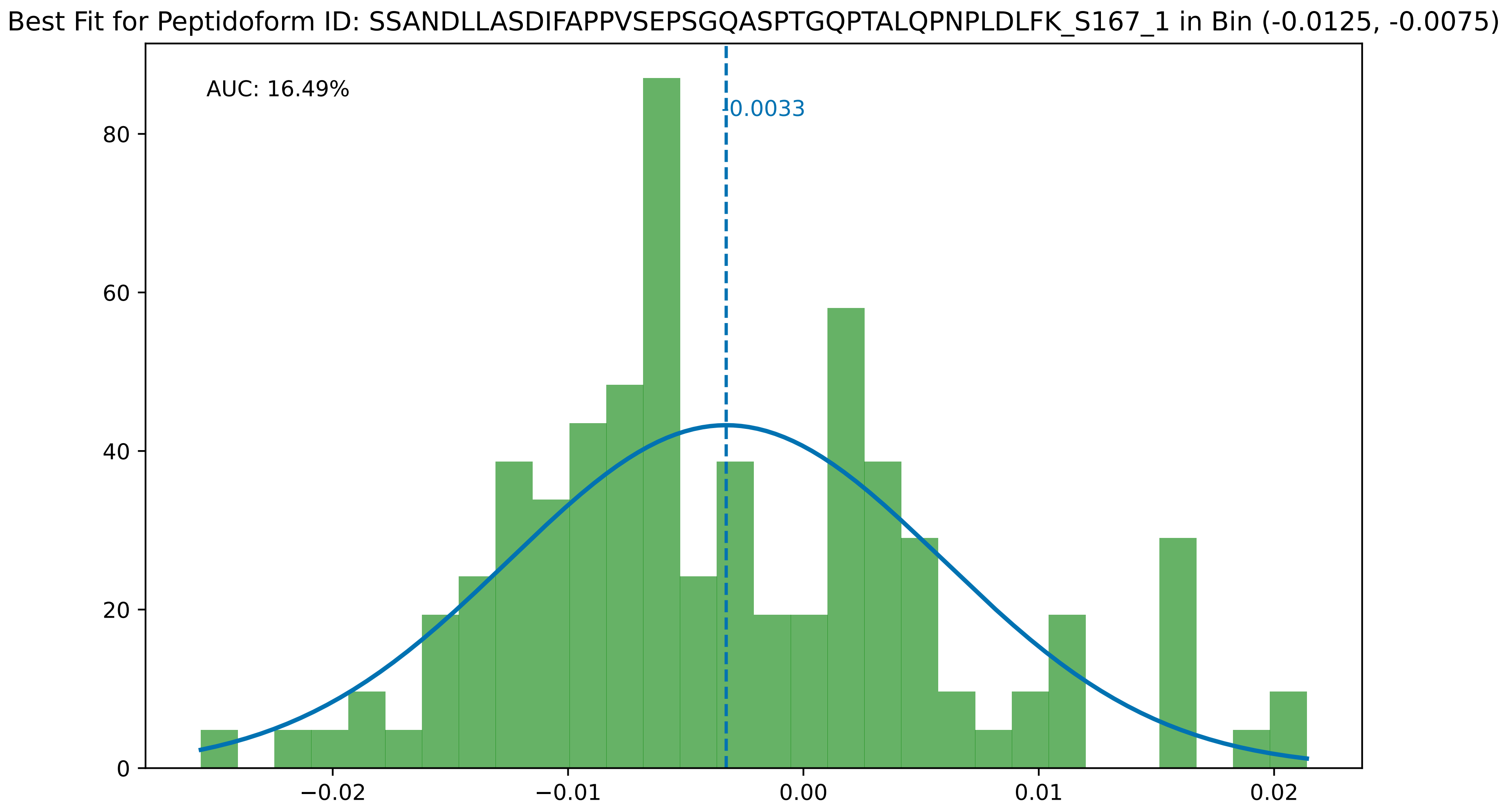
Best Fit for Peptidoform ID: SSANDLLASDIFAPPVSEPSGQASPTGQPTALQPNPLDLFK_S167_1 in Bin (-0.0125, -0.0075)
AUC: 16.49%
-0.0033
0
20
40
60
80
−0.02
−0.01
0.00
0.01
0.02

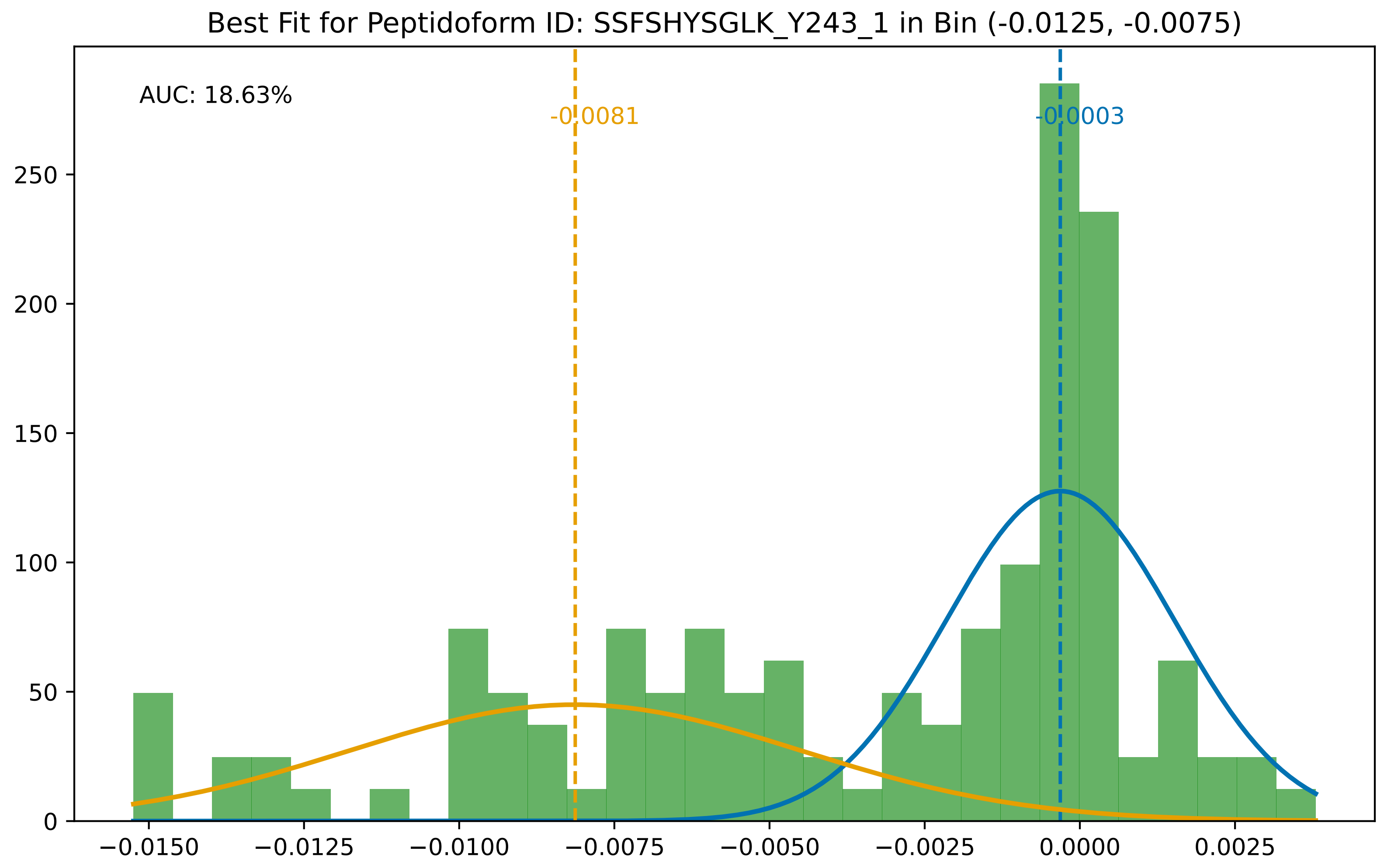

Best Fit for Peptidoform ID: SSFSHYSGLK_Y243_1 in Bin (-0.0125, -0.0075)
AUC: 18.63%
-0.0081
-0.0003
0
50
100
150
200
250
−0.0150
−0.0125
−0.0100
−0.0075
−0.0050
−0.0025
0.0000
0.0025

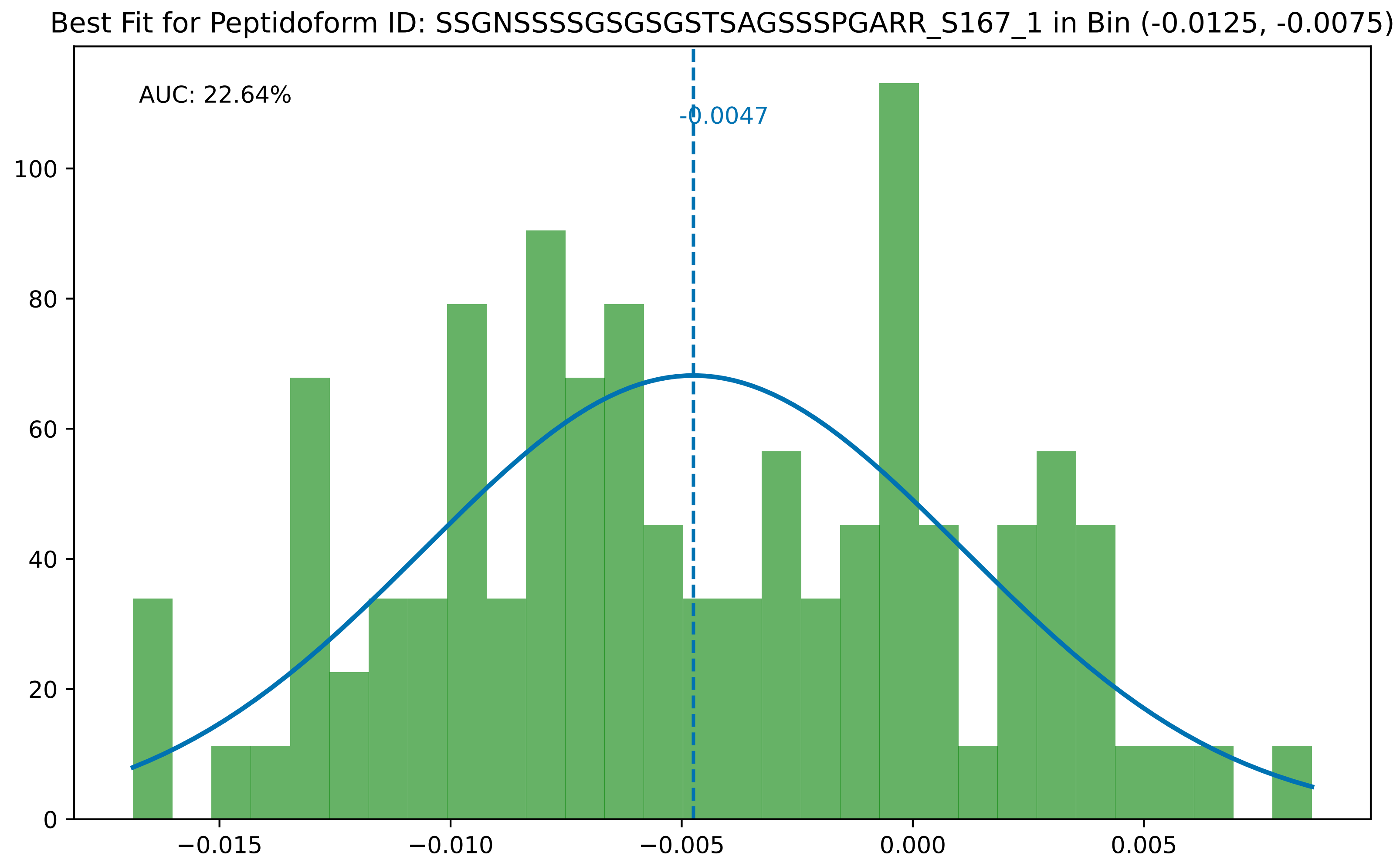
Best Fit for Peptidoform ID: SSGNSSSSGSGSGSTSAGSSSPGARR_S167_1 in Bin (-0.0125, -0.0075)
AUC: 22.64%
-0.0047
0
20
40
60
80
100
−0.015
−0.010
−0.005
0.000
0.005

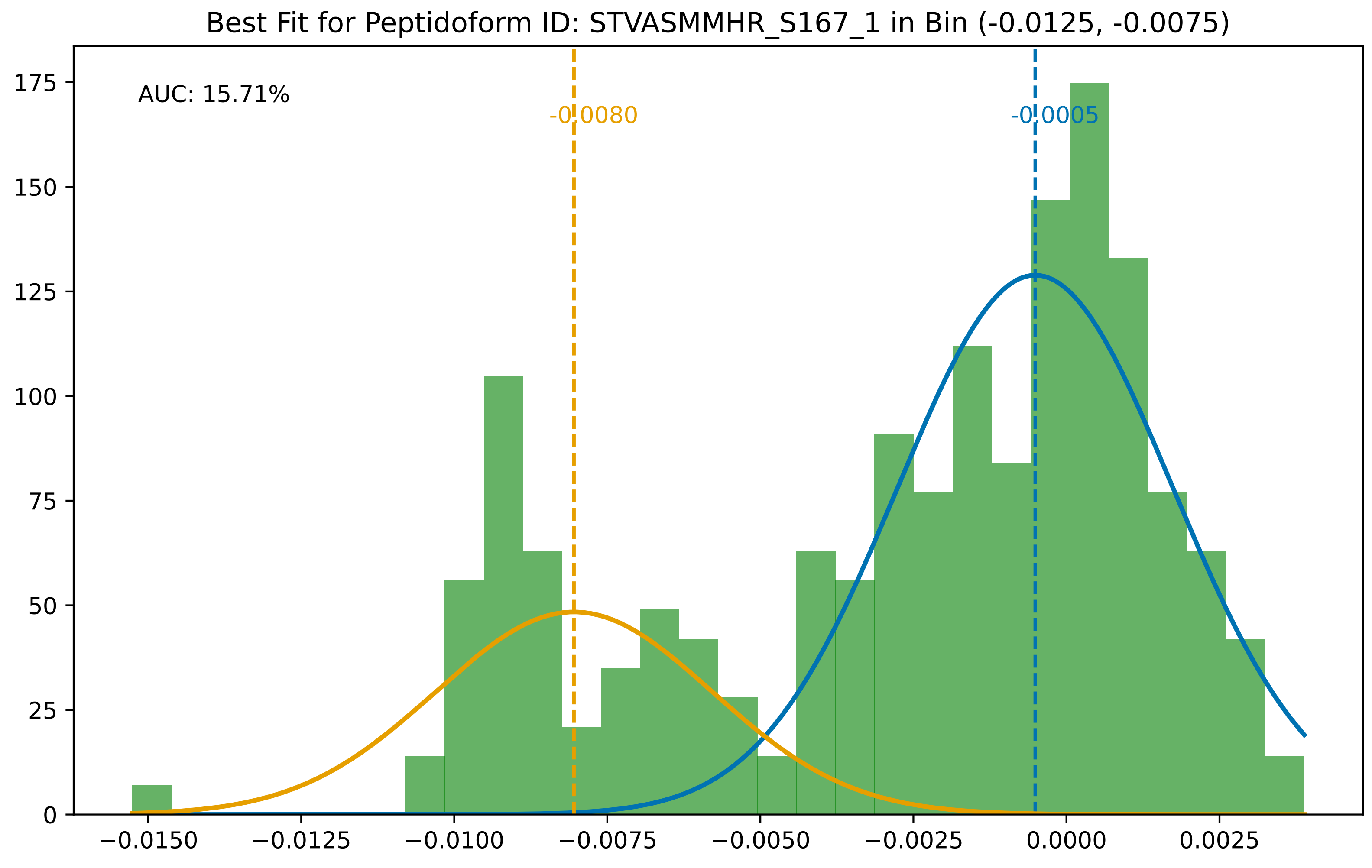

Best Fit for Peptidoform ID: STVASMMHR_S167_1 in Bin (-0.0125, -0.0075)
AUC: 15.71%
-0.0080
-0.0005
175
150
125
100
75
50
25
0
−0.0150
−0.0125
−0.0100
−0.0075
−0.0050
−0.0025
0.0000
0.0025

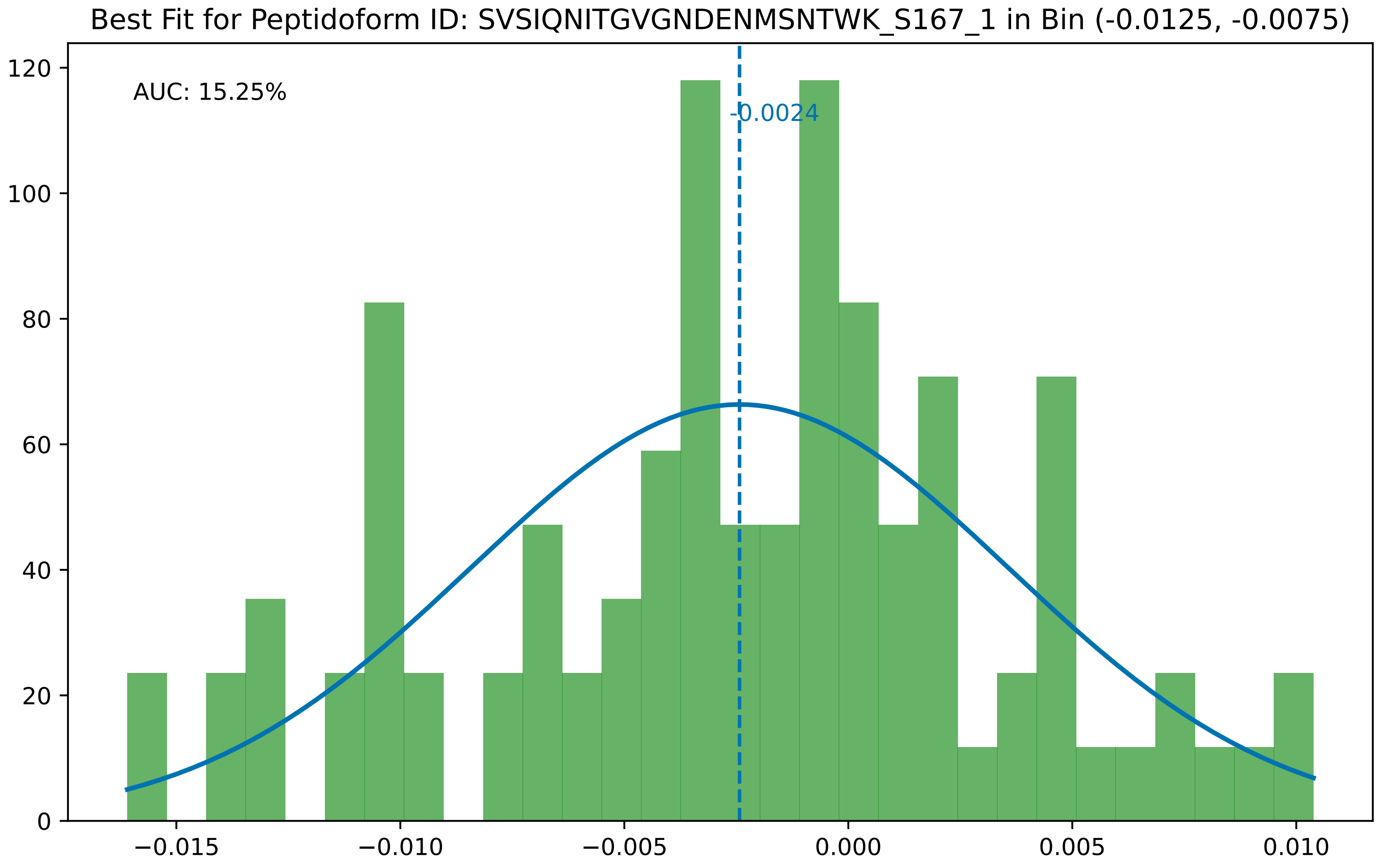

Best Fit for Peptidoform ID: SVSIQNITGVGNDENMSNTWK_S167_1 in Bin (-0.0125, -0.0075)
AUC: 15.25%
-0.0024
0
20
40
60
80
100
120
−0.015
−0.010
−0.005
0.000
0.005
0.010

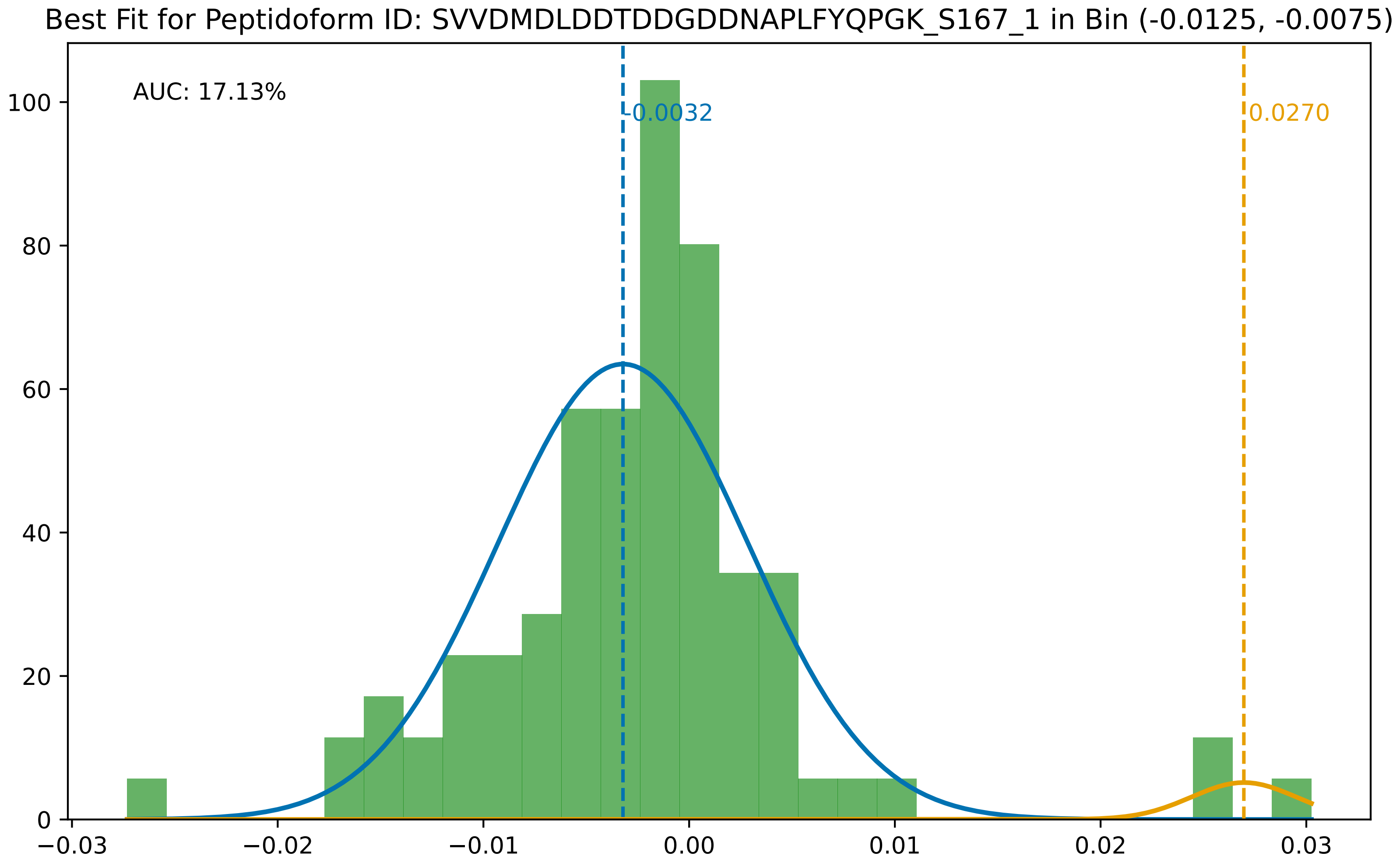

Best Fit for Peptidoform ID: SVVDMDLDDTDDGDDNAPLFYQPGK_S167_1 in Bin (-0.0125, -0.0075)
AUC: 17.13%
-0.0032
0.0270
100
80
60
40
20
0
−0.03
−0.02
−0.01
0.00
0.01
0.02
0.03

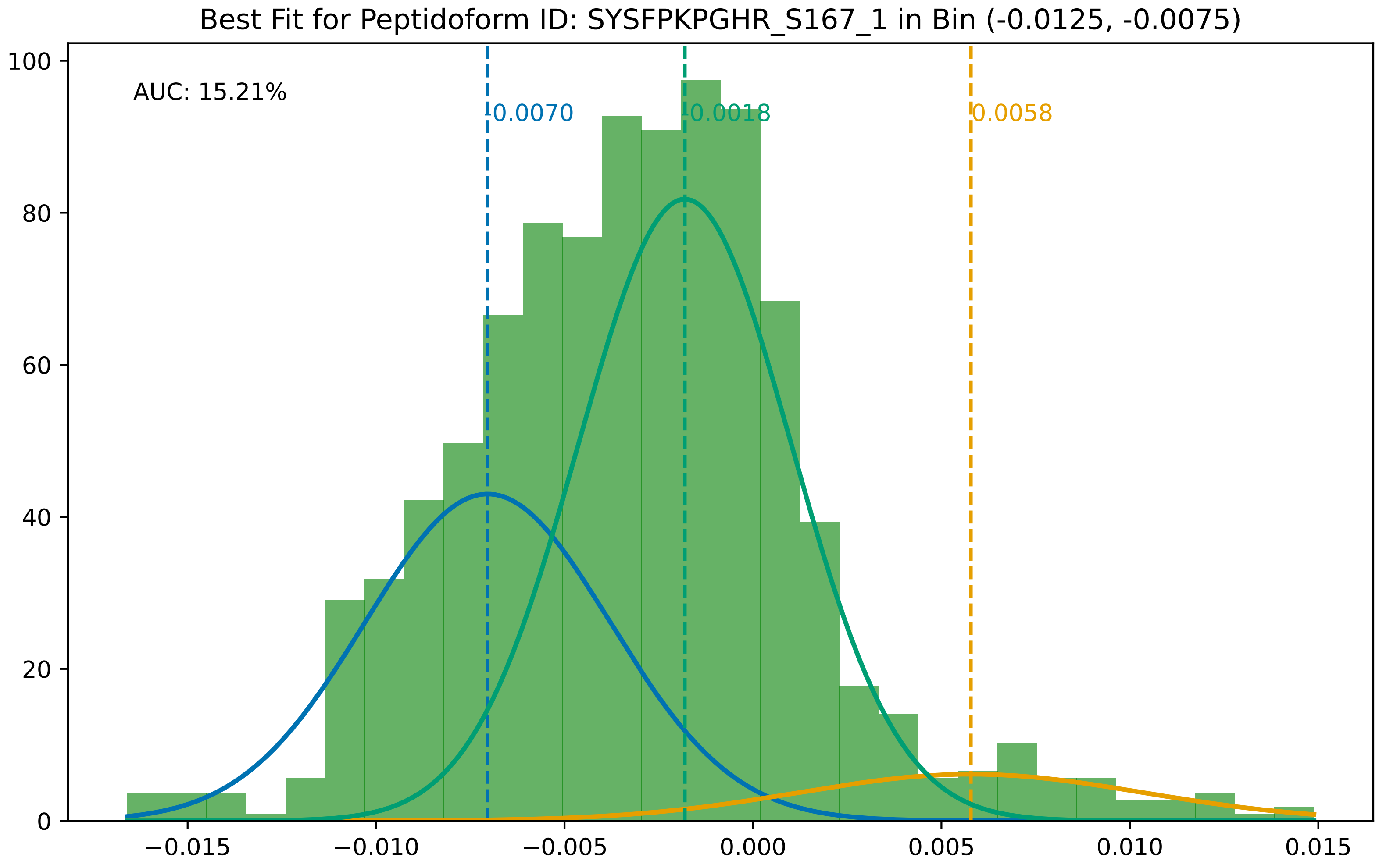
Best Fit for Peptidoform ID: SYSFPKPGHR_S167_1 in Bin (-0.0125, -0.0075)
AUC: 15.21%
-0.0070
-0.0018
0.0058
100
80
60
40
20
0
−0.015
−0.010
−0.005
0.000
0.005
0.010
0.015

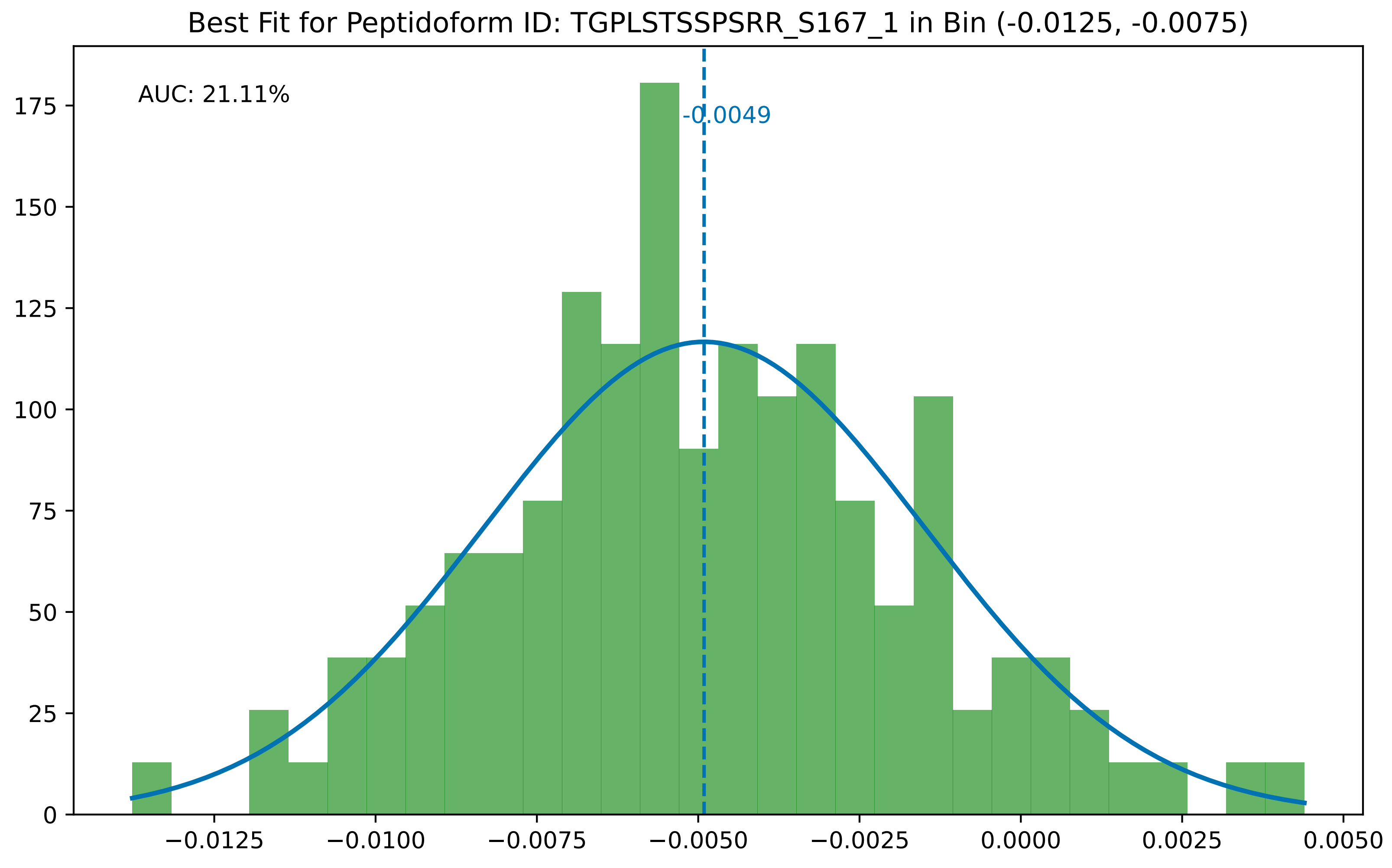

Best Fit for Peptidoform ID: TGPLSTSSPSRR_S167_1 in Bin (-0.0125, -0.0075)
AUC: 21.11%
-0.0049
0
25
50
75
100
125
150
175
−0.0125
−0.0100
−0.0075
−0.0050
−0.0025
0.0000
0.0025
0.0050

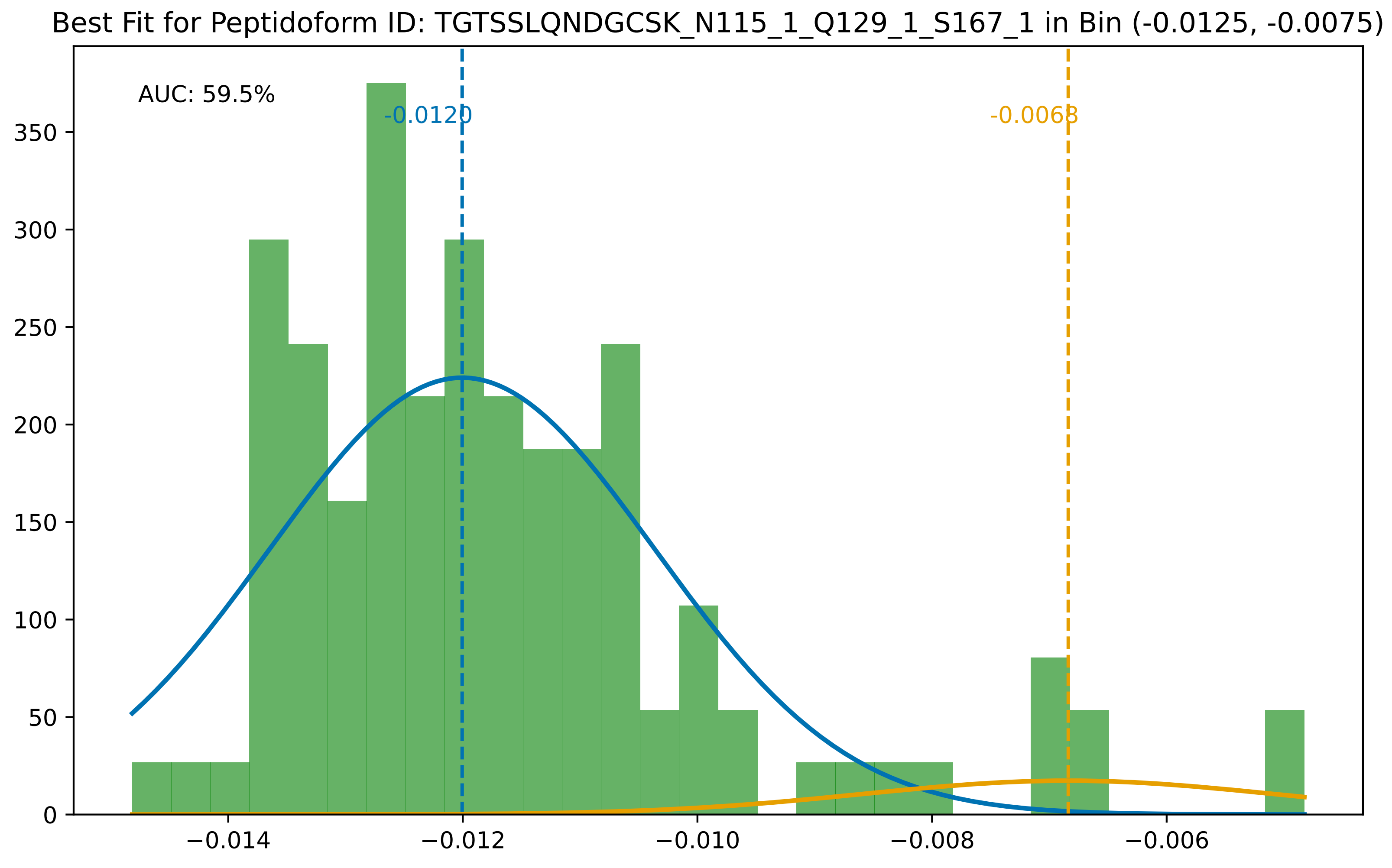

Best Fit for Peptidoform ID: TGTSSLQNDGCSK_N115_1_Q129_1_S167_1 in Bin (-0.0125, -0.0075)
AUC: 59.5%
-0.0120
-0.0068
0
50
100
150
200
250
300
350
−0.014
−0.012
−0.010
−0.008
−0.006

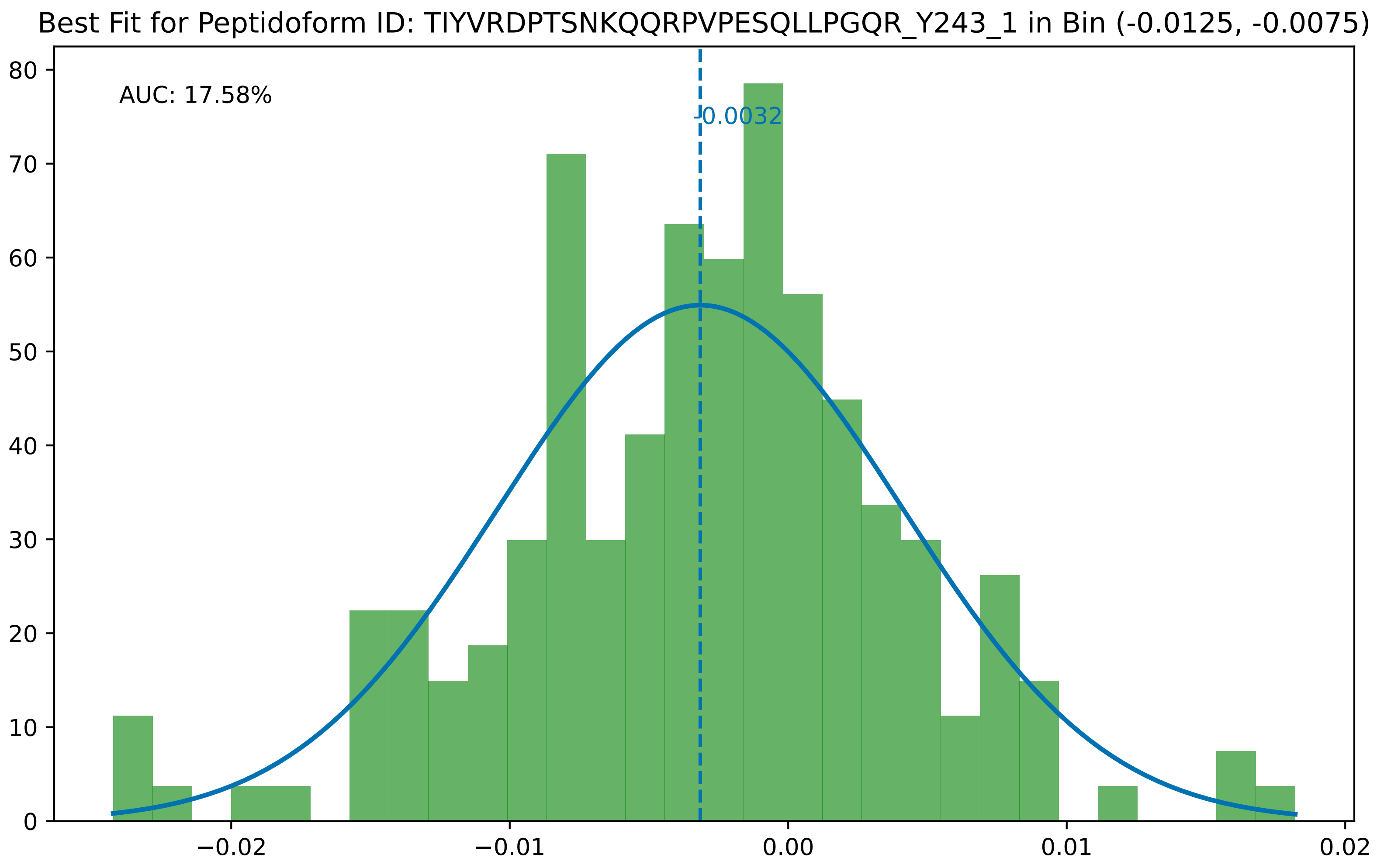

Best Fit for Peptidoform ID: TIYVRDPTSNKQQRPVPESQLLPGQR_Y243_1 in Bin (-0.0125, -0.0075)
AUC: 17.58%
-0.0032
80
70
60
50
40
30
20
10
0
−0.02
−0.01
0.00
0.01
0.02

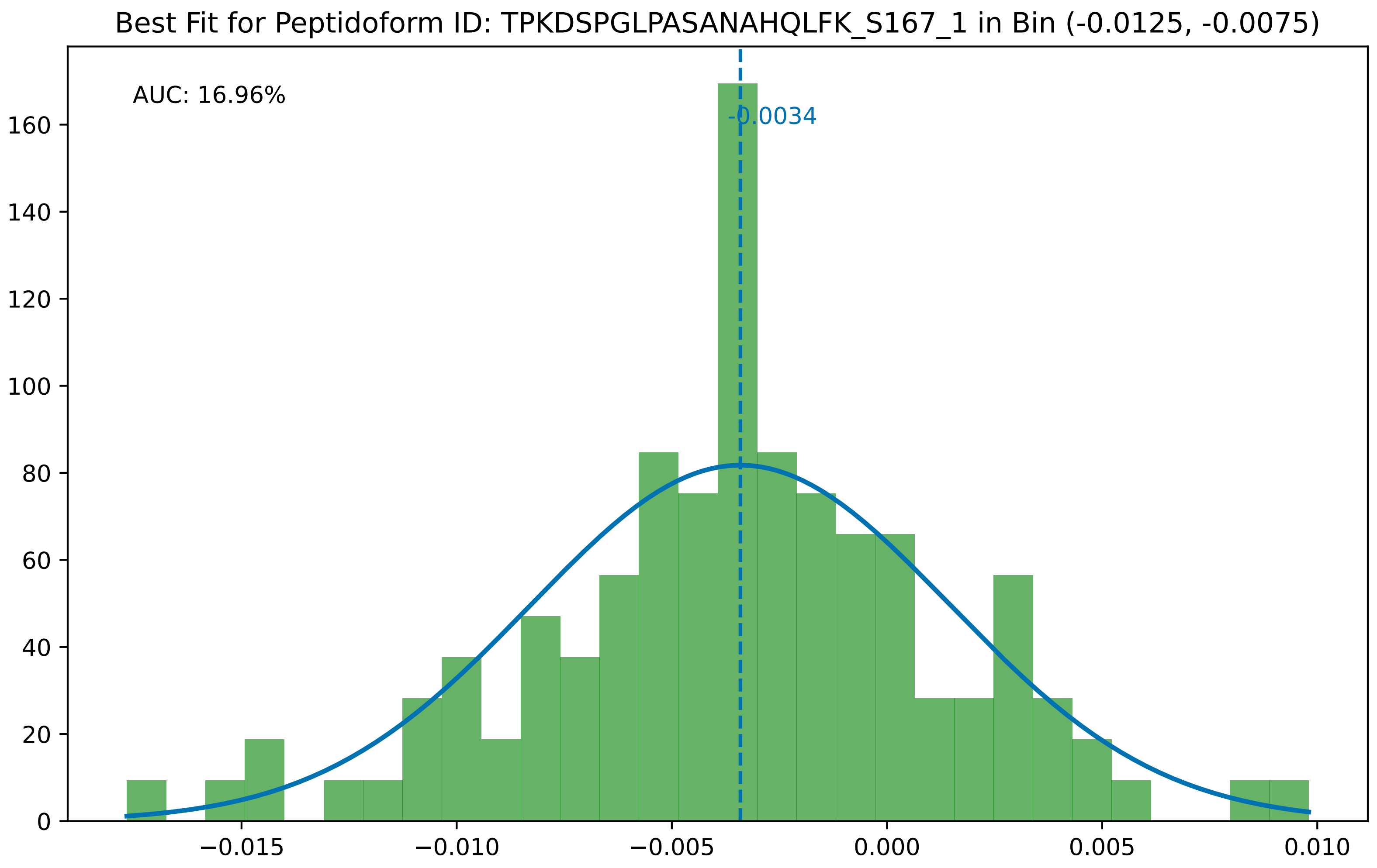

Best Fit for Peptidoform ID: TPKDSPGLPASANAHQLFK_S167_1 in Bin (-0.0125, -0.0075)
AUC: 16.96%
-0.0034
0
20
40
60
80
100
120
140
160
−0.015
−0.010
−0.005
0.000
0.005
0.010

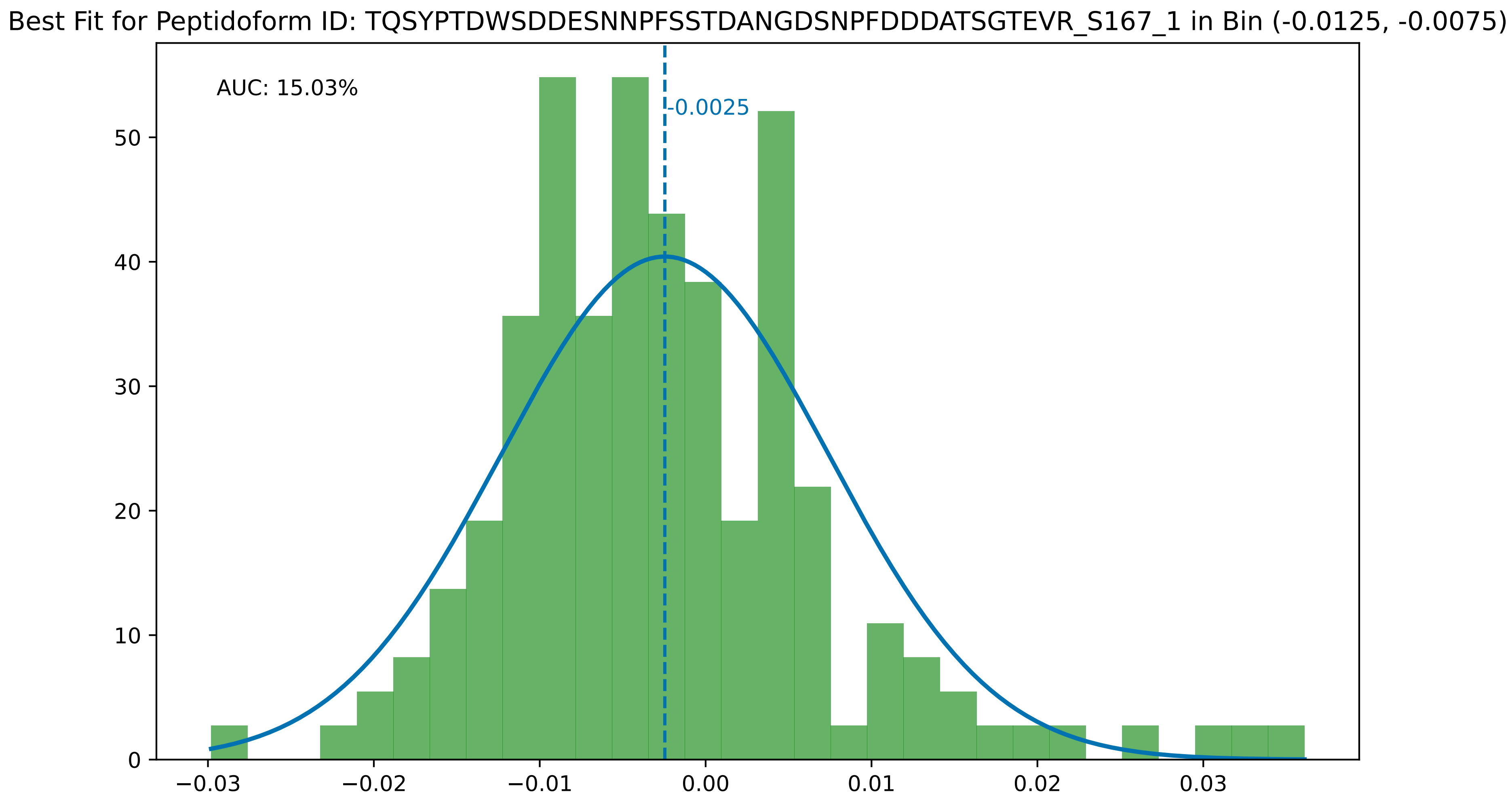

Best Fit for Peptidoform ID: TQSYPTDWSDDESNNPFSSTDANGDSNPFDDDATSGTEVR_S167_1 in Bin (-0.0125, -0.0075)
AUC: 15.03%
-0.0025
0
10
20
30
40
50
−0.03
−0.02
−0.01
0.00
0.01
0.02
0.03

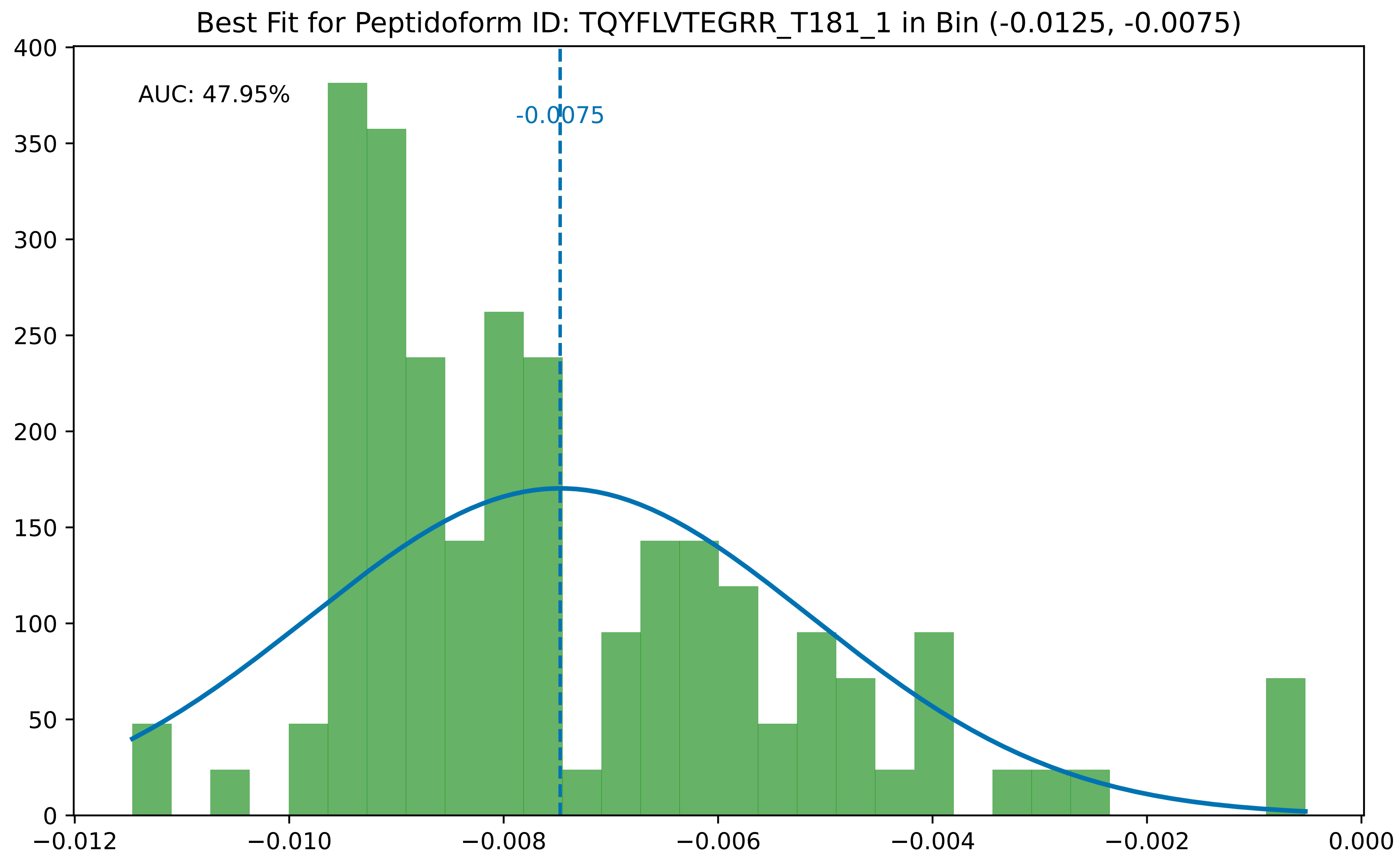

Best Fit for Peptidoform ID: TQYFLVTEGRR_T181_1 in Bin (-0.0125, -0.0075)
AUC: 47.95%
-0.0075
400
350
300
250
200
150
100
50
0
−0.012
−0.010
−0.008
−0.006
−0.004
−0.002
0.000

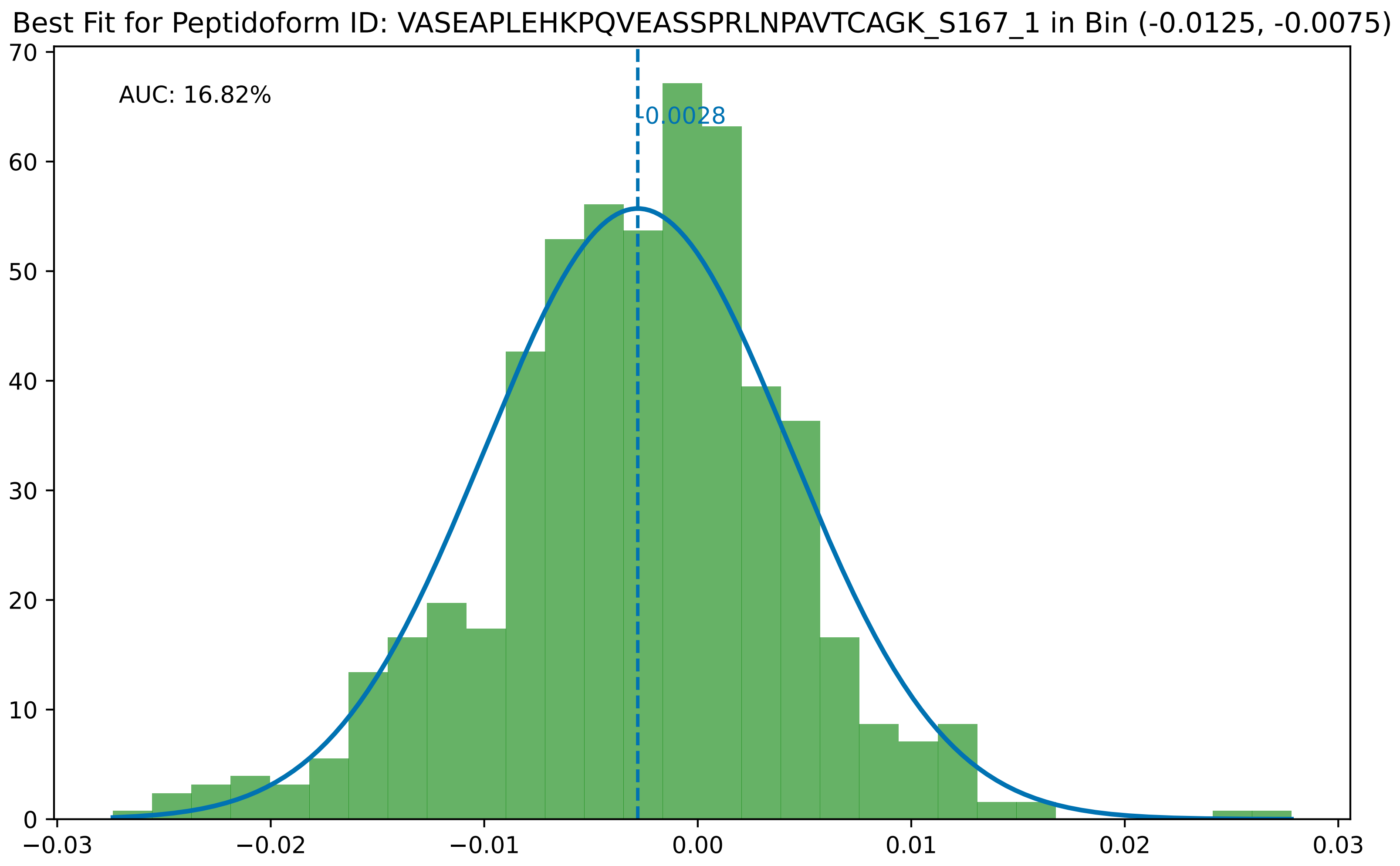
Best Fit for Peptidoform ID: VASEAPLEHKPQVEASSPRLNPAVTCAGK_S167_1 in Bin (-0.0125, -0.0075)
AUC: 16.82%
-0.0028
0
10
20
30
40
50
60
70
−0.03
−0.02
−0.01
0.00
0.01
0.02
0.03

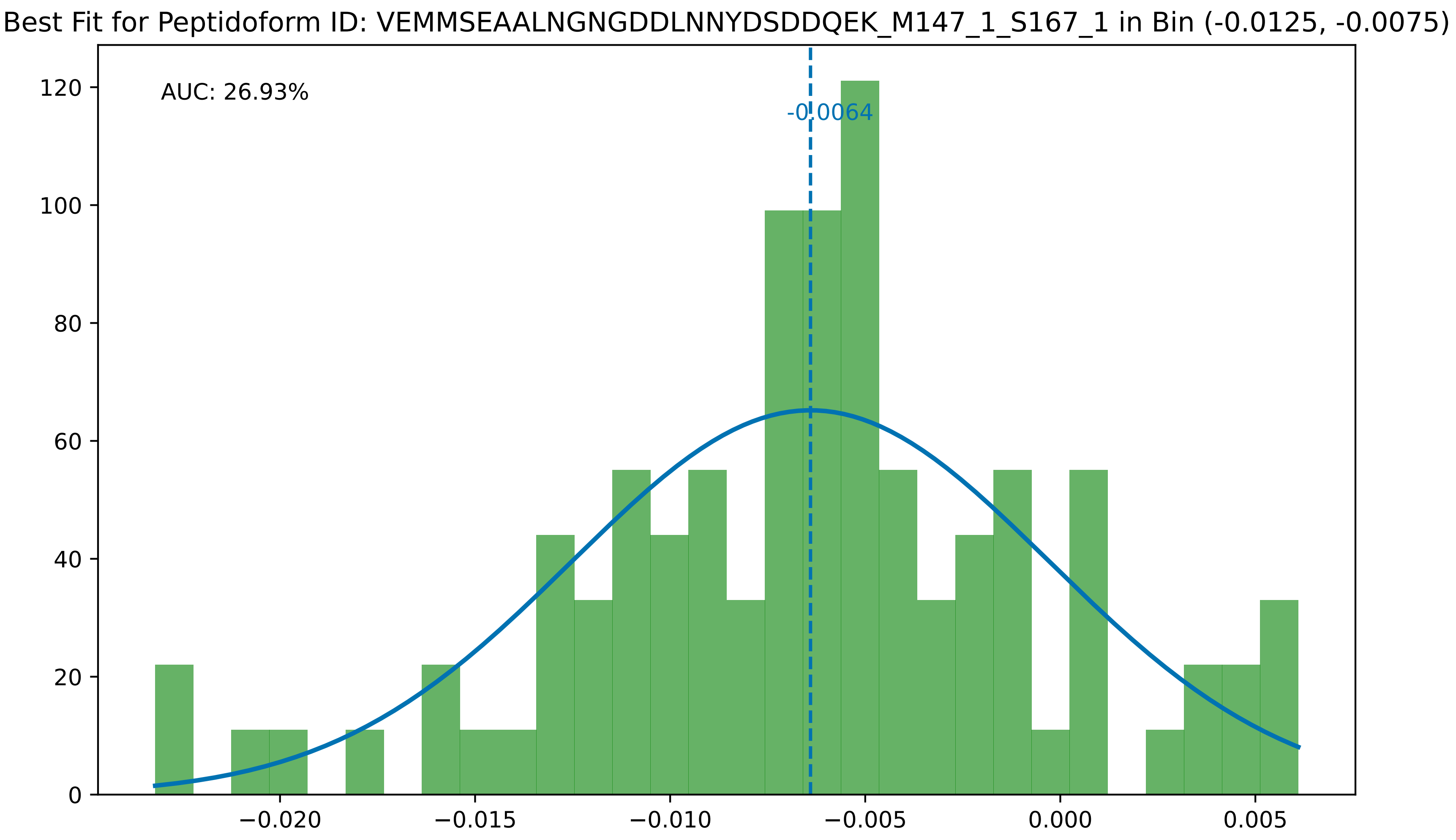
Best Fit for Peptidoform ID: VEMMSEAALNGNGDDLNNYDSDDQEK_M147_1_S167_1 in Bin (-0.0125, -0.0075)
AUC: 26.93%
-0.0064
0
20
40
60
80
100
120
−0.020
−0.015
−0.010
−0.005
0.000
0.005

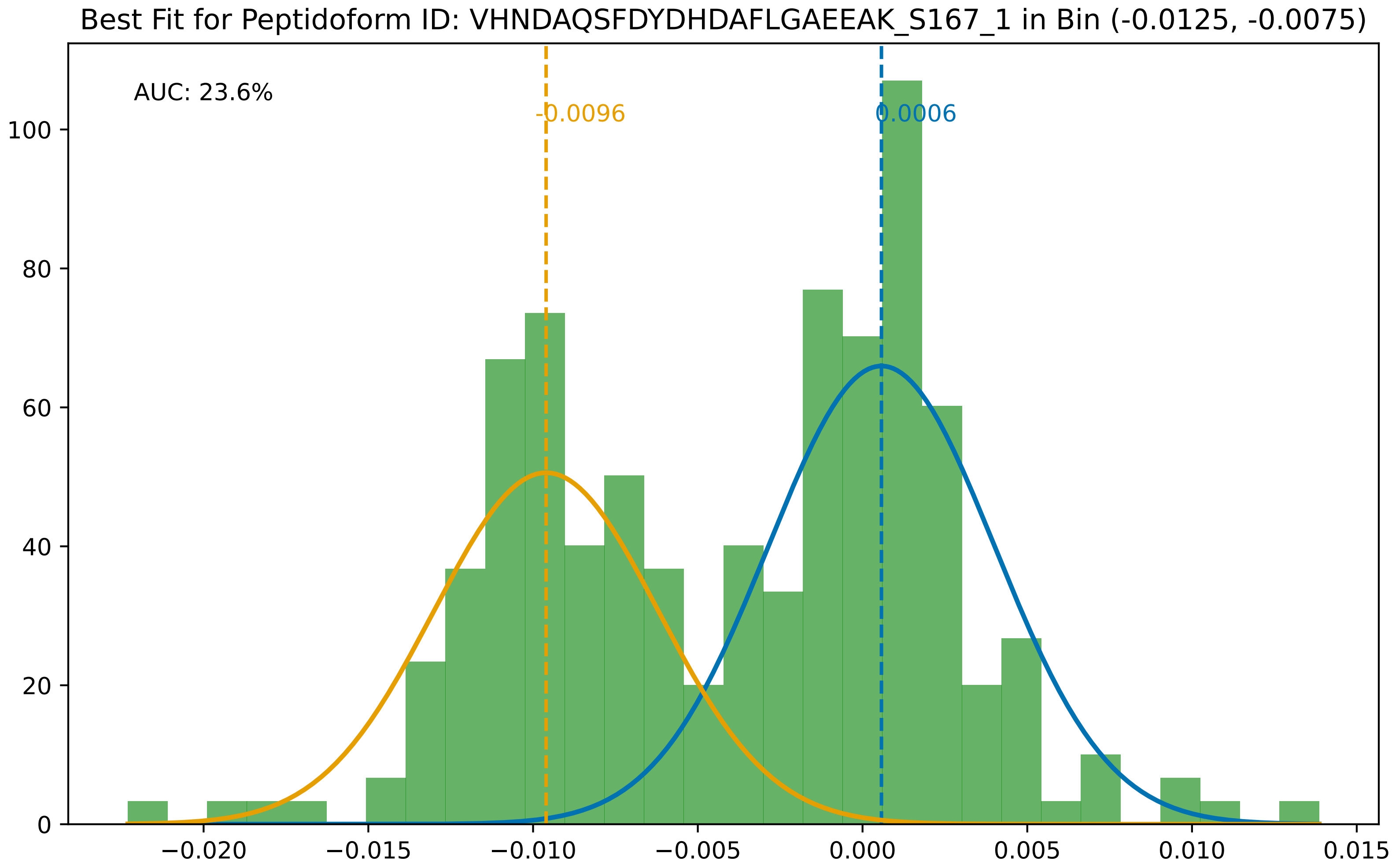

Best Fit for Peptidoform ID: VHNDAQSFDYDHDAFLGAEEAK_S167_1 in Bin (-0.0125, -0.0075)
AUC: 23.6%
-0.0096
0.0006
0
20
40
60
80
100
−0.020
−0.015
−0.010
−0.005
0.000
0.005
0.010
0.015

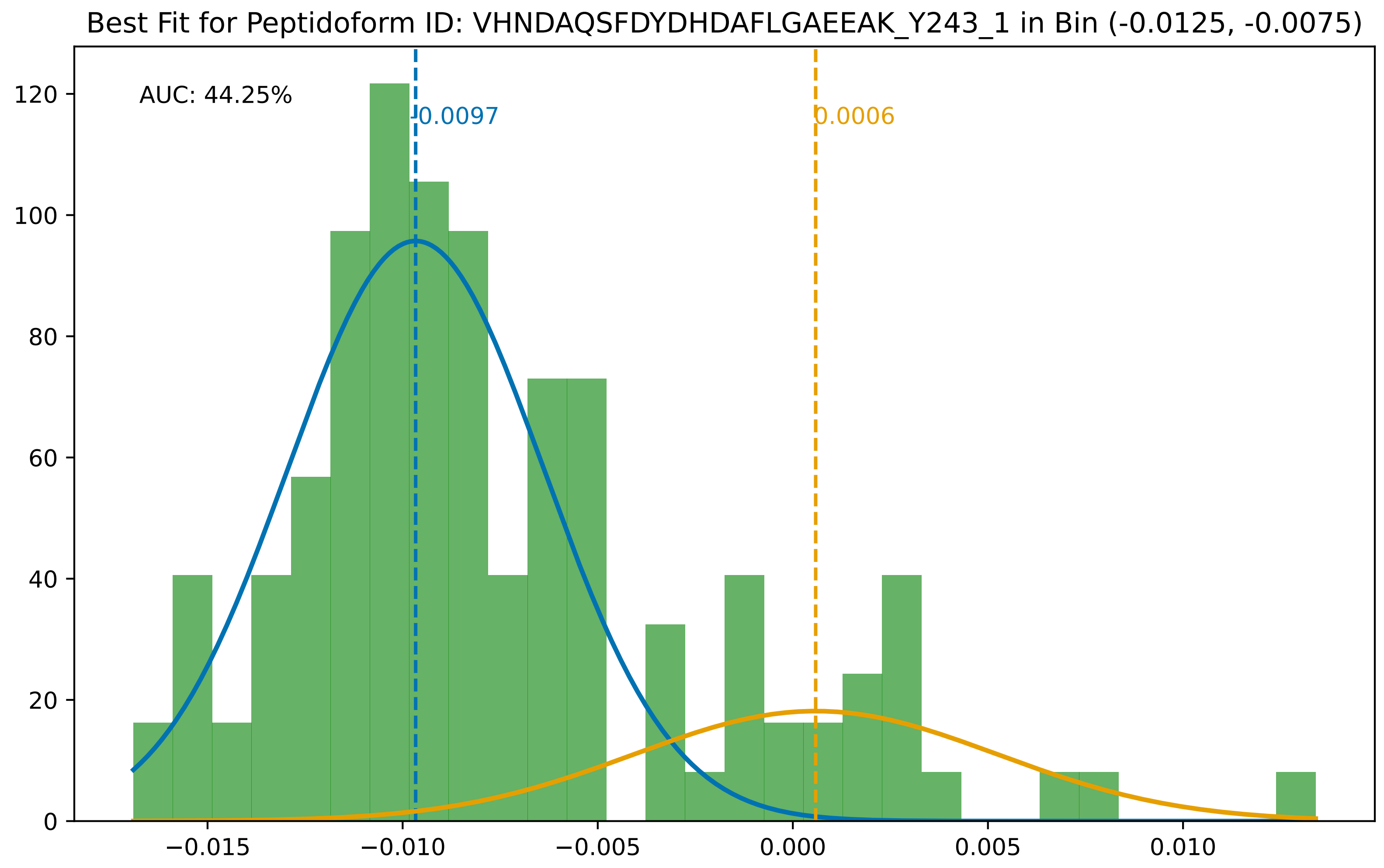
Best Fit for Peptidoform ID: VHNDAQSFDYDHDAFLGAEEAK_Y243_1 in Bin (-0.0125, -0.0075)
AUC: 44.25%
-0.0097
0.0006
0
20
40
60
80
100
120
−0.015
−0.010
−0.005
0.000
0.005
0.010

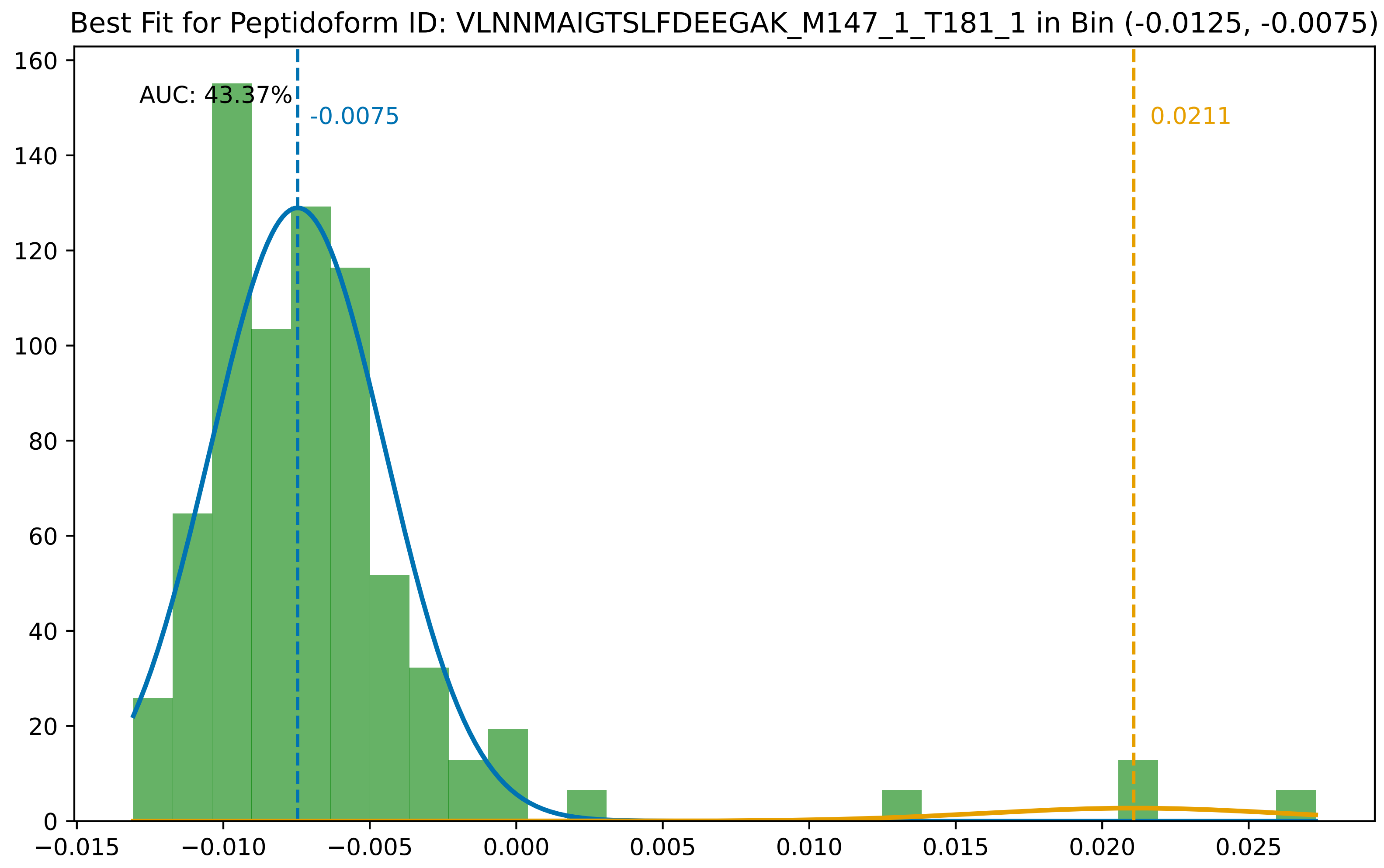

Best Fit for Peptidoform ID: VLNNMAIGTSLFDEEGAK_M147_1_T181_1 in Bin (-0.0125, -0.0075)
AUC: 43.37%
-0.0075
0.0211
0
20
40
60
80
100
120
140
160
−0.015
−0.010
−0.005
0.000
0.005
0.010
0.015
0.020
0.025

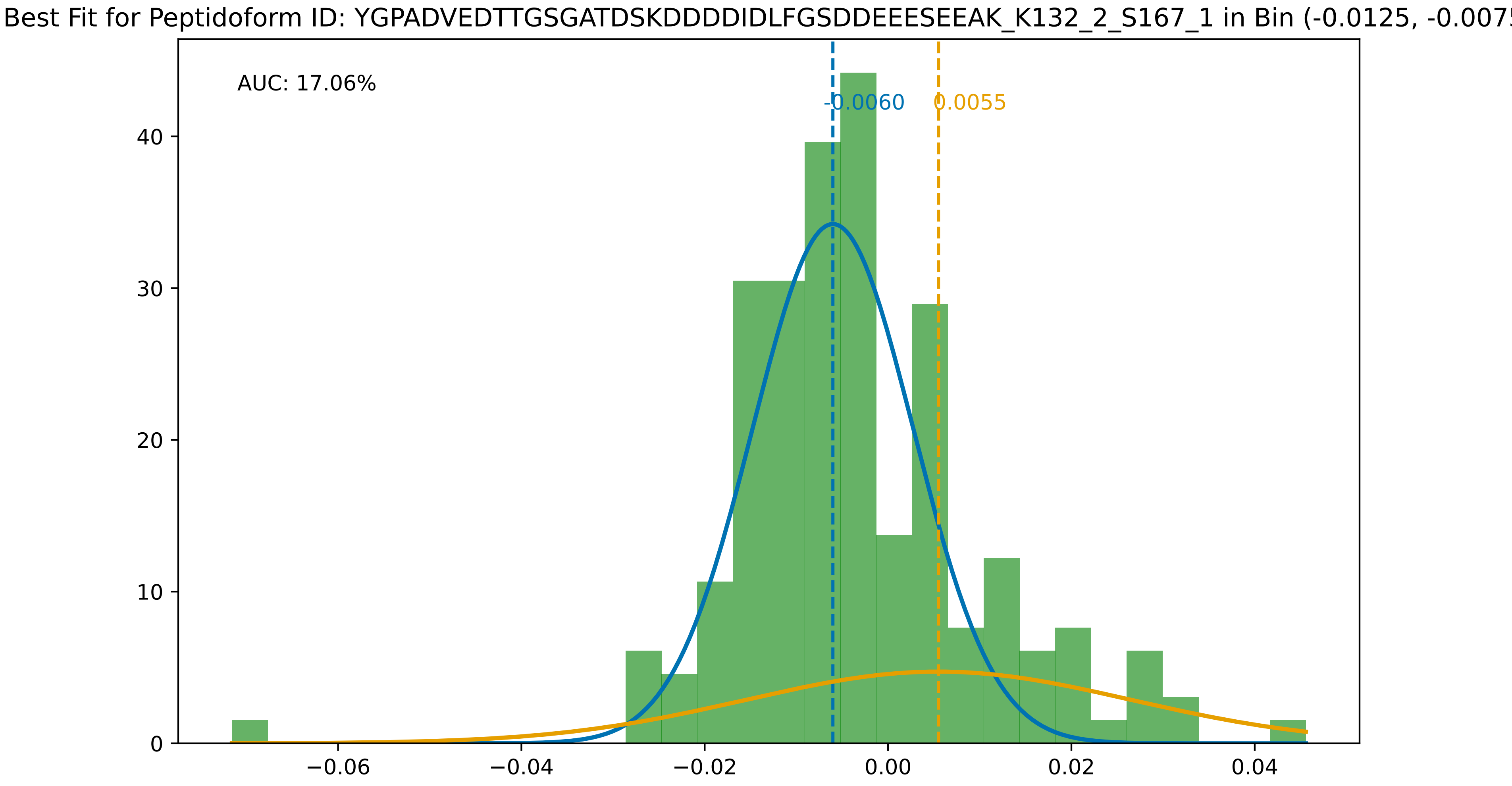

Best Fit for Peptidoform ID: YGPADVEDTTGSGATDSKDDDDIDLFGSDDEEESEEAK_K132_2_S167_1 in Bin (-0.0125, -0.0075
AUC: 17.06%
-0.0060
0.0055
0
10
20
30
40
−0.06
−0.04
−0.02
0.00
0.02
0.04

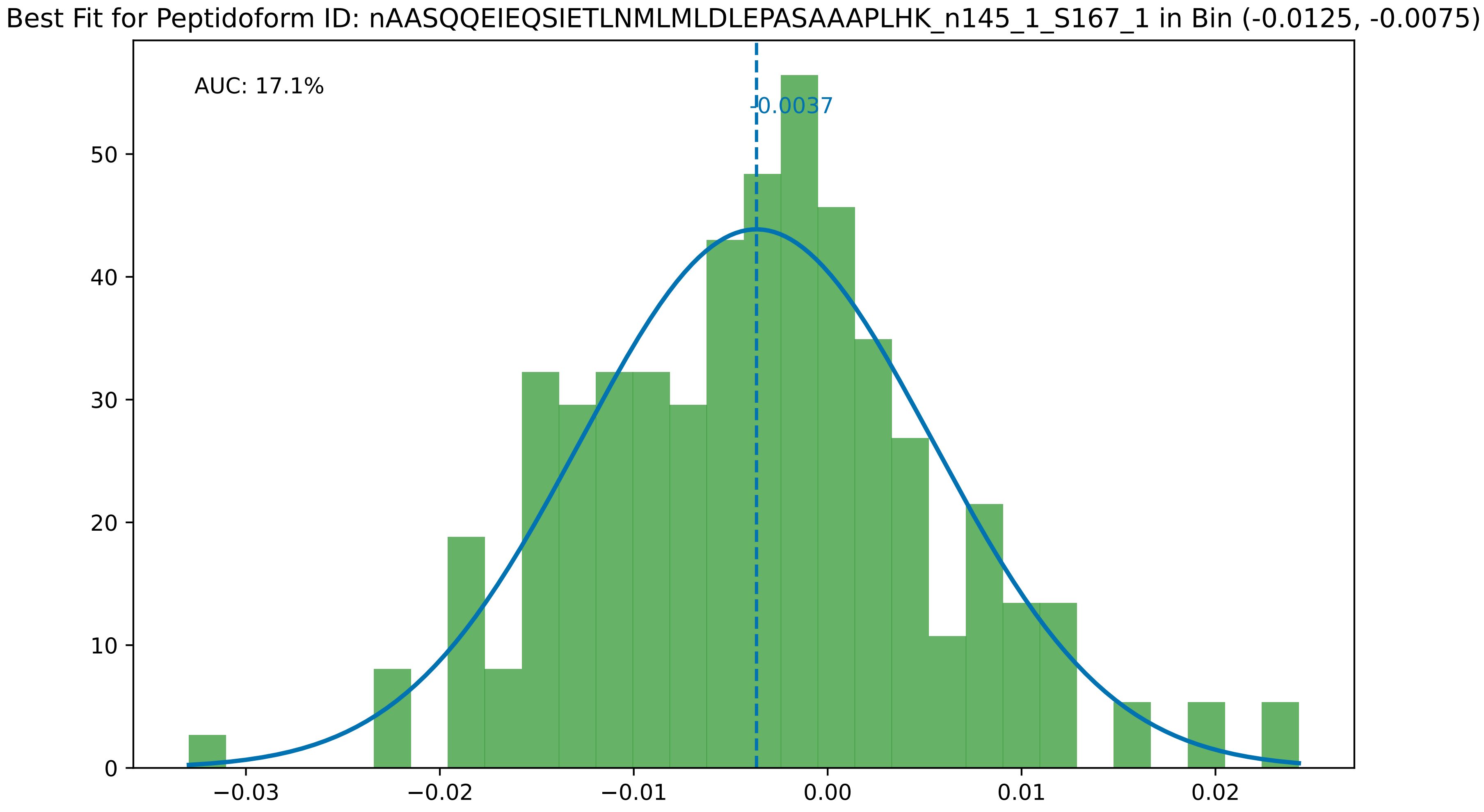

Best Fit for Peptidoform ID: nAASQQEIEQSIETLNMLMLDLEPASAAAPLHK_n145_1_S167_1 in Bin (-0.0125, -0.0075)
AUC: 17.1%
-0.0037
0
10
20
30
40
50
−0.03
−0.02
−0.01
0.00
0.01
0.02

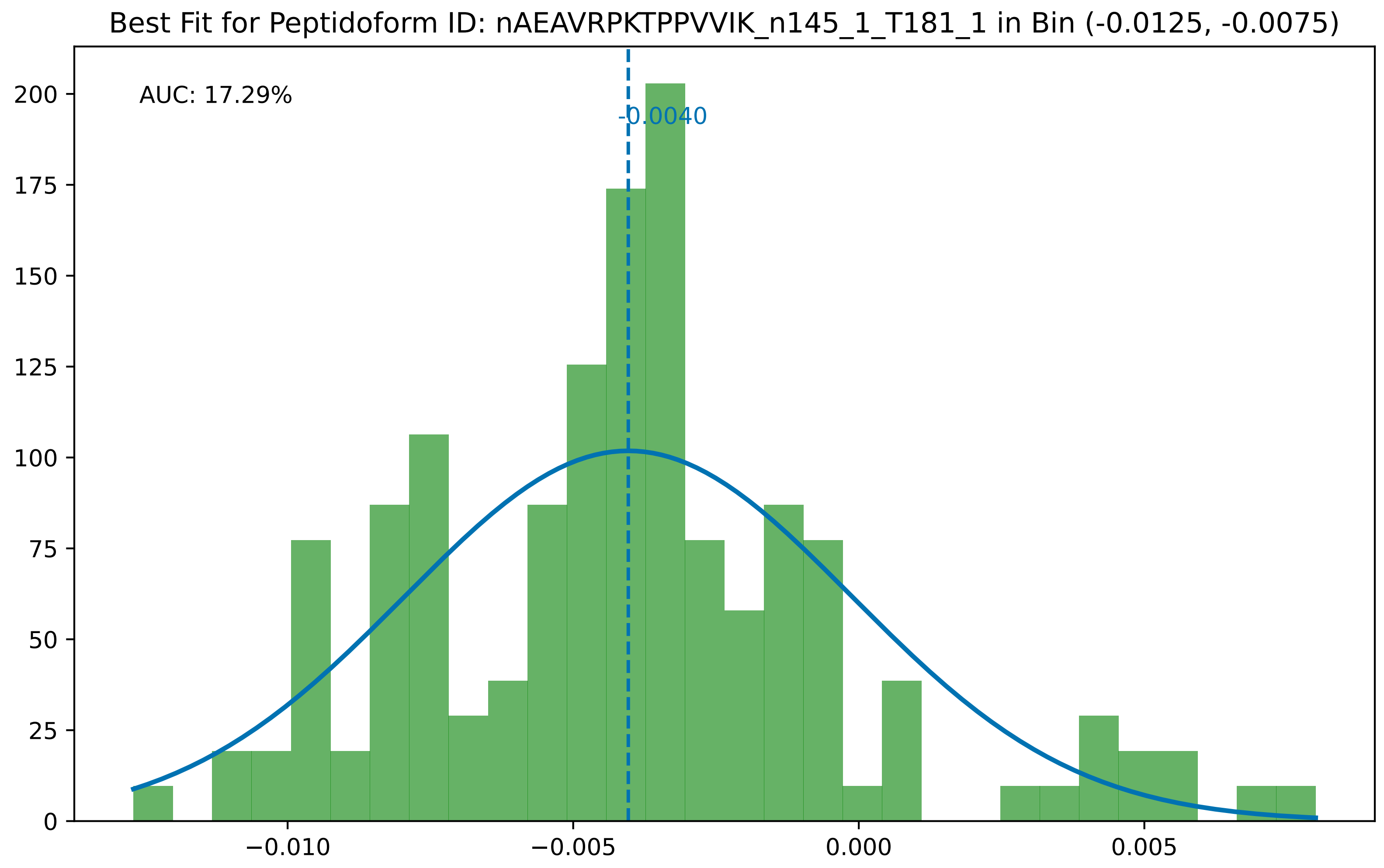
Best Fit for Peptidoform ID: nAEAVRPKTPPVVIK_n145_1_T181_1 in Bin (-0.0125, -0.0075)
AUC: 17.29%
-0.0040
0
25
50
75
100
125
150
175
200
−0.010
−0.005
0.000
0.005

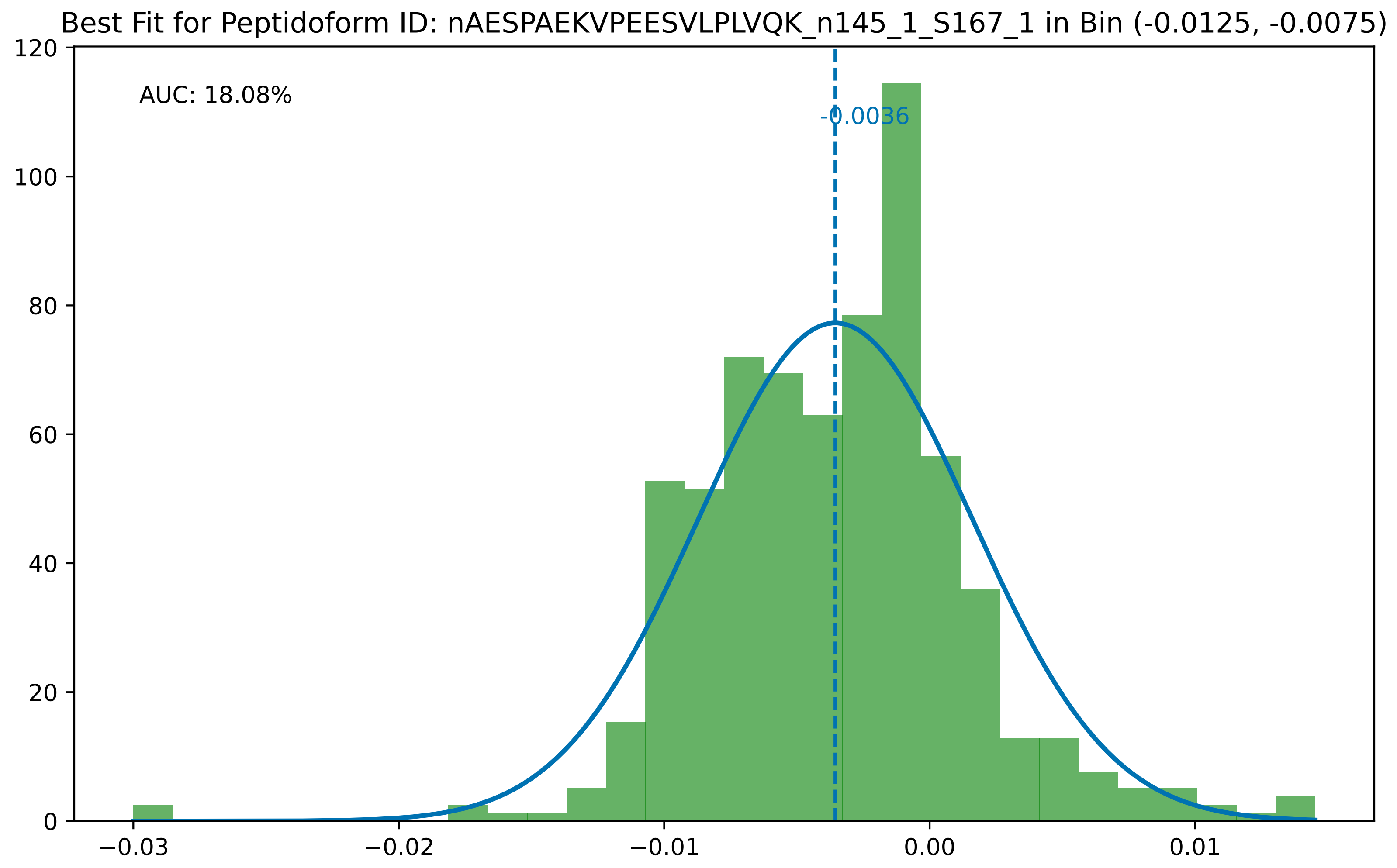

Best Fit for Peptidoform ID: nAESPAEKVPEESVLPLVQK_n145_1_S167_1 in Bin (-0.0125, -0.0075)
AUC: 18.08%
-0.0036
0
20
40
60
80
100
120
−0.03
−0.02
−0.01
0.00
0.01

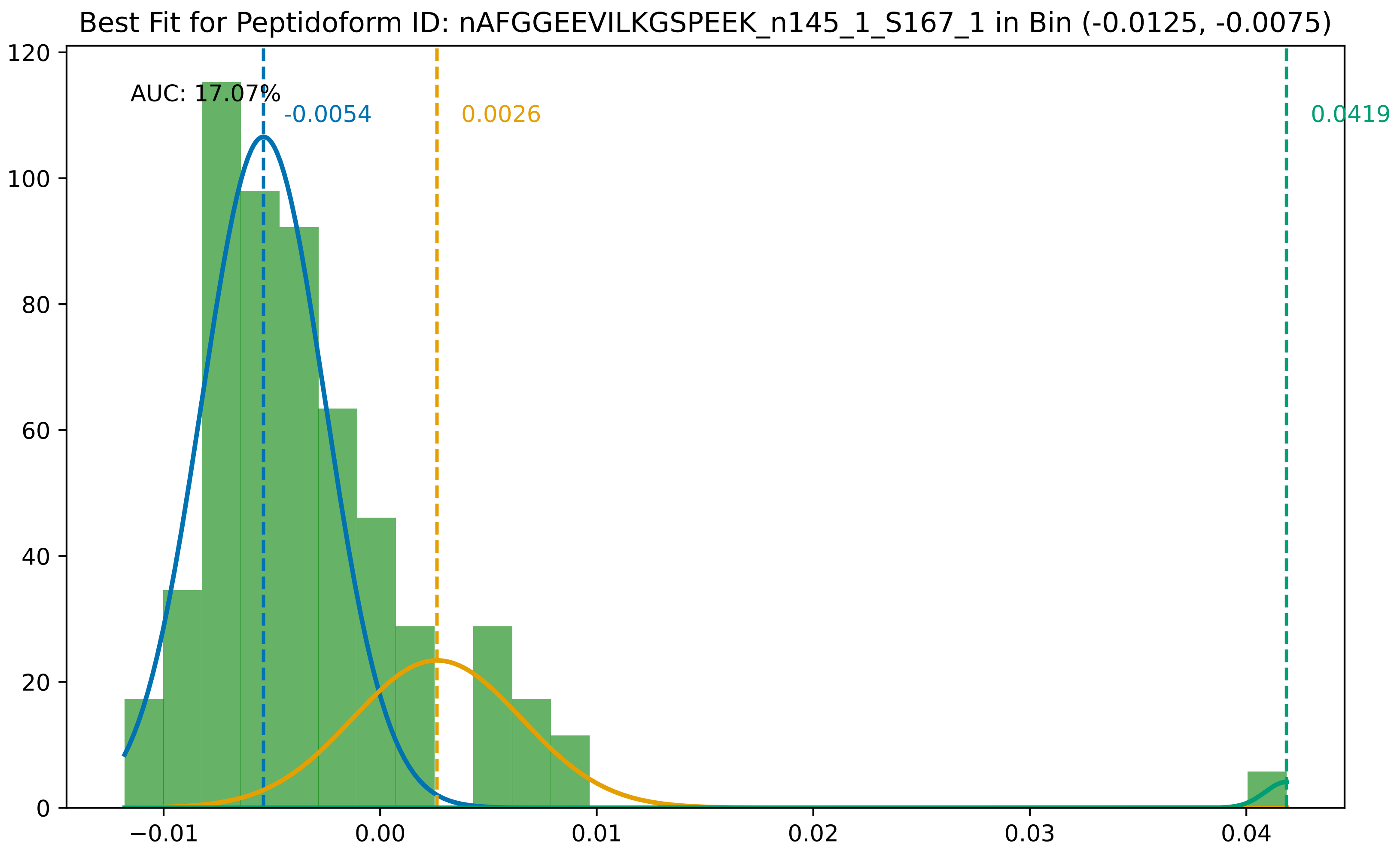

Best Fit for Peptidoform ID: nAFGGEEVILKGSPEEK_n145_1_S167_1 in Bin (-0.0125, -0.0075)
AUC: 17.07%
-0.0054
0.0026
0.0419
0
20
40
60
80
100
120
−0.01
0.00
0.01
0.02
0.03
0.04

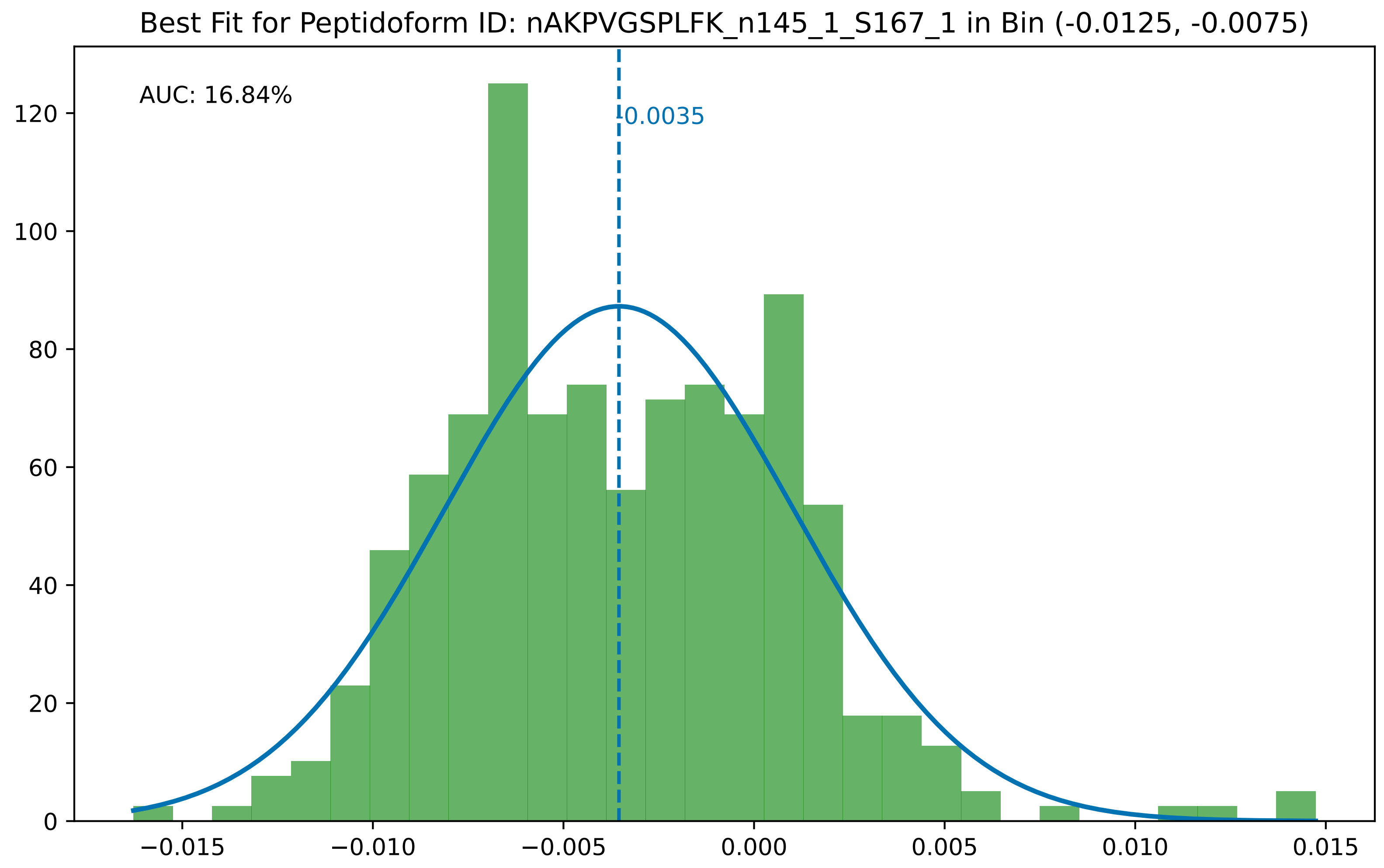

Best Fit for Peptidoform ID: nAKPVGSPLFK_n145_1_S167_1 in Bin (-0.0125, -0.0075)
AUC: 16.84%
-0.0035
0
20
40
60
80
100
120
−0.015
−0.010
−0.005
0.000
0.005
0.010
0.015

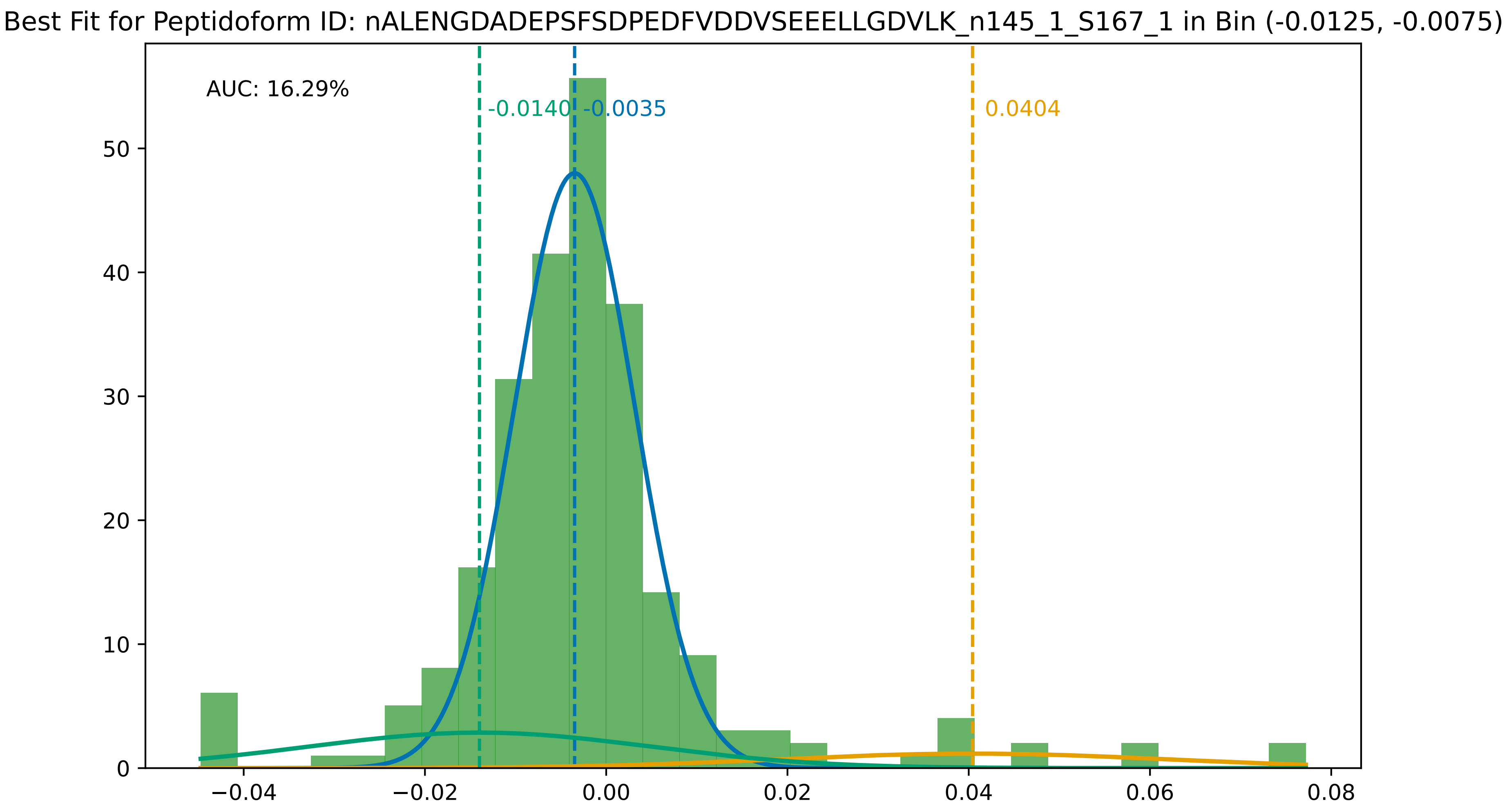
Best Fit for Peptidoform ID: nALENGDADEPSFSDPEDFVDDVSEEELLGDVLK_n145_1_S167_1 in Bin (-0.0125, -0.0075)
AUC: 16.29%
-0.0140
-0.0035
0.0404
0
10
20
30
40
50
−0.04
−0.02
0.00
0.02
0.04
0.06
0.08

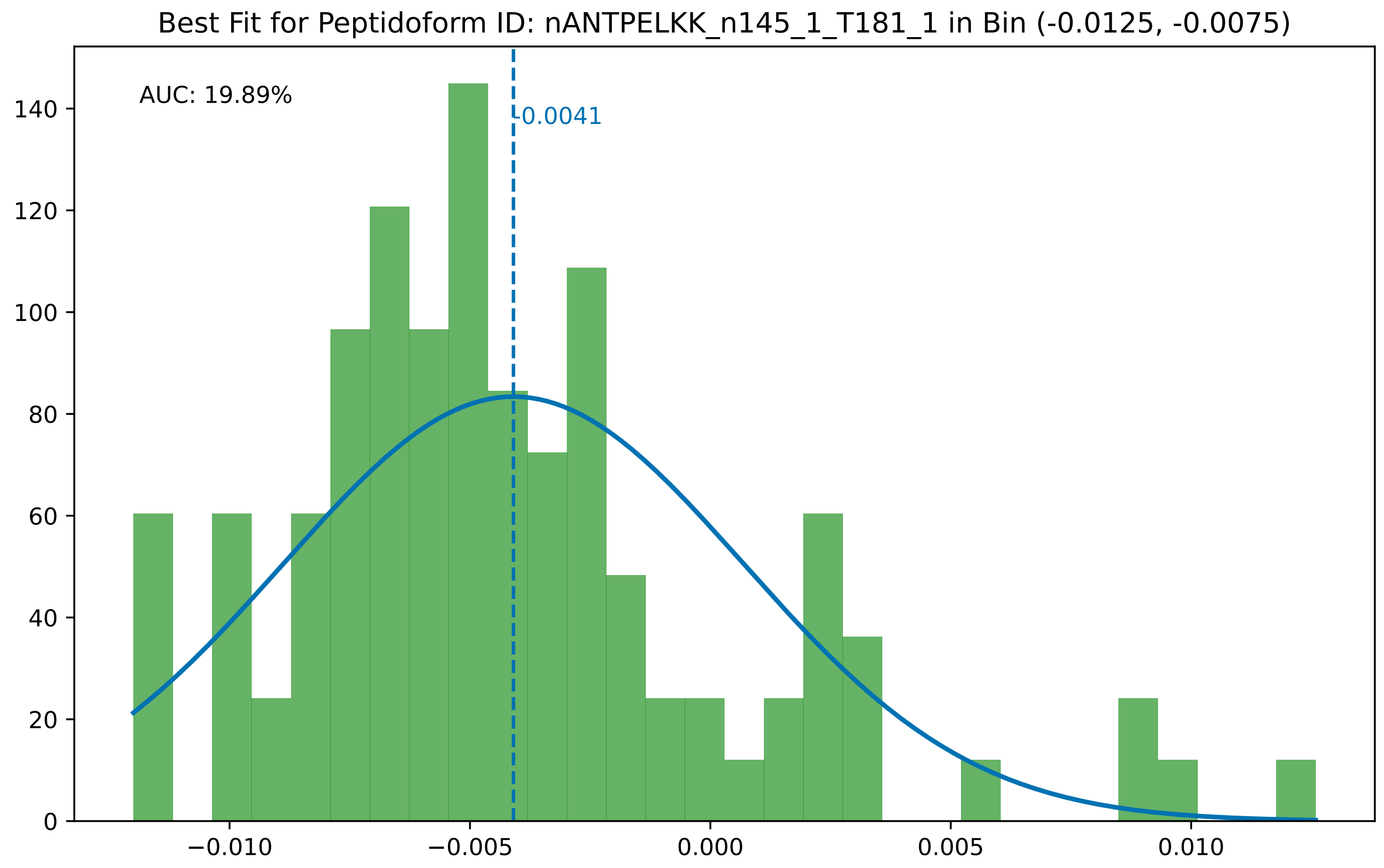

Best Fit for Peptidoform ID: nANTPELKK_n145_1_T181_1 in Bin (-0.0125, -0.0075)
AUC: 19.89%
-0.0041
0
20
40
60
80
100
120
140
−0.010
−0.005
0.000
0.005
0.010

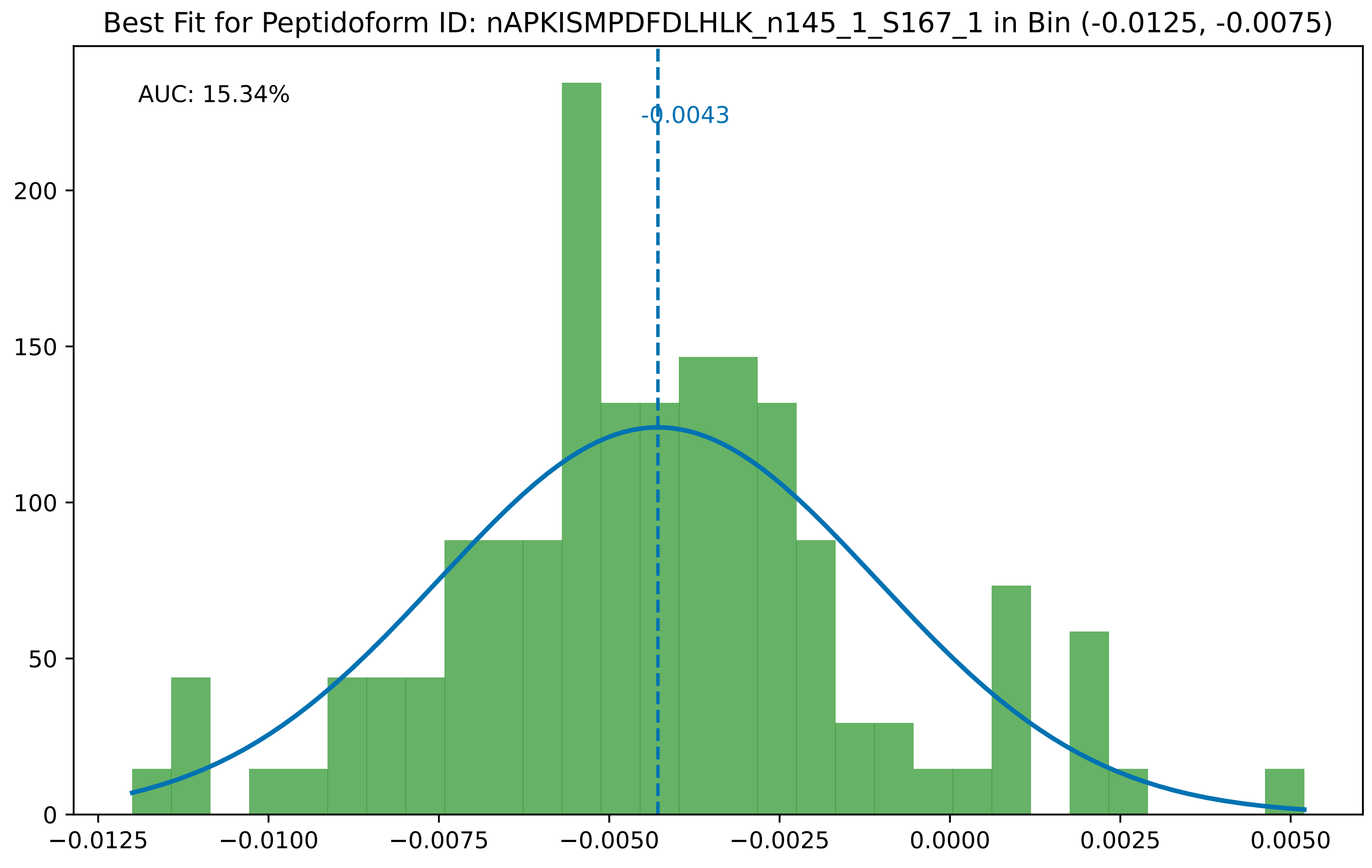

Best Fit for Peptidoform ID: nAPKISMPDFDLHLK_n145_1_S167_1 in Bin (-0.0125, -0.0075)
AUC: 15.34%
-0.0043
0
50
100
150
200
−0.0125
−0.0100
−0.0075
−0.0050
−0.0025
0.0000
0.0025
0.0050

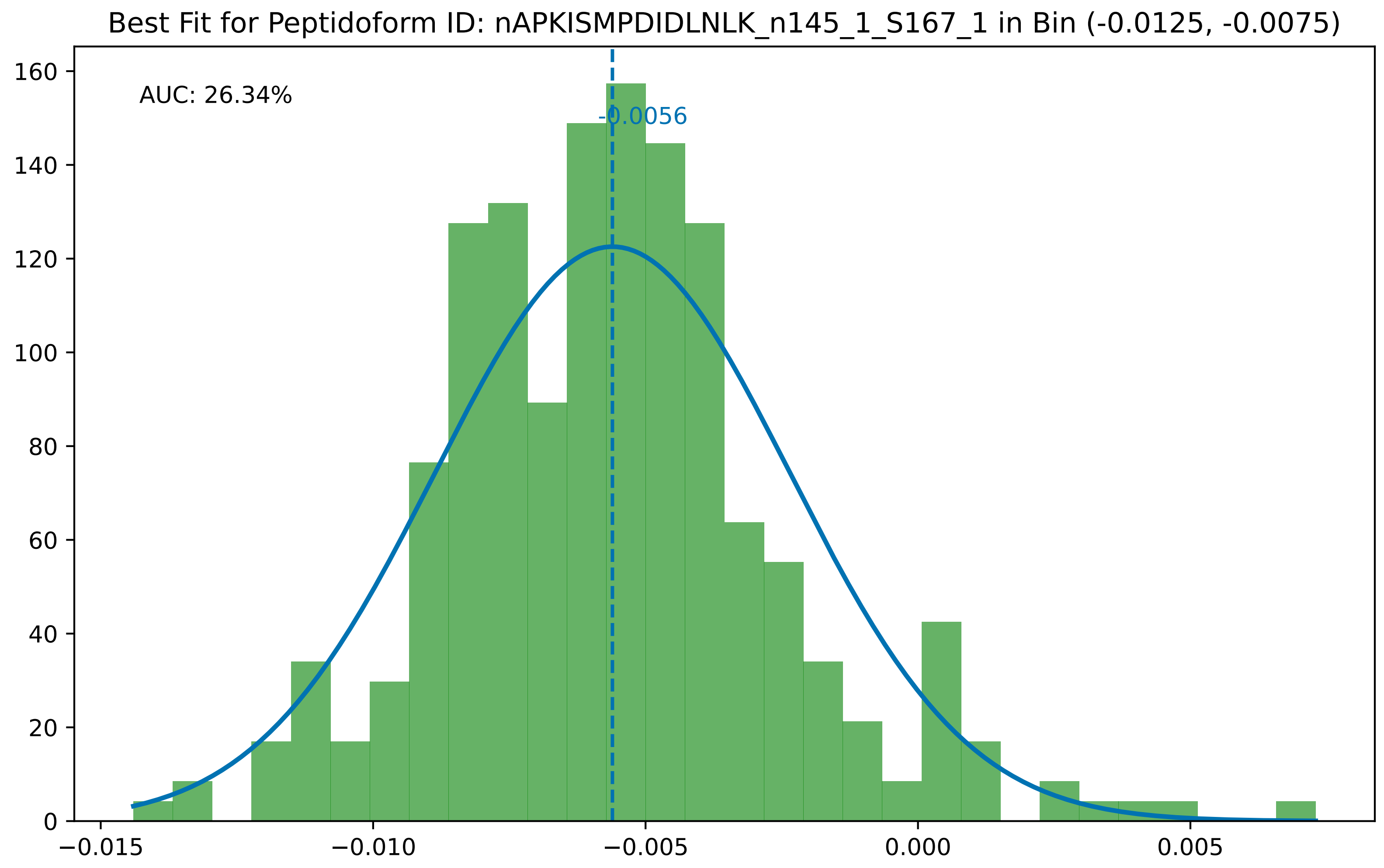
Best Fit for Peptidoform ID: nAPKISMPDIDLNLK_n145_1_S167_1 in Bin (-0.0125, -0.0075)
AUC: 26.34%
-0.0056
0
20
40
60
80
100
120
140
160
−0.015
−0.010
−0.005
0.000
0.005

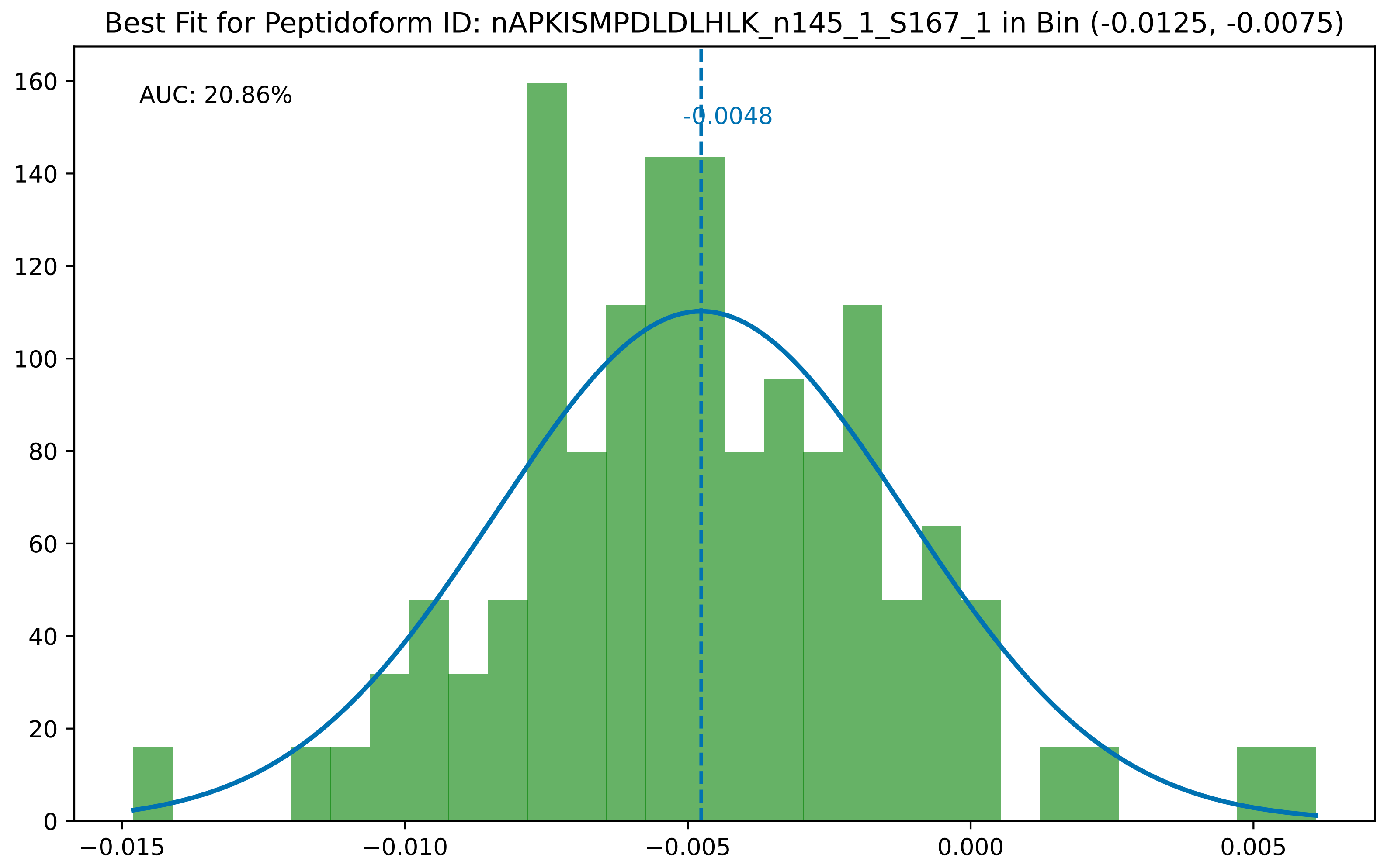
Best Fit for Peptidoform ID: nAPKISMPDLDLHLK_n145_1_S167_1 in Bin (-0.0125, -0.0075)
AUC: 20.86%
-0.0048
0
20
40
60
80
100
120
140
160
−0.015
−0.010
−0.005
0.000
0.005

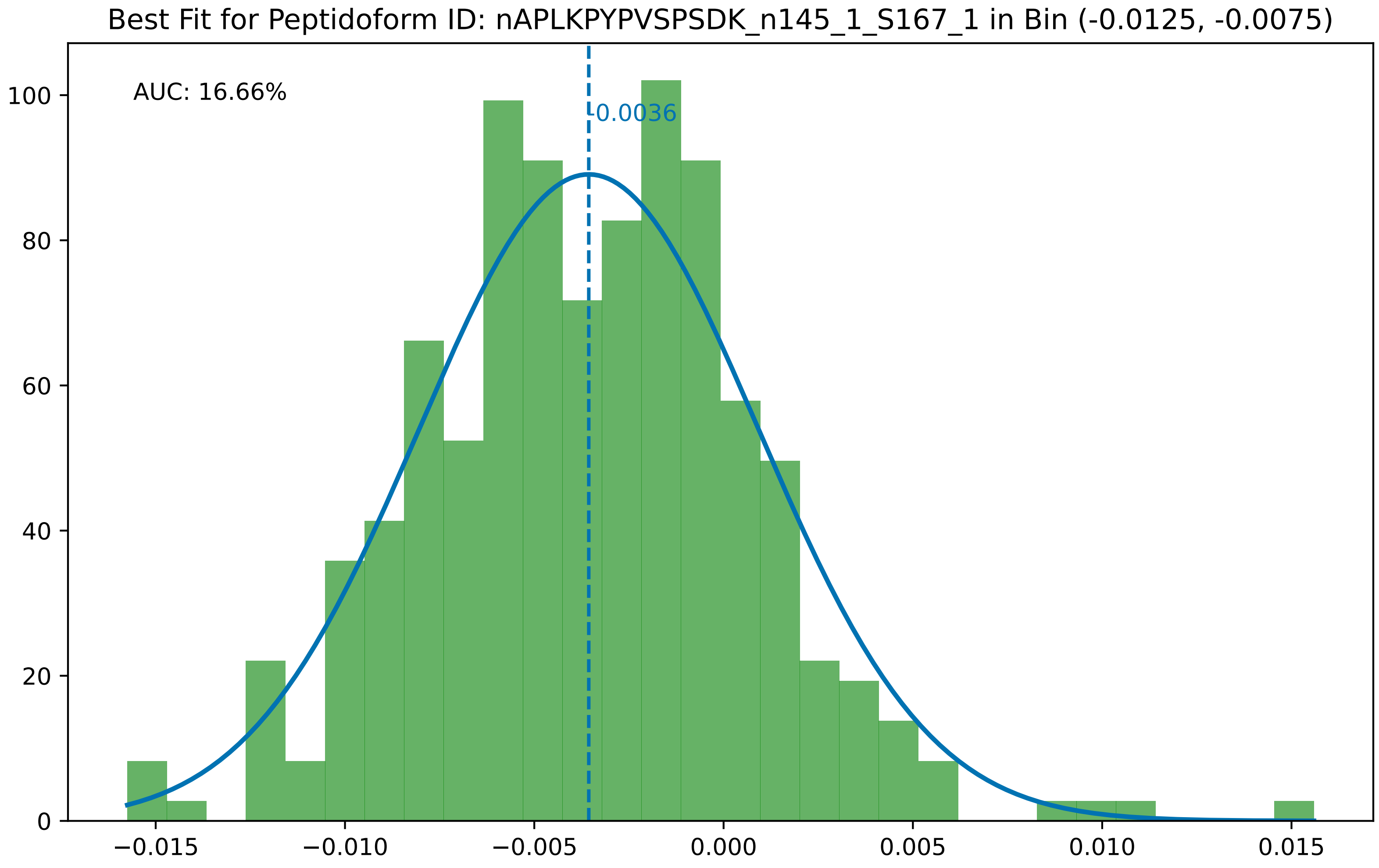
Best Fit for Peptidoform ID: nAPLKPYPVSPSDK_n145_1_S167_1 in Bin (-0.0125, -0.0075)
AUC: 16.66%
-0.0036
100
80
60
40
20
0
−0.015
−0.010
−0.005
0.000
0.005
0.010
0.015

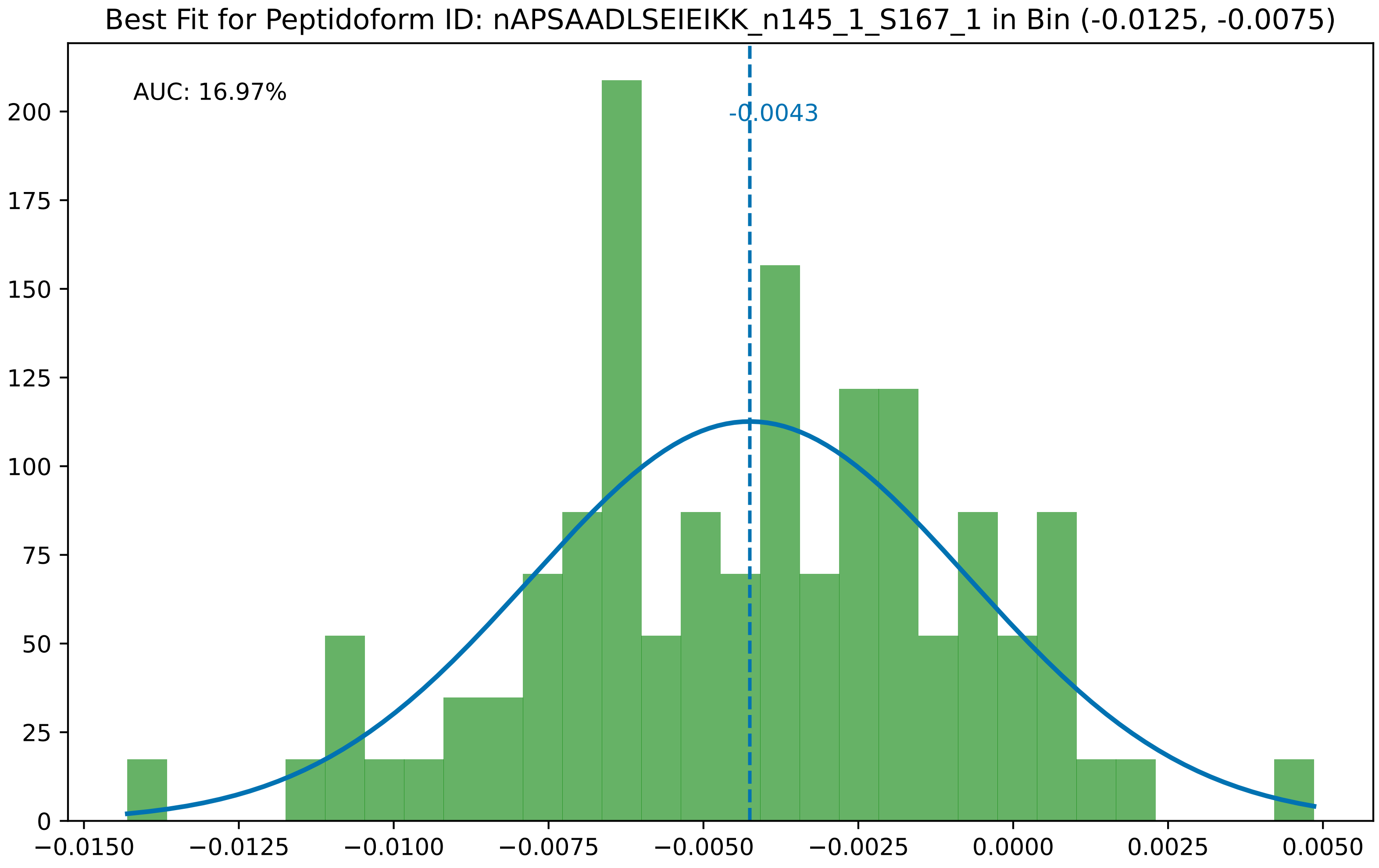

Best Fit for Peptidoform ID: nAPSAADLSEIEIKK_n145_1_S167_1 in Bin (-0.0125, -0.0075)
AUC: 16.97%
-0.0043
0
25
50
75
100
125
150
175
200
−0.0150
−0.0125
−0.0100
−0.0075
−0.0050
−0.0025
0.0000
0.0025
0.0050

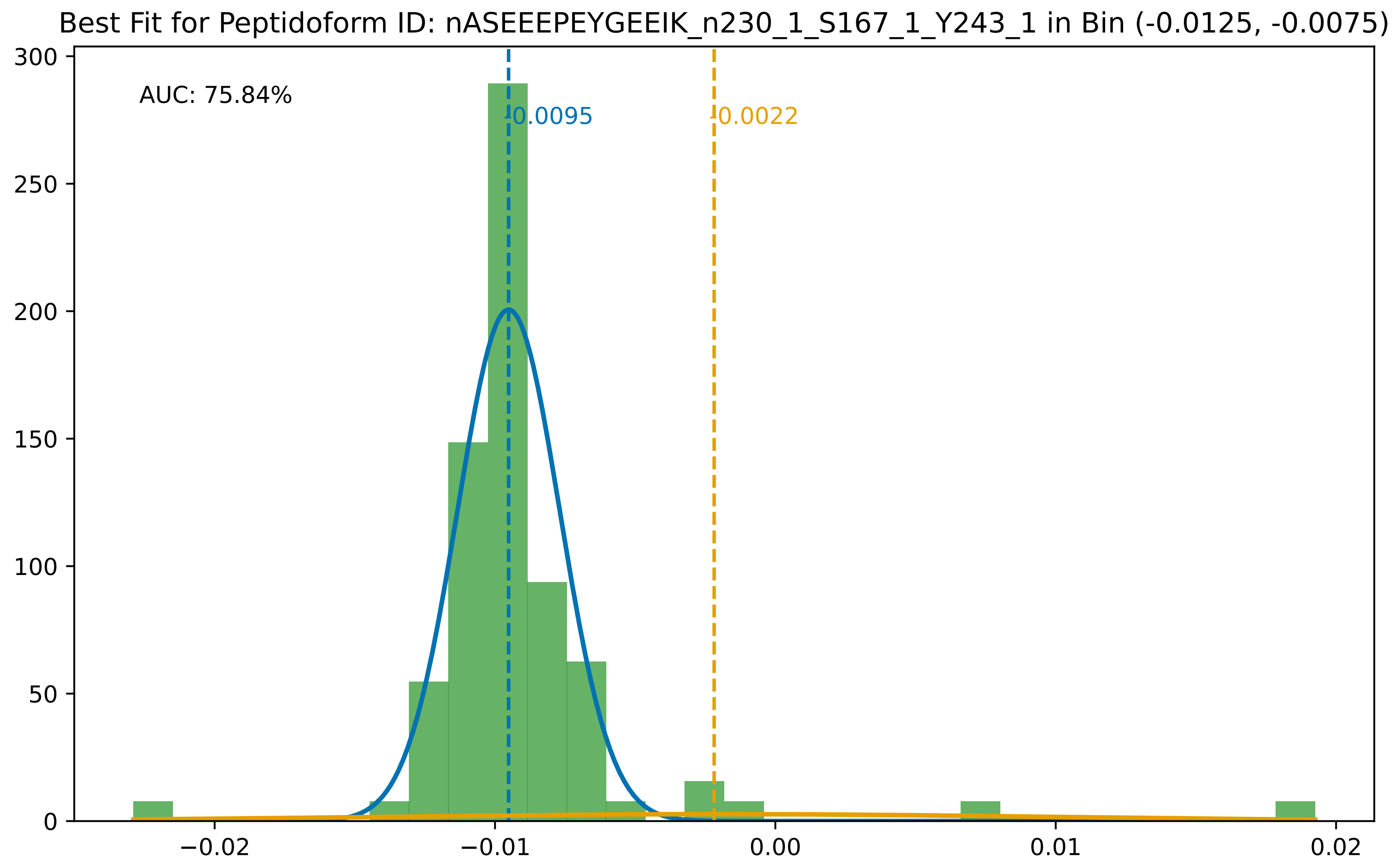
Best Fit for Peptidoform ID: nASEEEPEYGEEIK_n230_1_S167_1_Y243_1 in Bin (-0.0125, -0.0075)
AUC: 75.84%
-0.0095
-0.0022
300
250
200
150
100
50
0
−0.02
−0.01
0.00
0.01
0.02

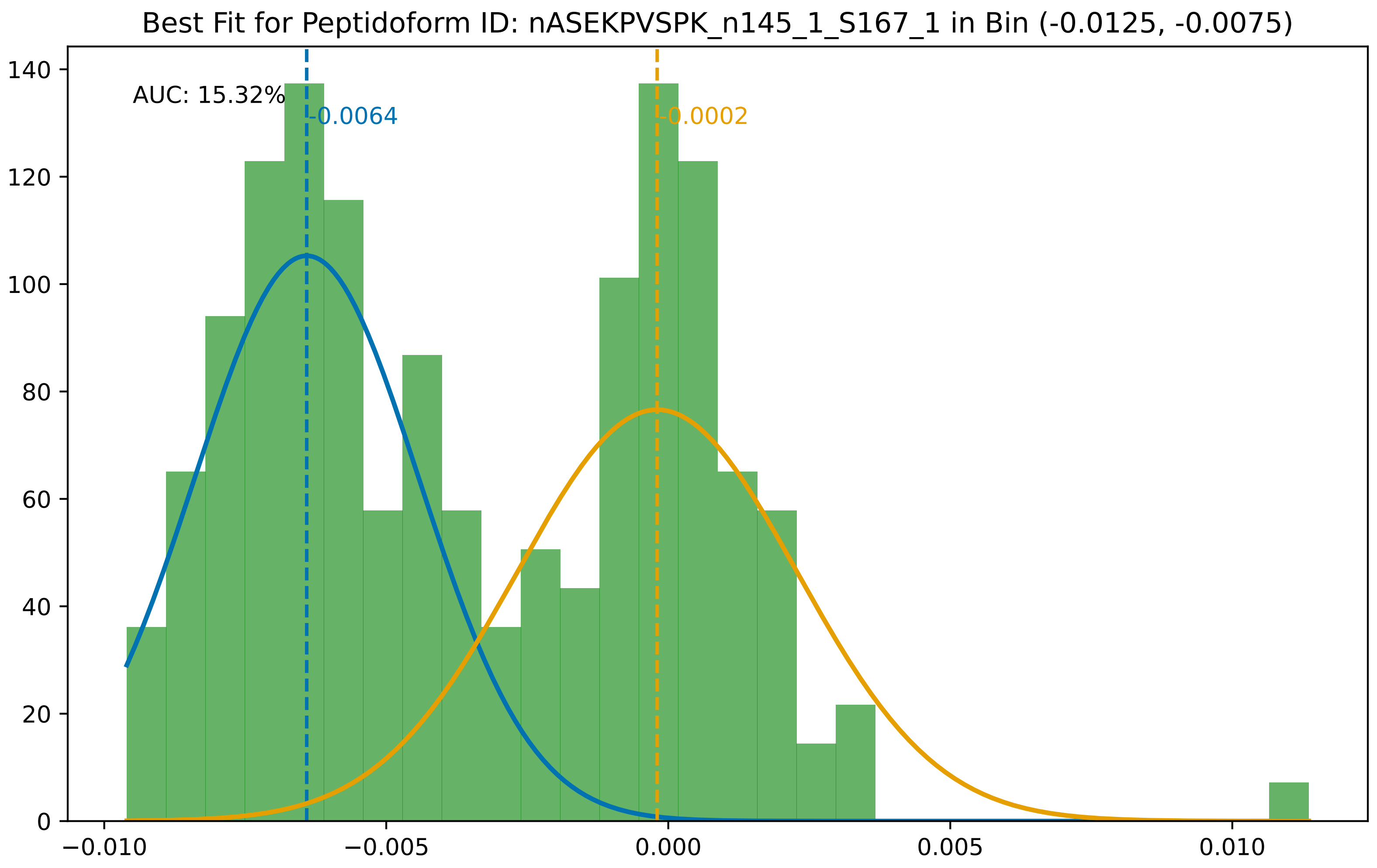

Best Fit for Peptidoform ID: nASEKPVSPK_n145_1_S167_1 in Bin (-0.0125, -0.0075)
AUC: 15.32%
-0.0064
-0.0002
0
20
40
60
80
100
120
140
−0.010
−0.005
0.000
0.005
0.010

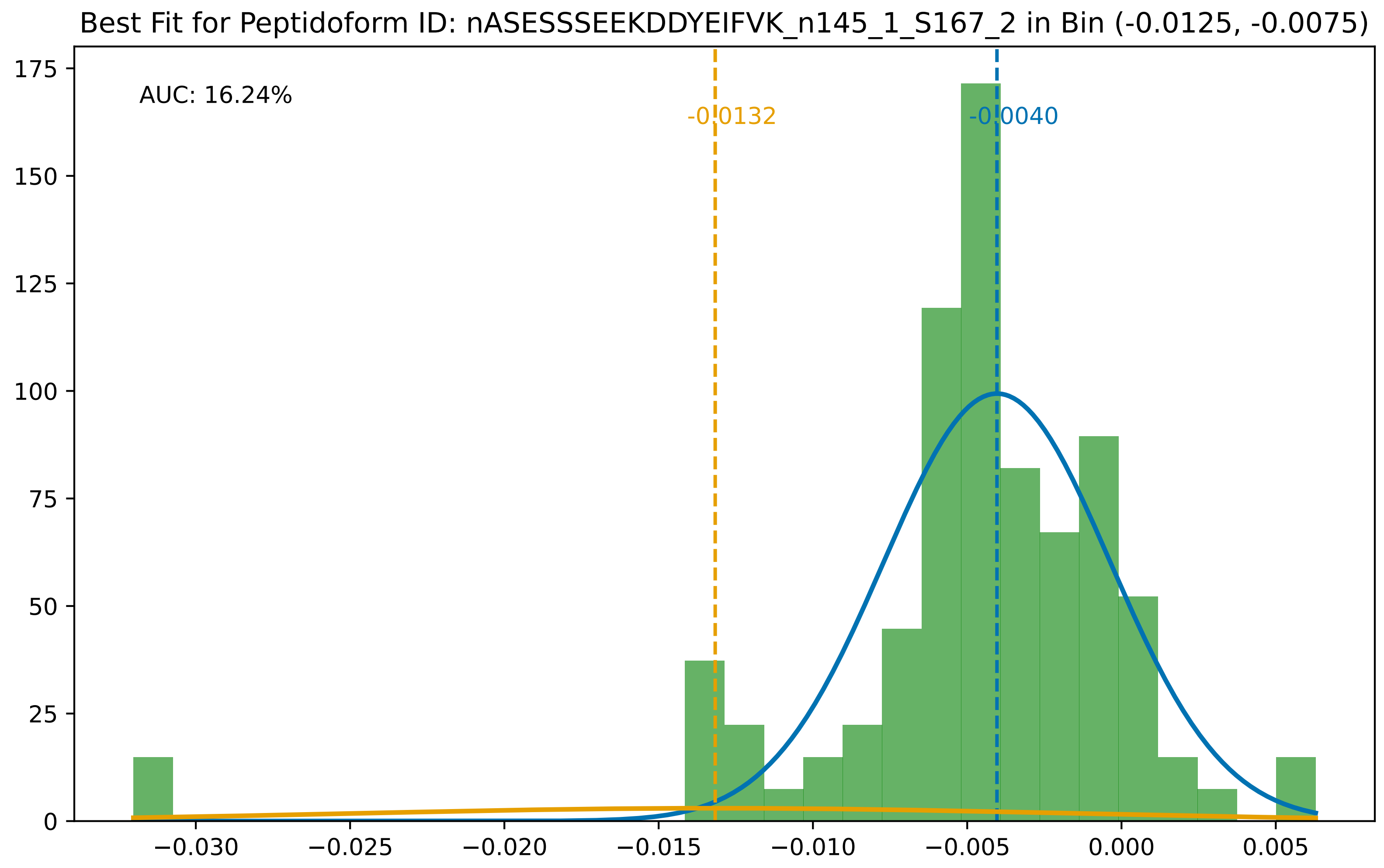

Best Fit for Peptidoform ID: nASESSSEEKDDYEIFVK_n145_1_S167_2 in Bin (-0.0125, -0.0075)
AUC: 16.24%
-0.0132
-0.0040
175
150
125
100
75
50
25
0
−0.030
−0.025
−0.020
−0.015
−0.010
−0.005
0.000
0.005

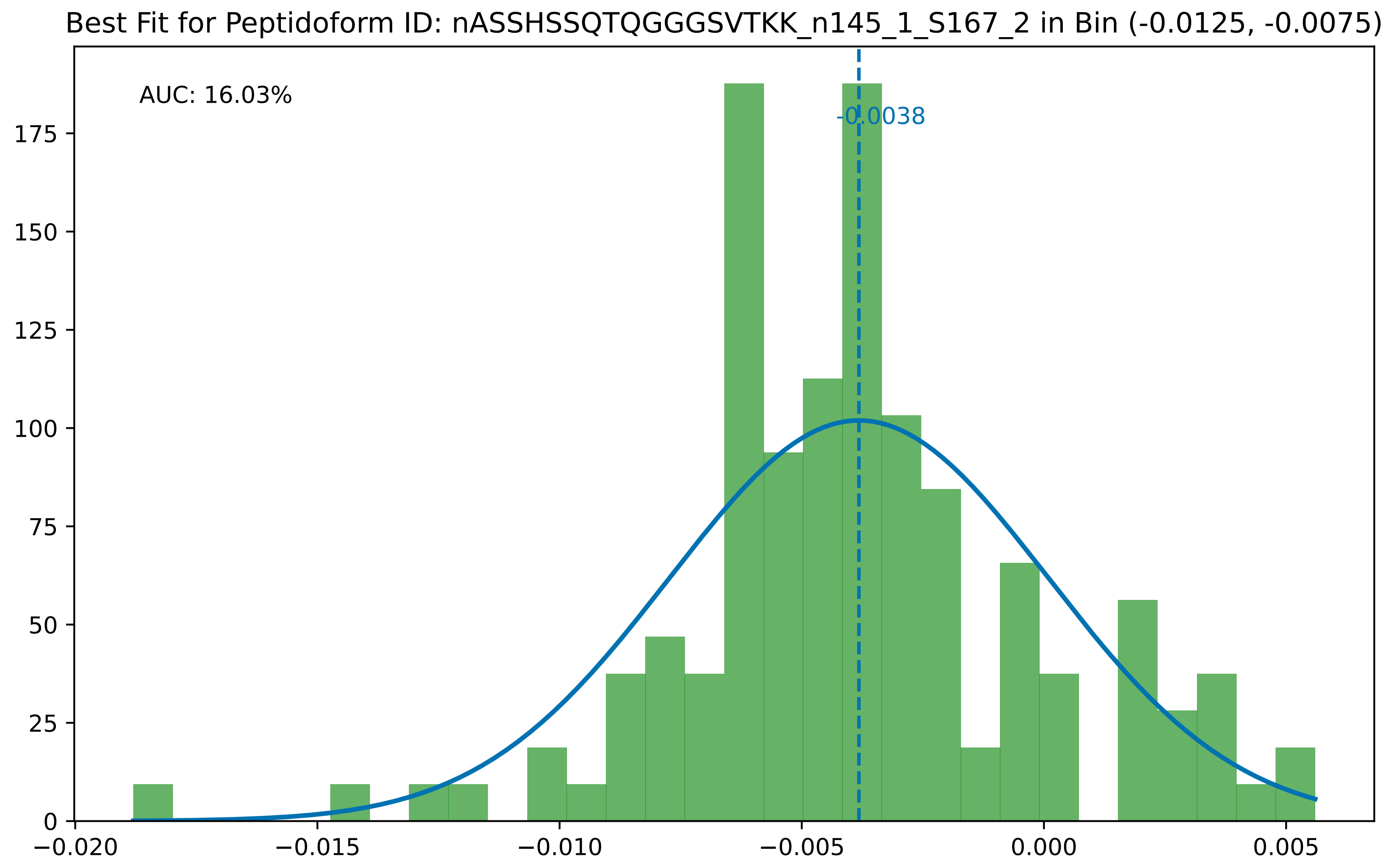

Best Fit for Peptidoform ID: nASSHSSQTQGGGSVTKK_n145_1_S167_2 in Bin (-0.0125, -0.0075)
AUC: 16.03%
-0.0038
0
25
50
75
100
125
150
175
−0.020
−0.015
−0.010
−0.005
0.000
0.005

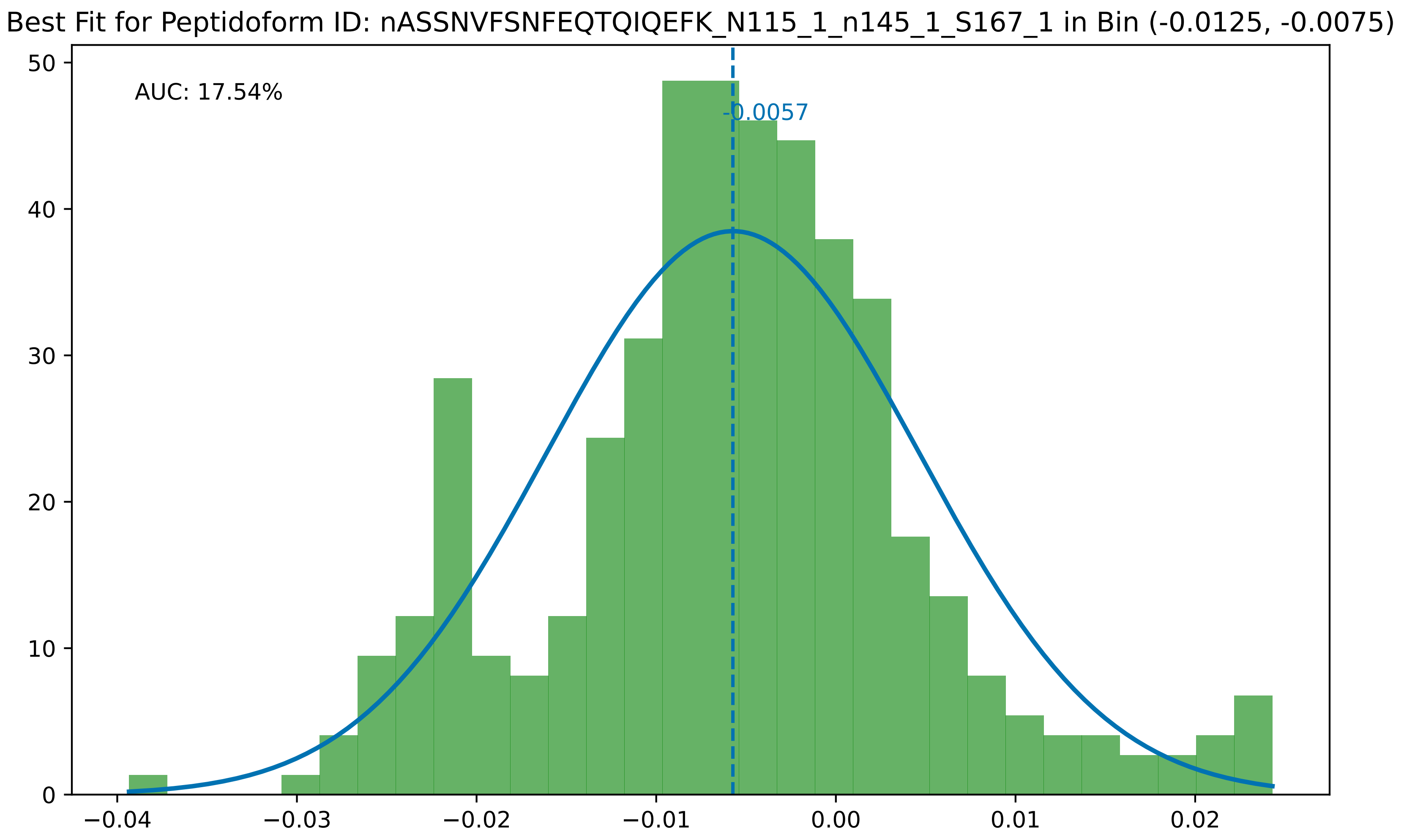

Best Fit for Peptidoform ID: nASSNVFSNFEQTQIQEFK_N115_1_n145_1_S167_1 in Bin (-0.0125, -0.0075)
AUC: 17.54%
-0.0057
0
10
20
30
40
50
−0.04
−0.03
−0.02
−0.01
0.00
0.01
0.02

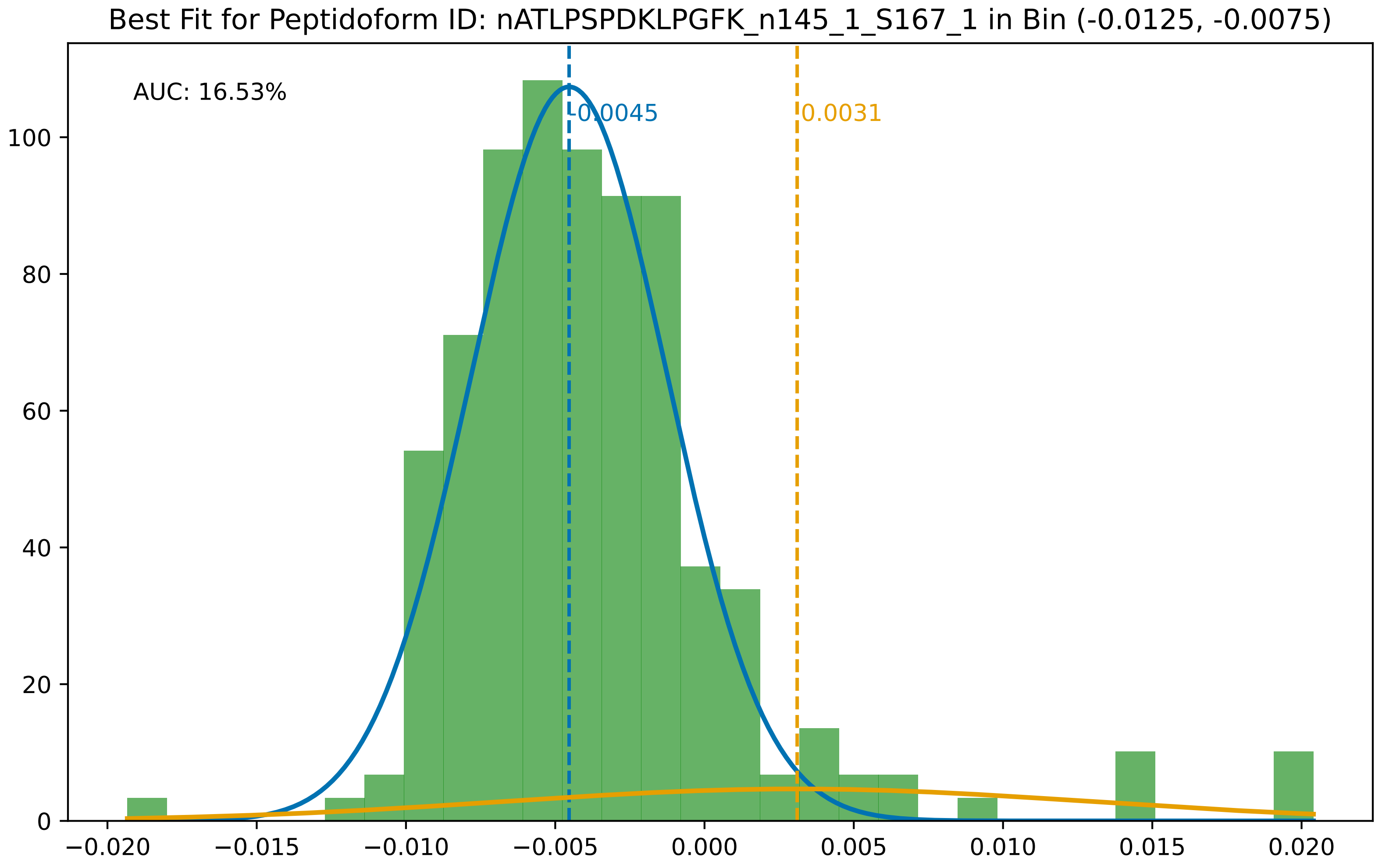

Best Fit for Peptidoform ID: nATLPSPDKLPGFK_n145_1_S167_1 in Bin (-0.0125, -0.0075)
AUC: 16.53%
-0.0045
0.0031
100
80
60
40
20
0
−0.020
−0.015
−0.010
−0.005
0.000
0.005
0.010
0.015
0.020

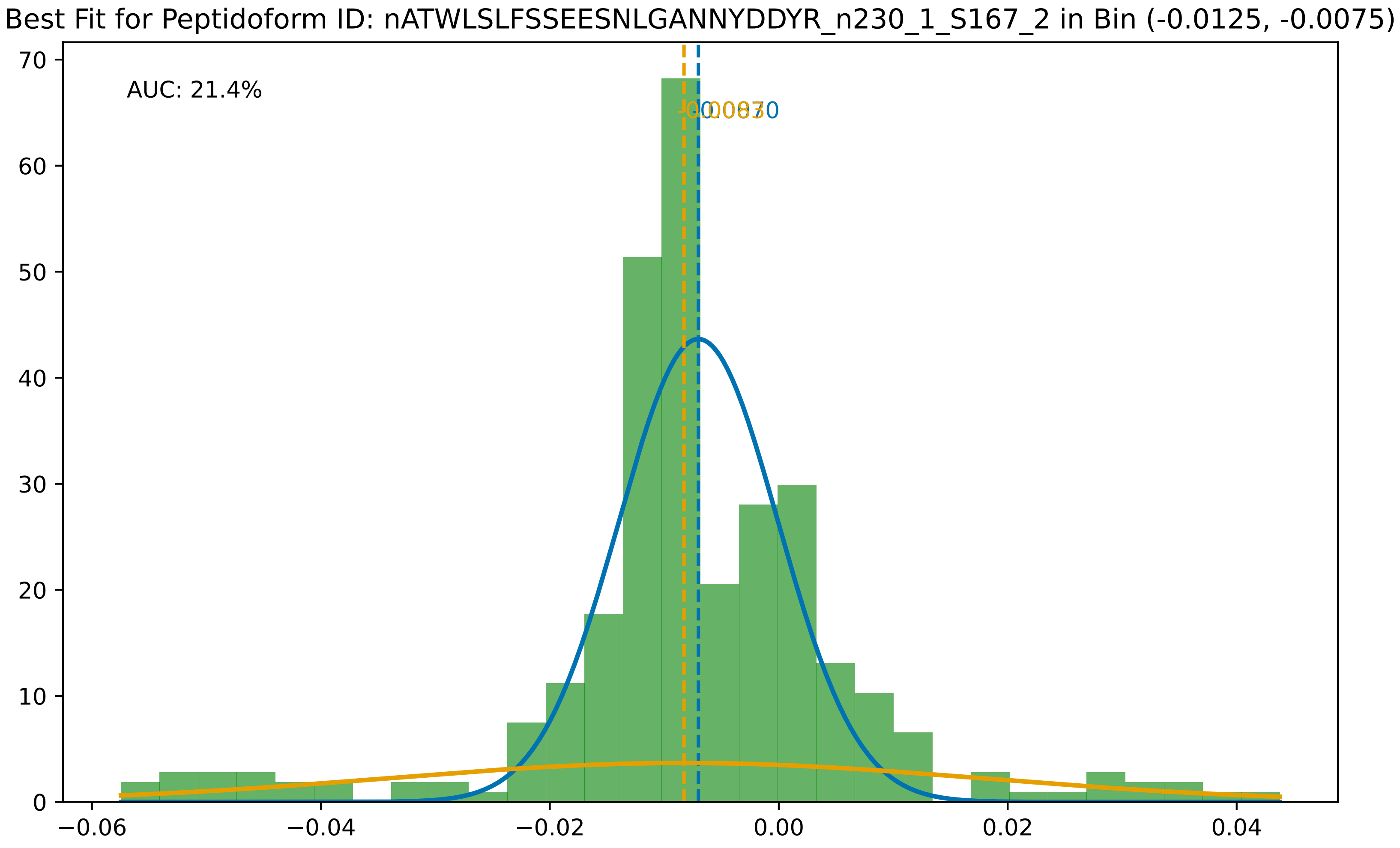

Best Fit for Peptidoform ID: nATWLSLFSSEESNLGANNYDDYR_n230_1_S167_2 in Bin (-0.0125, -0.0075)
AUC: 21.4%
70
60
50
40
30
20
10
0
−0.06
−0.04
−0.02
0.00
0.02
0.04

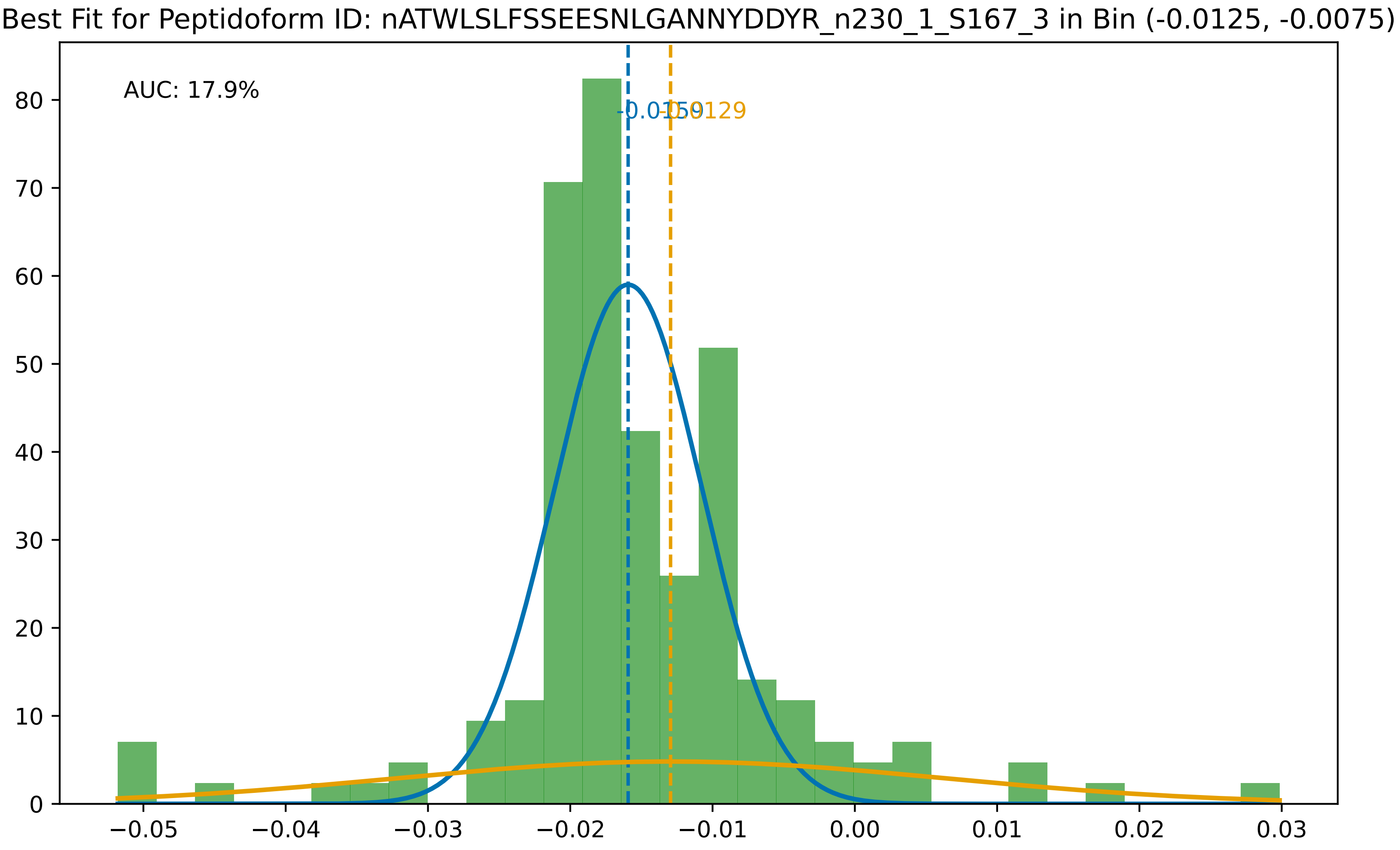

Best Fit for Peptidoform ID: nATWLSLFSSEESNLGANNYDDYR_n230_1_S167_3 in Bin (-0.0125, -0.0075)
AUC: 17.9%
-0.0159
-0.0129
0
10
20
30
40
50
60
70
80
−0.05
−0.04
−0.03
−0.02
−0.01
0.00
0.01
0.02
0.03

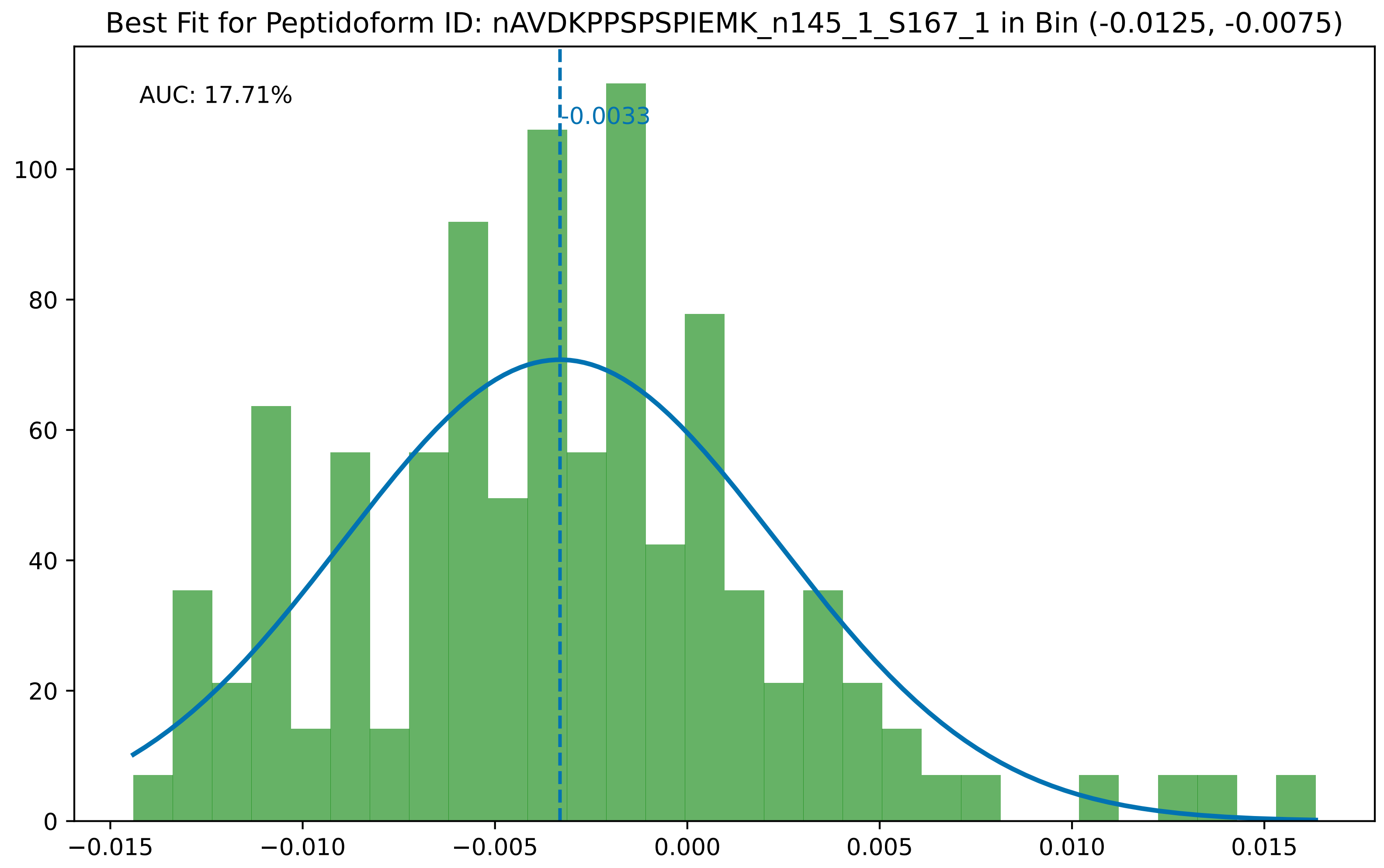
Best Fit for Peptidoform ID: nAVDKPPSPSPIEMK_n145_1_S167_1 in Bin (-0.0125, -0.0075)
AUC: 17.71%
-0.0033
0
20
40
60
80
100
−0.015
−0.010
−0.005
0.000
0.005
0.010
0.015

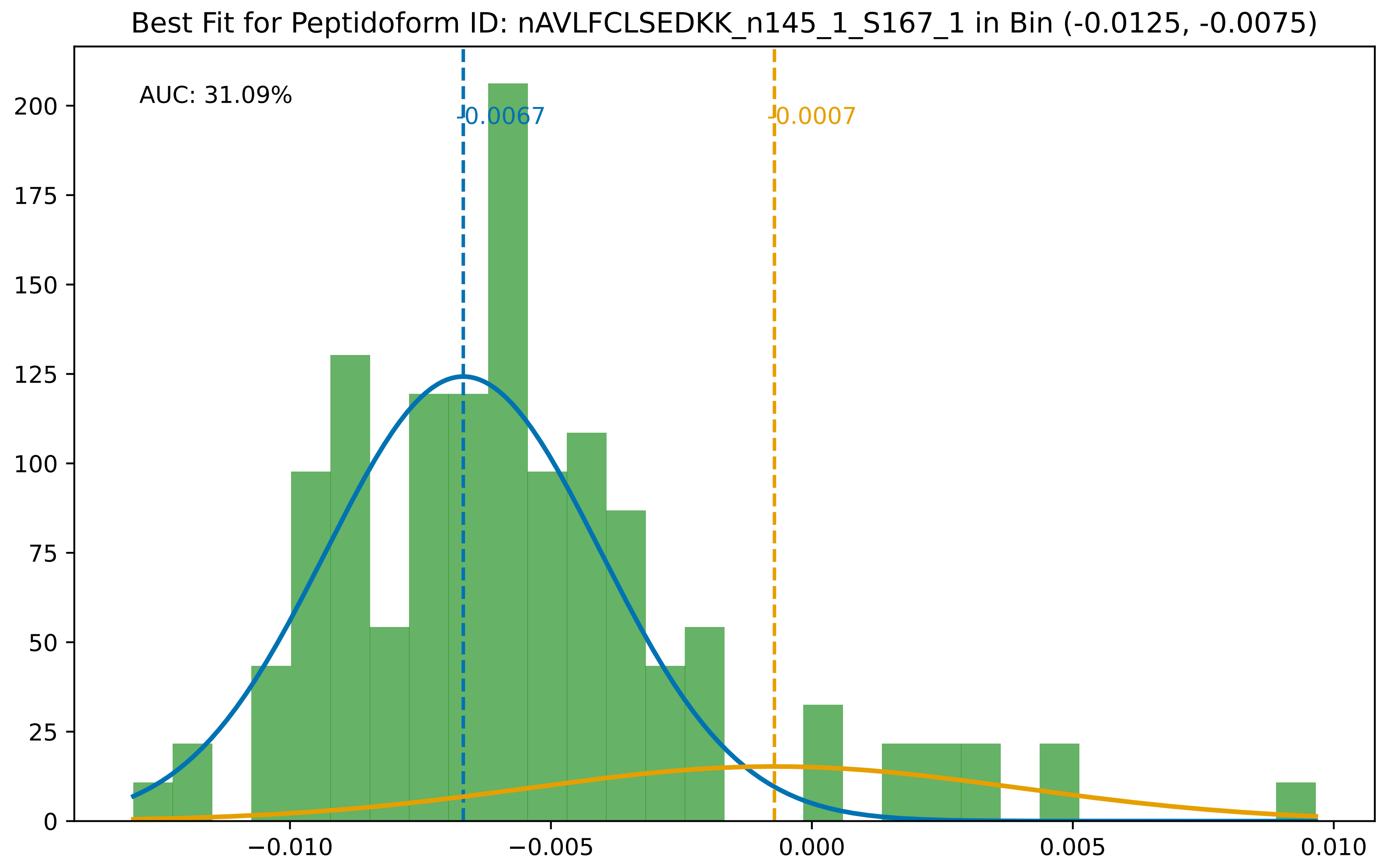

Best Fit for Peptidoform ID: nAVLFCLSEDKK_n145_1_S167_1 in Bin (-0.0125, -0.0075)
AUC: 31.09%
-0.0067
-0.0007
200
175
150
125
100
75
50
25
0
−0.010
−0.005
0.000
0.005
0.010

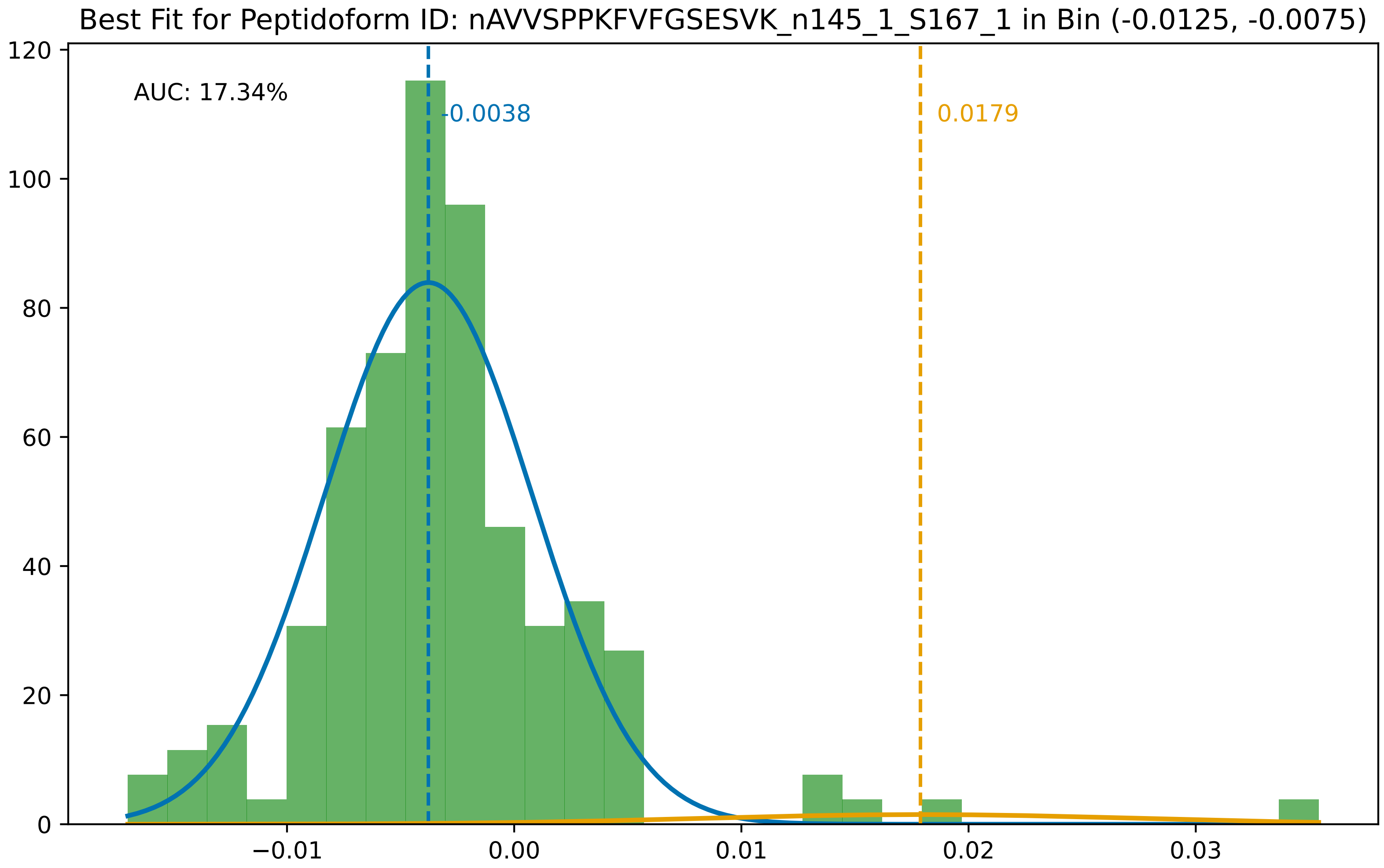
Best Fit for Peptidoform ID: nAVVSPPKFVFGSESVK_n145_1_S167_1 in Bin (-0.0125, -0.0075)
AUC: 17.34%
-0.0038
0.0179
0
20
40
60
80
100
120
−0.01
0.00
0.01
0.02
0.03

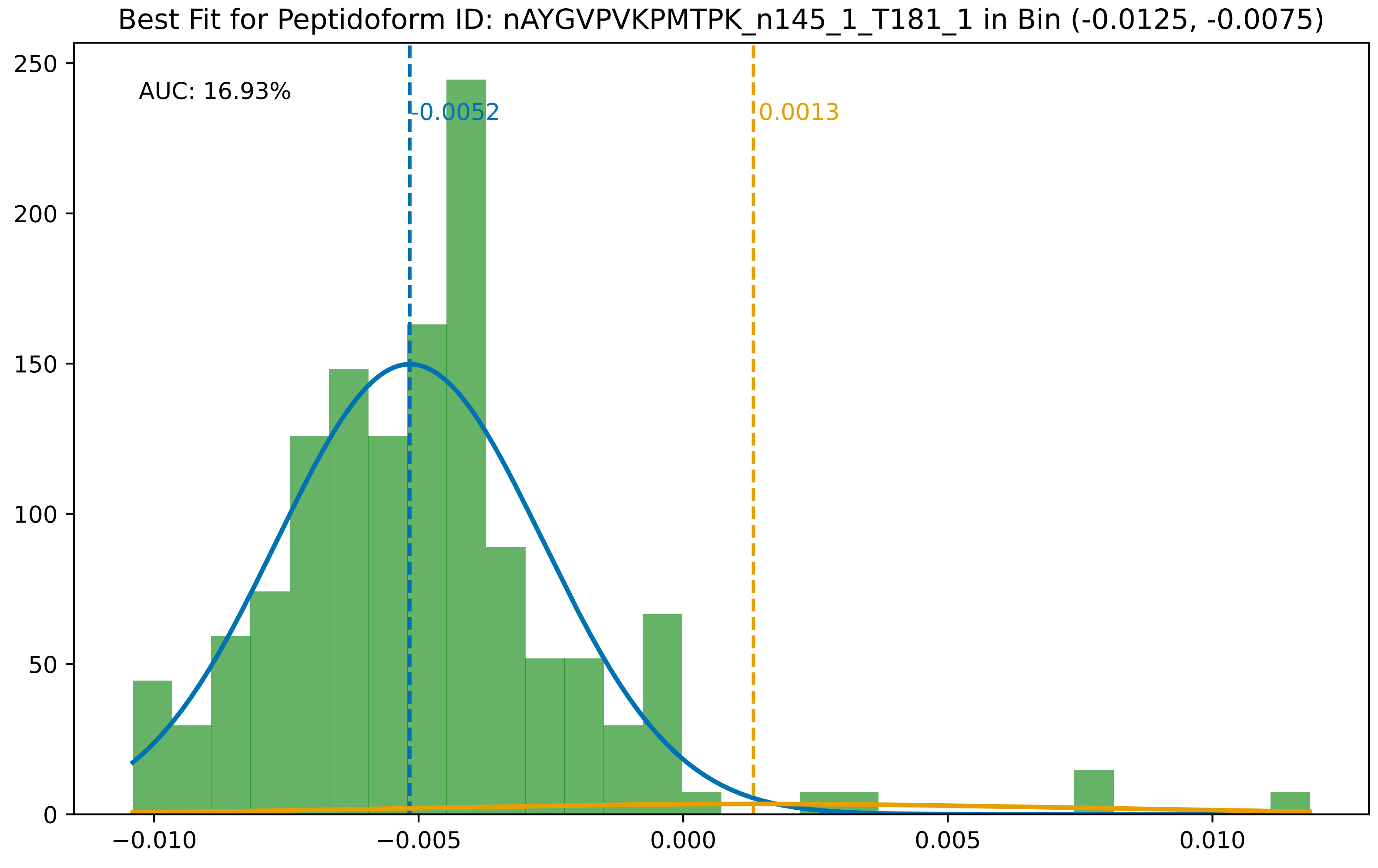

Best Fit for Peptidoform ID: nAYGVPVKPMTPK_n145_1_T181_1 in Bin (-0.0125, -0.0075)
AUC: 16.93%
-0.0052
0.0013
250
200
150
100
50
0
−0.010
−0.005
0.000
0.005
0.010

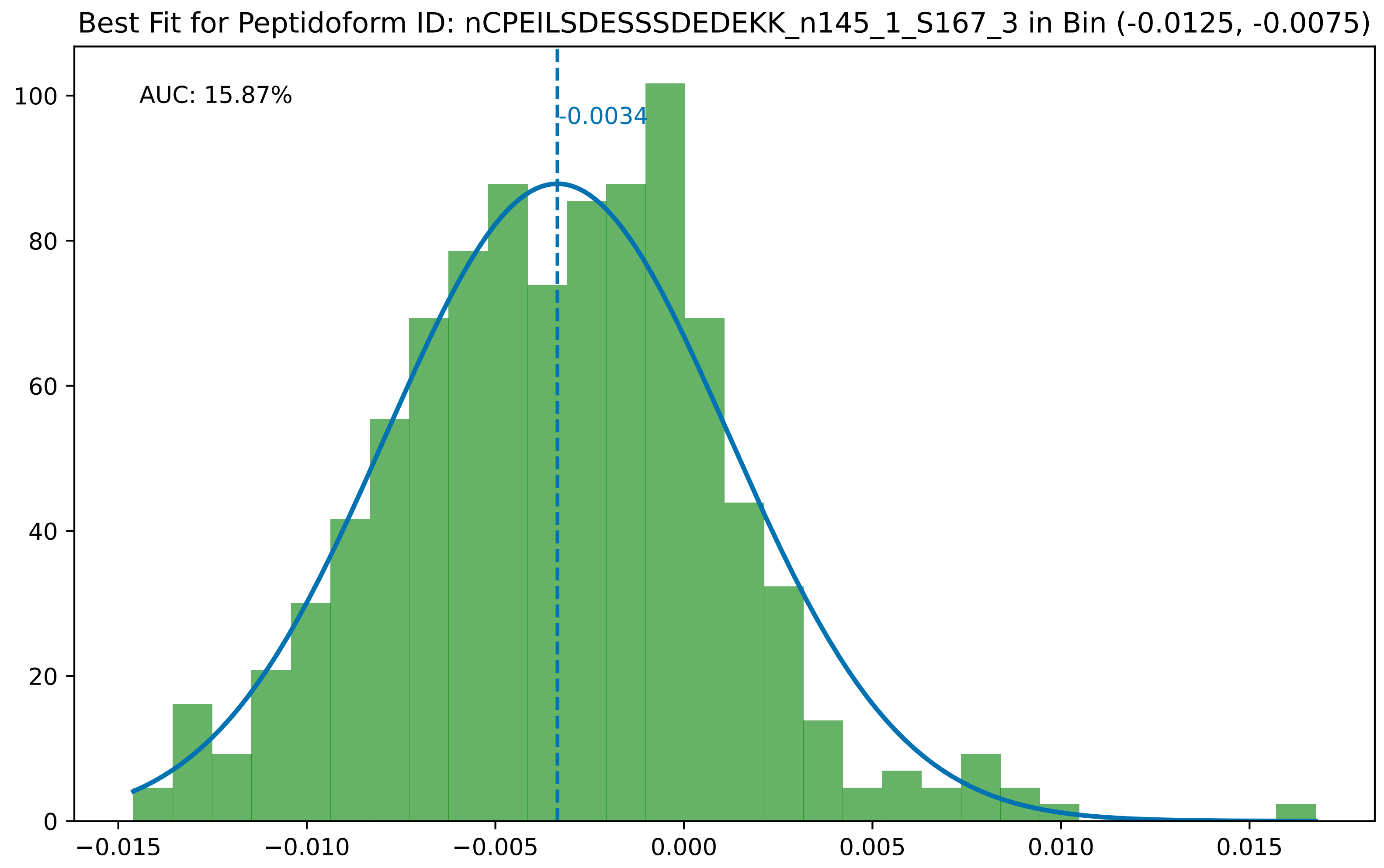

Best Fit for Peptidoform ID: nCPEILSDESSSDEDEKK_n145_1_S167_3 in Bin (-0.0125, -0.0075)
AUC: 15.87%
-0.0034
0
20
40
60
80
100
−0.015
−0.010
−0.005
0.000
0.005
0.010
0.015

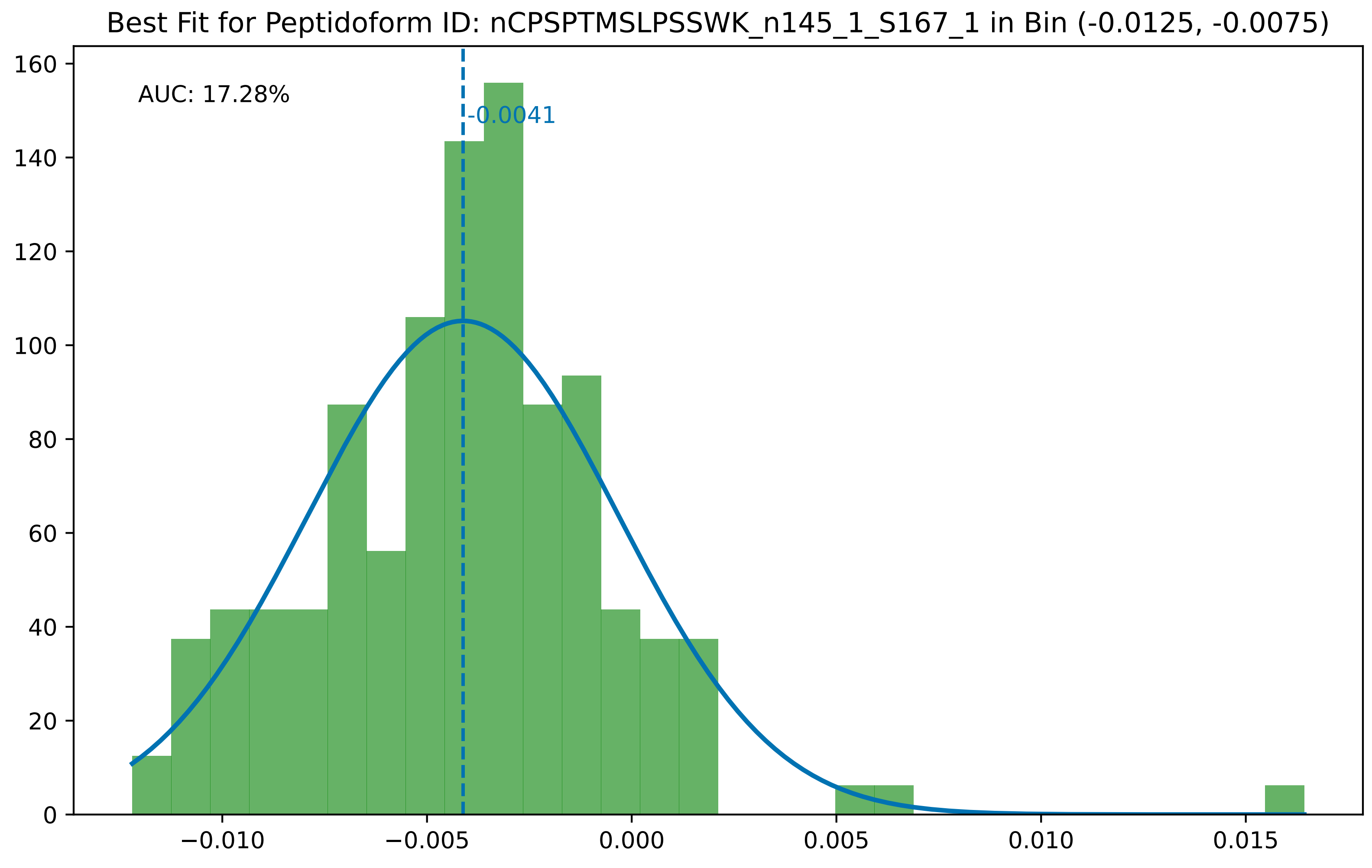

Best Fit for Peptidoform ID: nCPSPTMSLPSSWK_n145_1_S167_1 in Bin (-0.0125, -0.0075)
AUC: 17.28%
-0.0041
0
20
40
60
80
100
120
140
160
−0.010
−0.005
0.000
0.005
0.010
0.015

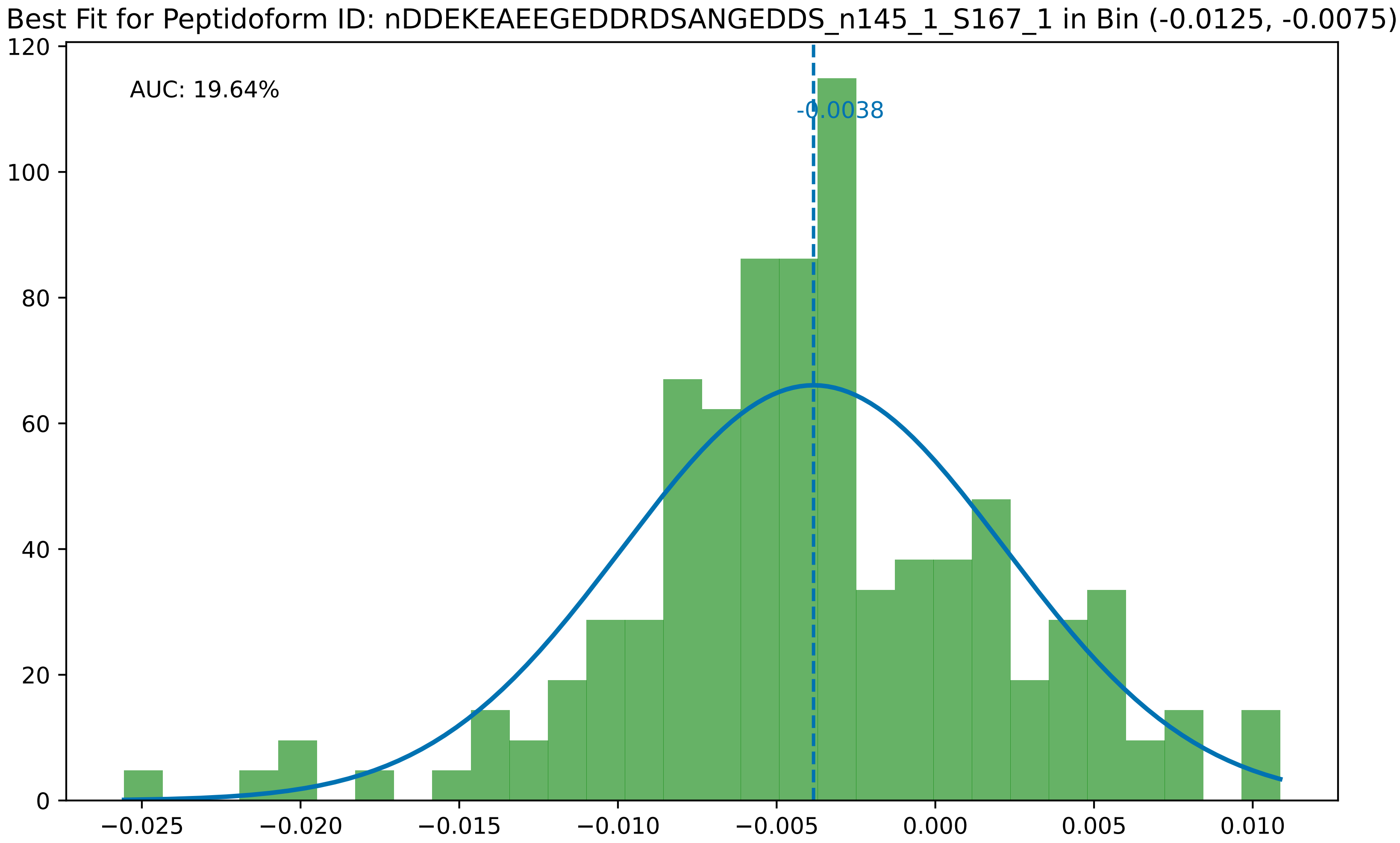

Best Fit for Peptidoform ID: nDDEKEAEEGEDDRDSANGEDDS_n145_1_S167_1 in Bin (-0.0125, -0.0075)
AUC: 19.64%
-0.0038
0
20
40
60
80
100
120
−0.025
−0.020
−0.015
−0.010
−0.005
0.000
0.005
0.010

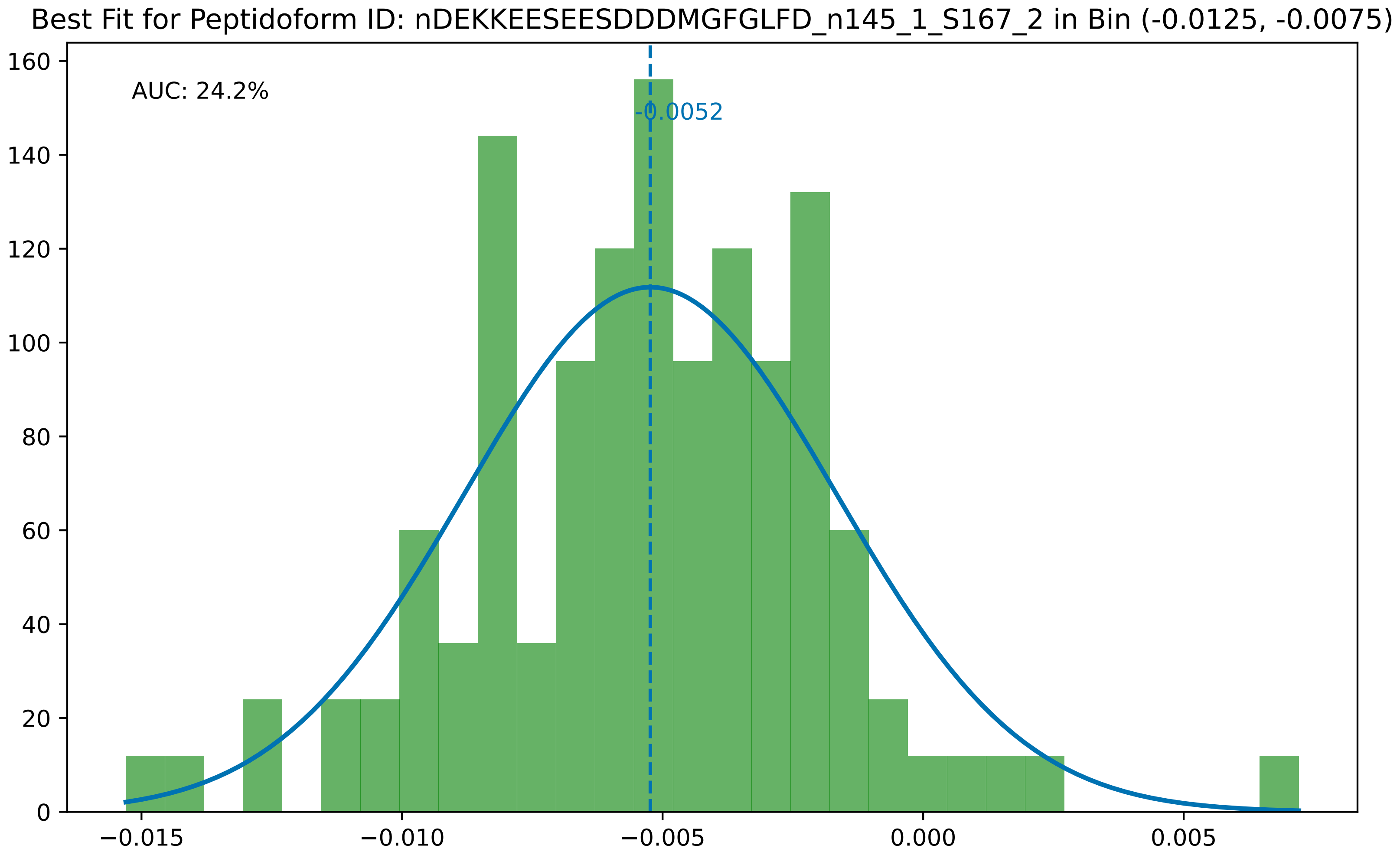
Best Fit for Peptidoform ID: nDEKKEESEESDDDMGFGLFD_n145_1_S167_2 in Bin (-0.0125, -0.0075)
AUC: 24.2%
-0.0052
0
20
40
60
80
100
120
140
160
−0.015
−0.010
−0.005
0.000
0.005

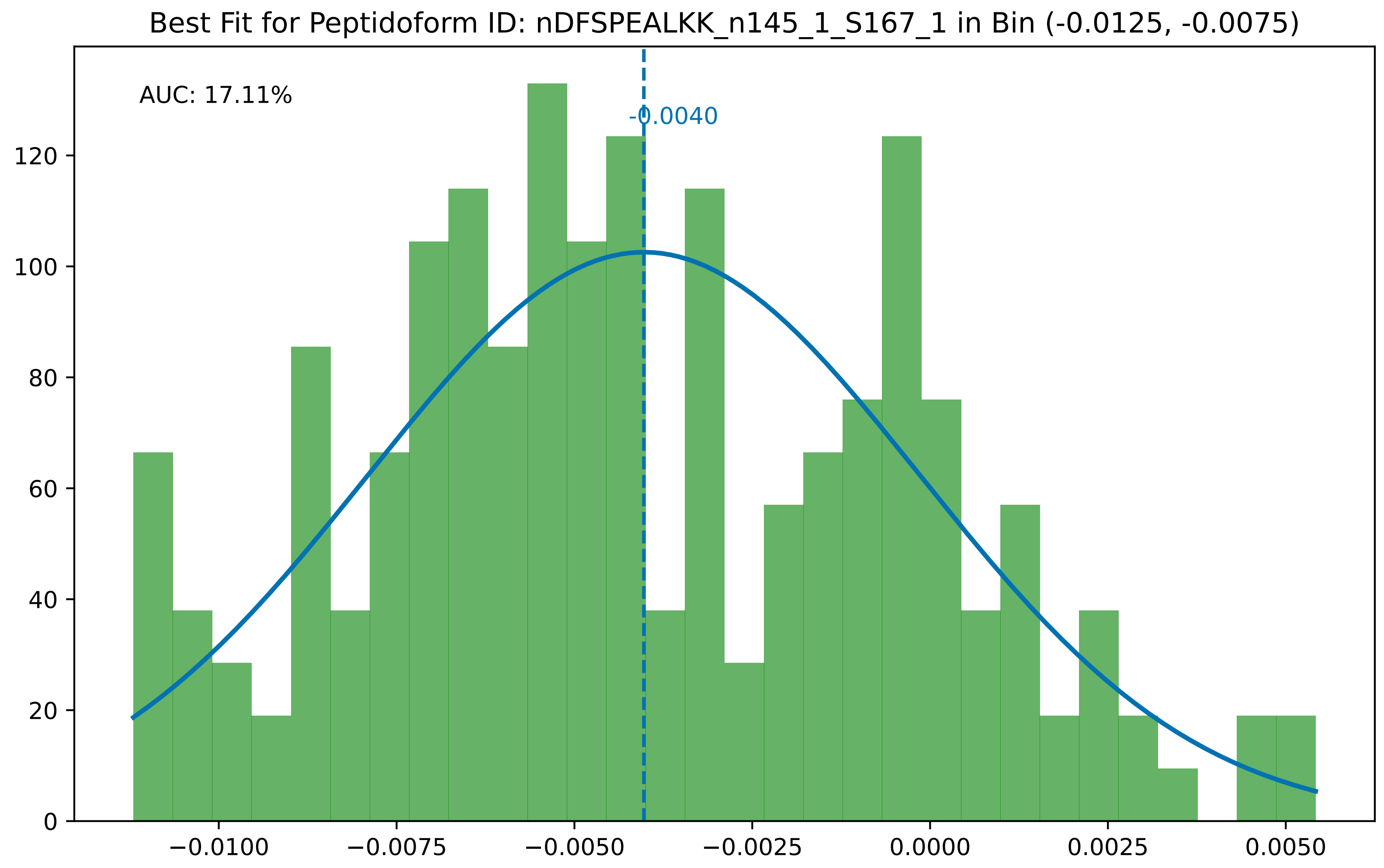

Best Fit for Peptidoform ID: nDFSPEALKK_n145_1_S167_1 in Bin (-0.0125, -0.0075)
AUC: 17.11%
-0.0040
0
20
40
60
80
100
120
−0.0100
−0.0075
−0.0050
−0.0025
0.0000
0.0025
0.0050

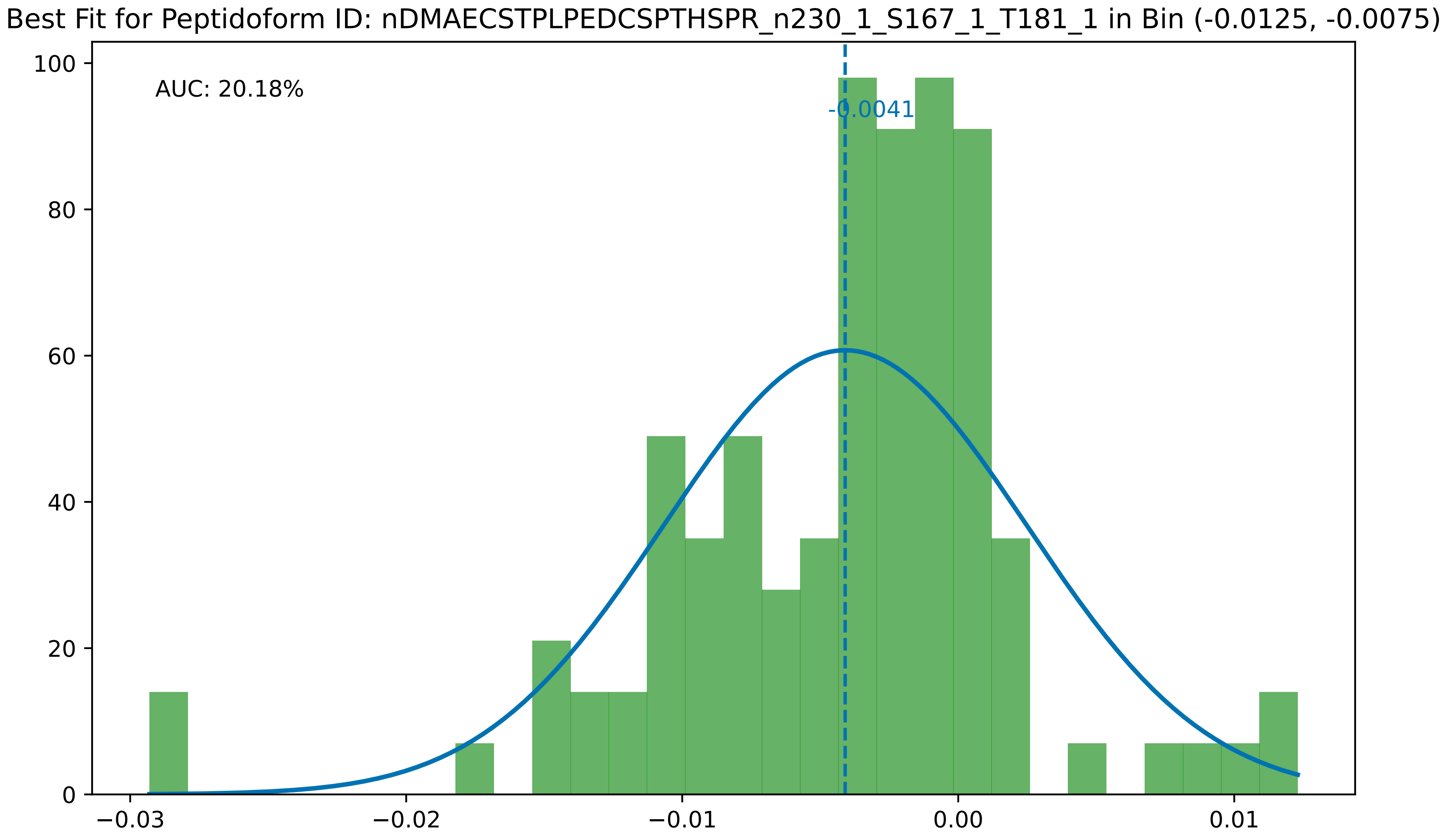

Best Fit for Peptidoform ID: nDMAECSTPLPEDCSPTHSPR_n230_1_S167_1_T181_1 in Bin (-0.0125, -0.0075)
AUC: 20.18%
-0.0041
100
80
60
40
20
0
−0.03
−0.02
−0.01
0.00
0.01

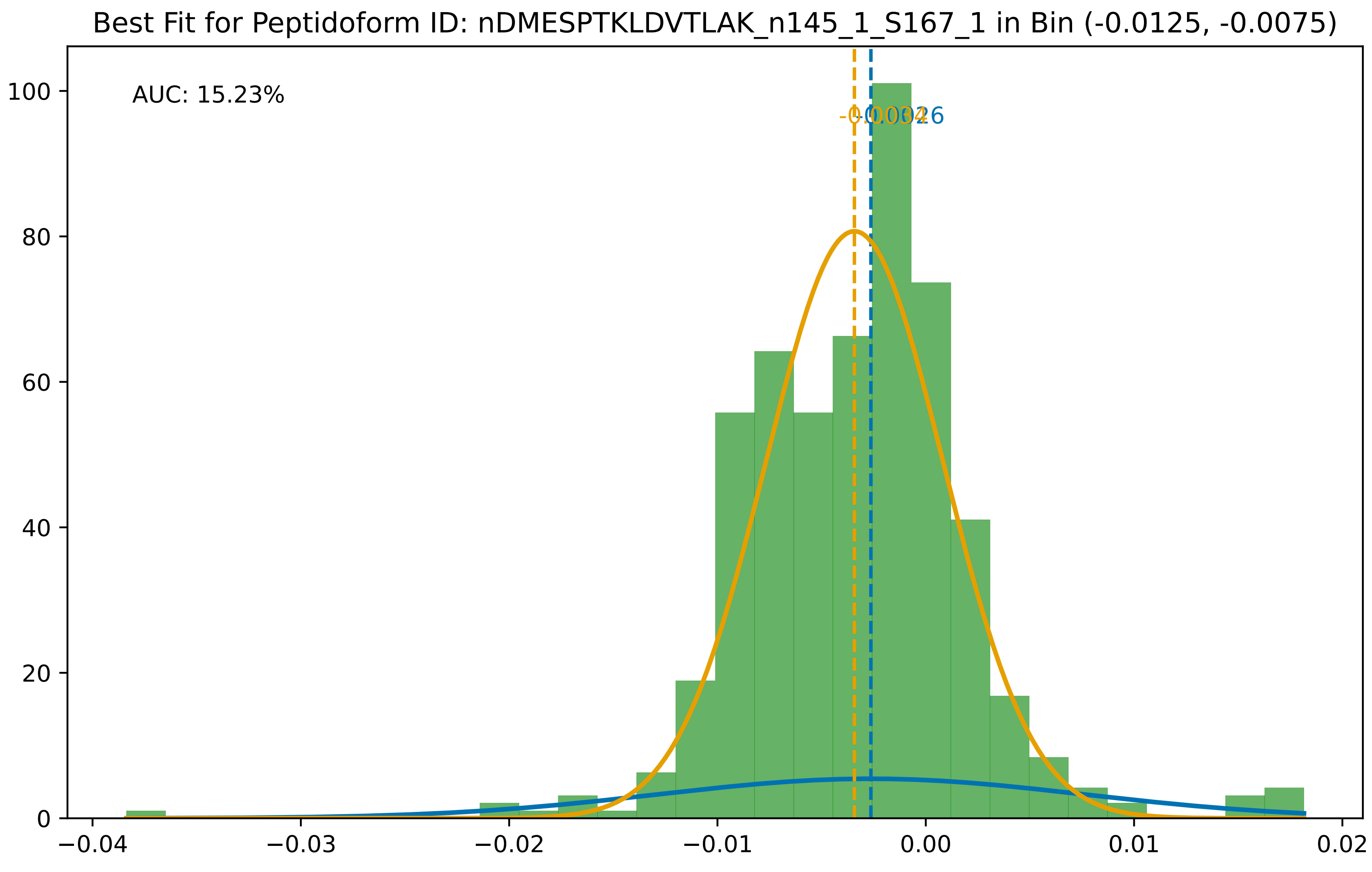

Best Fit for Peptidoform ID: nDMESPTKLDVTLAK_n145_1_S167_1 in Bin (-0.0125, -0.0075)
AUC: 15.23%
-0.0034
-0.0026
100
80
60
40
20
0
−0.04
−0.03
−0.02
−0.01
0.00
0.01
0.02

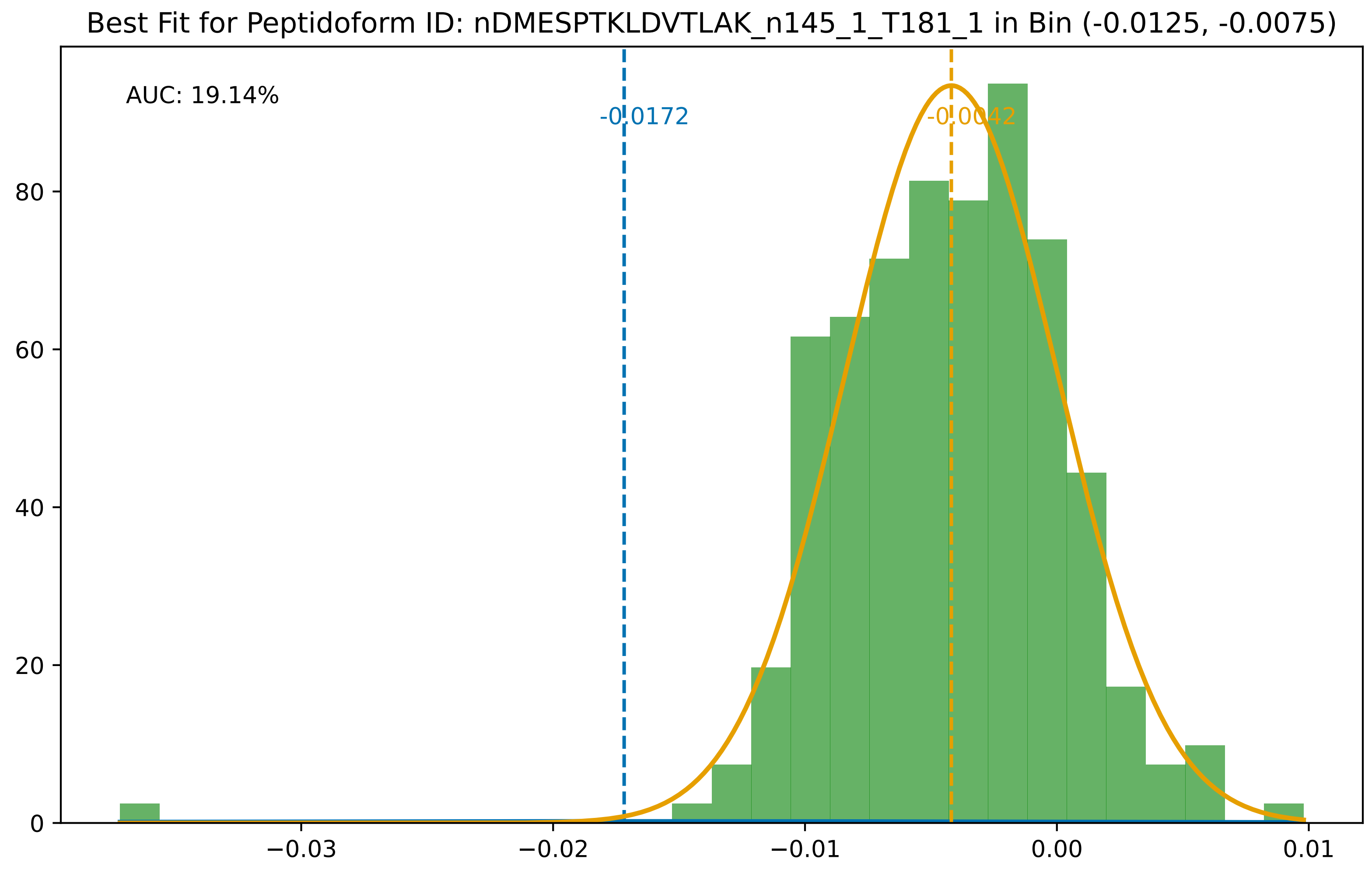

Best Fit for Peptidoform ID: nDMESPTKLDVTLAK_n145_1_T181_1 in Bin (-0.0125, -0.0075)
AUC: 19.14%
-0.0172
-0.0042
0
20
40
60
80
−0.03
−0.02
−0.01
0.00
0.01

Best Fit for Peptidoform ID: nDMSPLSETEMALGKDVTPPPETEVVLIK_n145_1_T181_1 in Bin (-0.0125, -0.0075)

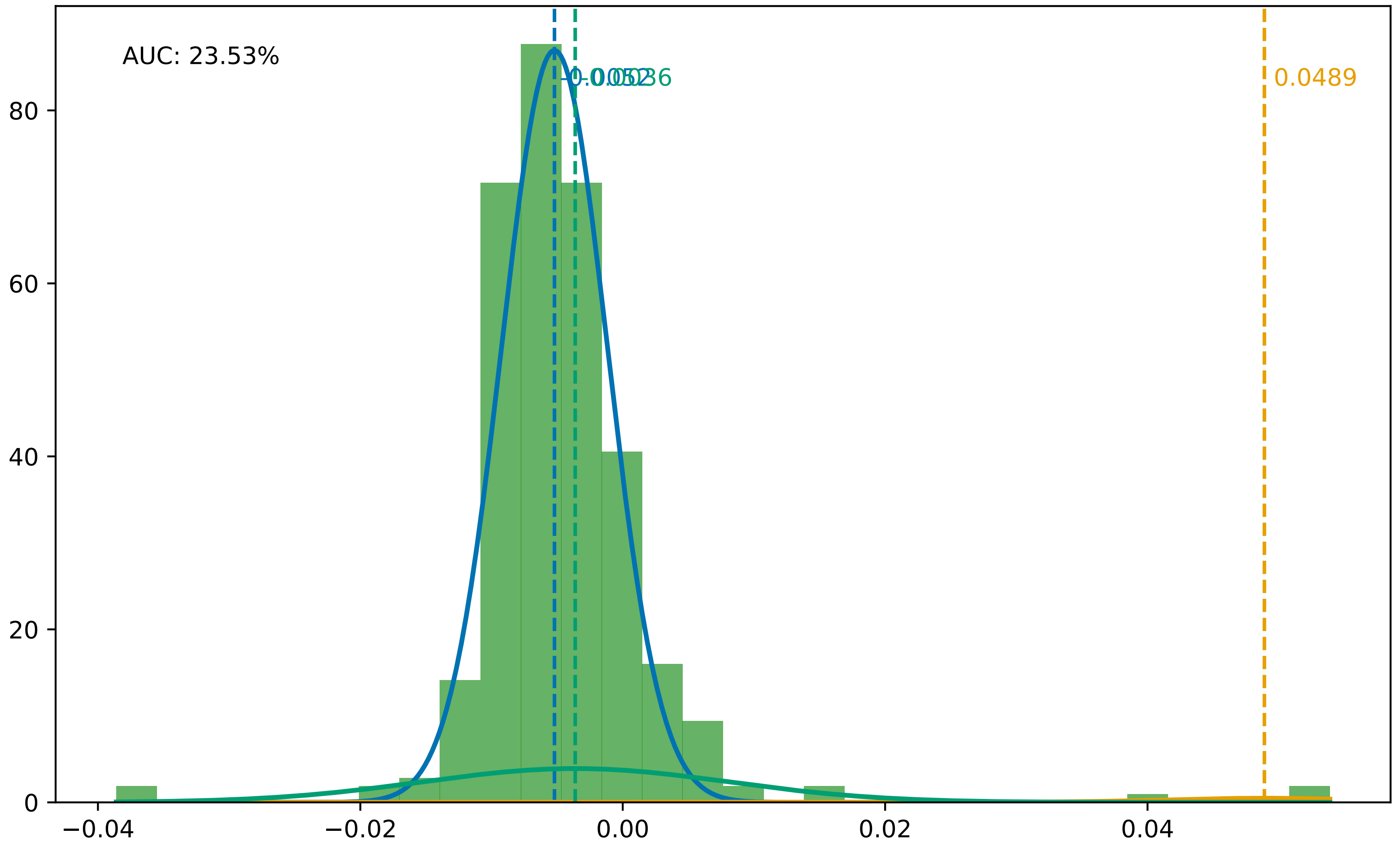

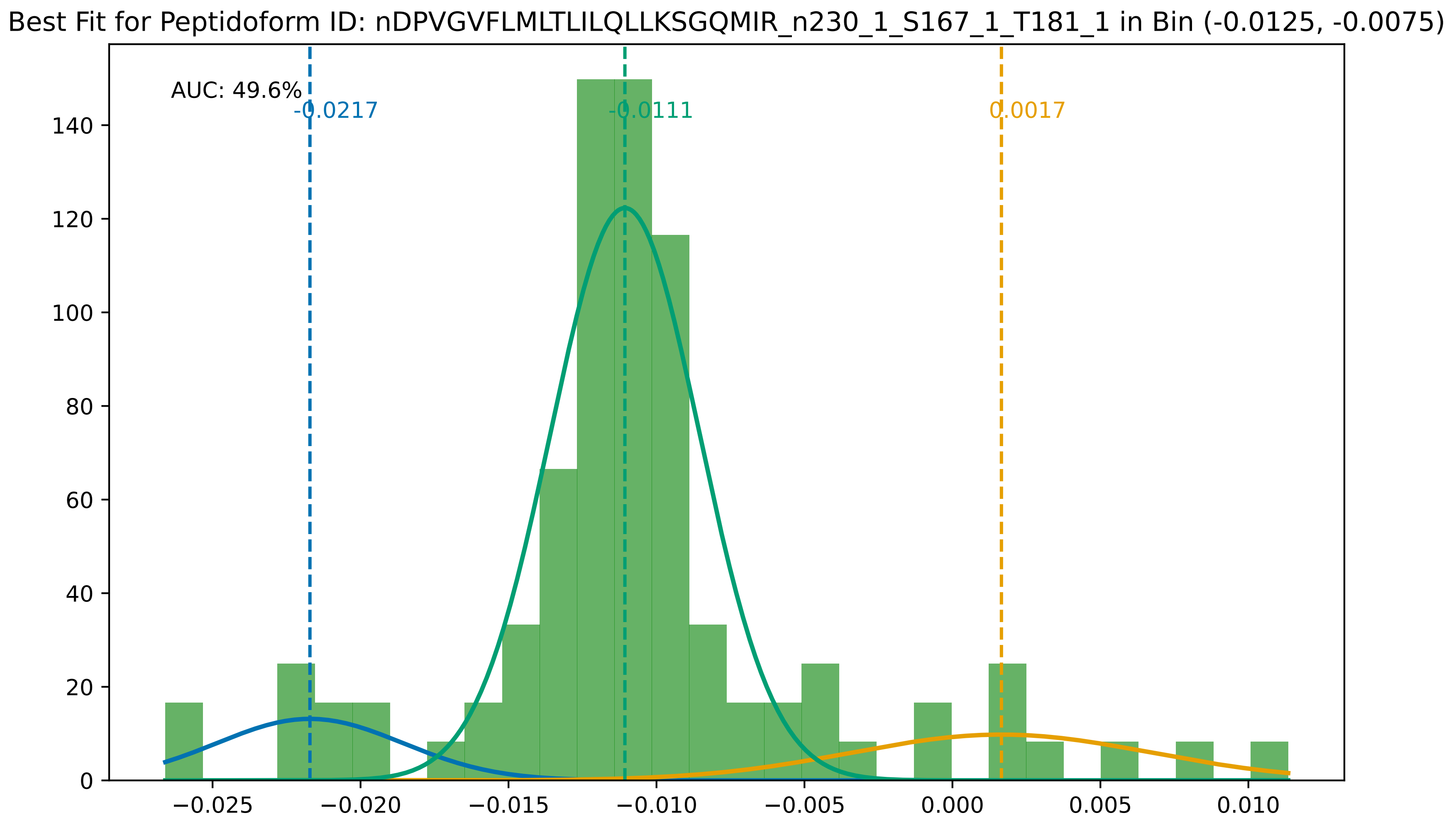

Best Fit for Peptidoform ID: nDPVGVFLMLTLILQLLKSGQMIR_n230_1_S167_1_T181_1 in Bin (-0.0125, -0.0075)
AUC: 49.6%
-0.0217
-0.0111
0.0017
0
20
40
60
80
100
120
140
−0.025
−0.020
−0.015
−0.010
−0.005
0.000
0.005
0.010

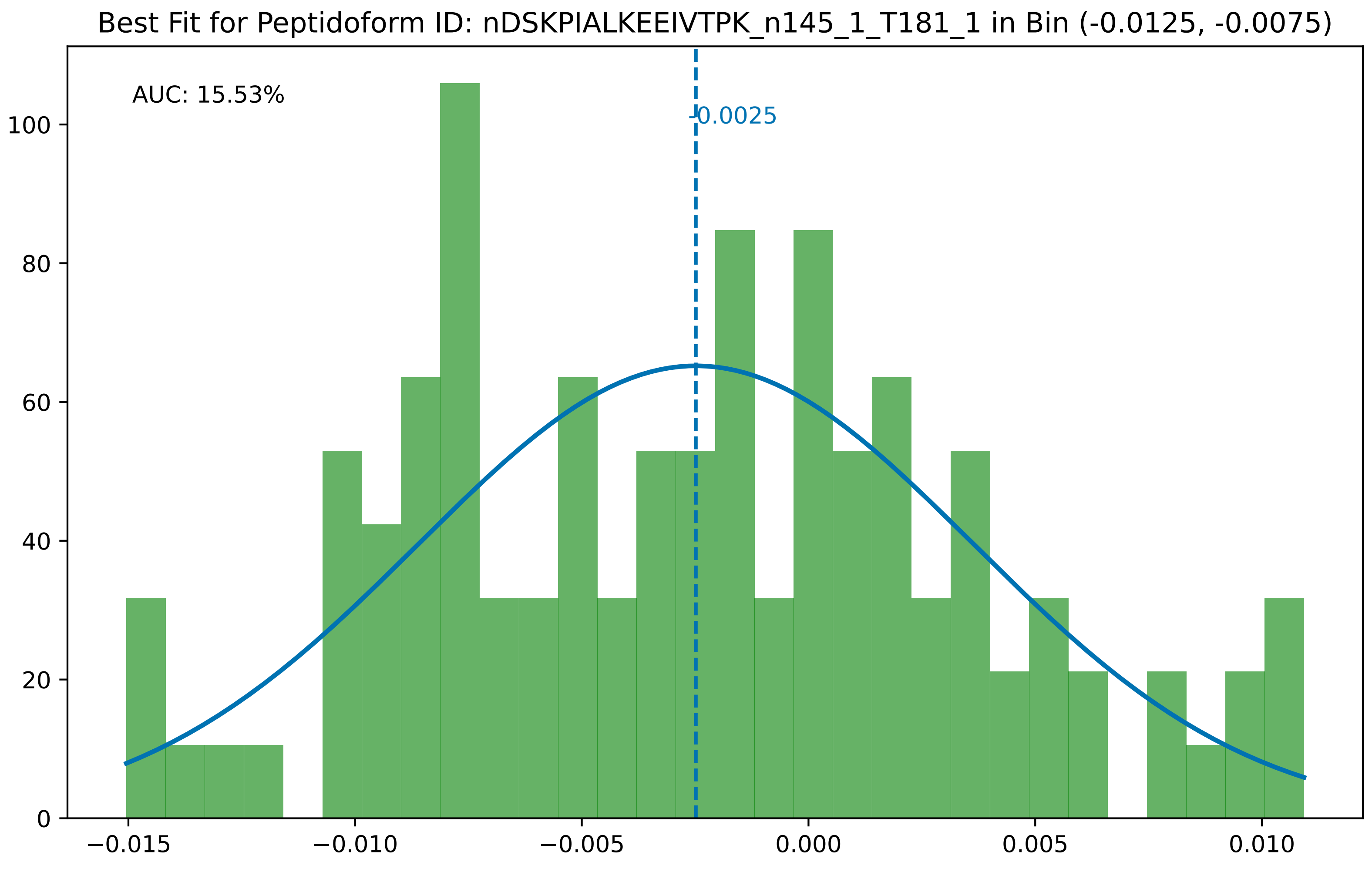

Best Fit for Peptidoform ID: nDSKPIALKEEIVTPK_n145_1_T181_1 in Bin (-0.0125, -0.0075)
AUC: 15.53%
-0.0025
0
20
40
60
80
100
−0.015
−0.010
−0.005
0.000
0.005
0.010

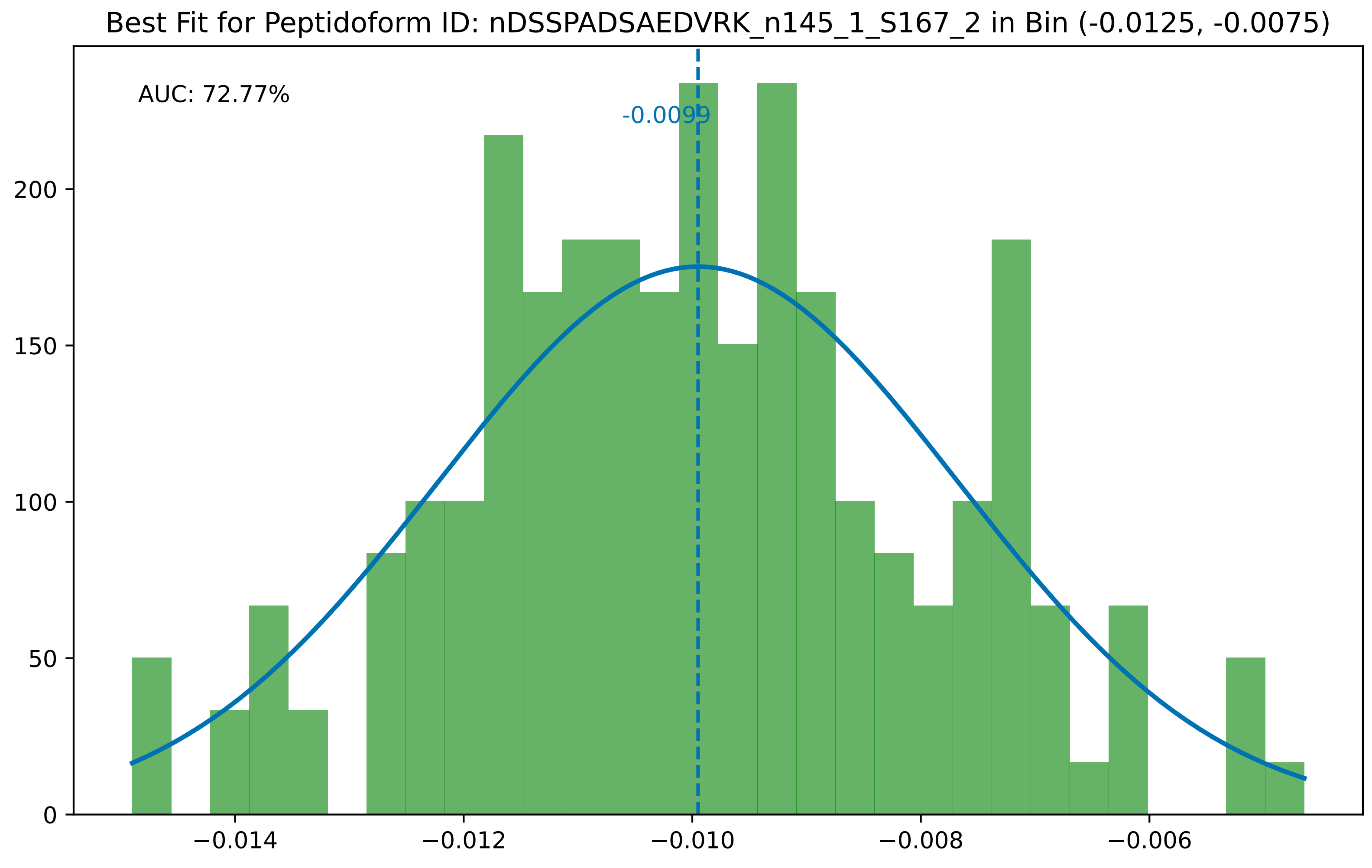

Best Fit for Peptidoform ID: nDSSPADSAEDVRK_n145_1_S167_2 in Bin (-0.0125, -0.0075)
AUC: 72.77%
-0.0099
0
50
100
150
200
−0.014
−0.012
−0.010
−0.008
−0.006

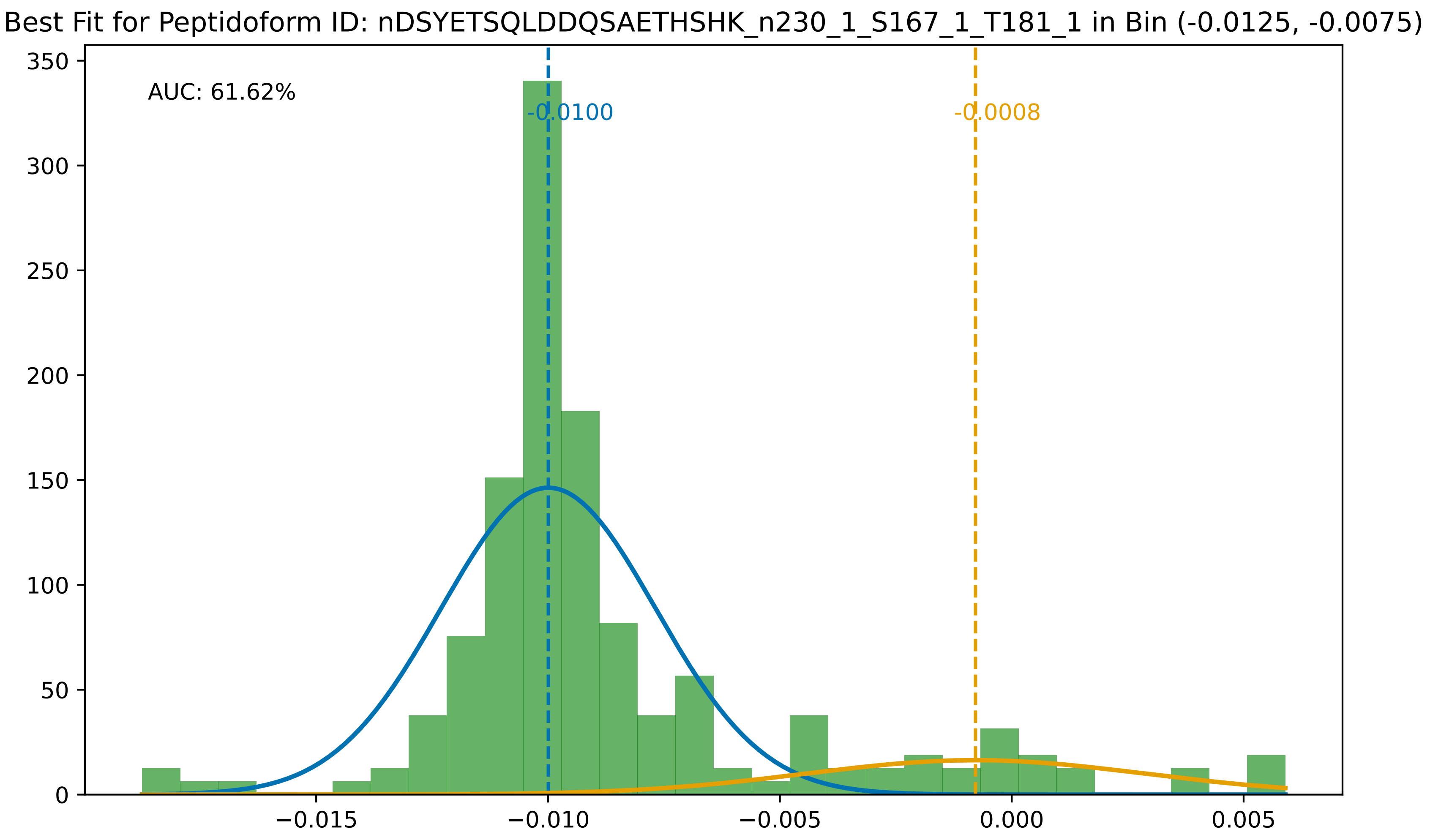
Best Fit for Peptidoform ID: nDSYETSQLDDQSAETHSHK_n230_1_S167_1_T181_1 in Bin (-0.0125, -0.0075)
AUC: 61.62%
-0.0100
-0.0008
350
300
250
200
150
100
50
0
−0.015
−0.010
−0.005
0.000
0.005

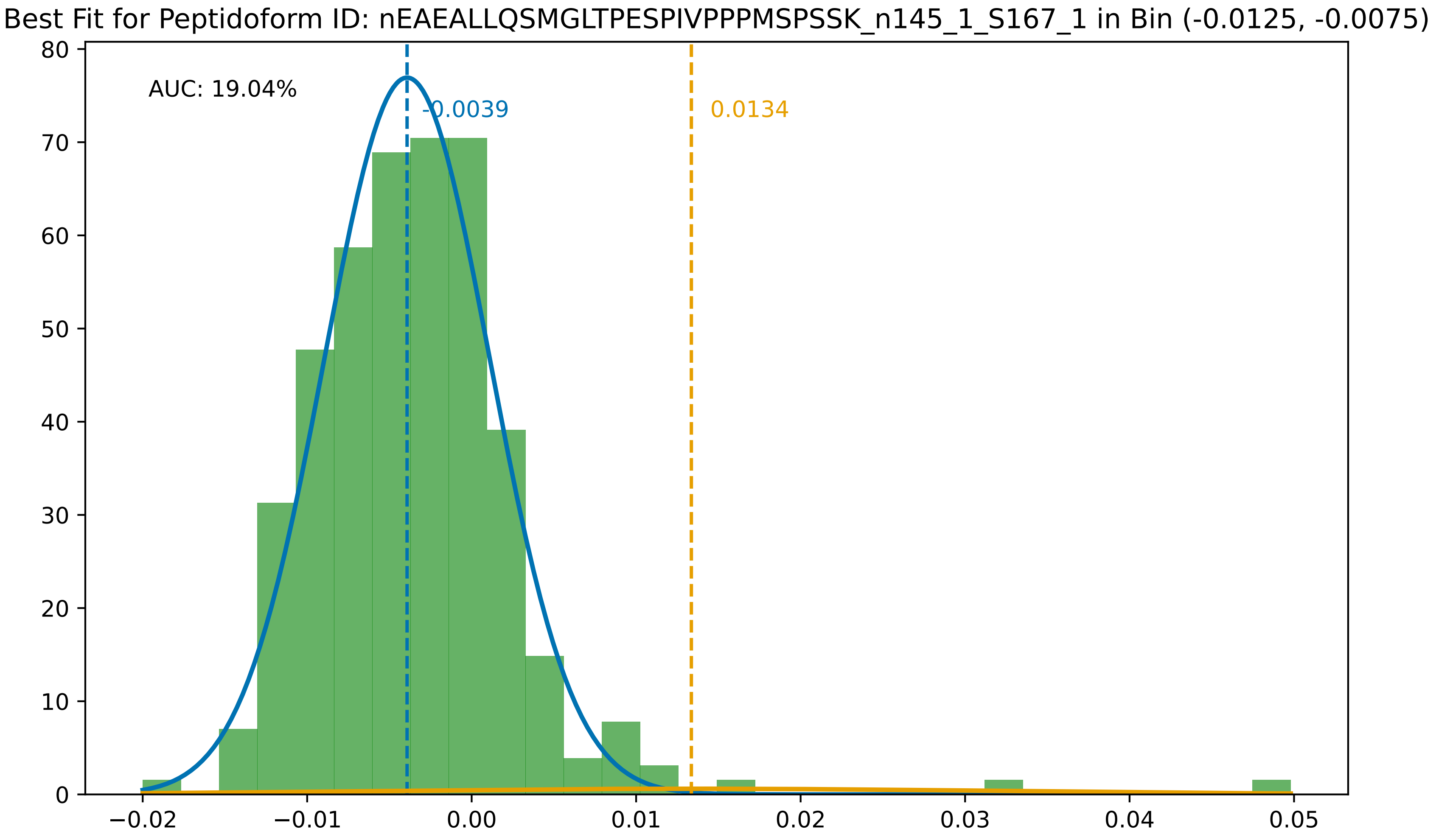

Best Fit for Peptidoform ID: nEAEALLQSMGLTPESPIVPPPMSPSSK_n145_1_S167_1 in Bin (-0.0125, -0.0075)
AUC: 19.04%
-0.0039
0.0134
80
70
60
50
40
30
20
10
0
−0.02
−0.01
0.00
0.01
0.02
0.03
0.04
0.05

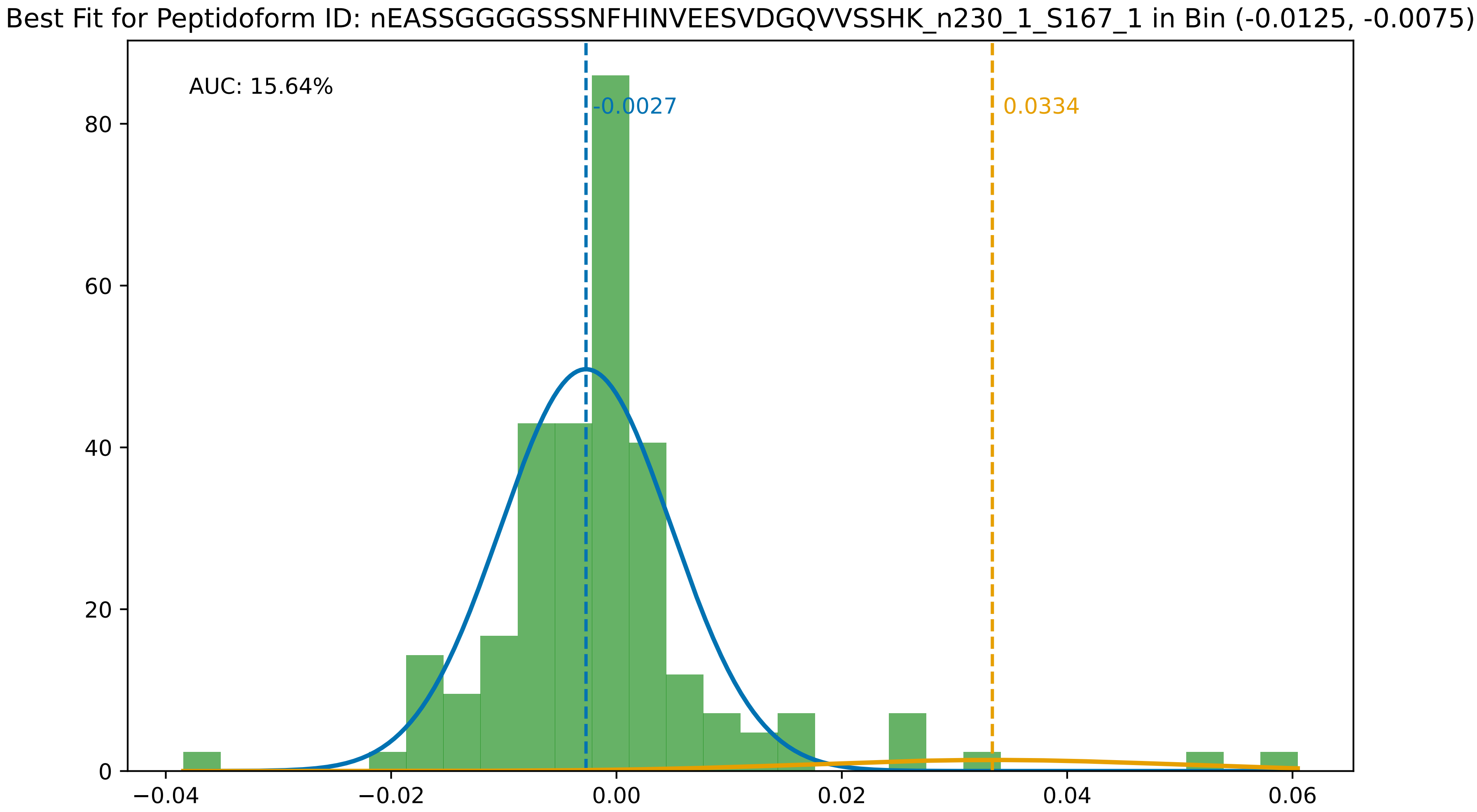

Best Fit for Peptidoform ID: nEASSGGGGSSSNFHINVEESVDGQVVSSHK_n230_1_S167_1 in Bin (-0.0125, -0.0075)
AUC: 15.64%
-0.0027
0.0334
0
20
40
60
80
−0.04
−0.02
0.00
0.02
0.04
0.06

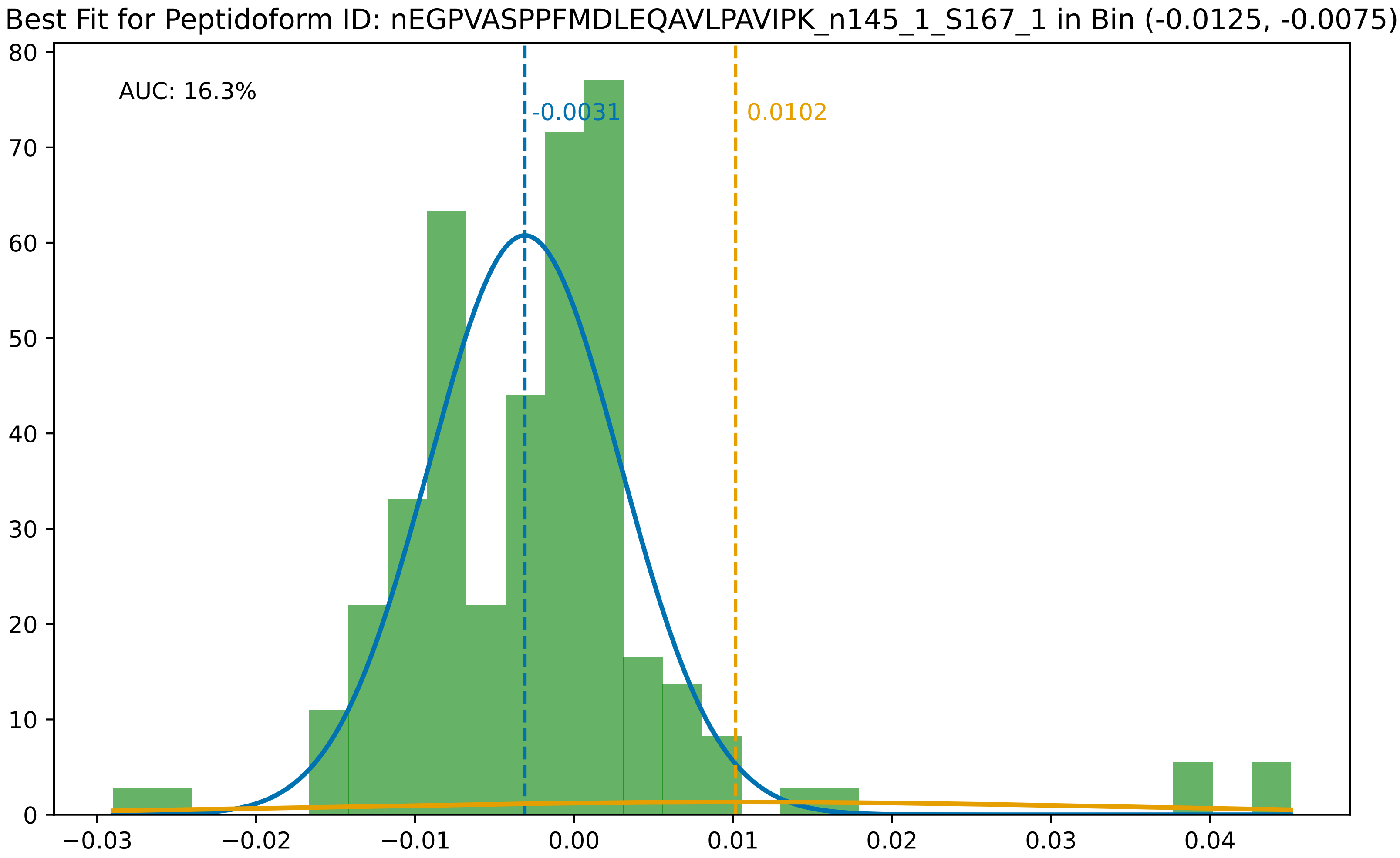
Best Fit for Peptidoform ID: nEGPVASPPFMDLEQAVLPAVIPK_n145_1_S167_1 in Bin (-0.0125, -0.0075)
AUC: 16.3%
-0.0031
0.0102
0
10
20
30
40
50
60
70
80
−0.03
−0.02
−0.01
0.00
0.01
0.02
0.03
0.04

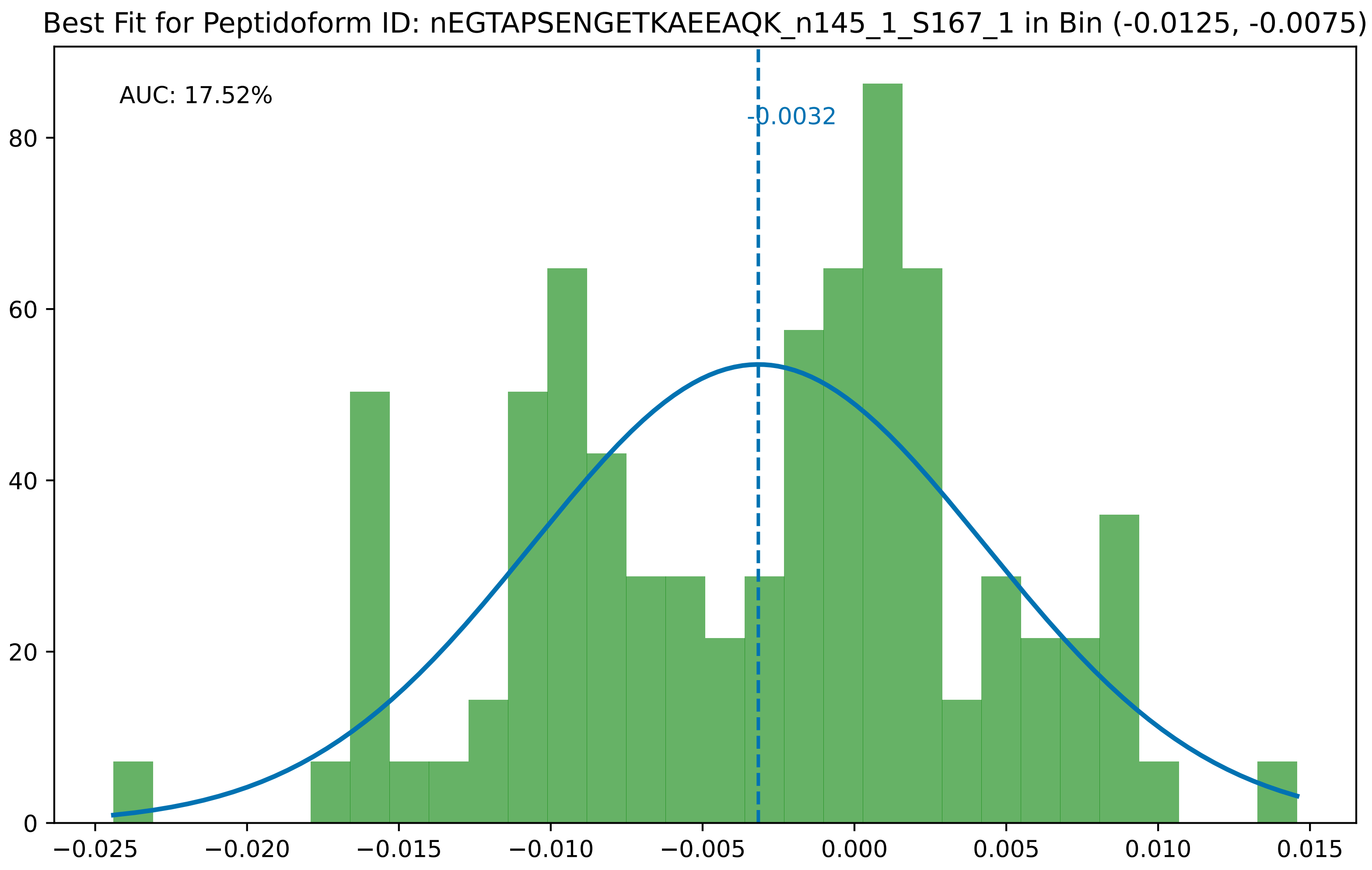
Best Fit for Peptidoform ID: nEGTAPSENGETKAEEAQK_n145_1_S167_1 in Bin (-0.0125, -0.0075)
AUC: 17.52%
-0.0032
0
20
40
60
80
−0.025
−0.020
−0.015
−0.010
−0.005
0.000
0.005
0.010
0.015

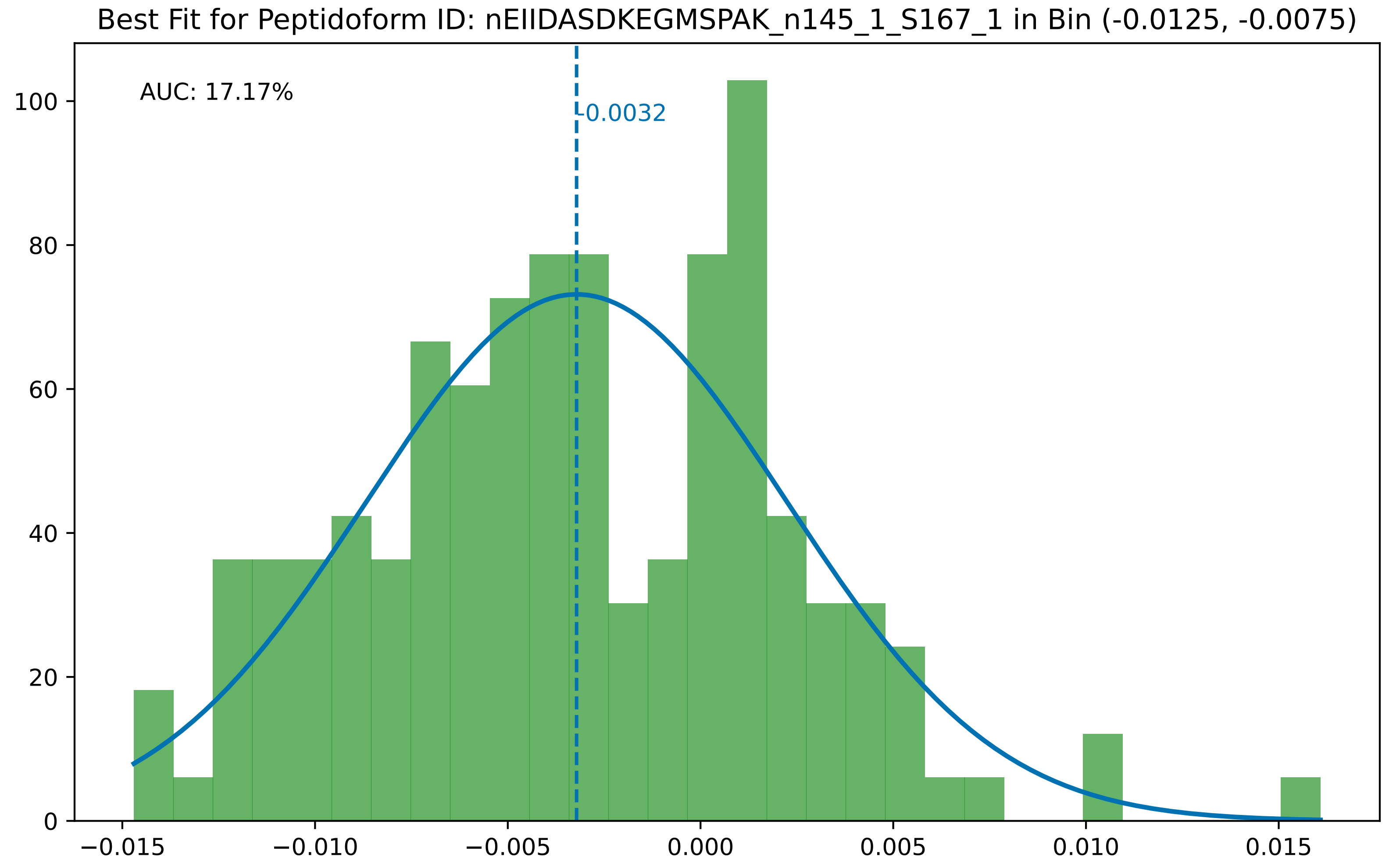
Best Fit for Peptidoform ID: nEIIDASDKEGMSPAK_n145_1_S167_1 in Bin (-0.0125, -0.0075)
AUC: 17.17%
-0.0032
0
20
40
60
80
100
−0.015
−0.010
−0.005
0.000
0.005
0.010
0.015

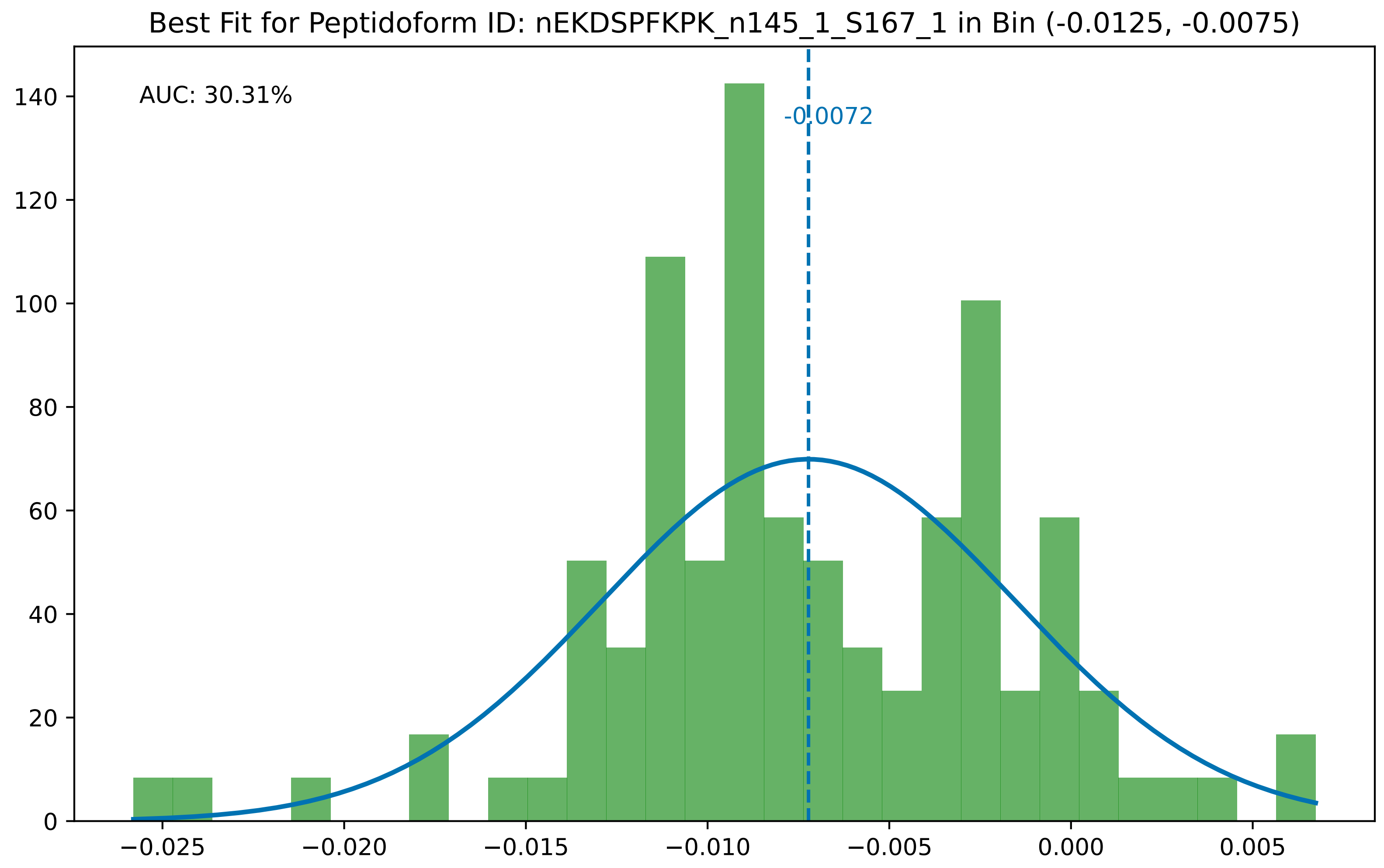

Best Fit for Peptidoform ID: nEKDSPFKPK_n145_1_S167_1 in Bin (-0.0125, -0.0075)
AUC: 30.31%
-0.0072
0
20
40
60
80
100
120
140
−0.025
−0.020
−0.015
−0.010
−0.005
0.000
0.005

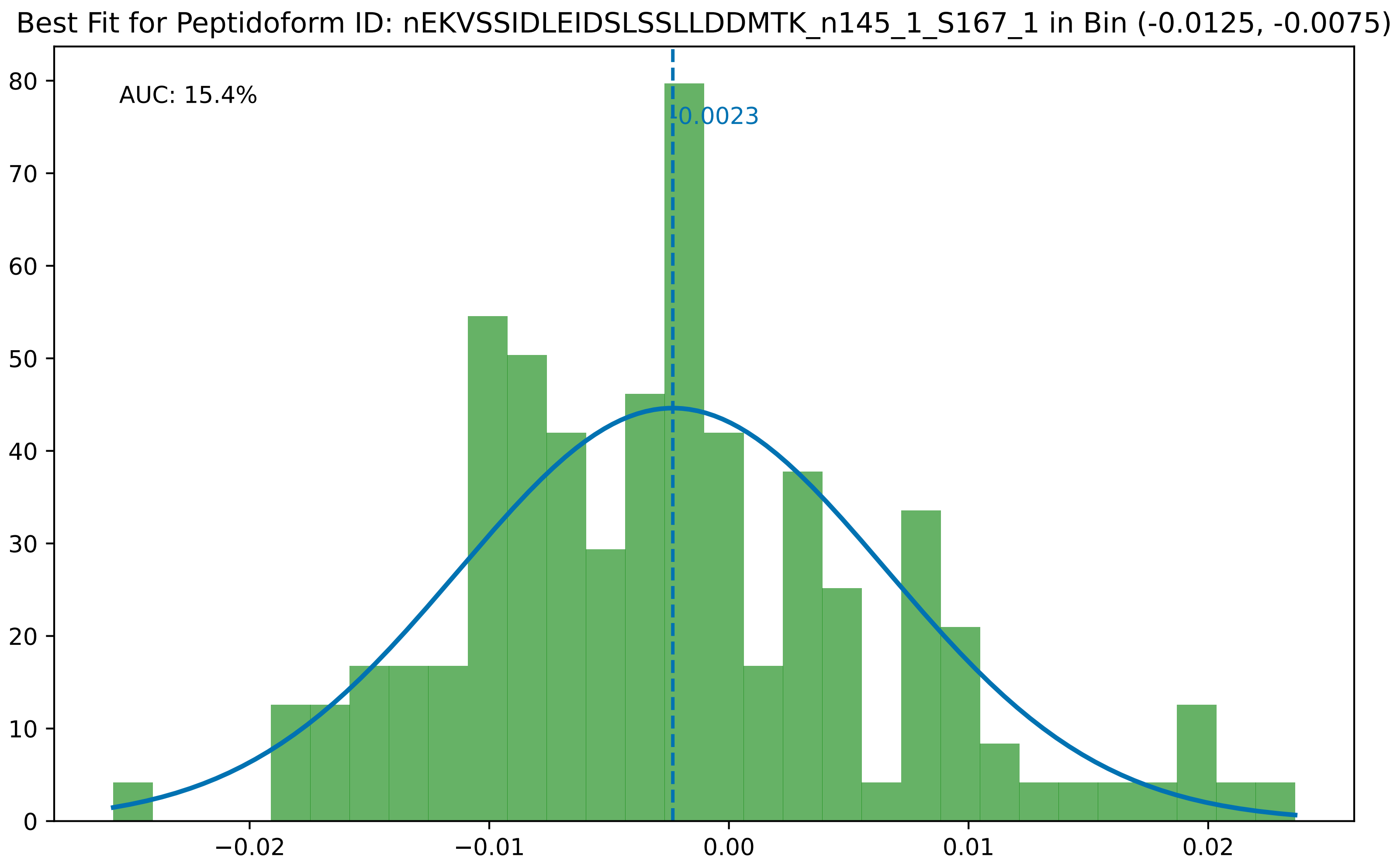

Best Fit for Peptidoform ID: nEKVSSIDLEIDSLSSLLDDMTK_n145_1_S167_1 in Bin (-0.0125, -0.0075)
AUC: 15.4%
-0.0023
0
10
20
30
40
50
60
70
80
−0.02
−0.01
0.00
0.01
0.02

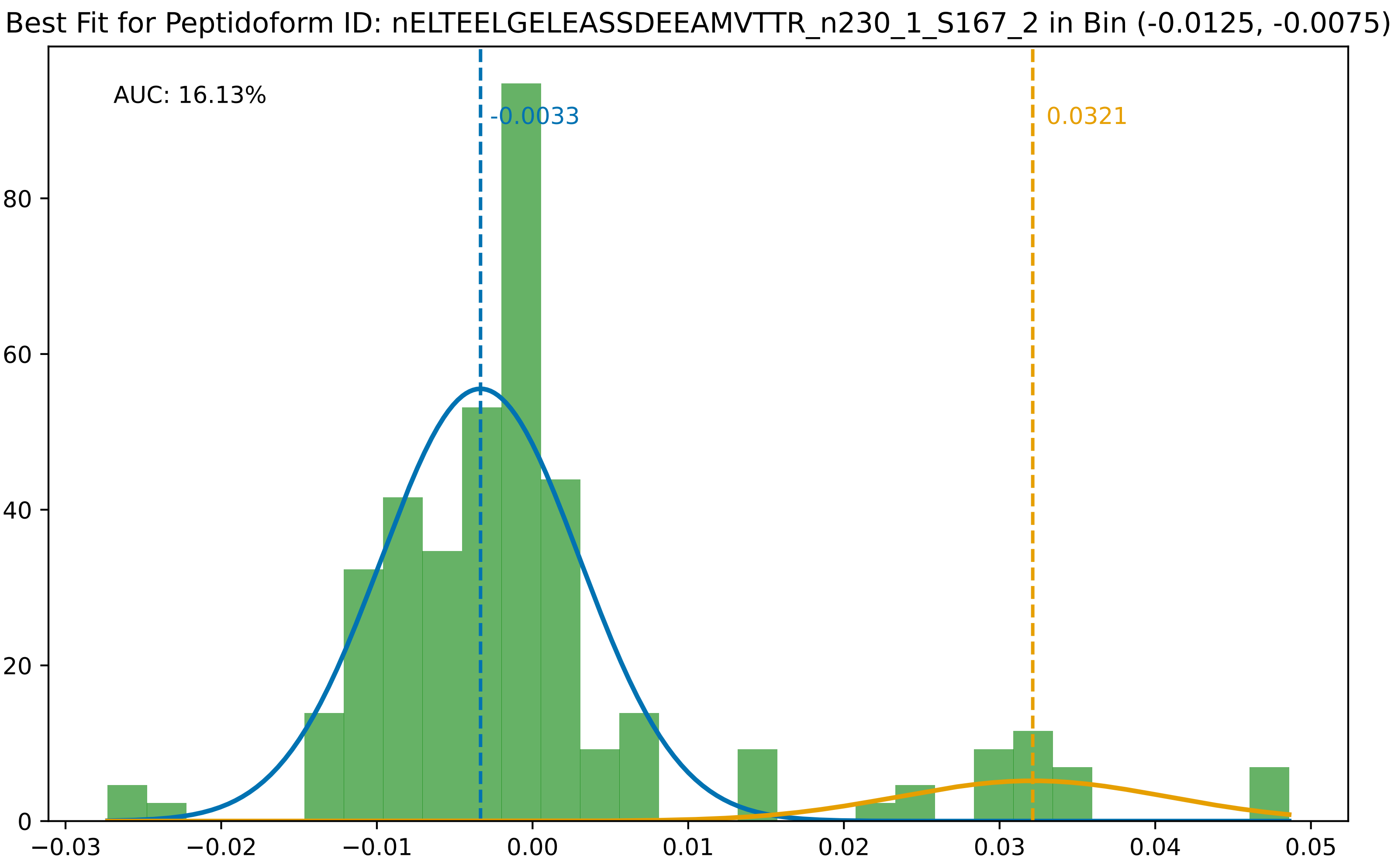

Best Fit for Peptidoform ID: nELTEELGELEASSDEEAMVTTR_n230_1_S167_2 in Bin (-0.0125, -0.0075)
AUC: 16.13%
-0.0033
0.0321
0
20
40
60
80
−0.03
−0.02
−0.01
0.00
0.01
0.02
0.03
0.04
0.05

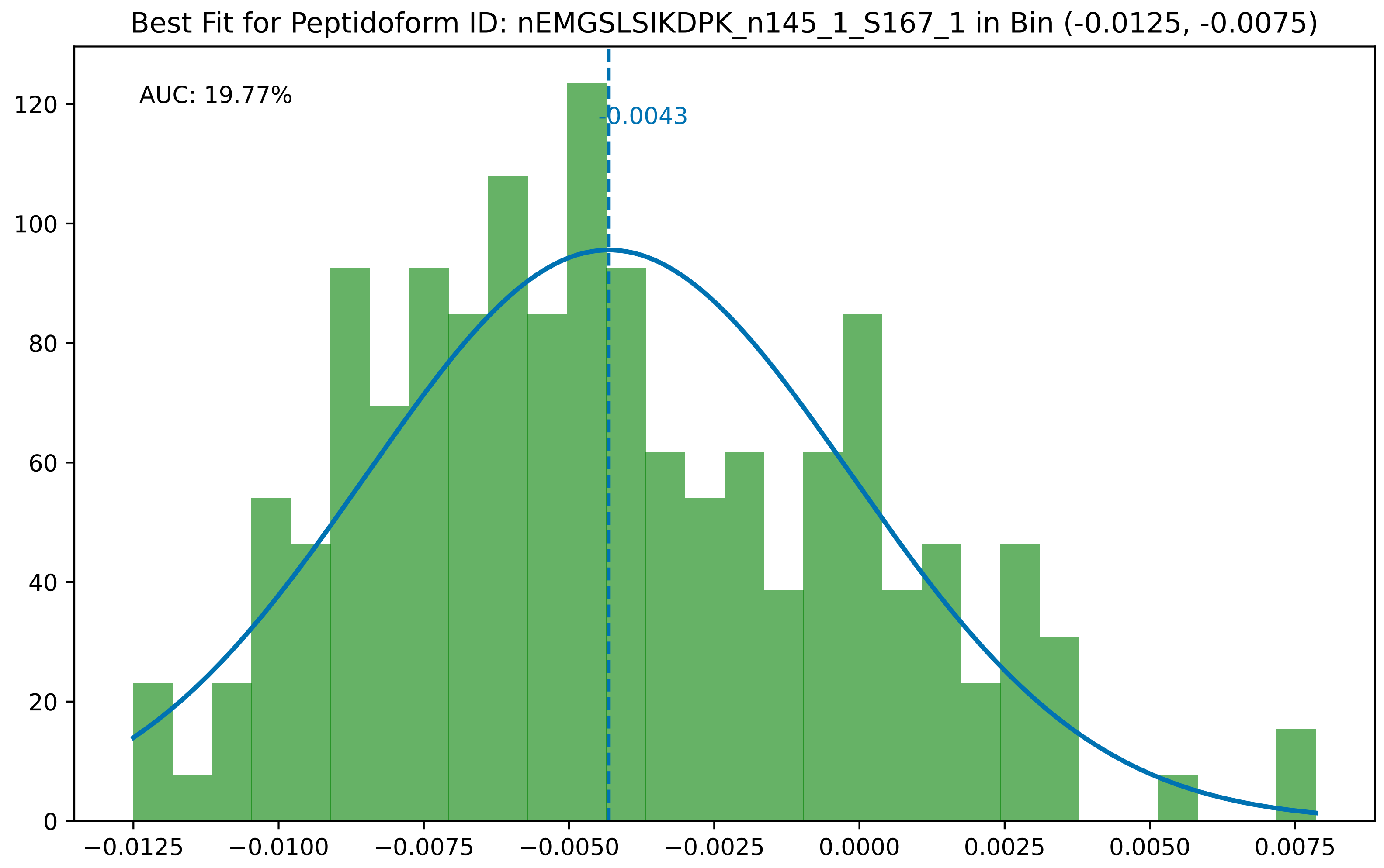

Best Fit for Peptidoform ID: nEMGSLSIKDPK_n145_1_S167_1 in Bin (-0.0125, -0.0075)
AUC: 19.77%
-0.0043
0
20
40
60
80
100
120
−0.0125
−0.0100
−0.0075
−0.0050
−0.0025
0.0000
0.0025
0.0050
0.0075

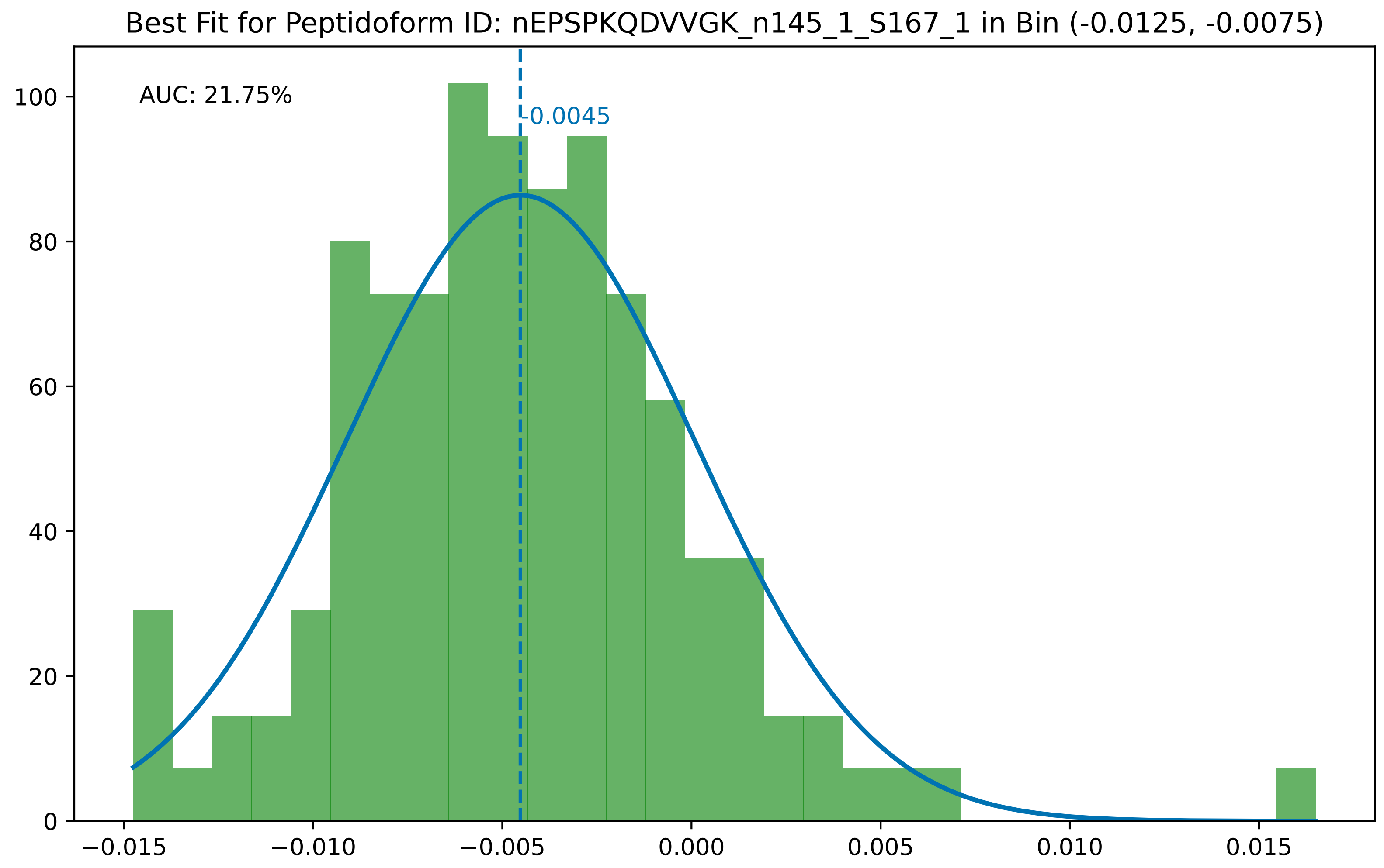

Best Fit for Peptidoform ID: nEPSPKQDVVGK_n145_1_S167_1 in Bin (-0.0125, -0.0075)
AUC: 21.75%
-0.0045
0
20
40
60
80
100
−0.015
−0.010
−0.005
0.000
0.005
0.010
0.015

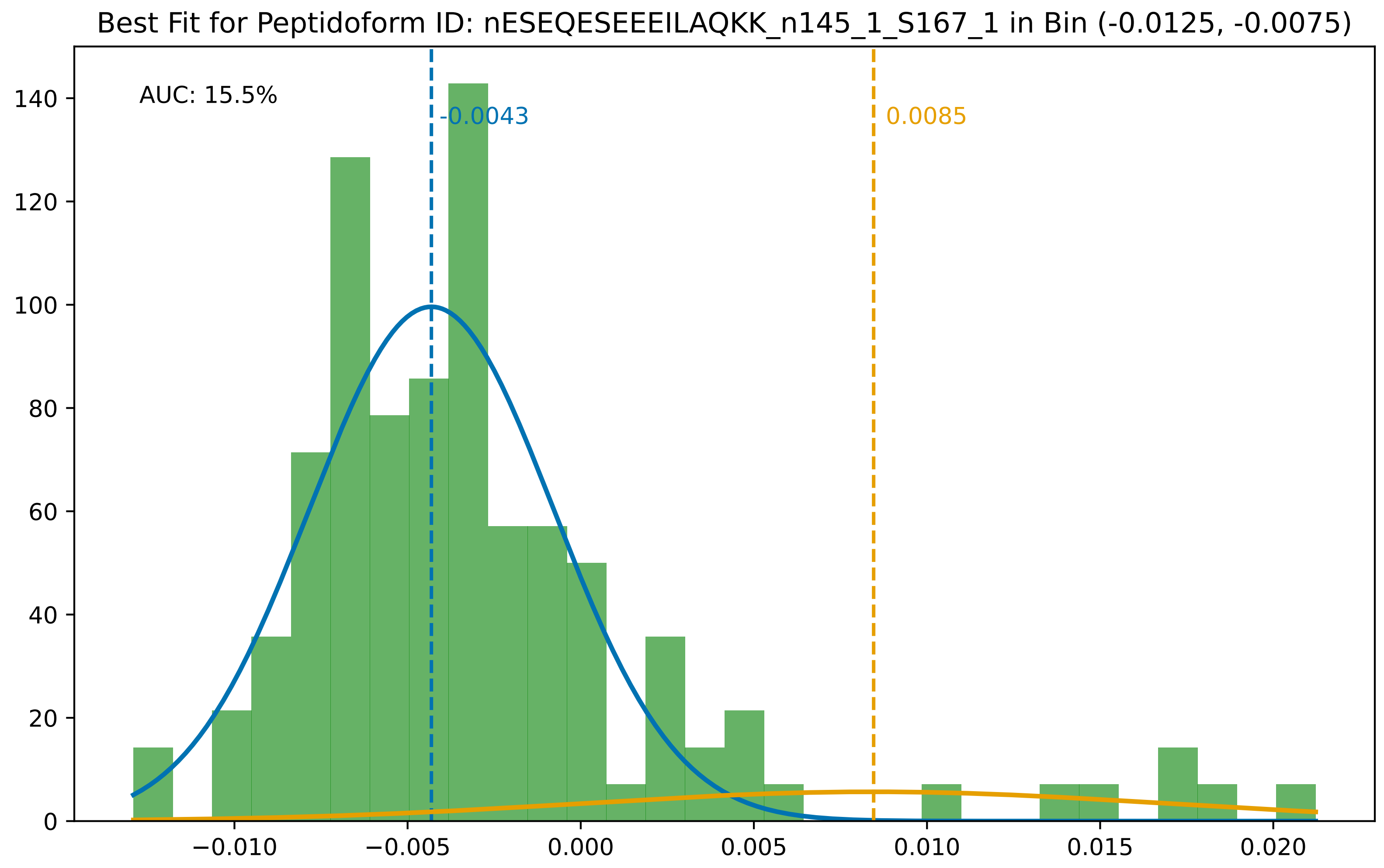

Best Fit for Peptidoform ID: nESEQESEEEILAQKK_n145_1_S167_1 in Bin (-0.0125, -0.0075)
AUC: 15.5%
-0.0043
0.0085
0
20
40
60
80
100
120
140
−0.010
−0.005
0.000
0.005
0.010
0.015
0.020

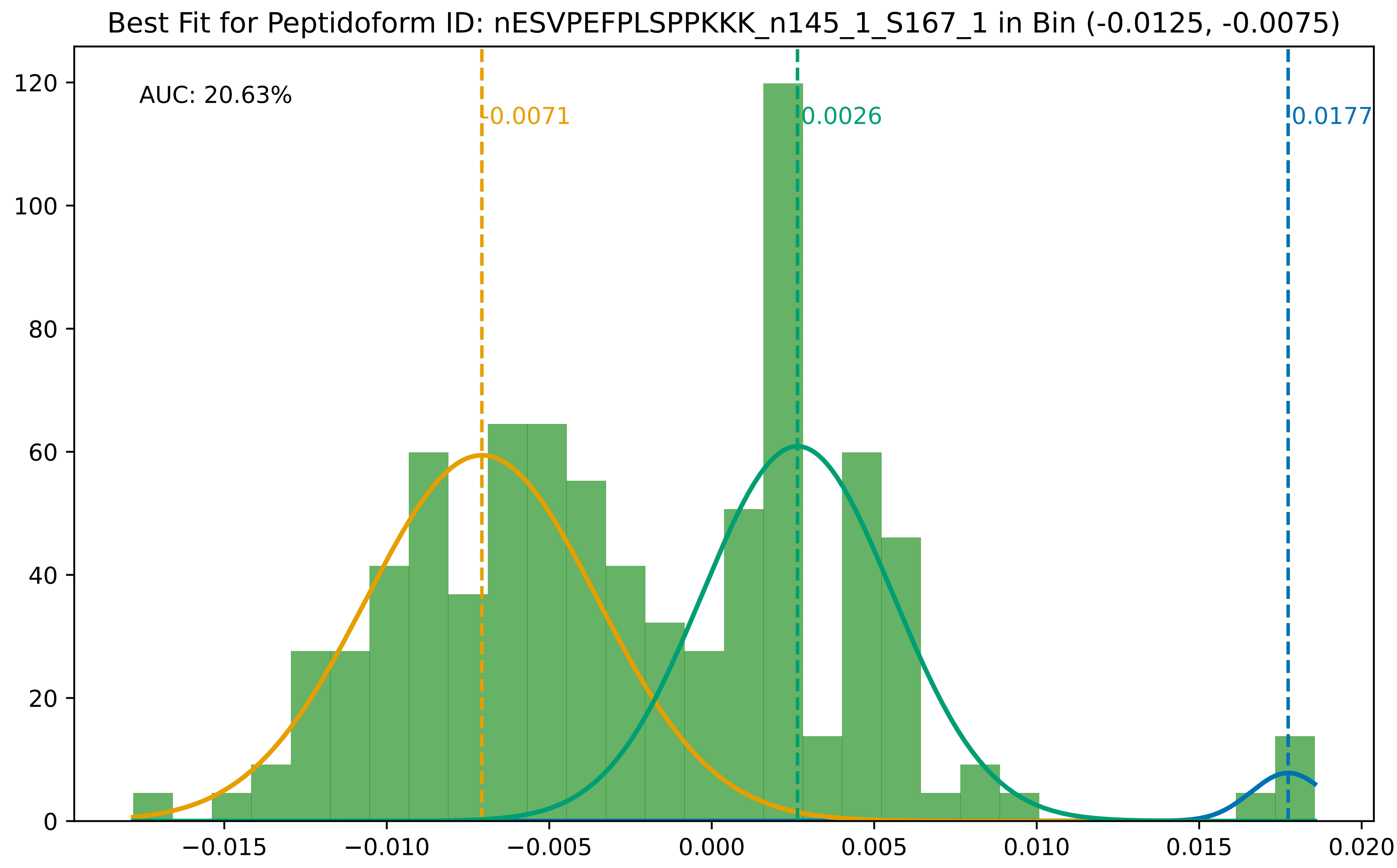

Best Fit for Peptidoform ID: nESVPEFPLSPPKKK_n145_1_S167_1 in Bin (-0.0125, -0.0075)
AUC: 20.63%
-0.0071
0.0026
0.0177
0
20
40
60
80
100
120
−0.015
−0.010
−0.005
0.000
0.005
0.010
0.015
0.020

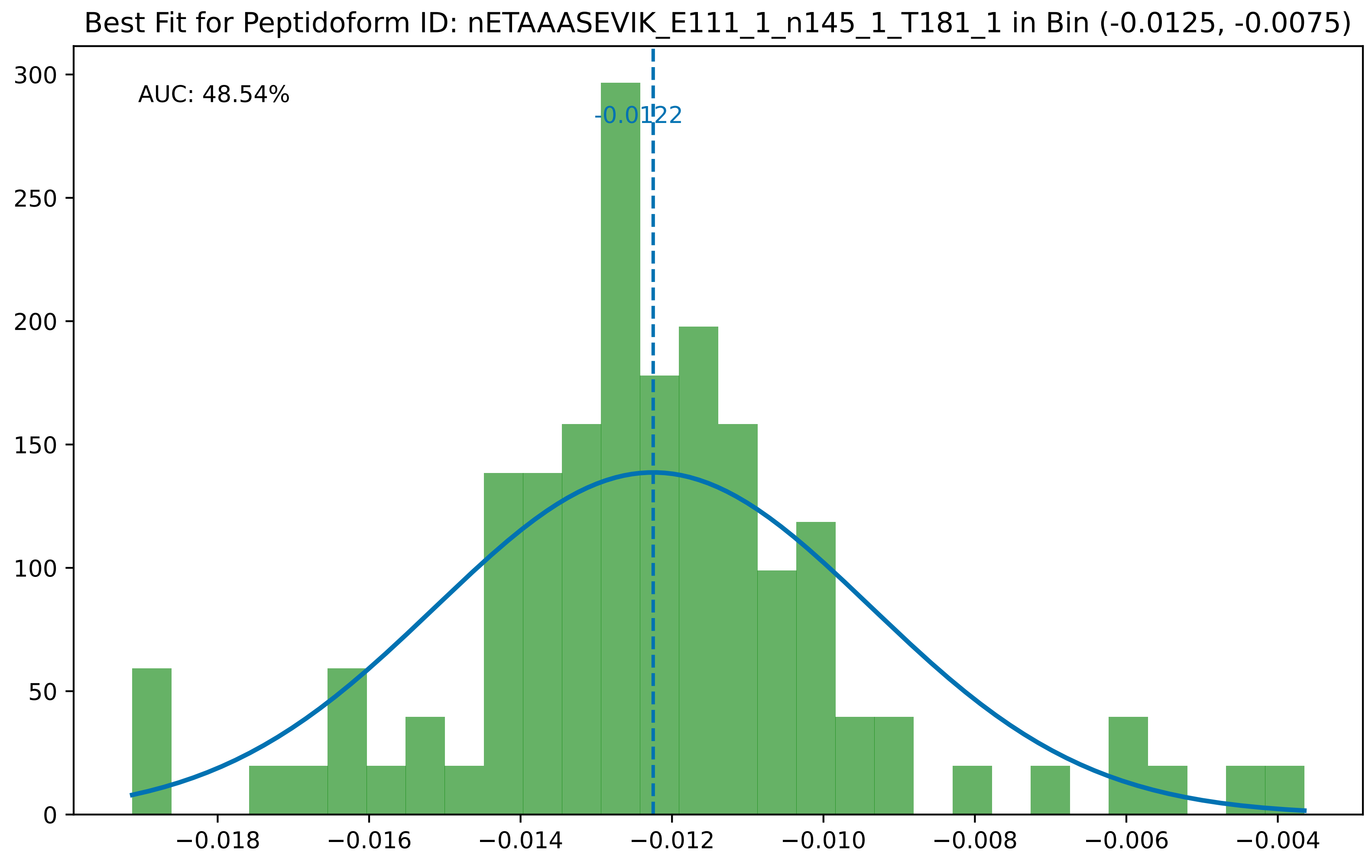
Best Fit for Peptidoform ID: nETAAASEVIK_E111_1_n145_1_T181_1 in Bin (-0.0125, -0.0075)
AUC: 48.54%
-0.0122
0
50
100
150
200
250
300
−0.018
−0.016
−0.014
−0.012
−0.010
−0.008
−0.006
−0.004

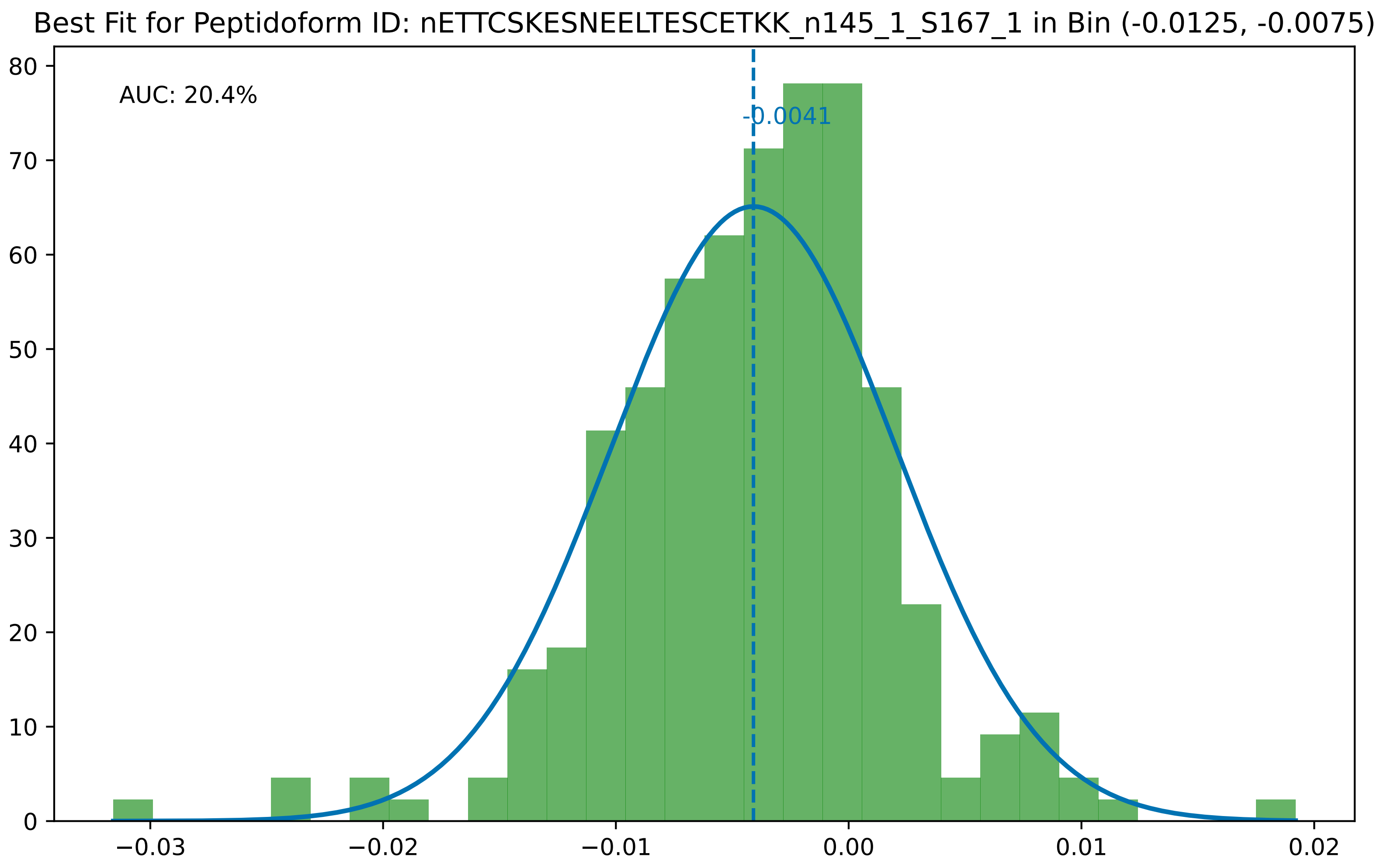

Best Fit for Peptidoform ID: nETTCSKESNEELTESCETKK_n145_1_S167_1 in Bin (-0.0125, -0.0075)
AUC: 20.4%
-0.0041
80
70
60
50
40
30
20
10
0
−0.03
−0.02
−0.01
0.00
0.01
0.02

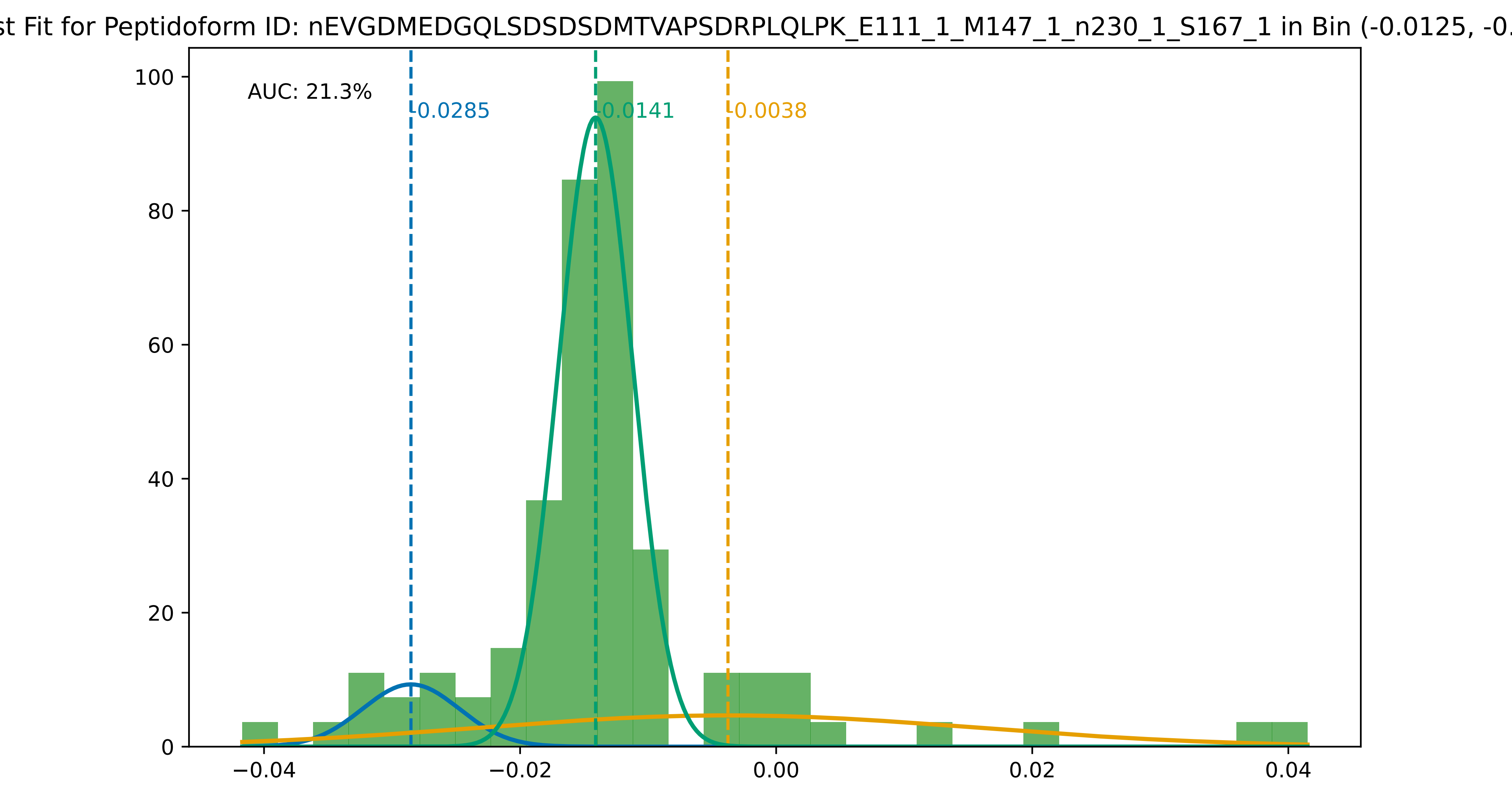
st Fit for Peptidoform ID: nEVGDMEDGQLSDSDSDMTVAPSDRPLQLPK_E111_1_M147_1_n230_1_S167_1 in Bin (-0.0125, -0.
AUC: 21.3%
-0.0285
-0.0141
-0.0038
100
80
60
40
20
0
−0.04
−0.02
0.00
0.02
0.04

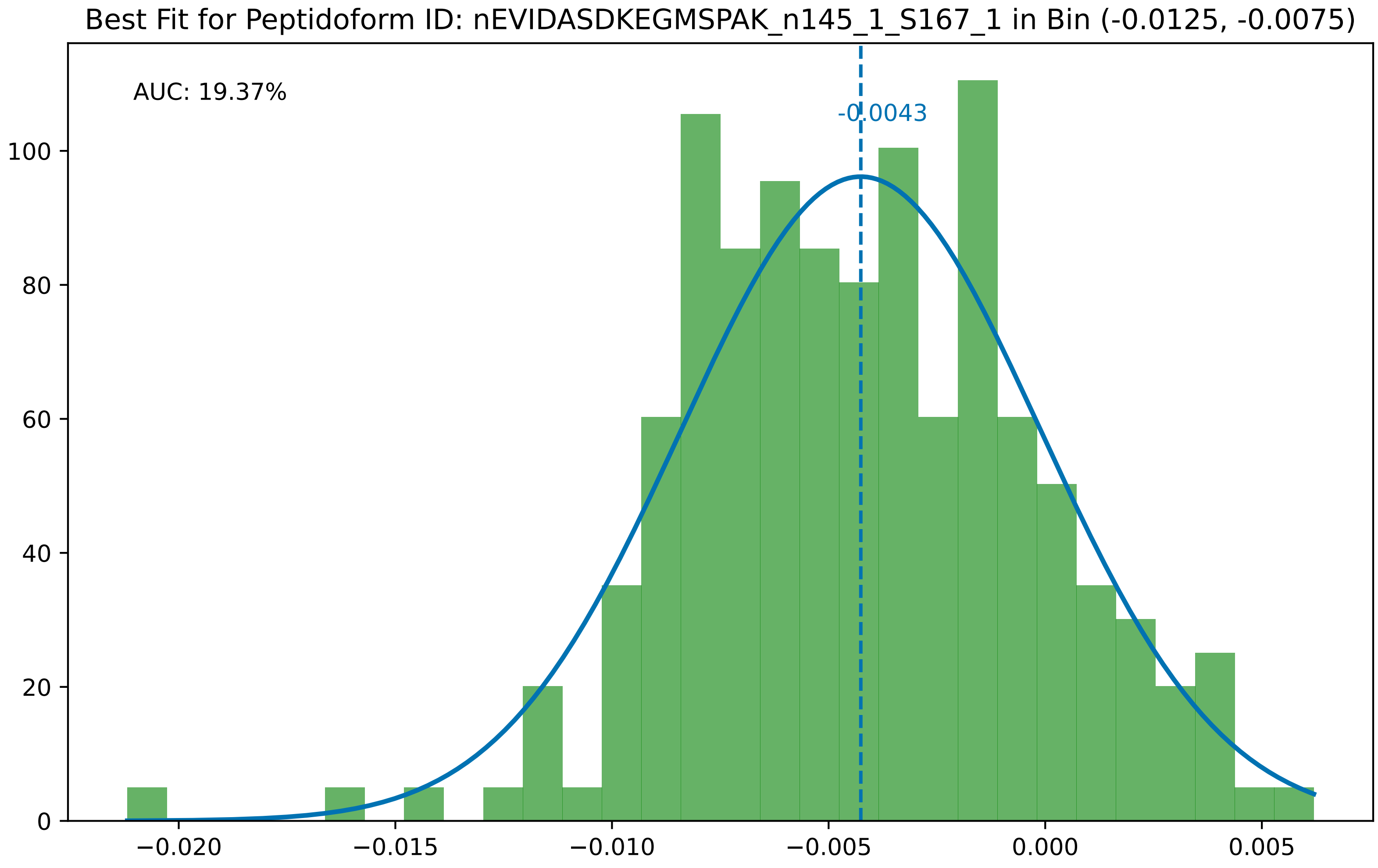

Best Fit for Peptidoform ID: nEVIDASDKEGMSPAK_n145_1_S167_1 in Bin (-0.0125, -0.0075)
AUC: 19.37%
-0.0043
0
20
40
60
80
100
−0.020
−0.015
−0.010
−0.005
0.000
0.005

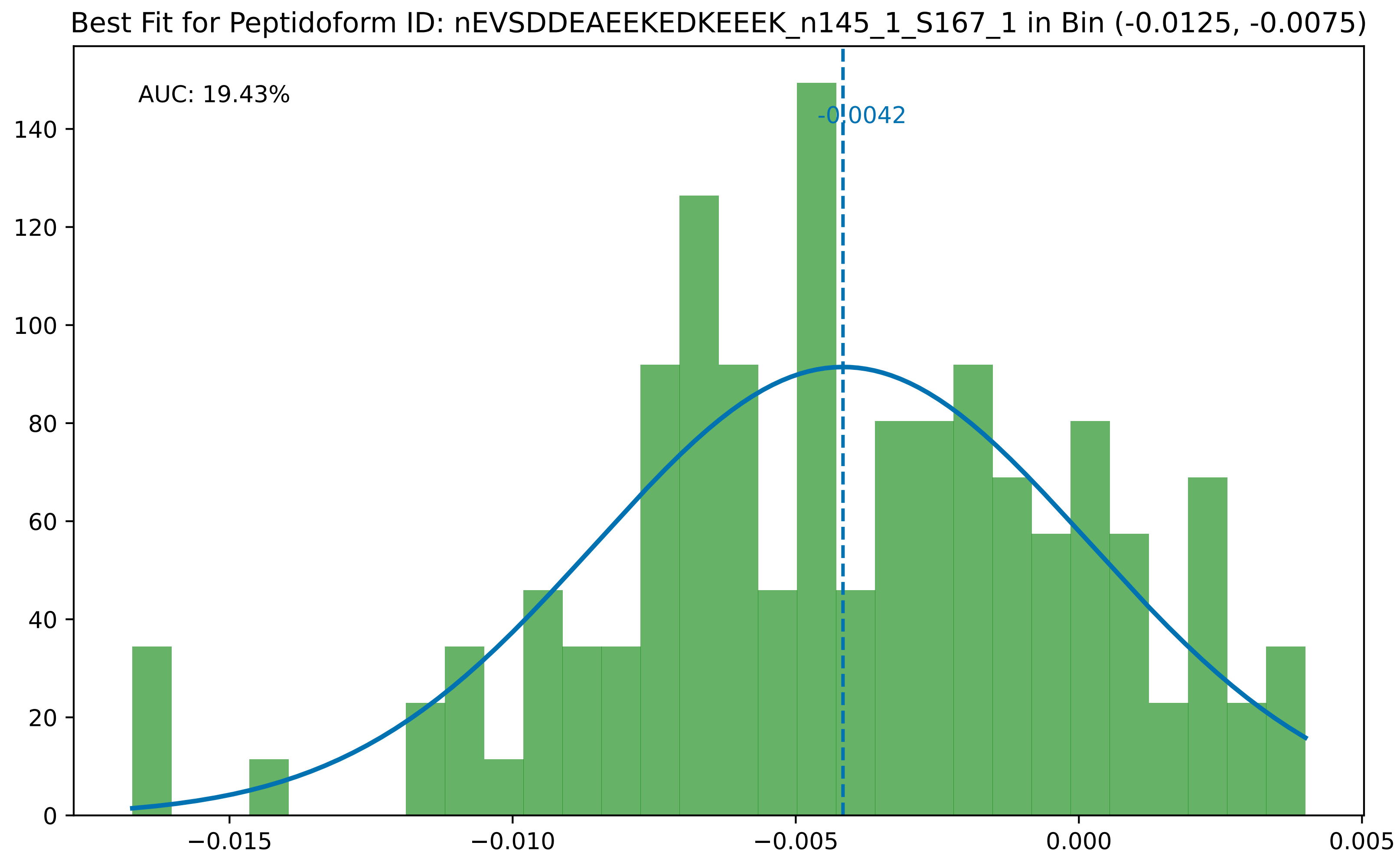

Best Fit for Peptidoform ID: nEVSDDEAEEKEDKEEEK_n145_1_S167_1 in Bin (-0.0125, -0.0075)
AUC: 19.43%
-0.0042
0
20
40
60
80
100
120
140
−0.015
−0.010
−0.005
0.000
0.005

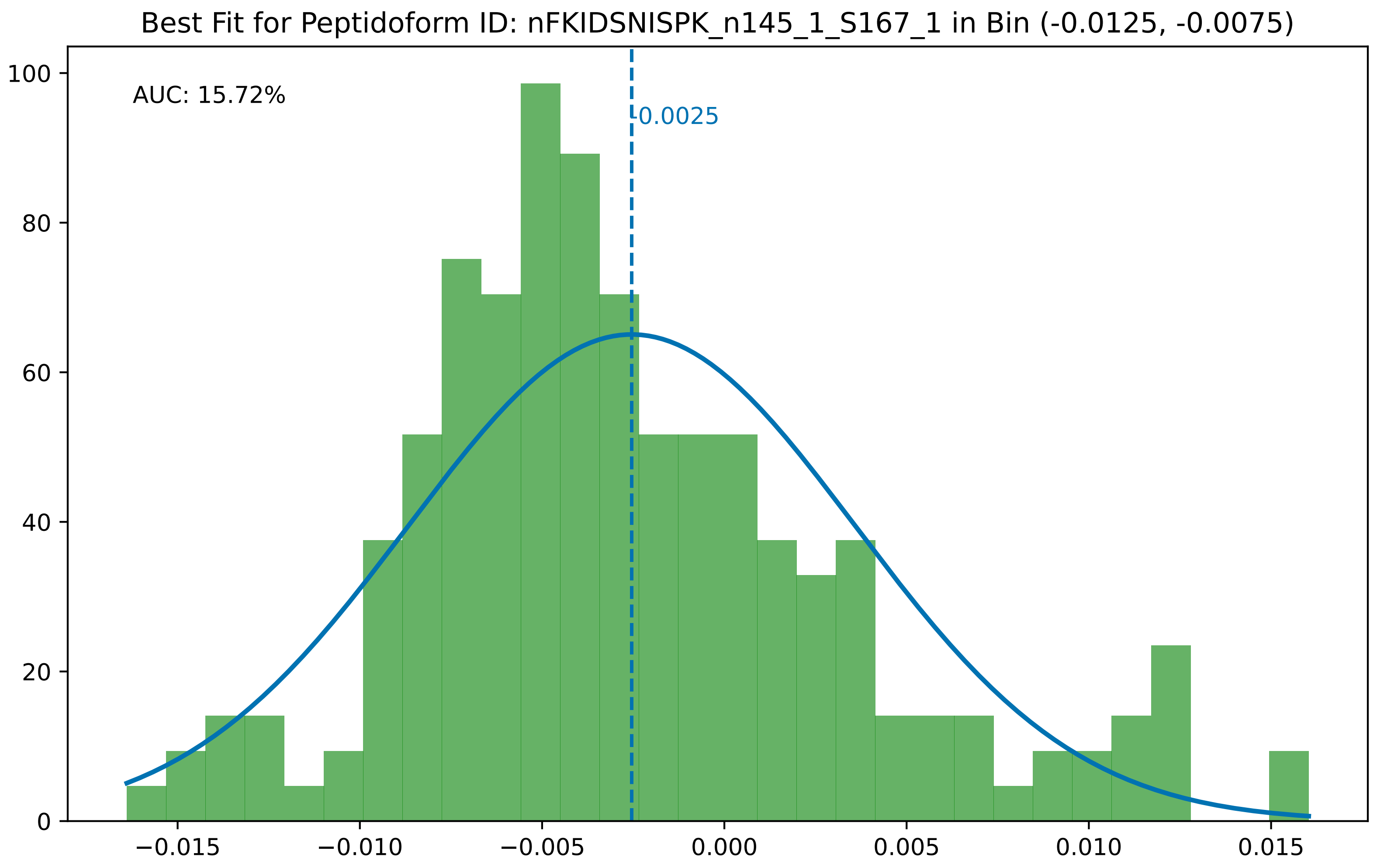
Best Fit for Peptidoform ID: nFKIDSNISPK_n145_1_S167_1 in Bin (-0.0125, -0.0075)
AUC: 15.72%
-0.0025
0
20
40
60
80
100
−0.015
−0.010
−0.005
0.000
0.005
0.010
0.015

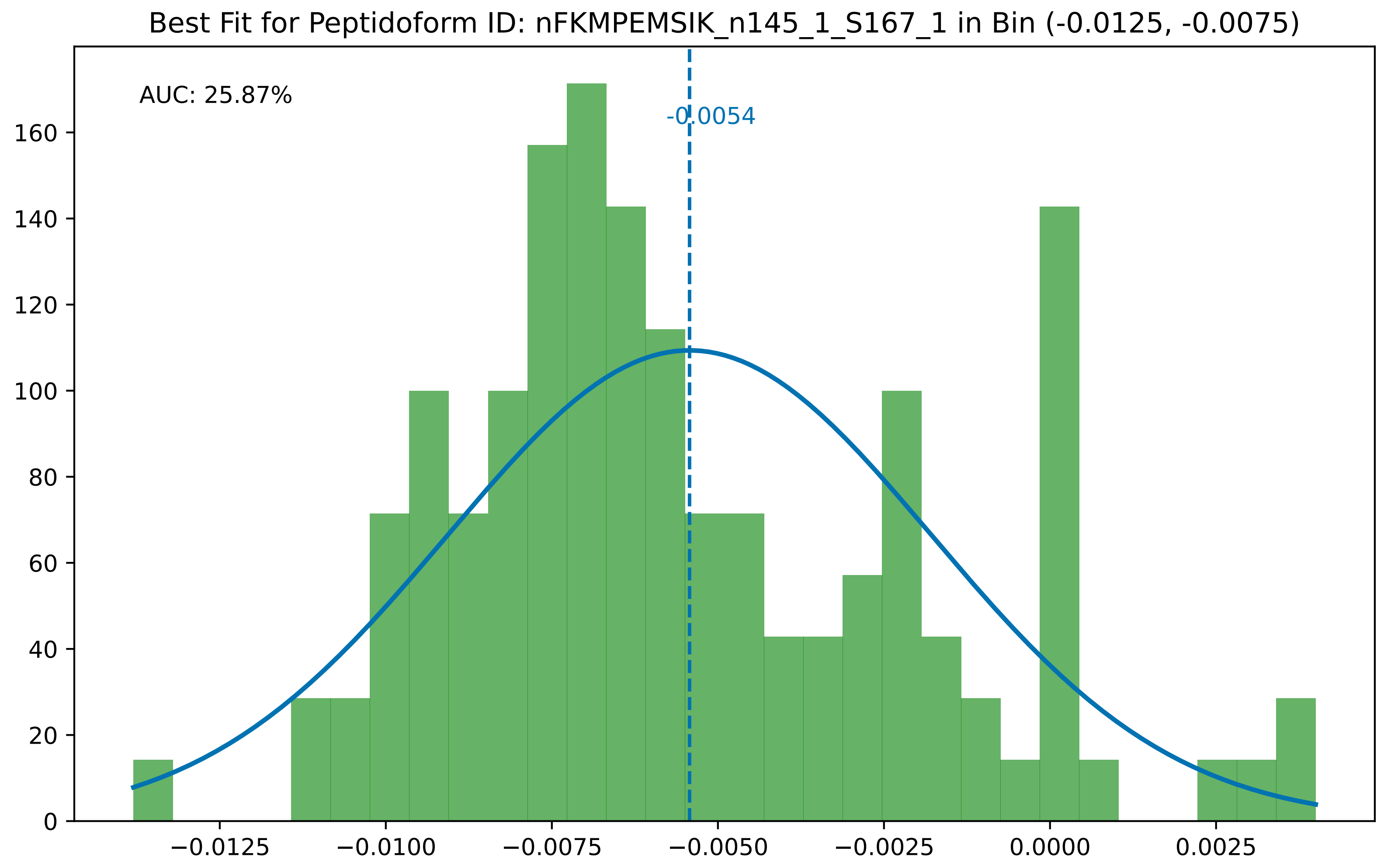
Best Fit for Peptidoform ID: nFKMPEMSIK_n145_1_S167_1 in Bin (-0.0125, -0.0075)
AUC: 25.87%
-0.0054
0
20
40
60
80
100
120
140
160
−0.0125
−0.0100
−0.0075
−0.0050
−0.0025
0.0000
0.0025

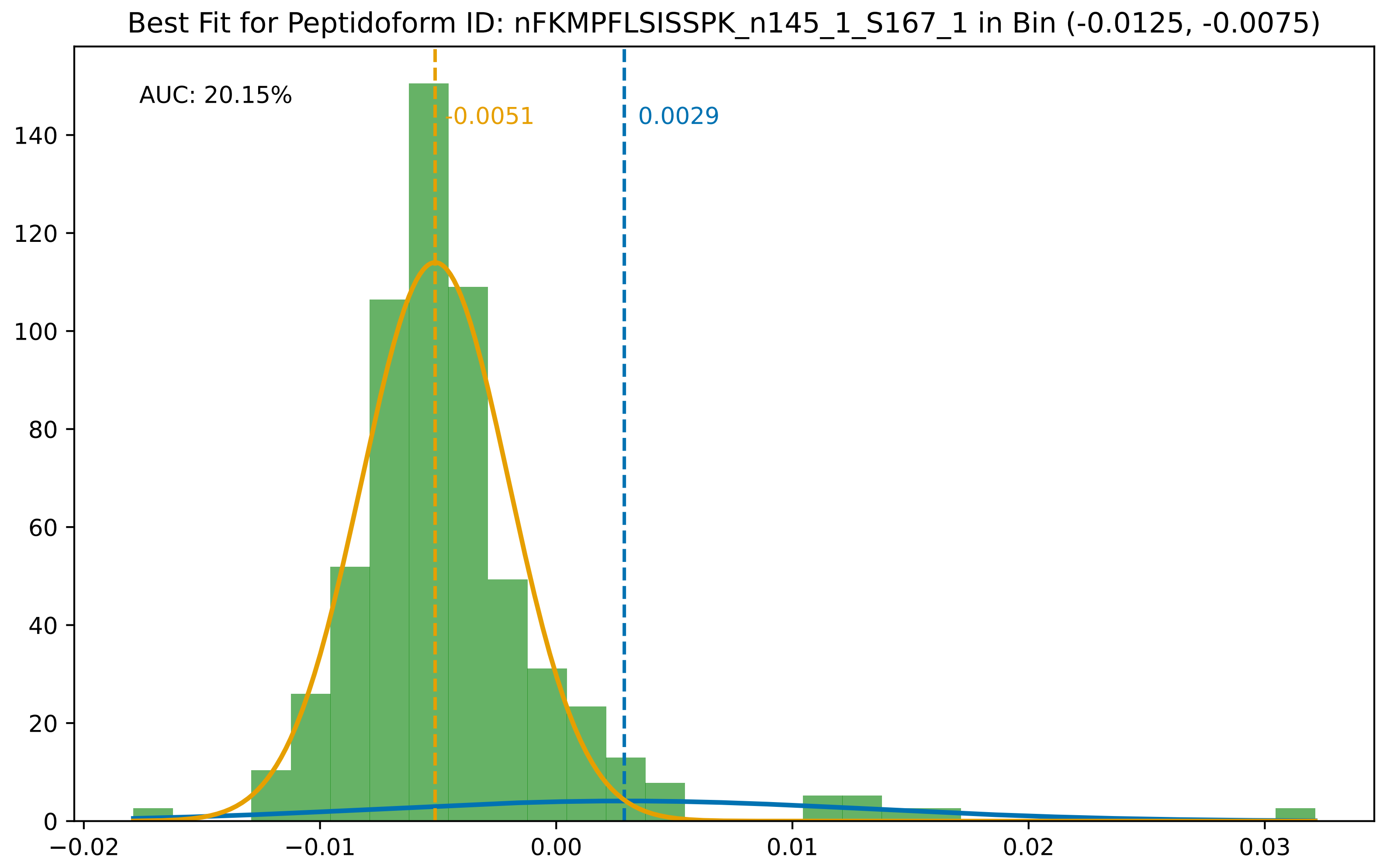

Best Fit for Peptidoform ID: nFKMPFLSISSPK_n145_1_S167_1 in Bin (-0.0125, -0.0075)
AUC: 20.15%
-0.0051
0.0029
140
120
100
80
60
40
20
0
−0.02
−0.01
0.00
0.01
0.02
0.03

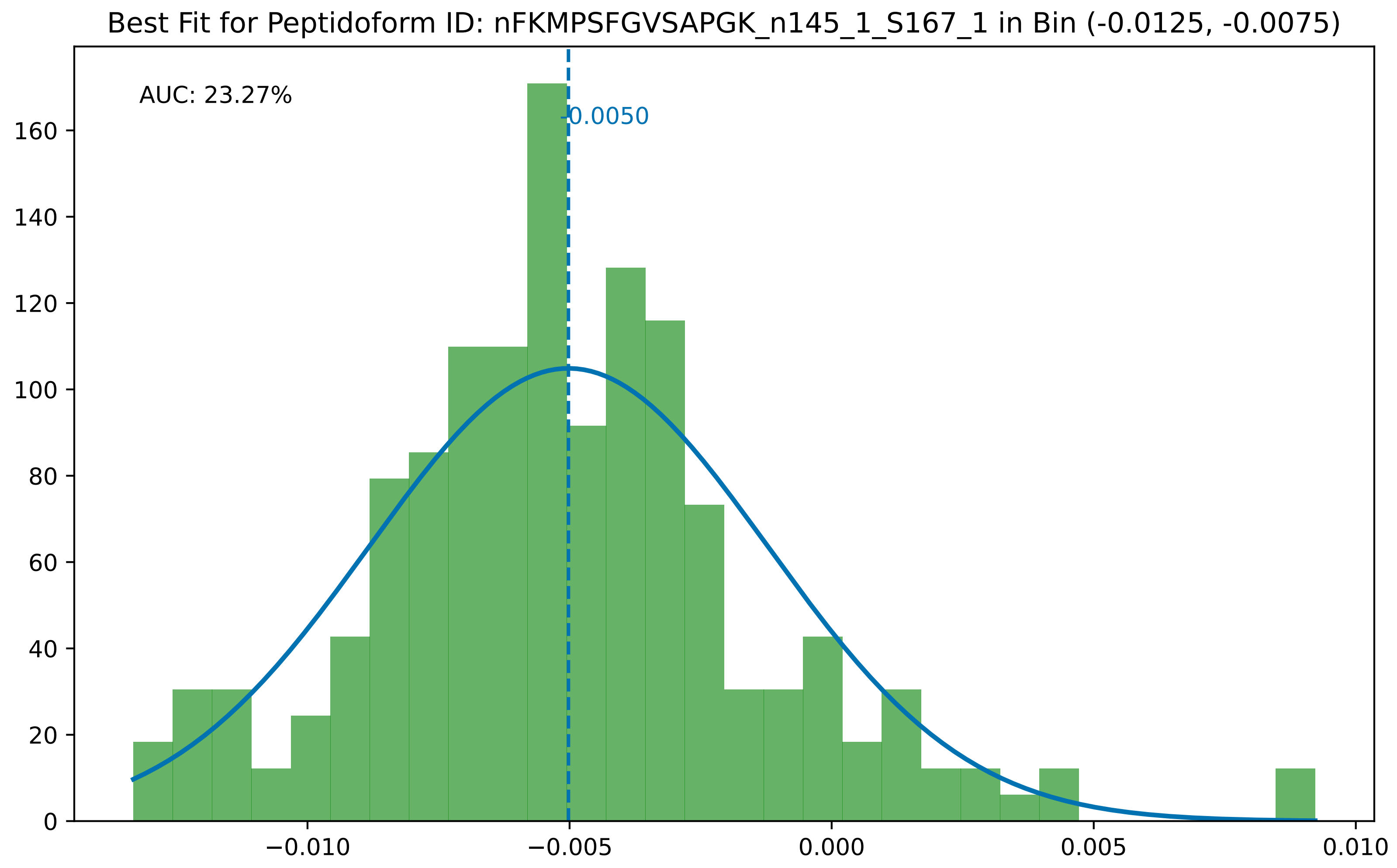

Best Fit for Peptidoform ID: nFKMPSFGVSAPGK_n145_1_S167_1 in Bin (-0.0125, -0.0075)
AUC: 23.27%
-0.0050
0
20
40
60
80
100
120
140
160
−0.010
−0.005
0.000
0.005
0.010

Best Fit for Peptidoform ID: nFKTQPVTFDEIQEVEEEGVSPMEEEK_n145_1_T181_1 in Bin (-0.0125, -0.0075)

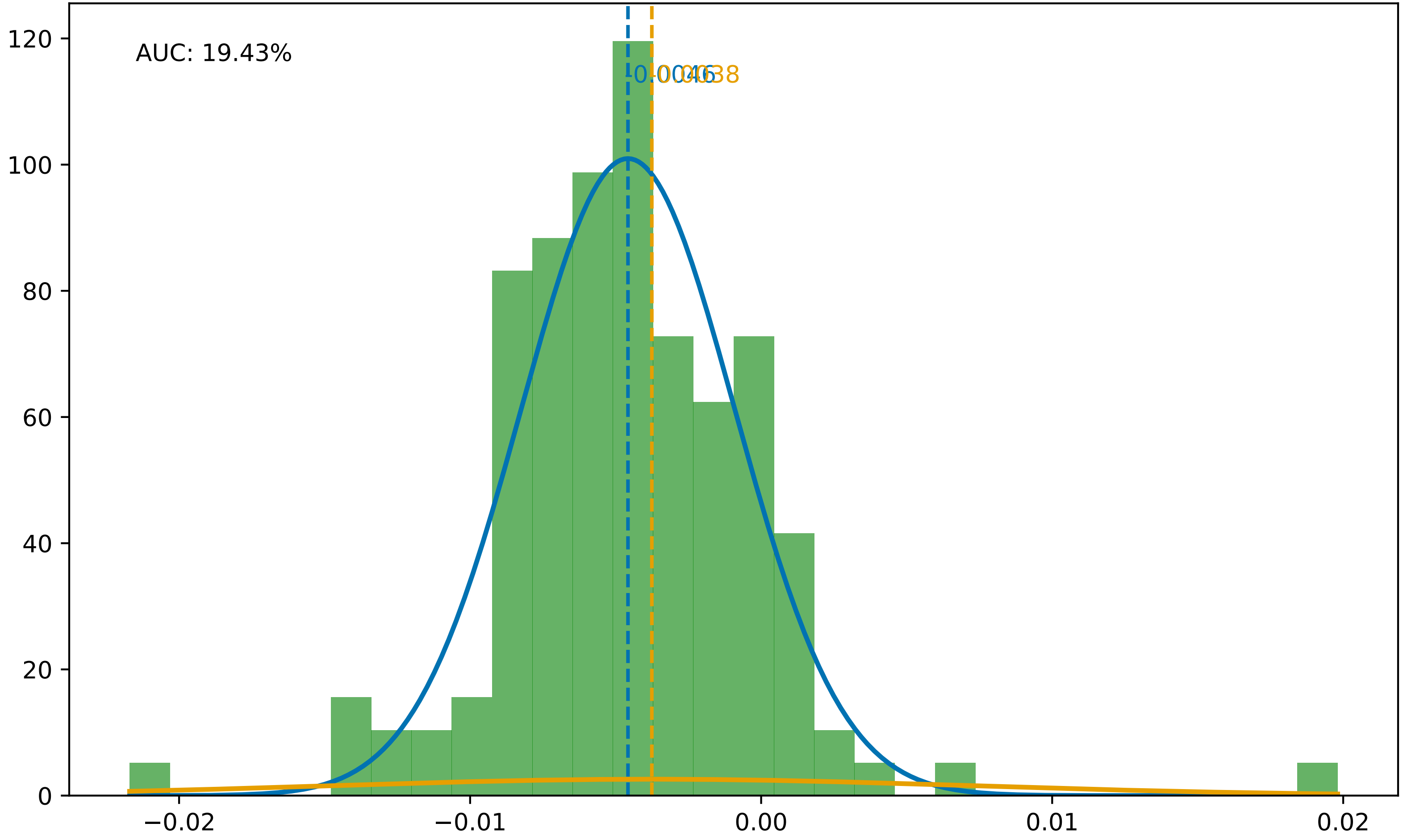

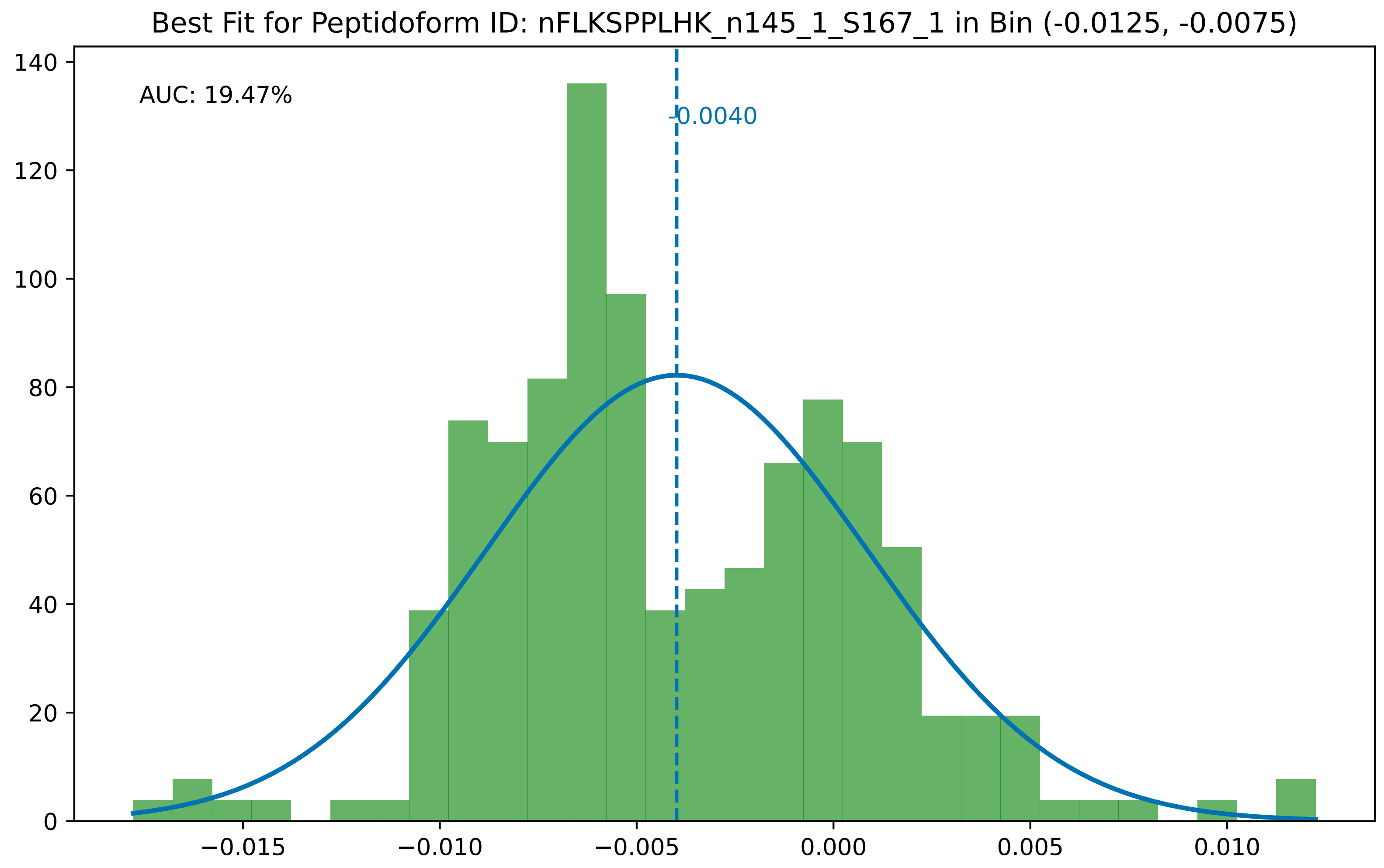
Best Fit for Peptidoform ID: nFLKSPPLHK_n145_1_S167_1 in Bin (-0.0125, -0.0075)
AUC: 19.47%
-0.0040
0
20
40
60
80
100
120
140
−0.015
−0.010
−0.005
0.000
0.005
0.010

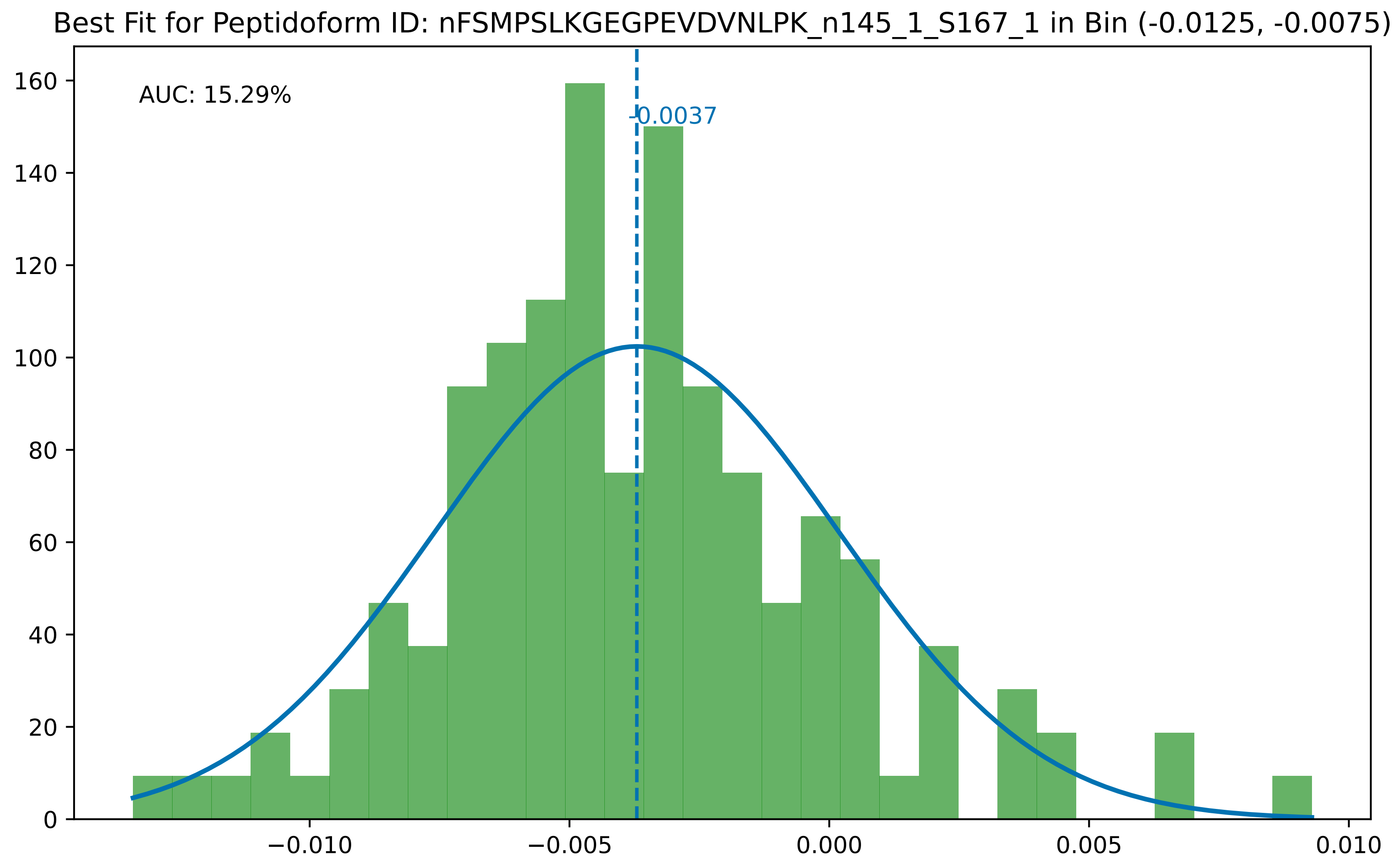
Best Fit for Peptidoform ID: nFSMPSLKGEGPEVDVNLPK_n145_1_S167_1 in Bin (-0.0125, -0.0075)
AUC: 15.29%
-0.0037
0
20
40
60
80
100
120
140
160
−0.010
−0.005
0.000
0.005
0.010

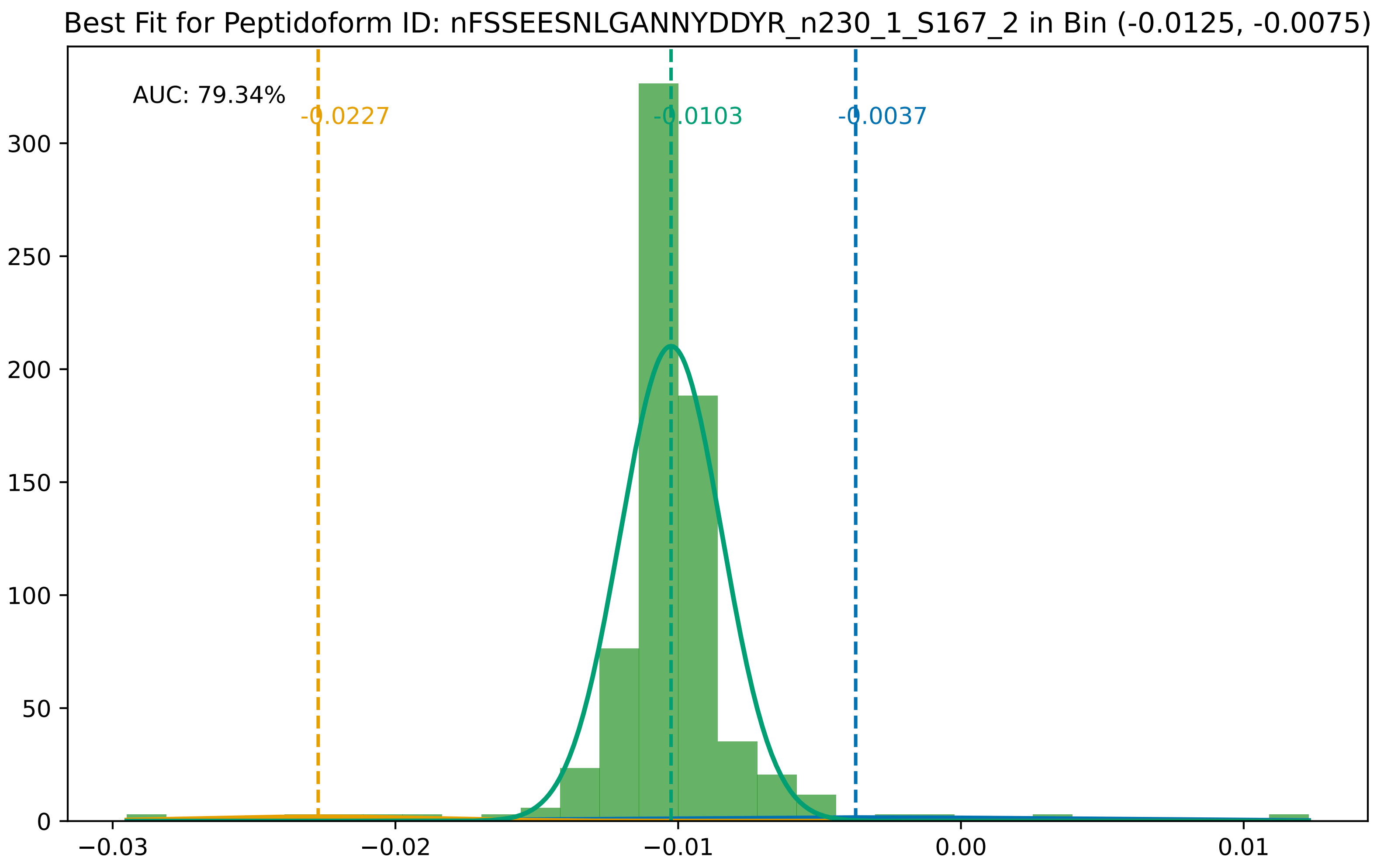

Best Fit for Peptidoform ID: nFSSEESNLGANNYDDYR_n230_1_S167_2 in Bin (-0.0125, -0.0075)
AUC: 79.34%
-0.0227
-0.0103
-0.0037
300
250
200
150
100
50
0
−0.03
−0.02
−0.01
0.00
0.01

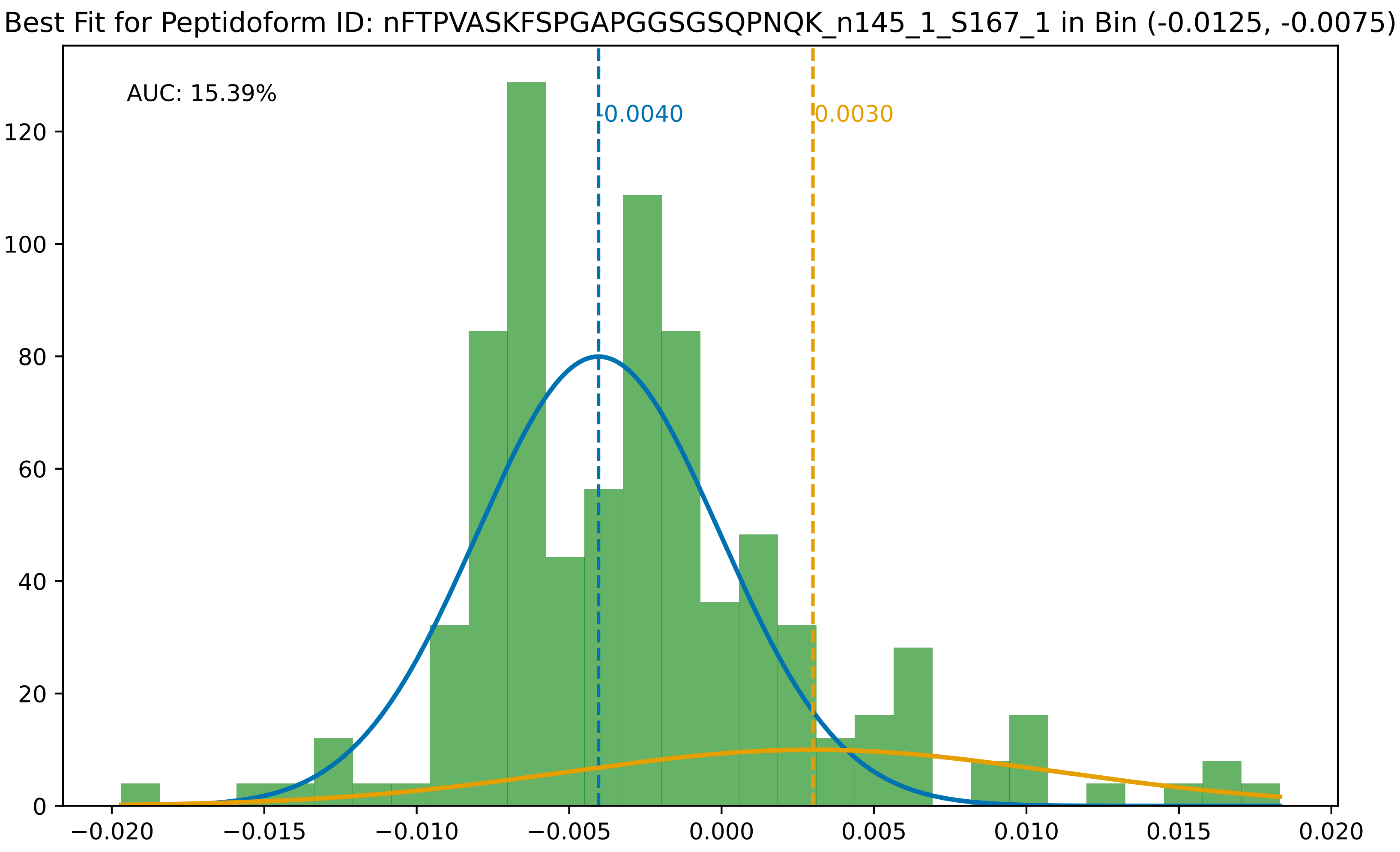
Best Fit for Peptidoform ID: nFTPVASKFSPGAPGGSGSQPNQK_n145_1_S167_1 in Bin (-0.0125, -0.0075)
AUC: 15.39%
-0.0040
0.0030
0
20
40
60
80
100
120
−0.020
−0.015
−0.010
−0.005
0.000
0.005
0.010
0.015
0.020

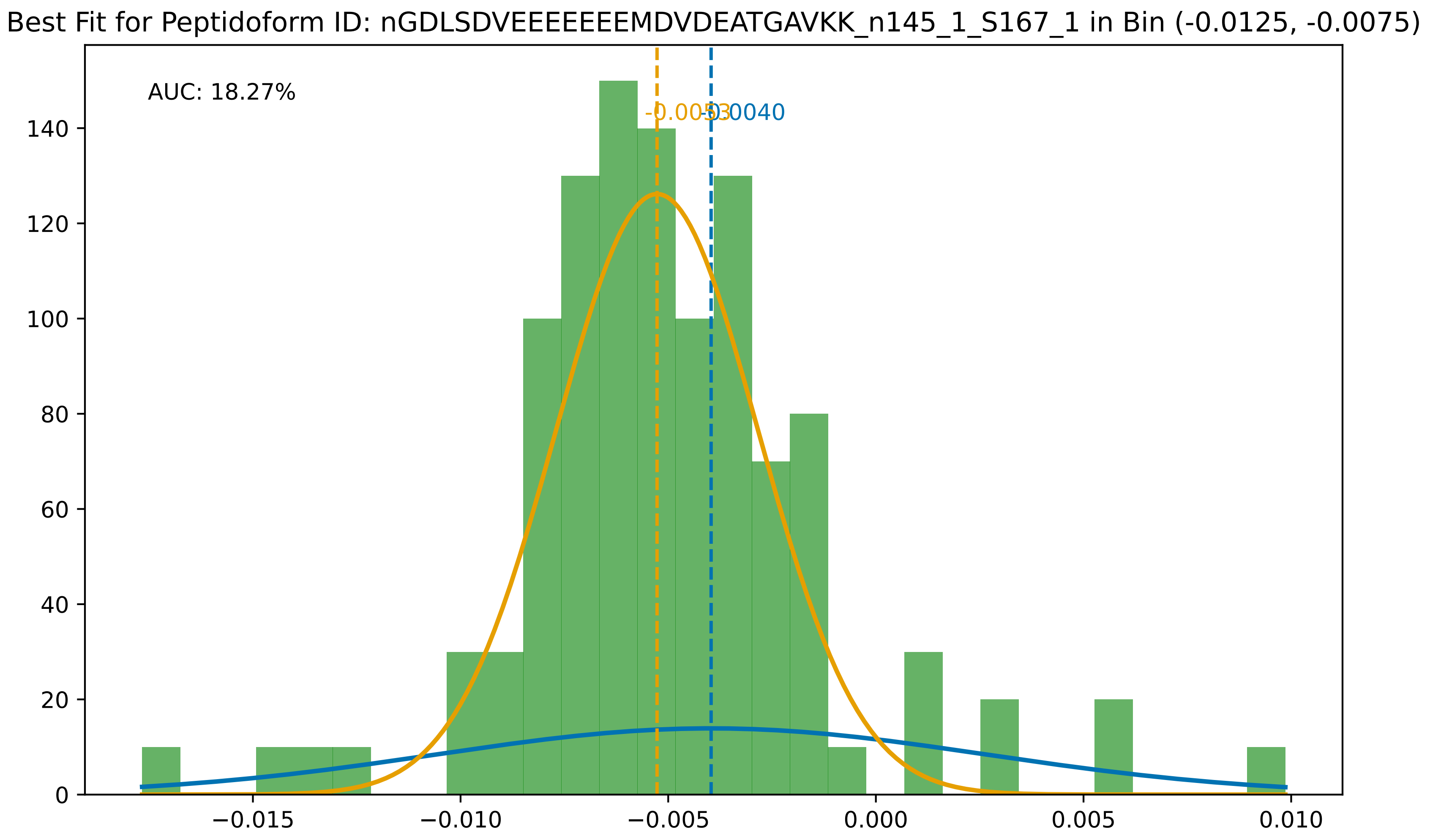
Best Fit for Peptidoform ID: nGDLSDVEEEEEEEMDVDEATGAVKK_n145_1_S167_1 in Bin (-0.0125, -0.0075)
AUC: 18.27%
-0.0053
-0.0040
0
20
40
60
80
100
120
140
−0.015
−0.010
−0.005
0.000
0.005
0.010

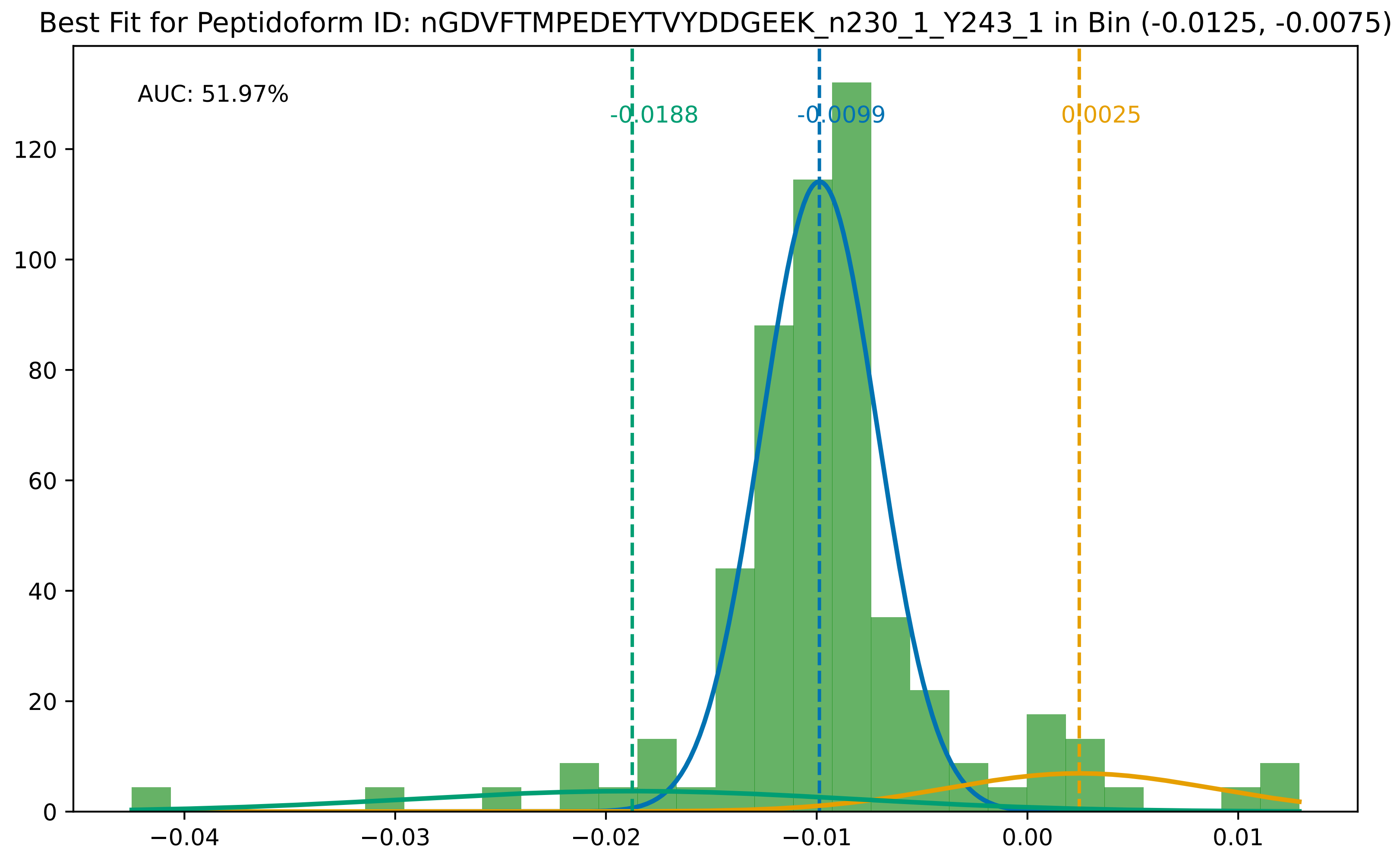
Best Fit for Peptidoform ID: nGDVFTMPEDEYTVYDDGEEK_n230_1_Y243_1 in Bin (-0.0125, -0.0075)
AUC: 51.97%
-0.0188
-0.0099
0.0025
0
20
40
60
80
100
120
−0.04
−0.03
−0.02
−0.01
0.00
0.01

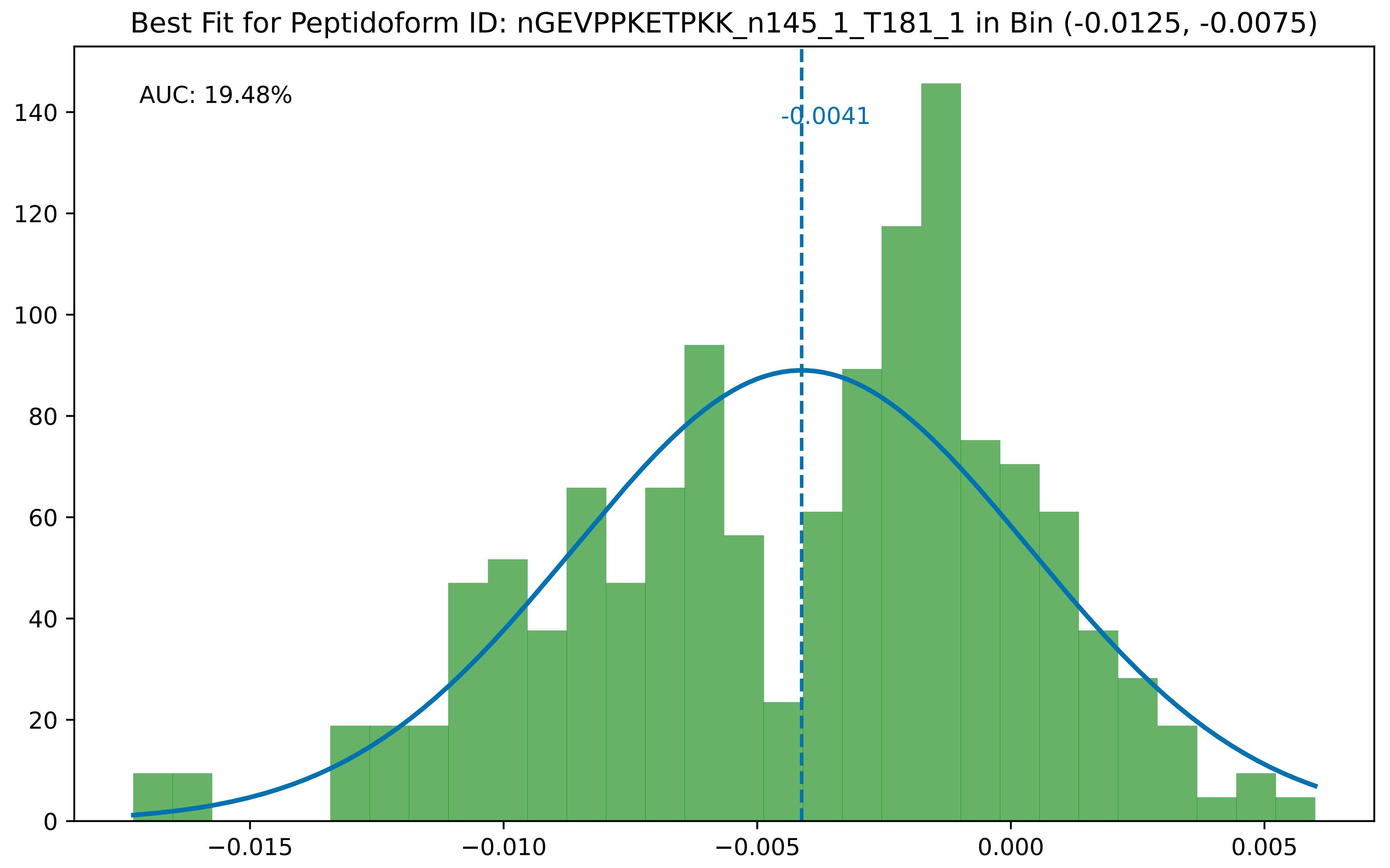

Best Fit for Peptidoform ID: nGEVPPKETPKK_n145_1_T181_1 in Bin (-0.0125, -0.0075)
AUC: 19.48%
-0.0041
0
20
40
60
80
100
120
140
−0.015
−0.010
−0.005
0.000
0.005

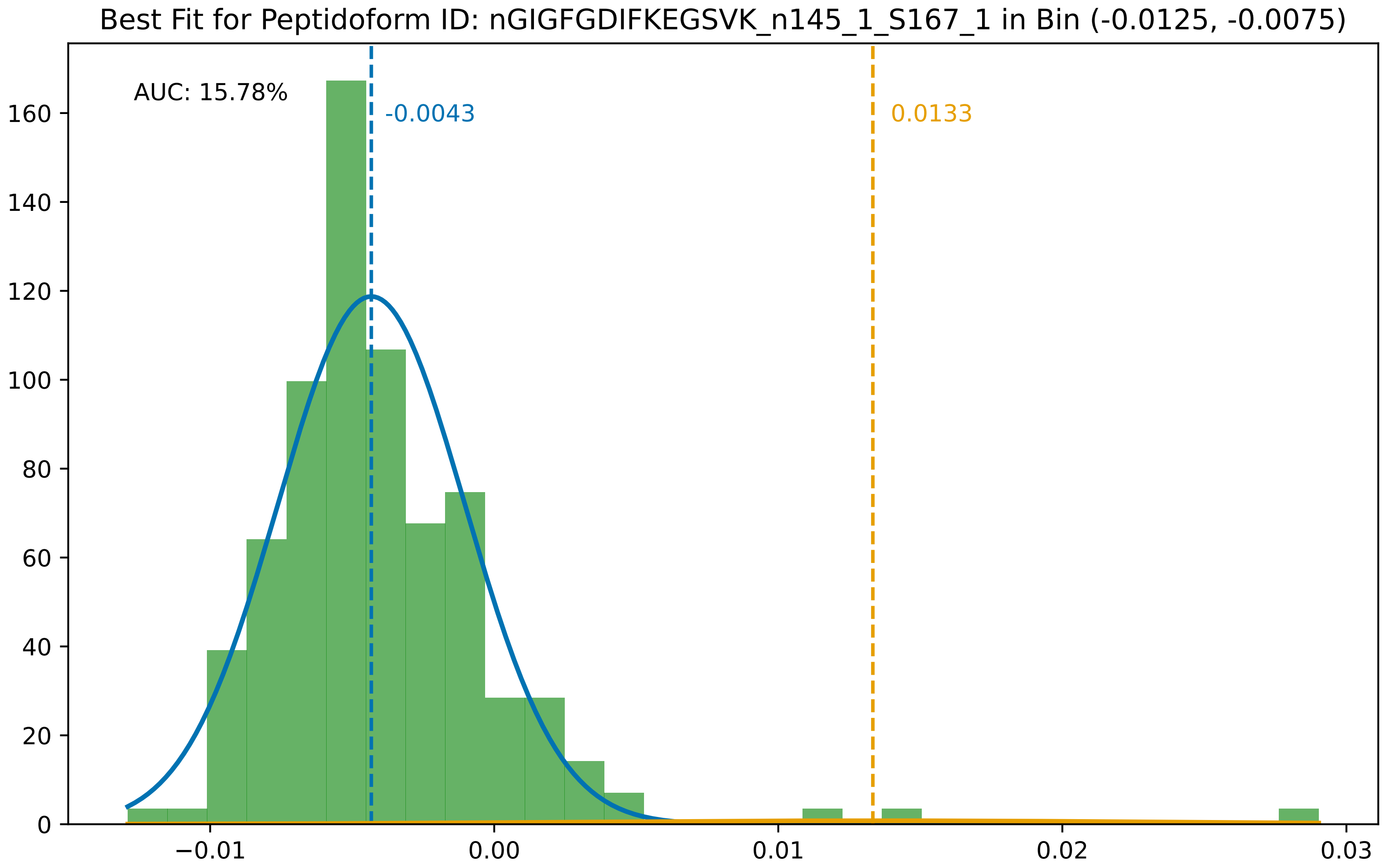

Best Fit for Peptidoform ID: nGIGFGDIFKEGSVK_n145_1_S167_1 in Bin (-0.0125, -0.0075)
AUC: 15.78%
-0.0043
0.0133
0
20
40
60
80
100
120
140
160
−0.01
0.00
0.01
0.02
0.03

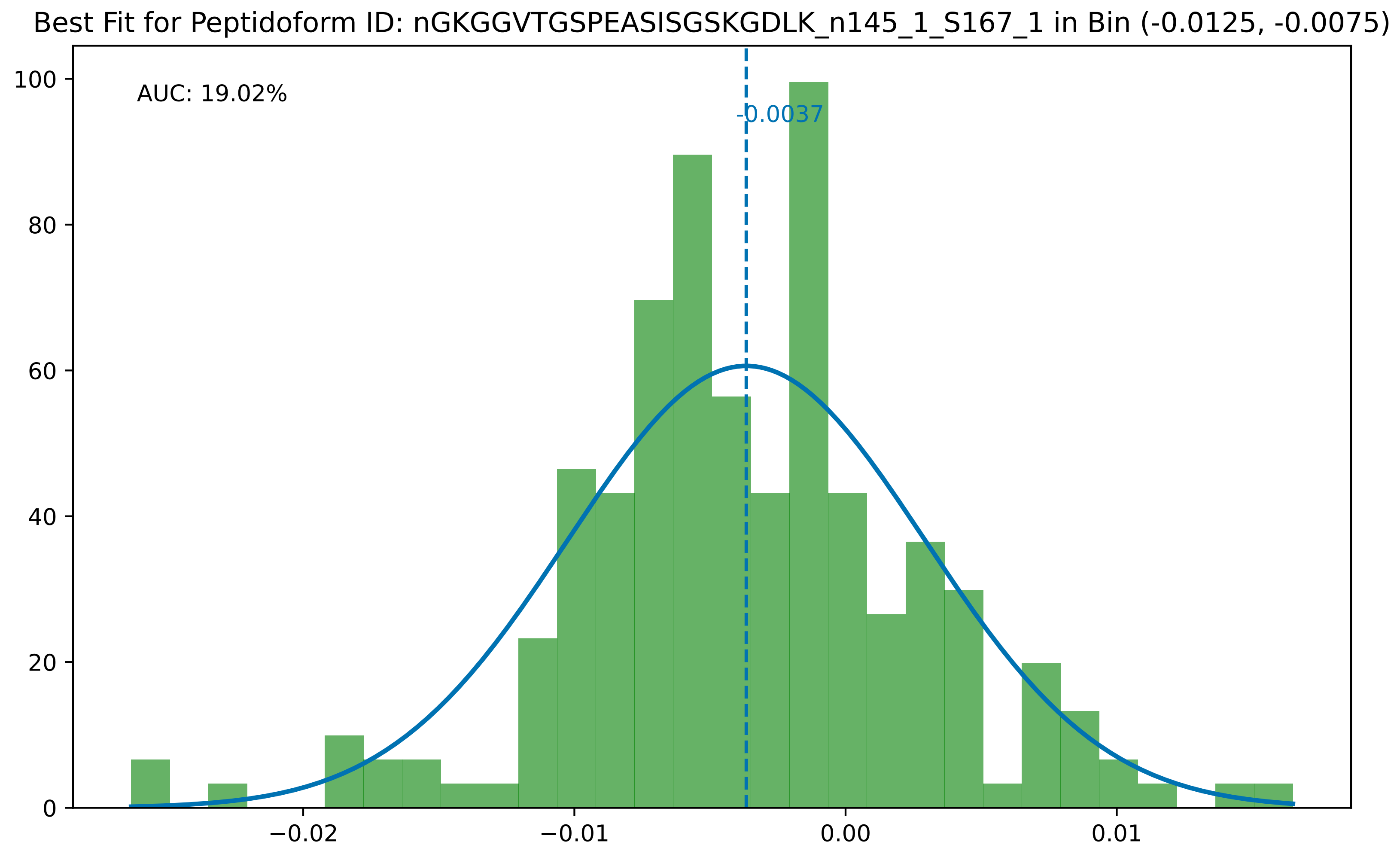

Best Fit for Peptidoform ID: nGKGGVTGSPEASISGSKGDLK_n145_1_S167_1 in Bin (-0.0125, -0.0075)
AUC: 19.02%
-0.0037
100
80
60
40
20
0
−0.02
−0.01
0.00
0.01

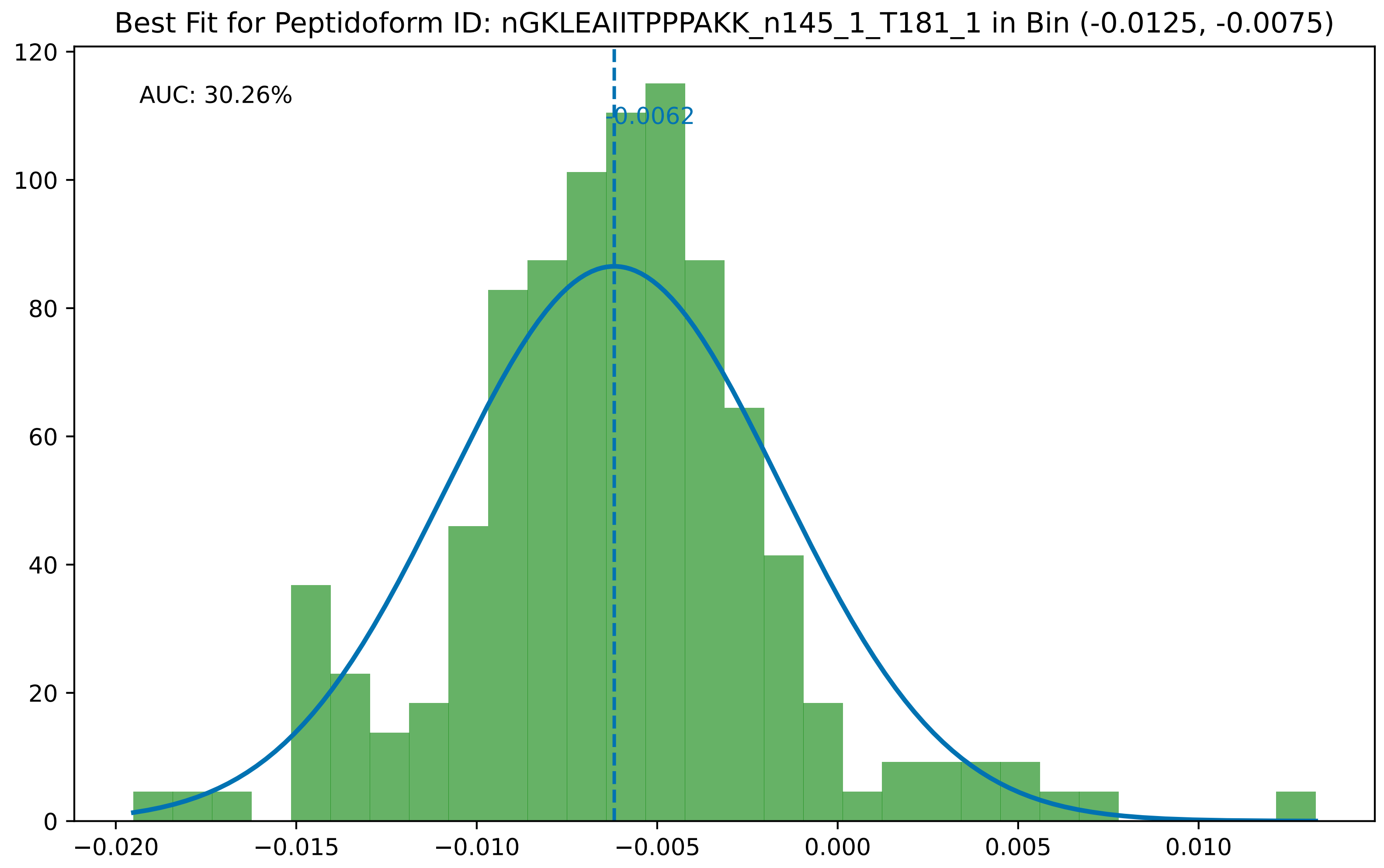
Best Fit for Peptidoform ID: nGKLEAIITPPPAKK_n145_1_T181_1 in Bin (-0.0125, -0.0075)
AUC: 30.26%
-0.0062
120
100
80
60
40
20
0
−0.020
−0.015
−0.010
−0.005
0.000
0.005
0.010

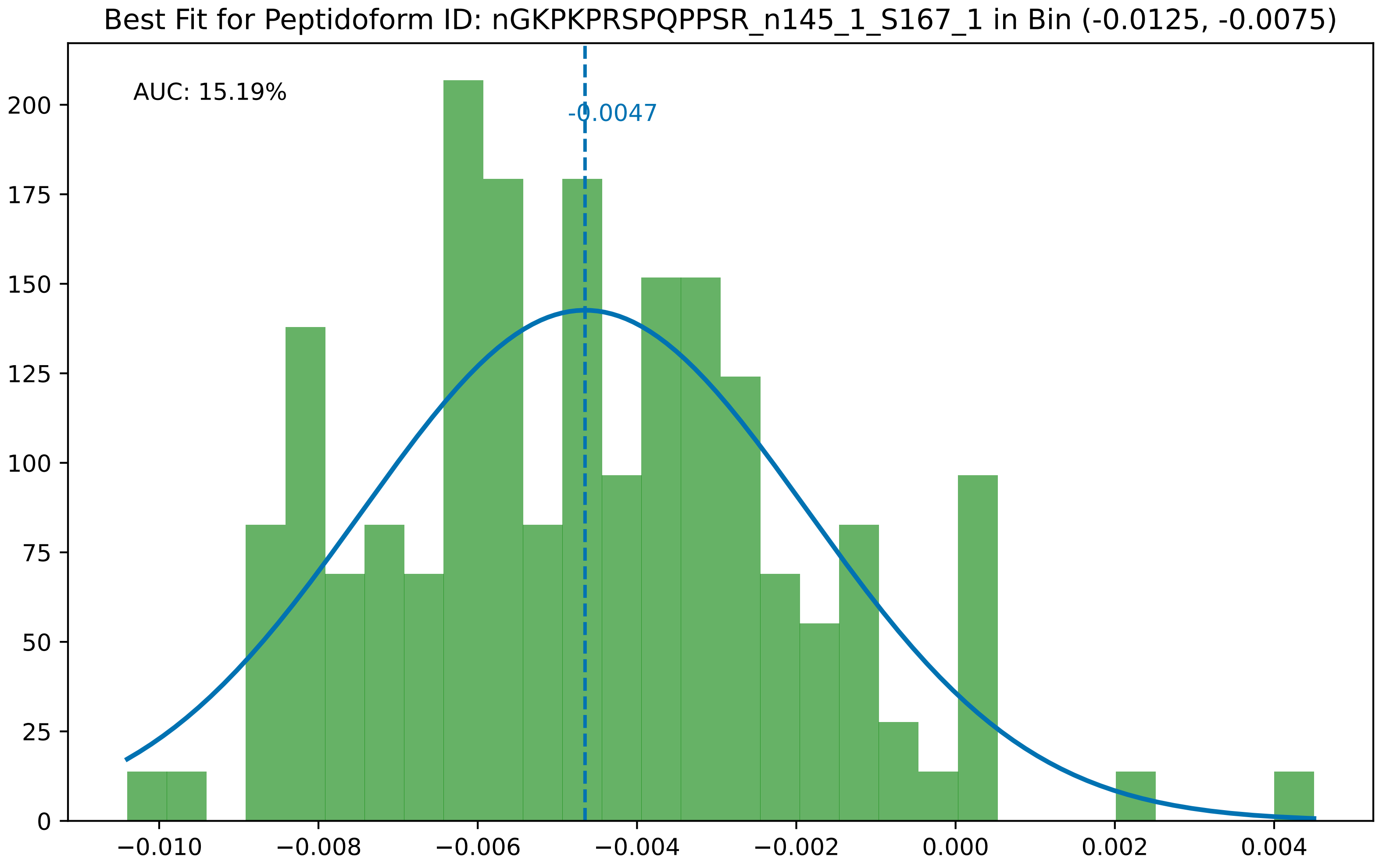

Best Fit for Peptidoform ID: nGKPKPRSPQPPSR_n145_1_S167_1 in Bin (-0.0125, -0.0075)
AUC: 15.19%
-0.0047
0
25
50
75
100
125
150
175
200
−0.010
−0.008
−0.006
−0.004
−0.002
0.000
0.002
0.004

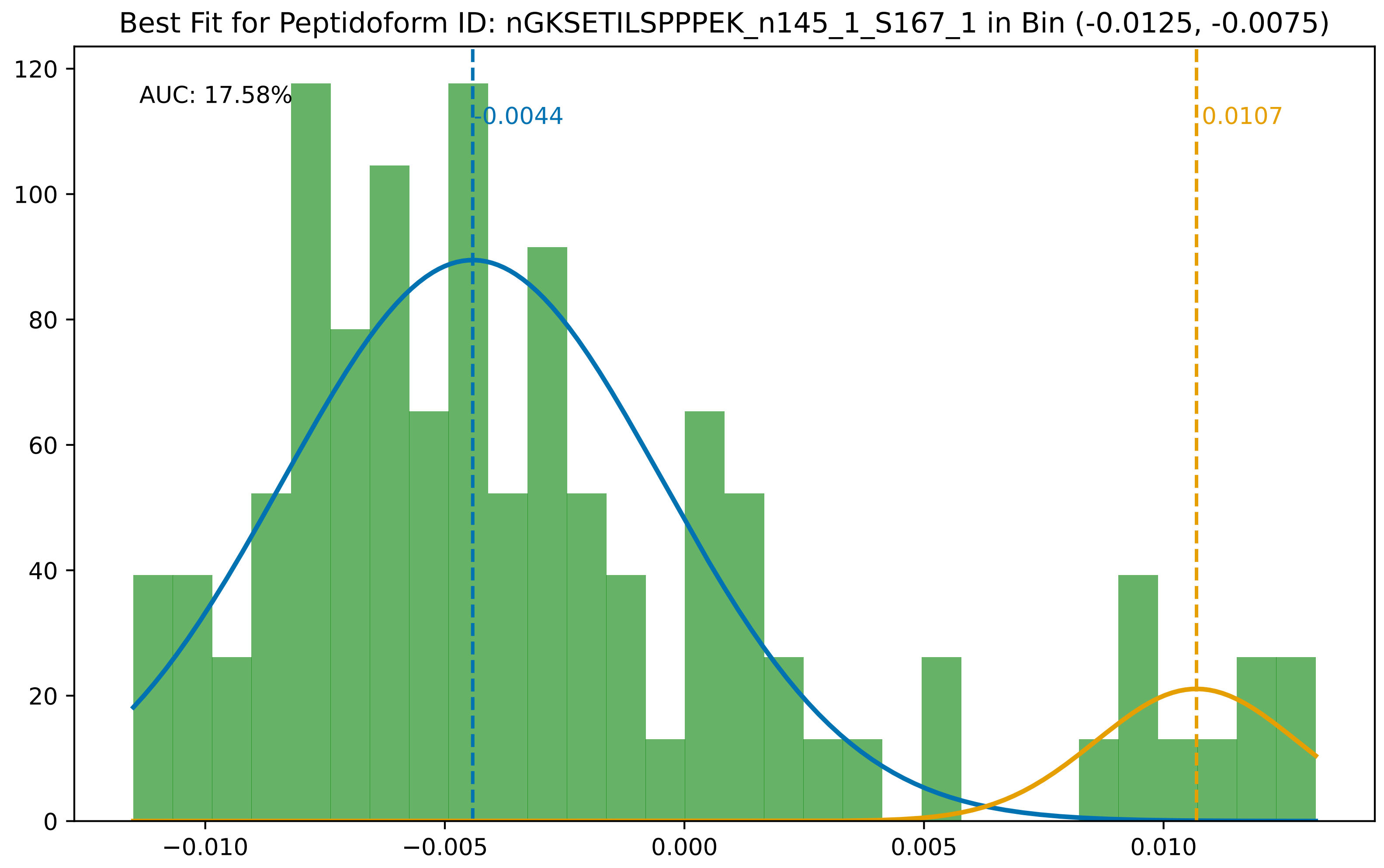
Best Fit for Peptidoform ID: nGKSETILSPPPEK_n145_1_S167_1 in Bin (-0.0125, -0.0075)
AUC: 17.58%
-0.0044
0.0107
0
20
40
60
80
100
120
−0.010
−0.005
0.000
0.005
0.010

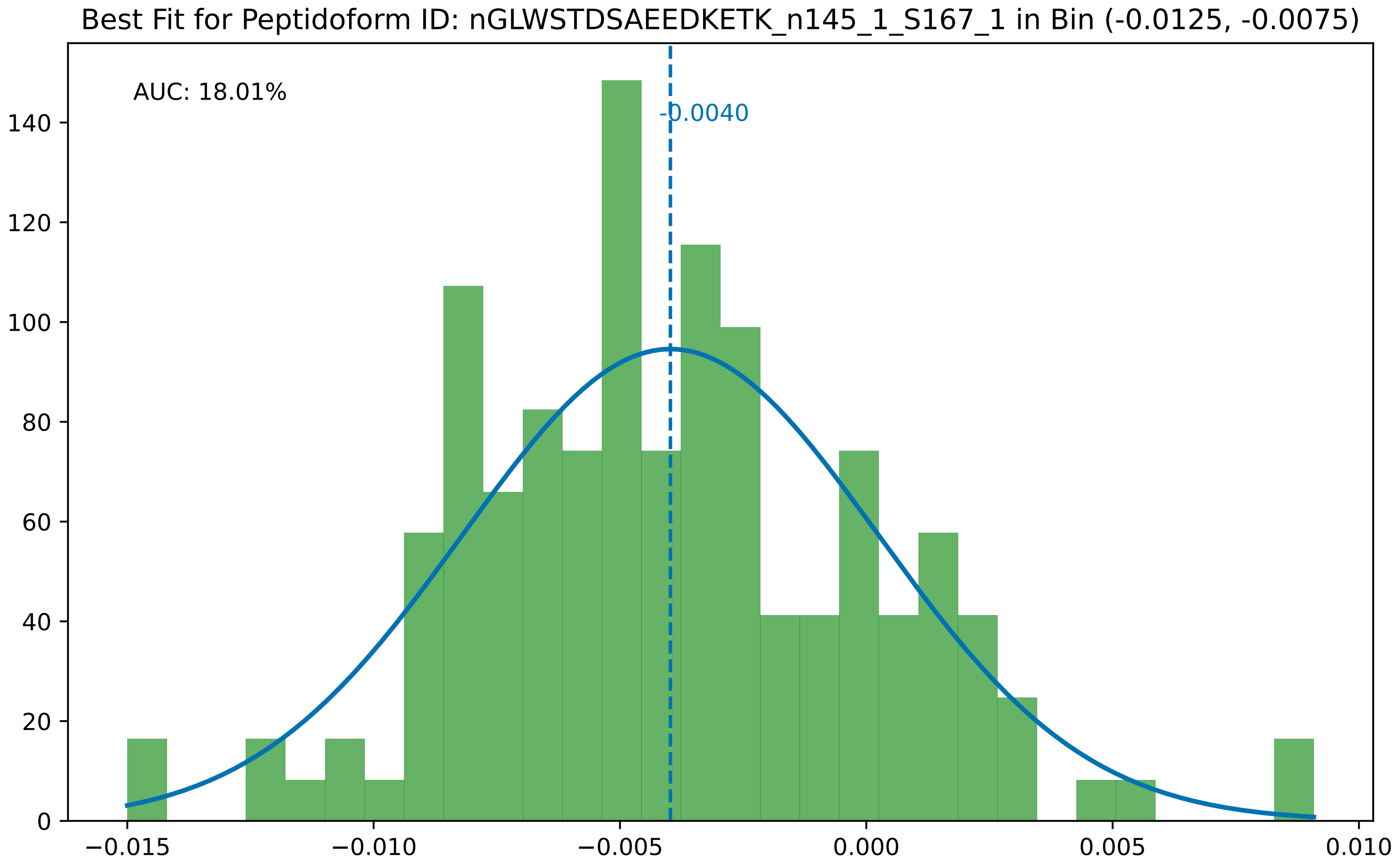
Best Fit for Peptidoform ID: nGLWSTDSAEEDKETK_n145_1_S167_1 in Bin (-0.0125, -0.0075)
AUC: 18.01%
-0.0040
0
20
40
60
80
100
120
140
−0.015
−0.010
−0.005
0.000
0.005
0.010

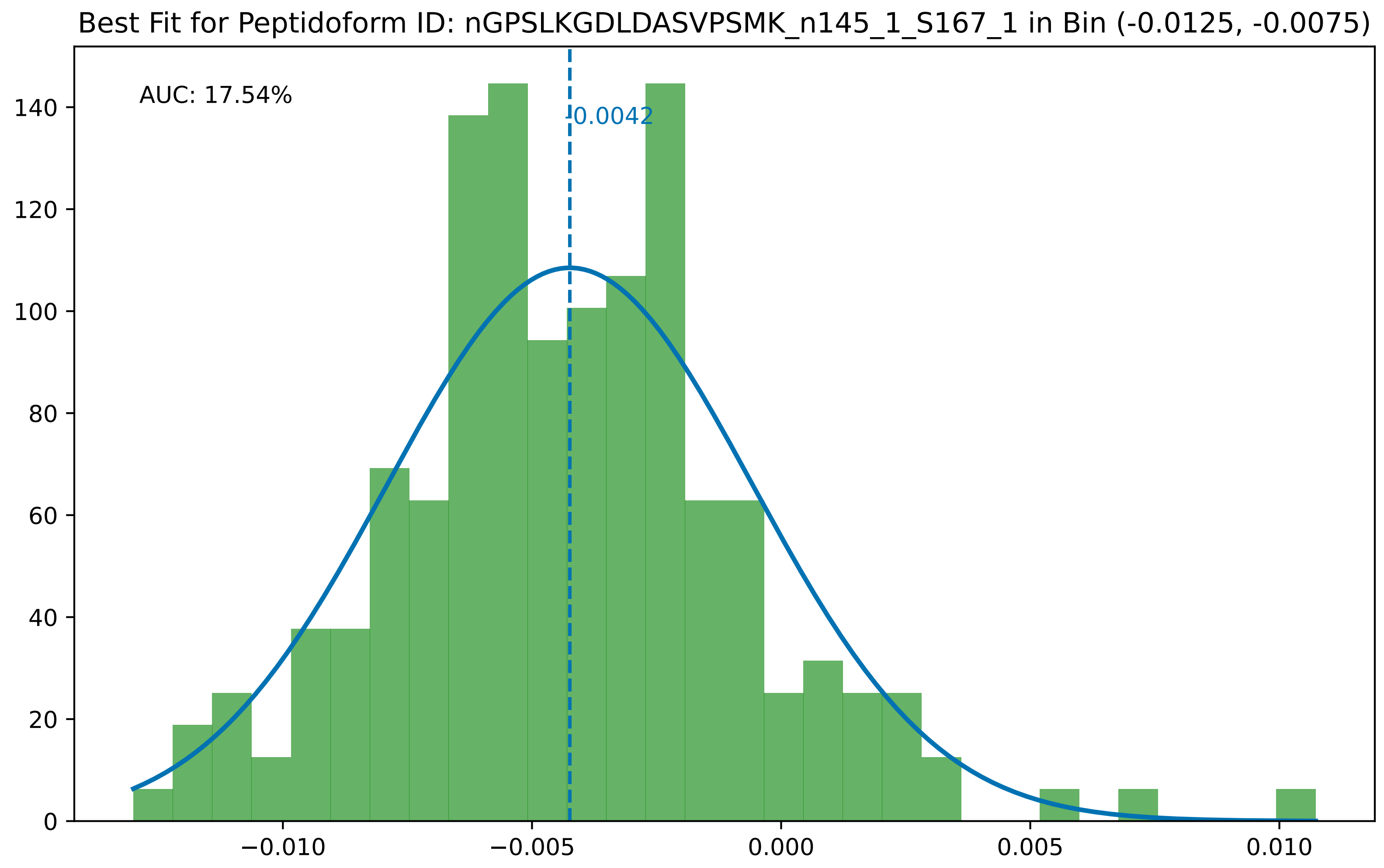
Best Fit for Peptidoform ID: nGPSLKGDLDASVPSMK_n145_1_S167_1 in Bin (-0.0125, -0.0075)
AUC: 17.54%
-0.0042
0
20
40
60
80
100
120
140
−0.010
−0.005
0.000
0.005
0.010

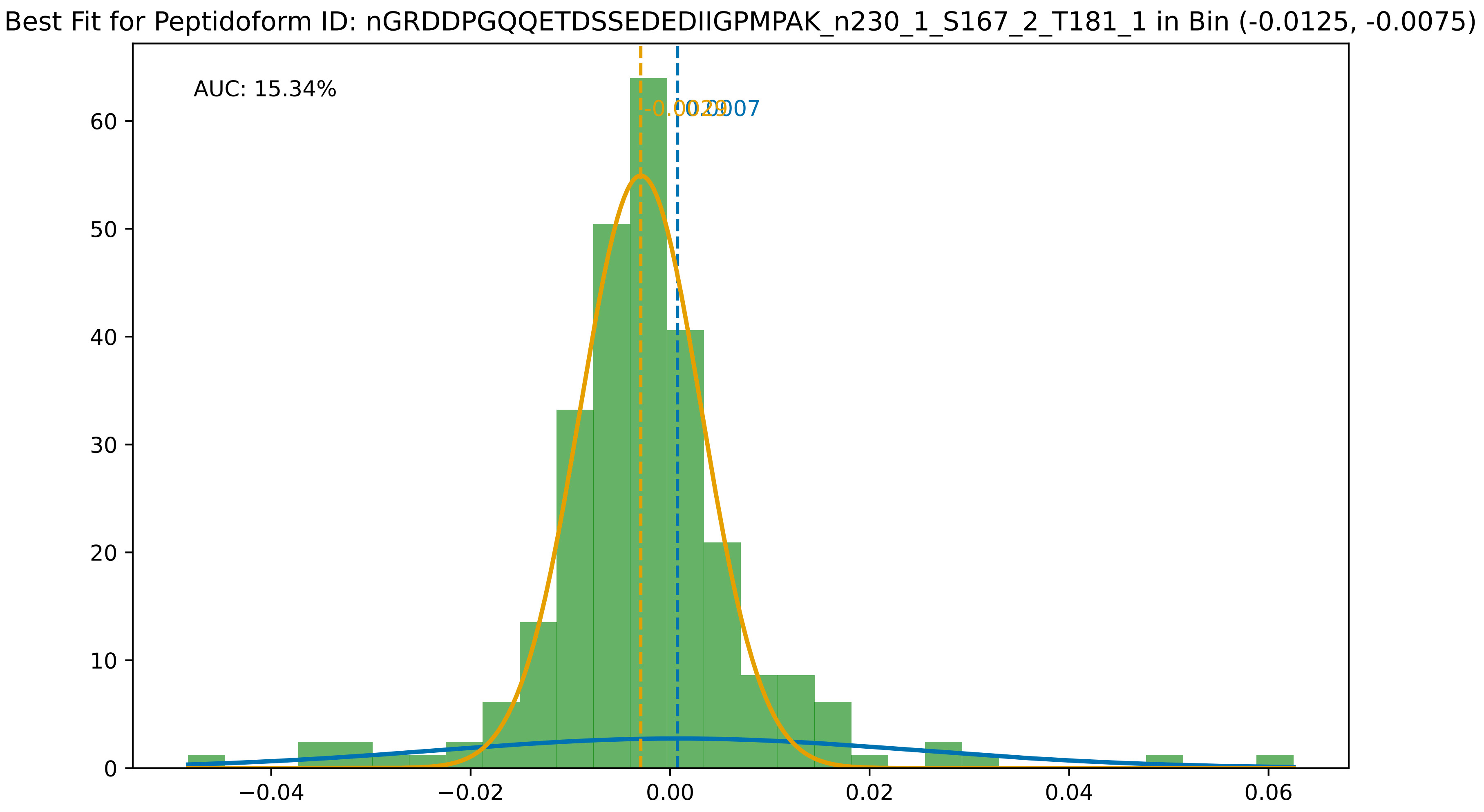
Best Fit for Peptidoform ID: nGRDDPGQQETDSSEDEDIIGPMPAK_n230_1_S167_2_T181_1 in Bin (-0.0125, -0.0075)
AUC: 15.34%
-0.0029
0.0007
0
10
20
30
40
50
60
−0.04
−0.02
0.00
0.02
0.04
0.06

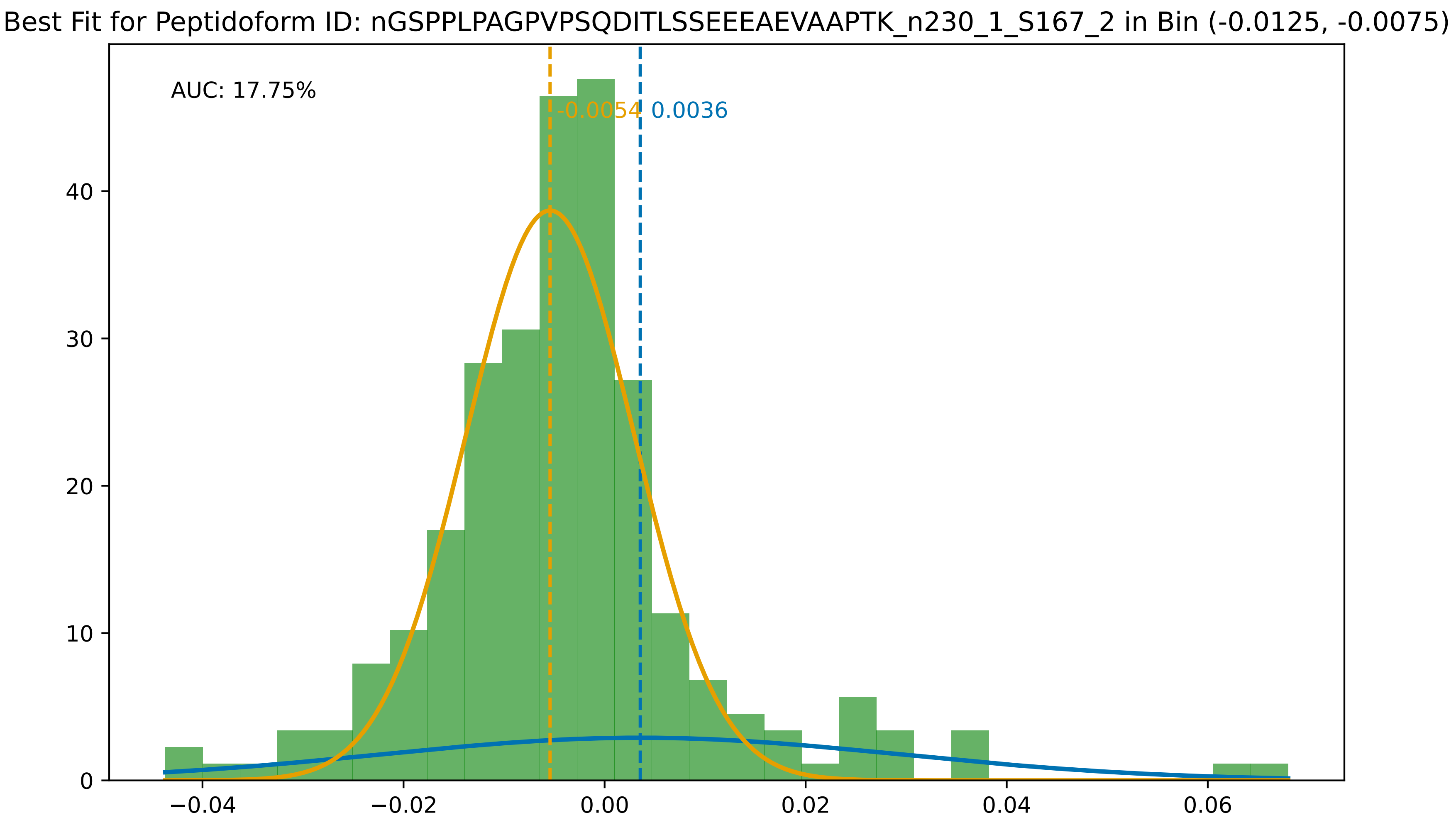

Best Fit for Peptidoform ID: nGSPPLPAGPVPSQDITLSSEEEAEVAAPTK_n230_1_S167_2 in Bin (-0.0125, -0.0075)
AUC: 17.75%
-0.0054
0.0036
0
10
20
30
40
−0.04
−0.02
0.00
0.02
0.04
0.06

for Peptidoform ID: nGTAGNALMDGASQLMGENRTMTIHNGMFFSTYDRDN_M147_1_n230_1_S167_1_T181_1 in Bin (-0.0125

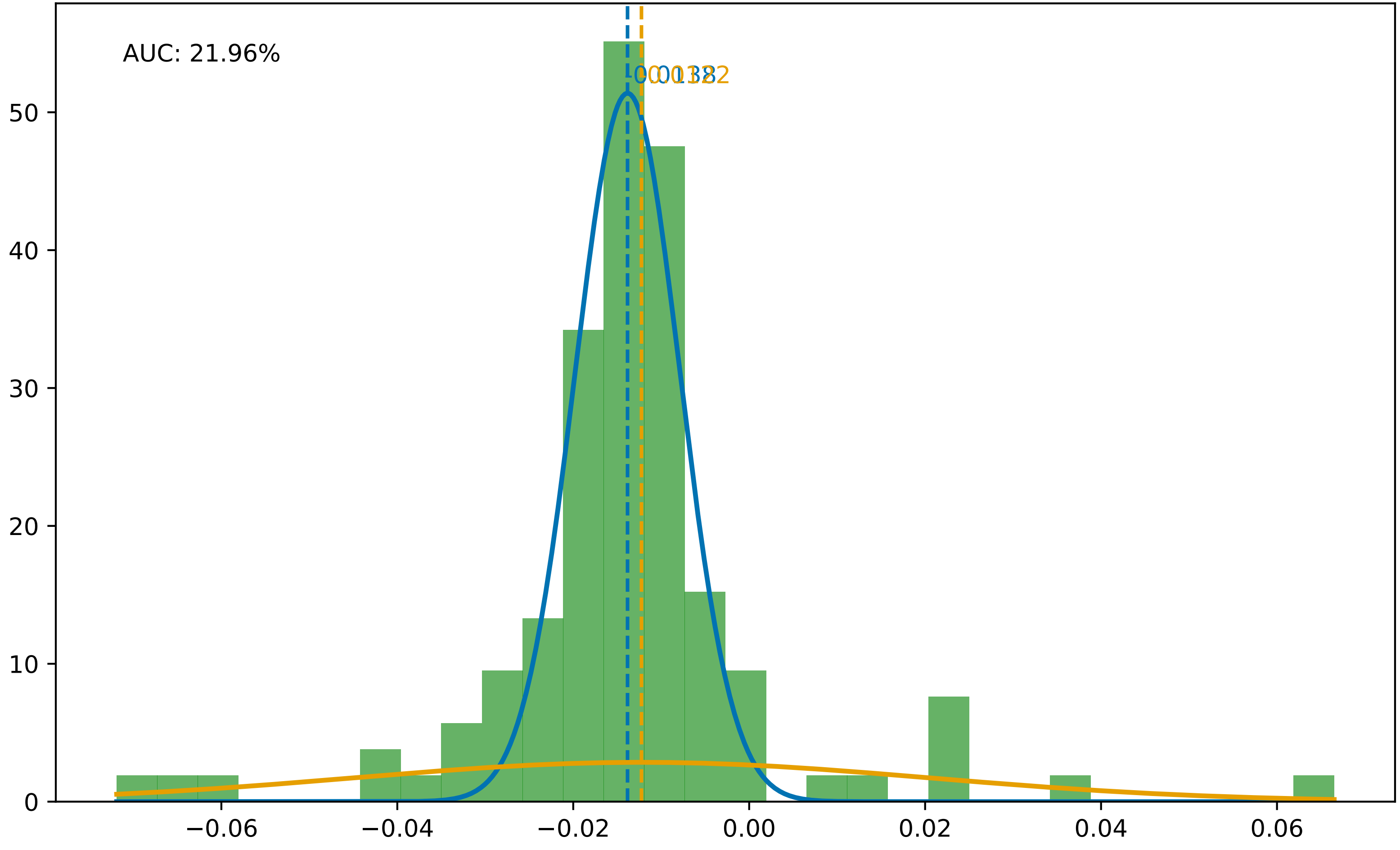

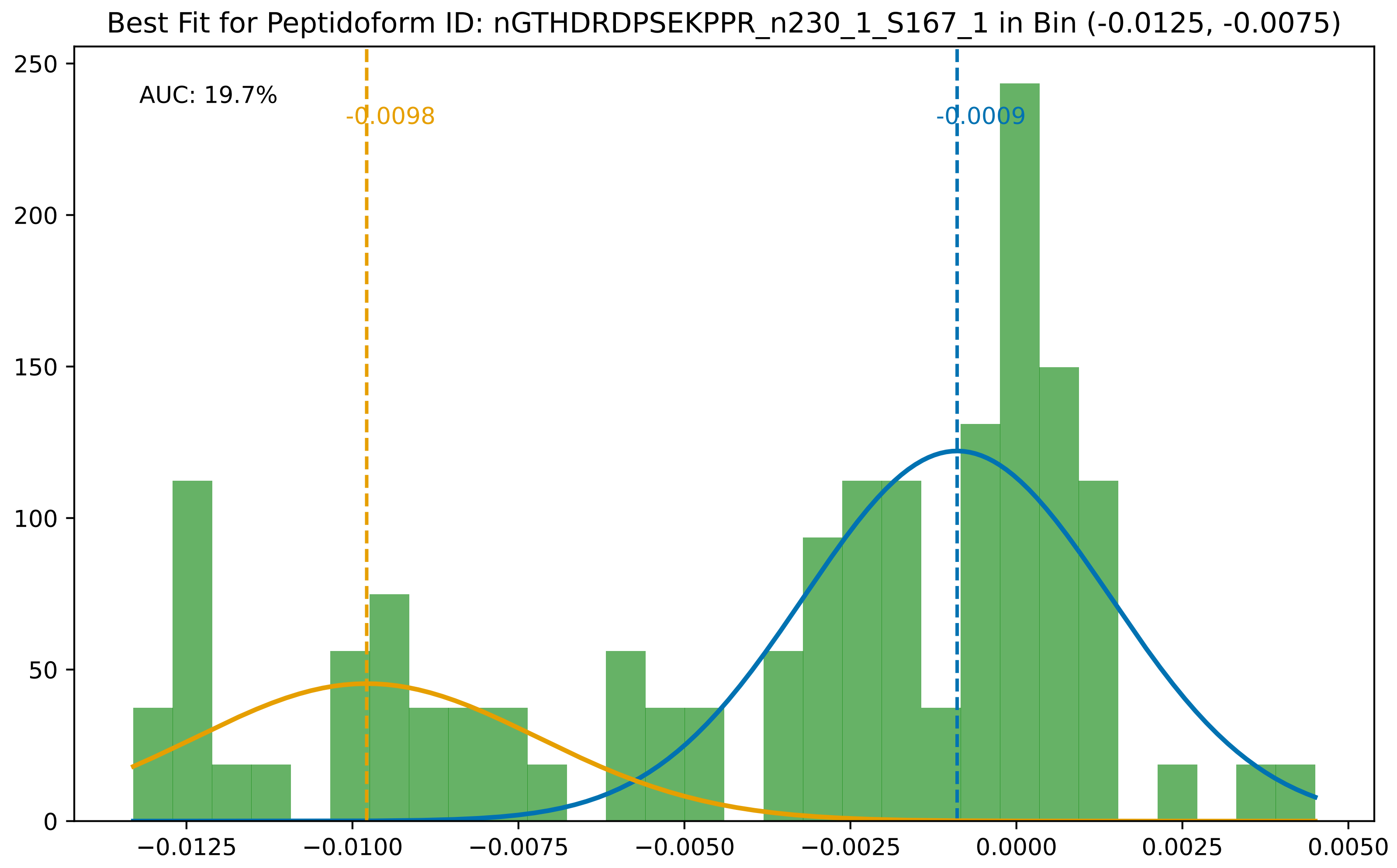

Best Fit for Peptidoform ID: nGTHDRDPSEKPPR_n230_1_S167_1 in Bin (-0.0125, -0.0075)
AUC: 19.7%
-0.0098
-0.0009
250
200
150
100
50
0
−0.0125
−0.0100
−0.0075
−0.0050
−0.0025
0.0000
0.0025
0.0050

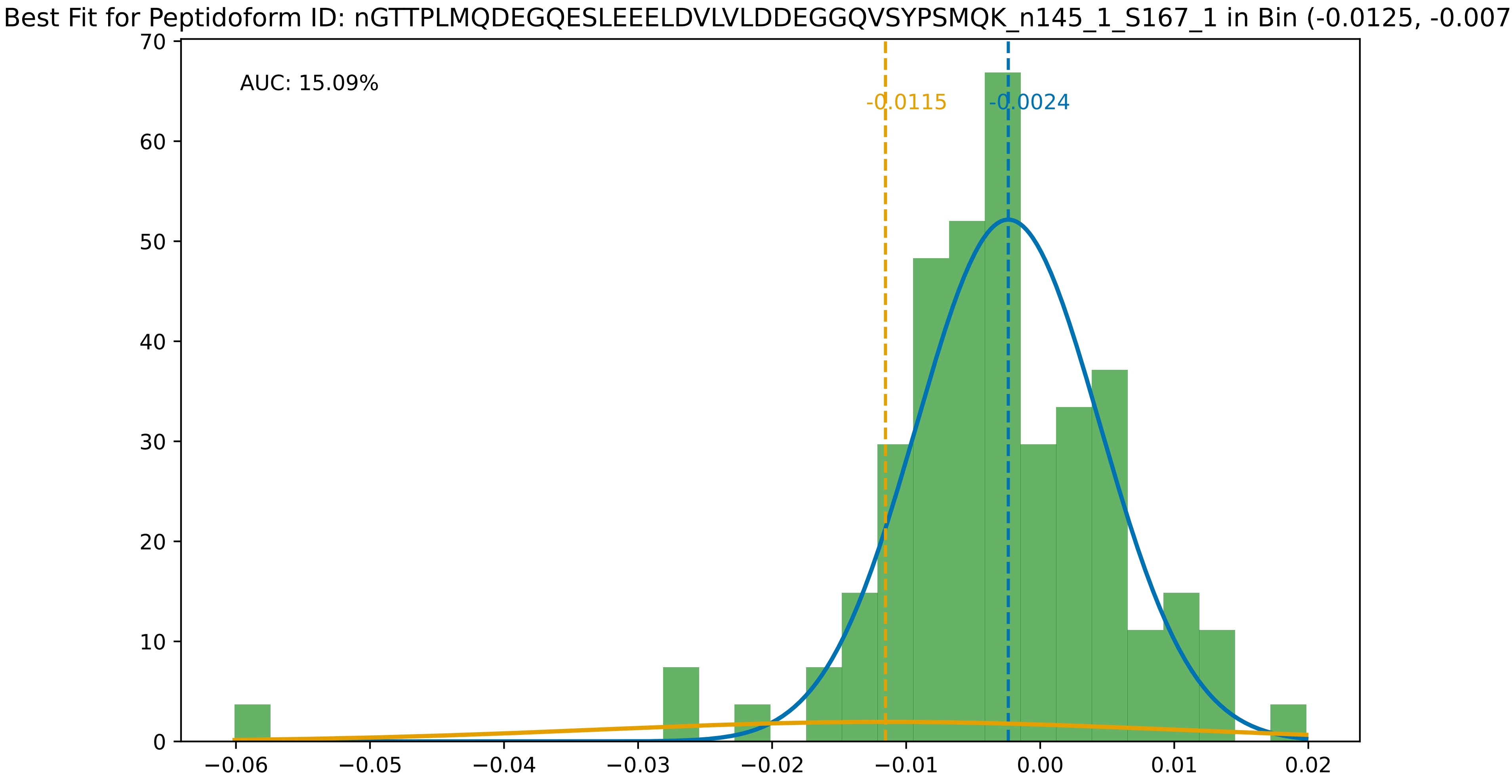
Best Fit for Peptidoform ID: nGTTPLMQDEGQESLEEELDVLVLDDEGGQVSYPSMQK_n145_1_S167_1 in Bin (-0.0125, -0.007
AUC: 15.09%
-0.0115
-0.0024
0
10
20
30
40
50
60
70
−0.06
−0.05
−0.04
−0.03
−0.02
−0.01
0.00
0.01
0.02

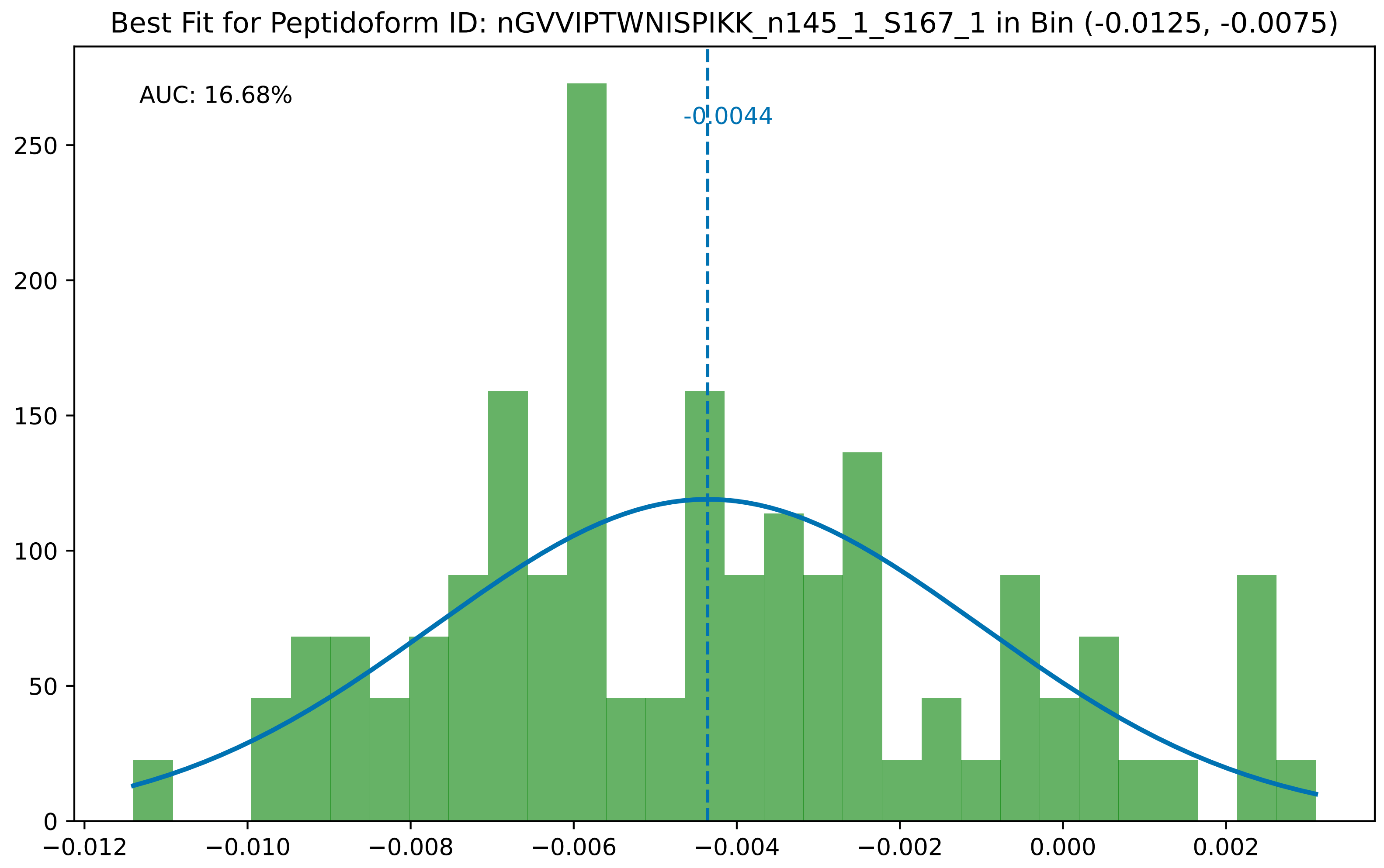
Best Fit for Peptidoform ID: nGVVIPTWNISPIKK_n145_1_S167_1 in Bin (-0.0125, -0.0075)
AUC: 16.68%
-0.0044
0
50
100
150
200
250
−0.012
−0.010
−0.008
−0.006
−0.004
−0.002
0.000
0.002

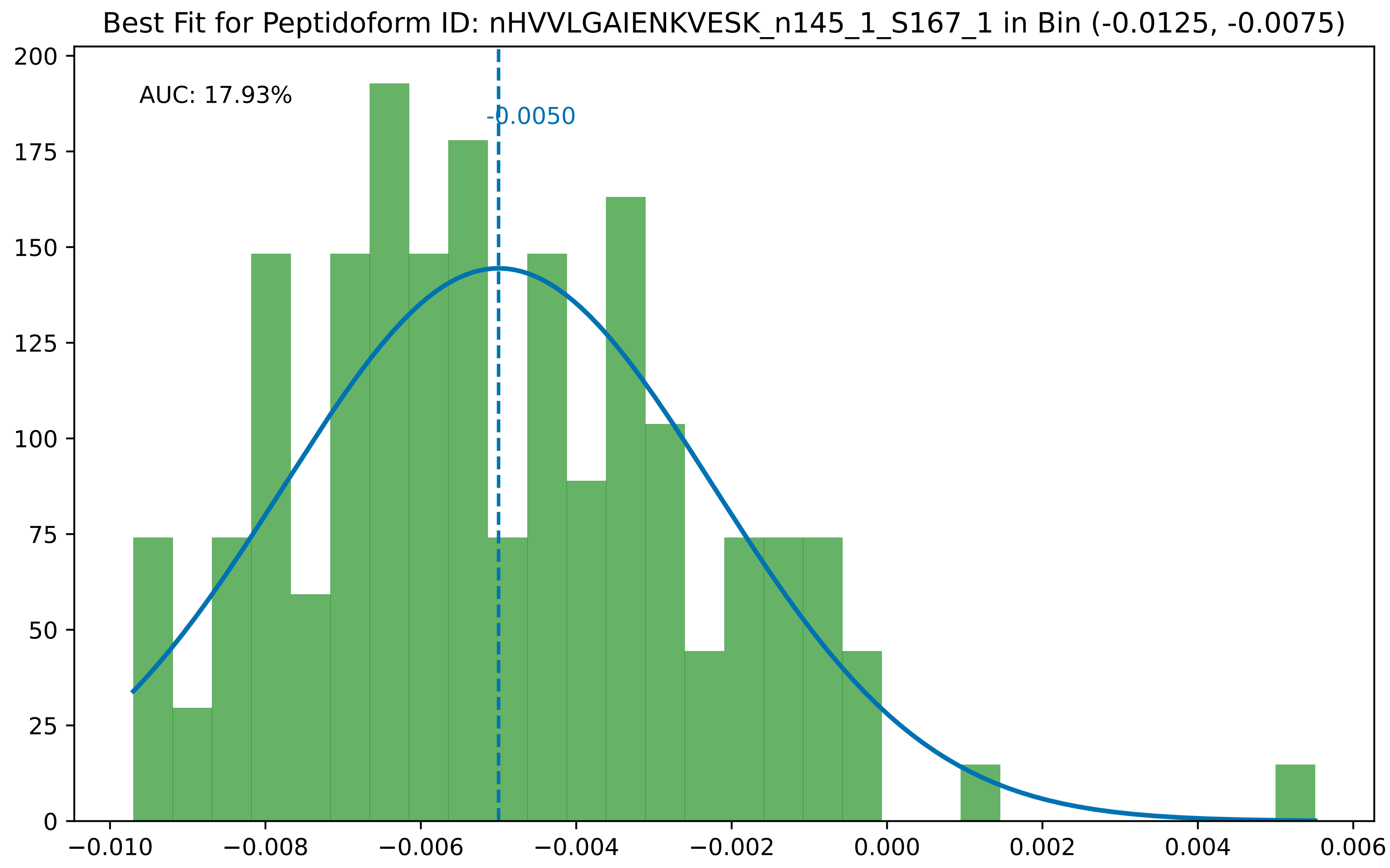

Best Fit for Peptidoform ID: nHVVLGAIENKVESK_n145_1_S167_1 in Bin (-0.0125, -0.0075)
AUC: 17.93%
-0.0050
200
175
150
125
100
75
50
25
0
−0.010
−0.008
−0.006
−0.004
−0.002
0.000
0.002
0.004
0.006

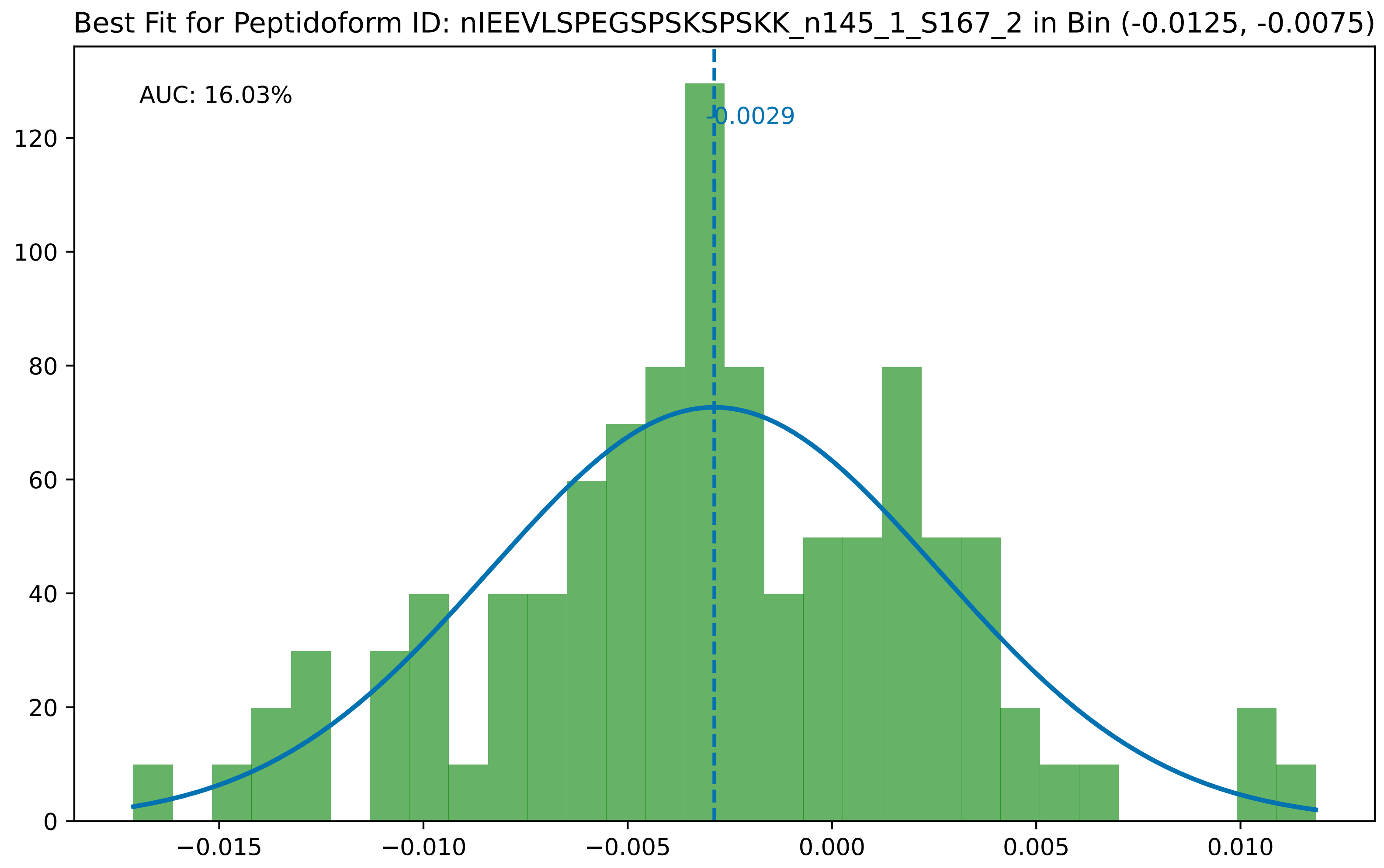

Best Fit for Peptidoform ID: nIEEVLSPEGSPSKSPSKK_n145_1_S167_2 in Bin (-0.0125, -0.0075)
AUC: 16.03%
-0.0029
0
20
40
60
80
100
120
−0.015
−0.010
−0.005
0.000
0.005
0.010

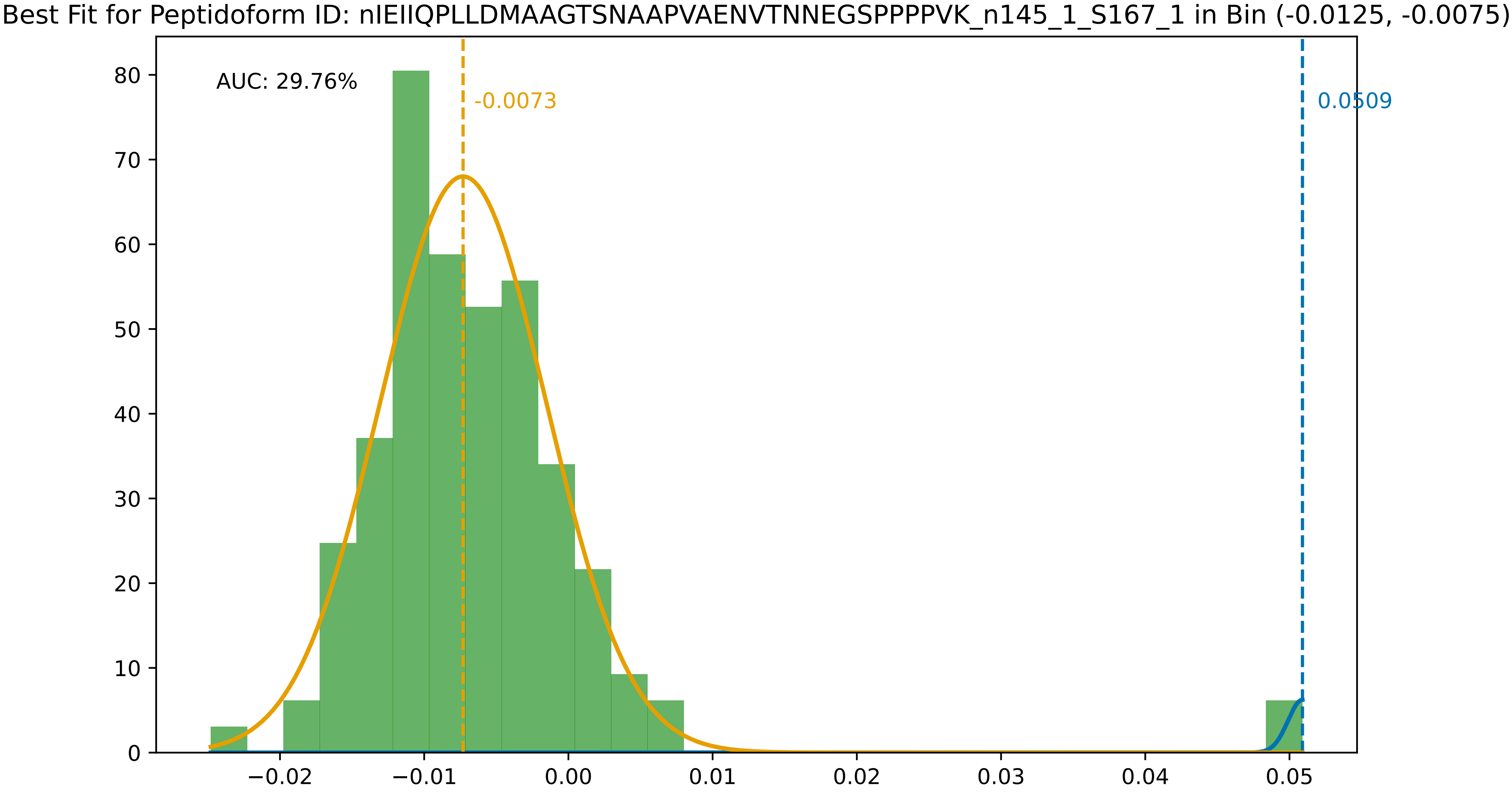

Best Fit for Peptidoform ID: nIEIIQPLLDMAAGTSNAAPVAENVTNNEGSPPPPVK_n145_1_S167_1 in Bin (-0.0125, -0.0075)
AUC: 29.76%
-0.0073
0.0509
0
10
20
30
40
50
60
70
80
−0.02
−0.01
0.00
0.01
0.02
0.03
0.04
0.05

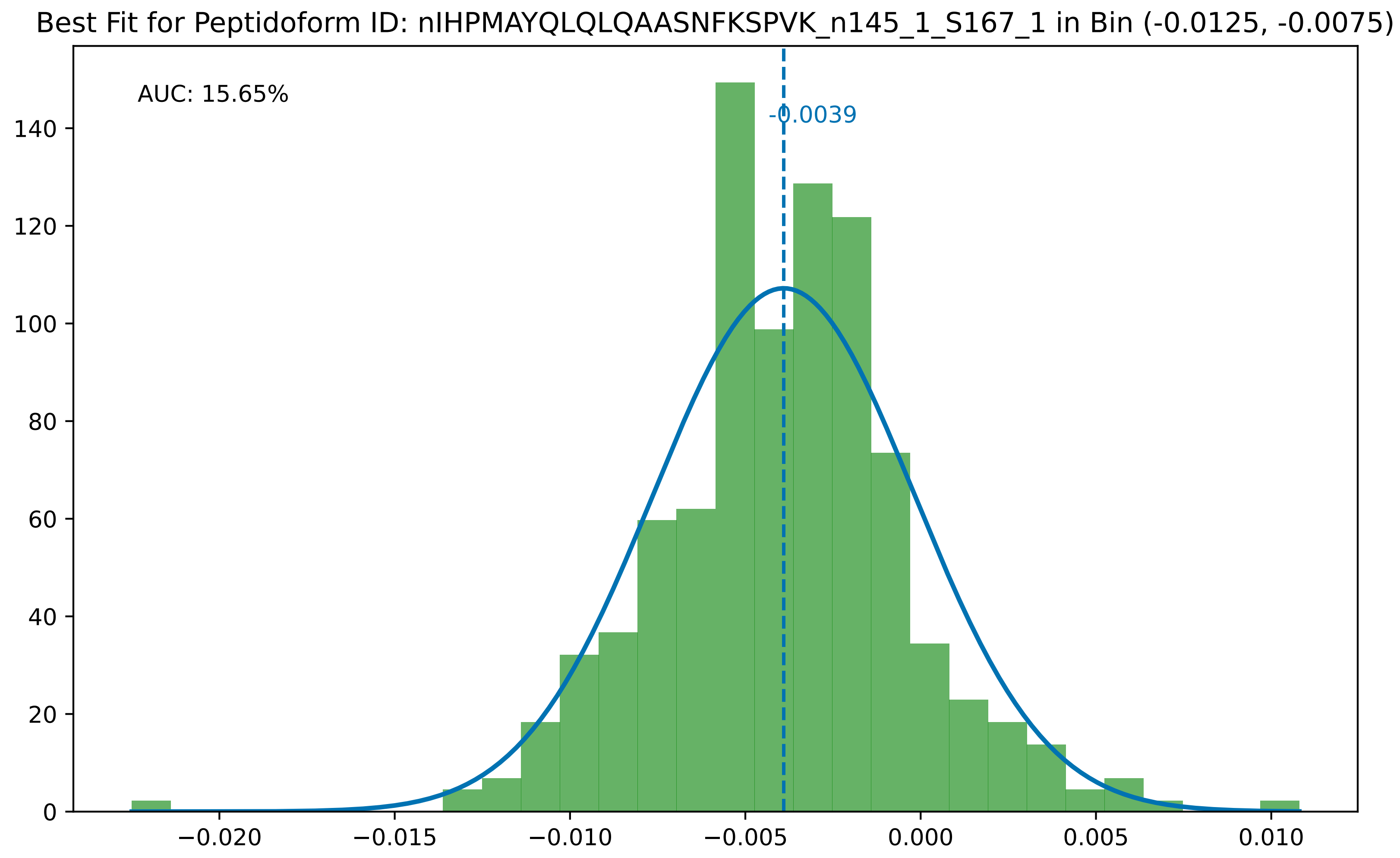

Best Fit for Peptidoform ID: nIHPMAYQLQLQAASNFKSPVK_n145_1_S167_1 in Bin (-0.0125, -0.0075)
AUC: 15.65%
-0.0039
0
20
40
60
80
100
120
140
−0.020
−0.015
−0.010
−0.005
0.000
0.005
0.010

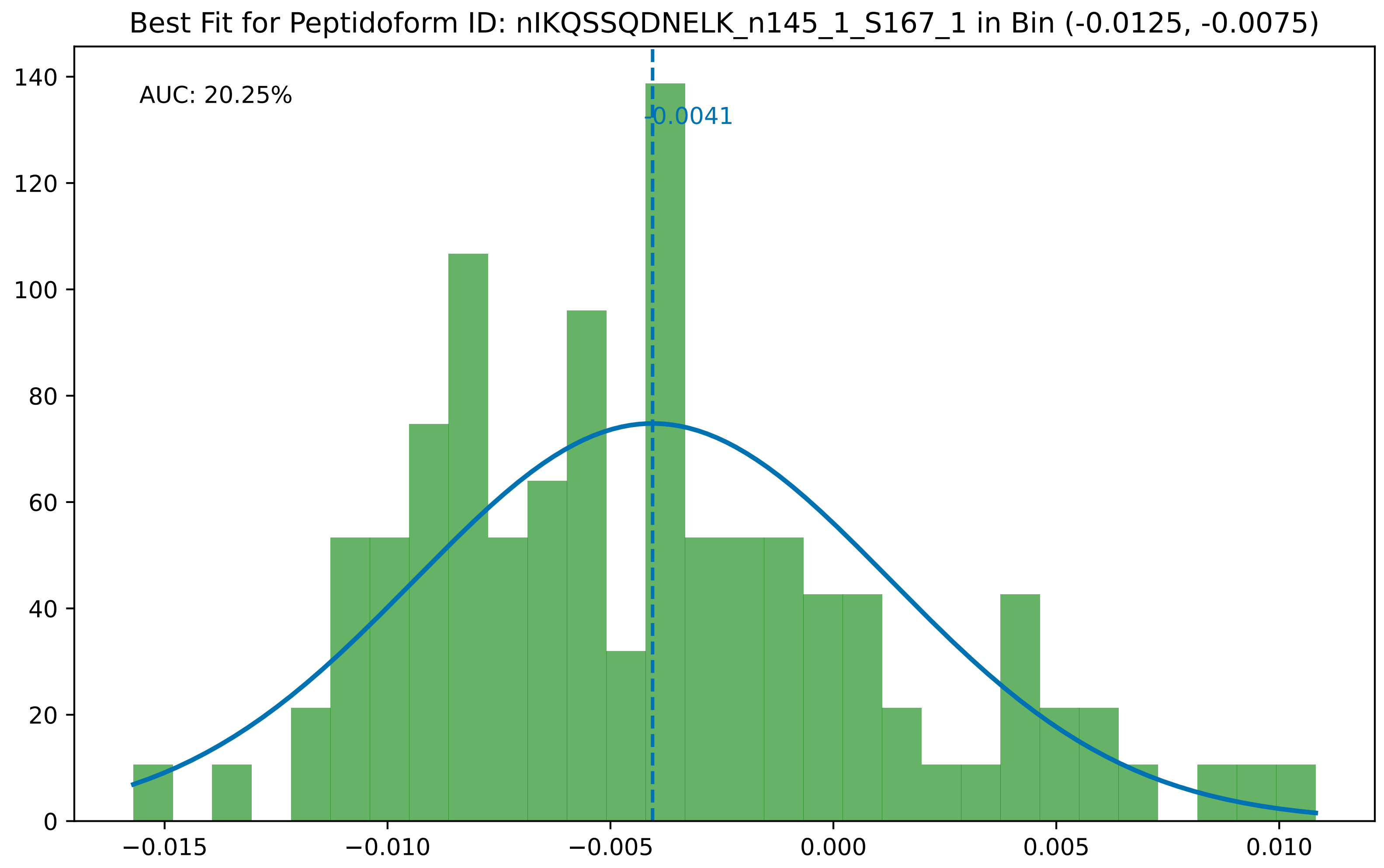
Best Fit for Peptidoform ID: nIKQSSQDNELK_n145_1_S167_1 in Bin (-0.0125, -0.0075)
AUC: 20.25%
-0.0041
0
20
40
60
80
100
120
140
−0.015
−0.010
−0.005
0.000
0.005
0.010

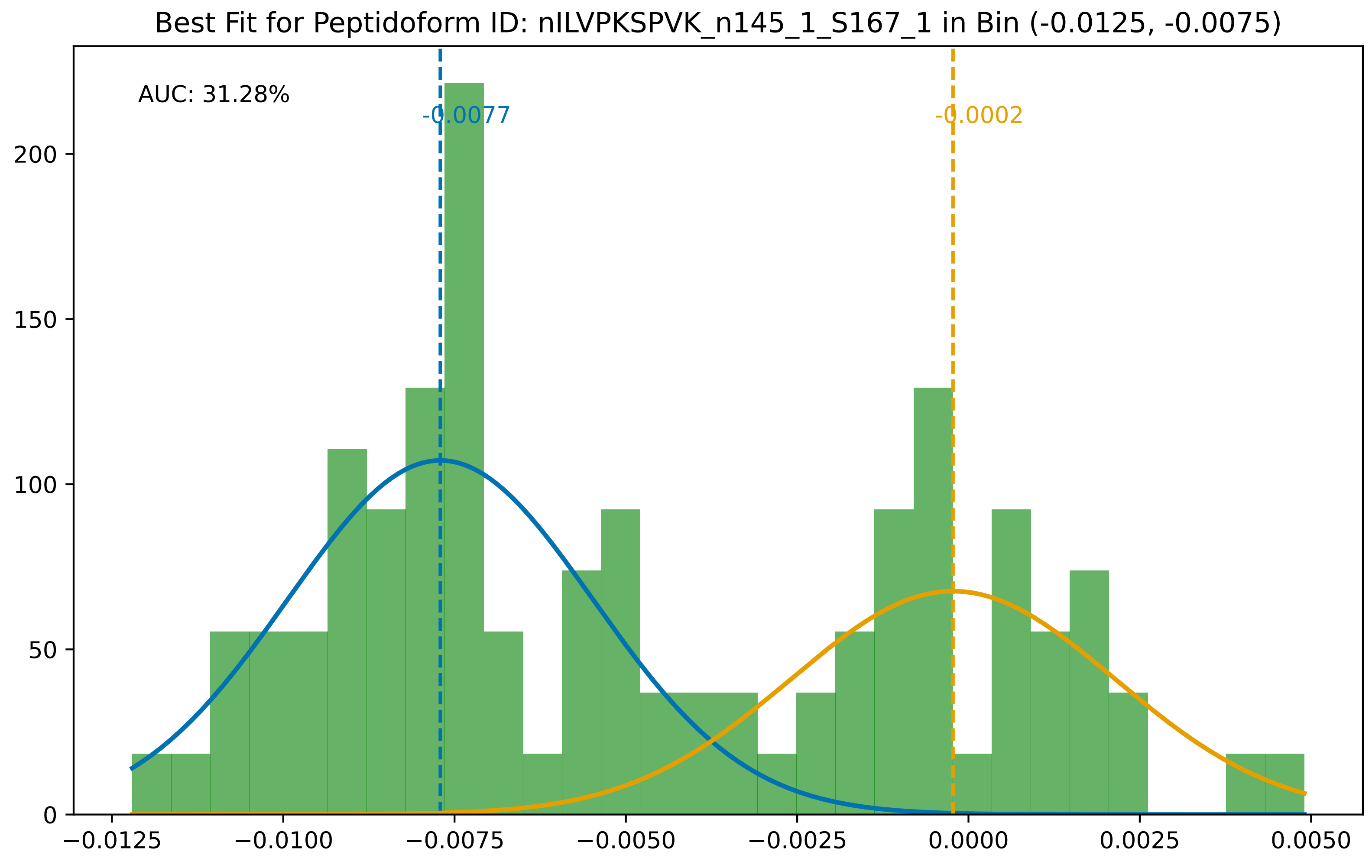

Best Fit for Peptidoform ID: nILVPKSPVK_n145_1_S167_1 in Bin (-0.0125, -0.0075)
AUC: 31.28%
-0.0077
-0.0002
0
50
100
150
200
−0.0125
−0.0100
−0.0075
−0.0050
−0.0025
0.0000
0.0025
0.0050

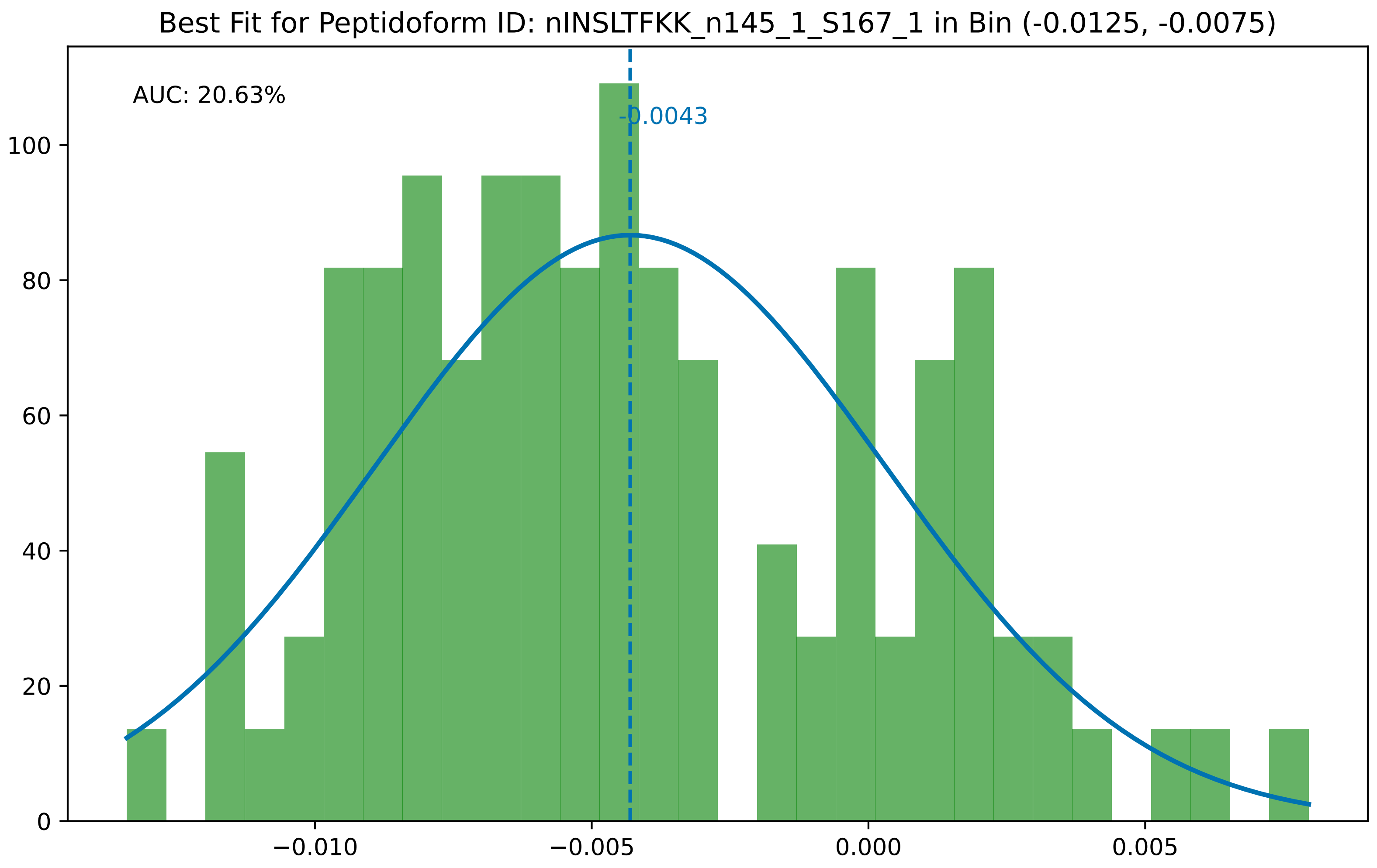

Best Fit for Peptidoform ID: nINSLTFKK_n145_1_S167_1 in Bin (-0.0125, -0.0075)
AUC: 20.63%
-0.0043
100
80
60
40
20
0
−0.010
−0.005
0.000
0.005

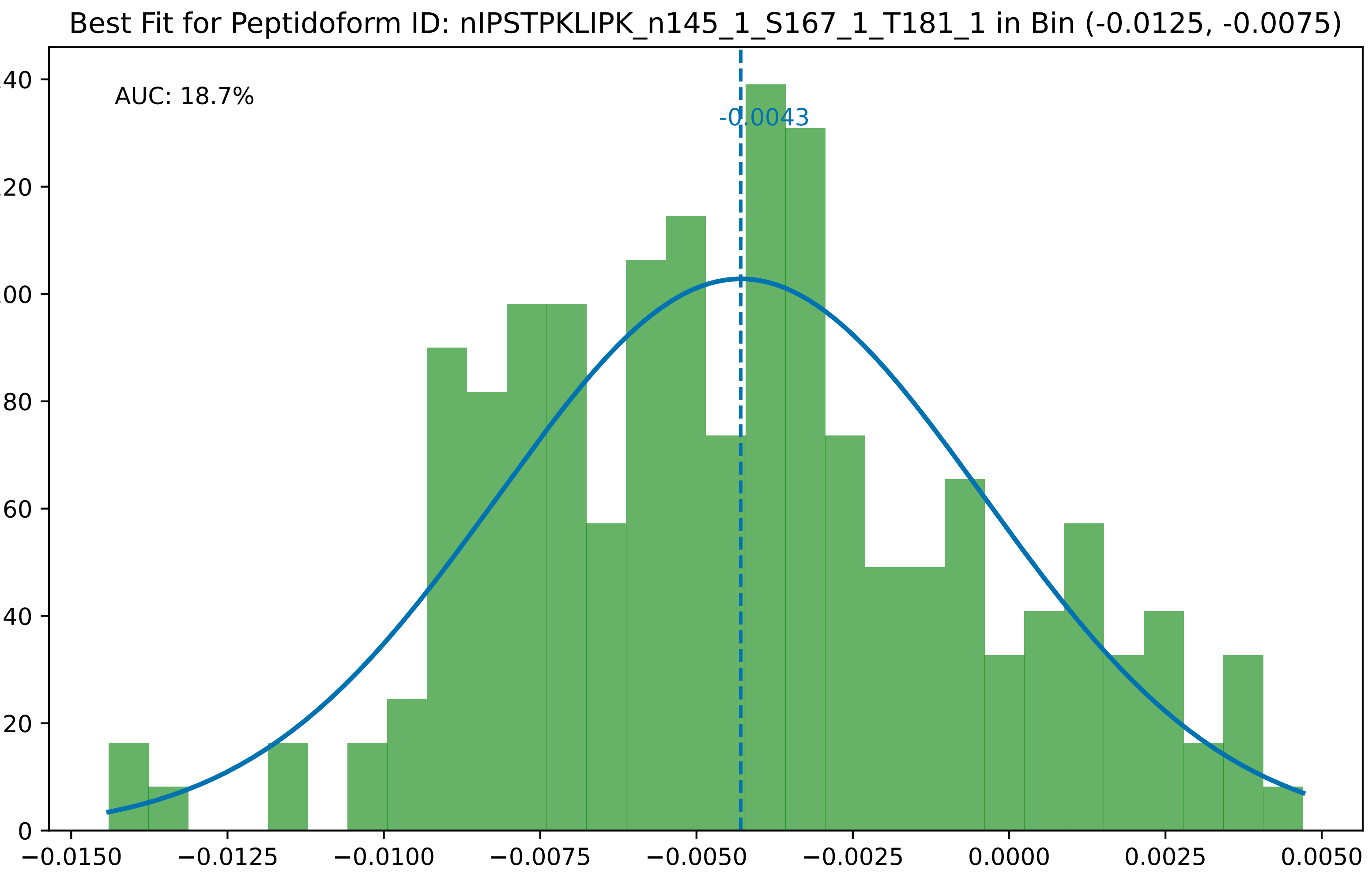
Best Fit for Peptidoform ID: nIPSTPKLIPK_n145_1_S167_1_T181_1 in Bin (-0.0125, -0.0075)
AUC: 18.7%
-0.0043
0
20
40
60
80
100
120
140
−0.0150
−0.0125
−0.0100
−0.0075
−0.0050
−0.0025
0.0000
0.0025
0.0050

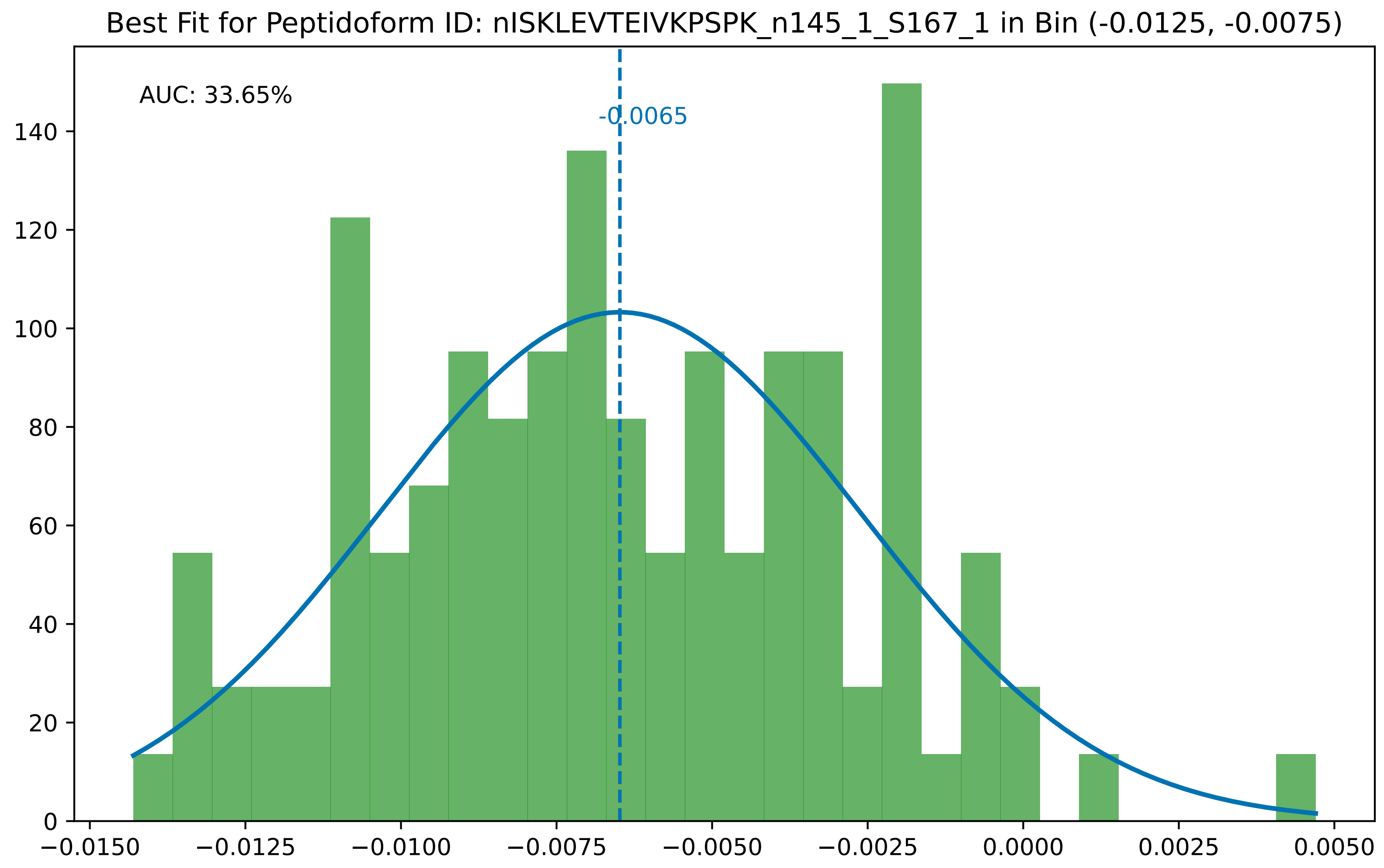
Best Fit for Peptidoform ID: nISKLEVTEIVKPSPK_n145_1_S167_1 in Bin (-0.0125, -0.0075)
AUC: 33.65%
-0.0065
0
20
40
60
80
100
120
140
−0.0150
−0.0125
−0.0100
−0.0075
−0.0050
−0.0025
0.0000
0.0025
0.0050

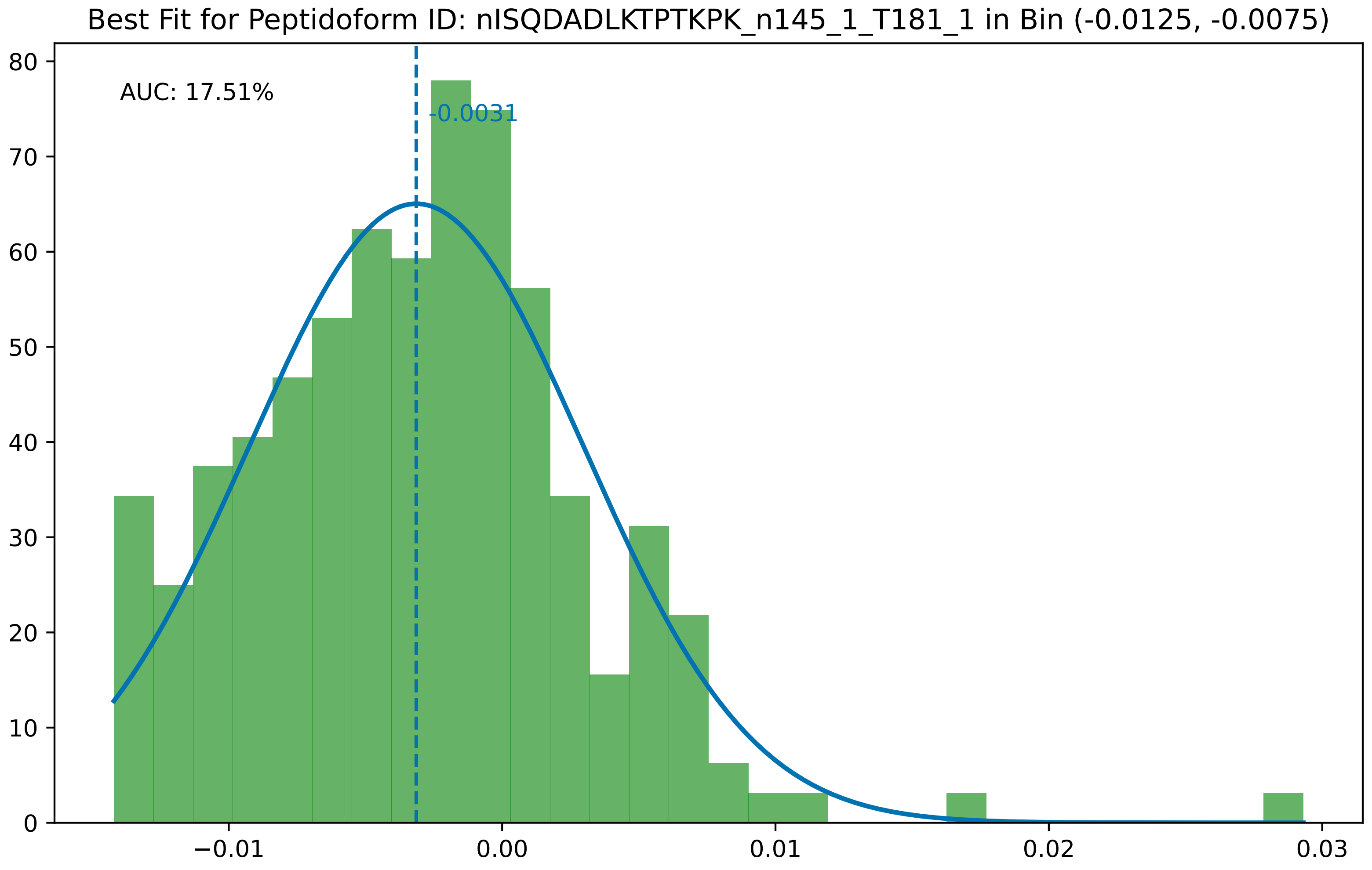
Best Fit for Peptidoform ID: nISQDADLKTPTKPK_n145_1_T181_1 in Bin (-0.0125, -0.0075)
AUC: 17.51%
-0.0031
0
10
20
30
40
50
60
70
80
−0.01
0.00
0.01
0.02
0.03

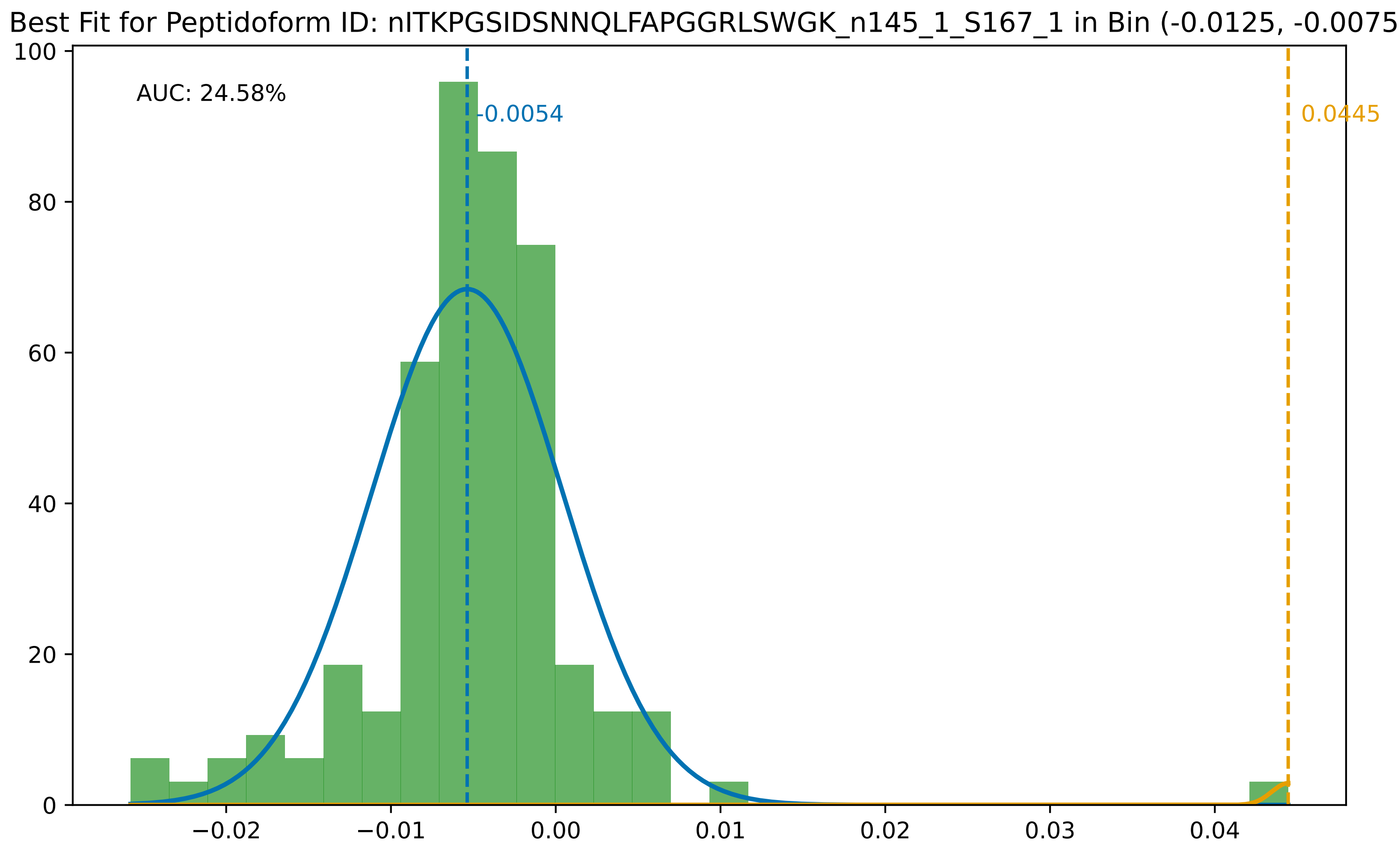

Best Fit for Peptidoform ID: nITKPGSIDSNNQLFAPGGRLSWGK_n145_1_S167_1 in Bin (-0.0125, -0.0075)
AUC: 24.58%
-0.0054
0.0445
100
80
60
40
20
0
−0.02
−0.01
0.00
0.01
0.02
0.03
0.04

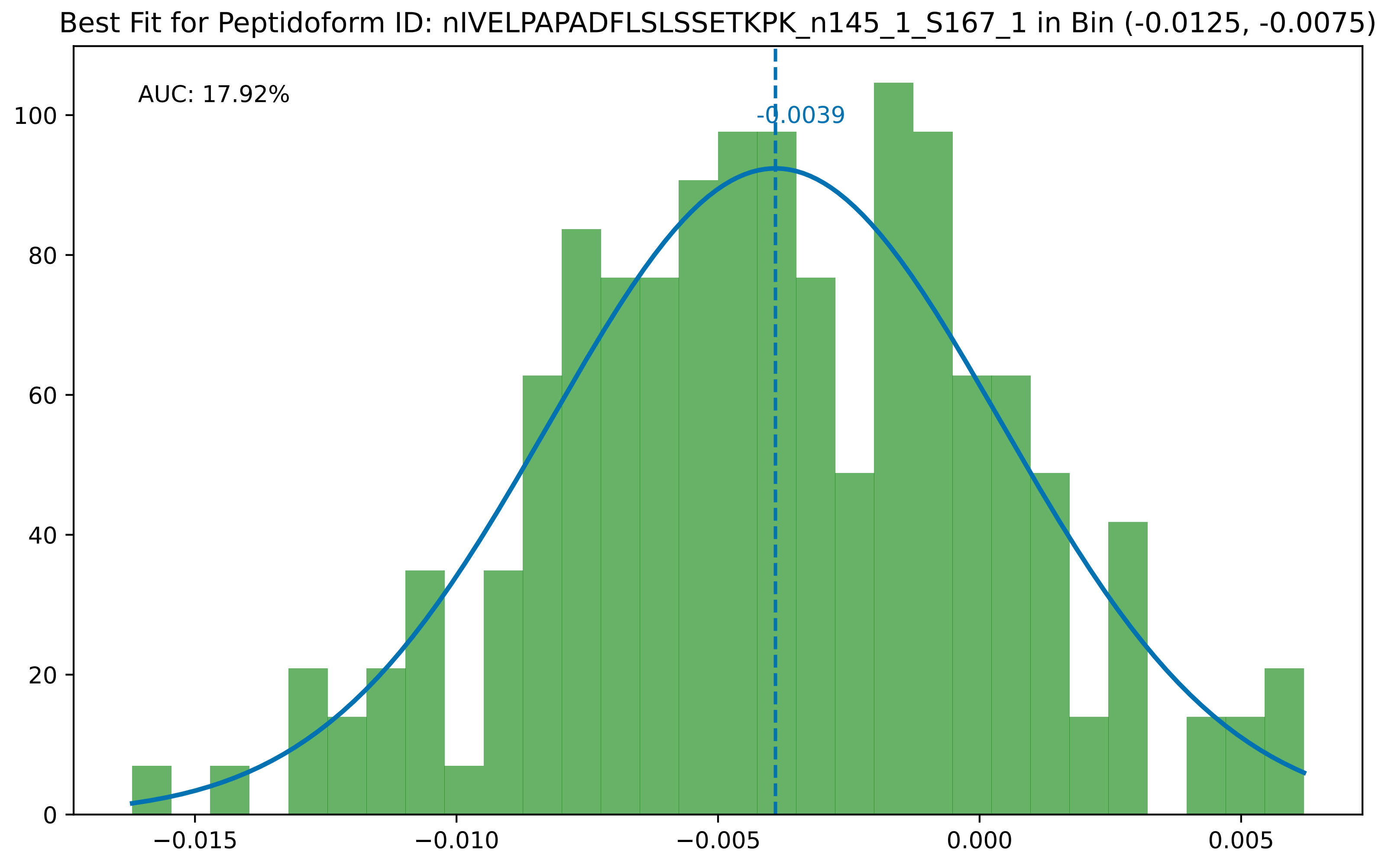
Best Fit for Peptidoform ID: nIVELPAPADFLSLSSETKPK_n145_1_S167_1 in Bin (-0.0125, -0.0075)
AUC: 17.92%
-0.0039
0
20
40
60
80
100
−0.015
−0.010
−0.005
0.000
0.005

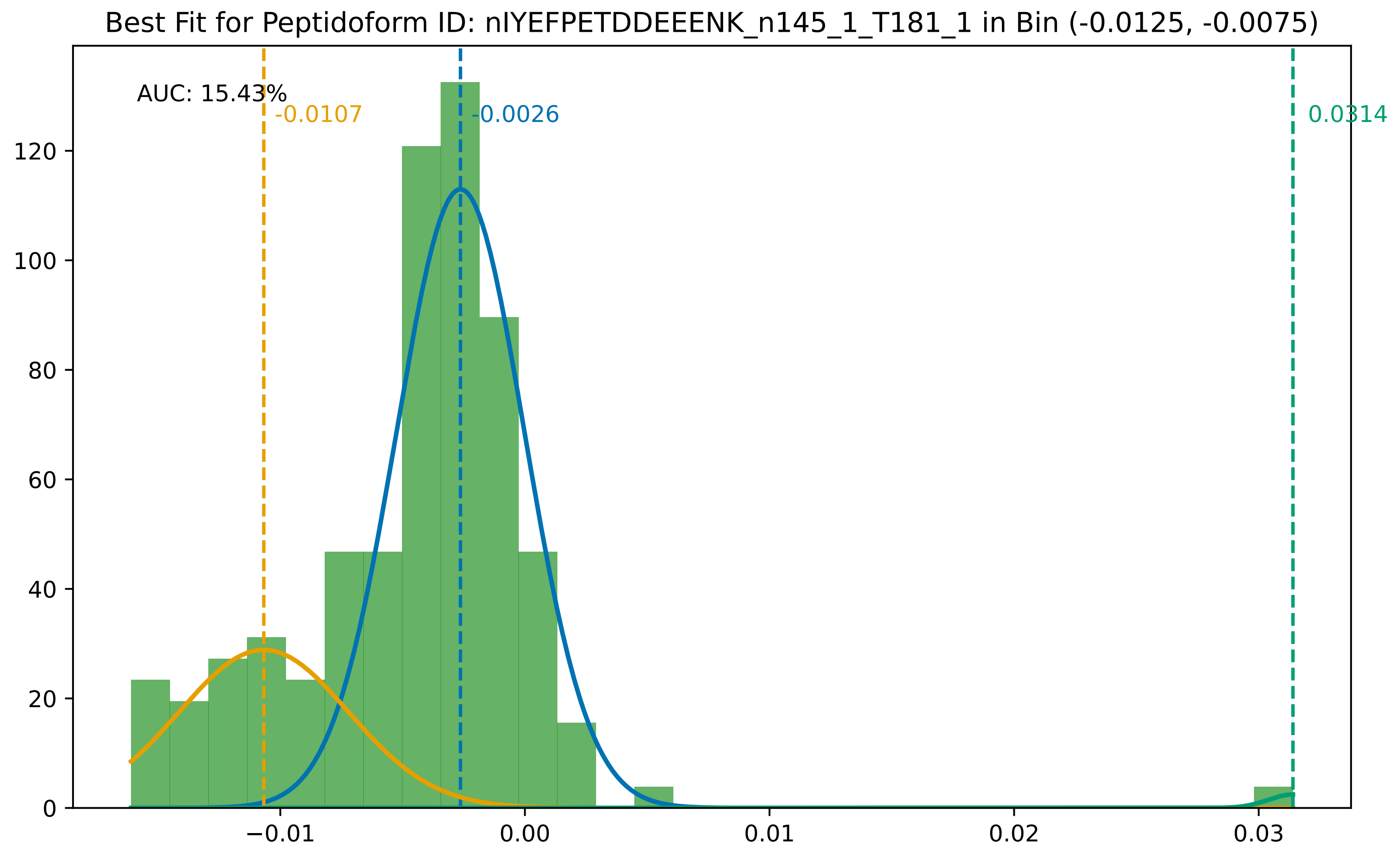
Best Fit for Peptidoform ID: nIYEFPETDDEEENK_n145_1_T181_1 in Bin (-0.0125, -0.0075)
AUC: 15.43%
-0.0107
-0.0026
0.0314
0
20
40
60
80
100
120
−0.01
0.00
0.01
0.02
0.03

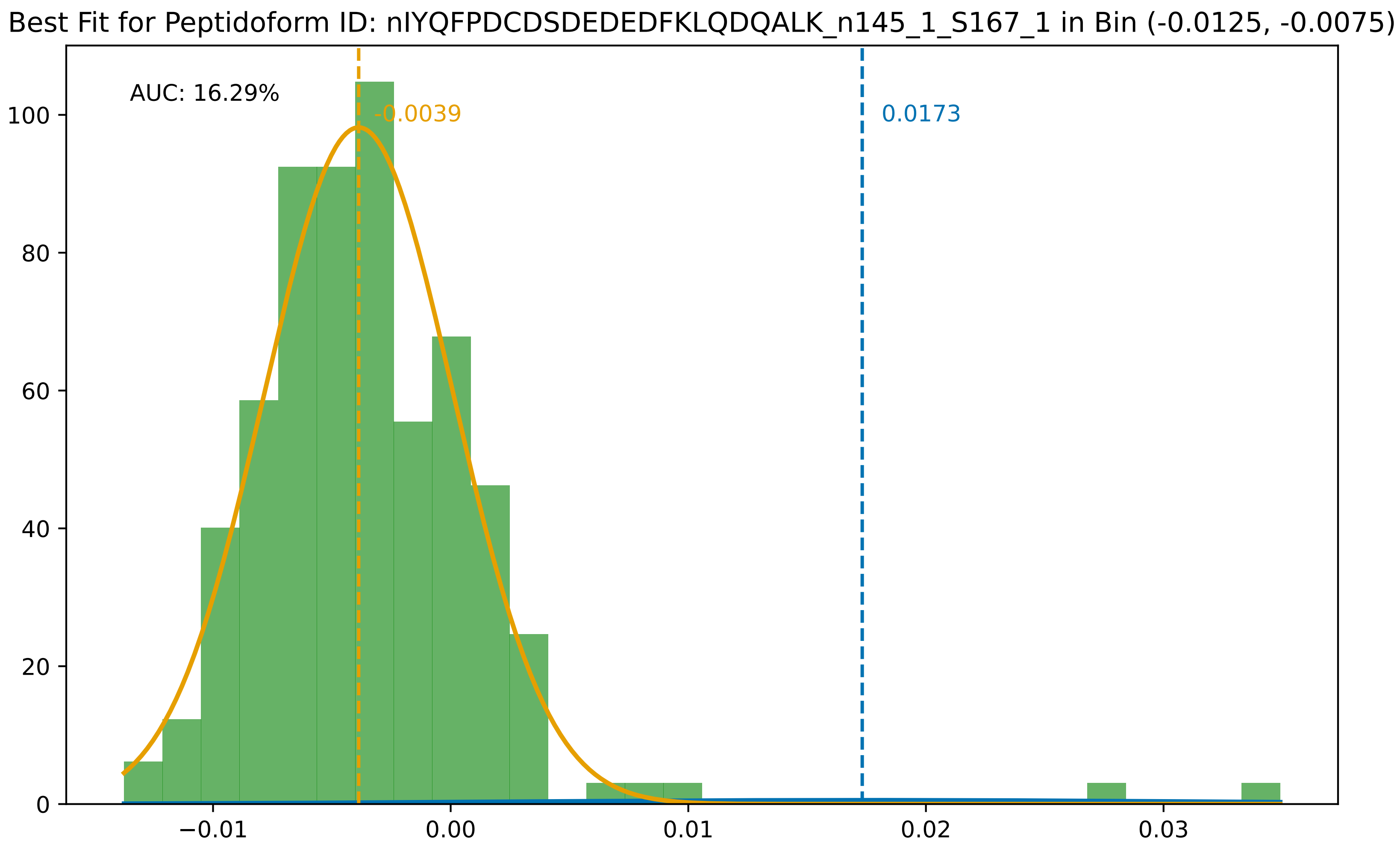

Best Fit for Peptidoform ID: nIYQFPDCDSDEDEDFKLQDQALK_n145_1_S167_1 in Bin (-0.0125, -0.0075)
AUC: 16.29%
-0.0039
0.0173
0
20
40
60
80
100
−0.01
0.00
0.01
0.02
0.03

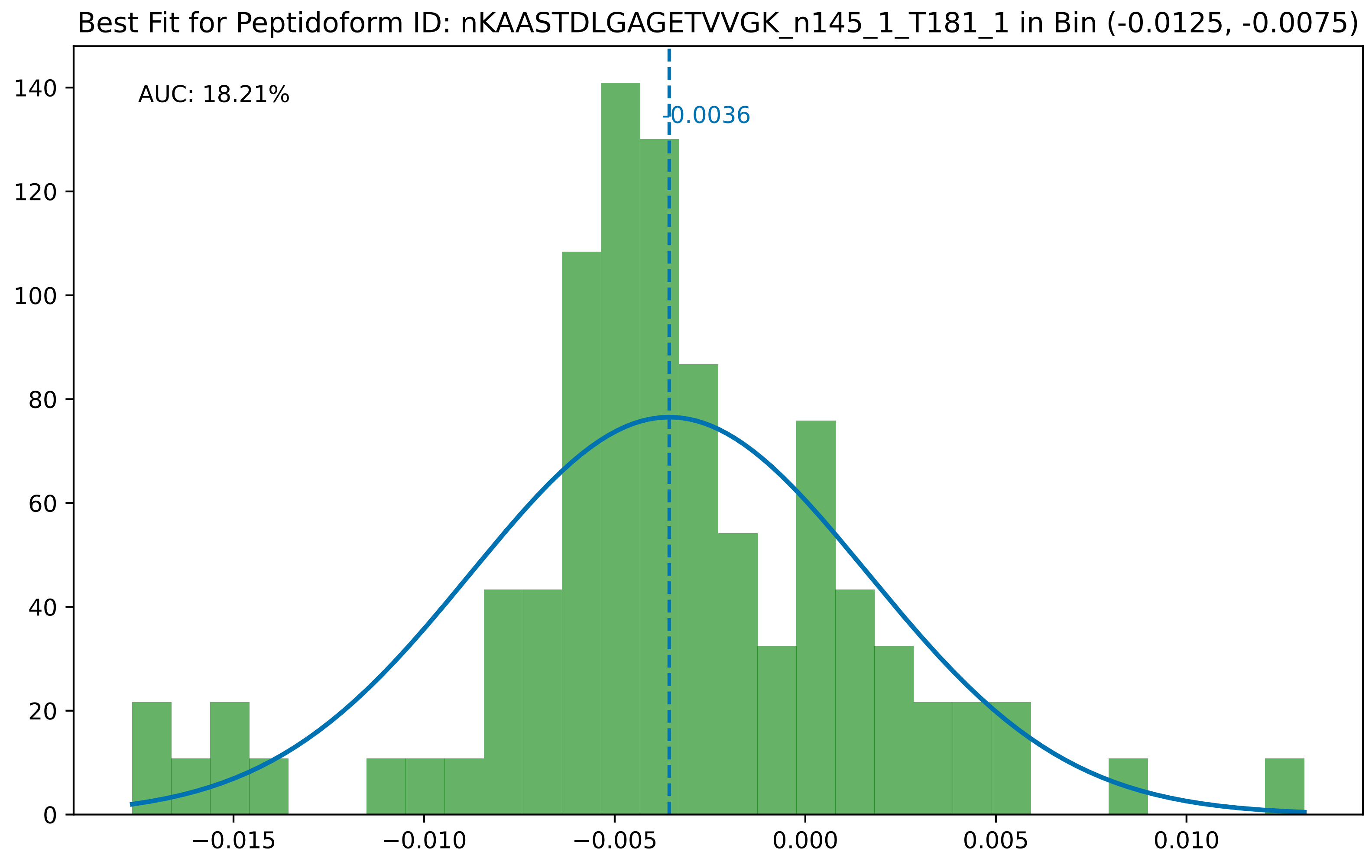

Best Fit for Peptidoform ID: nKAASTDLGAGETVVGK_n145_1_T181_1 in Bin (-0.0125, -0.0075)
AUC: 18.21%
-0.0036
0
20
40
60
80
100
120
140
−0.015
−0.010
−0.005
0.000
0.005
0.010

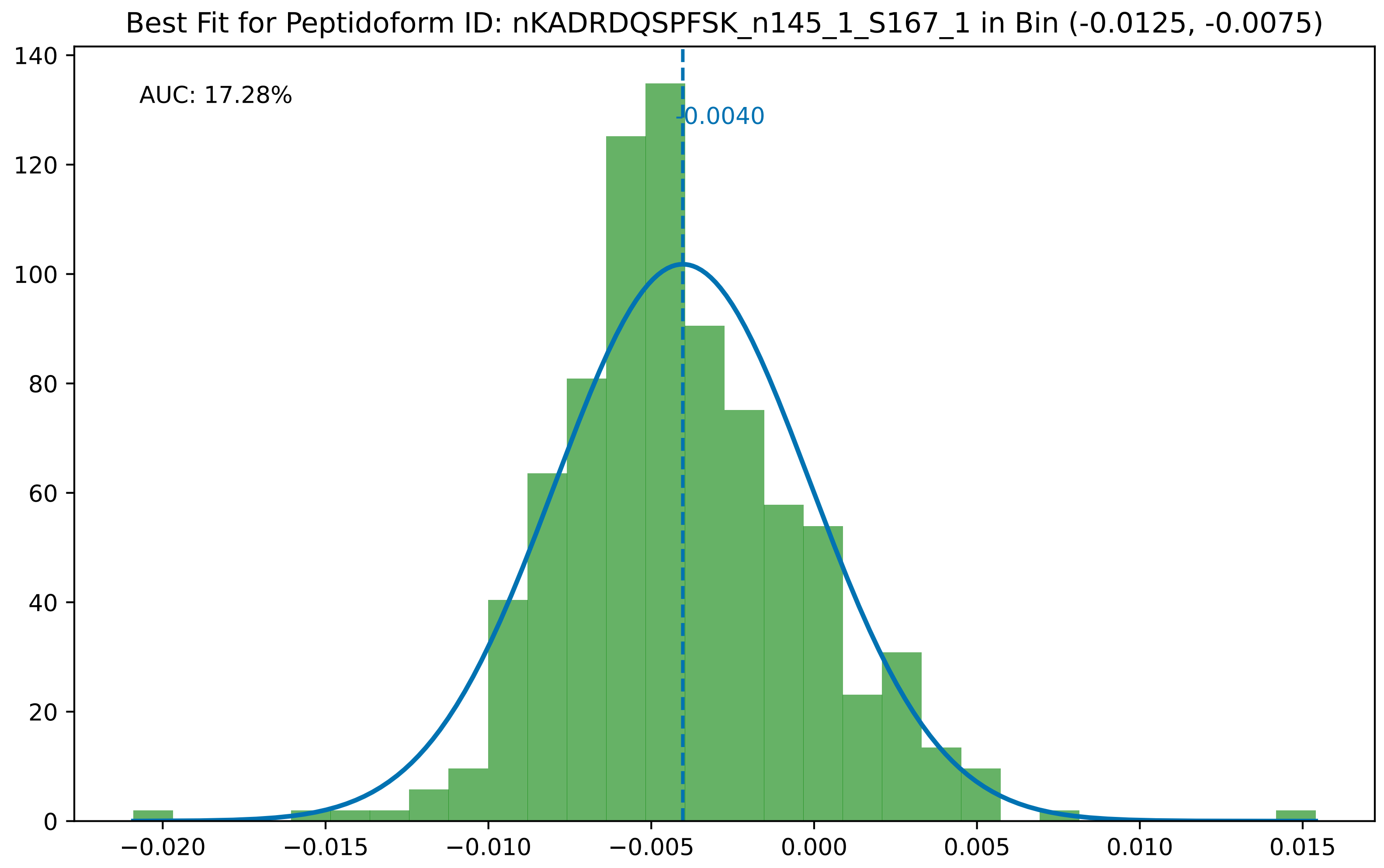

Best Fit for Peptidoform ID: nKADRDQSPFSK_n145_1_S167_1 in Bin (-0.0125, -0.0075)
AUC: 17.28%
-0.0040
0
20
40
60
80
100
120
140
−0.020
−0.015
−0.010
−0.005
0.000
0.005
0.010
0.015

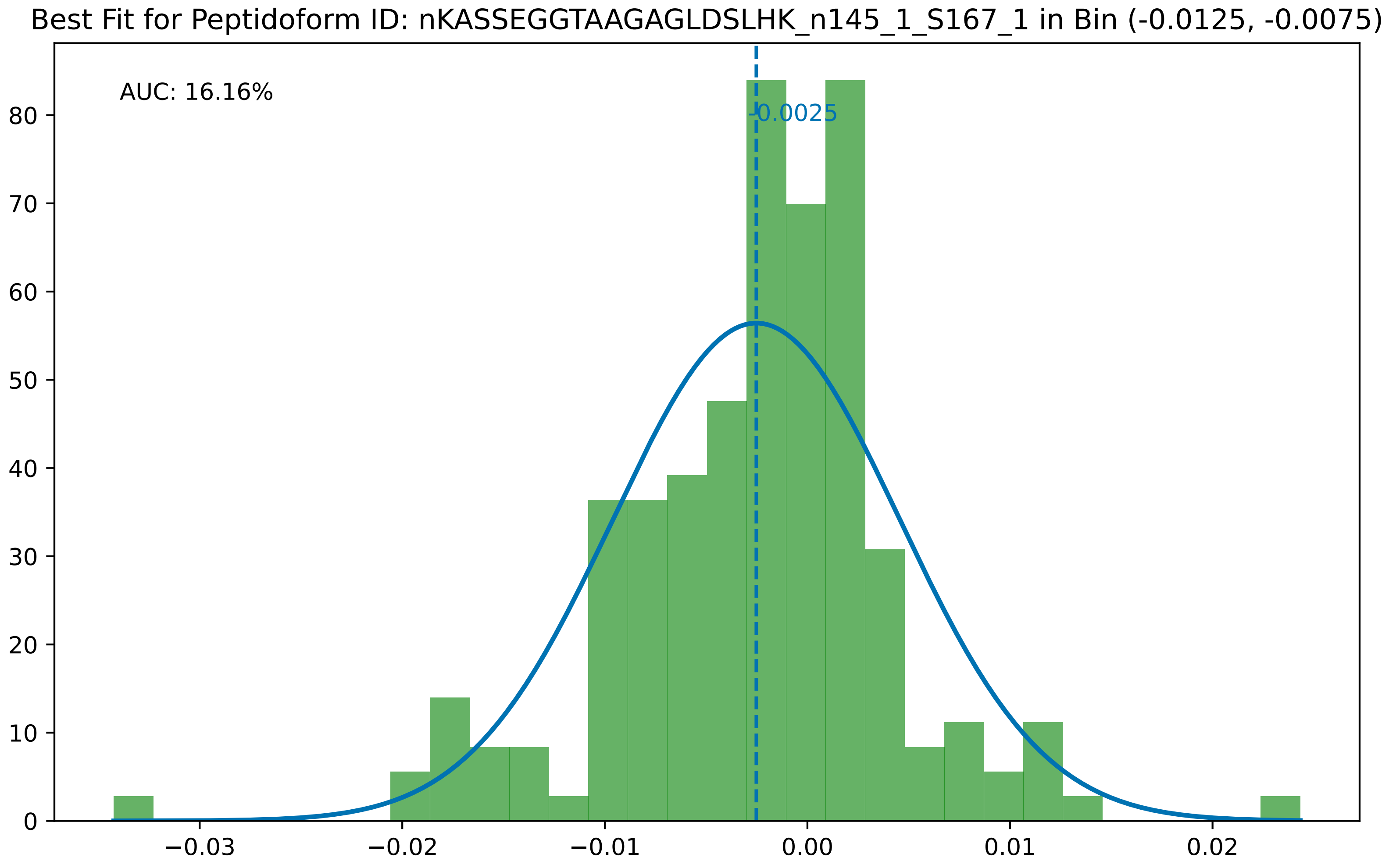

Best Fit for Peptidoform ID: nKASSEGGTAAGAGLDSLHK_n145_1_S167_1 in Bin (-0.0125, -0.0075)
AUC: 16.16%
-0.0025
0
10
20
30
40
50
60
70
80
−0.03
−0.02
−0.01
0.00
0.01
0.02

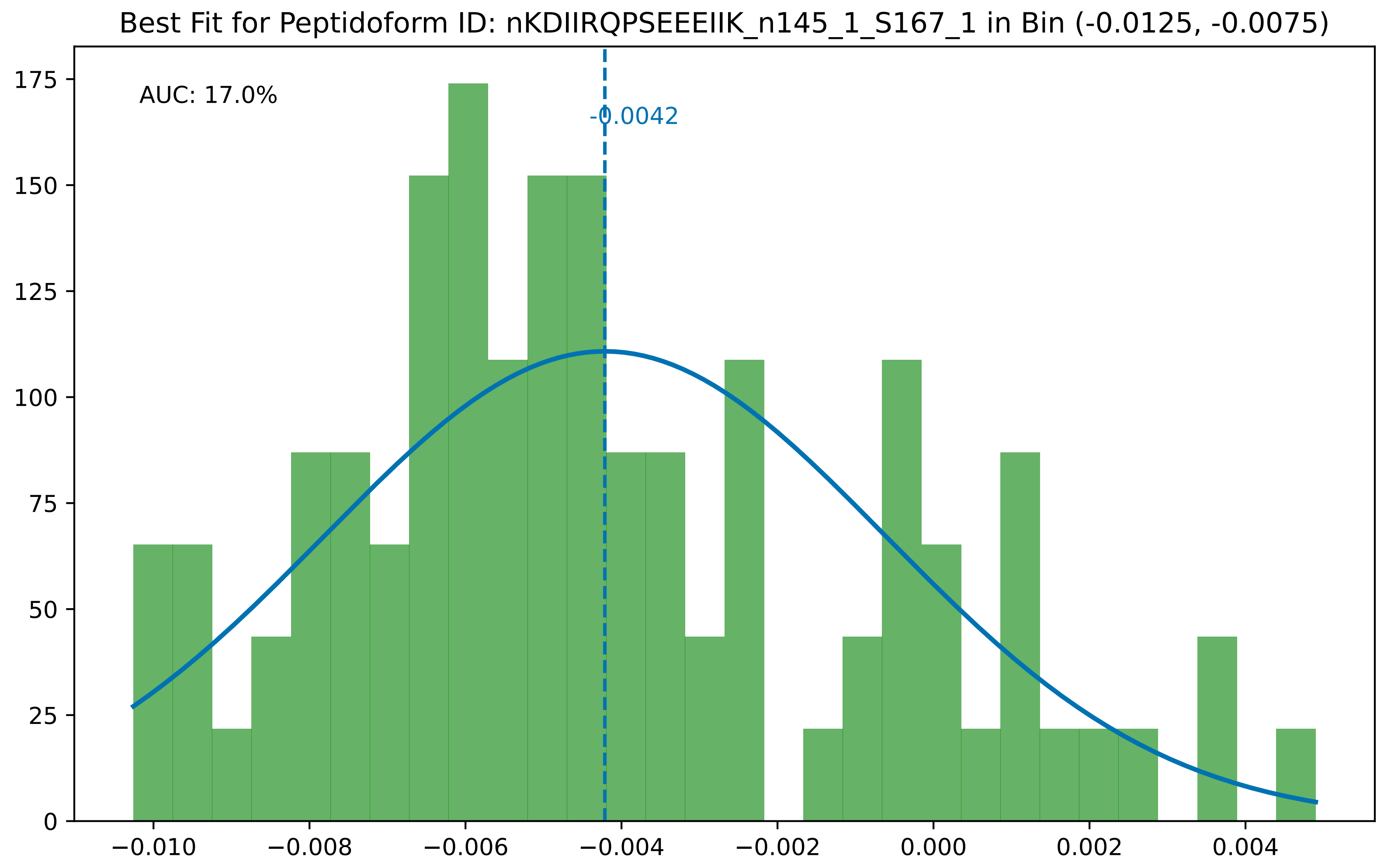
Best Fit for Peptidoform ID: nKDIIRQPSEEEIIK_n145_1_S167_1 in Bin (-0.0125, -0.0075)
AUC: 17.0%
-0.0042
0
25
50
75
100
125
150
175
−0.010
−0.008
−0.006
−0.004
−0.002
0.000
0.002
0.004

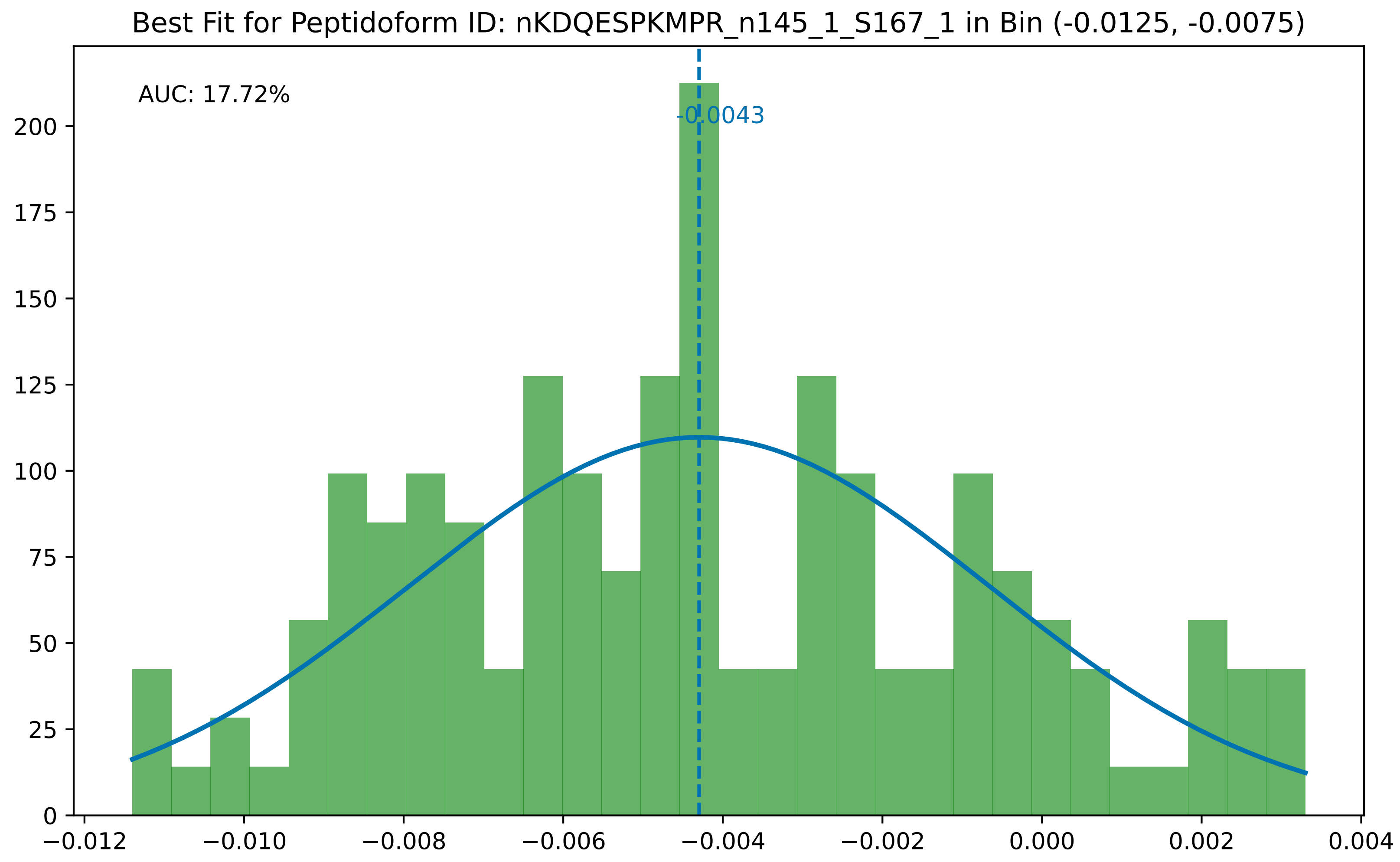

Best Fit for Peptidoform ID: nKDQESPKMPR_n145_1_S167_1 in Bin (-0.0125, -0.0075)
AUC: 17.72%
-0.0043
0
25
50
75
100
125
150
175
200
−0.012
−0.010
−0.008
−0.006
−0.004
−0.002
0.000
0.002
0.004

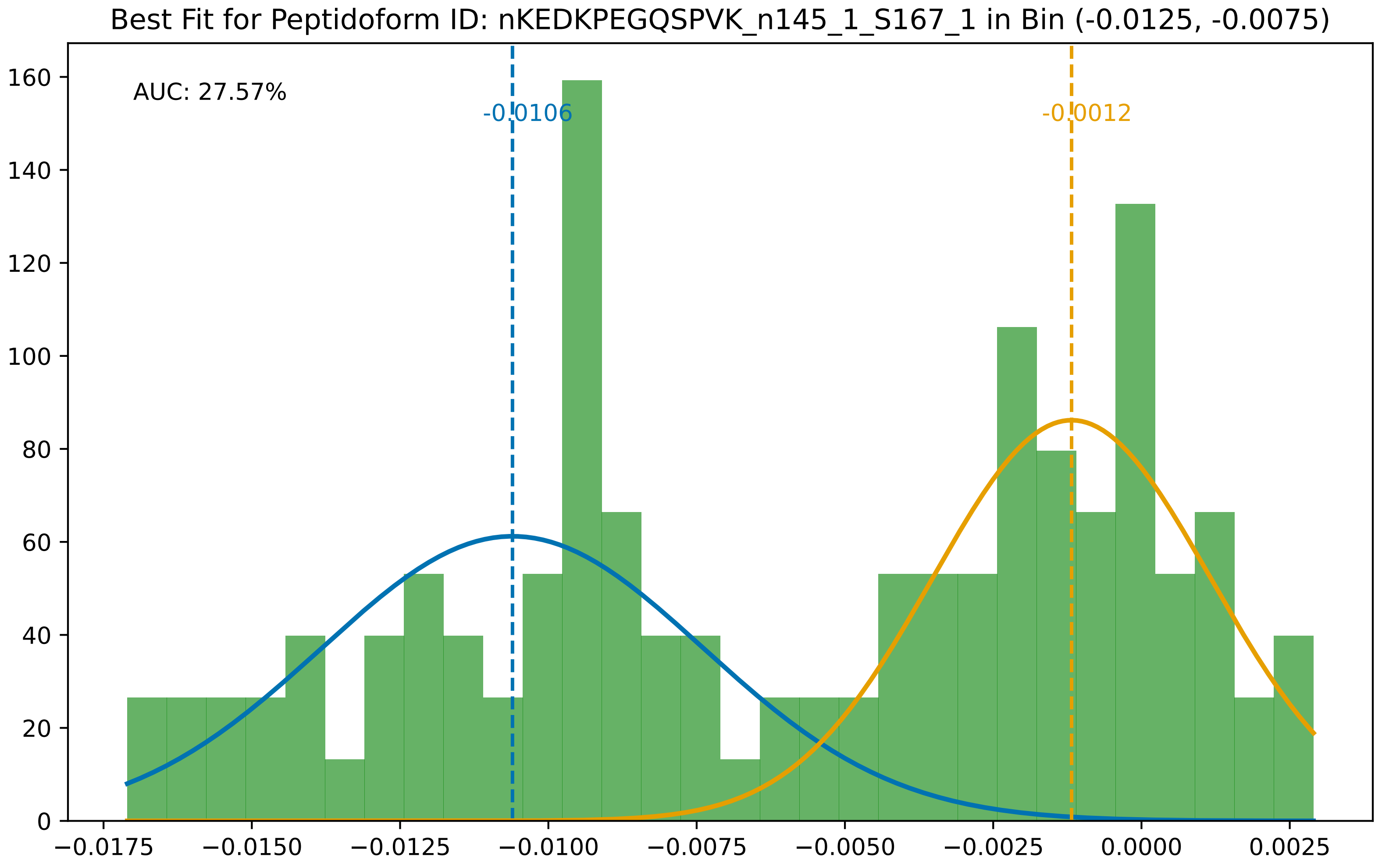

Best Fit for Peptidoform ID: nKEDKPEGQSPVK_n145_1_S167_1 in Bin (-0.0125, -0.0075)
AUC: 27.57%
-0.0106
-0.0012
0
20
40
60
80
100
120
140
160
−0.0175
−0.0150
−0.0125
−0.0100
−0.0075
−0.0050
−0.0025
0.0000
0.0025

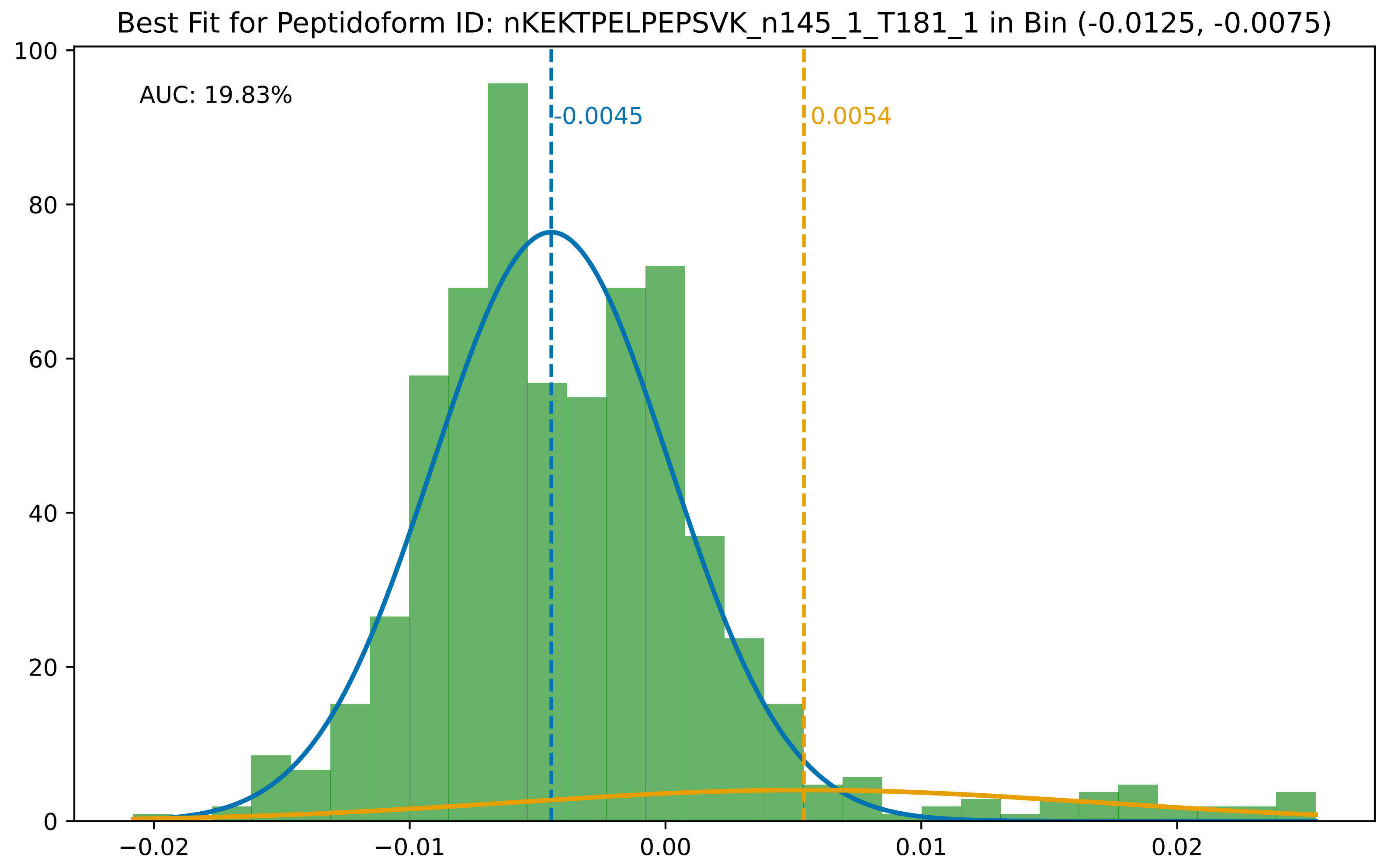

Best Fit for Peptidoform ID: nKEKTPELPEPSVK_n145_1_T181_1 in Bin (-0.0125, -0.0075)
AUC: 19.83%
-0.0045
0.0054
100
80
60
40
20
0
−0.02
−0.01
0.00
0.01
0.02

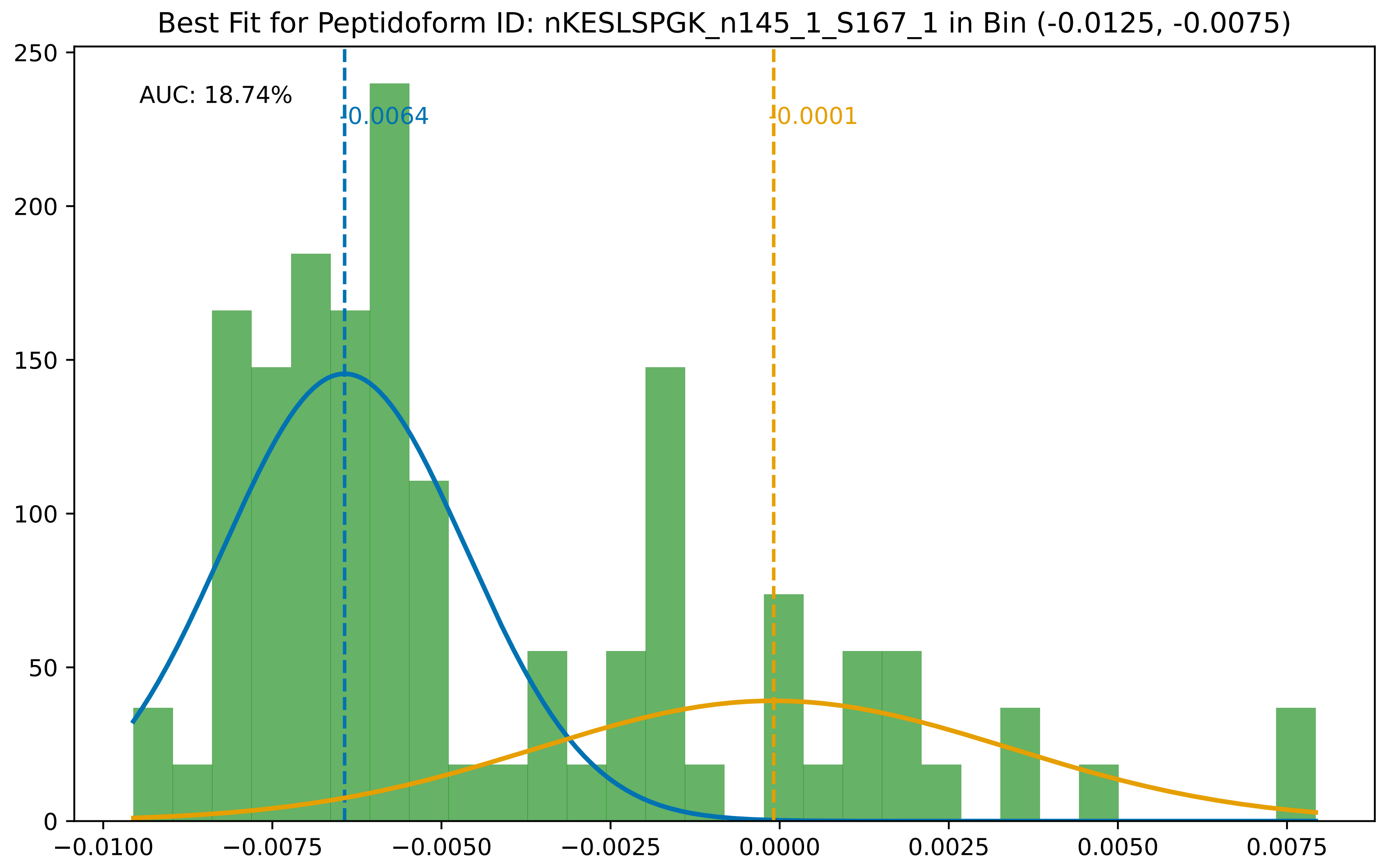

Best Fit for Peptidoform ID: nKESLSPGK_n145_1_S167_1 in Bin (-0.0125, -0.0075)
AUC: 18.74%
0.0064
0.0001
250
200
150
100
50
0
−0.0100
−0.0075
−0.0050
−0.0025
0.0000
0.0025
0.0050
0.0075

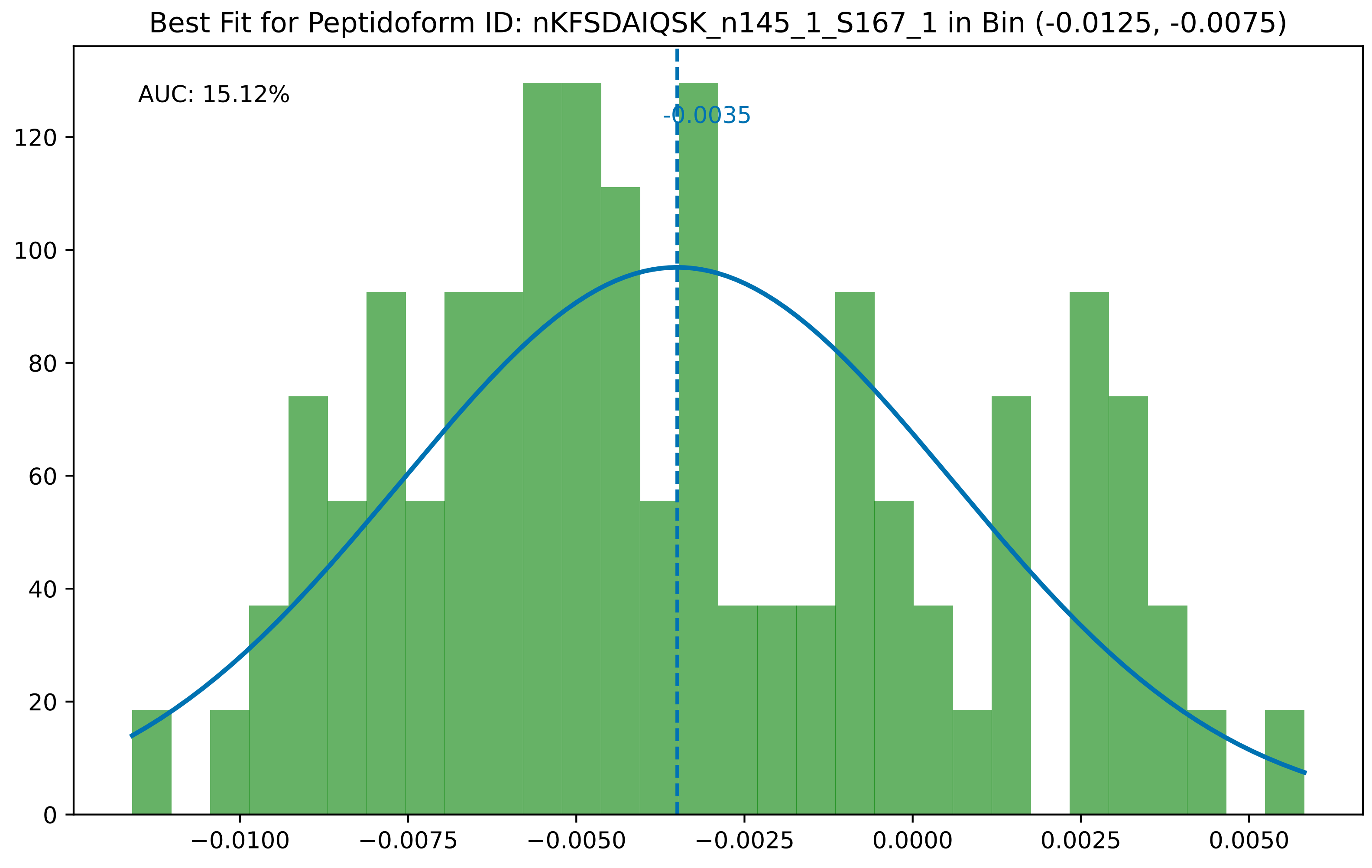

Best Fit for Peptidoform ID: nKFSDAIQSK_n145_1_S167_1 in Bin (-0.0125, -0.0075)
AUC: 15.12%
-0.0035
0
20
40
60
80
100
120
−0.0100
−0.0075
−0.0050
−0.0025
0.0000
0.0025
0.0050

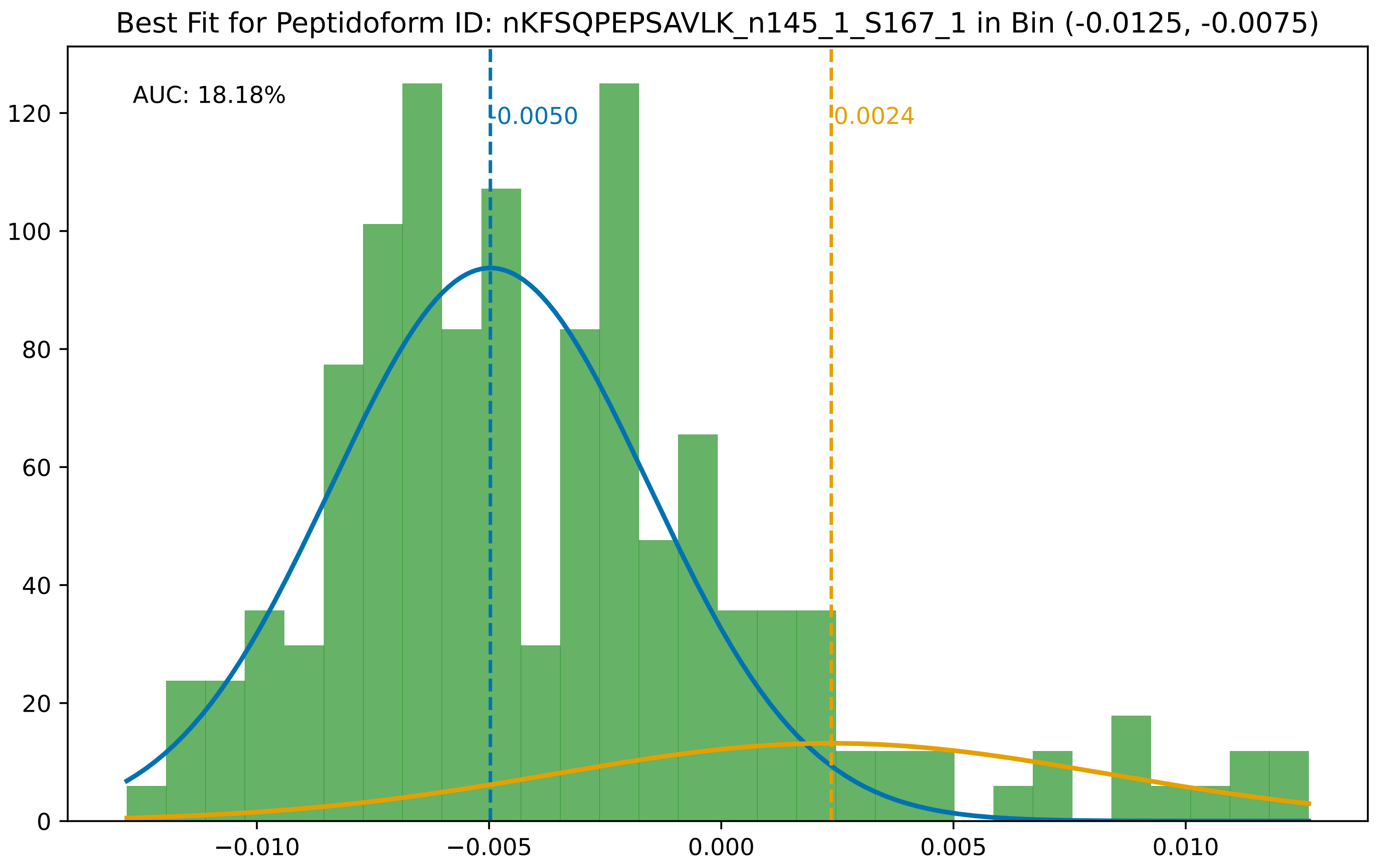

Best Fit for Peptidoform ID: nKFSQPEPSAVLK_n145_1_S167_1 in Bin (-0.0125, -0.0075)
AUC: 18.18%
-0.0050
0.0024
0
20
40
60
80
100
120
−0.010
−0.005
0.000
0.005
0.010

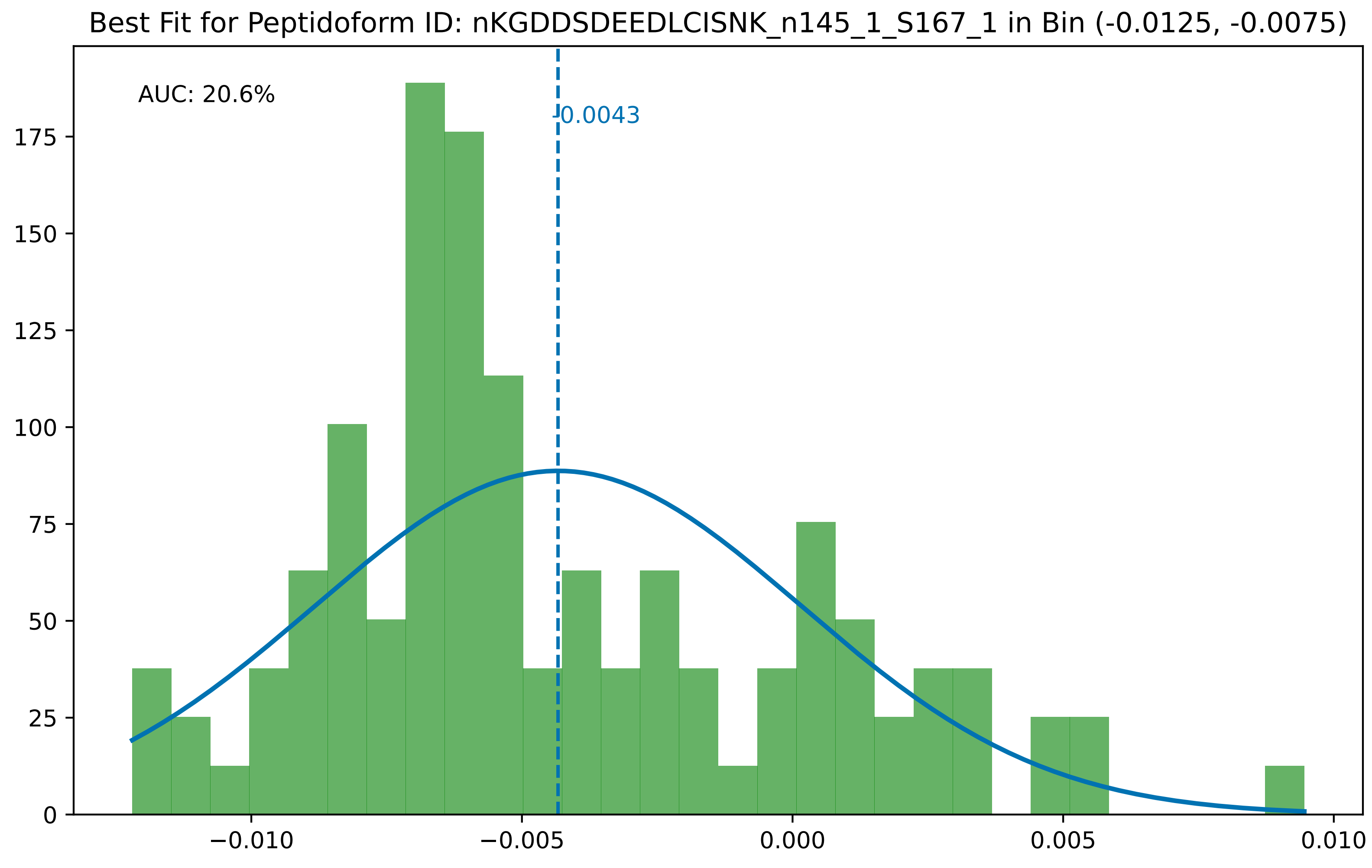

Best Fit for Peptidoform ID: nKGDDSDEEDLCISNK_n145_1_S167_1 in Bin (-0.0125, -0.0075)
AUC: 20.6%
-0.0043
0
25
50
75
100
125
150
175
−0.010
−0.005
0.000
0.005
0.010

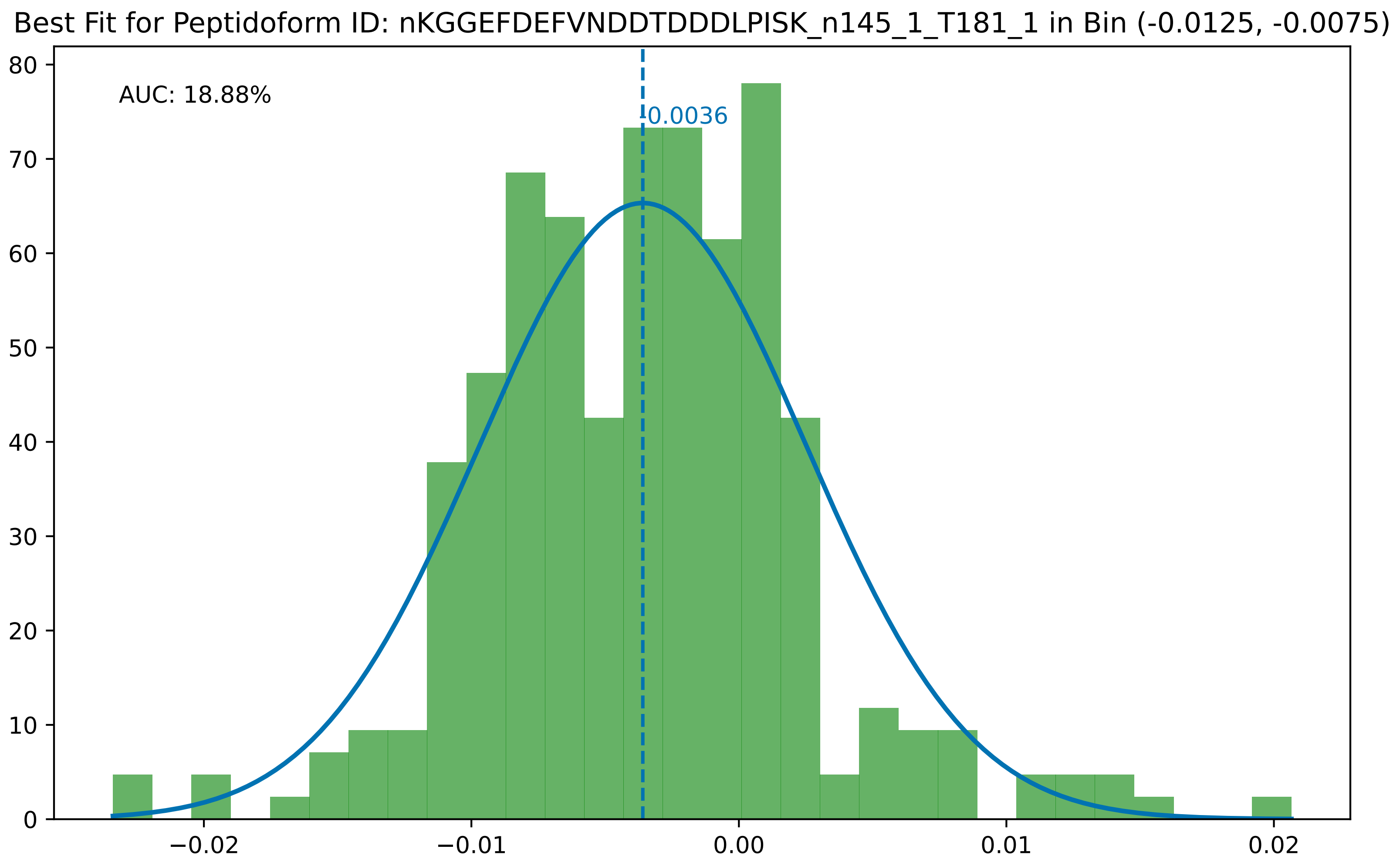

Best Fit for Peptidoform ID: nKGGEFDEFVNDDTDDDLPISK_n145_1_T181_1 in Bin (-0.0125, -0.0075)
AUC: 18.88%
-0.0036
0
10
20
30
40
50
60
70
80
−0.02
−0.01
0.00
0.01
0.02

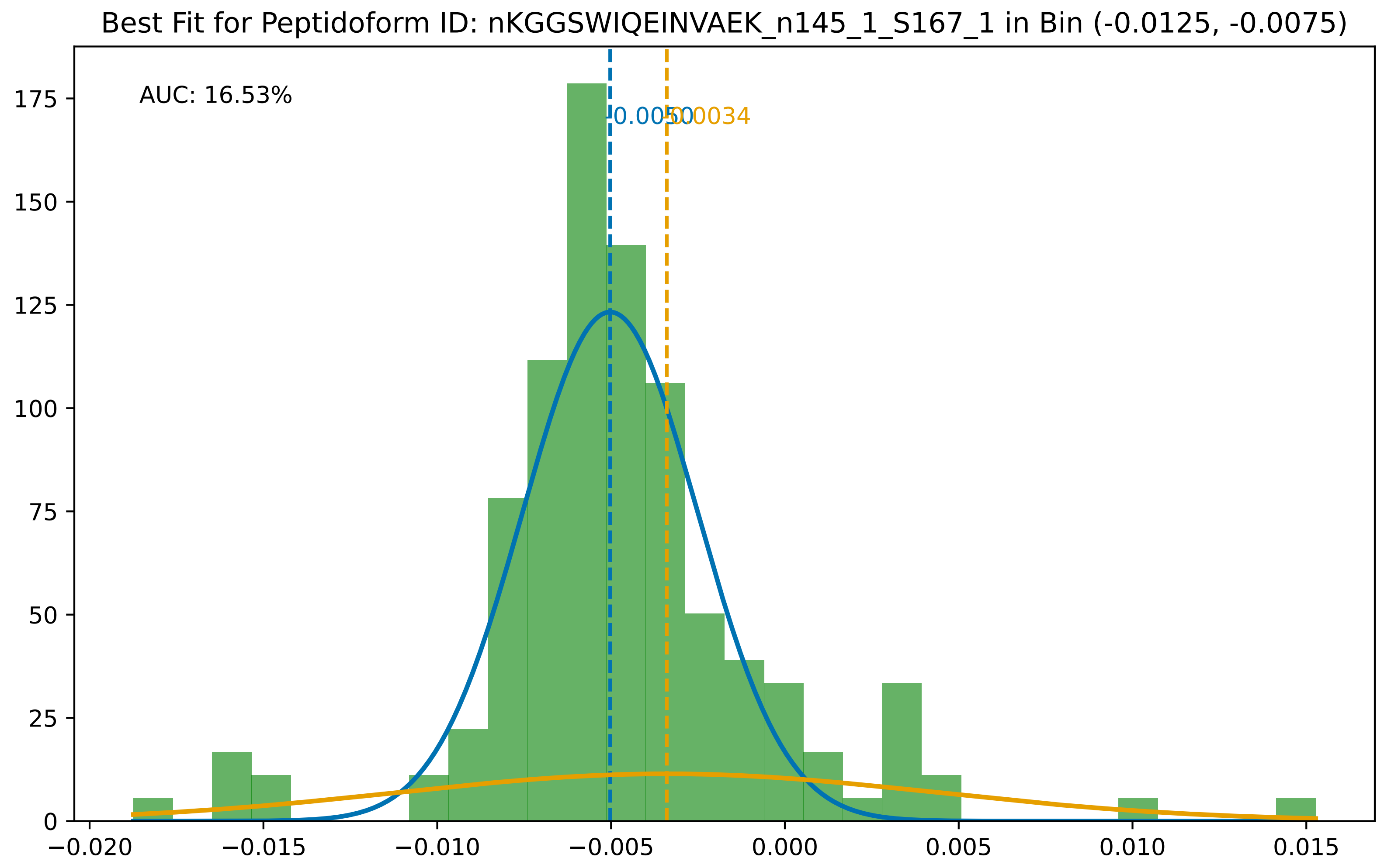

Best Fit for Peptidoform ID: nKGGSWIQEINVAEK_n145_1_S167_1 in Bin (-0.0125, -0.0075)
AUC: 16.53%
-0.0050
-0.0034
0
25
50
75
100
125
150
175
−0.020
−0.015
−0.010
−0.005
0.000
0.005
0.010
0.015

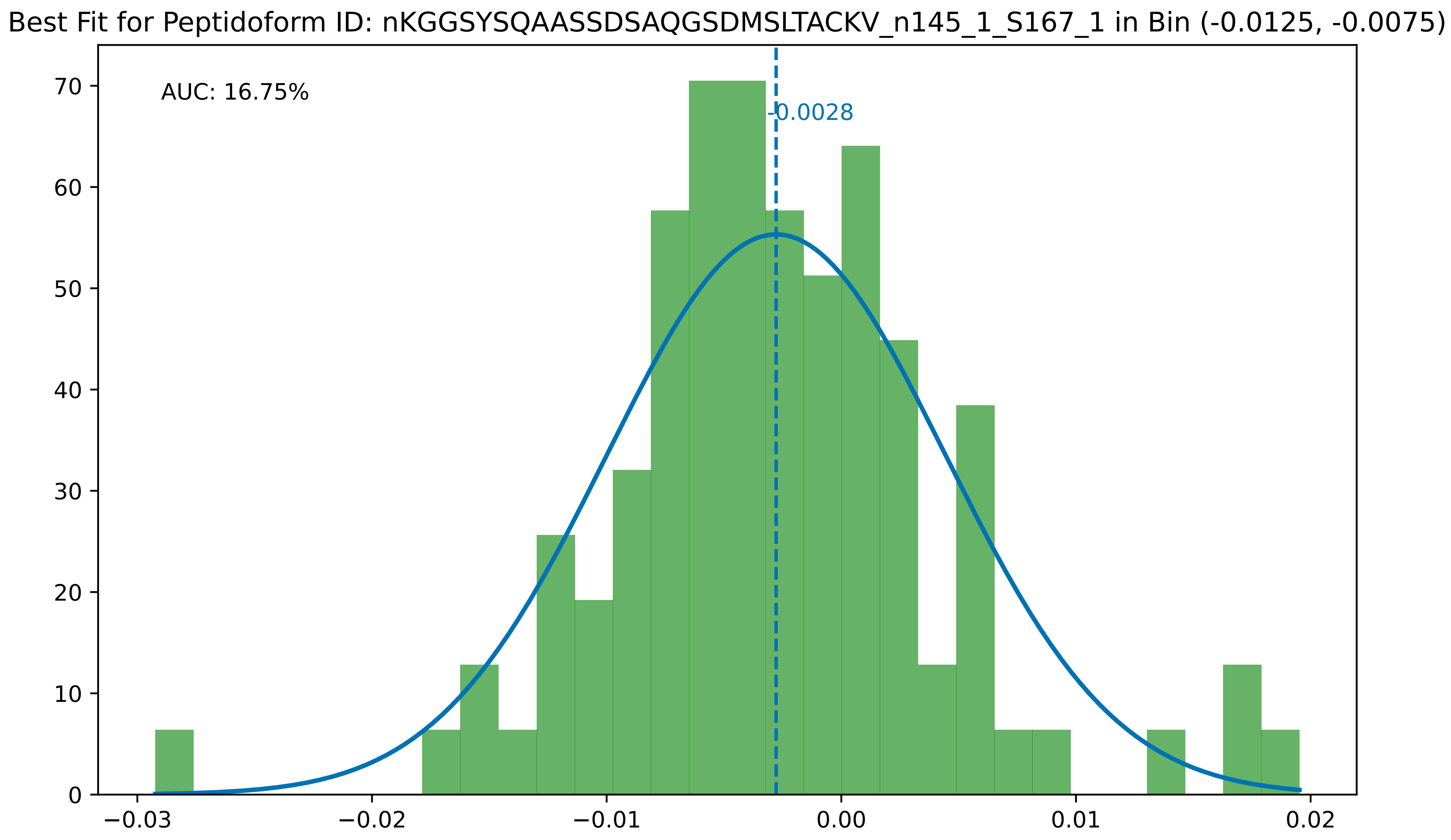

Best Fit for Peptidoform ID: nKGGSYSQAASSDSAQGSDMSLTACKV_n145_1_S167_1 in Bin (-0.0125, -0.0075)
AUC: 16.75%
-0.0028
70
60
50
40
30
20
10
0
−0.03
−0.02
−0.01
0.00
0.01
0.02

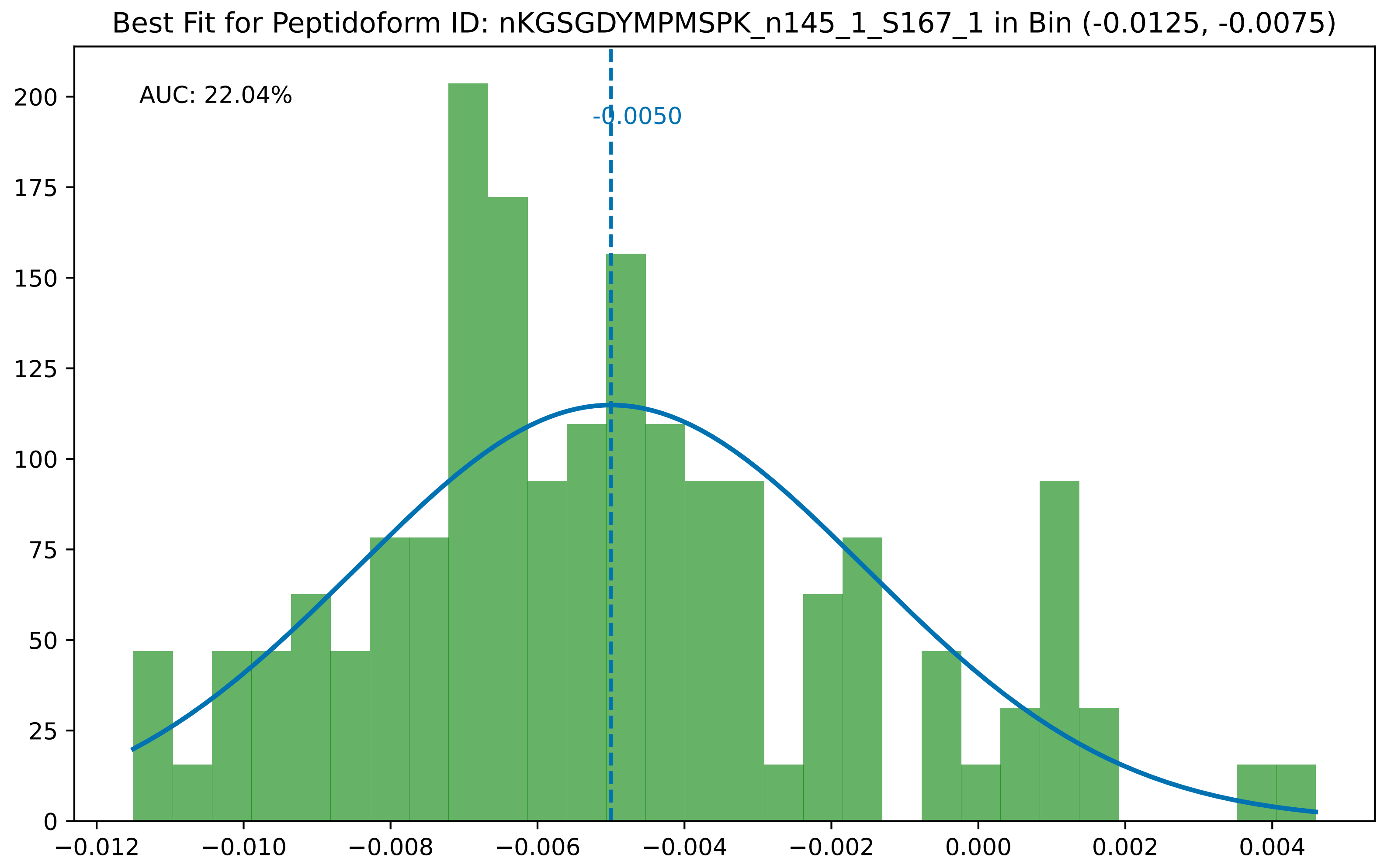

Best Fit for Peptidoform ID: nKGSGDYMPMSPK_n145_1_S167_1 in Bin (-0.0125, -0.0075)
AUC: 22.04%
-0.0050
0
25
50
75
100
125
150
175
200
−0.012
−0.010
−0.008
−0.006
−0.004
−0.002
0.000
0.002
0.004

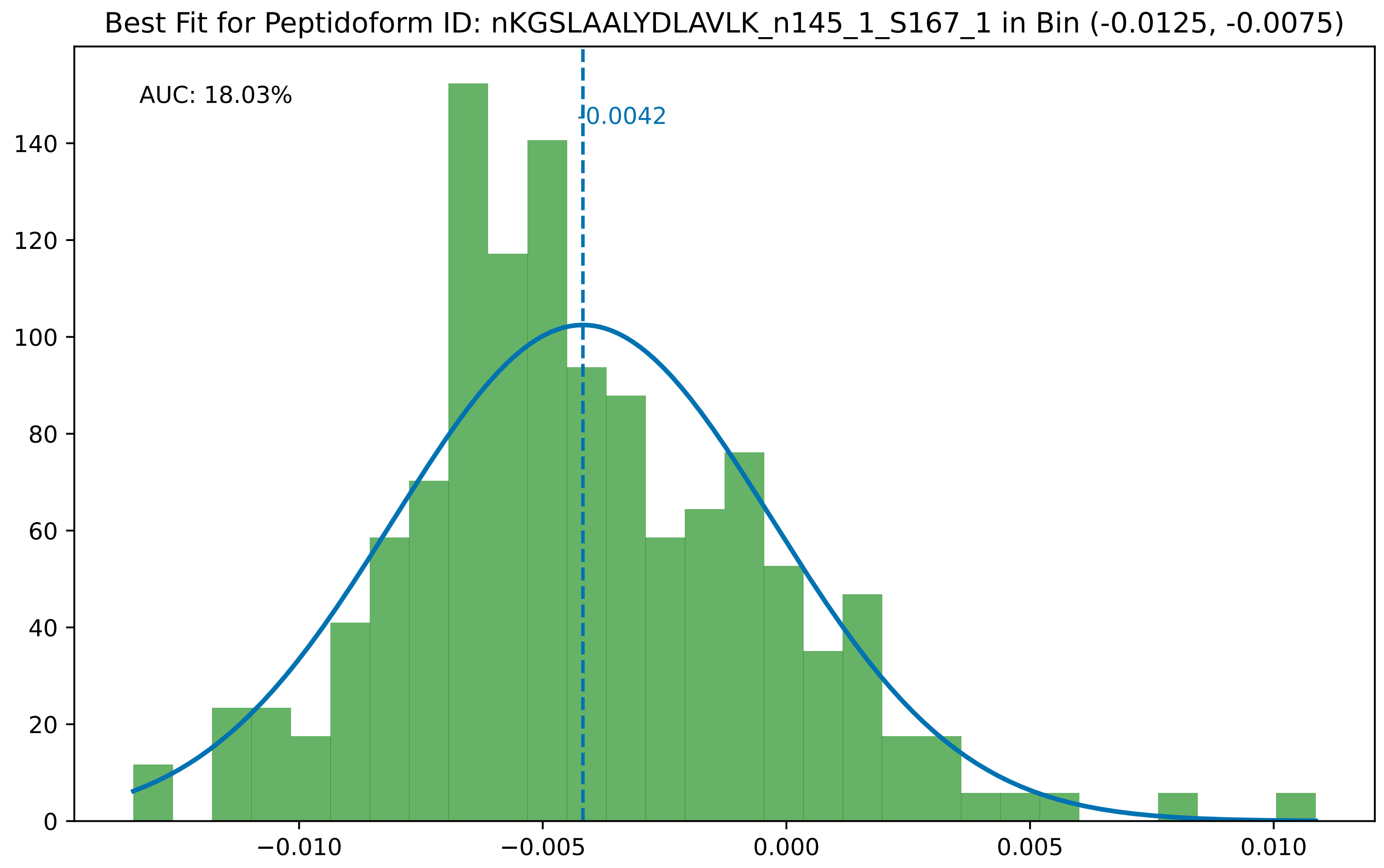

Best Fit for Peptidoform ID: nKGSLAALYDLAVLK_n145_1_S167_1 in Bin (-0.0125, -0.0075)
AUC: 18.03%
-0.0042
0
20
40
60
80
100
120
140
−0.010
−0.005
0.000
0.005
0.010

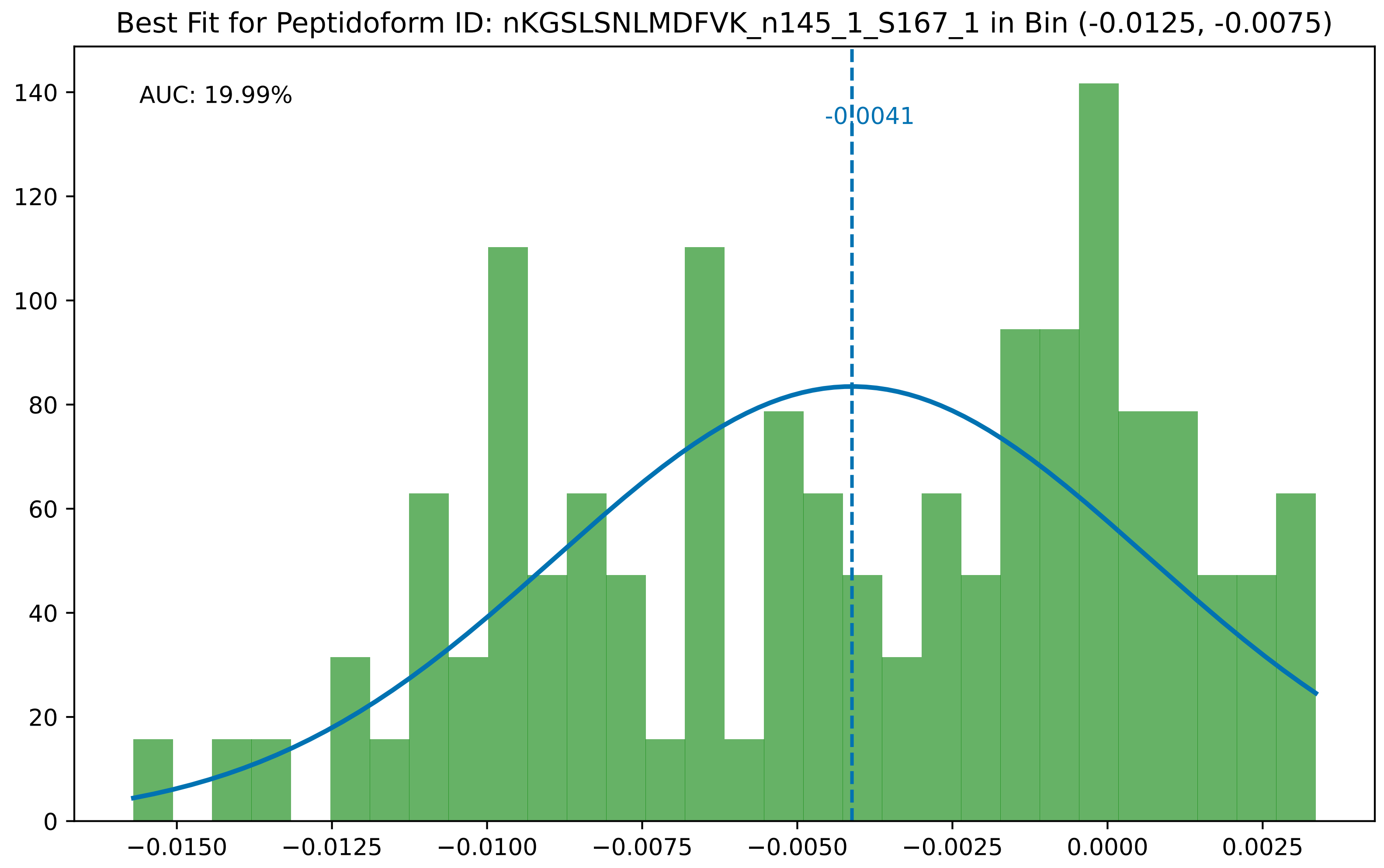

Best Fit for Peptidoform ID: nKGSLSNLMDFVK_n145_1_S167_1 in Bin (-0.0125, -0.0075)
AUC: 19.99%
-0.0041
0
20
40
60
80
100
120
140
−0.0150
−0.0125
−0.0100
−0.0075
−0.0050
−0.0025
0.0000
0.0025

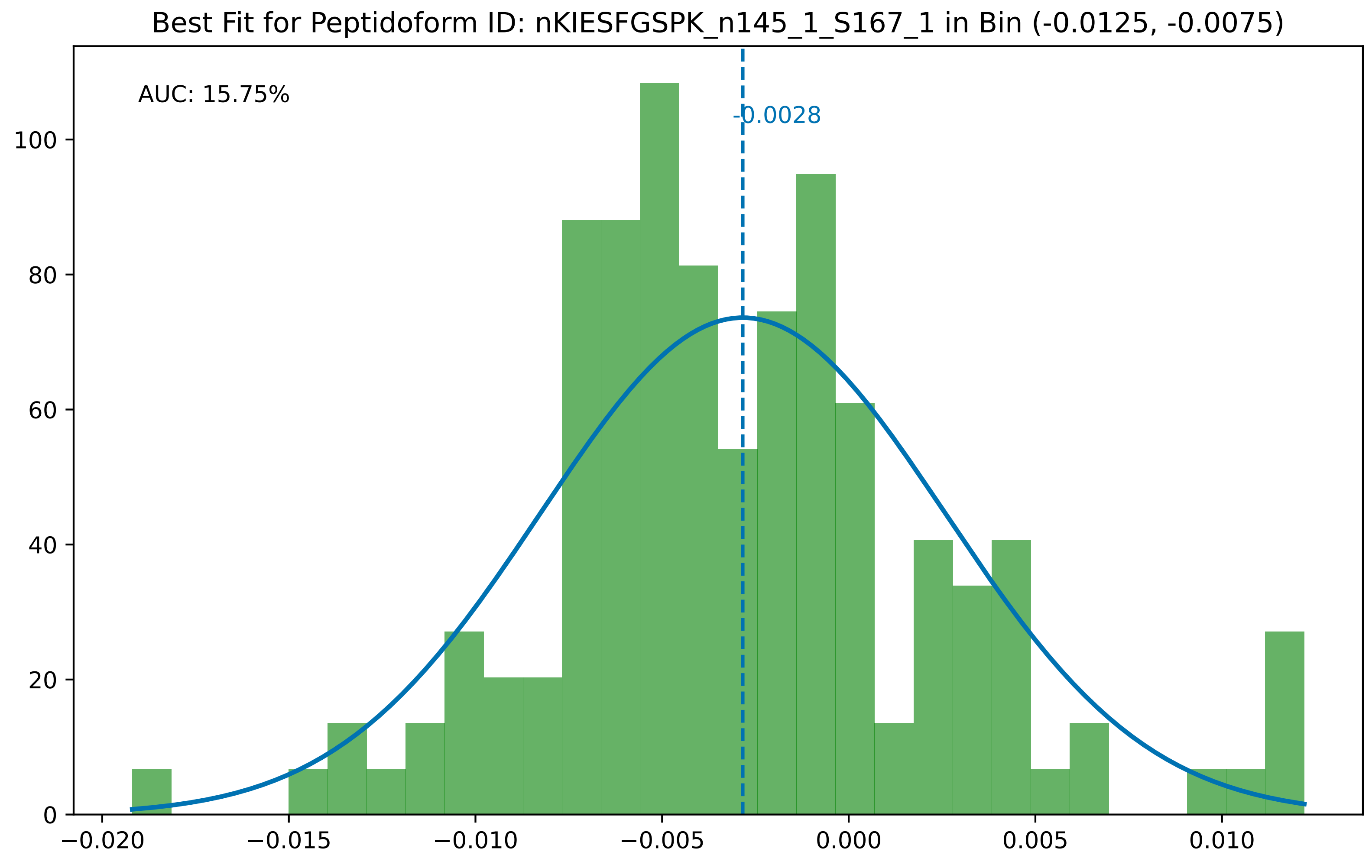

Best Fit for Peptidoform ID: nKIESFGSPK_n145_1_S167_1 in Bin (-0.0125, -0.0075)
AUC: 15.75%
-0.0028
0
20
40
60
80
100
−0.020
−0.015
−0.010
−0.005
0.000
0.005
0.010

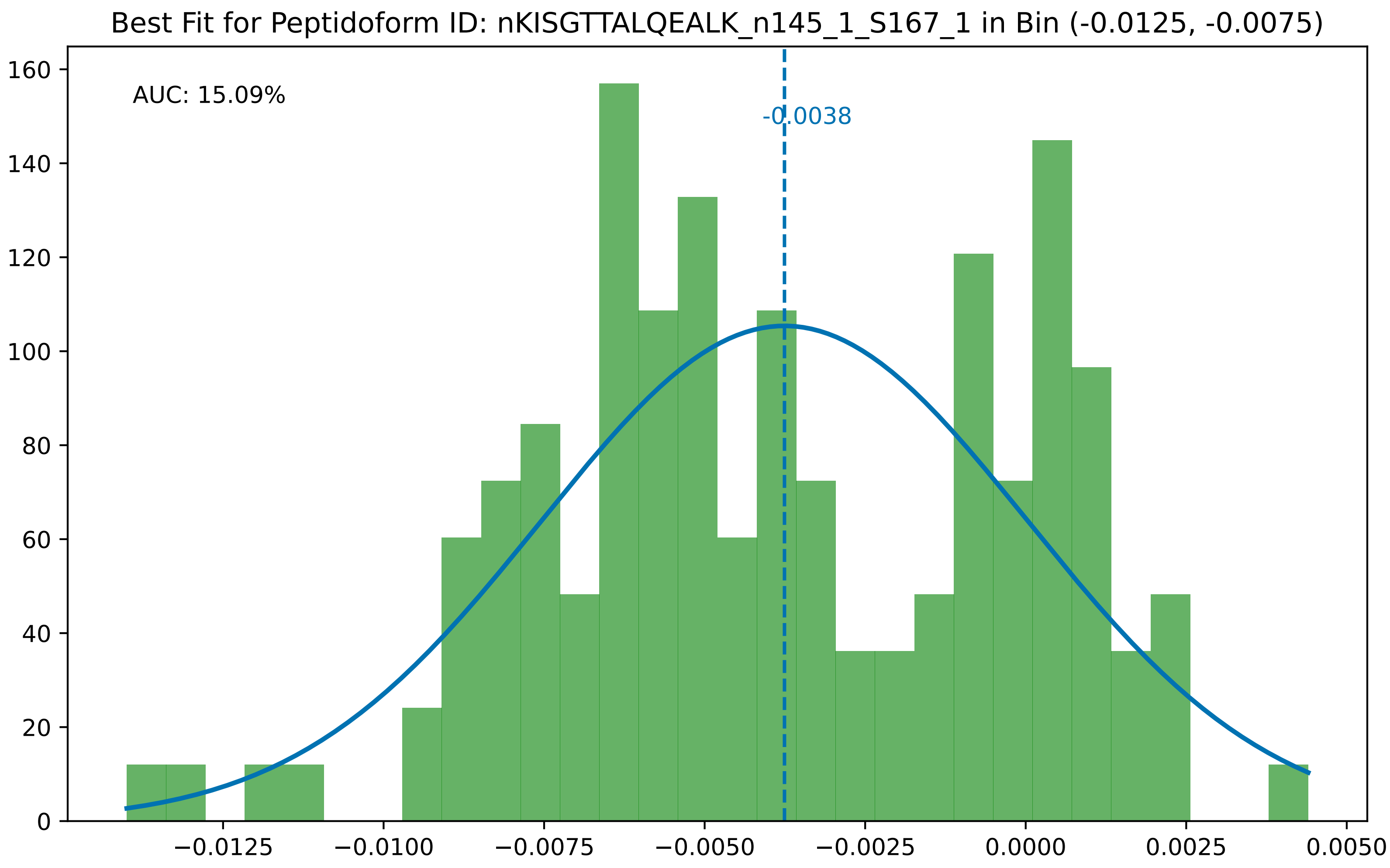

Best Fit for Peptidoform ID: nKISGTTALQEALK_n145_1_S167_1 in Bin (-0.0125, -0.0075)
AUC: 15.09%
-0.0038
160
140
120
100
80
60
40
20
0
−0.0125
−0.0100
−0.0075
−0.0050
−0.0025
0.0000
0.0025
0.0050

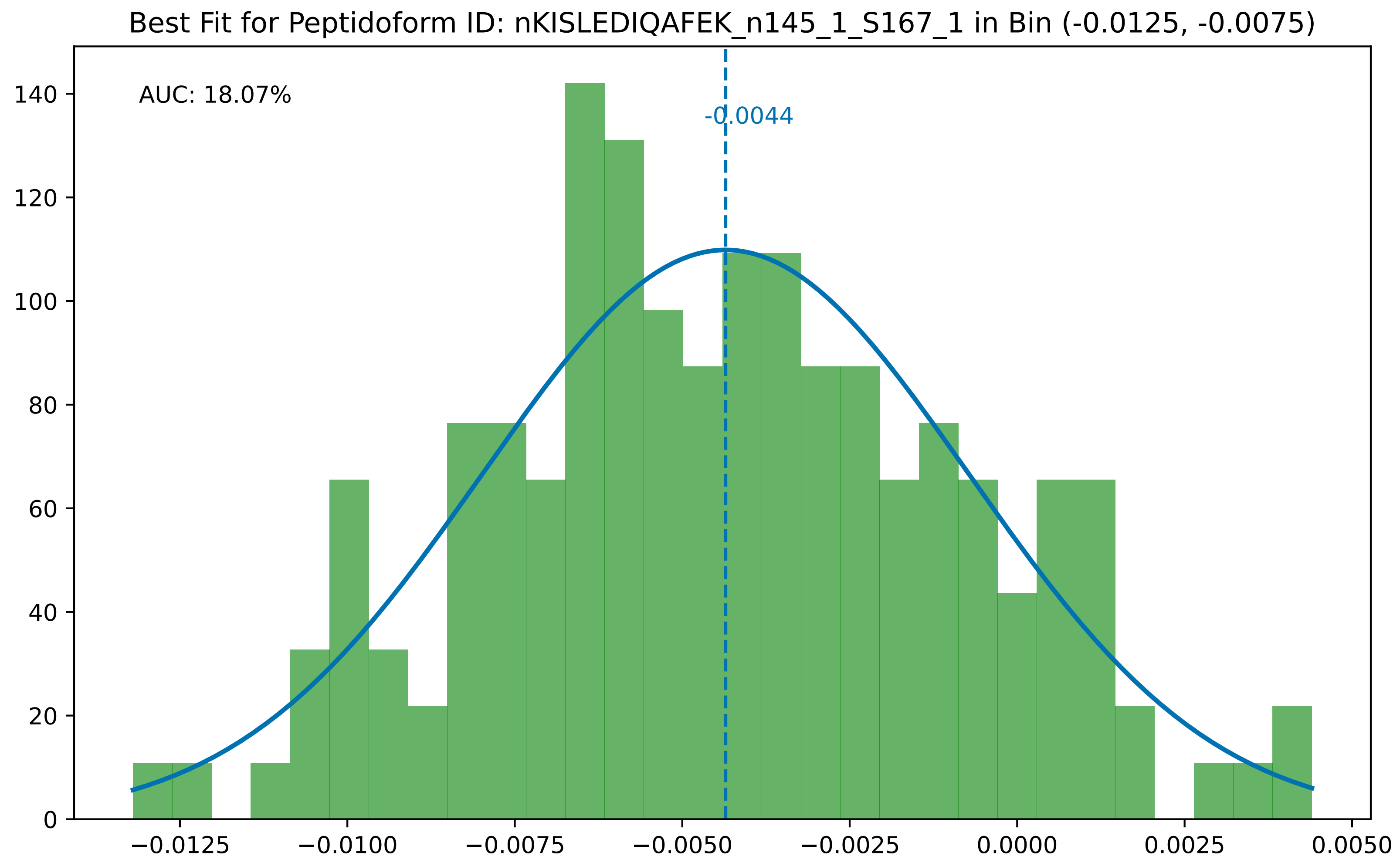

Best Fit for Peptidoform ID: nKISLEDIQAFEK_n145_1_S167_1 in Bin (-0.0125, -0.0075)
AUC: 18.07%
-0.0044
0
20
40
60
80
100
120
140
−0.0125
−0.0100
−0.0075
−0.0050
−0.0025
0.0000
0.0025
0.0050

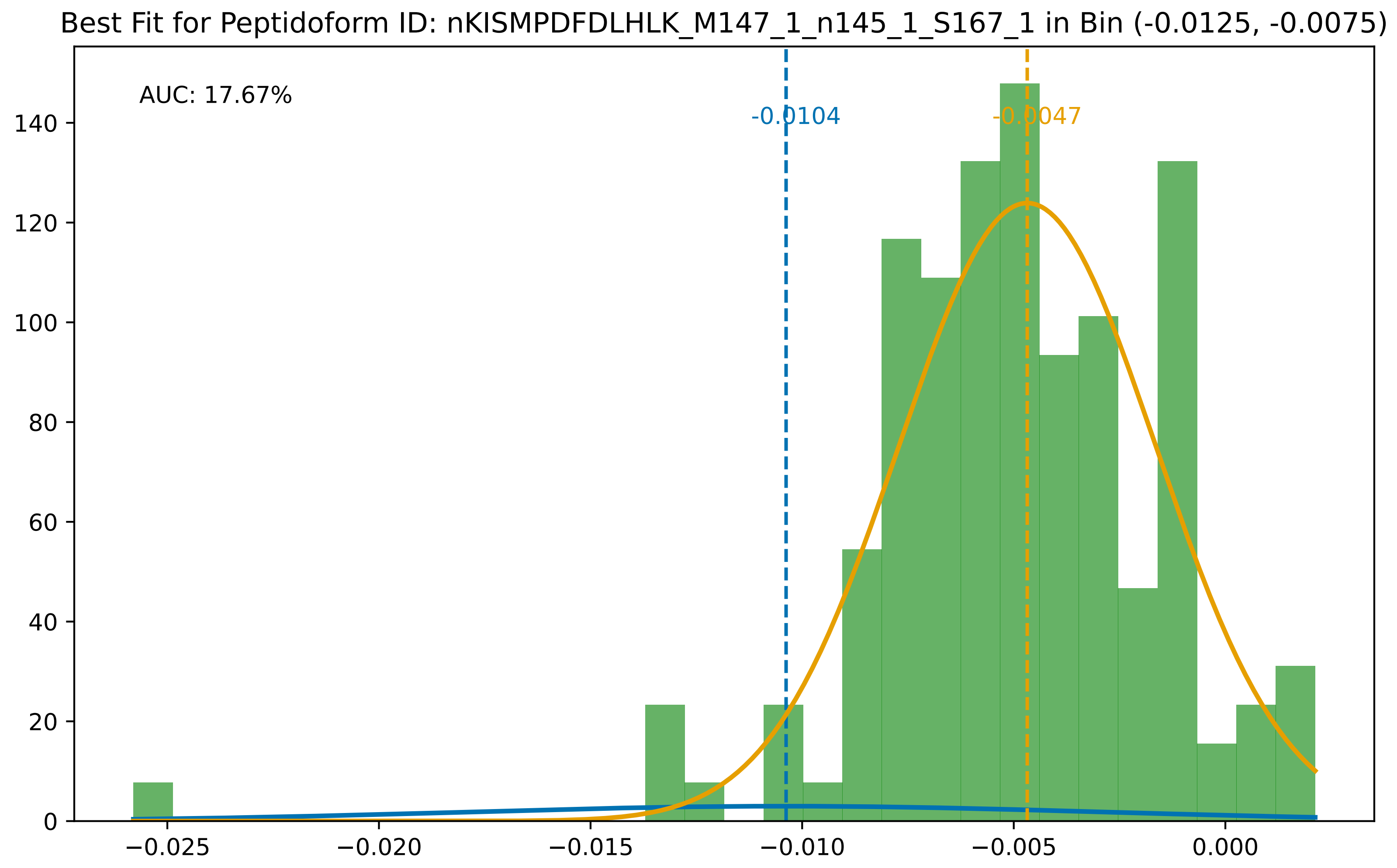

Best Fit for Peptidoform ID: nKISMPDFDLHLK_M147_1_n145_1_S167_1 in Bin (-0.0125, -0.0075)
AUC: 17.67%
-0.0104
-0.0047
0
20
40
60
80
100
120
140
−0.025
−0.020
−0.015
−0.010
−0.005
0.000

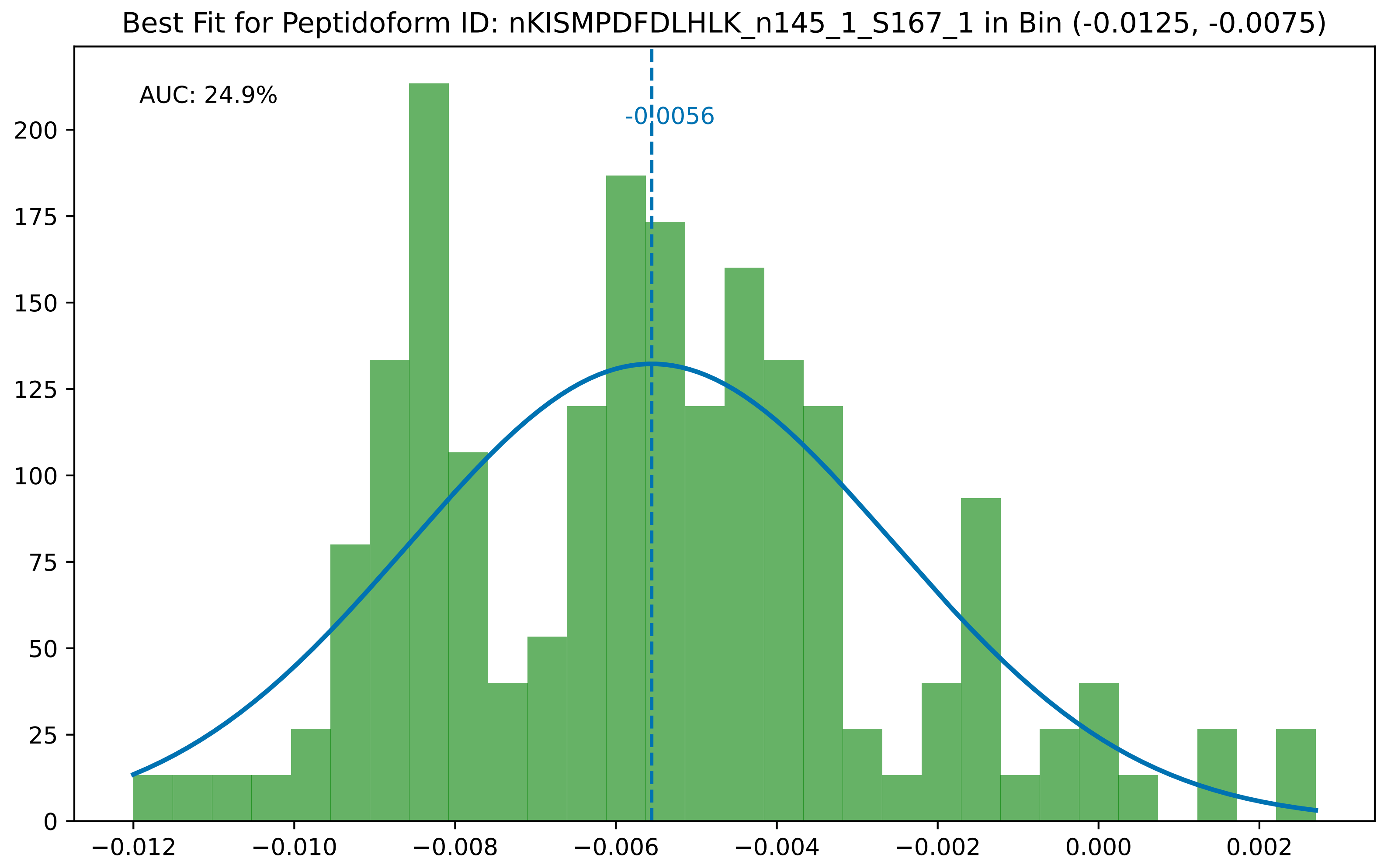

Best Fit for Peptidoform ID: nKISMPDFDLHLK_n145_1_S167_1 in Bin (-0.0125, -0.0075)
AUC: 24.9%
-0.0056
0
25
50
75
100
125
150
175
200
−0.012
−0.010
−0.008
−0.006
−0.004
−0.002
0.000
0.002

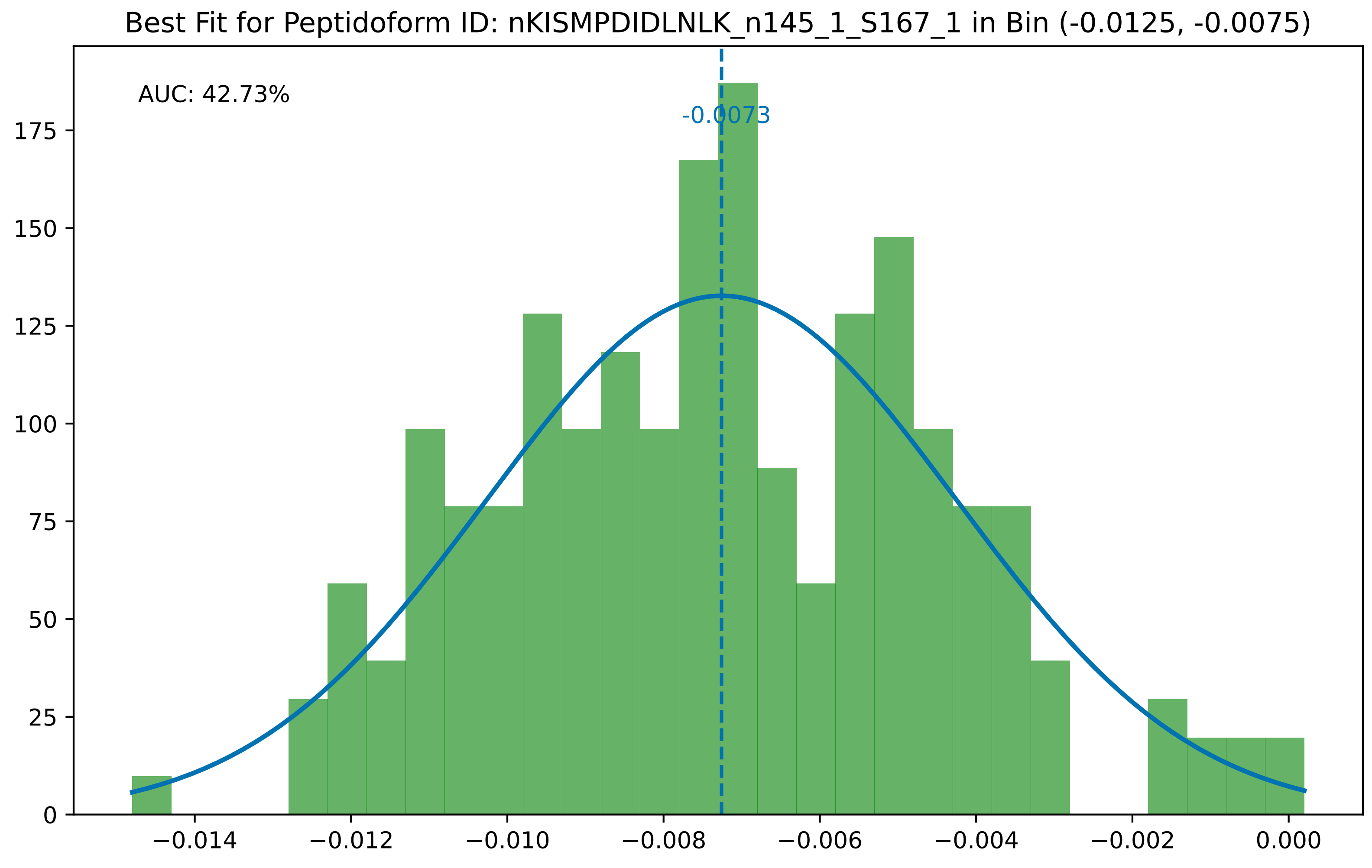
Best Fit for Peptidoform ID: nKISMPDIDLNLK_n145_1_S167_1 in Bin (-0.0125, -0.0075)
AUC: 42.73%
-0.0073
0
25
50
75
100
125
150
175
−0.014
−0.012
−0.010
−0.008
−0.006
−0.004
−0.002
0.000

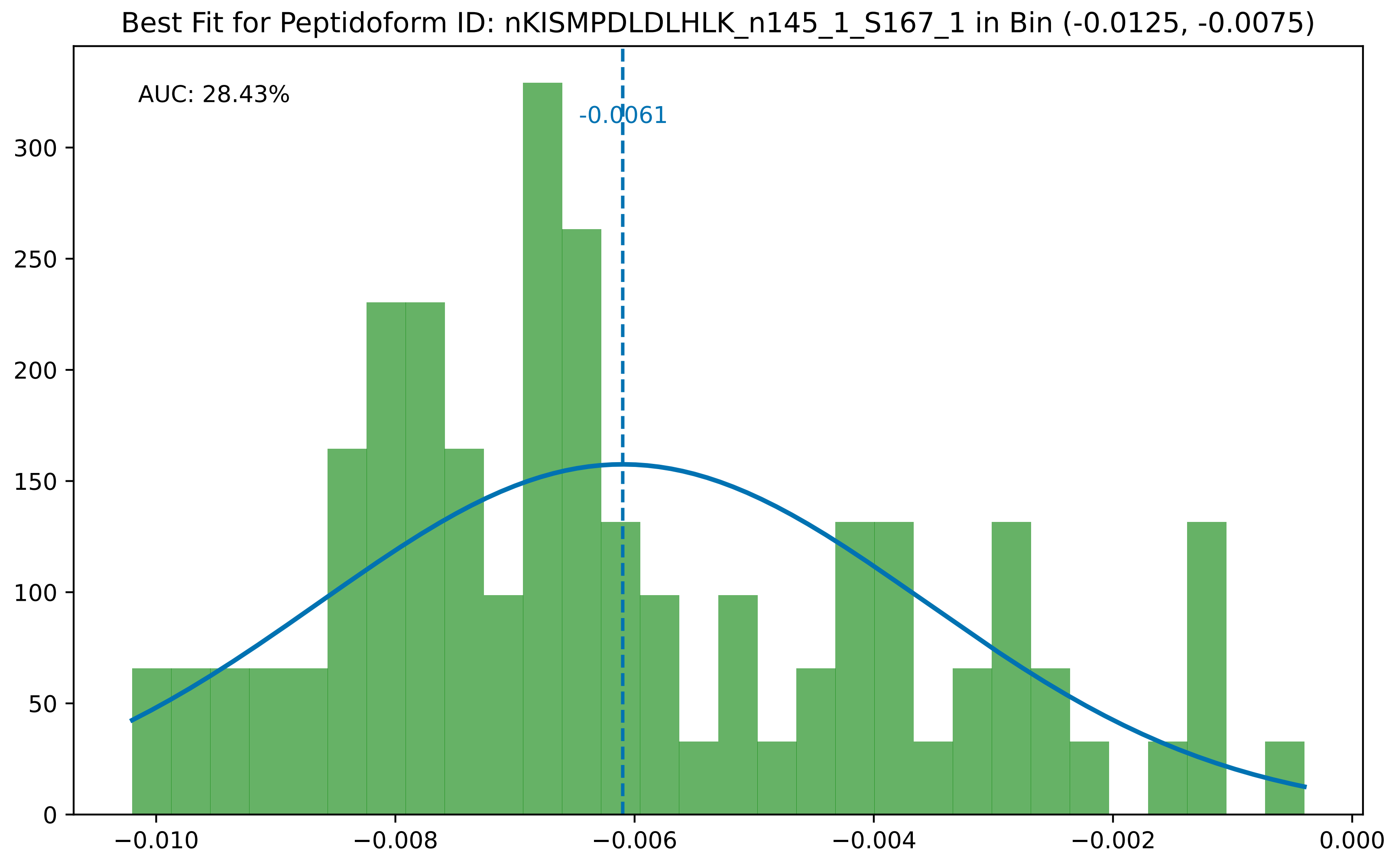

Best Fit for Peptidoform ID: nKISMPDLDLHLK_n145_1_S167_1 in Bin (-0.0125, -0.0075)
AUC: 28.43%
-0.0061
0
50
100
150
200
250
300
−0.010
−0.008
−0.006
−0.004
−0.002
0.000

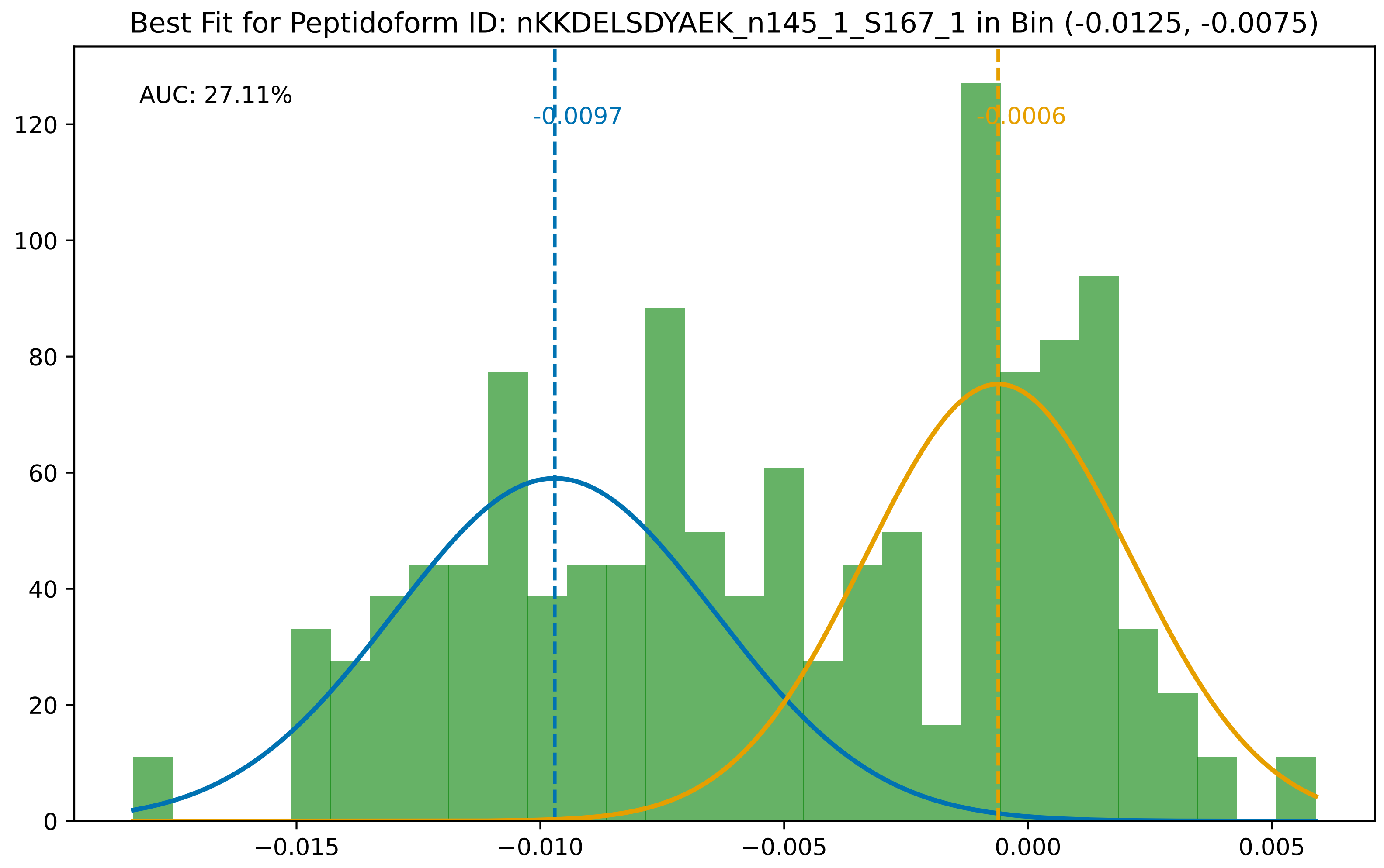
Best Fit for Peptidoform ID: nKKDELSDYAEK_n145_1_S167_1 in Bin (-0.0125, -0.0075)
AUC: 27.11%
-0.0097
-0.0006
0
20
40
60
80
100
120
−0.015
−0.010
−0.005
0.000
0.005

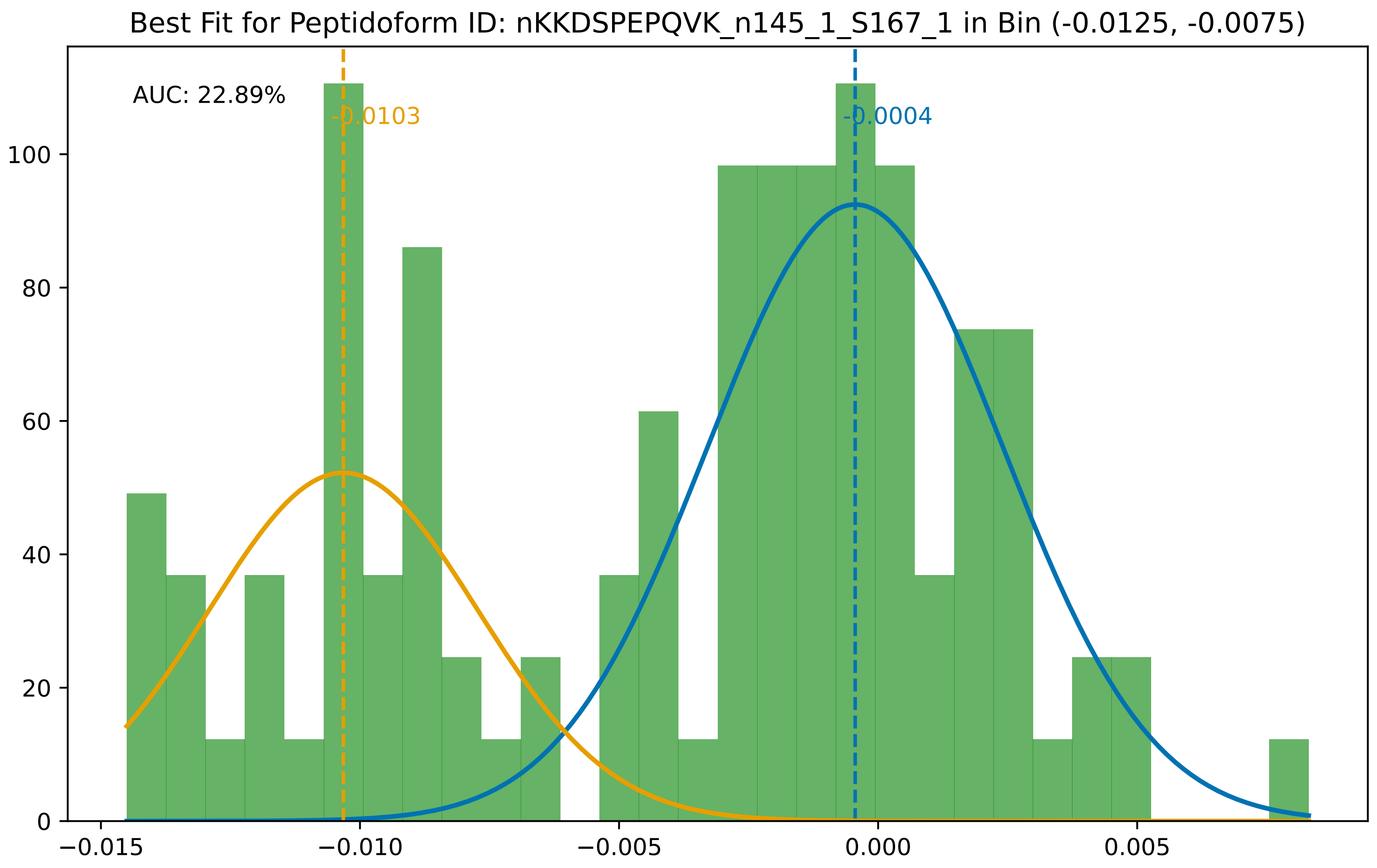
Best Fit for Peptidoform ID: nKKDSPEPQVK_n145_1_S167_1 in Bin (-0.0125, -0.0075)
AUC: 22.89%
-0.0103
-0.0004
0
20
40
60
80
100
−0.015
−0.010
−0.005
0.000
0.005

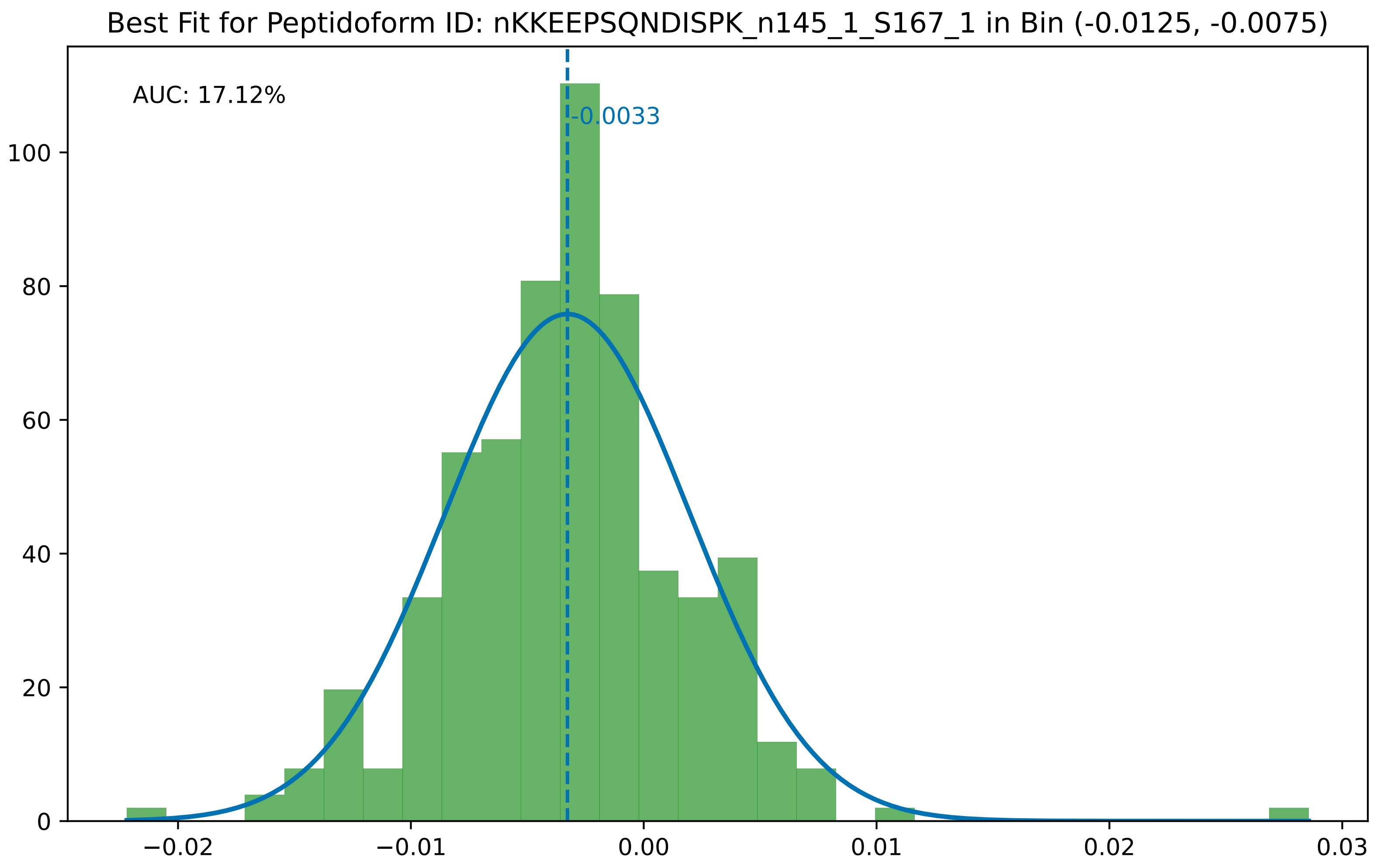
Best Fit for Peptidoform ID: nKKEEPSQNDISPK_n145_1_S167_1 in Bin (-0.0125, -0.0075)
AUC: 17.12%
-0.0033
0
20
40
60
80
100
−0.02
−0.01
0.00
0.01
0.02
0.03

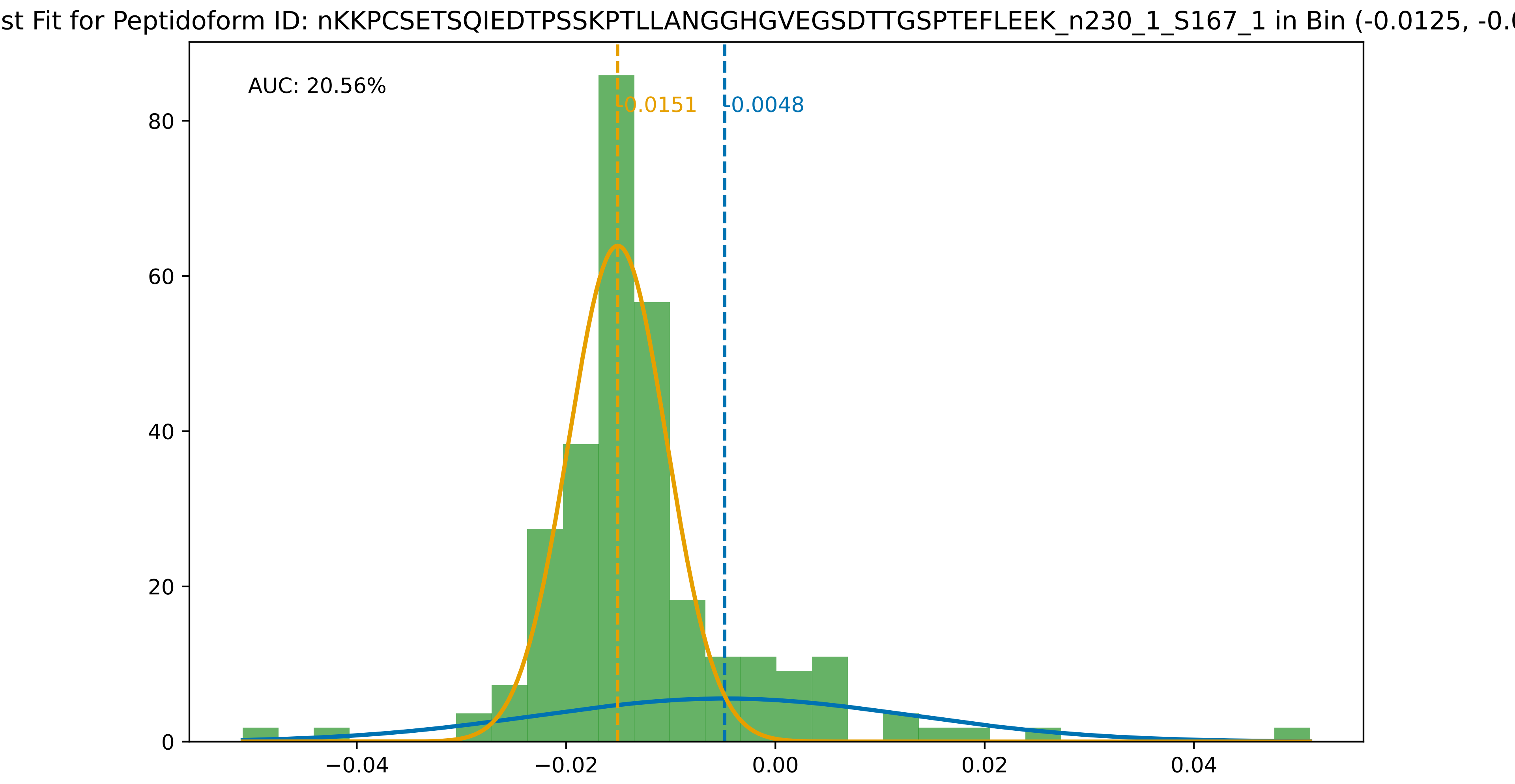

st Fit for Peptidoform ID: nKKPCSETSQIEDTPSSKPTLLANGGHGVEGSDTTGSPTEFLEEK_n230_1_S167_1 in Bin (-0.0125, -0.0
AUC: 20.56%
-0.0151
-0.0048
0
20
40
60
80
−0.04
−0.02
0.00
0.02
0.04

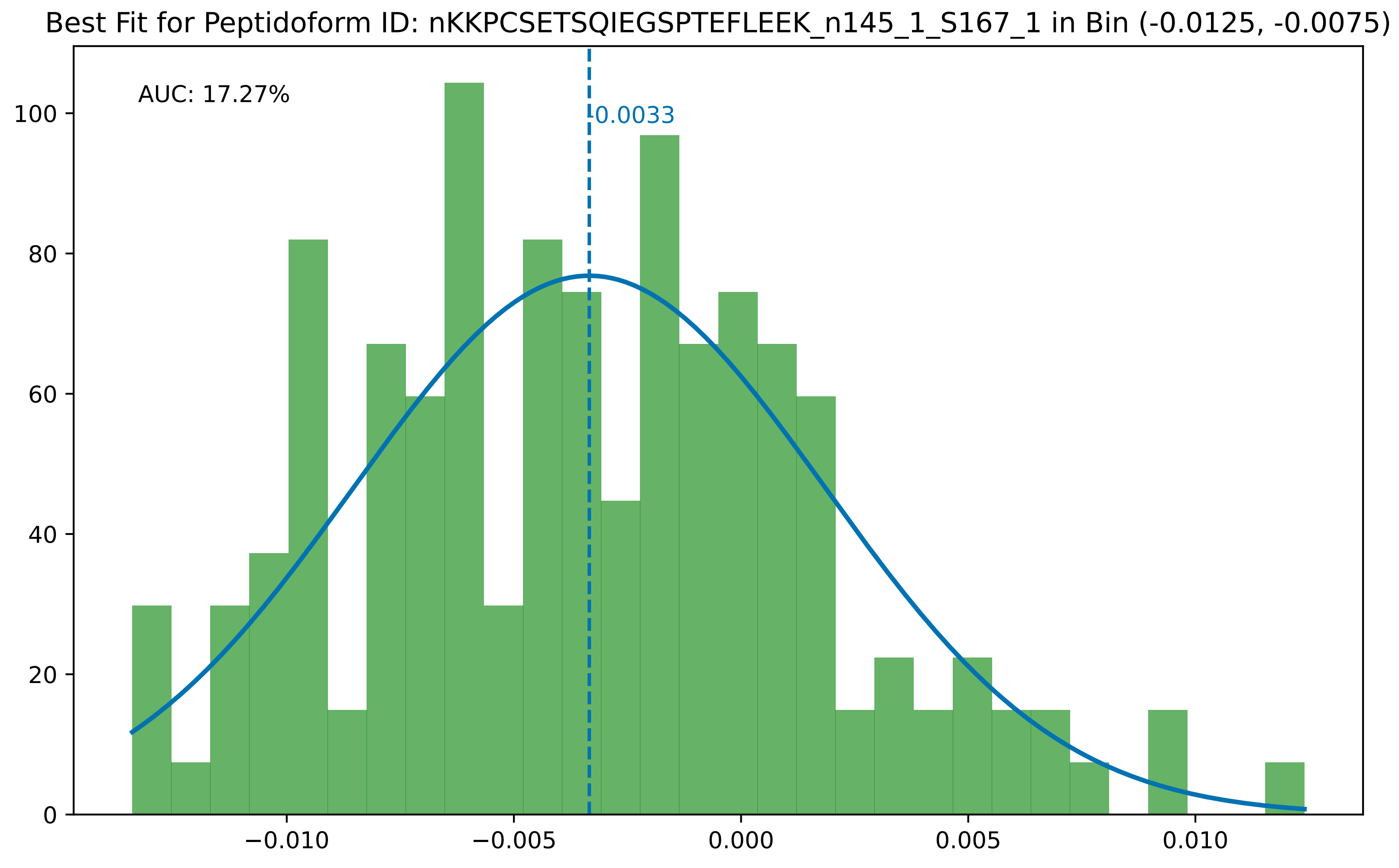
Best Fit for Peptidoform ID: nKKPCSETSQIEGSPTEFLEEK_n145_1_S167_1 in Bin (-0.0125, -0.0075)
AUC: 17.27%
-0.0033
0
20
40
60
80
100
−0.010
−0.005
0.000
0.005
0.010

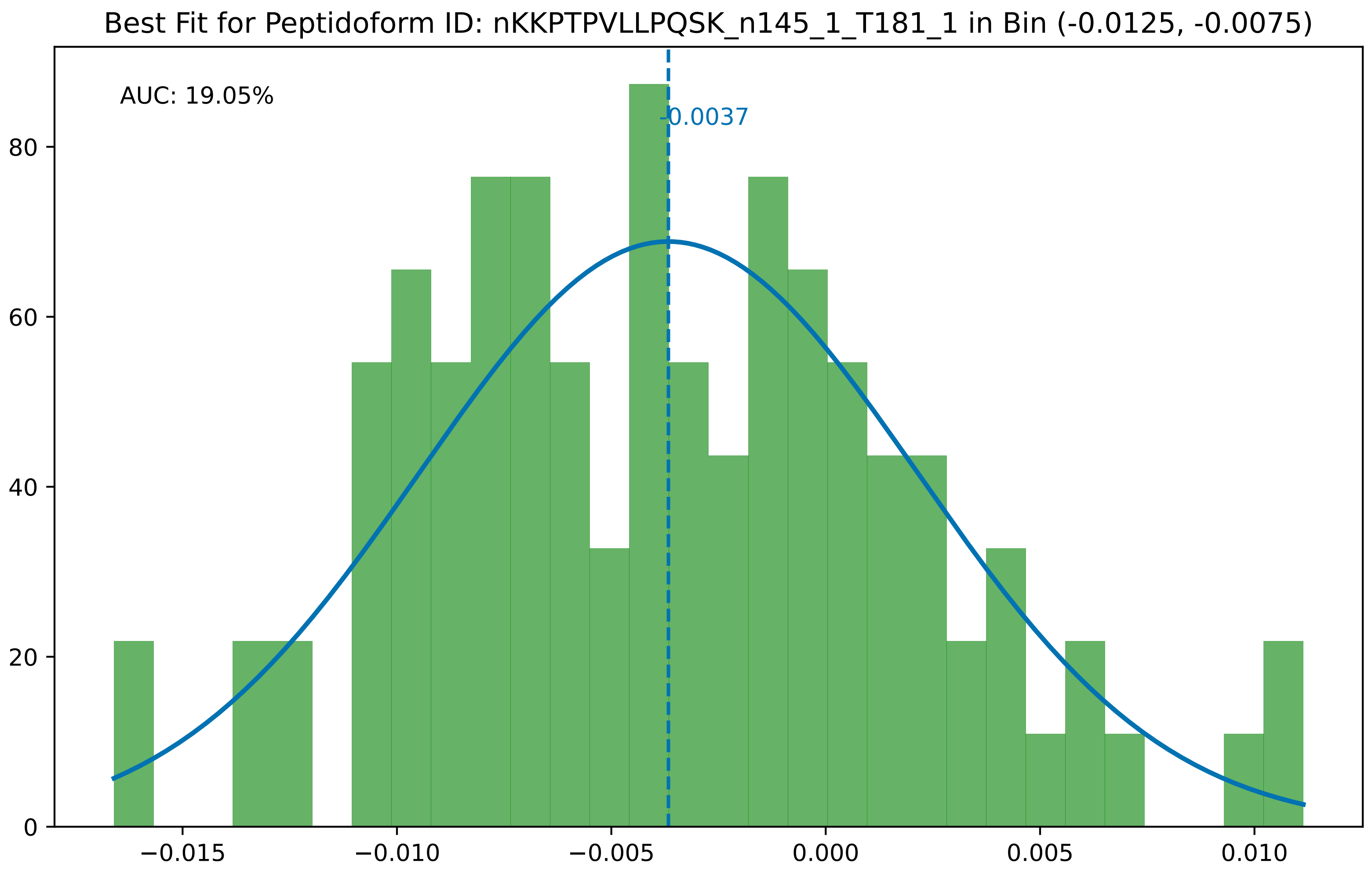

Best Fit for Peptidoform ID: nKKPTPVLLPQSK_n145_1_T181_1 in Bin (-0.0125, -0.0075)
AUC: 19.05%
-0.0037
0
20
40
60
80
−0.015
−0.010
−0.005
0.000
0.005
0.010

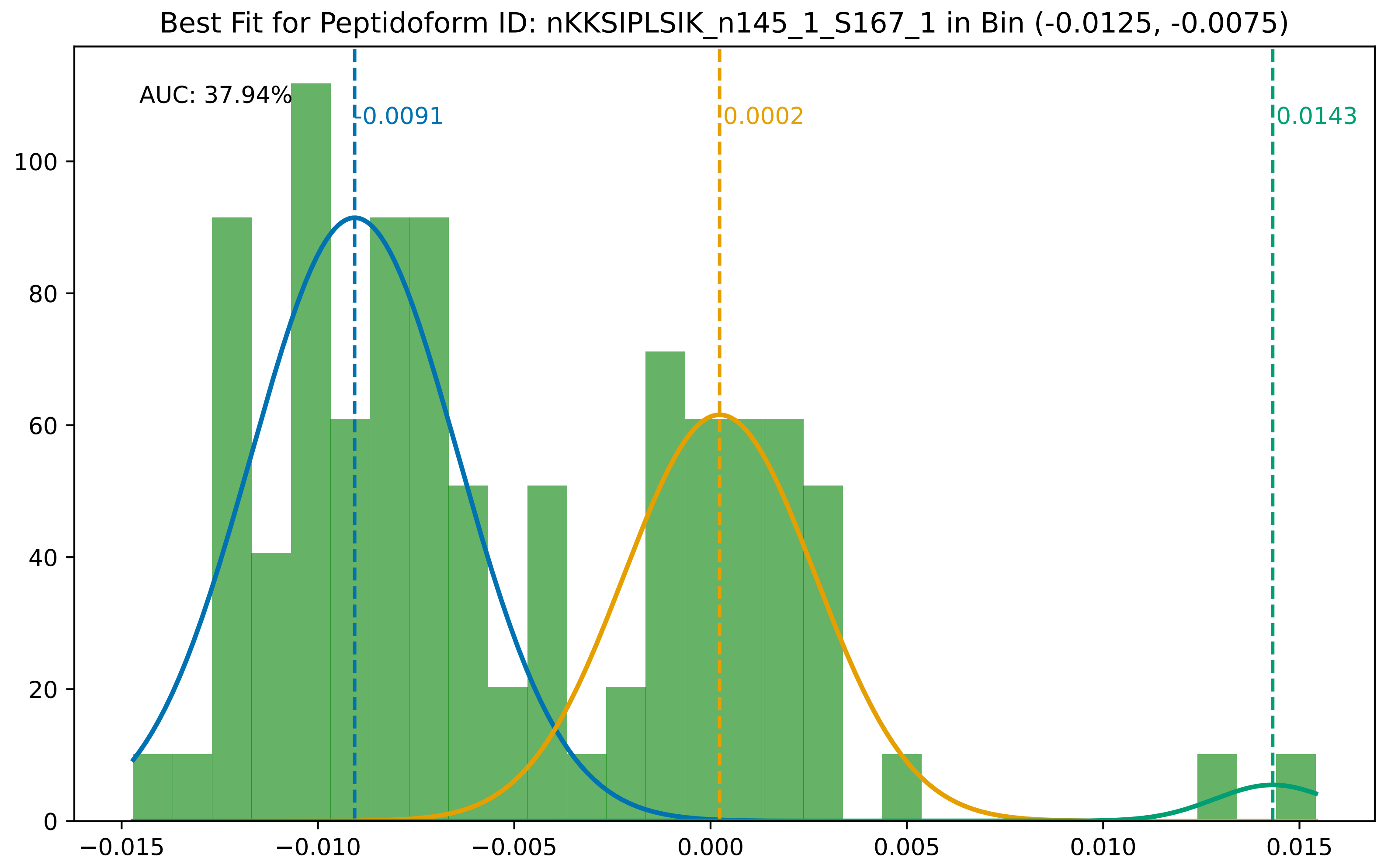
Best Fit for Peptidoform ID: nKKSIPLSIK_n145_1_S167_1 in Bin (-0.0125, -0.0075)
AUC: 37.94%
-0.0091
0.0002
0.0143
0
20
40
60
80
100
−0.015
−0.010
−0.005
0.000
0.005
0.010
0.015

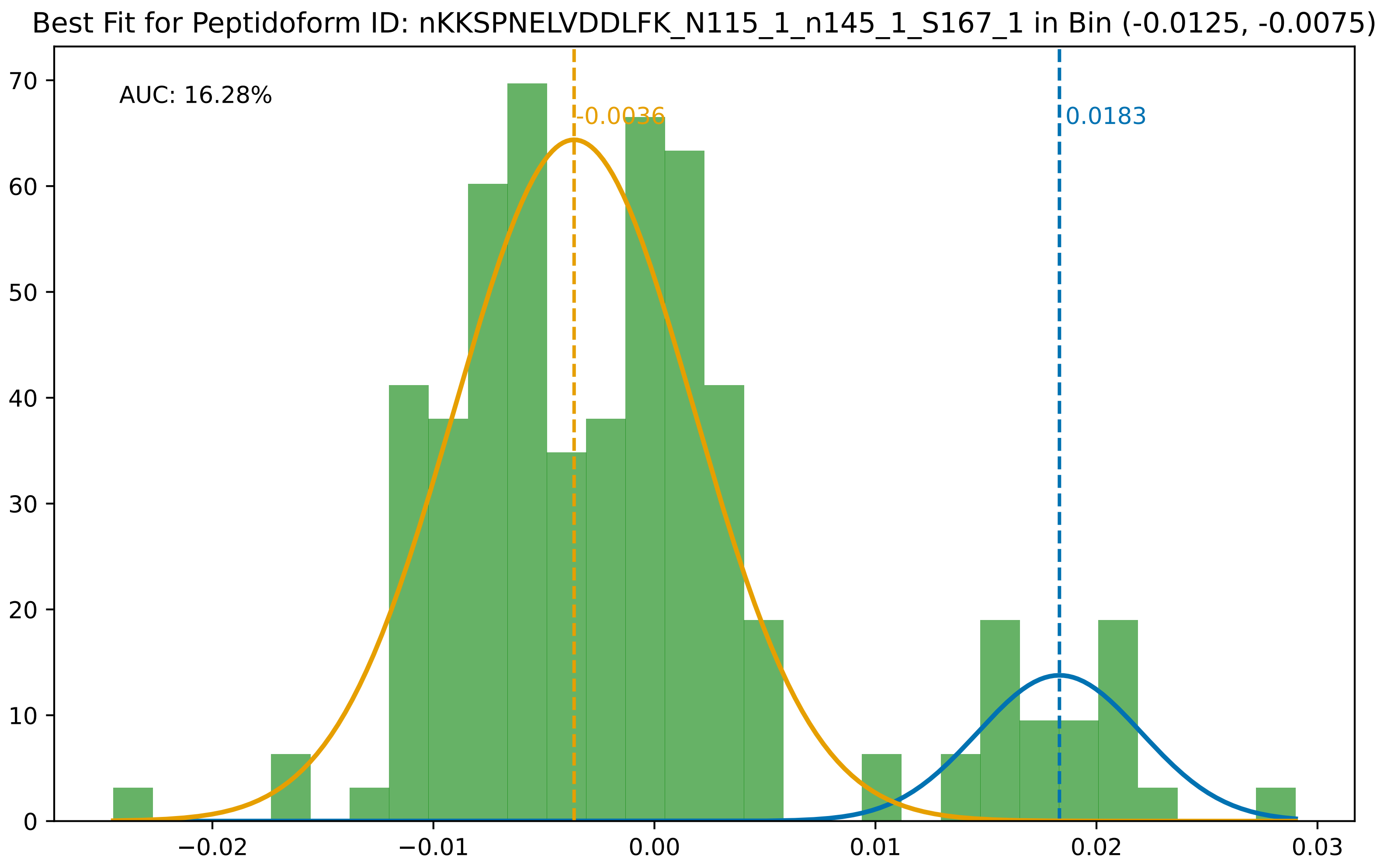

Best Fit for Peptidoform ID: nKKSPNELVDDLFK_N115_1_n145_1_S167_1 in Bin (-0.0125, -0.0075)
AUC: 16.28%
-0.0036
0.0183
0
10
20
30
40
50
60
70
−0.02
−0.01
0.00
0.01
0.02
0.03

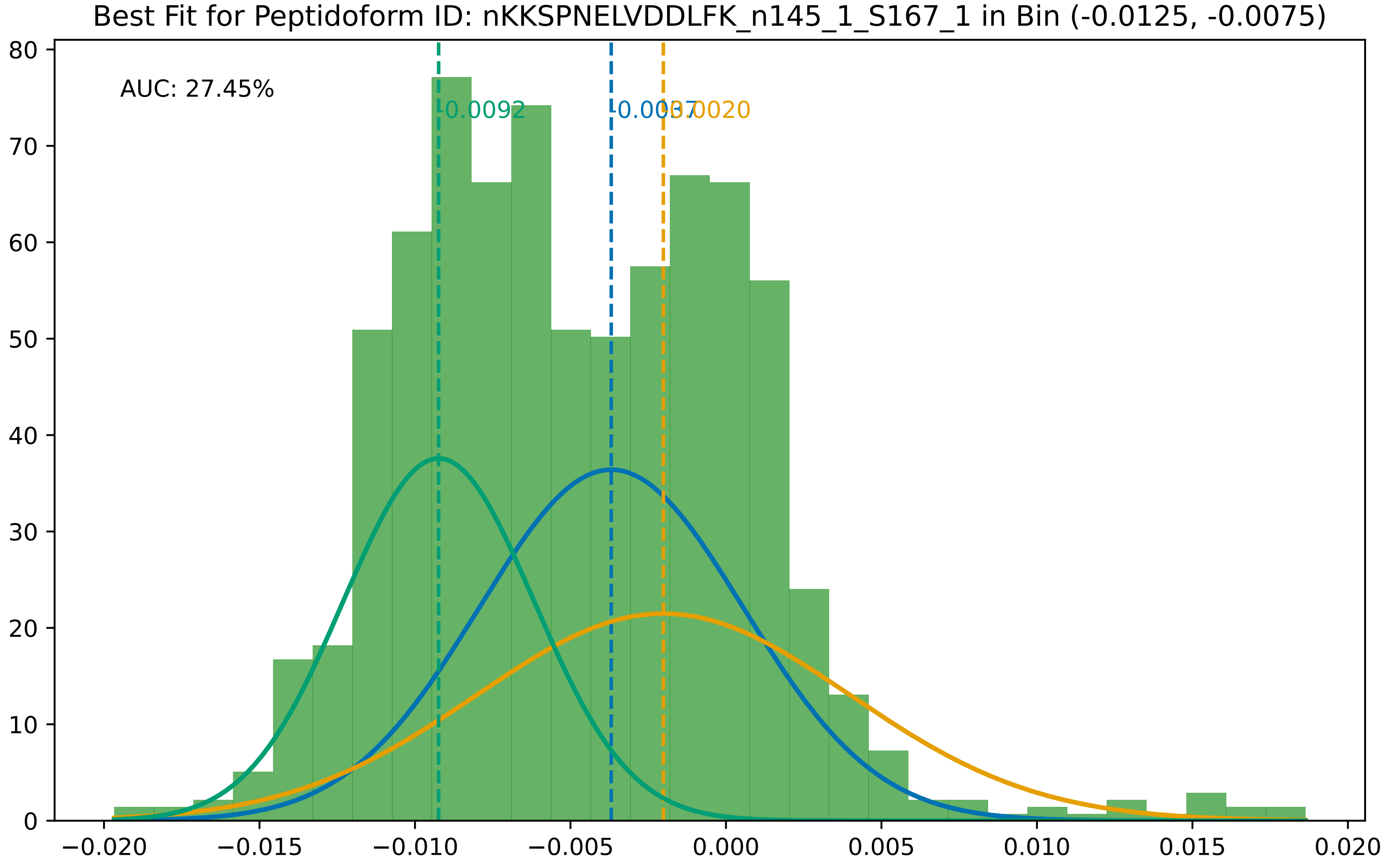

Best Fit for Peptidoform ID: nKKSPNELVDDLFK_n145_1_S167_1 in Bin (-0.0125, -0.0075)
AUC: 27.45%
-0.0092
-0.0037
-0.0020
80
70
60
50
40
30
20
10
0
−0.020
−0.015
−0.010
−0.005
0.000
0.005
0.010
0.015
0.020

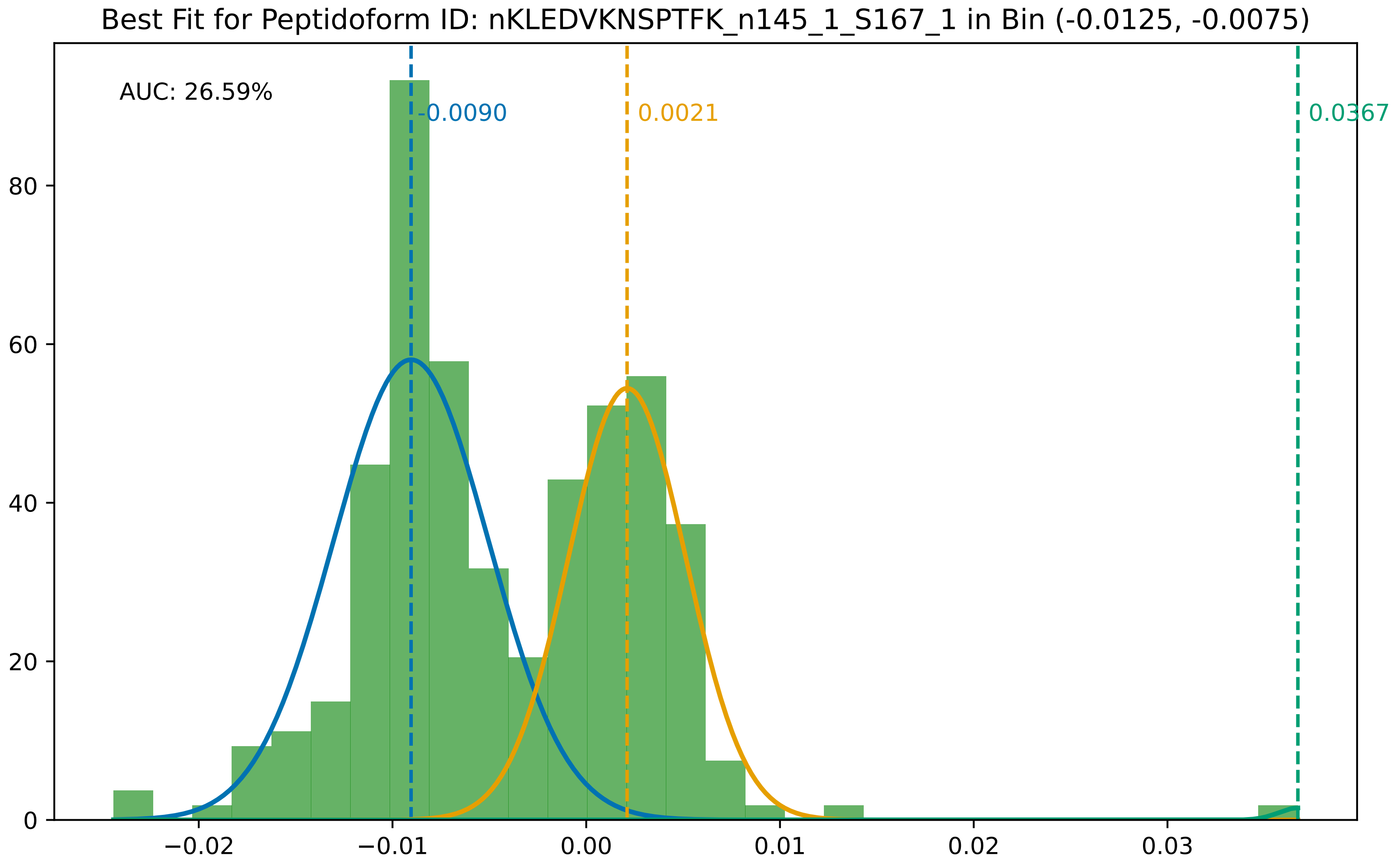

Best Fit for Peptidoform ID: nKLEDVKNSPTFK_n145_1_S167_1 in Bin (-0.0125, -0.0075)
AUC: 26.59%
-0.0090
0.0021
0.0367
0
20
40
60
80
−0.02
−0.01
0.00
0.01
0.02
0.03

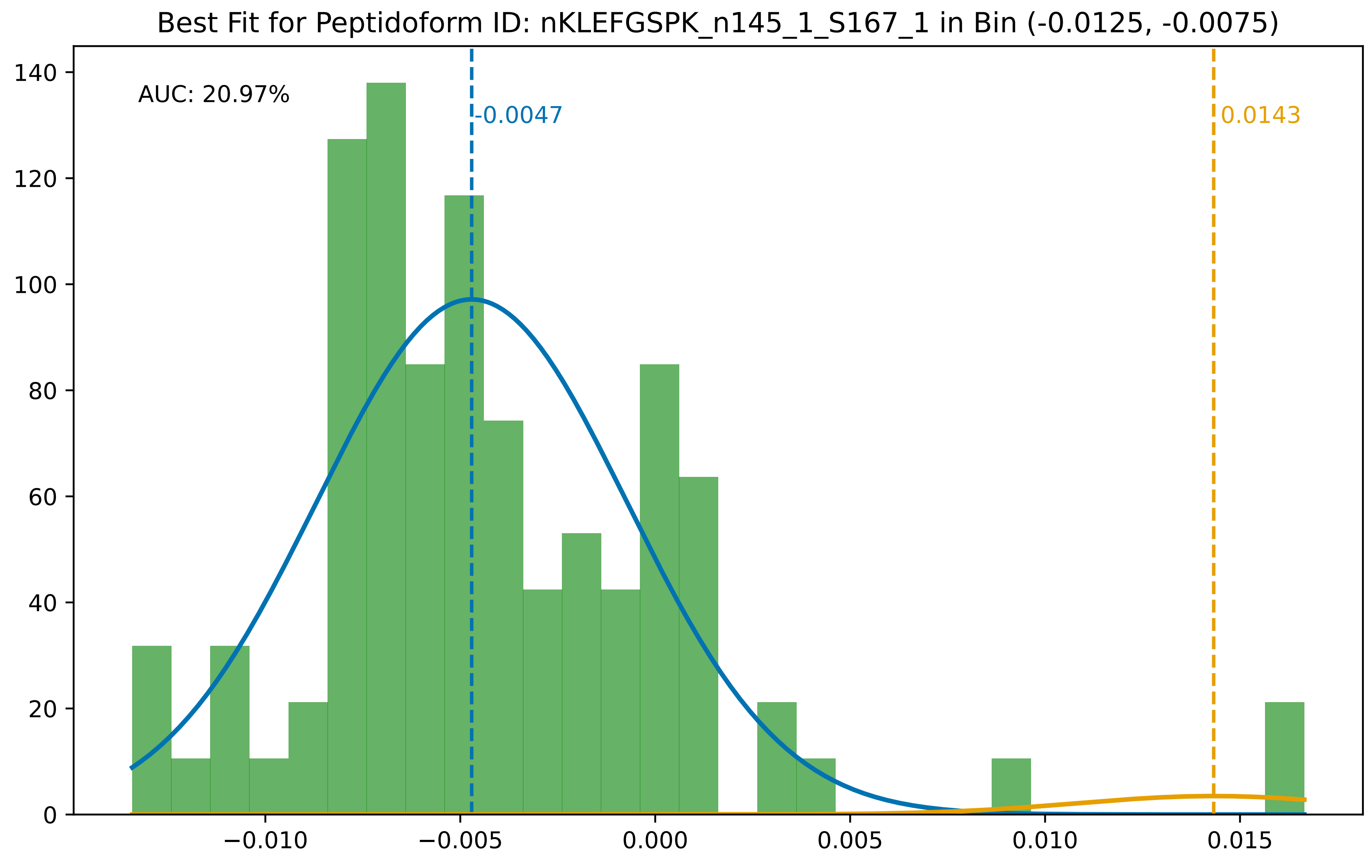

Best Fit for Peptidoform ID: nKLEFGSPK_n145_1_S167_1 in Bin (-0.0125, -0.0075)
AUC: 20.97%
-0.0047
0.0143
0
20
40
60
80
100
120
140
−0.010
−0.005
0.000
0.005
0.010
0.015

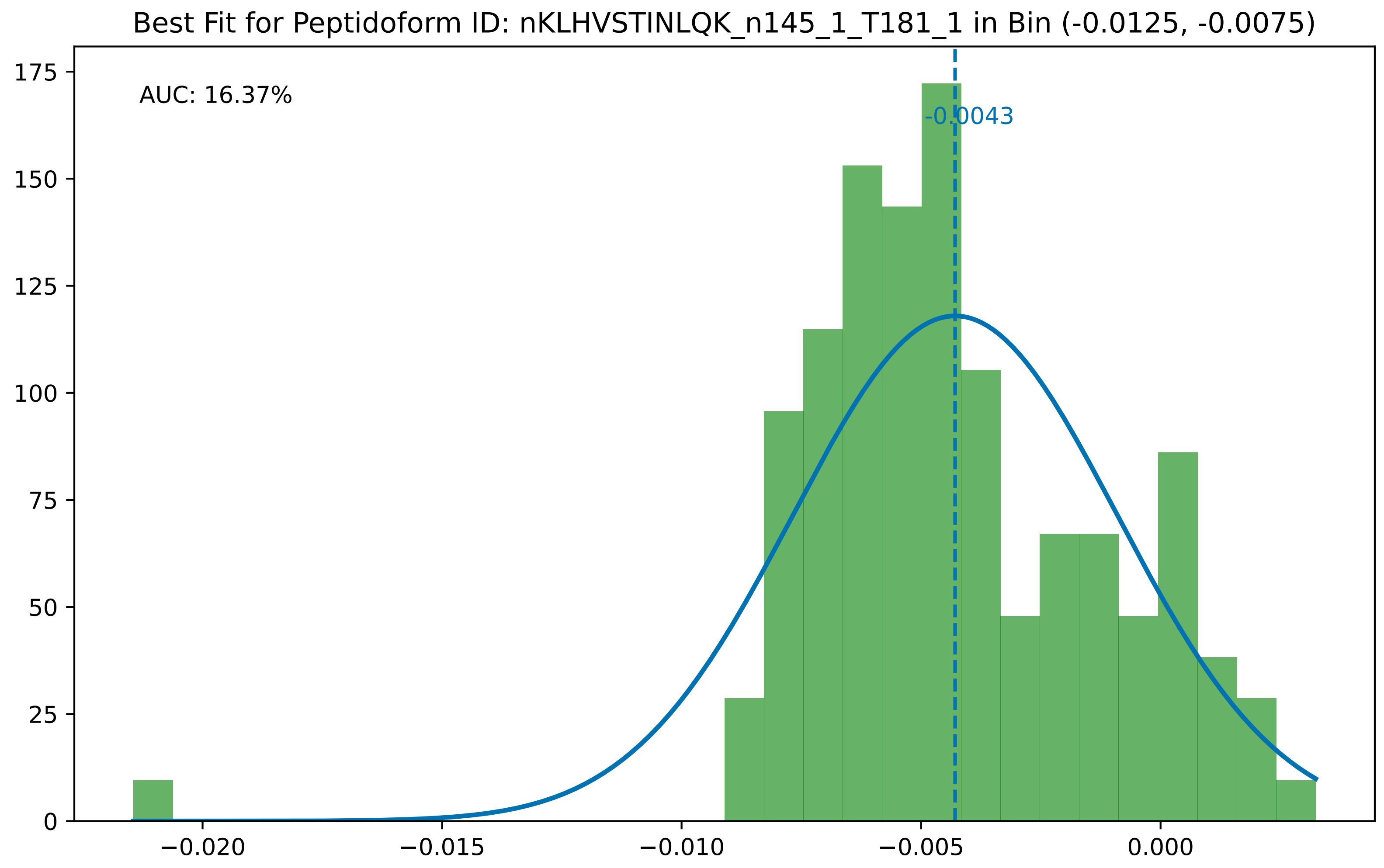

Best Fit for Peptidoform ID: nKLHVSTINLQK_n145_1_T181_1 in Bin (-0.0125, -0.0075)
AUC: 16.37%
-0.0043
0
25
50
75
100
125
150
175
−0.020
−0.015
−0.010
−0.005
0.000

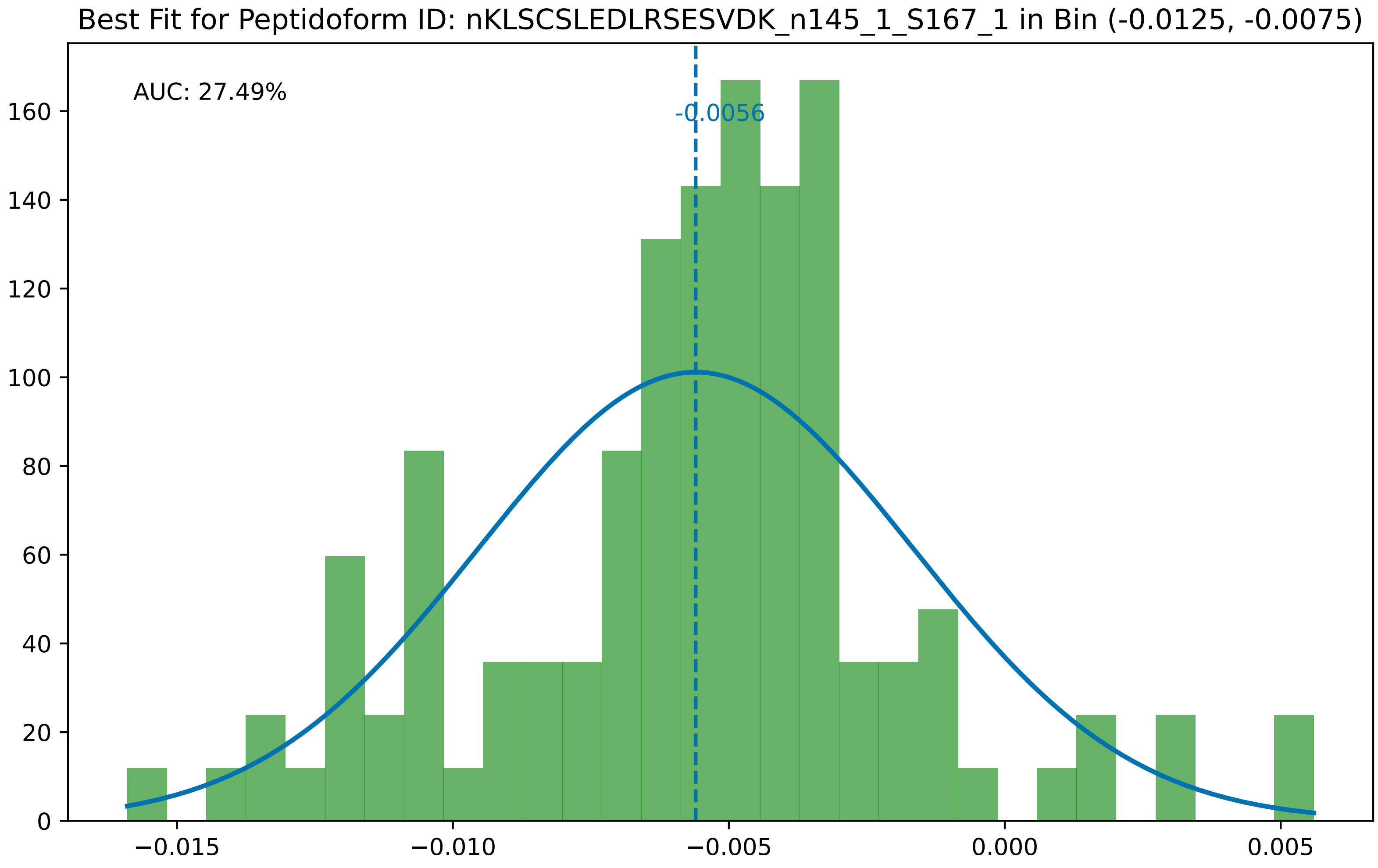

Best Fit for Peptidoform ID: nKLSCSLEDLRSESVDK_n145_1_S167_1 in Bin (-0.0125, -0.0075)
AUC: 27.49%
-0.0056
0
20
40
60
80
100
120
140
160
−0.015
−0.010
−0.005
0.000
0.005

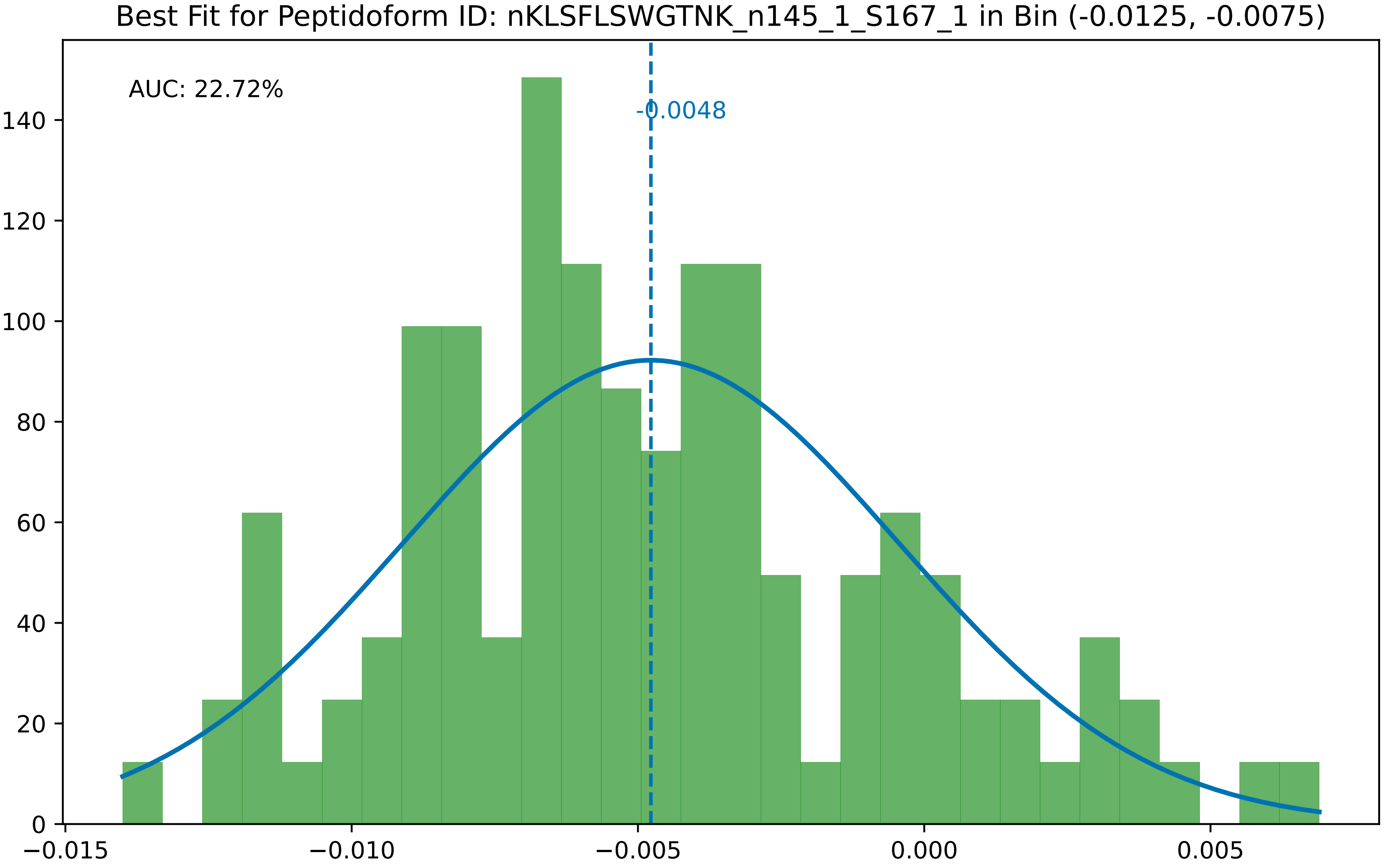

Best Fit for Peptidoform ID: nKLSFLSWGTNK_n145_1_S167_1 in Bin (-0.0125, -0.0075)
AUC: 22.72%
-0.0048
0
20
40
60
80
100
120
140
−0.015
−0.010
−0.005
0.000
0.005

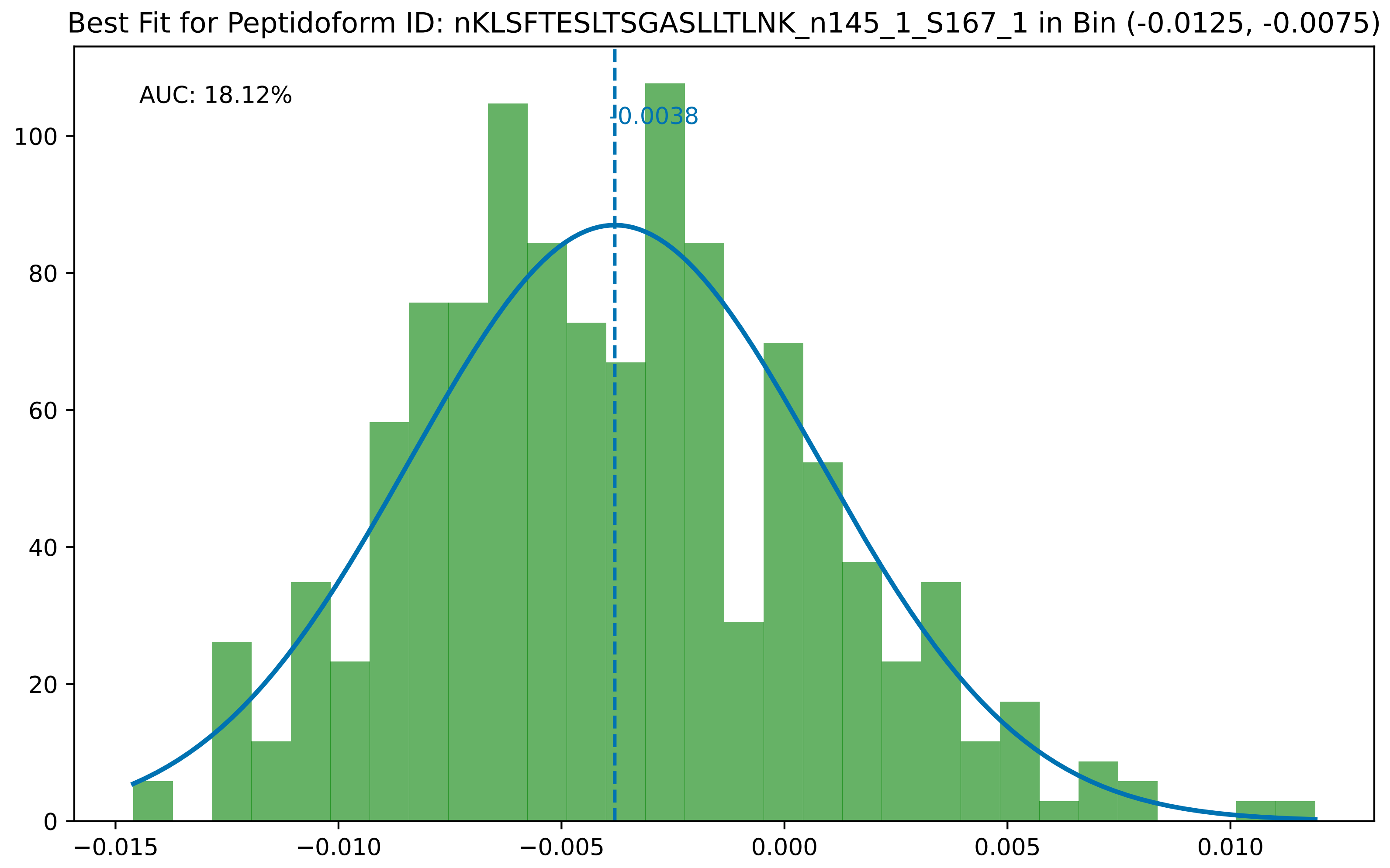

Best Fit for Peptidoform ID: nKLSFTESLTSGASLLTLNK_n145_1_S167_1 in Bin (-0.0125, -0.0075)
AUC: 18.12%
-0.0038
0
20
40
60
80
100
−0.015
−0.010
−0.005
0.000
0.005
0.010

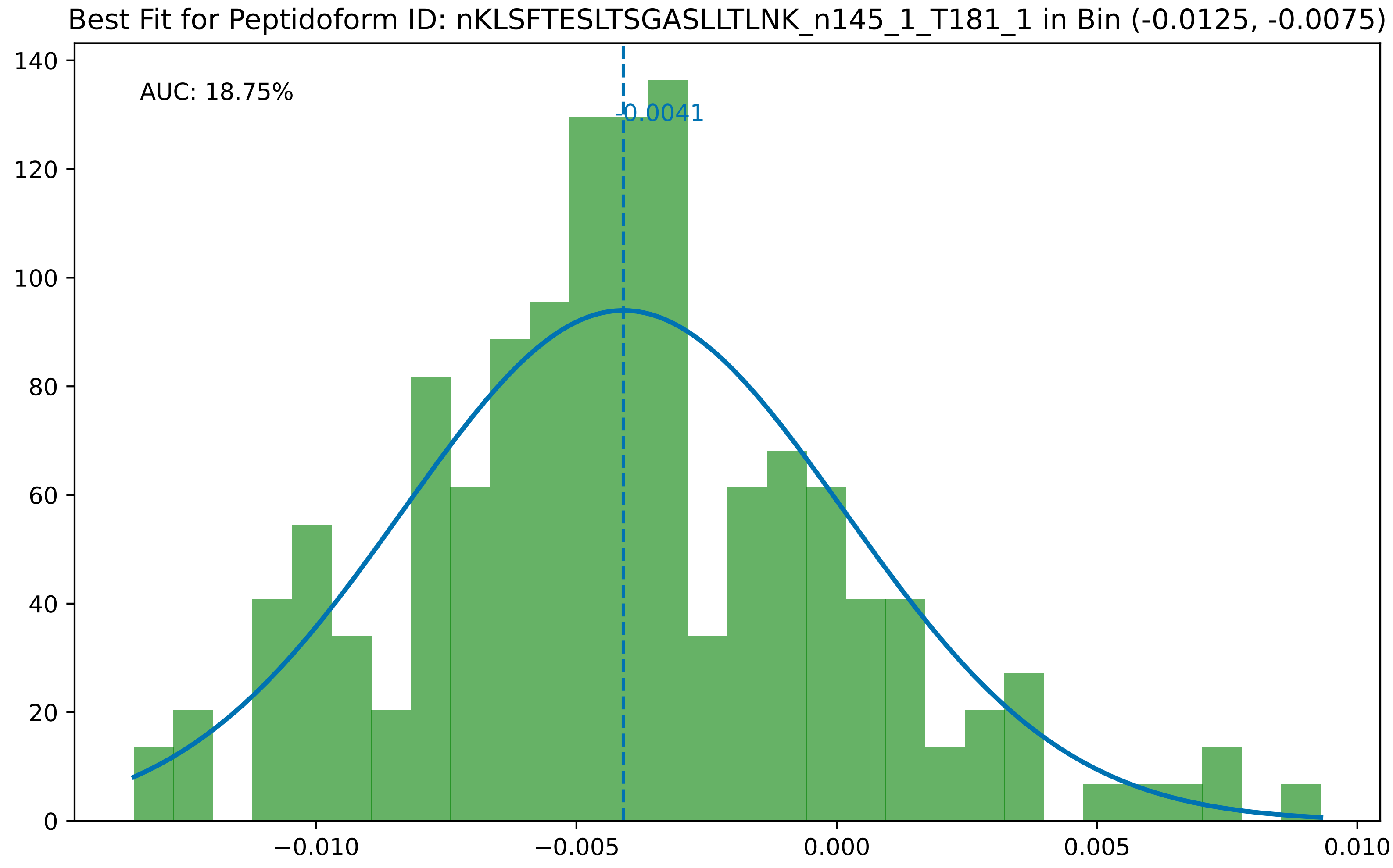

Best Fit for Peptidoform ID: nKLSFTESLTSGASLLTLNK_n145_1_T181_1 in Bin (-0.0125, -0.0075)
AUC: 18.75%
-0.0041
0
20
40
60
80
100
120
140
−0.010
−0.005
0.000
0.005
0.010

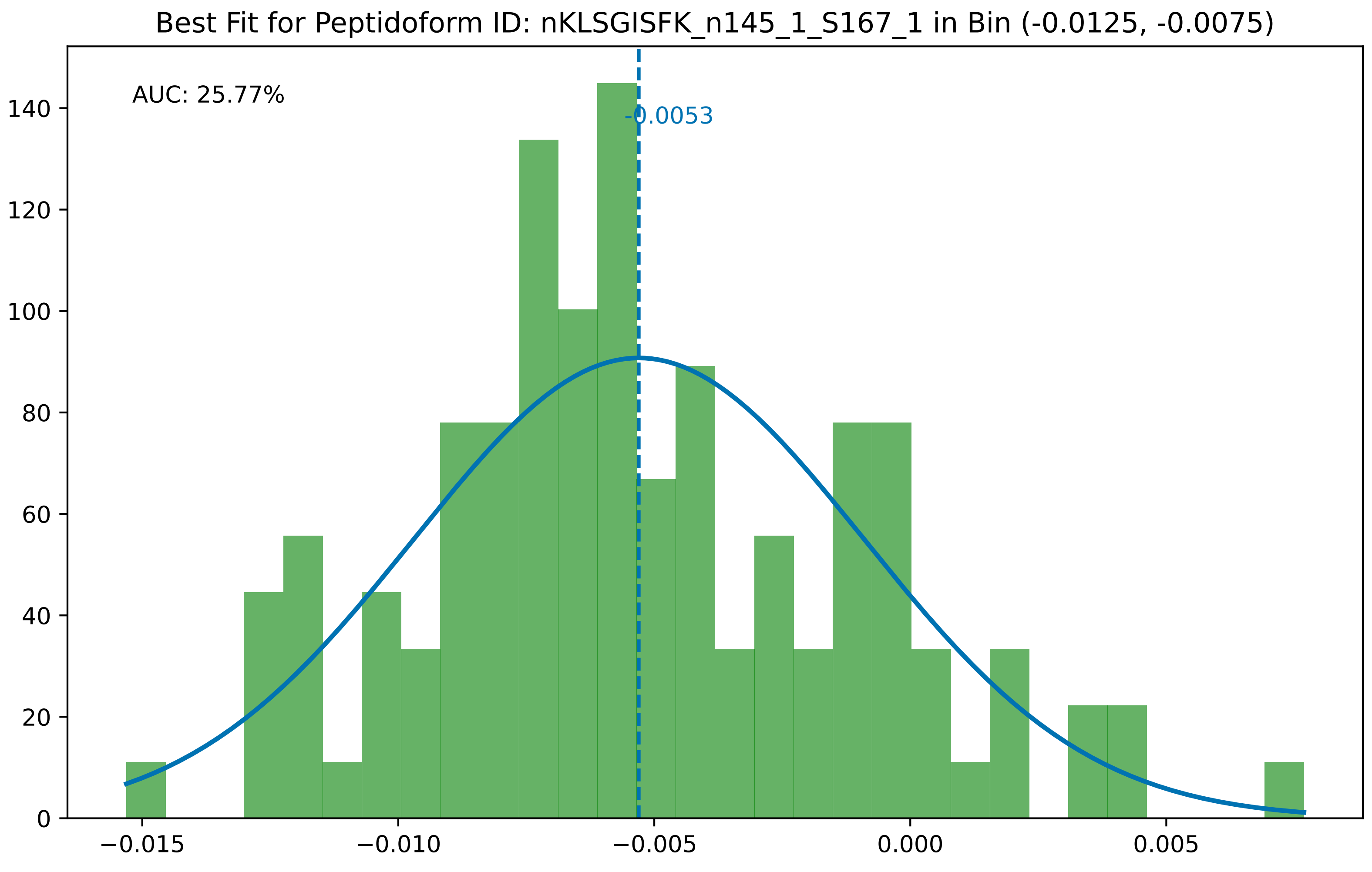

Best Fit for Peptidoform ID: nKLSGISFK_n145_1_S167_1 in Bin (-0.0125, -0.0075)
AUC: 25.77%
-0.0053
0
20
40
60
80
100
120
140
−0.015
−0.010
−0.005
0.000
0.005

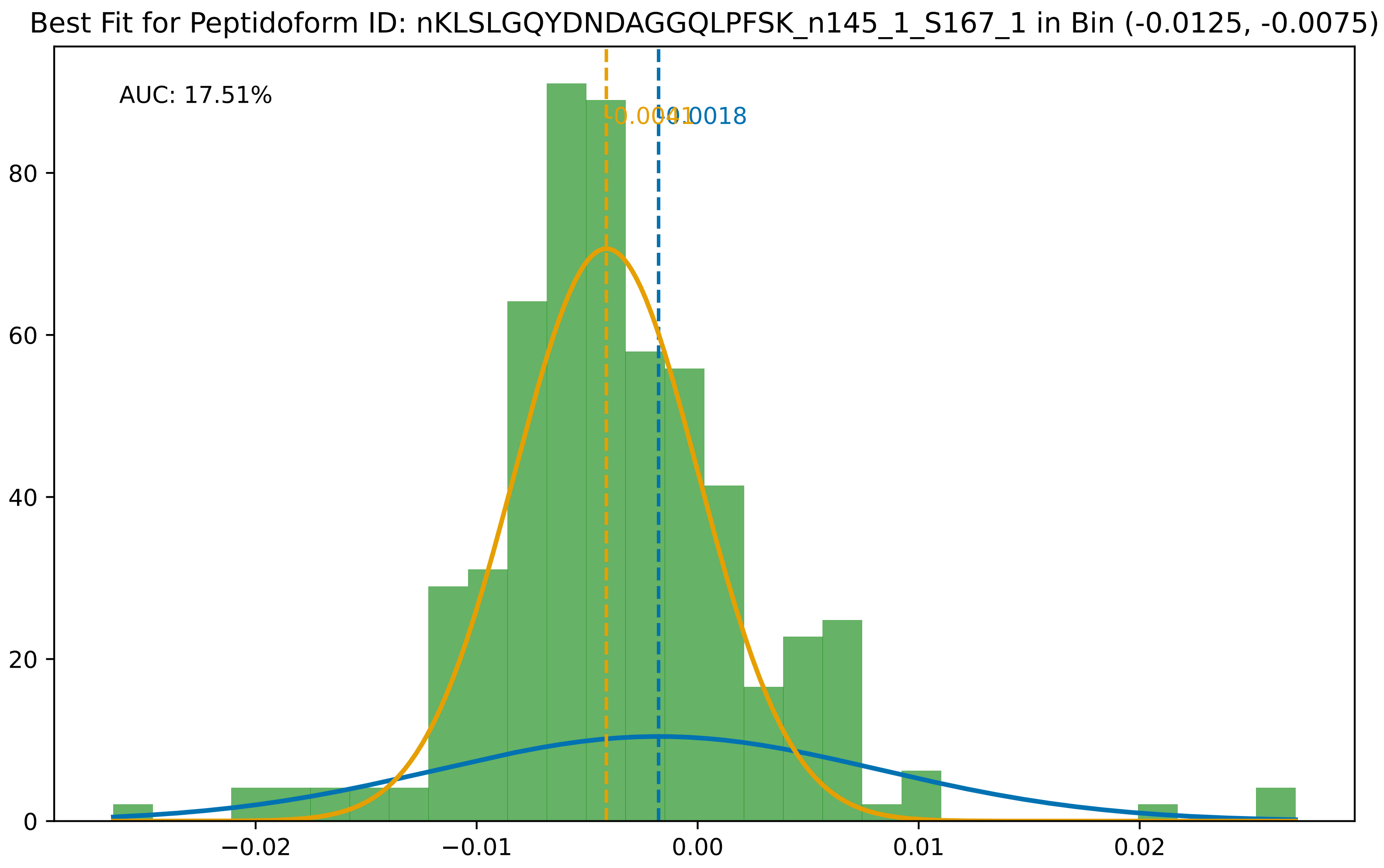
Best Fit for Peptidoform ID: nKLSLGQYDNDAGGQLPFSK_n145_1_S167_1 in Bin (-0.0125, -0.0075)
AUC: 17.51%
-0.0041
-0.0018
80
60
40
20
0
−0.02
−0.01
0.00
0.01
0.02

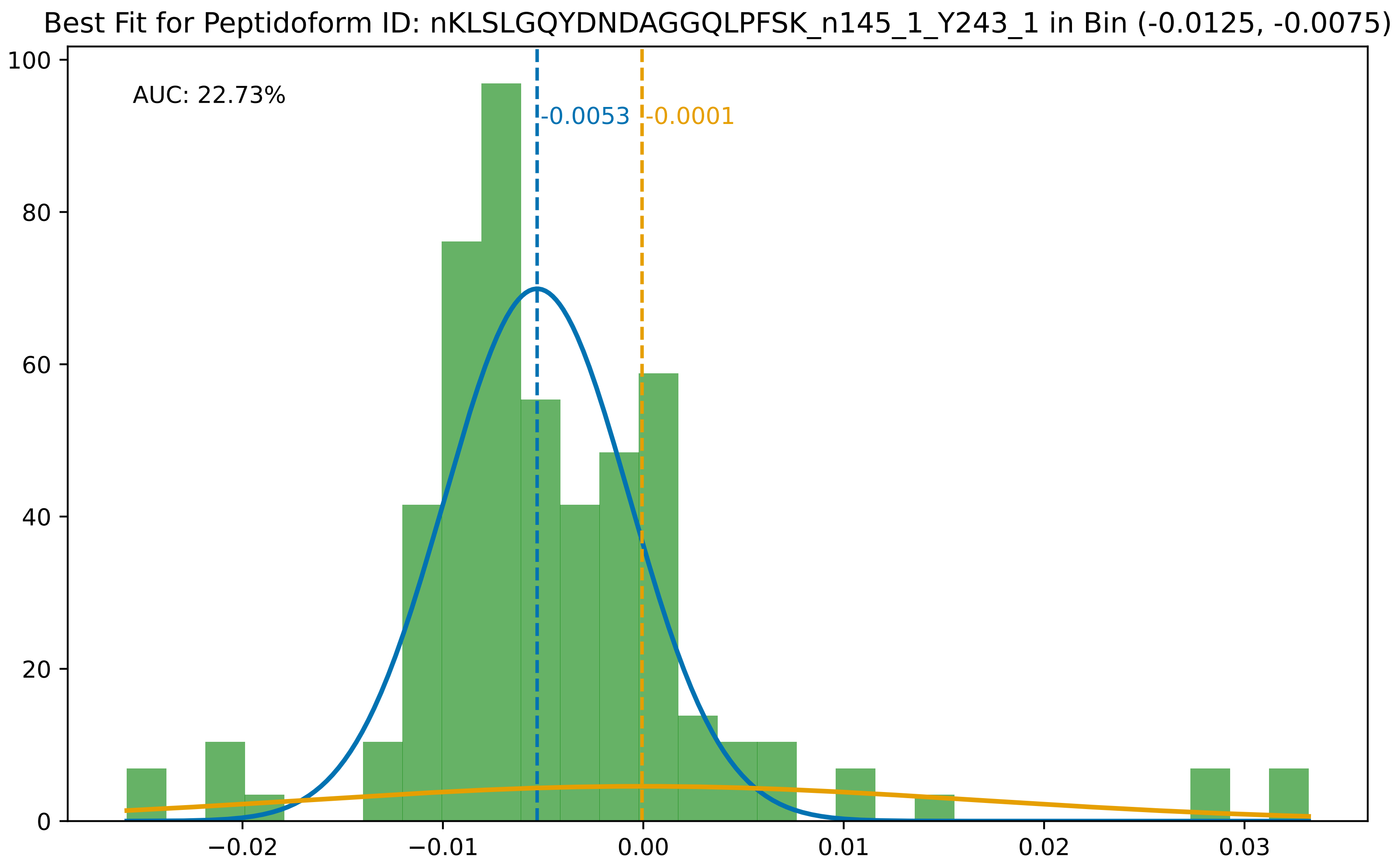
Best Fit for Peptidoform ID: nKLSLGQYDNDAGGQLPFSK_n145_1_Y243_1 in Bin (-0.0125, -0.0075)
AUC: 22.73%
-0.0053
-0.0001
100
80
60
40
20
0
−0.02
−0.01
0.00
0.01
0.02
0.03

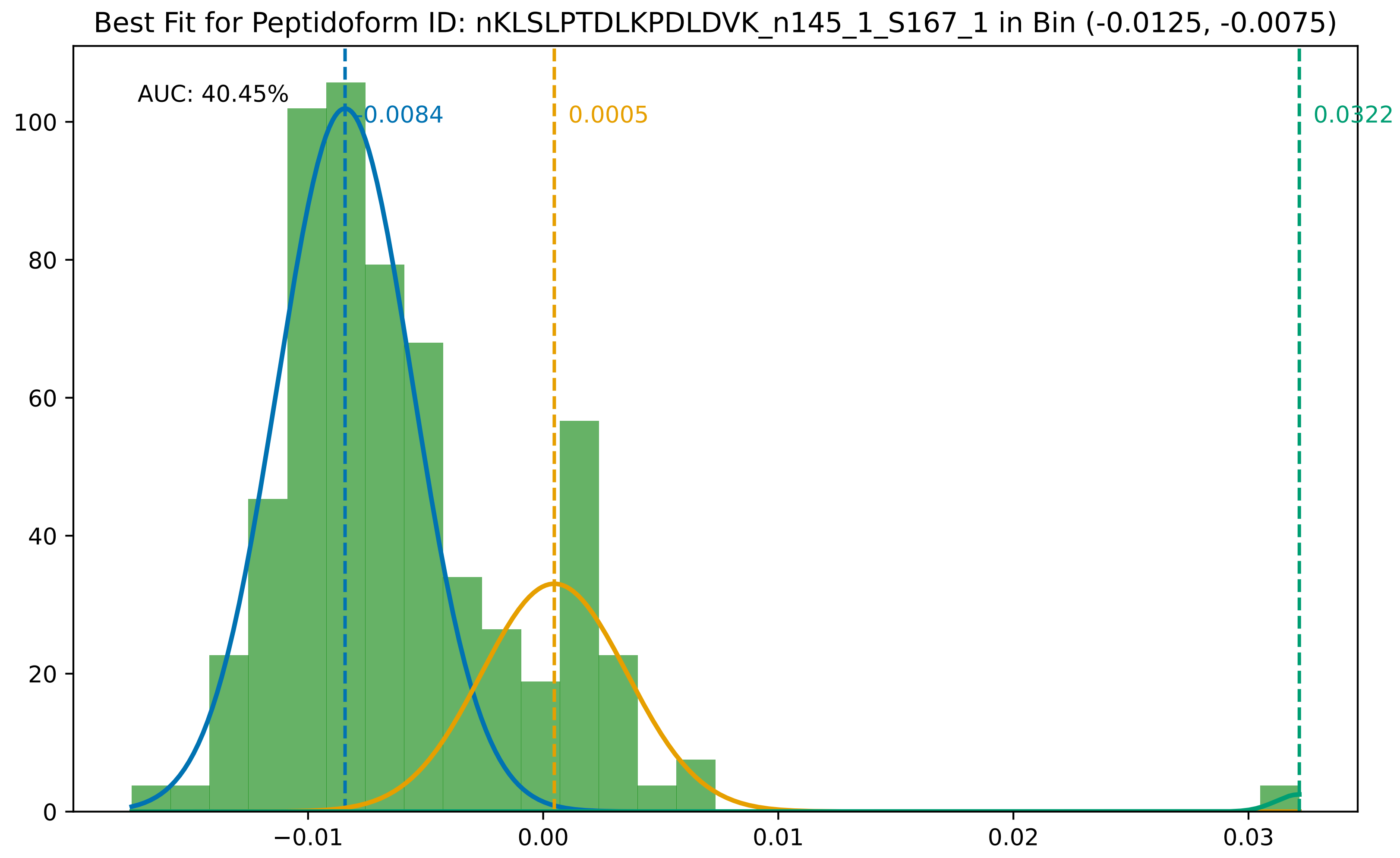

Best Fit for Peptidoform ID: nKLSLPTDLKPDLDVK_n145_1_S167_1 in Bin (-0.0125, -0.0075)
AUC: 40.45%
-0.0084
0.0005
0.0322
0
20
40
60
80
100
−0.01
0.00
0.01
0.02
0.03

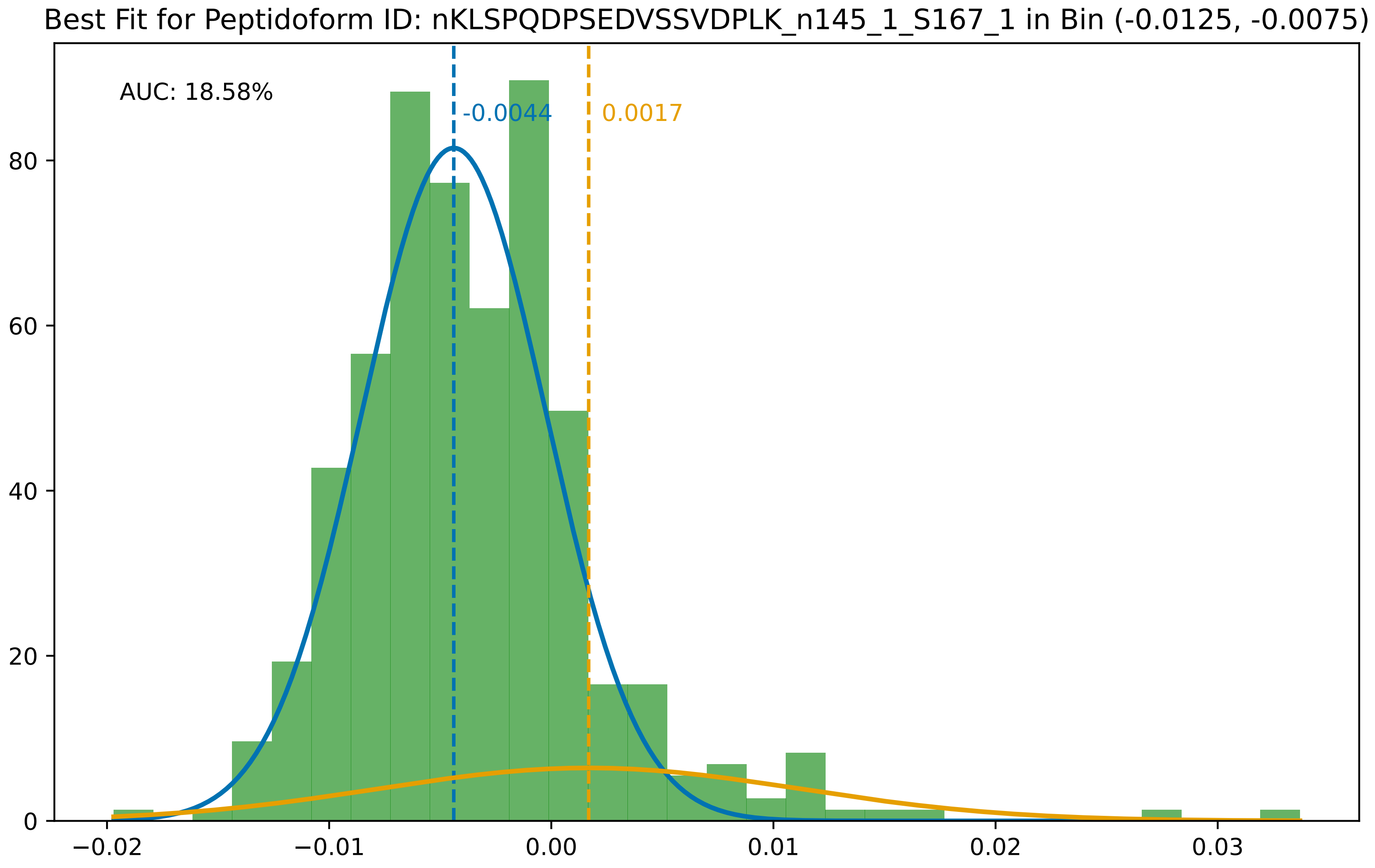

Best Fit for Peptidoform ID: nKLSPQDPSEDVSSVDPLK_n145_1_S167_1 in Bin (-0.0125, -0.0075)
AUC: 18.58%
-0.0044
0.0017
0
20
40
60
80
−0.02
−0.01
0.00
0.01
0.02
0.03

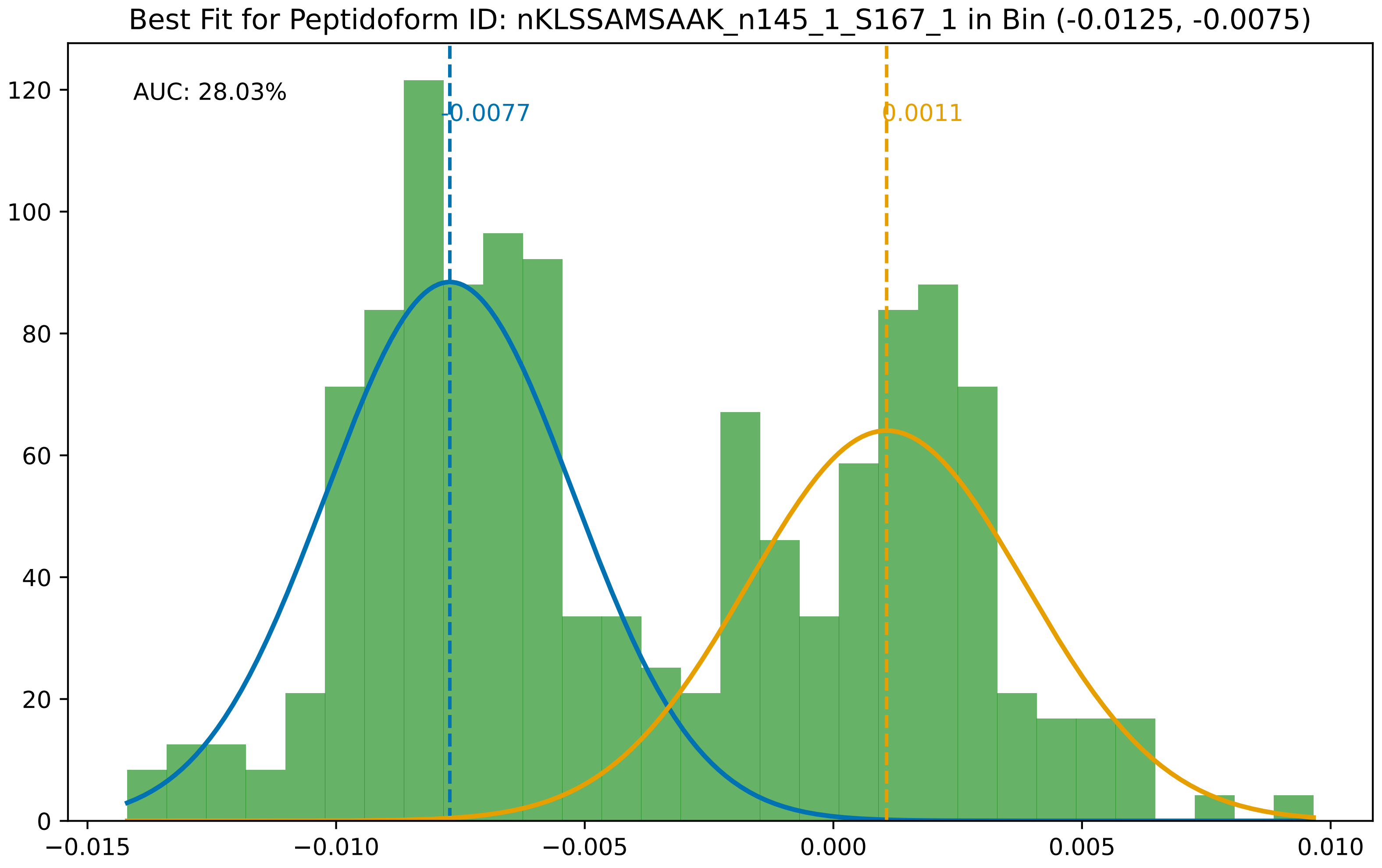
Best Fit for Peptidoform ID: nKLSSAMSAAK_n145_1_S167_1 in Bin (-0.0125, -0.0075)
AUC: 28.03%
-0.0077
0.0011
0
20
40
60
80
100
120
−0.015
−0.010
−0.005
0.000
0.005
0.010

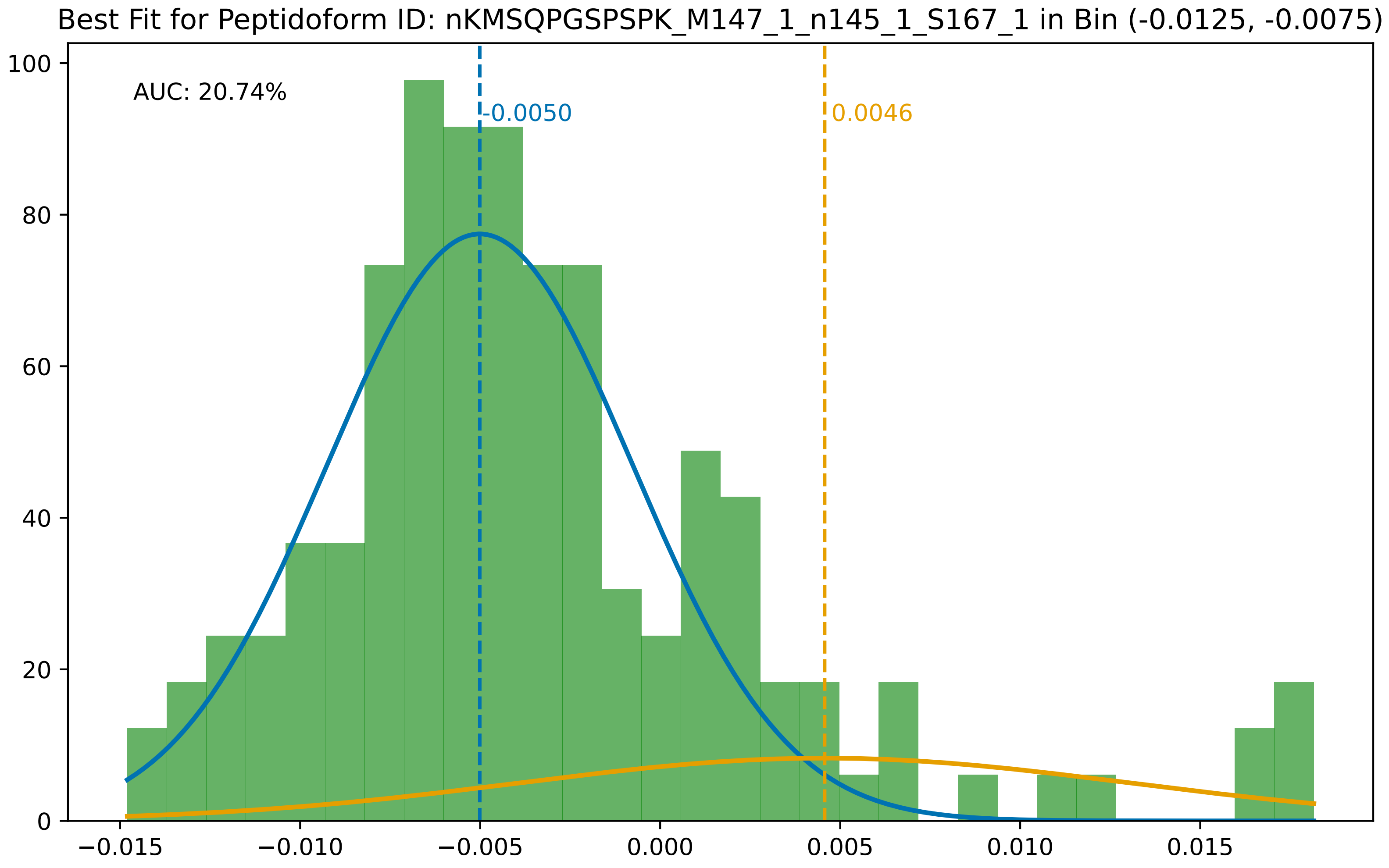

Best Fit for Peptidoform ID: nKMSQPGSPSPK_M147_1_n145_1_S167_1 in Bin (-0.0125, -0.0075)
AUC: 20.74%
-0.0050
0.0046
100
80
60
40
20
0
−0.015
−0.010
−0.005
0.000
0.005
0.010
0.015

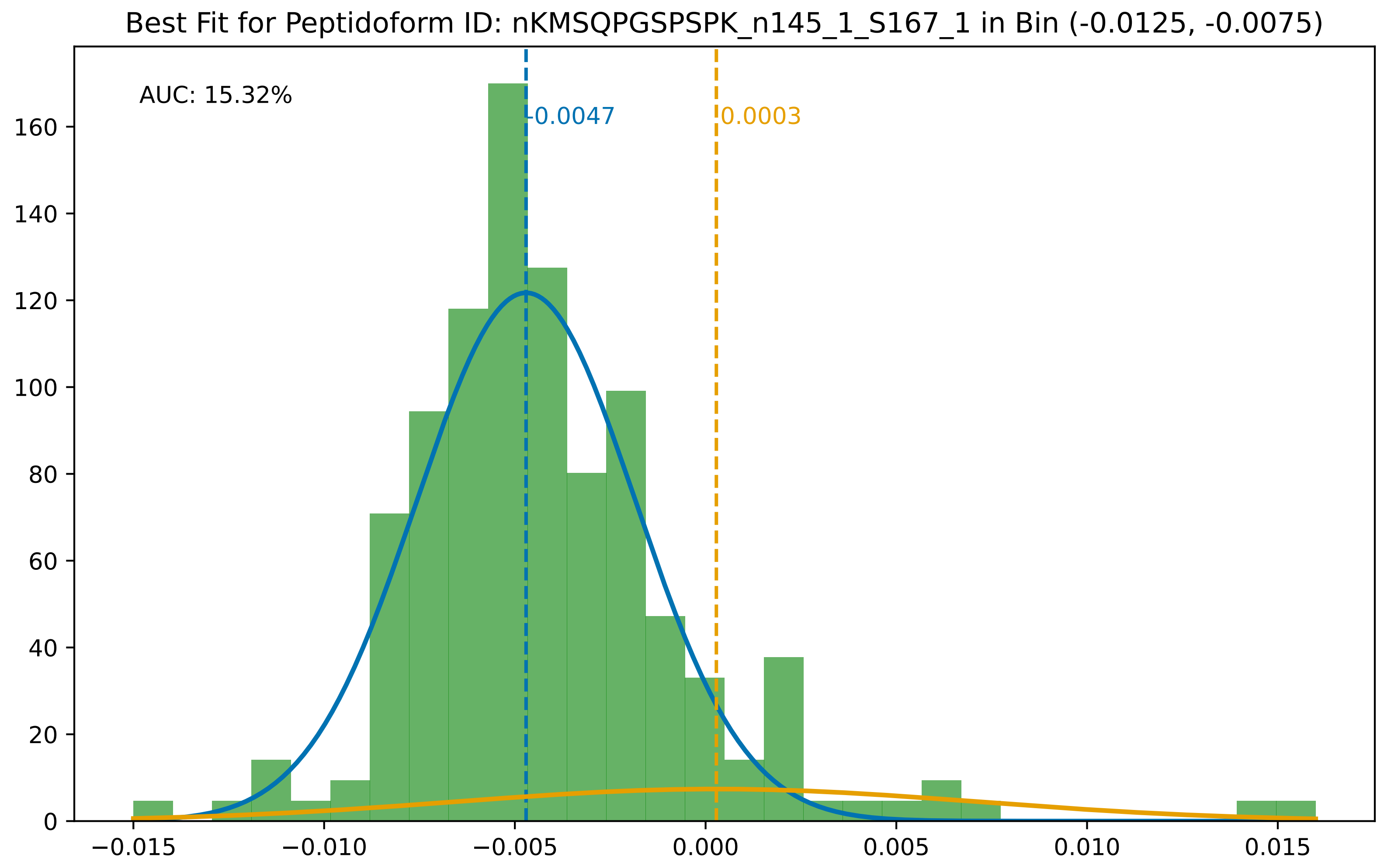

Best Fit for Peptidoform ID: nKMSQPGSPSPK_n145_1_S167_1 in Bin (-0.0125, -0.0075)
AUC: 15.32%
-0.0047
0.0003
0
20
40
60
80
100
120
140
160
−0.015
−0.010
−0.005
0.000
0.005
0.010
0.015

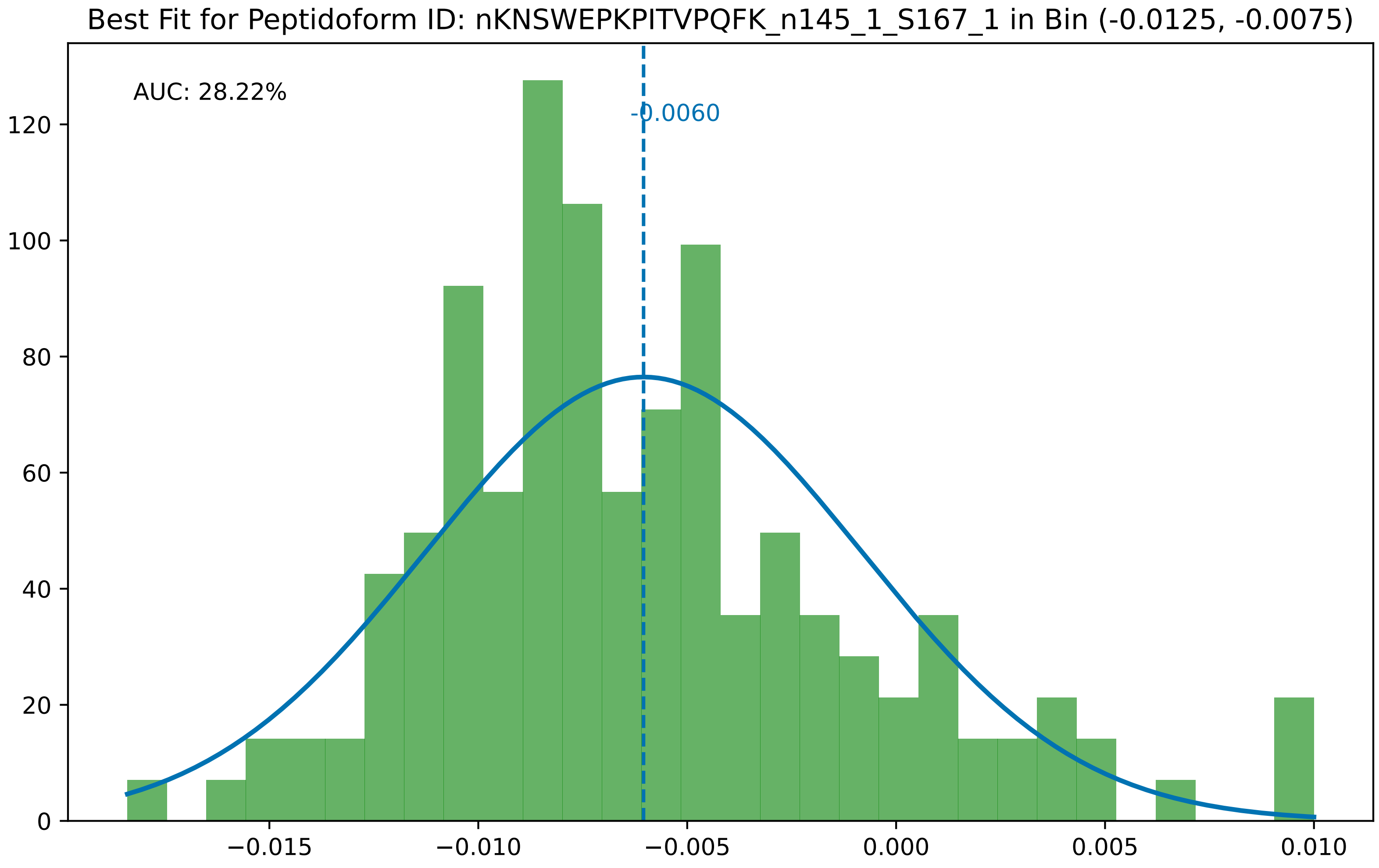
Best Fit for Peptidoform ID: nKNSWEPKPITVPQFK_n145_1_S167_1 in Bin (-0.0125, -0.0075)
AUC: 28.22%
-0.0060
0
20
40
60
80
100
120
−0.015
−0.010
−0.005
0.000
0.005
0.010

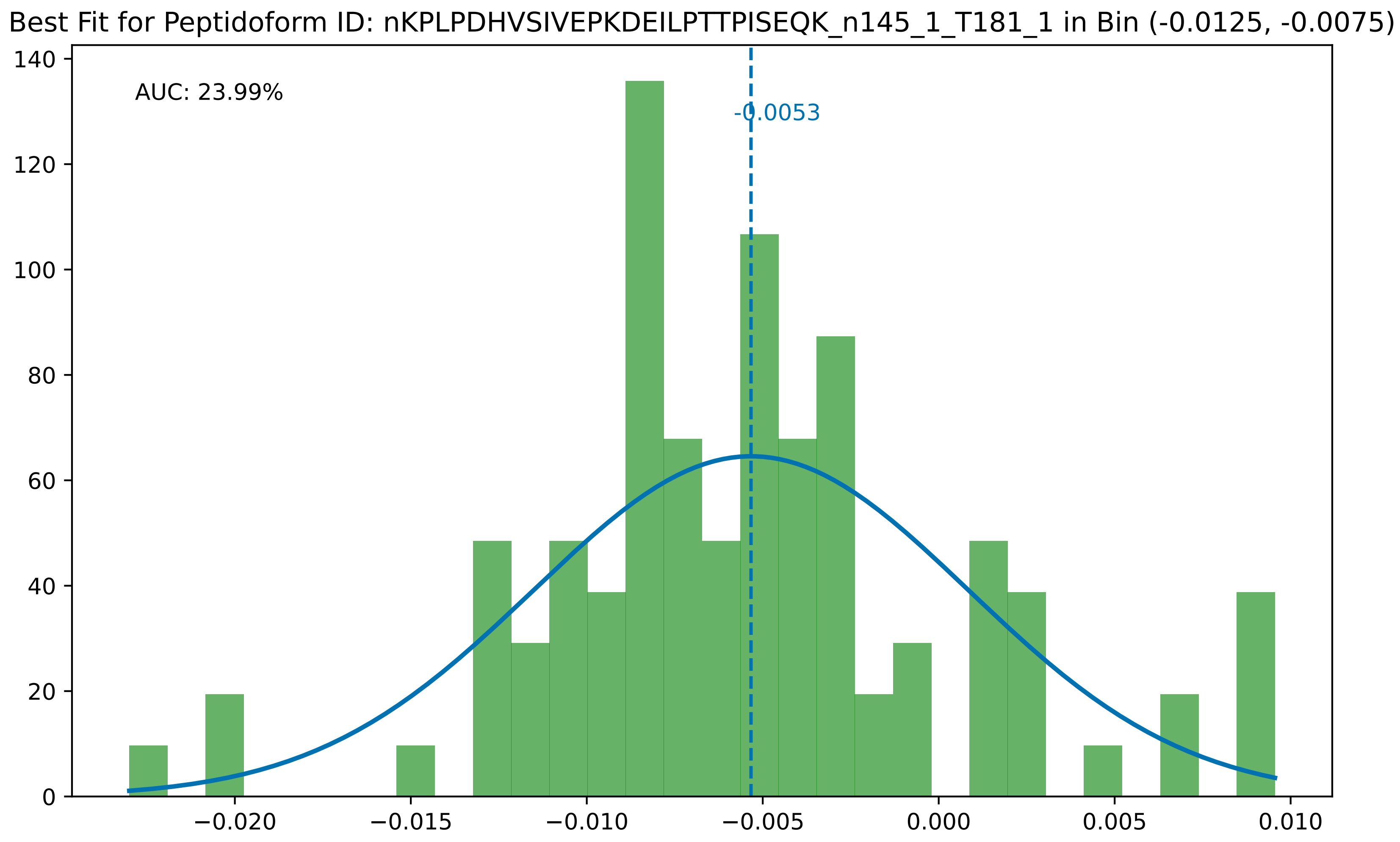

Best Fit for Peptidoform ID: nKPLPDHVSIVEPKDEILPTTPISEQK_n145_1_T181_1 in Bin (-0.0125, -0.0075)
AUC: 23.99%
-0.0053
0
20
40
60
80
100
120
140
−0.020
−0.015
−0.010
−0.005
0.000
0.005
0.010

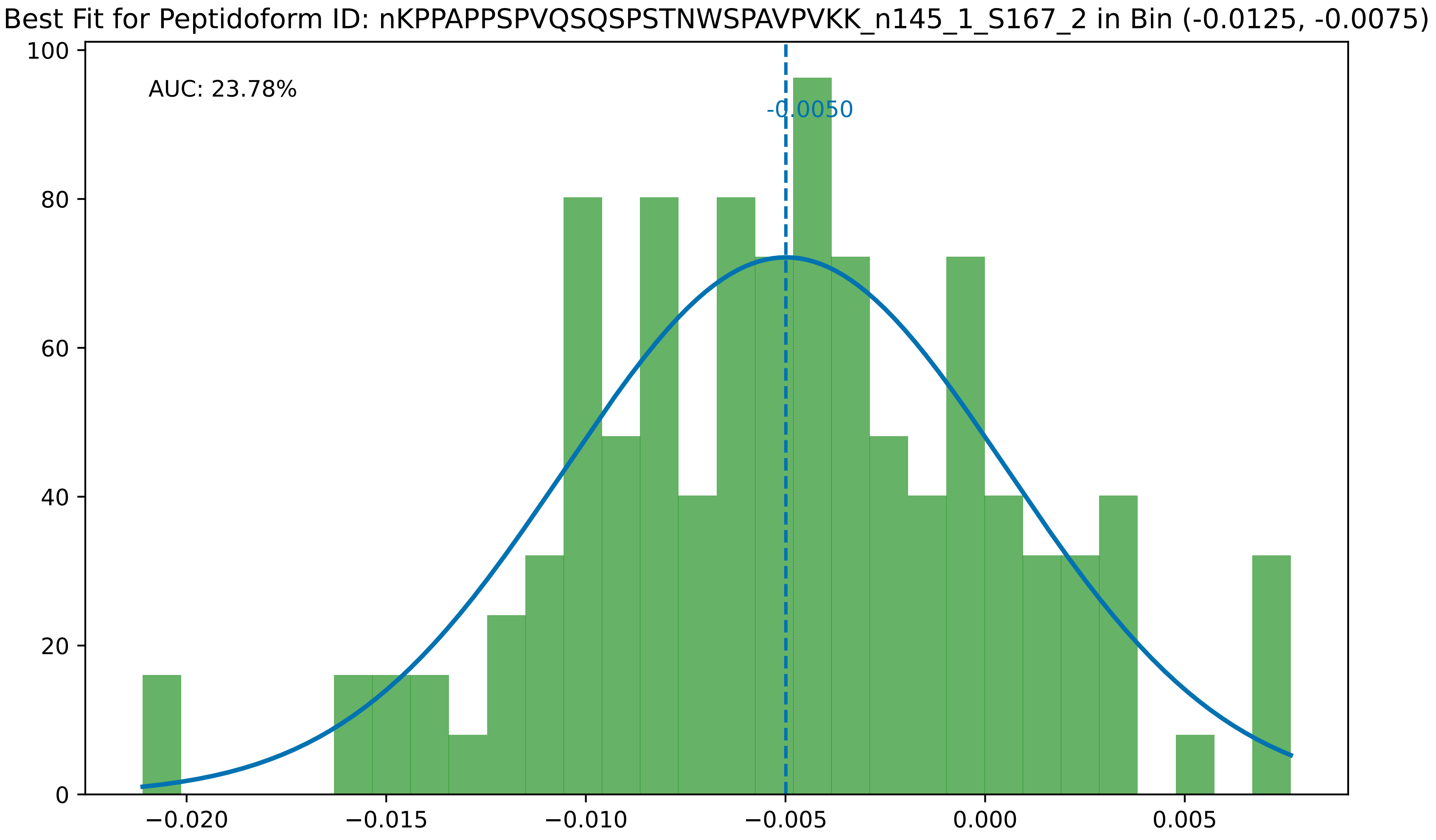

Best Fit for Peptidoform ID: nKPPAPPSPVQSQSPSTNWSPAVPVKK_n145_1_S167_2 in Bin (-0.0125, -0.0075)
AUC: 23.78%
-0.0050
0
20
40
60
80
100
−0.020
−0.015
−0.010
−0.005
0.000
0.005

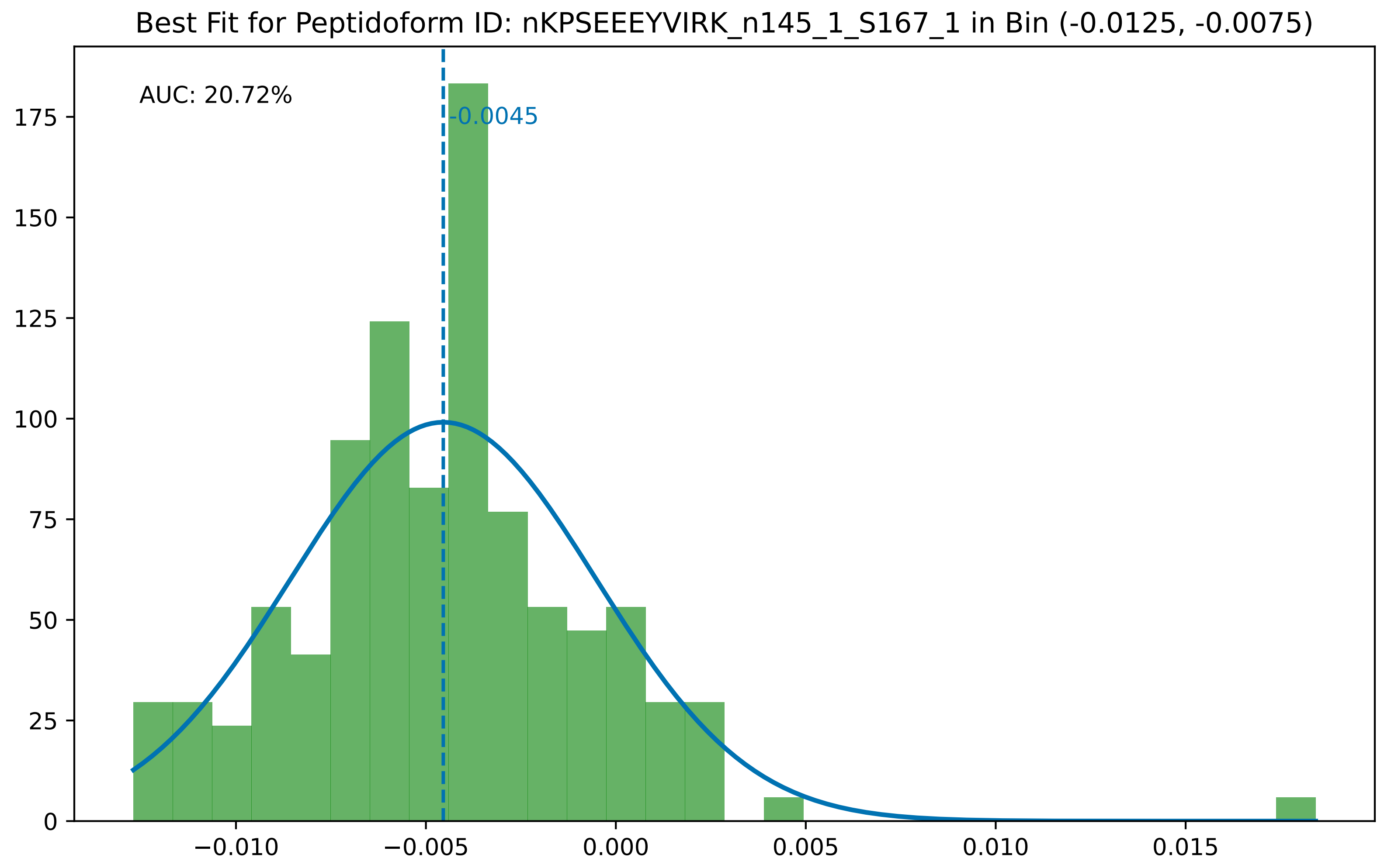

Best Fit for Peptidoform ID: nKPSEEEYVIRK_n145_1_S167_1 in Bin (-0.0125, -0.0075)
AUC: 20.72%
-0.0045
0
25
50
75
100
125
150
175
−0.010
−0.005
0.000
0.005
0.010
0.015

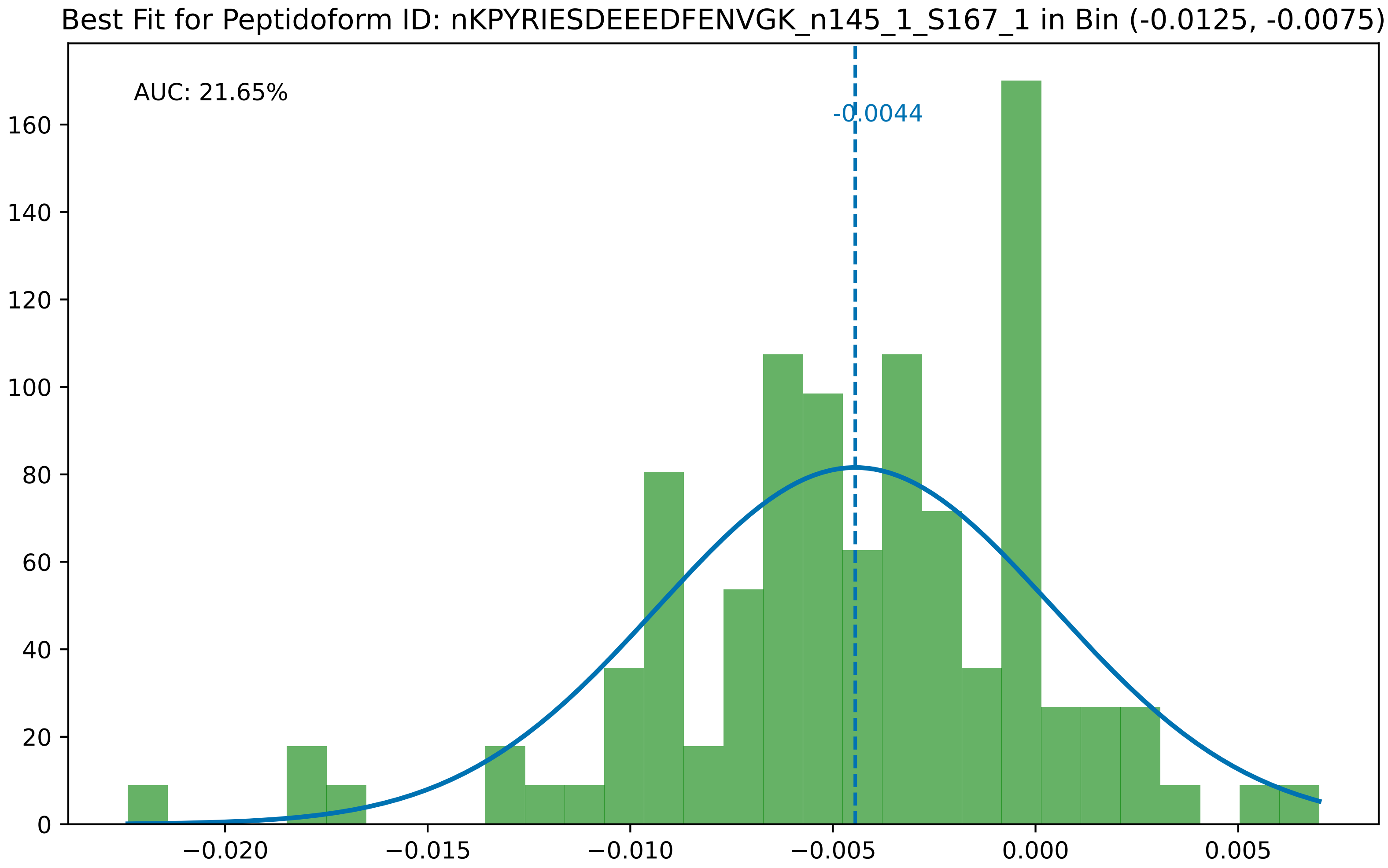

Best Fit for Peptidoform ID: nKPYRIESDEEEDFENVGK_n145_1_S167_1 in Bin (-0.0125, -0.0075)
AUC: 21.65%
-0.0044
0
20
40
60
80
100
120
140
160
−0.020
−0.015
−0.010
−0.005
0.000
0.005

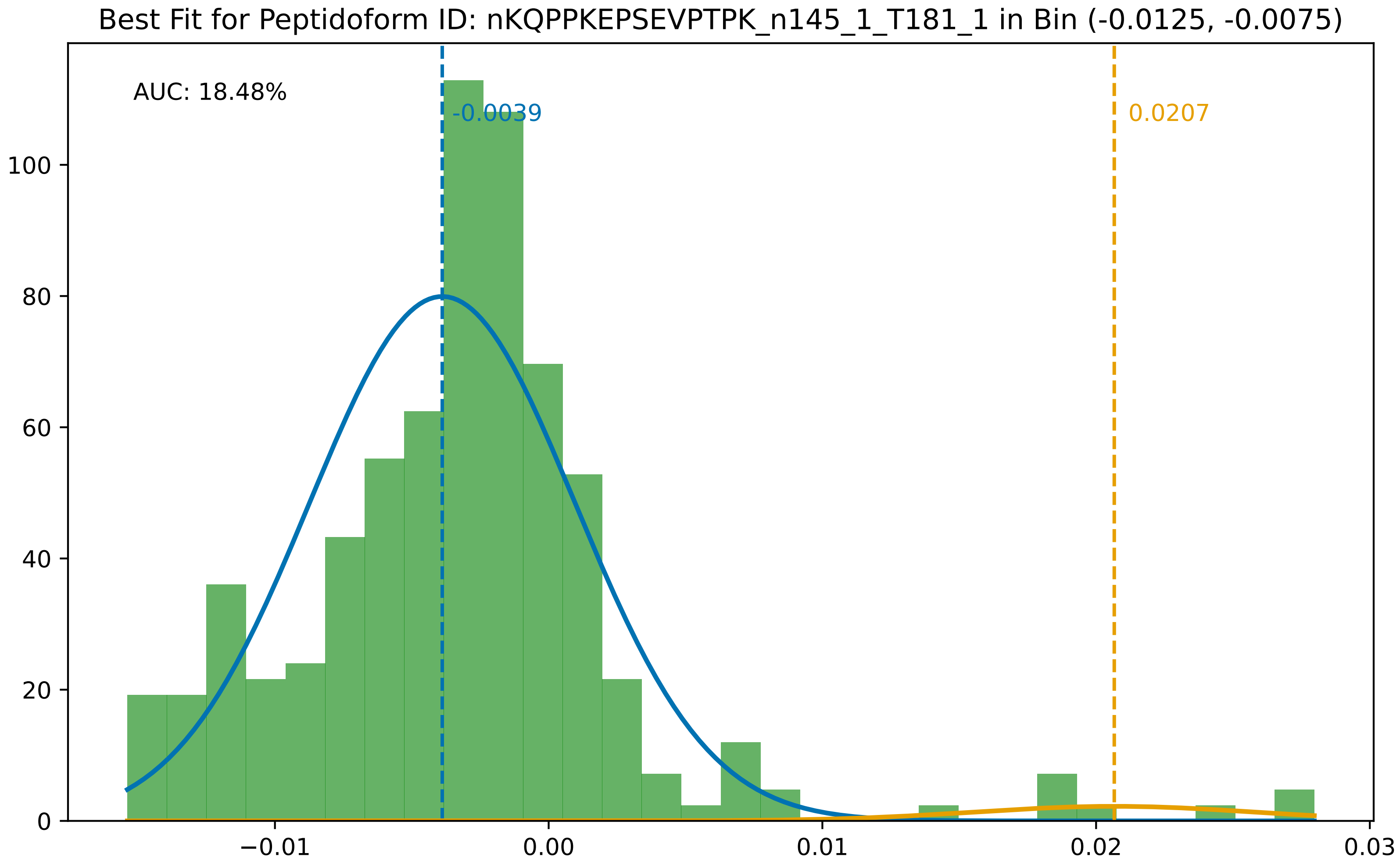

Best Fit for Peptidoform ID: nKQPPKEPSEVPTPK_n145_1_T181_1 in Bin (-0.0125, -0.0075)
AUC: 18.48%
-0.0039
0.0207
100
80
60
40
20
0
−0.01
0.00
0.01
0.02
0.03

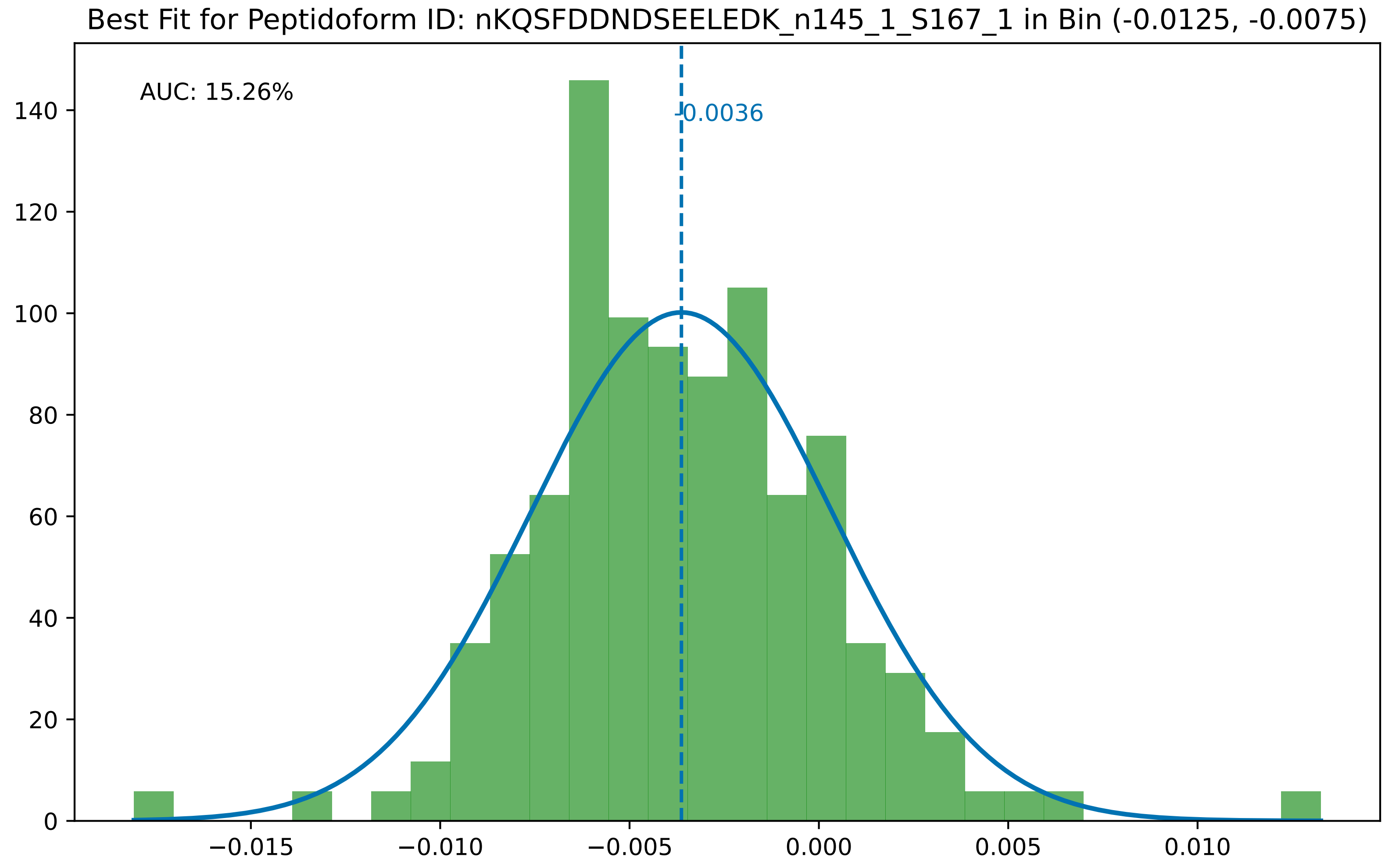
Best Fit for Peptidoform ID: nKQSFDDNDSEELEDK_n145_1_S167_1 in Bin (-0.0125, -0.0075)
AUC: 15.26%
-0.0036
0
20
40
60
80
100
120
140
−0.015
−0.010
−0.005
0.000
0.005
0.010

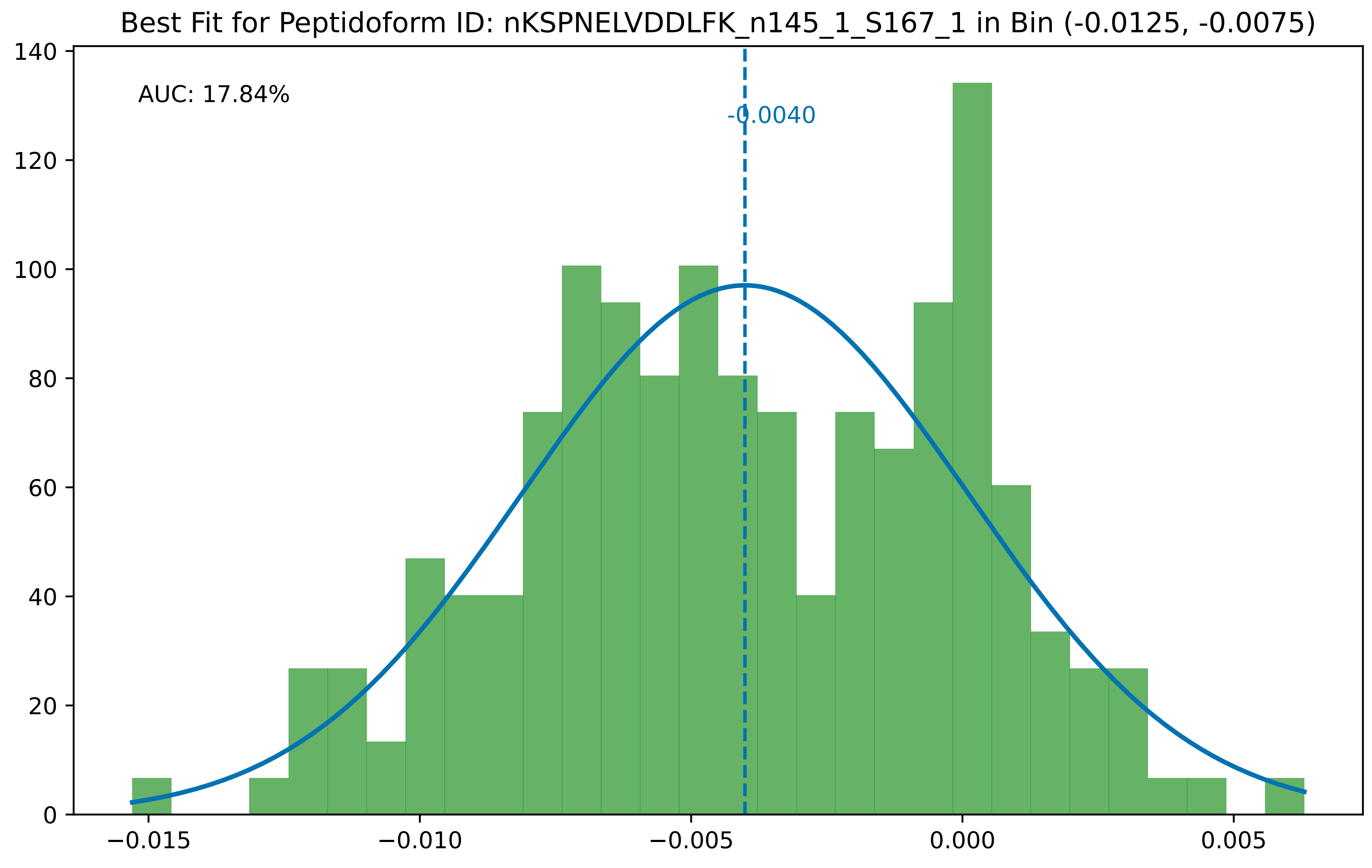

Best Fit for Peptidoform ID: nKSPNELVDDLFK_n145_1_S167_1 in Bin (-0.0125, -0.0075)
AUC: 17.84%
-0.0040
0
20
40
60
80
100
120
140
−0.015
−0.010
−0.005
0.000
0.005

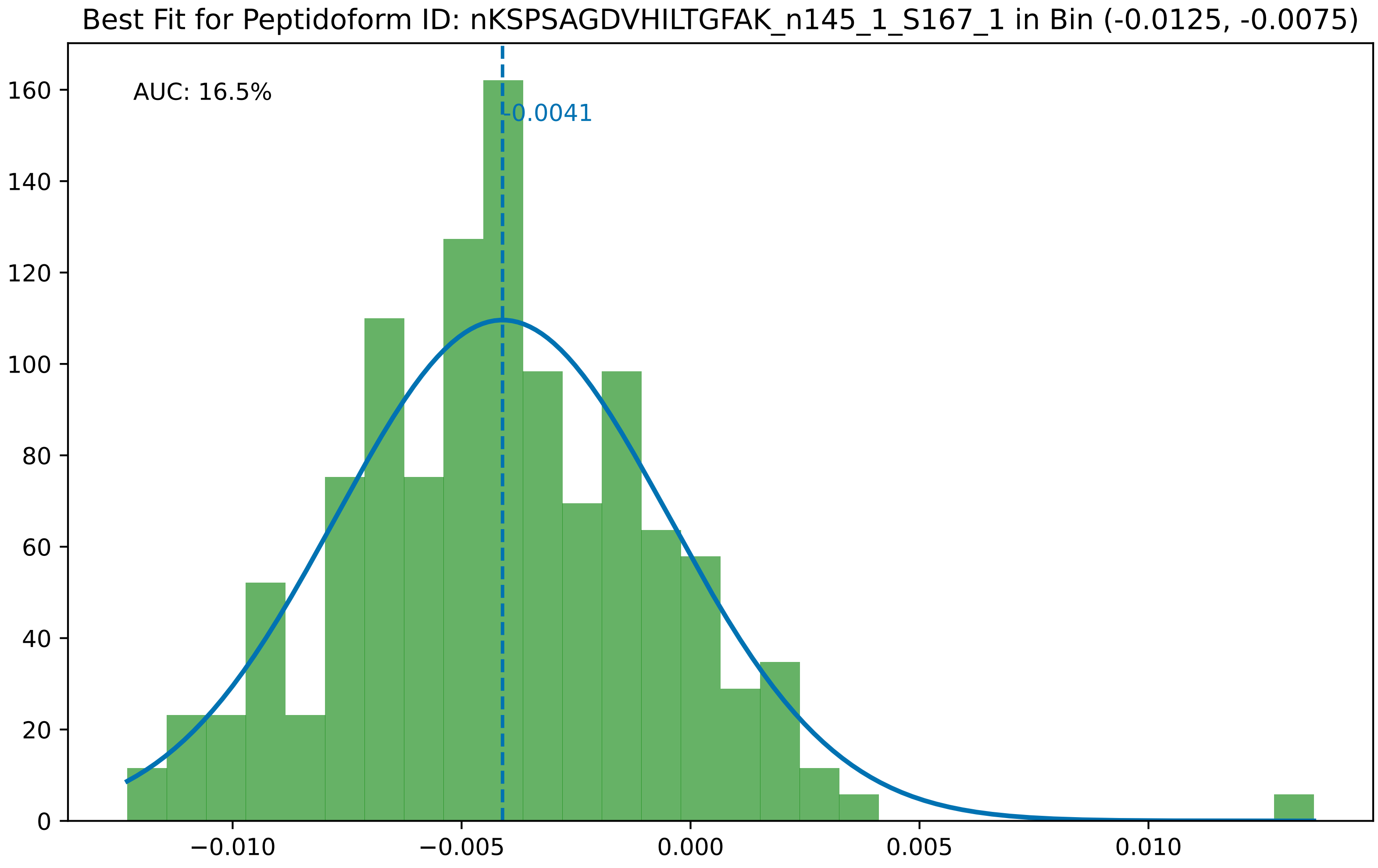

Best Fit for Peptidoform ID: nKSPSAGDVHILTGFAK_n145_1_S167_1 in Bin (-0.0125, -0.0075)
AUC: 16.5%
-0.0041
0
20
40
60
80
100
120
140
160
−0.010
−0.005
0.000
0.005
0.010

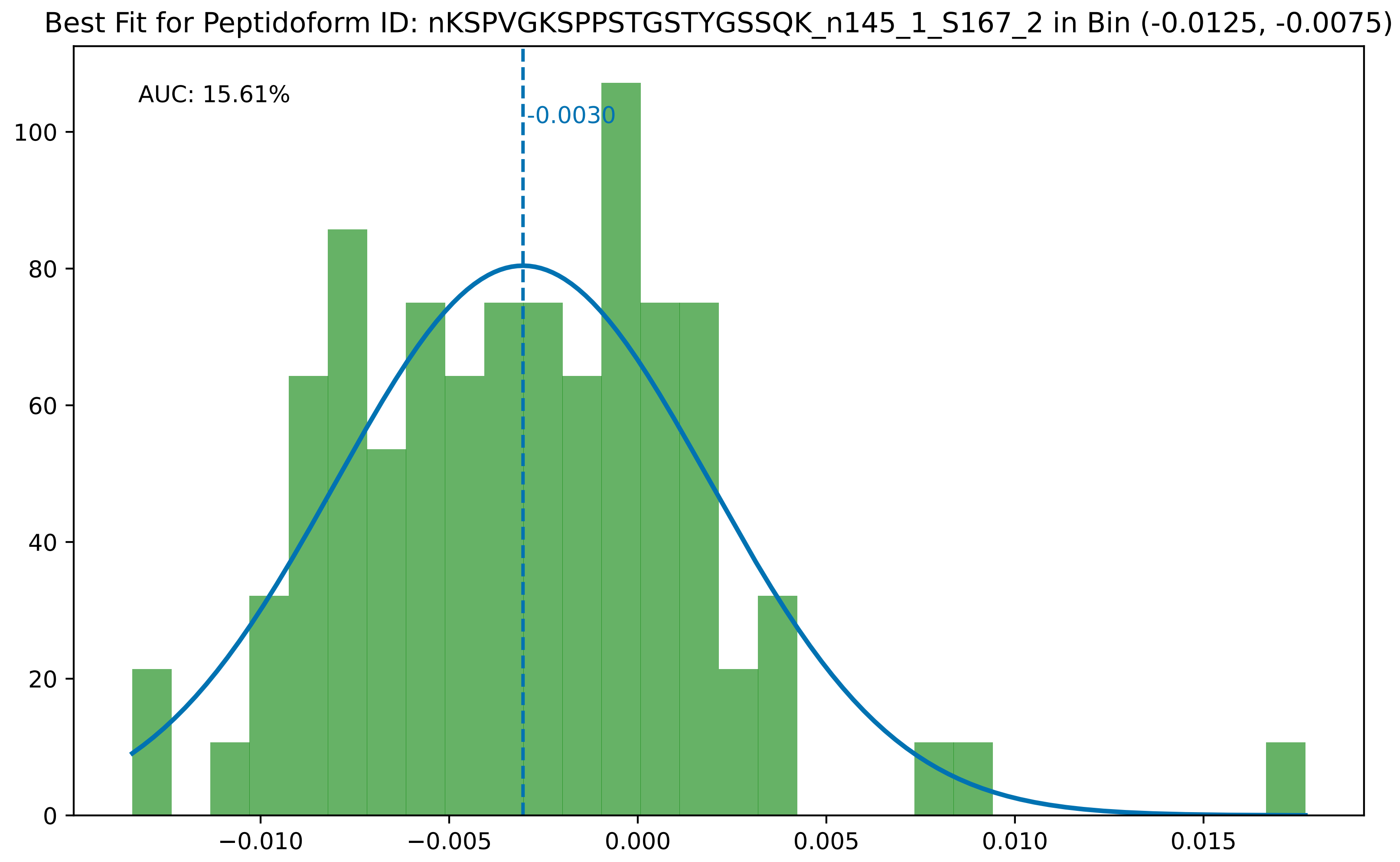

Best Fit for Peptidoform ID: nKSPVGKSPPSTGSTYGSSQK_n145_1_S167_2 in Bin (-0.0125, -0.0075)
AUC: 15.61%
-0.0030
0
20
40
60
80
100
−0.010
−0.005
0.000
0.005
0.010
0.015

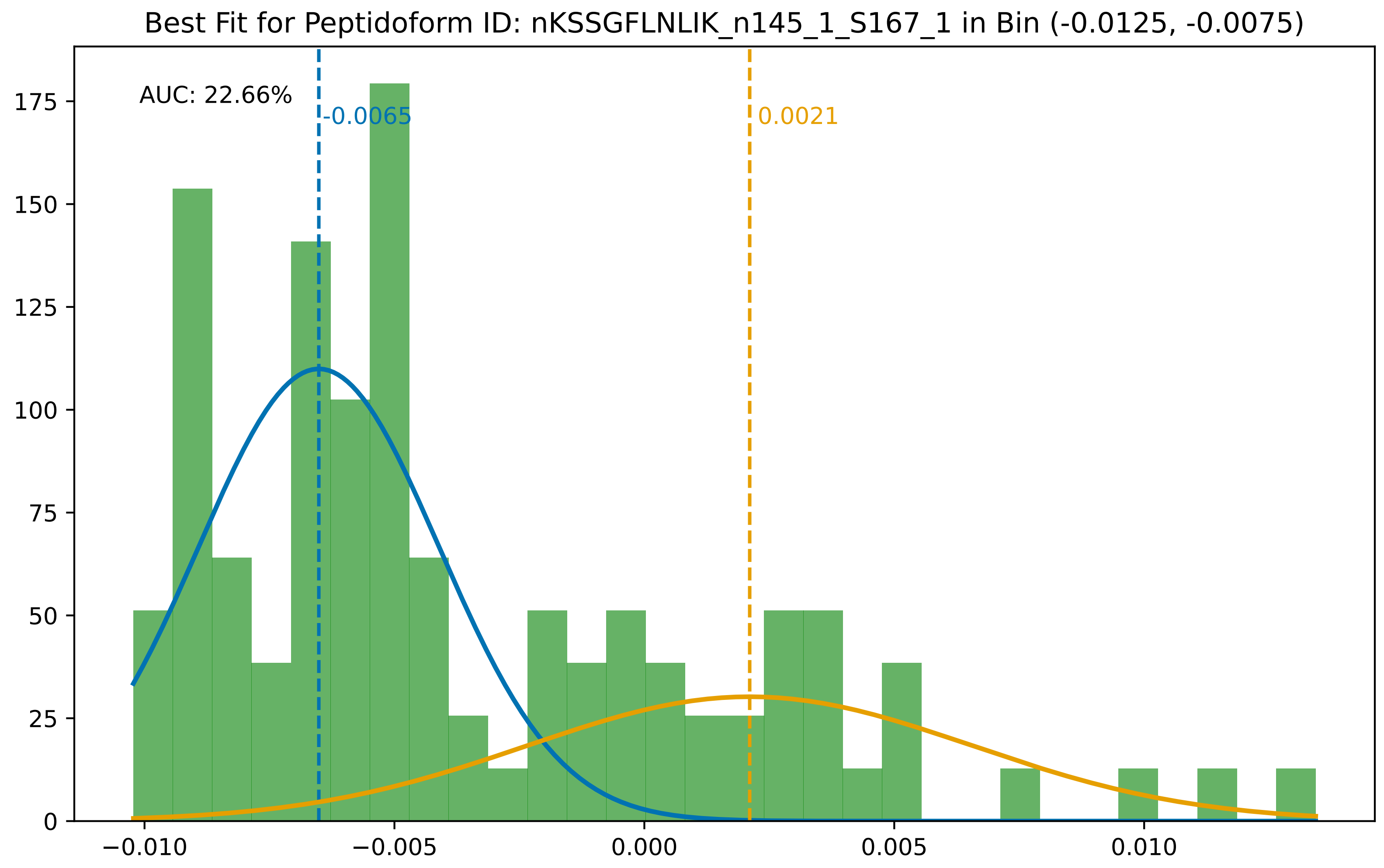
Best Fit for Peptidoform ID: nKSSGFLNLIK_n145_1_S167_1 in Bin (-0.0125, -0.0075)
AUC: 22.66%
-0.0065
0.0021
0
25
50
75
100
125
150
175
−0.010
−0.005
0.000
0.005
0.010

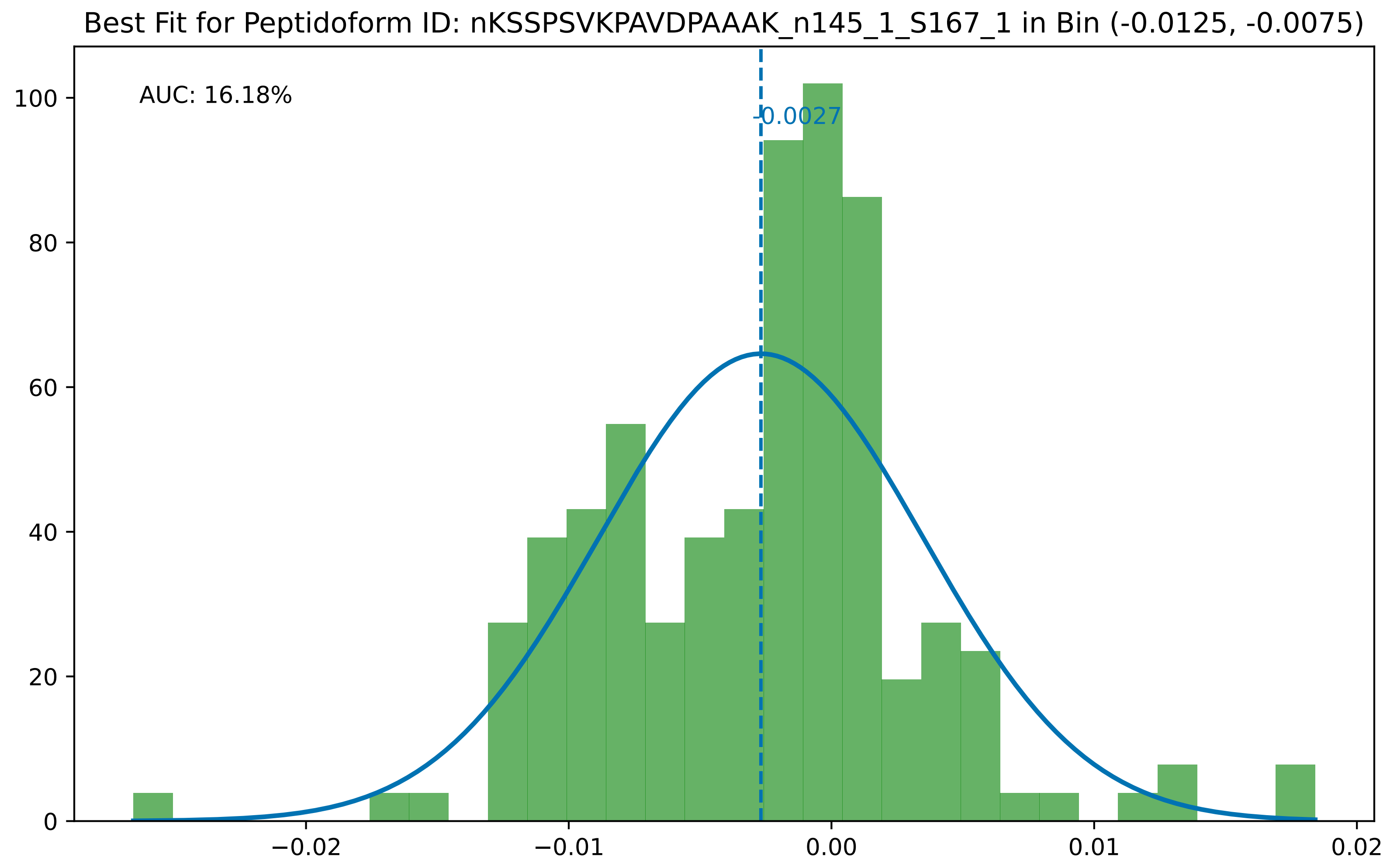

Best Fit for Peptidoform ID: nKSSPSVKPAVDPAAAK_n145_1_S167_1 in Bin (-0.0125, -0.0075)
AUC: 16.18%
-0.0027
0
20
40
60
80
100
−0.02
−0.01
0.00
0.01
0.02

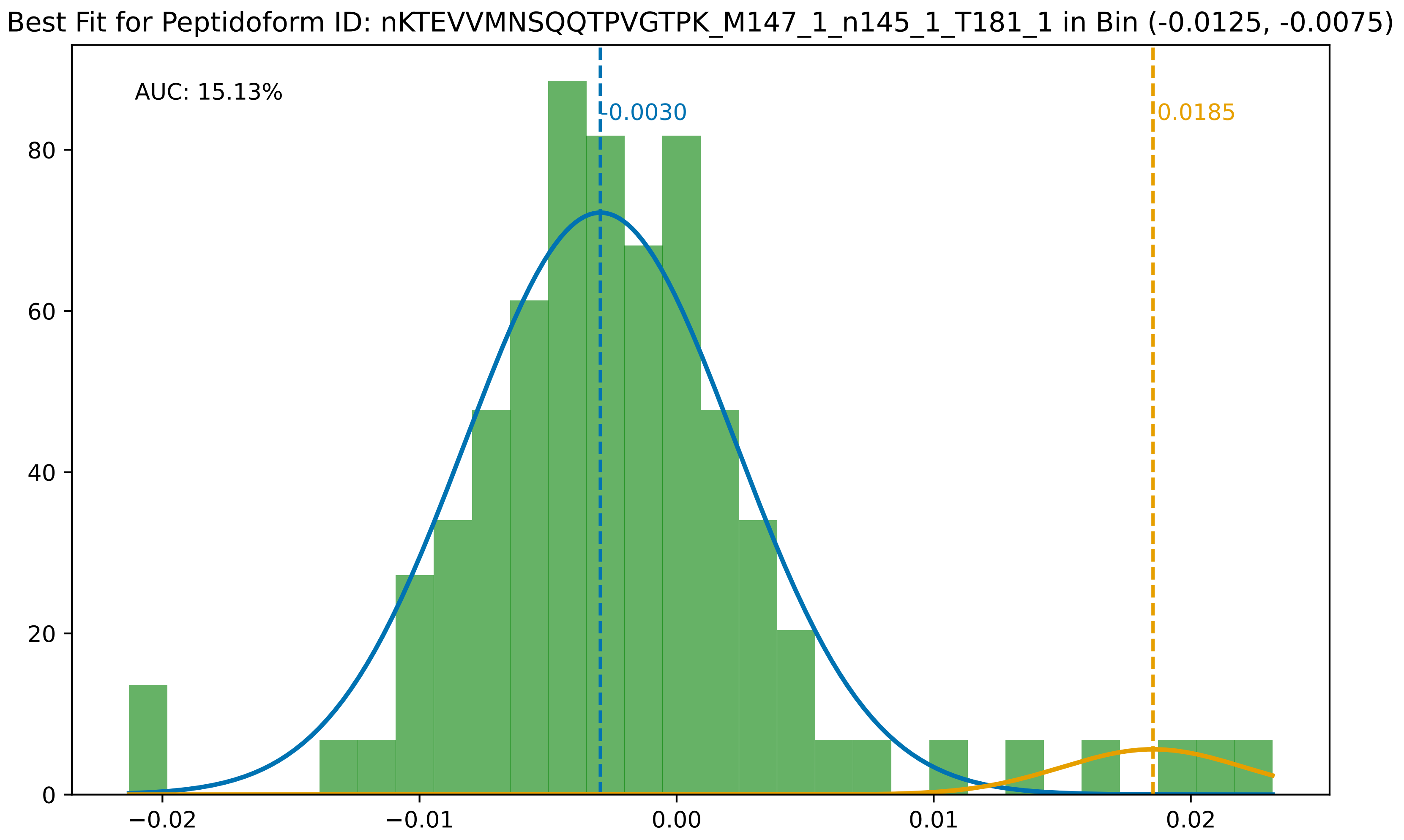
Best Fit for Peptidoform ID: nKTEVVMNSQQTPVGTPK_M147_1_n145_1_T181_1 in Bin (-0.0125, -0.0075)
AUC: 15.13%
-0.0030
0.0185
0
20
40
60
80
−0.02
−0.01
0.00
0.01
0.02

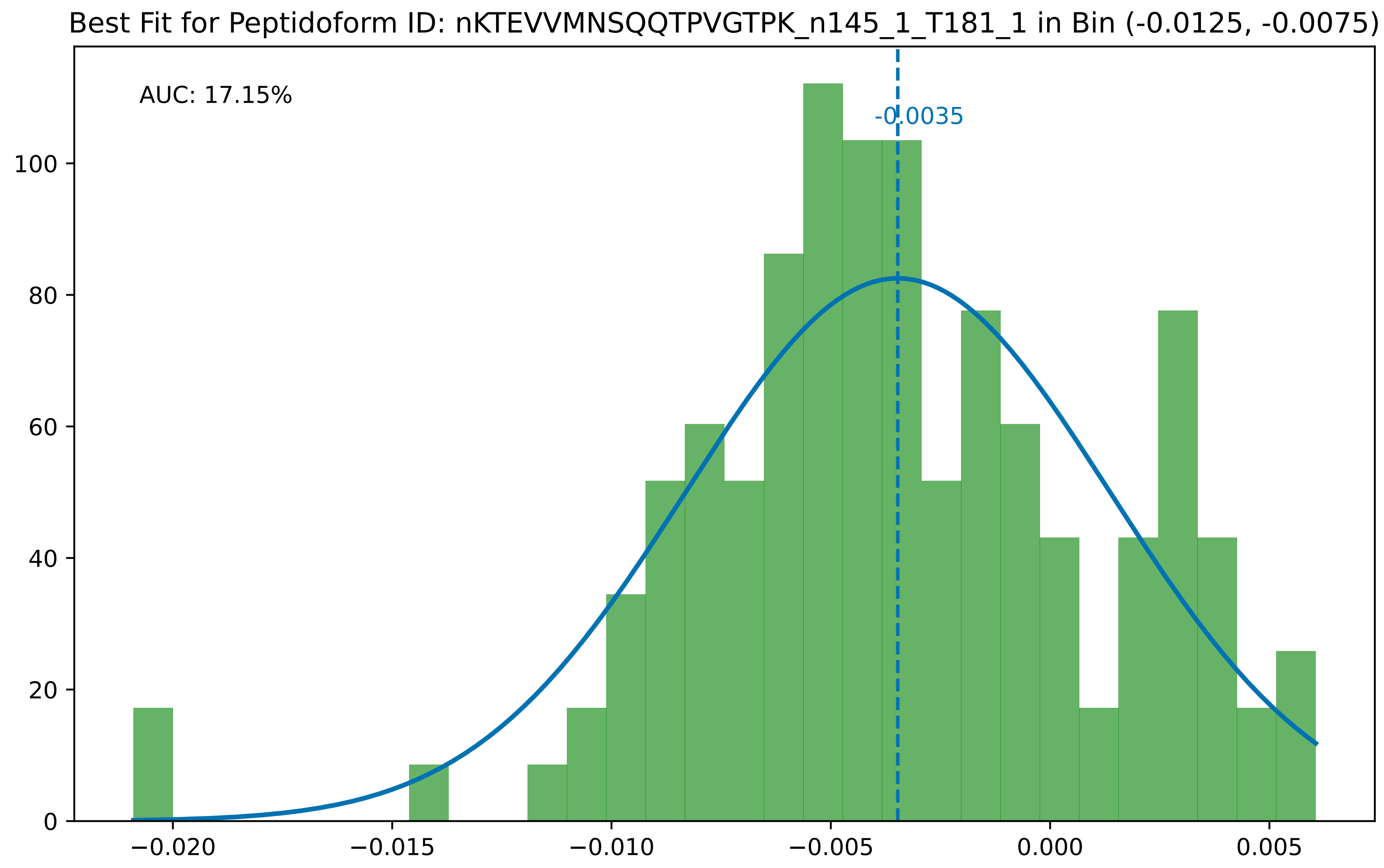
Best Fit for Peptidoform ID: nKTEVVMNSQQTPVGTPK_n145_1_T181_1 in Bin (-0.0125, -0.0075)
AUC: 17.15%
-0.0035
0
20
40
60
80
100
−0.020
−0.015
−0.010
−0.005
0.000
0.005

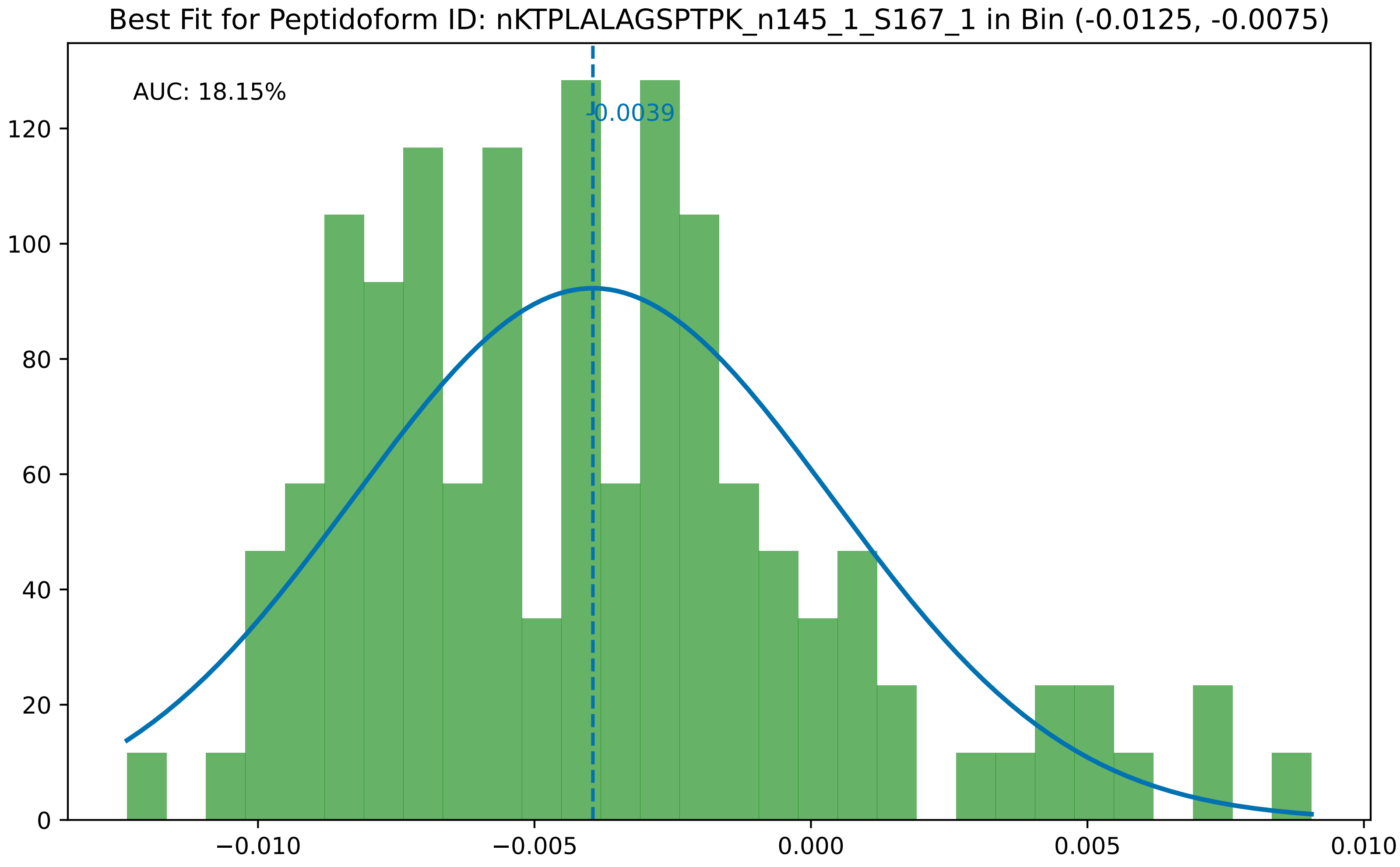

Best Fit for Peptidoform ID: nKTPLALAGSPTPK_n145_1_S167_1 in Bin (-0.0125, -0.0075)
AUC: 18.15%
-0.0039
0
20
40
60
80
100
120
−0.010
−0.005
0.000
0.005
0.010

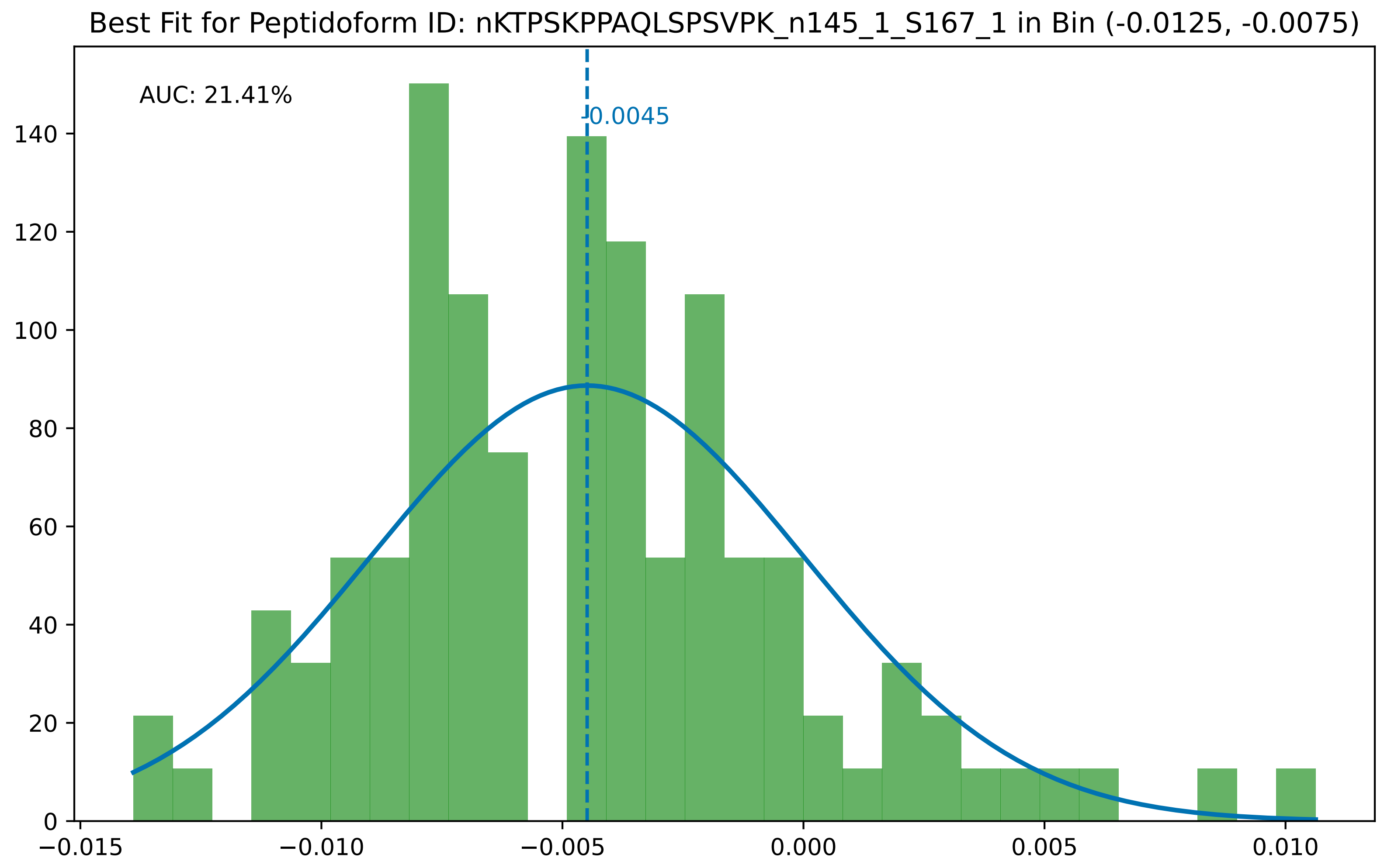
Best Fit for Peptidoform ID: nKTPSKPPAQLSPSVPK_n145_1_S167_1 in Bin (-0.0125, -0.0075)
AUC: 21.41%
-0.0045
0
20
40
60
80
100
120
140
−0.015
−0.010
−0.005
0.000
0.005
0.010

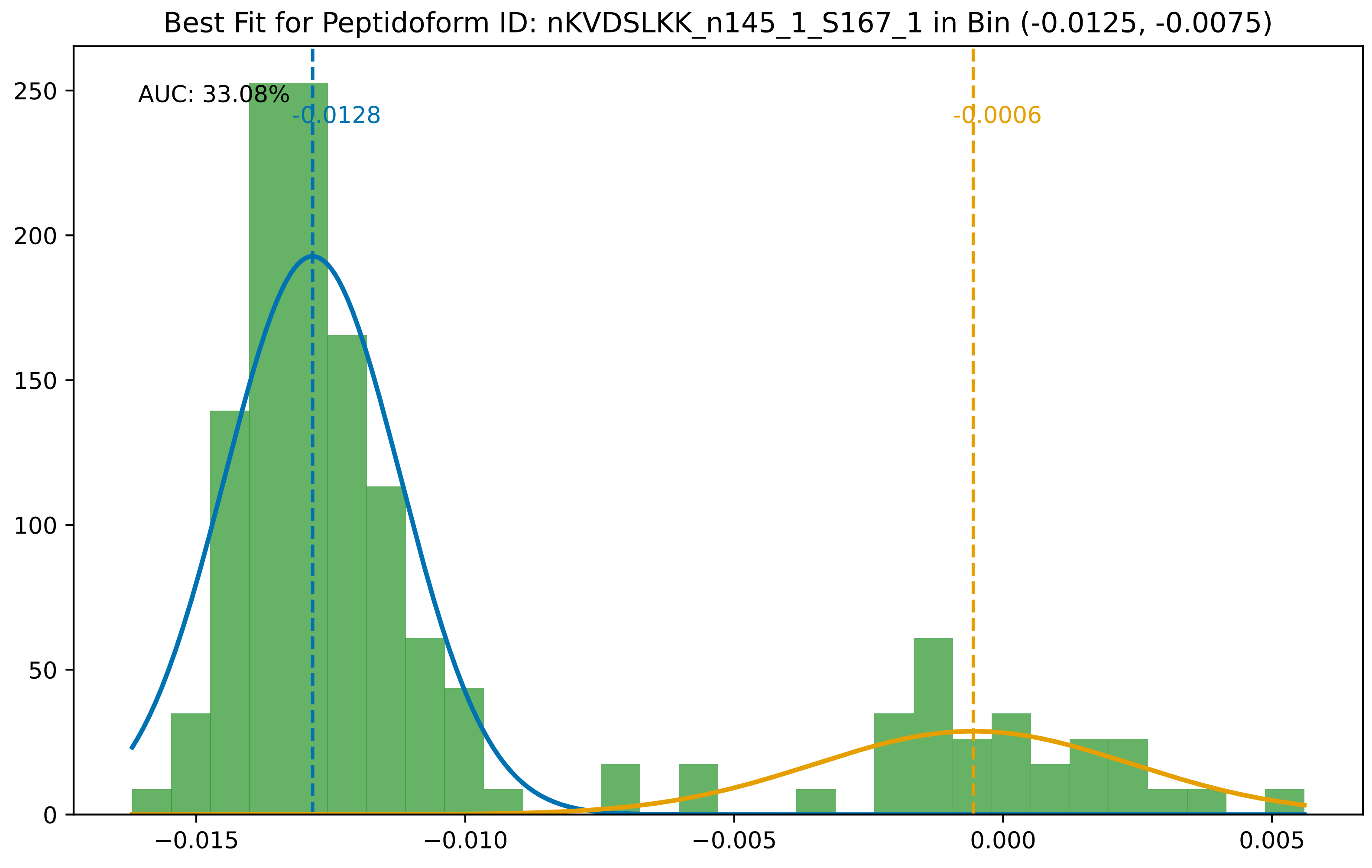

Best Fit for Peptidoform ID: nKVDSLKK_n145_1_S167_1 in Bin (-0.0125, -0.0075)
AUC: 33.08%
-0.0128
-0.0006
0
50
100
150
200
250
−0.015
−0.010
−0.005
0.000
0.005

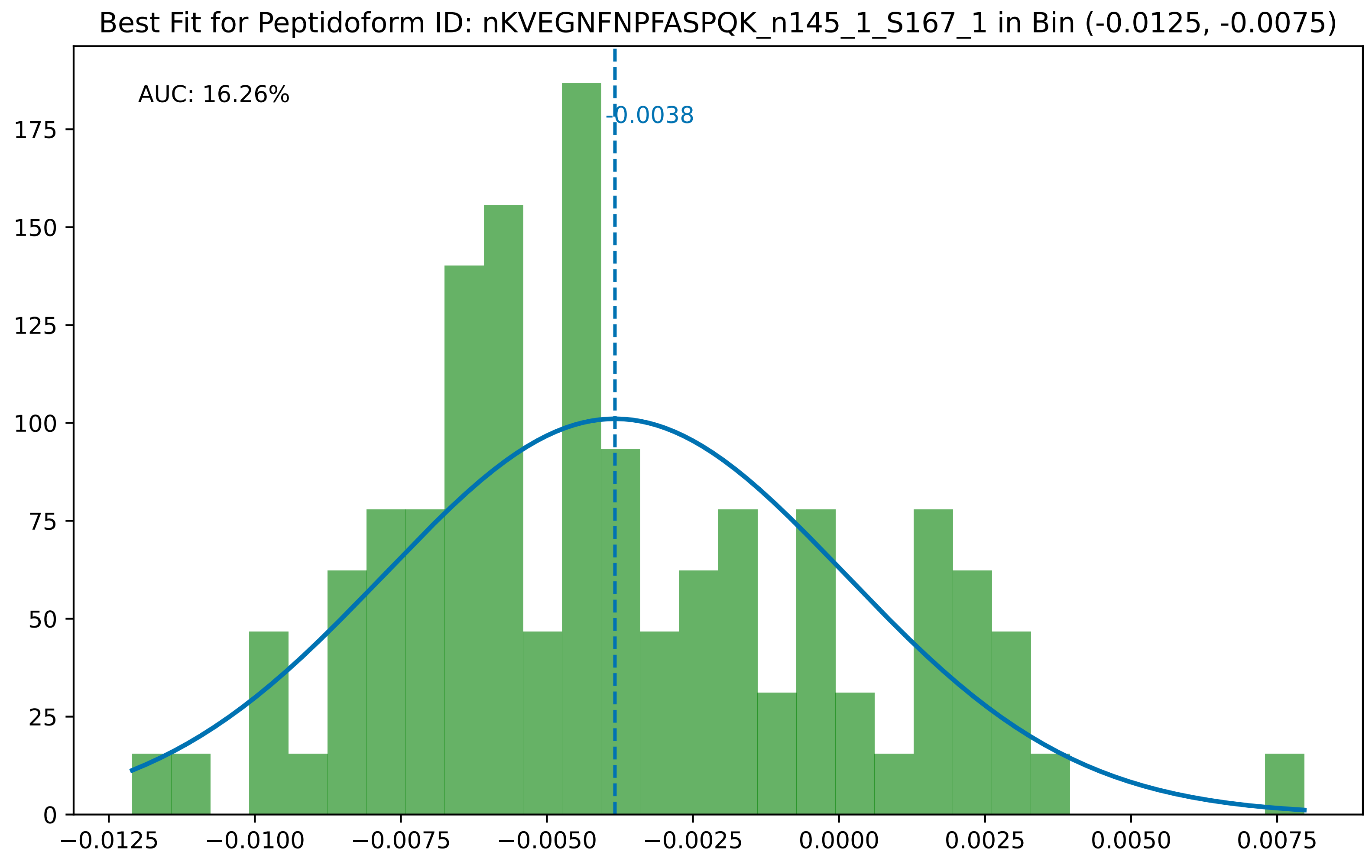

Best Fit for Peptidoform ID: nKVEGNFNPFASPQK_n145_1_S167_1 in Bin (-0.0125, -0.0075)
AUC: 16.26%
-0.0038
0
25
50
75
100
125
150
175
−0.0125
−0.0100
−0.0075
−0.0050
−0.0025
0.0000
0.0025
0.0050
0.0075

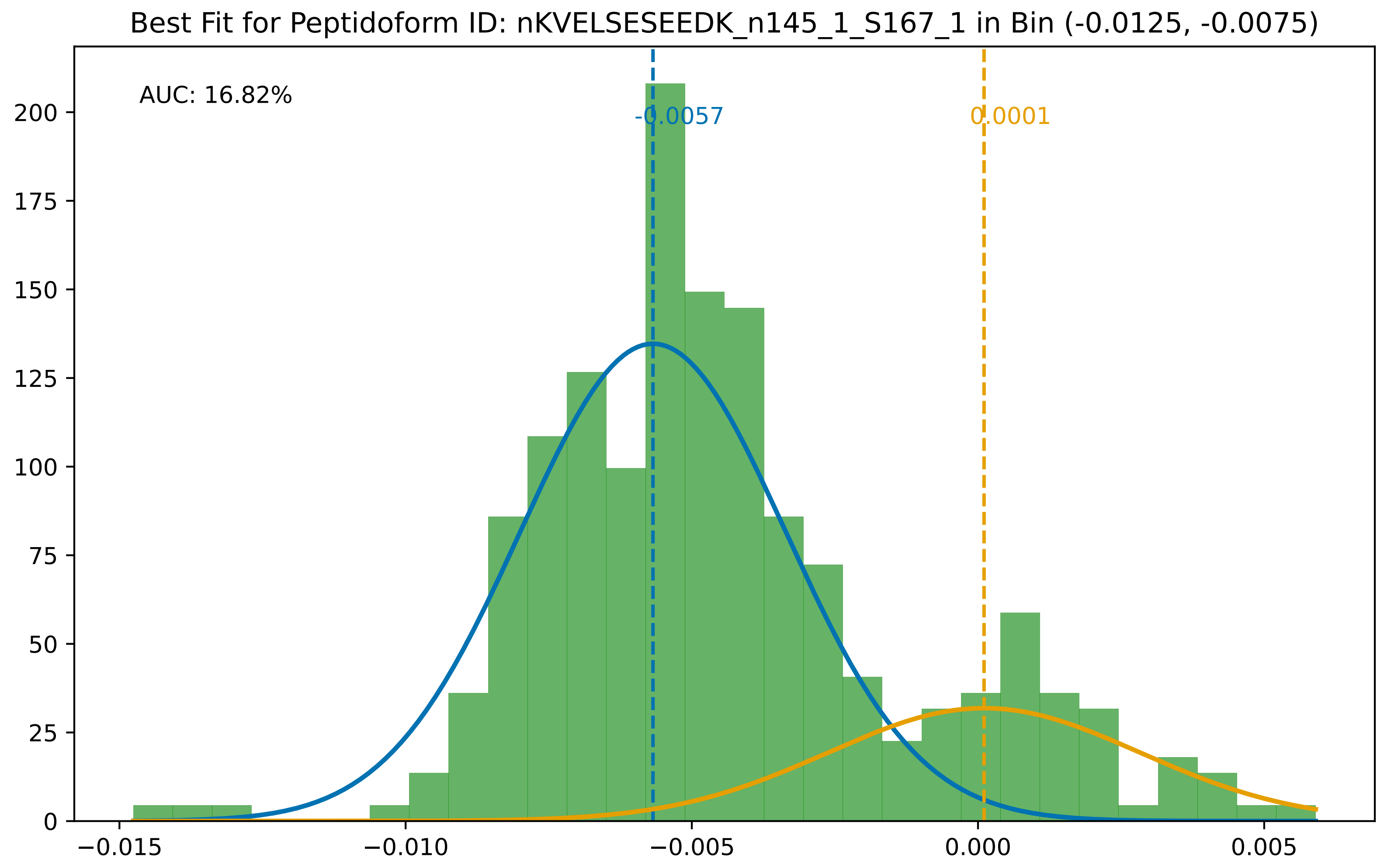
Best Fit for Peptidoform ID: nKVELSESEEDK_n145_1_S167_1 in Bin (-0.0125, -0.0075)
AUC: 16.82%
-0.0057
0.0001
0
25
50
75
100
125
150
175
200
−0.015
−0.010
−0.005
0.000
0.005

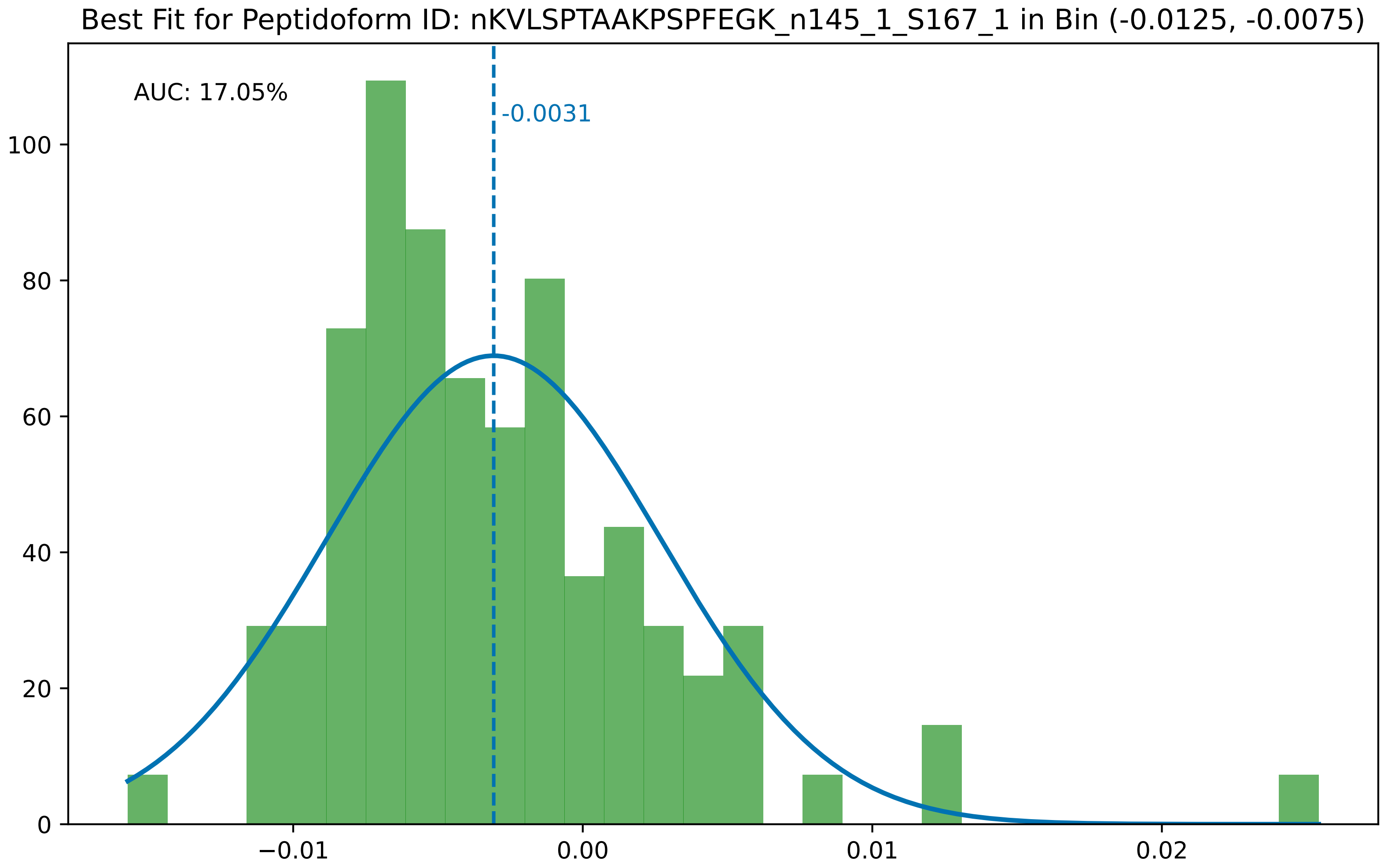
Best Fit for Peptidoform ID: nKVLSPTAAKPSPFEGK_n145_1_S167_1 in Bin (-0.0125, -0.0075)
AUC: 17.05%
-0.0031
0
20
40
60
80
100
−0.01
0.00
0.01
0.02

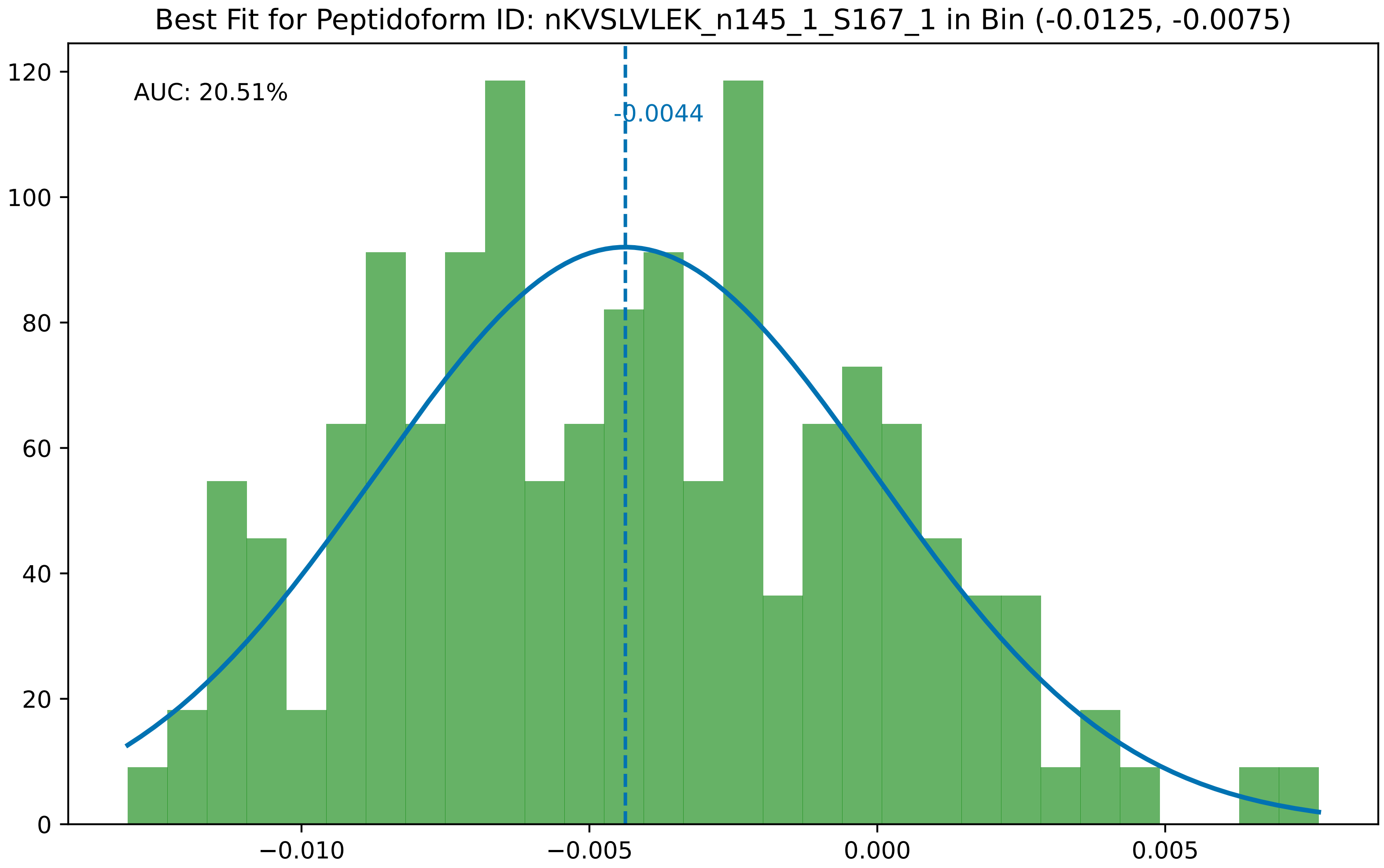

Best Fit for Peptidoform ID: nKVSLVLEK_n145_1_S167_1 in Bin (-0.0125, -0.0075)
AUC: 20.51%
-0.0044
120
100
80
60
40
20
0
−0.010
−0.005
0.000
0.005

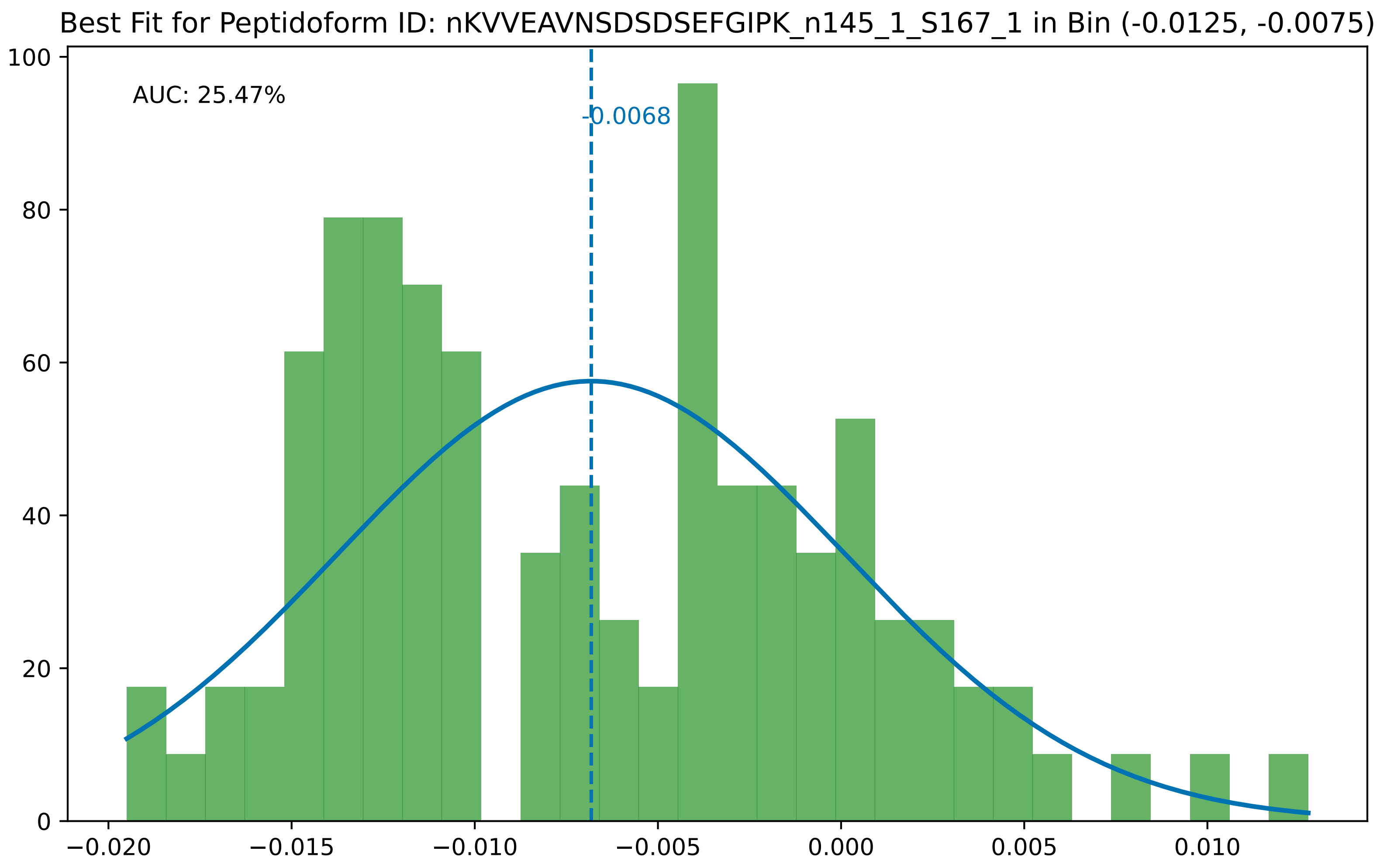

Best Fit for Peptidoform ID: nKVVEAVNSDSDSEFGIPK_n145_1_S167_1 in Bin (-0.0125, -0.0075)
AUC: 25.47%
-0.0068
0
20
40
60
80
100
−0.020
−0.015
−0.010
−0.005
0.000
0.005
0.010

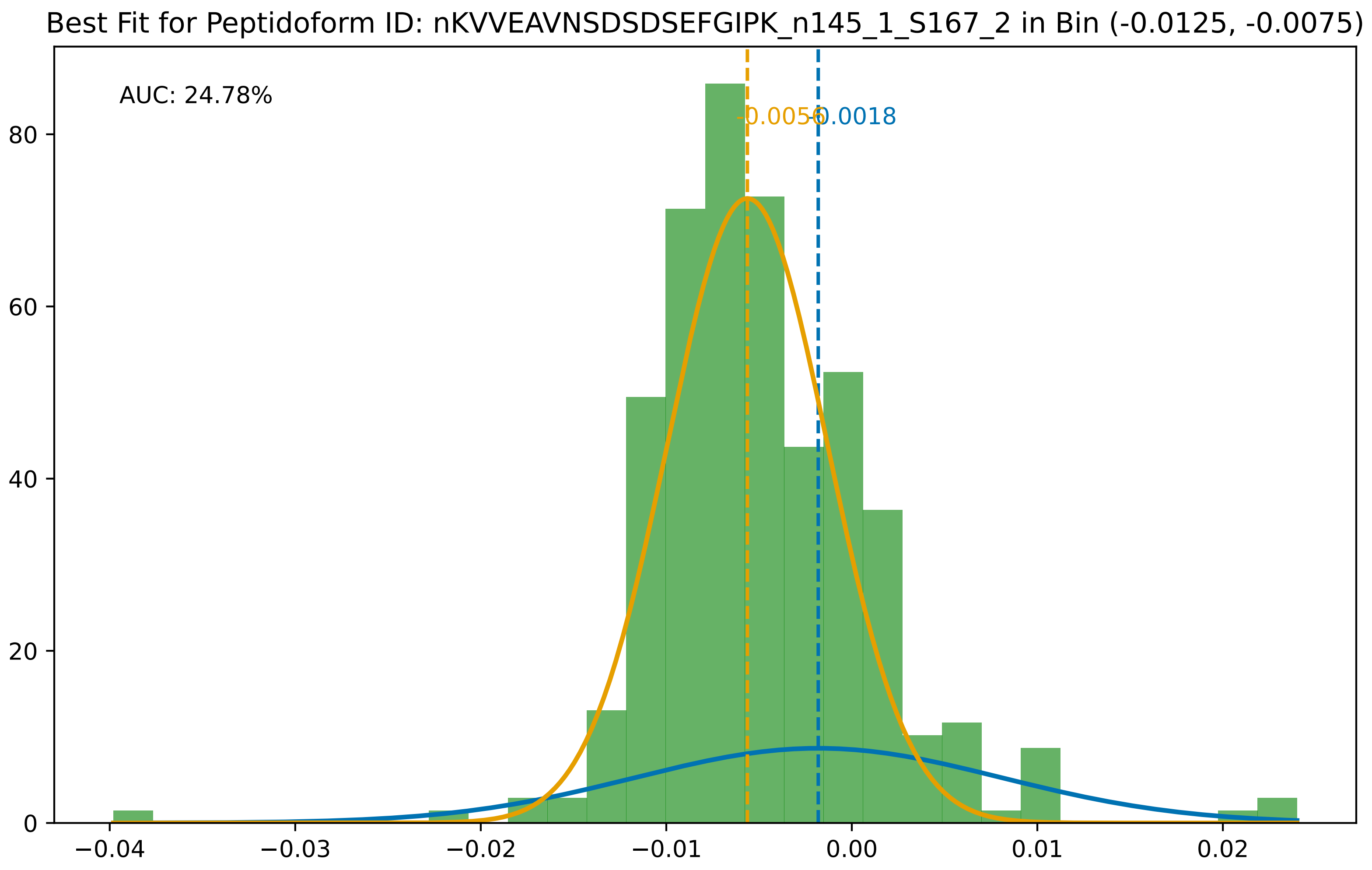
Best Fit for Peptidoform ID: nKVVEAVNSDSDSEFGIPK_n145_1_S167_2 in Bin (-0.0125, -0.0075)
AUC: 24.78%
-0.0056
-0.0018
0
20
40
60
80
−0.04
−0.03
−0.02
−0.01
0.00
0.01
0.02

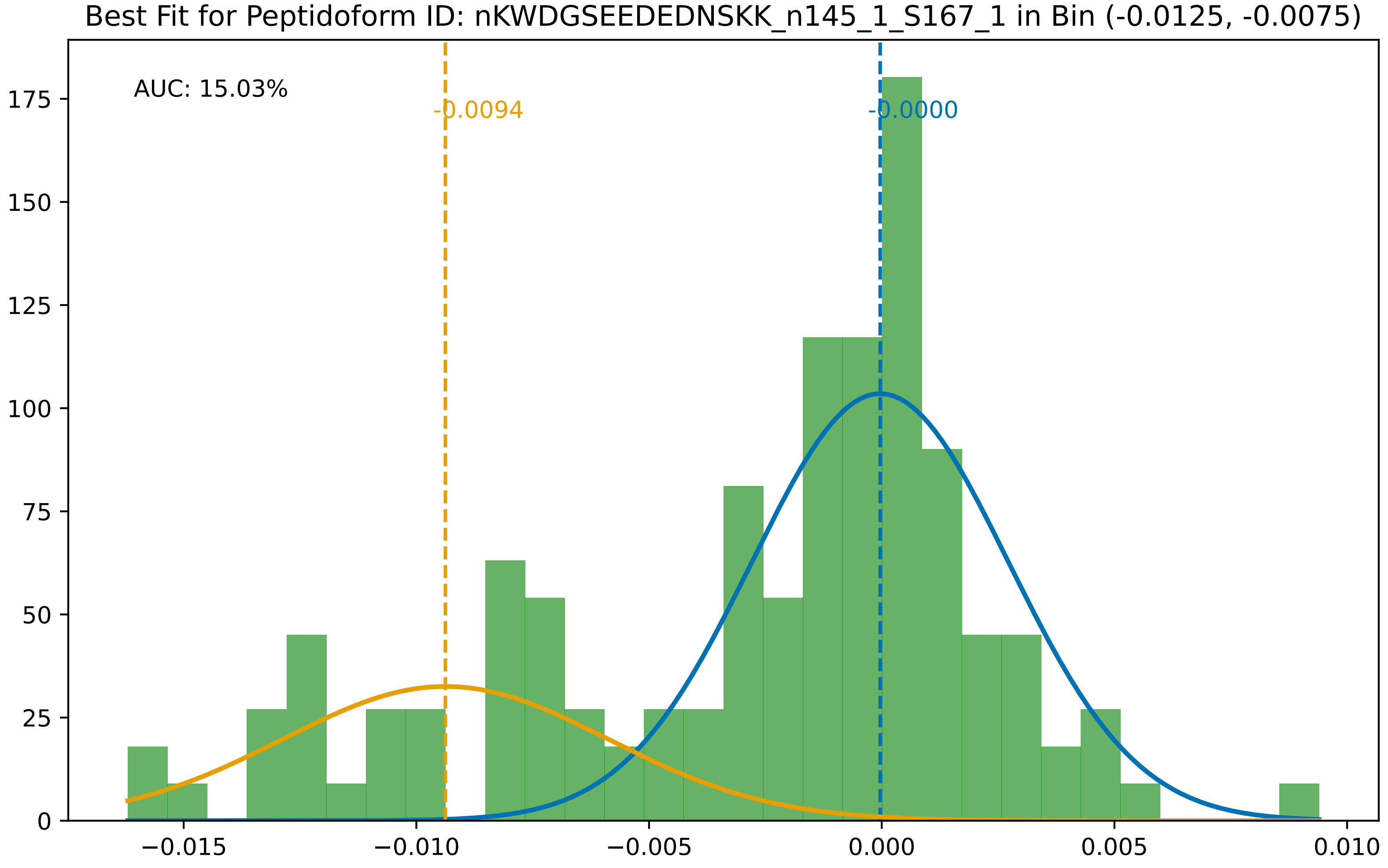

Best Fit for Peptidoform ID: nKWDGSEEDEDNSKK_n145_1_S167_1 in Bin (-0.0125, -0.0075)
AUC: 15.03%
-0.0094
-0.0000
0
25
50
75
100
125
150
175
−0.015
−0.010
−0.005
0.000
0.005
0.010

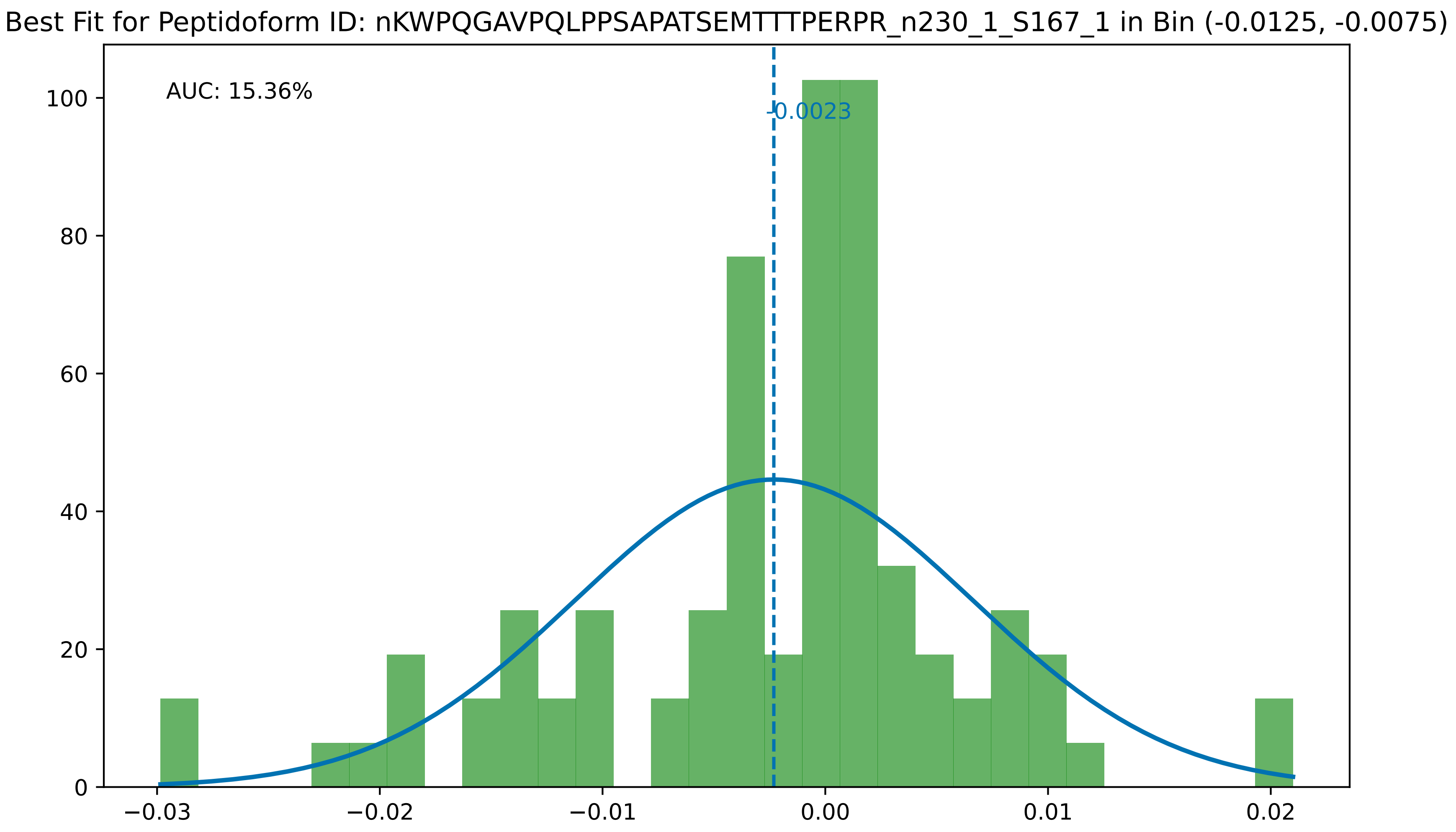

Best Fit for Peptidoform ID: nKWPQGAVPQLPPSAPATSEMTTTPERPR_n230_1_S167_1 in Bin (-0.0125, -0.0075)
AUC: 15.36%
-0.0023
0
20
40
60
80
100
−0.03
−0.02
−0.01
0.00
0.01
0.02

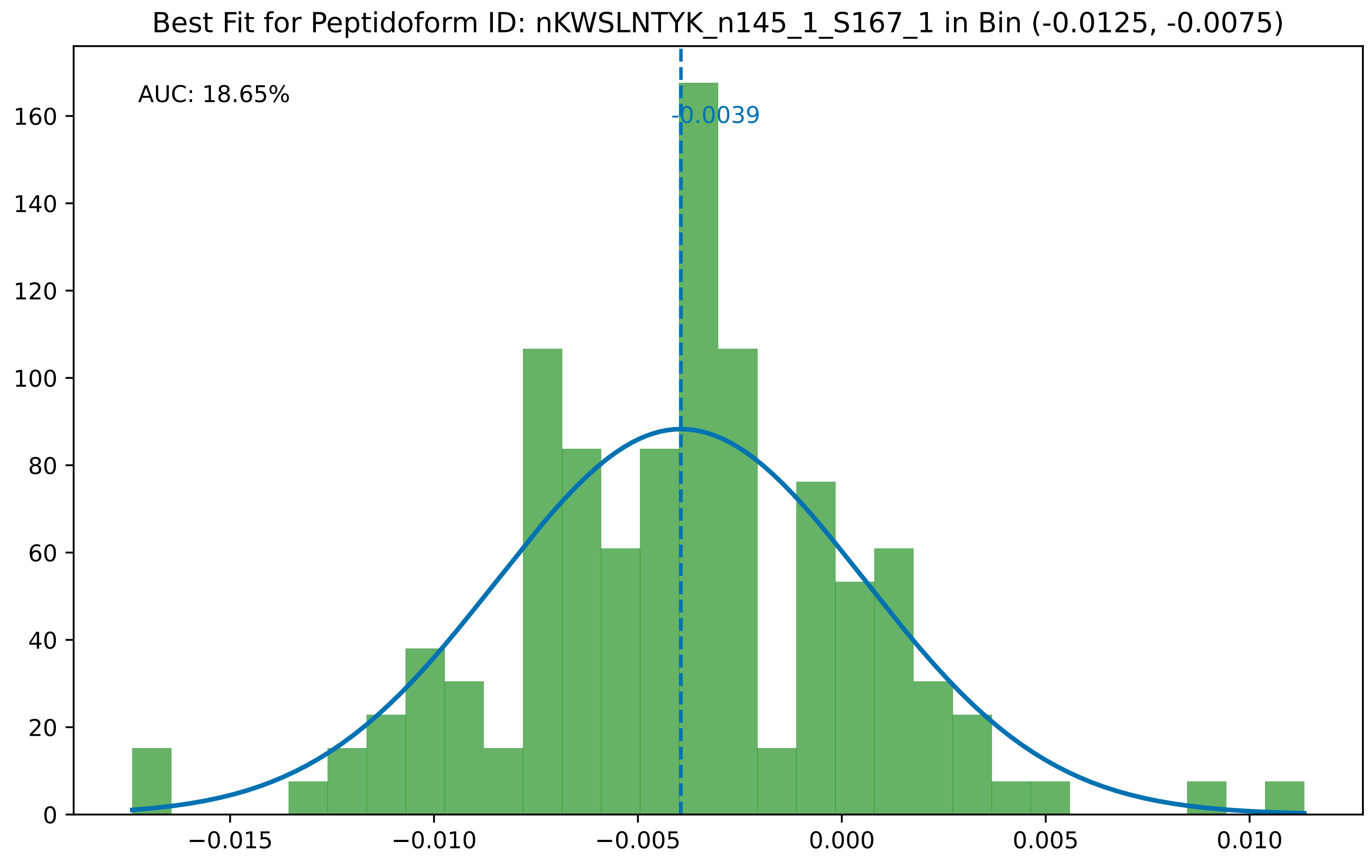

Best Fit for Peptidoform ID: nKWSLNTYK_n145_1_S167_1 in Bin (-0.0125, -0.0075)
AUC: 18.65%
-0.0039
0
20
40
60
80
100
120
140
160
−0.015
−0.010
−0.005
0.000
0.005
0.010

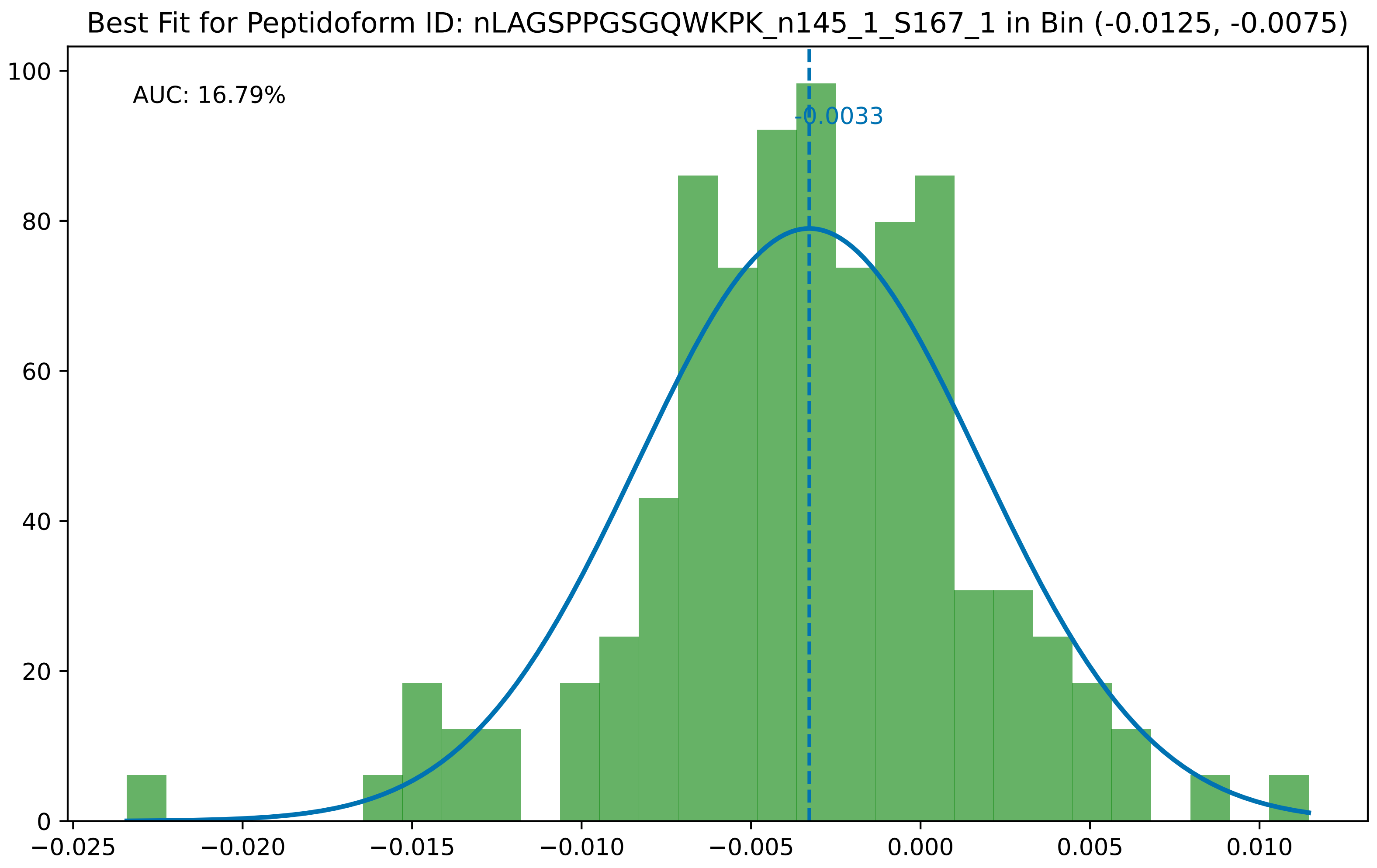

Best Fit for Peptidoform ID: nLAGSPPGSGQWKPK_n145_1_S167_1 in Bin (-0.0125, -0.0075)
AUC: 16.79%
-0.0033
0
20
40
60
80
100
−0.025
−0.020
−0.015
−0.010
−0.005
0.000
0.005
0.010

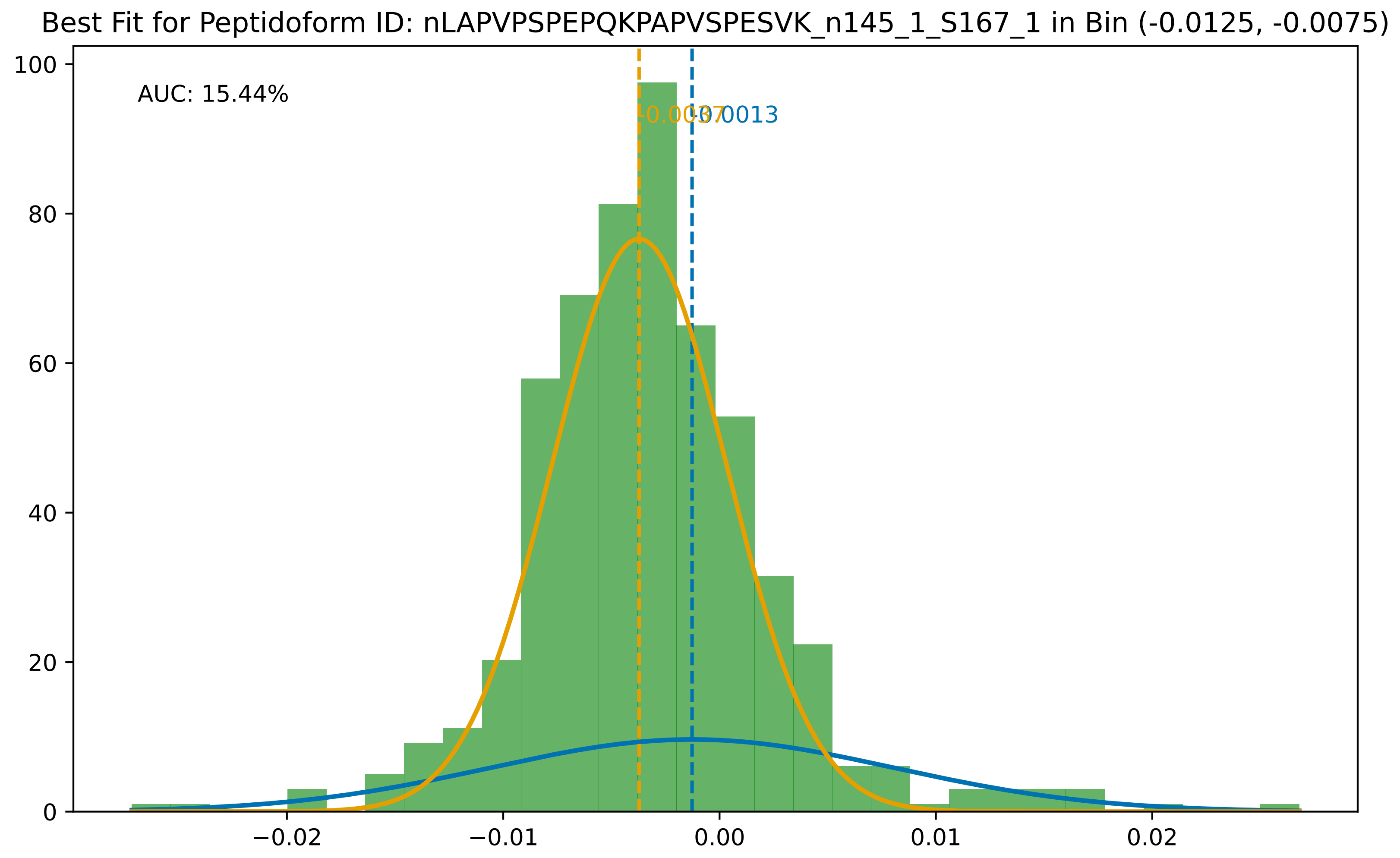

Best Fit for Peptidoform ID: nLAPVPSPEPQKPAPVSPESVK_n145_1_S167_1 in Bin (-0.0125, -0.0075)
AUC: 15.44%
-0.0037
-0.0013
100
80
60
40
20
0
−0.02
−0.01
0.00
0.01
0.02

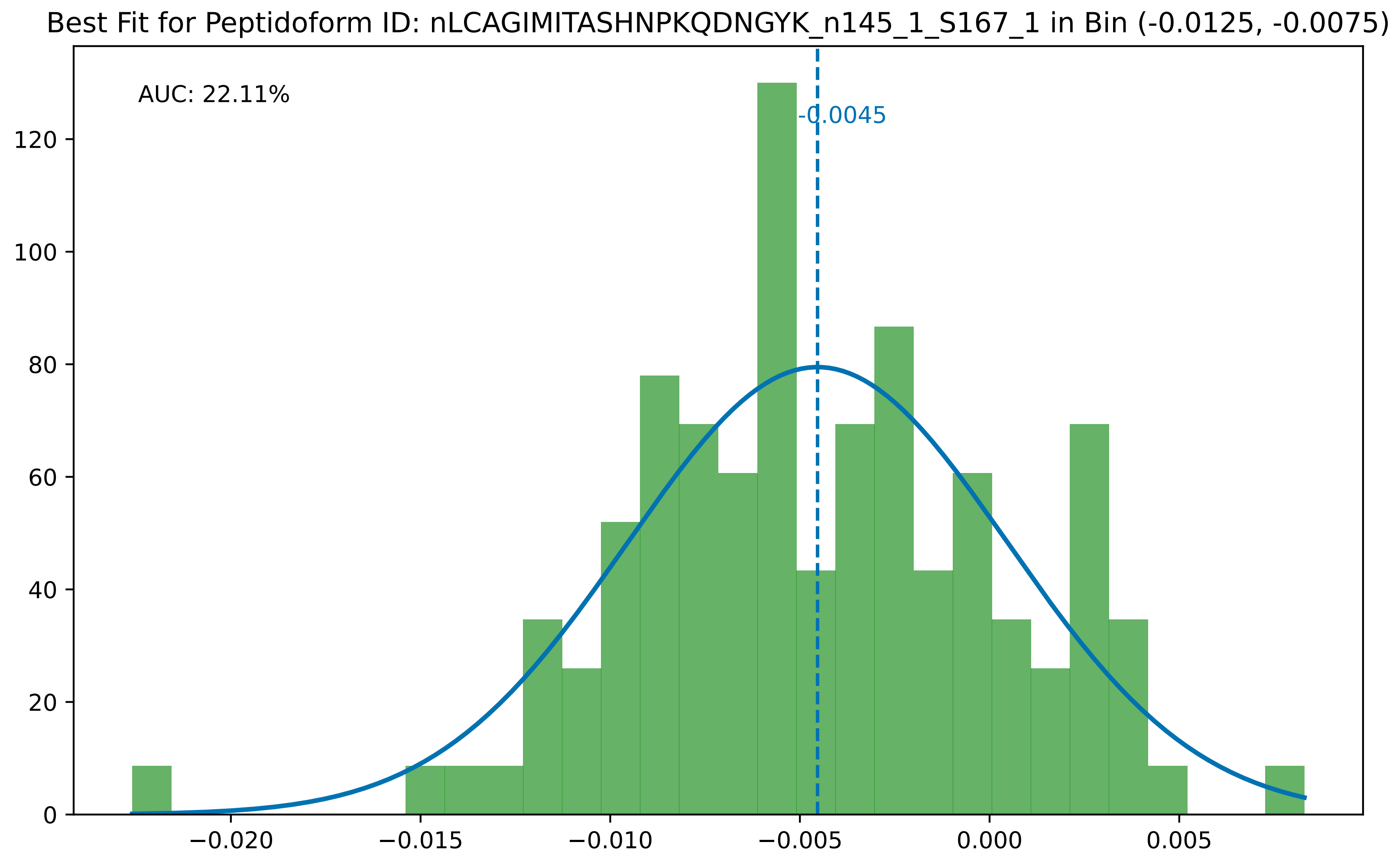
Best Fit for Peptidoform ID: nLCAGIMITASHNPKQDNGYK_n145_1_S167_1 in Bin (-0.0125, -0.0075)
AUC: 22.11%
-0.0045
0
20
40
60
80
100
120
−0.020
−0.015
−0.010
−0.005
0.000
0.005

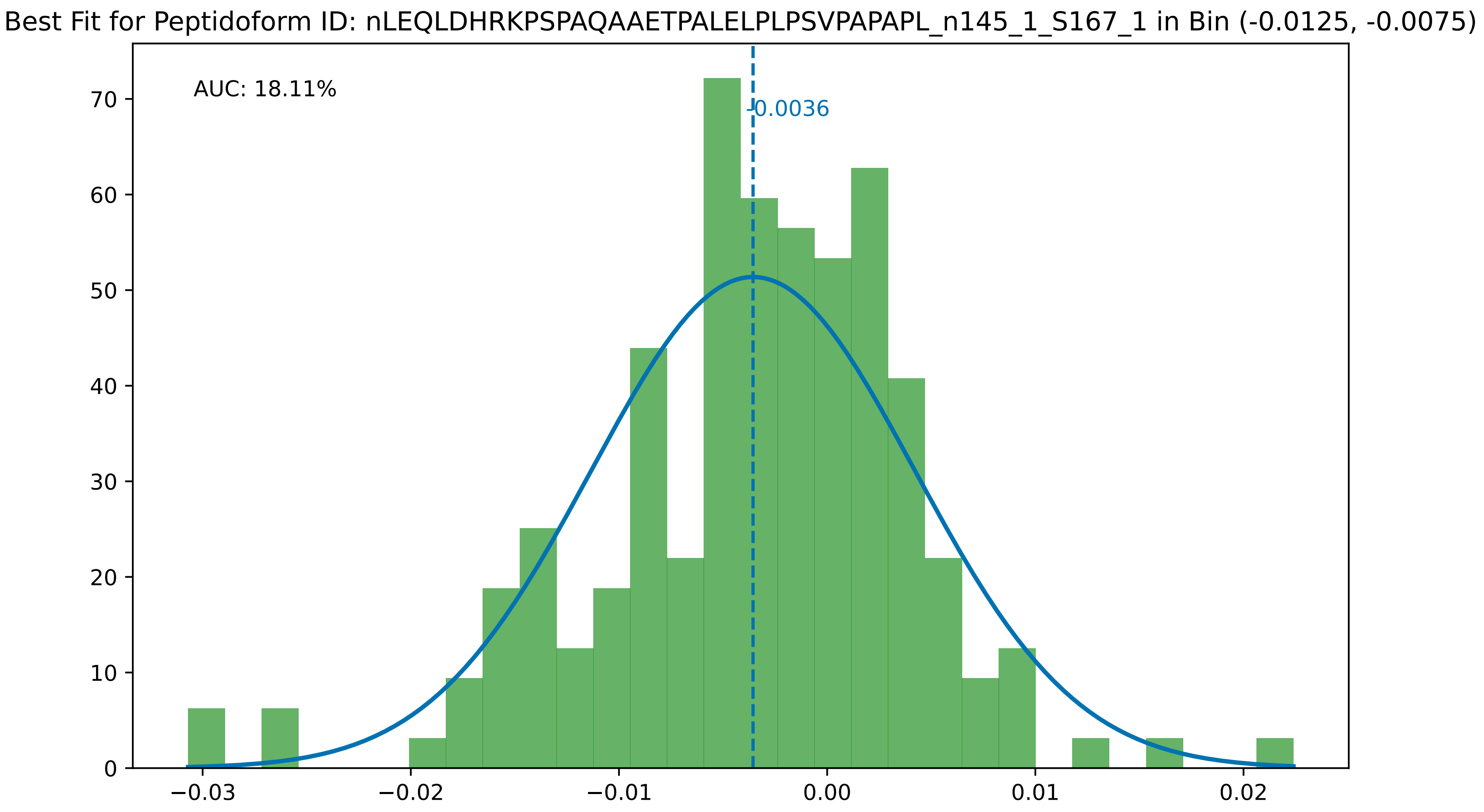

Best Fit for Peptidoform ID: nLEQLDHRKPSPAQAAETPALELPLPSVPAPAPL_n145_1_S167_1 in Bin (-0.0125, -0.0075)
AUC: 18.11%
-0.0036
0
10
20
30
40
50
60
70
−0.03
−0.02
−0.01
0.00
0.01
0.02

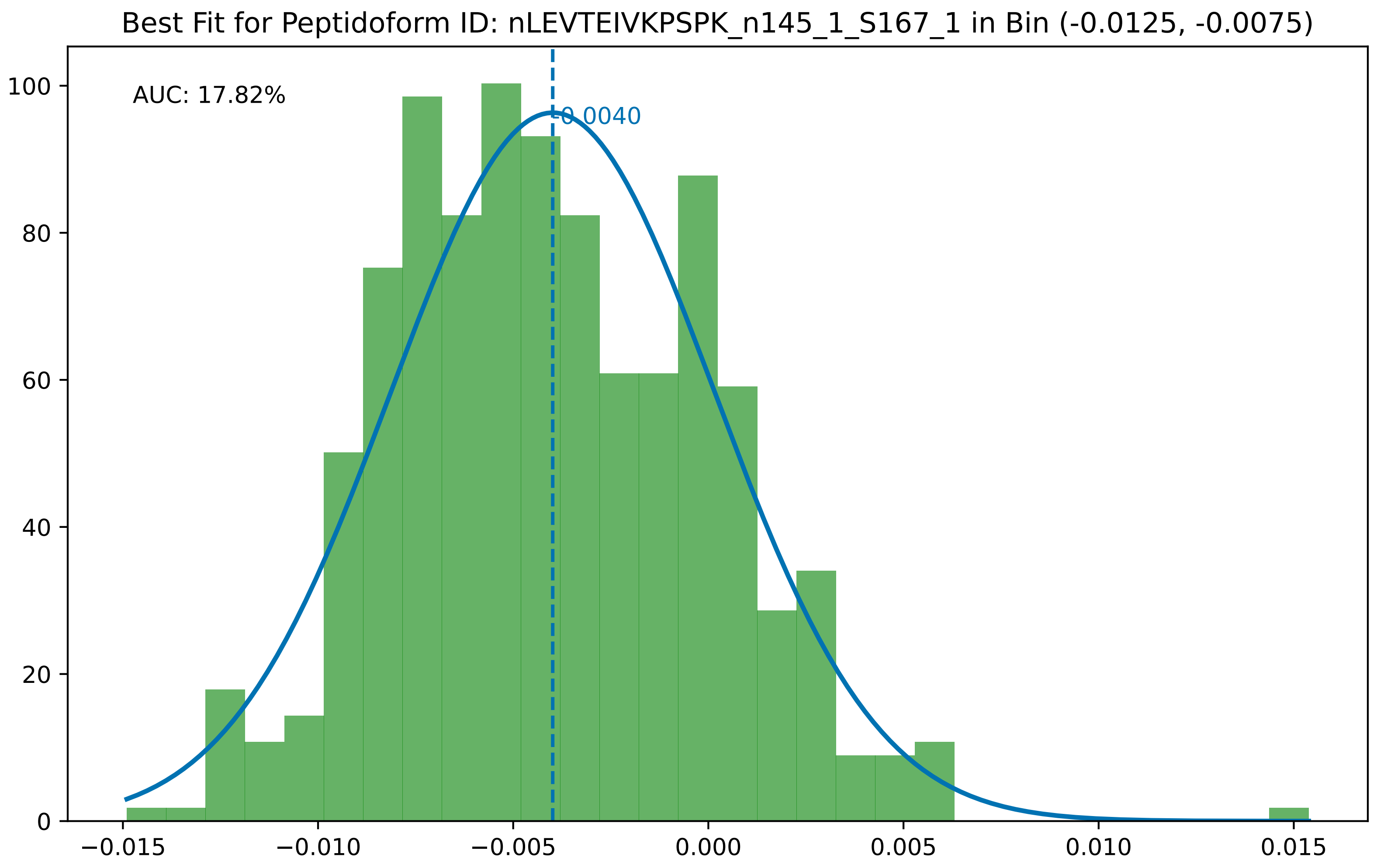

Best Fit for Peptidoform ID: nLEVTEIVKPSPK_n145_1_S167_1 in Bin (-0.0125, -0.0075)
AUC: 17.82%
-0.0040
0
20
40
60
80
100
−0.015
−0.010
−0.005
0.000
0.005
0.010
0.015

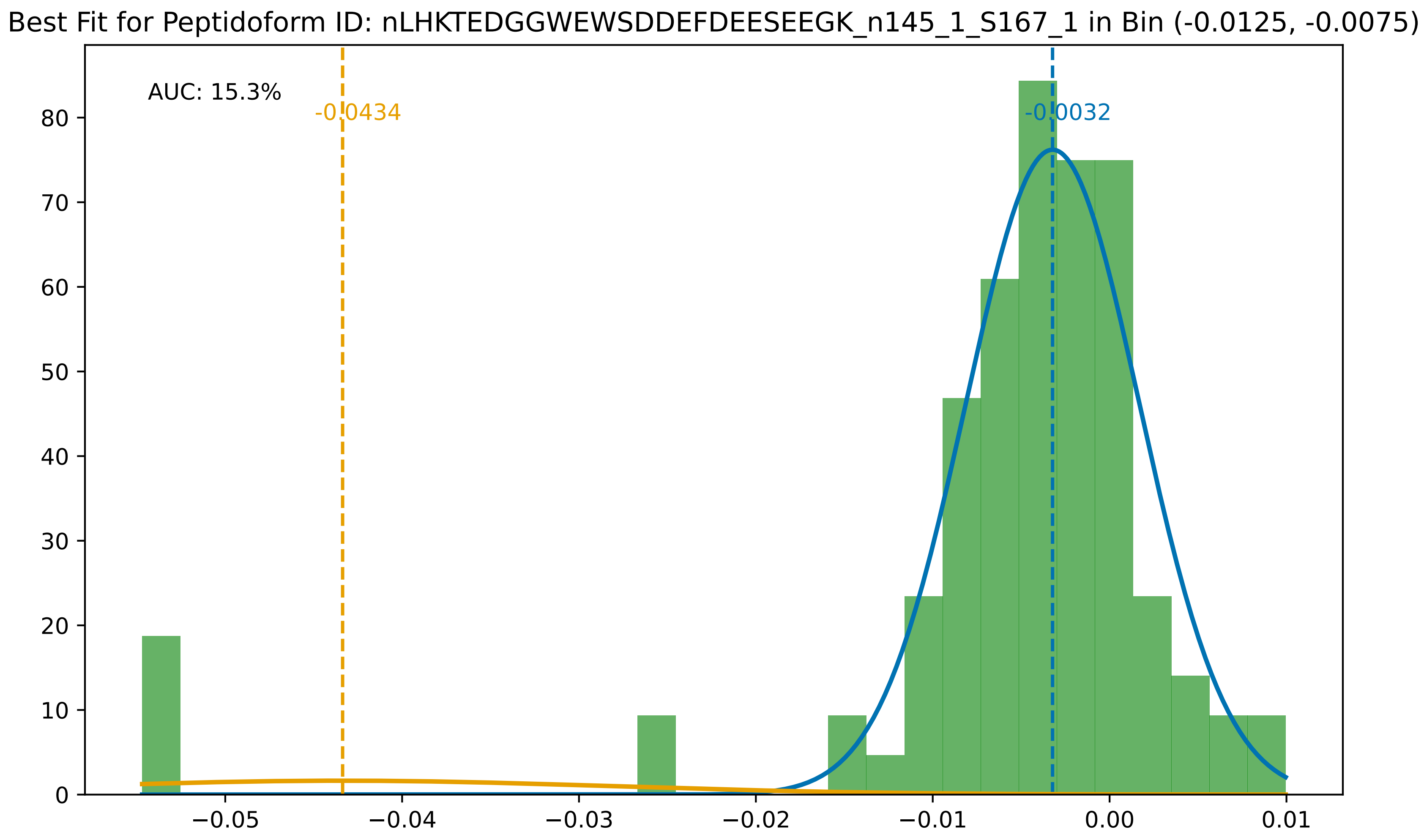

Best Fit for Peptidoform ID: nLHKTEDGGWEWSDDEFDEESEEGK_n145_1_S167_1 in Bin (-0.0125, -0.0075)
AUC: 15.3%
-0.0434
-0.0032
0
10
20
30
40
50
60
70
80
−0.05
−0.04
−0.03
−0.02
−0.01
0.00
0.01

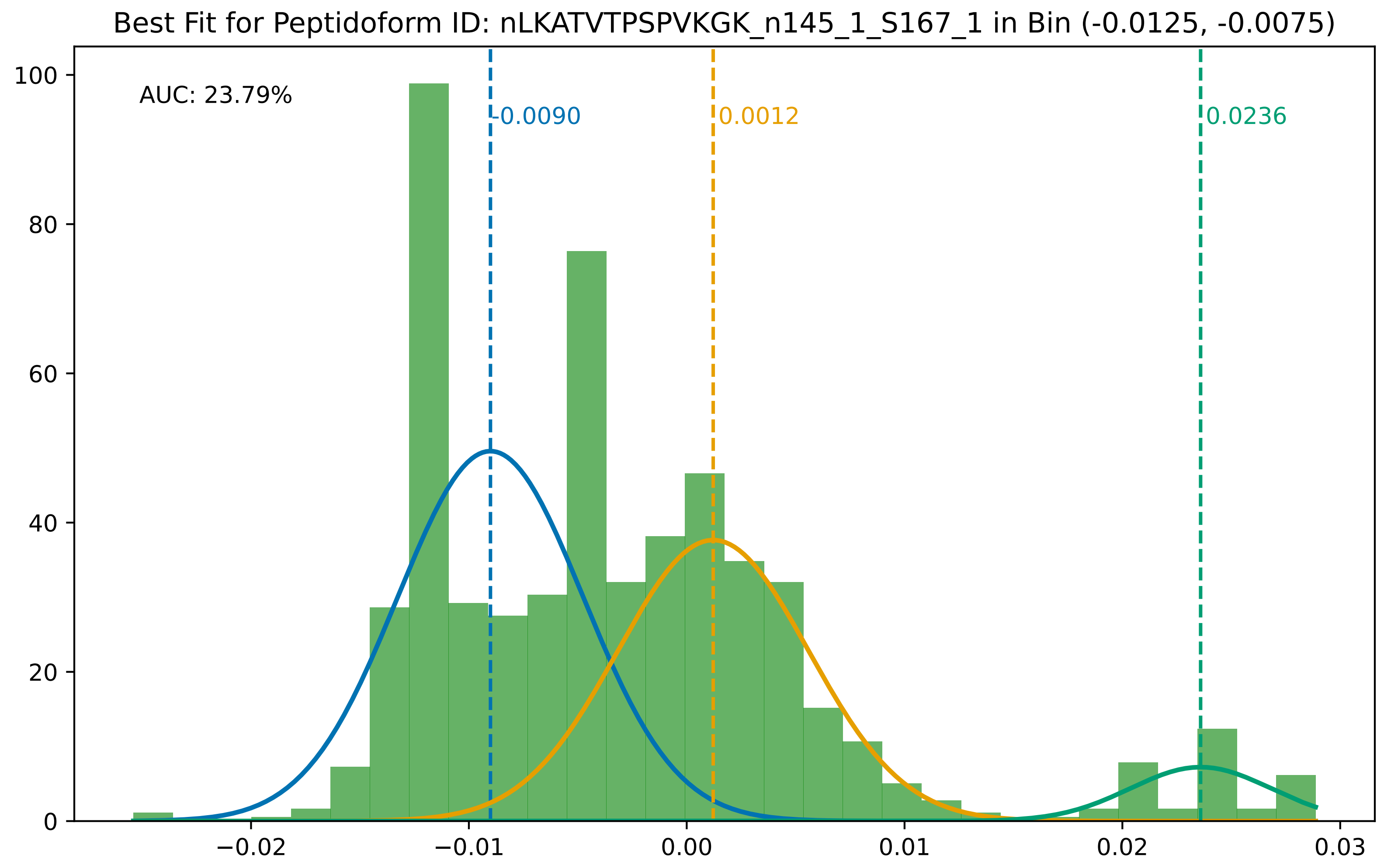

Best Fit for Peptidoform ID: nLKATVTPSPVKGK_n145_1_S167_1 in Bin (-0.0125, -0.0075)
AUC: 23.79%
-0.0090
0.0012
0.0236
100
80
60
40
20
0
−0.02
−0.01
0.00
0.01
0.02
0.03

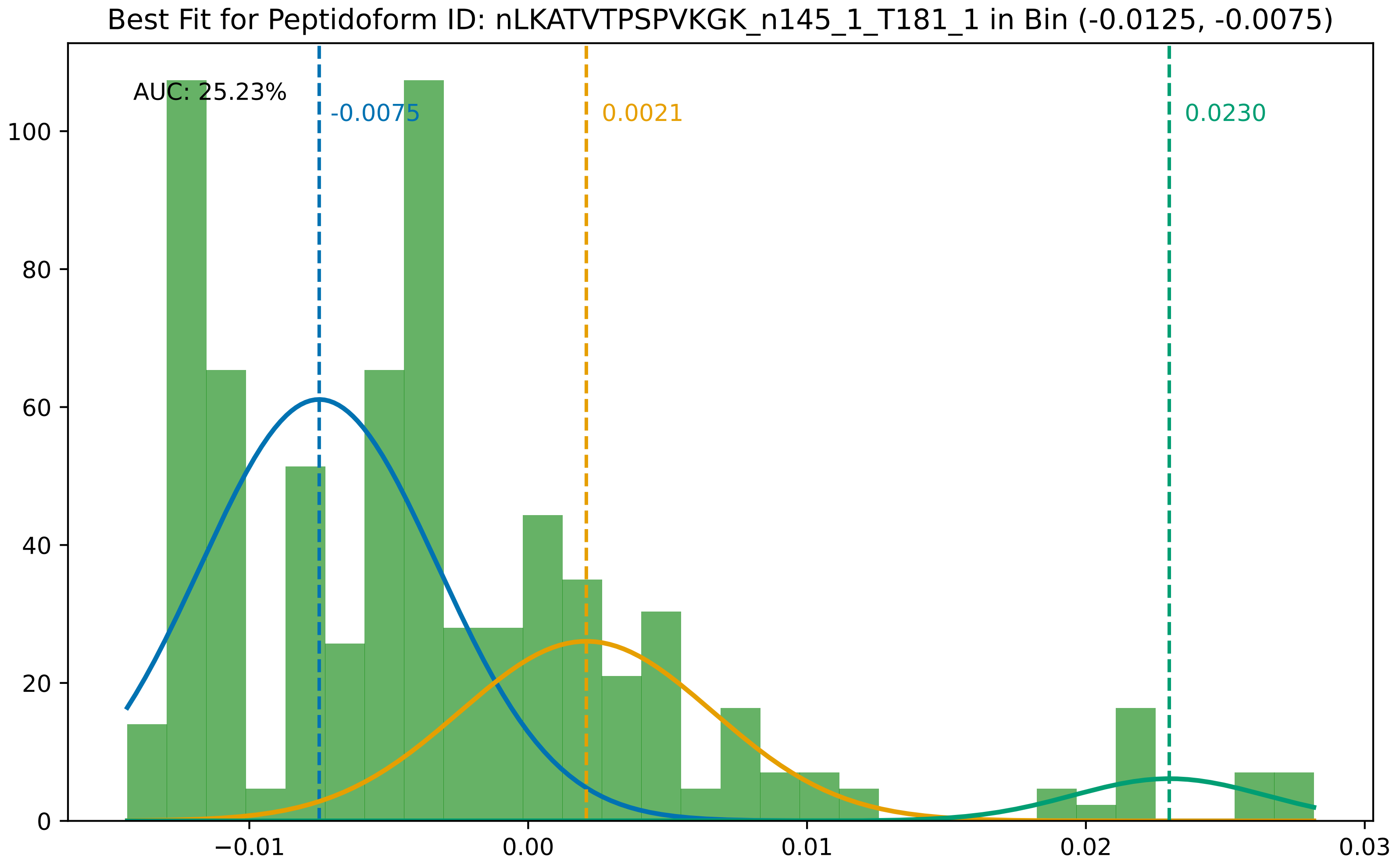
Best Fit for Peptidoform ID: nLKATVTPSPVKGK_n145_1_T181_1 in Bin (-0.0125, -0.0075)
AUC: 25.23%
-0.0075
0.0021
0.0230
100
80
60
40
20
0
−0.01
0.00
0.01
0.02
0.03

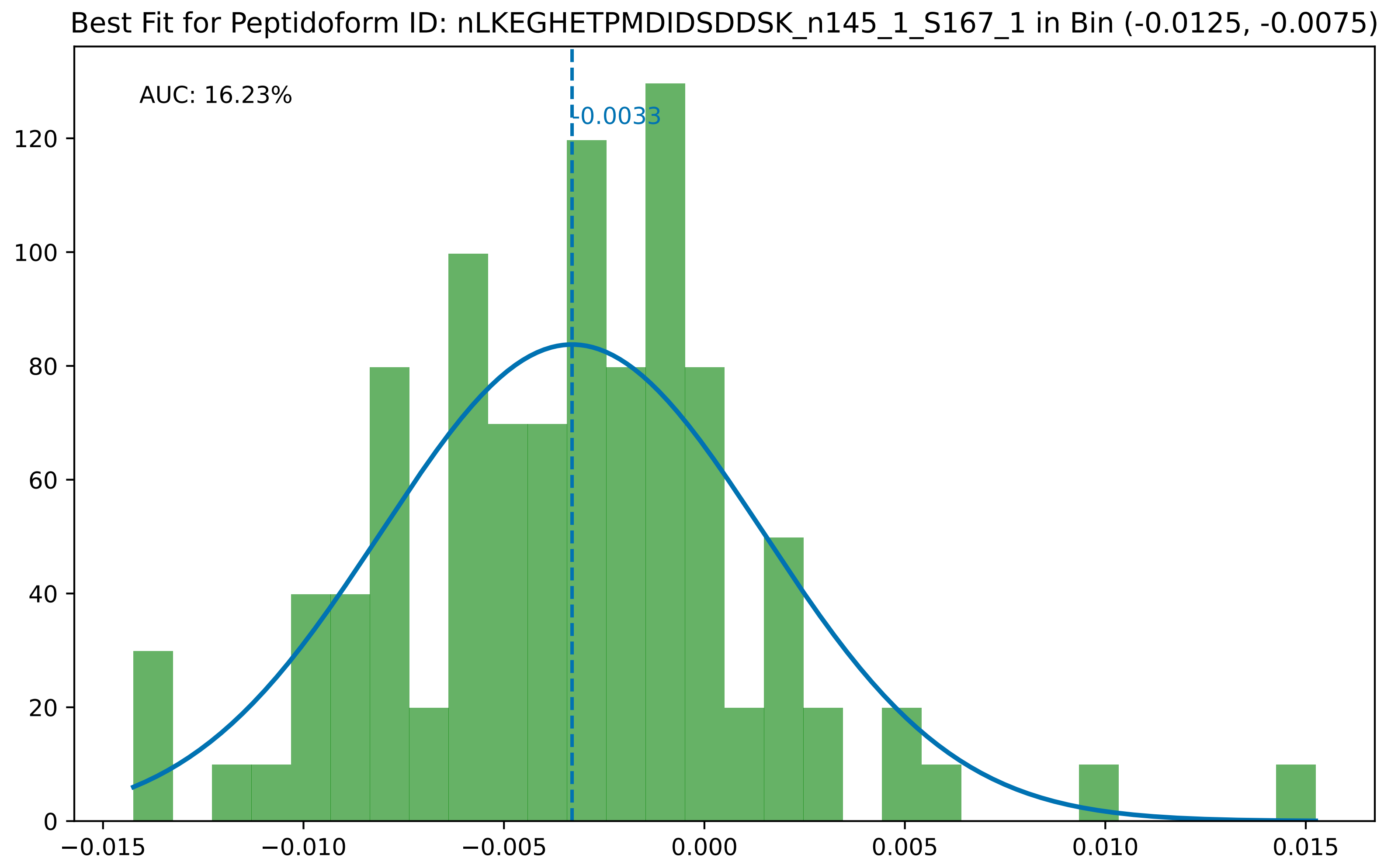
Best Fit for Peptidoform ID: nLKEGHETPMDIDSDDSK_n145_1_S167_1 in Bin (-0.0125, -0.0075)
AUC: 16.23%
-0.0033
0
20
40
60
80
100
120
−0.015
−0.010
−0.005
0.000
0.005
0.010
0.015

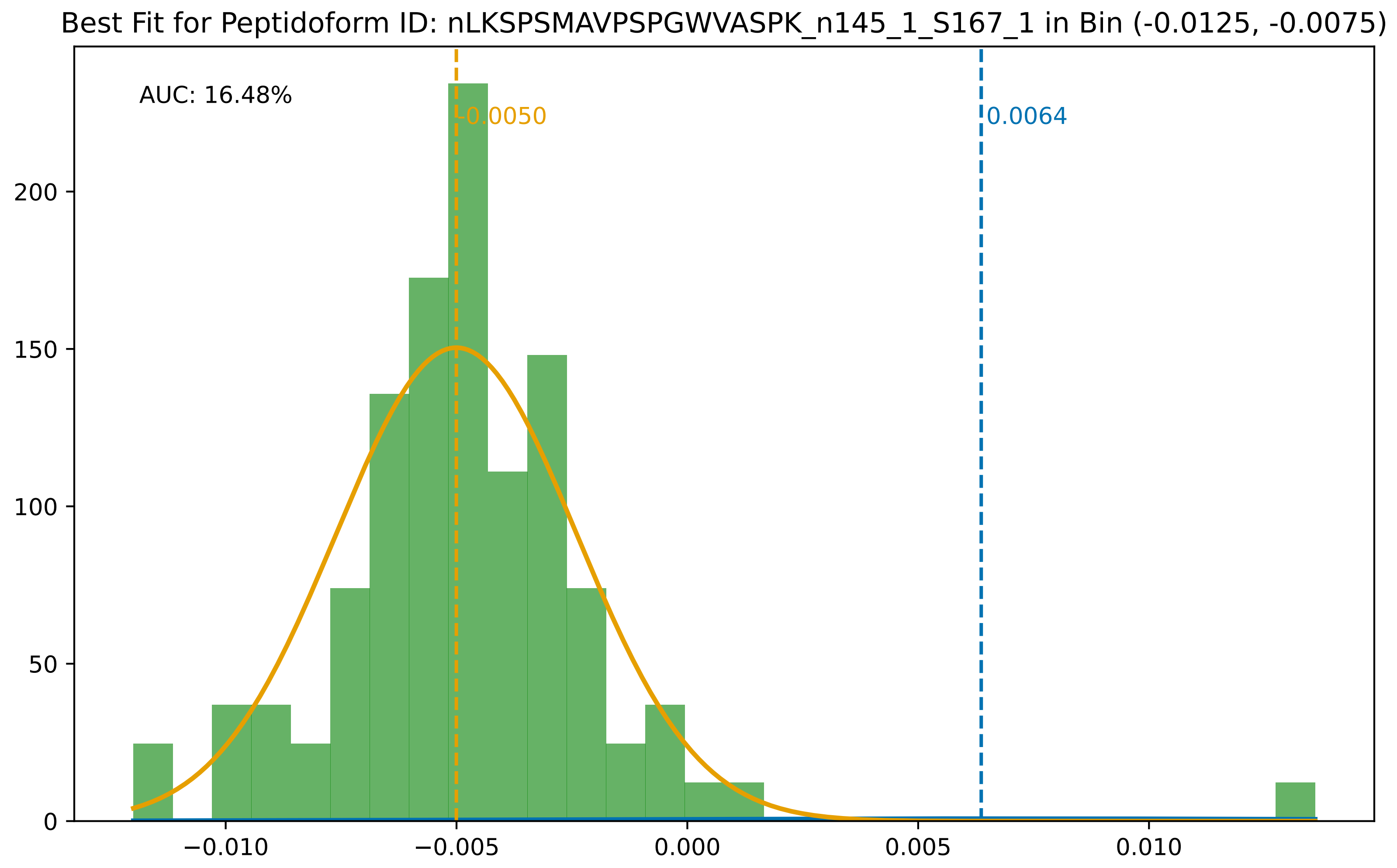

Best Fit for Peptidoform ID: nLKSPSMAVPSPGWVASPK_n145_1_S167_1 in Bin (-0.0125, -0.0075)
AUC: 16.48%
-0.0050
0.0064
0
50
100
150
200
−0.010
−0.005
0.000
0.005
0.010

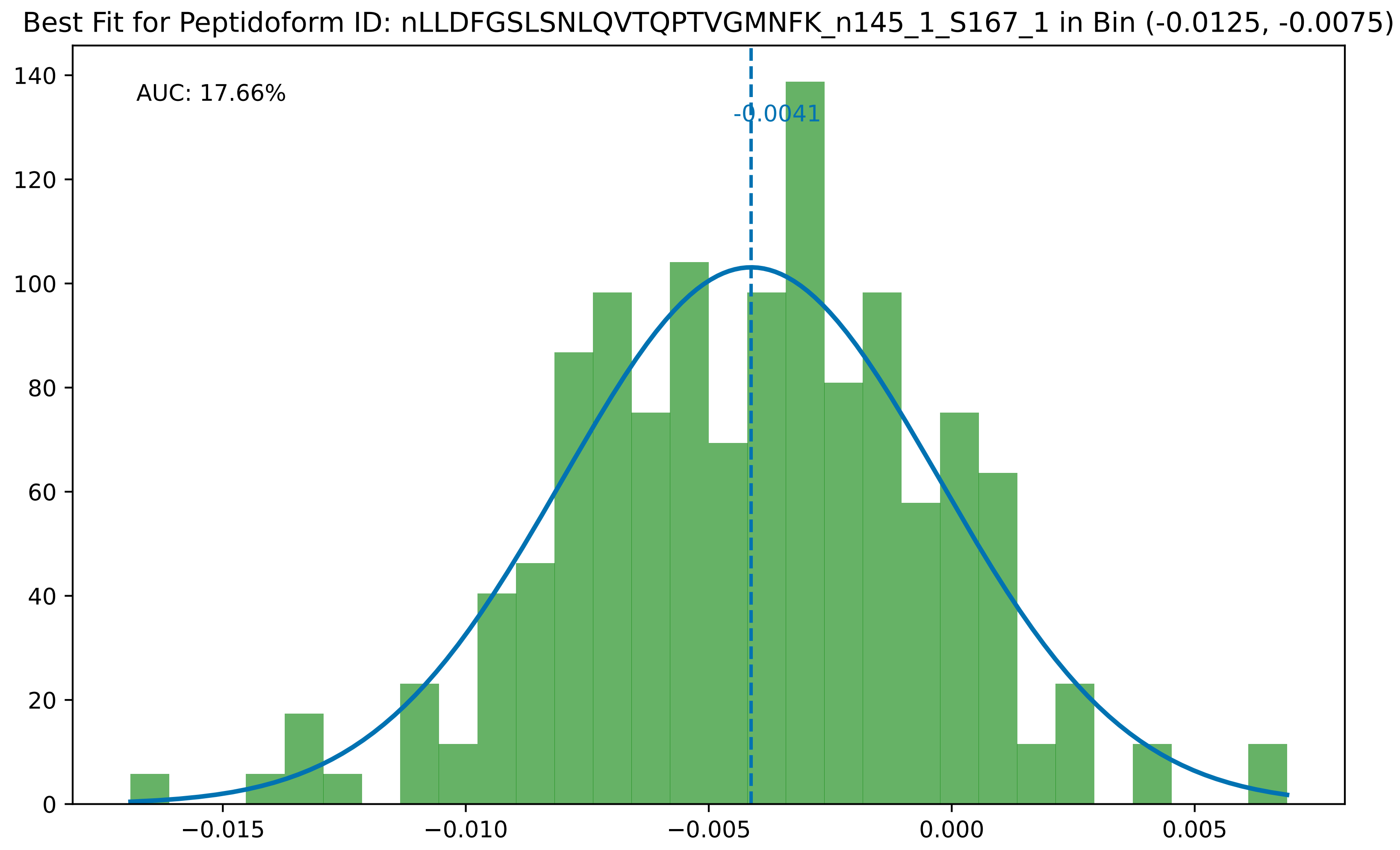
Best Fit for Peptidoform ID: nLLDFGSLSNLQVTQPTVGMNFK_n145_1_S167_1 in Bin (-0.0125, -0.0075)
AUC: 17.66%
-0.0041
0
20
40
60
80
100
120
140
−0.015
−0.010
−0.005
0.000
0.005

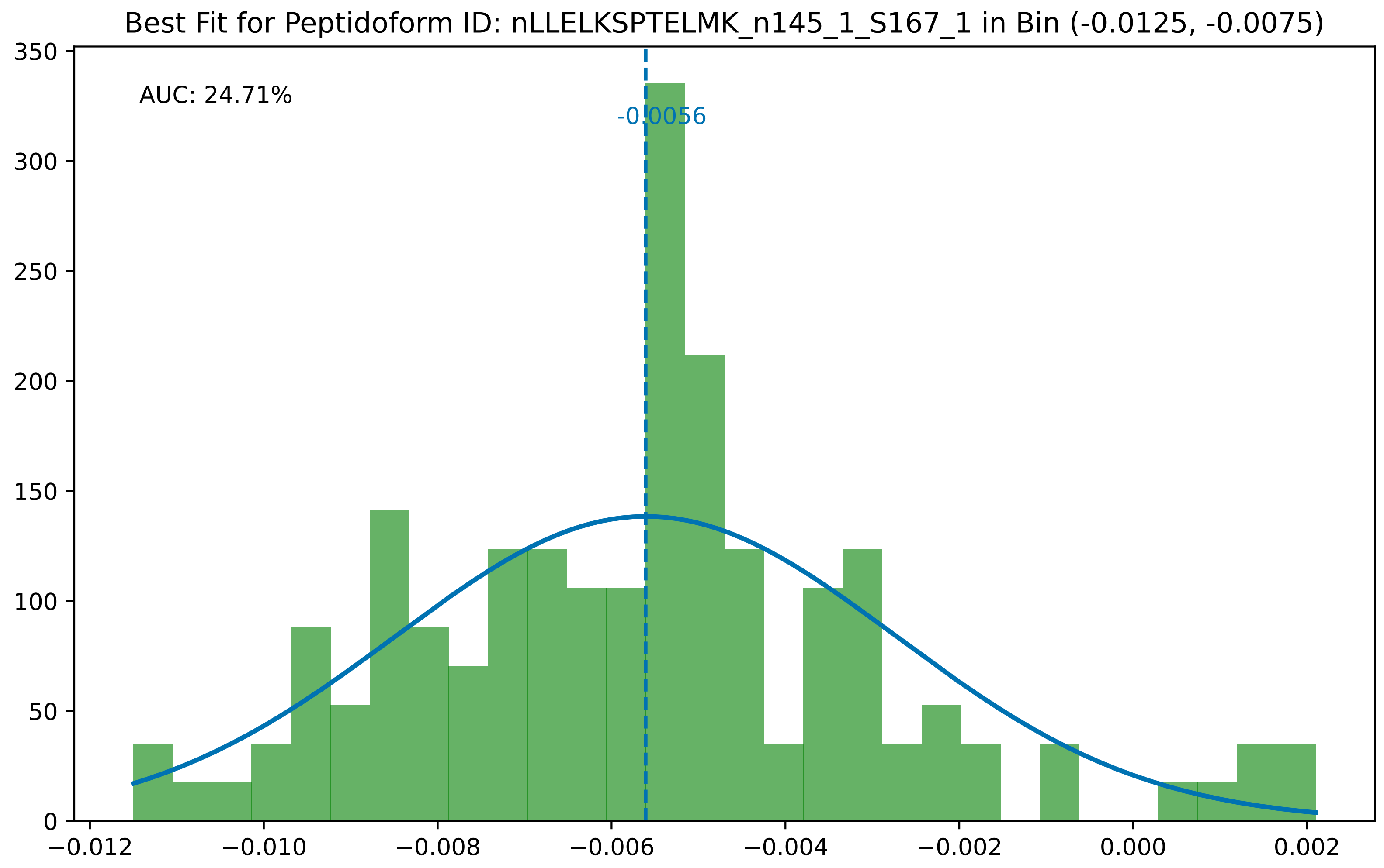

Best Fit for Peptidoform ID: nLLELKSPTELMK_n145_1_S167_1 in Bin (-0.0125, -0.0075)
AUC: 24.71%
-0.0056
0
50
100
150
200
250
300
350
−0.012
−0.010
−0.008
−0.006
−0.004
−0.002
0.000
0.002

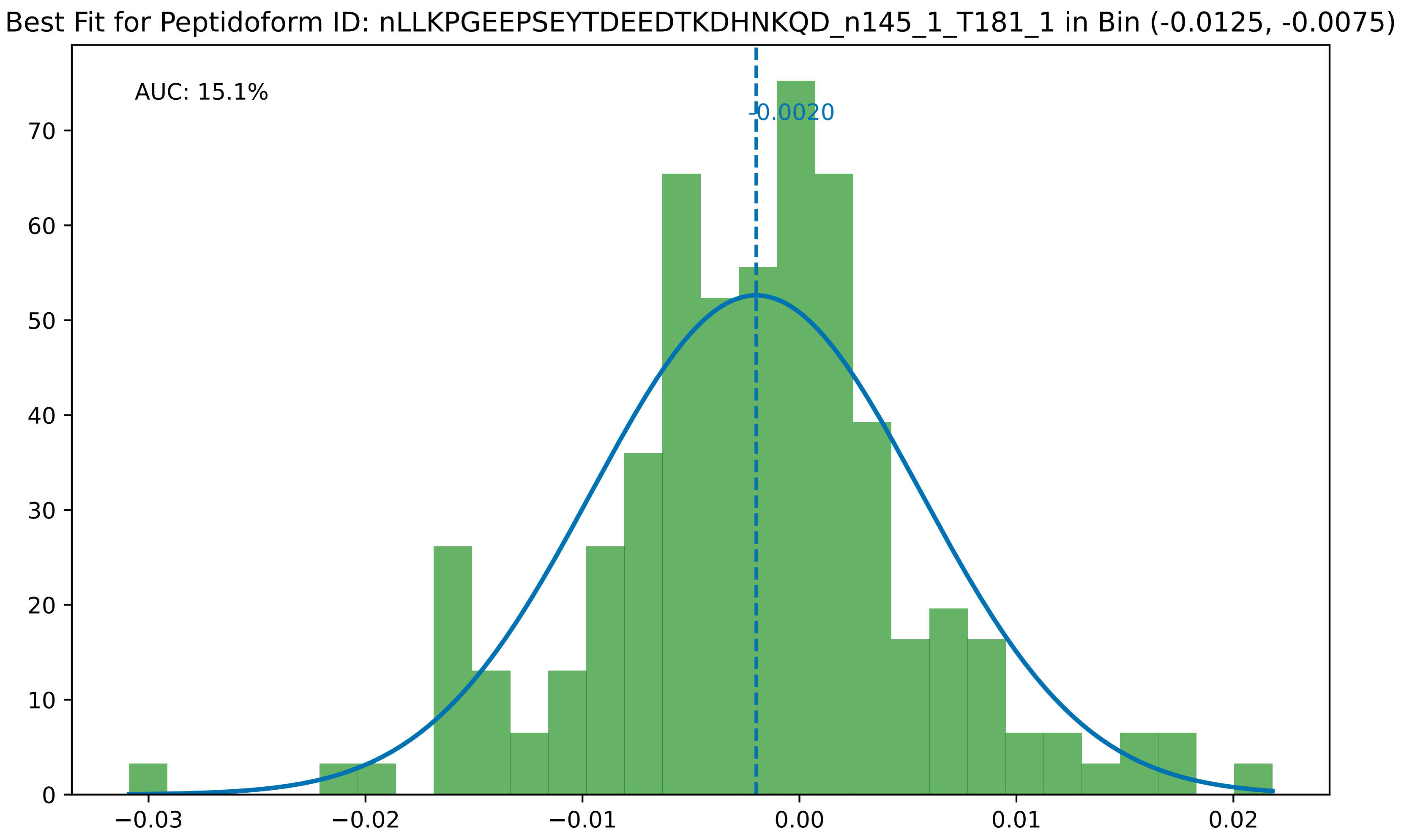

Best Fit for Peptidoform ID: nLLKPGEEPSEYTDEEDTKDHNKQD_n145_1_T181_1 in Bin (-0.0125, -0.0075)
AUC: 15.1%
-0.0020
70
60
50
40
30
20
10
0
−0.03
−0.02
−0.01
0.00
0.01
0.02

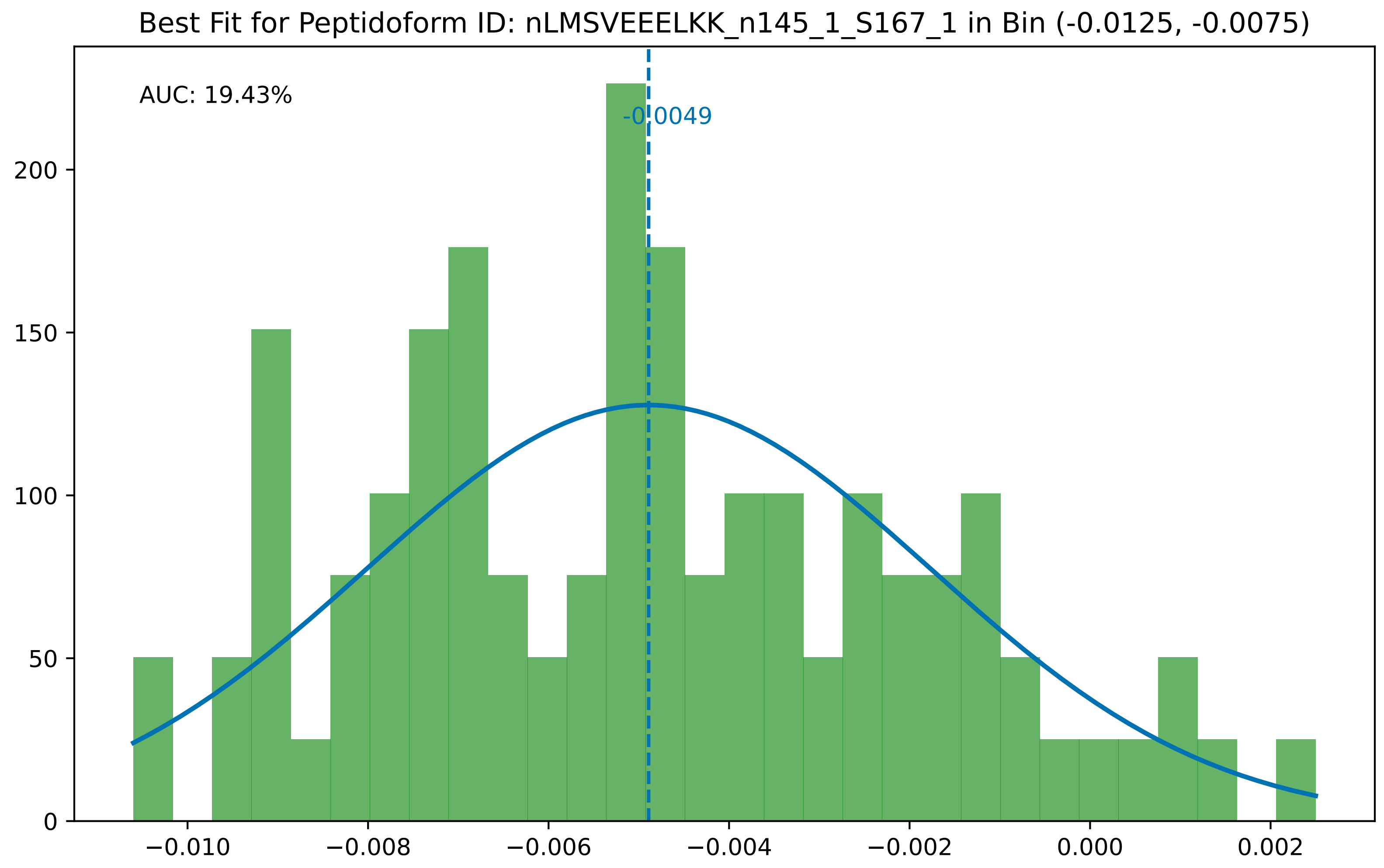
Best Fit for Peptidoform ID: nLMSVEEELKK_n145_1_S167_1 in Bin (-0.0125, -0.0075)
AUC: 19.43%
-0.0049
0
50
100
150
200
−0.010
−0.008
−0.006
−0.004
−0.002
0.000
0.002

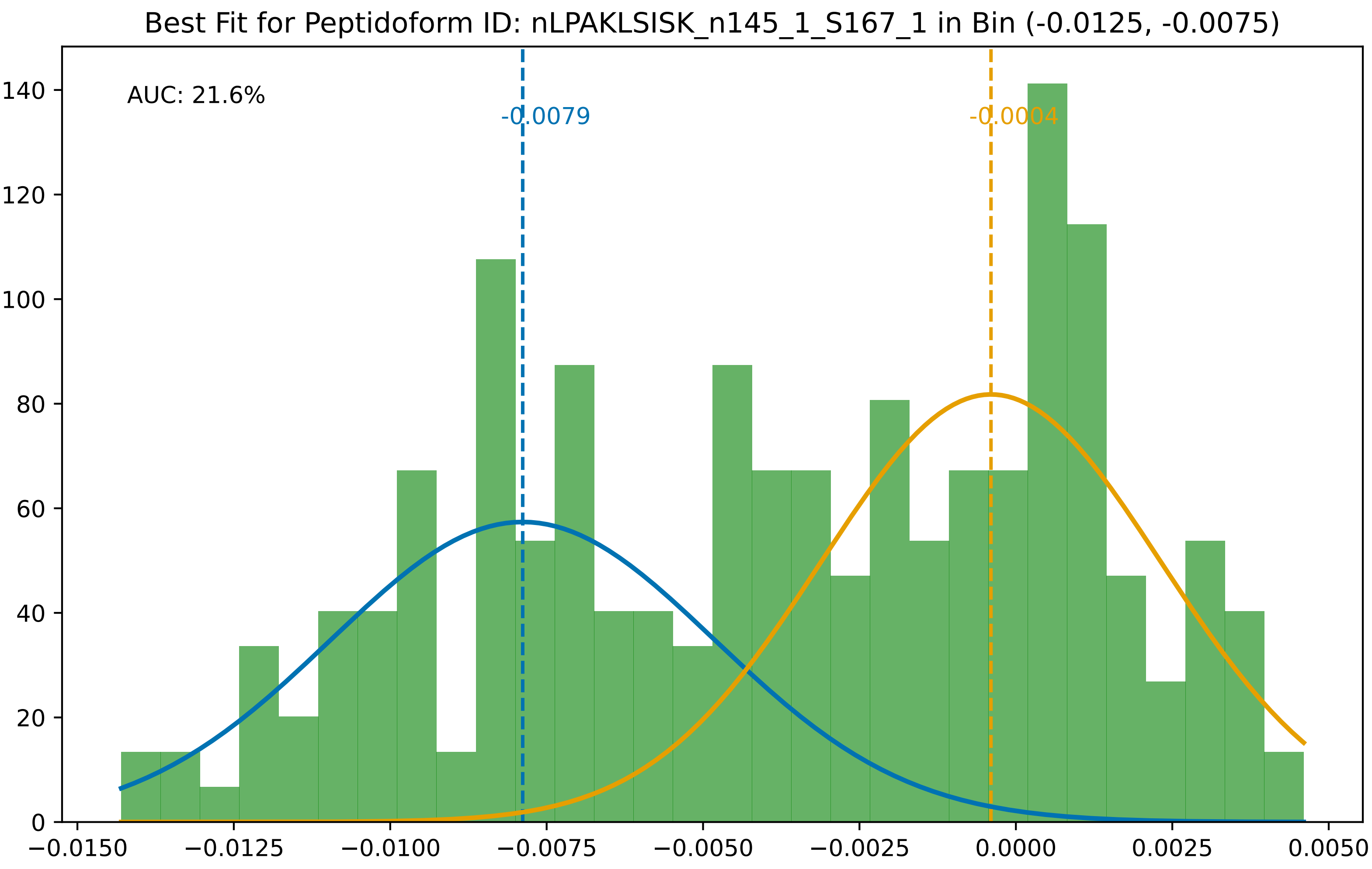

Best Fit for Peptidoform ID: nLPAKLSISK_n145_1_S167_1 in Bin (-0.0125, -0.0075)
AUC: 21.6%
-0.0079
-0.0004
0
20
40
60
80
100
120
140
−0.0150
−0.0125
−0.0100
−0.0075
−0.0050
−0.0025
0.0000
0.0025
0.0050

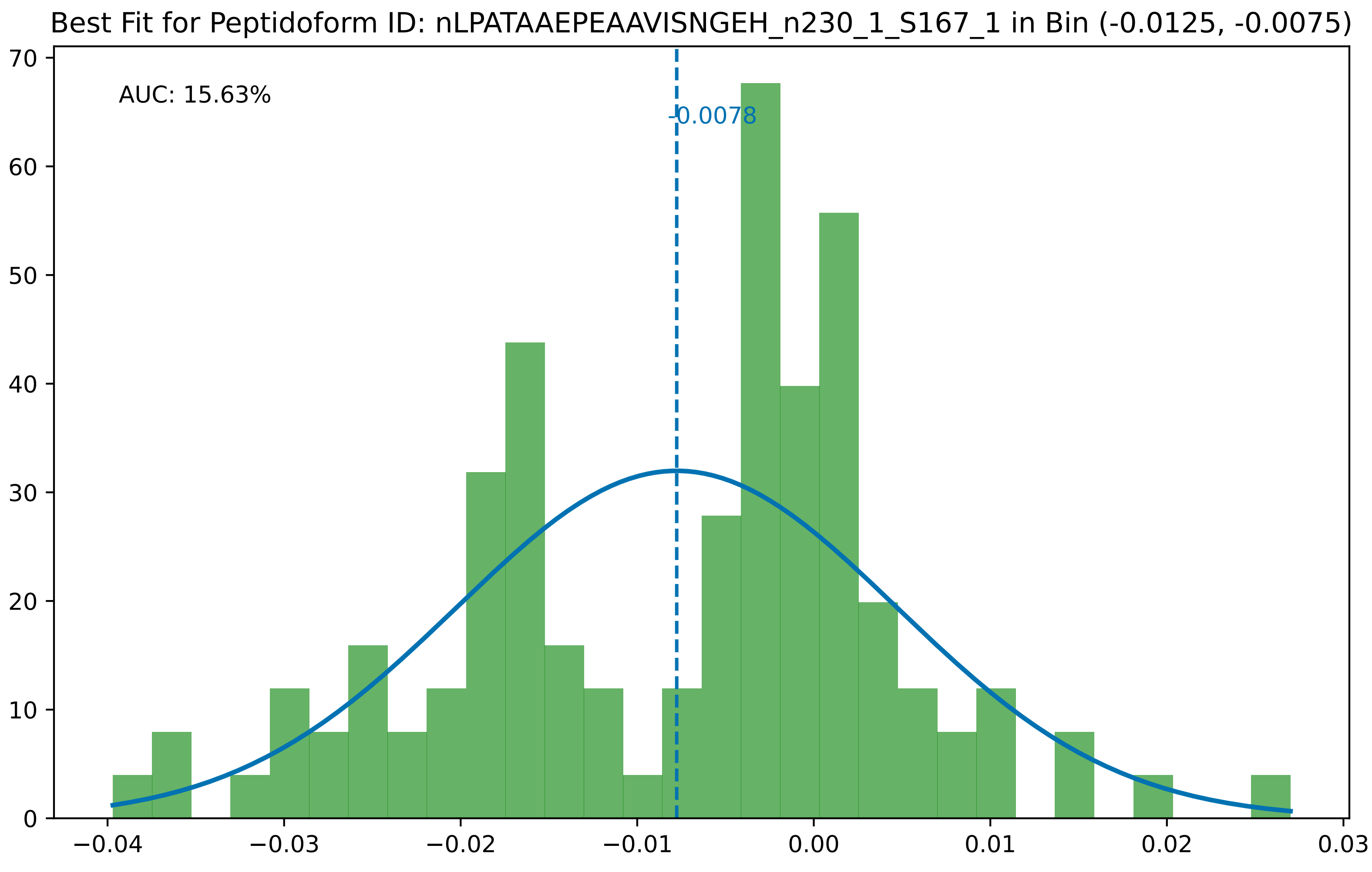
Best Fit for Peptidoform ID: nLPATAAEPEAAVISNGEH_n230_1_S167_1 in Bin (-0.0125, -0.0075)
AUC: 15.63%
-0.0078
70
60
50
40
30
20
10
0
−0.04
−0.03
−0.02
−0.01
0.00
0.01
0.02
0.03

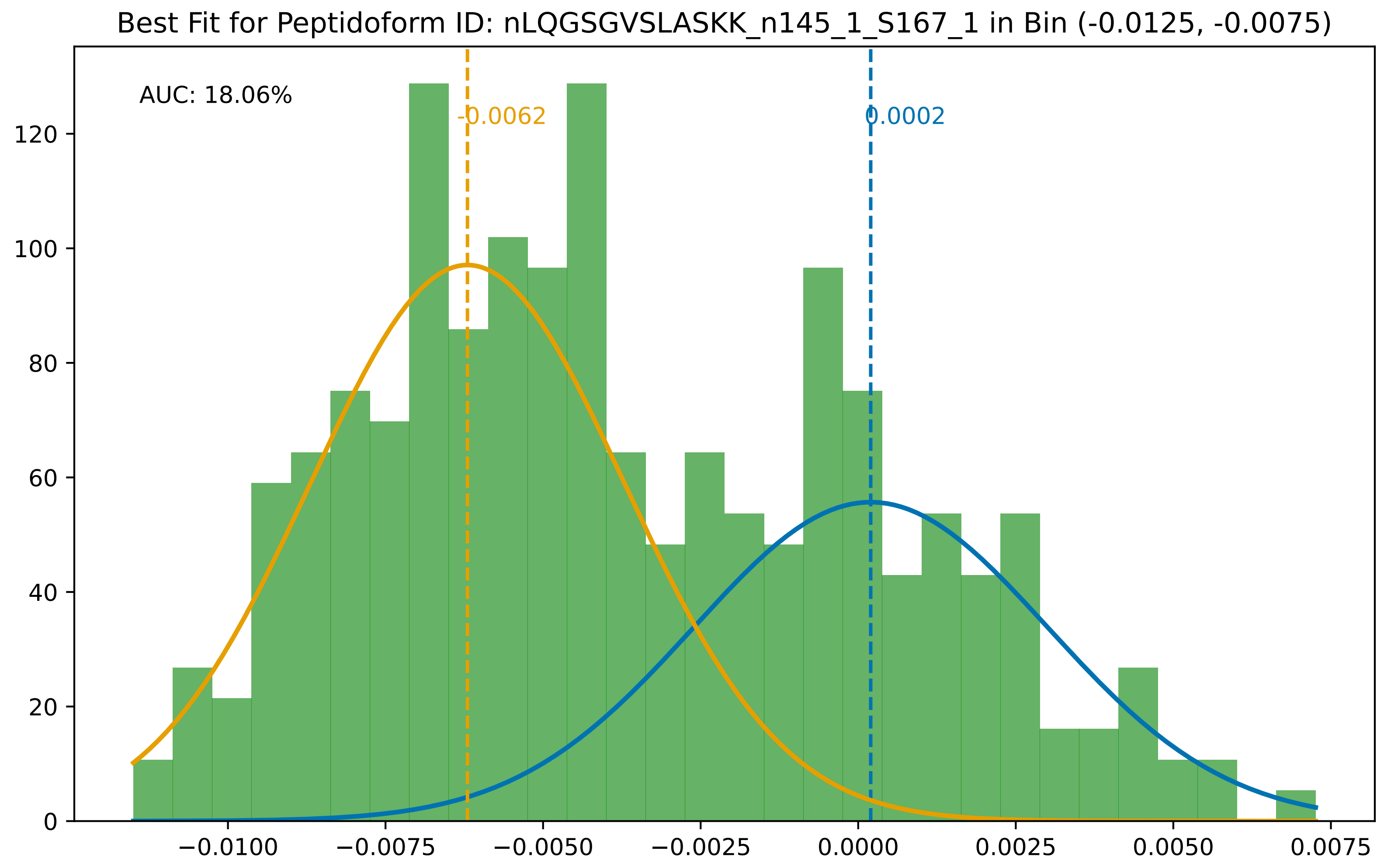

Best Fit for Peptidoform ID: nLQGSGVSLASKK_n145_1_S167_1 in Bin (-0.0125, -0.0075)
AUC: 18.06%
-0.0062
0.0002
0
20
40
60
80
100
120
−0.0100
−0.0075
−0.0050
−0.0025
0.0000
0.0025
0.0050
0.0075

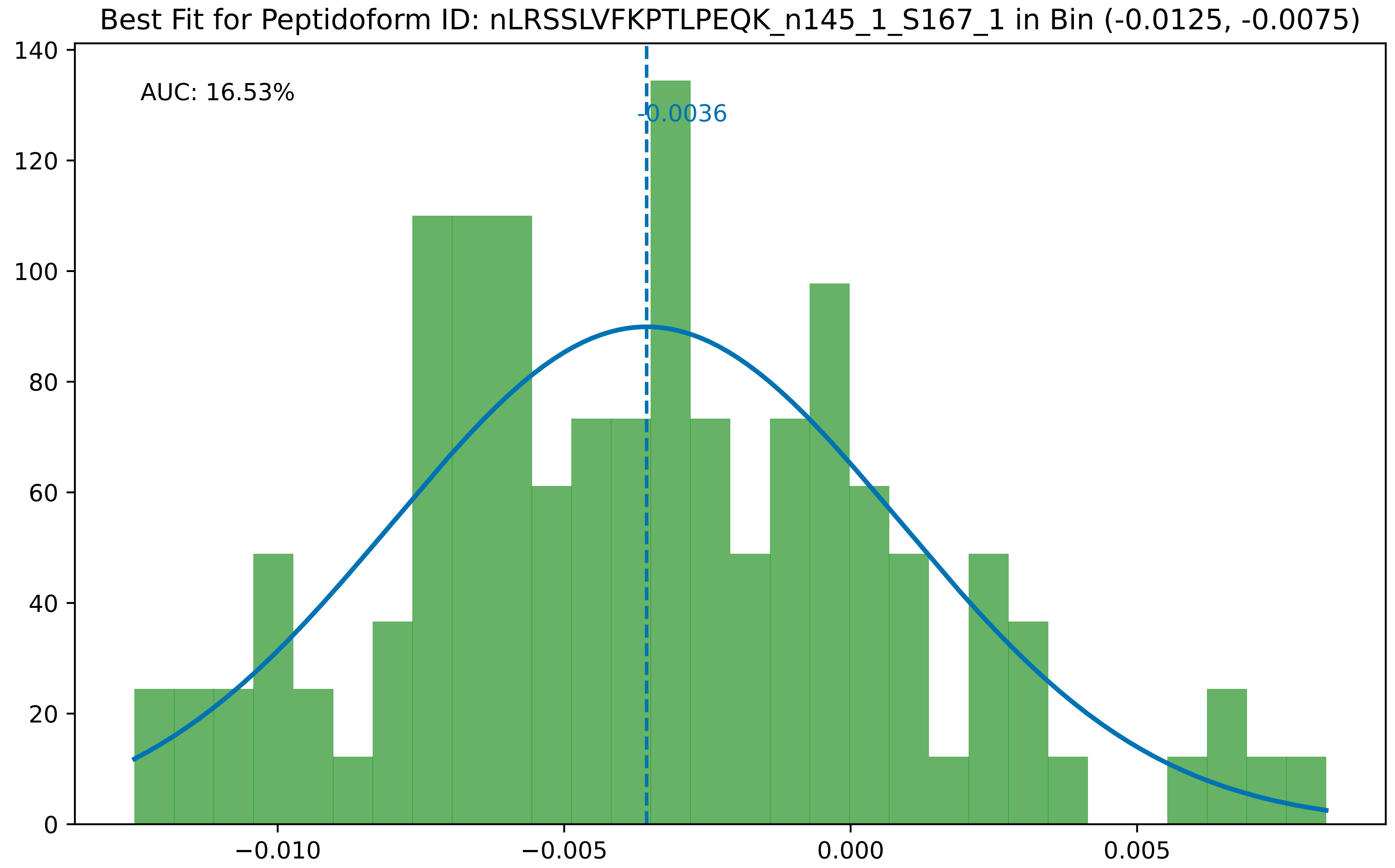

Best Fit for Peptidoform ID: nLRSSLVFKPTLPEQK_n145_1_S167_1 in Bin (-0.0125, -0.0075)
AUC: 16.53%
-0.0036
0
20
40
60
80
100
120
140
−0.010
−0.005
0.000
0.005

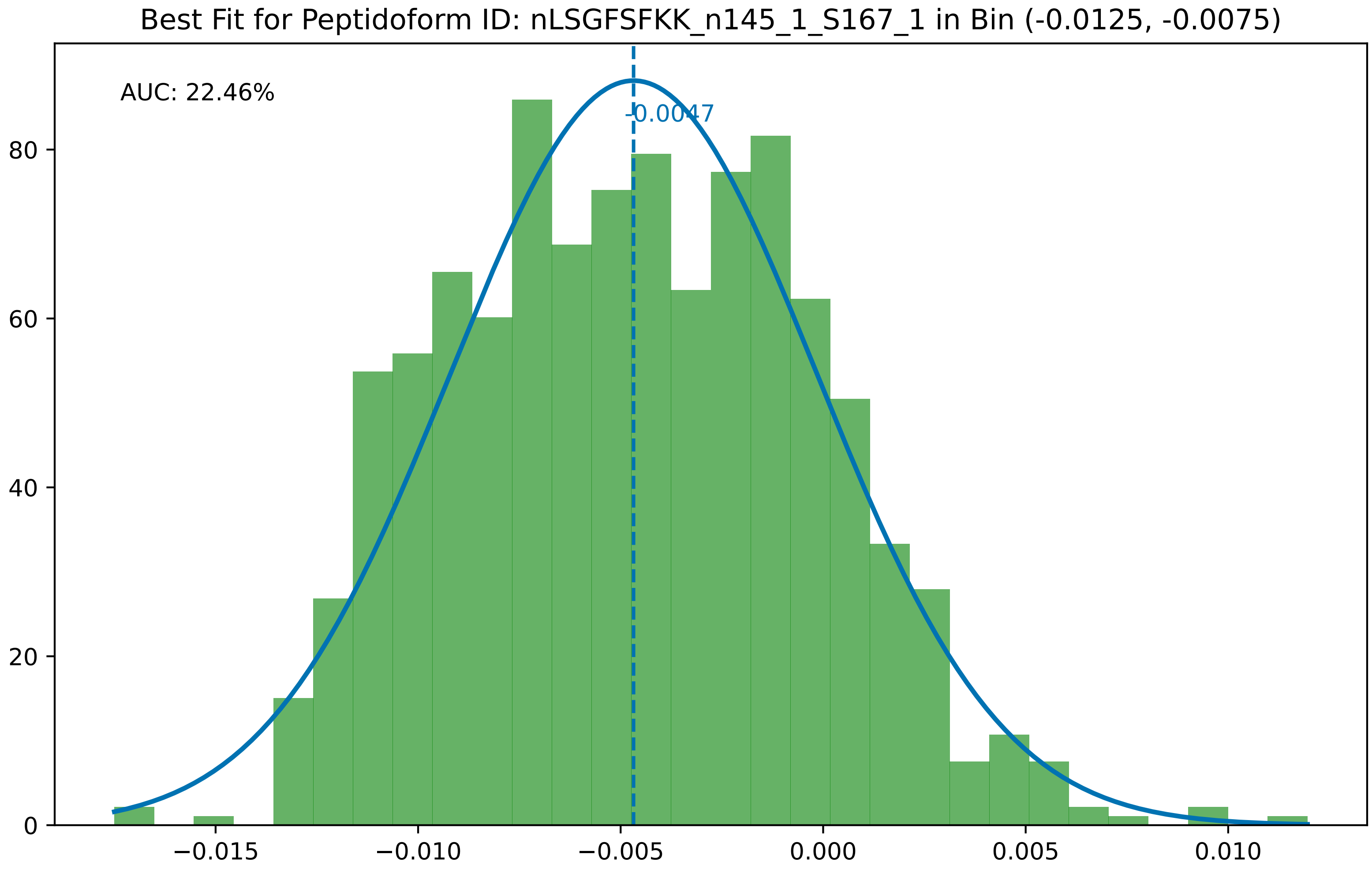

Best Fit for Peptidoform ID: nLSGFSFKK_n145_1_S167_1 in Bin (-0.0125, -0.0075)
AUC: 22.46%
-0.0047
0
20
40
60
80
−0.015
−0.010
−0.005
0.000
0.005
0.010

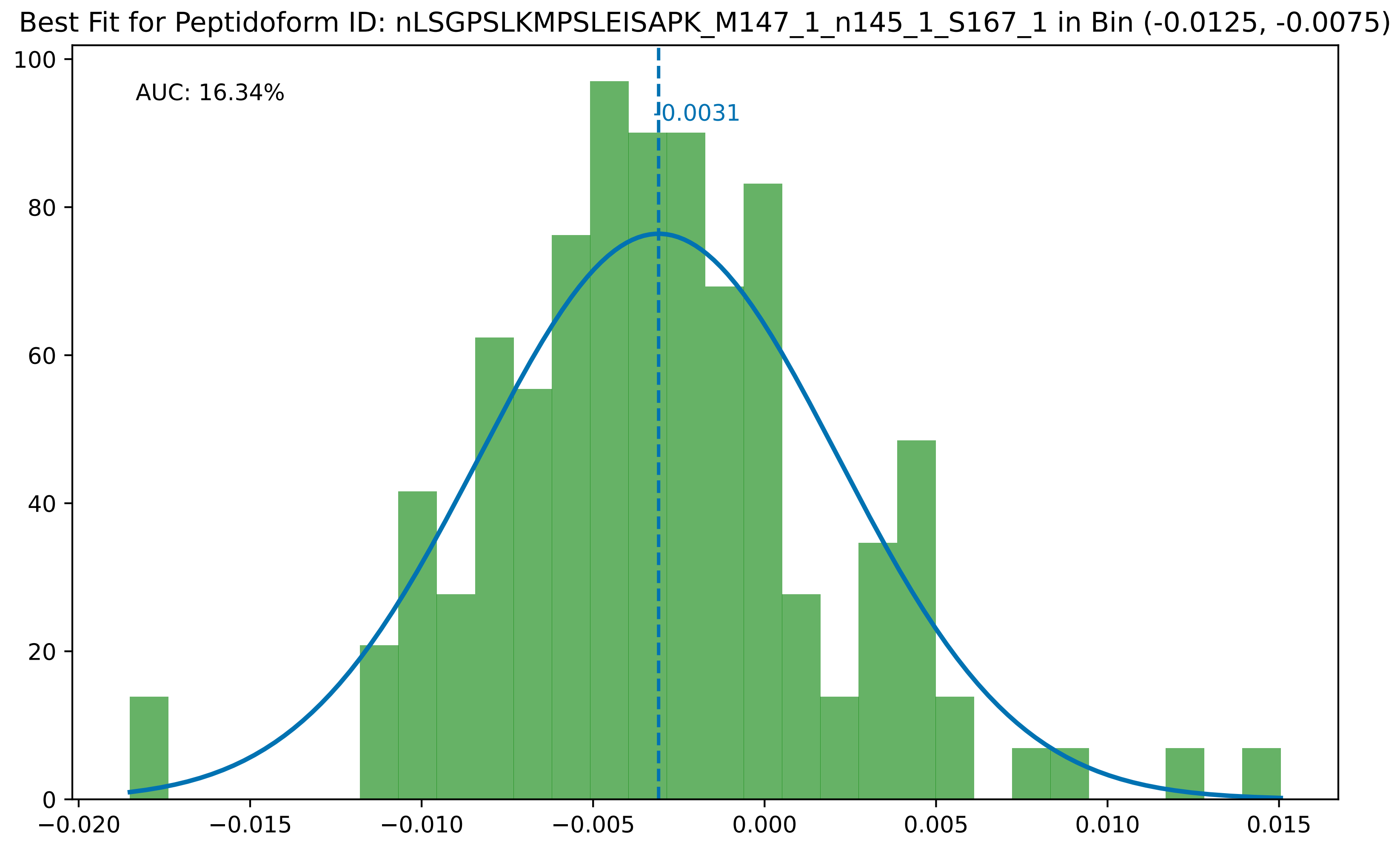
Best Fit for Peptidoform ID: nLSGPSLKMPSLEISAPK_M147_1_n145_1_S167_1 in Bin (-0.0125, -0.0075)
AUC: 16.34%
-0.0031
0
20
40
60
80
100
−0.020
−0.015
−0.010
−0.005
0.000
0.005
0.010
0.015

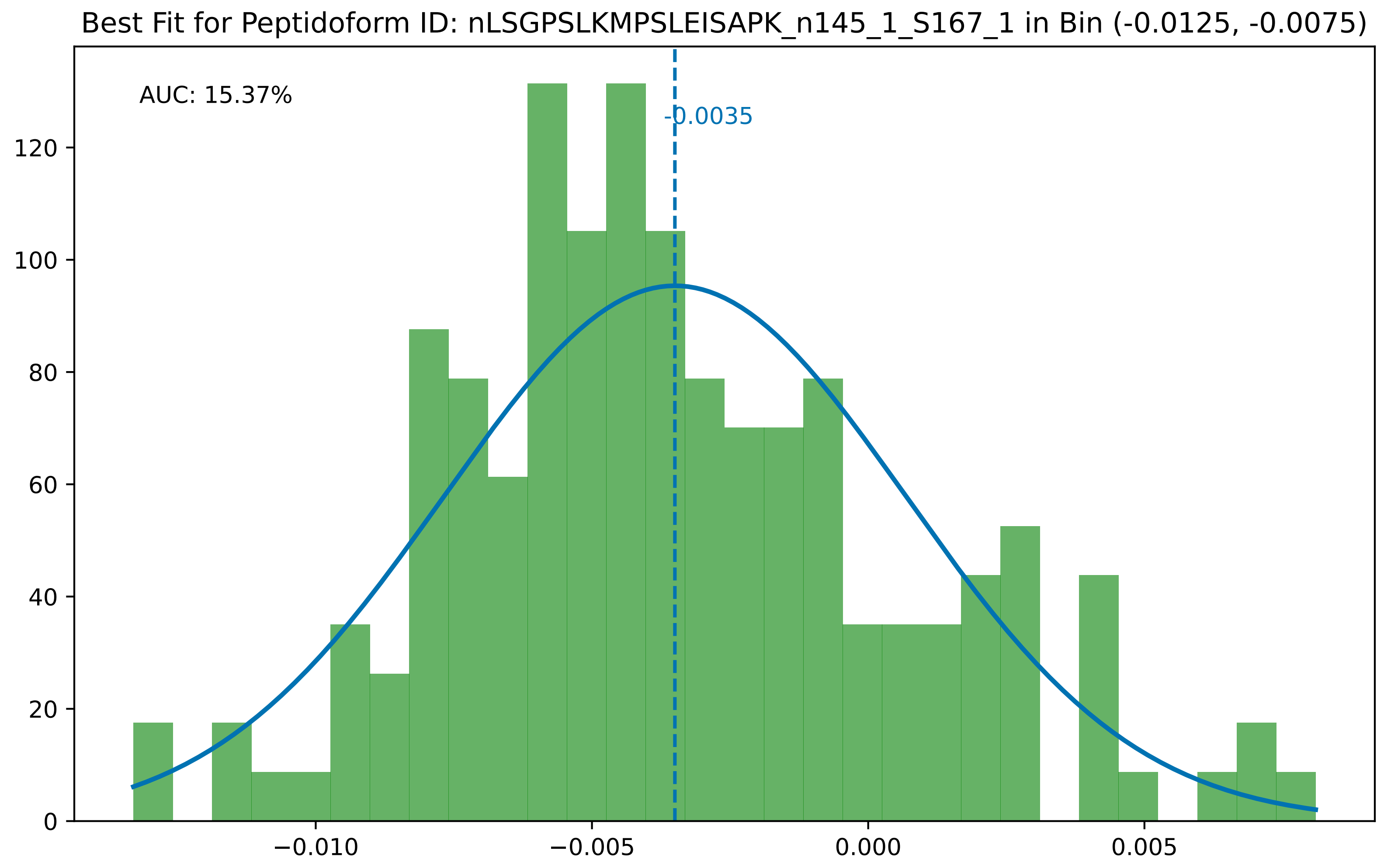

Best Fit for Peptidoform ID: nLSGPSLKMPSLEISAPK_n145_1_S167_1 in Bin (-0.0125, -0.0075)
AUC: 15.37%
-0.0035
0
20
40
60
80
100
120
−0.010
−0.005
0.000
0.005

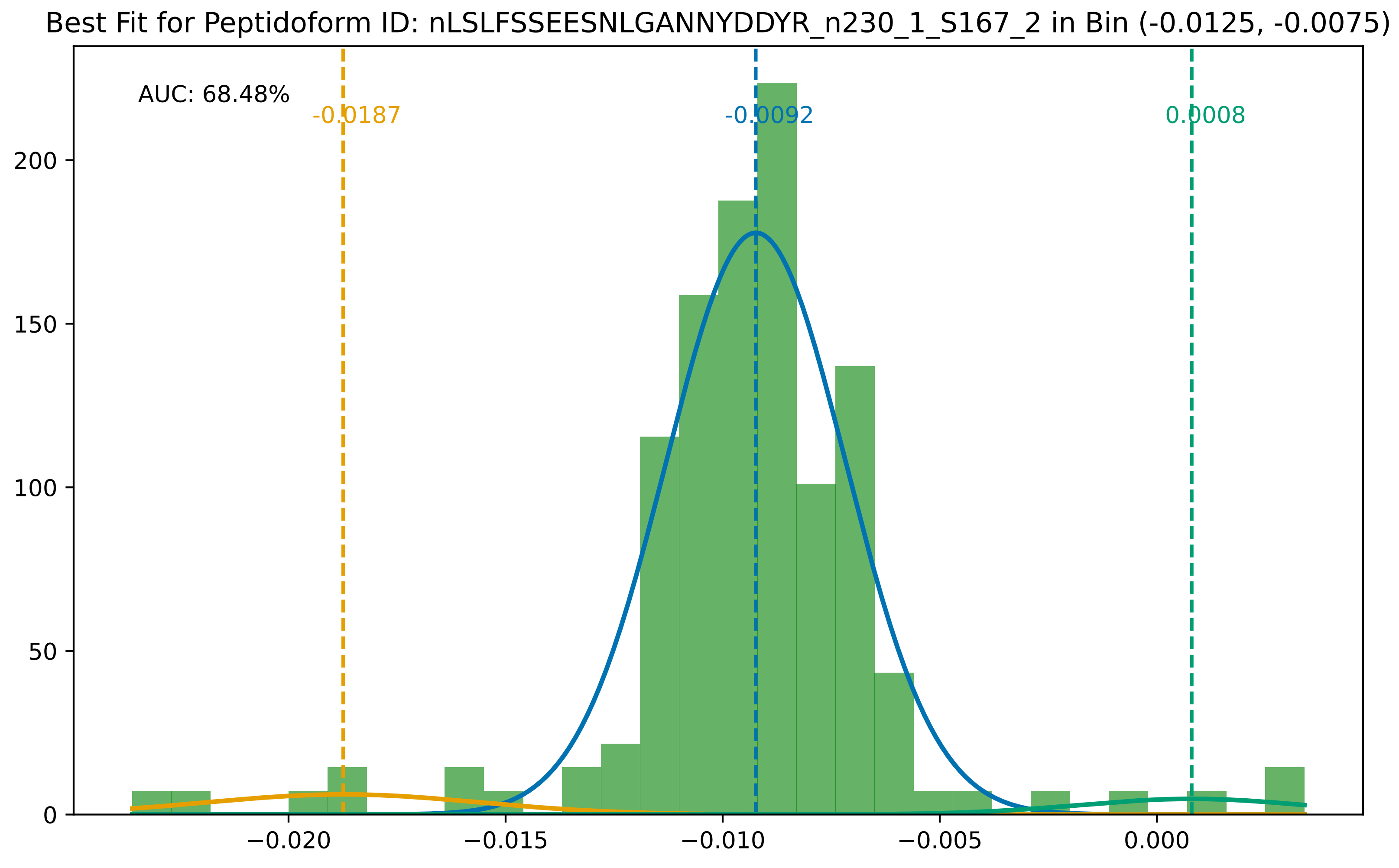
Best Fit for Peptidoform ID: nLSLFSSEESNLGANNYDDYR_n230_1_S167_2 in Bin (-0.0125, -0.0075)
AUC: 68.48%
-0.0187
-0.0092
0.0008
200
150
100
50
0
−0.020
−0.015
−0.010
−0.005
0.000

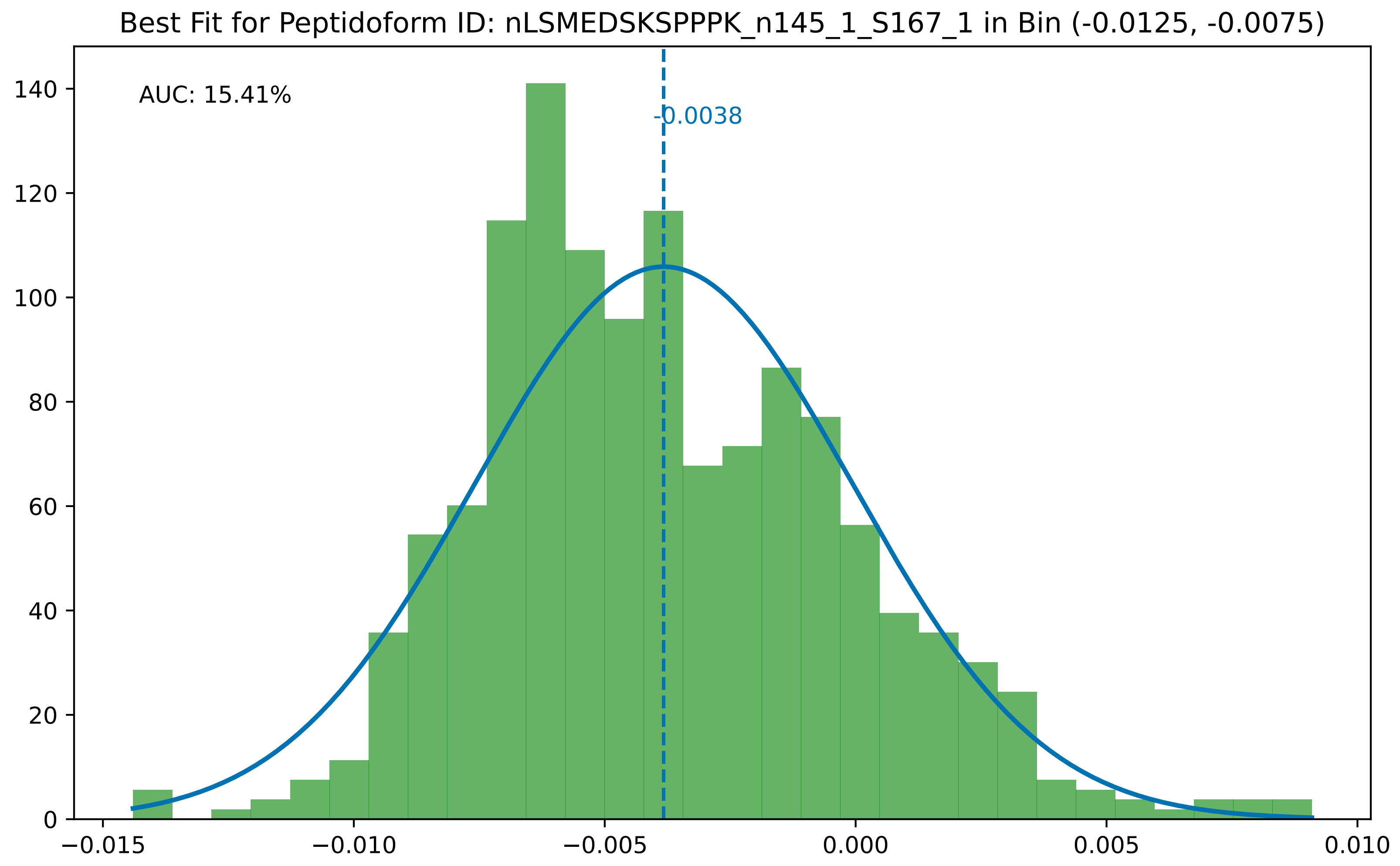

Best Fit for Peptidoform ID: nLSMEDSKSPPPK_n145_1_S167_1 in Bin (-0.0125, -0.0075)
AUC: 15.41%
-0.0038
0
20
40
60
80
100
120
140
−0.015
−0.010
−0.005
0.000
0.005
0.010

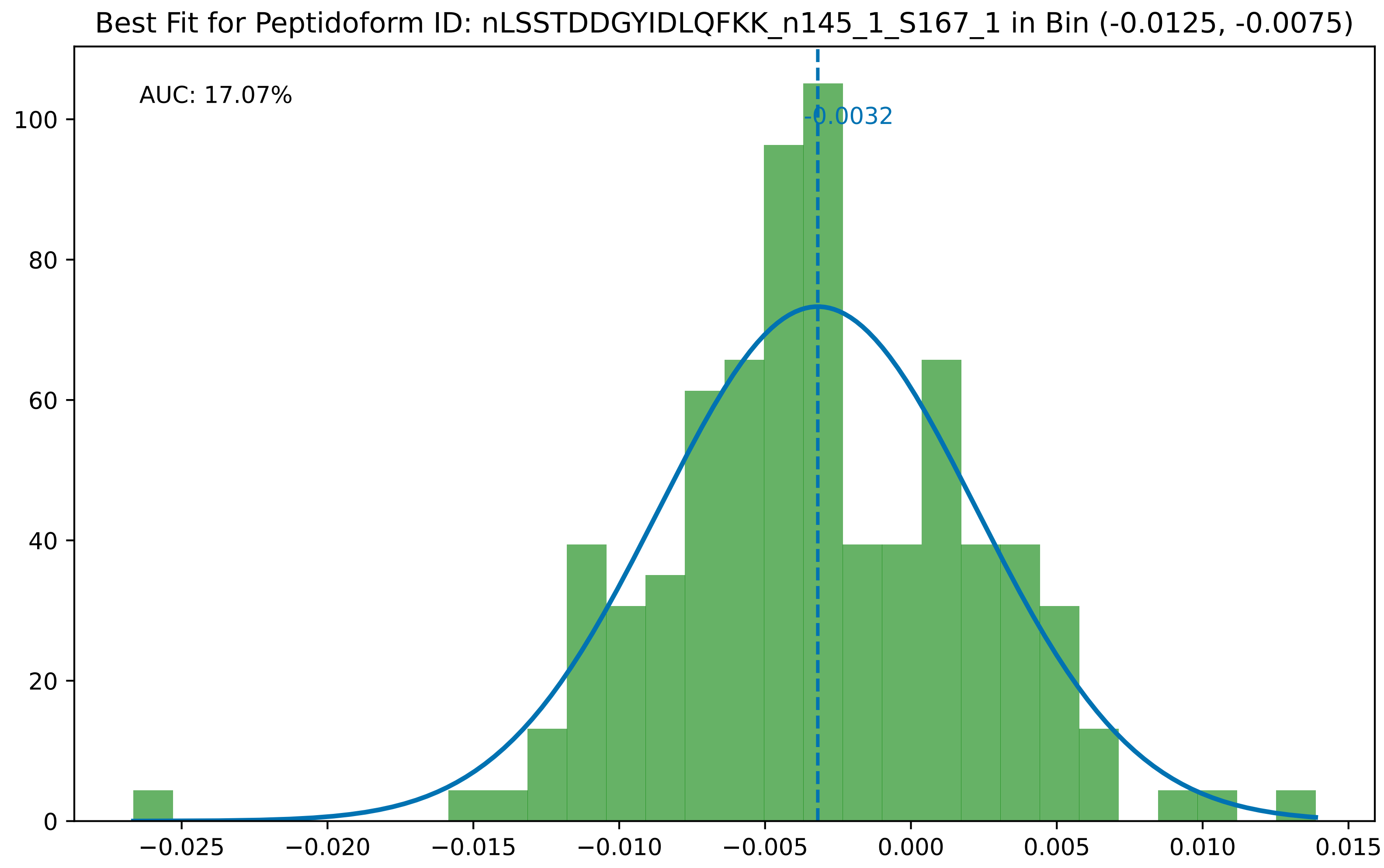

Best Fit for Peptidoform ID: nLSSTDDGYIDLQFKK_n145_1_S167_1 in Bin (-0.0125, -0.0075)
AUC: 17.07%
-0.0032
0
20
40
60
80
100
−0.025
−0.020
−0.015
−0.010
−0.005
0.000
0.005
0.010
0.015

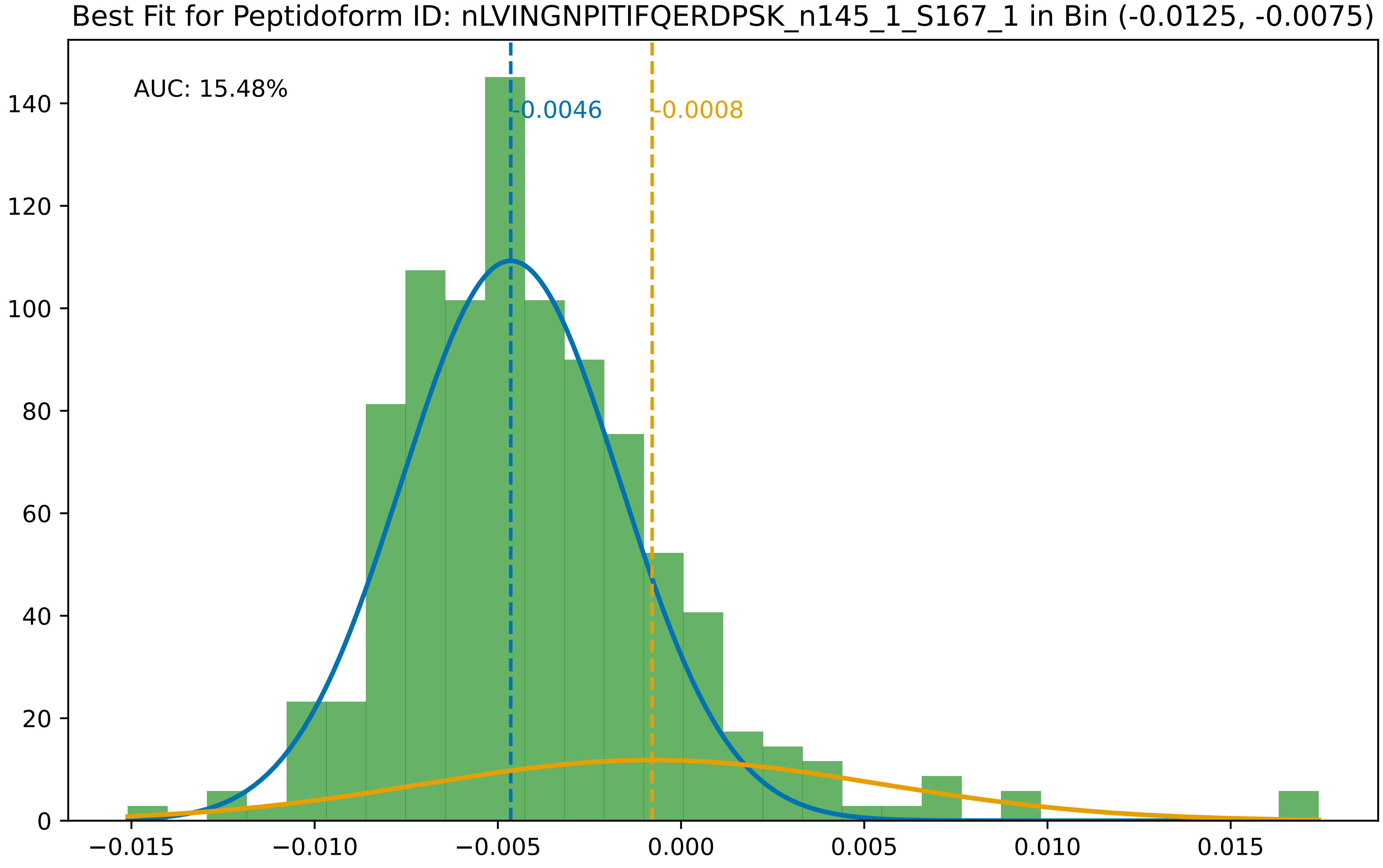

Best Fit for Peptidoform ID: nLVINGNPITIFQERDPSK_n145_1_S167_1 in Bin (-0.0125, -0.0075)
AUC: 15.48%
-0.0046
-0.0008
0
20
40
60
80
100
120
140
−0.015
−0.010
−0.005
0.000
0.005
0.010
0.015

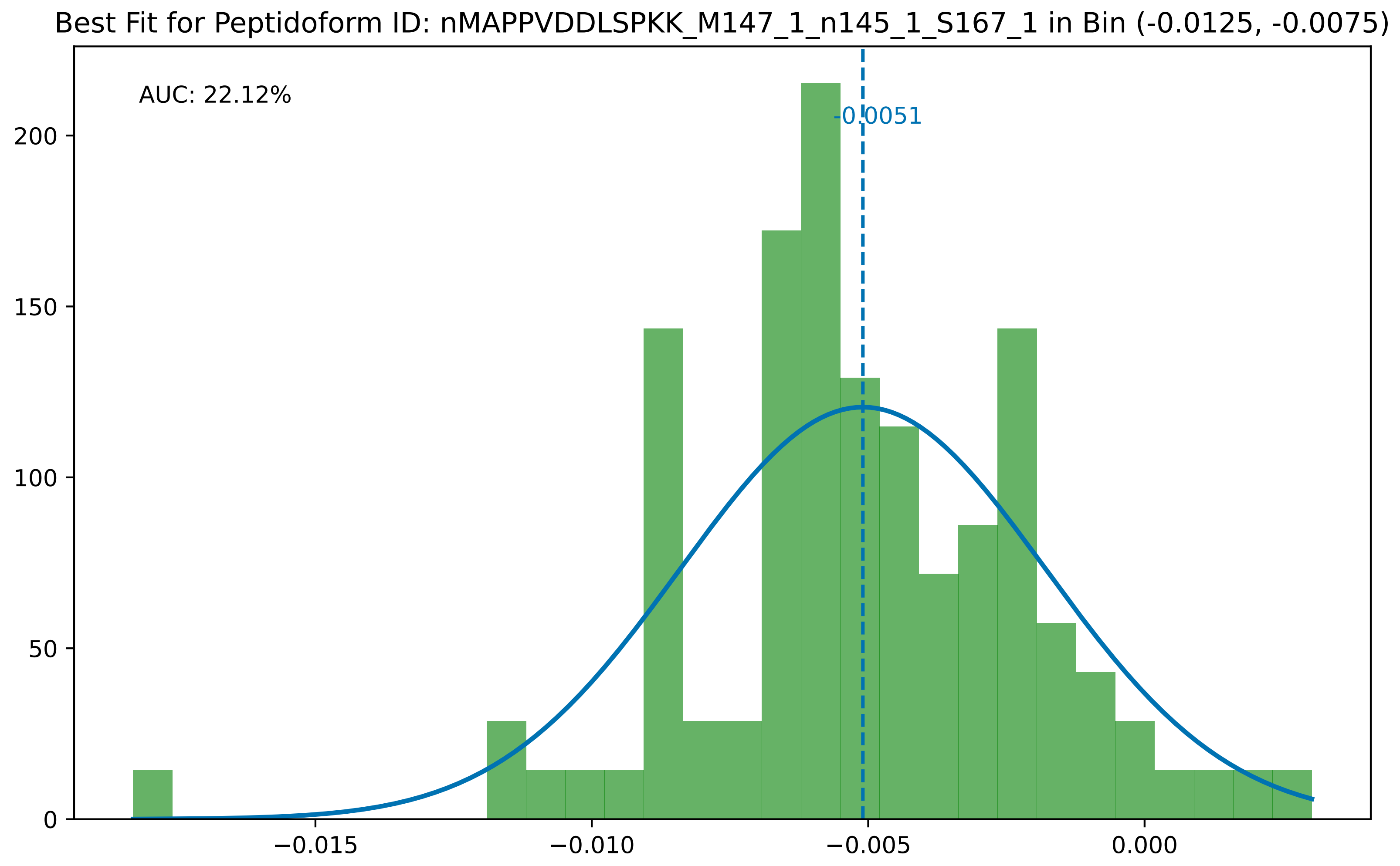

Best Fit for Peptidoform ID: nMAPPVDDLSPKK_M147_1_n145_1_S167_1 in Bin (-0.0125, -0.0075)
AUC: 22.12%
-0.0051
200
150
100
50
0
−0.015
−0.010
−0.005
0.000

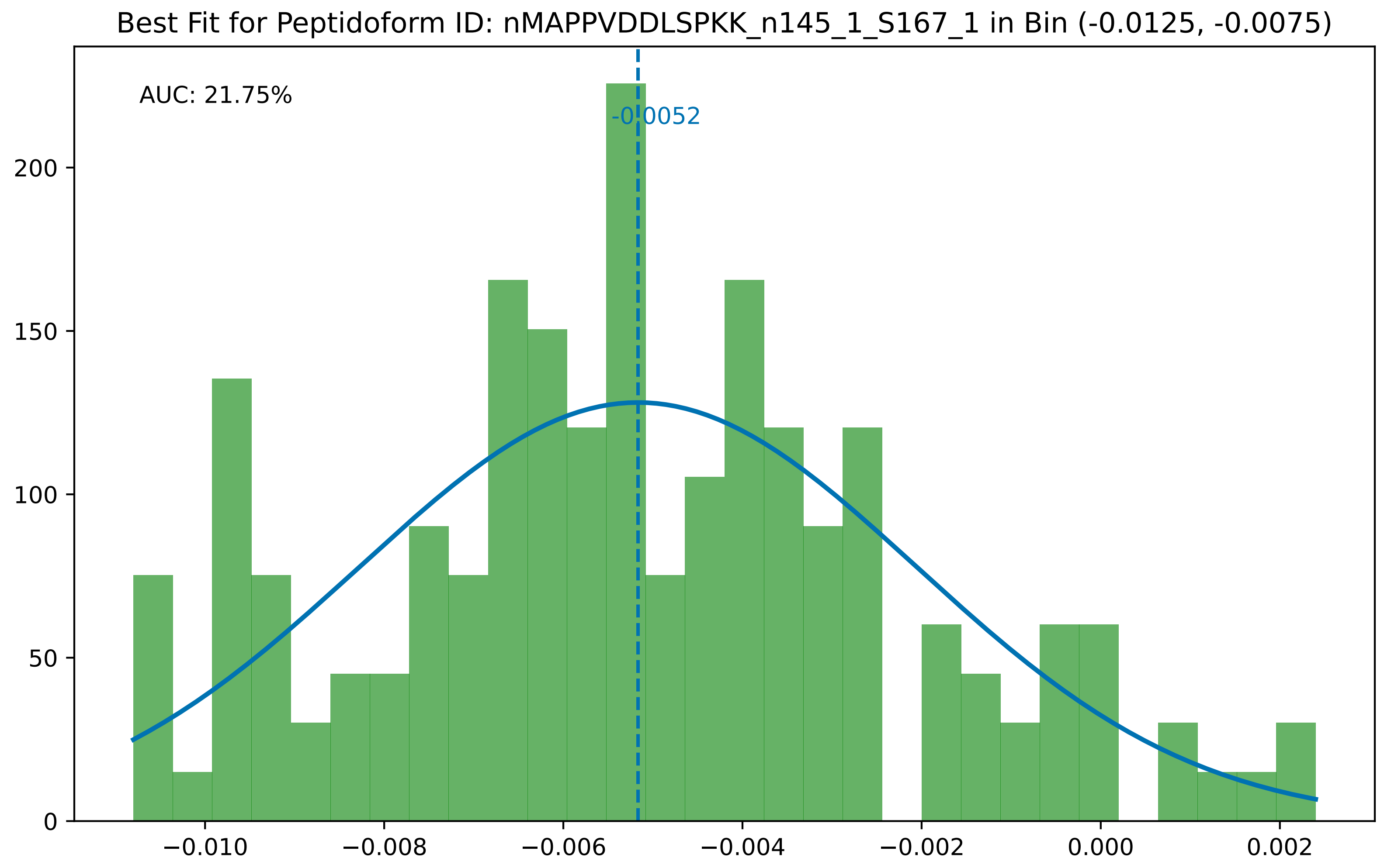
Best Fit for Peptidoform ID: nMAPPVDDLSPKK_n145_1_S167_1 in Bin (-0.0125, -0.0075)
AUC: 21.75%
-0.0052
0
50
100
150
200
−0.010
−0.008
−0.006
−0.004
−0.002
0.000
0.002

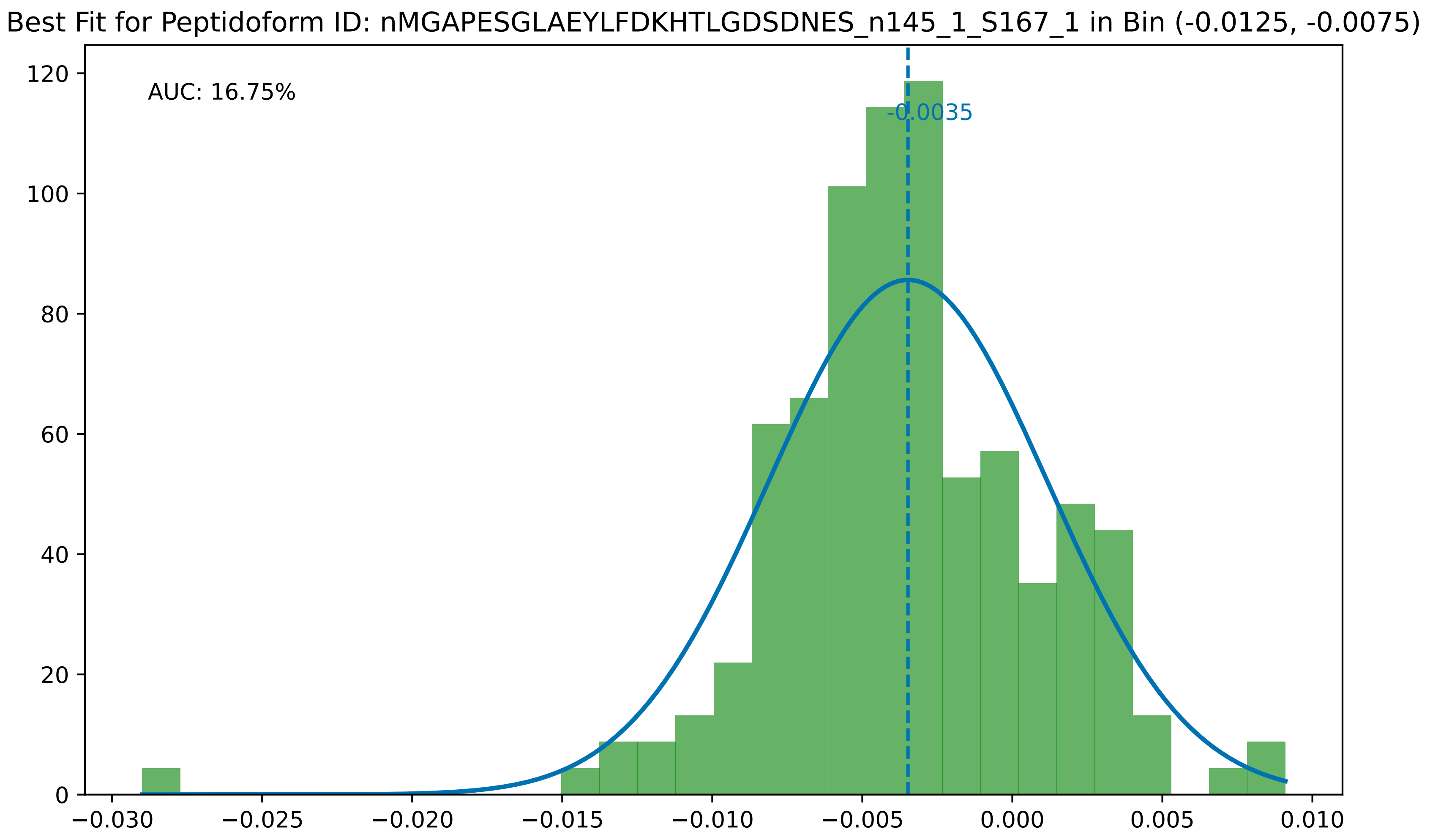

Best Fit for Peptidoform ID: nMGAPESGLAEYLFDKHTLGDSDNES_n145_1_S167_1 in Bin (-0.0125, -0.0075)
AUC: 16.75%
-0.0035
0
20
40
60
80
100
120
−0.030
−0.025
−0.020
−0.015
−0.010
−0.005
0.000
0.005
0.010

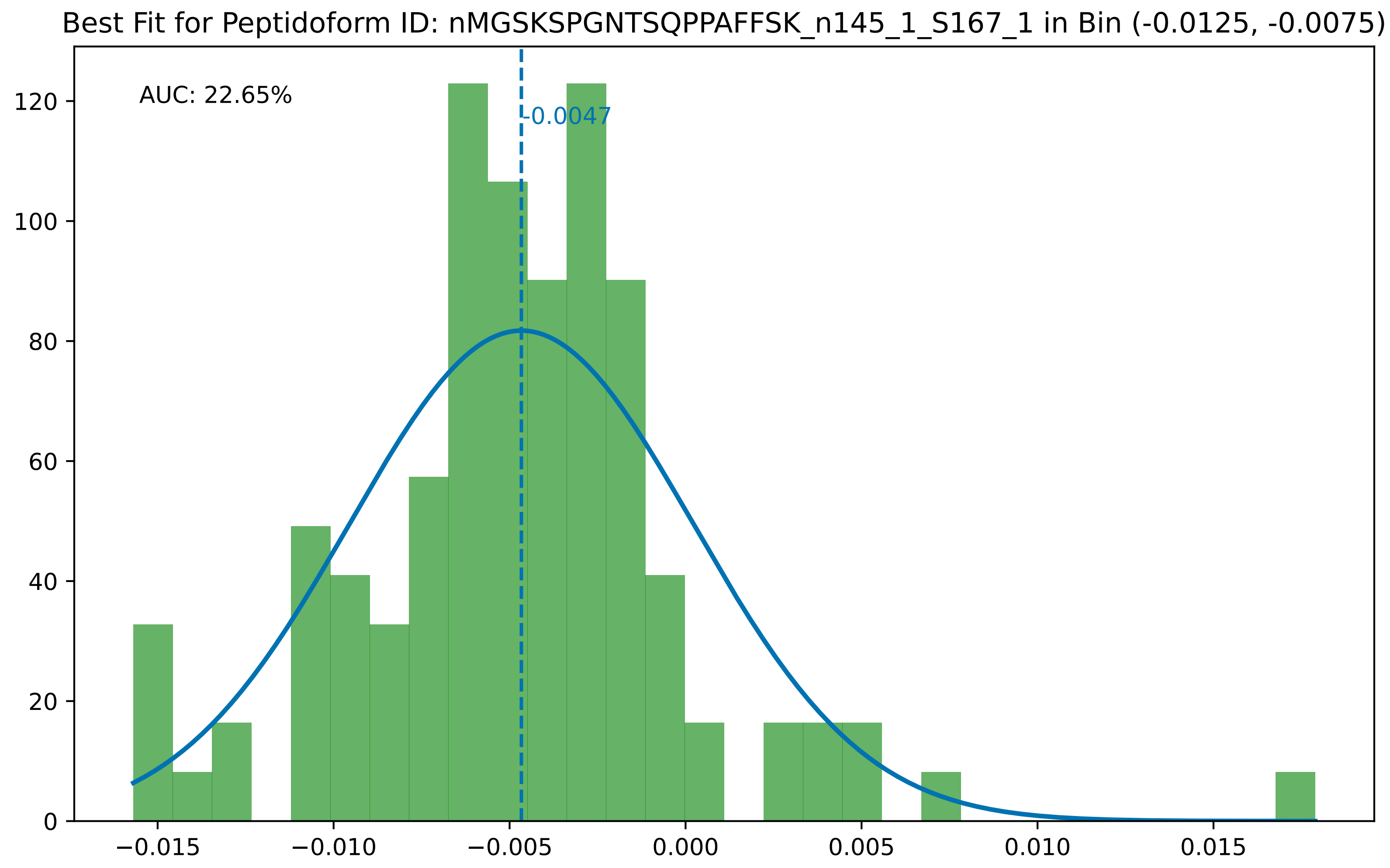
Best Fit for Peptidoform ID: nMGSKSPGNTSQPPAFFSK_n145_1_S167_1 in Bin (-0.0125, -0.0075)
AUC: 22.65%
-0.0047
0
20
40
60
80
100
120
−0.015
−0.010
−0.005
0.000
0.005
0.010
0.015

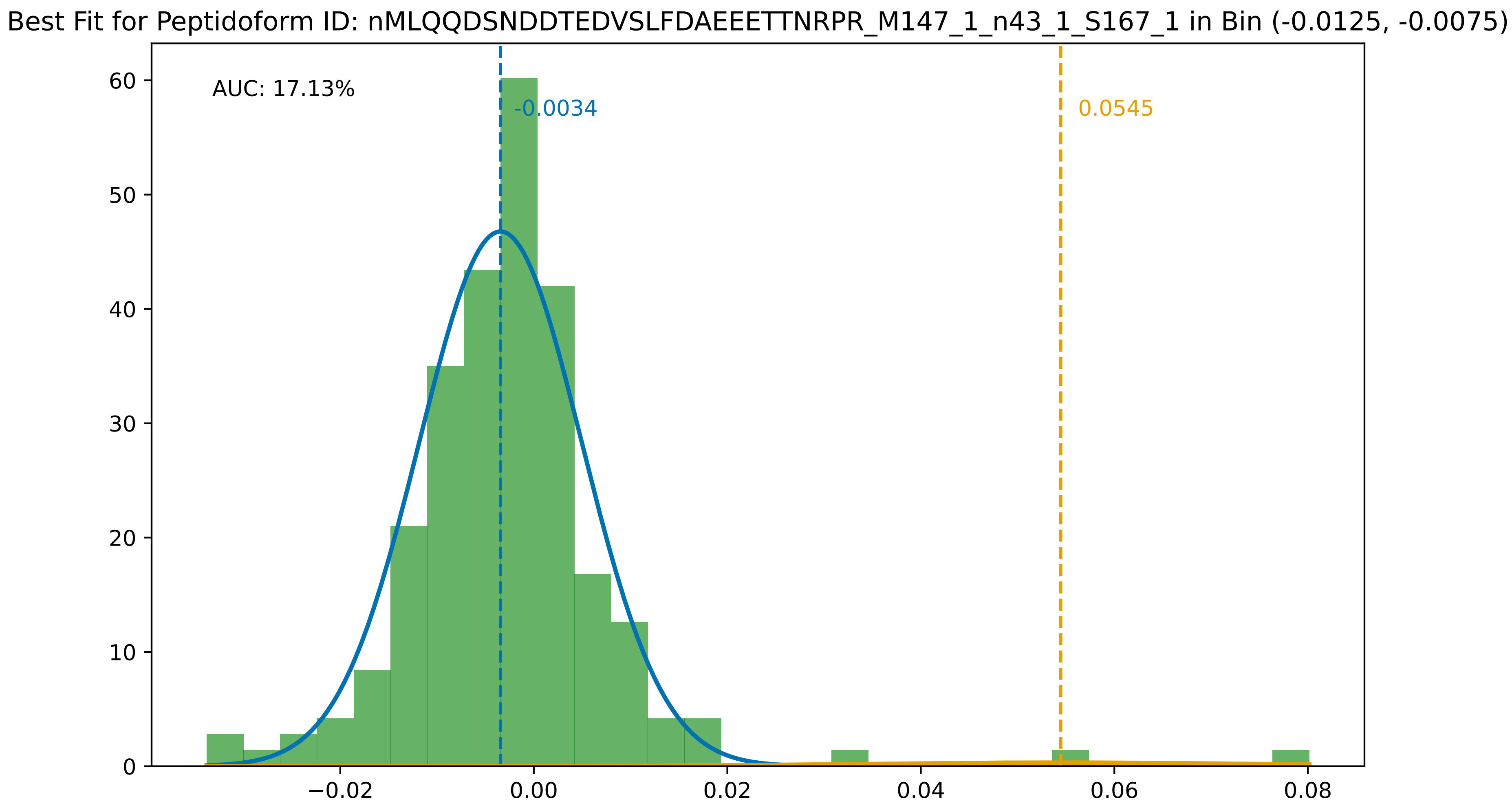

Best Fit for Peptidoform ID: nMLQQDSNDDTEDVSLFDAEEETTNRPR_M147_1_n43_1_S167_1 in Bin (-0.0125, -0.0075)
AUC: 17.13%
-0.0034
0.0545
0
10
20
30
40
50
60
−0.02
0.00
0.02
0.04
0.06
0.08

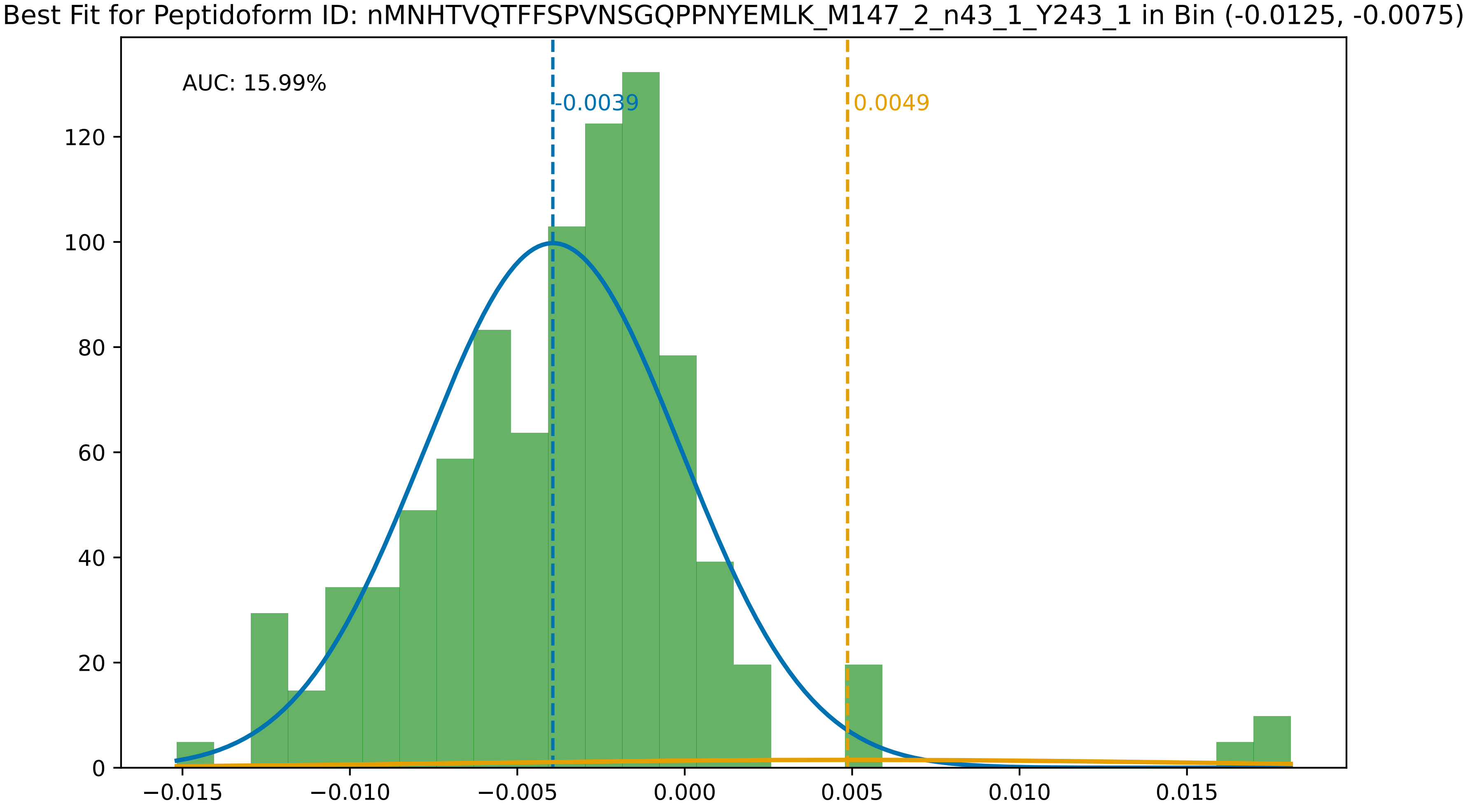
Best Fit for Peptidoform ID: nMNHTVQTFFSPVNSGQPPNYEMLK_M147_2_n43_1_Y243_1 in Bin (-0.0125, -0.0075)
AUC: 15.99%
-0.0039
0.0049
0
20
40
60
80
100
120
−0.015
−0.010
−0.005
0.000
0.005
0.010
0.015

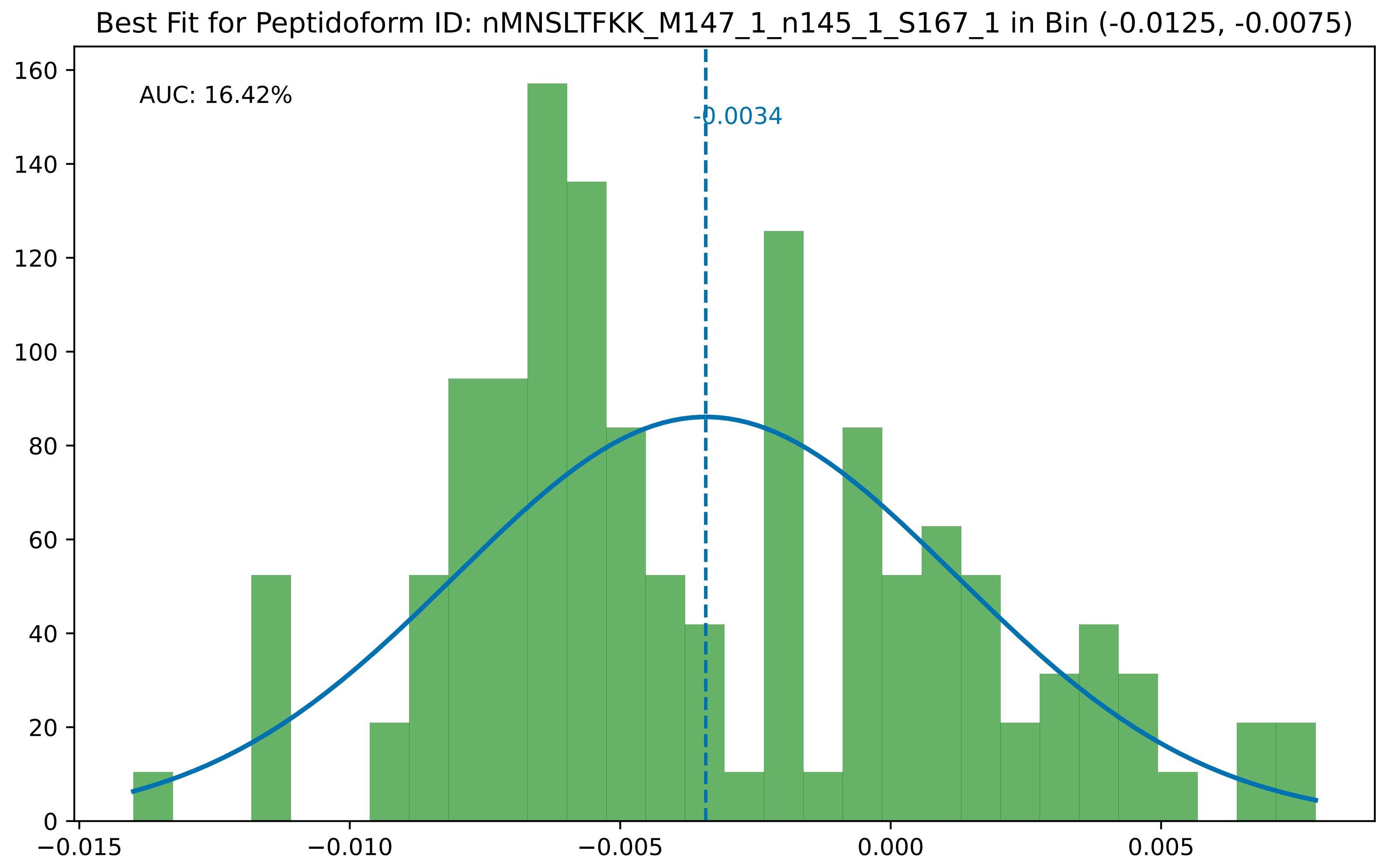

Best Fit for Peptidoform ID: nMNSLTFKK_M147_1_n145_1_S167_1 in Bin (-0.0125, -0.0075)
AUC: 16.42%
-0.0034
0
20
40
60
80
100
120
140
160
−0.015
−0.010
−0.005
0.000
0.005

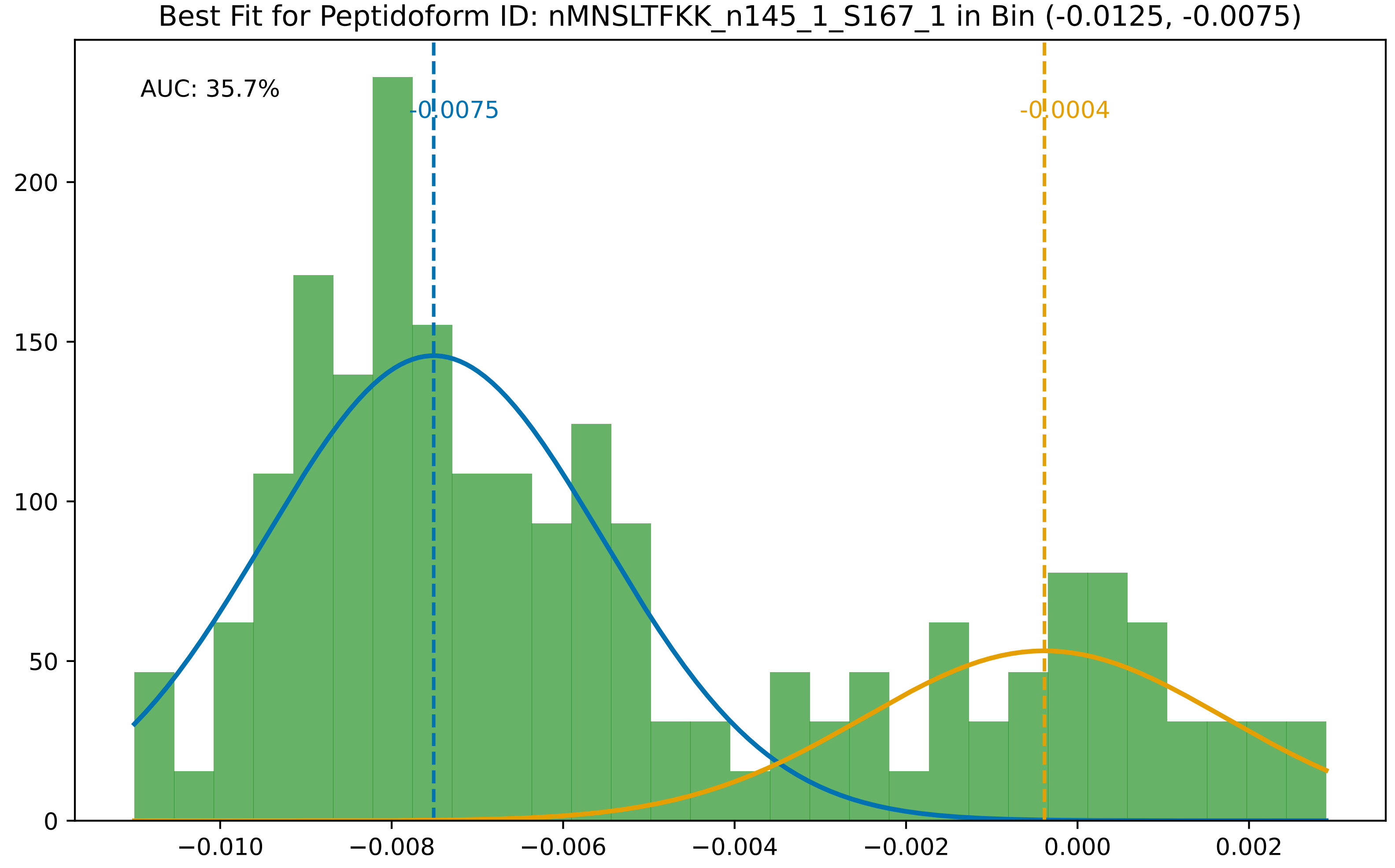
Best Fit for Peptidoform ID: nMNSLTFKK_n145_1_S167_1 in Bin (-0.0125, -0.0075)
AUC: 35.7%
-0.0075
-0.0004
0
50
100
150
200
−0.010
−0.008
−0.006
−0.004
−0.002
0.000
0.002

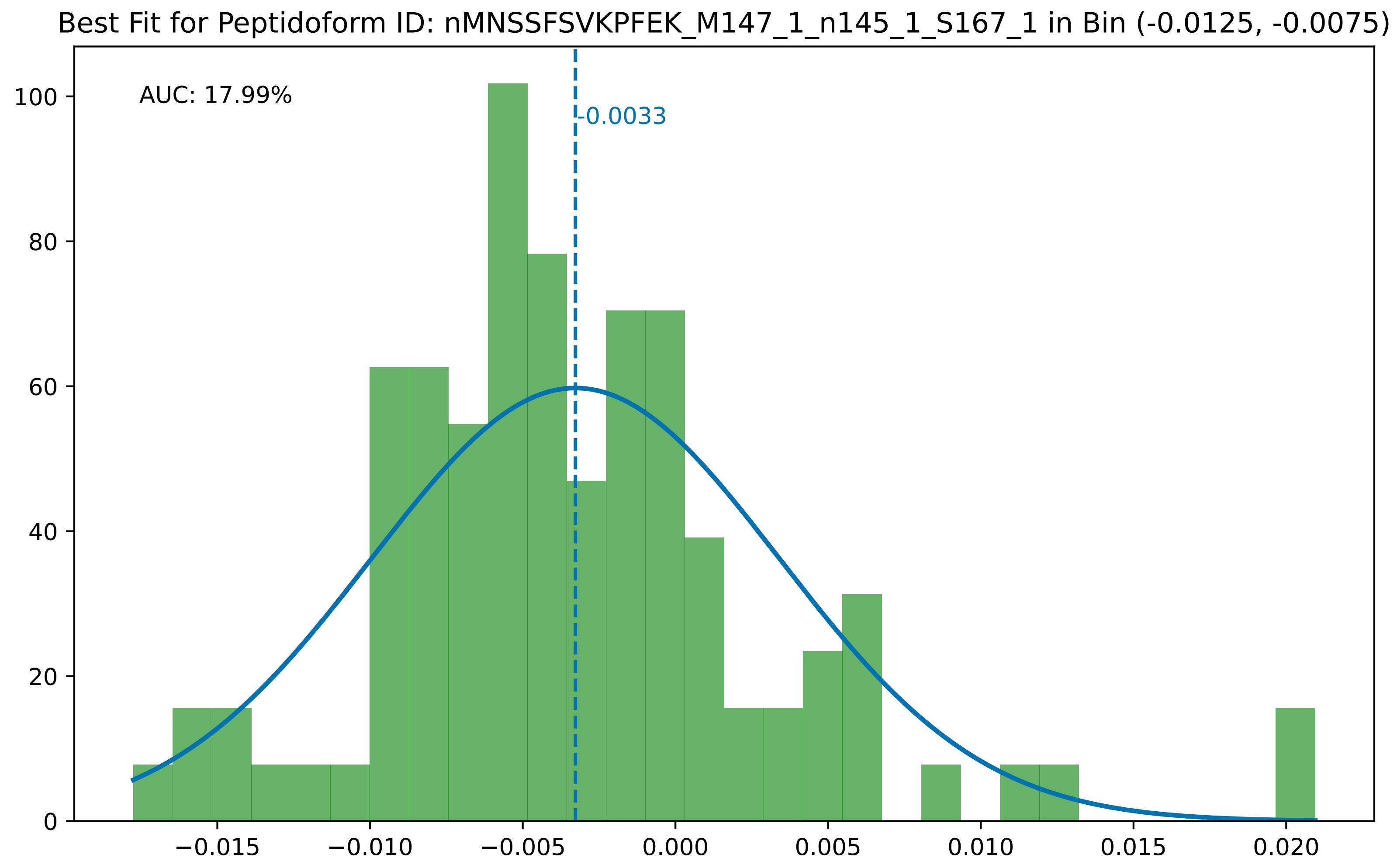

Best Fit for Peptidoform ID: nMNSSFSVKPFEK_M147_1_n145_1_S167_1 in Bin (-0.0125, -0.0075)
AUC: 17.99%
-0.0033
0
20
40
60
80
100
−0.015
−0.010
−0.005
0.000
0.005
0.010
0.015
0.020

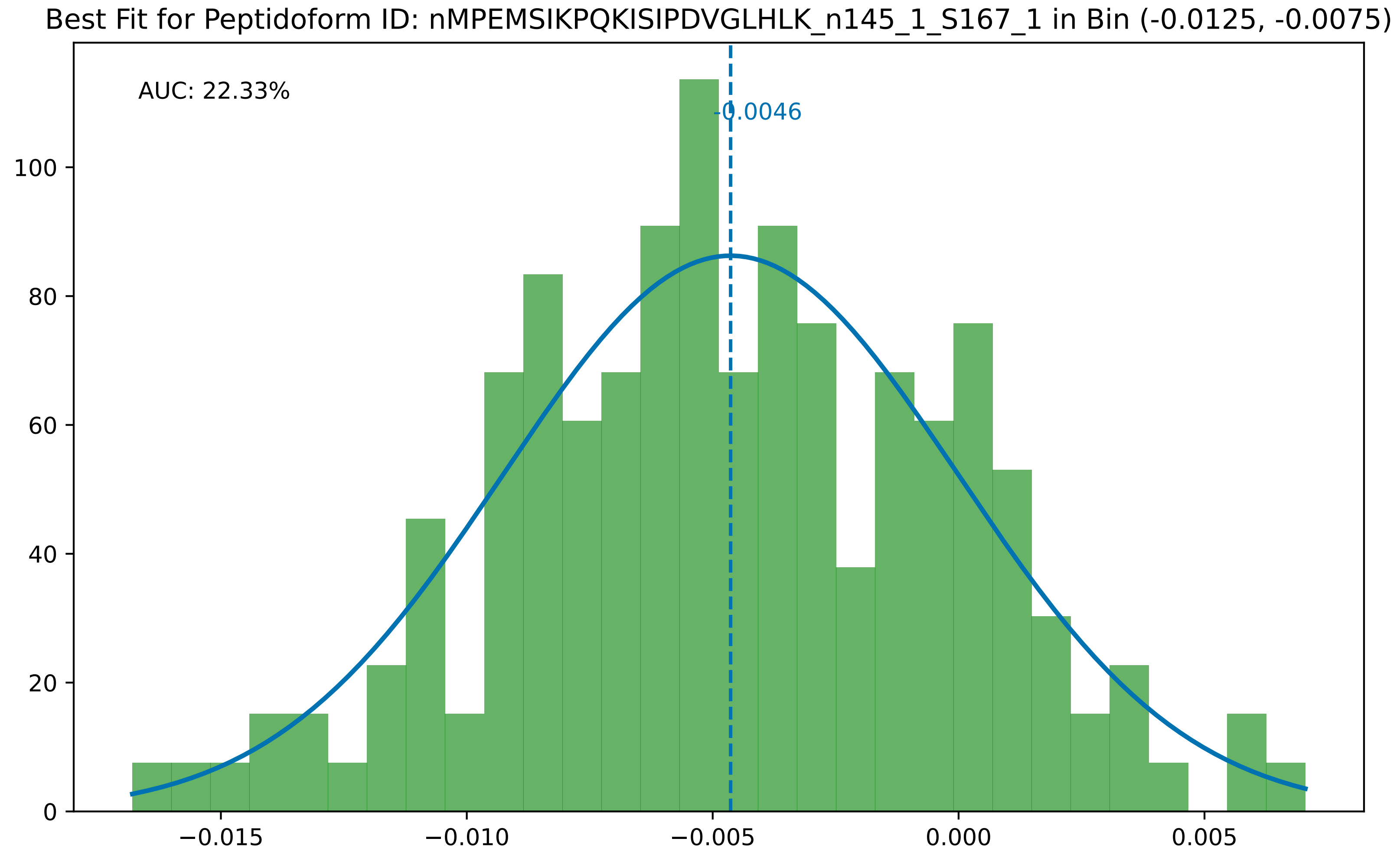

Best Fit for Peptidoform ID: nMPEMSIKPQKISIPDVGLHLK_n145_1_S167_1 in Bin (-0.0125, -0.0075)
AUC: 22.33%
-0.0046
0
20
40
60
80
100
−0.015
−0.010
−0.005
0.000
0.005

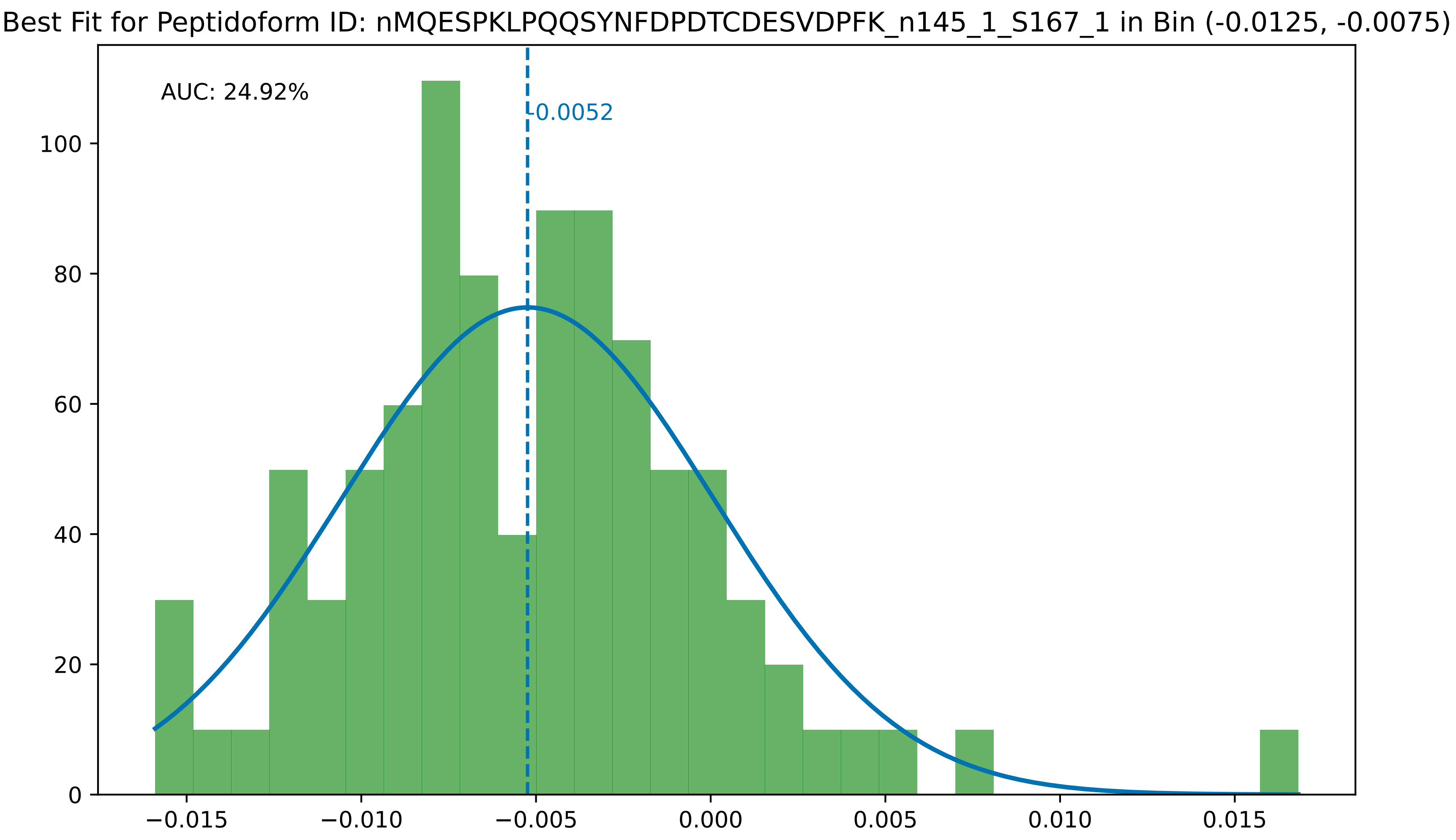

Best Fit for Peptidoform ID: nMQESPKLPQQSYNFDPDTCDESVDPFK_n145_1_S167_1 in Bin (-0.0125, -0.0075)
AUC: 24.92%
-0.0052
0
20
40
60
80
100
−0.015
−0.010
−0.005
0.000
0.005
0.010
0.015

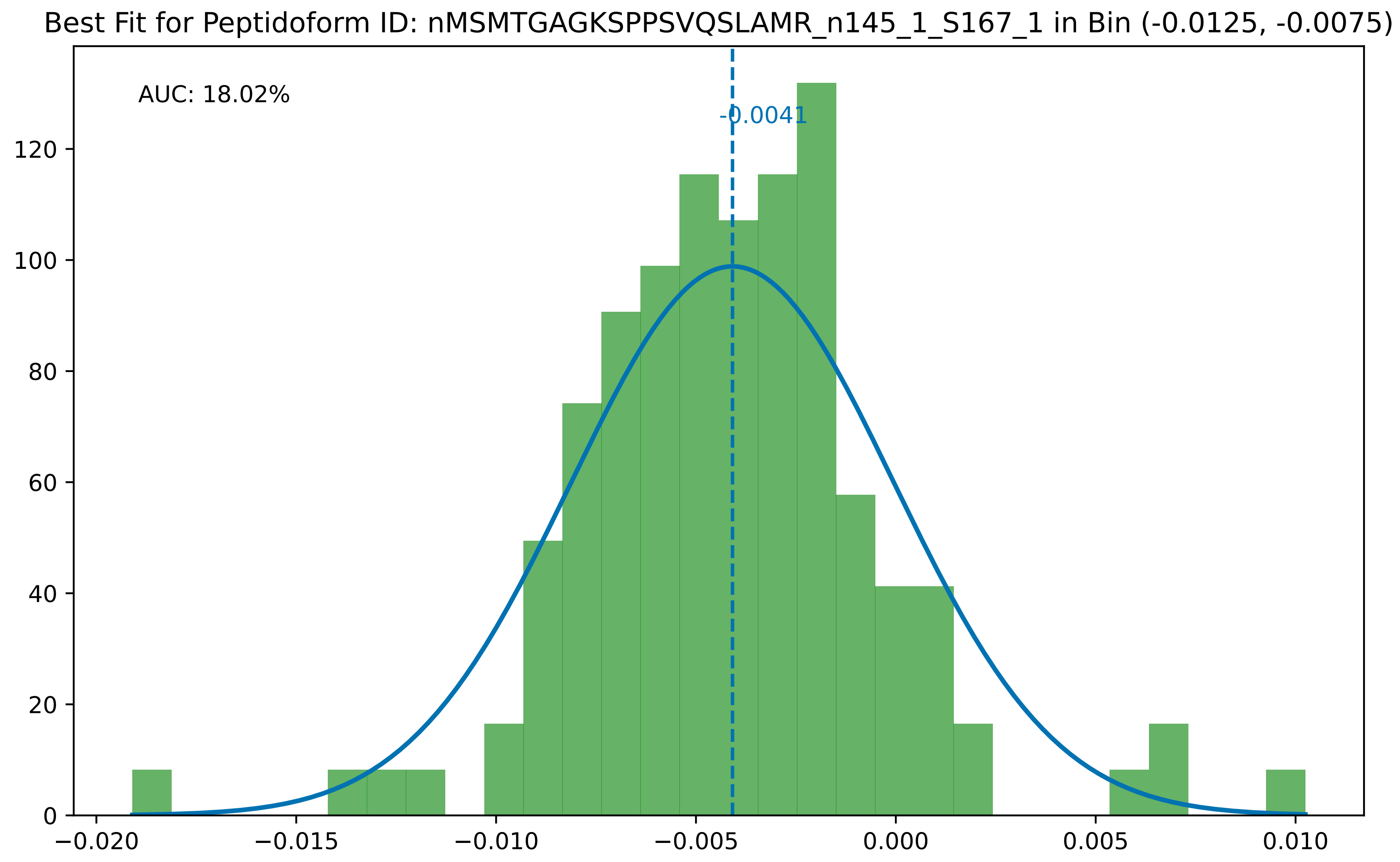
Best Fit for Peptidoform ID: nMSMTGAGKSPPSVQSLAMR_n145_1_S167_1 in Bin (-0.0125, -0.0075)
AUC: 18.02%
-0.0041
0
20
40
60
80
100
120
−0.020
−0.015
−0.010
−0.005
0.000
0.005
0.010

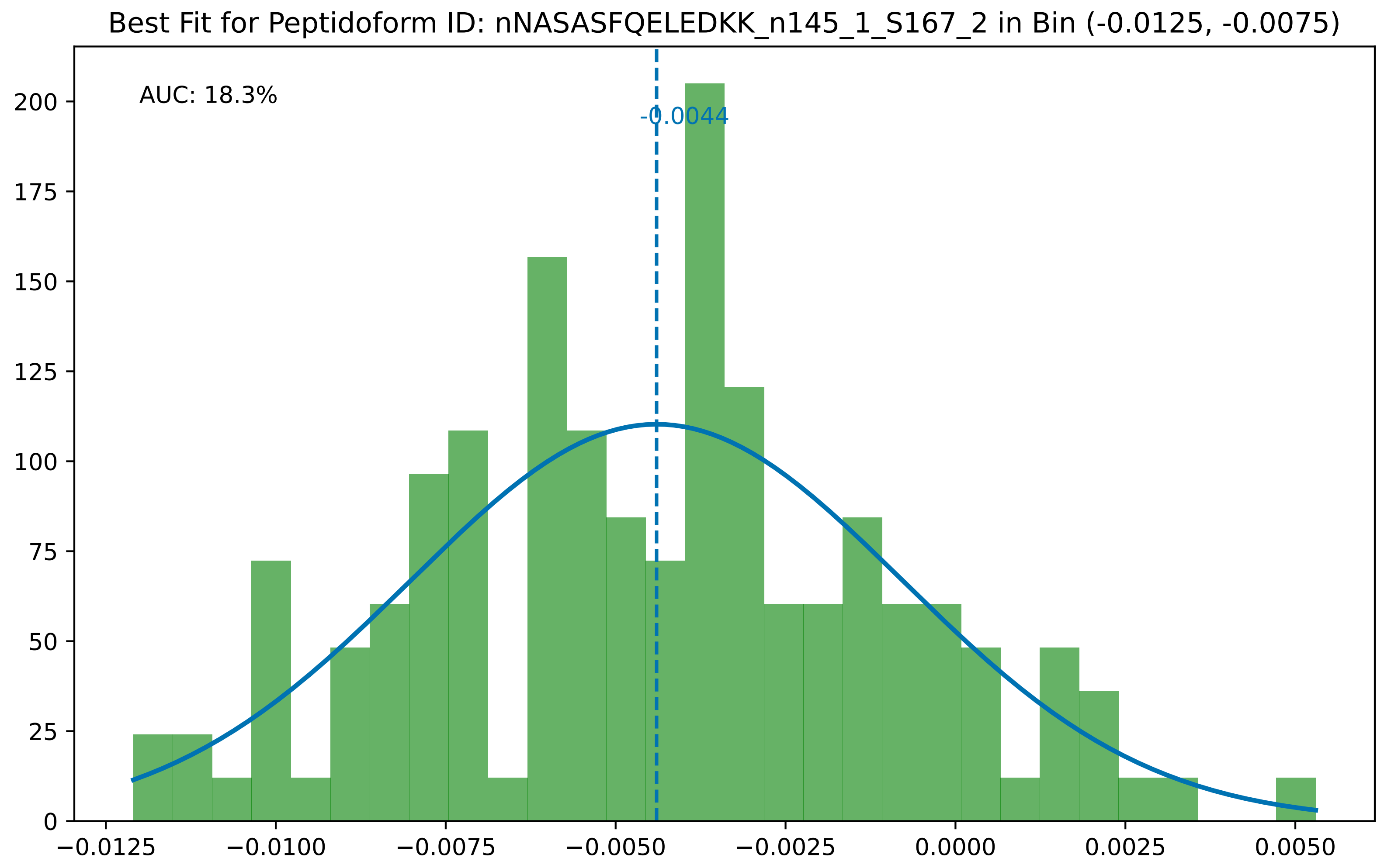

Best Fit for Peptidoform ID: nNASASFQELEDKK_n145_1_S167_2 in Bin (-0.0125, -0.0075)
AUC: 18.3%
-0.0044
0
25
50
75
100
125
150
175
200
−0.0125
−0.0100
−0.0075
−0.0050
−0.0025
0.0000
0.0025
0.0050

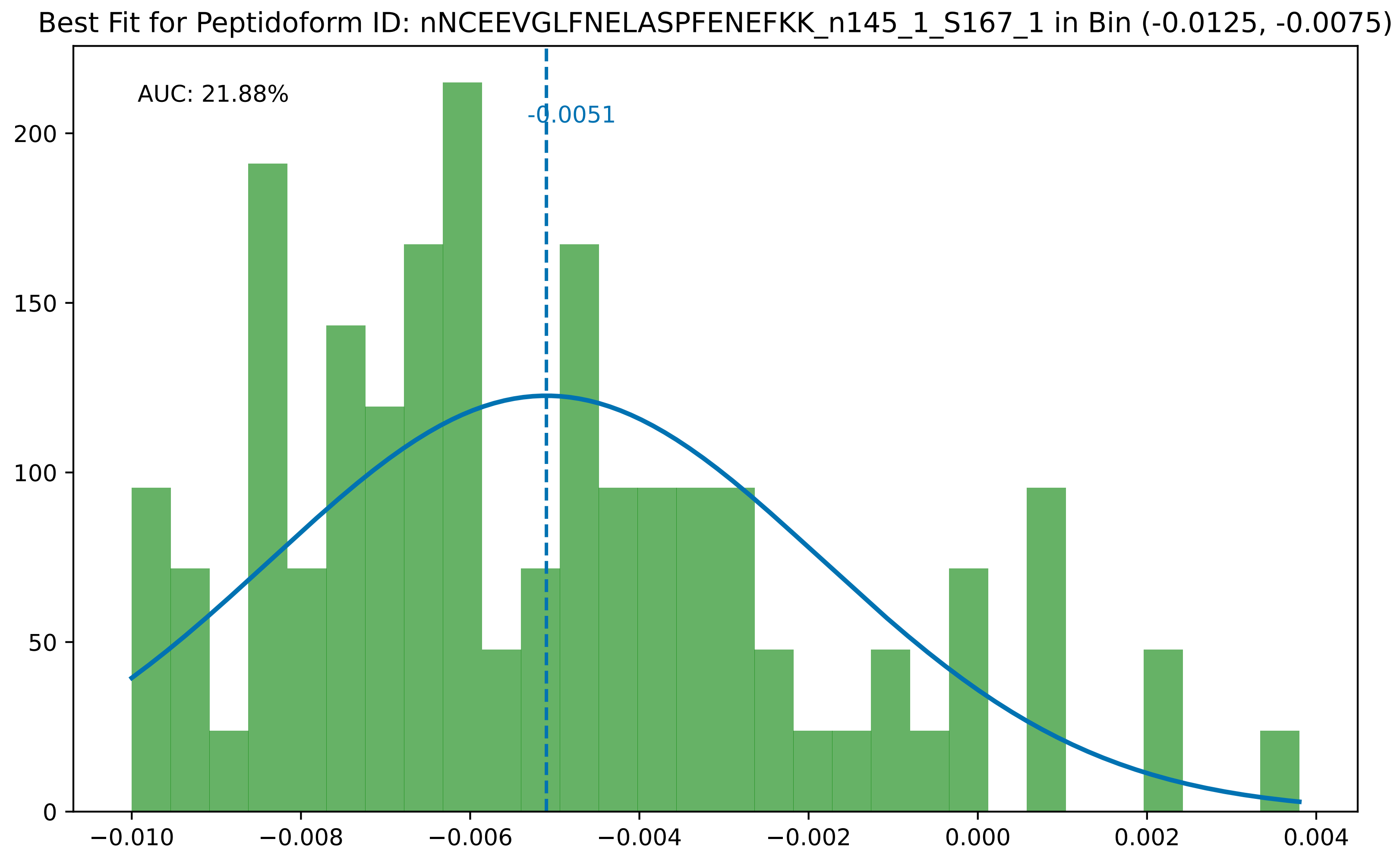

Best Fit for Peptidoform ID: nNCEEVGLFNELASPFENEFKK_n145_1_S167_1 in Bin (-0.0125, -0.0075)
AUC: 21.88%
-0.0051
0
50
100
150
200
−0.010
−0.008
−0.006
−0.004
−0.002
0.000
0.002
0.004

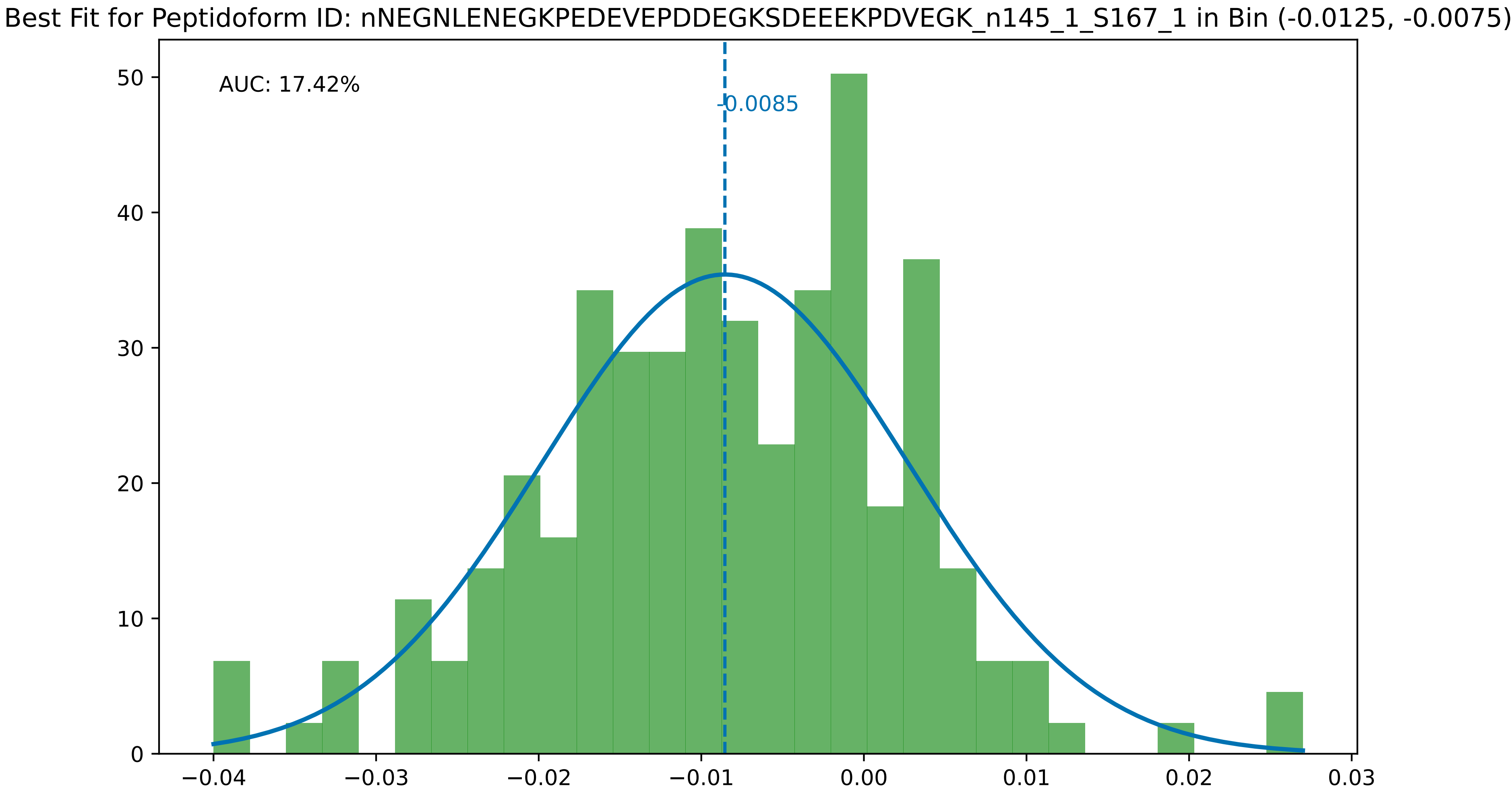
Best Fit for Peptidoform ID: nNEGNLENEGKPEDEVEPDDEGKSDEEEKPDVEGK_n145_1_S167_1 in Bin (-0.0125, -0.0075)
AUC: 17.42%
-0.0085
0
10
20
30
40
50
−0.04
−0.03
−0.02
−0.01
0.00
0.01
0.02
0.03

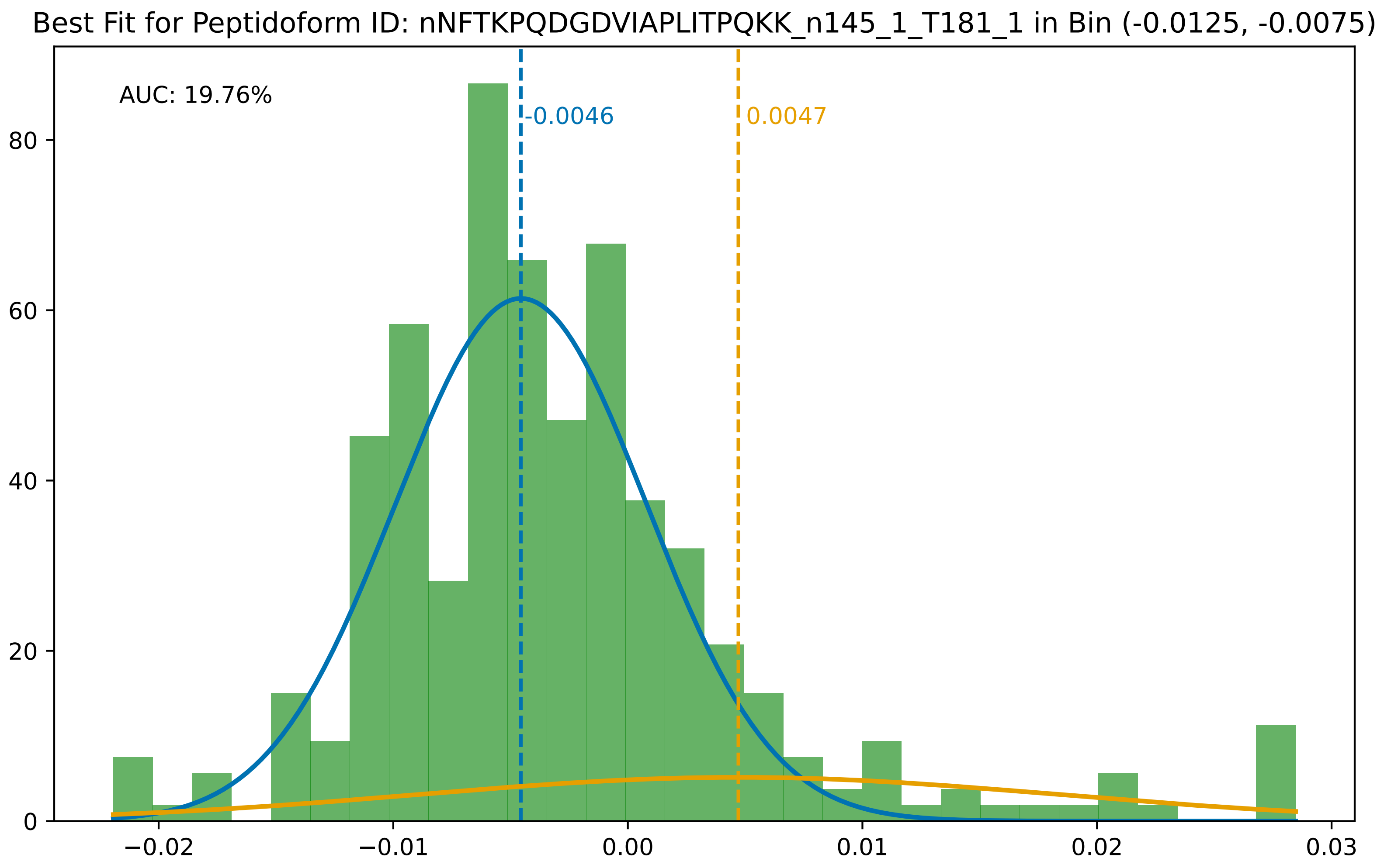

Best Fit for Peptidoform ID: nNFTKPQDGDVIAPLITPQKK_n145_1_T181_1 in Bin (-0.0125, -0.0075)
AUC: 19.76%
-0.0046
0.0047
0
20
40
60
80
−0.02
−0.01
0.00
0.01
0.02
0.03

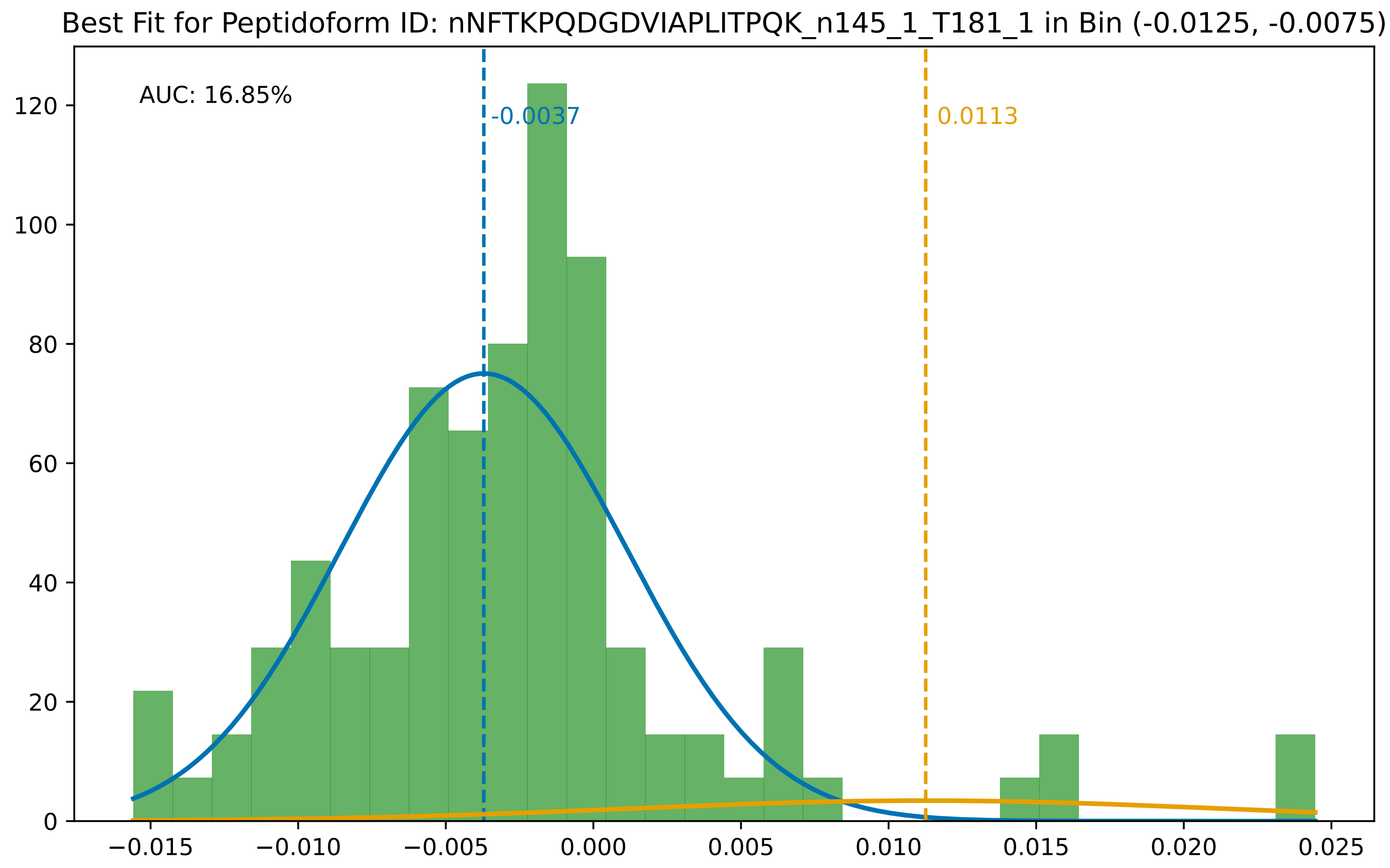
Best Fit for Peptidoform ID: nNFTKPQDGDVIAPLITPQK_n145_1_T181_1 in Bin (-0.0125, -0.0075)
AUC: 16.85%
-0.0037
0.0113
0
20
40
60
80
100
120
−0.015
−0.010
−0.005
0.000
0.005
0.010
0.015
0.020
0.025

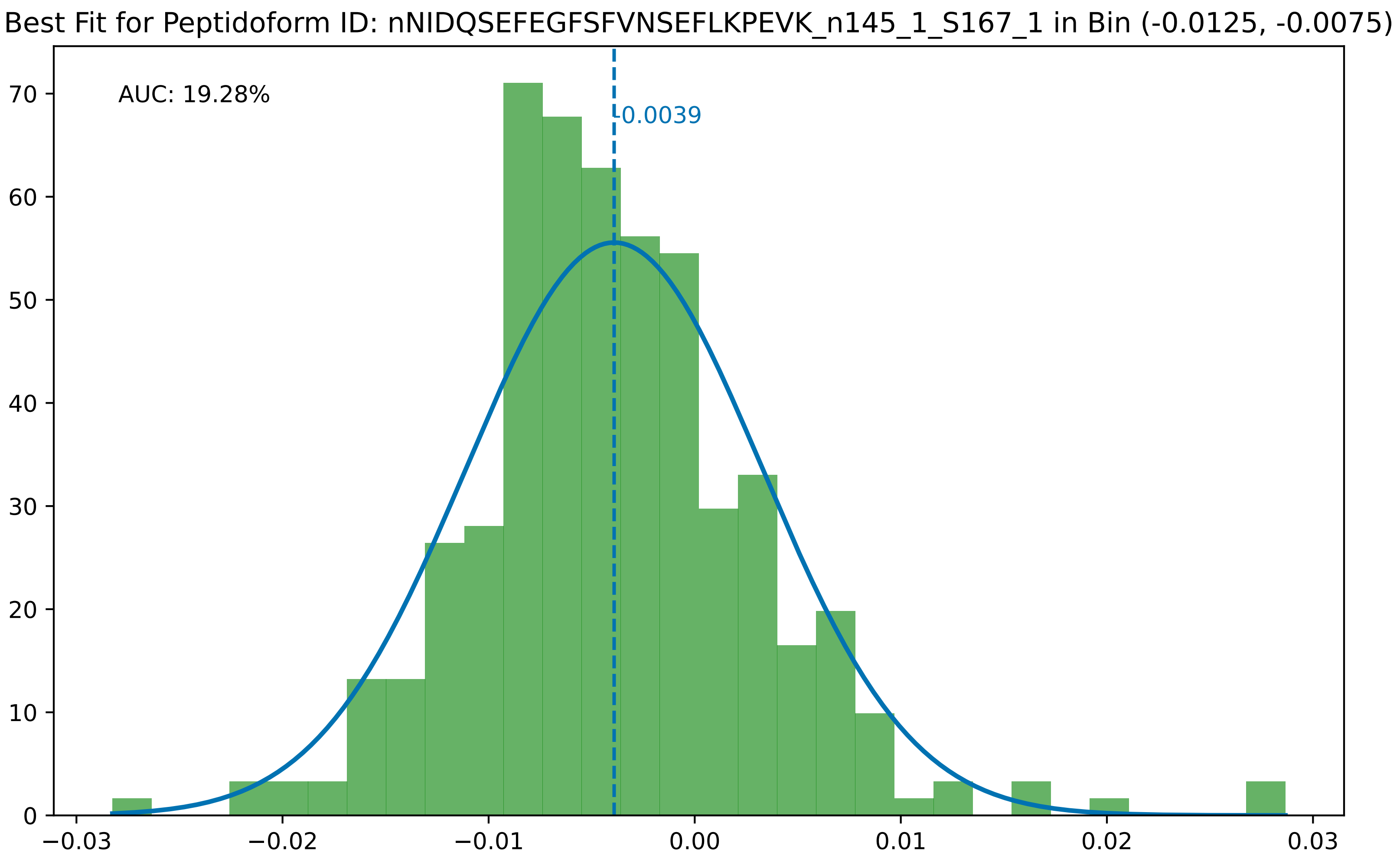

Best Fit for Peptidoform ID: nNIDQSEFEGFSFVNSEFLKPEVK_n145_1_S167_1 in Bin (-0.0125, -0.0075)
AUC: 19.28%
-0.0039
0
10
20
30
40
50
60
70
−0.03
−0.02
−0.01
0.00
0.01
0.02
0.03

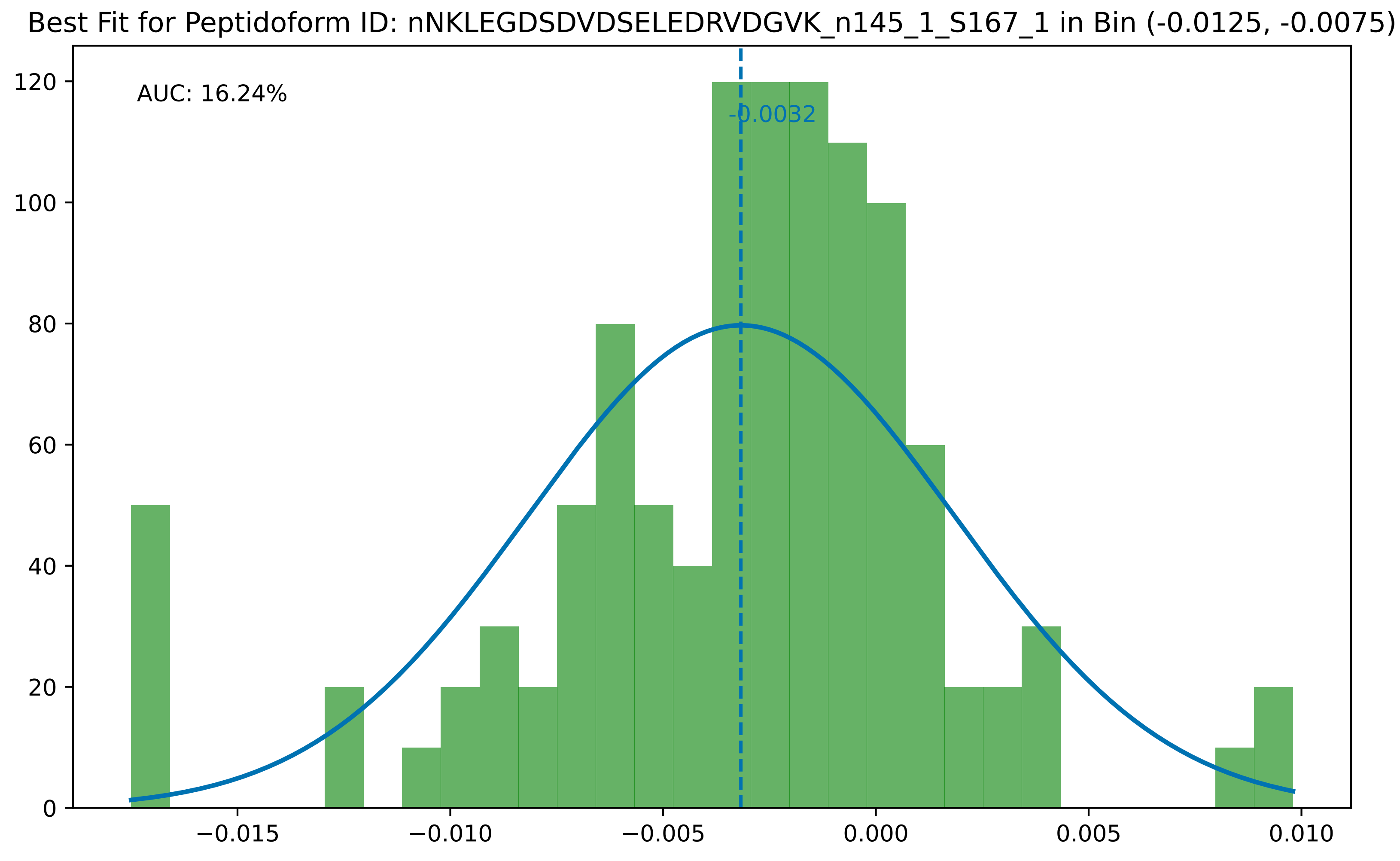

Best Fit for Peptidoform ID: nNKLEGDSDVDSELEDRVDGVK_n145_1_S167_1 in Bin (-0.0125, -0.0075)
AUC: 16.24%
-0.0032
0
20
40
60
80
100
120
−0.015
−0.010
−0.005
0.000
0.005
0.010

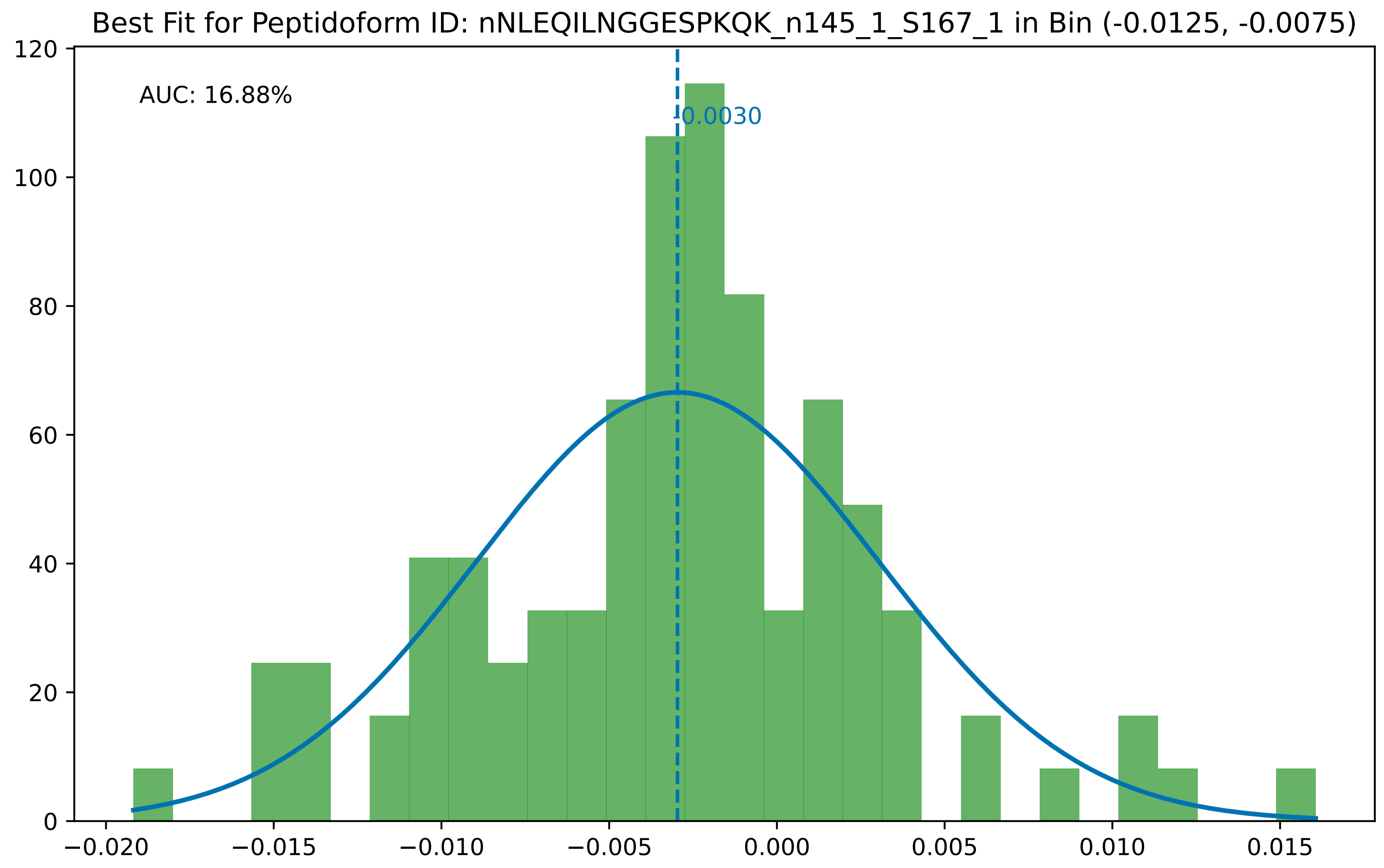

Best Fit for Peptidoform ID: nNLEQILNGGESPKQK_n145_1_S167_1 in Bin (-0.0125, -0.0075)
AUC: 16.88%
-0.0030
0
20
40
60
80
100
120
−0.020
−0.015
−0.010
−0.005
0.000
0.005
0.010
0.015

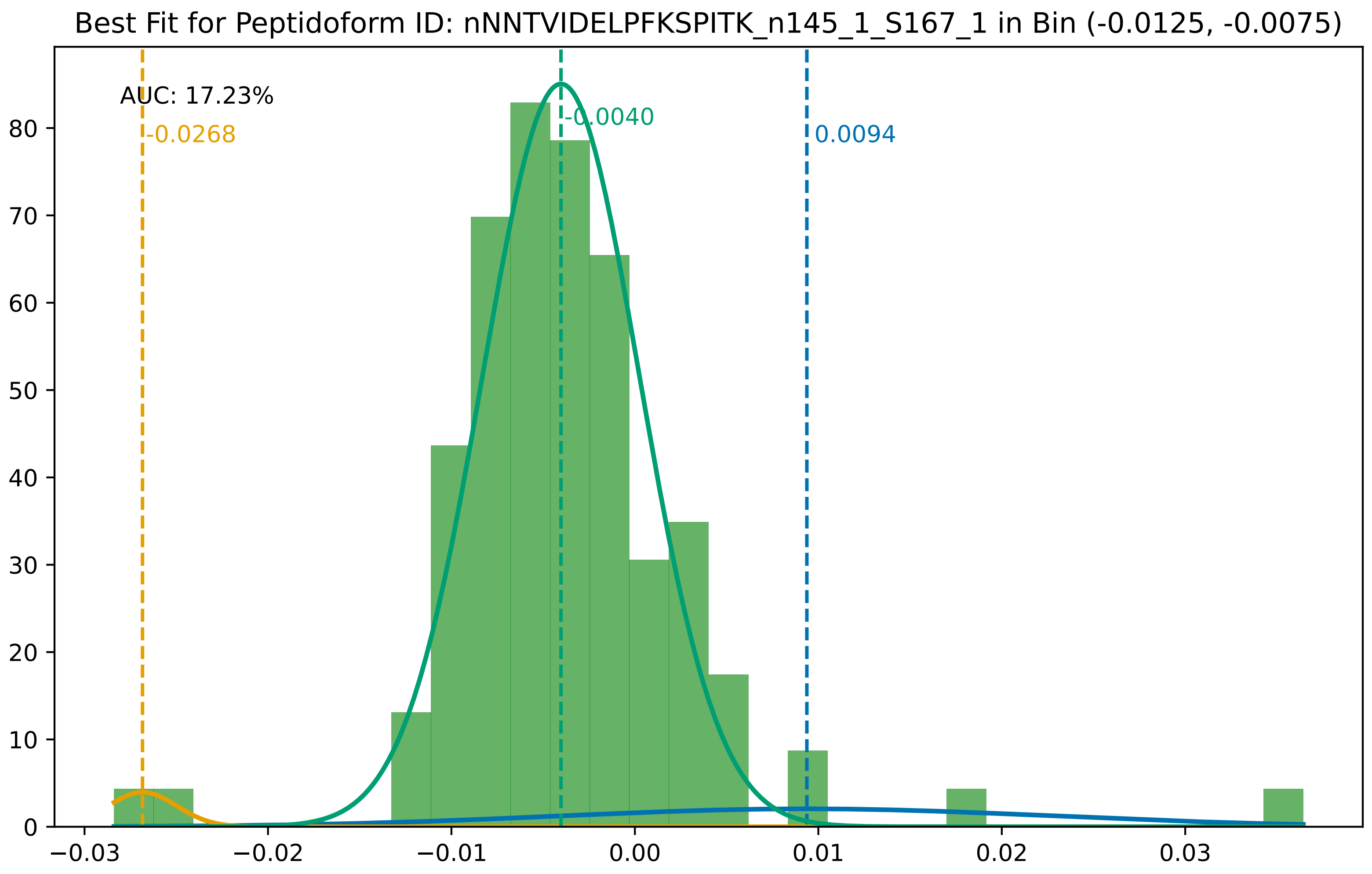

Best Fit for Peptidoform ID: nNNTVIDELPFKSPITK_n145_1_S167_1 in Bin (-0.0125, -0.0075)
AUC: 17.23%
-0.0268
-0.0040
0.0094
0
10
20
30
40
50
60
70
80
−0.03
−0.02
−0.01
0.00
0.01
0.02
0.03

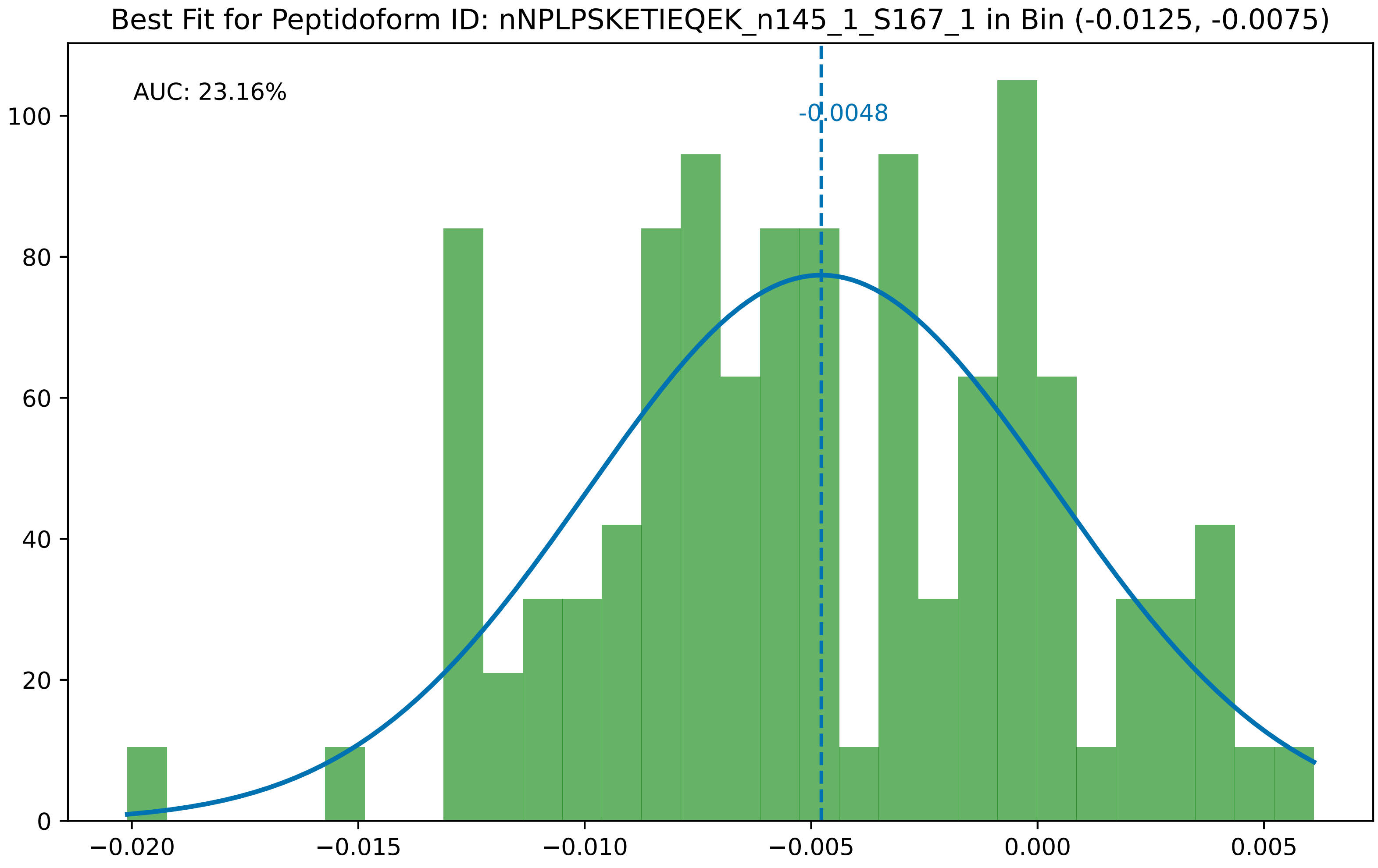

Best Fit for Peptidoform ID: nNPLPSKETIEQEK_n145_1_S167_1 in Bin (-0.0125, -0.0075)
AUC: 23.16%
-0.0048
0
20
40
60
80
100
−0.020
−0.015
−0.010
−0.005
0.000
0.005

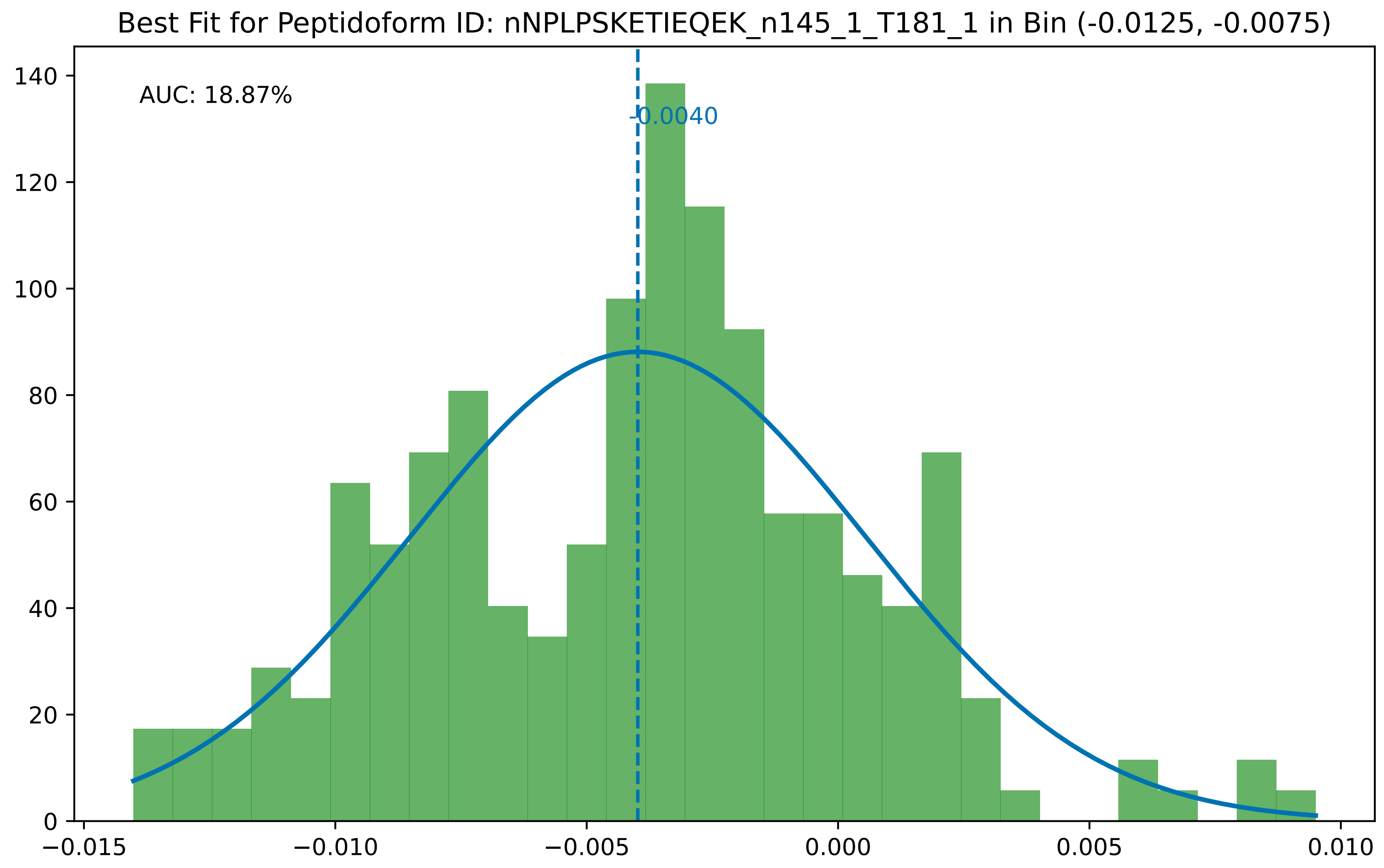
Best Fit for Peptidoform ID: nNPLPSKETIEQEK_n145_1_T181_1 in Bin (-0.0125, -0.0075)
AUC: 18.87%
-0.0040
0
20
40
60
80
100
120
140
−0.015
−0.010
−0.005
0.000
0.005
0.010

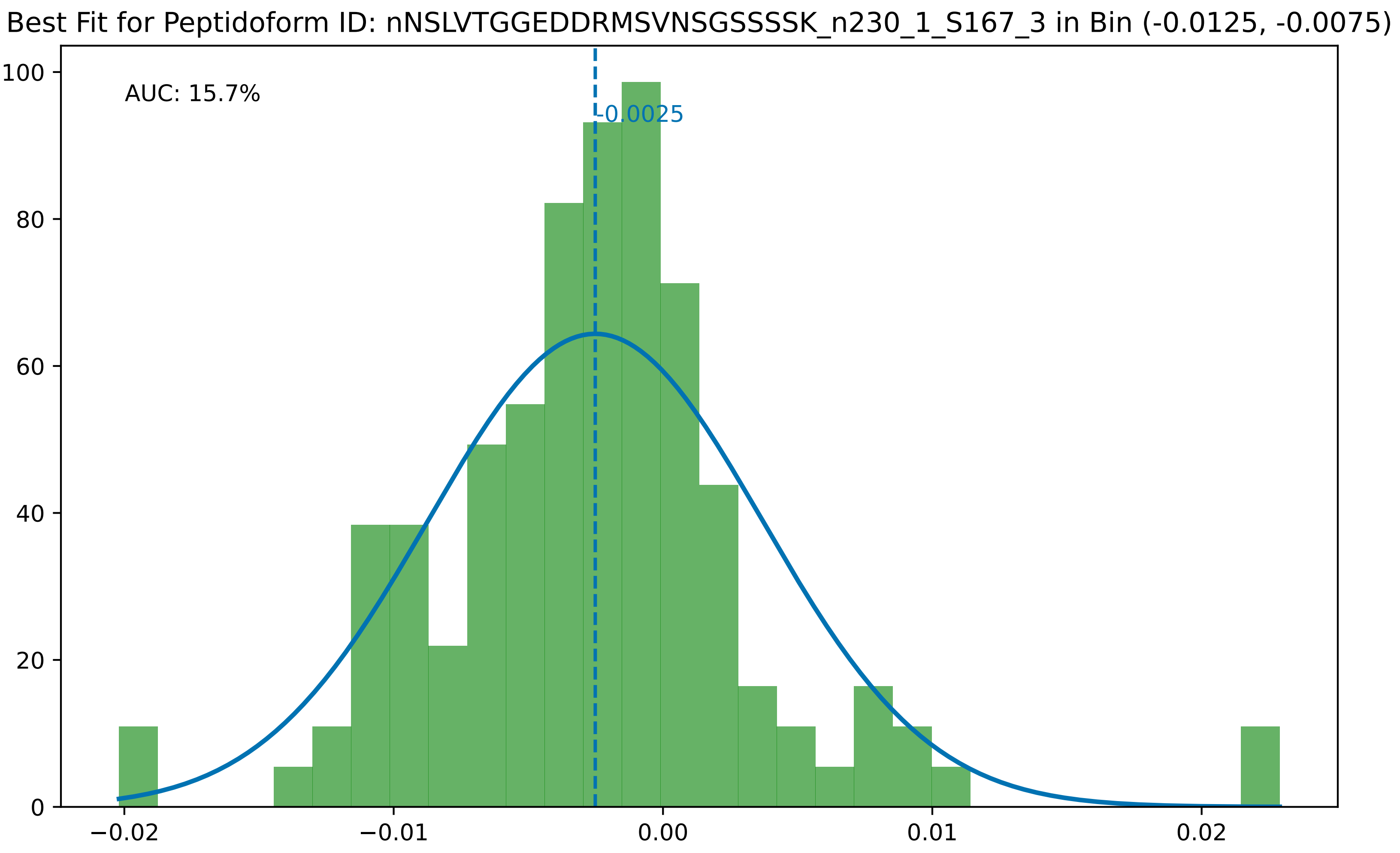
Best Fit for Peptidoform ID: nNSLVTGGEDDRMSVNSGSSSSK_n230_1_S167_3 in Bin (-0.0125, -0.0075)
AUC: 15.7%
-0.0025
0
20
40
60
80
100
−0.02
−0.01
0.00
0.01
0.02

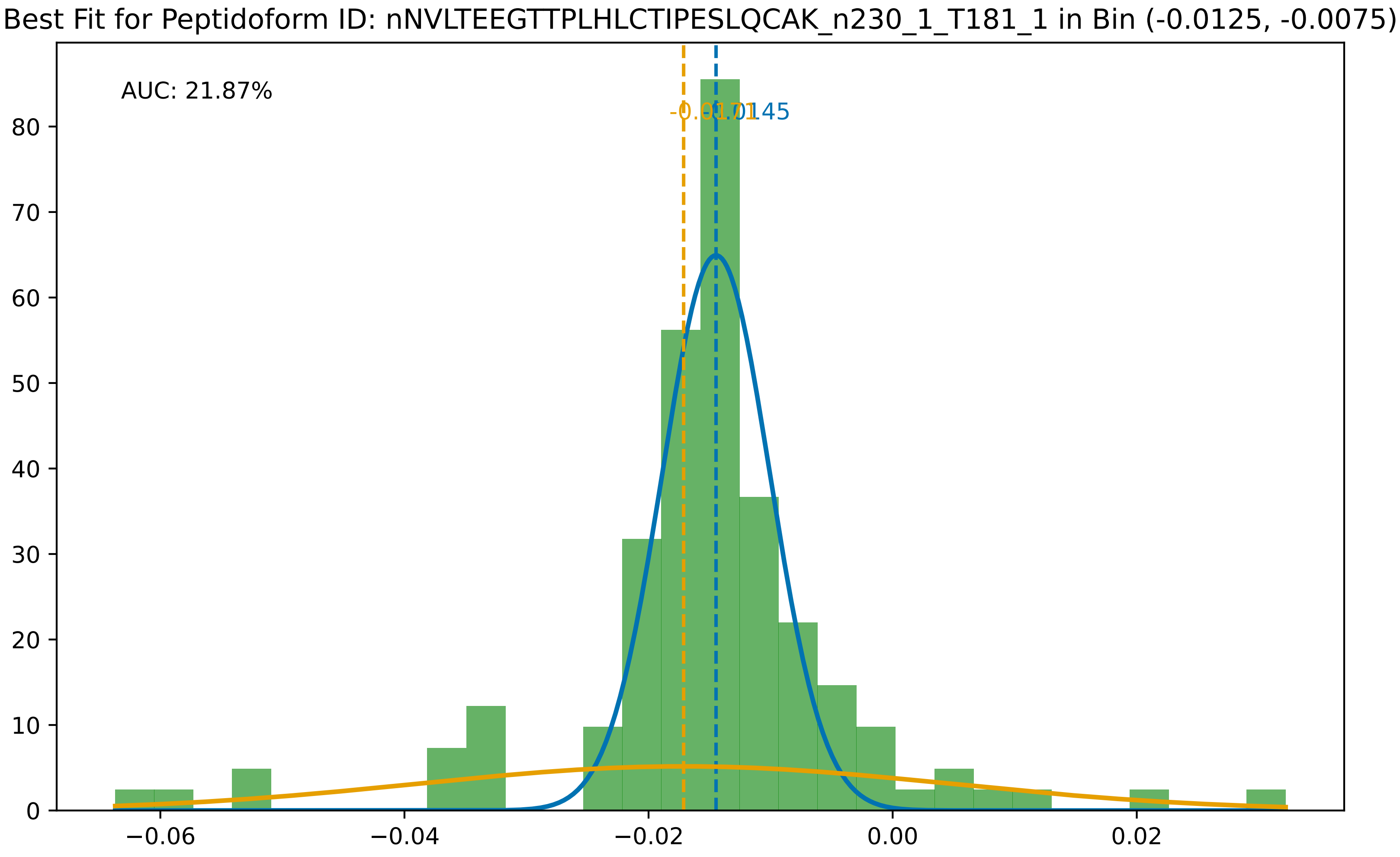
Best Fit for Peptidoform ID: nNVLTEEGTTPLHLCTIPESLQCAK_n230_1_T181_1 in Bin (-0.0125, -0.0075)
AUC: 21.87%
-0.0145
0
10
20
30
40
50
60
70
80
−0.06
−0.04
−0.02
0.00
0.02

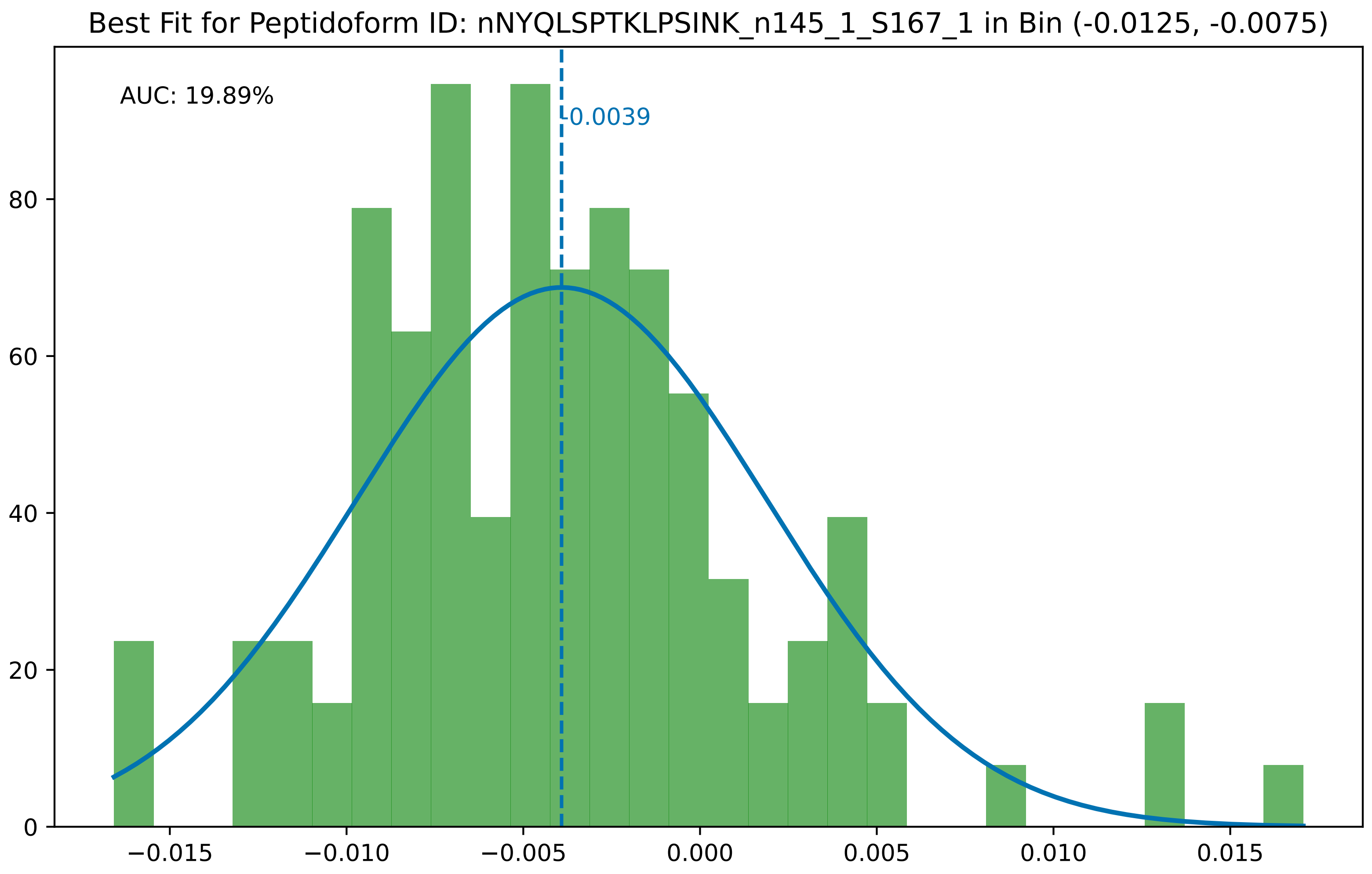

Best Fit for Peptidoform ID: nNYQLSPTKLPSINK_n145_1_S167_1 in Bin (-0.0125, -0.0075)
AUC: 19.89%
-0.0039
0
20
40
60
80
−0.015
−0.010
−0.005
0.000
0.005
0.010
0.015

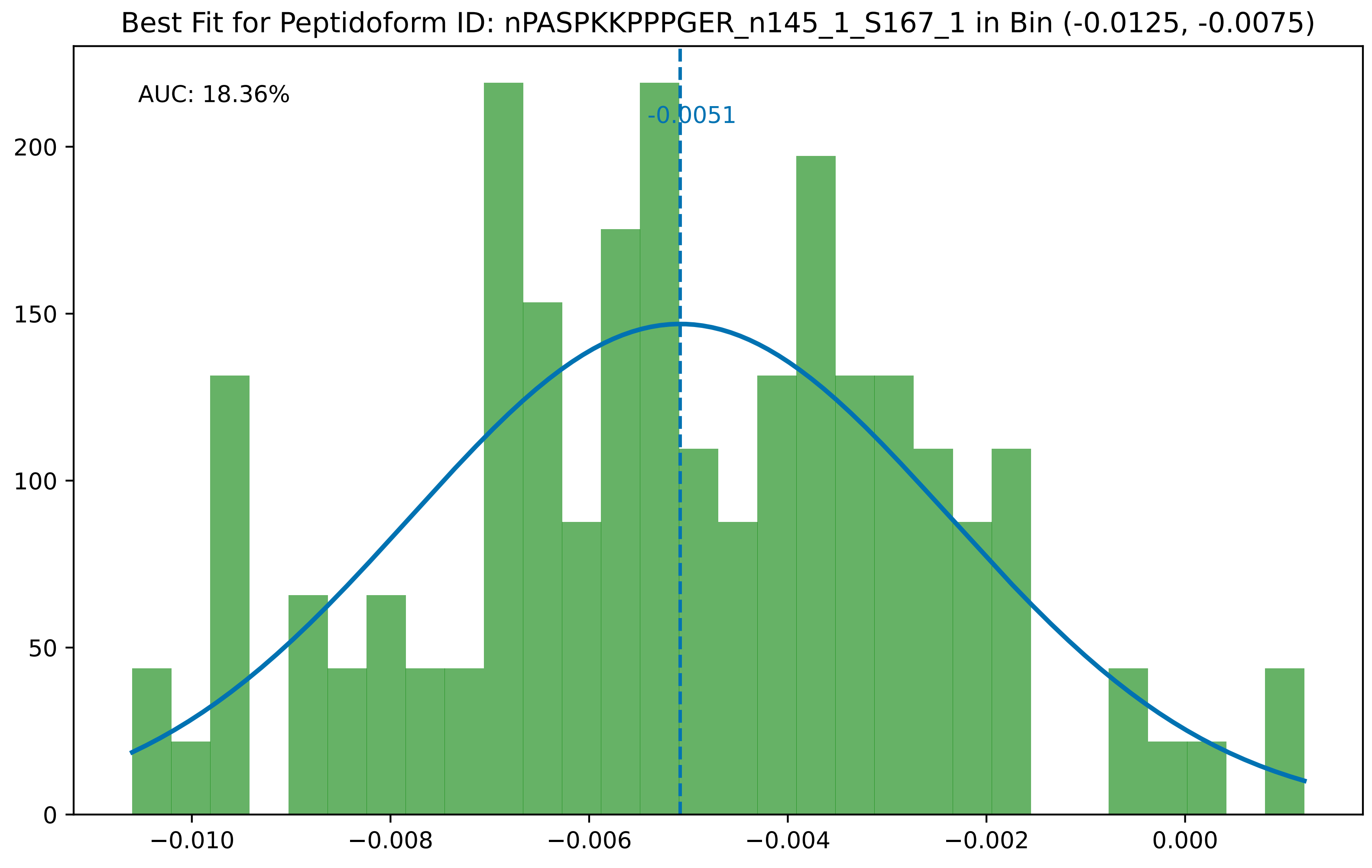

Best Fit for Peptidoform ID: nPASPKKPPPGER_n145_1_S167_1 in Bin (-0.0125, -0.0075)
AUC: 18.36%
-0.0051
0
50
100
150
200
−0.010
−0.008
−0.006
−0.004
−0.002
0.000

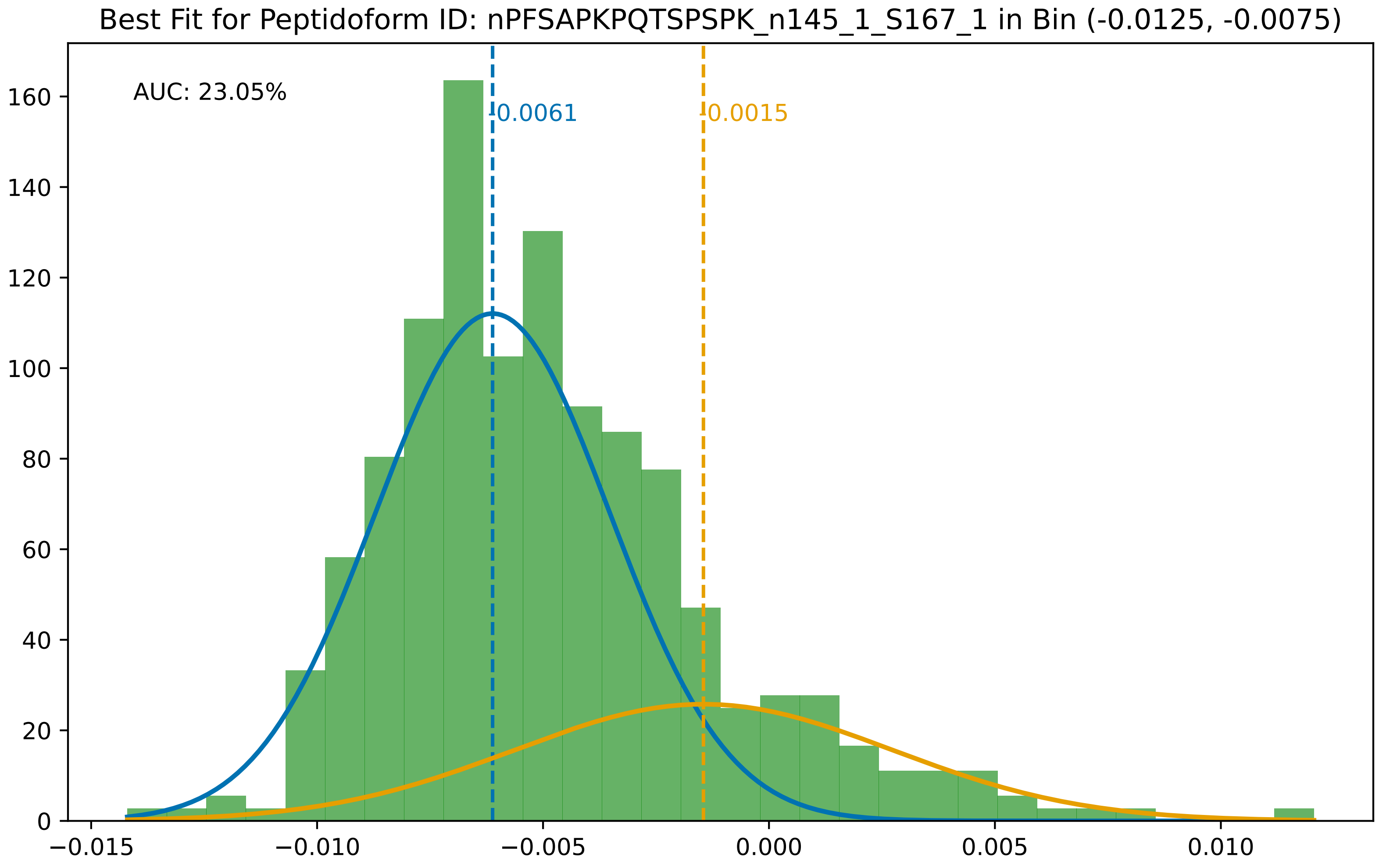

Best Fit for Peptidoform ID: nPFSAPKPQTSPSPK_n145_1_S167_1 in Bin (-0.0125, -0.0075)
AUC: 23.05%
-0.0061
-0.0015
0
20
40
60
80
100
120
140
160
−0.015
−0.010
−0.005
0.000
0.005
0.010

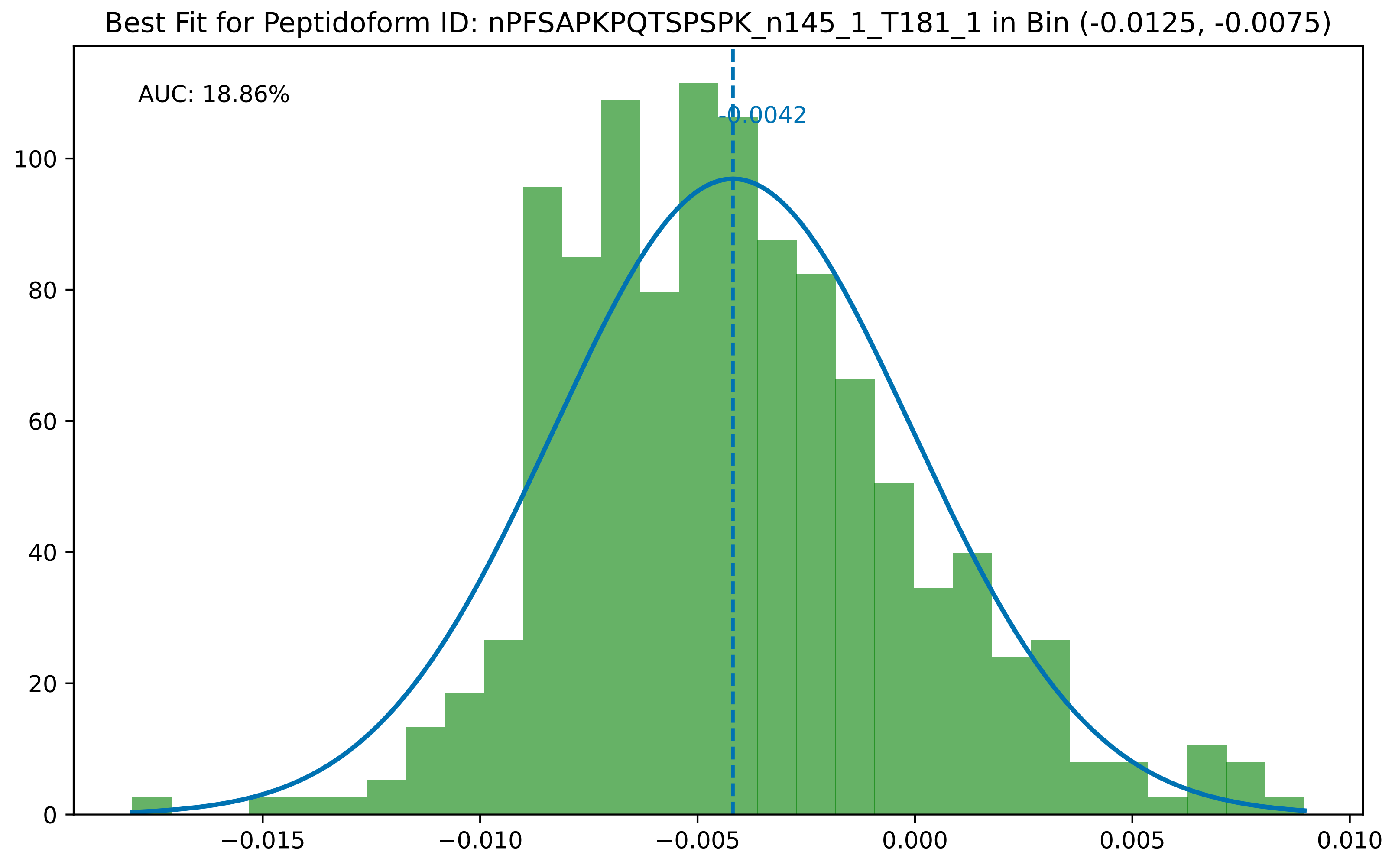

Best Fit for Peptidoform ID: nPFSAPKPQTSPSPK_n145_1_T181_1 in Bin (-0.0125, -0.0075)
AUC: 18.86%
-0.0042
0
20
40
60
80
100
−0.015
−0.010
−0.005
0.000
0.005
0.010

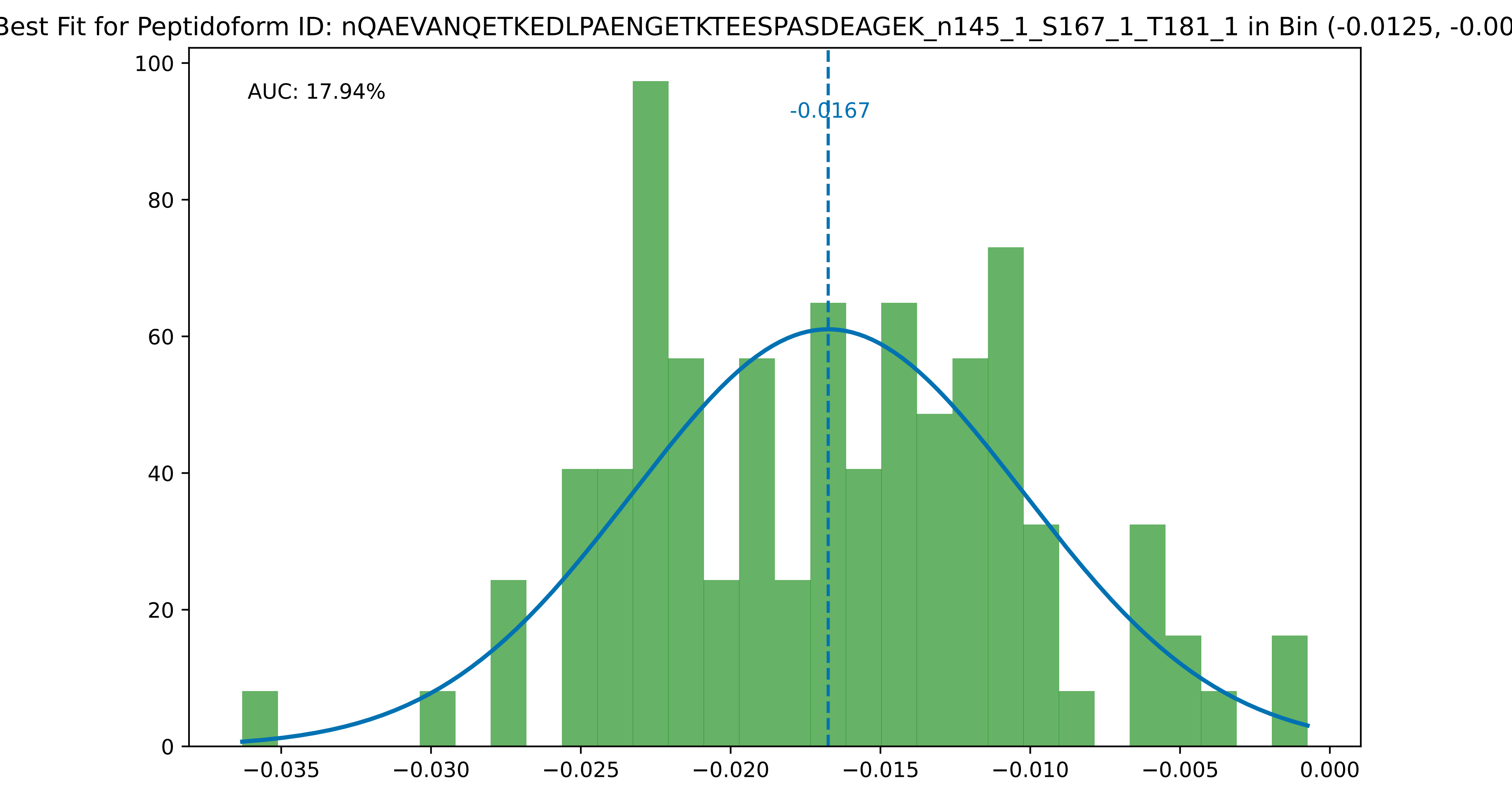

Best Fit for Peptidoform ID: nQAEVANQETKEDLPAENGETKTEESPASDEAGEK_n145_1_S167_1_T181_1 in Bin (-0.0125, -0.00
AUC: 17.94%
-0.0167
100
80
60
40
20
0
−0.035
−0.030
−0.025
−0.020
−0.015
−0.010
−0.005
0.000

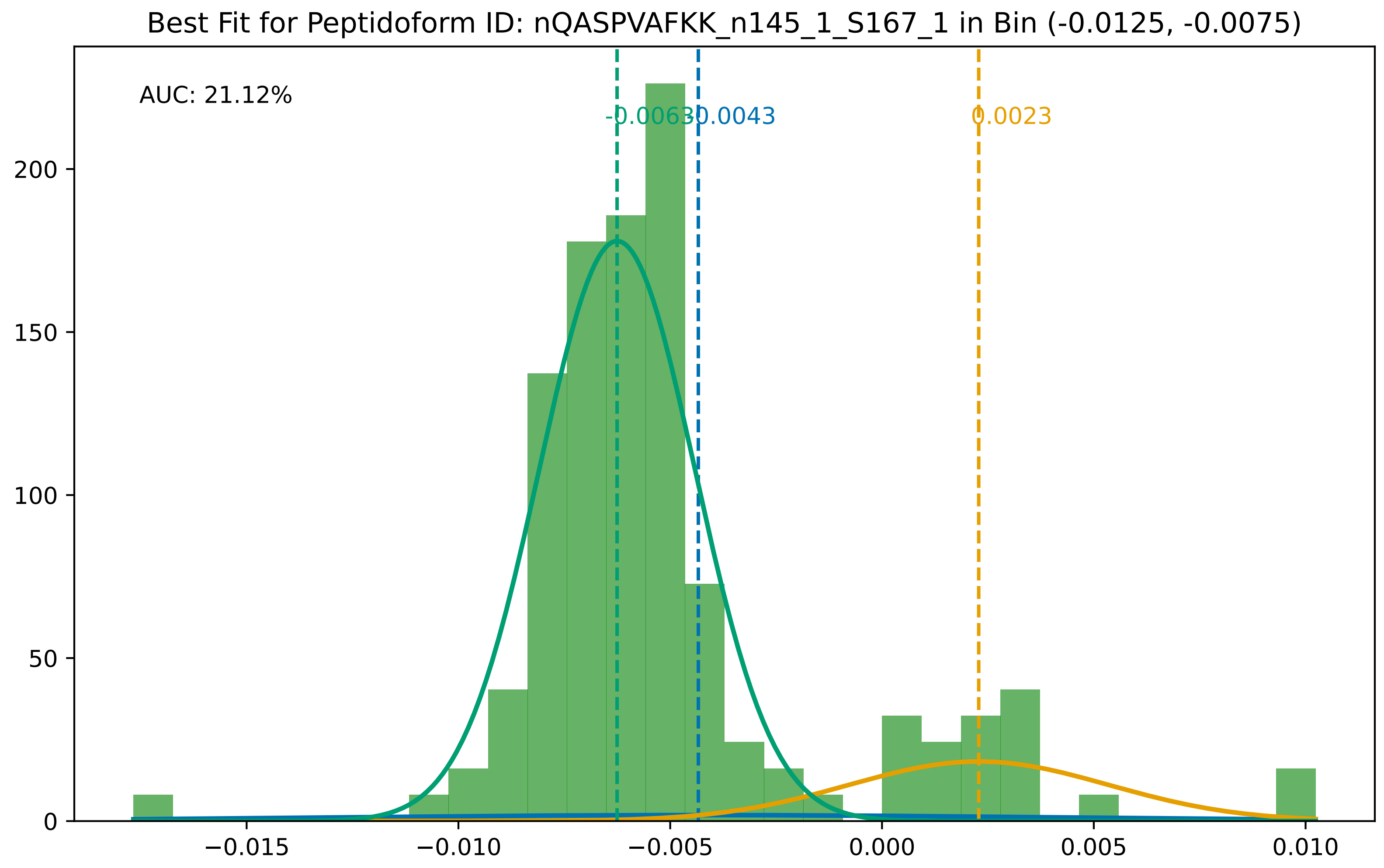

Best Fit for Peptidoform ID: nQASPVAFKK_n145_1_S167_1 in Bin (-0.0125, -0.0075)
AUC: 21.12%
-0.0063
-0.0043
0.0023
0
50
100
150
200
−0.015
−0.010
−0.005
0.000
0.005
0.010

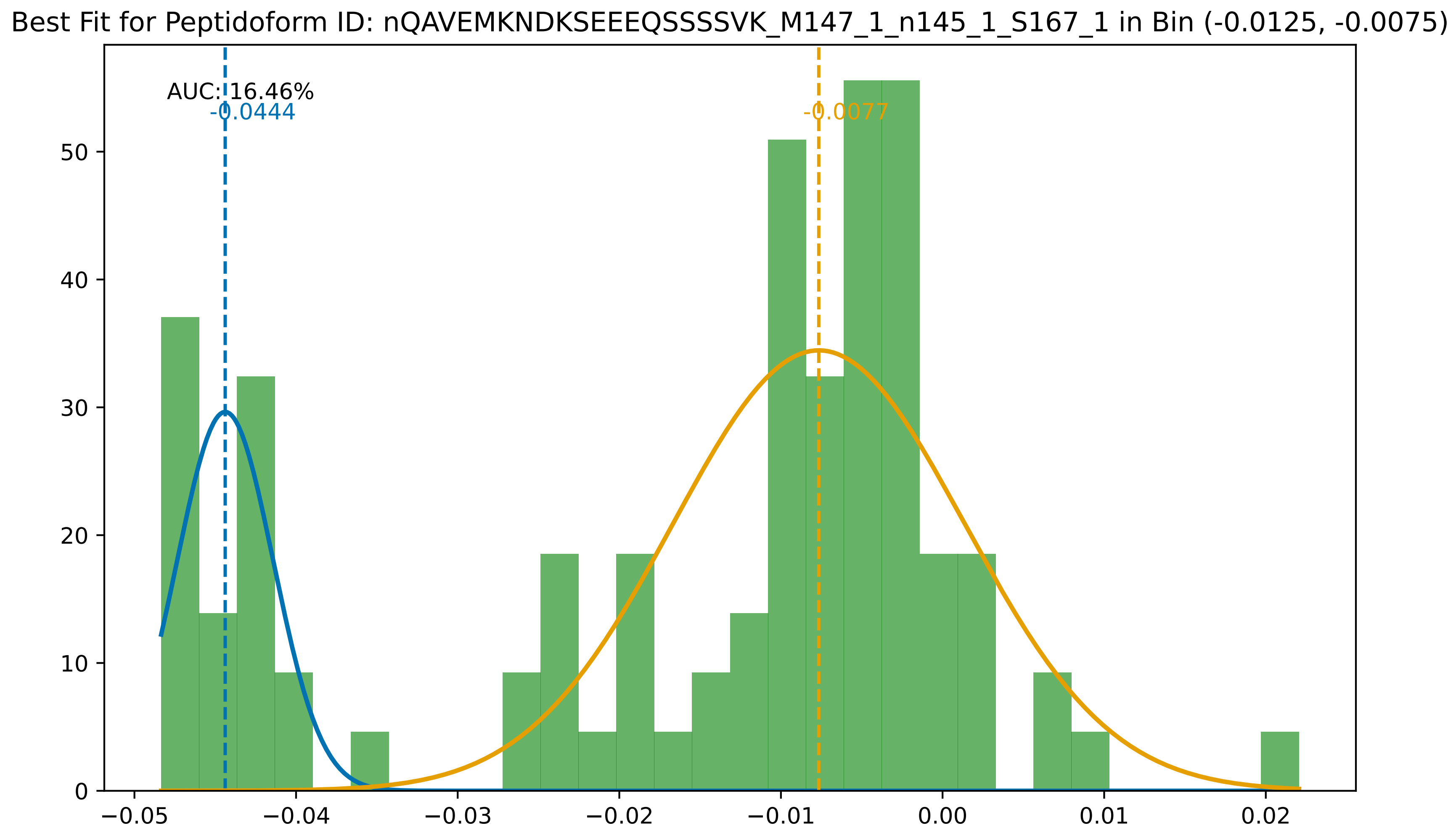

Best Fit for Peptidoform ID: nQAVEMKNDKSEEEQSSSSVK_M147_1_n145_1_S167_1 in Bin (-0.0125, -0.0075)
AUC: 16.46%
-0.0444
-0.0077
0
10
20
30
40
50
−0.05
−0.04
−0.03
−0.02
−0.01
0.00
0.01
0.02

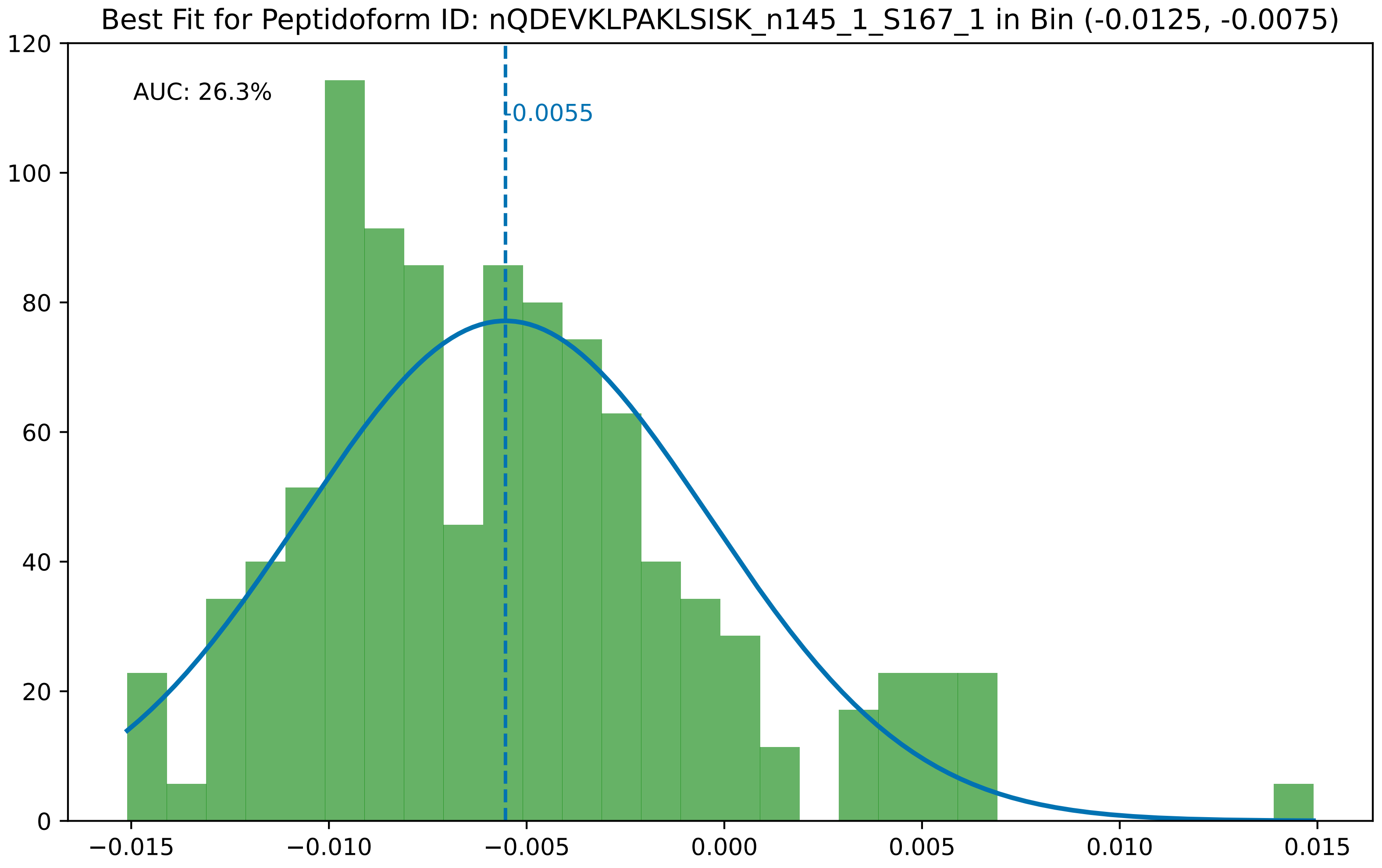
Best Fit for Peptidoform ID: nQDEVKLPAKLSISK_n145_1_S167_1 in Bin (-0.0125, -0.0075)
AUC: 26.3%
-0.0055
120
100
80
60
40
20
0
−0.015
−0.010
−0.005
0.000
0.005
0.010
0.015

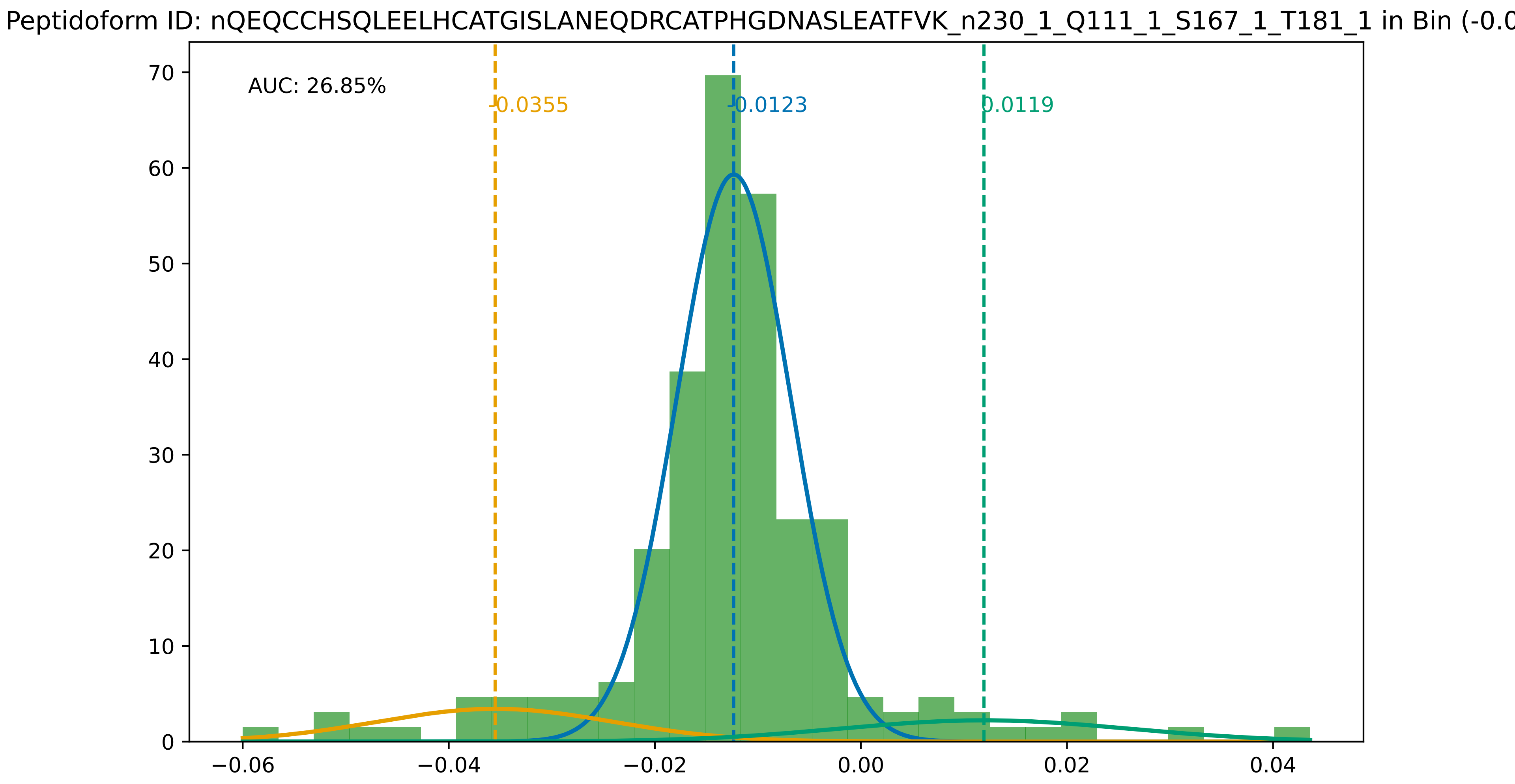

Peptidoform ID: nQEQCCHSQLEELHCATGISLANEQDRCATPHGDNASLEATFVK_n230_1_Q111_1_S167_1_T181_1 in Bin (-0.0
AUC: 26.85%
-0.0355
-0.0123
0.0119
0
10
20
30
40
50
60
70
−0.06
−0.04
−0.02
0.00
0.02
0.04

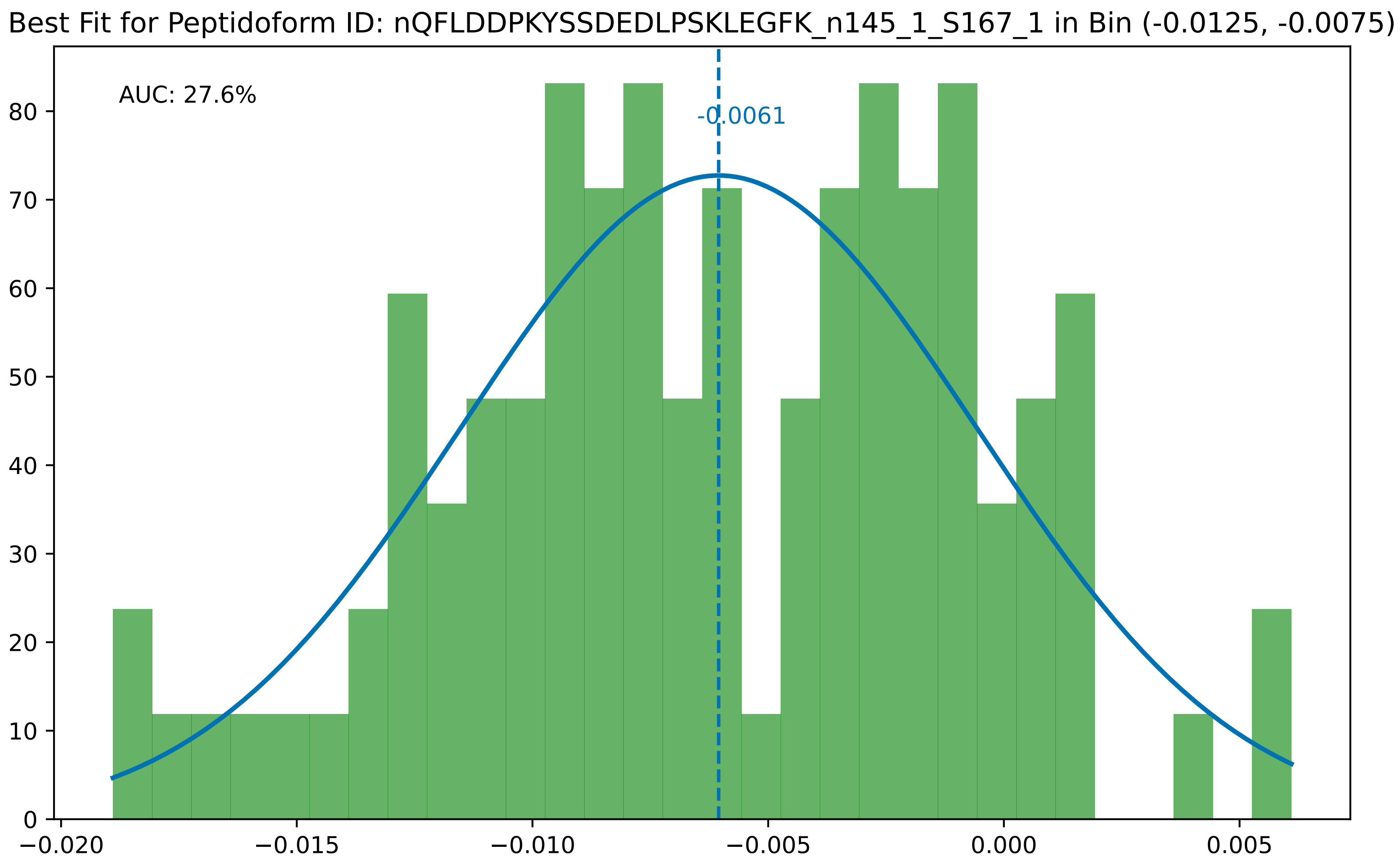

Best Fit for Peptidoform ID: nQFLDDPKYSSDEDLPSKLEGFK_n145_1_S167_1 in Bin (-0.0125, -0.0075)
AUC: 27.6%
-0.0061
0
10
20
30
40
50
60
70
80
−0.020
−0.015
−0.010
−0.005
0.000
0.005

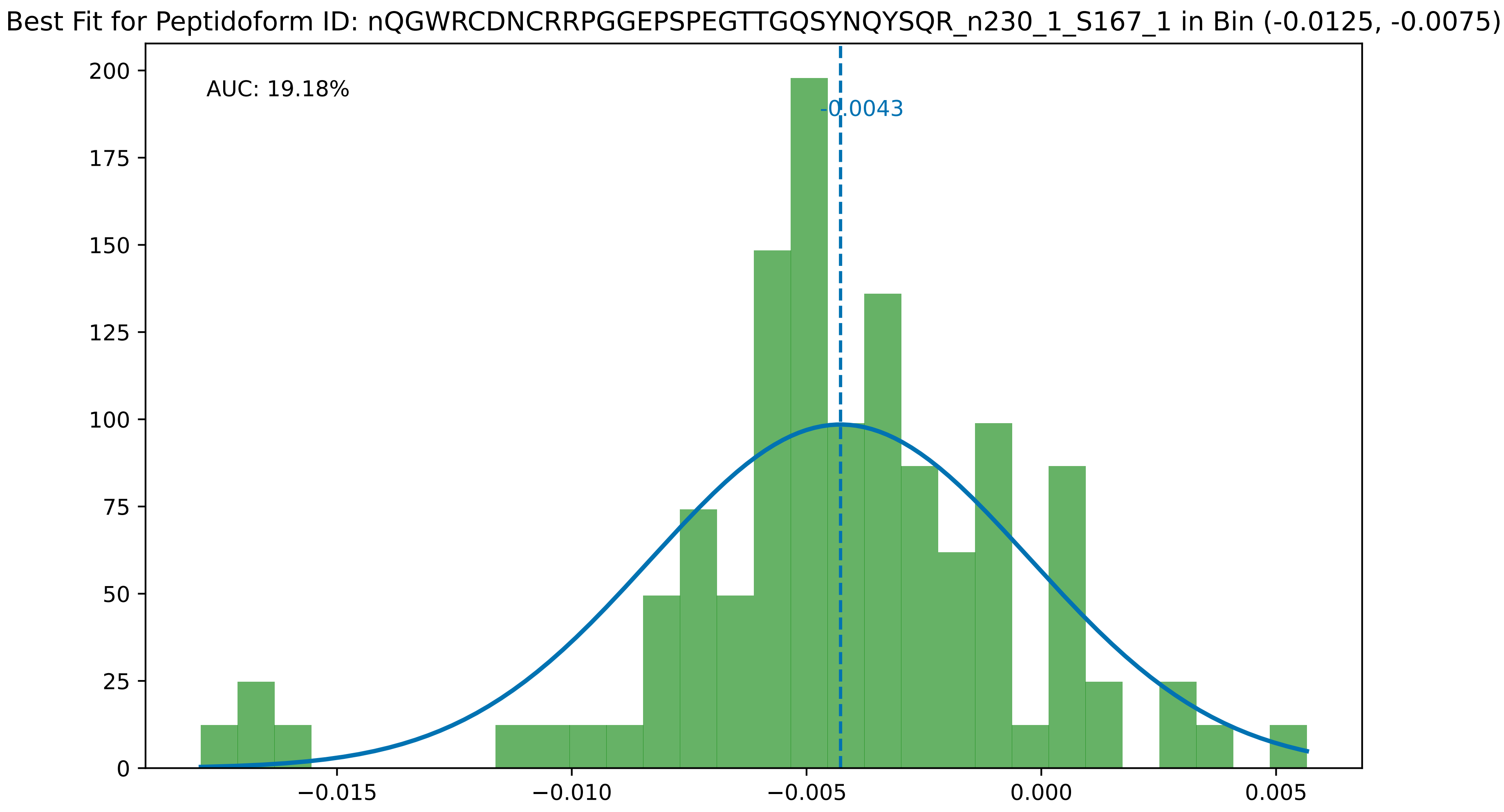

Best Fit for Peptidoform ID: nQGWRCDNCRRPGGEPSPEGTTGQSYNQYSQR_n230_1_S167_1 in Bin (-0.0125, -0.0075)
AUC: 19.18%
-0.0043
200
175
150
125
100
75
50
25
0
−0.015
−0.010
−0.005
0.000
0.005

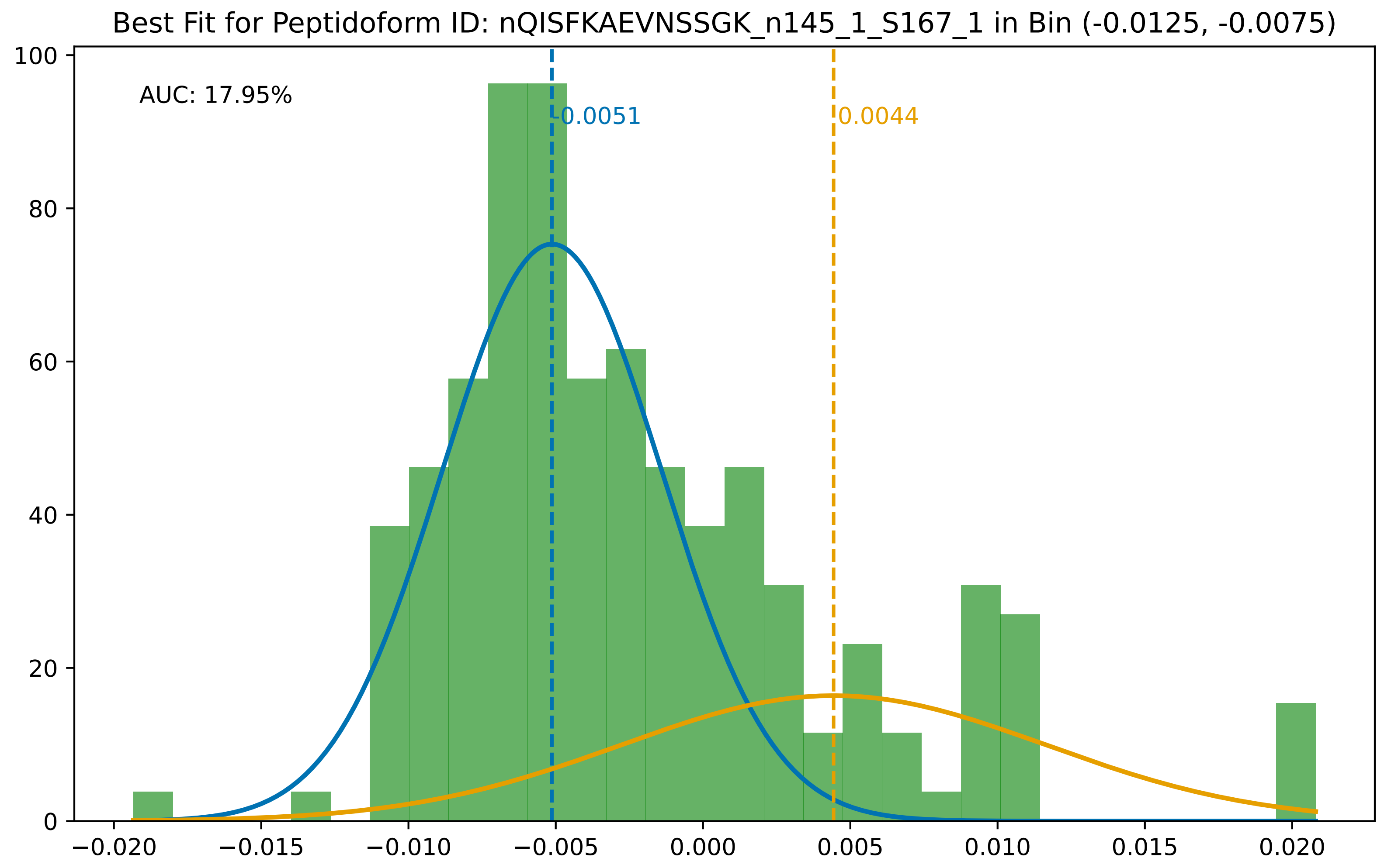

Best Fit for Peptidoform ID: nQISFKAEVNSSGK_n145_1_S167_1 in Bin (-0.0125, -0.0075)
AUC: 17.95%
-0.0051
0.0044
100
80
60
40
20
0
−0.020
−0.015
−0.010
−0.005
0.000
0.005
0.010
0.015
0.020

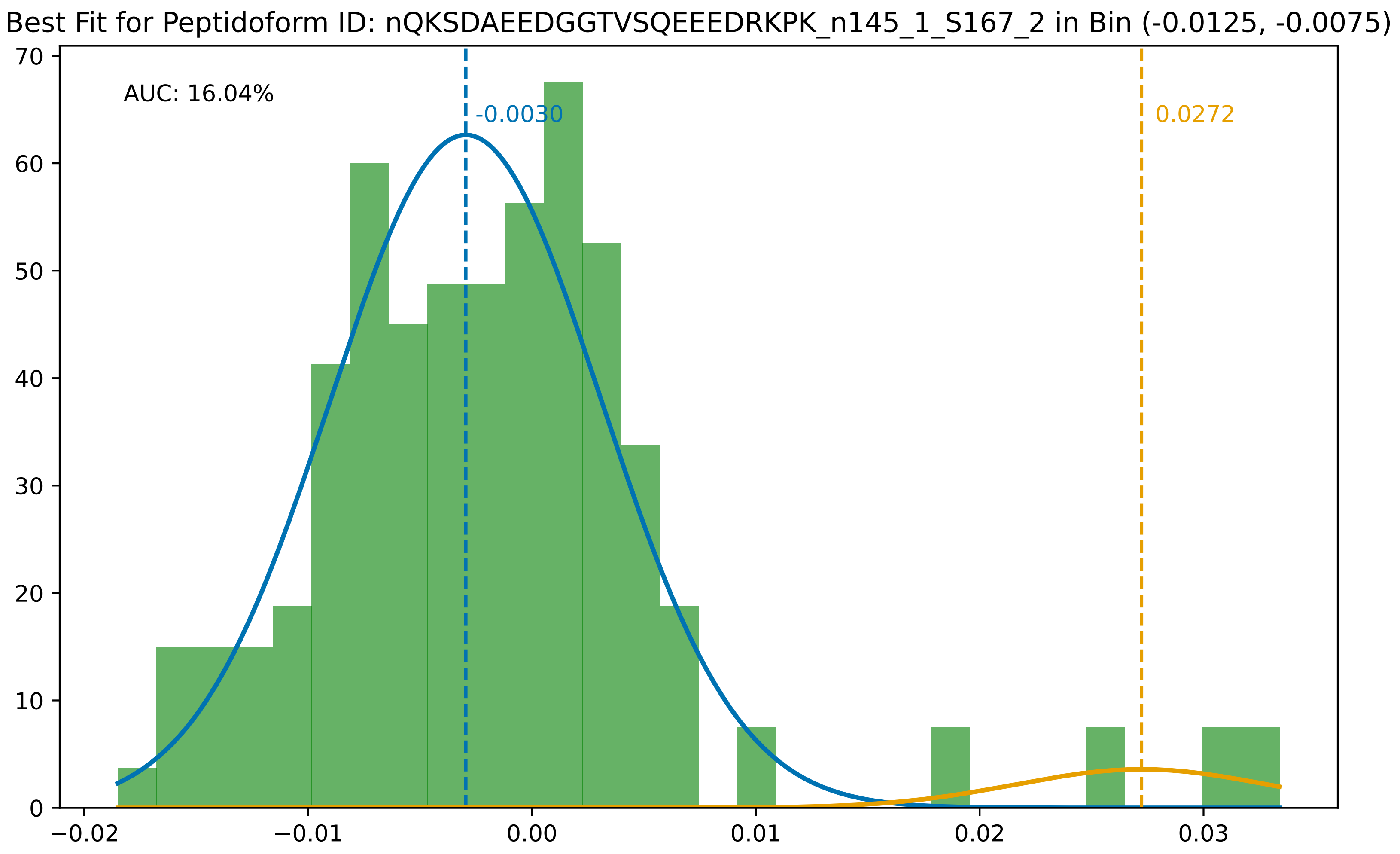

Best Fit for Peptidoform ID: nQKSDAEEDGGTVSQEEEDRKPK_n145_1_S167_2 in Bin (-0.0125, -0.0075)
AUC: 16.04%
-0.0030
0.0272
70
60
50
40
30
20
10
0
−0.02
−0.01
0.00
0.01
0.02
0.03

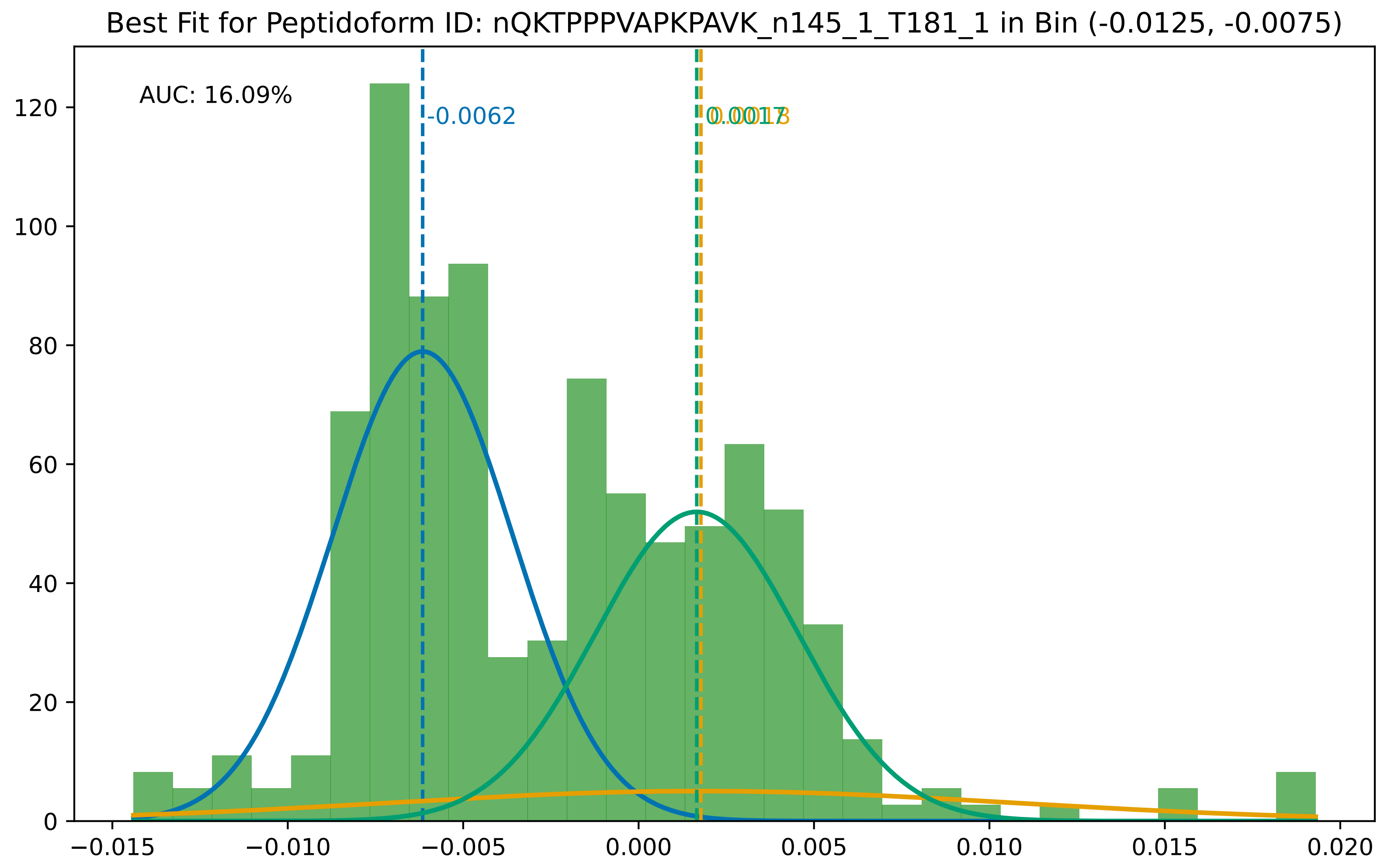

Best Fit for Peptidoform ID: nQKTPPPVAPKPAVK_n145_1_T181_1 in Bin (-0.0125, -0.0075)
AUC: 16.09%
-0.0062
0.0017
0.0018
0
20
40
60
80
100
120
−0.015
−0.010
−0.005
0.000
0.005
0.010
0.015
0.020

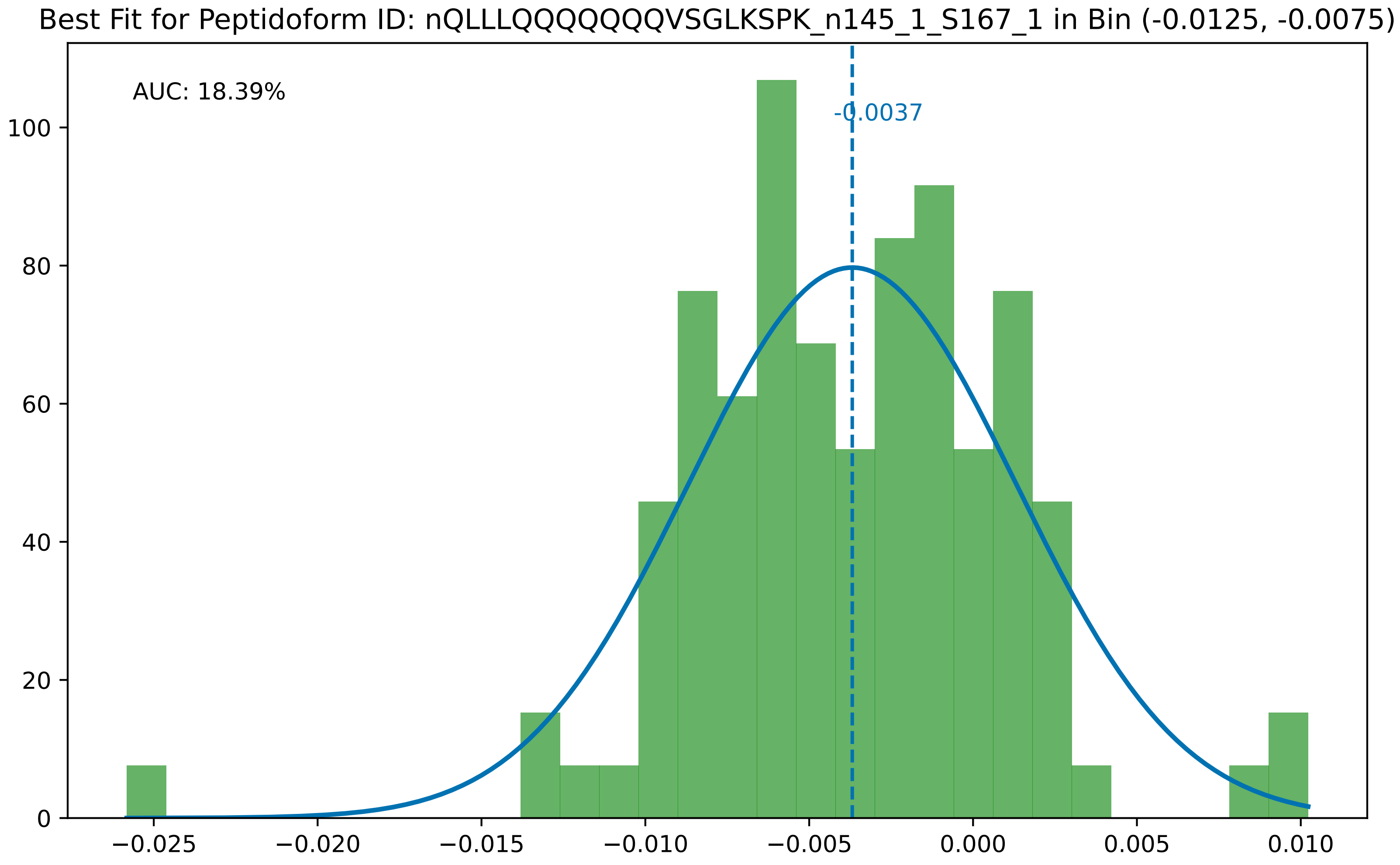
Best Fit for Peptidoform ID: nQLLLQQQQQQVSGLKSPK_n145_1_S167_1 in Bin (-0.0125, -0.0075)
AUC: 18.39%
-0.0037
0
20
40
60
80
100
−0.025
−0.020
−0.015
−0.010
−0.005
0.000
0.005
0.010

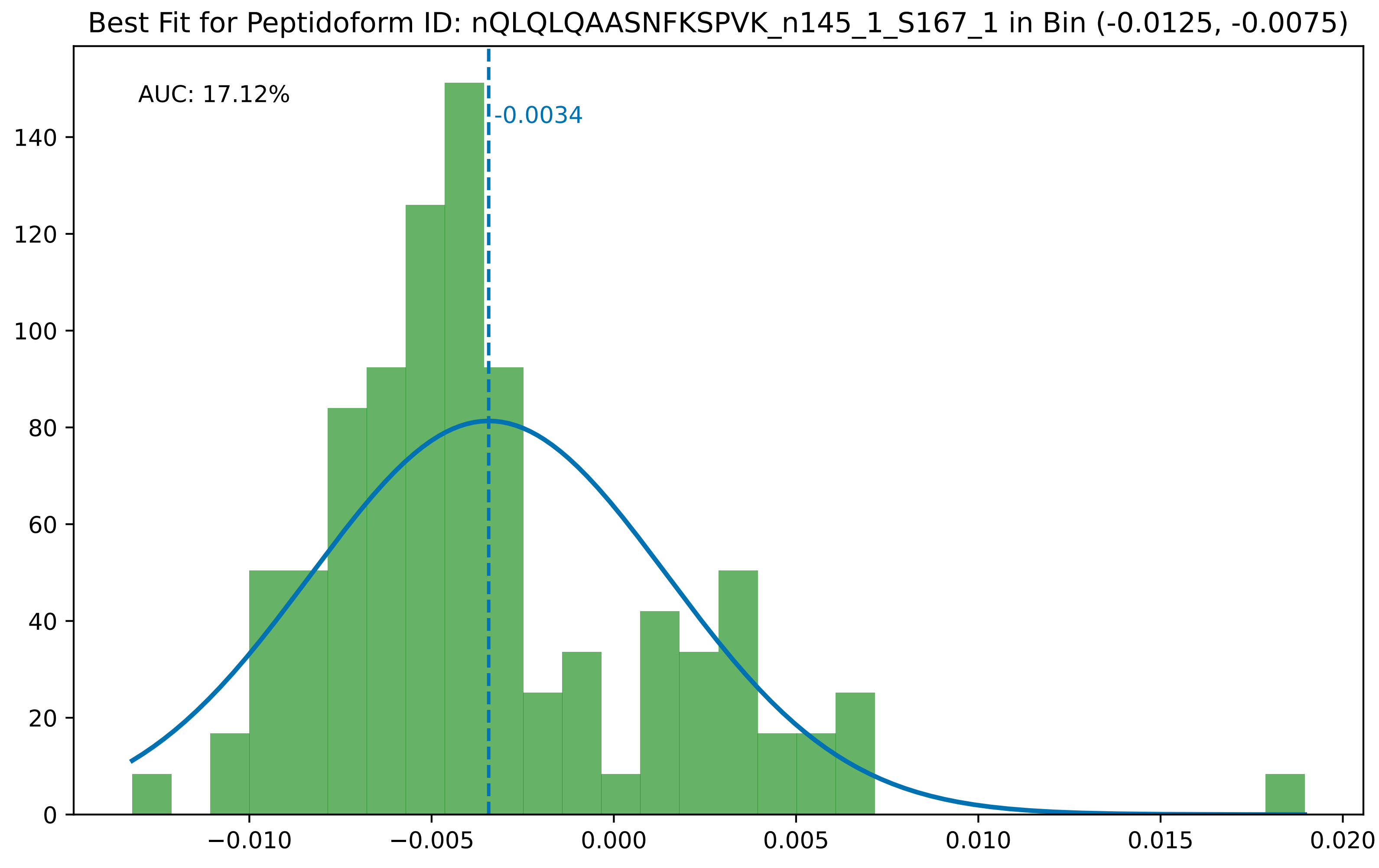

Best Fit for Peptidoform ID: nQLQLQAASNFKSPVK_n145_1_S167_1 in Bin (-0.0125, -0.0075)
AUC: 17.12%
-0.0034
0
20
40
60
80
100
120
140
−0.010
−0.005
0.000
0.005
0.010
0.015
0.020

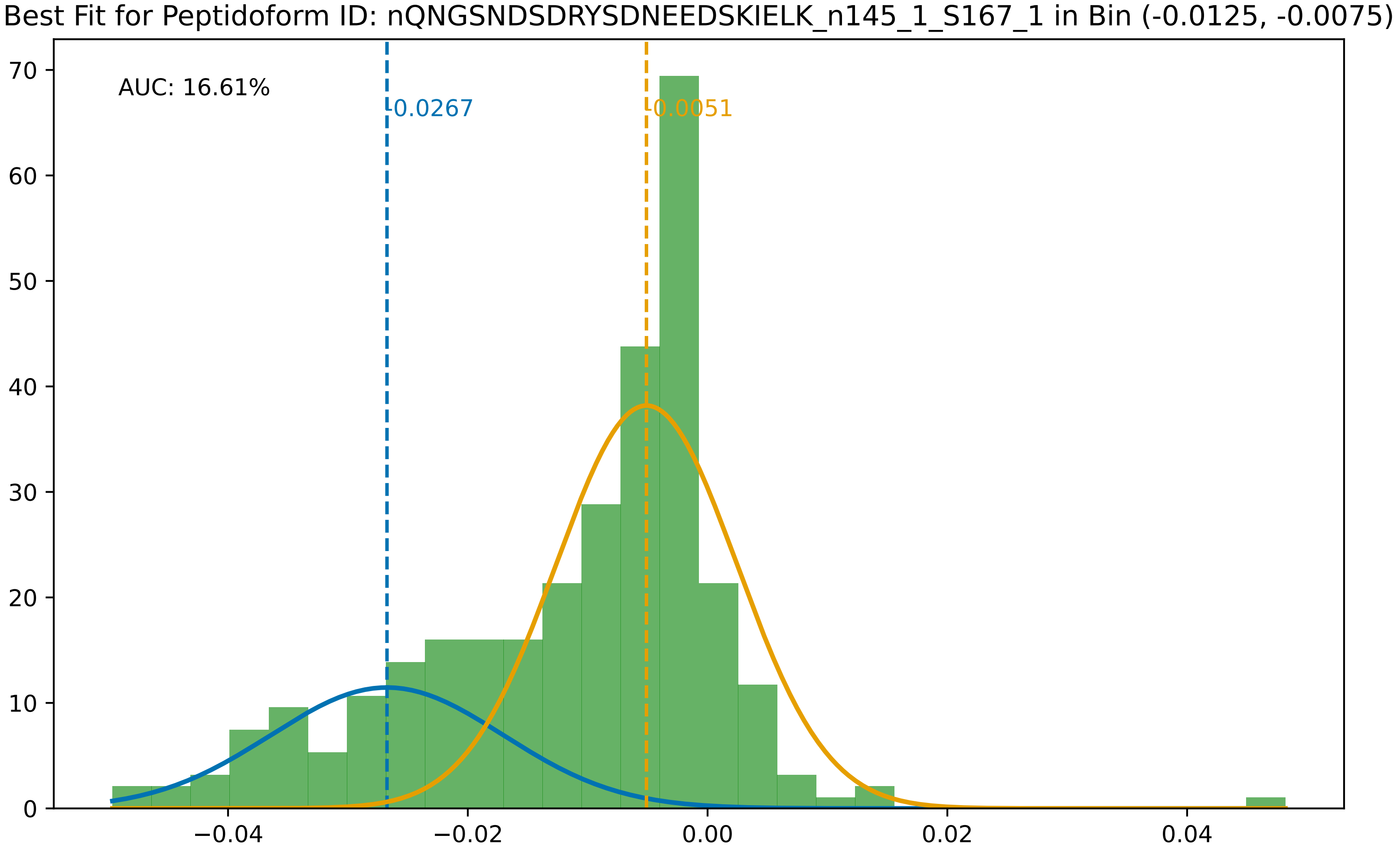

Best Fit for Peptidoform ID: nQNGSNDSDRYSDNEEDSKIELK_n145_1_S167_1 in Bin (-0.0125, -0.0075)
AUC: 16.61%
-0.0267
-0.0051
70
60
50
40
30
20
10
0
−0.04
−0.02
0.00
0.02
0.04

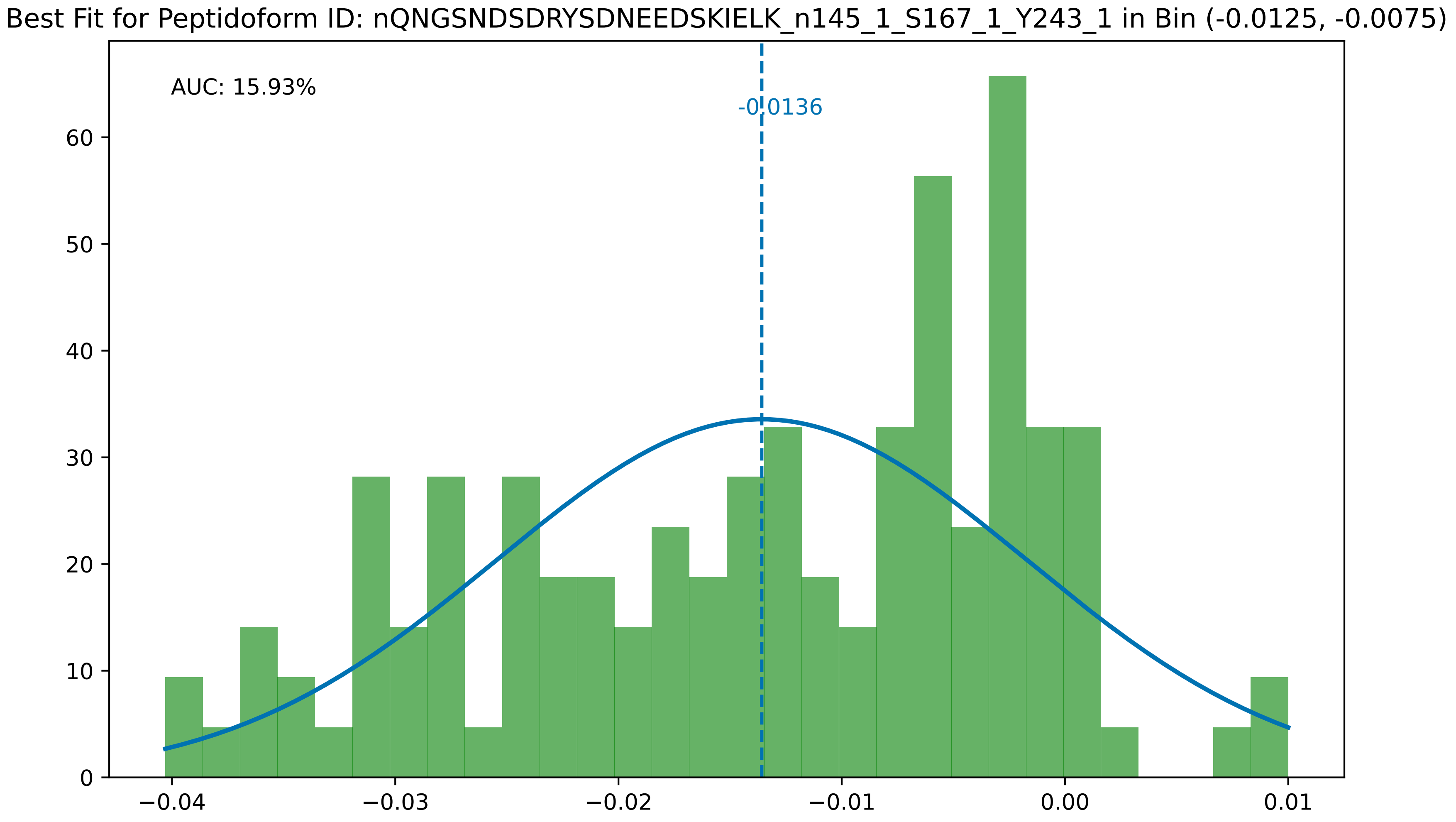

Best Fit for Peptidoform ID: nQNGSNDSDRYSDNEEDSKIELK_n145_1_S167_1_Y243_1 in Bin (-0.0125, -0.0075)
AUC: 15.93%
-0.0136
0
10
20
30
40
50
60
−0.04
−0.03
−0.02
−0.01
0.00
0.01

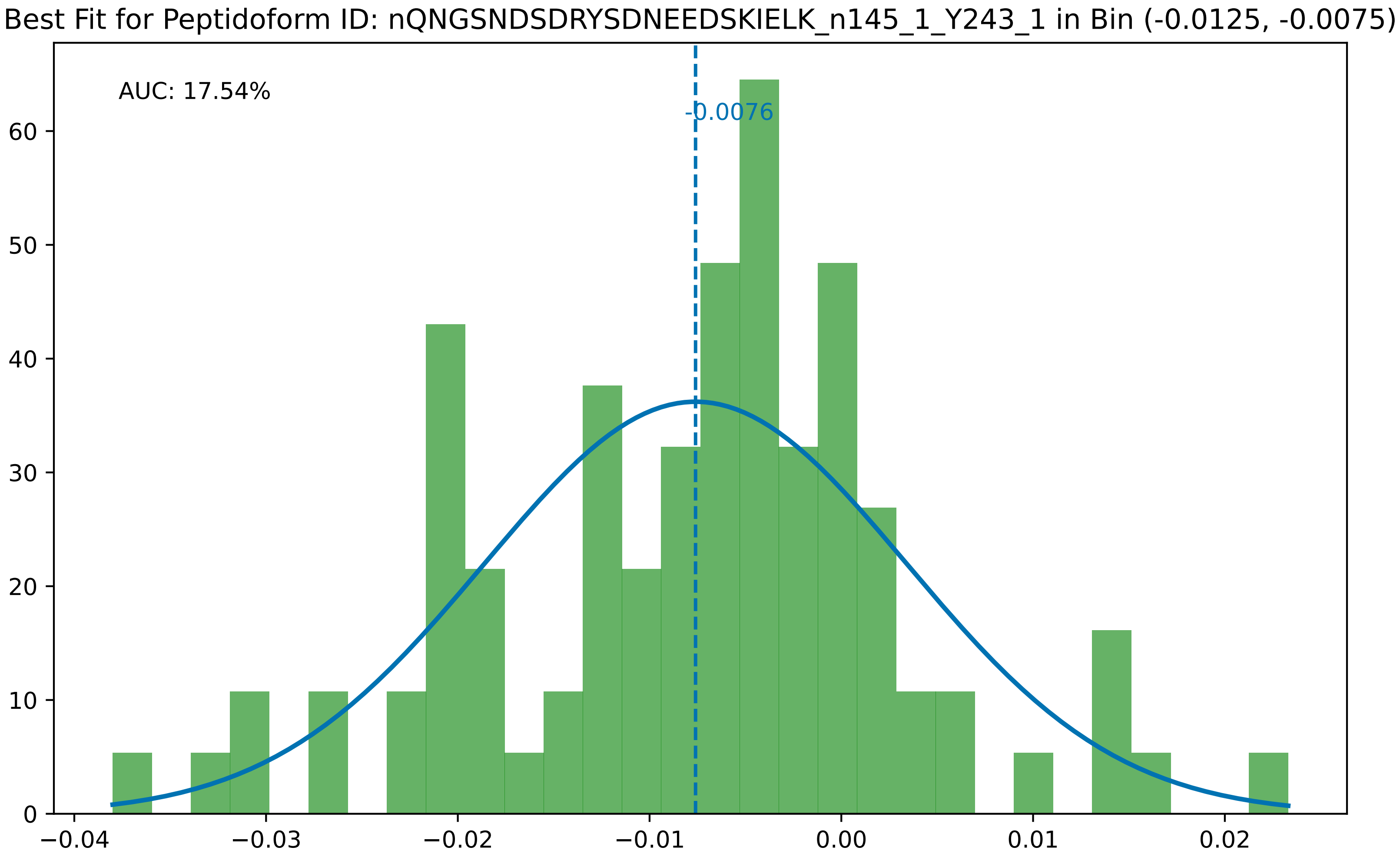

Best Fit for Peptidoform ID: nQNGSNDSDRYSDNEEDSKIELK_n145_1_Y243_1 in Bin (-0.0125, -0.0075)
AUC: 17.54%
-0.0076
0
10
20
30
40
50
60
−0.04
−0.03
−0.02
−0.01
0.00
0.01
0.02

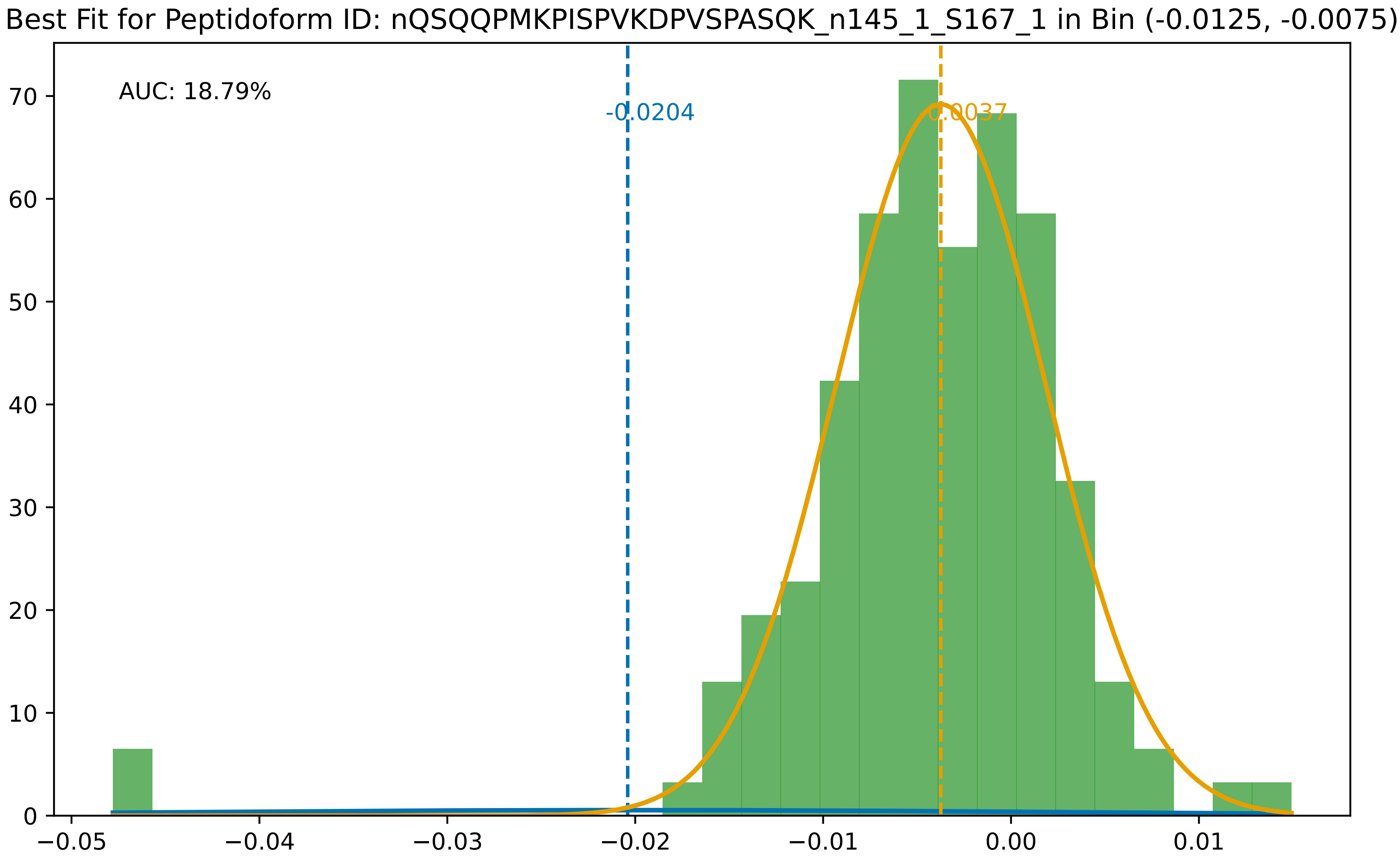

Best Fit for Peptidoform ID: nQSQQPMKPISPVKDPVSPASQK_n145_1_S167_1 in Bin (-0.0125, -0.0075)
AUC: 18.79%
-0.0204
-0.0037
0
10
20
30
40
50
60
70
−0.05
−0.04
−0.03
−0.02
−0.01
0.00
0.01

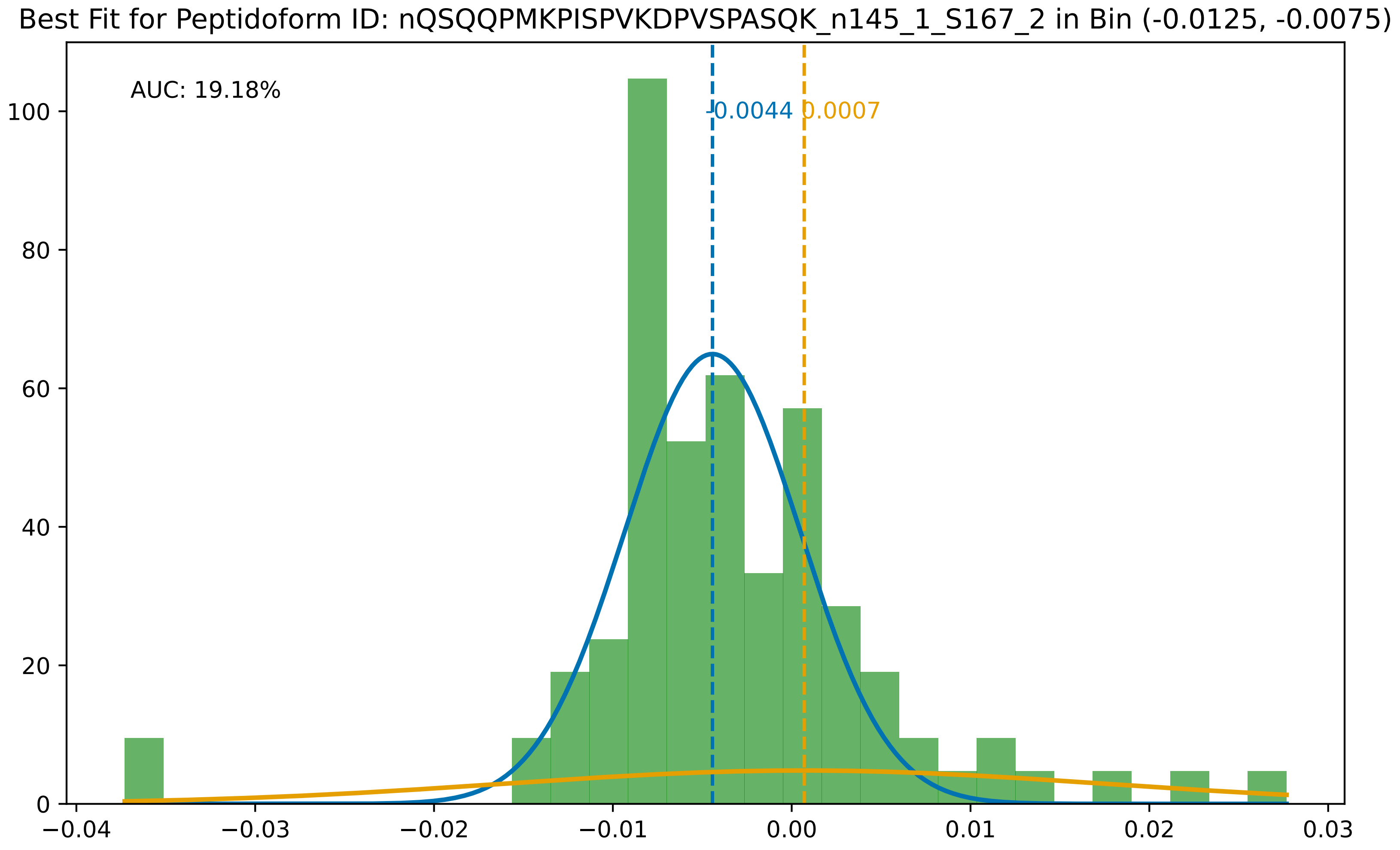

Best Fit for Peptidoform ID: nQSQQPMKPISPVKDPVSPASQK_n145_1_S167_2 in Bin (-0.0125, -0.0075)
AUC: 19.18%
-0.0044
0.0007
0
20
40
60
80
100
−0.04
−0.03
−0.02
−0.01
0.00
0.01
0.02
0.03

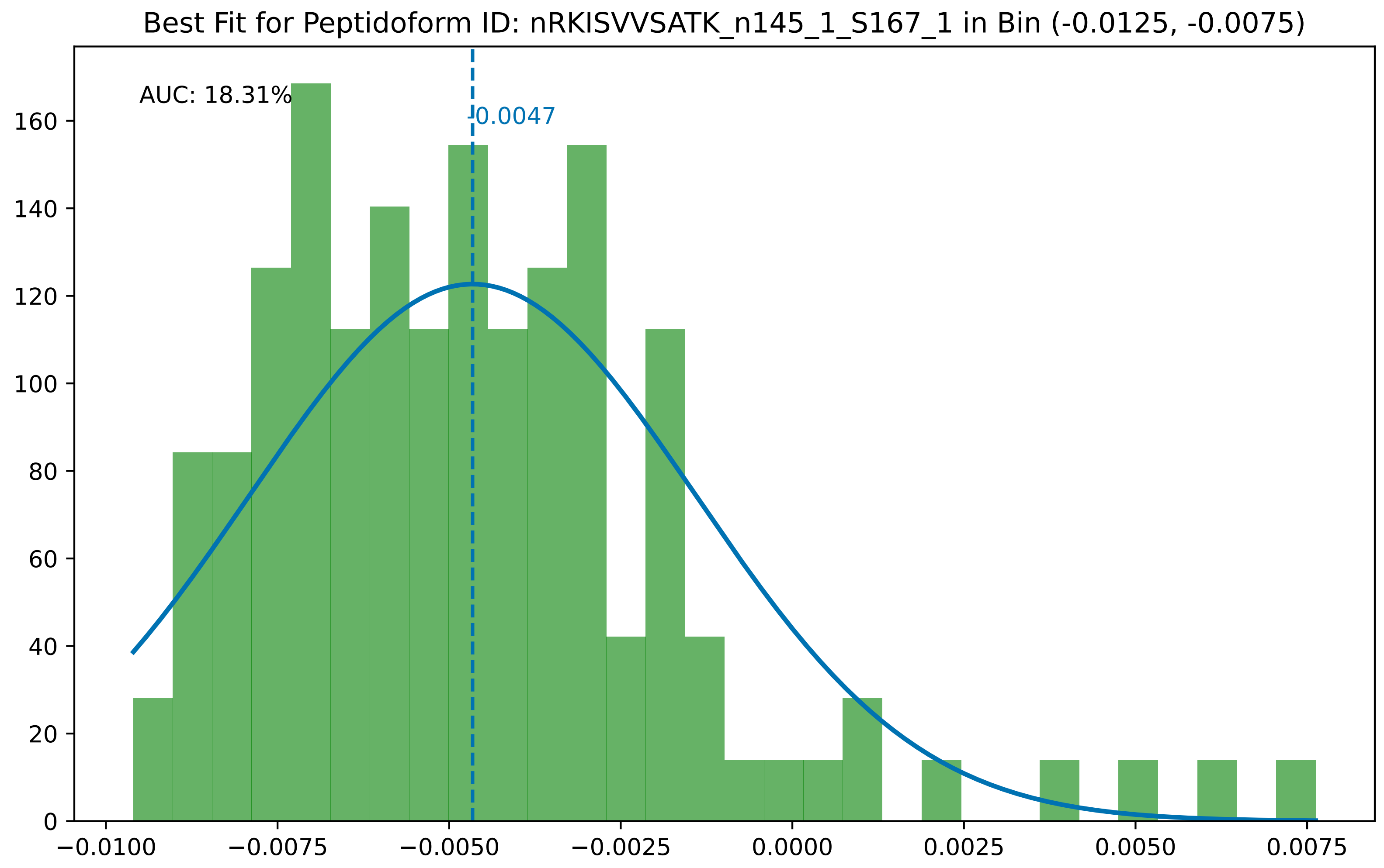

Best Fit for Peptidoform ID: nRKISVVSATK_n145_1_S167_1 in Bin (-0.0125, -0.0075)
AUC: 18.31%
-0.0047
0
20
40
60
80
100
120
140
160
−0.0100
−0.0075
−0.0050
−0.0025
0.0000
0.0025
0.0050
0.0075

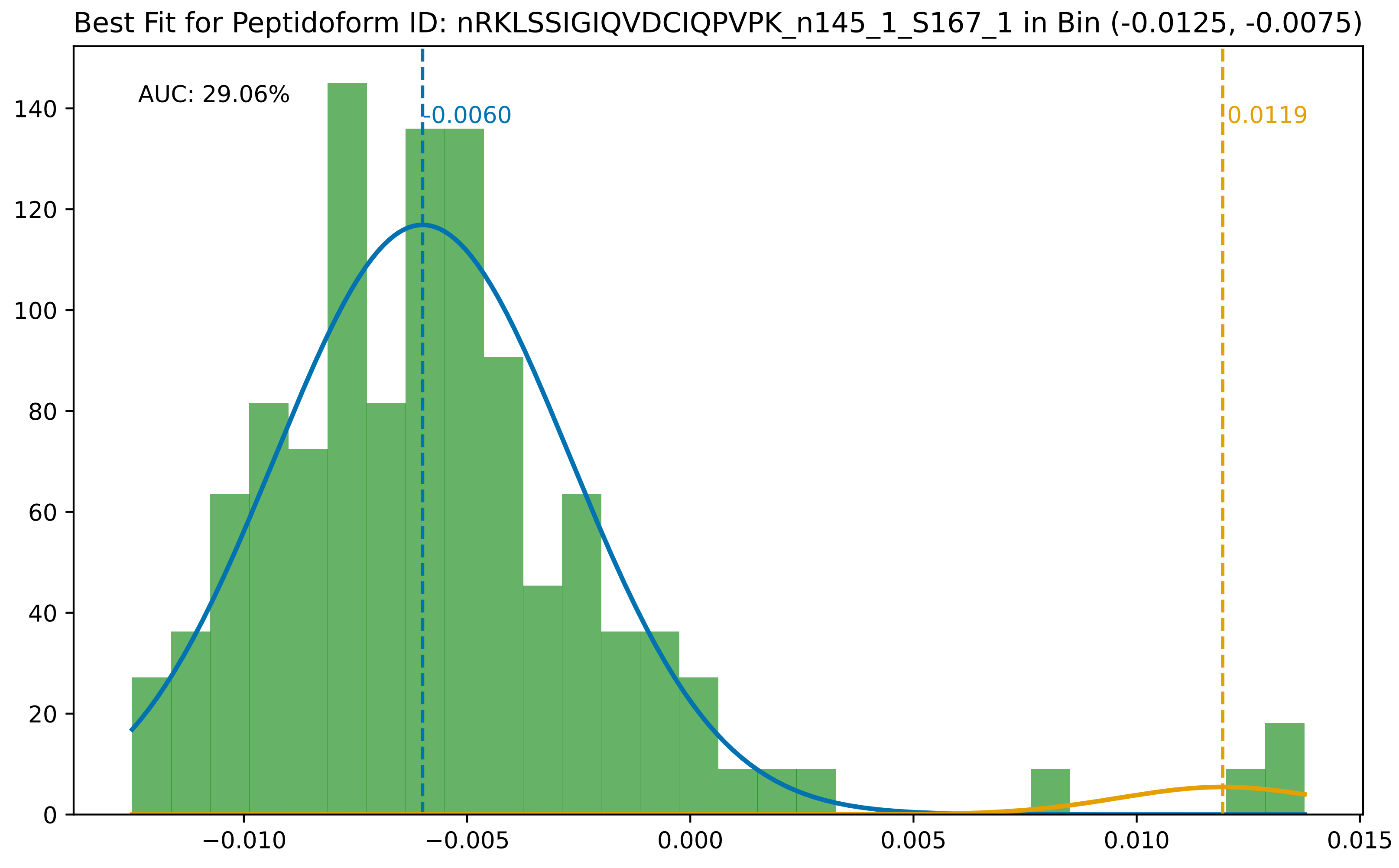

Best Fit for Peptidoform ID: nRKLSSIGIQVDCIQPVPK_n145_1_S167_1 in Bin (-0.0125, -0.0075)
AUC: 29.06%
-0.0060
0.0119
0
20
40
60
80
100
120
140
−0.010
−0.005
0.000
0.005
0.010
0.015

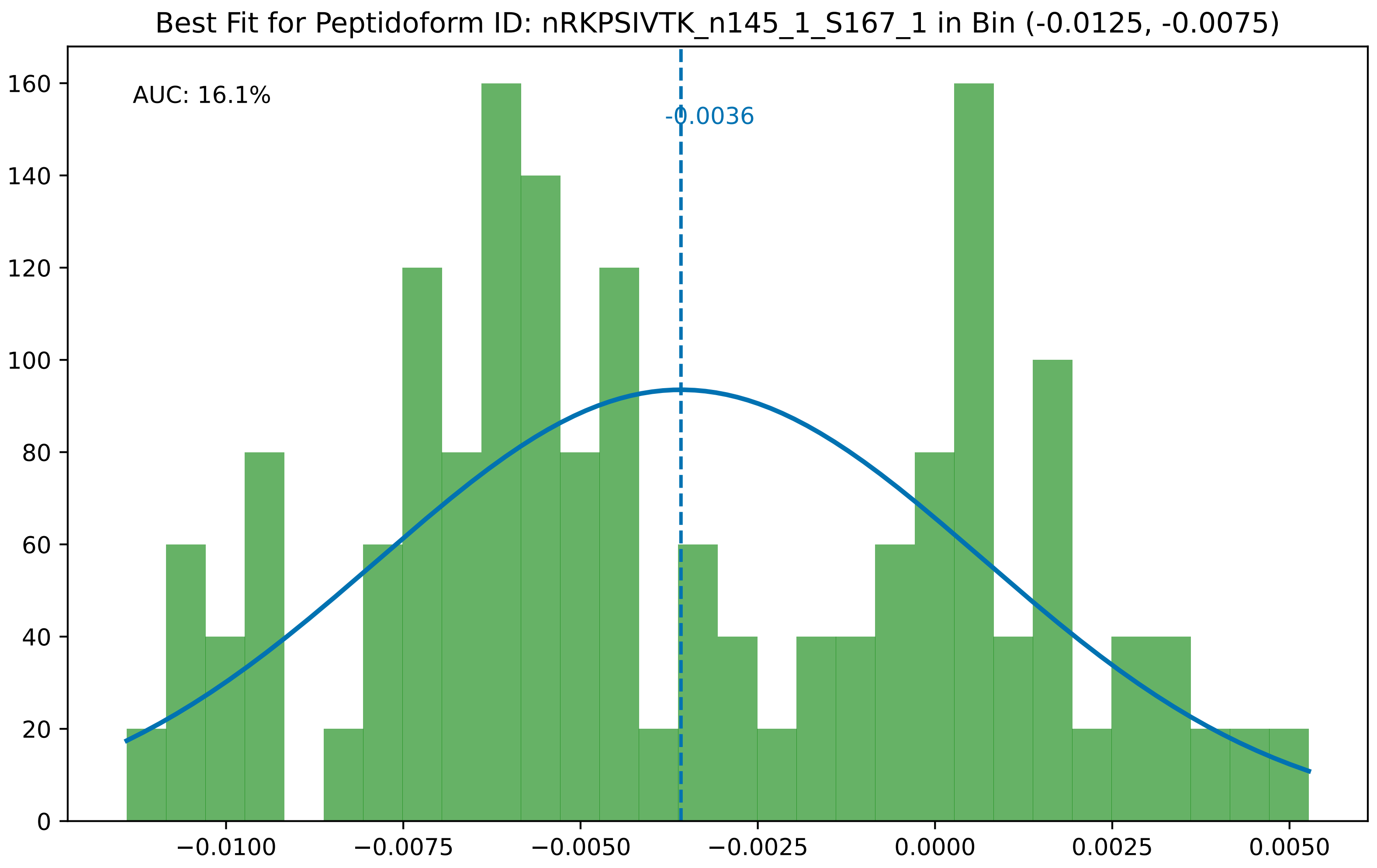
Best Fit for Peptidoform ID: nRKPSIVTK_n145_1_S167_1 in Bin (-0.0125, -0.0075)
AUC: 16.1%
-0.0036
0
20
40
60
80
100
120
140
160
−0.0100
−0.0075
−0.0050
−0.0025
0.0000
0.0025
0.0050

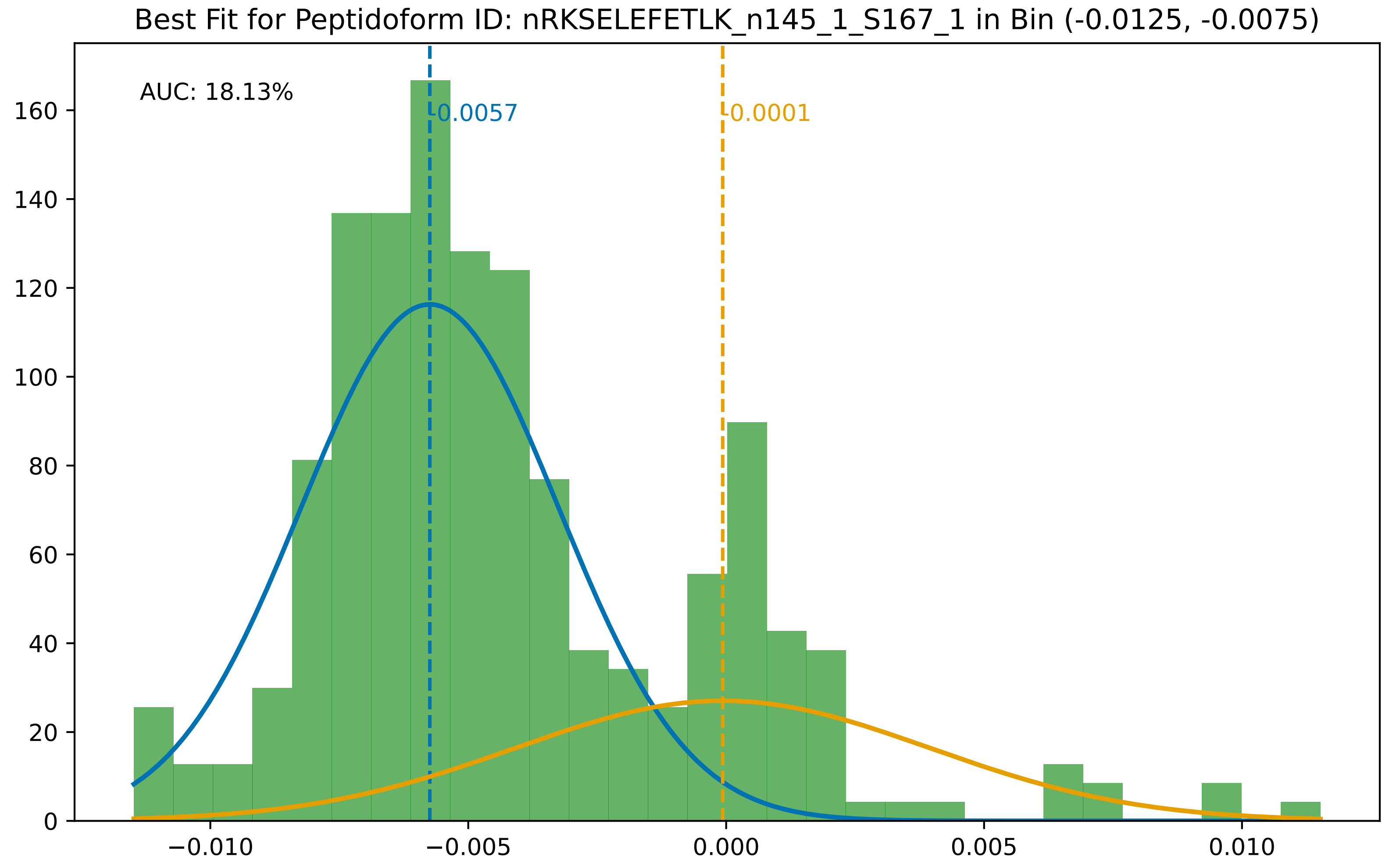
Best Fit for Peptidoform ID: nRKSELEFETLK_n145_1_S167_1 in Bin (-0.0125, -0.0075)
AUC: 18.13%
-0.0057
-0.0001
0
20
40
60
80
100
120
140
160
−0.010
−0.005
0.000
0.005
0.010

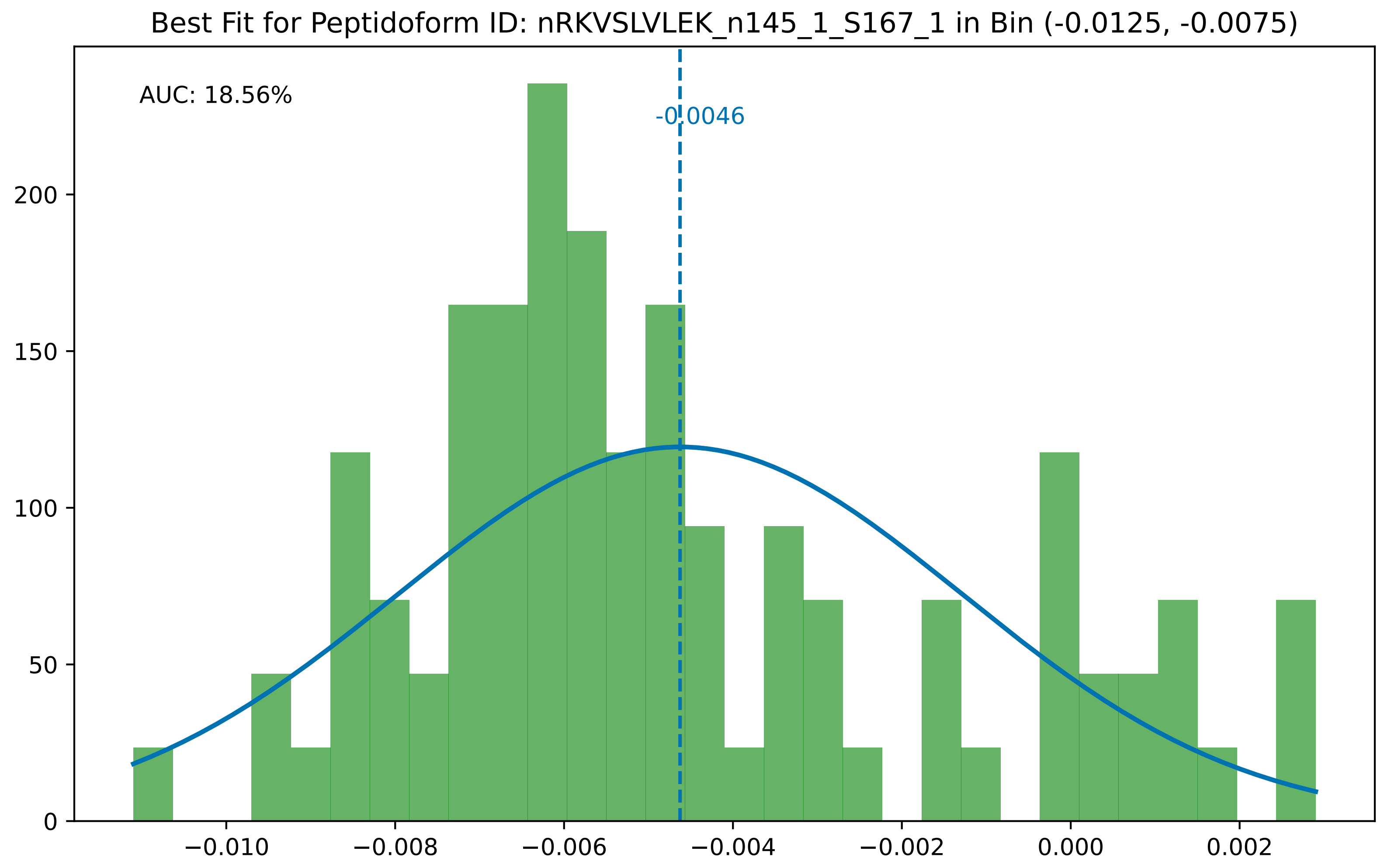

Best Fit for Peptidoform ID: nRKVSLVLEK_n145_1_S167_1 in Bin (-0.0125, -0.0075)
AUC: 18.56%
-0.0046
0
50
100
150
200
−0.010
−0.008
−0.006
−0.004
−0.002
0.000
0.002

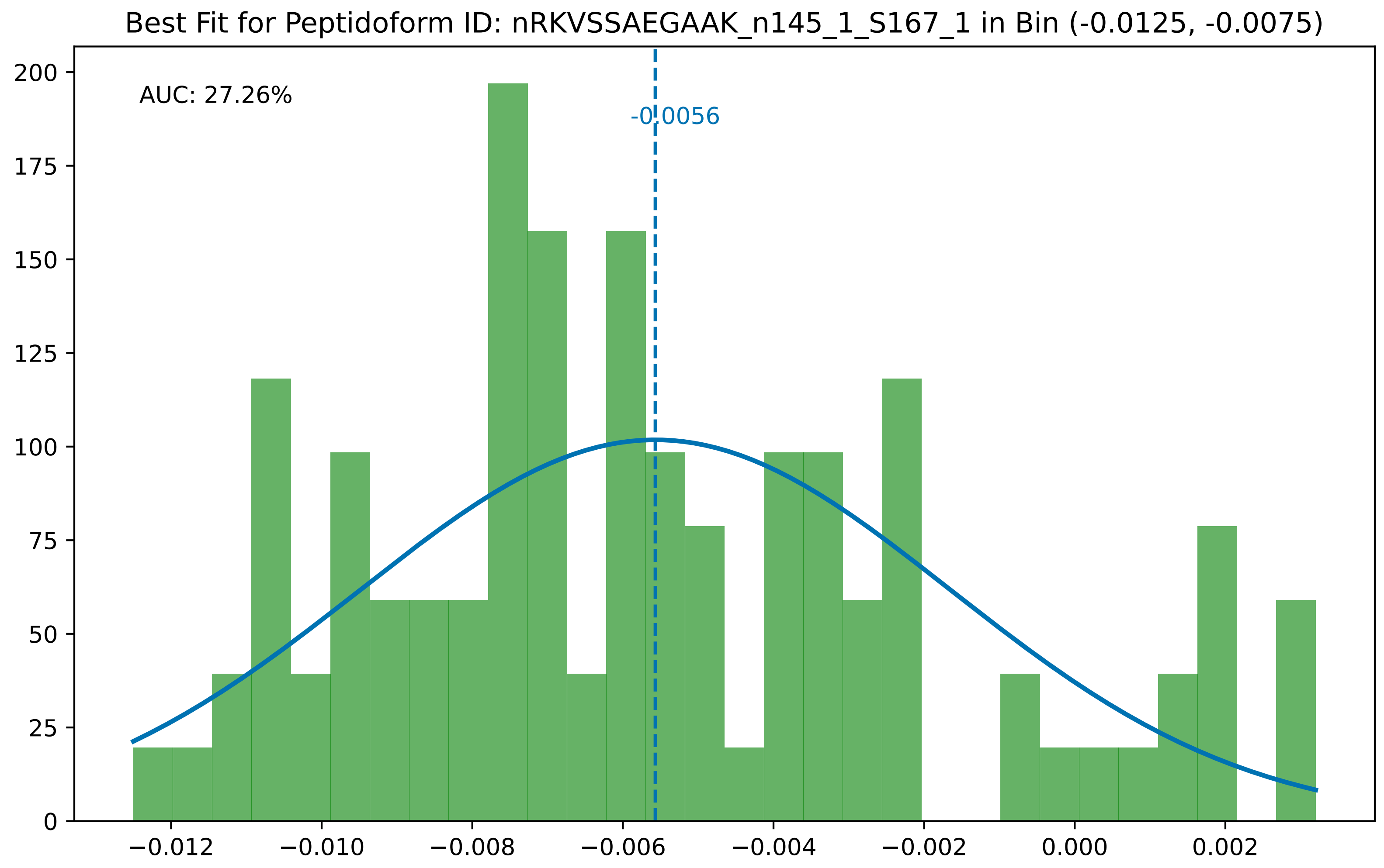
Best Fit for Peptidoform ID: nRKVSSAEGAAK_n145_1_S167_1 in Bin (-0.0125, -0.0075)
AUC: 27.26%
-0.0056
0
25
50
75
100
125
150
175
200
−0.012
−0.010
−0.008
−0.006
−0.004
−0.002
0.000
0.002

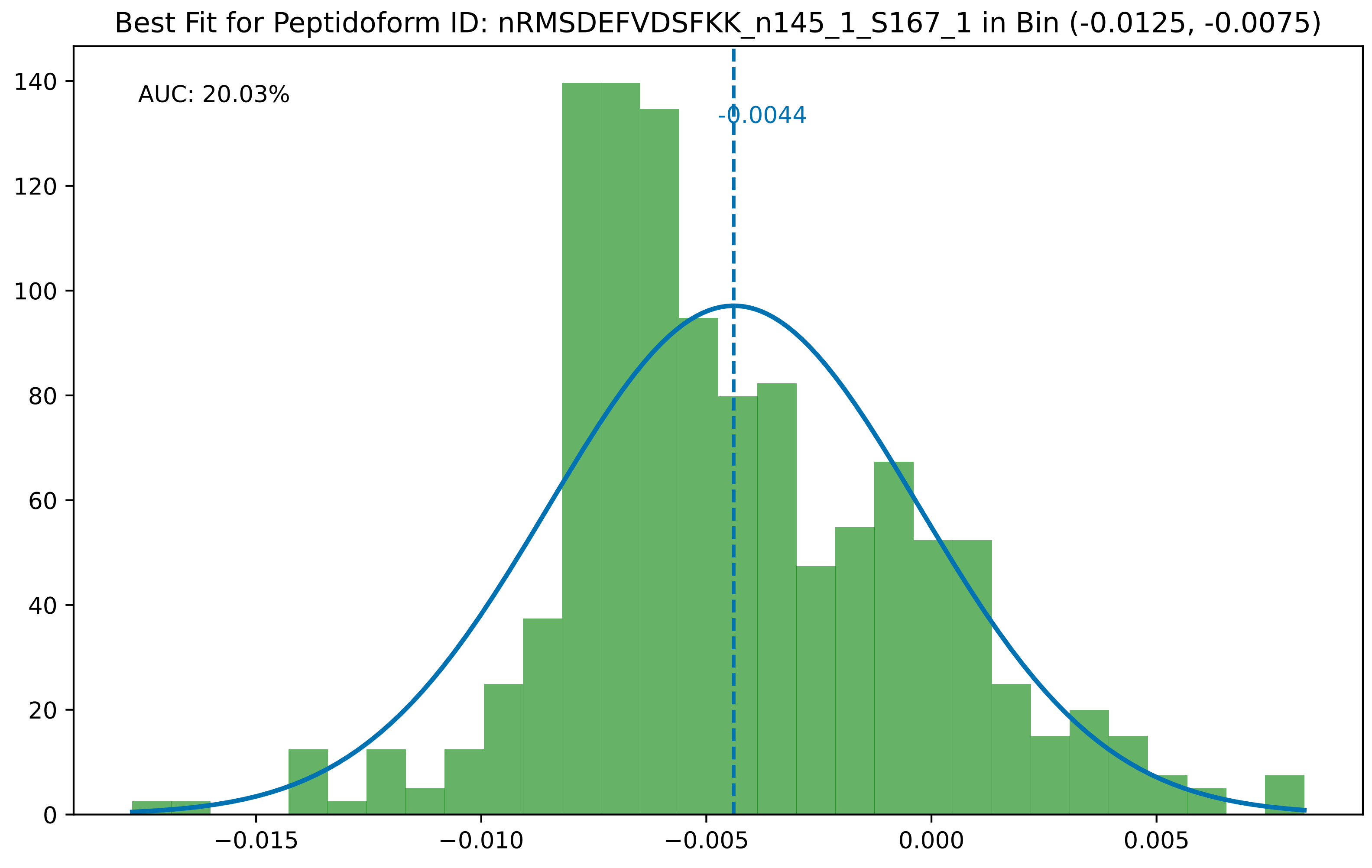
Best Fit for Peptidoform ID: nRMSDEFVDSFKK_n145_1_S167_1 in Bin (-0.0125, -0.0075)
AUC: 20.03%
-0.0044
0
20
40
60
80
100
120
140
−0.015
−0.010
−0.005
0.000
0.005

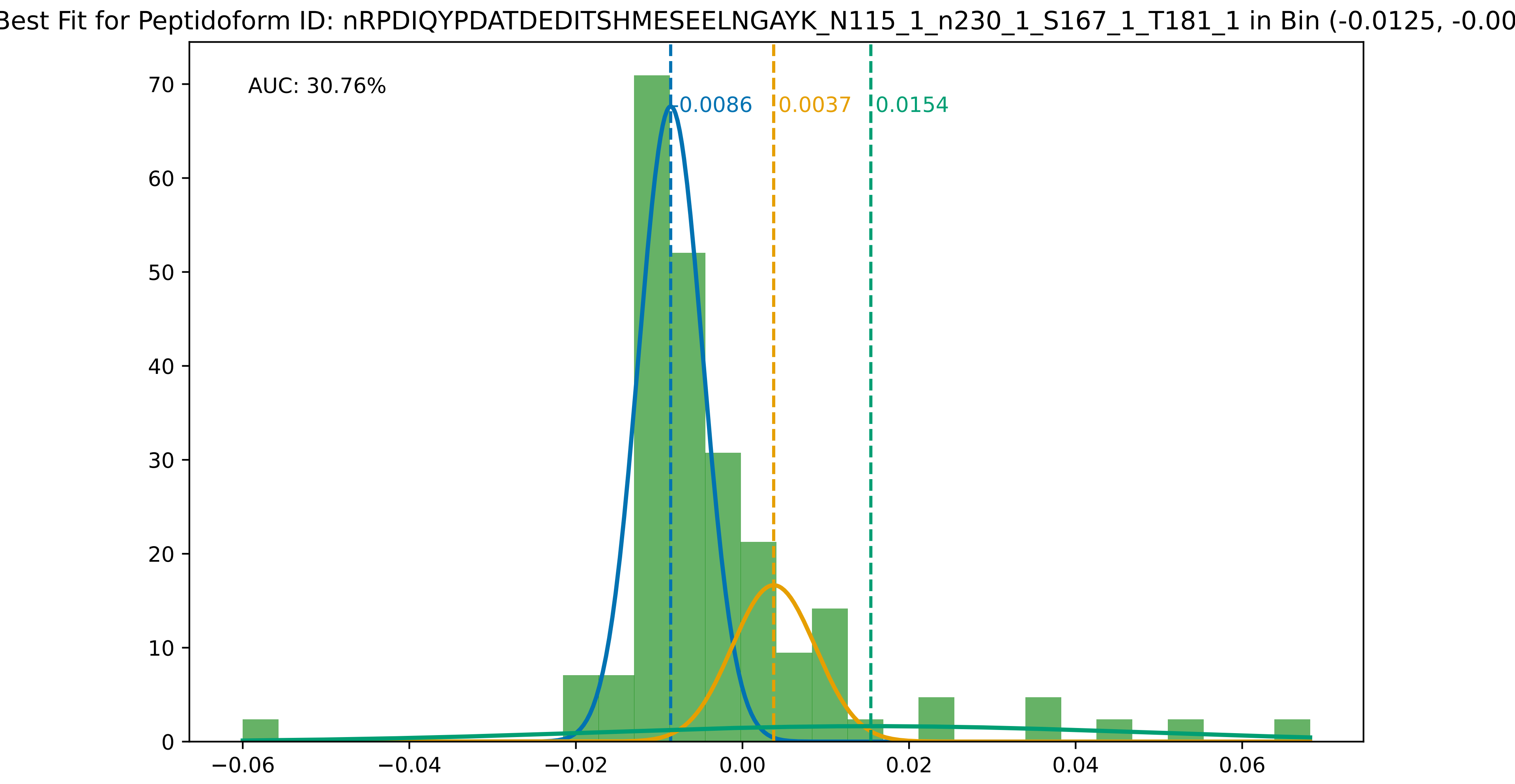
Best Fit for Peptidoform ID: nRPDIQYPDATDEDITSHMESEELNGAYK_N115_1_n230_1_S167_1_T181_1 in Bin (-0.0125, -0.00
AUC: 30.76%
-0.0086
0.0037
0.0154
0
10
20
30
40
50
60
70
−0.06
−0.04
−0.02
0.00
0.02
0.04
0.06

Best Fit for Peptidoform ID: nRPDIQYPDATDEDITSHMESEELNGAYK_n230_1_S167_1_T181_1 in Bin (-0.0125, -0.0075)

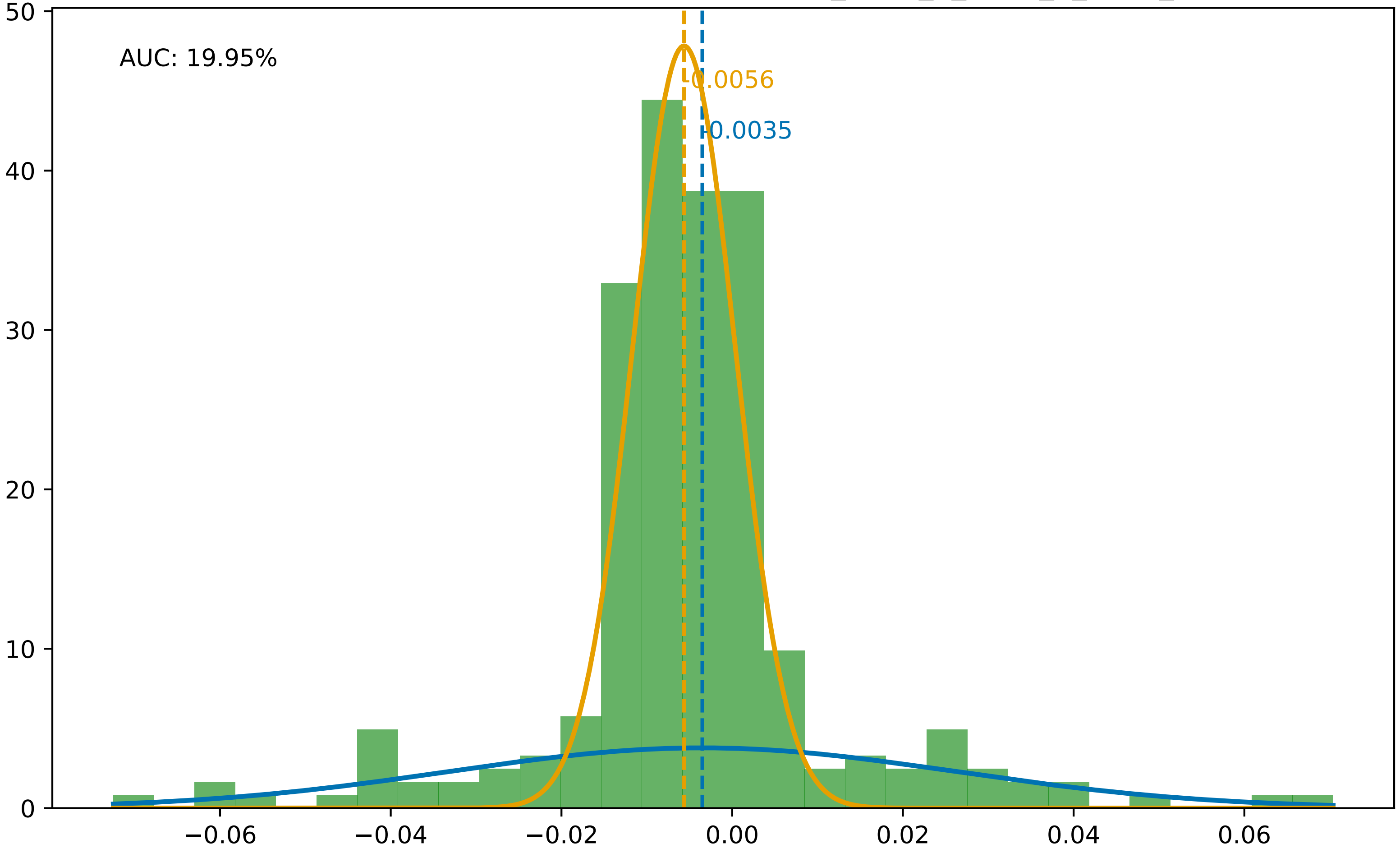

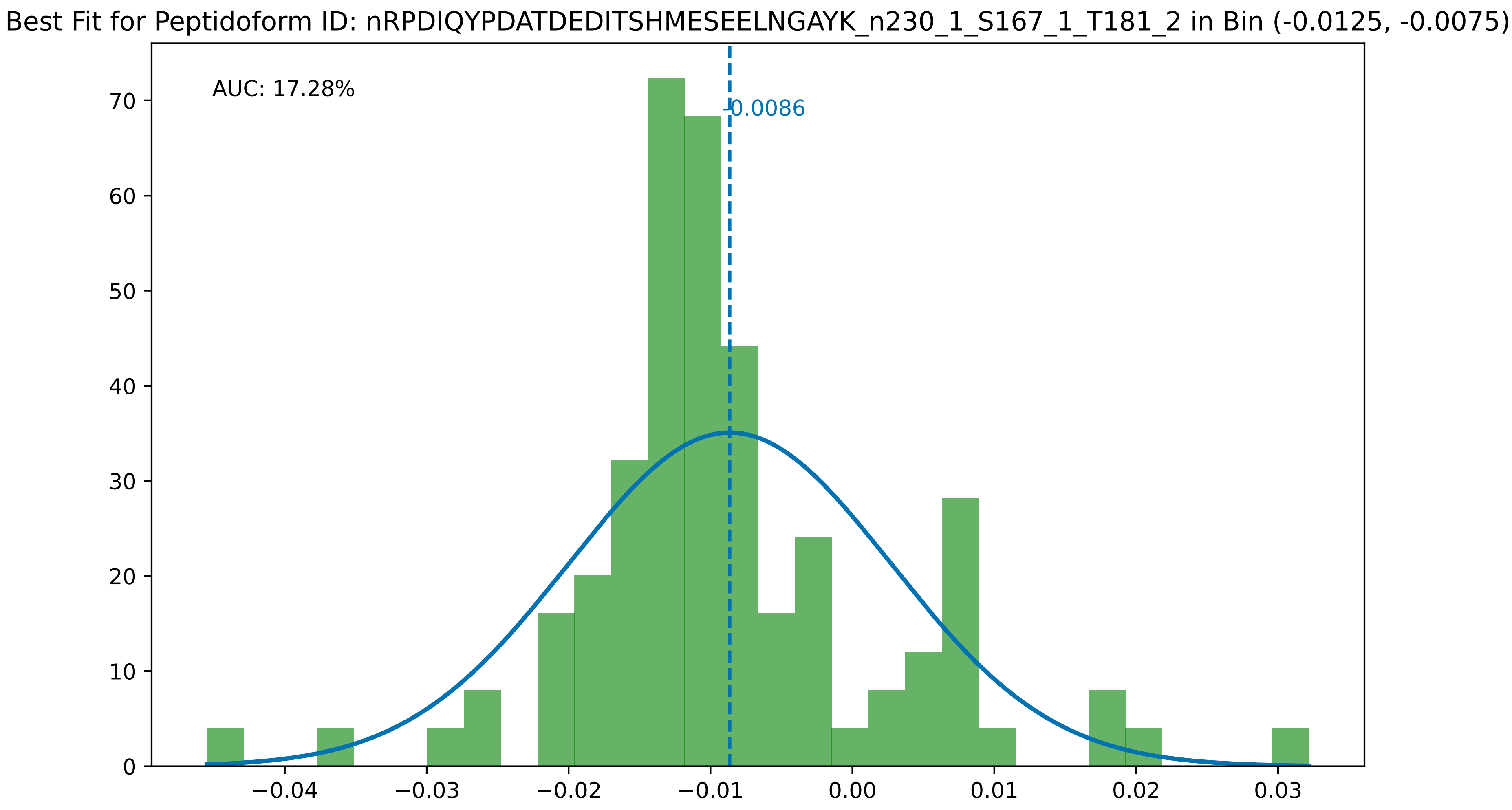

Best Fit for Peptidoform ID: nRPDIQYPDATDEDITSHMESEELNGAYK_n230_1_S167_1_T181_2 in Bin (-0.0125, -0.0075)
AUC: 17.28%
-0.0086
0
10
20
30
40
50
60
70
−0.04
−0.03
−0.02
−0.01
0.00
0.01
0.02
0.03

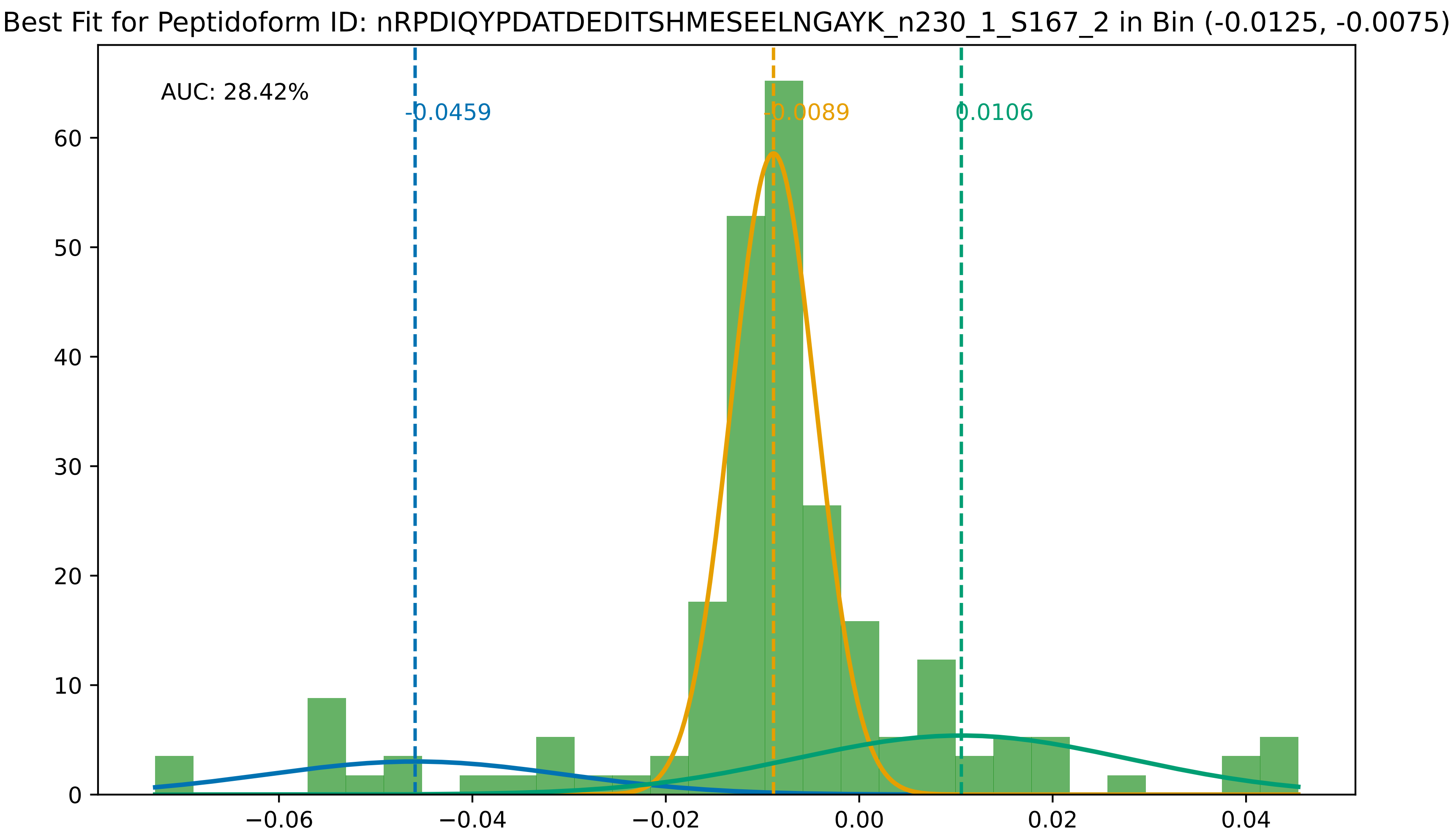

Best Fit for Peptidoform ID: nRPDIQYPDATDEDITSHMESEELNGAYK_n230_1_S167_2 in Bin (-0.0125, -0.0075)
AUC: 28.42%
-0.0459
-0.0089
0.0106
0
10
20
30
40
50
60
−0.06
−0.04
−0.02
0.00
0.02
0.04

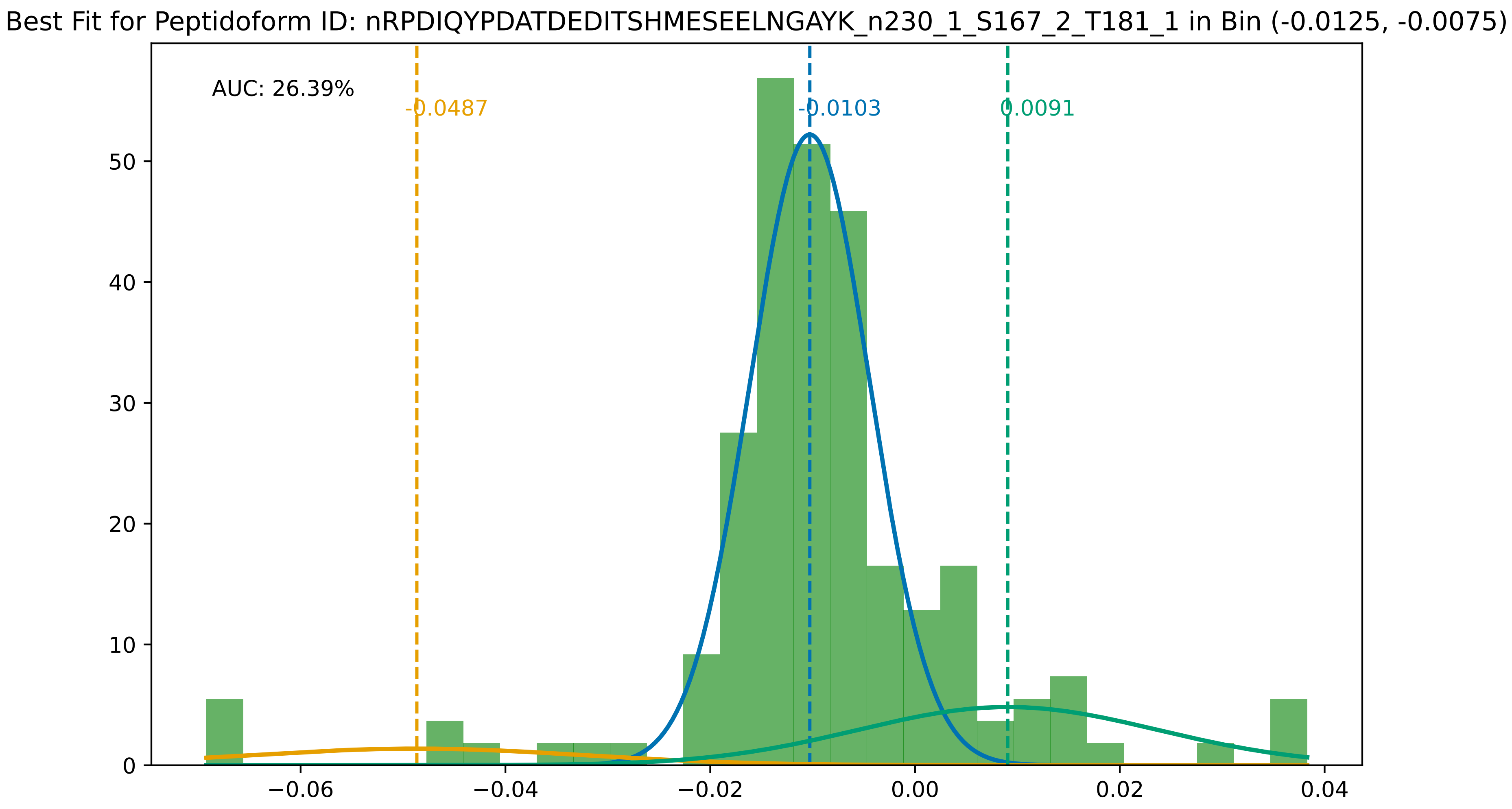
Best Fit for Peptidoform ID: nRPDIQYPDATDEDITSHMESEELNGAYK_n230_1_S167_2_T181_1 in Bin (-0.0125, -0.0075)
AUC: 26.39%
-0.0487
-0.0103
0.0091
0
10
20
30
40
50
−0.06
−0.04
−0.02
0.00
0.02
0.04

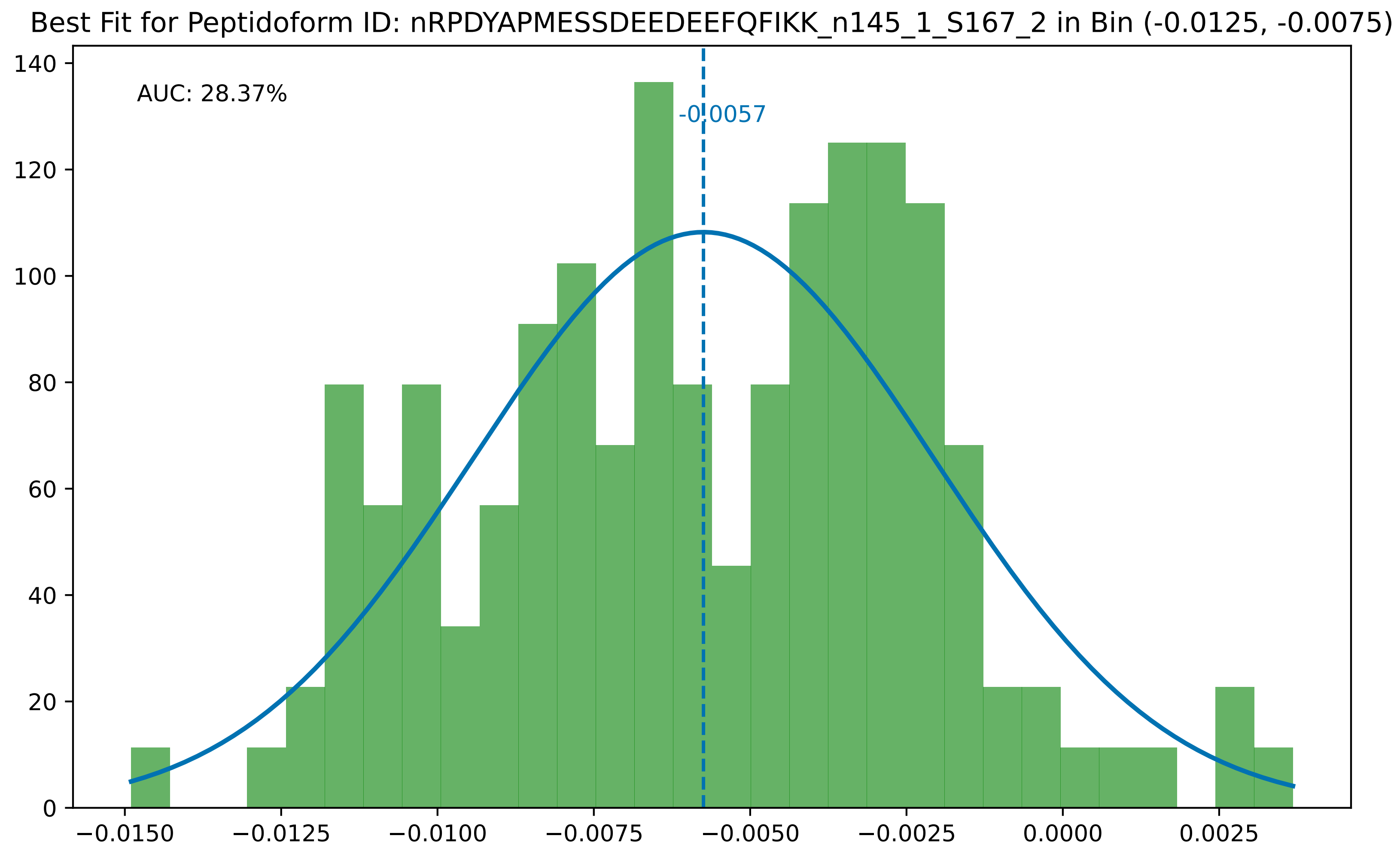

Best Fit for Peptidoform ID: nRPDYAPMESSDEEDEEFQFIKK_n145_1_S167_2 in Bin (-0.0125, -0.0075)
AUC: 28.37%
-0.0057
0
20
40
60
80
100
120
140
−0.0150
−0.0125
−0.0100
−0.0075
−0.0050
−0.0025
0.0000
0.0025

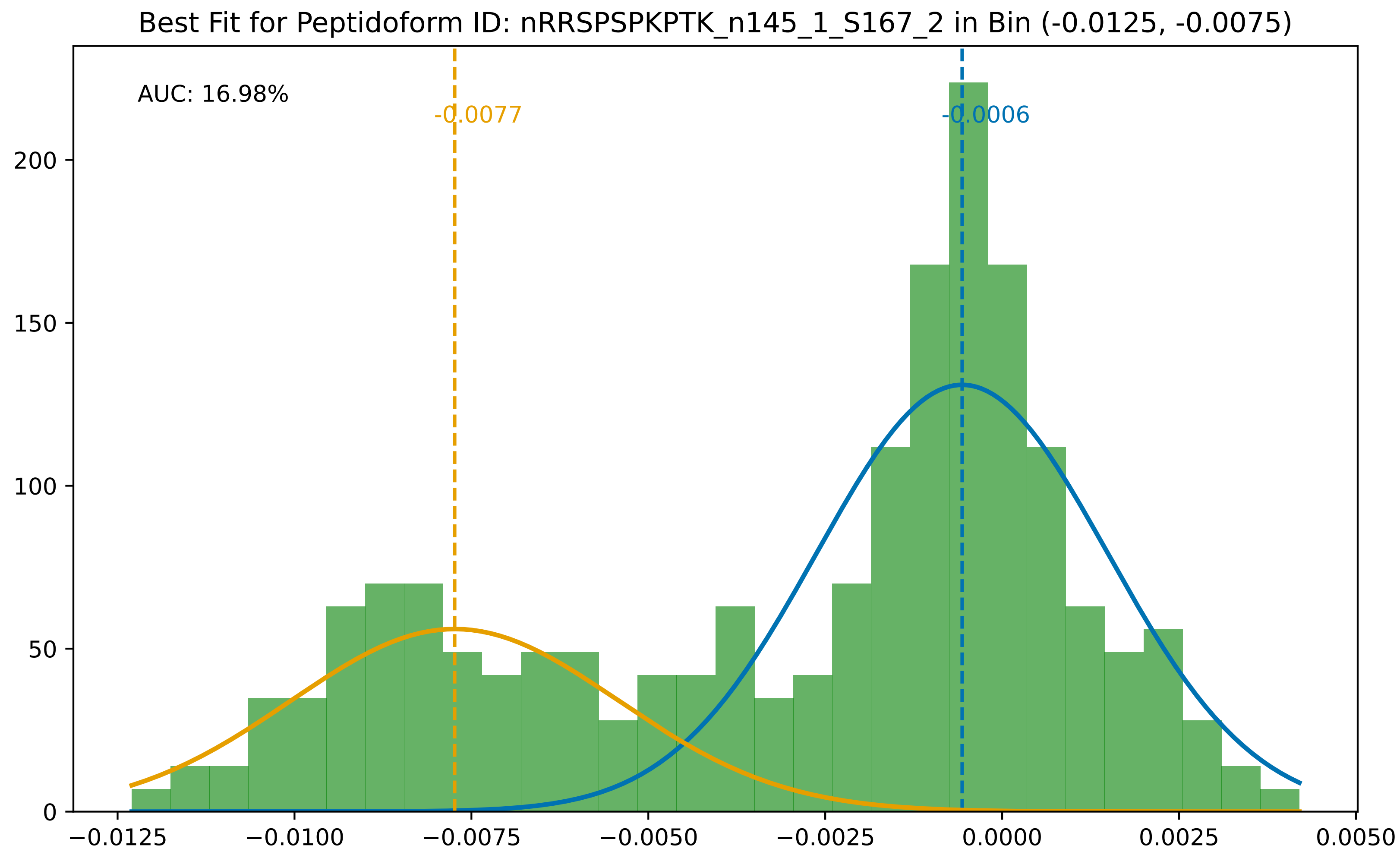
Best Fit for Peptidoform ID: nRRSPSPKPTK_n145_1_S167_2 in Bin (-0.0125, -0.0075)
AUC: 16.98%
-0.0077
-0.0006
200
150
100
50
0
−0.0125
−0.0100
−0.0075
−0.0050
−0.0025
0.0000
0.0025
0.0050

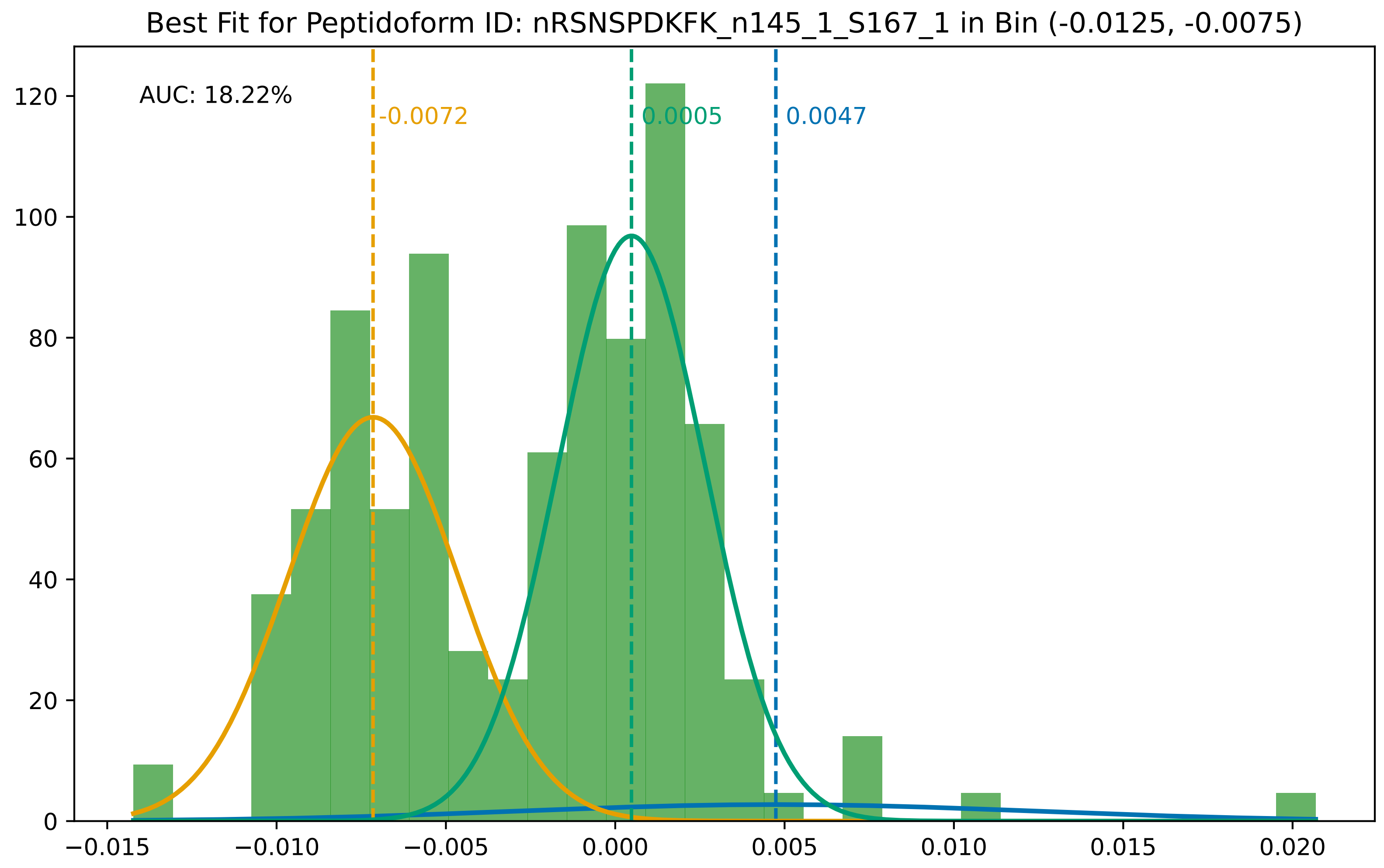
Best Fit for Peptidoform ID: nRSNSPDKFK_n145_1_S167_1 in Bin (-0.0125, -0.0075)
AUC: 18.22%
-0.0072
0.0005
0.0047
0
20
40
60
80
100
120
−0.015
−0.010
−0.005
0.000
0.005
0.010
0.015
0.020

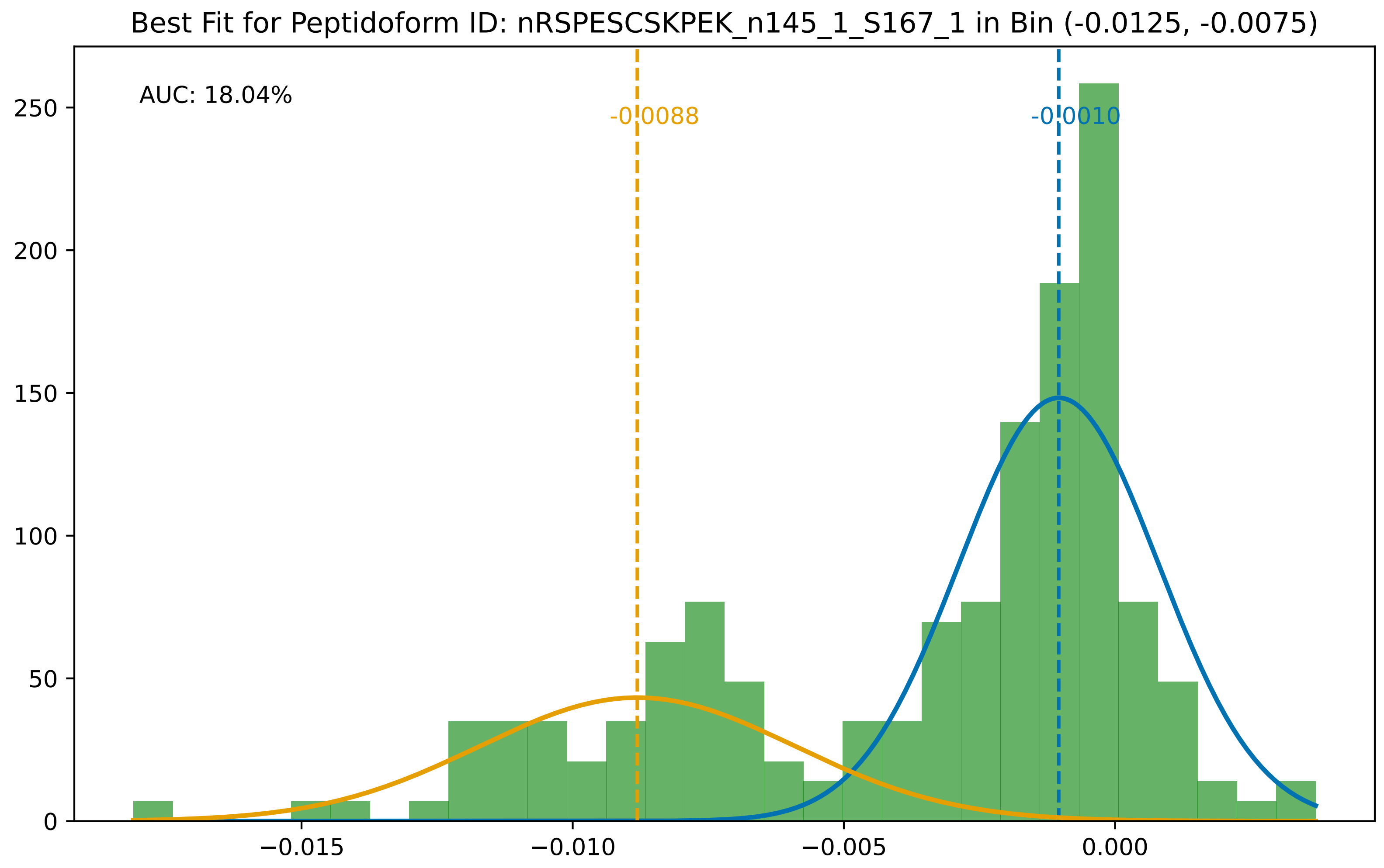

Best Fit for Peptidoform ID: nRSPESCSKPEK_n145_1_S167_1 in Bin (-0.0125, -0.0075)
AUC: 18.04%
-0.0088
-0.0010
0
50
100
150
200
250
−0.015
−0.010
−0.005
0.000

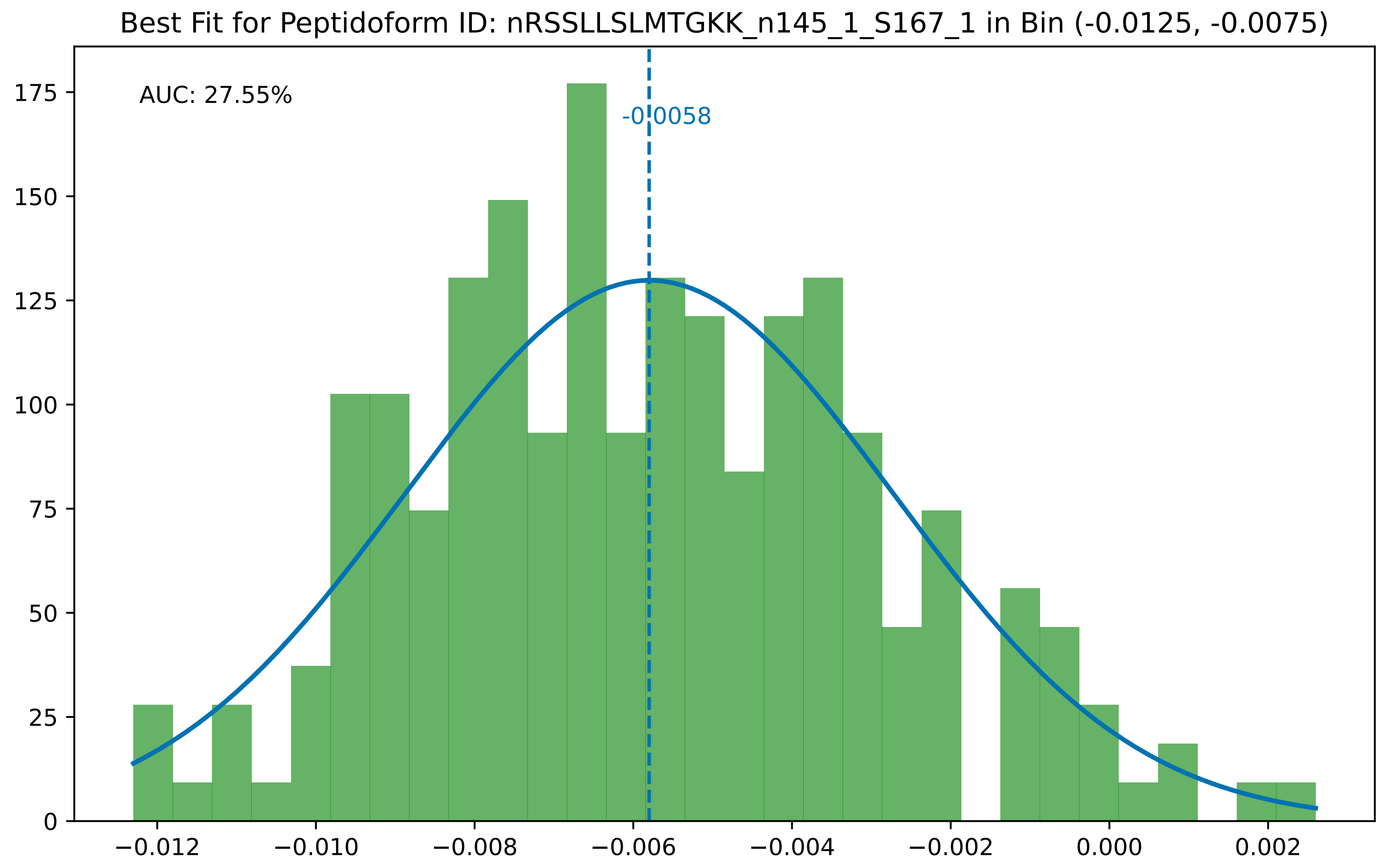

Best Fit for Peptidoform ID: nRSSLLSLMTGKK_n145_1_S167_1 in Bin (-0.0125, -0.0075)
AUC: 27.55%
-0.0058
175
150
125
100
75
50
25
0
−0.012
−0.010
−0.008
−0.006
−0.004
−0.002
0.000
0.002

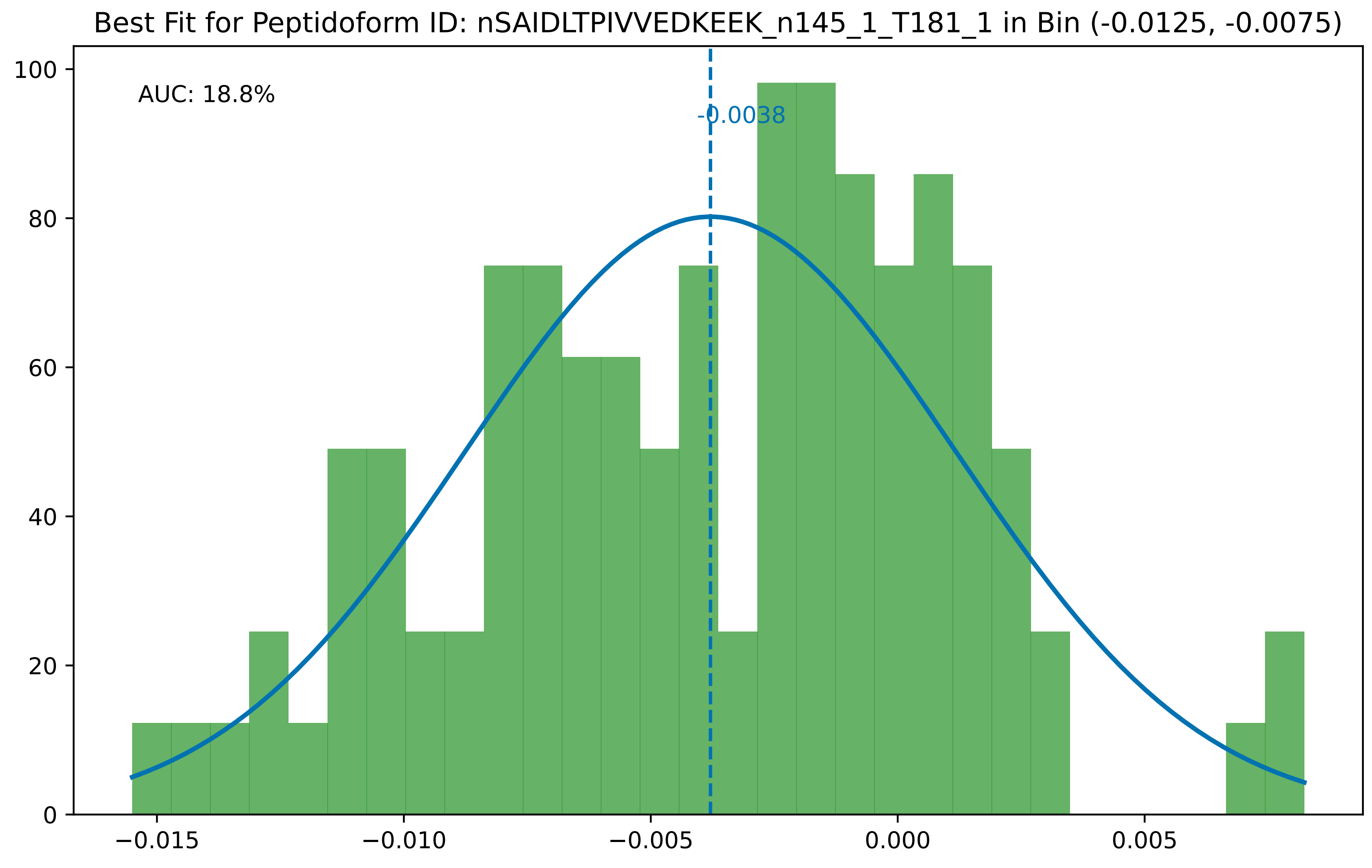

Best Fit for Peptidoform ID: nSAIDLTPIVVEDKEEK_n145_1_T181_1 in Bin (-0.0125, -0.0075)
AUC: 18.8%
-0.0038
0
20
40
60
80
100
−0.015
−0.010
−0.005
0.000
0.005

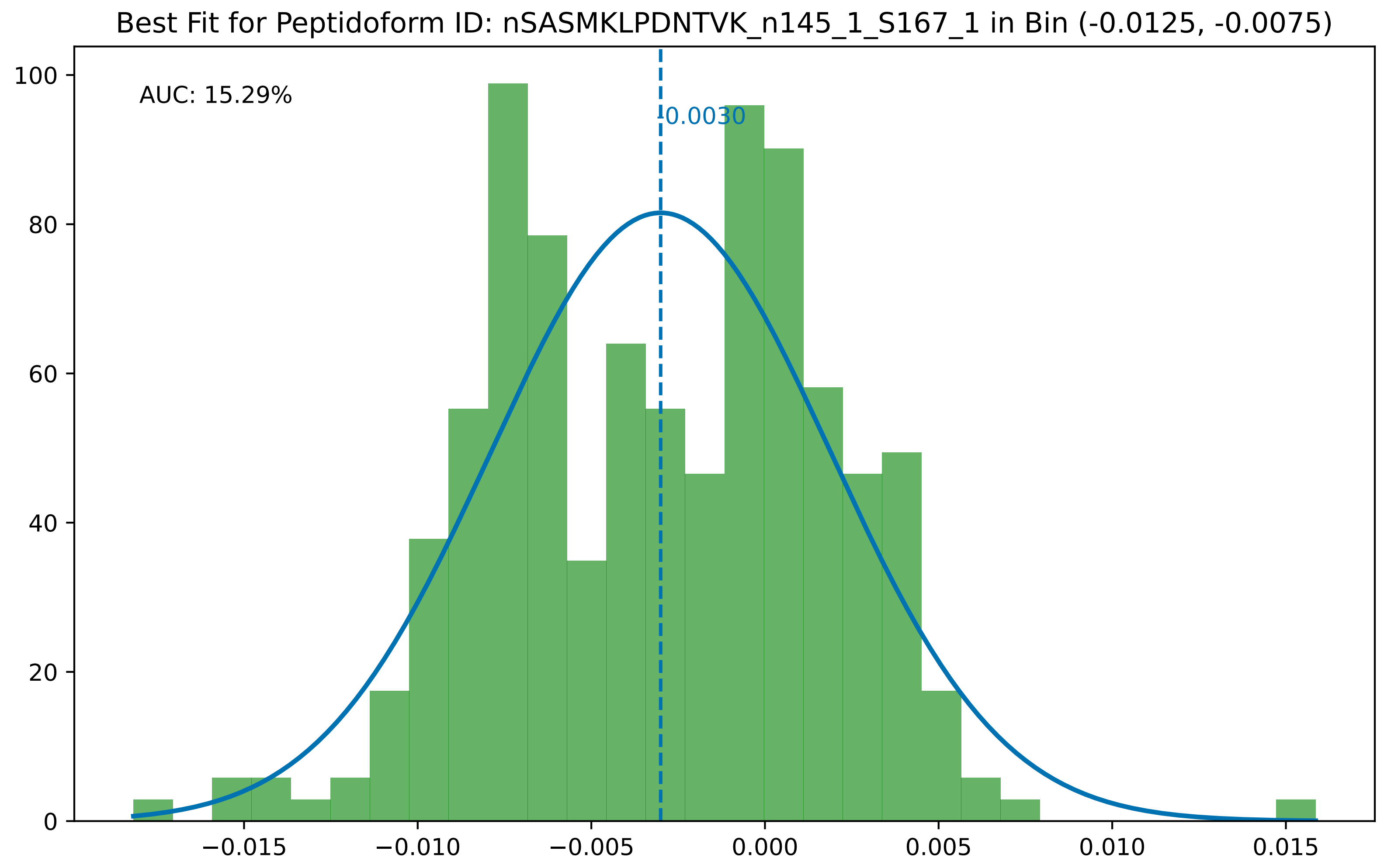

Best Fit for Peptidoform ID: nSASMKLPDNTVK_n145_1_S167_1 in Bin (-0.0125, -0.0075)
AUC: 15.29%
-0.0030
0
20
40
60
80
100
−0.015
−0.010
−0.005
0.000
0.005
0.010
0.015

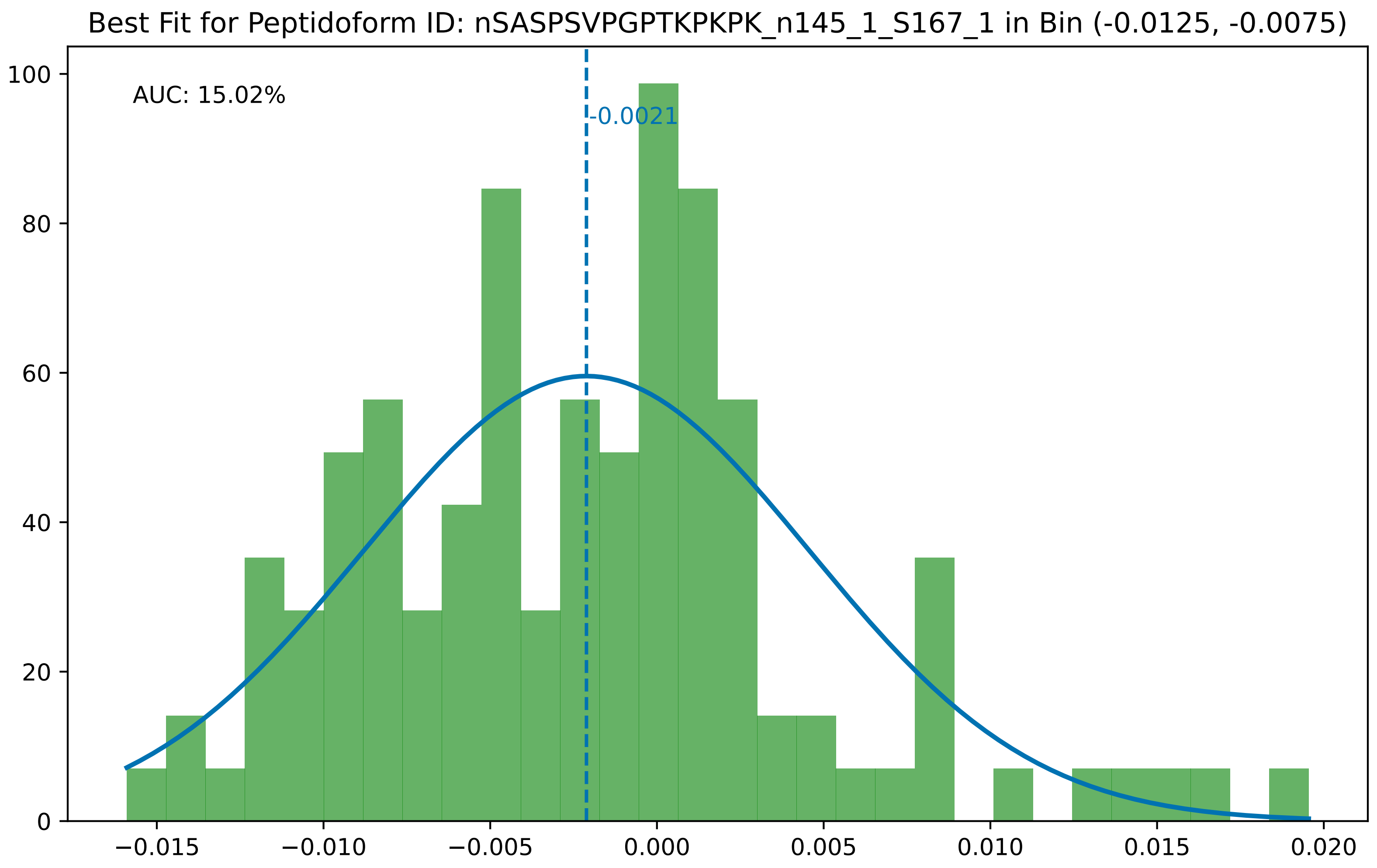

Best Fit for Peptidoform ID: nSASPSVPGPTKPKPK_n145_1_S167_1 in Bin (-0.0125, -0.0075)
AUC: 15.02%
-0.0021
0
20
40
60
80
100
−0.015
−0.010
−0.005
0.000
0.005
0.010
0.015
0.020

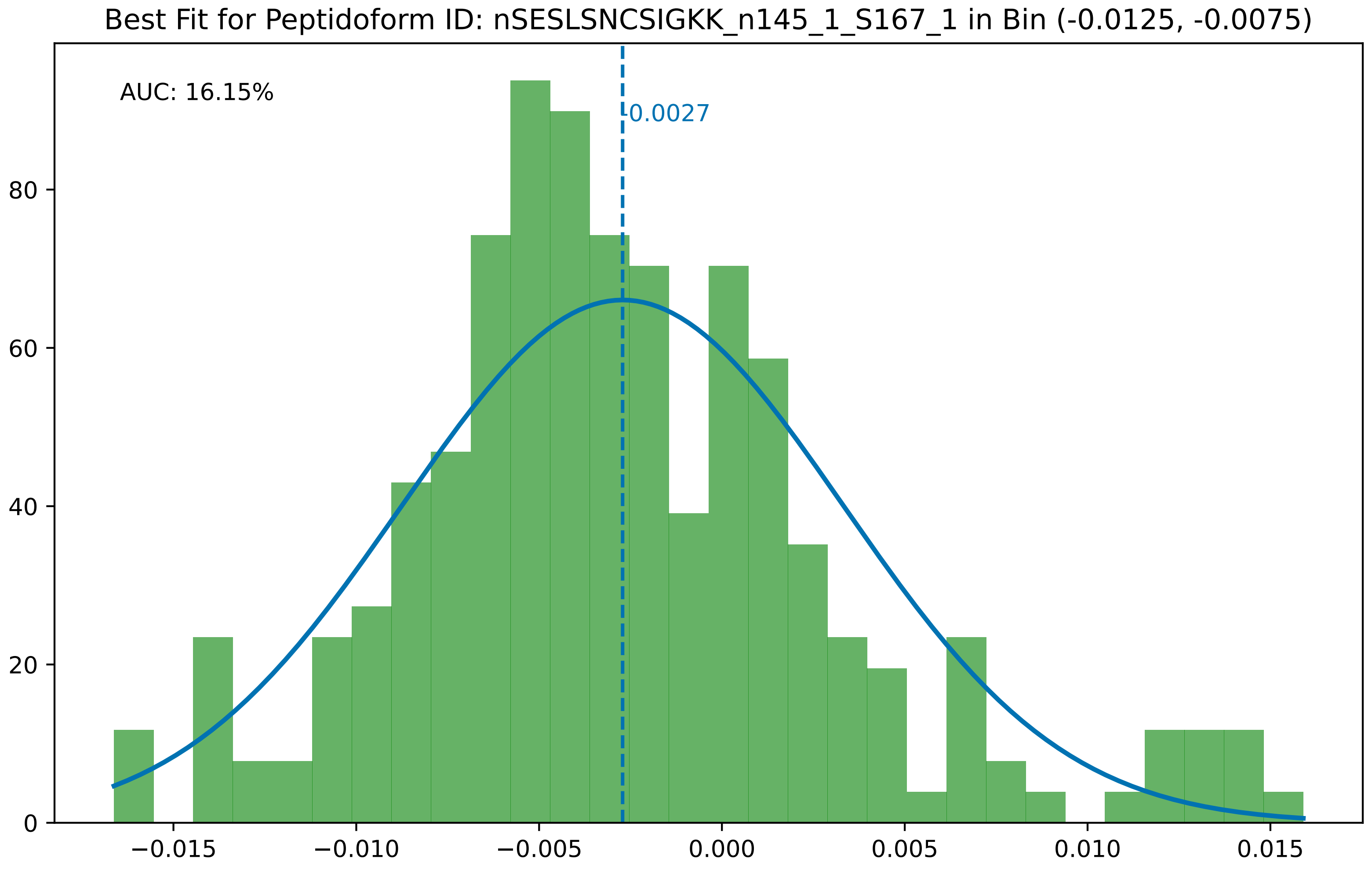

Best Fit for Peptidoform ID: nSESLSNCSIGKK_n145_1_S167_1 in Bin (-0.0125, -0.0075)
AUC: 16.15%
-0.0027
0
20
40
60
80
−0.015
−0.010
−0.005
0.000
0.005
0.010
0.015

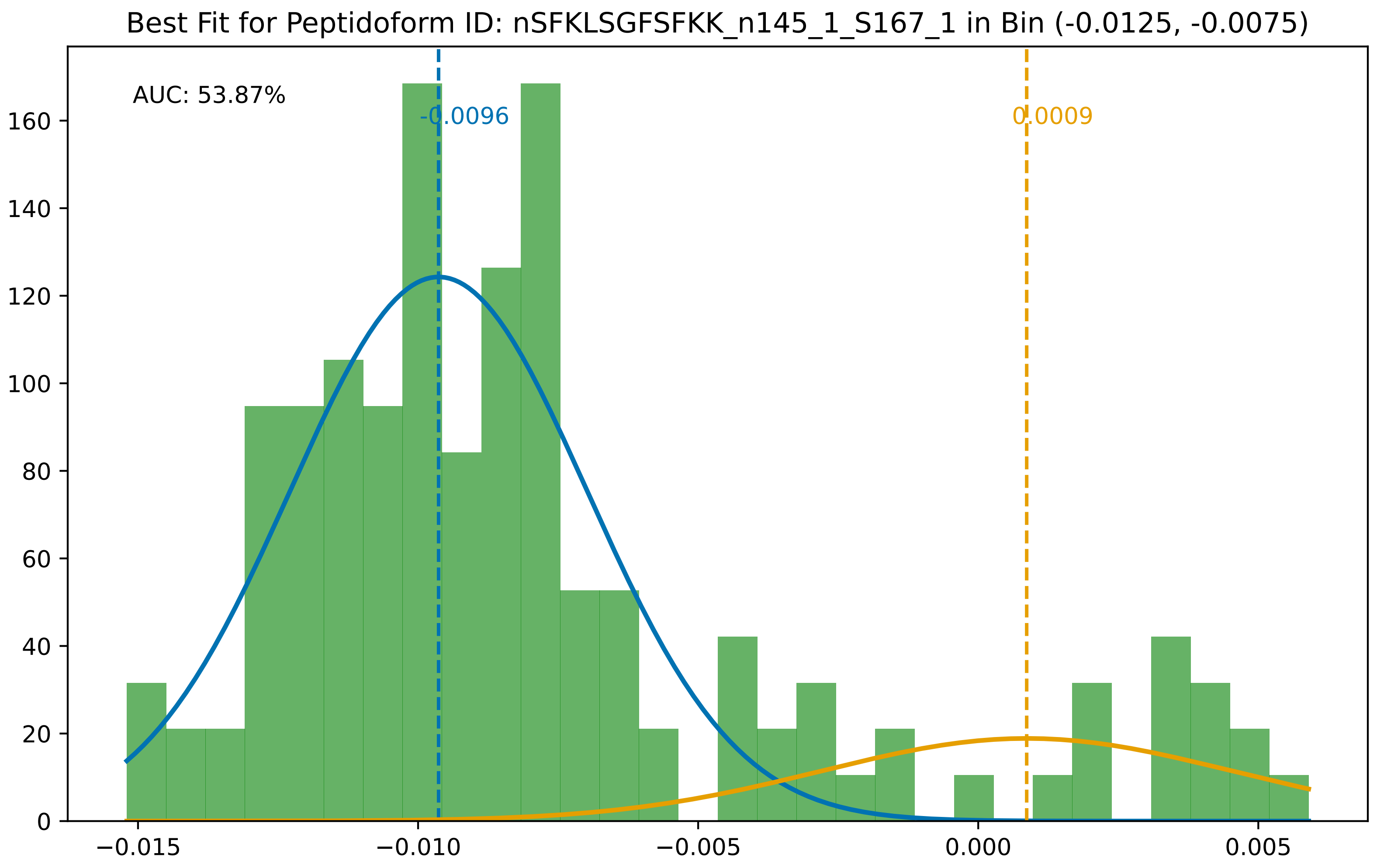
Best Fit for Peptidoform ID: nSFKLSGFSFKK_n145_1_S167_1 in Bin (-0.0125, -0.0075)
AUC: 53.87%
-0.0096
0.0009
0
20
40
60
80
100
120
140
160
−0.015
−0.010
−0.005
0.000
0.005

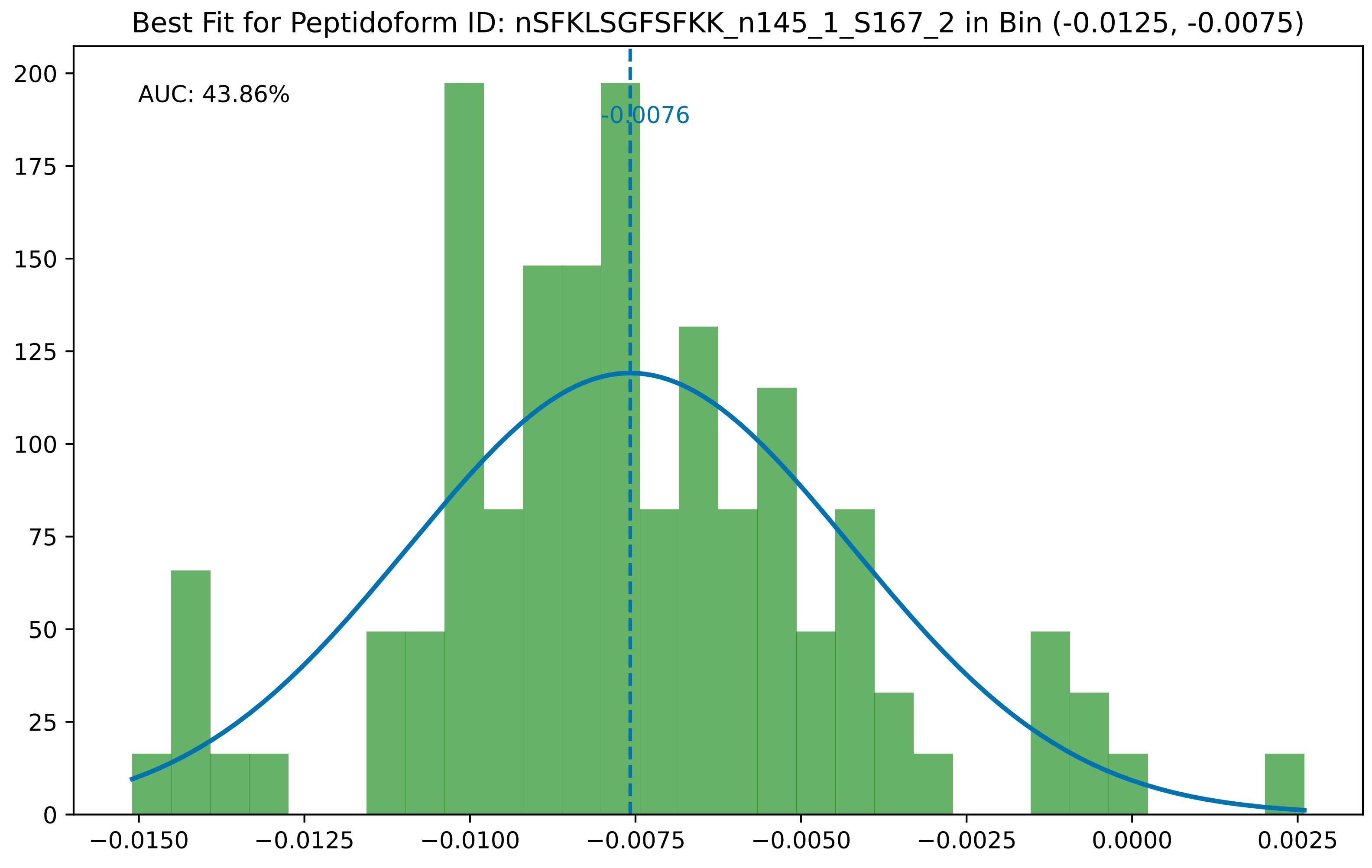
Best Fit for Peptidoform ID: nSFKLSGFSFKK_n145_1_S167_2 in Bin (-0.0125, -0.0075)
AUC: 43.86%
-0.0076
200
175
150
125
100
75
50
25
0
−0.0150
−0.0125
−0.0100
−0.0075
−0.0050
−0.0025
0.0000
0.0025

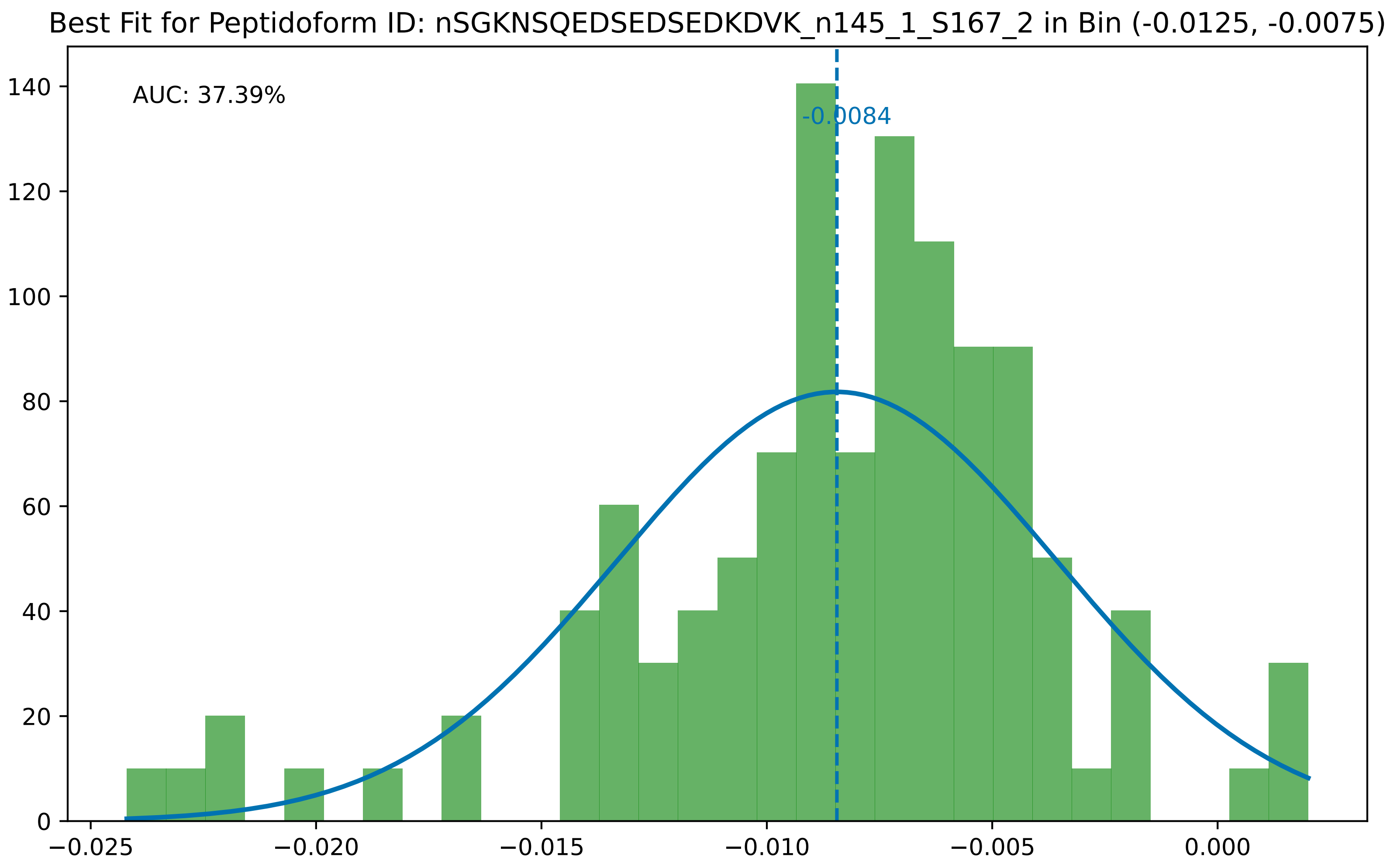

Best Fit for Peptidoform ID: nSGKNSQEDSEDSEDKDVK_n145_1_S167_2 in Bin (-0.0125, -0.0075)
AUC: 37.39%
-0.0084
0
20
40
60
80
100
120
140
−0.025
−0.020
−0.015
−0.010
−0.005
0.000

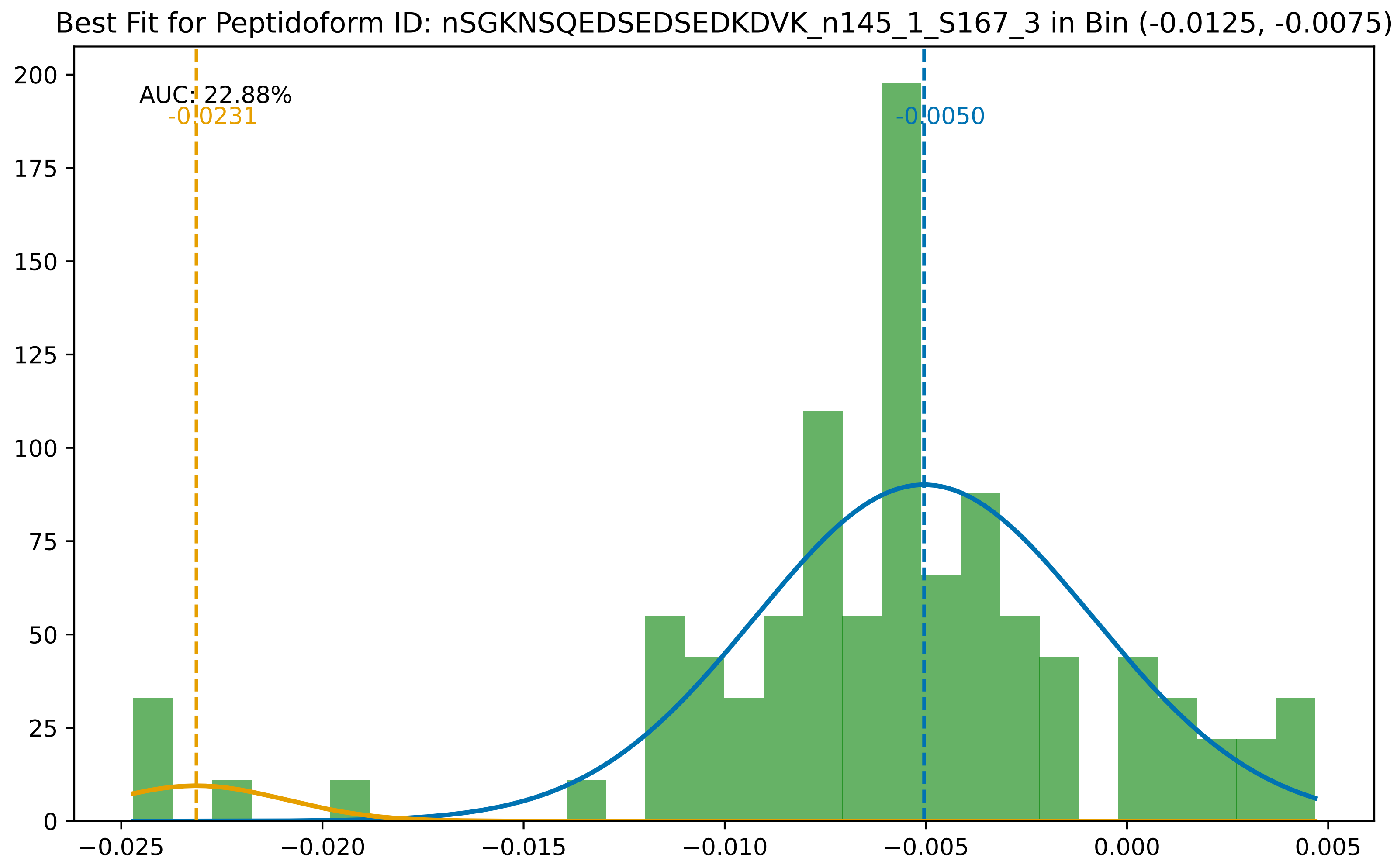

Best Fit for Peptidoform ID: nSGKNSQEDSEDSEDKDVK_n145_1_S167_3 in Bin (-0.0125, -0.0075)
AUC: 22.88%
-0.0231
-0.0050
200
175
150
125
100
75
50
25
0
−0.025
−0.020
−0.015
−0.010
−0.005
0.000
0.005

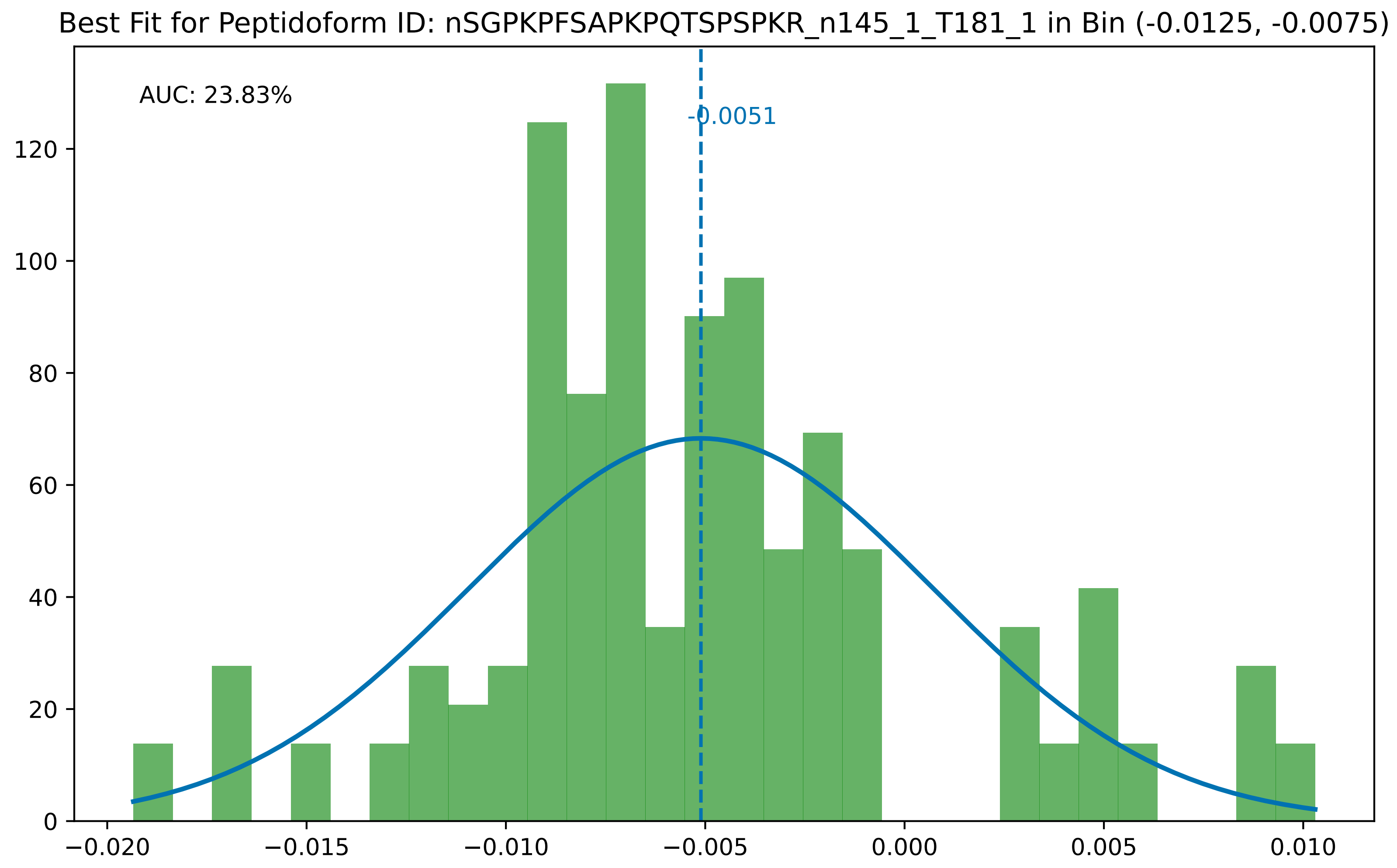

Best Fit for Peptidoform ID: nSGPKPFSAPKPQTSPSPKR_n145_1_T181_1 in Bin (-0.0125, -0.0075)
AUC: 23.83%
-0.0051
0
20
40
60
80
100
120
−0.020
−0.015
−0.010
−0.005
0.000
0.005
0.010

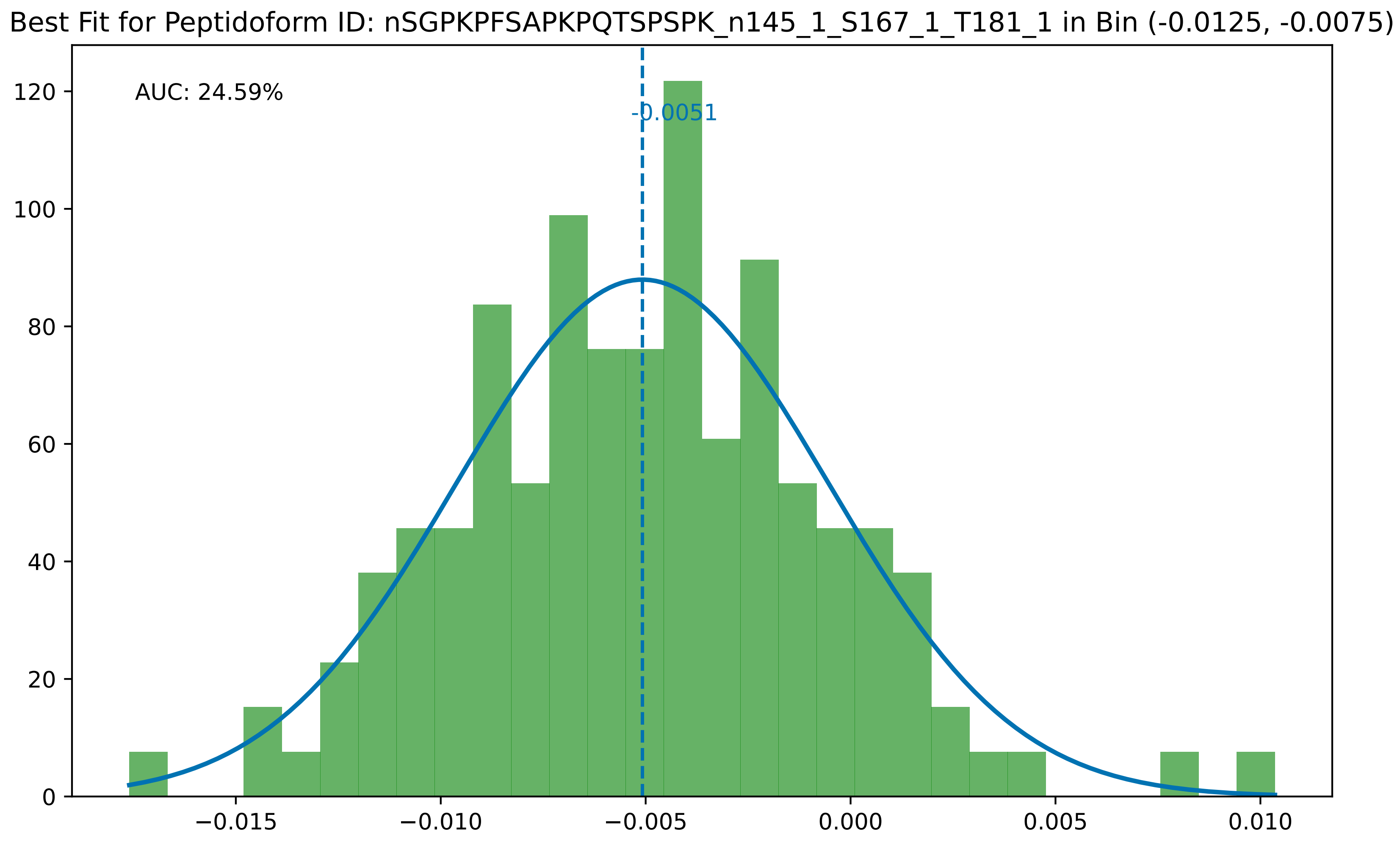
Best Fit for Peptidoform ID: nSGPKPFSAPKPQTSPSPK_n145_1_S167_1_T181_1 in Bin (-0.0125, -0.0075)
AUC: 24.59%
-0.0051
0
20
40
60
80
100
120
−0.015
−0.010
−0.005
0.000
0.005
0.010

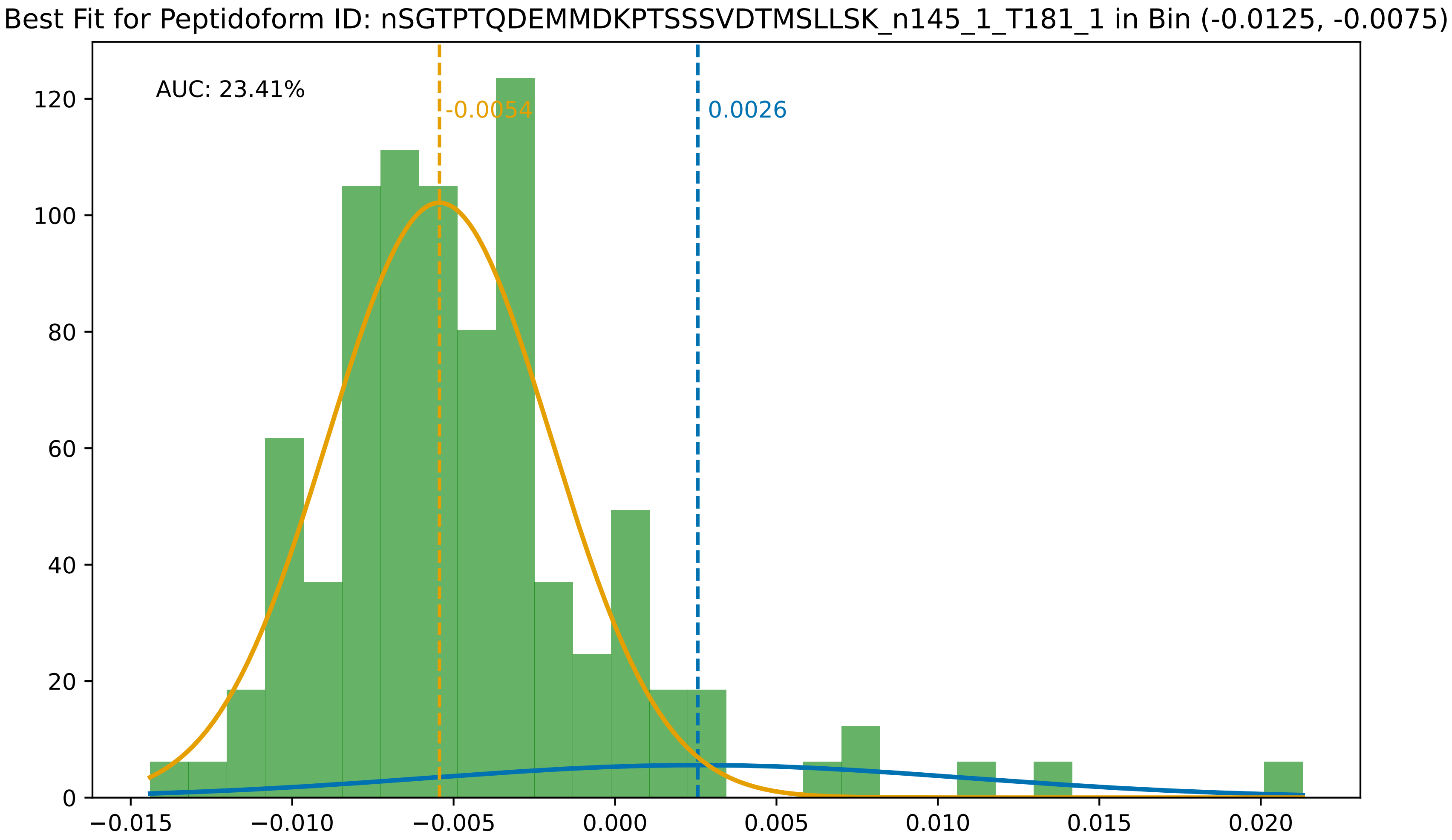

Best Fit for Peptidoform ID: nSGTPTQDEMMDKPTSSSVDTMSLLSK_n145_1_T181_1 in Bin (-0.0125, -0.0075)
AUC: 23.41%
-0.0054
0.0026
0
20
40
60
80
100
120
−0.015
−0.010
−0.005
0.000
0.005
0.010
0.015
0.020

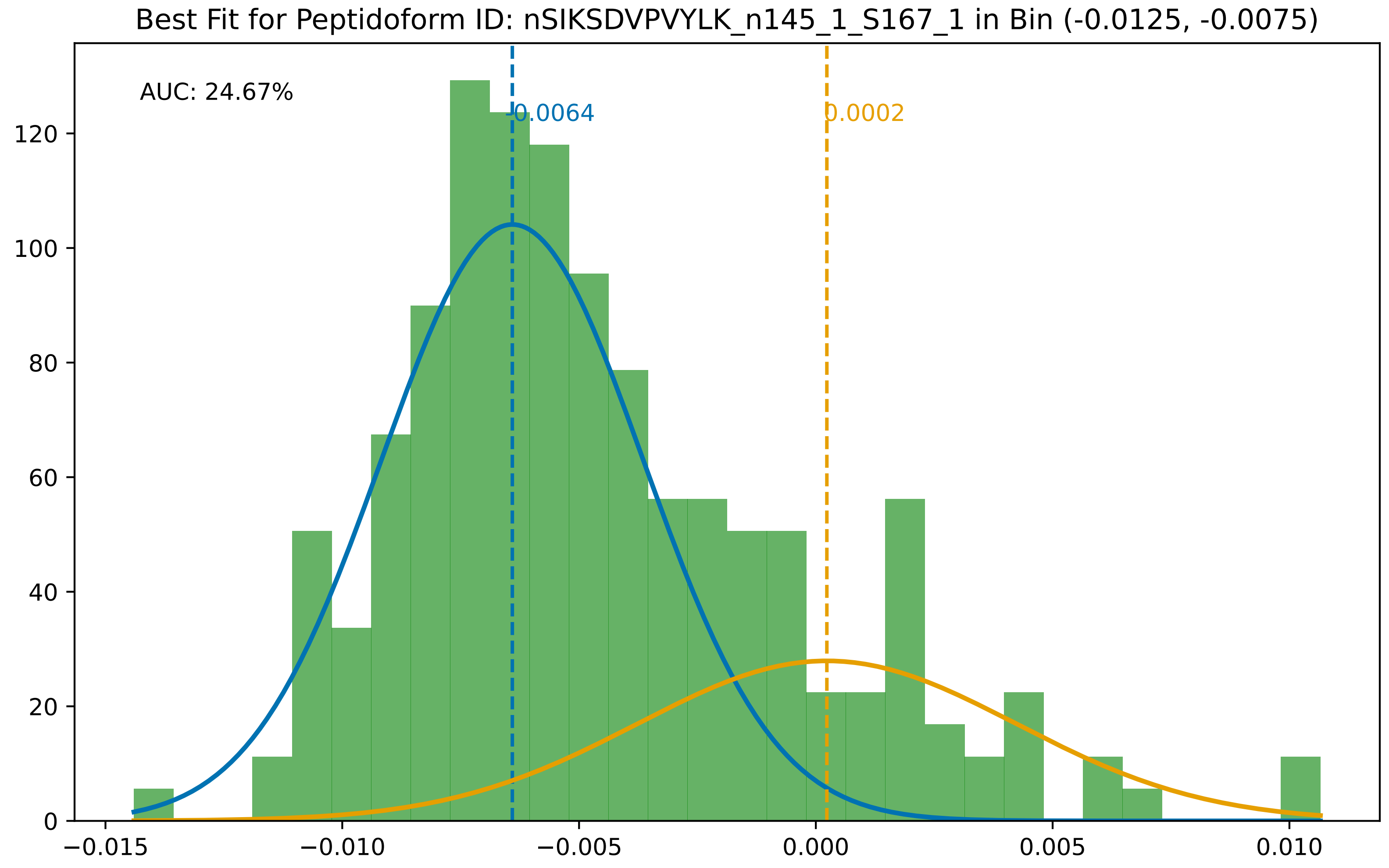

Best Fit for Peptidoform ID: nSIKSDVPVYLK_n145_1_S167_1 in Bin (-0.0125, -0.0075)
AUC: 24.67%
-0.0064
0.0002
0
20
40
60
80
100
120
−0.015
−0.010
−0.005
0.000
0.005
0.010

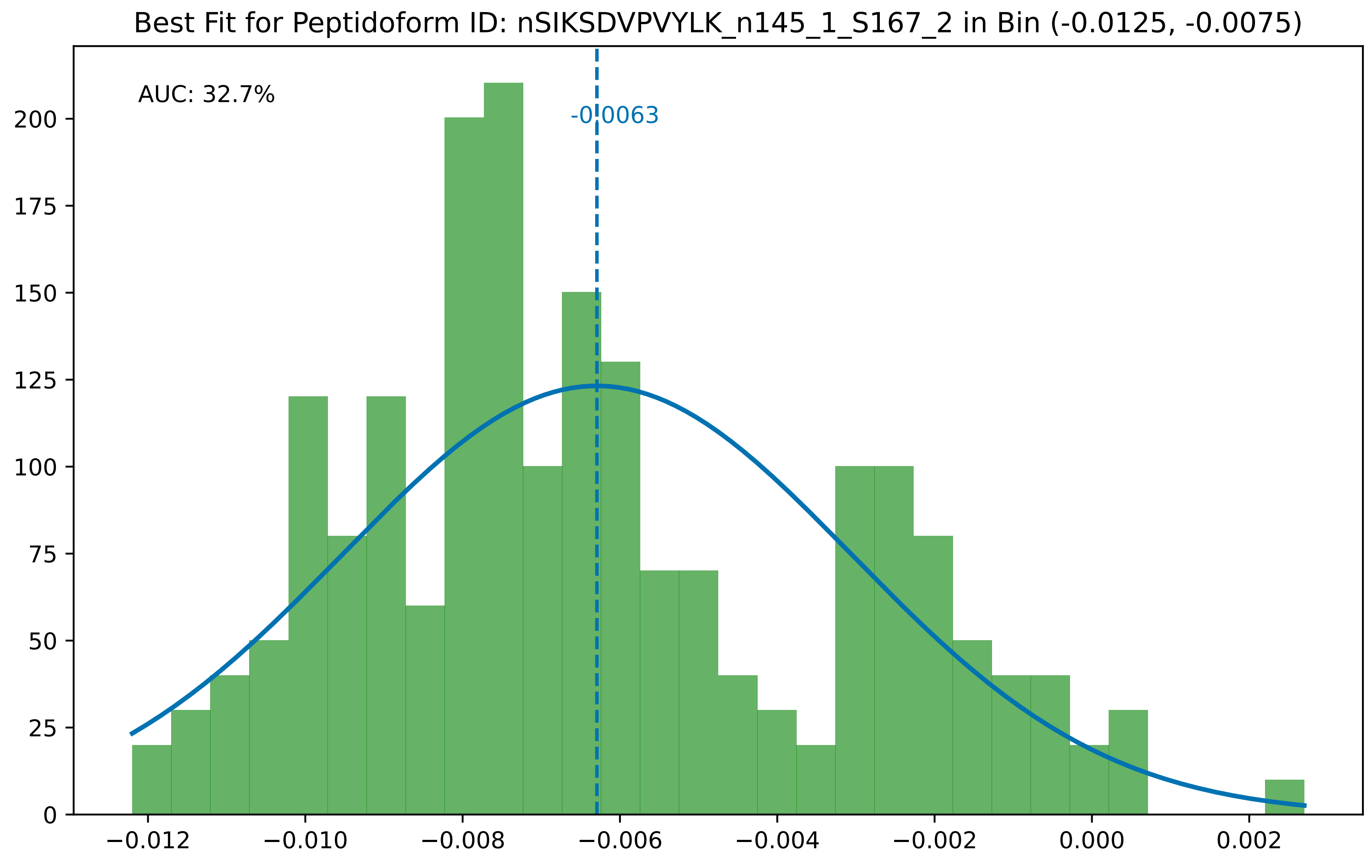

Best Fit for Peptidoform ID: nSIKSDVPVYLK_n145_1_S167_2 in Bin (-0.0125, -0.0075)
AUC: 32.7%
-0.0063
0
25
50
75
100
125
150
175
200
−0.012
−0.010
−0.008
−0.006
−0.004
−0.002
0.000
0.002

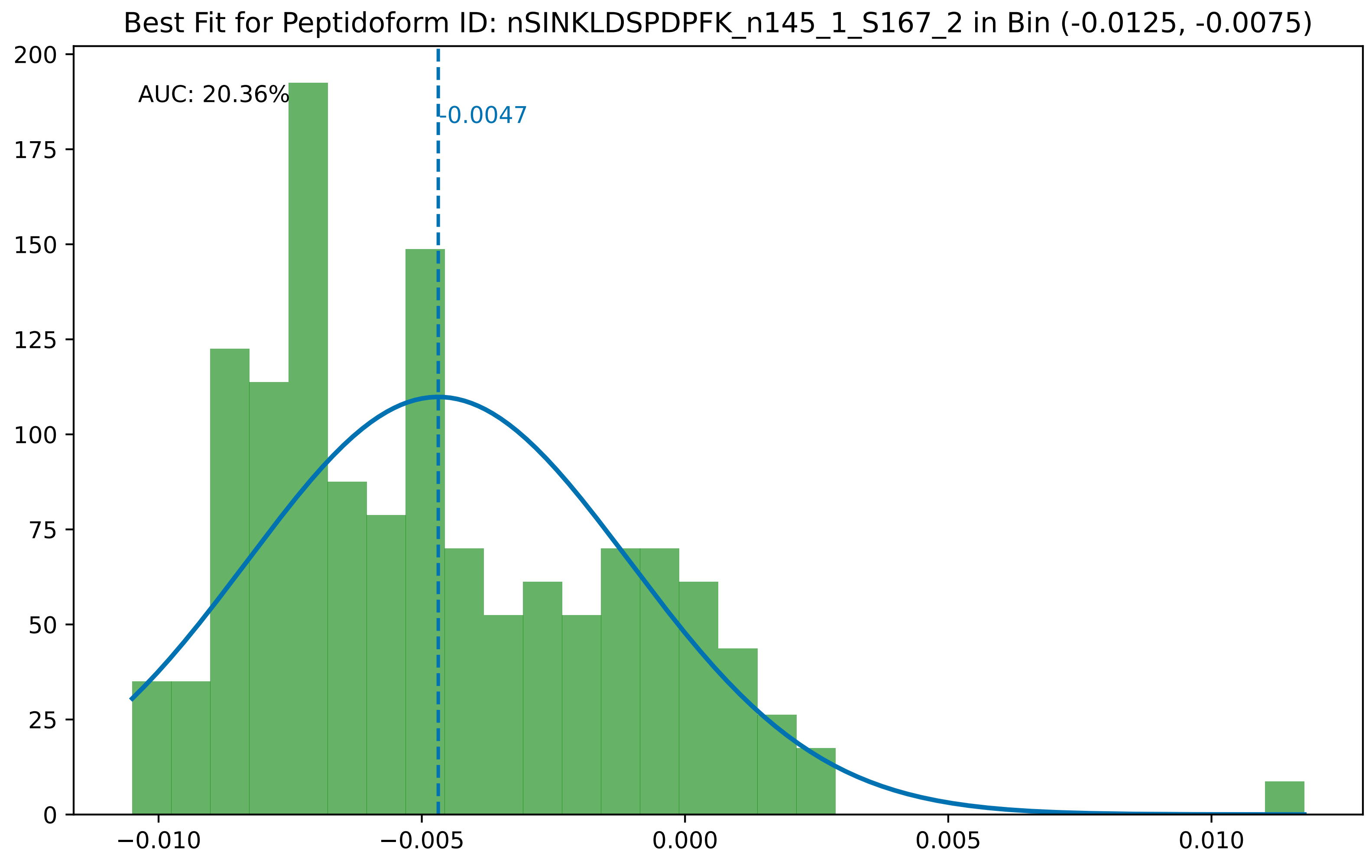

Best Fit for Peptidoform ID: nSINKLDSPDPFK_n145_1_S167_2 in Bin (-0.0125, -0.0075)
AUC: 20.36%
-0.0047
0
25
50
75
100
125
150
175
200
−0.010
−0.005
0.000
0.005
0.010

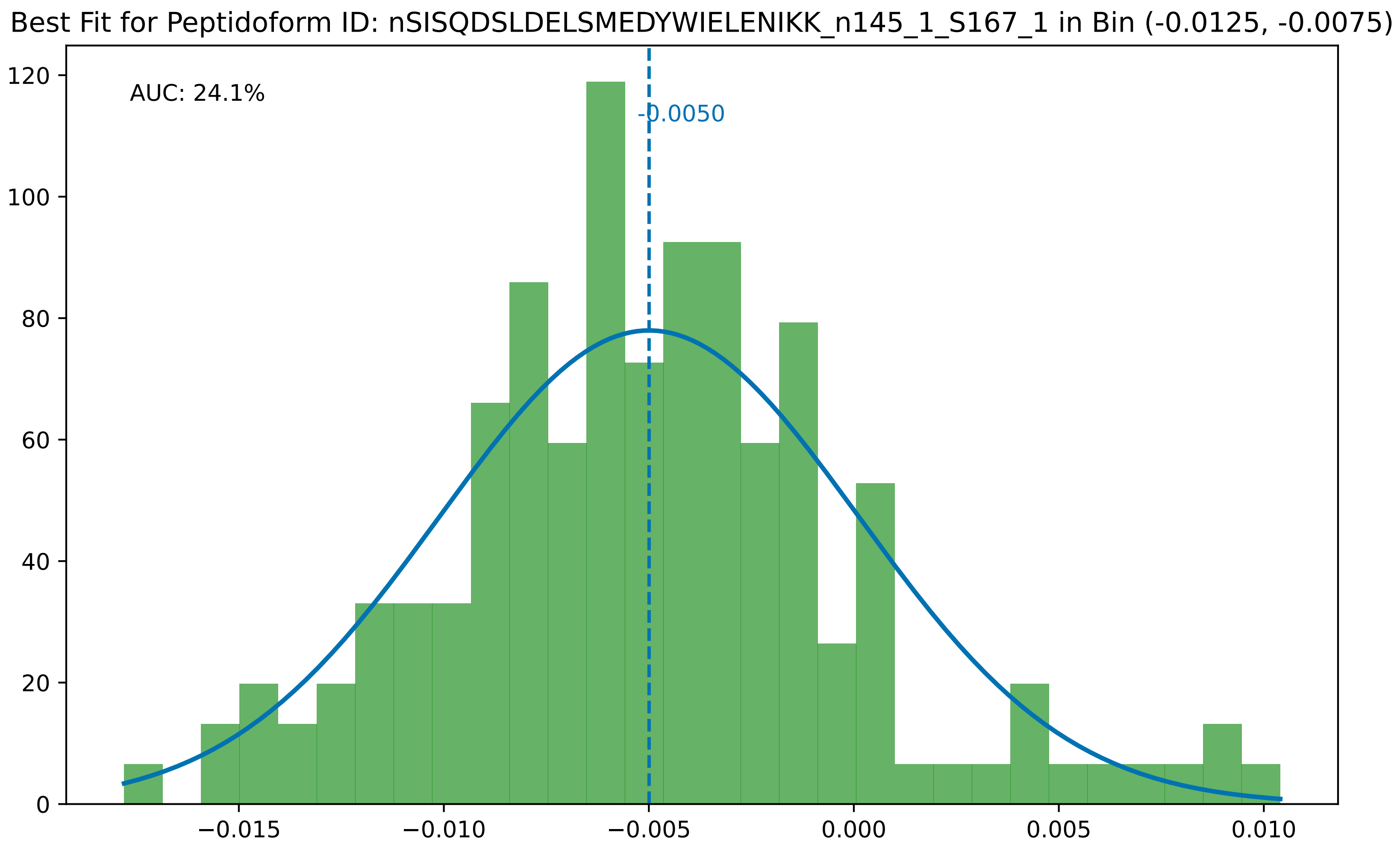
Best Fit for Peptidoform ID: nSISQDSLDELSMEDYWIELENIKK_n145_1_S167_1 in Bin (-0.0125, -0.0075)
AUC: 24.1%
-0.0050
120
100
80
60
40
20
0
−0.015
−0.010
−0.005
0.000
0.005
0.010

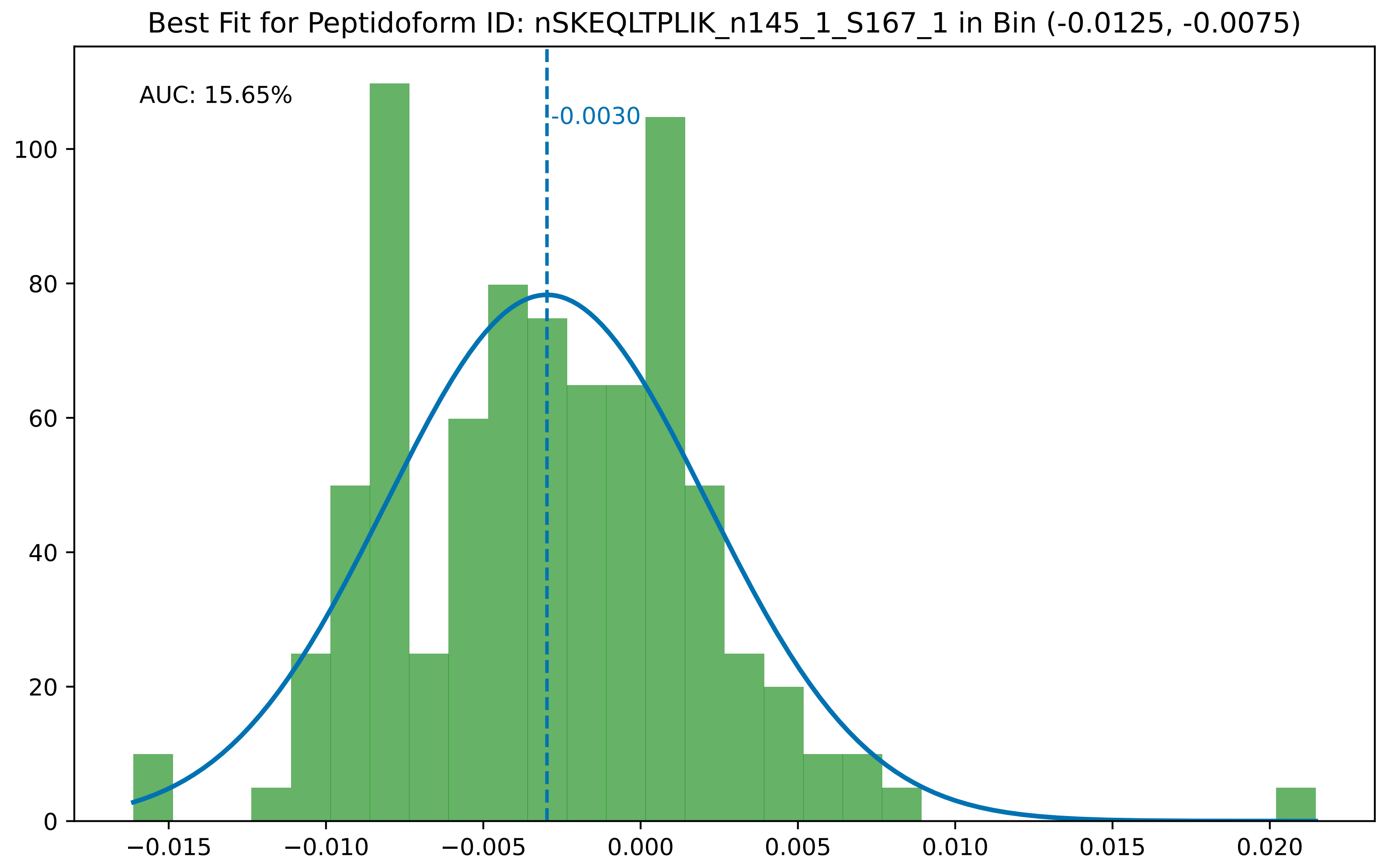

Best Fit for Peptidoform ID: nSKEQLTPLIK_n145_1_S167_1 in Bin (-0.0125, -0.0075)
AUC: 15.65%
-0.0030
0
20
40
60
80
100
−0.015
−0.010
−0.005
0.000
0.005
0.010
0.015
0.020

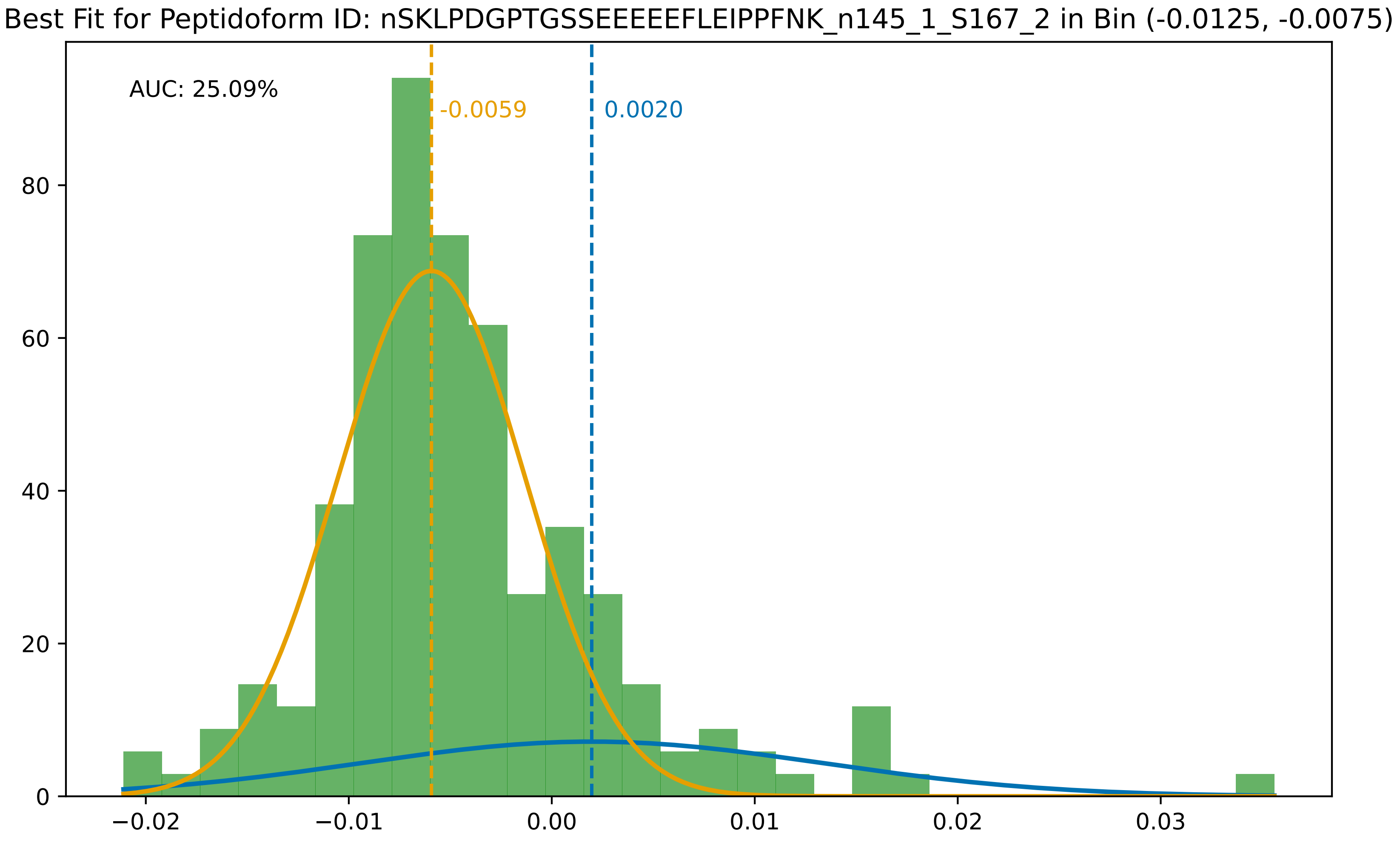

Best Fit for Peptidoform ID: nSKLPDGPTGSSEEEEEFLEIPPFNK_n145_1_S167_2 in Bin (-0.0125, -0.0075)
AUC: 25.09%
-0.0059
0.0020
80
60
40
20
0
−0.02
−0.01
0.00
0.01
0.02
0.03

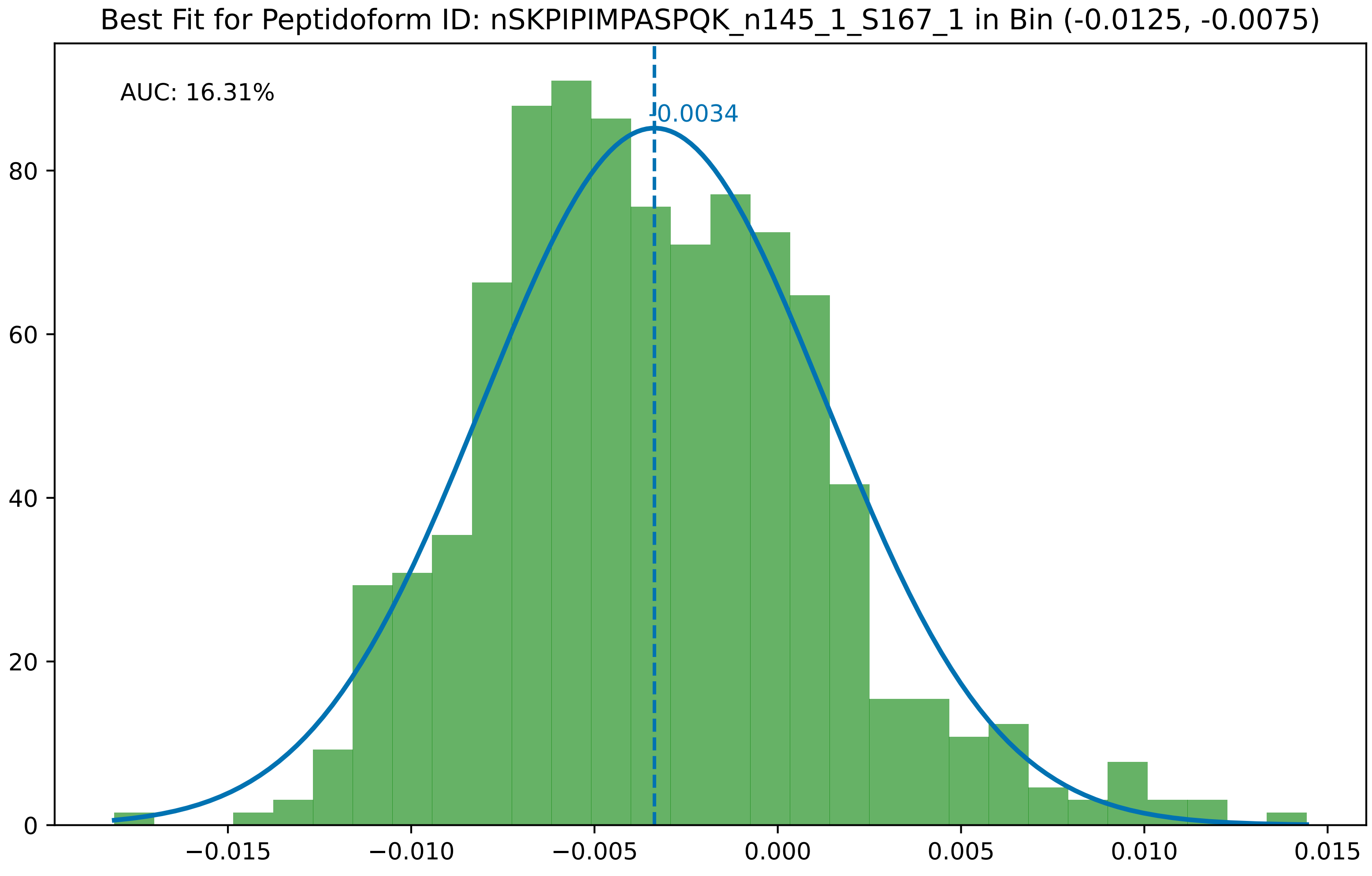
Best Fit for Peptidoform ID: nSKPIPIMPASPQK_n145_1_S167_1 in Bin (-0.0125, -0.0075)
AUC: 16.31%
0.0034
0
20
40
60
80
−0.015
−0.010
−0.005
0.000
0.005
0.010
0.015

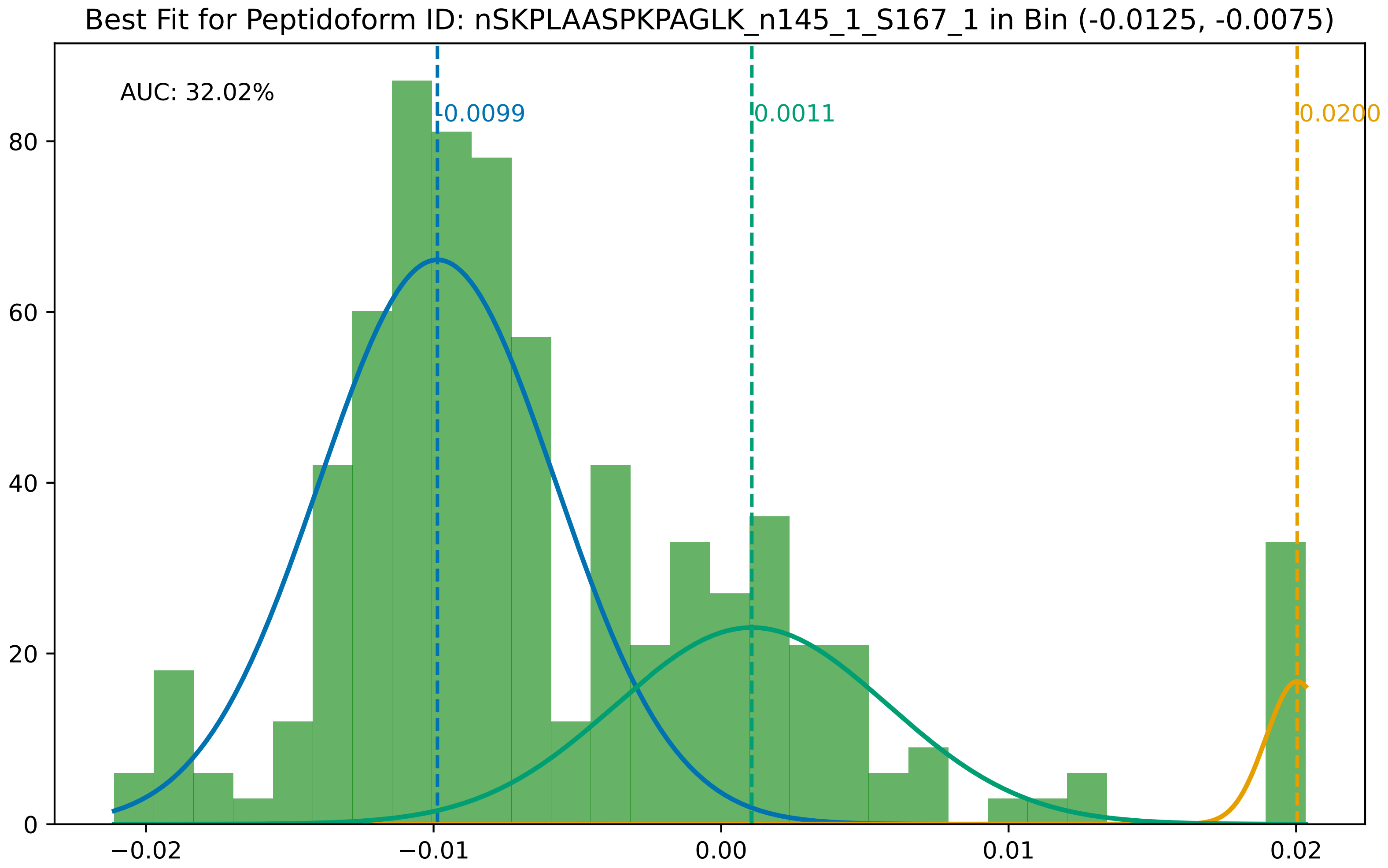
Best Fit for Peptidoform ID: nSKPLAASPKPAGLK_n145_1_S167_1 in Bin (-0.0125, -0.0075)
AUC: 32.02%
0.0099
0.0011
0.0200
0
20
40
60
80
−0.02
−0.01
0.00
0.01
0.02

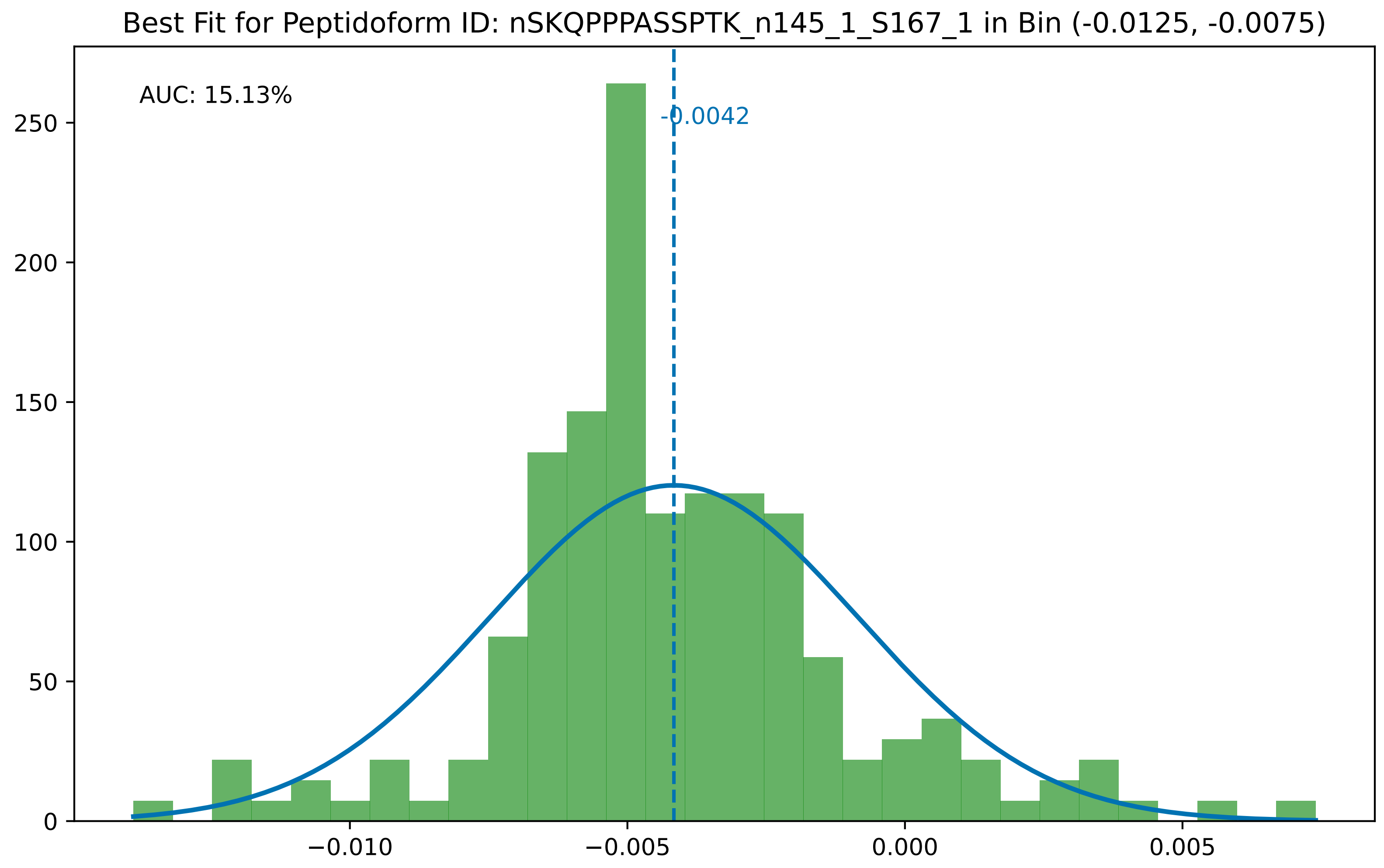

Best Fit for Peptidoform ID: nSKQPPPASSPTK_n145_1_S167_1 in Bin (-0.0125, -0.0075)
AUC: 15.13%
-0.0042
0
50
100
150
200
250
−0.010
−0.005
0.000
0.005

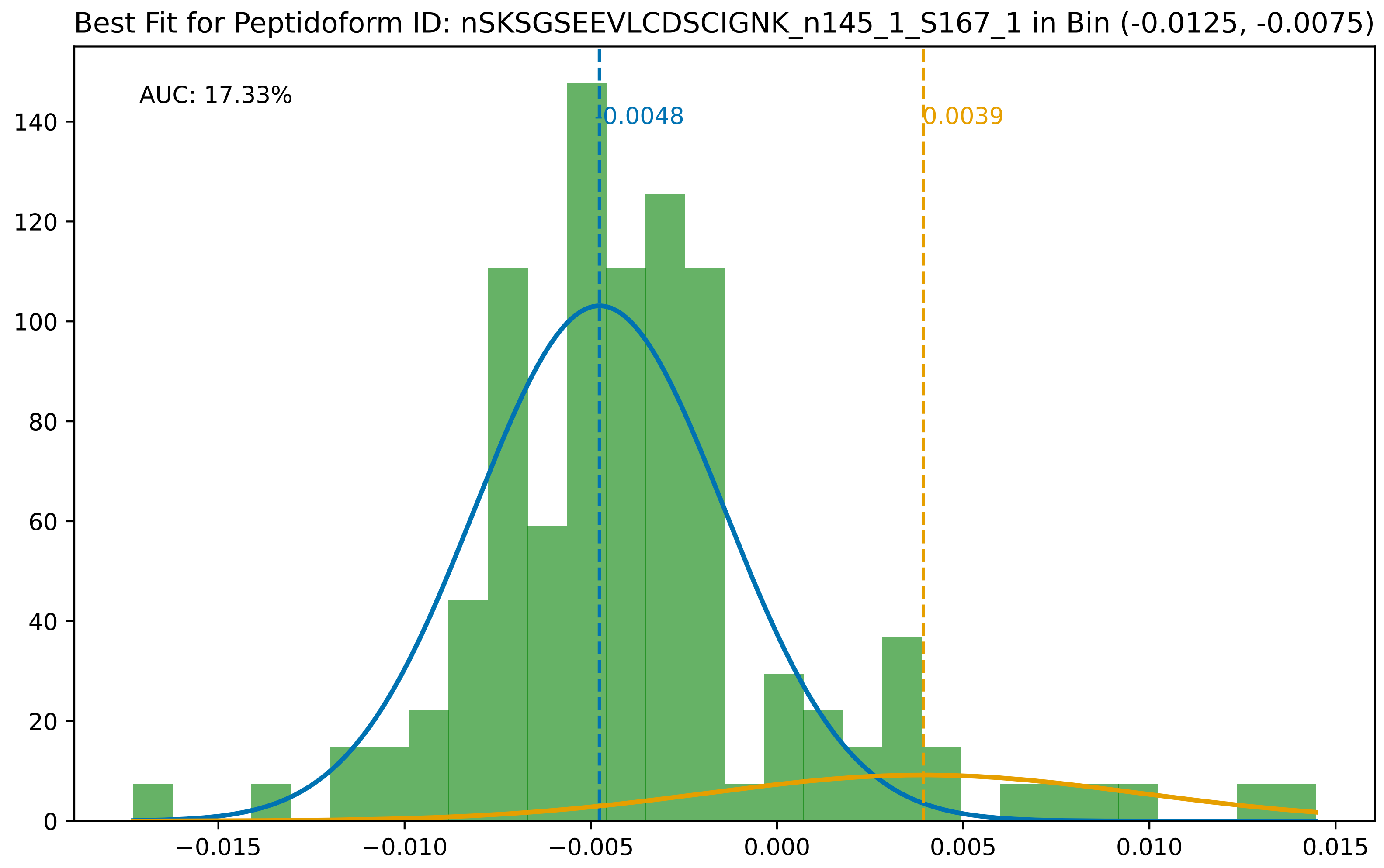
Best Fit for Peptidoform ID: nSKSGSEEVLCDSCIGNK_n145_1_S167_1 in Bin (-0.0125, -0.0075)
AUC: 17.33%
-0.0048
0.0039
0
20
40
60
80
100
120
140
−0.015
−0.010
−0.005
0.000
0.005
0.010
0.015

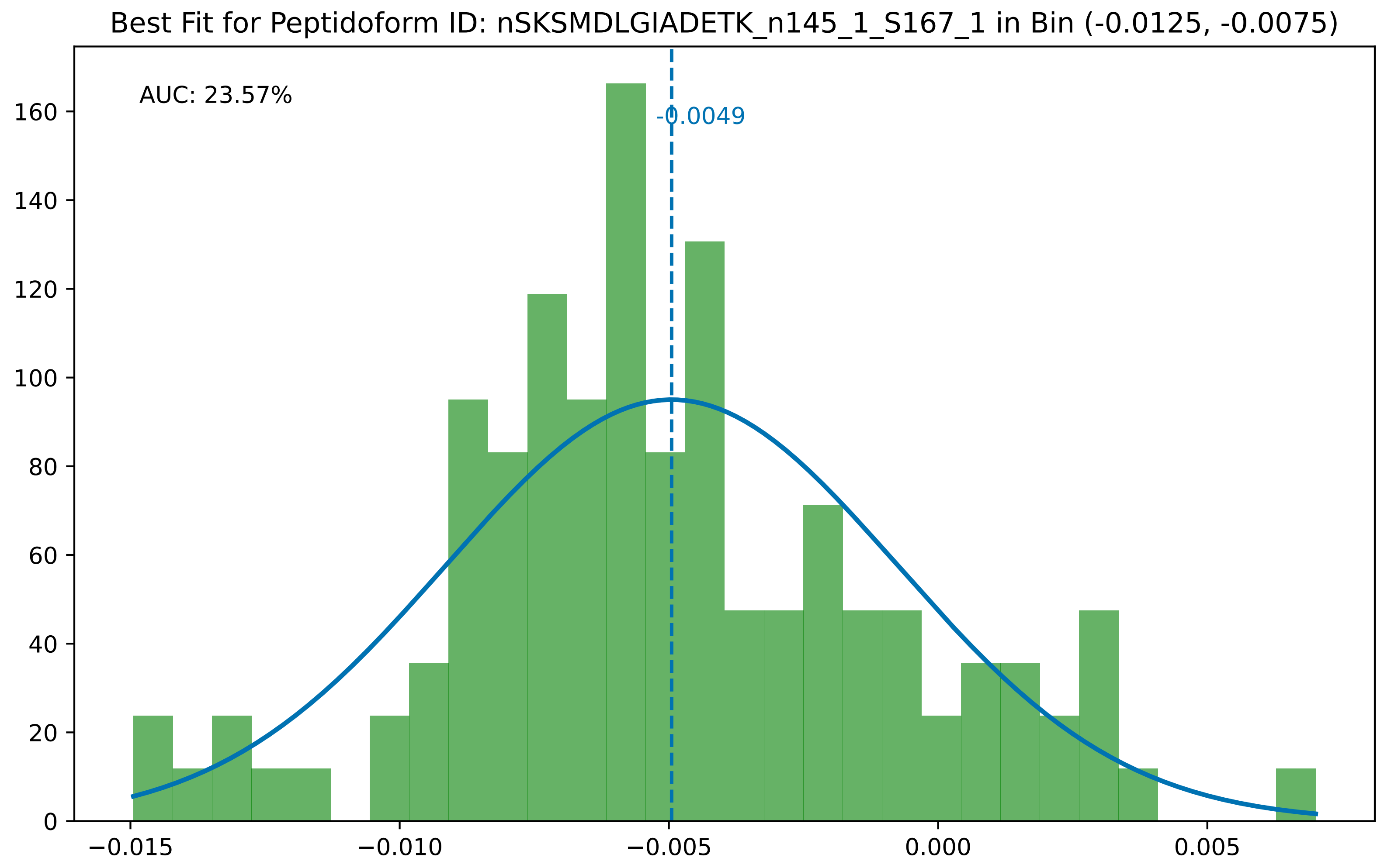

Best Fit for Peptidoform ID: nSKSMDLGIADETK_n145_1_S167_1 in Bin (-0.0125, -0.0075)
AUC: 23.57%
-0.0049
0
20
40
60
80
100
120
140
160
−0.015
−0.010
−0.005
0.000
0.005

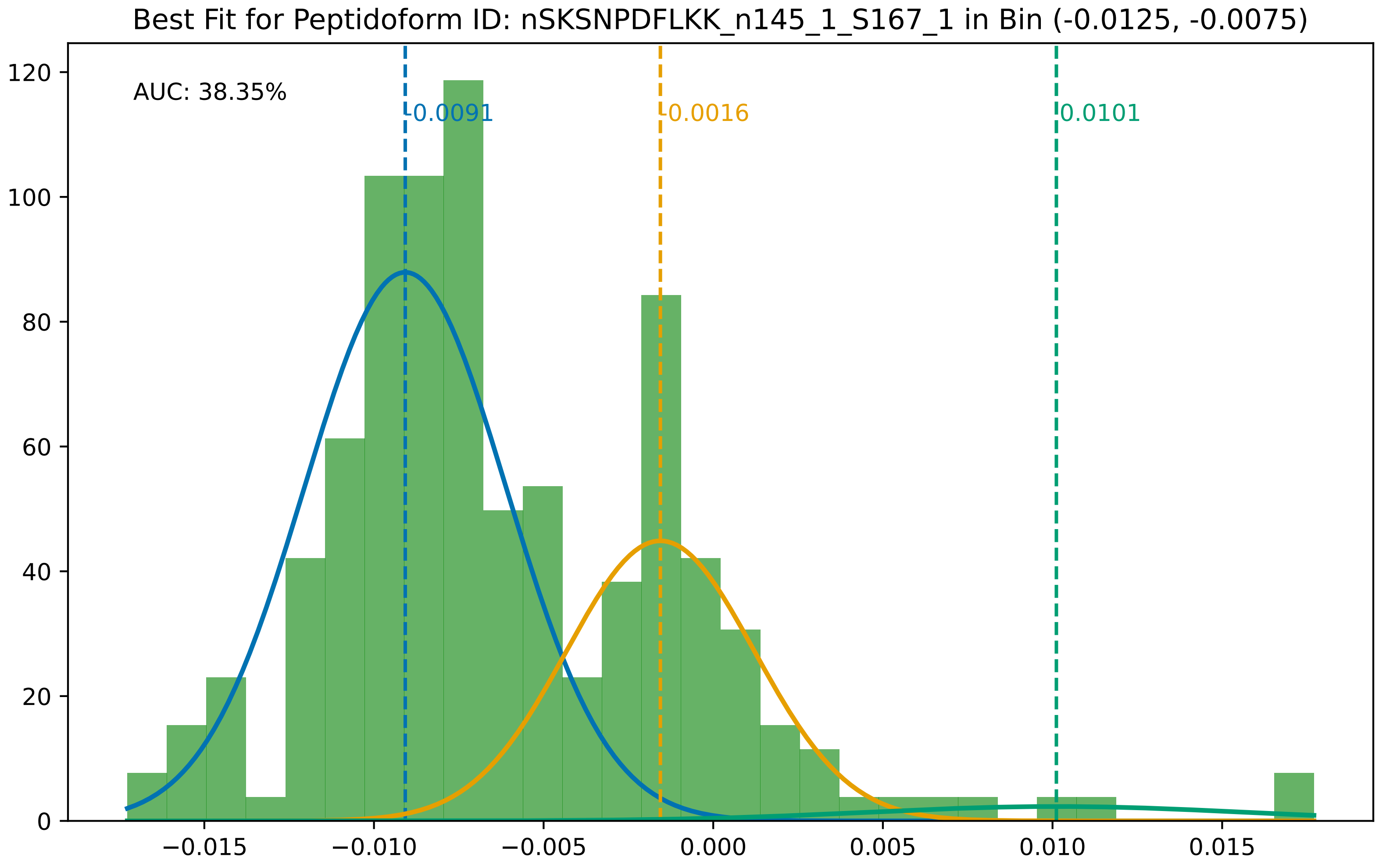

Best Fit for Peptidoform ID: nSKSNPDFLKK_n145_1_S167_1 in Bin (-0.0125, -0.0075)
AUC: 38.35%
-0.0091
-0.0016
0.0101
0
20
40
60
80
100
120
−0.015
−0.010
−0.005
0.000
0.005
0.010
0.015

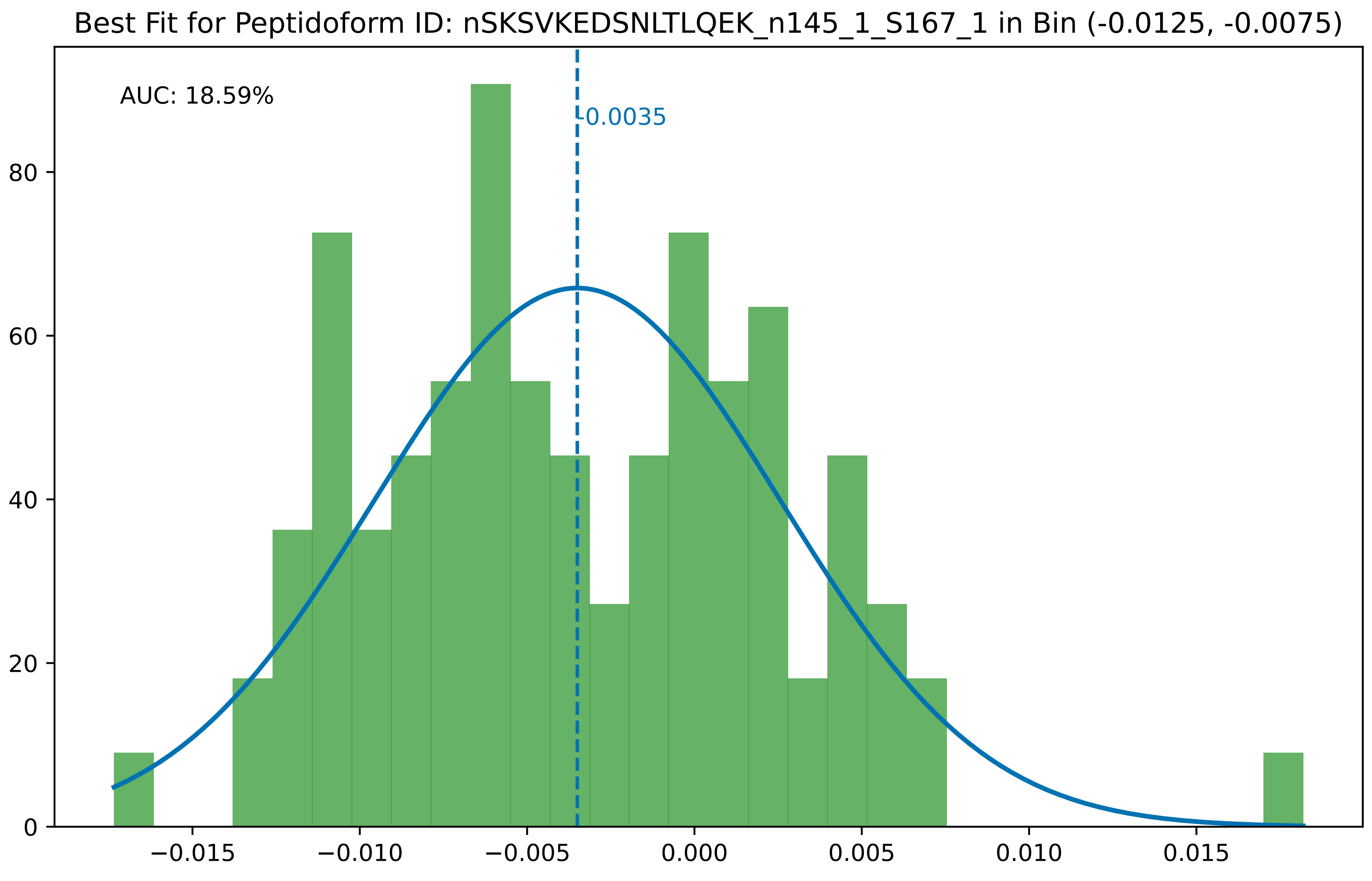
Best Fit for Peptidoform ID: nSKSVKEDSNLTLQEK_n145_1_S167_1 in Bin (-0.0125, -0.0075)
AUC: 18.59%
-0.0035
0
20
40
60
80
−0.015
−0.010
−0.005
0.000
0.005
0.010
0.015

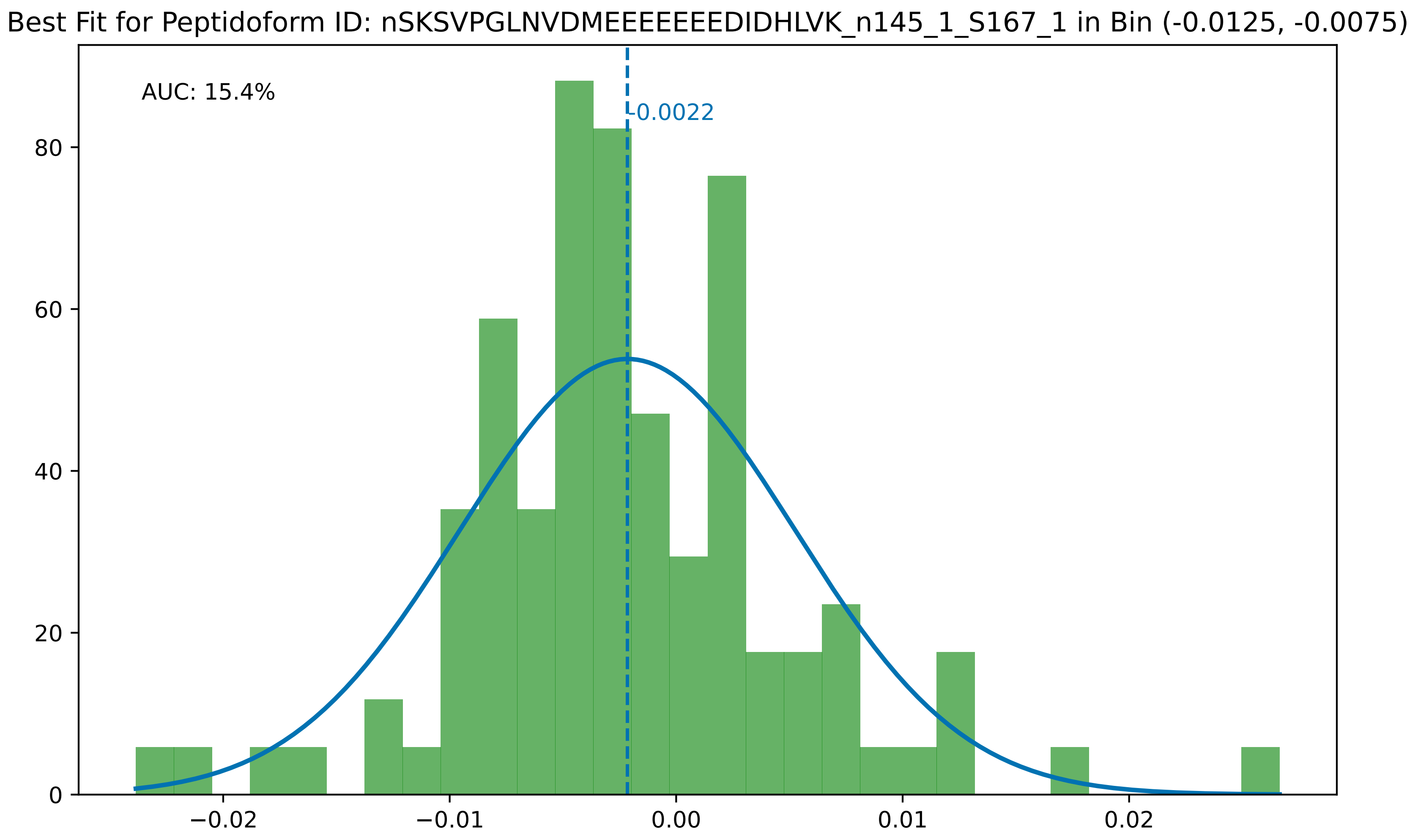

Best Fit for Peptidoform ID: nSKSVPGLNVDMEEEEEEEDIDHLVK_n145_1_S167_1 in Bin (-0.0125, -0.0075)
AUC: 15.4%
-0.0022
0
20
40
60
80
−0.02
−0.01
0.00
0.01
0.02

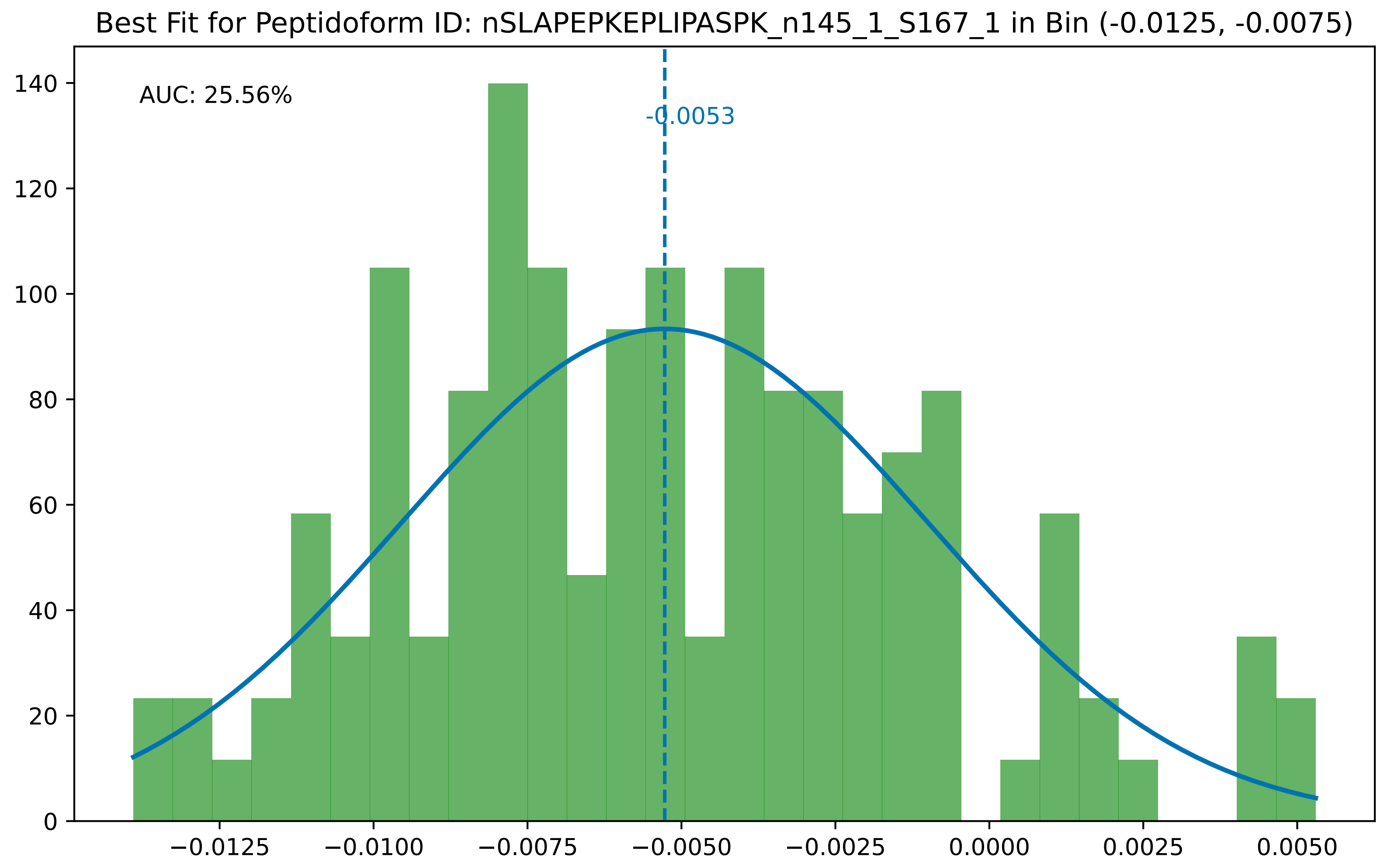

Best Fit for Peptidoform ID: nSLAPEPKEPLIPASPK_n145_1_S167_1 in Bin (-0.0125, -0.0075)
AUC: 25.56%
-0.0053
0
20
40
60
80
100
120
140
−0.0125
−0.0100
−0.0075
−0.0050
−0.0025
0.0000
0.0025
0.0050

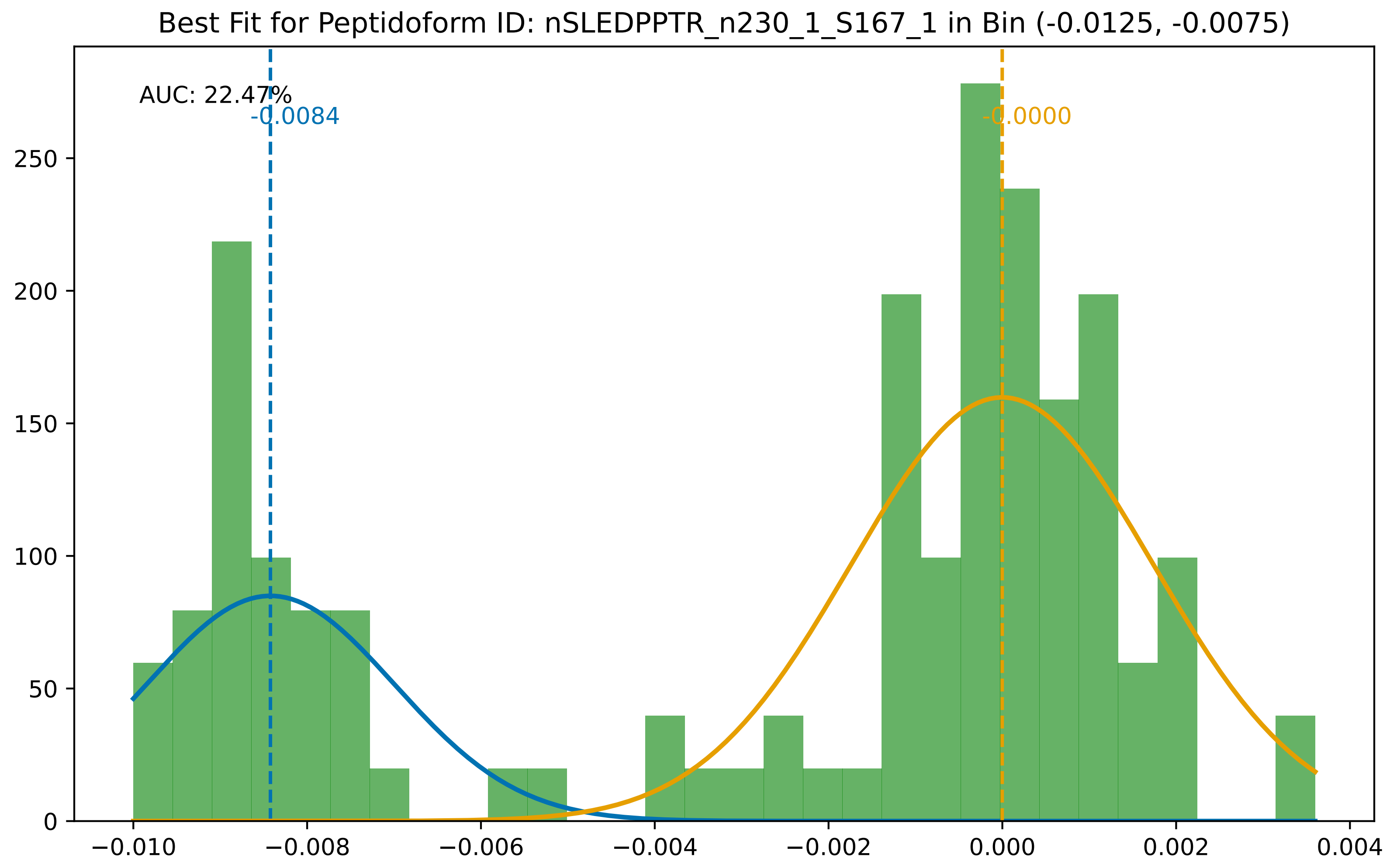
Best Fit for Peptidoform ID: nSLEDPPTR_n230_1_S167_1 in Bin (-0.0125, -0.0075)
AUC: 22.47%
-0.0084
-0.0000
250
200
150
100
50
0
−0.010
−0.008
−0.006
−0.004
−0.002
0.000
0.002
0.004

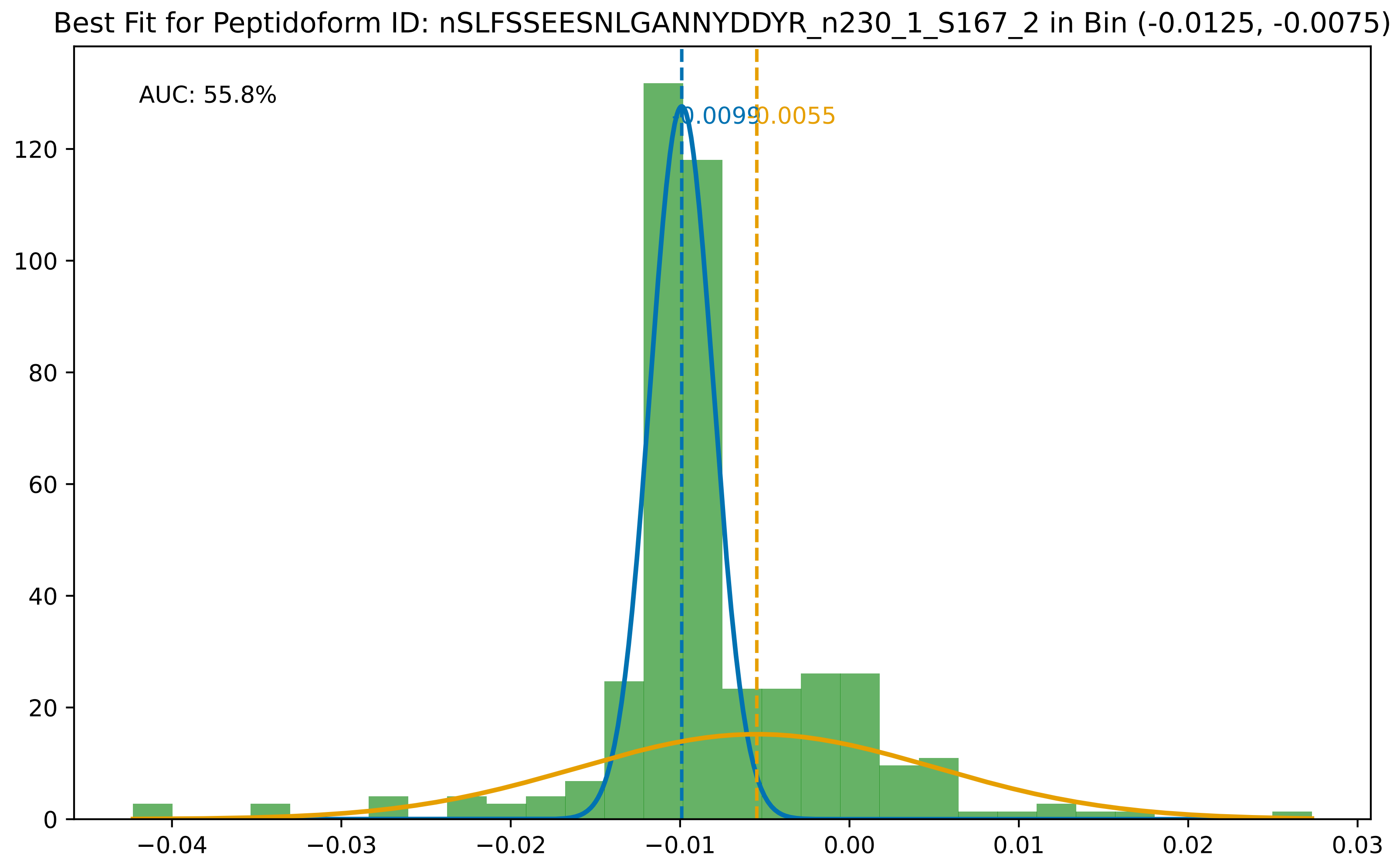

Best Fit for Peptidoform ID: nSLFSSEESNLGANNYDDYR_n230_1_S167_2 in Bin (-0.0125, -0.0075)
AUC: 55.8%
-0.0099
-0.0055
0
20
40
60
80
100
120
−0.04
−0.03
−0.02
−0.01
0.00
0.01
0.02
0.03

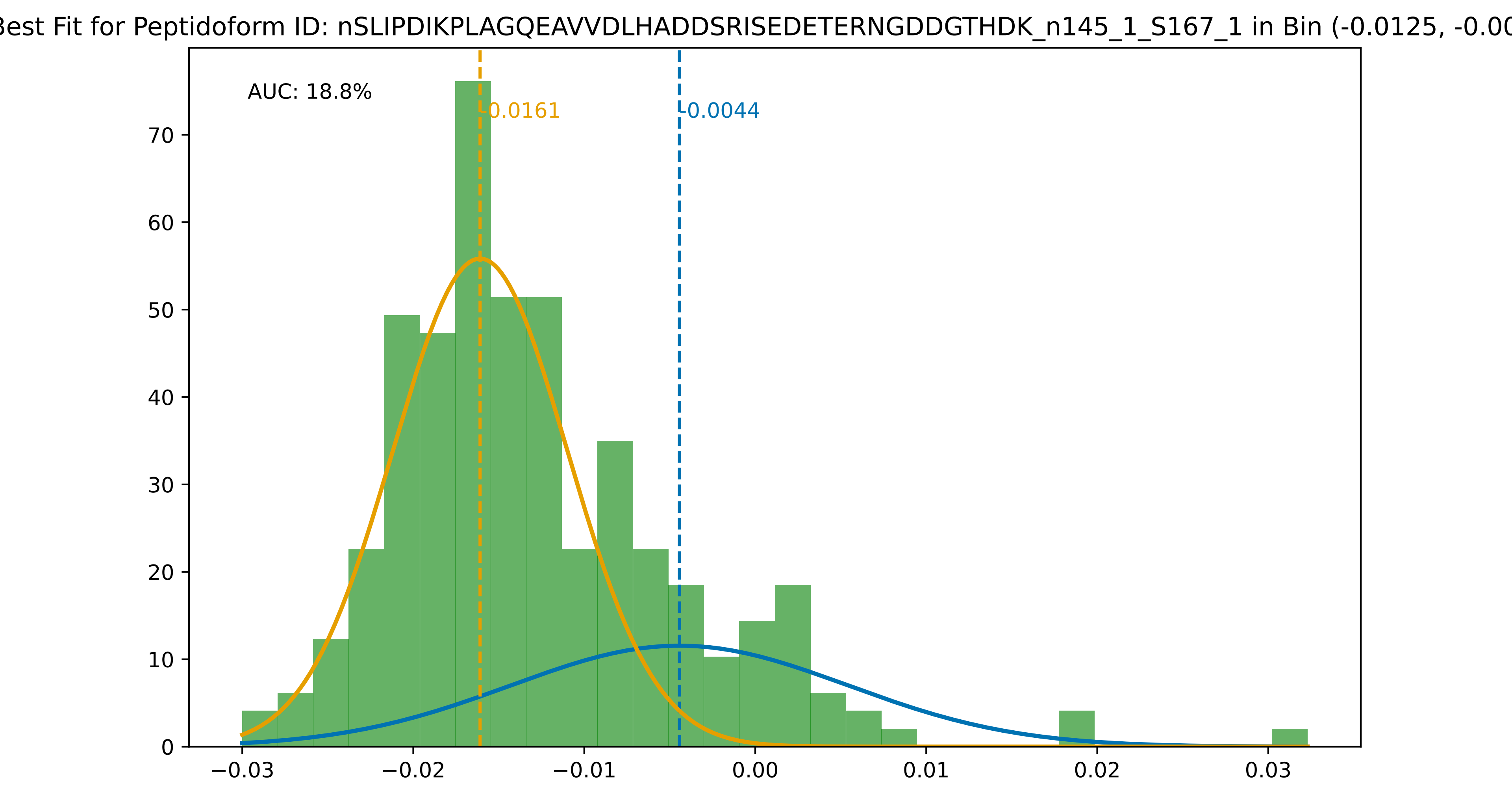
Best Fit for Peptidoform ID: nSLIPDIKPLAGQEAVVDLHADDSRISEDETERNGDDGTHDK_n145_1_S167_1 in Bin (-0.0125, -0.00
AUC: 18.8%
-0.0161
-0.0044
0
10
20
30
40
50
60
70
−0.03
−0.02
−0.01
0.00
0.01
0.02
0.03

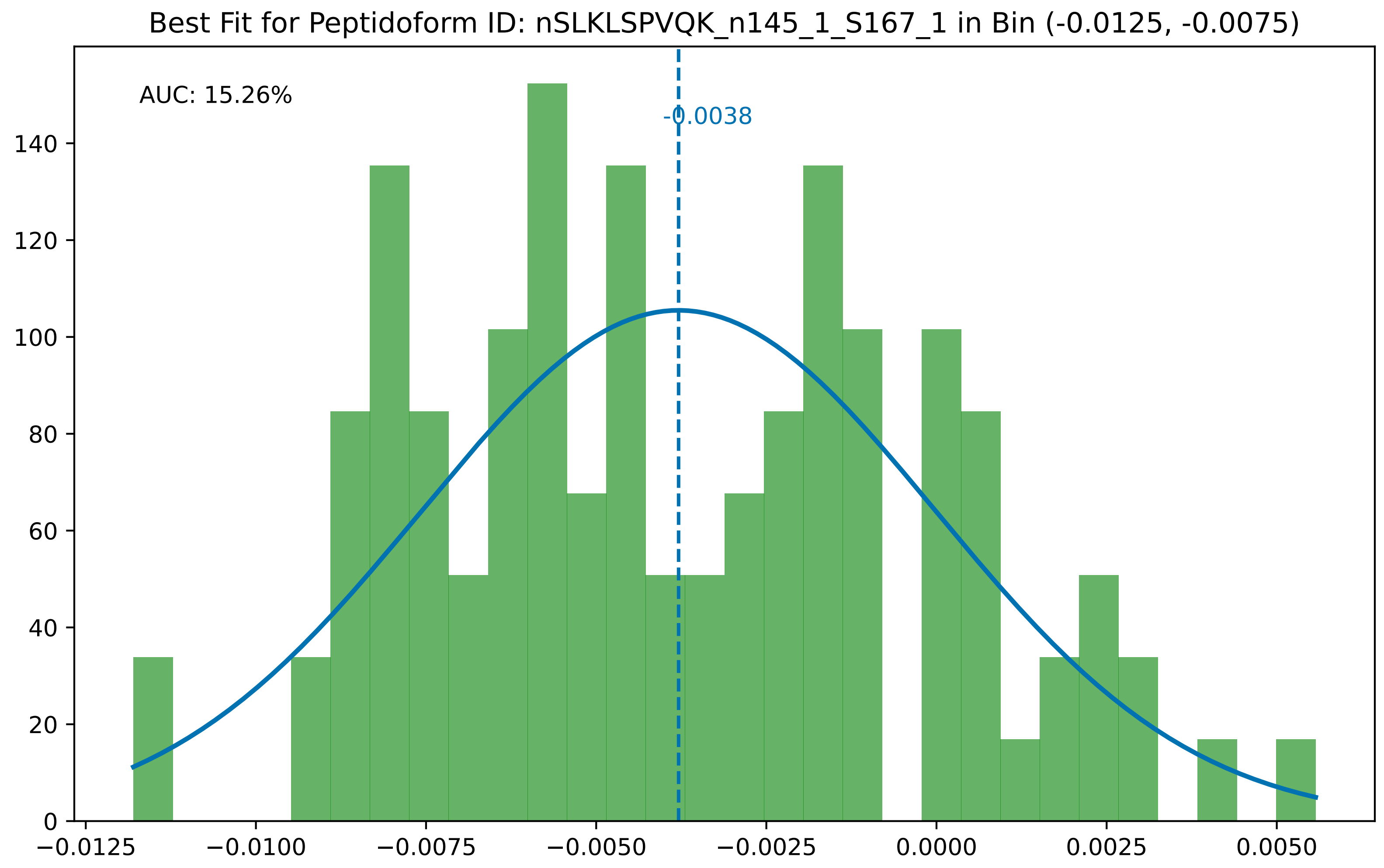
Best Fit for Peptidoform ID: nSLKLSPVQK_n145_1_S167_1 in Bin (-0.0125, -0.0075)
AUC: 15.26%
-0.0038
0
20
40
60
80
100
120
140
−0.0125
−0.0100
−0.0075
−0.0050
−0.0025
0.0000
0.0025
0.0050

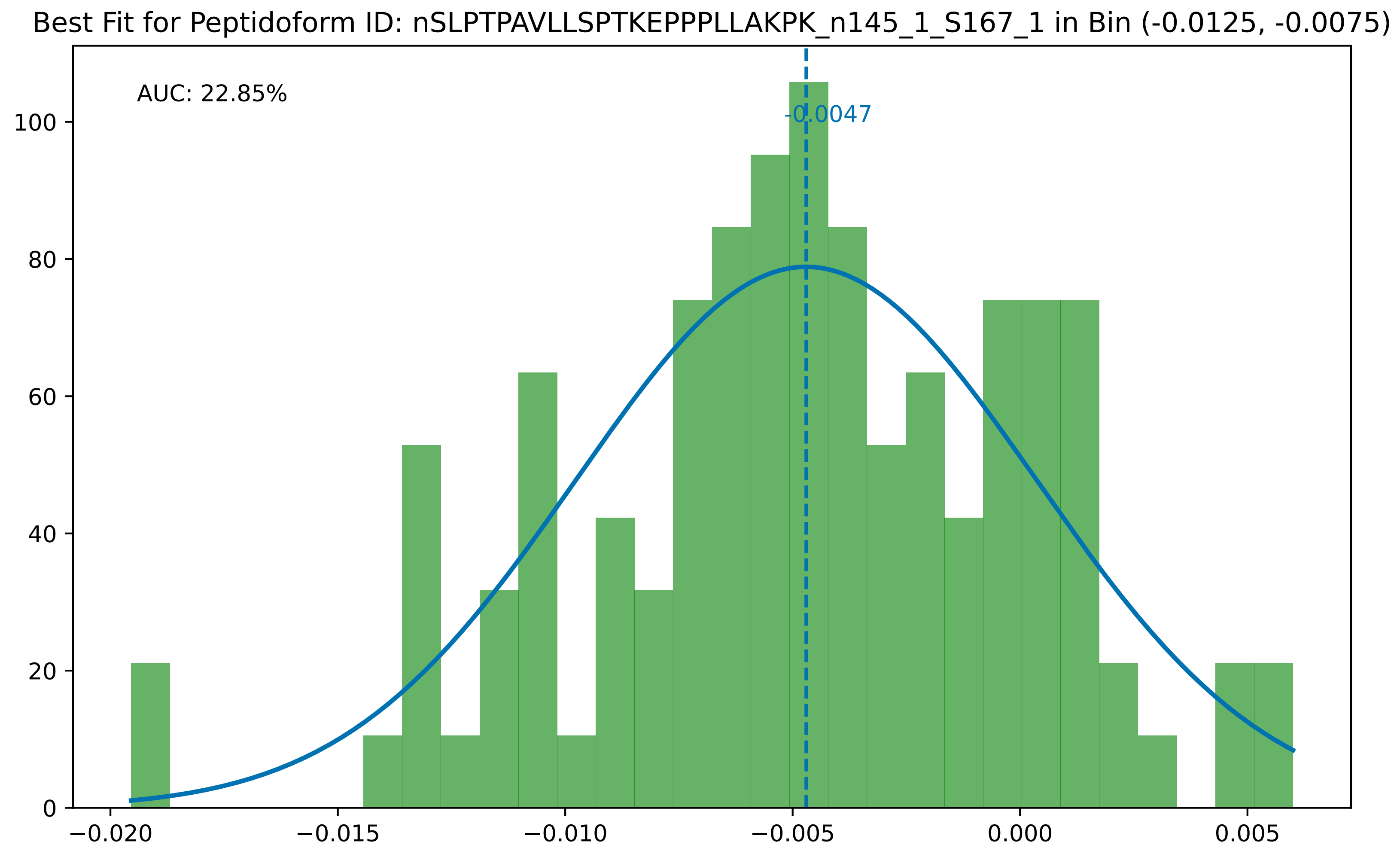

Best Fit for Peptidoform ID: nSLPTPAVLLSPTKEPPPLLAKPK_n145_1_S167_1 in Bin (-0.0125, -0.0075)
AUC: 22.85%
-0.0047
0
20
40
60
80
100
−0.020
−0.015
−0.010
−0.005
0.000
0.005

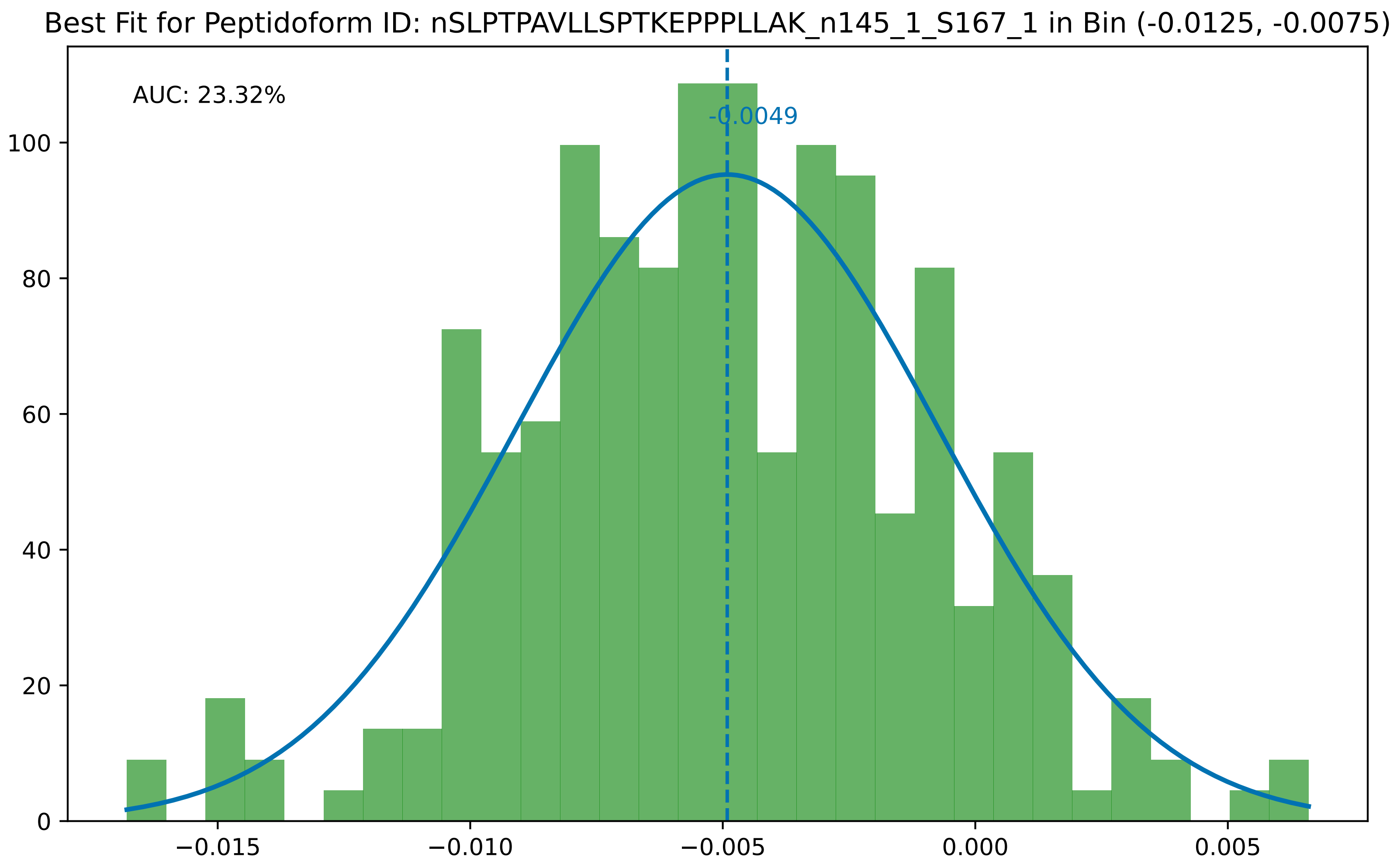

Best Fit for Peptidoform ID: nSLPTPAVLLSPTKEPPPLLAK_n145_1_S167_1 in Bin (-0.0125, -0.0075)
AUC: 23.32%
-0.0049
0
20
40
60
80
100
−0.015
−0.010
−0.005
0.000
0.005

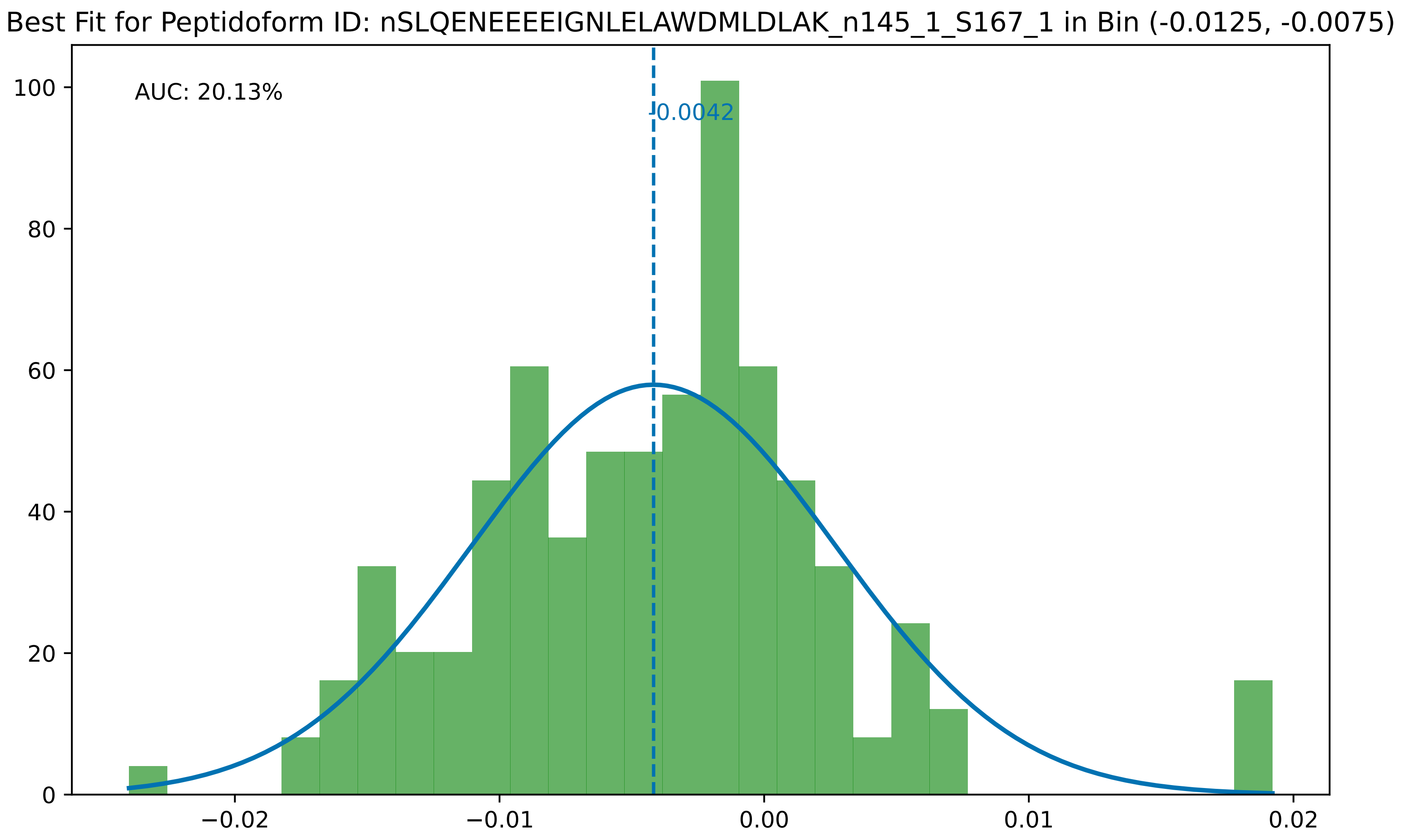

Best Fit for Peptidoform ID: nSLQENEEEEIGNLELAWDMLDLAK_n145_1_S167_1 in Bin (-0.0125, -0.0075)
AUC: 20.13%
-0.0042
100
80
60
40
20
0
−0.02
−0.01
0.00
0.01
0.02

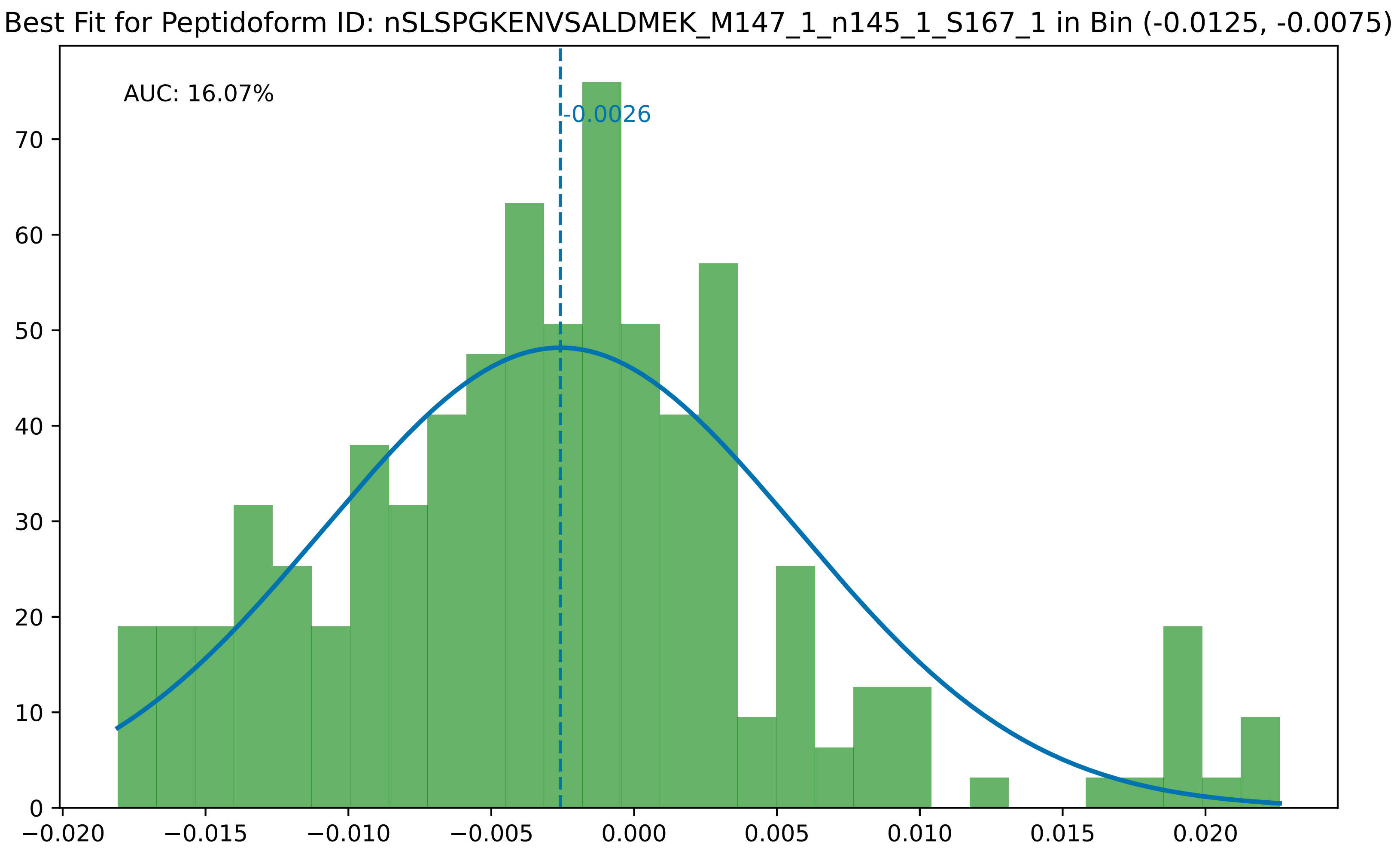

Best Fit for Peptidoform ID: nSLSPGKENVSALDMEK_M147_1_n145_1_S167_1 in Bin (-0.0125, -0.0075)
AUC: 16.07%
-0.0026
0
10
20
30
40
50
60
70
−0.020
−0.015
−0.010
−0.005
0.000
0.005
0.010
0.015
0.020

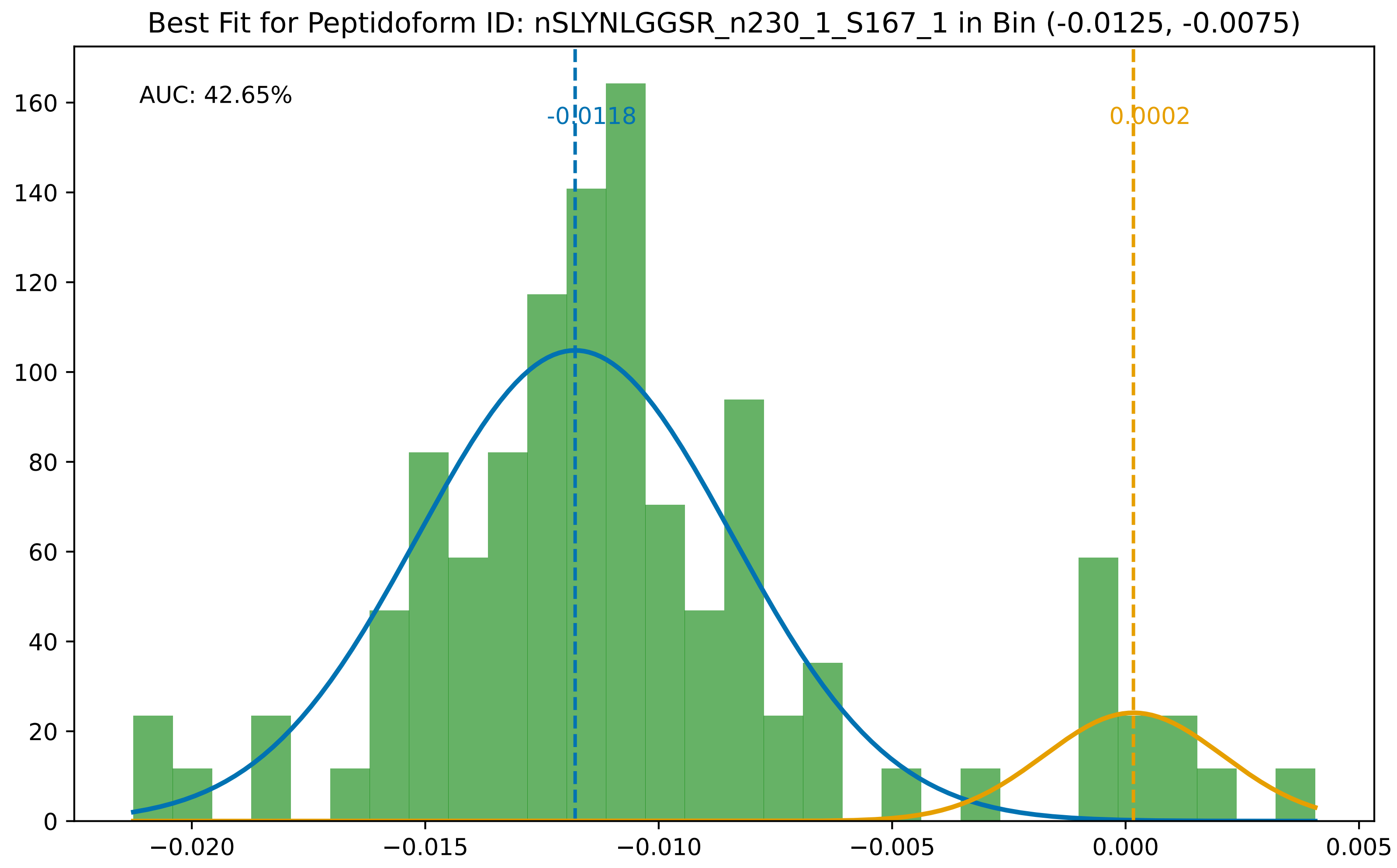

Best Fit for Peptidoform ID: nSLYNLGGSR_n230_1_S167_1 in Bin (-0.0125, -0.0075)
AUC: 42.65%
-0.0118
0.0002
0
20
40
60
80
100
120
140
160
−0.020
−0.015
−0.010
−0.005
0.000
0.005

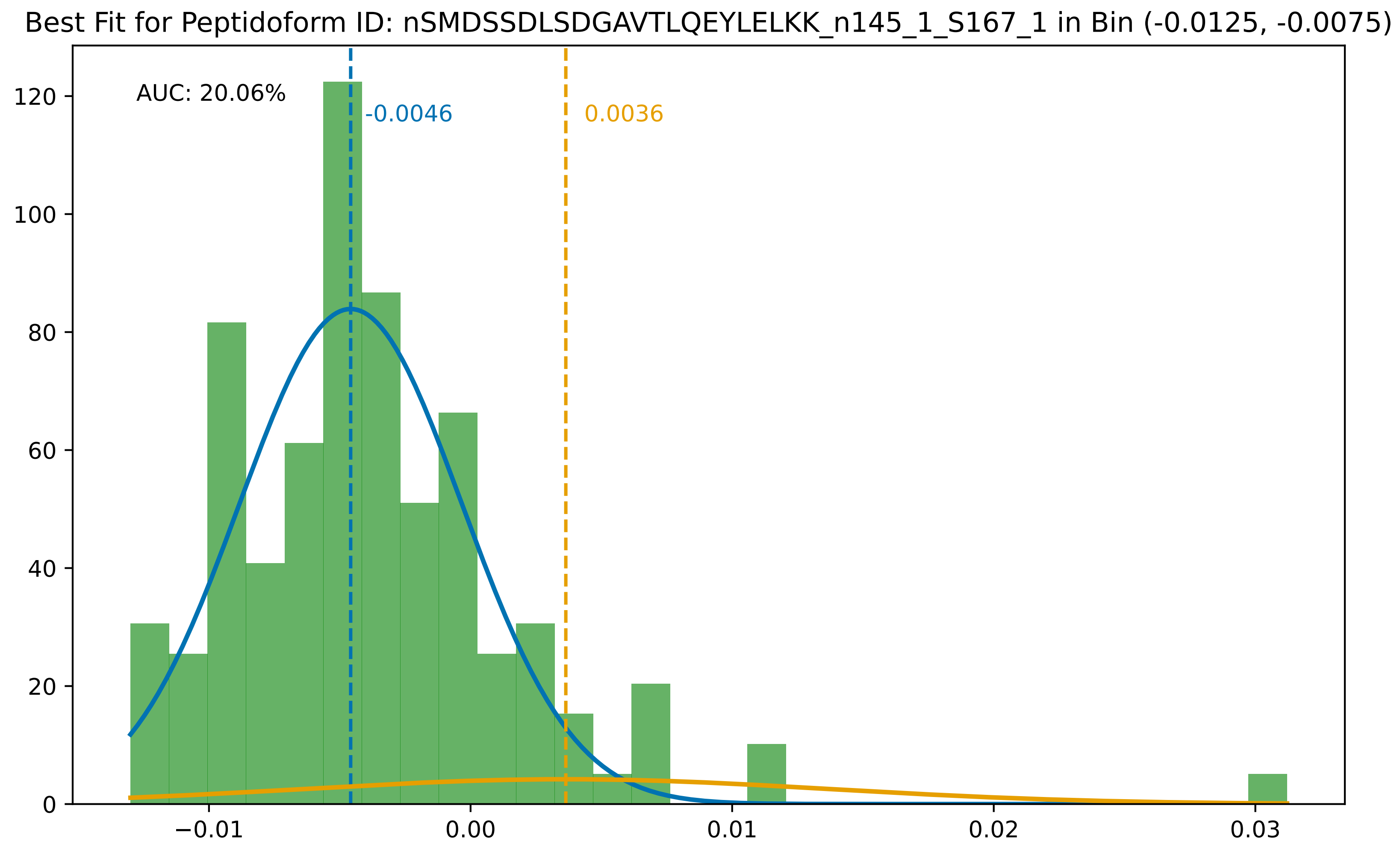

Best Fit for Peptidoform ID: nSMDSSDLSDGAVTLQEYLELKK_n145_1_S167_1 in Bin (-0.0125, -0.0075)
AUC: 20.06%
-0.0046
0.0036
0
20
40
60
80
100
120
−0.01
0.00
0.01
0.02
0.03

Best Fit for Peptidoform ID: nSMSVDLSHIPLKDPLLFK_n145_1_S167_1 in Bin (-0.0125, -0.0075)

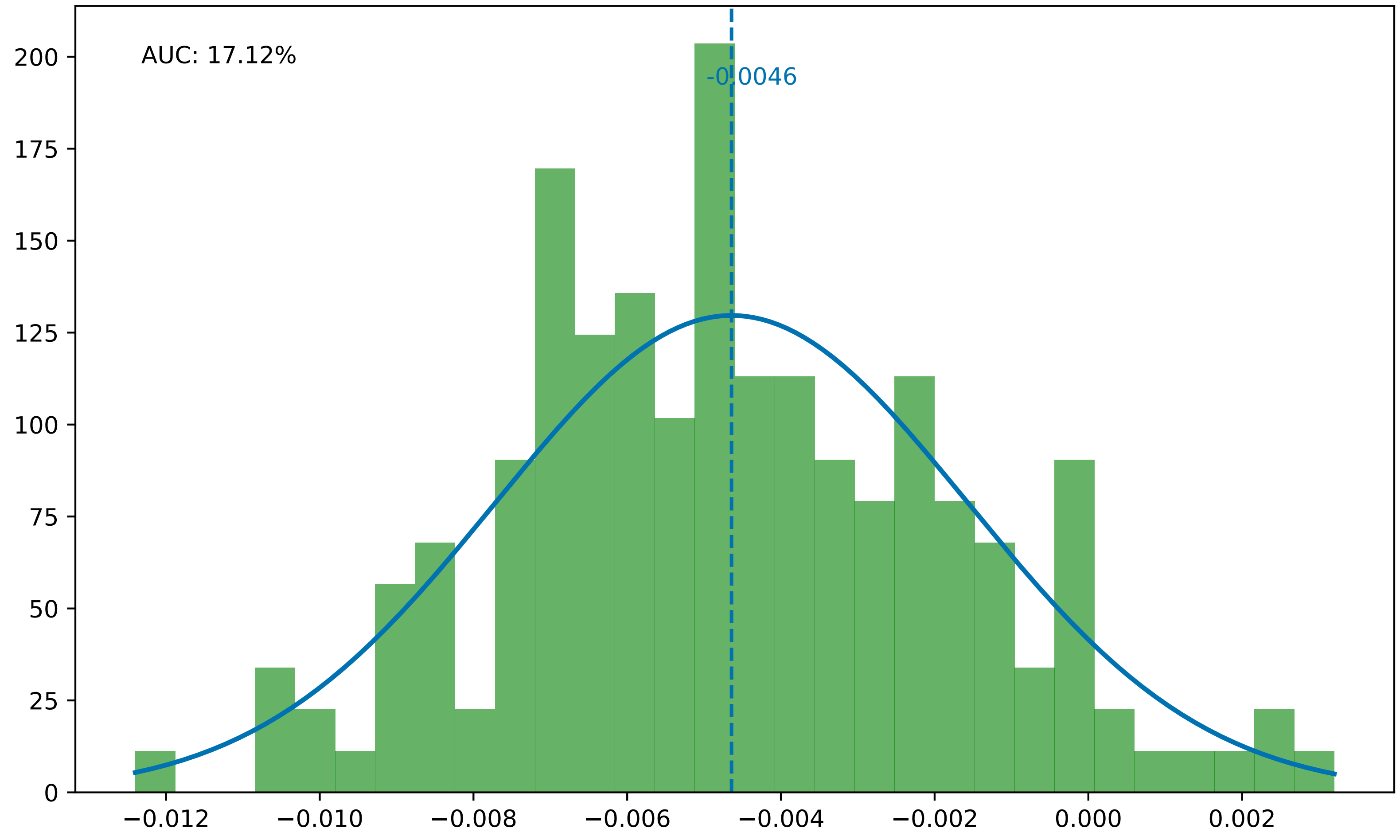

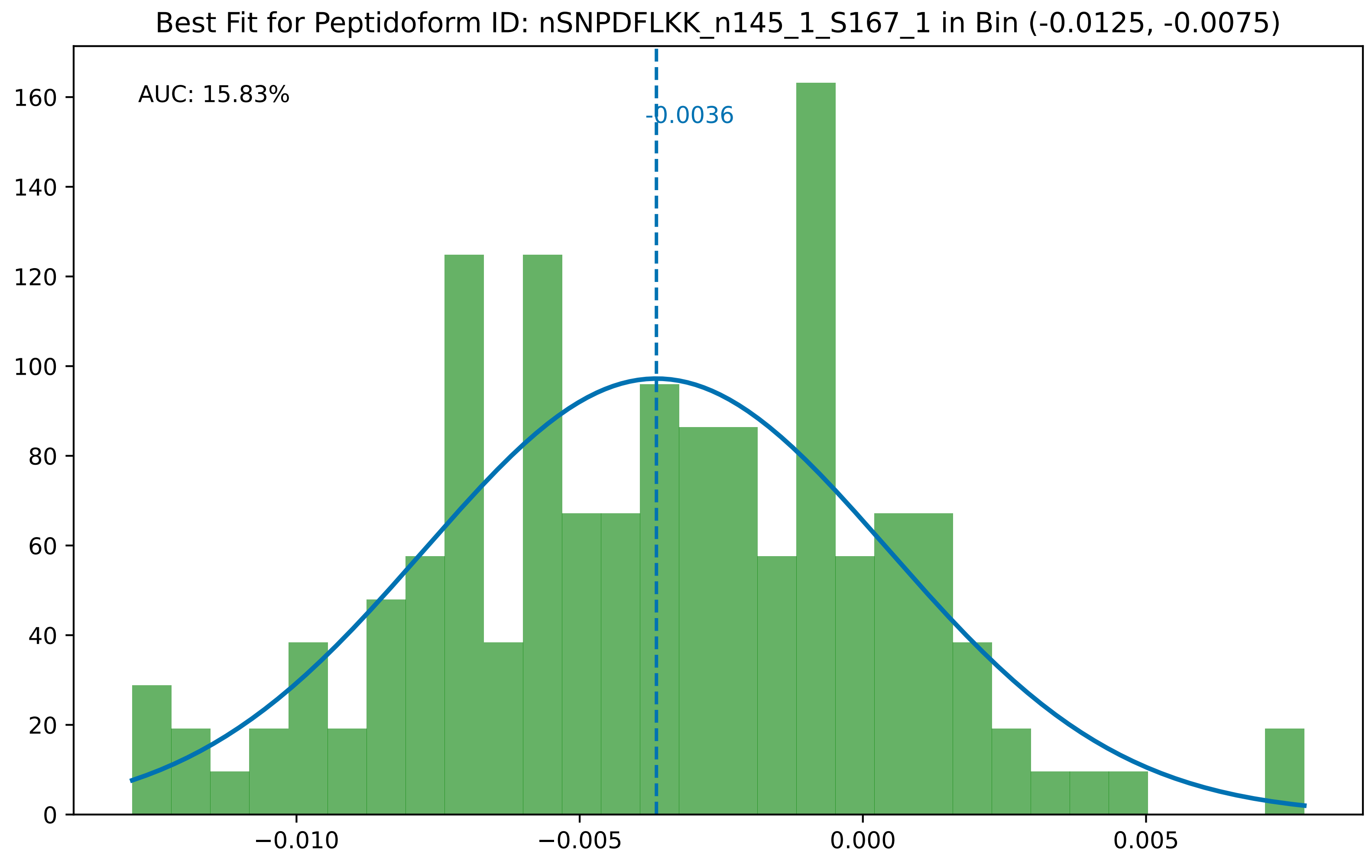
Best Fit for Peptidoform ID: nSNPDFLKK_n145_1_S167_1 in Bin (-0.0125, -0.0075)
AUC: 15.83%
-0.0036
0
20
40
60
80
100
120
140
160
−0.010
−0.005
0.000
0.005

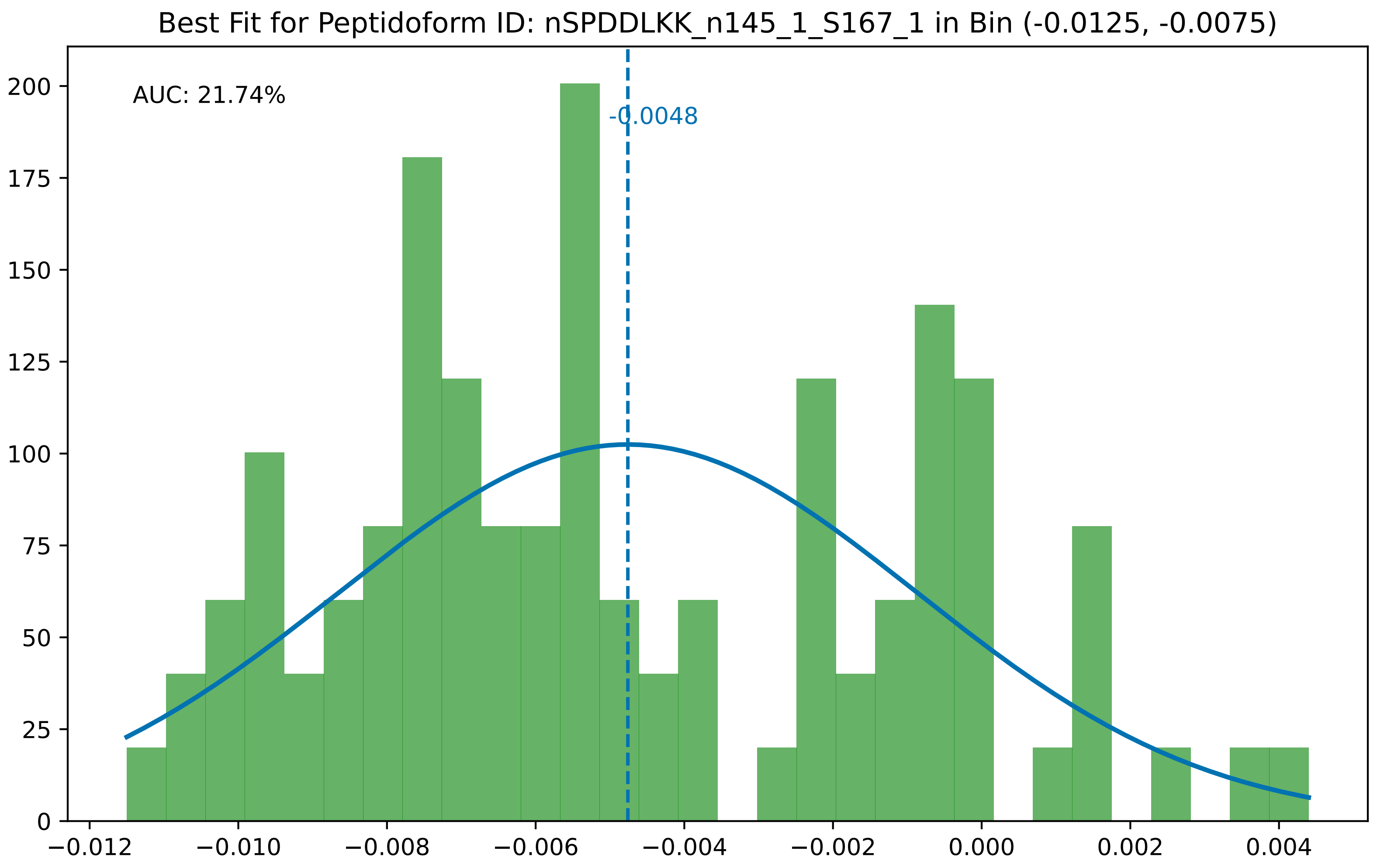
Best Fit for Peptidoform ID: nSPDDLKK_n145_1_S167_1 in Bin (-0.0125, -0.0075)
AUC: 21.74%
-0.0048
200
175
150
125
100
75
50
25
0
−0.012
−0.010
−0.008
−0.006
−0.004
−0.002
0.000
0.002
0.004

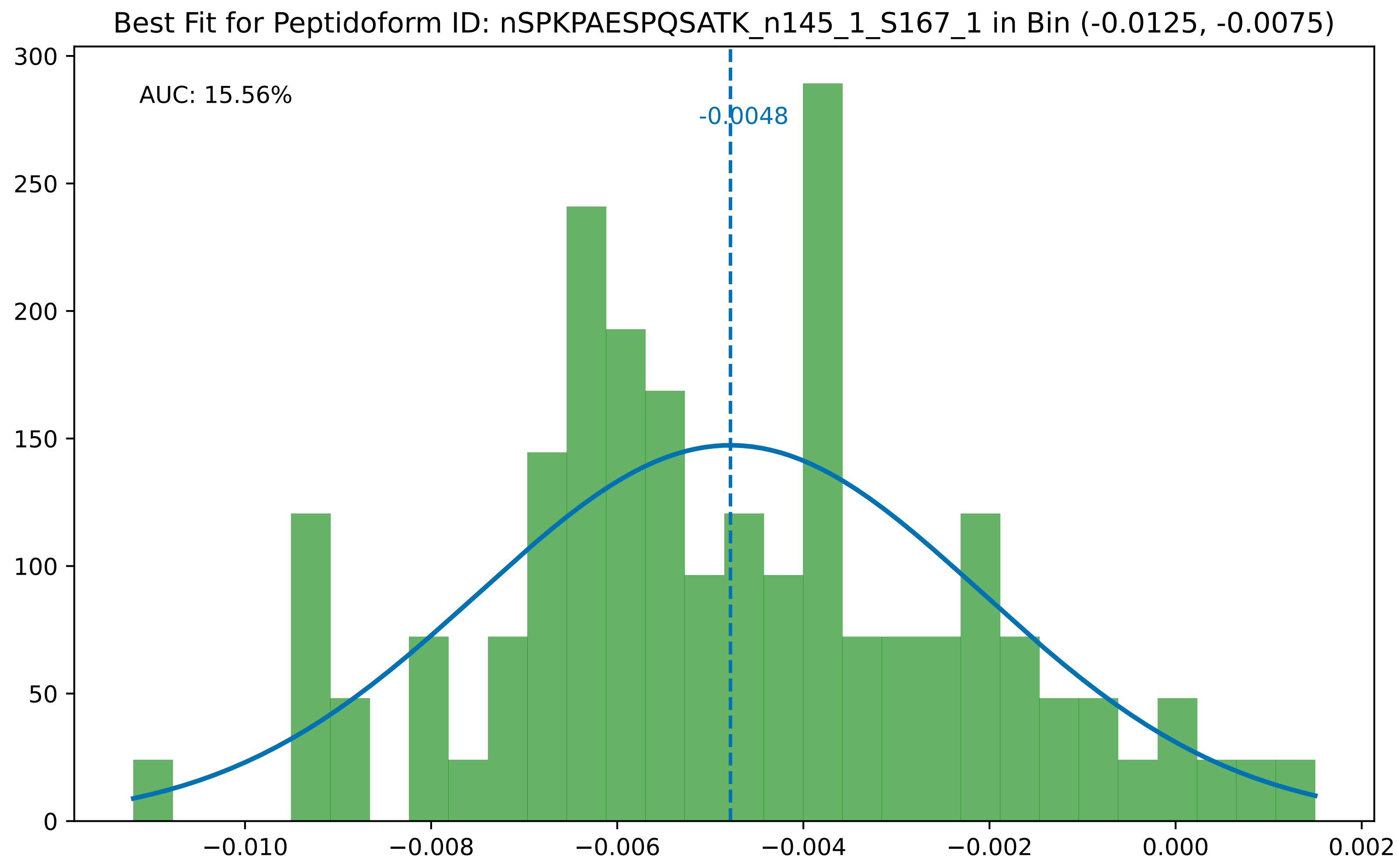
Best Fit for Peptidoform ID: nSPKPAESPQSATK_n145_1_S167_1 in Bin (-0.0125, -0.0075)
AUC: 15.56%
-0.0048
300
250
200
150
100
50
0
−0.010
−0.008
−0.006
−0.004
−0.002
0.000
0.002

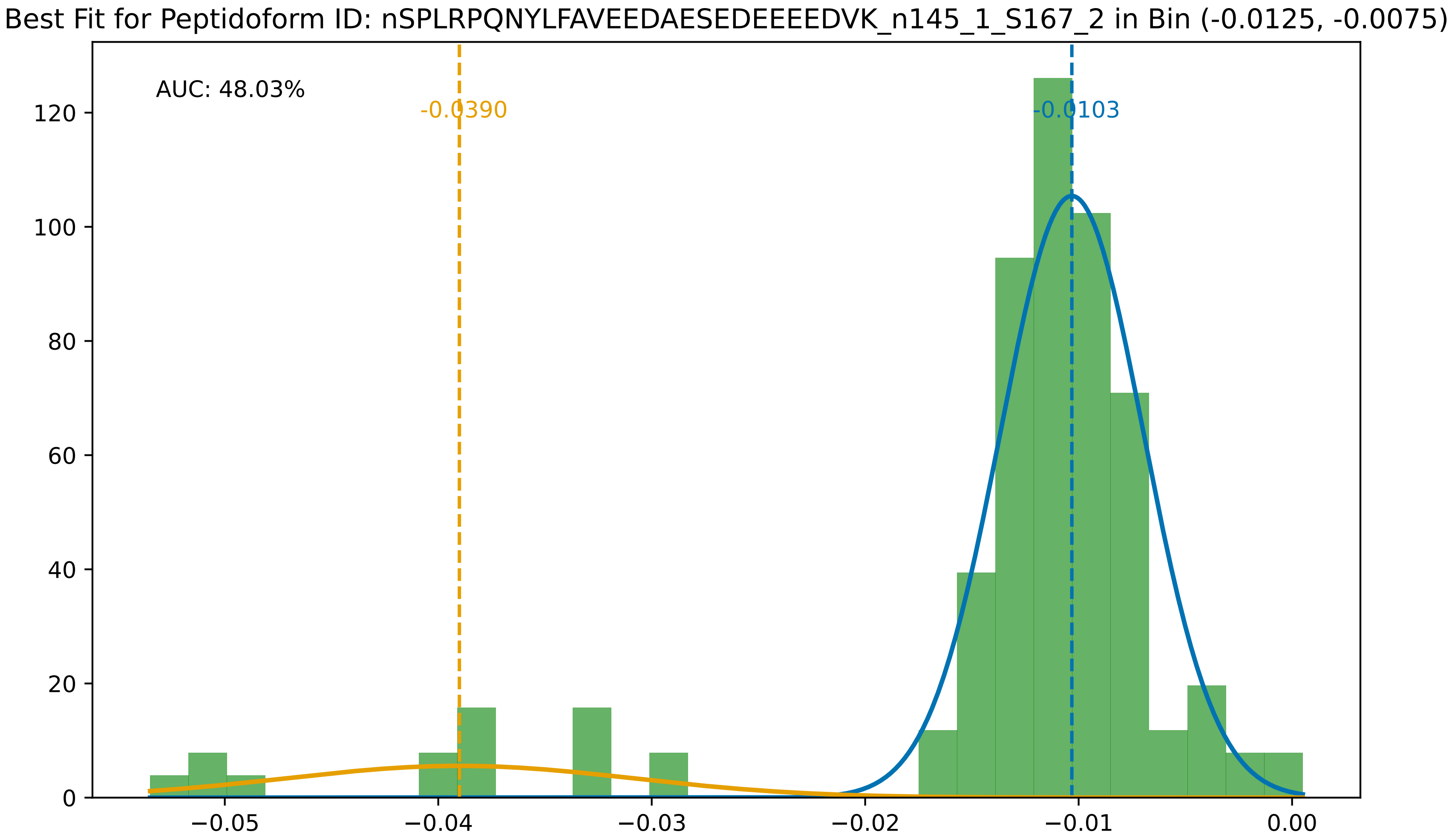
Best Fit for Peptidoform ID: nSPLRPQNYLFAVEEDAESEDEEEEDVK_n145_1_S167_2 in Bin (-0.0125, -0.0075)
AUC: 48.03%
-0.0390
-0.0103
0
20
40
60
80
100
120
−0.05
−0.04
−0.03
−0.02
−0.01
0.00

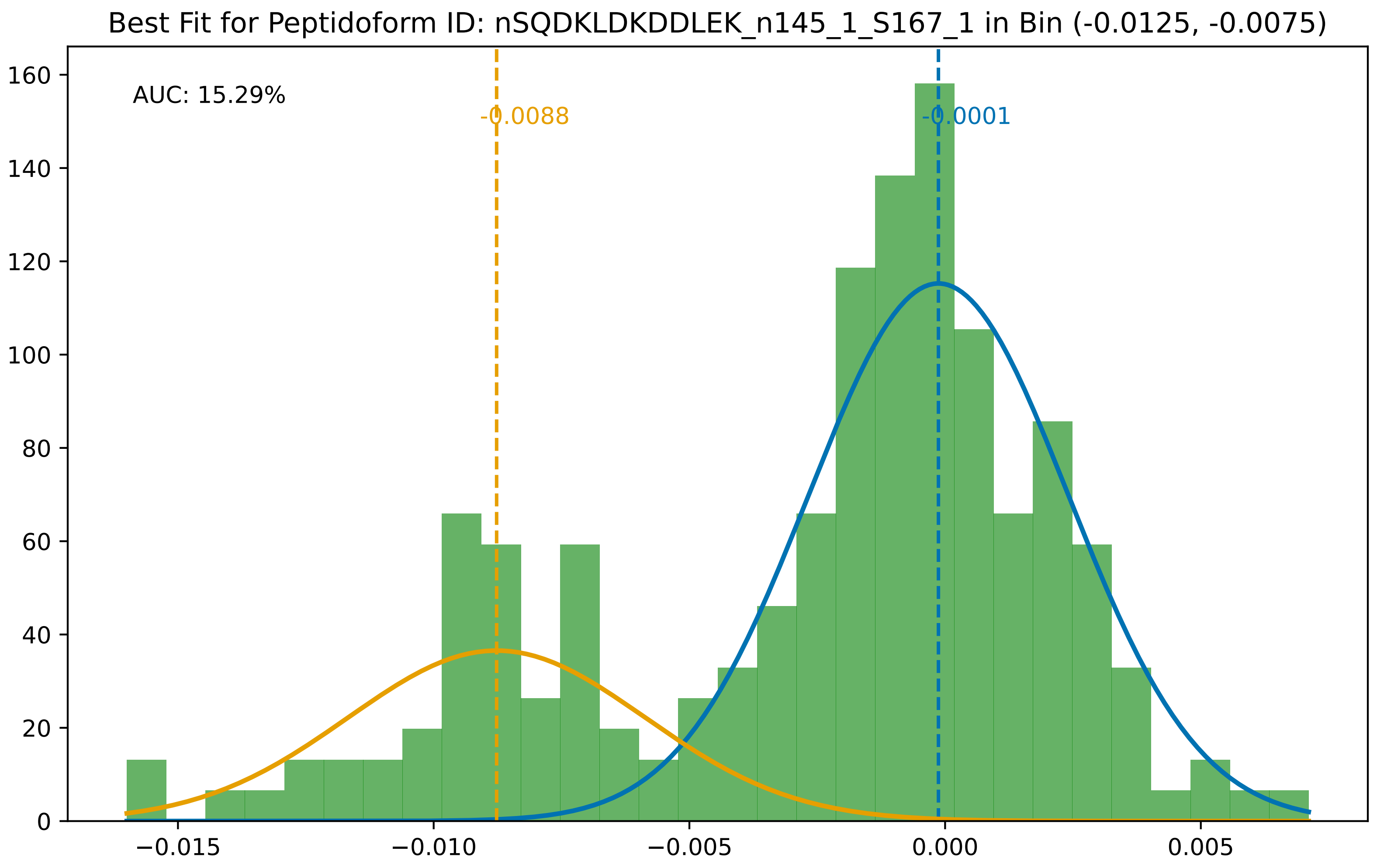
Best Fit for Peptidoform ID: nSQDKLDKDDLEK_n145_1_S167_1 in Bin (-0.0125, -0.0075)
AUC: 15.29%
-0.0088
-0.0001
0
20
40
60
80
100
120
140
160
−0.015
−0.010
−0.005
0.000
0.005

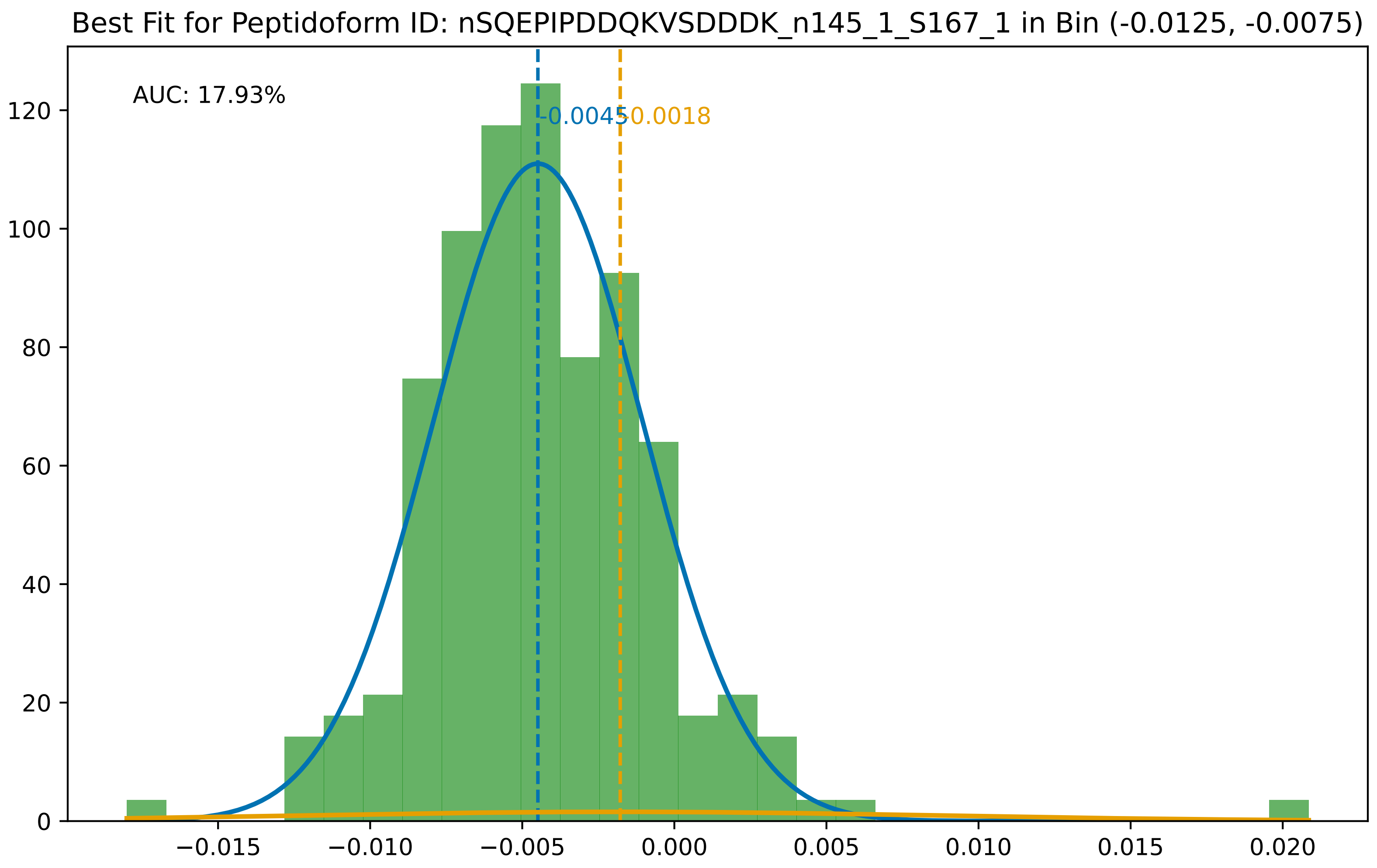
Best Fit for Peptidoform ID: nSQEPIPDDQKVSDDDK_n145_1_S167_1 in Bin (-0.0125, -0.0075)
AUC: 17.93%
-0.0045
0.0018
0
20
40
60
80
100
120
−0.015
−0.010
−0.005
0.000
0.005
0.010
0.015
0.020

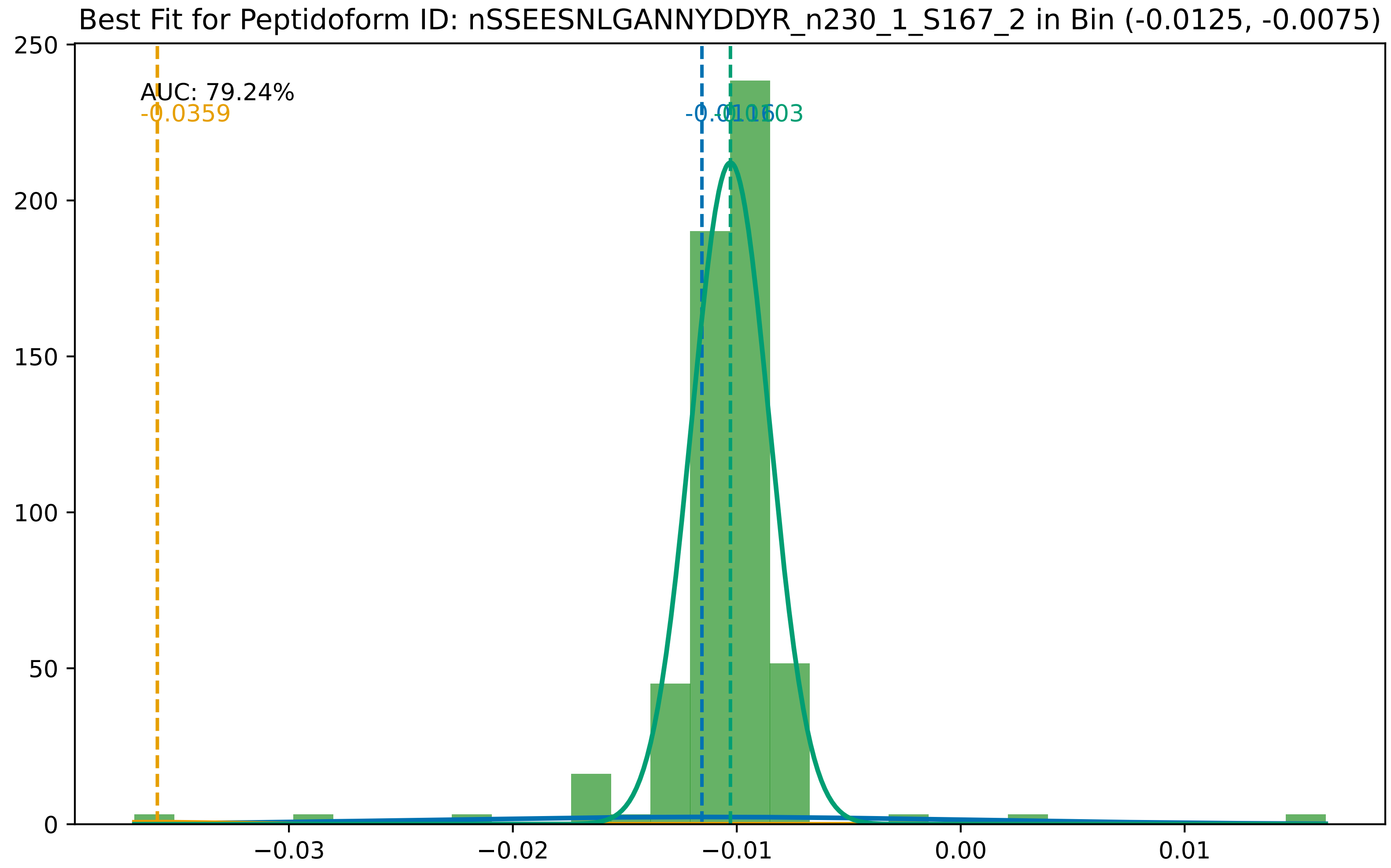
Best Fit for Peptidoform ID: nSSEESNLGANNYDDYR_n230_1_S167_2 in Bin (-0.0125, -0.0075)
AUC: 79.24%
-0.0359
250
200
150
100
50
0
−0.03
−0.02
−0.01
0.00
0.01

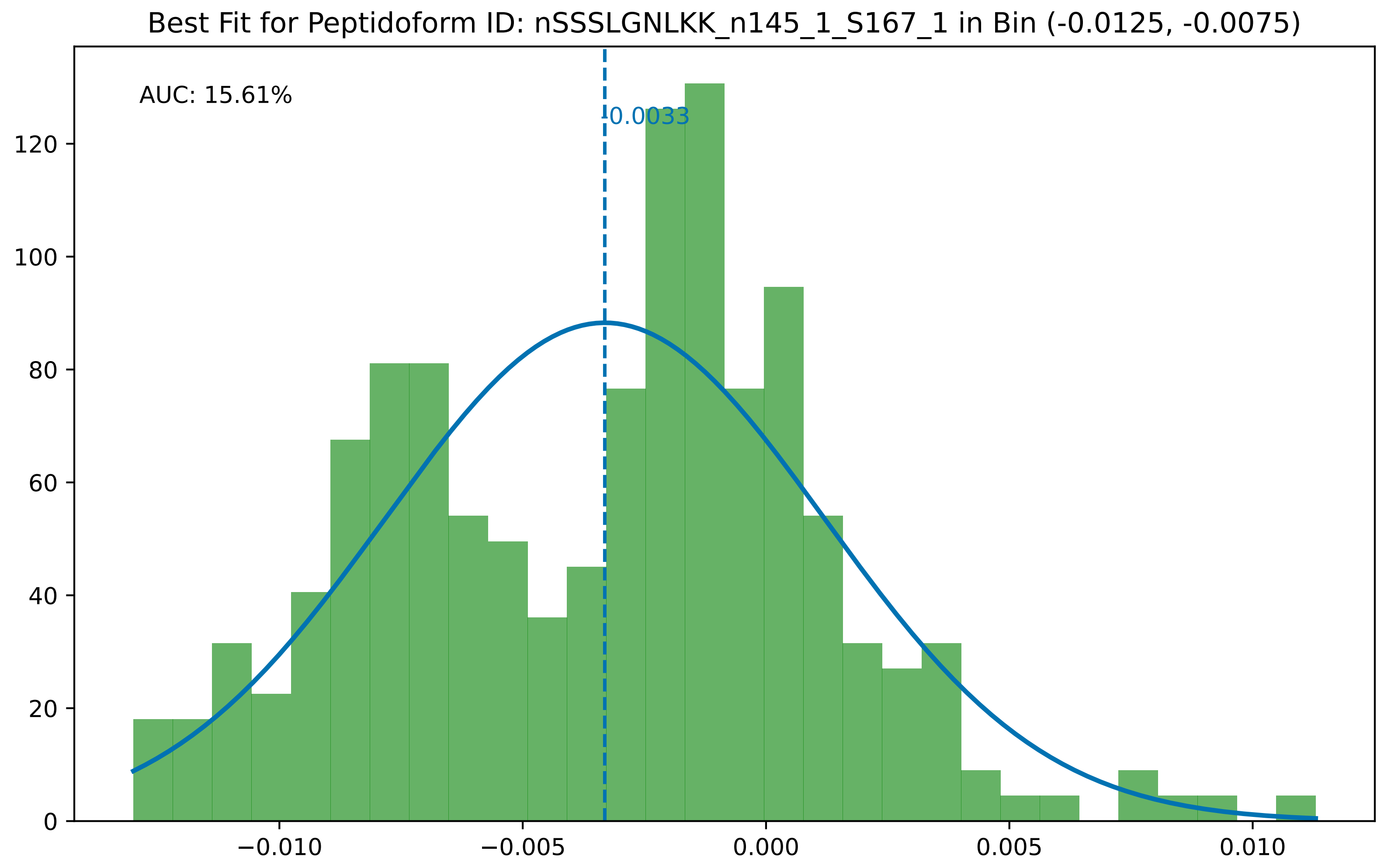

Best Fit for Peptidoform ID: nSSSLGNLKK_n145_1_S167_1 in Bin (-0.0125, -0.0075)
AUC: 15.61%
-0.0033
0
20
40
60
80
100
120
−0.010
−0.005
0.000
0.005
0.010

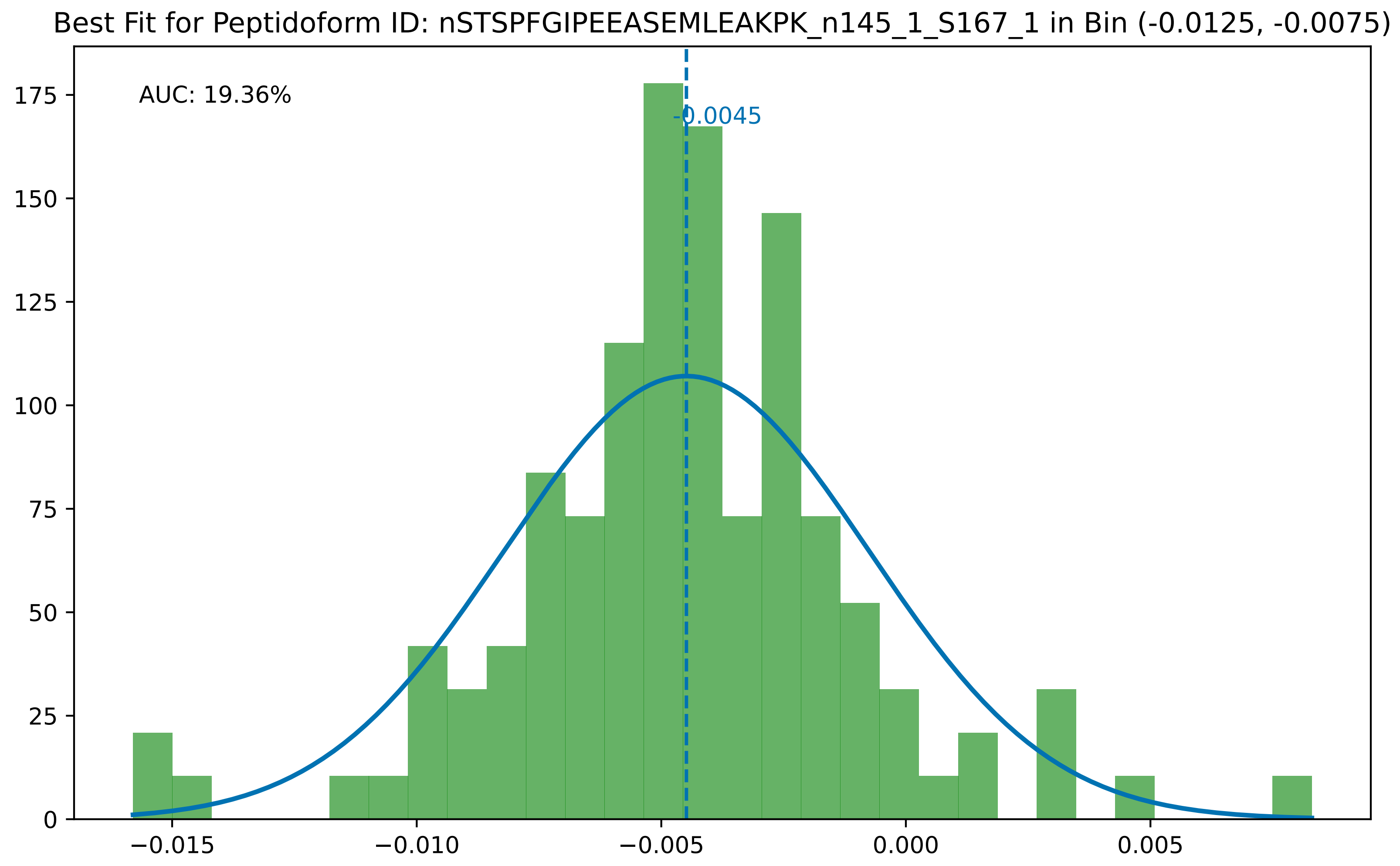

Best Fit for Peptidoform ID: nSTSPFGIPEEASEMLEAKPK_n145_1_S167_1 in Bin (-0.0125, -0.0075)
AUC: 19.36%
-0.0045
175
150
125
100
75
50
25
0
−0.015
−0.010
−0.005
0.000
0.005

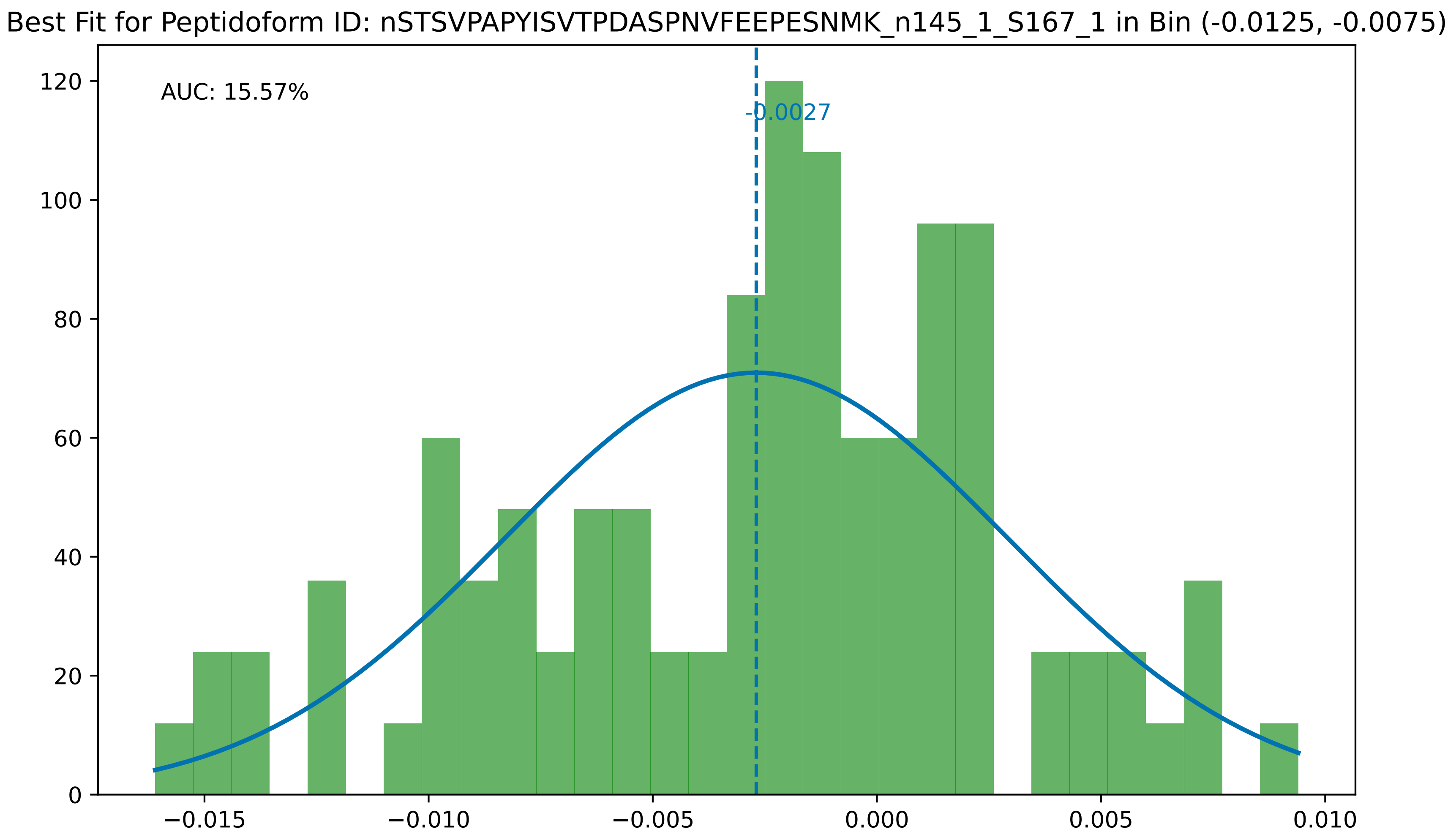

Best Fit for Peptidoform ID: nSTSVPAPYISVTPDASPNVFEEPESNMK_n145_1_S167_1 in Bin (-0.0125, -0.0075)
AUC: 15.57%
-0.0027
0
20
40
60
80
100
120
−0.015
−0.010
−0.005
0.000
0.005
0.010

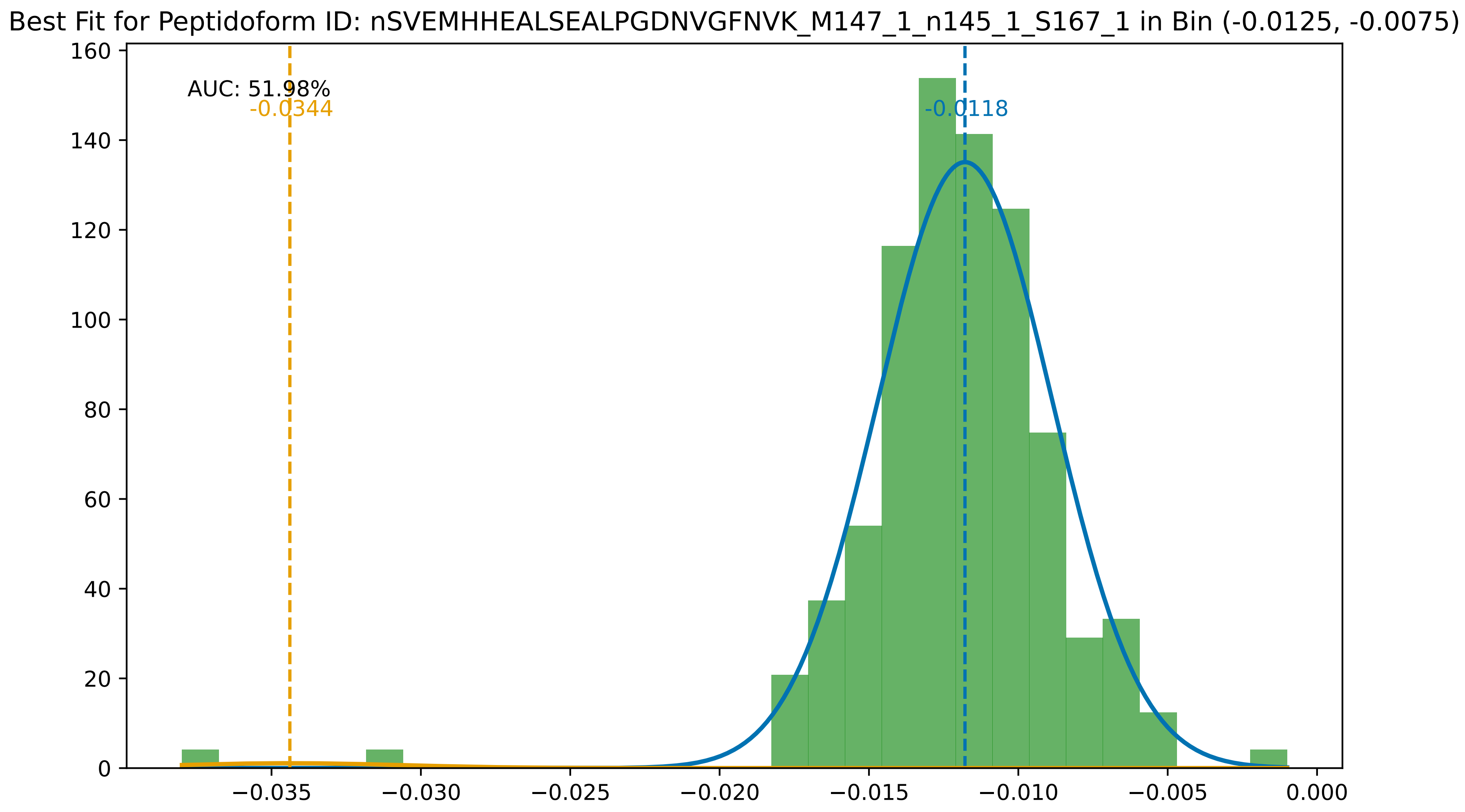
Best Fit for Peptidoform ID: nSVEMHHEALSEALPGDNVGFNVK_M147_1_n145_1_S167_1 in Bin (-0.0125, -0.0075)
AUC: 51.98%
-0.0344
-0.0118
0
20
40
60
80
100
120
140
160
−0.035
−0.030
−0.025
−0.020
−0.015
−0.010
−0.005
0.000

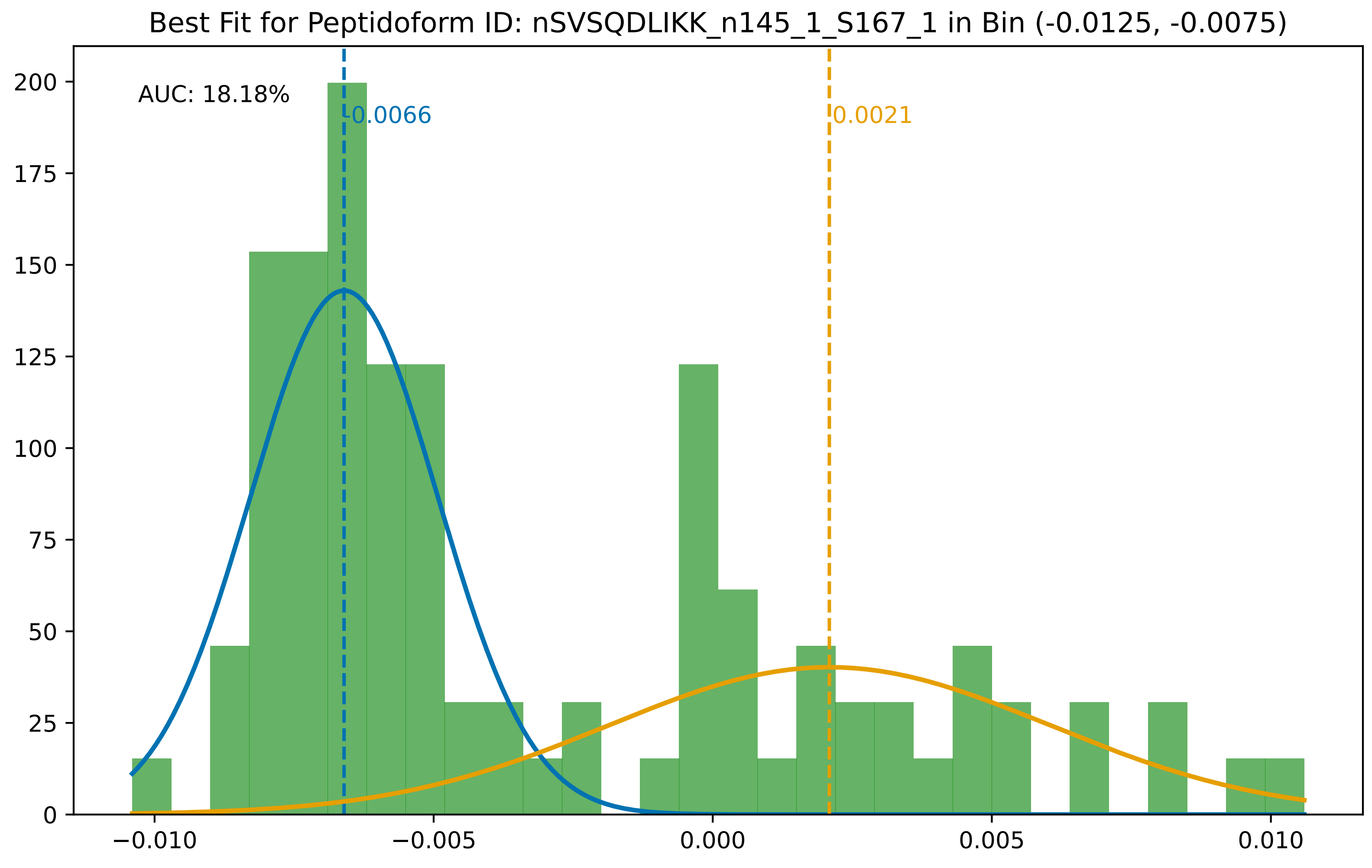

Best Fit for Peptidoform ID: nSVSQDLIKK_n145_1_S167_1 in Bin (-0.0125, -0.0075)
AUC: 18.18%
-0.0066
0.0021
200
175
150
125
100
75
50
25
0
−0.010
−0.005
0.000
0.005
0.010

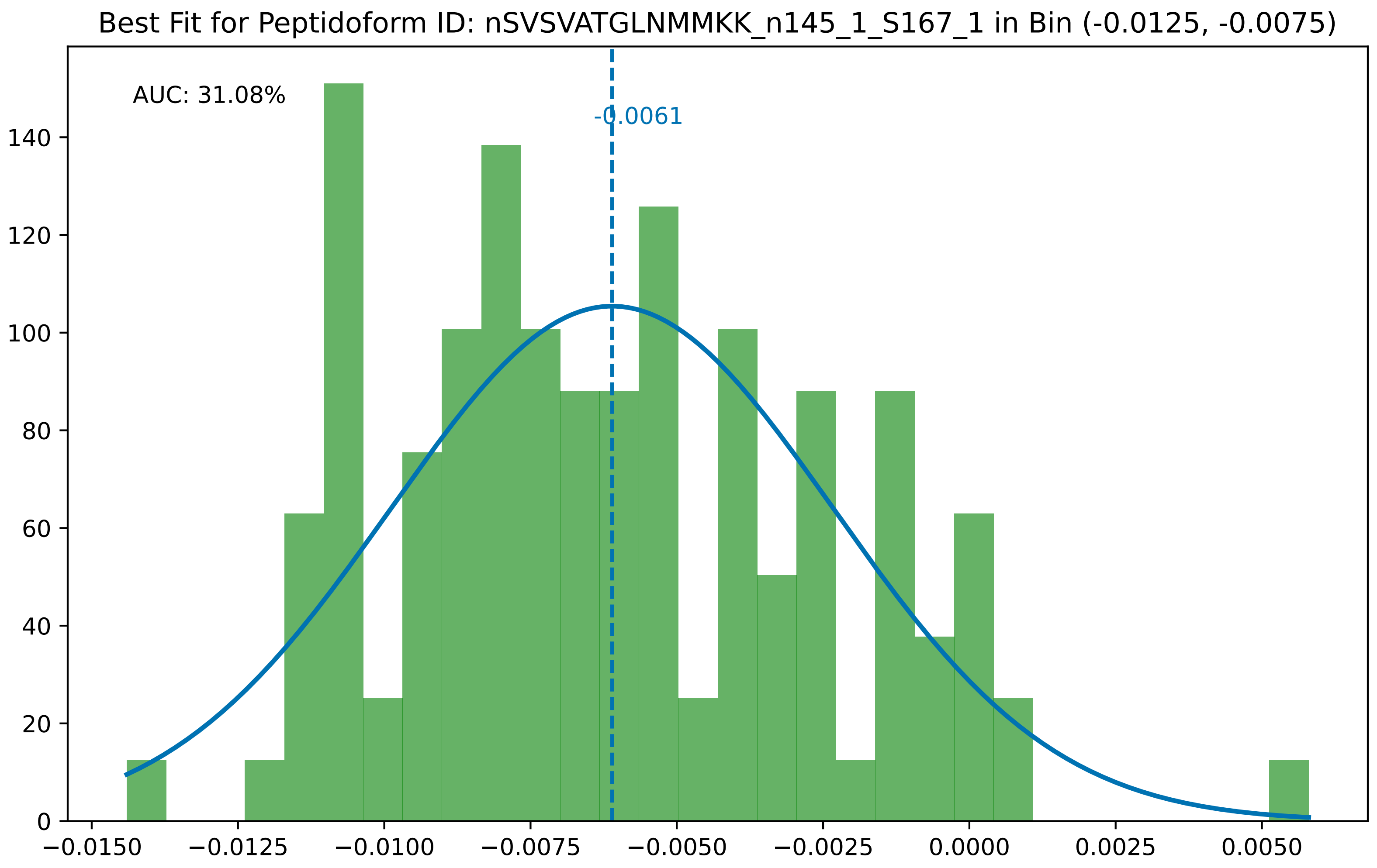

Best Fit for Peptidoform ID: nSVSVATGLNMMKK_n145_1_S167_1 in Bin (-0.0125, -0.0075)
AUC: 31.08%
-0.0061
0
20
40
60
80
100
120
140
−0.0150
−0.0125
−0.0100
−0.0075
−0.0050
−0.0025
0.0000
0.0025
0.0050

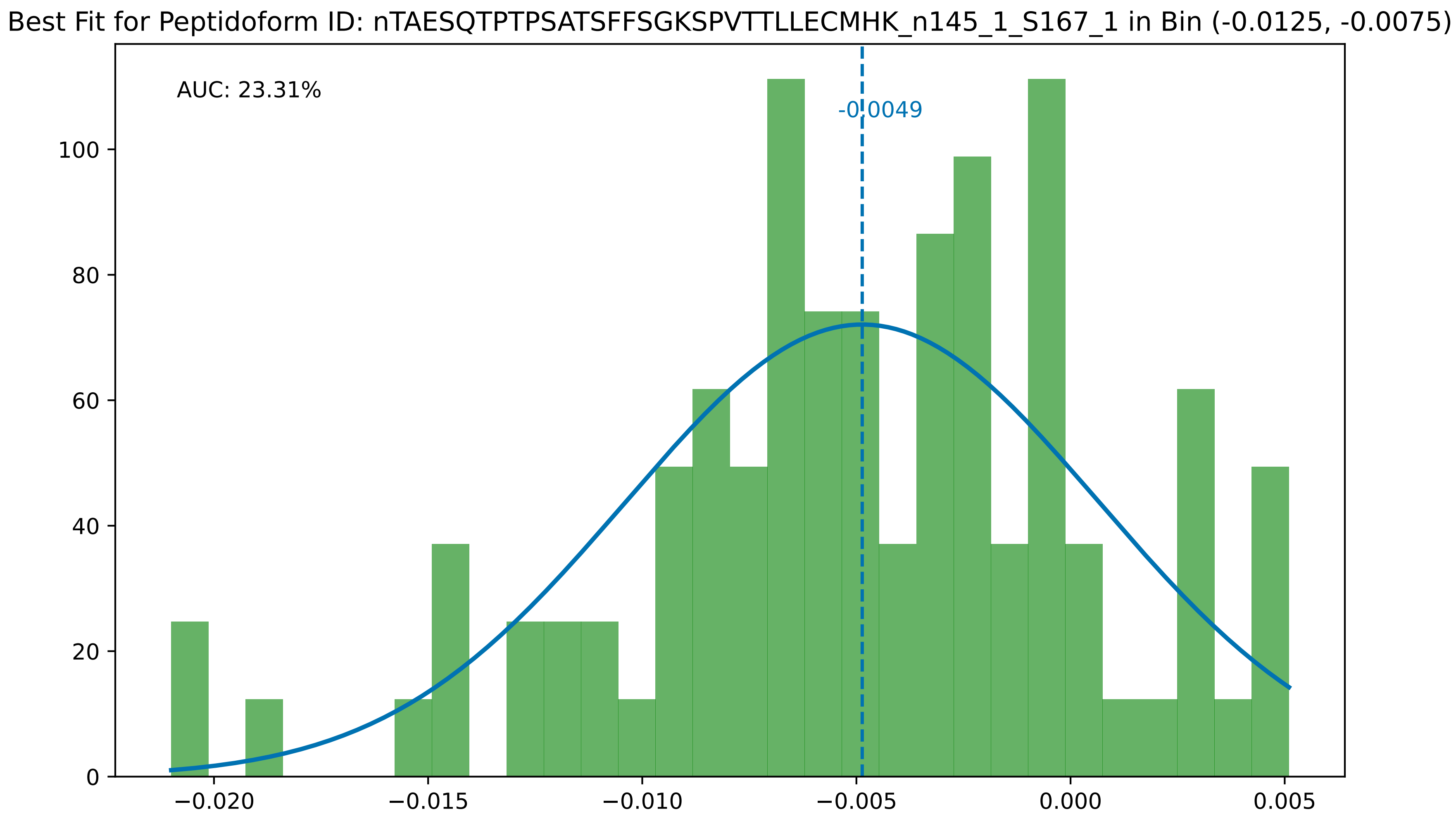

Best Fit for Peptidoform ID: nTAESQTPTPSATSFFSGKSPVTTLLECMHK_n145_1_S167_1 in Bin (-0.0125, -0.0075)
AUC: 23.31%
-0.0049
0
20
40
60
80
100
−0.020
−0.015
−0.010
−0.005
0.000
0.005

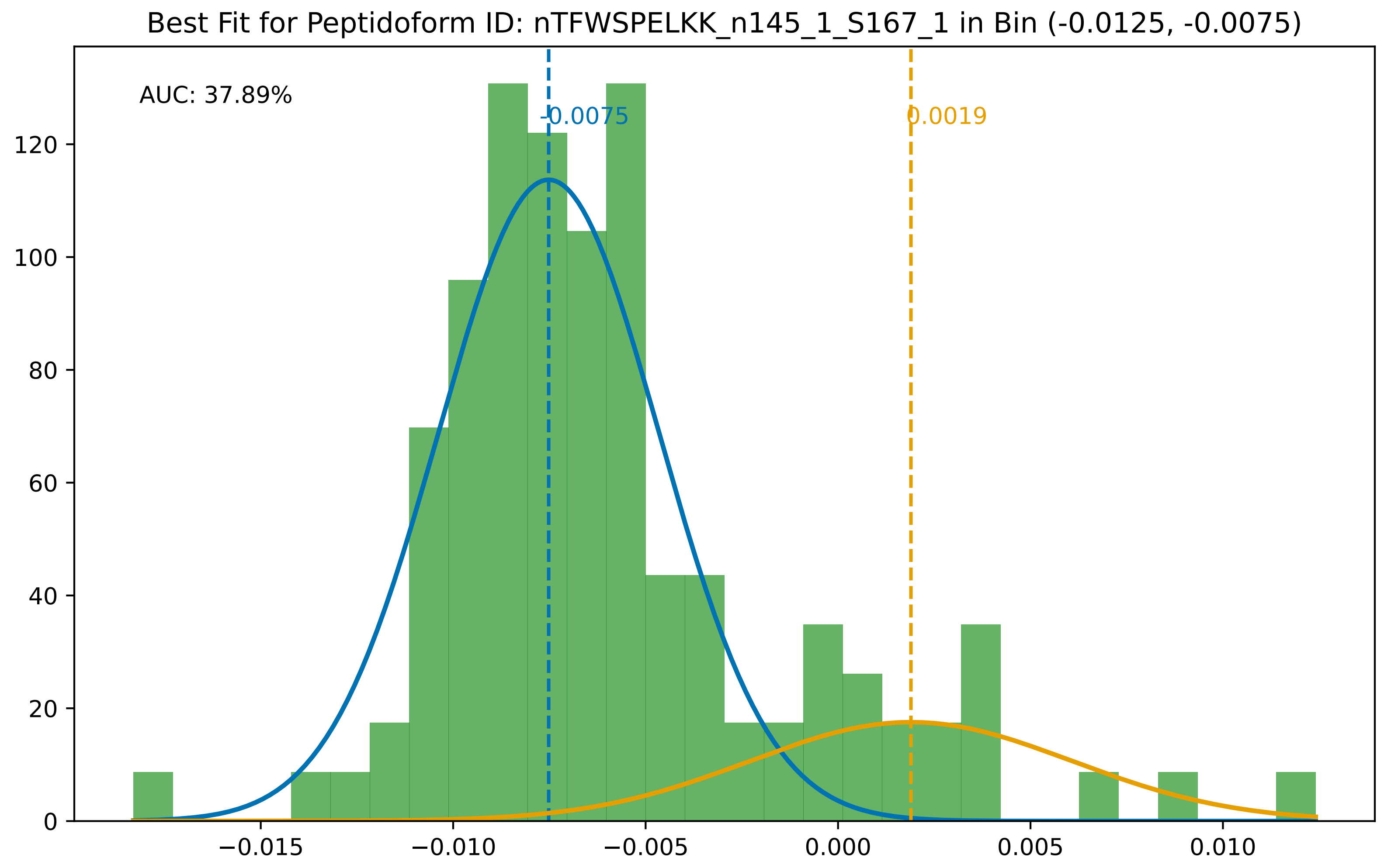

Best Fit for Peptidoform ID: nTFWSPELKK_n145_1_S167_1 in Bin (-0.0125, -0.0075)
AUC: 37.89%
-0.0075
0.0019
0
20
40
60
80
100
120
−0.015
−0.010
−0.005
0.000
0.005
0.010

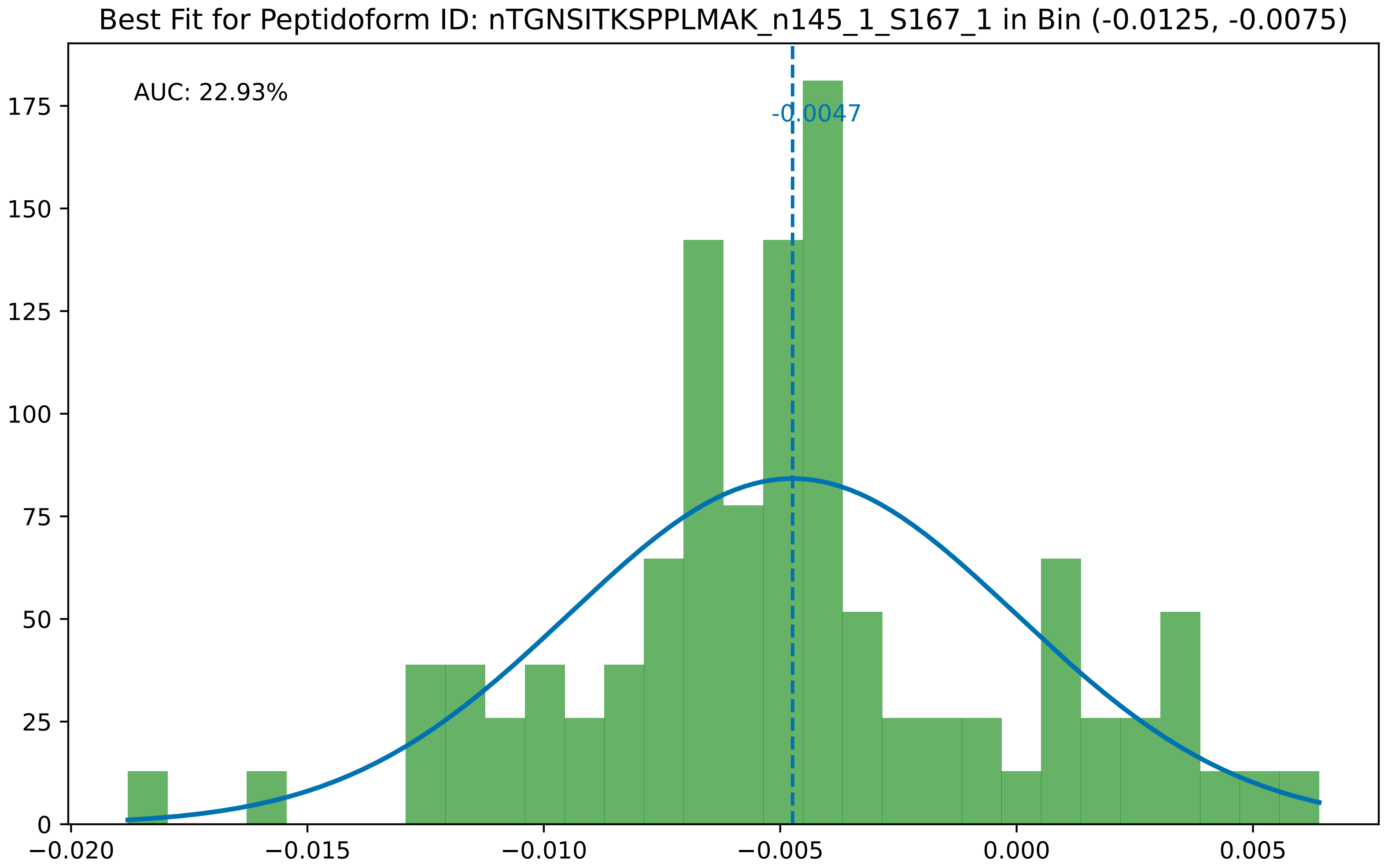
Best Fit for Peptidoform ID: nTGNSITKSPPLMAK_n145_1_S167_1 in Bin (-0.0125, -0.0075)
AUC: 22.93%
-0.0047
0
25
50
75
100
125
150
175
−0.020
−0.015
−0.010
−0.005
0.000
0.005

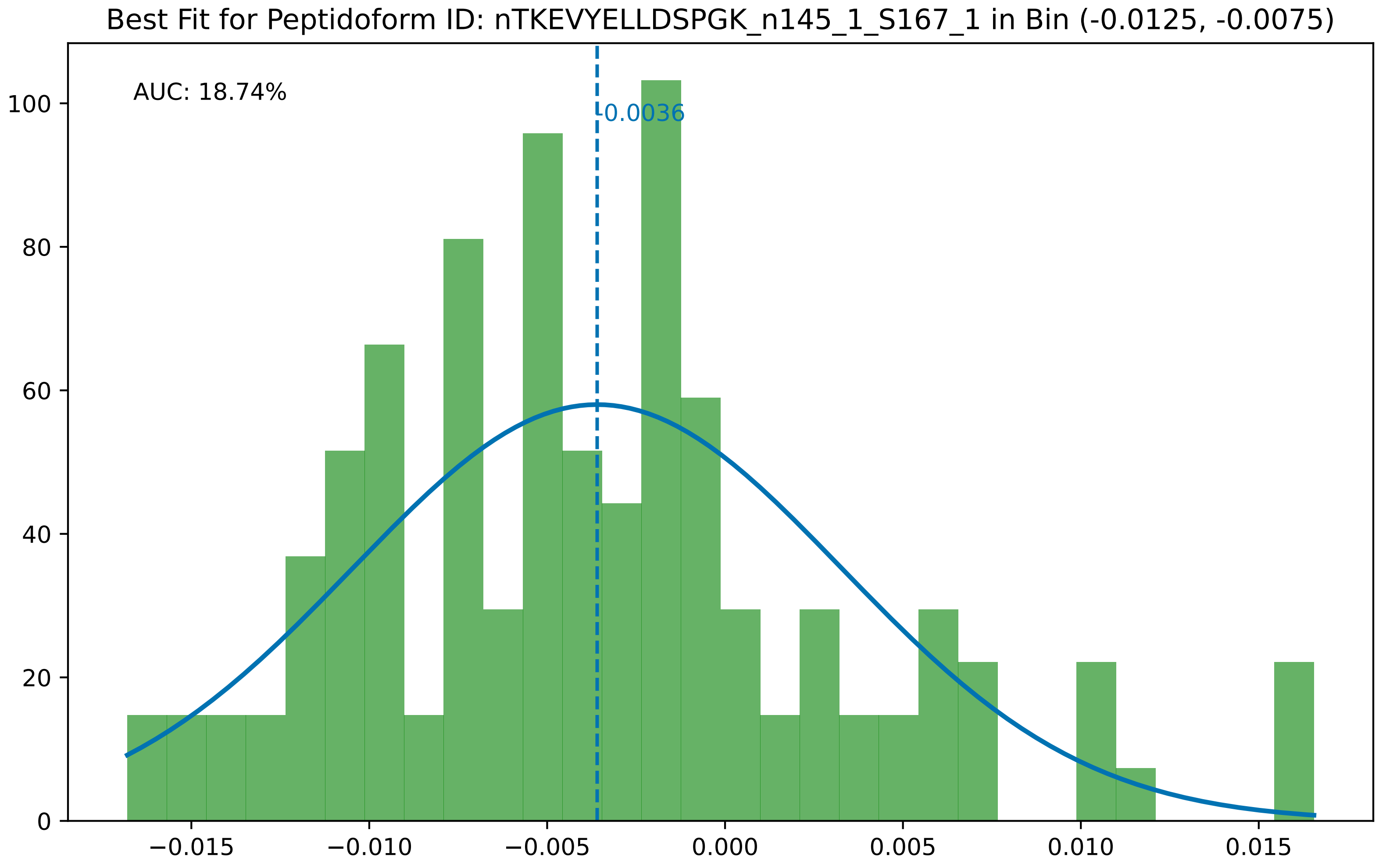

Best Fit for Peptidoform ID: nTKEVYELLDSPGK_n145_1_S167_1 in Bin (-0.0125, -0.0075)
AUC: 18.74%
-0.0036
0
20
40
60
80
100
−0.015
−0.010
−0.005
0.000
0.005
0.010
0.015

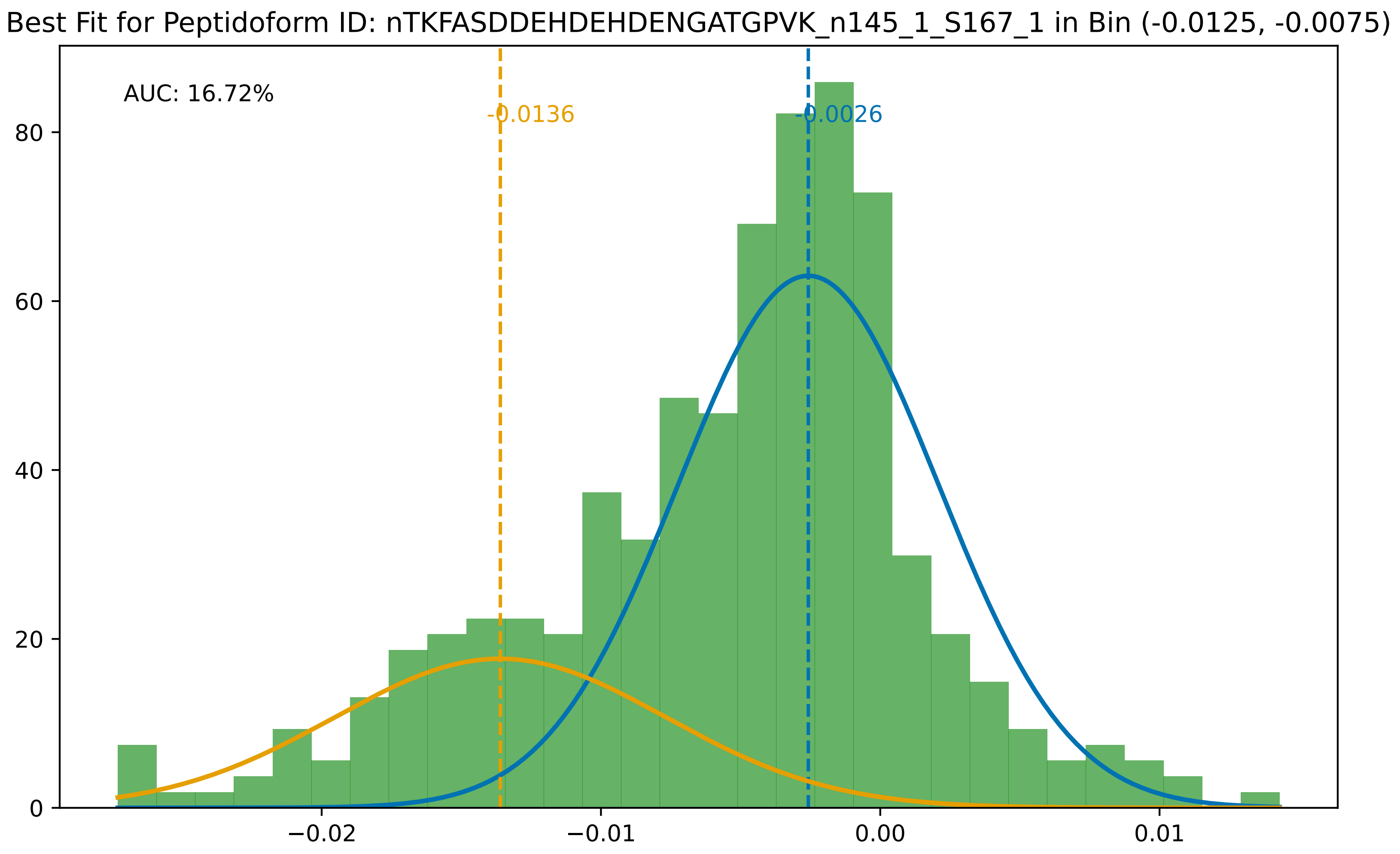
Best Fit for Peptidoform ID: nTKFASDDEHDEHDENGATGPVK_n145_1_S167_1 in Bin (-0.0125, -0.0075)
AUC: 16.72%
-0.0136
-0.0026
80
60
40
20
0
−0.02
−0.01
0.00
0.01

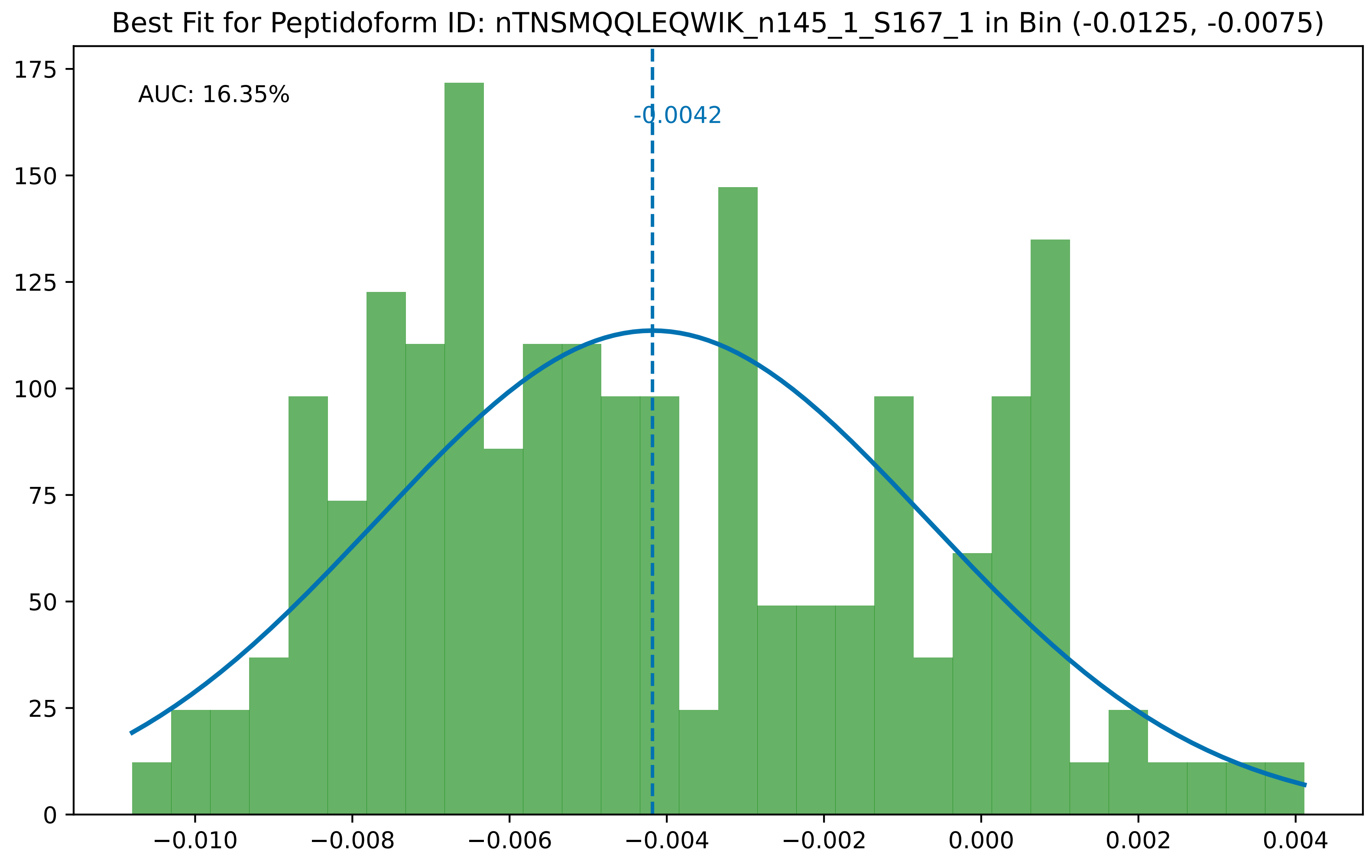

Best Fit for Peptidoform ID: nTNSMQQLEQWIK_n145_1_S167_1 in Bin (-0.0125, -0.0075)
AUC: 16.35%
-0.0042
175
150
125
100
75
50
25
0
−0.010
−0.008
−0.006
−0.004
−0.002
0.000
0.002
0.004

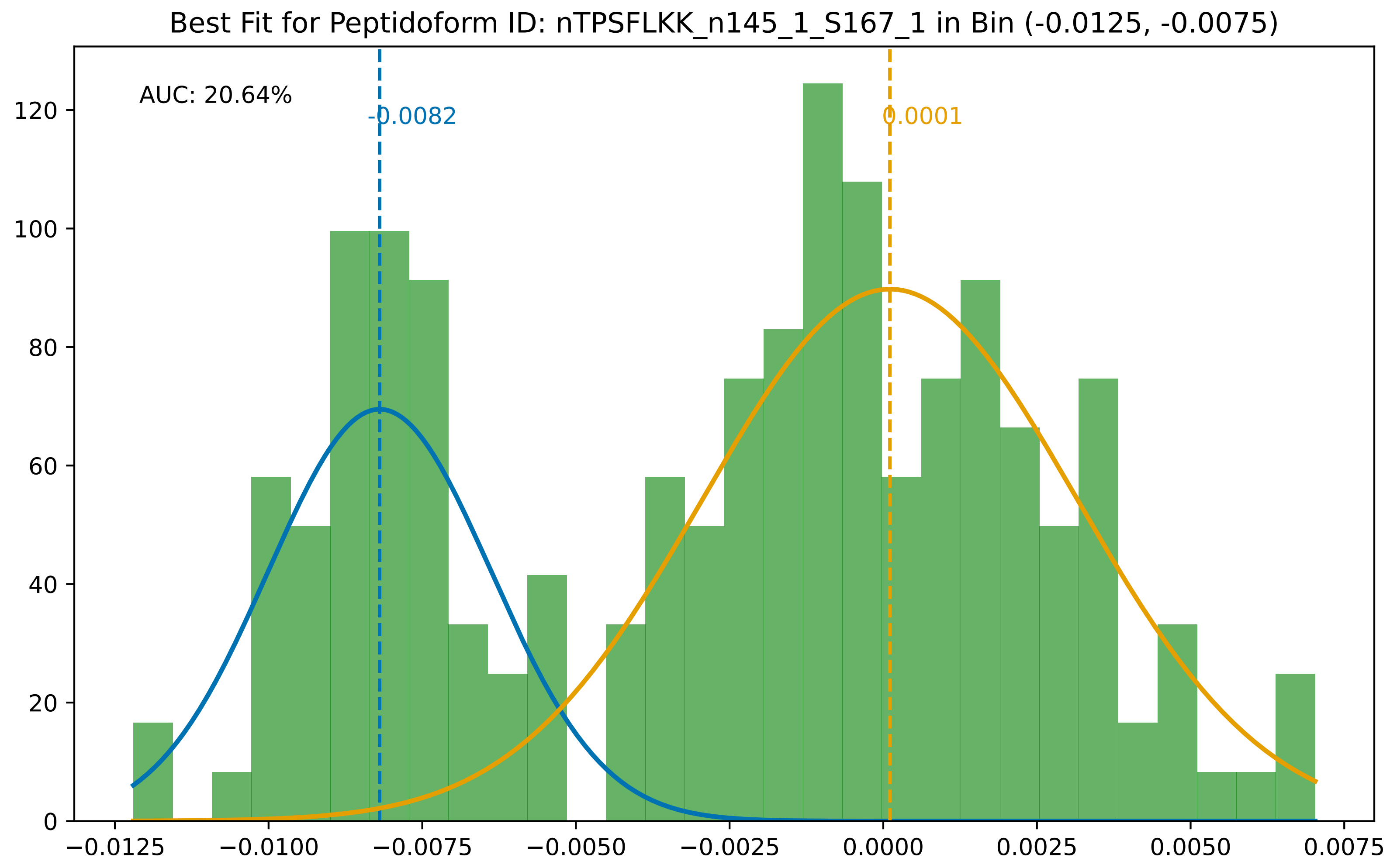

Best Fit for Peptidoform ID: nTPSFLKK_n145_1_S167_1 in Bin (-0.0125, -0.0075)
AUC: 20.64%
-0.0082
0.0001
0
20
40
60
80
100
120
−0.0125
−0.0100
−0.0075
−0.0050
−0.0025
0.0000
0.0025
0.0050
0.0075

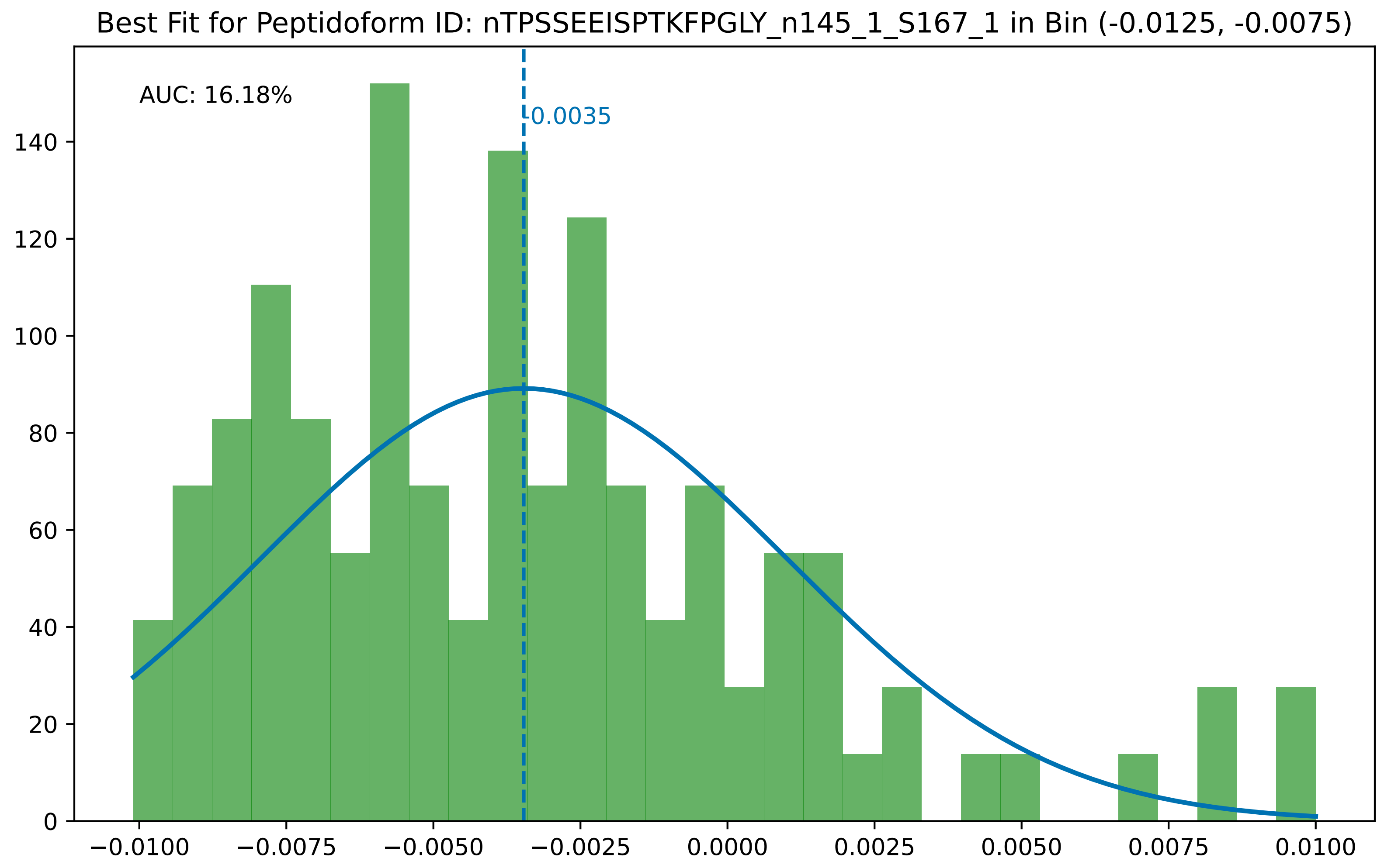

Best Fit for Peptidoform ID: nTPSSEEISPTKFPGLY_n145_1_S167_1 in Bin (-0.0125, -0.0075)
AUC: 16.18%
-0.0035
0
20
40
60
80
100
120
140
−0.0100
−0.0075
−0.0050
−0.0025
0.0000
0.0025
0.0050
0.0075
0.0100

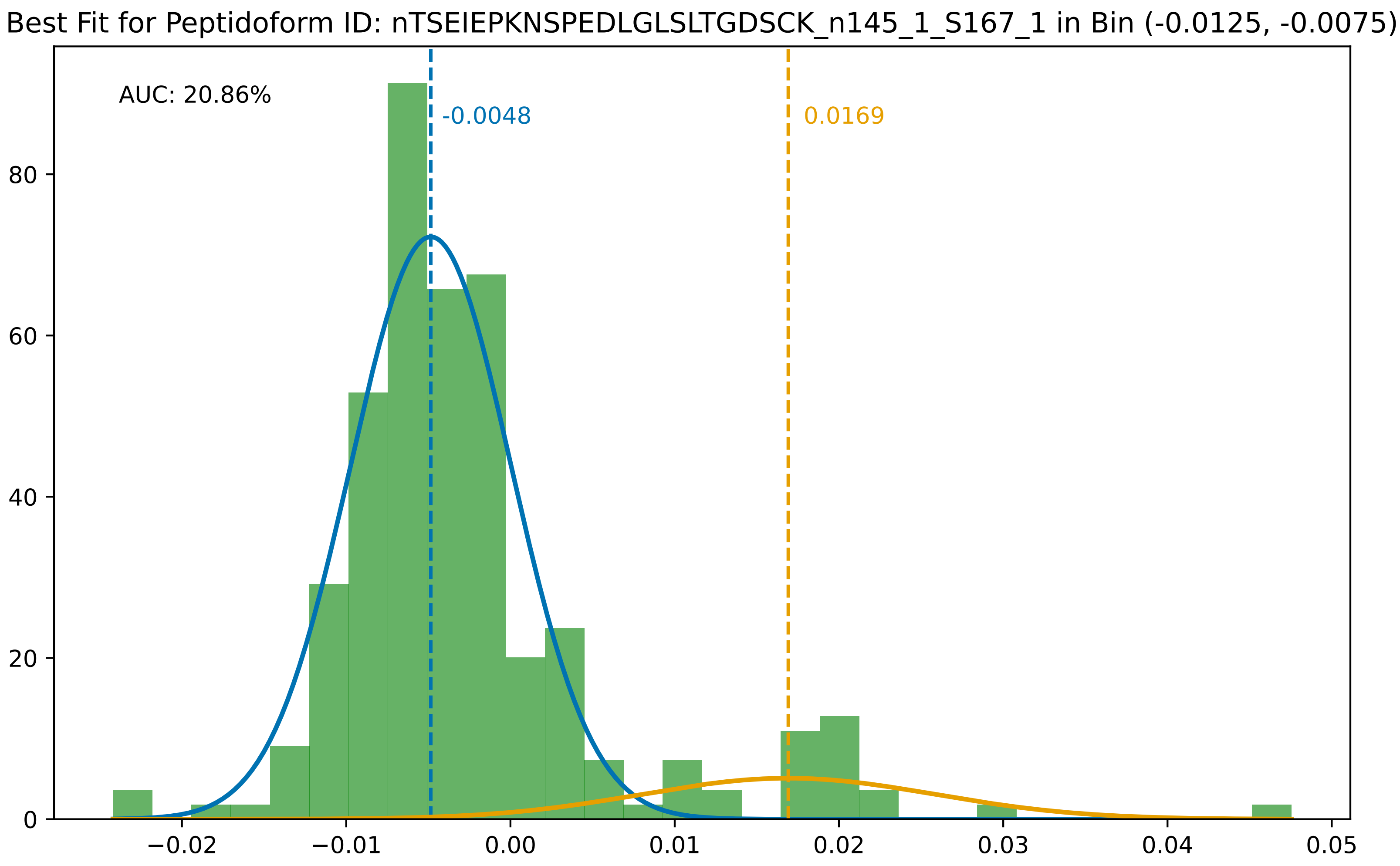

Best Fit for Peptidoform ID: nTSEIEPKNSPEDLGLSLTGDSCK_n145_1_S167_1 in Bin (-0.0125, -0.0075)
AUC: 20.86%
-0.0048
0.0169
0
20
40
60
80
−0.02
−0.01
0.00
0.01
0.02
0.03
0.04
0.05

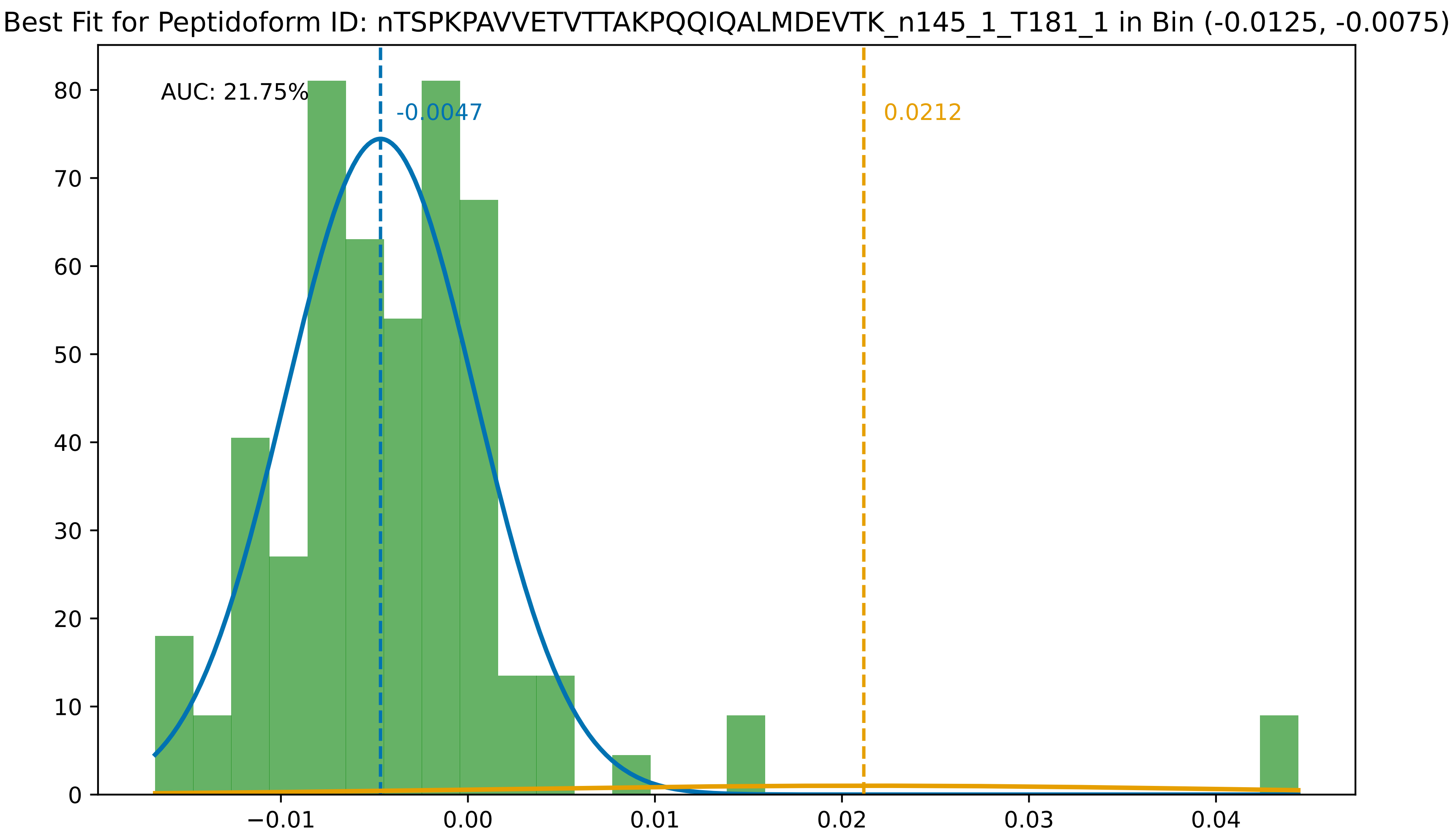
Best Fit for Peptidoform ID: nTSPKPAVVETVTTAKPQQIQALMDEVTK_n145_1_T181_1 in Bin (-0.0125, -0.0075)
AUC: 21.75%
-0.0047
0.0212
0
10
20
30
40
50
60
70
80
−0.01
0.00
0.01
0.02
0.03
0.04

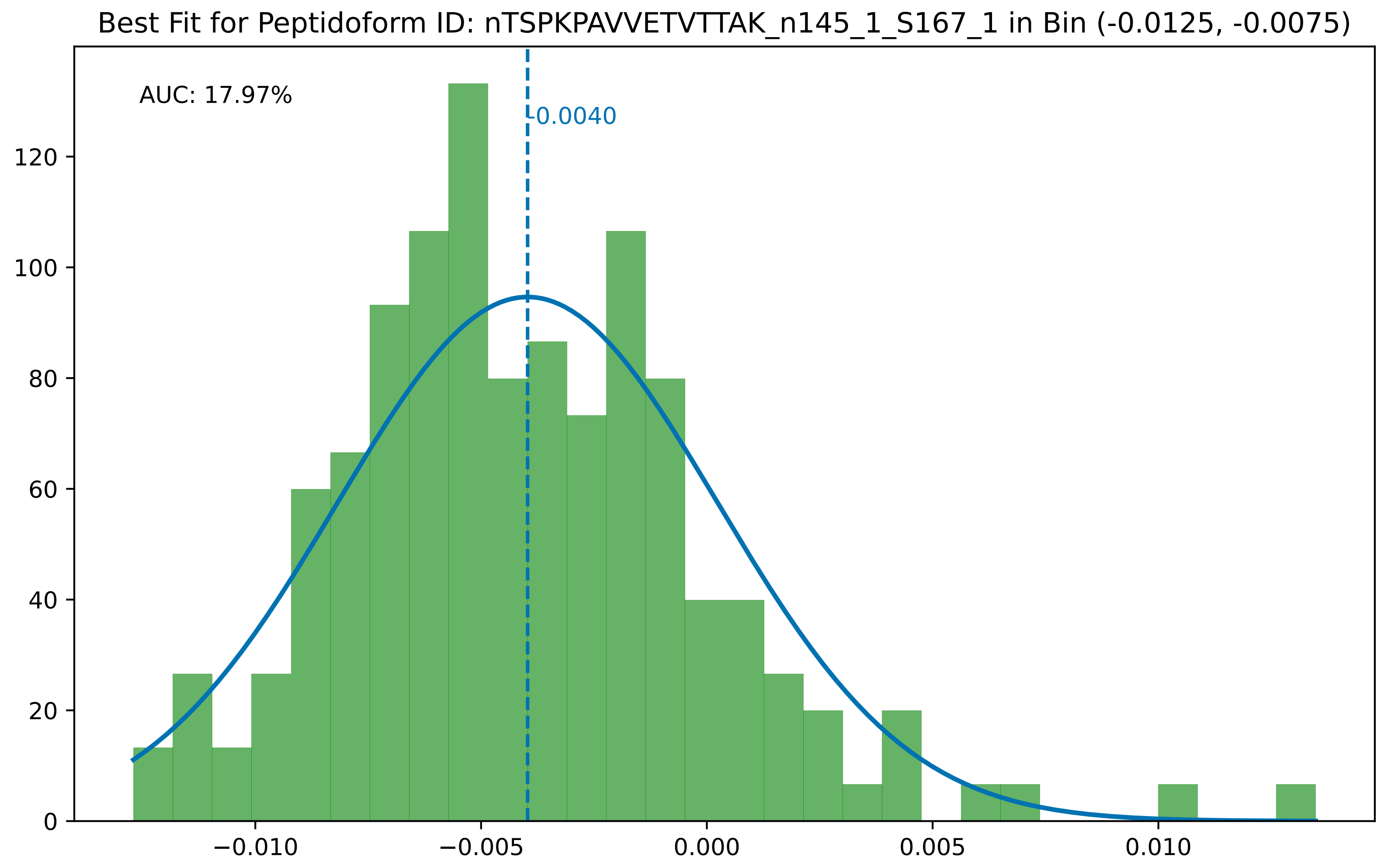

Best Fit for Peptidoform ID: nTSPKPAVVETVTTAK_n145_1_S167_1 in Bin (-0.0125, -0.0075)
AUC: 17.97%
-0.0040
0
20
40
60
80
100
120
−0.010
−0.005
0.000
0.005
0.010

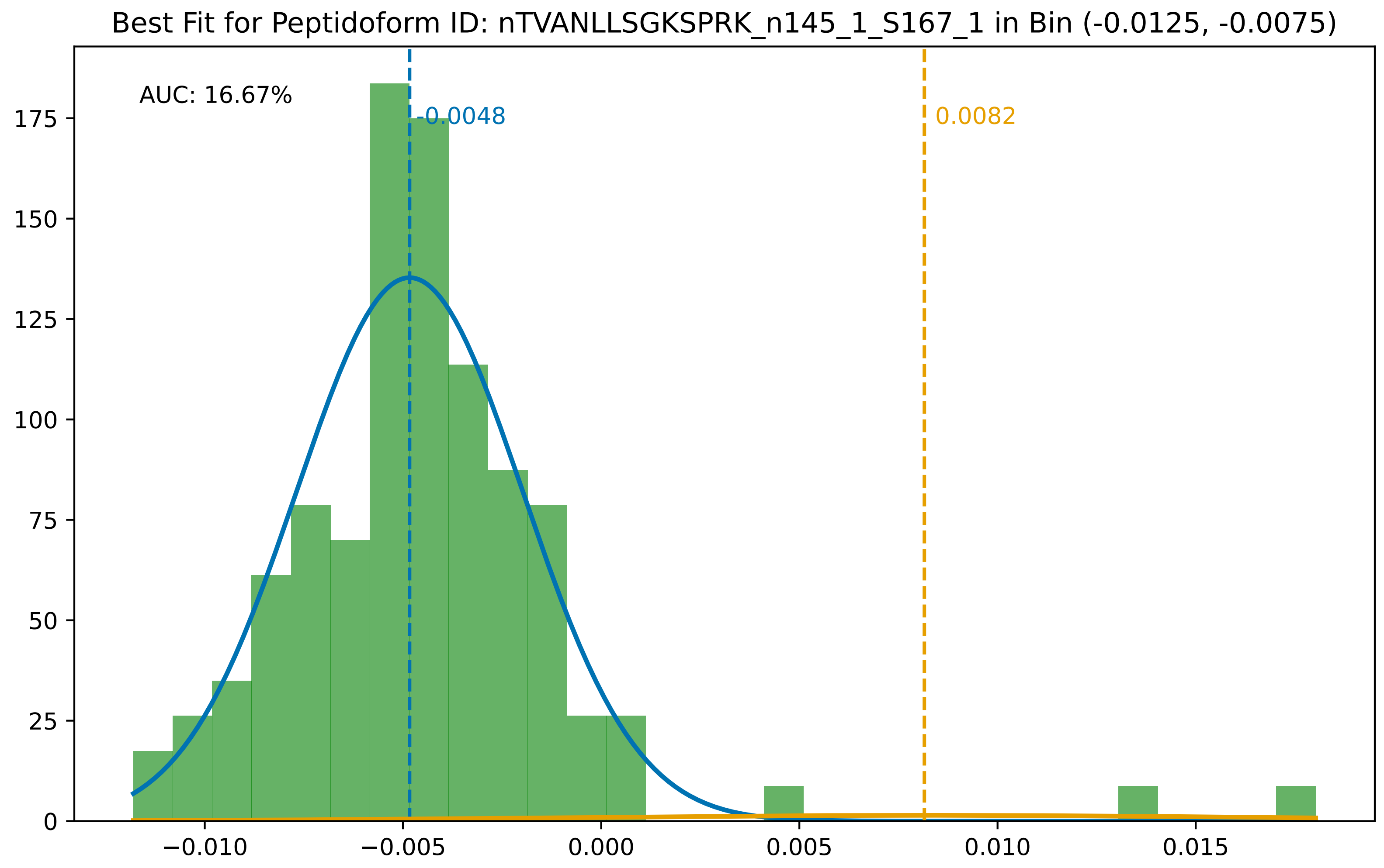

Best Fit for Peptidoform ID: nTVANLLSGKSPRK_n145_1_S167_1 in Bin (-0.0125, -0.0075)
AUC: 16.67%
-0.0048
0.0082
175
150
125
100
75
50
25
0
−0.010
−0.005
0.000
0.005
0.010
0.015

Best Fit for Peptidoform ID: nTVITEEFKVPDK_n145_1_T181_1 in Bin (-0.0125, -0.0075)

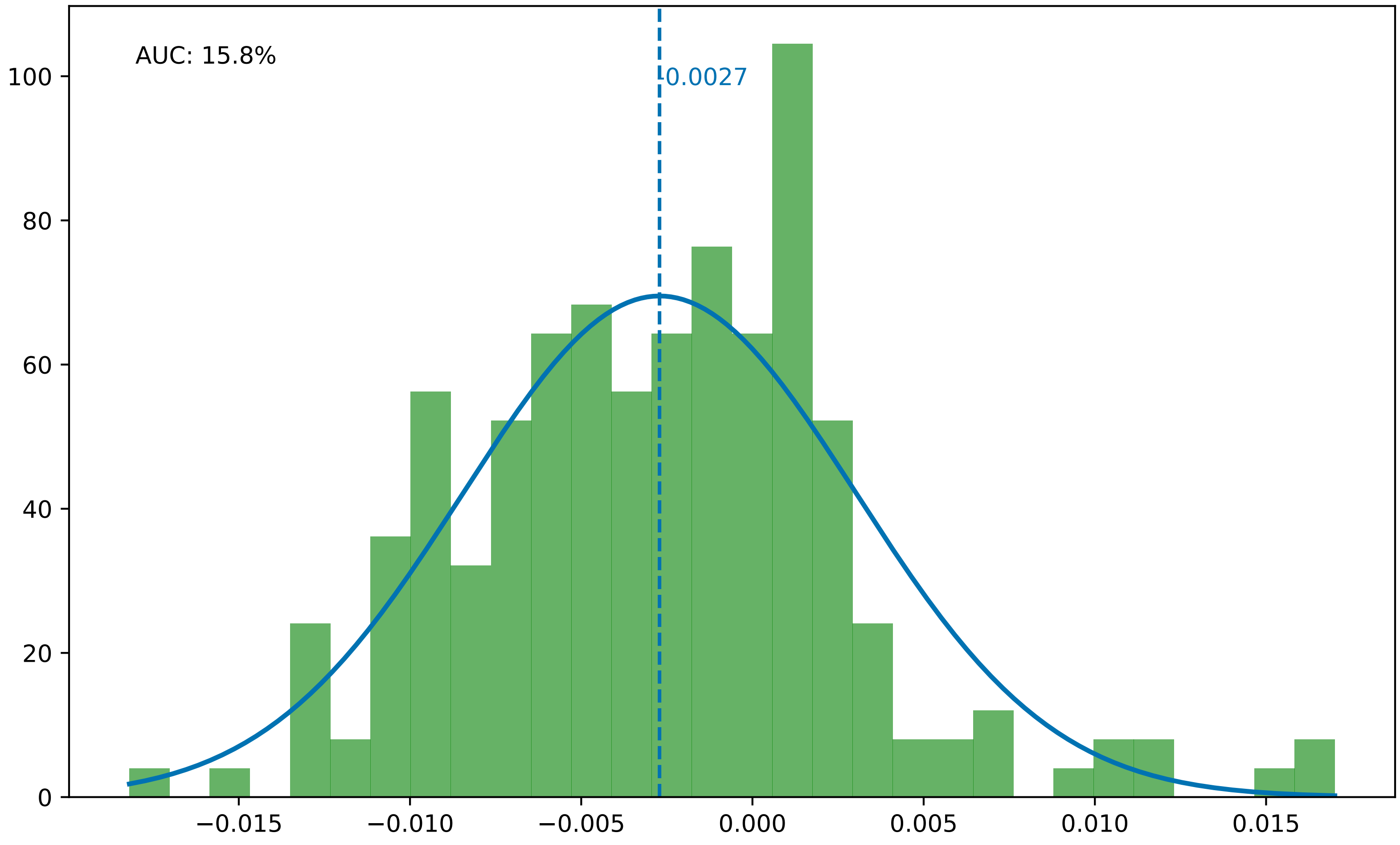

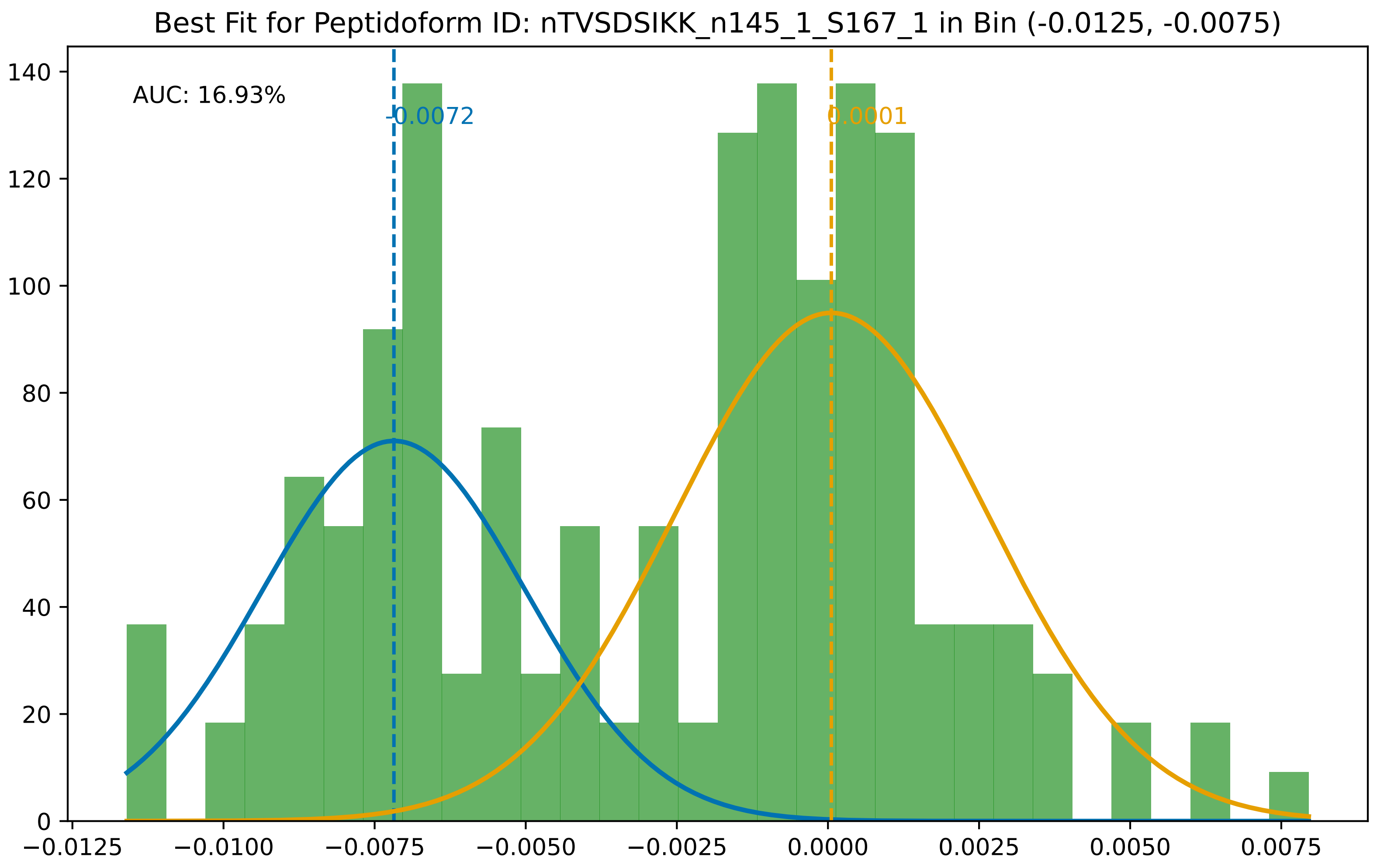

Best Fit for Peptidoform ID: nTVSDSIKK_n145_1_S167_1 in Bin (-0.0125, -0.0075)
AUC: 16.93%
-0.0072
0.0001
0
20
40
60
80
100
120
140
−0.0125
−0.0100
−0.0075
−0.0050
−0.0025
0.0000
0.0025
0.0050
0.0075

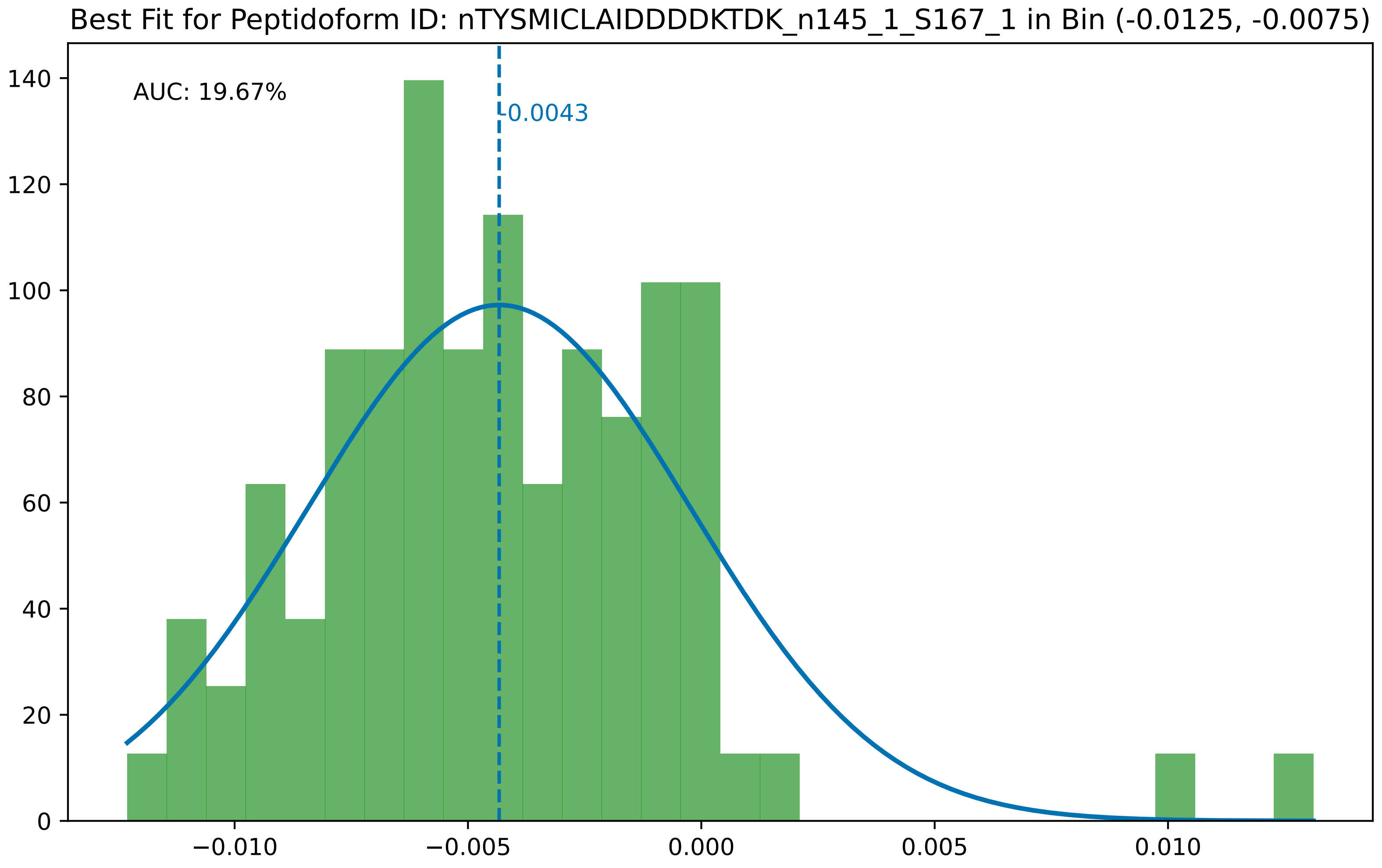

Best Fit for Peptidoform ID: nTYSMICLAIDDDDKTDK_n145_1_S167_1 in Bin (-0.0125, -0.0075)
AUC: 19.67%
-0.0043
0
20
40
60
80
100
120
140
−0.010
−0.005
0.000
0.005
0.010

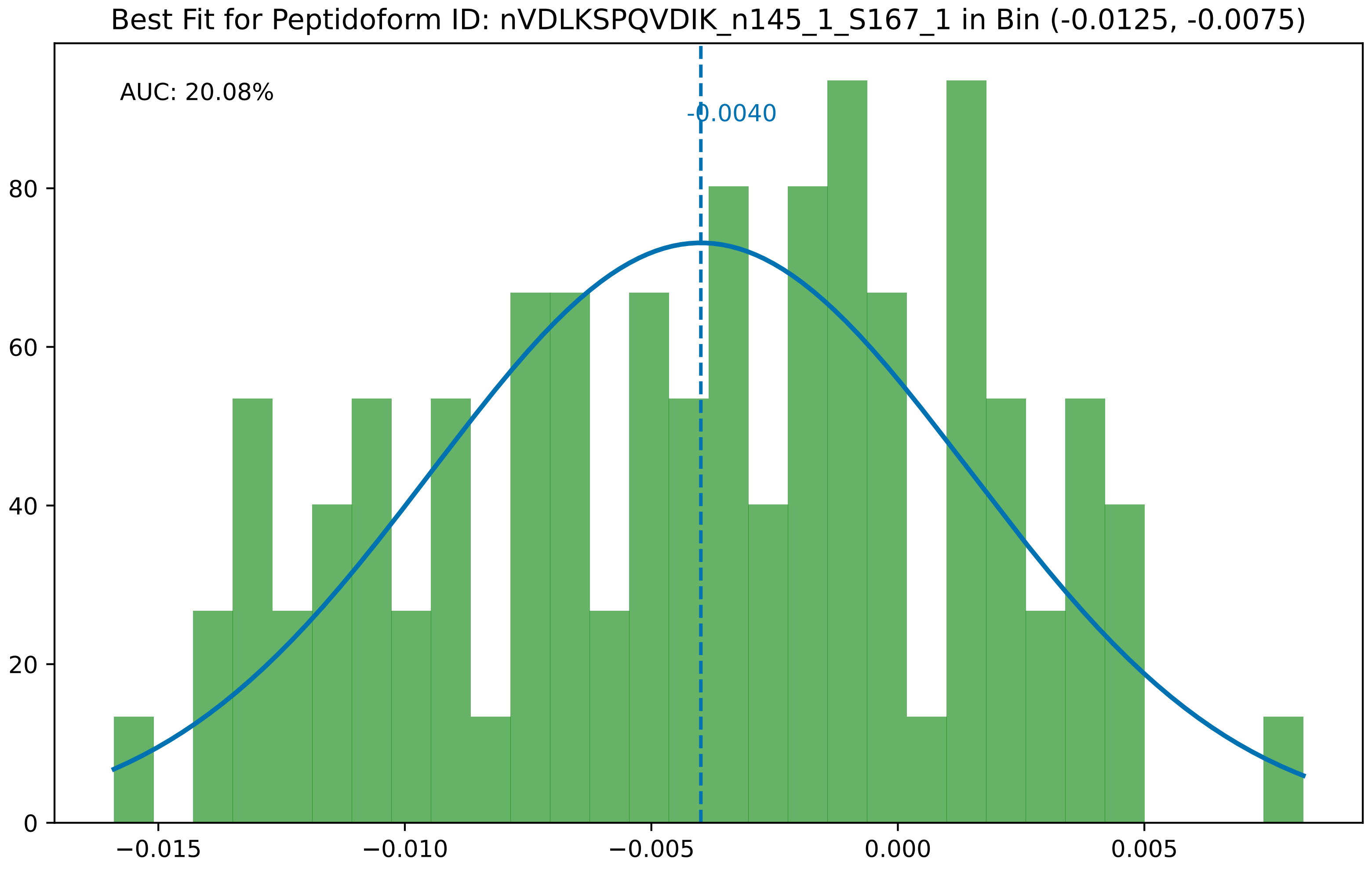
Best Fit for Peptidoform ID: nVDLKSPQVDIK_n145_1_S167_1 in Bin (-0.0125, -0.0075)
AUC: 20.08%
-0.0040
0
20
40
60
80
−0.015
−0.010
−0.005
0.000
0.005

Best Fit for Peptidoform ID: nVEMYSGSDDDDDFNKLPK_M147_1_n145_1_S167_1 in Bin (-0.0125, -0.0075)

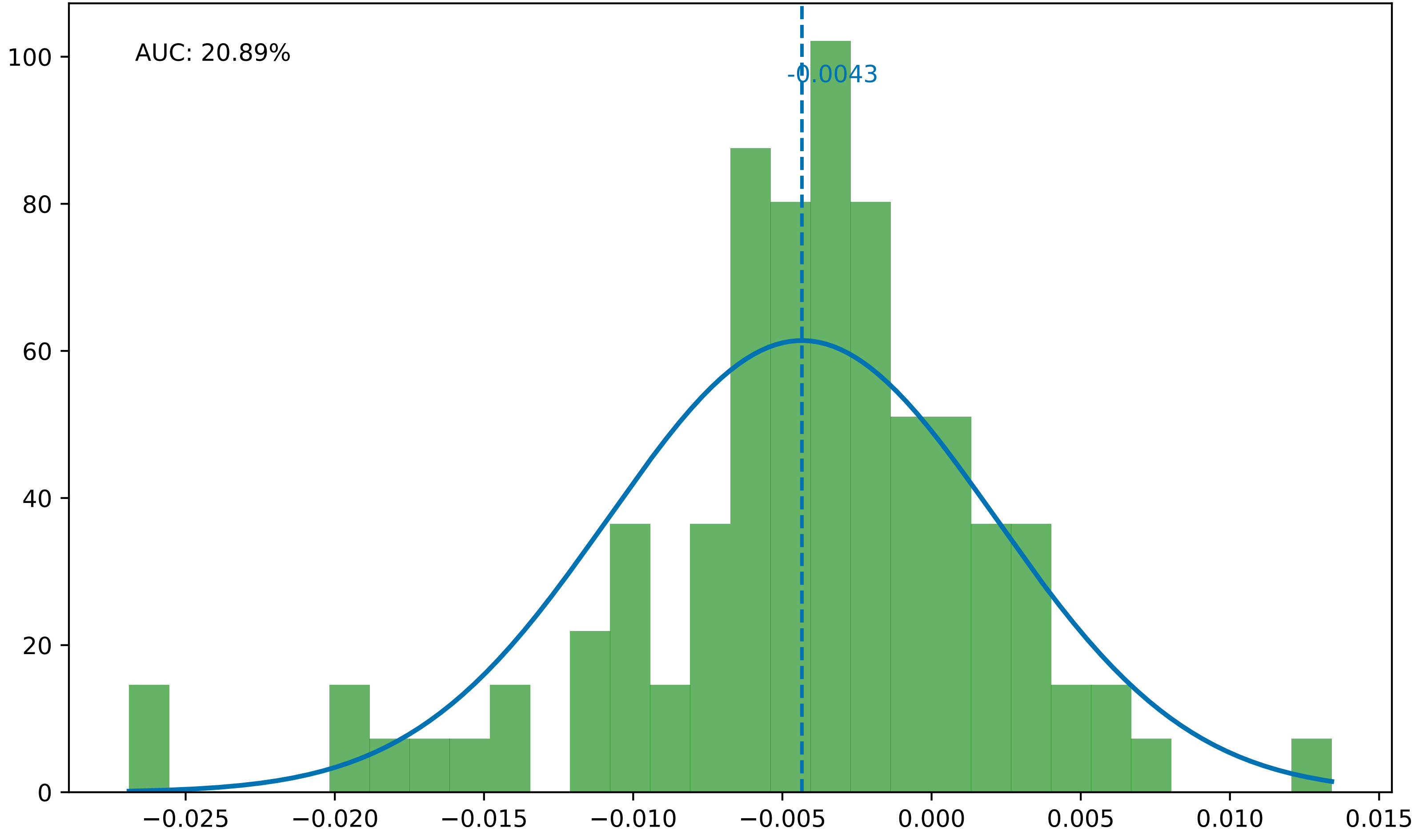

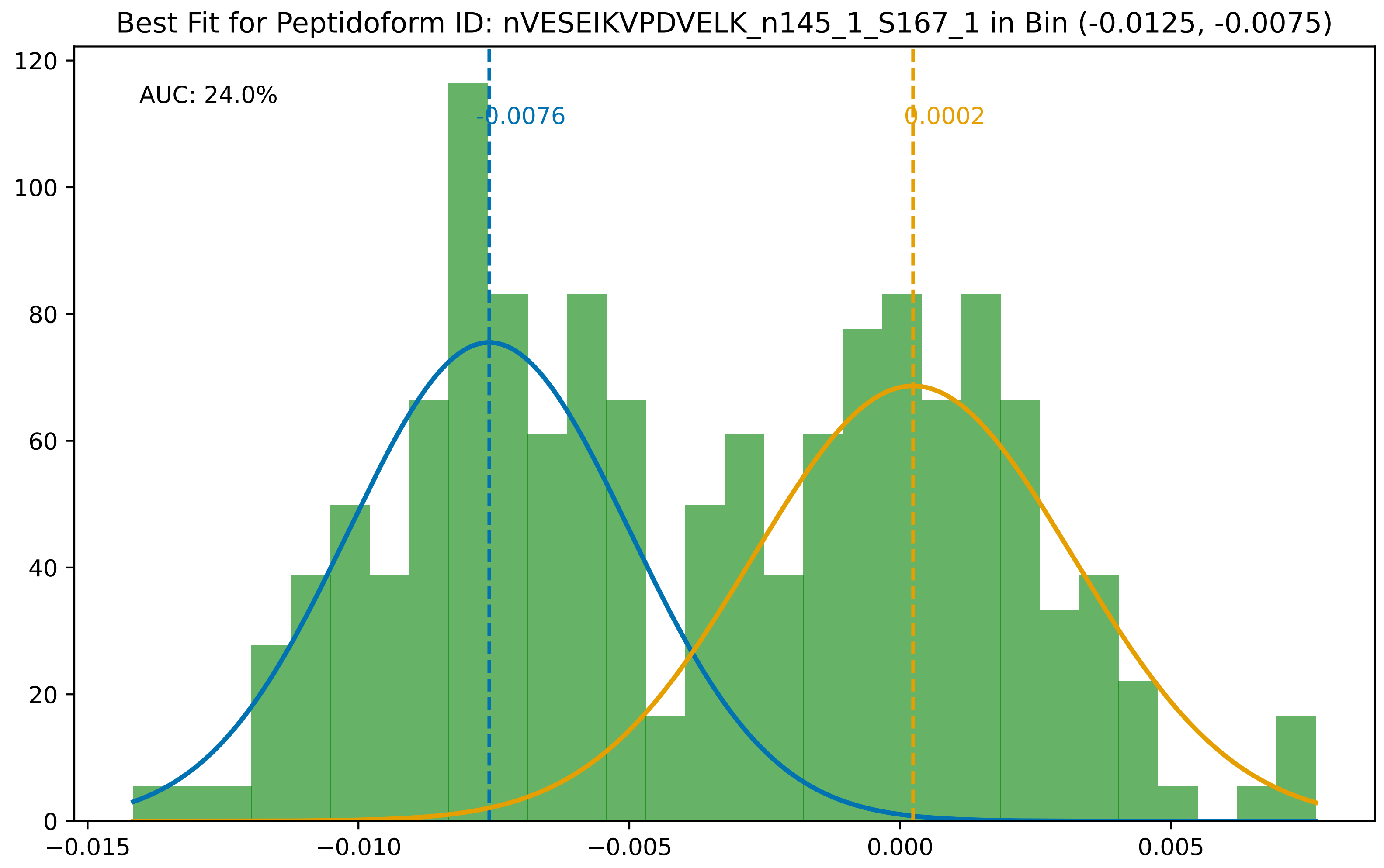
Best Fit for Peptidoform ID: nVESEIKVPDVELK_n145_1_S167_1 in Bin (-0.0125, -0.0075)
AUC: 24.0%
-0.0076
0.0002
0
20
40
60
80
100
120
−0.015
−0.010
−0.005
0.000
0.005

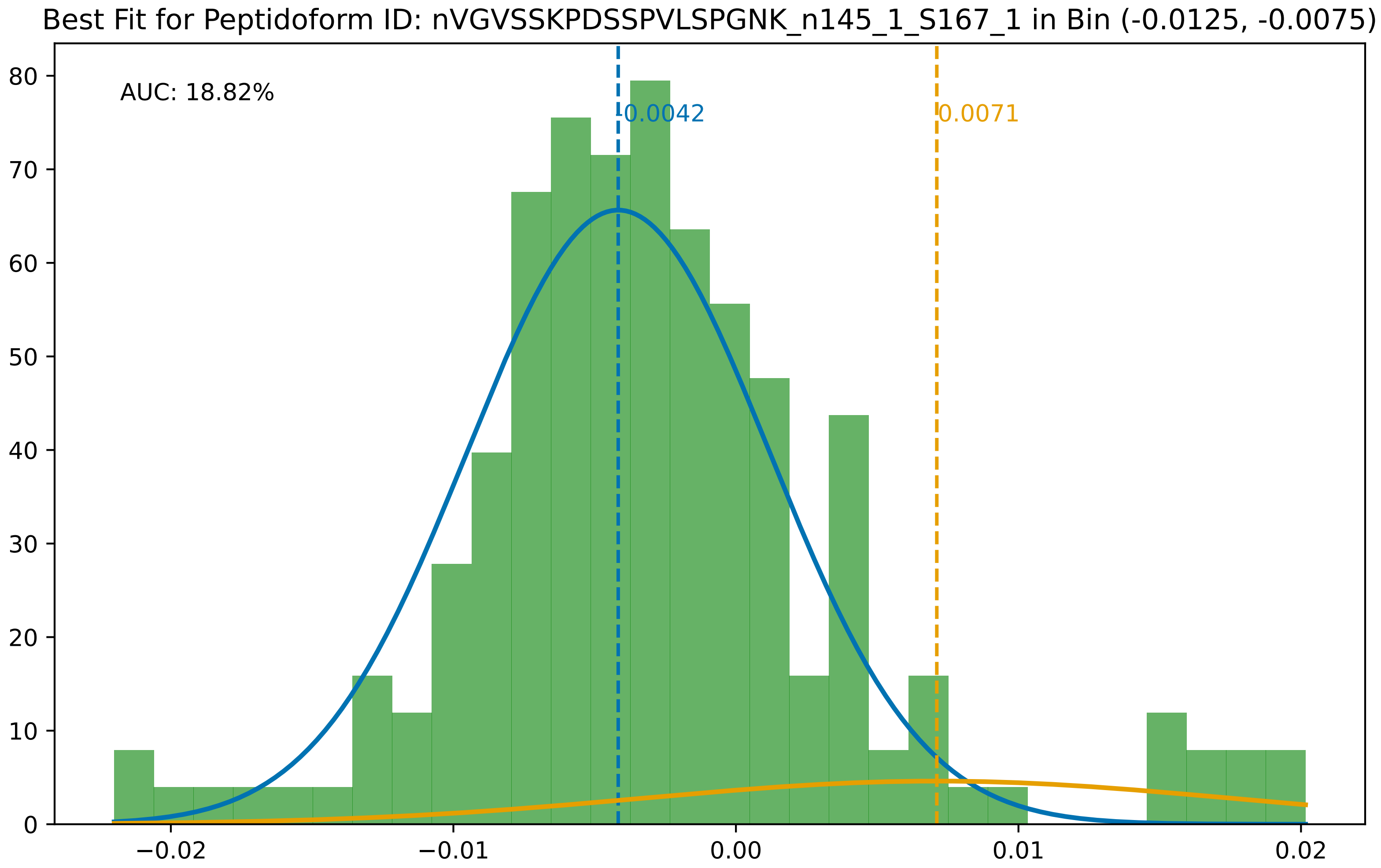
Best Fit for Peptidoform ID: nVGVSSKPDSSPVLSPGNK_n145_1_S167_1 in Bin (-0.0125, -0.0075)
AUC: 18.82%
0.0042
0.0071
80
70
60
50
40
30
20
10
0
−0.02
−0.01
0.00
0.01
0.02

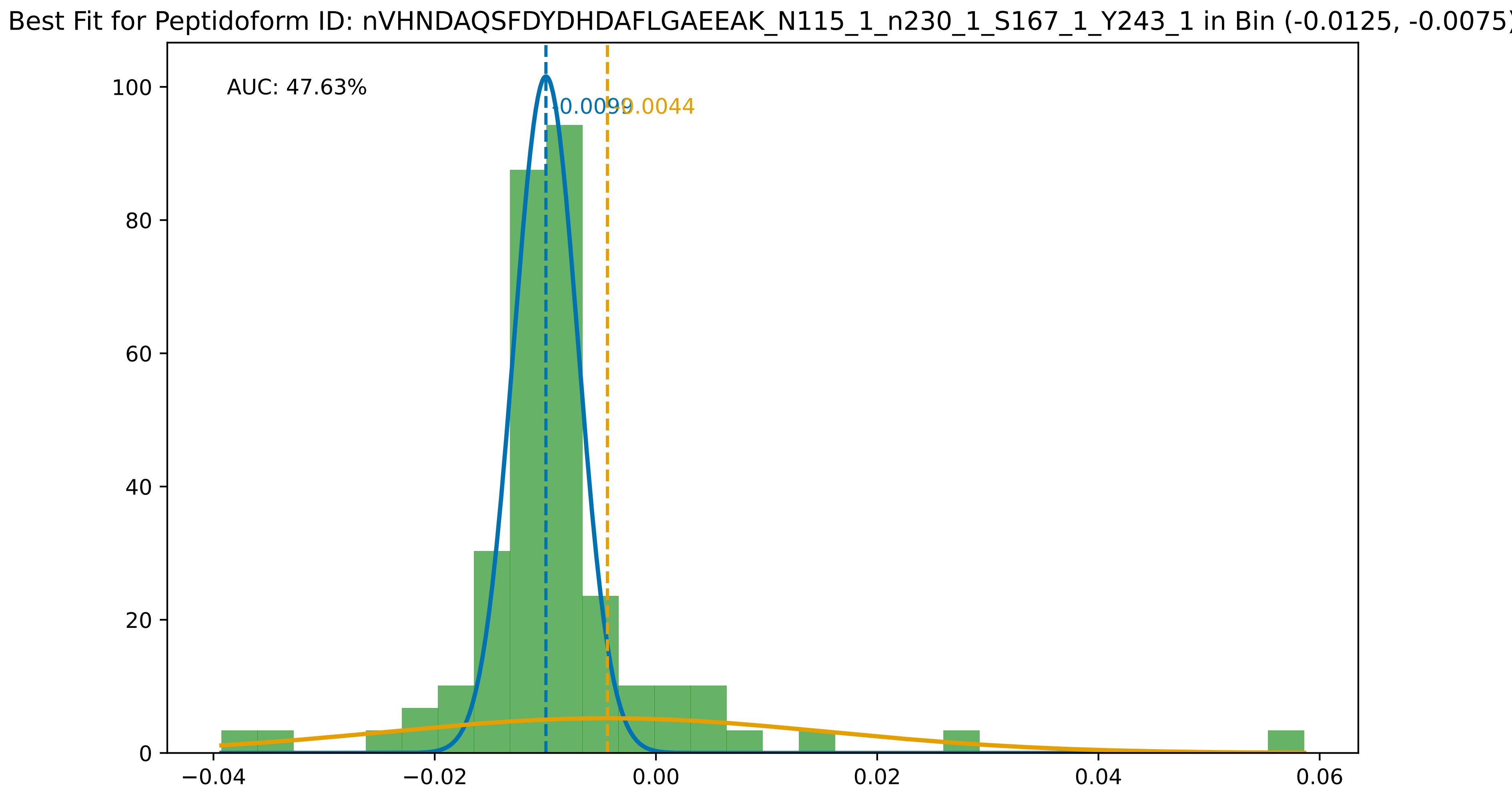
Best Fit for Peptidoform ID: nVHNDAQSFDYDHDAFLGAEEAK_N115_1_n230_1_S167_1_Y243_1 in Bin (-0.0125, -0.0075)
AUC: 47.63%
-0.0099
-0.0044
0
20
40
60
80
100
−0.04
−0.02
0.00
0.02
0.04
0.06

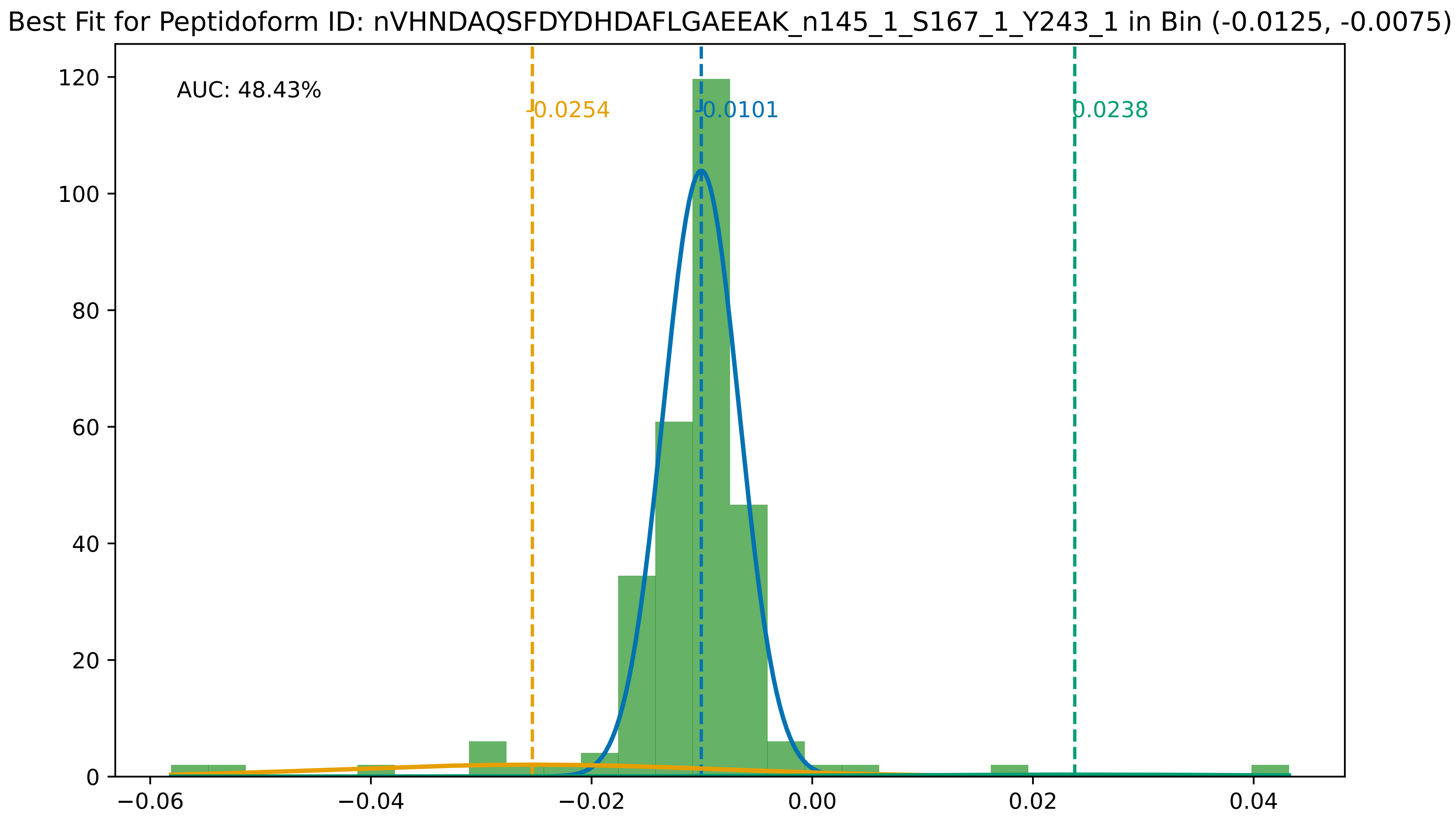

Best Fit for Peptidoform ID: nVHNDAQSFDYDHDAFLGAEEAK_n145_1_S167_1_Y243_1 in Bin (-0.0125, -0.0075)
AUC: 48.43%
-0.0254
-0.0101
0.0238
0
20
40
60
80
100
120
−0.06
−0.04
−0.02
0.00
0.02
0.04

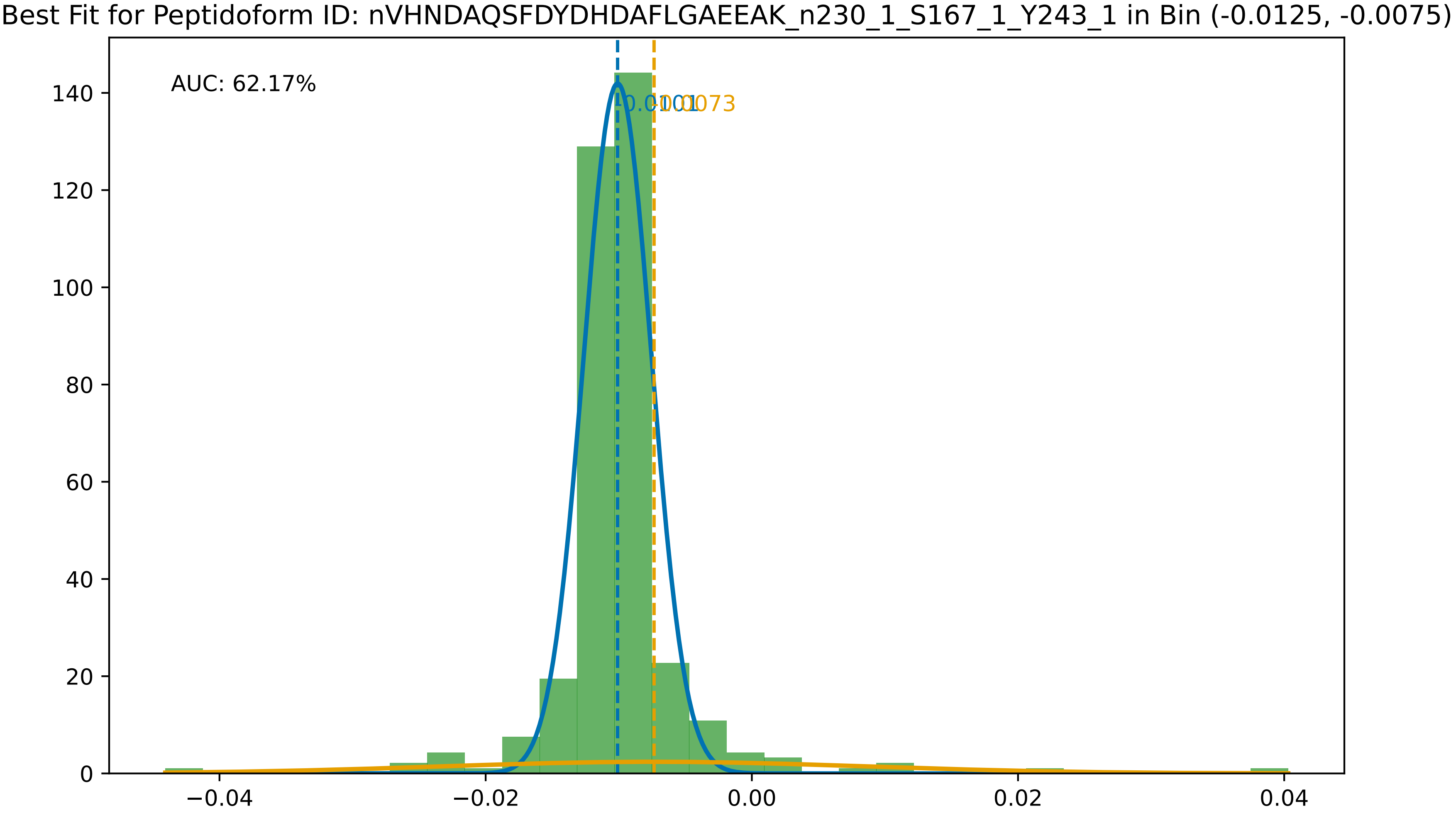

Best Fit for Peptidoform ID: nVHNDAQSFDYDHDAFLGAEEAK_n230_1_S167_1_Y243_1 in Bin (-0.0125, -0.0075)
AUC: 62.17%
-0.0101
-0.0073
0
20
40
60
80
100
120
140
−0.04
−0.02
0.00
0.02
0.04

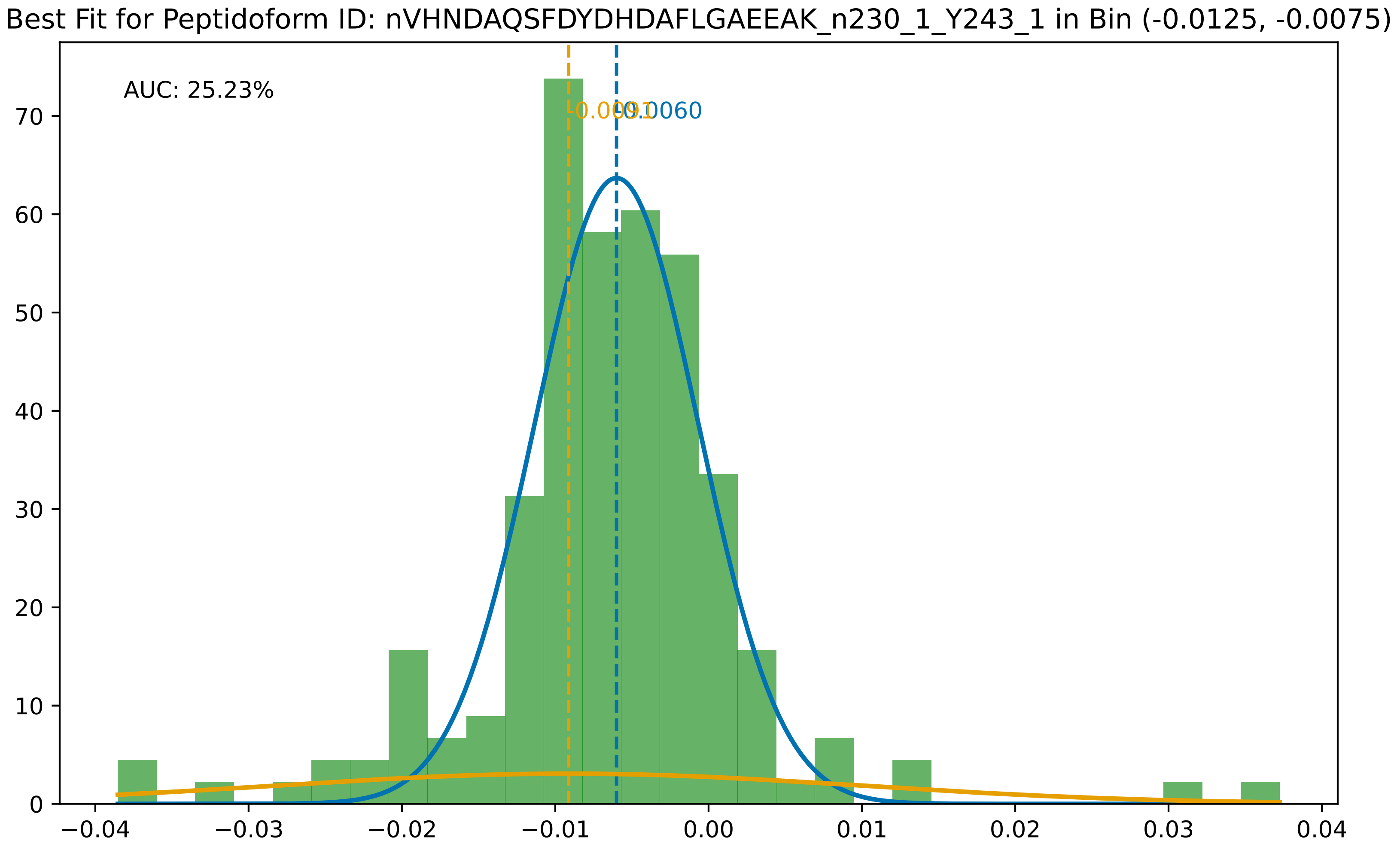
Best Fit for Peptidoform ID: nVHNDAQSFDYDHDAFLGAEEAK_n230_1_Y243_1 in Bin (-0.0125, -0.0075)
AUC: 25.23%
0
10
20
30
40
50
60
70
−0.04
−0.03
−0.02
−0.01
0.00
0.01
0.02
0.03
0.04

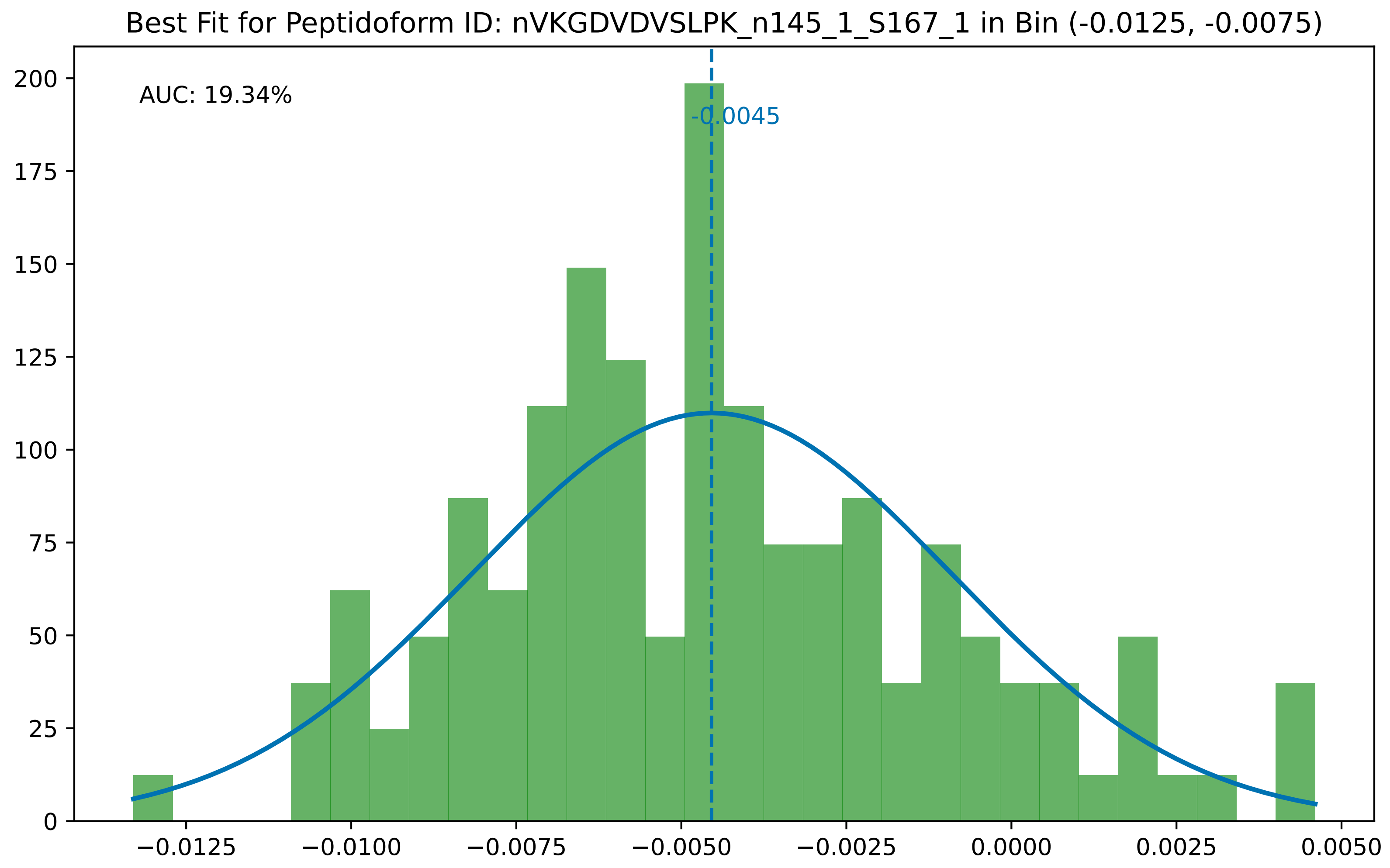

Best Fit for Peptidoform ID: nVKGDVDVSLPK_n145_1_S167_1 in Bin (-0.0125, -0.0075)
AUC: 19.34%
-0.0045
0
25
50
75
100
125
150
175
200
−0.0125
−0.0100
−0.0075
−0.0050
−0.0025
0.0000
0.0025
0.0050

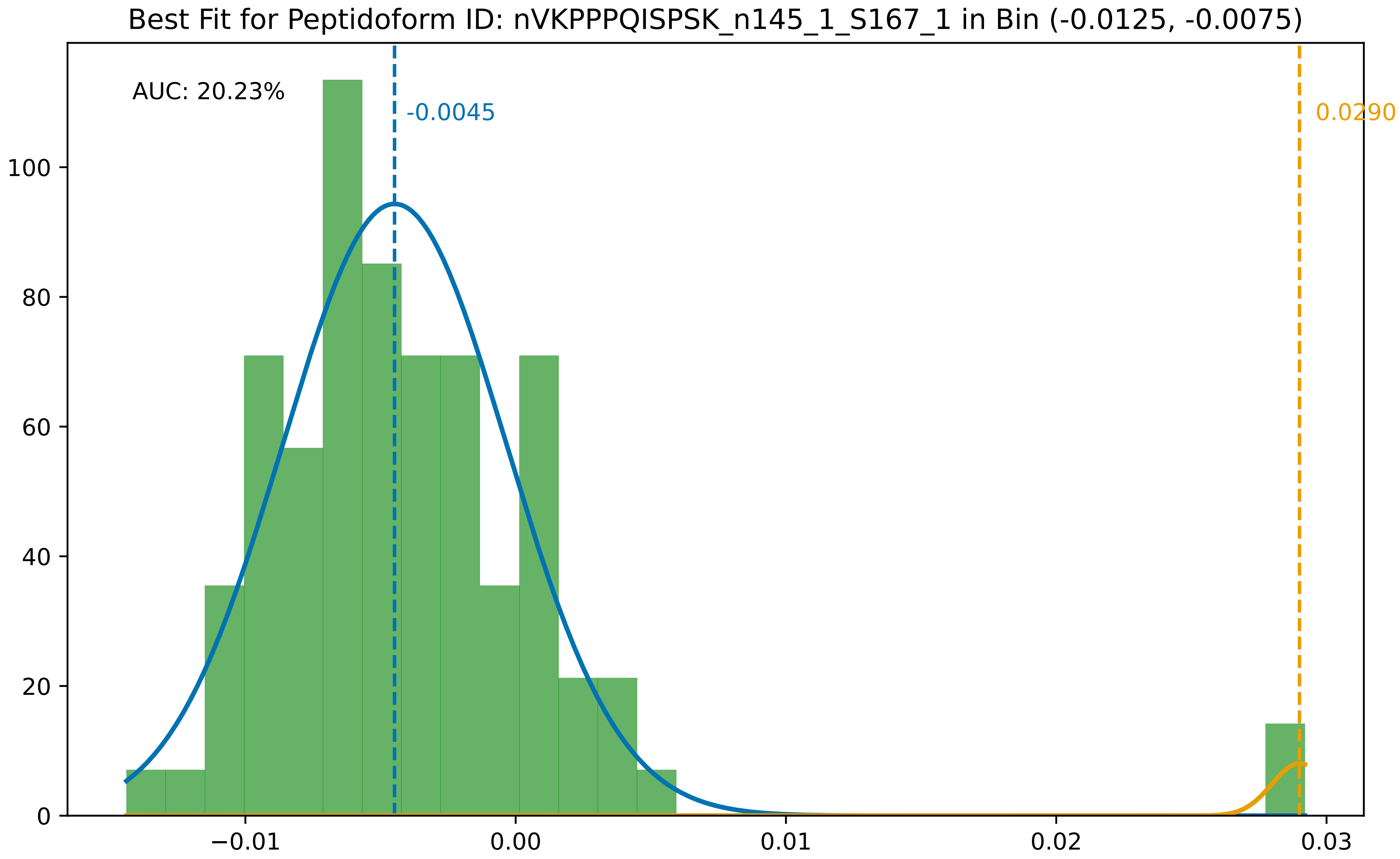
Best Fit for Peptidoform ID: nVKPPPQISPSK_n145_1_S167_1 in Bin (-0.0125, -0.0075)
AUC: 20.23%
-0.0045
0.0290
0
20
40
60
80
100
−0.01
0.00
0.01
0.02
0.03

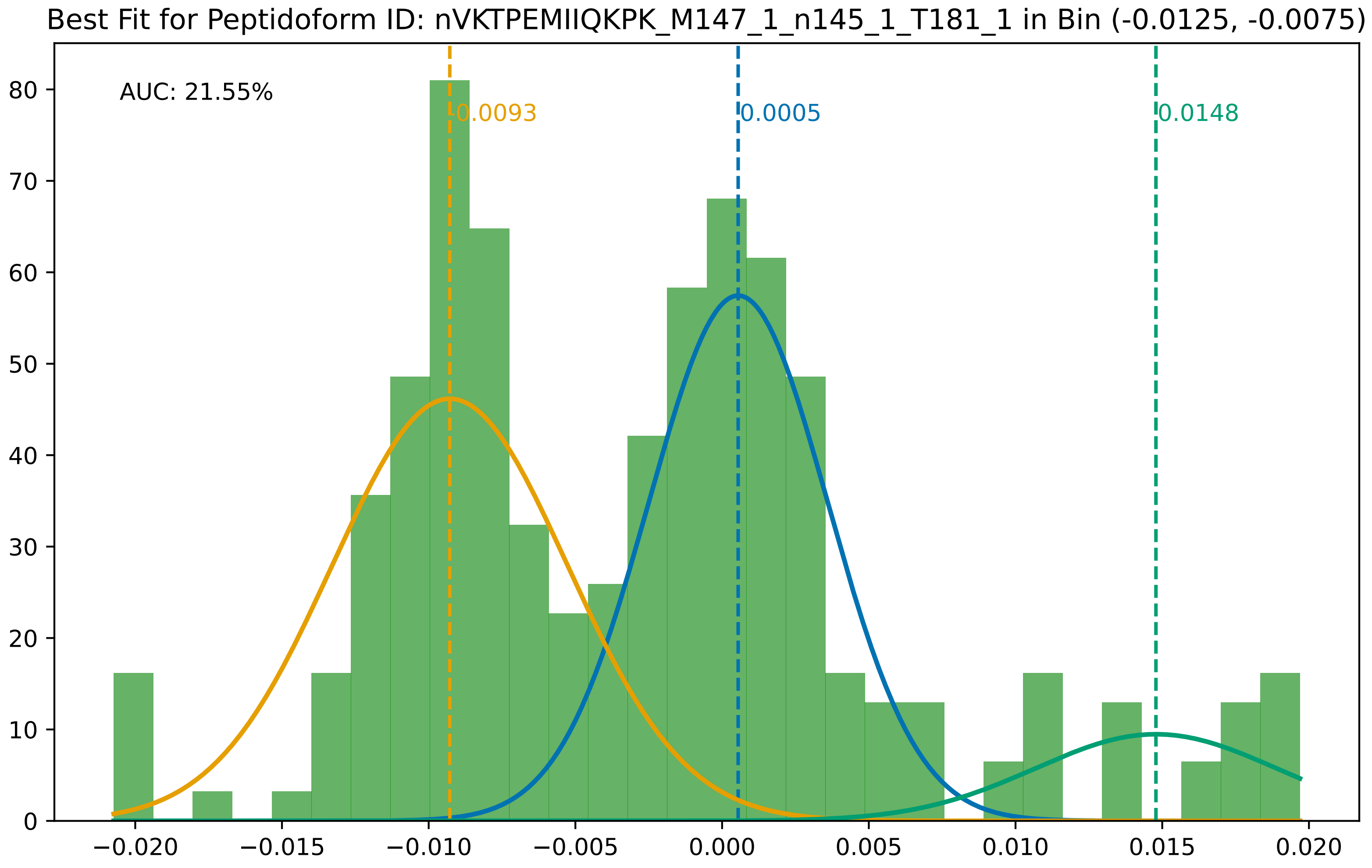
Best Fit for Peptidoform ID: nVKTPEMIIQKPK_M147_1_n145_1_T181_1 in Bin (-0.0125, -0.0075)
AUC: 21.55%
0.0093
0.0005
0.0148
0
10
20
30
40
50
60
70
80
−0.020
−0.015
−0.010
−0.005
0.000
0.005
0.010
0.015
0.020

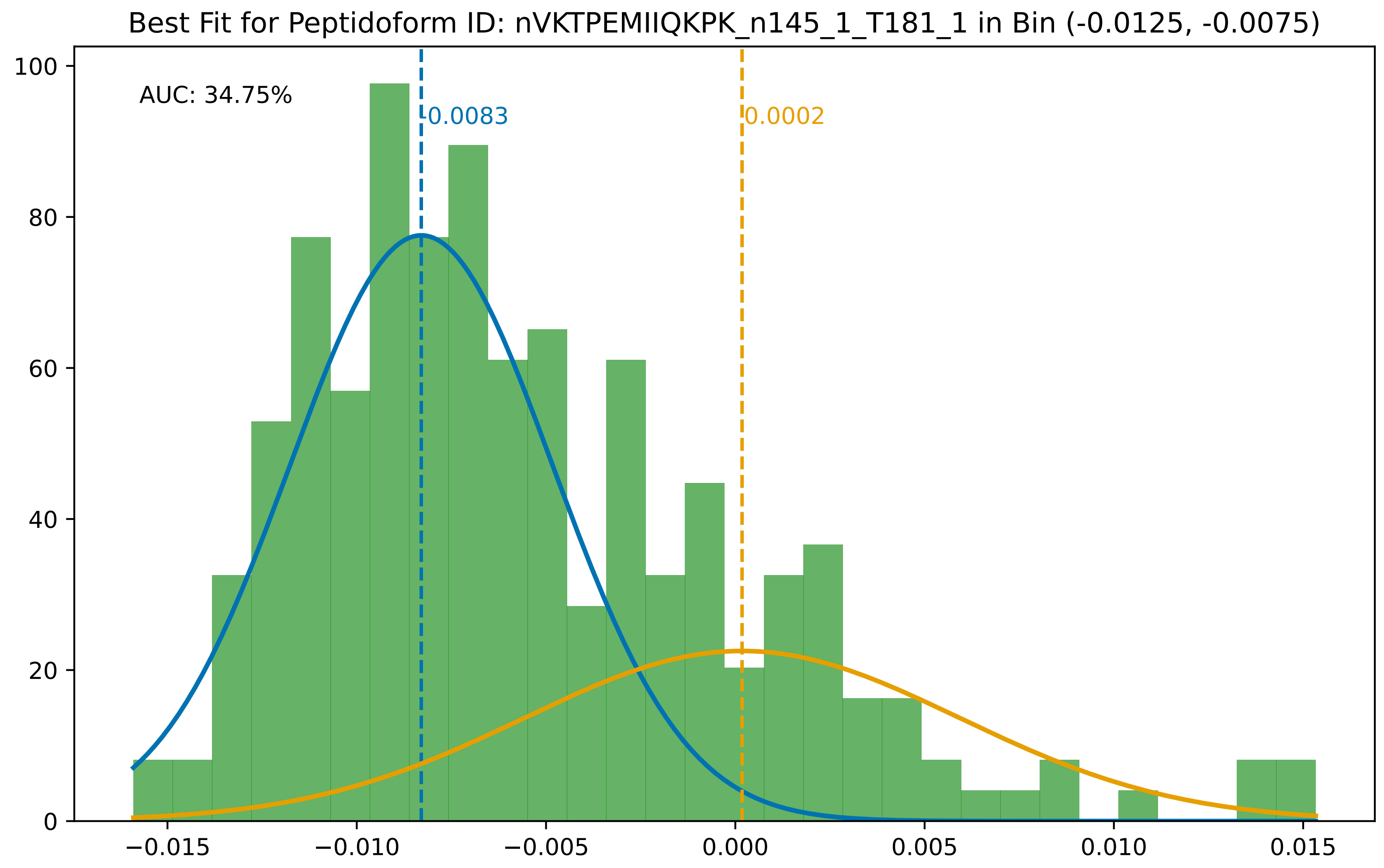
Best Fit for Peptidoform ID: nVKTPEMIIQKPK_n145_1_T181_1 in Bin (-0.0125, -0.0075)
AUC: 34.75%
0.0083
0.0002
100
80
60
40
20
0
−0.015
−0.010
−0.005
0.000
0.005
0.010
0.015

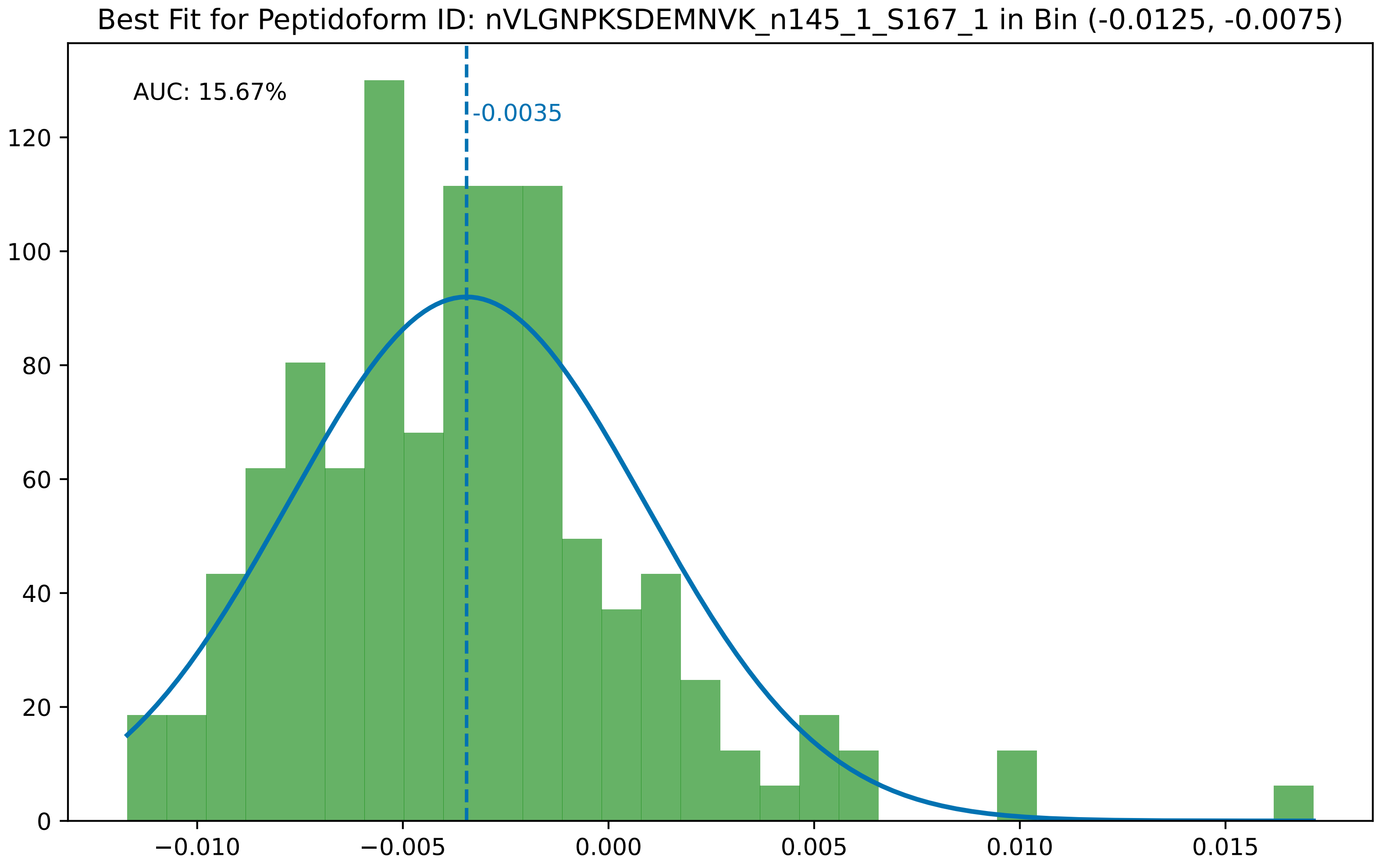
Best Fit for Peptidoform ID: nVLGNPKSDEMNVK_n145_1_S167_1 in Bin (-0.0125, -0.0075)
AUC: 15.67%
-0.0035
0
20
40
60
80
100
120
−0.010
−0.005
0.000
0.005
0.010
0.015

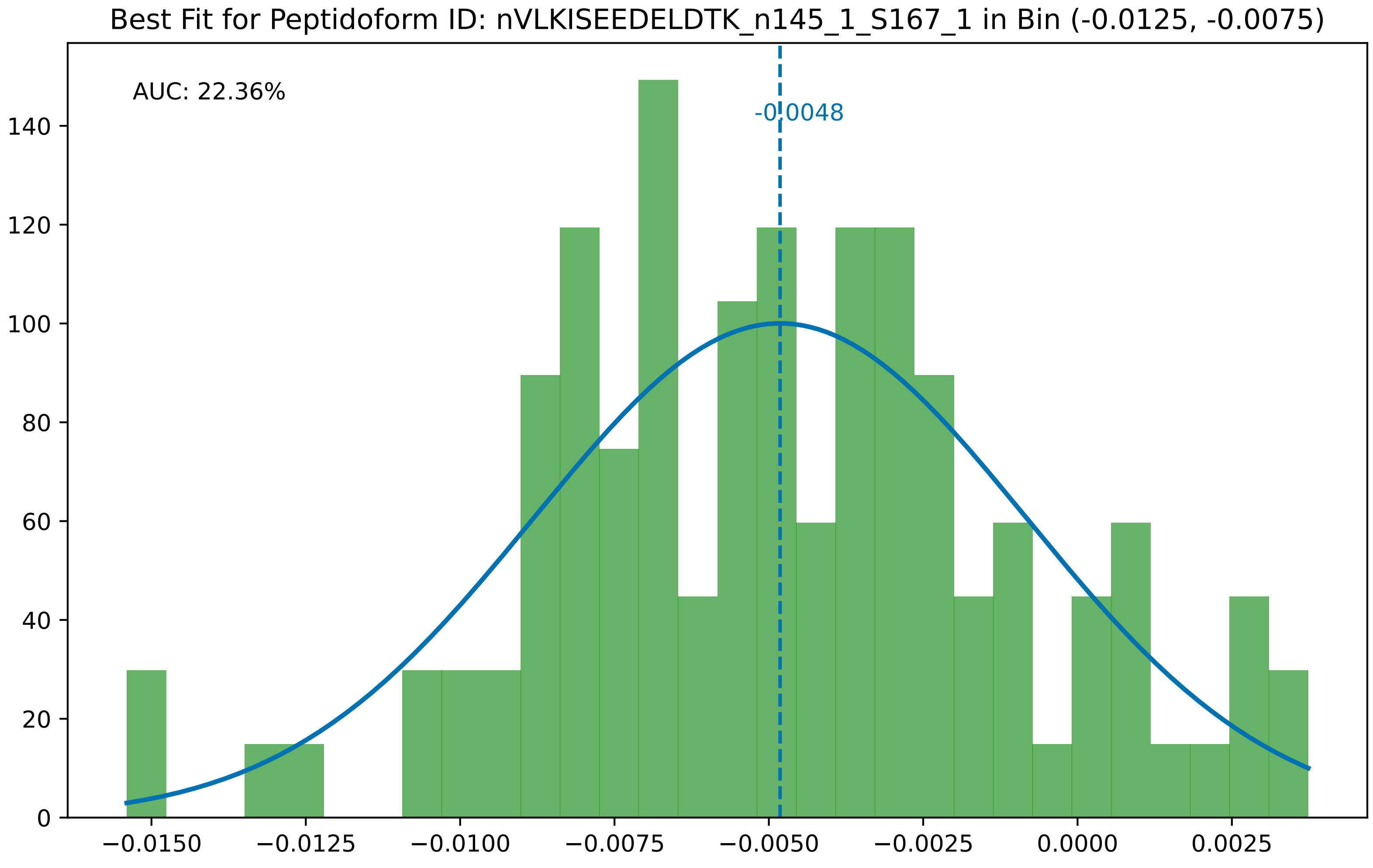

Best Fit for Peptidoform ID: nVLKISEEDELDTK_n145_1_S167_1 in Bin (-0.0125, -0.0075)
AUC: 22.36%
-0.0048
0
20
40
60
80
100
120
140
−0.0150
−0.0125
−0.0100
−0.0075
−0.0050
−0.0025
0.0000
0.0025

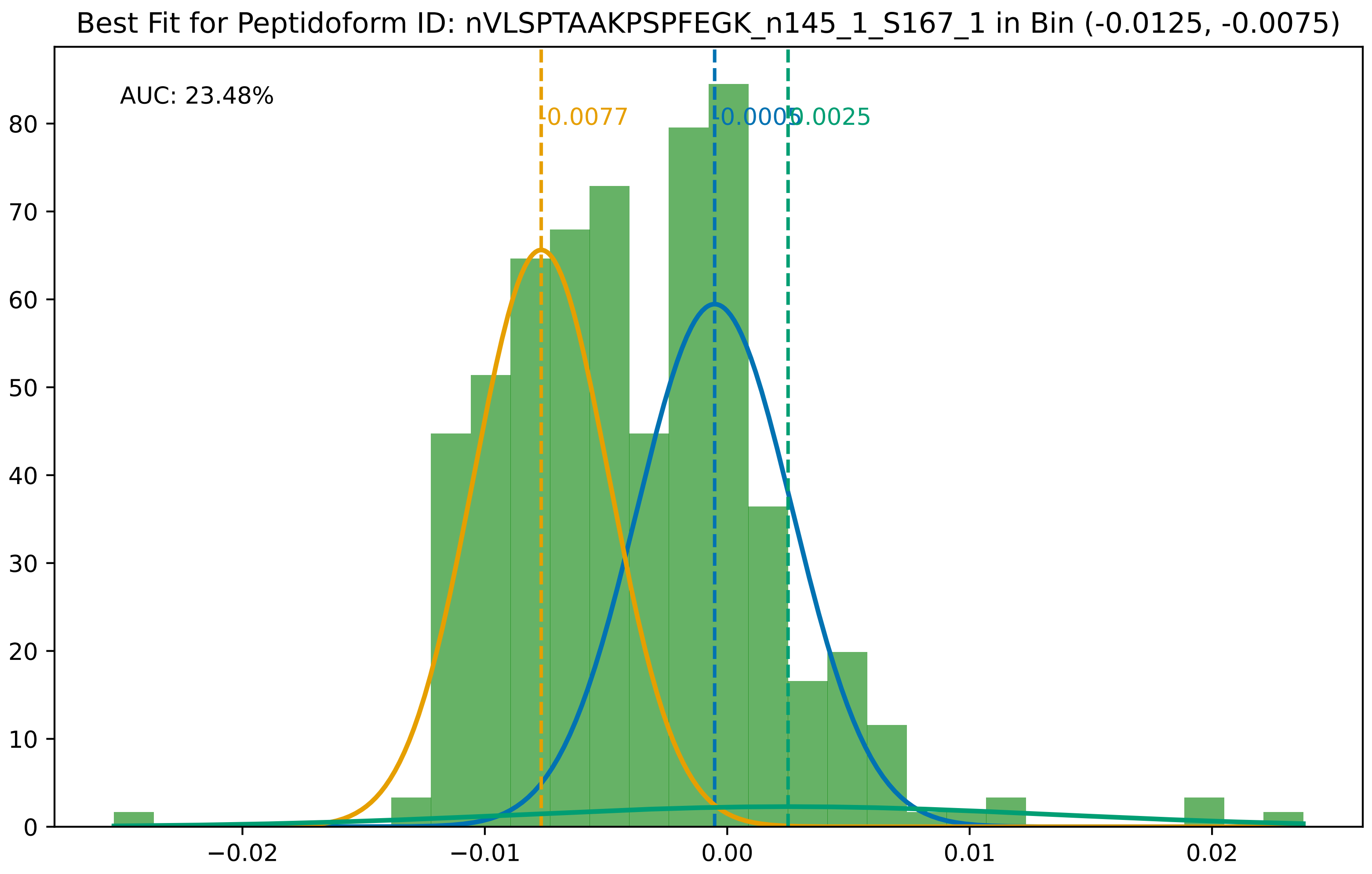

Best Fit for Peptidoform ID: nVLSPTAAKPSPFEGK_n145_1_S167_1 in Bin (-0.0125, -0.0075)
AUC: 23.48%
-0.0077
-0.0005
0.0025
0
10
20
30
40
50
60
70
80
−0.02
−0.01
0.00
0.01
0.02

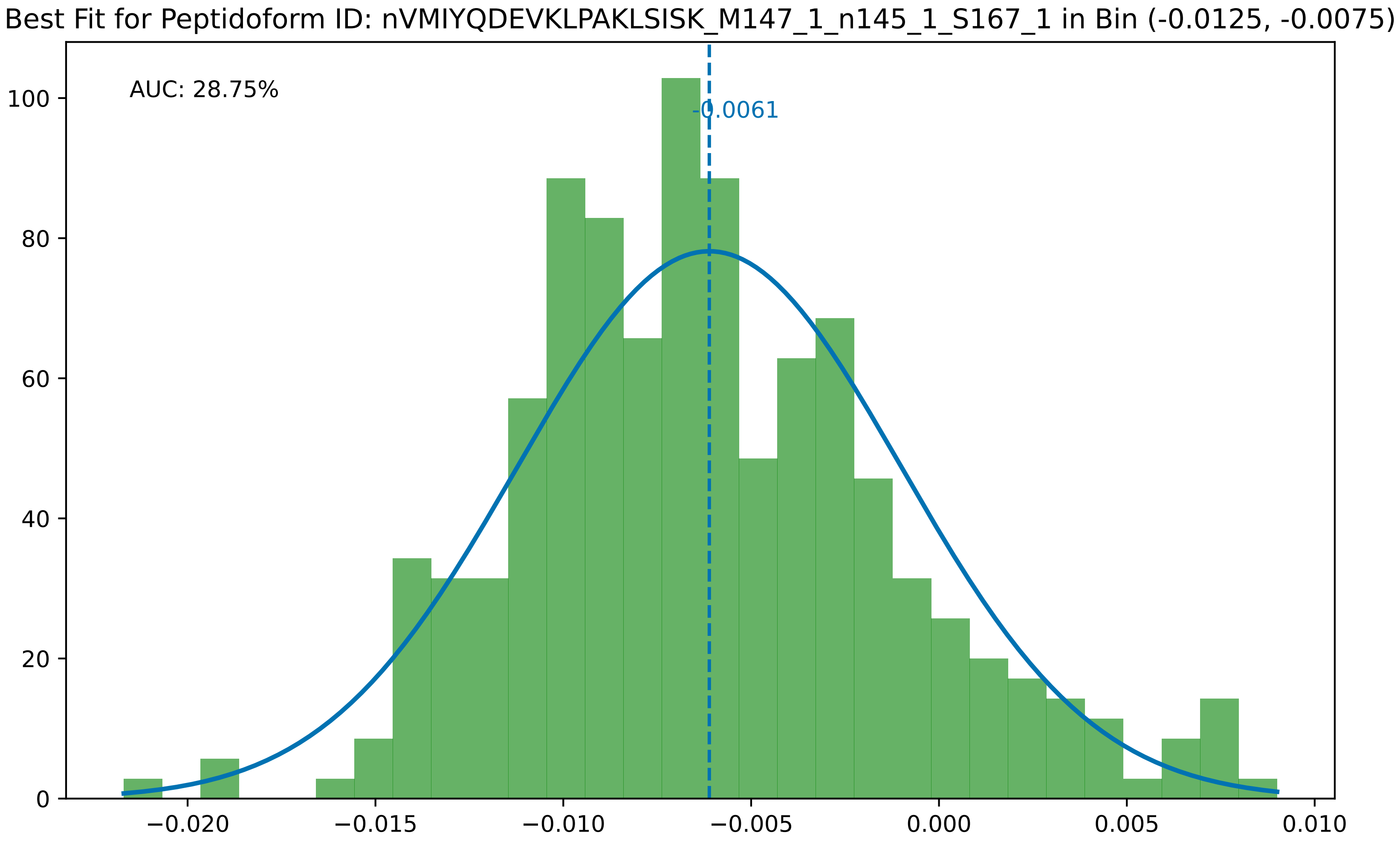

Best Fit for Peptidoform ID: nVMIYQDEVKLPAKLSISK_M147_1_n145_1_S167_1 in Bin (-0.0125, -0.0075)
AUC: 28.75%
-0.0061
0
20
40
60
80
100
−0.020
−0.015
−0.010
−0.005
0.000
0.005
0.010

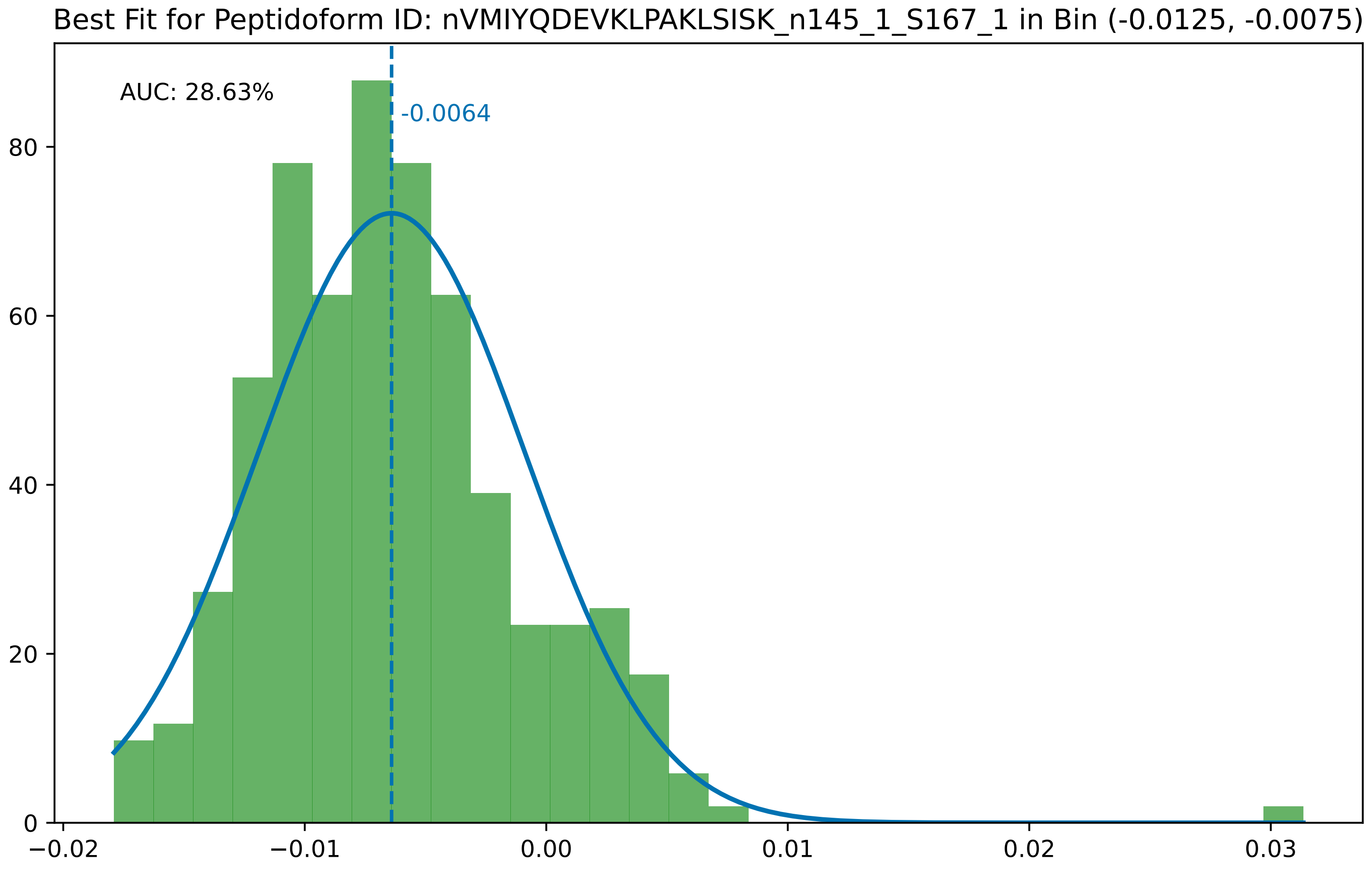

Best Fit for Peptidoform ID: nVMIYQDEVKLPAKLSISK_n145_1_S167_1 in Bin (-0.0125, -0.0075)
AUC: 28.63%
-0.0064
0
20
40
60
80
−0.02
−0.01
0.00
0.01
0.02
0.03

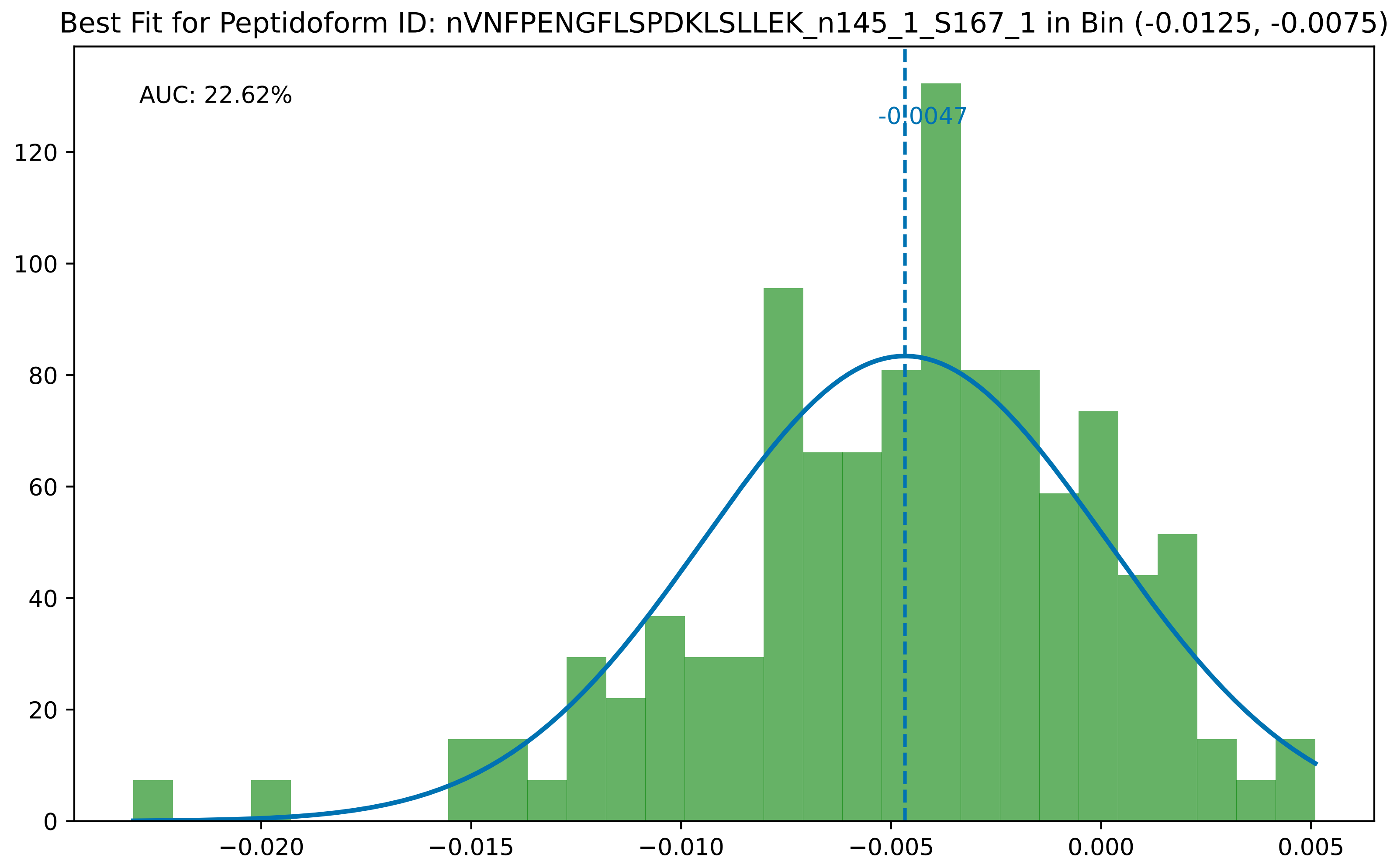

Best Fit for Peptidoform ID: nVNFPENGFLSPDKLSLLEK_n145_1_S167_1 in Bin (-0.0125, -0.0075)
AUC: 22.62%
-0.0047
0
20
40
60
80
100
120
−0.020
−0.015
−0.010
−0.005
0.000
0.005

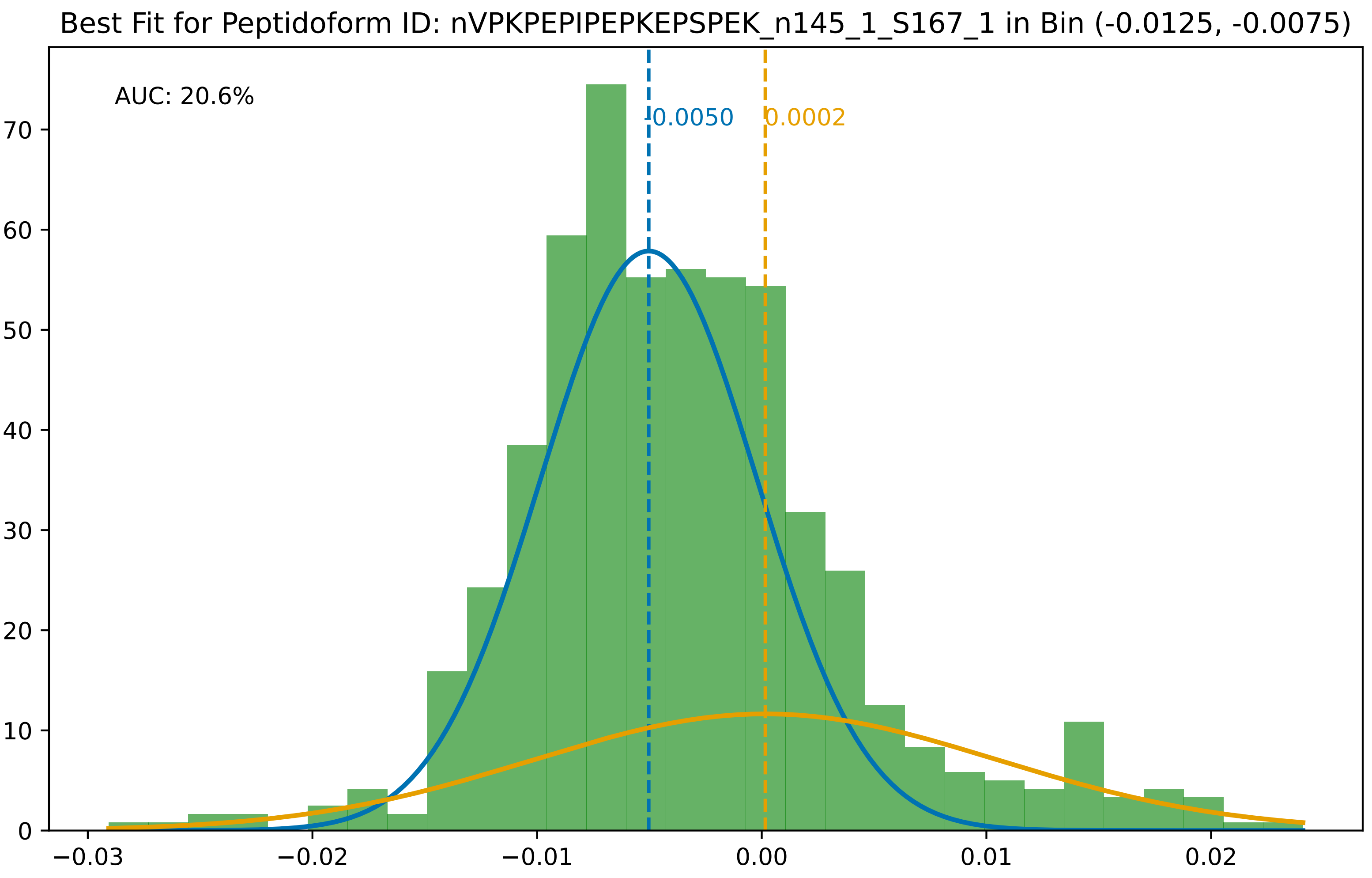
Best Fit for Peptidoform ID: nVPKPEPIPEPKEPSPEK_n145_1_S167_1 in Bin (-0.0125, -0.0075)
AUC: 20.6%
-0.0050
0.0002
0
10
20
30
40
50
60
70
−0.03
−0.02
−0.01
0.00
0.01
0.02

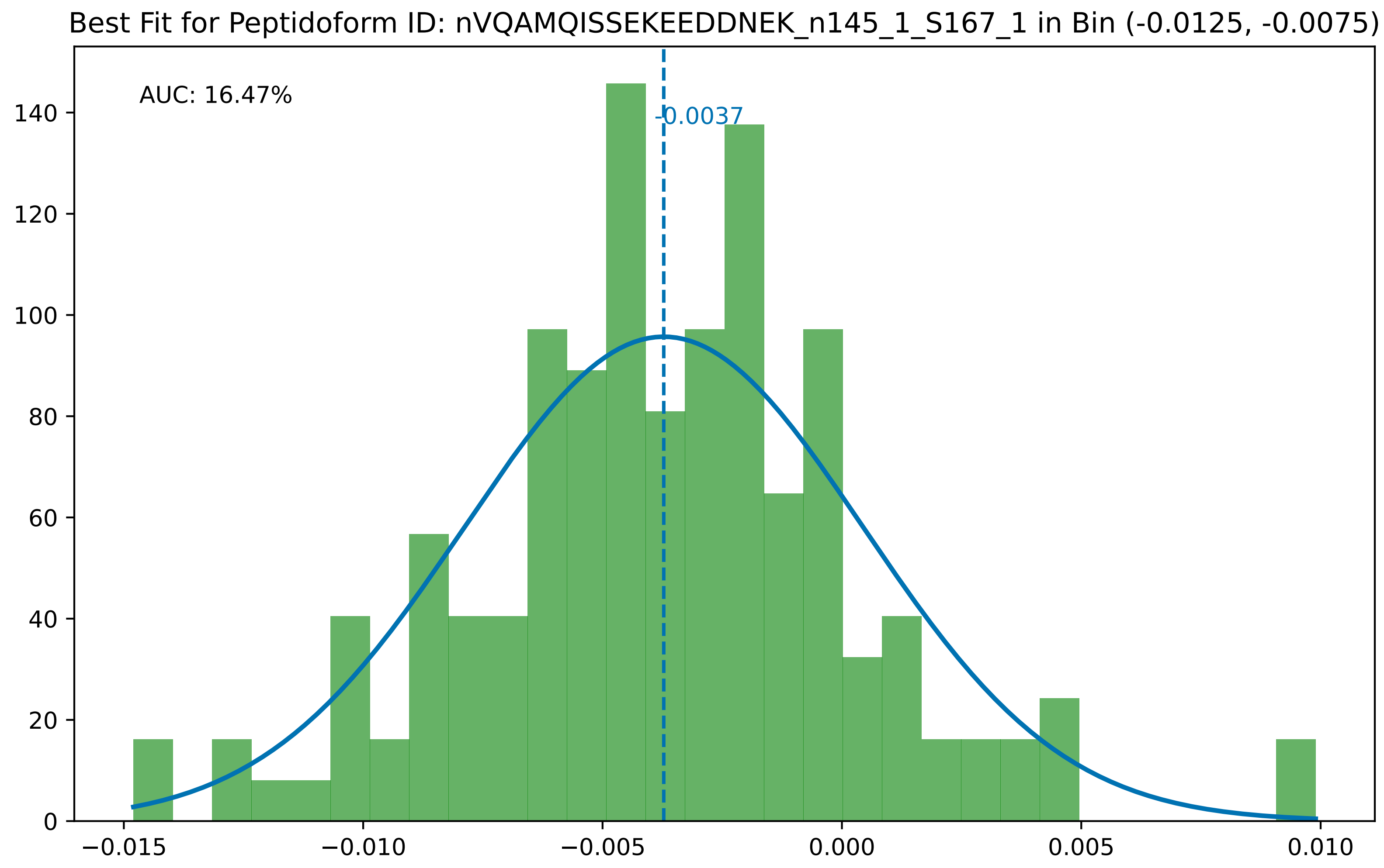

Best Fit for Peptidoform ID: nVQAMQISSEKEEDDNEK_n145_1_S167_1 in Bin (-0.0125, -0.0075)
AUC: 16.47%
-0.0037
0
20
40
60
80
100
120
140
−0.015
−0.010
−0.005
0.000
0.005
0.010

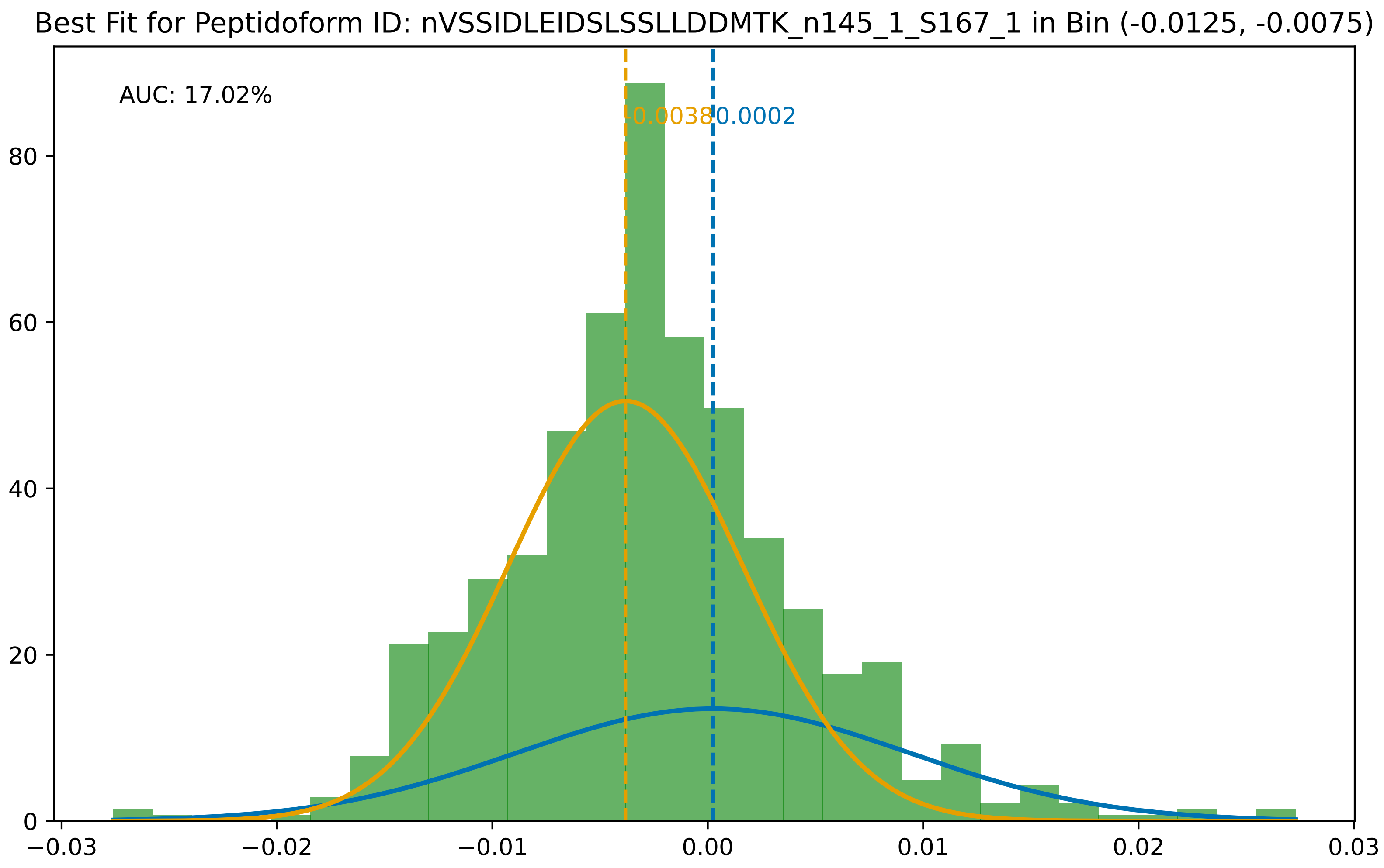
Best Fit for Peptidoform ID: nVSSIDLEIDSLSSLLDDMTK_n145_1_S167_1 in Bin (-0.0125, -0.0075)
AUC: 17.02%
-0.0038
0.0002
0
20
40
60
80
−0.03
−0.02
−0.01
0.00
0.01
0.02
0.03

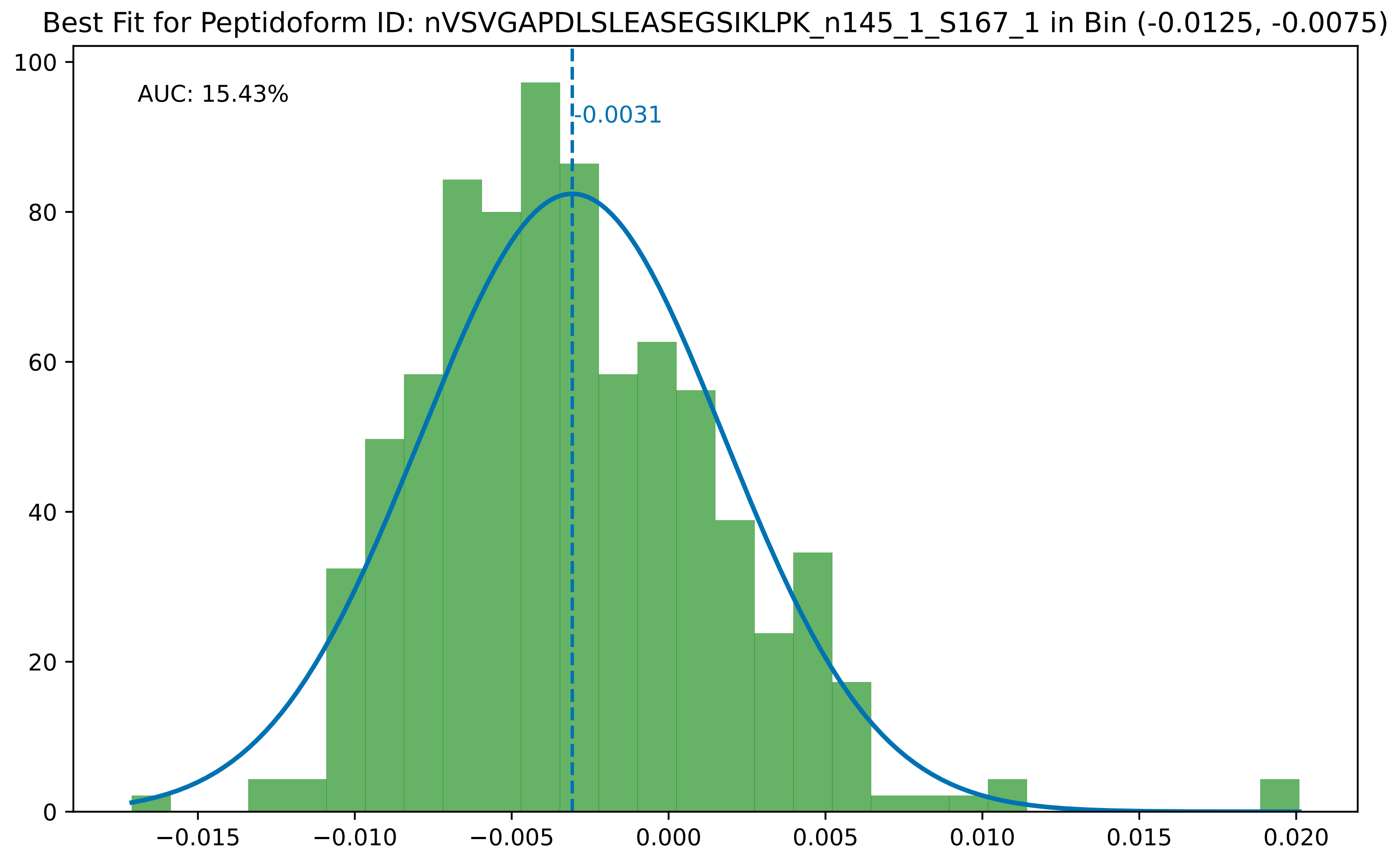

Best Fit for Peptidoform ID: nVSVGAPDLSLEASEGSIKLPK_n145_1_S167_1 in Bin (-0.0125, -0.0075)
AUC: 15.43%
-0.0031
100
80
60
40
20
0
−0.015
−0.010
−0.005
0.000
0.005
0.010
0.015
0.020

Best Fit for Peptidoform ID: nVVEAVNSDSDSEFGIPKK_n145_1_S167_2 in Bin (-0.0125, -0.0075)

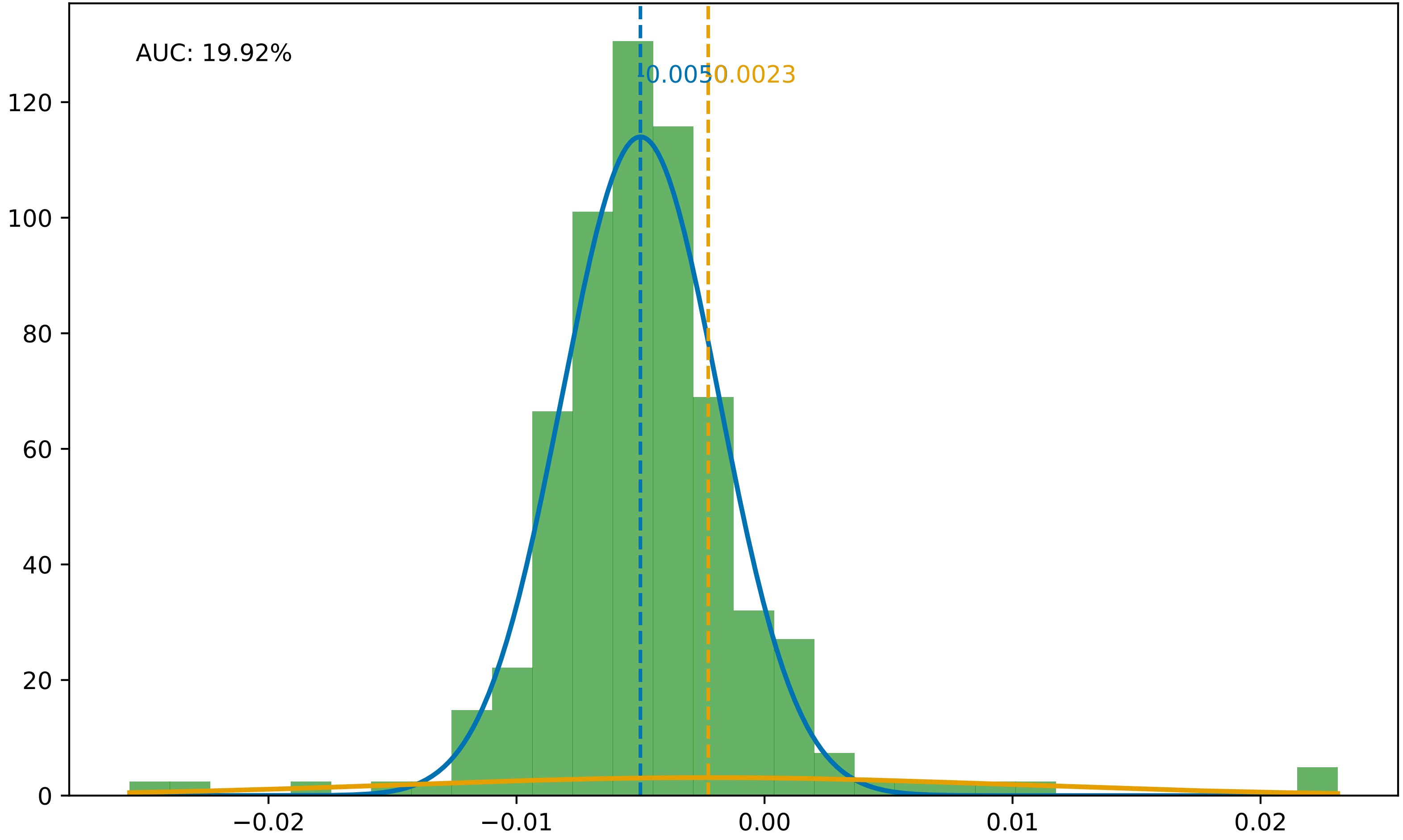

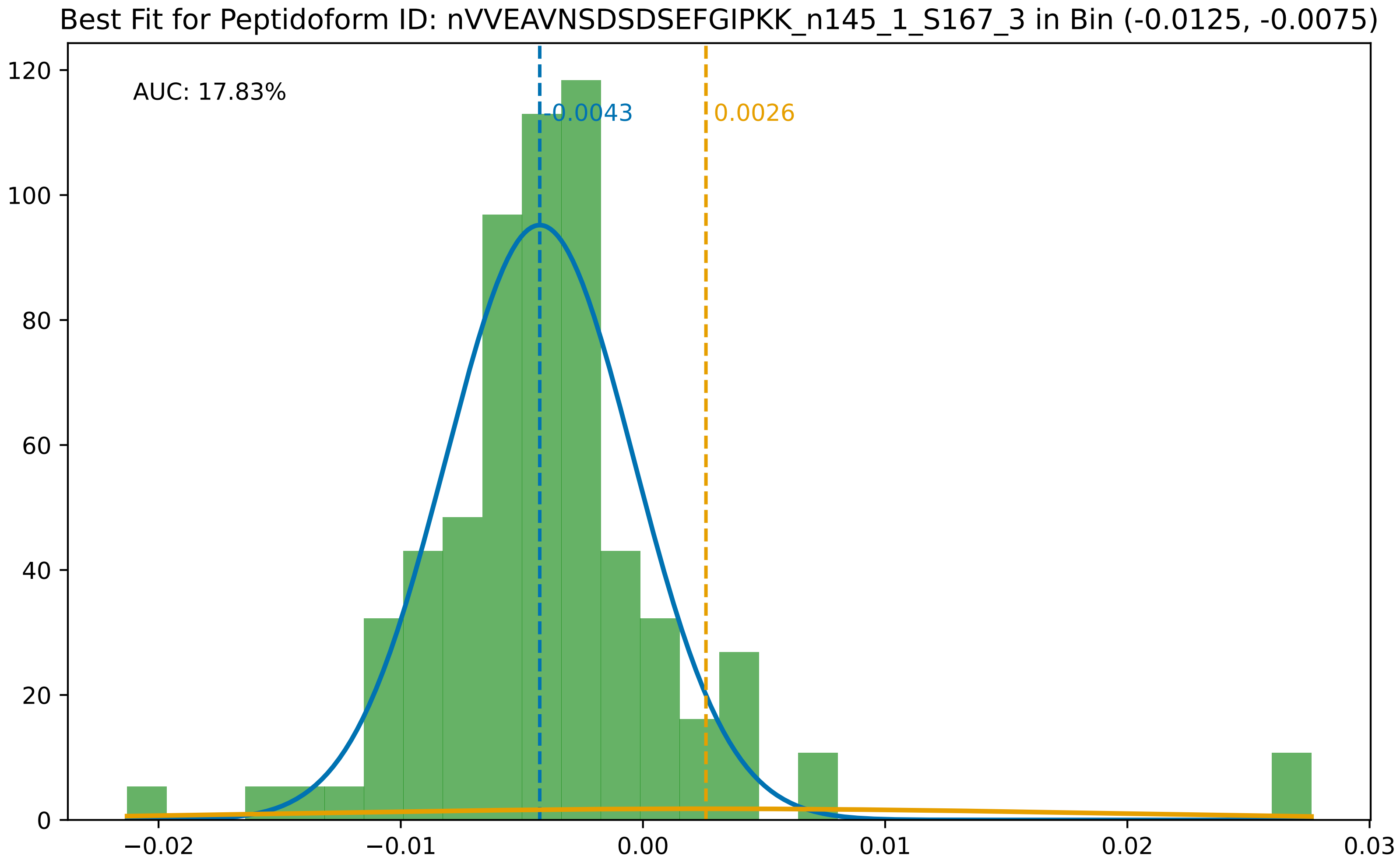
Best Fit for Peptidoform ID: nVVEAVNSDSDSEFGIPKK_n145_1_S167_3 in Bin (-0.0125, -0.0075)
AUC: 17.83%
-0.0043
0.0026
120
100
80
60
40
20
0
−0.02
−0.01
0.00
0.01
0.02
0.03

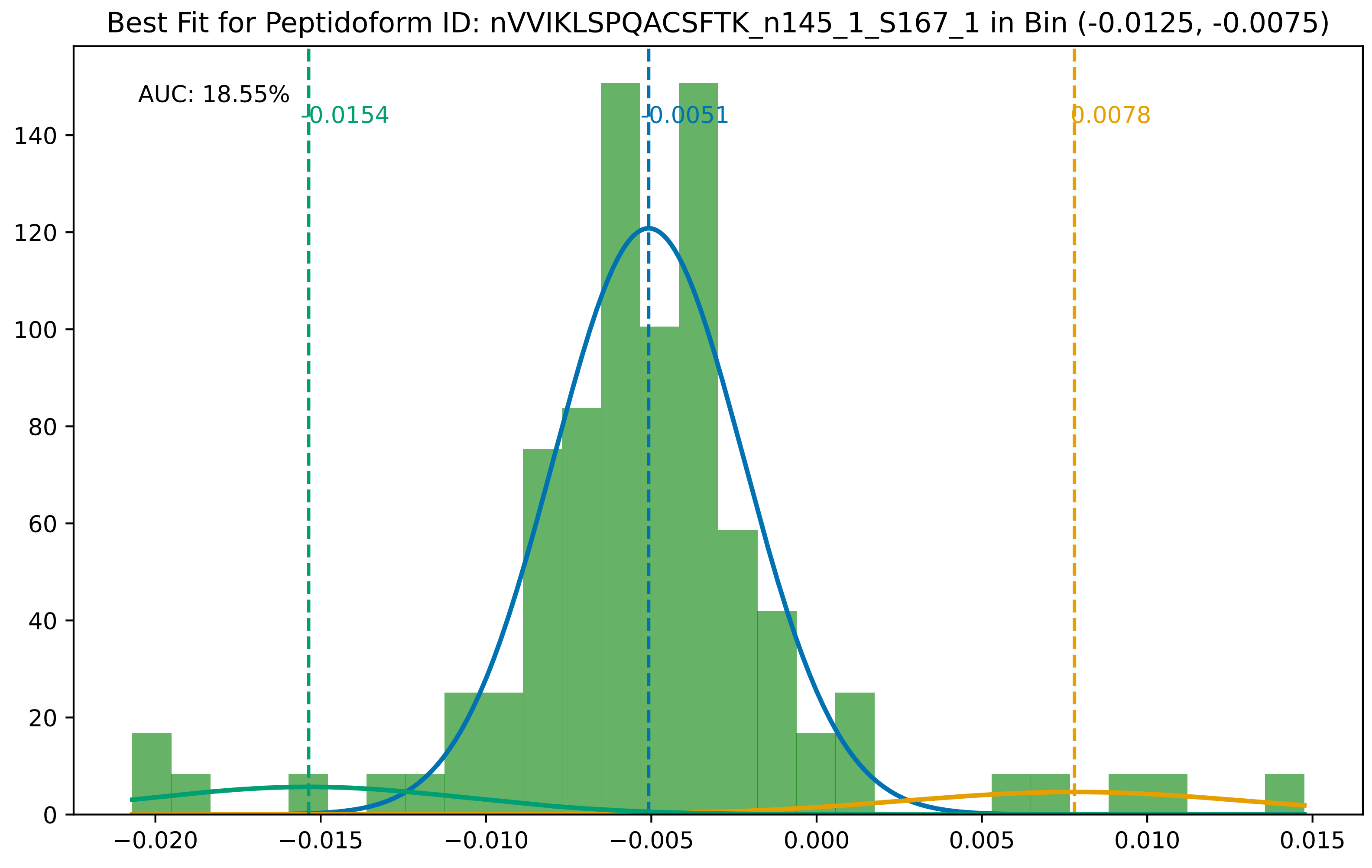

Best Fit for Peptidoform ID: nVVIKLSPQACSFTK_n145_1_S167_1 in Bin (-0.0125, -0.0075)
AUC: 18.55%
-0.0154
-0.0051
0.0078
0
20
40
60
80
100
120
140
−0.020
−0.015
−0.010
−0.005
0.000
0.005
0.010
0.015

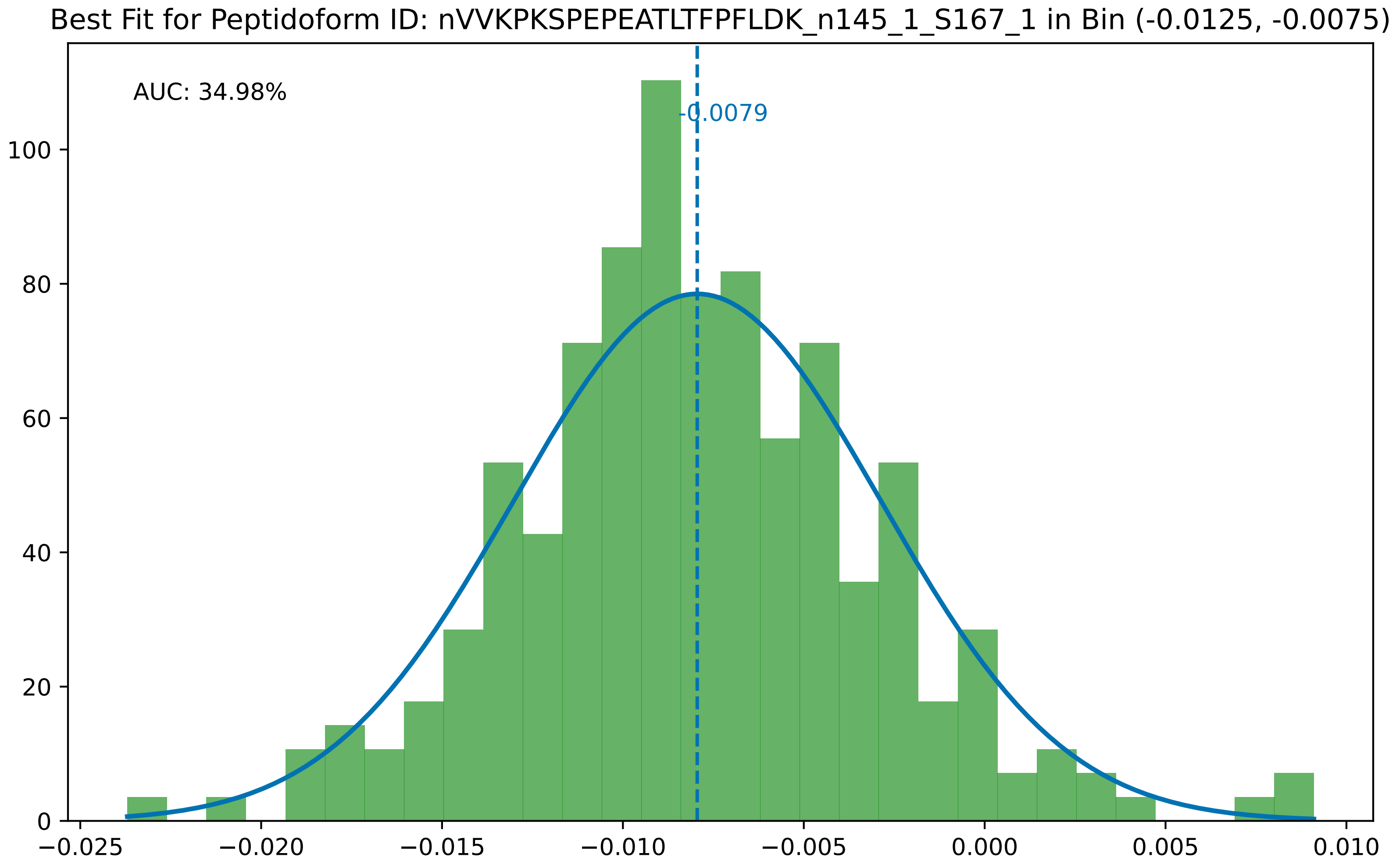
Best Fit for Peptidoform ID: nVVKPKSPEPEATLTFPFLDK_n145_1_S167_1 in Bin (-0.0125, -0.0075)
AUC: 34.98%
-0.0079
100
80
60
40
20
0
−0.025
−0.020
−0.015
−0.010
−0.005
0.000
0.005
0.010

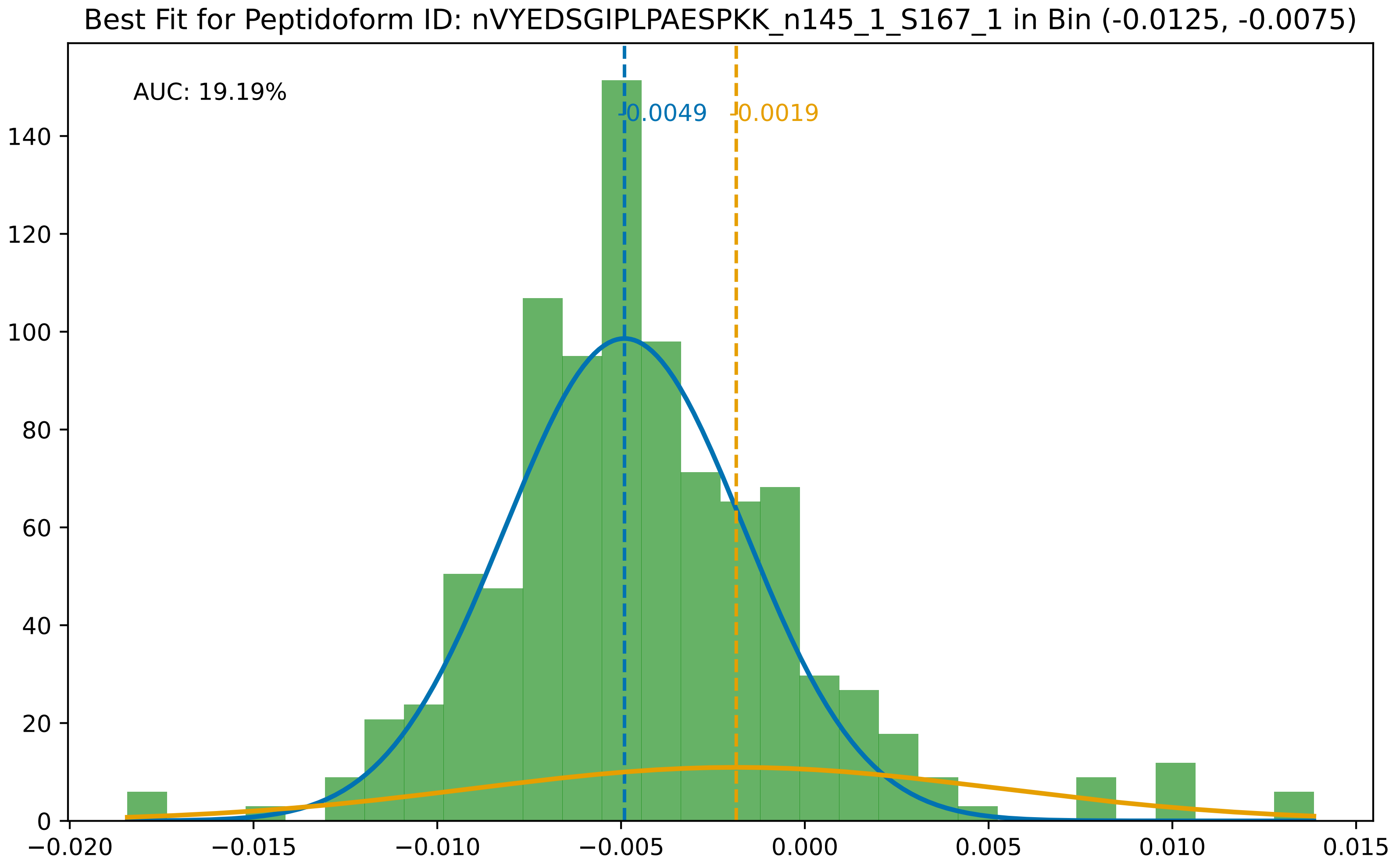

Best Fit for Peptidoform ID: nVYEDSGIPLPAESPKK_n145_1_S167_1 in Bin (-0.0125, -0.0075)
AUC: 19.19%
-0.0049
-0.0019
0
20
40
60
80
100
120
140
−0.020
−0.015
−0.010
−0.005
0.000
0.005
0.010
0.015

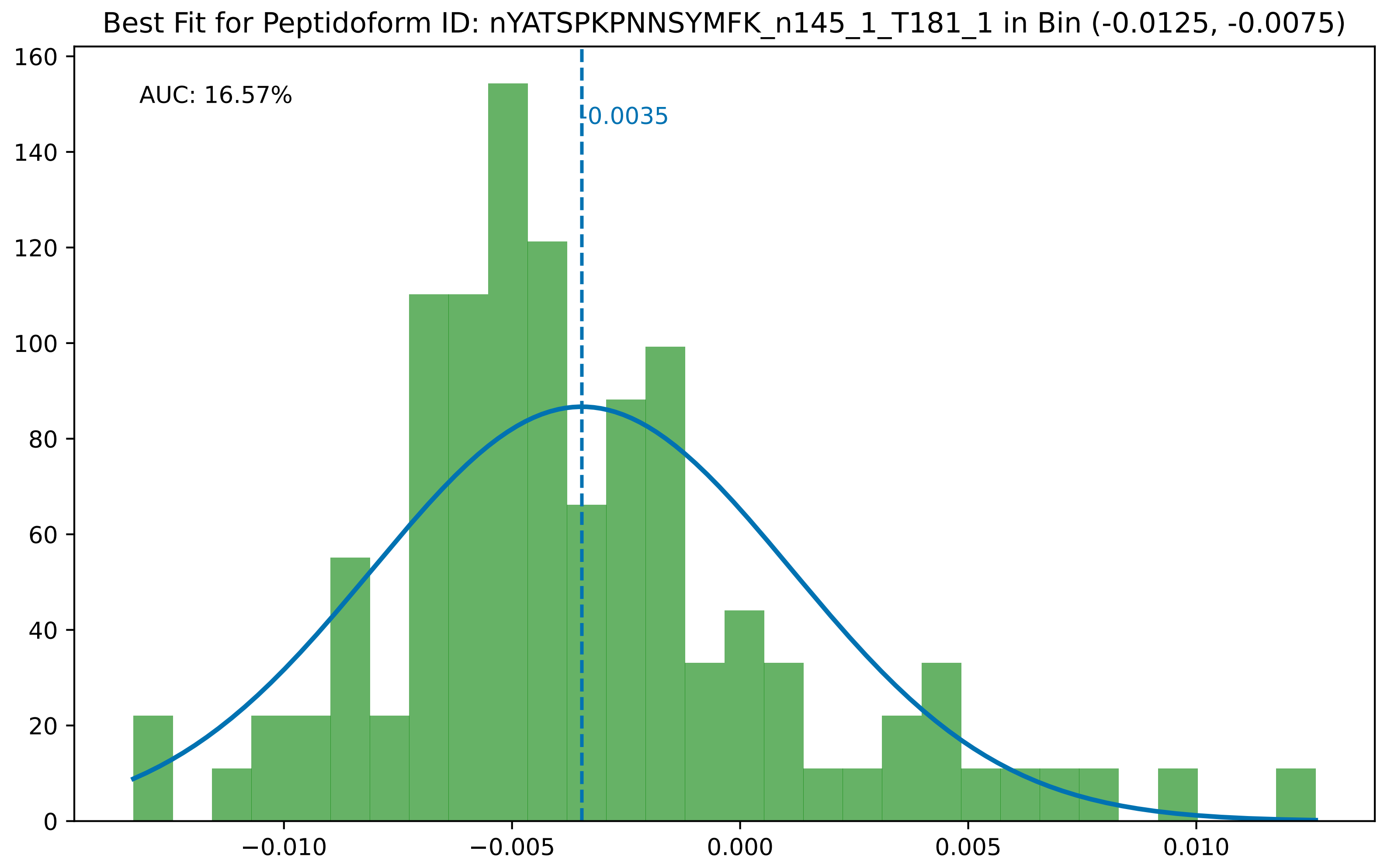

Best Fit for Peptidoform ID: nYATSPKPNNSYMFK_n145_1_T181_1 in Bin (-0.0125, -0.0075)
AUC: 16.57%
-0.0035
0
20
40
60
80
100
120
140
160
−0.010
−0.005
0.000
0.005
0.010

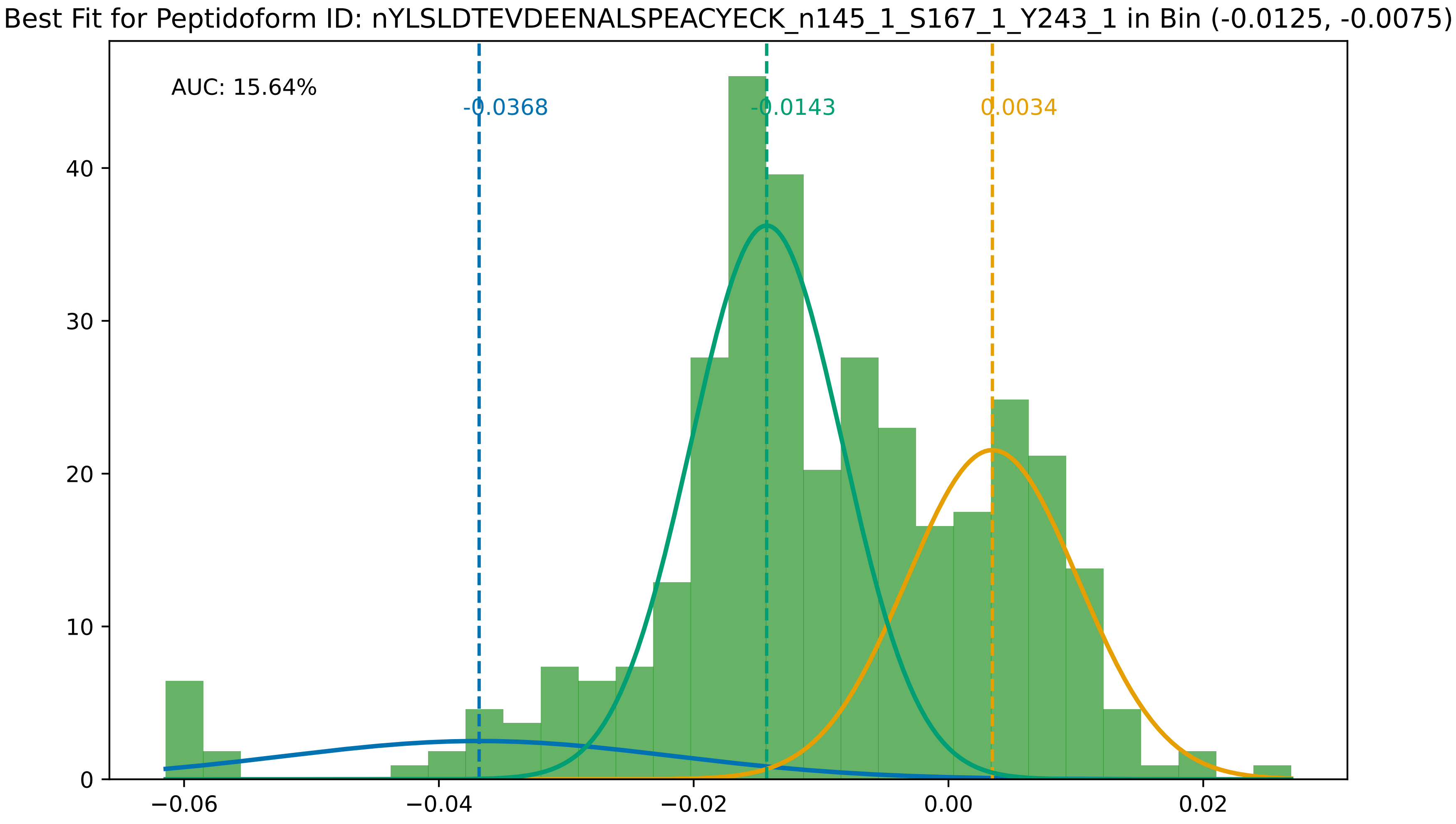

Best Fit for Peptidoform ID: nYLSLDTEVDEENALSPEACYECK_n145_1_S167_1_Y243_1 in Bin (-0.0125, -0.0075)
AUC: 15.64%
-0.0368
-0.0143
0.0034
0
10
20
30
40
−0.06
−0.04
−0.02
0.00
0.02

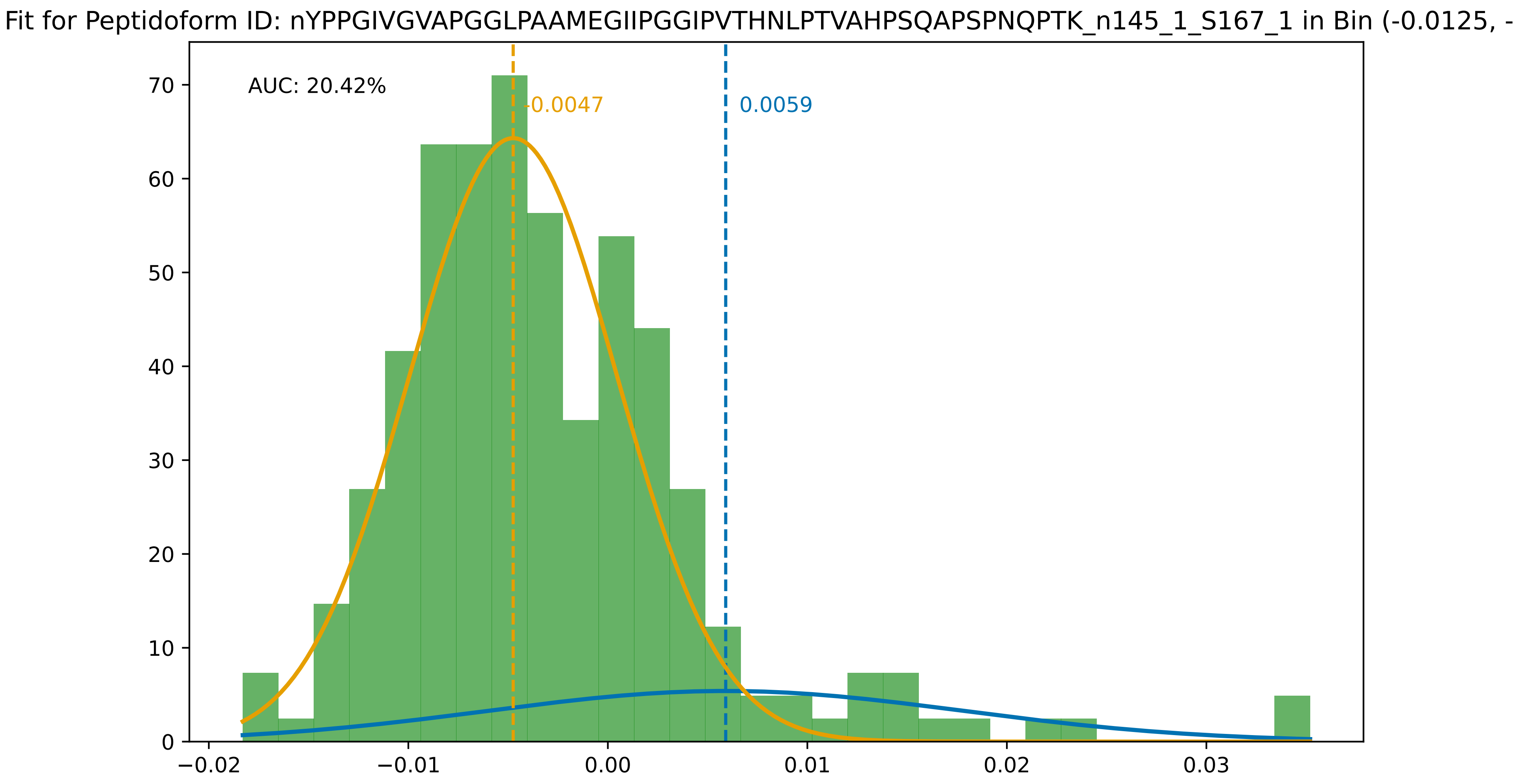

Fit for Peptidoform ID: nYPPGIVGVAPGGLPAAMEGIIPGGIPVTHNLPTVAHPSQAPSPNQPTK_n145_1_S167_1 in Bin (-0.0125, -
AUC: 20.42%
-0.0047
0.0059
0
10
20
30
40
50
60
70
−0.02
−0.01
0.00
0.01
0.02
0.03

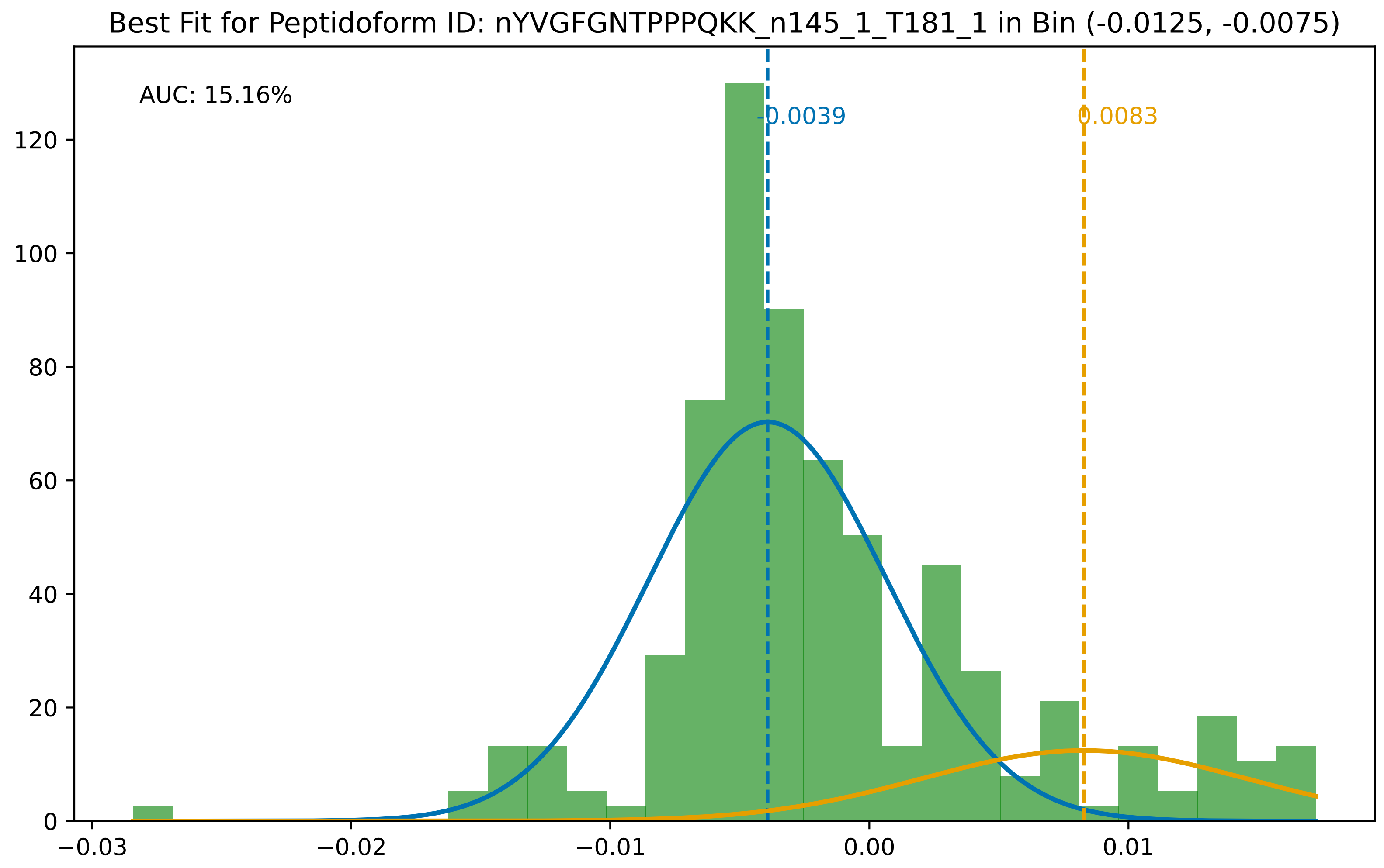

Best Fit for Peptidoform ID: nYVGFGNTPPPQKK_n145_1_T181_1 in Bin (-0.0125, -0.0075)
AUC: 15.16%
-0.0039
0.0083
0
20
40
60
80
100
120
−0.03
−0.02
−0.01
0.00
0.01
