## Supplementary File 4 for "Searching for Sulfotyrosines (sY) in a HA(pY)STACK": histograms_post_auc_filtering_bin_-0.0175_-0.0125.pdf

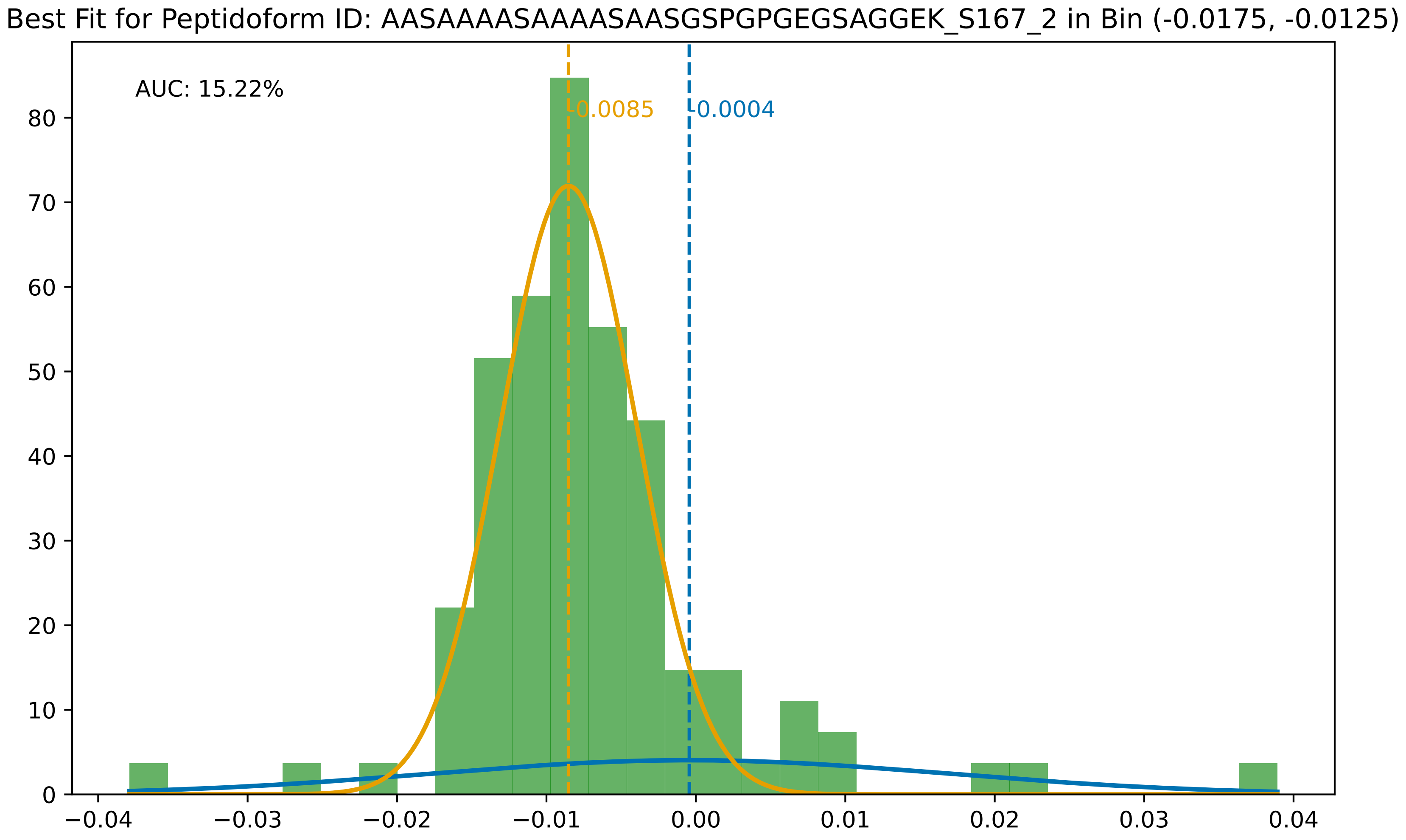

Best Fit for Peptidoform ID: AASAAAASAAAASAASGSPGPGEGSAGGEK_S167_2 in Bin (-0.0175, -0.0125)
AUC: 15.22%
-0.0085
-0.0004
0
10
20
30
40
50
60
70
80
−0.04
−0.03
−0.02
−0.01
0.00
0.01
0.02
0.03
0.04

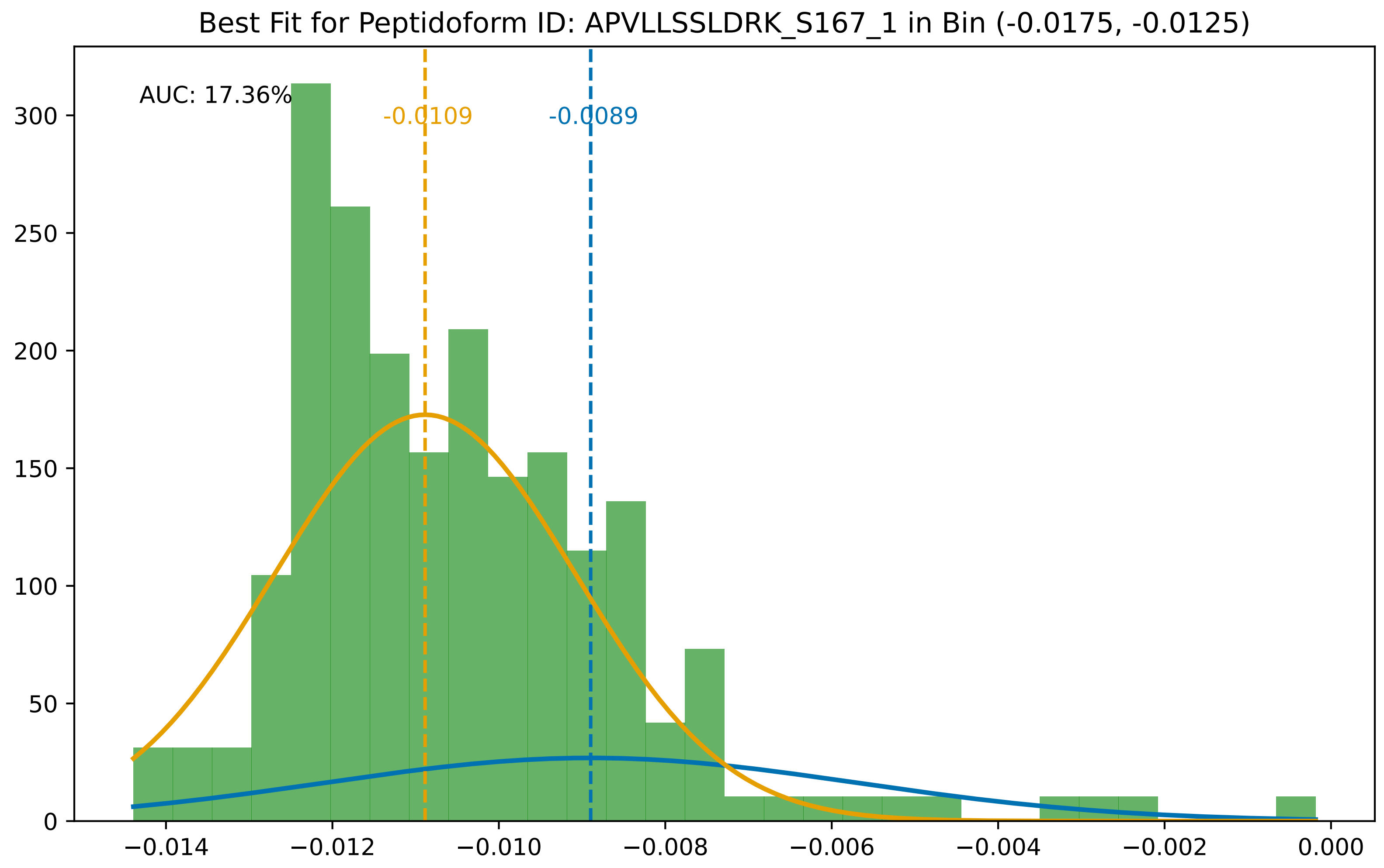
Best Fit for Peptidoform ID: APVLLSSLDRK_S167_1 in Bin (-0.0175, -0.0125)
AUC: 17.36%
-0.0109
-0.0089
0
50
100
150
200
250
300
−0.014
−0.012
−0.010
−0.008
−0.006
−0.004
−0.002
0.000

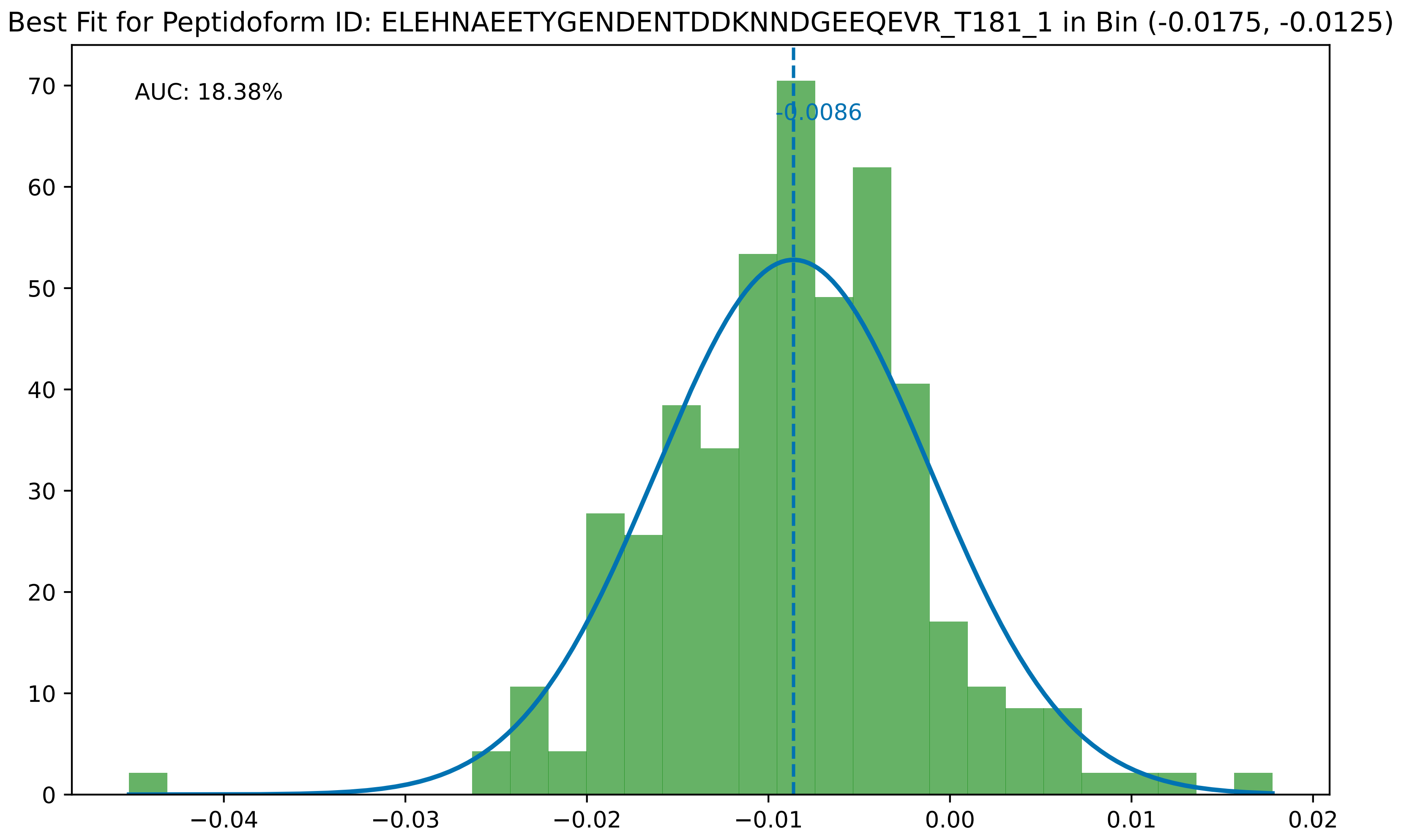
Best Fit for Peptidoform ID: ELEHNAEETYGENDENTDDKNNDGEEQEVR_T181_1 in Bin (-0.0175, -0.0125)
AUC: 18.38%
-0.0086
0
10
20
30
40
50
60
70
−0.04
−0.03
−0.02
−0.01
0.00
0.01
0.02

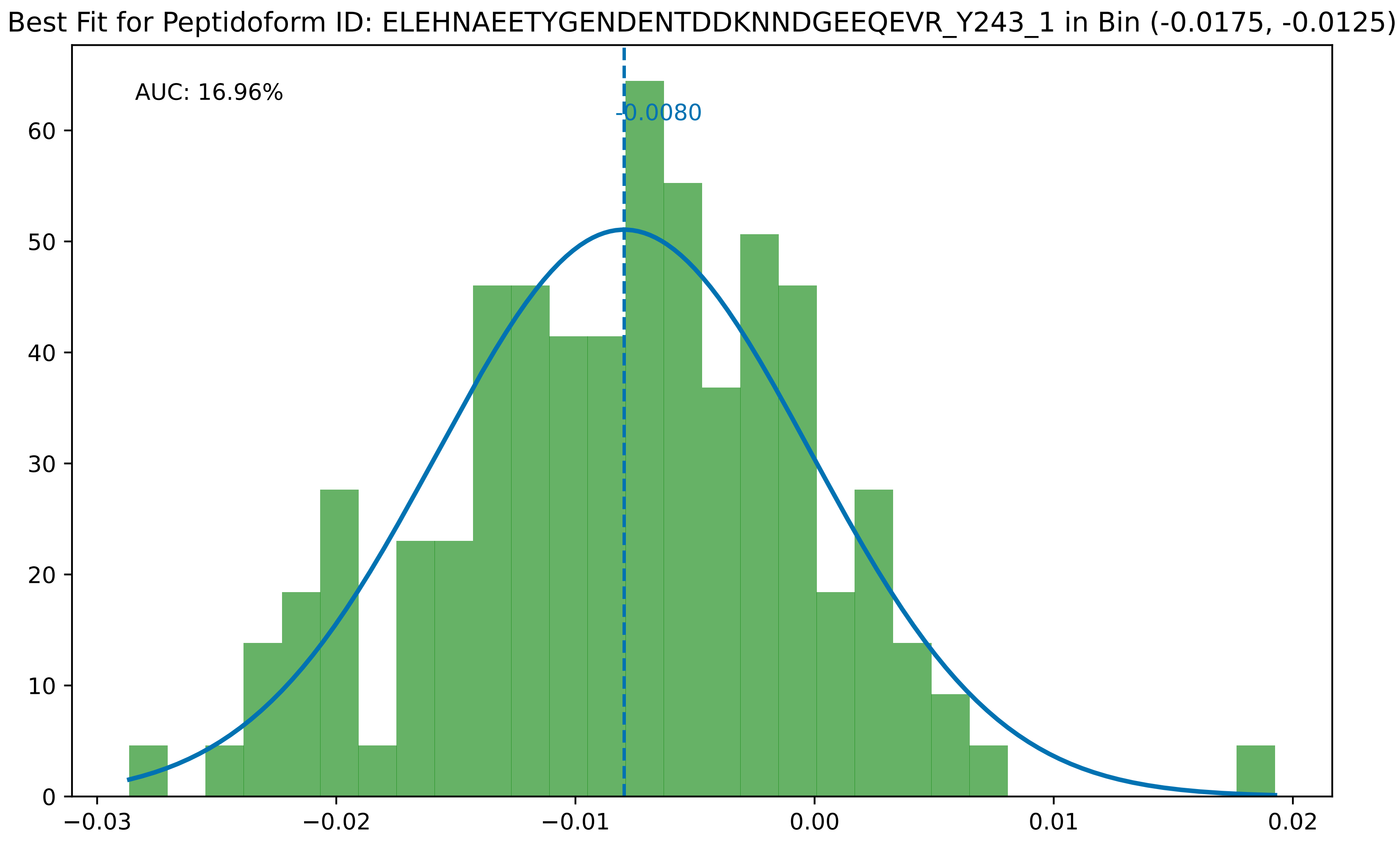
Best Fit for Peptidoform ID: ELEHNAEETYGENDENTDDKNNDGEEQEVR_Y243_1 in Bin (-0.0175, -0.0125)
AUC: 16.96%
-0.0080
0
10
20
30
40
50
60
−0.03
−0.02
−0.01
0.00
0.01
0.02

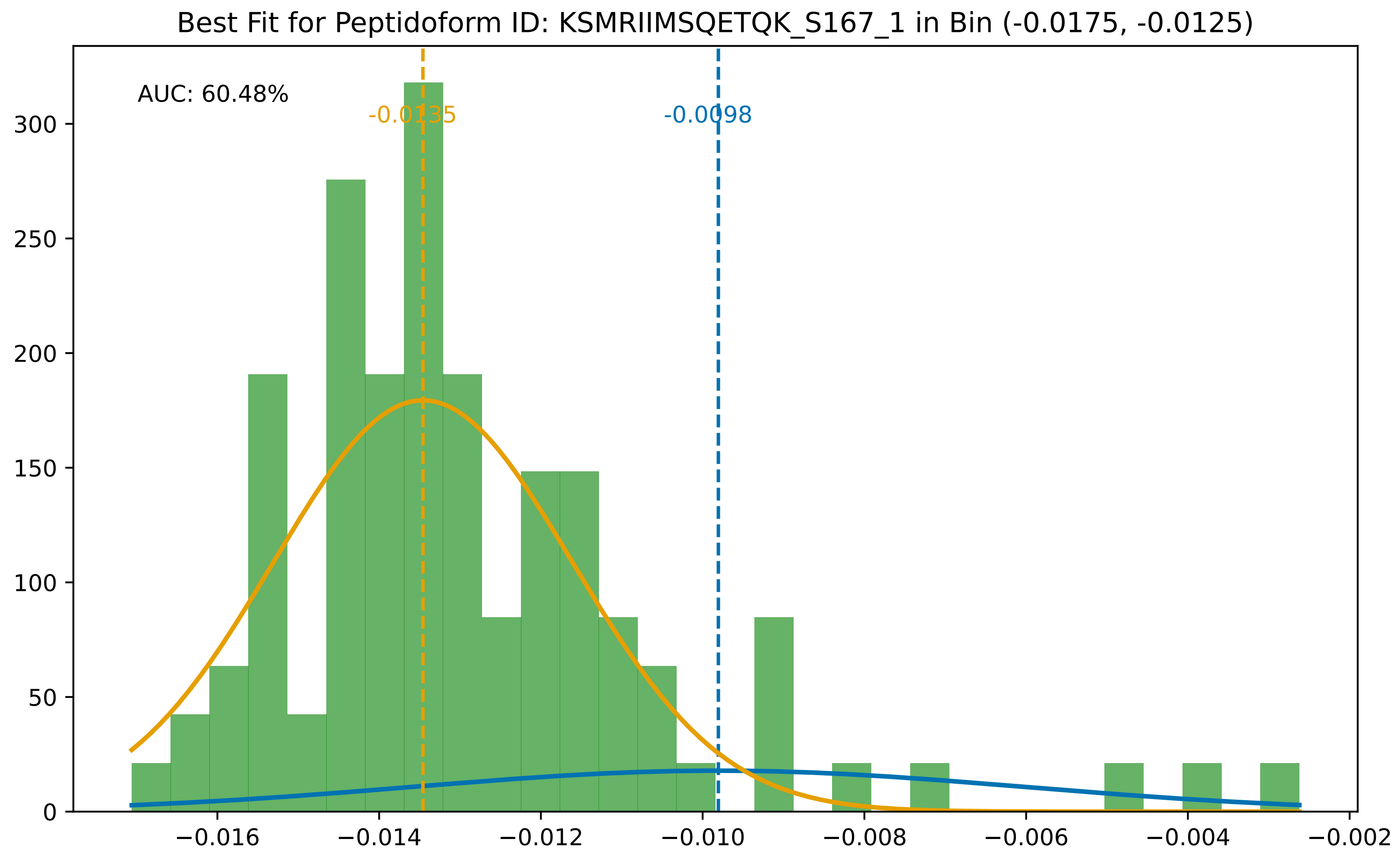
Best Fit for Peptidoform ID: KSMRIIMSQETQK_S167_1 in Bin (-0.0175, -0.0125)
AUC: 60.48%
-0.0135
-0.0098
0
50
100
150
200
250
300
−0.016
−0.014
−0.012
−0.010
−0.008
−0.006
−0.004
−0.002

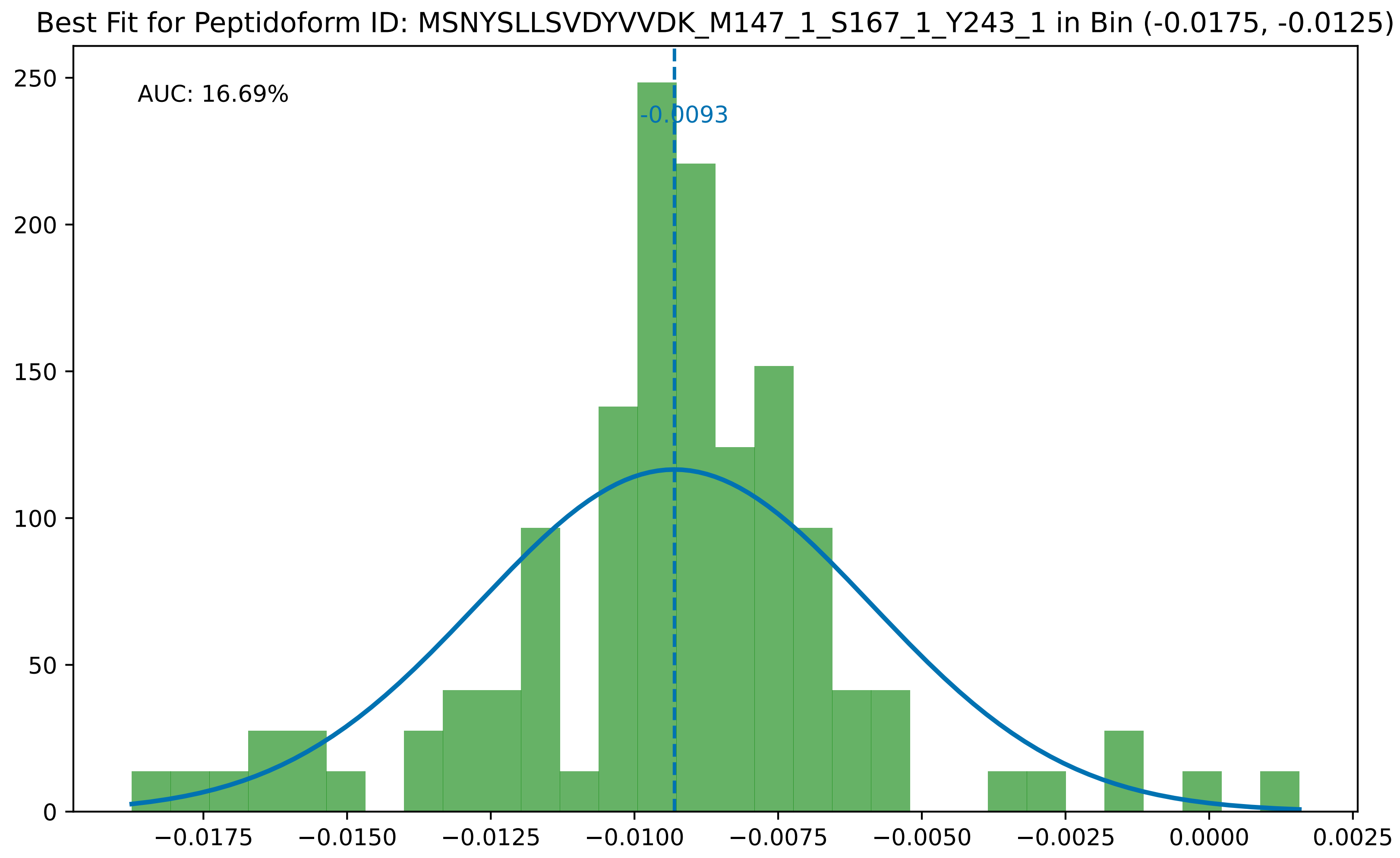

Best Fit for Peptidoform ID: MSNYSLLSVDYVVDK_M147_1_S167_1_Y243_1 in Bin (-0.0175, -0.0125)
AUC: 16.69%
-0.0093
250
200
150
100
50
0
−0.0175
−0.0150
−0.0125
−0.0100
−0.0075
−0.0050
−0.0025
0.0000
0.0025

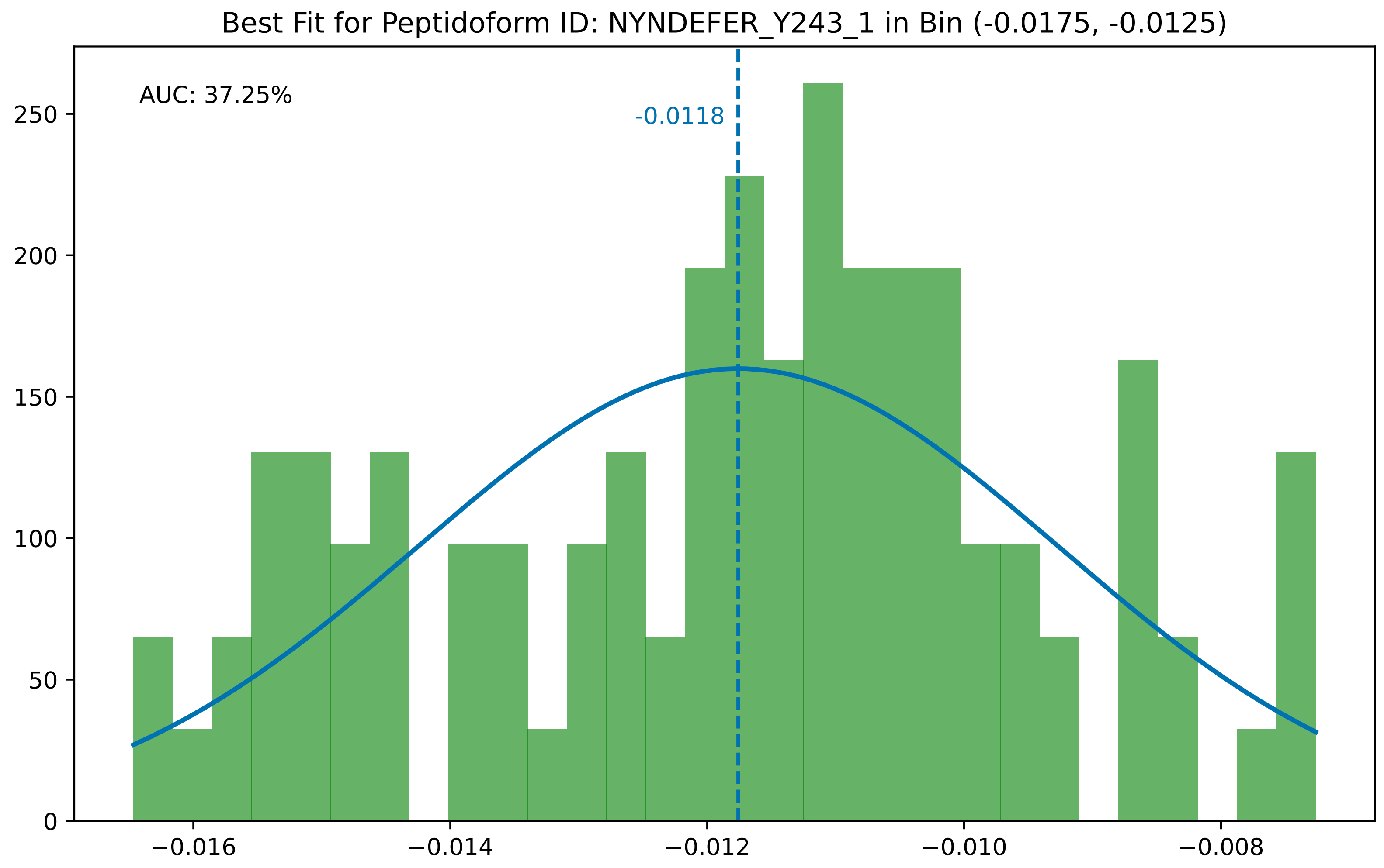

Best Fit for Peptidoform ID: NYNDEFER_Y243_1 in Bin (-0.0175, -0.0125)
AUC: 37.25%
-0.0118
0
50
100
150
200
250
−0.016
−0.014
−0.012
−0.010
−0.008

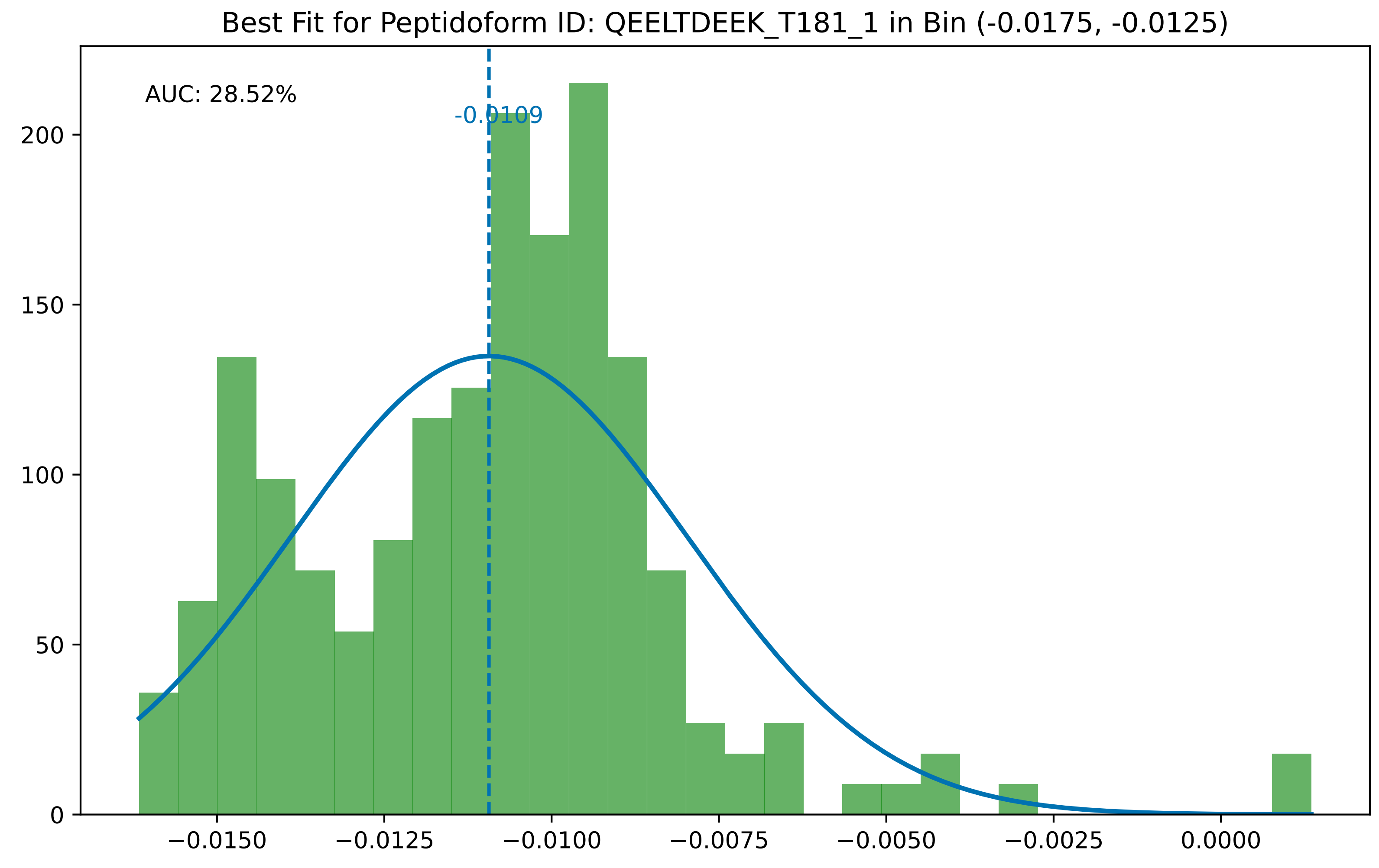

Best Fit for Peptidoform ID: QEELTDEEK_T181_1 in Bin (-0.0175, -0.0125)
AUC: 28.52%
-0.0109
0
50
100
150
200
−0.0150
−0.0125
−0.0100
−0.0075
−0.0050
−0.0025
0.0000

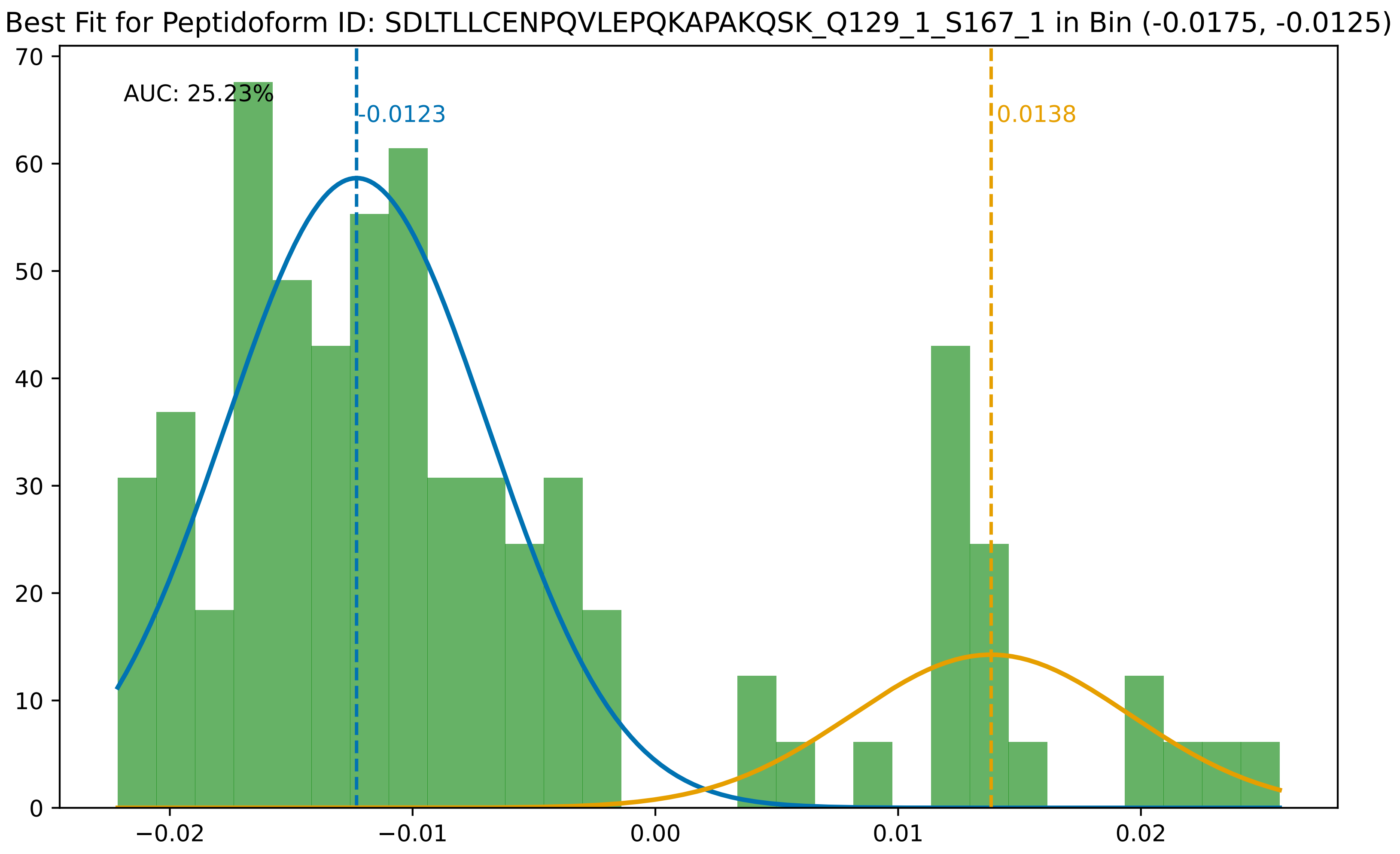

Best Fit for Peptidoform ID: SDLTLLCENPQVLEPQKAPAKQSK_Q129_1_S167_1 in Bin (-0.0175, -0.0125)
AUC: 25.23%
-0.0123
0.0138
70
60
50
40
30
20
10
0
−0.02
−0.01
0.00
0.01
0.02

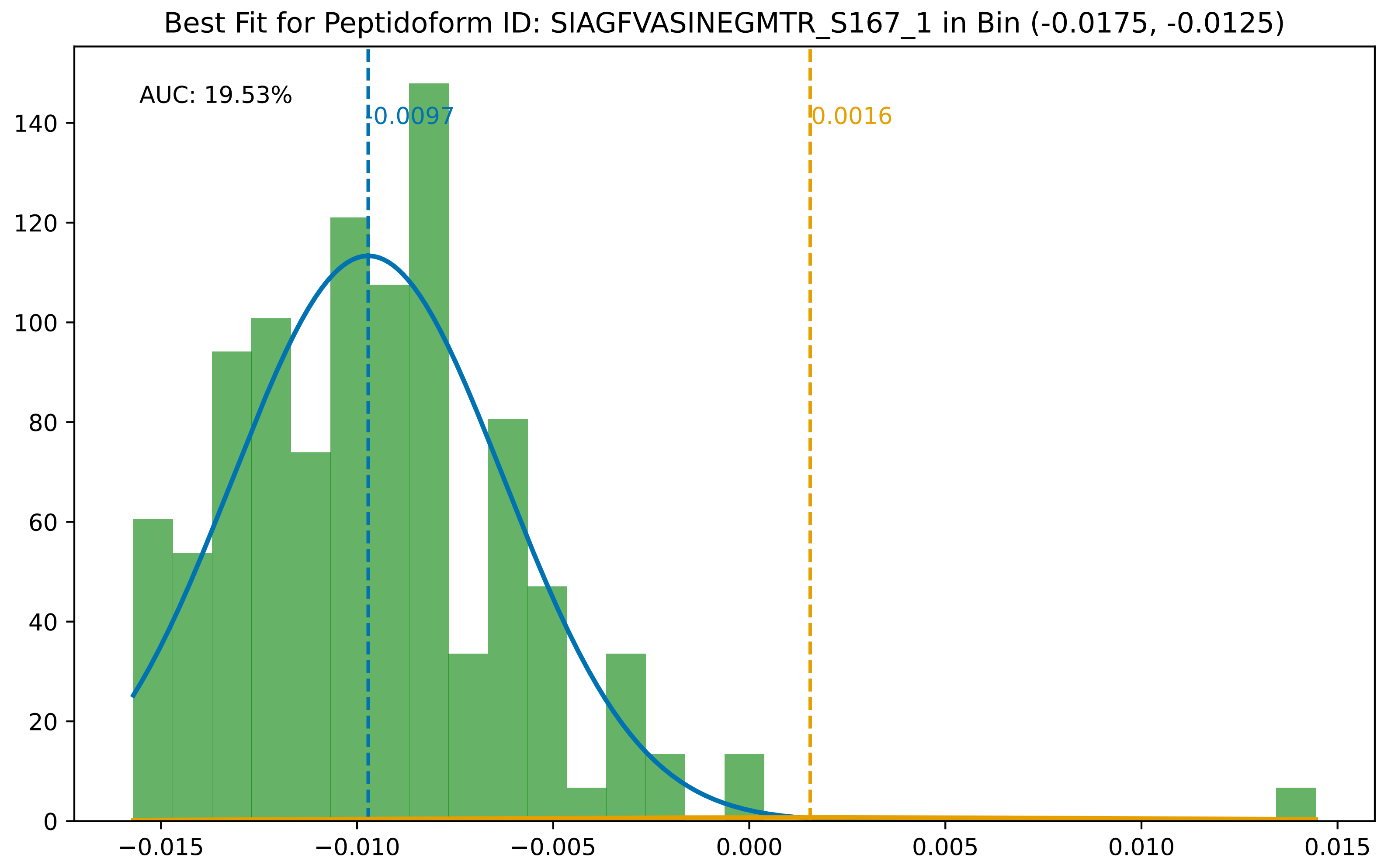

Best Fit for Peptidoform ID: SIAGFVASINEGMTR_S167_1 in Bin (-0.0175, -0.0125)
AUC: 19.53%
-0.0097
0.0016
0
20
40
60
80
100
120
140
−0.015
−0.010
−0.005
0.000
0.005
0.010
0.015

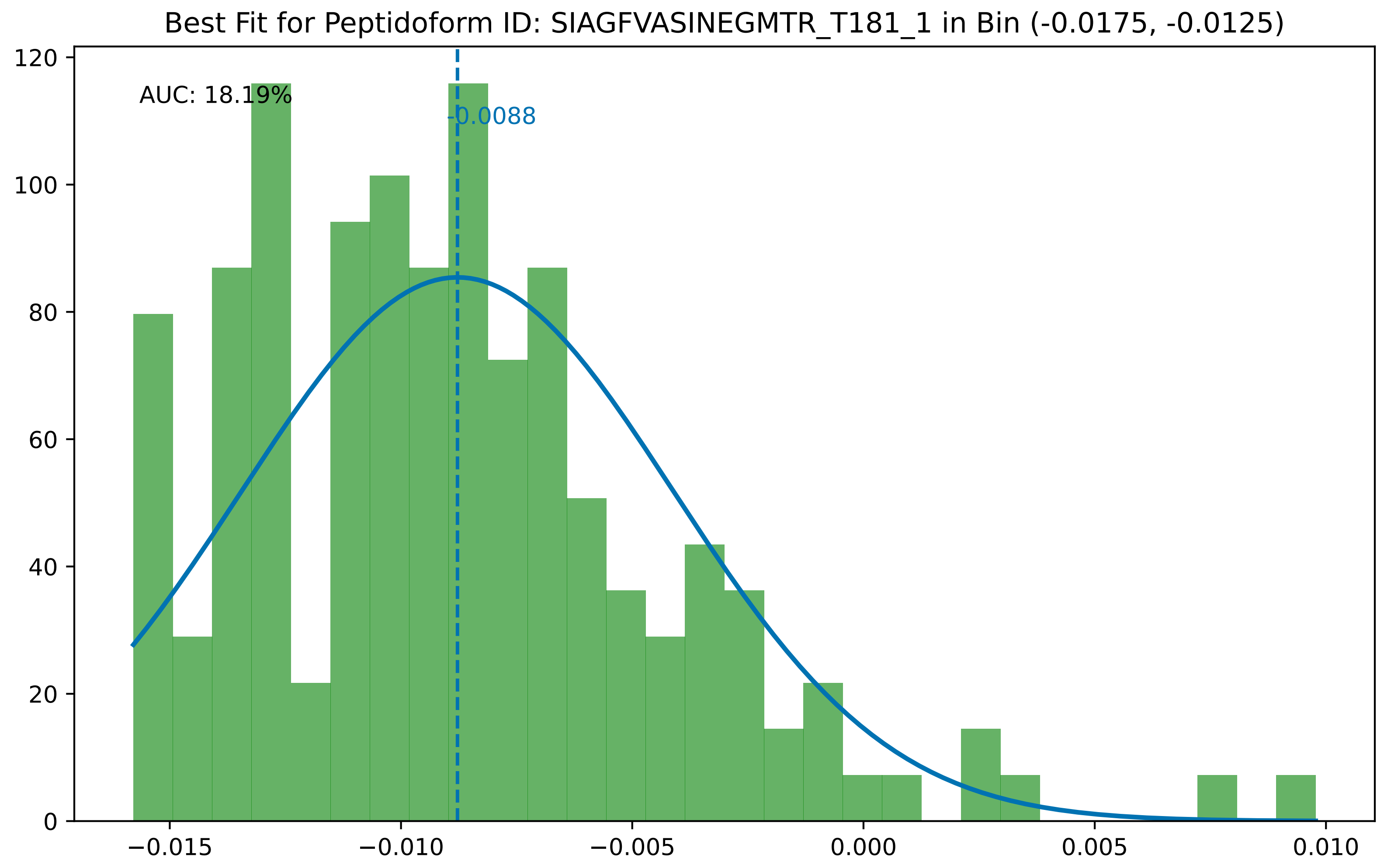
Best Fit for Peptidoform ID: SIAGFVASINEGMTR_T181_1 in Bin (-0.0175, -0.0125)
AUC: 18.19%
-0.0088
0
20
40
60
80
100
120
−0.015
−0.010
−0.005
0.000
0.005
0.010

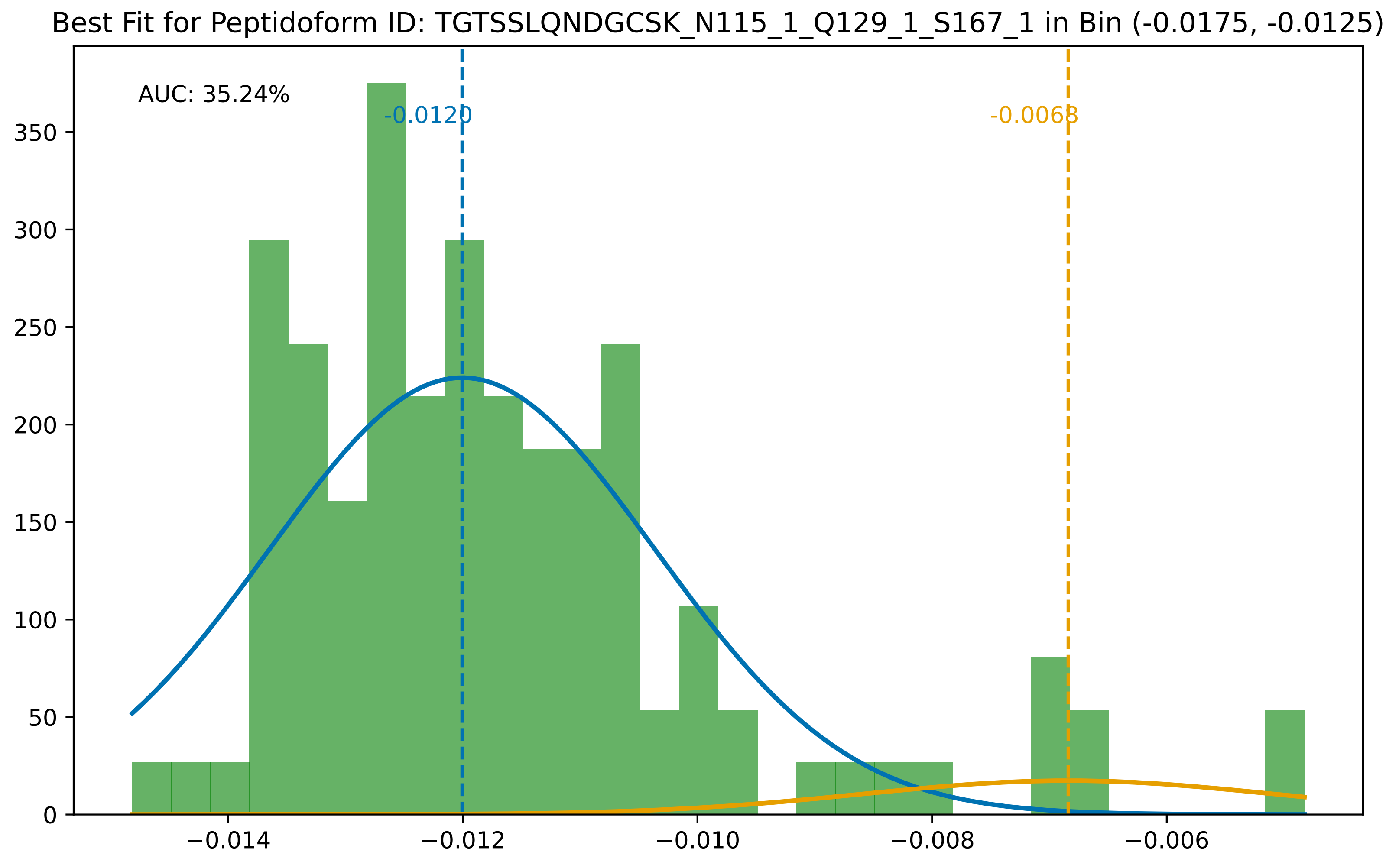
Best Fit for Peptidoform ID: TGTSSLQNDGCSK_N115_1_Q129_1_S167_1 in Bin (-0.0175, -0.0125)
AUC: 35.24%
-0.0120
-0.0068
350
300
250
200
150
100
50
0
−0.014
−0.012
−0.010
−0.008
−0.006

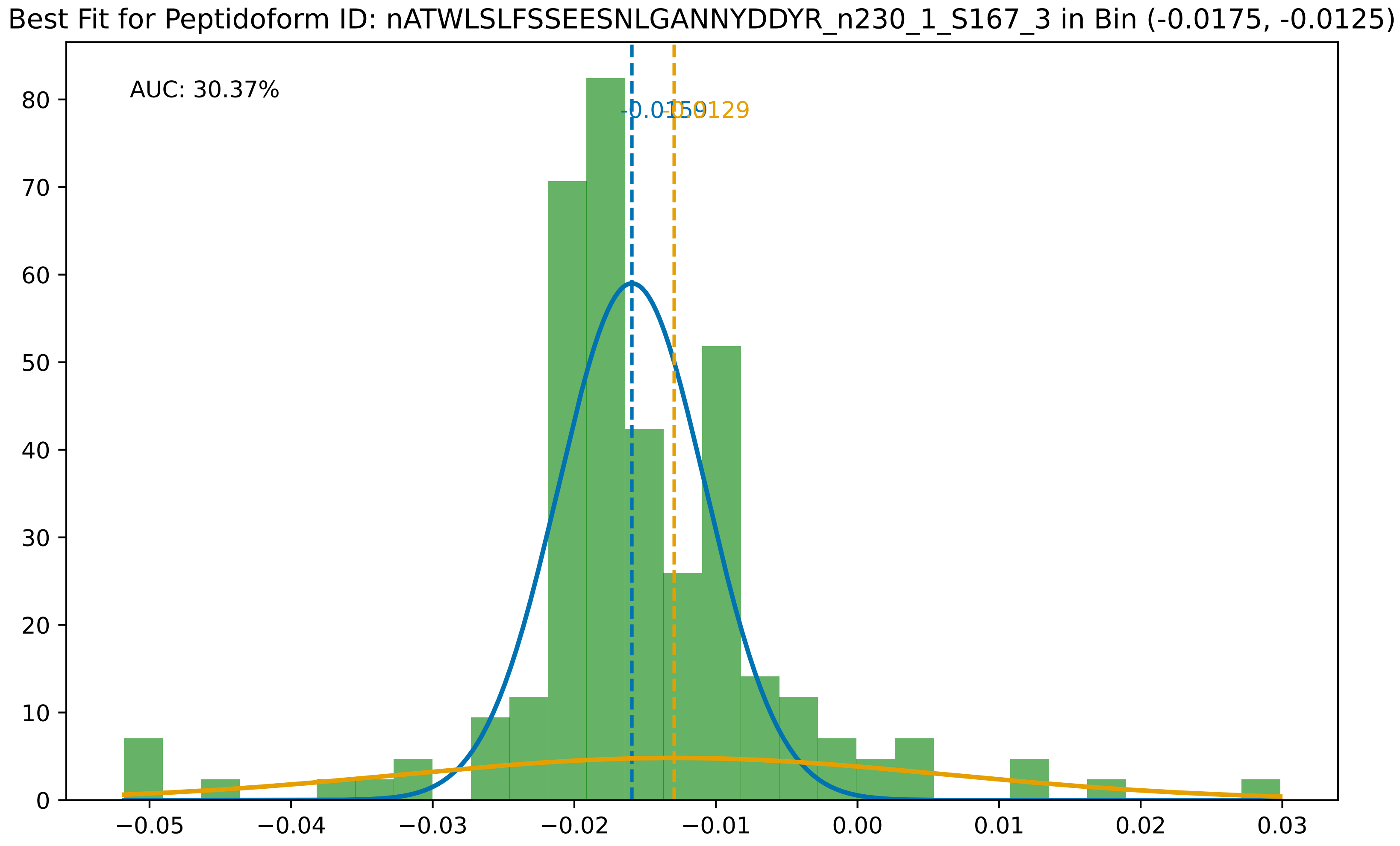

Best Fit for Peptidoform ID: nATWLSLFSSEESNLGANNYDDYR_n230_1_S167_3 in Bin (-0.0175, -0.0125)
AUC: 30.37%
-0.0159
-0.0129
0
10
20
30
40
50
60
70
80
−0.05
−0.04
−0.03
−0.02
−0.01
0.00
0.01
0.02
0.03

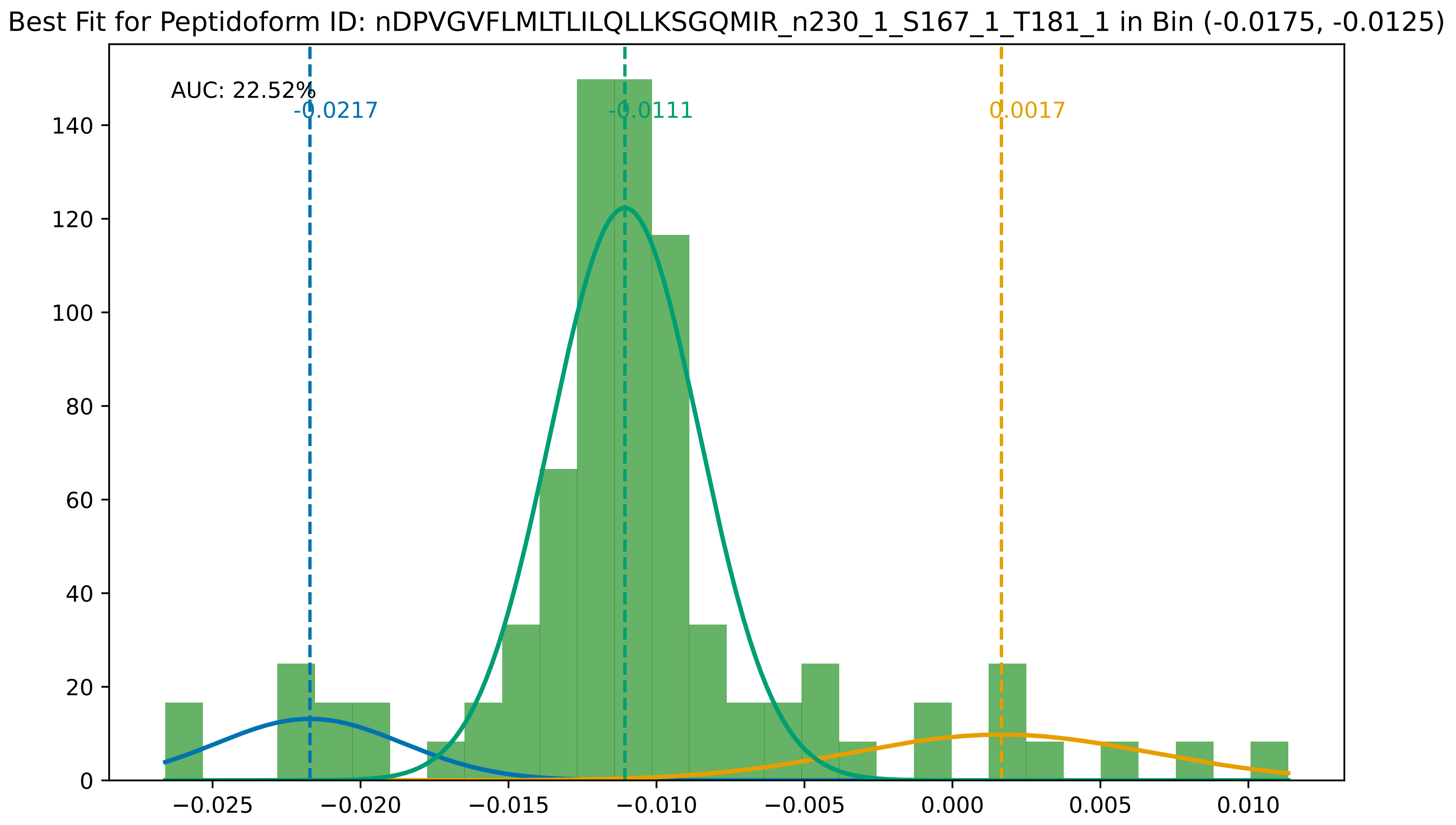
Best Fit for Peptidoform ID: nDPVGVFLMLTLILQLLKSGQMIR_n230_1_S167_1_T181_1 in Bin (-0.0175, -0.0125)
AUC: 22.52%
-0.0217
-0.0111
0.0017
0
20
40
60
80
100
120
140
−0.025
−0.020
−0.015
−0.010
−0.005
0.000
0.005
0.010

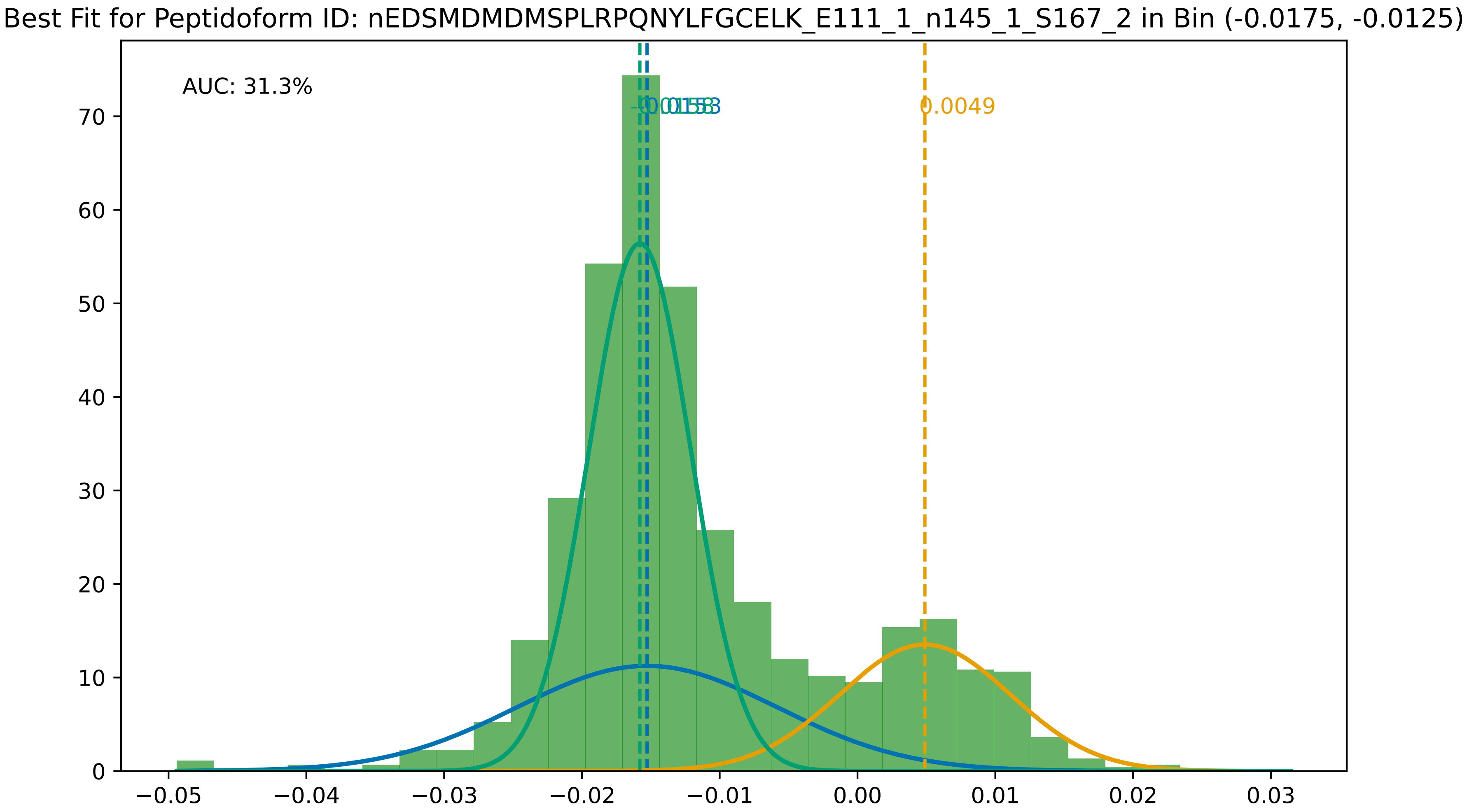

Best Fit for Peptidoform ID: nEDSMDMDMSPLRPQNYLFGCELK_E111_1_n145_1_S167_2 in Bin (-0.0175, -0.0125)
AUC: 31.3%
0.0049
0
10
20
30
40
50
60
70
−0.05
−0.04
−0.03
−0.02
−0.01
0.00
0.01
0.02
0.03

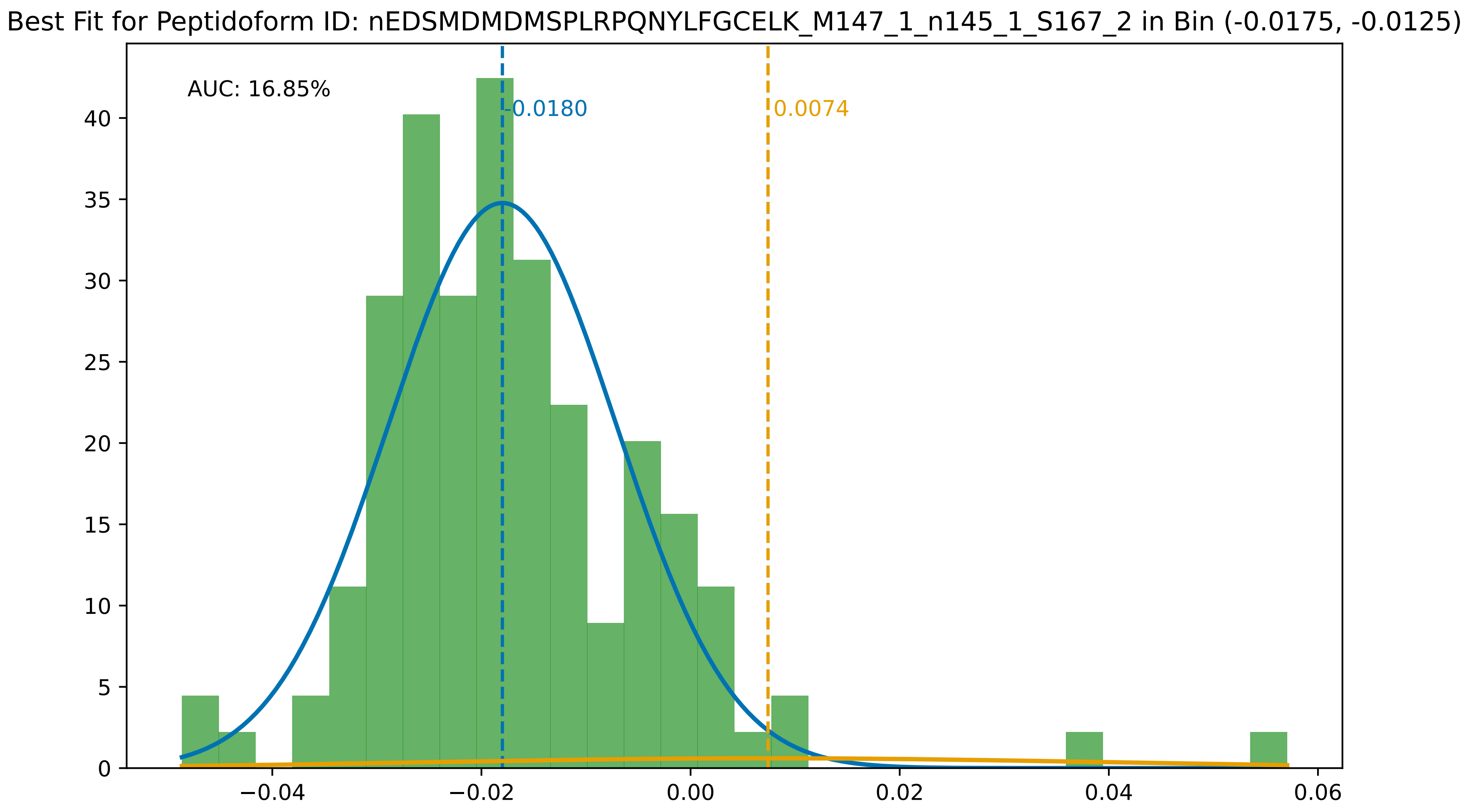

Best Fit for Peptidoform ID: nEDSMDMDMSPLRPQNYLFGCELK_M147_1_n145_1_S167_2 in Bin (-0.0175, -0.0125)
AUC: 16.85%
-0.0180
0.0074
0
5
10
15
20
25
30
35
40
−0.04
−0.02
0.00
0.02
0.04
0.06

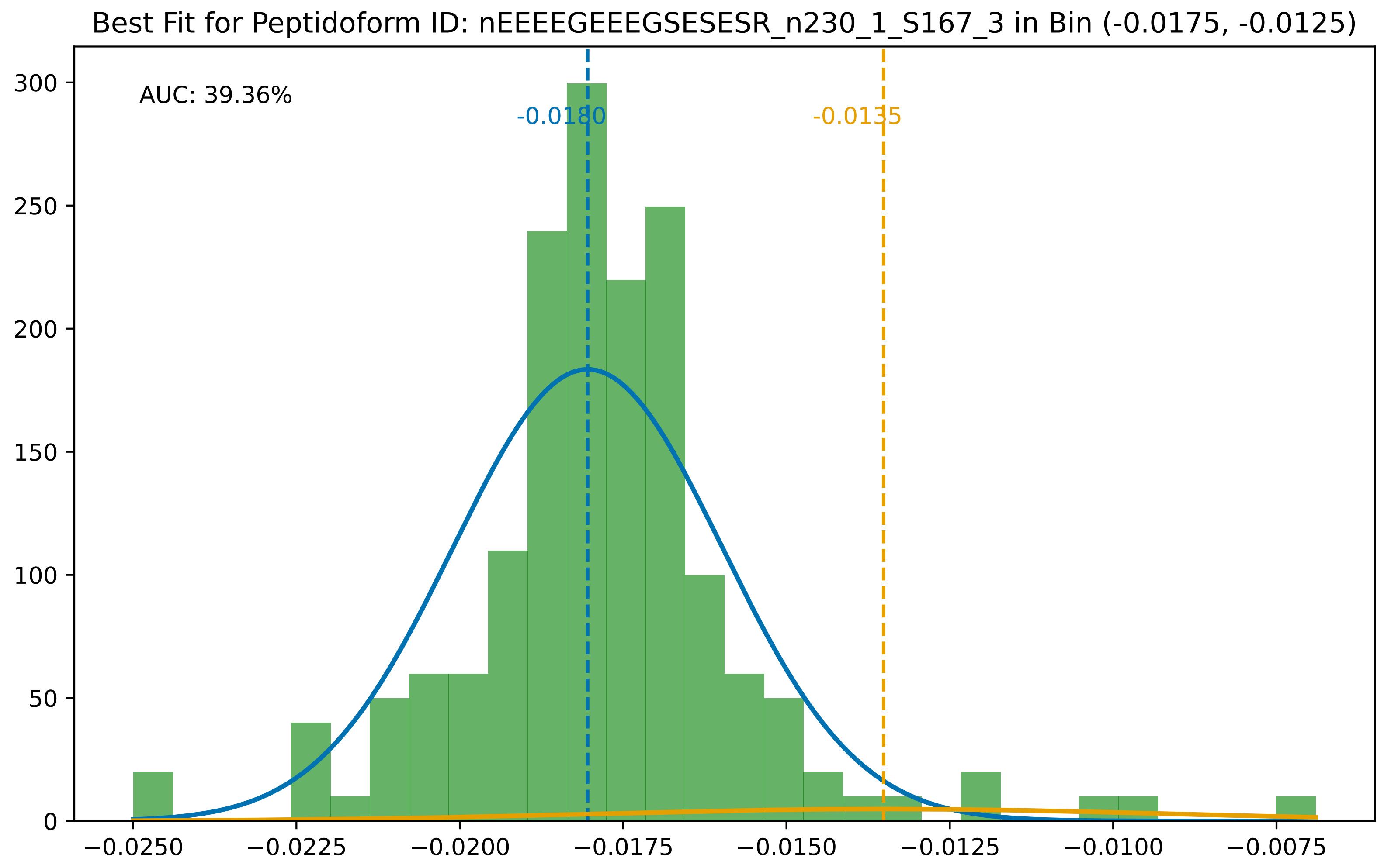
Best Fit for Peptidoform ID: nEEEEGEEEGSESESR_n230_1_S167_3 in Bin (-0.0175, -0.0125)
AUC: 39.36%
-0.0180
-0.0135
300
250
200
150
100
50
0
−0.0250
−0.0225
−0.0200
−0.0175
−0.0150
−0.0125
−0.0100
−0.0075

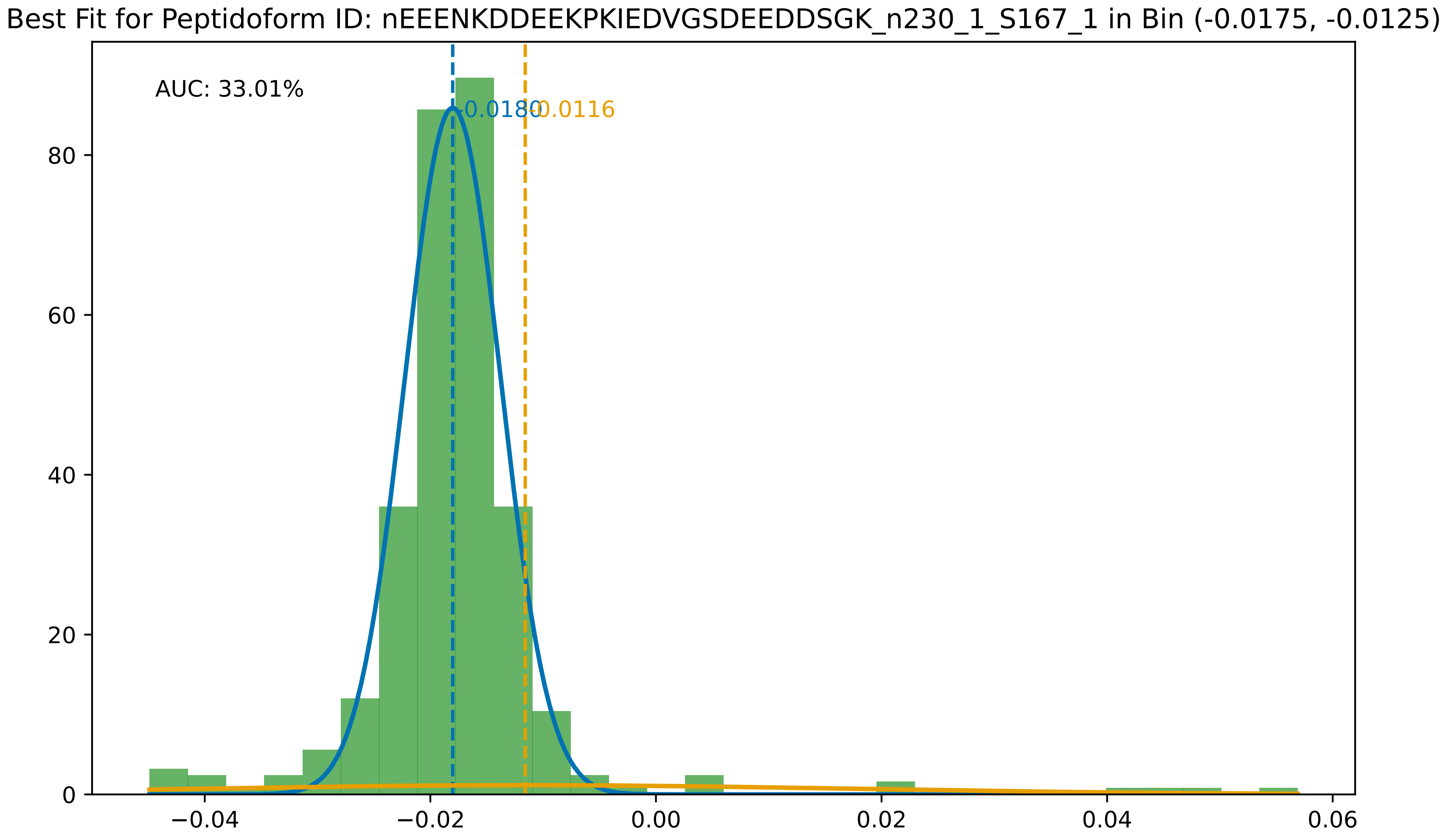
Best Fit for Peptidoform ID: nEEENKDDEEKPKIEDVGSDEEDDSGK_n230_1_S167_1 in Bin (-0.0175, -0.0125)
AUC: 33.01%
-0.0180
-0.0116
0
20
40
60
80
−0.04
−0.02
0.00
0.02
0.04
0.06

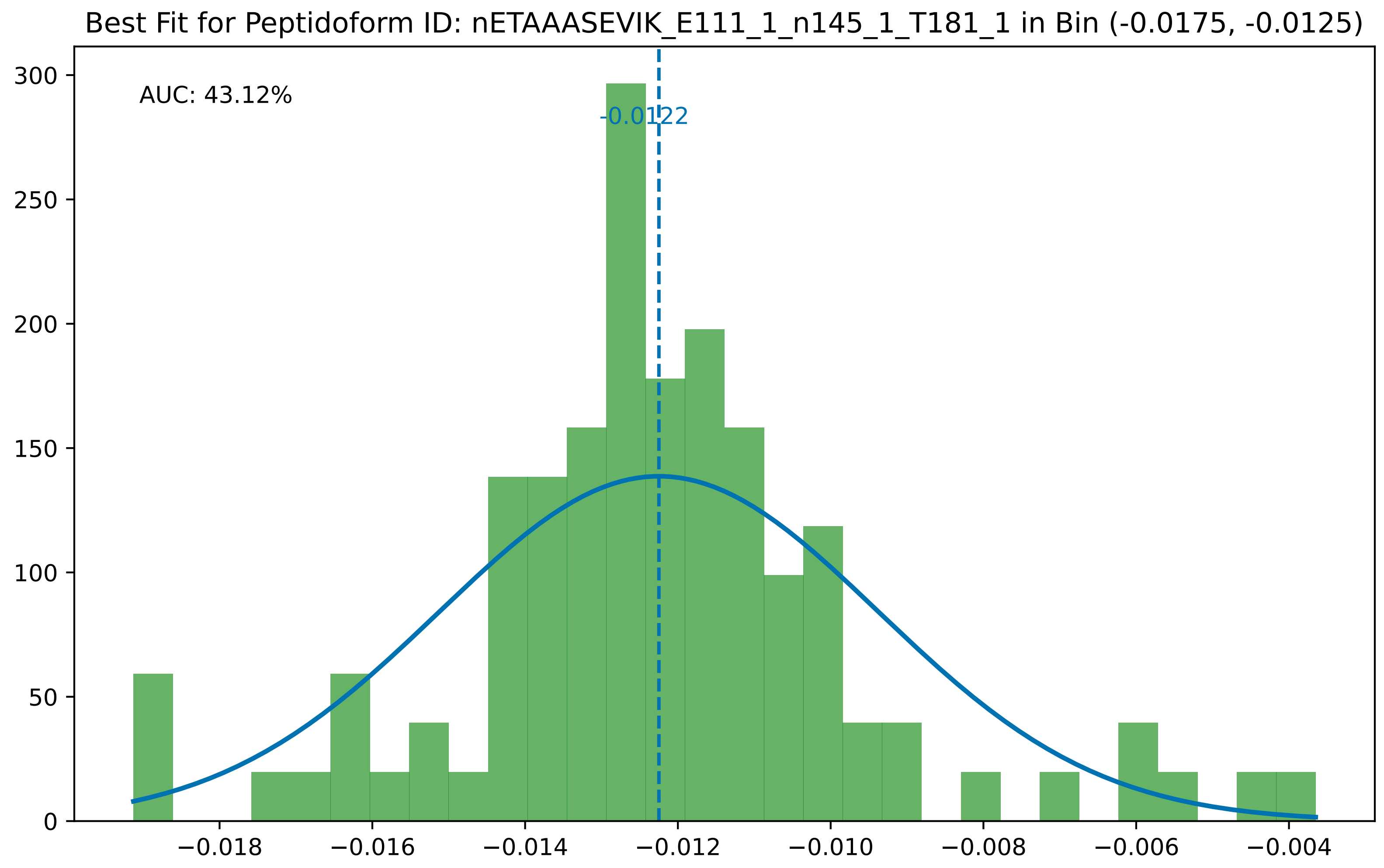
Best Fit for Peptidoform ID: nETAAASEVIK_E111_1_n145_1_T181_1 in Bin (-0.0175, -0.0125)
AUC: 43.12%
-0.0122
300
250
200
150
100
50
0
−0.018
−0.016
−0.014
−0.012
−0.010
−0.008
−0.006
−0.004

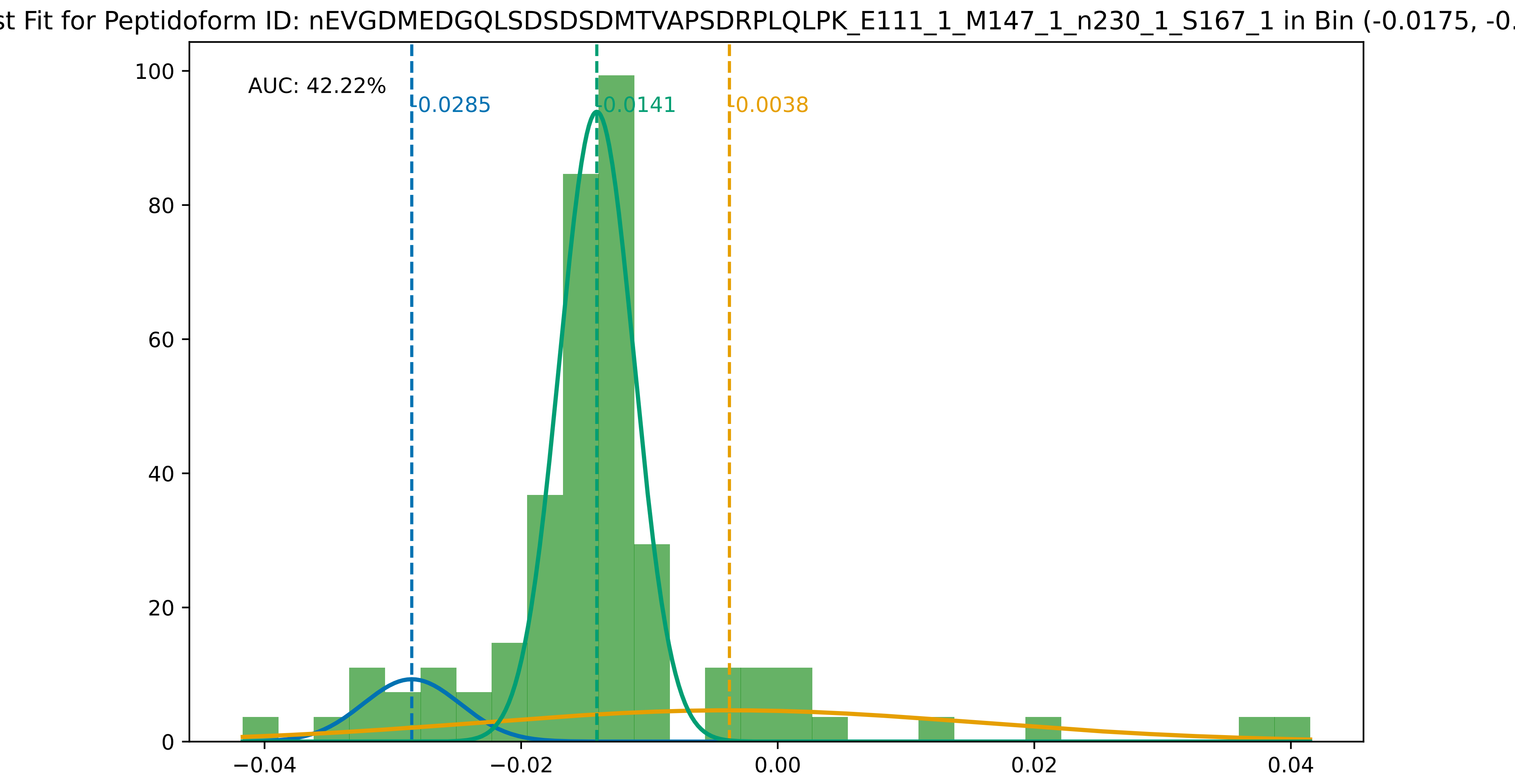
st Fit for Peptidoform ID: nEVGDMEDGQLSDSDSDMTVAPSDRPLQLPK_E111_1_M147_1_n230_1_S167_1 in Bin (-0.0175, -0.
AUC: 42.22%
-0.0285
-0.0141
-0.0038
100
80
60
40
20
0
−0.04
−0.02
0.00
0.02
0.04

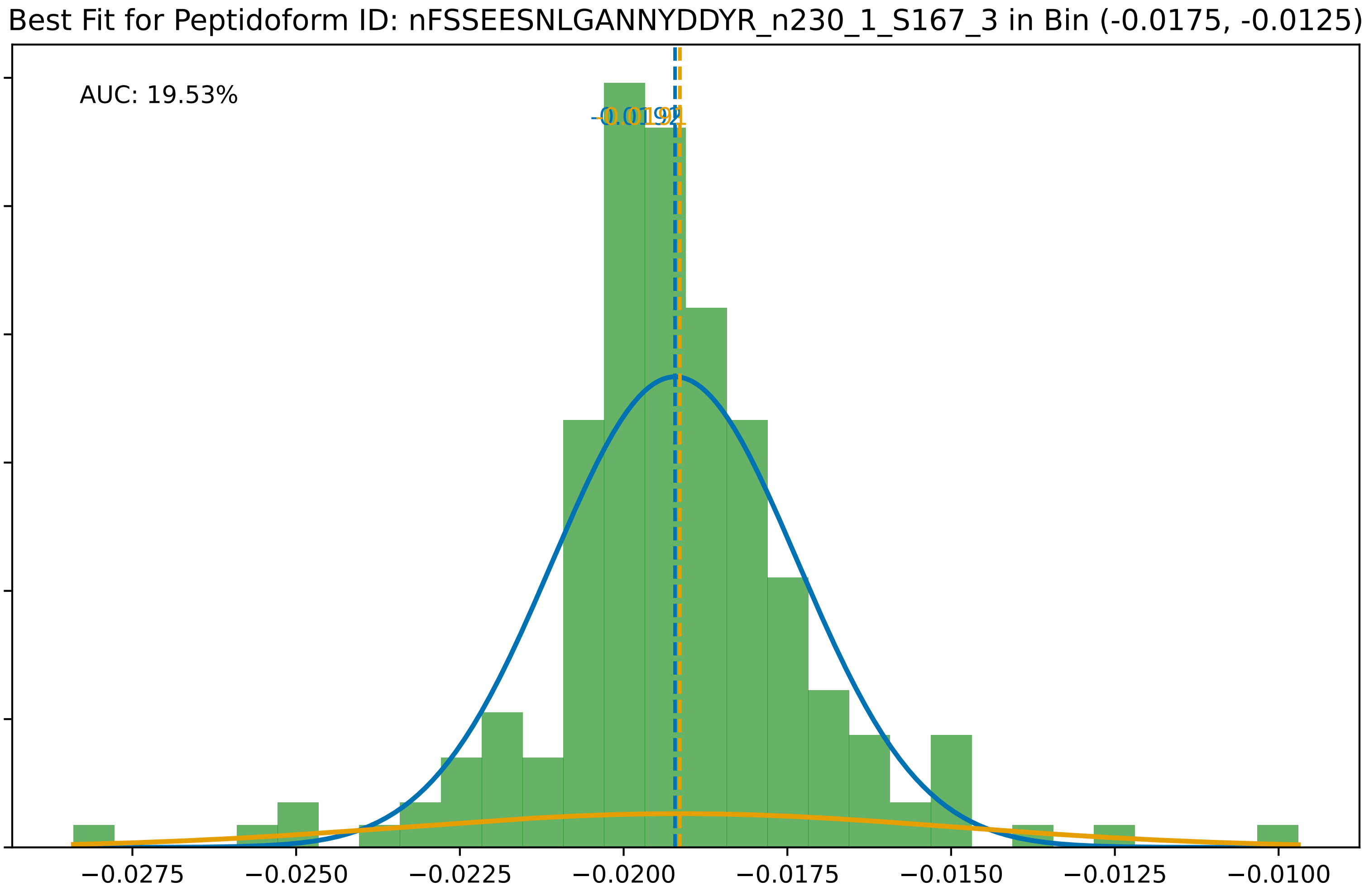
Best Fit for Peptidoform ID: nFSSEESNLGANNYDDYR_n230_1_S167_3 in Bin (-0.0175, -0.0125)
AUC: 19.53%
300
250
200
150
100
50
0
−0.0275
−0.0250
−0.0225
−0.0200
−0.0175
−0.0150
−0.0125
−0.0100

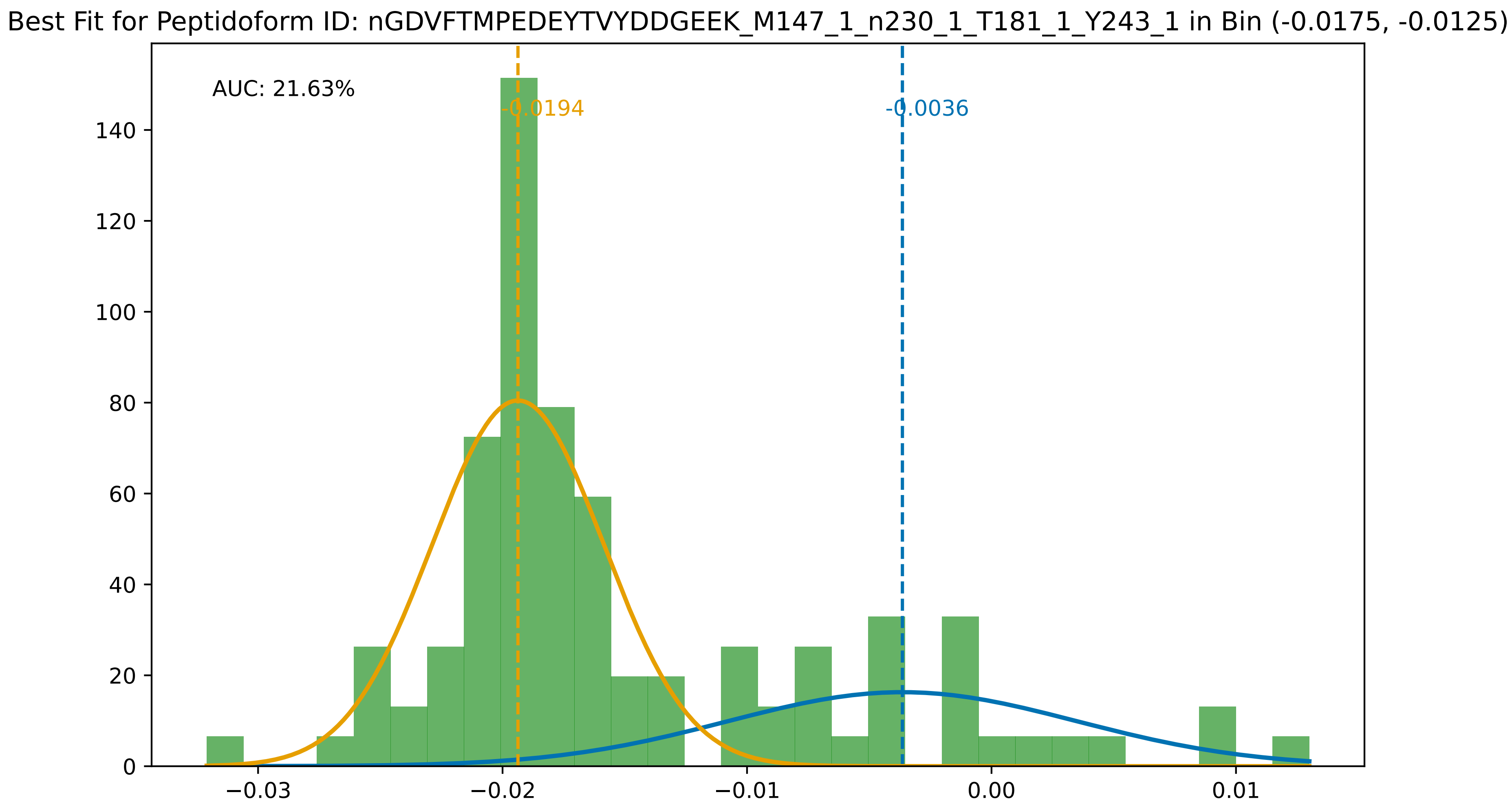

Best Fit for Peptidoform ID: nGDVFTMPEDEYTVYDDGEEK_M147_1_n230_1_T181_1_Y243_1 in Bin (-0.0175, -0.0125)
AUC: 21.63%
-0.0194
-0.0036
0
20
40
60
80
100
120
140
−0.03
−0.02
−0.01
0.00
0.01

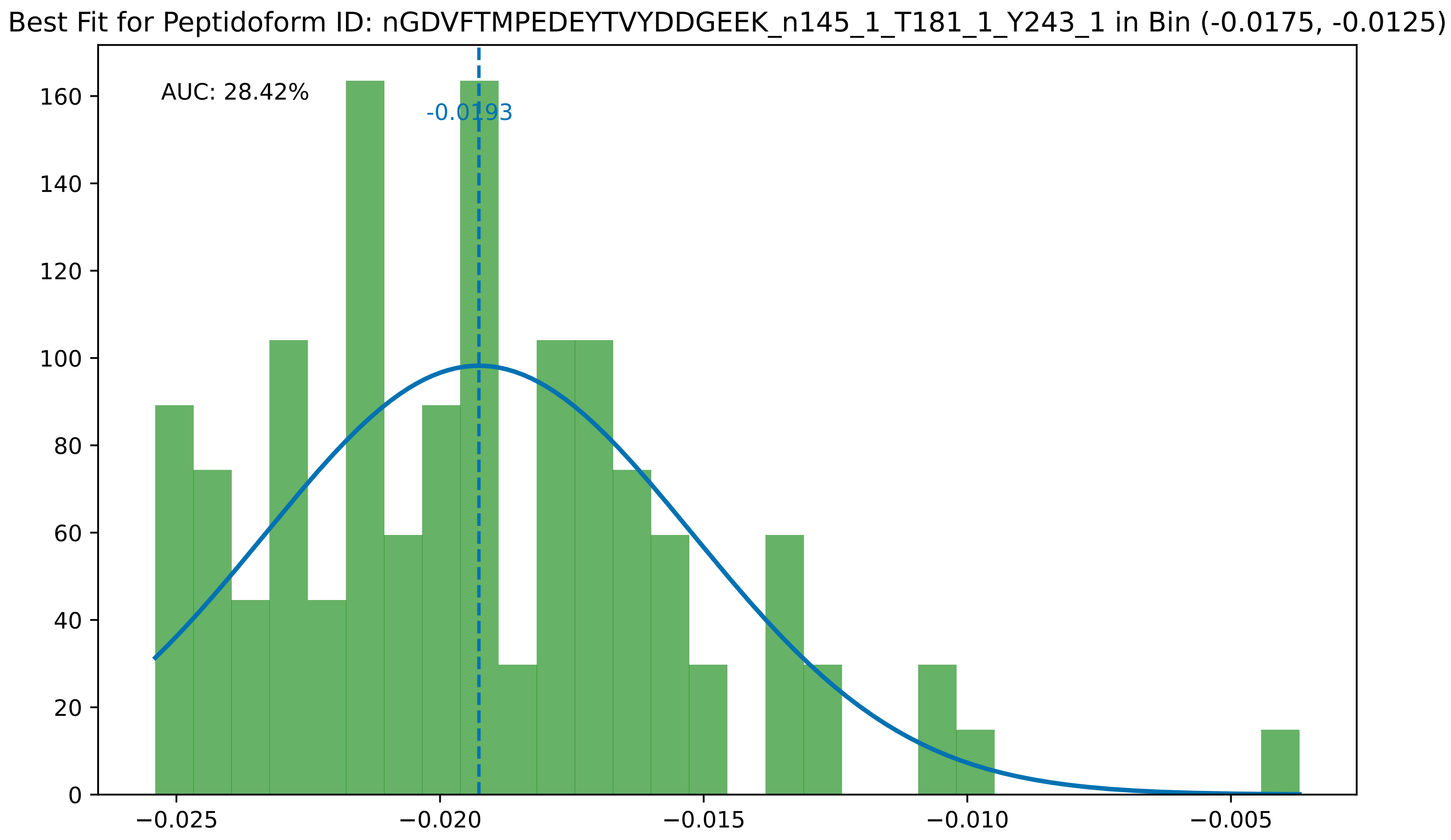

Best Fit for Peptidoform ID: nGDVFTMPEDEYTVYDDGEEK_n145_1_T181_1_Y243_1 in Bin (-0.0175, -0.0125)
AUC: 28.42%
-0.0193
0
20
40
60
80
100
120
140
160
−0.025
−0.020
−0.015
−0.010
−0.005

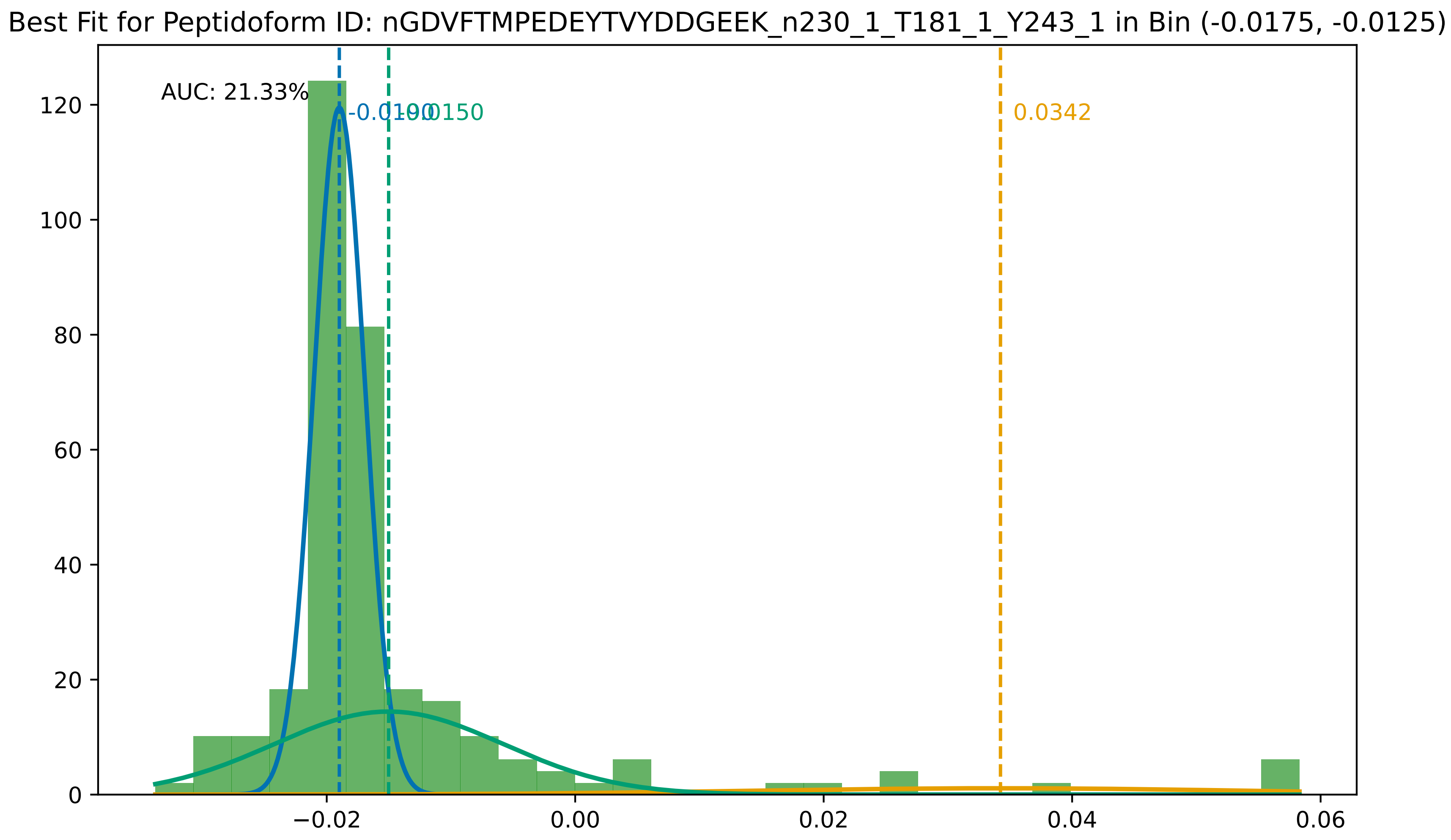

Best Fit for Peptidoform ID: nGDVFTMPEDEYTVYDDGEEK_n230_1_T181_1_Y243_1 in Bin (-0.0175, -0.0125)
AUC: 21.33%
-0.0190
-0.0150
0.0342
0
20
40
60
80
100
120
−0.02
0.00
0.02
0.04
0.06

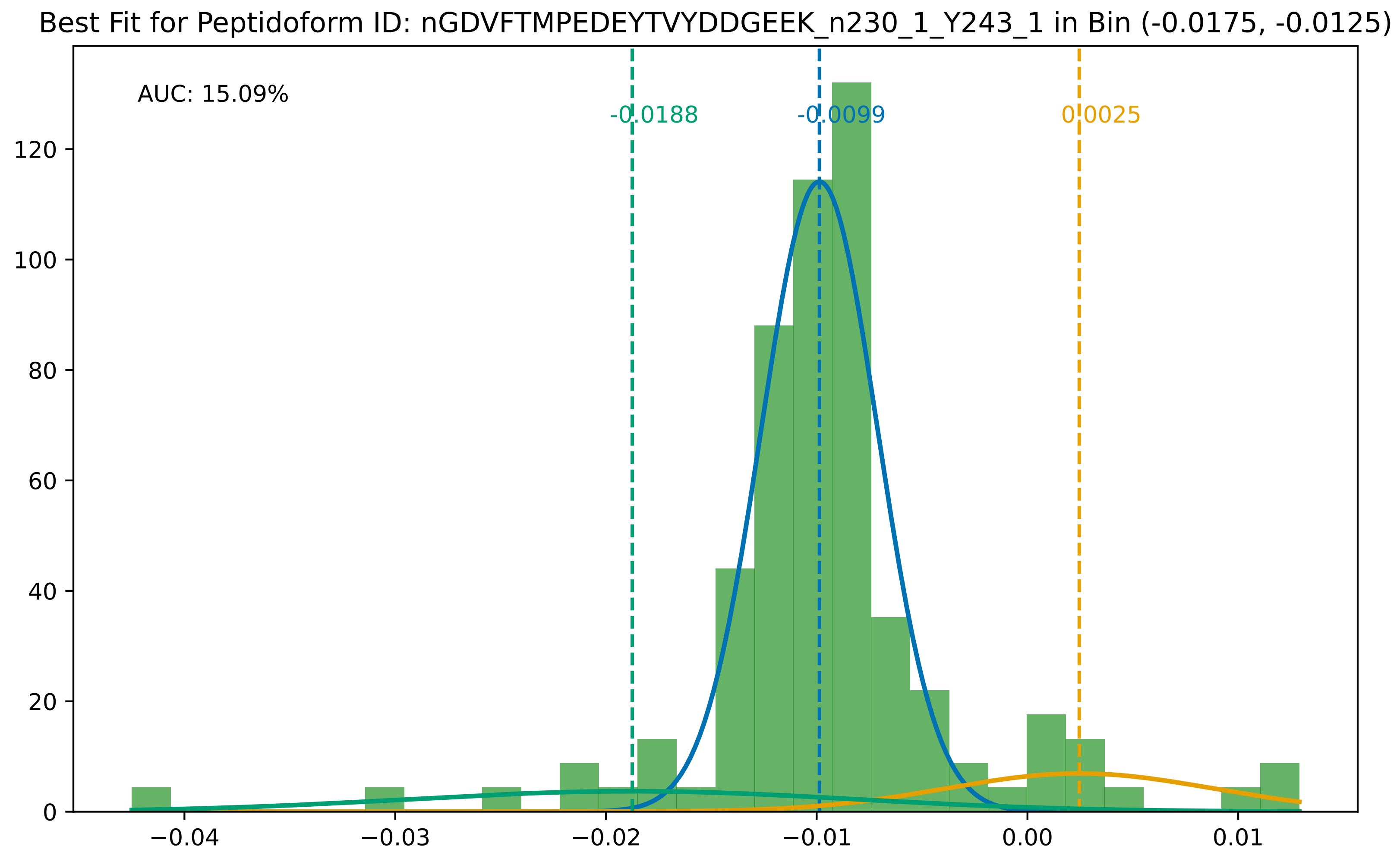

Best Fit for Peptidoform ID: nGDVFTMPEDEYTVYDDGEEK_n230_1_Y243_1 in Bin (-0.0175, -0.0125)
AUC: 15.09%
-0.0188
-0.0099
0.0025
0
20
40
60
80
100
120
−0.04
−0.03
−0.02
−0.01
0.00
0.01

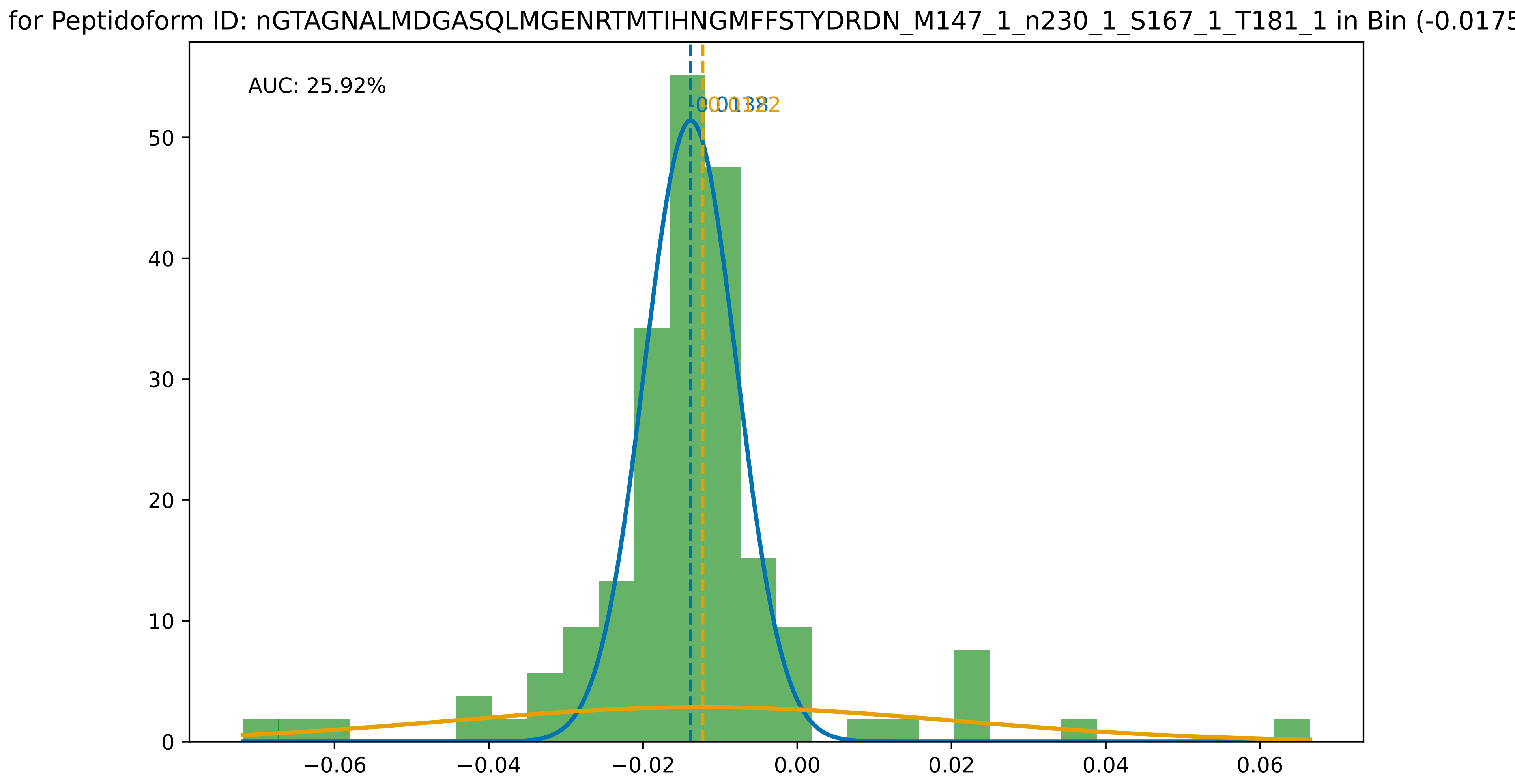
for Peptidoform ID: nGTAGNALMDGASQLMGENRTMTIHNGMFFSTYDRDN_M147_1_n230_1_S167_1_T181_1 in Bin (-0.0175
AUC: 25.92%
0.0138
0.0122
0
10
20
30
40
50
−0.06
−0.04
−0.02
0.00
0.02
0.04
0.06

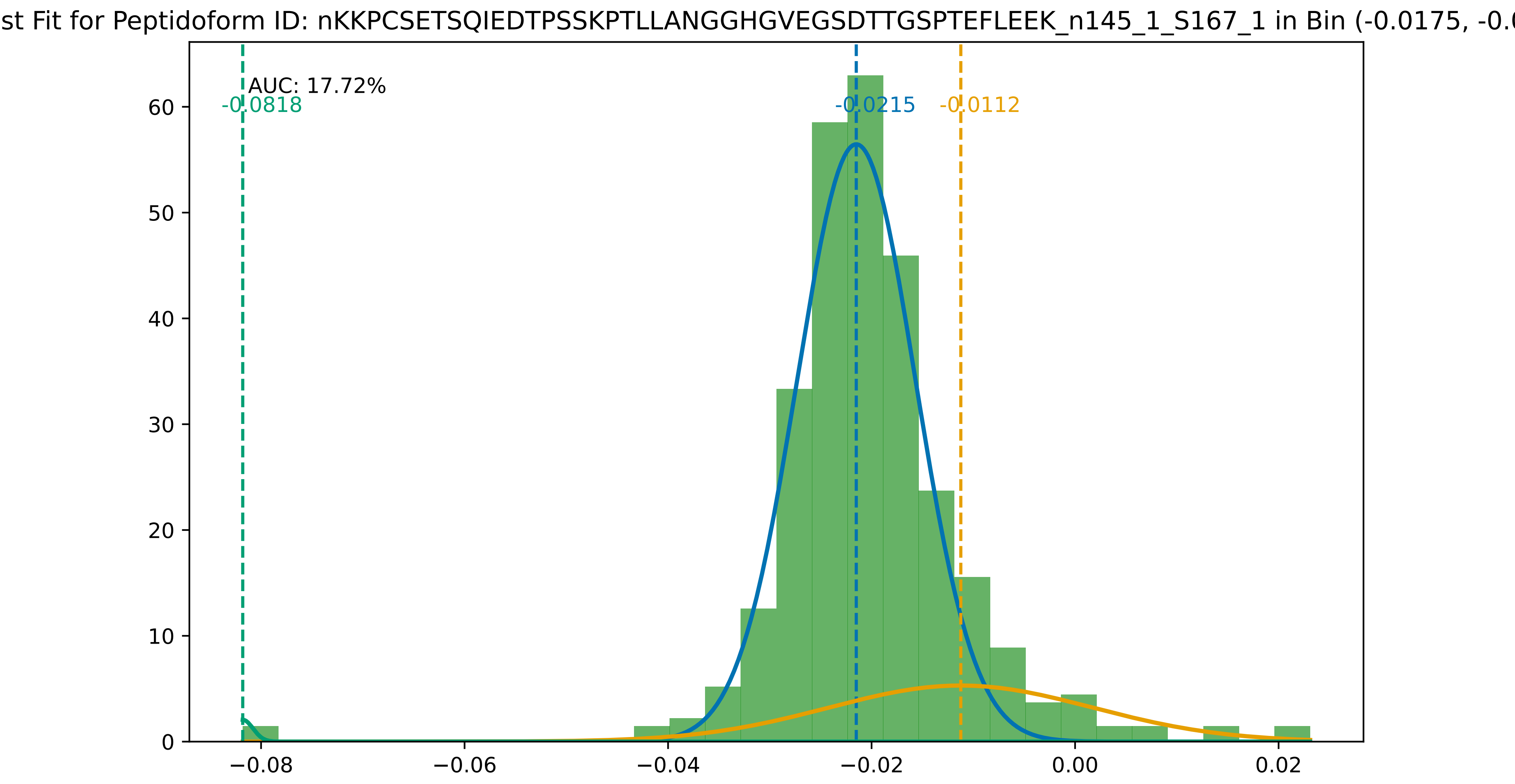

st Fit for Peptidoform ID: nKKPCSETSQIEDTPSSKPTLLANGGHGVEGSDTTGSPTEFLEEK_n145_1_S167_1 in Bin (-0.0175, -0.0
AUC: 17.72%
-0.0818
-0.0215
-0.0112
0
10
20
30
40
50
60
−0.08
−0.06
−0.04
−0.02
0.00
0.02

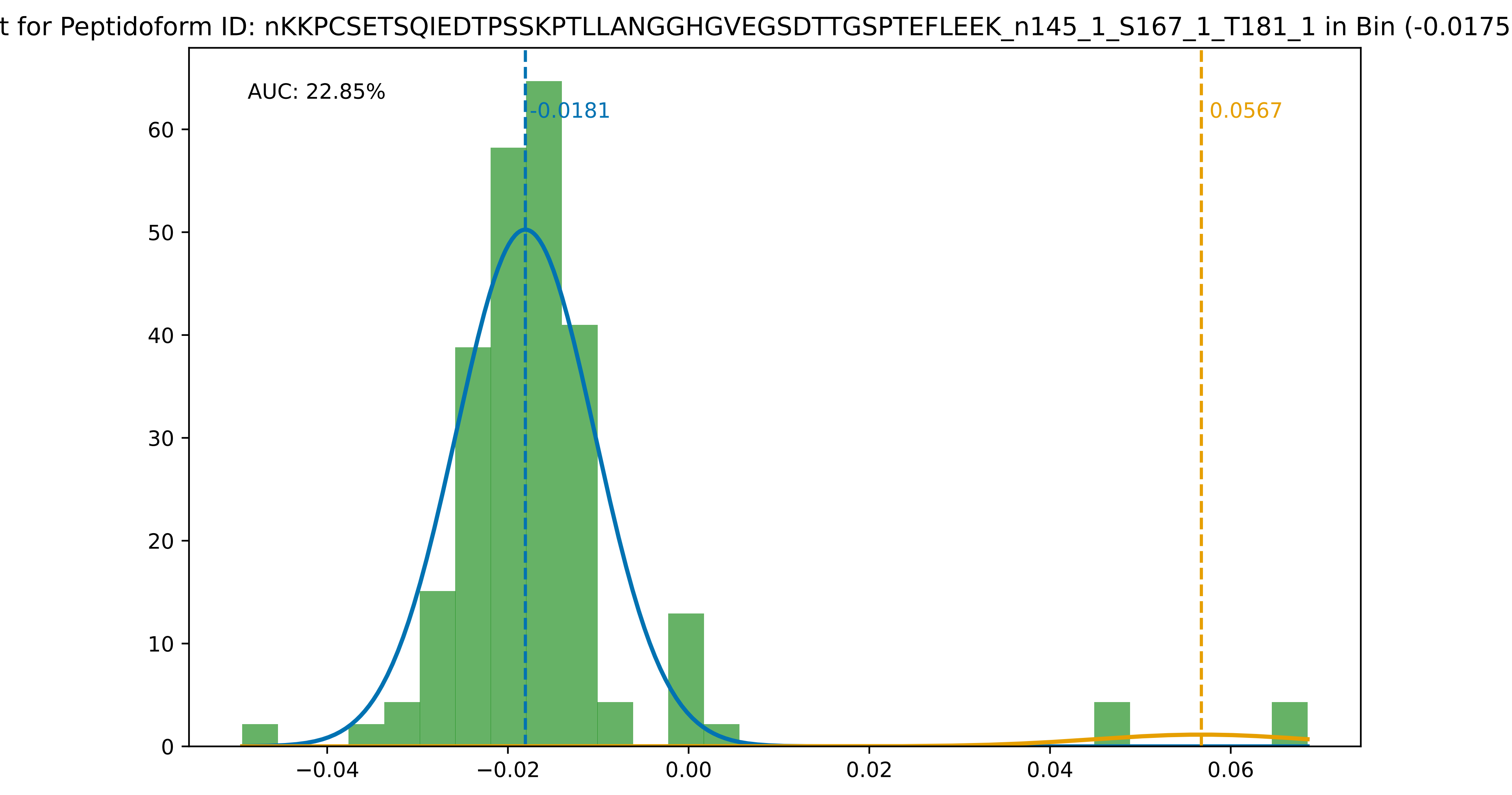
t for Peptidoform ID: nKKPCSETSQIEDTPSSKPTLLANGGHGVEGSDTTGSPTEFLEEK_n145_1_S167_1_T181_1 in Bin (-0.0175
AUC: 22.85%
-0.0181
0.0567
0
10
20
30
40
50
60
−0.04
−0.02
0.00
0.02
0.04
0.06

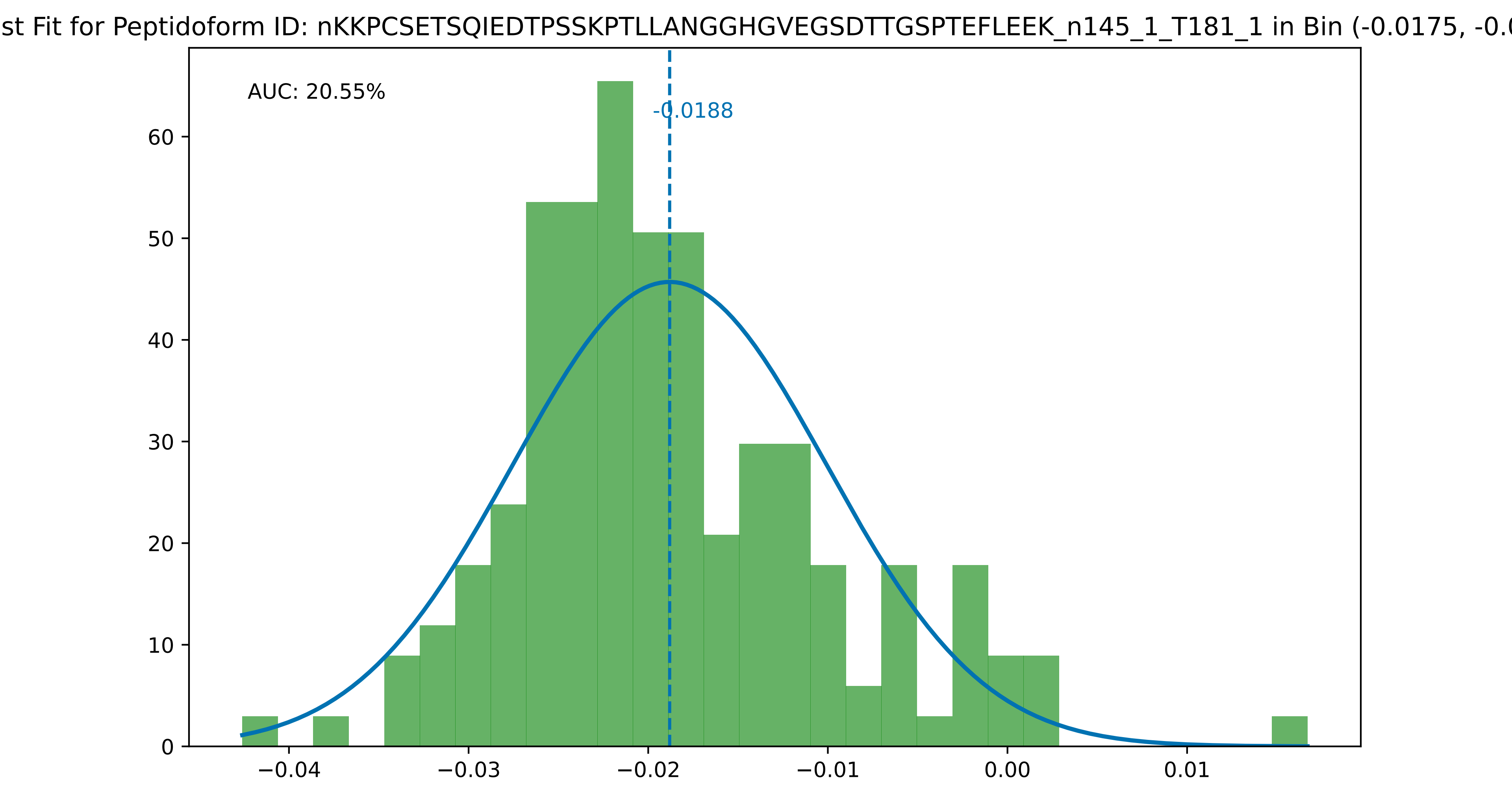
st Fit for Peptidoform ID: nKKPCSETSQIEDTPSSKPTLLANGGHGVEGSDTTGSPTEFLEEK_n145_1_T181_1 in Bin (-0.0175, -0.0
AUC: 20.55%
-0.0188
0
10
20
30
40
50
60
−0.04
−0.03
−0.02
−0.01
0.00
0.01

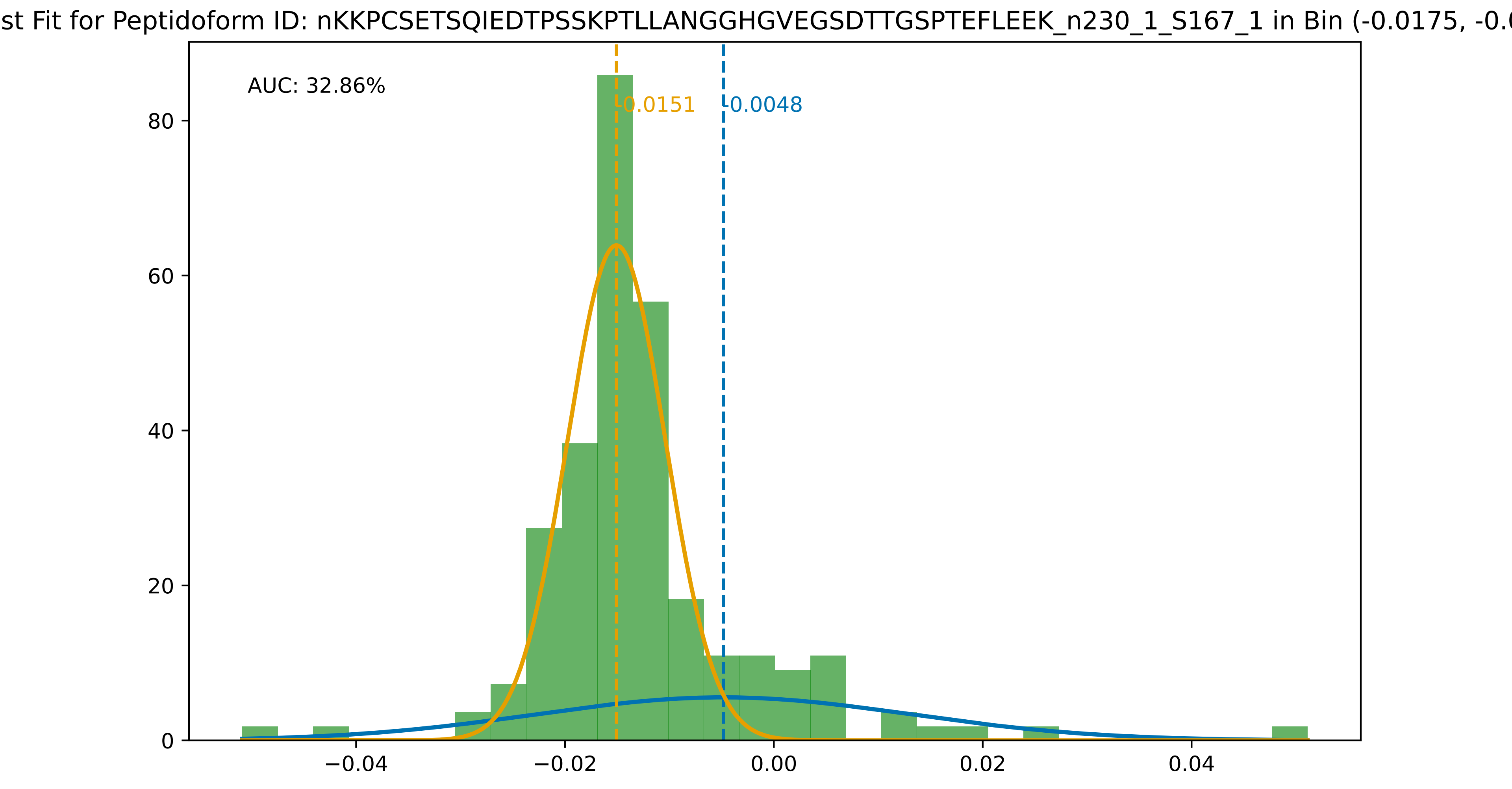
st Fit for Peptidoform ID: nKKPCSETSQIEDTPSSKPTLLANGGHGVEGSDTTGSPTEFLEEK_n230_1_S167_1 in Bin (-0.0175, -0.0
AUC: 32.86%
-0.0151
-0.0048
0
20
40
60
80
−0.04
−0.02
0.00
0.02
0.04

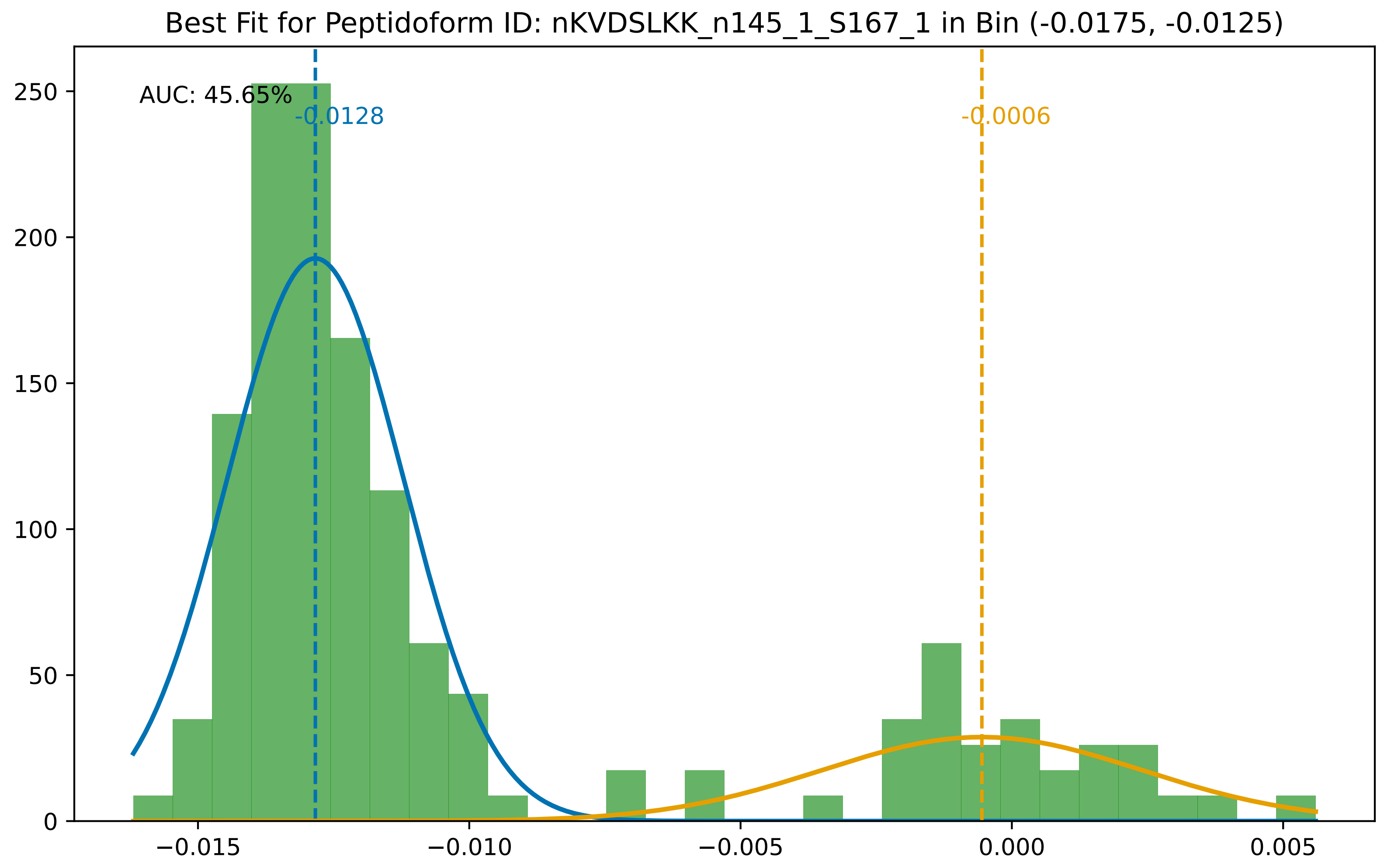

Best Fit for Peptidoform ID: nKVDSLKK_n145_1_S167_1 in Bin (-0.0175, -0.0125)
AUC: 45.65%
-0.0128
-0.0006
250
200
150
100
50
0
−0.015
−0.010
−0.005
0.000
0.005

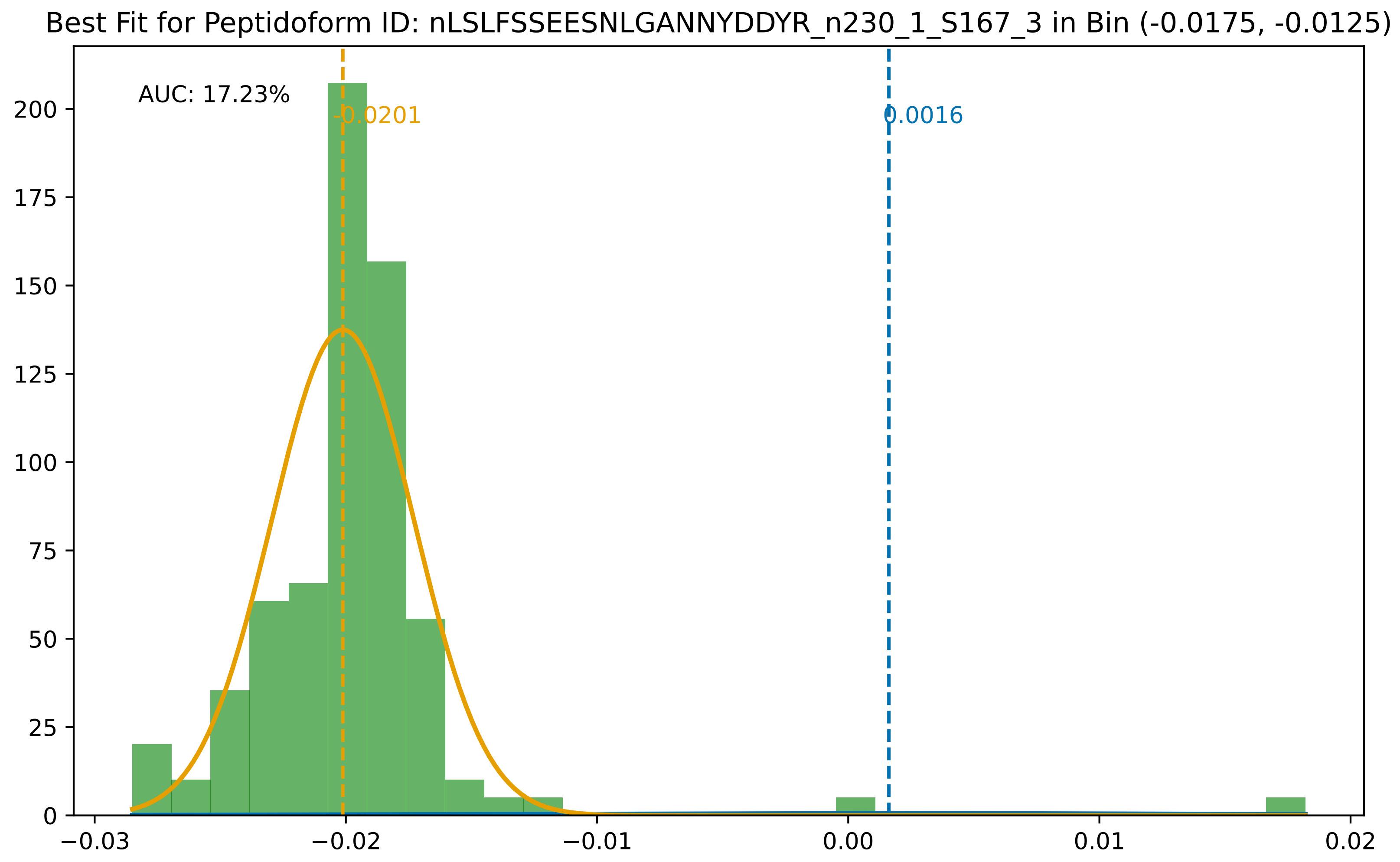

Best Fit for Peptidoform ID: nLSLFSSEESNLGANNYDDYR_n230_1_S167_3 in Bin (-0.0175, -0.0125)
AUC: 17.23%
-0.0201
0.0016
0
25
50
75
100
125
150
175
200
−0.03
−0.02
−0.01
0.00
0.01
0.02

for Peptidoform ID: nMPEDEYTVYDDGEEKNNATVHEQVGGPSLTSDLQAQSK_M147_1_n230_1_S167_1_T181_1 in Bin (-0.017

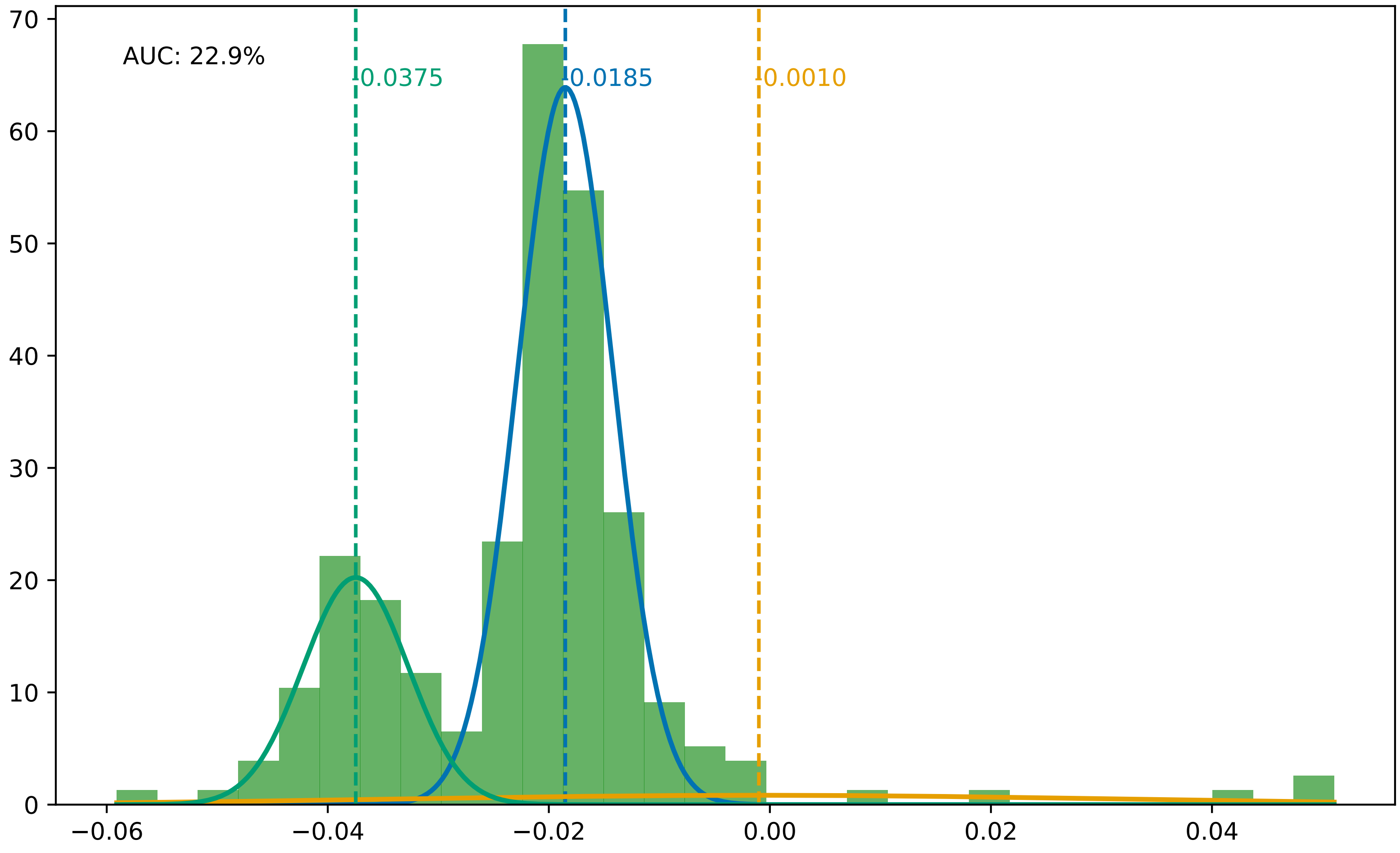

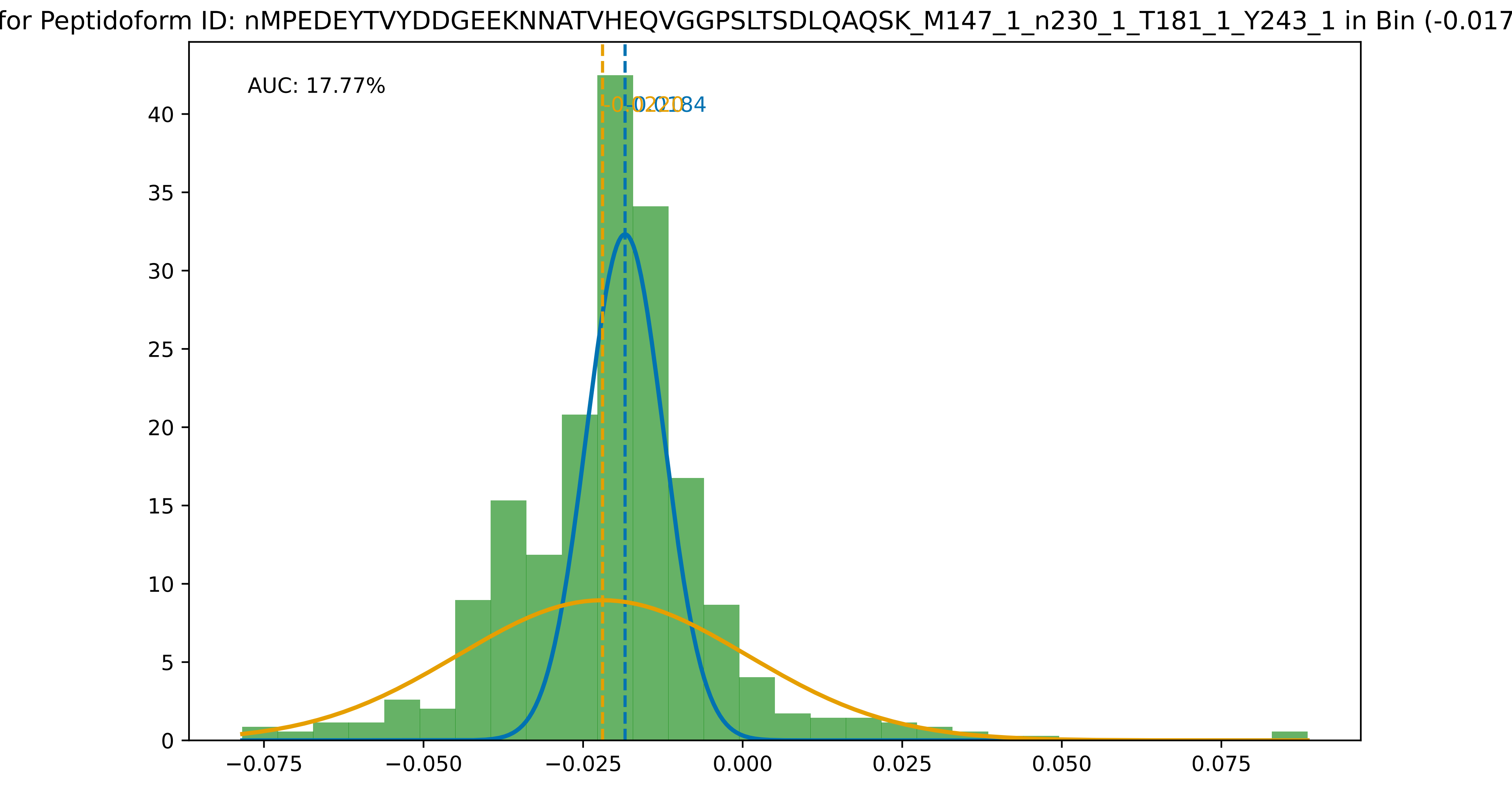
for Peptidoform ID: nMPEDEYTVYDDGEEKNNATVHEQVGGPSLTSDLQAQSK_M147_1_n230_1_T181_1_Y243_1 in Bin (-0.017
AUC: 17.77%
0
5
10
15
20
25
30
35
40
−0.075
−0.050
−0.025
0.000
0.025
0.050
0.075

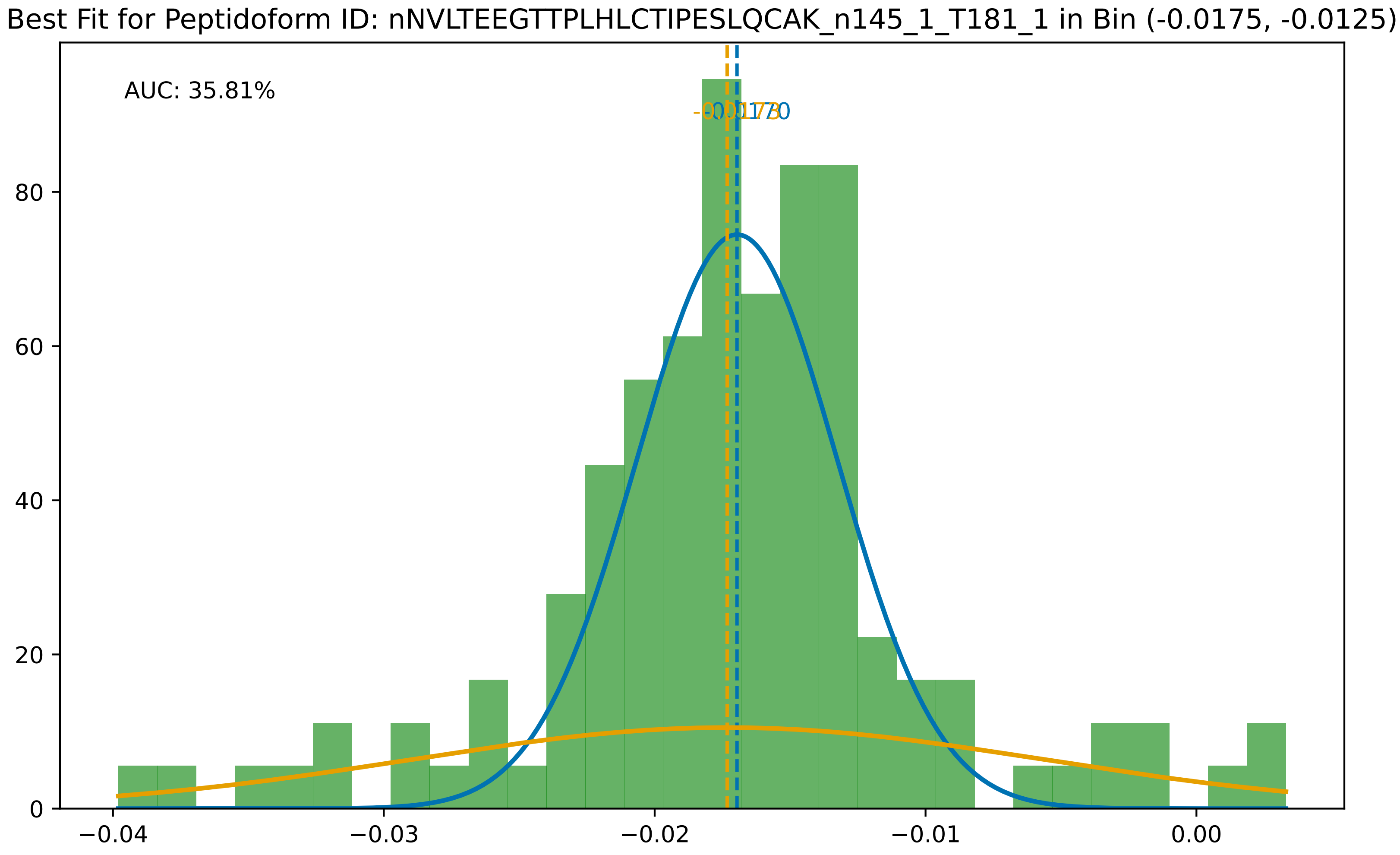

Best Fit for Peptidoform ID: nNVLTEEGTTPLHLCTIPESLQCAK_n145_1_T181_1 in Bin (-0.0175, -0.0125)
AUC: 35.81%
80
60
40
20
0
−0.04
−0.03
−0.02
−0.01
0.00

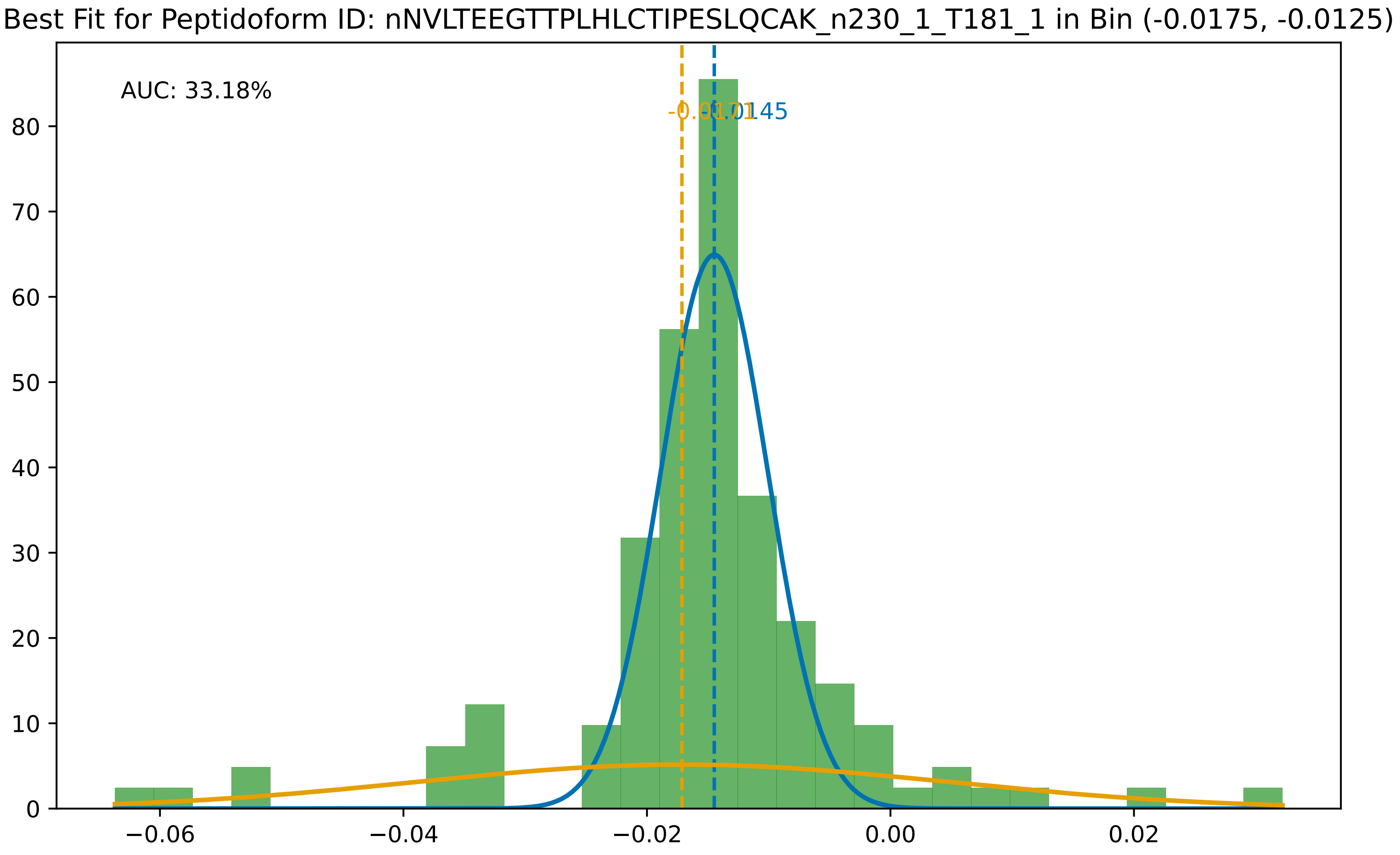
Best Fit for Peptidoform ID: nNVLTEEGTTPLHLCTIPESLQCAK_n230_1_T181_1 in Bin (-0.0175, -0.0125)
AUC: 33.18%
-0.0171
-0.0145
0
10
20
30
40
50
60
70
80
−0.06
−0.04
−0.02
0.00
0.02

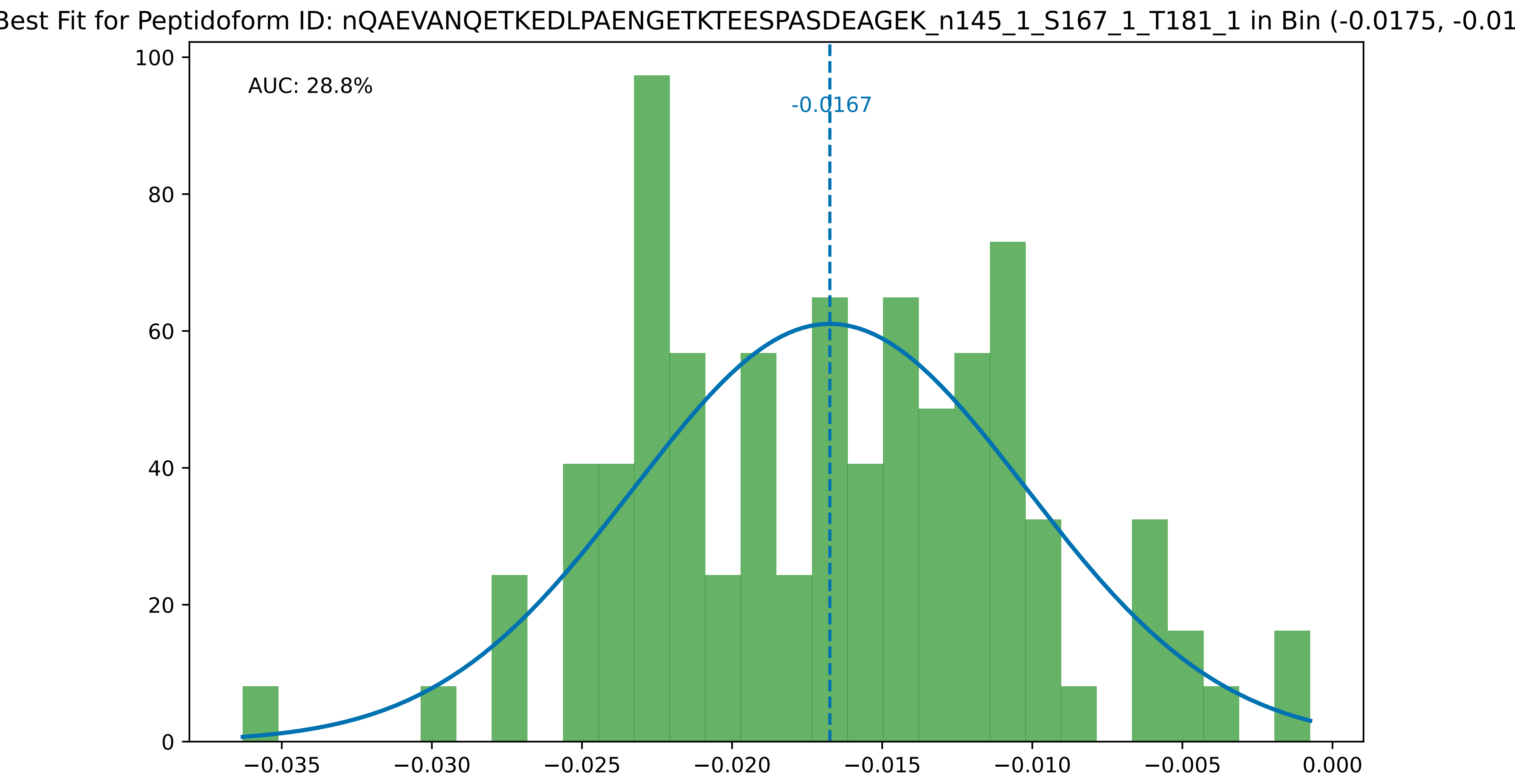
Best Fit for Peptidoform ID: nQAEVANQETKEDLPAENGETKTEESPASDEAGEK_n145_1_S167_1_T181_1 in Bin (-0.0175, -0.01
AUC: 28.8%
-0.0167
100
80
60
40
20
0
−0.035
−0.030
−0.025
−0.020
−0.015
−0.010
−0.005
0.000

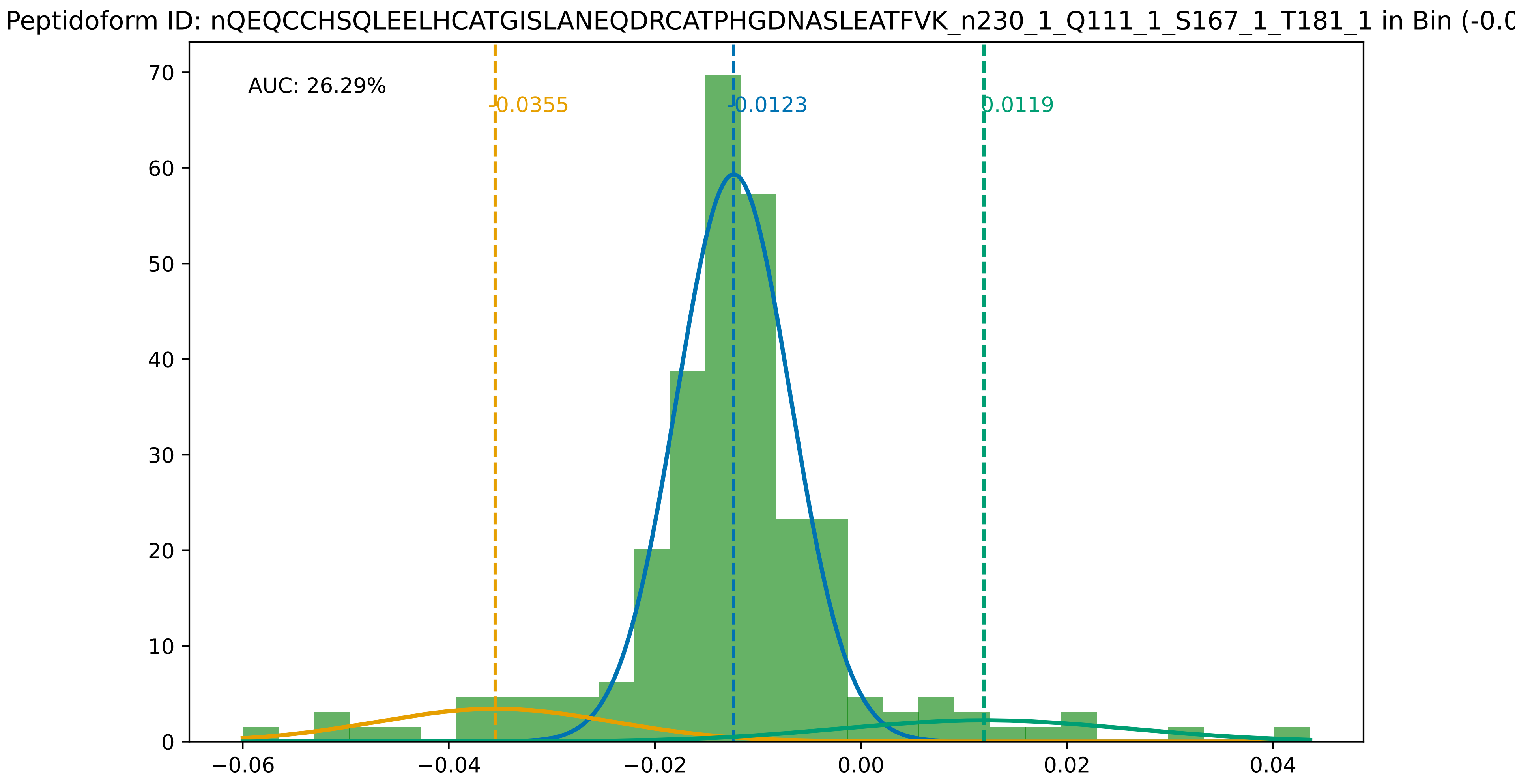

Peptidoform ID: nQEQCCHSQLEELHCATGISLANEQDRCATPHGDNASLEATFVK_n230_1_Q111_1_S167_1_T181_1 in Bin (-0.0
AUC: 26.29%
-0.0355
-0.0123
0.0119
0
10
20
30
40
50
60
70
−0.06
−0.04
−0.02
0.00
0.02
0.04

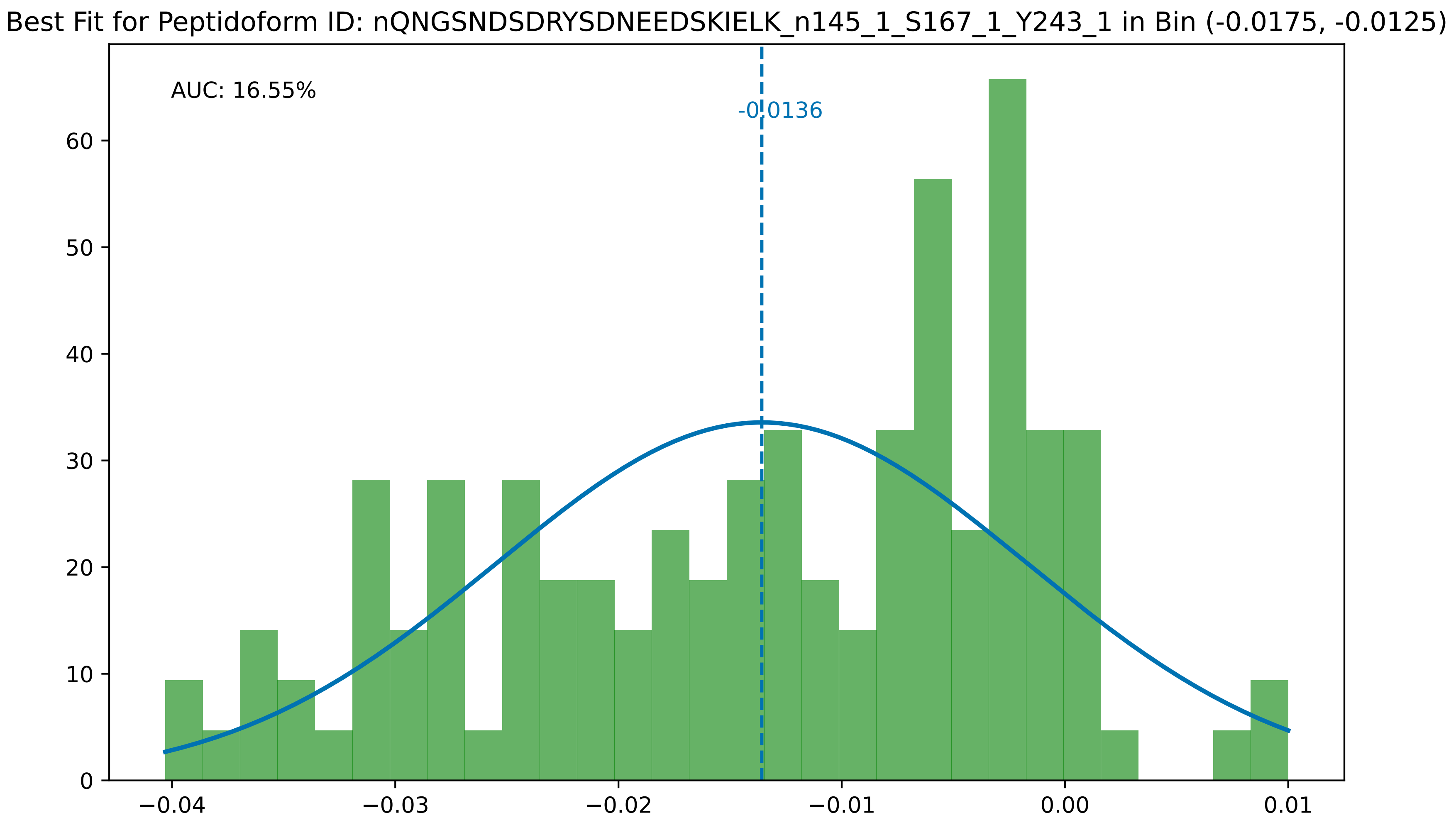
Best Fit for Peptidoform ID: nQNGSNDSDRYSDNEEDSKIELK_n145_1_S167_1_Y243_1 in Bin (-0.0175, -0.0125)
AUC: 16.55%
-0.0136
60
50
40
30
20
10
0
−0.04
−0.03
−0.02
−0.01
0.00
0.01

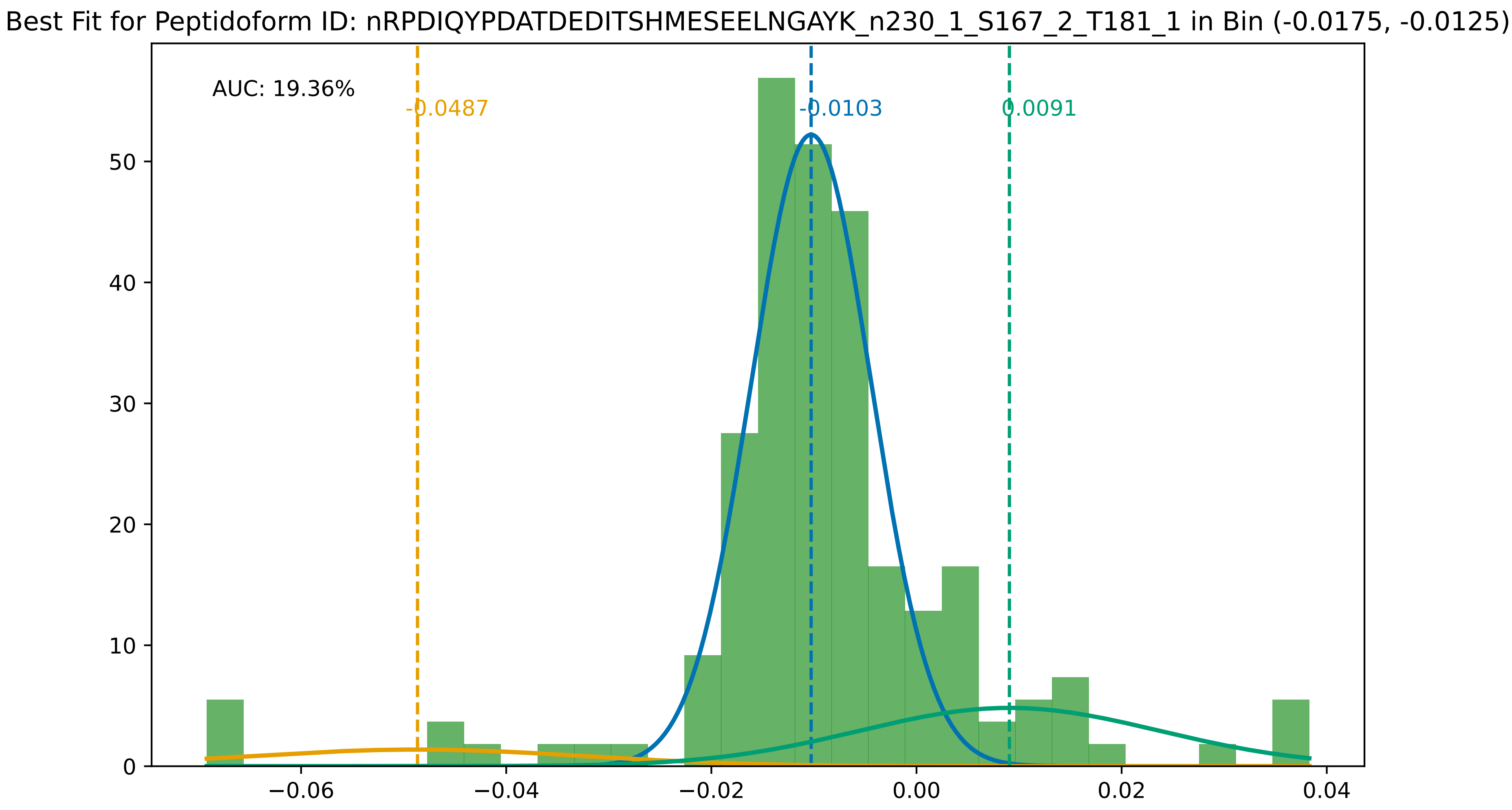
Best Fit for Peptidoform ID: nRPDIQYPDATDEDITSHMESEELNGAYK_n230_1_S167_2_T181_1 in Bin (-0.0175, -0.0125)
AUC: 19.36%
-0.0487
-0.0103
0.0091
0
10
20
30
40
50
−0.06
−0.04
−0.02
0.00
0.02
0.04

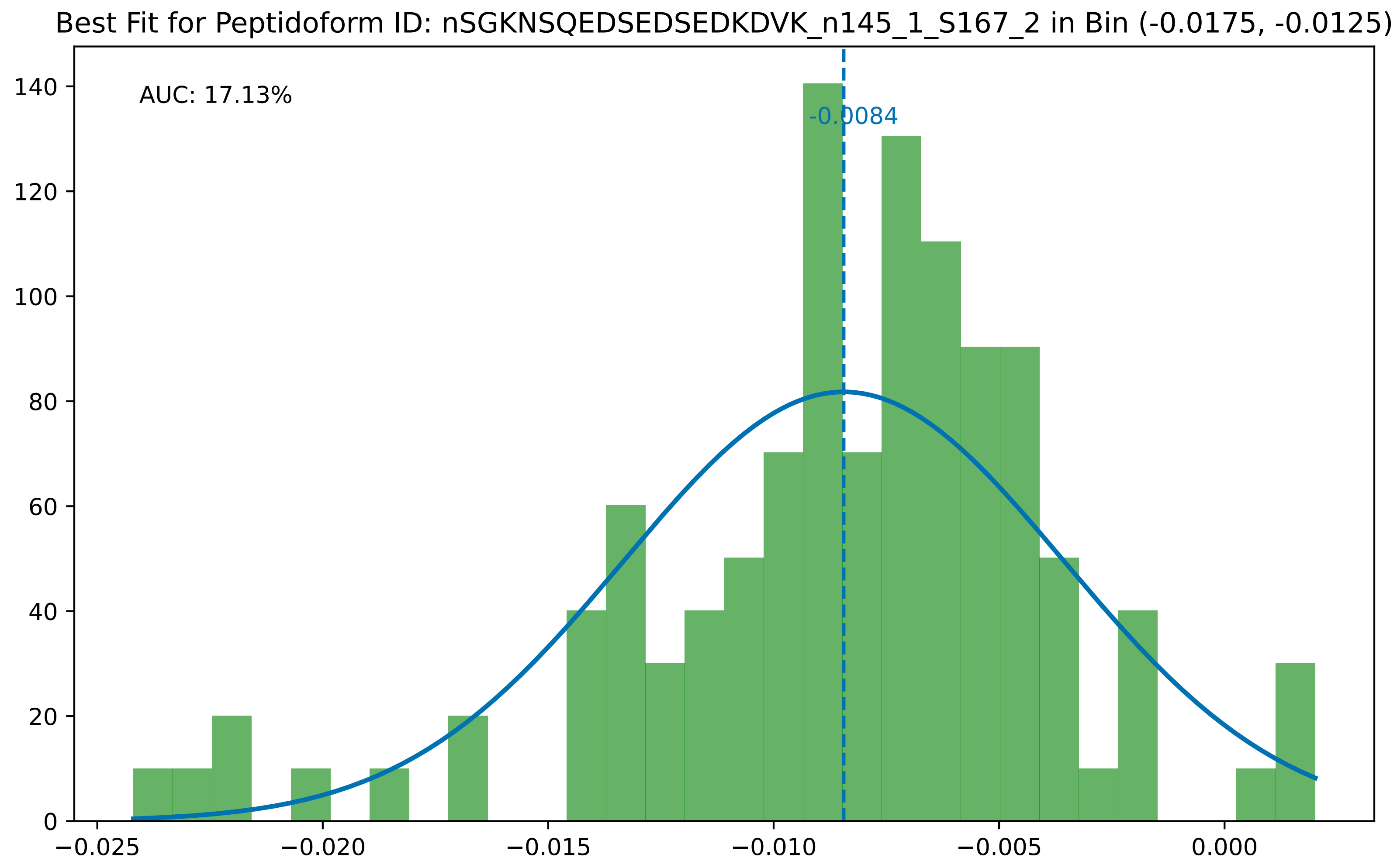
Best Fit for Peptidoform ID: nSGKNSQEDSEDSEDKDVK_n145_1_S167_2 in Bin (-0.0175, -0.0125)
AUC: 17.13%
-0.0084
0
20
40
60
80
100
120
140
−0.025
−0.020
−0.015
−0.010
−0.005
0.000

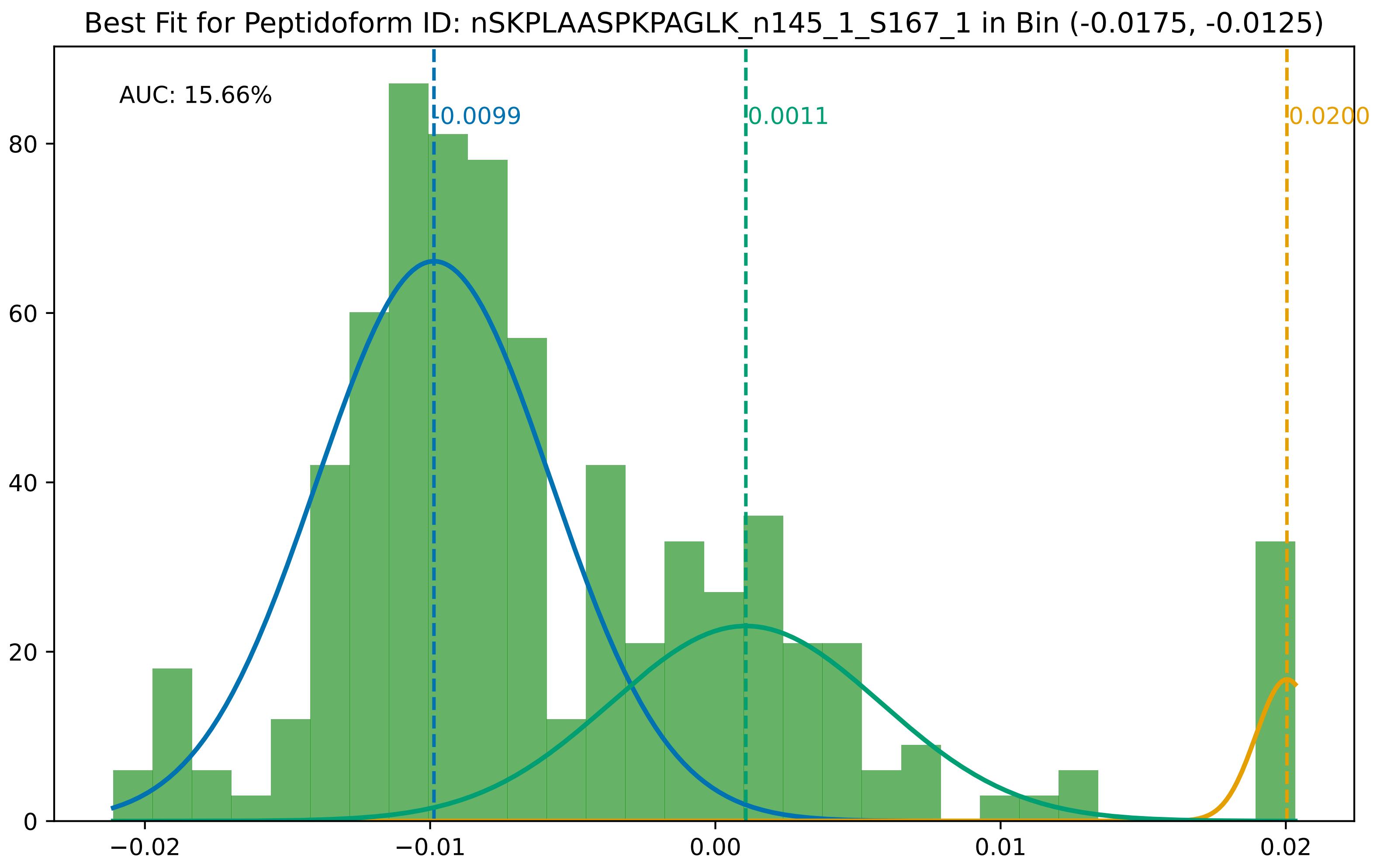

Best Fit for Peptidoform ID: nSKPLAASPKPAGLK_n145_1_S167_1 in Bin (-0.0175, -0.0125)
AUC: 15.66%
-0.0099
0.0011
0.0200
80
60
40
20
0
−0.02
−0.01
0.00
0.01
0.02

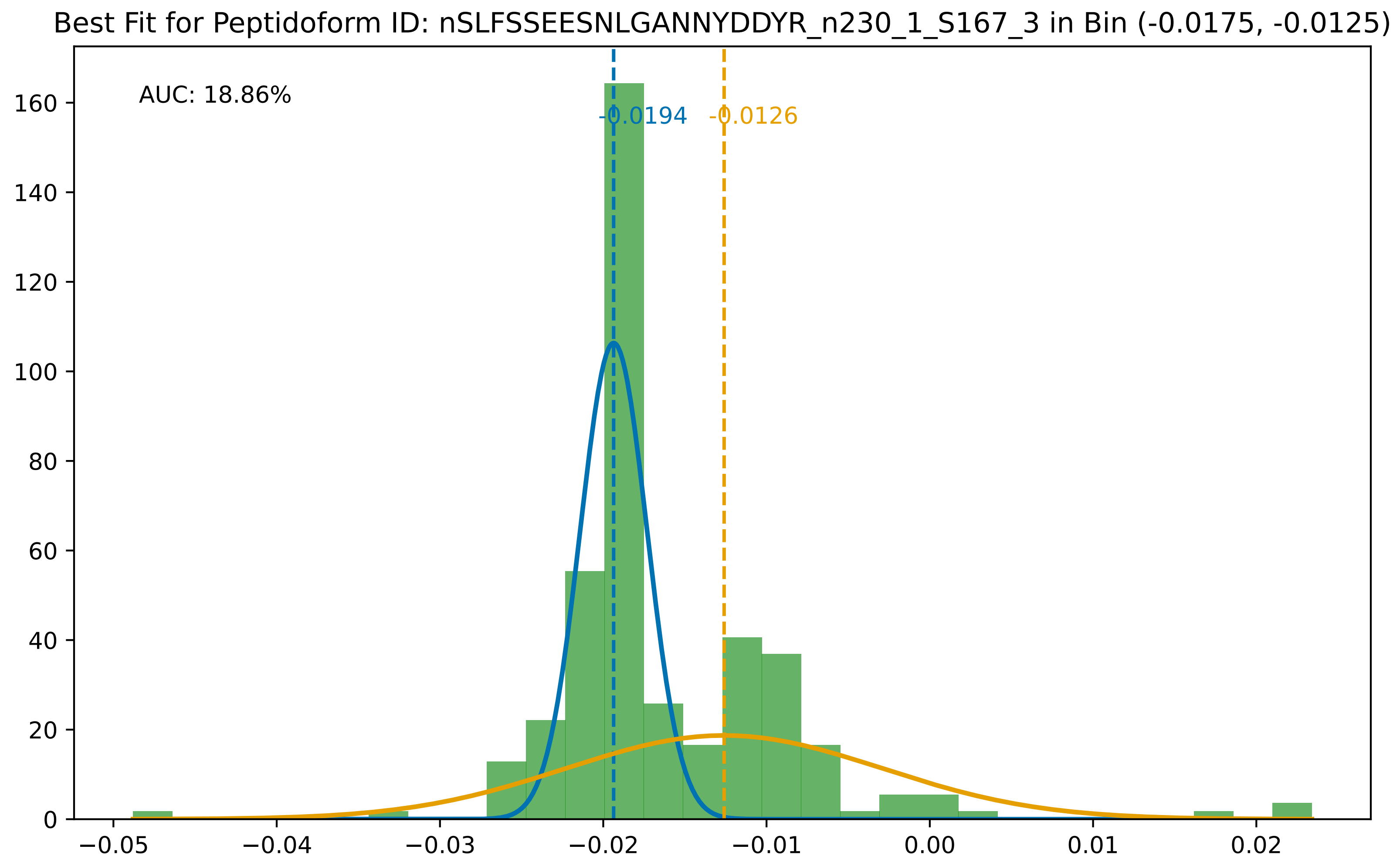

Best Fit for Peptidoform ID: nSLFSSEESNLGANNYDDYR_n230_1_S167_3 in Bin (-0.0175, -0.0125)
AUC: 18.86%
-0.0194
-0.0126
0
20
40
60
80
100
120
140
160
−0.05
−0.04
−0.03
−0.02
−0.01
0.00
0.01
0.02

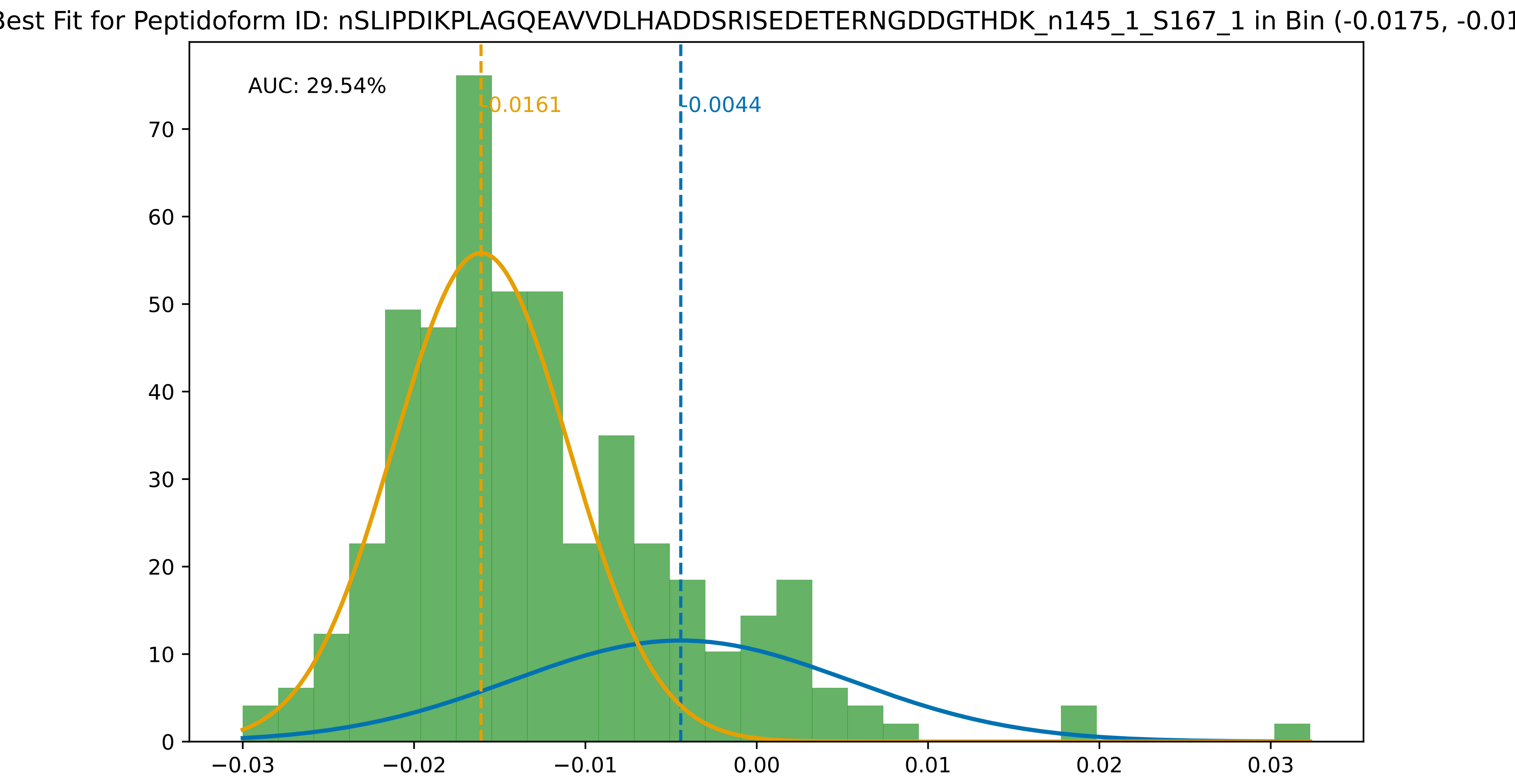

Best Fit for Peptidoform ID: nSLIPDIKPLAGQEAVVDLHADDSRISEDETERNGDDGTHDK_n145_1_S167_1 in Bin (-0.0175, -0.01
AUC: 29.54%
-0.0161
-0.0044
0
10
20
30
40
50
60
70
−0.03
−0.02
−0.01
0.00
0.01
0.02
0.03

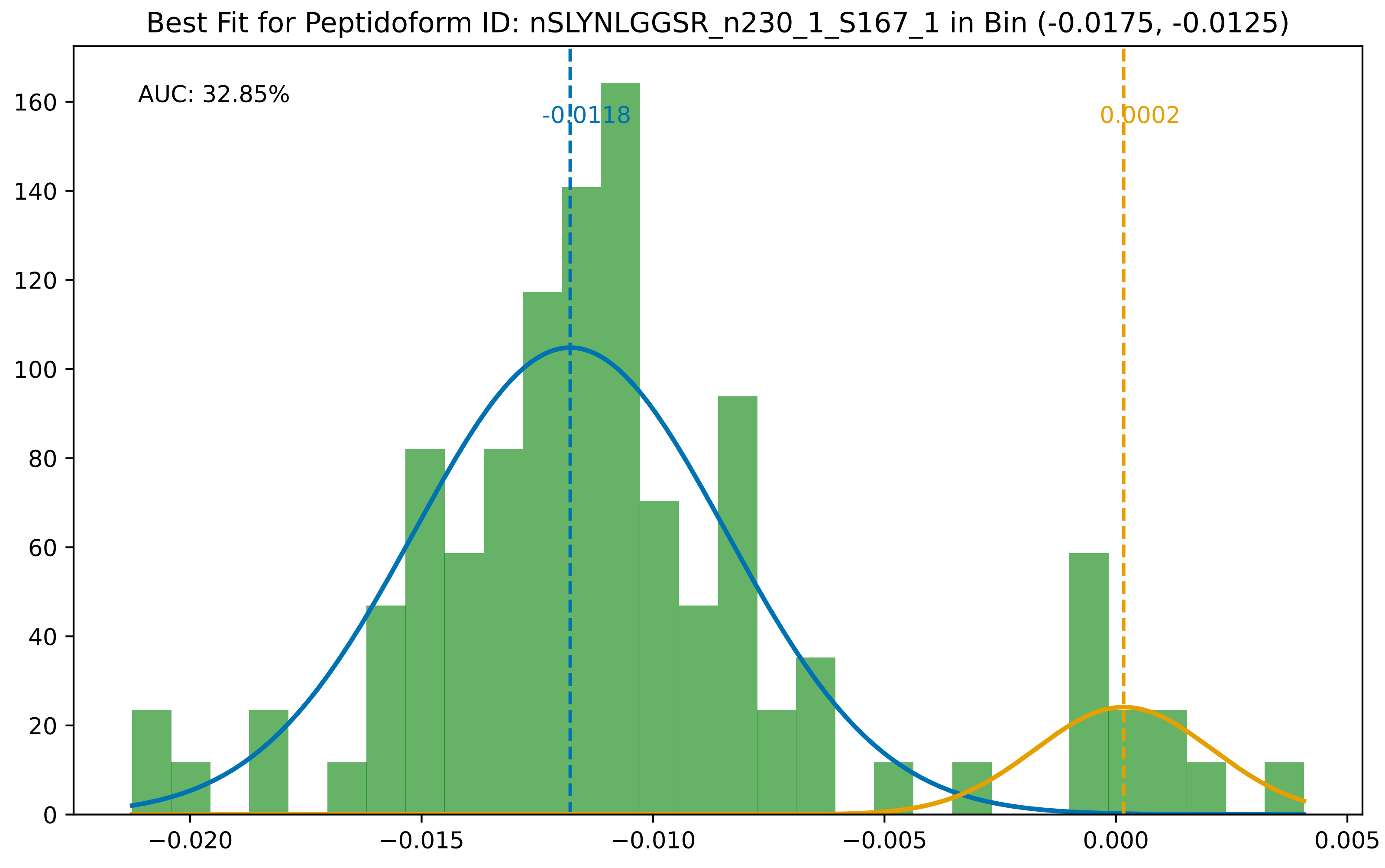
Best Fit for Peptidoform ID: nSLYNLGGSR_n230_1_S167_1 in Bin (-0.0175, -0.0125)
AUC: 32.85%
-0.0118
0.0002
0
20
40
60
80
100
120
140
160
−0.020
−0.015
−0.010
−0.005
0.000
0.005

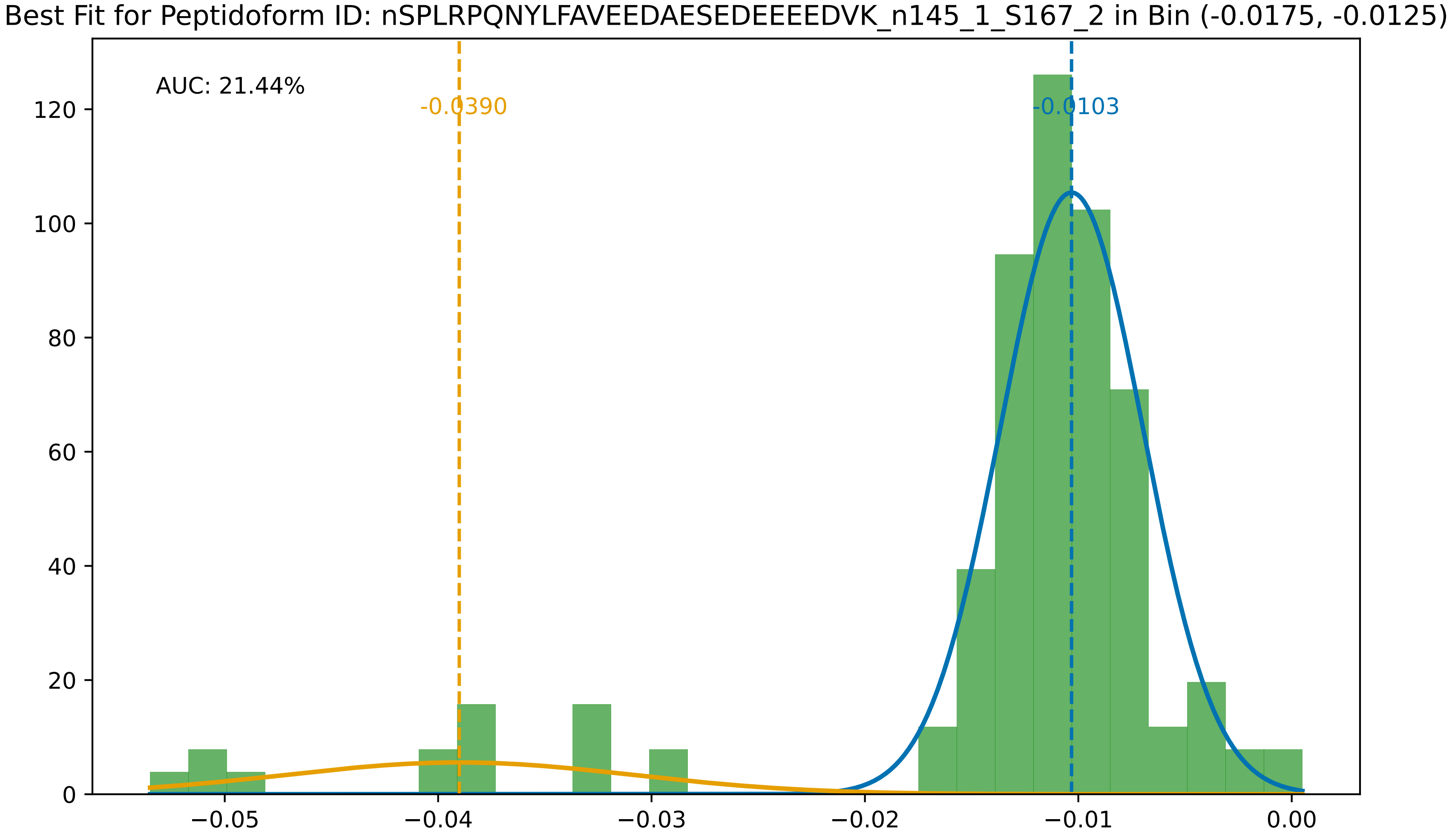

Best Fit for Peptidoform ID: nSPLRPQNYLFAVEEDAESEDEEEEDVK_n145_1_S167_2 in Bin (-0.0175, -0.0125)
AUC: 21.44%
-0.0390
-0.0103
0
20
40
60
80
100
120
−0.05
−0.04
−0.03
−0.02
−0.01
0.00

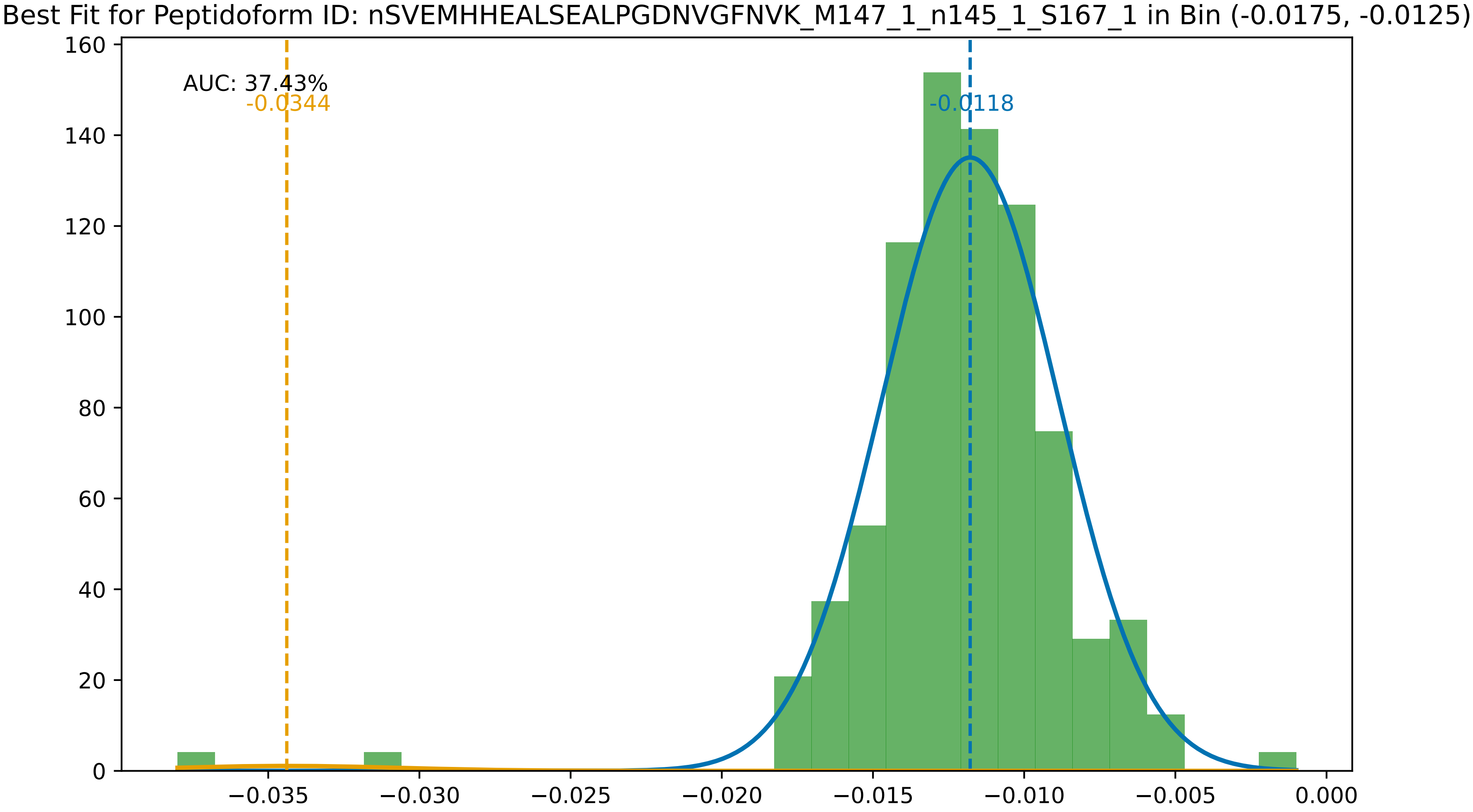

Best Fit for Peptidoform ID: nSVEMHHEALSEALPGDNVGFNVK_M147_1_n145_1_S167_1 in Bin (-0.0175, -0.0125)
AUC: 37.43%
-0.0344
-0.0118
0
20
40
60
80
100
120
140
160
−0.035
−0.030
−0.025
−0.020
−0.015
−0.010
−0.005
0.000

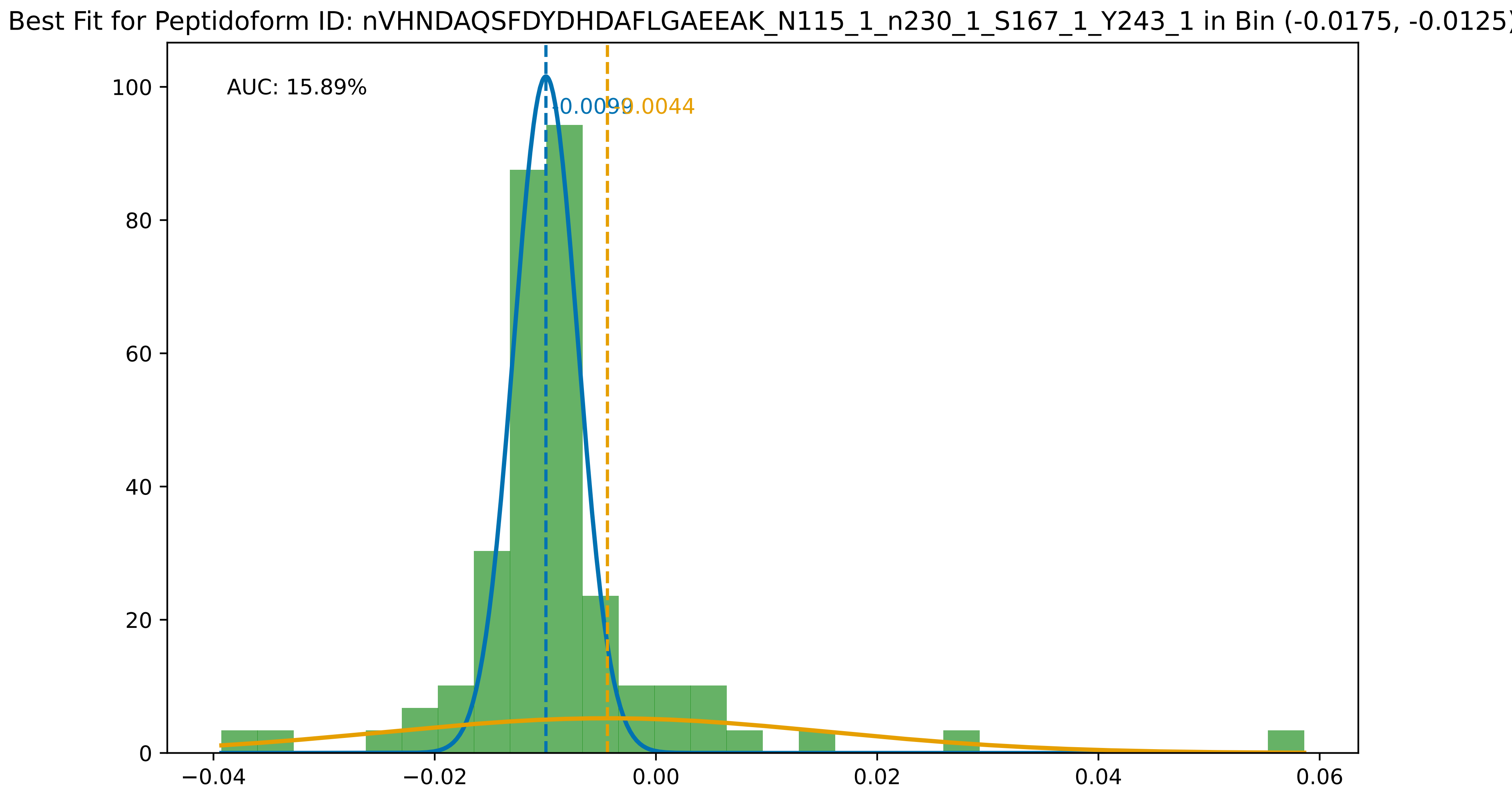
Best Fit for Peptidoform ID: nVHNDAQSFDYDHDAFLGAEEAK_N115_1_n230_1_S167_1_Y243_1 in Bin (-0.0175, -0.0125)
AUC: 15.89%
-0.0099
-0.0044
100
80
60
40
20
0
−0.04
−0.02
0.00
0.02
0.04
0.06

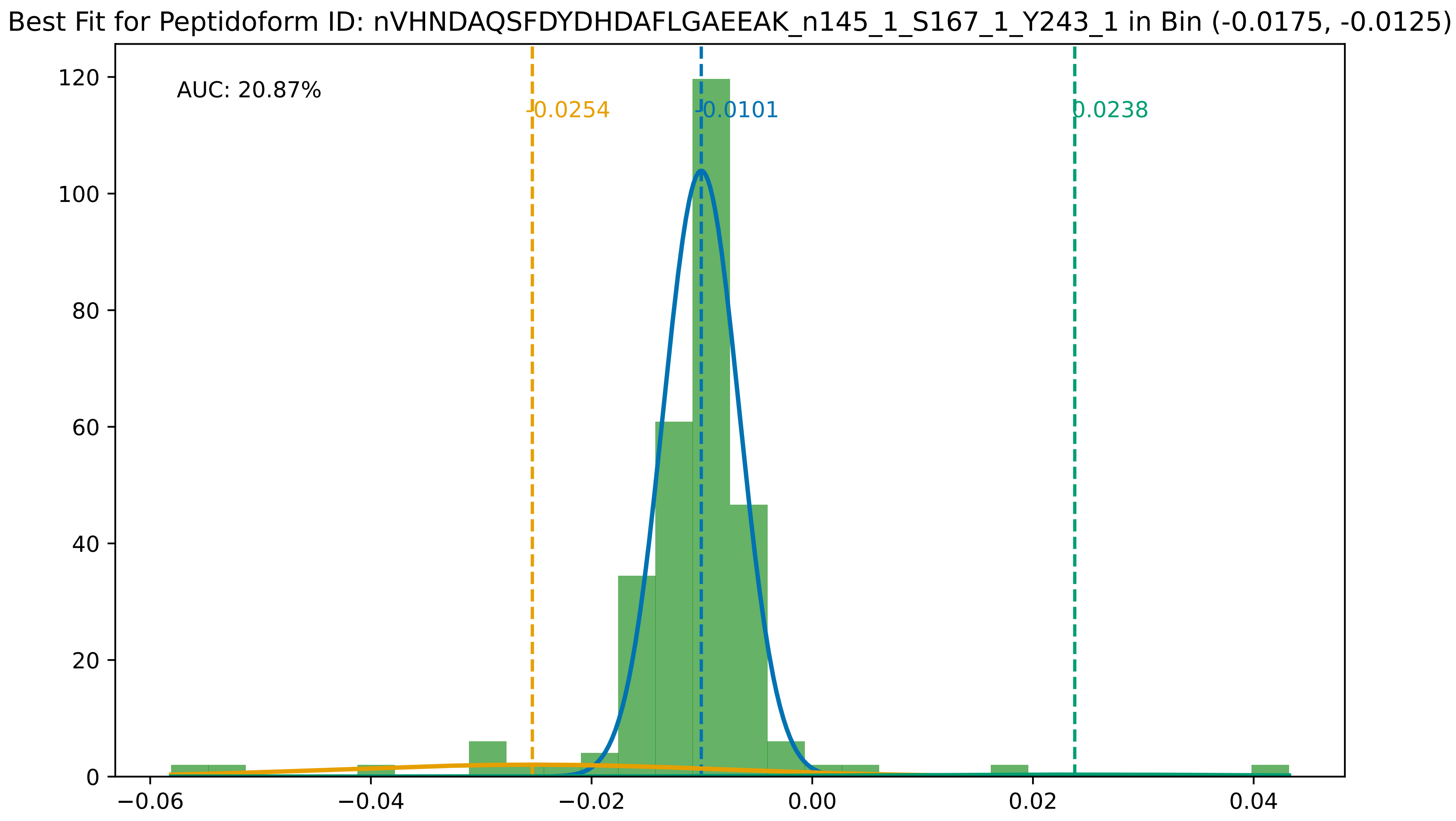

Best Fit for Peptidoform ID: nVHNDAQSFDYDHDAFLGAEEAK_n145_1_S167_1_Y243_1 in Bin (-0.0175, -0.0125)
AUC: 20.87%
-0.0254
-0.0101
0.0238
120
100
80
60
40
20
0
−0.06
−0.04
−0.02
0.00
0.02
0.04

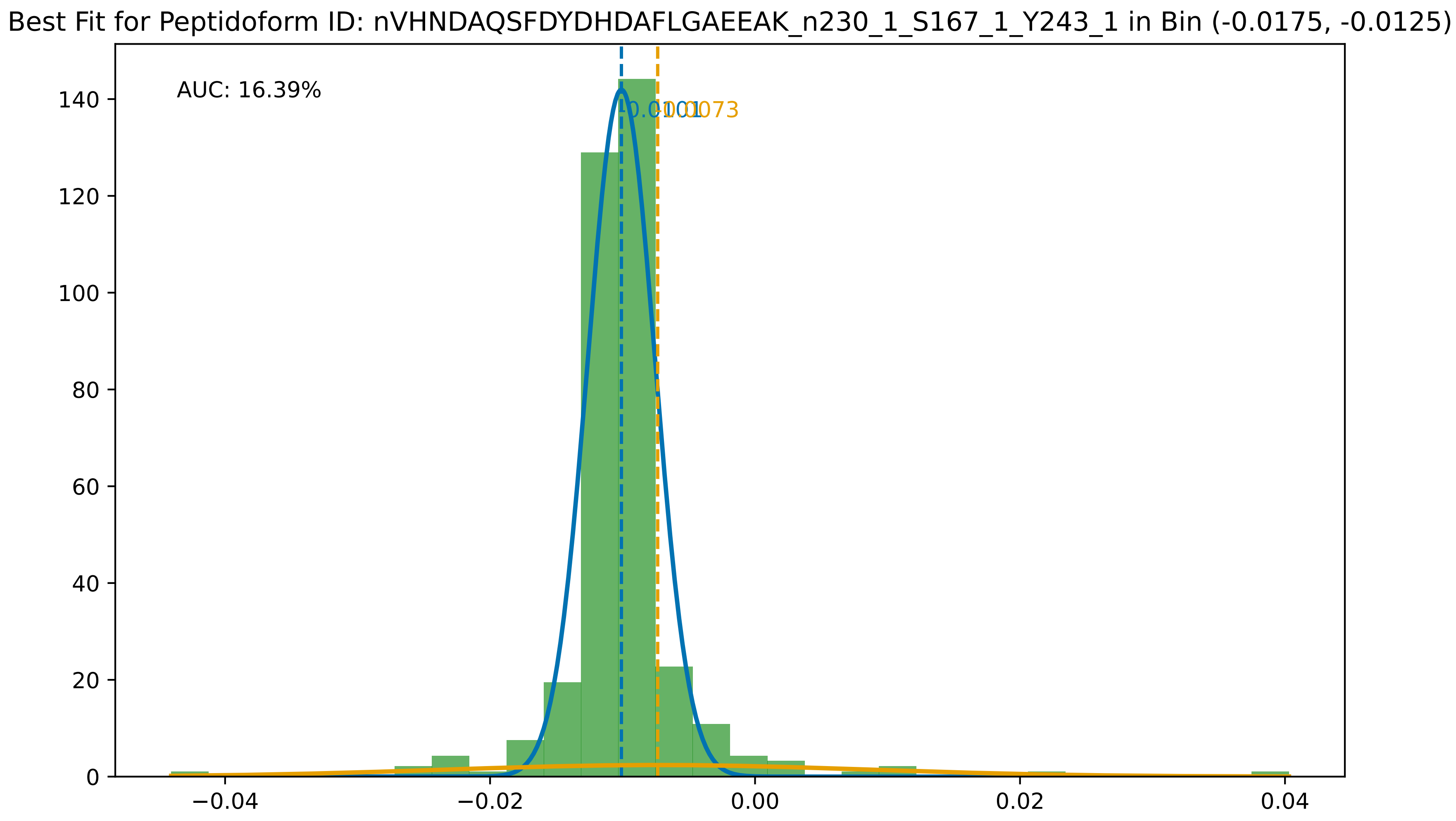

Best Fit for Peptidoform ID: nVHNDAQSFDYDHDAFLGAEEAK_n230_1_S167_1_Y243_1 in Bin (-0.0175, -0.0125)
AUC: 16.39%
-0.0101
-0.0073
0
20
40
60
80
100
120
140
−0.04
−0.02
0.00
0.02
0.04

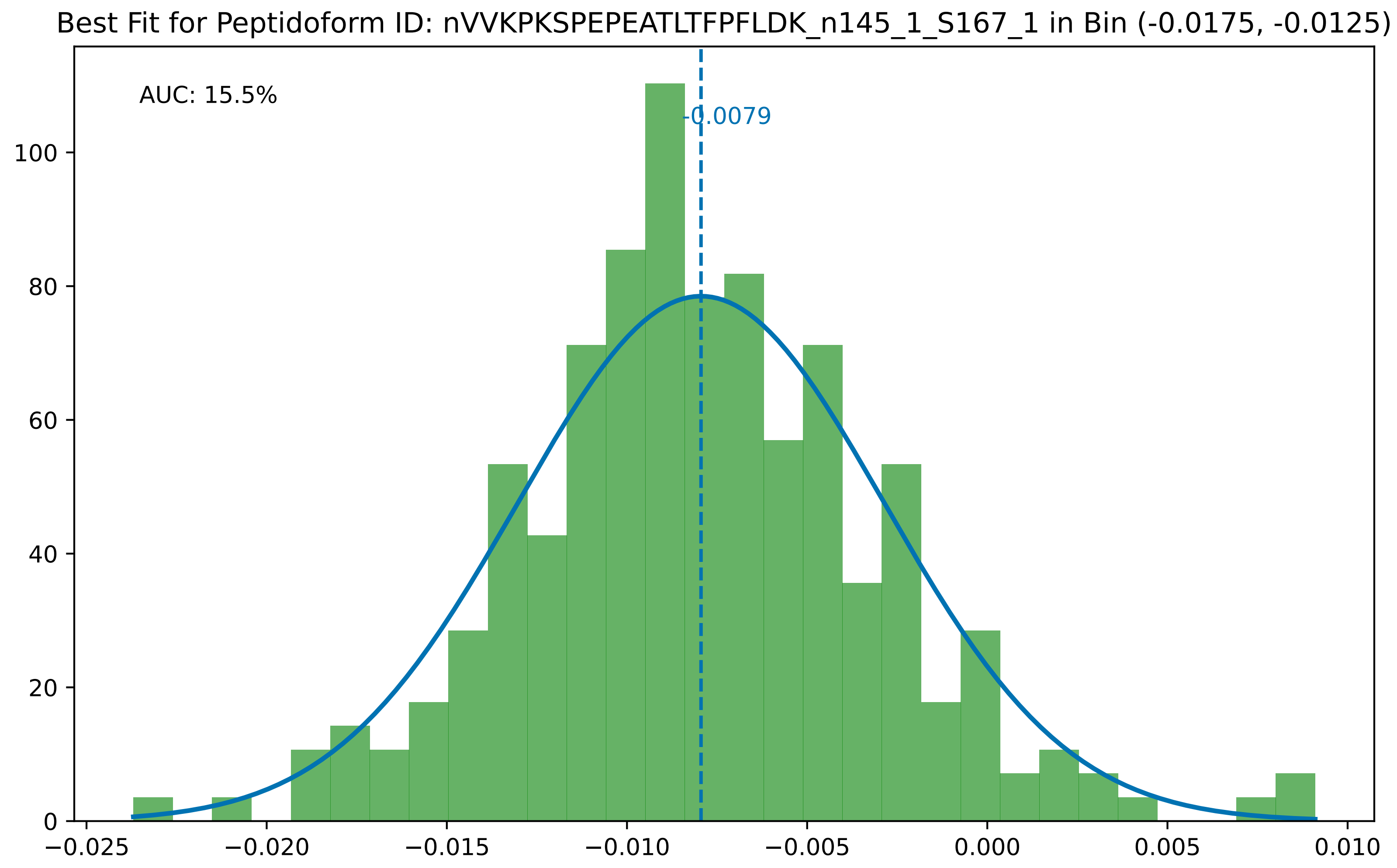
Best Fit for Peptidoform ID: nVVKPKSPEPEATLTFPFLDK_n145_1_S167_1 in Bin (-0.0175, -0.0125)
AUC: 15.5%
-0.0079
0
20
40
60
80
100
−0.025
−0.020
−0.015
−0.010
−0.005
0.000
0.005
0.010

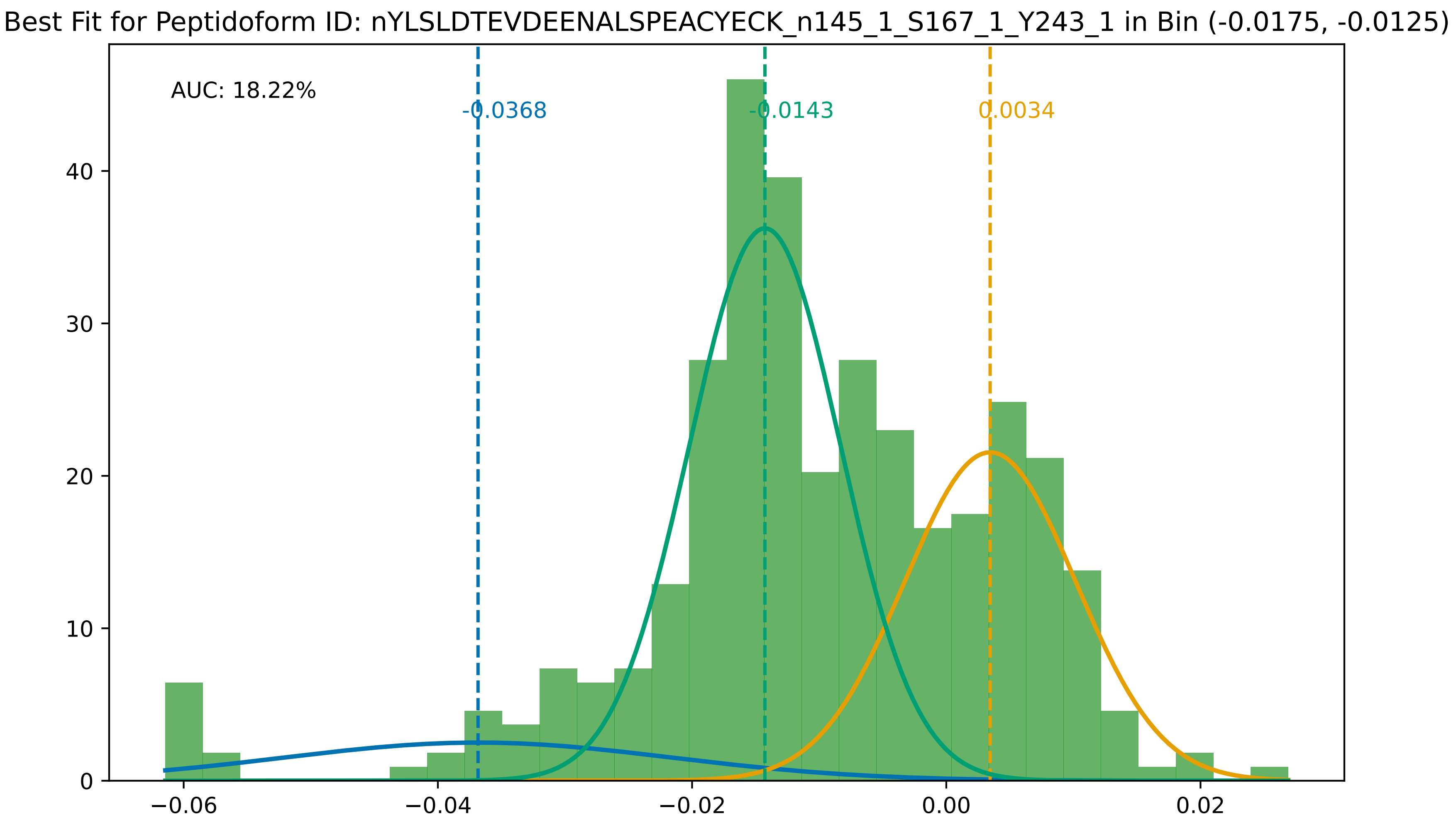
Best Fit for Peptidoform ID: nYLSLDTEVDEENALSPEACYECK_n145_1_S167_1_Y243_1 in Bin (-0.0175, -0.0125)
AUC: 18.22%
-0.0368
-0.0143
0.0034
0
10
20
30
40
−0.06
−0.04
−0.02
0.00
0.02
