## Supplementary File 4 for "Searching for Sulfotyrosines (sY) in a HA(pY)STACK": histograms_post_auc_filtering_bin_-0.0225_-0.0175.pdf

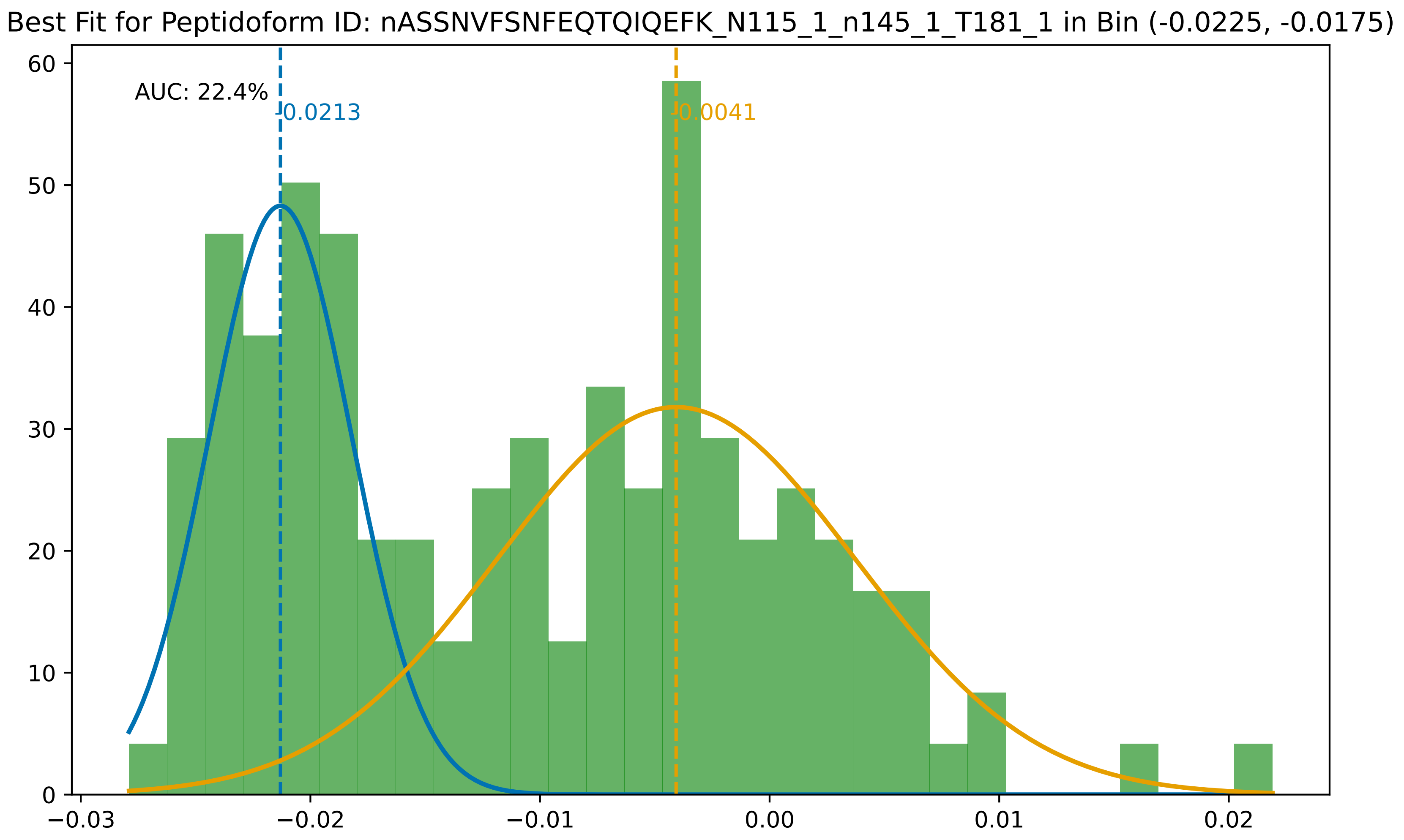

Best Fit for Peptidoform ID: nASSNVFSNFEQTQIQEFK_N115_1_n145_1_T181_1 in Bin (-0.0225, -0.0175)
AUC: 22.4%
-0.0213
-0.0041
60
50
40
30
20
10
0
−0.03
−0.02
−0.01
0.00
0.01
0.02

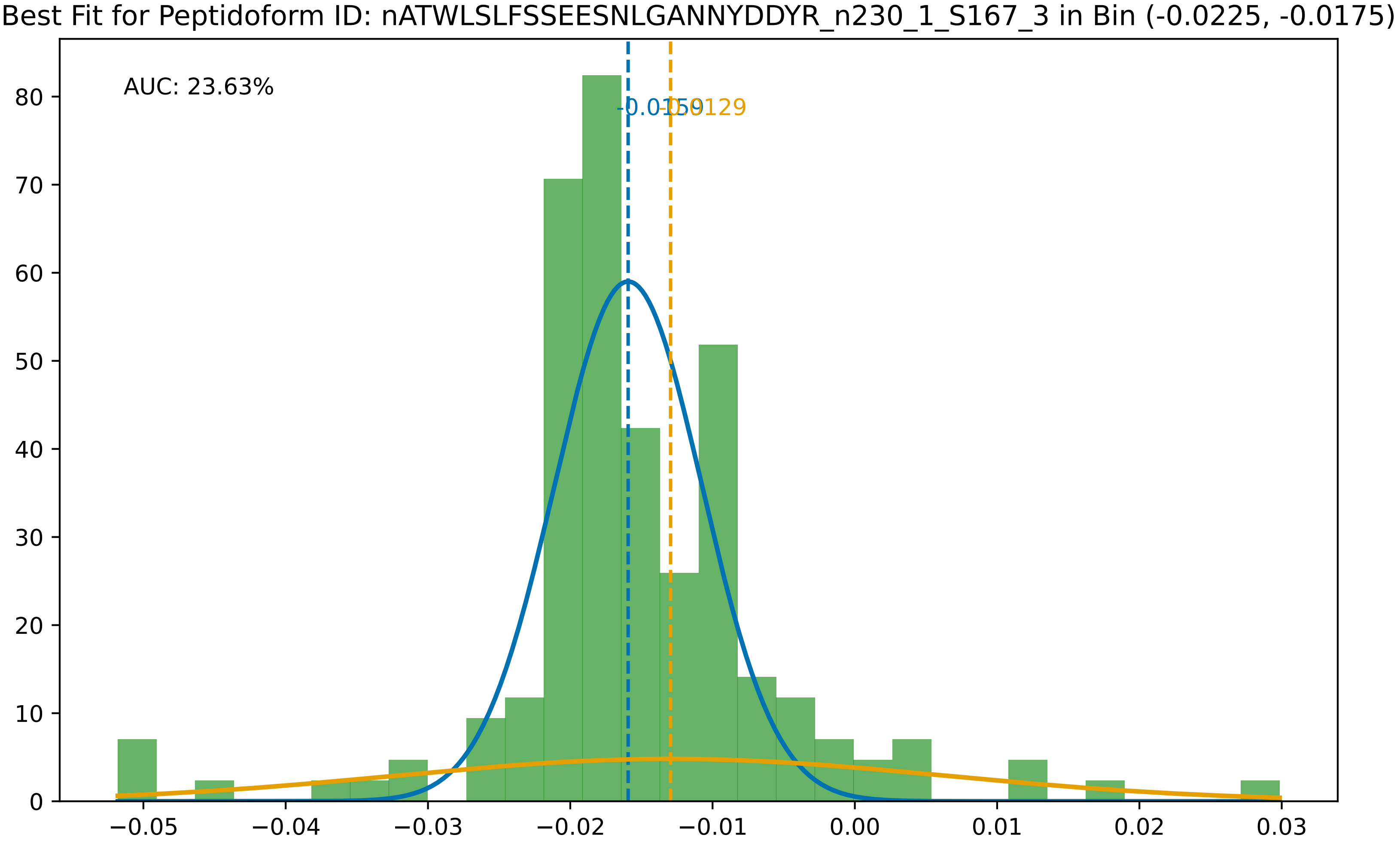
Best Fit for Peptidoform ID: nATWLSLFSSEESNLGANNYDDYR_n230_1_S167_3 in Bin (-0.0225, -0.0175)
AUC: 23.63%
-0.0159
-0.0129
0
10
20
30
40
50
60
70
80
−0.05
−0.04
−0.03
−0.02
−0.01
0.00
0.01
0.02
0.03

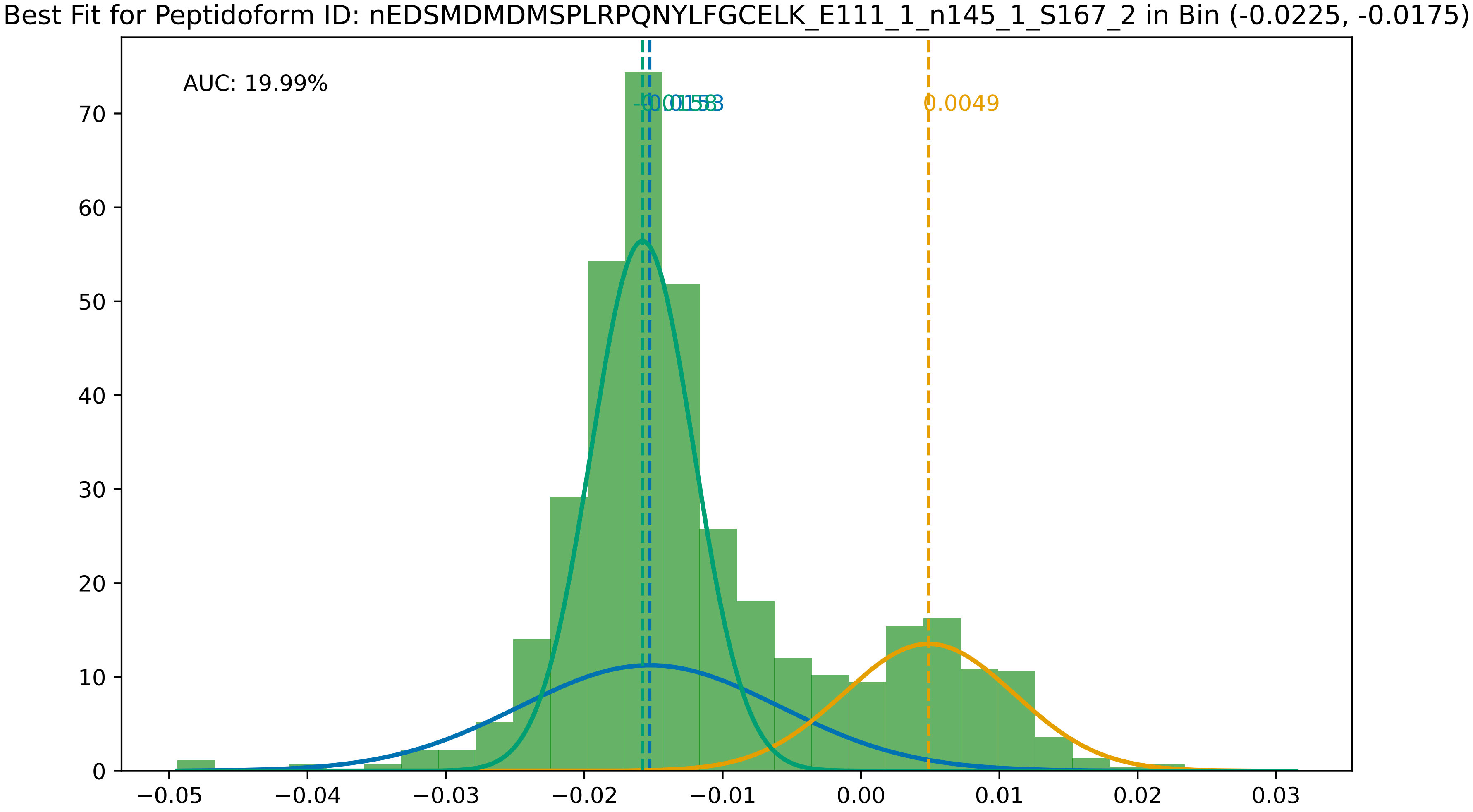

Best Fit for Peptidoform ID: nEDSMDMDMSPLRPQNYLFGCELK_E111_1_n145_1_S167_2 in Bin (-0.0225, -0.0175)
AUC: 19.99%
0.0049
0
10
20
30
40
50
60
70
−0.05
−0.04
−0.03
−0.02
−0.01
0.00
0.01
0.02
0.03

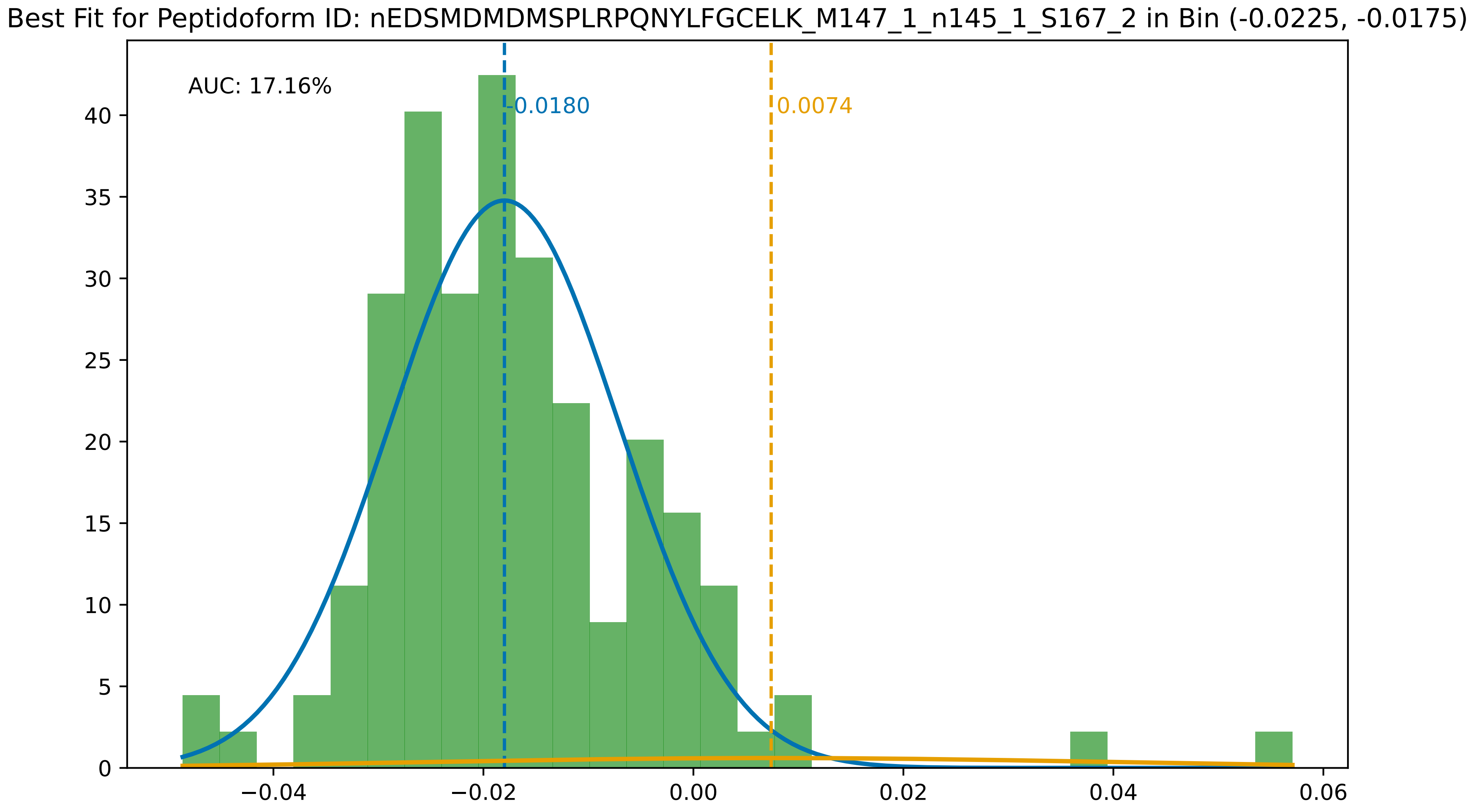
Best Fit for Peptidoform ID: nEDSMDMDMSPLRPQNYLFGCELK_M147_1_n145_1_S167_2 in Bin (-0.0225, -0.0175)
AUC: 17.16%
-0.0180
0.0074
0
5
10
15
20
25
30
35
40
−0.04
−0.02
0.00
0.02
0.04
0.06

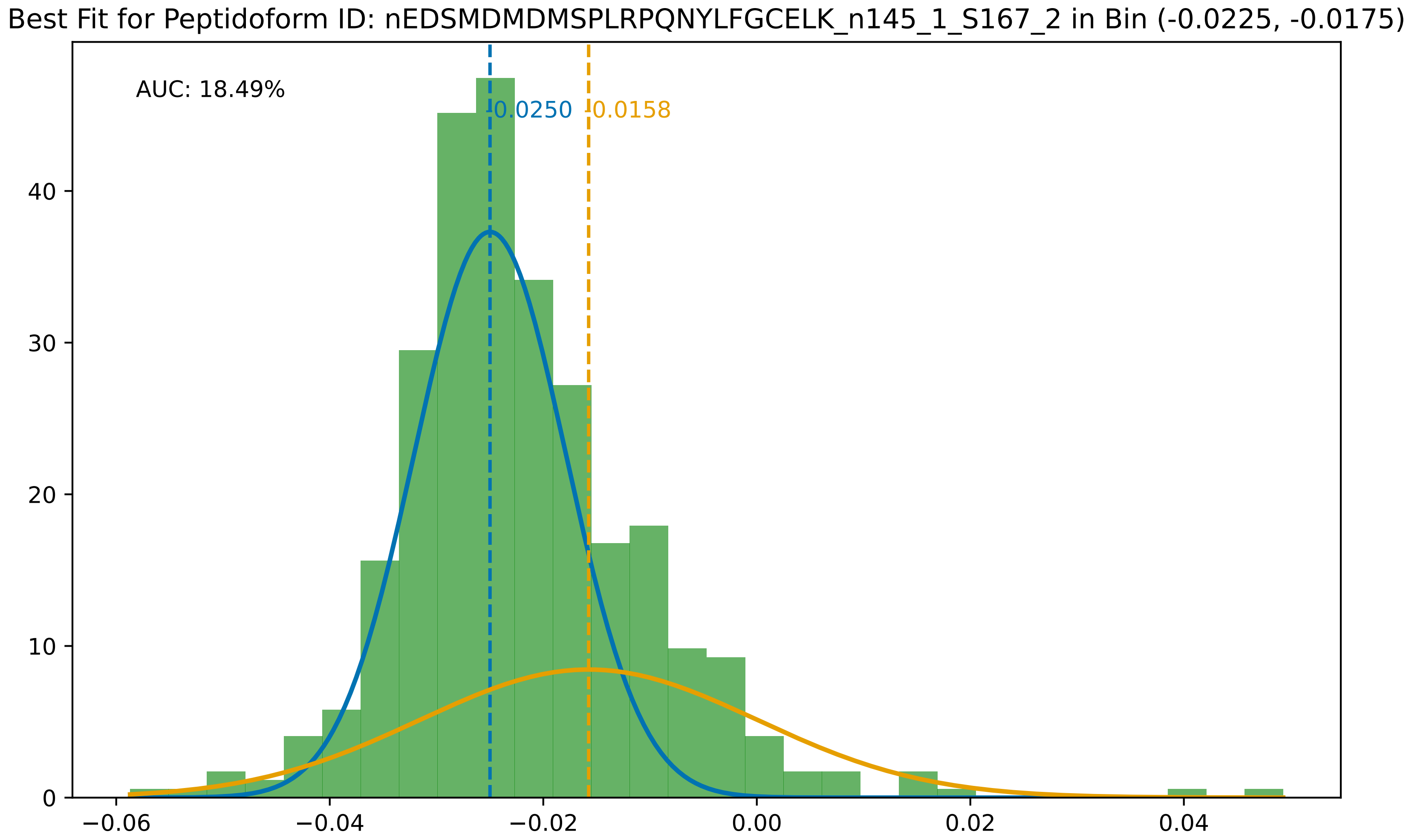
Best Fit for Peptidoform ID: nEDSMDMDMSPLRPQNYLFGCELK_n145_1_S167_2 in Bin (-0.0225, -0.0175)
AUC: 18.49%
-0.0250
-0.0158
0
10
20
30
40
−0.06
−0.04
−0.02
0.00
0.02
0.04

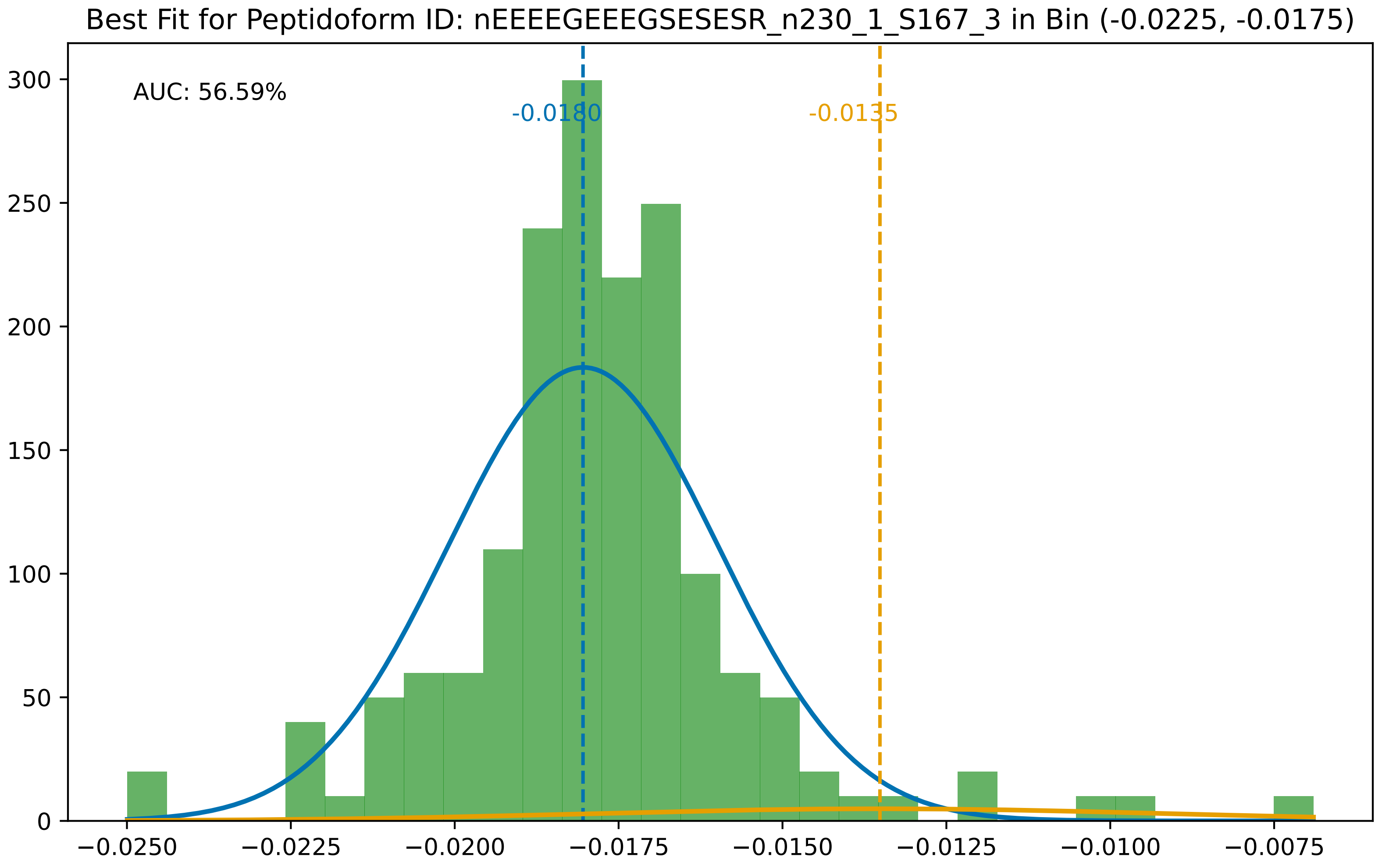
Best Fit for Peptidoform ID: nEEEEGEEEGSESESR_n230_1_S167_3 in Bin (-0.0225, -0.0175)
AUC: 56.59%
-0.0180
-0.0135
300
250
200
150
100
50
0
−0.0250
−0.0225
−0.0200
−0.0175
−0.0150
−0.0125
−0.0100
−0.0075

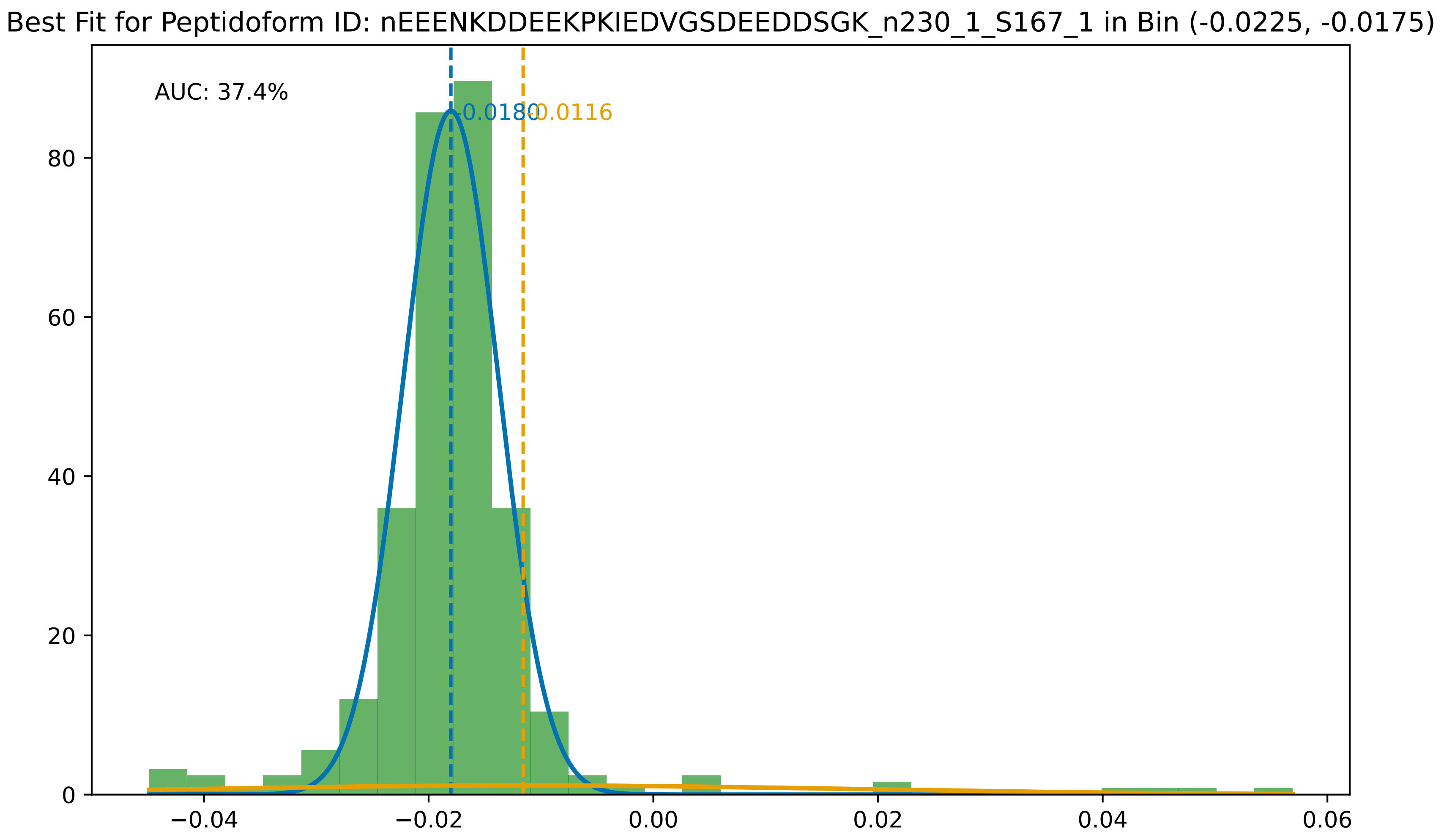

Best Fit for Peptidoform ID: nEEENKDDEEKPKIEDVGSDEEDDSGK_n230_1_S167_1 in Bin (-0.0225, -0.0175)
AUC: 37.4%
-0.0180
-0.0116
0
20
40
60
80
−0.04
−0.02
0.00
0.02
0.04
0.06

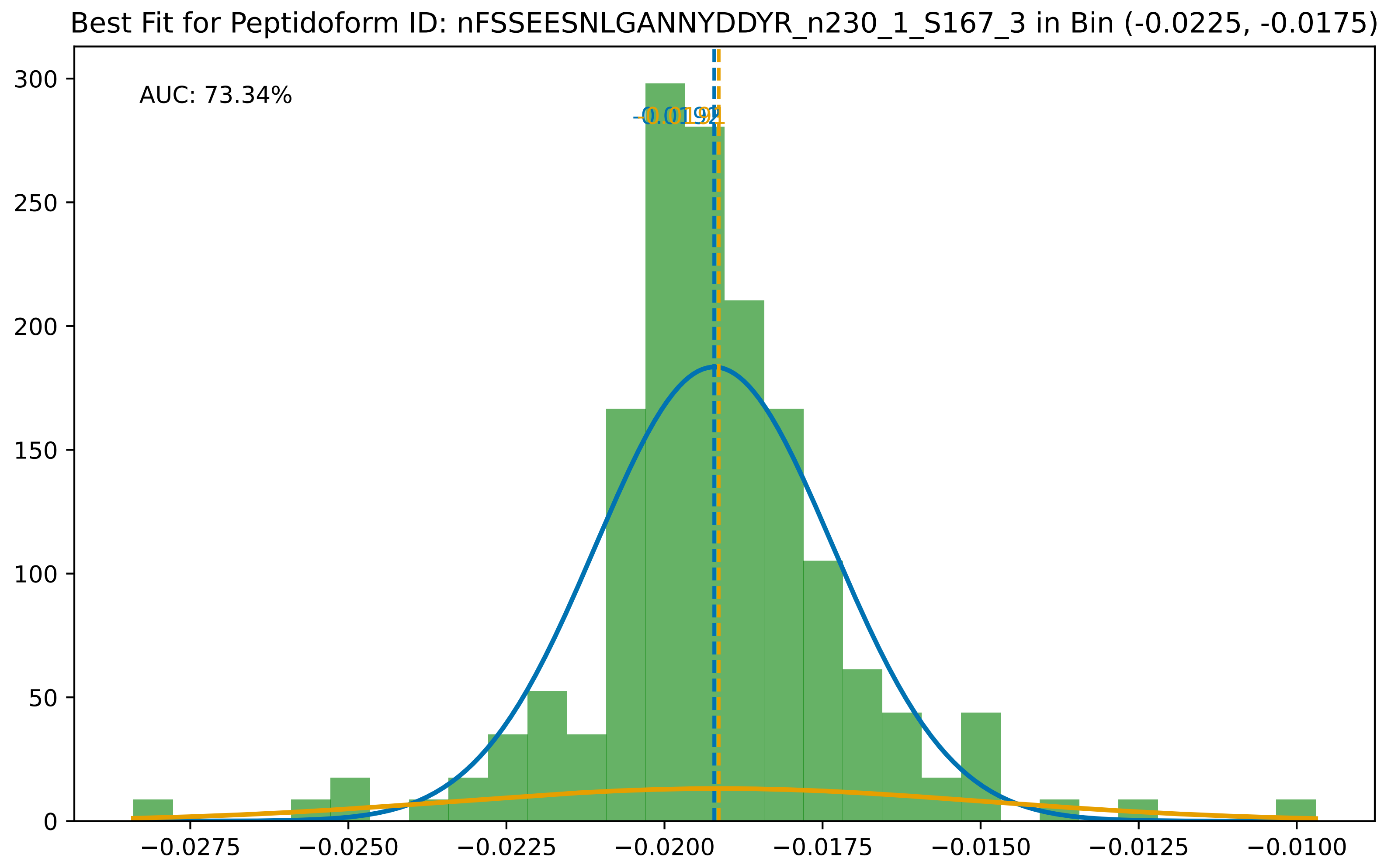

Best Fit for Peptidoform ID: nFSSEESNLGANNYDDYR_n230_1_S167_3 in Bin (-0.0225, -0.0175)
AUC: 73.34%
300
250
200
150
100
50
0
−0.0275
−0.0250
−0.0225
−0.0200
−0.0175
−0.0150
−0.0125
−0.0100

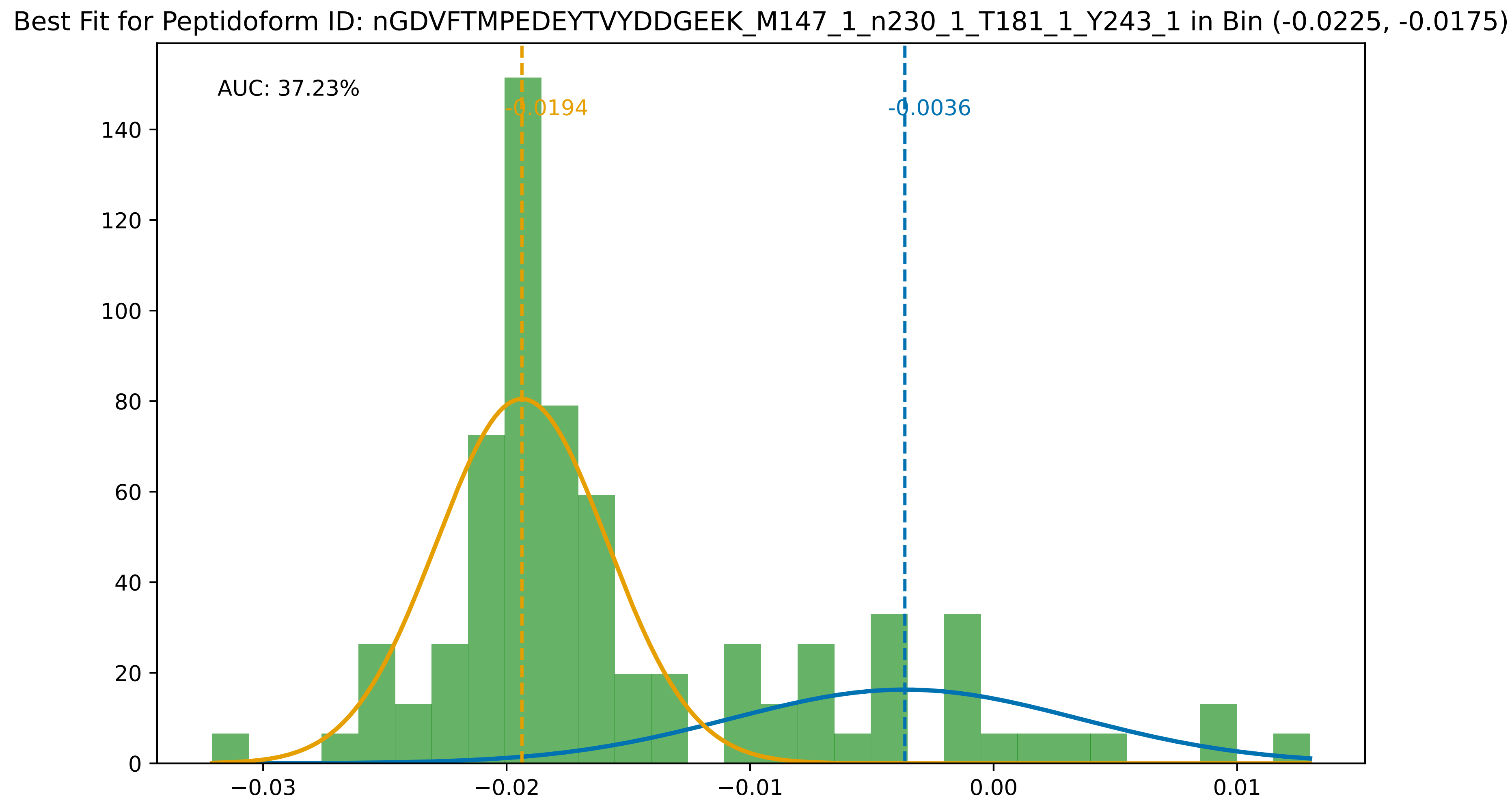
Best Fit for Peptidoform ID: nGDVFTMPEDEYTVYDDGEEK_M147_1_n230_1_T181_1_Y243_1 in Bin (-0.0225, -0.0175)
AUC: 37.23%
-0.0194
-0.0036
0
20
40
60
80
100
120
140
−0.03
−0.02
−0.01
0.00
0.01

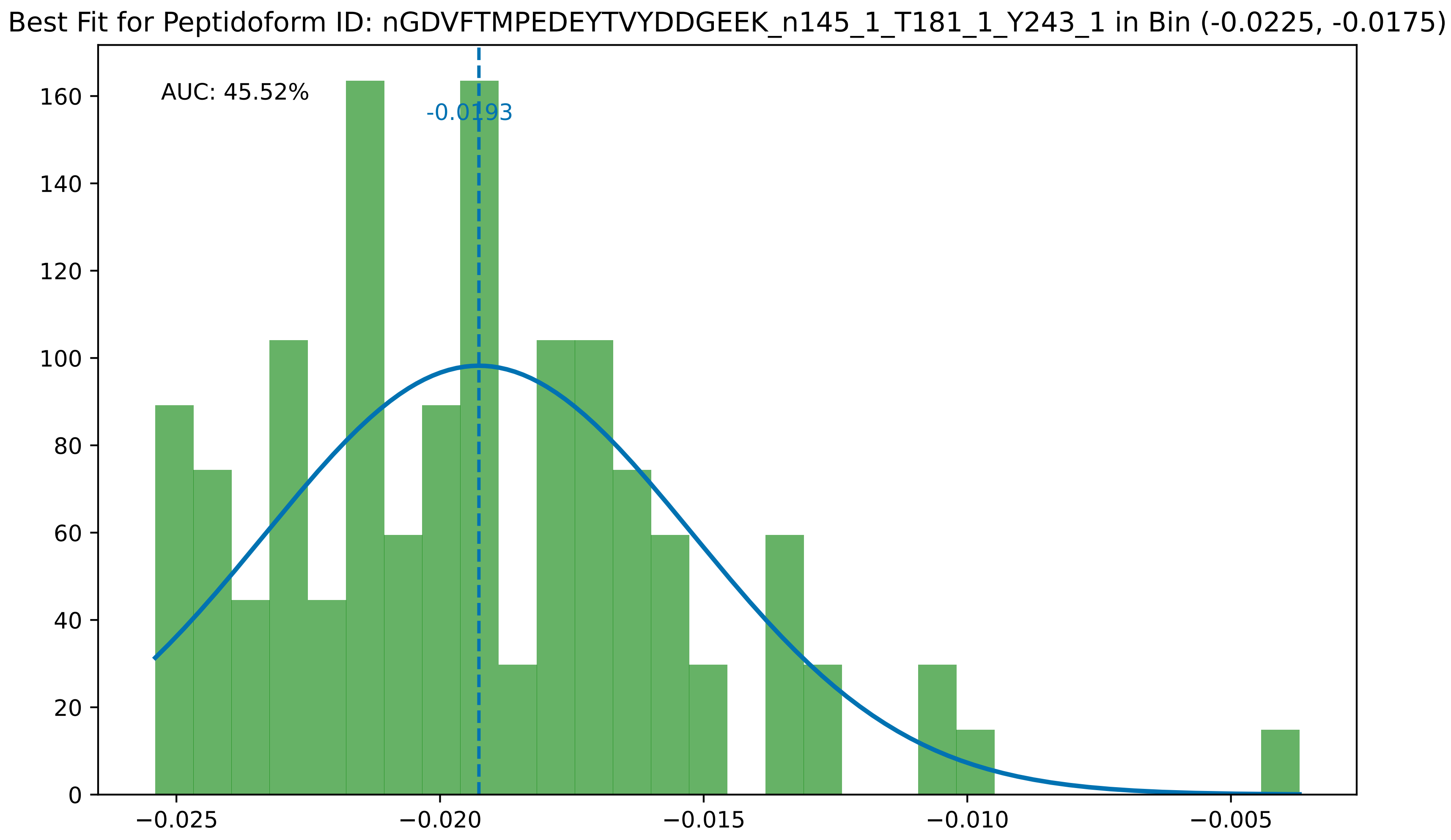

Best Fit for Peptidoform ID: nGDVFTMPEDEYTVYDDGEEK_n145_1_T181_1_Y243_1 in Bin (-0.0225, -0.0175)
AUC: 45.52%
-0.0193
0
20
40
60
80
100
120
140
160
−0.025
−0.020
−0.015
−0.010
−0.005

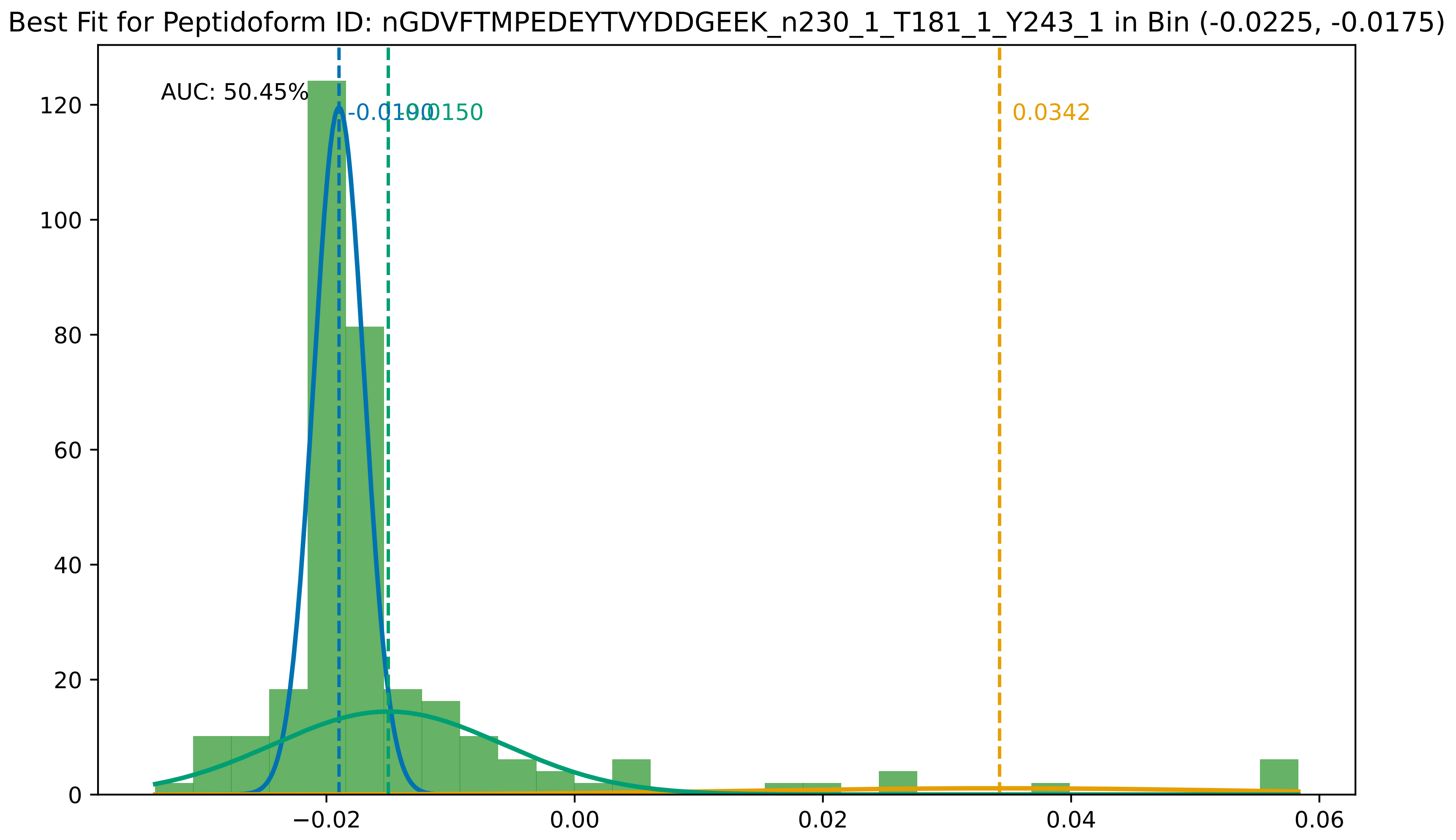

Best Fit for Peptidoform ID: nGDVFTMPEDEYTVYDDGEEK_n230_1_T181_1_Y243_1 in Bin (-0.0225, -0.0175)
AUC: 50.45%
-0.0190
-0.0150
0.0342
0
20
40
60
80
100
120
−0.02
0.00
0.02
0.04
0.06

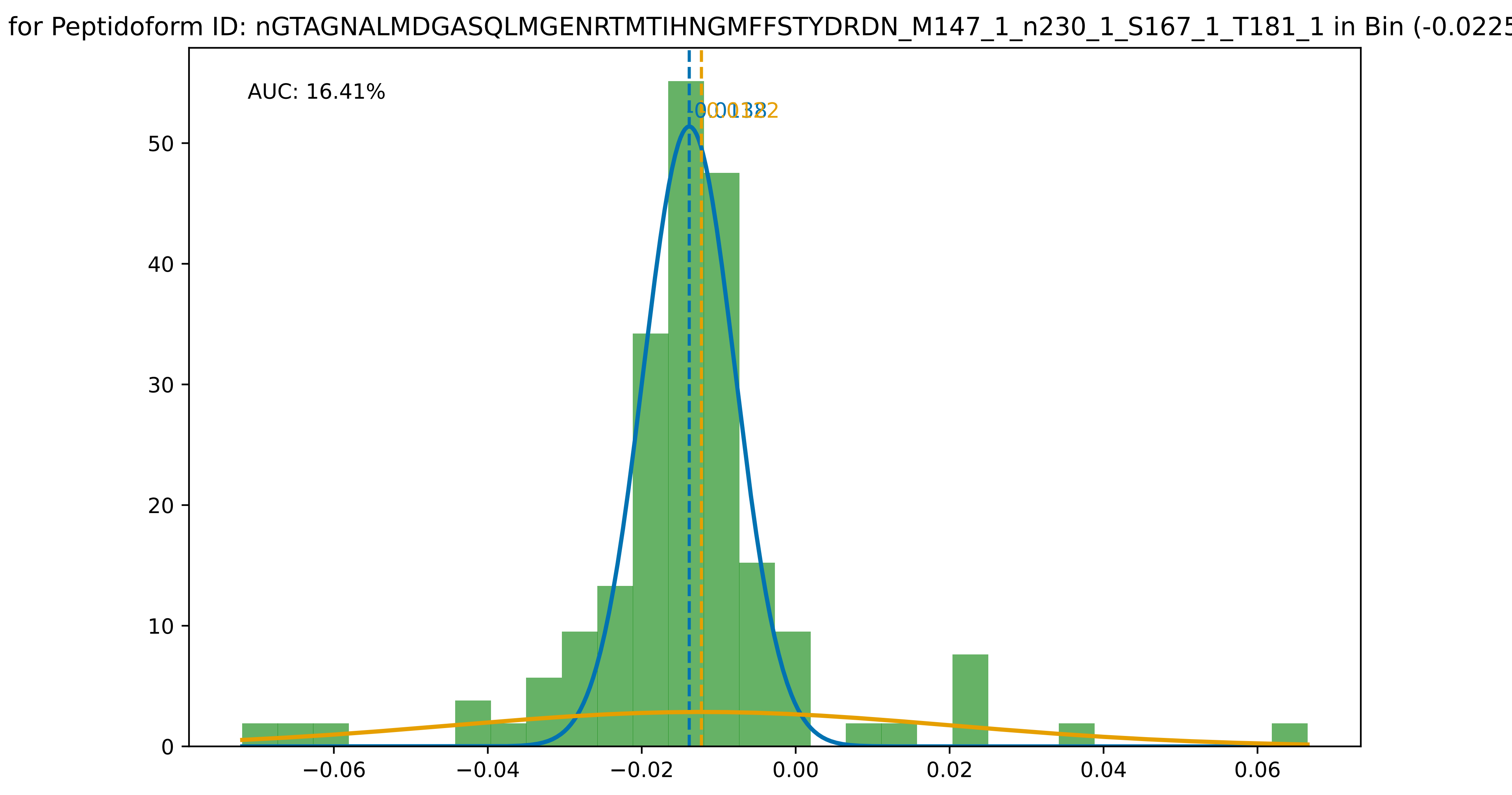
for Peptidoform ID: nGTAGNALMDGASQLMGENRTMTIHNGMFFSTYDRDN_M147_1_n230_1_S167_1_T181_1 in Bin (-0.0225
AUC: 16.41%
0.0138
0.0122
0
10
20
30
40
50
−0.06
−0.04
−0.02
0.00
0.02
0.04
0.06

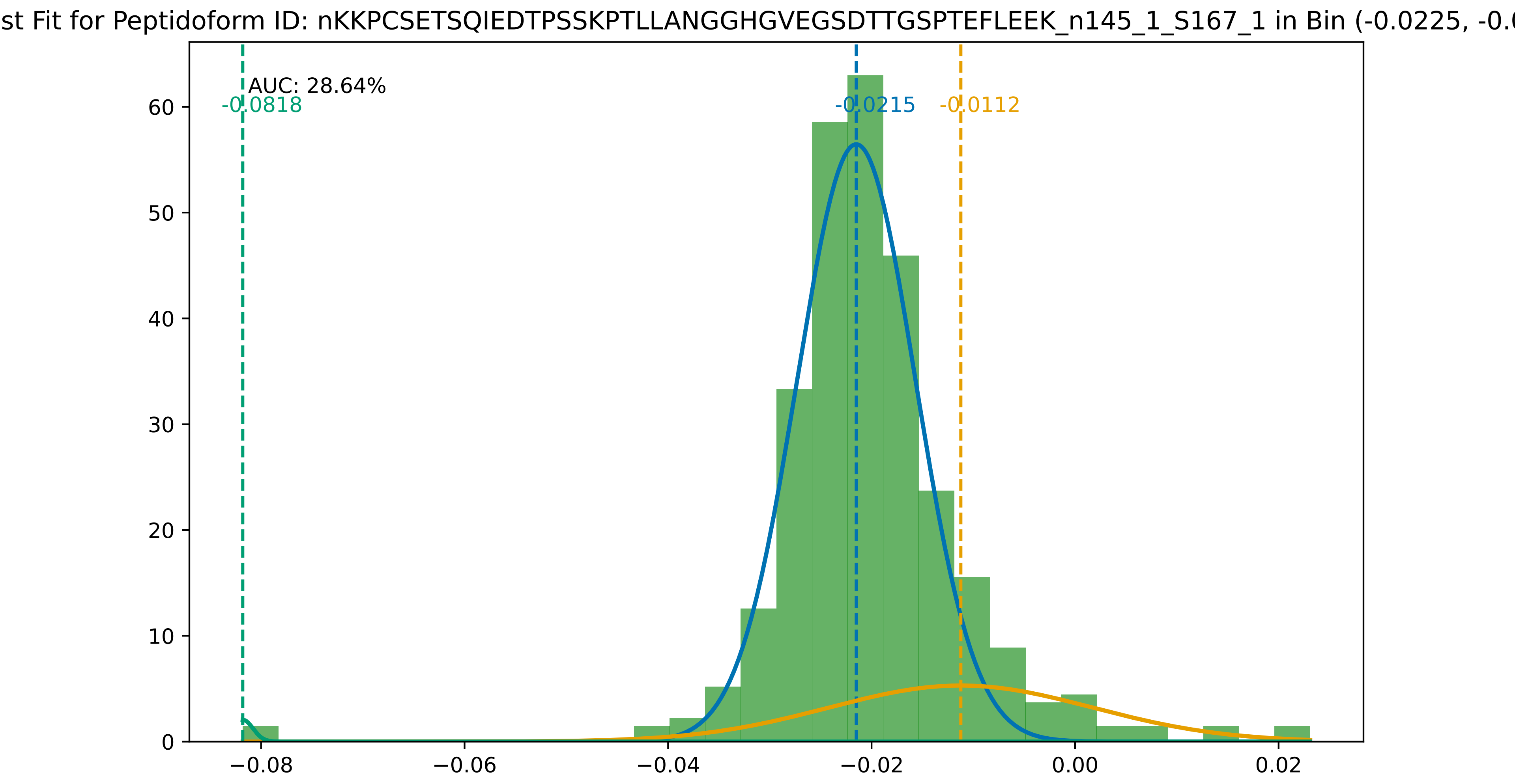
st Fit for Peptidoform ID: nKKPCSETSQIEDTPSSKPTLLANGGHGVEGSDTTGSPTEFLEEK_n145_1_S167_1 in Bin (-0.0225, -0.0
AUC: 28.64%
-0.0818
-0.0215
-0.0112
0
10
20
30
40
50
60
−0.08
−0.06
−0.04
−0.02
0.00
0.02

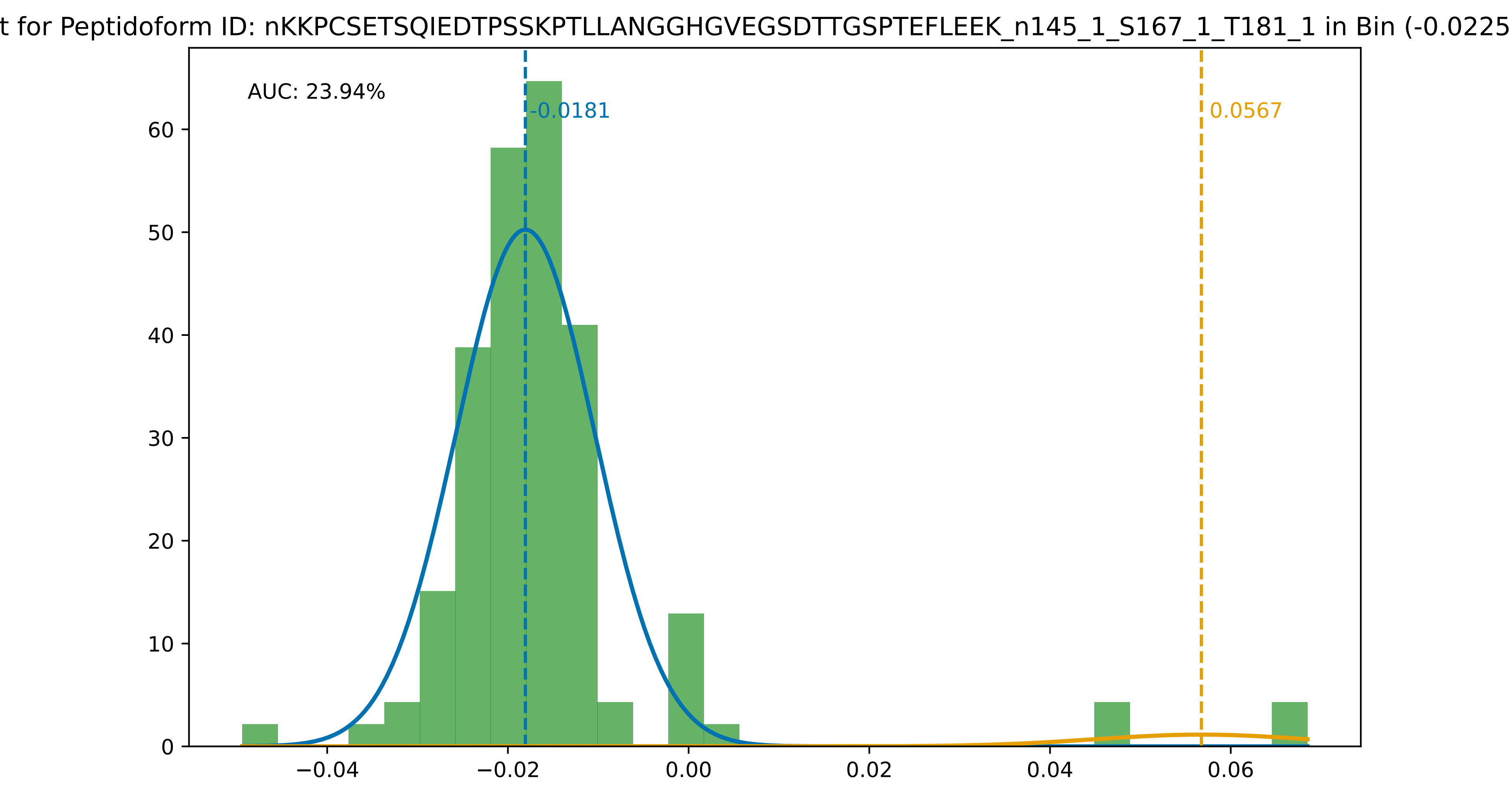

t for Peptidoform ID: nKKPCSETSQIEDTPSSKPTLLANGGHGVEGSDTTGSPTEFLEEK_n145_1_S167_1_T181_1 in Bin (-0.0225
AUC: 23.94%
-0.0181
0.0567
0
10
20
30
40
50
60
−0.04
−0.02
0.00
0.02
0.04
0.06

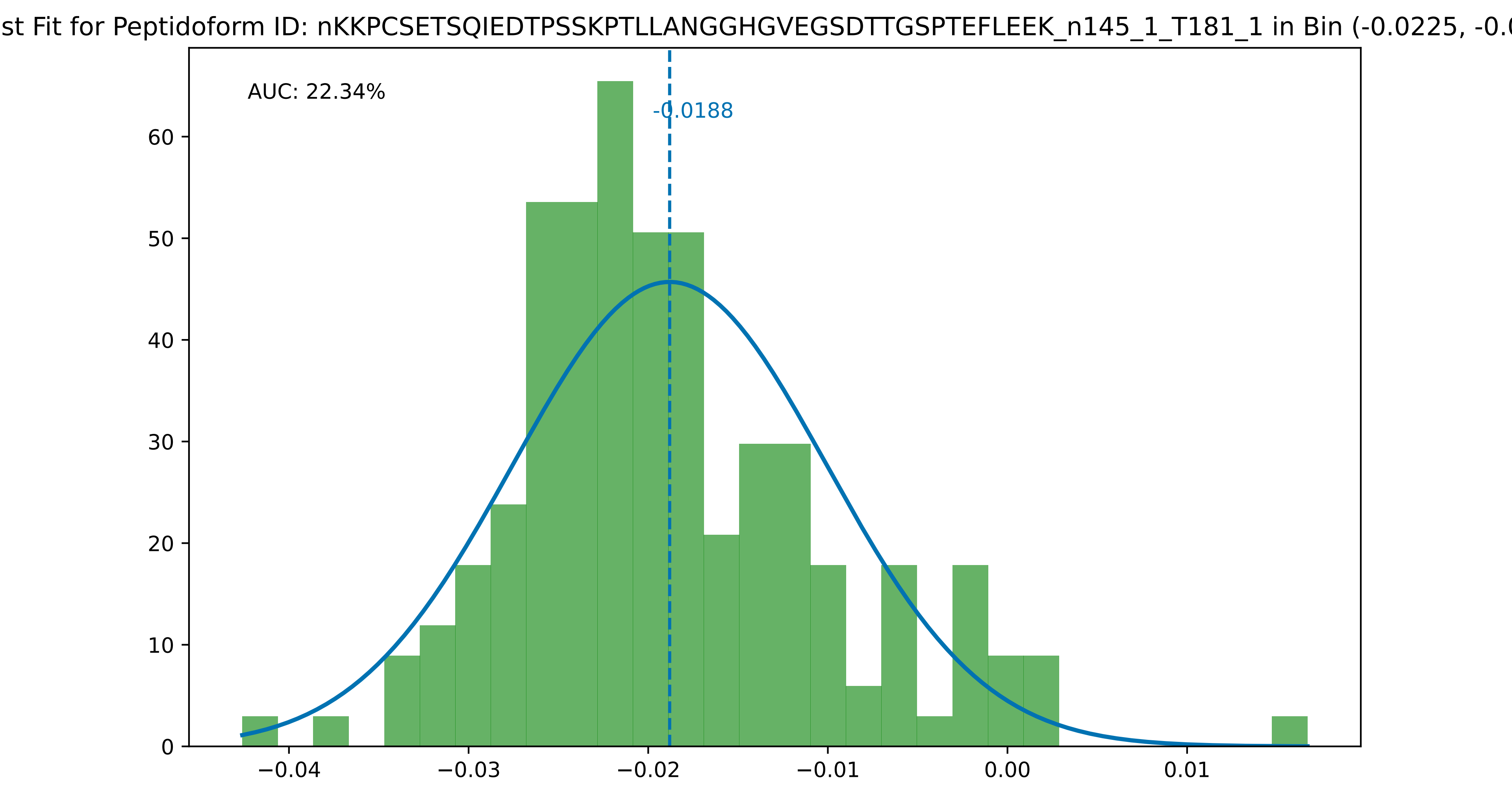

st Fit for Peptidoform ID: nKKPCSETSQIEDTPSSKPTLLANGGHGVEGSDTTGSPTEFLEEK_n145_1_T181_1 in Bin (-0.0225, -0.0
AUC: 22.34%
-0.0188
60
50
40
30
20
10
0
−0.04
−0.03
−0.02
−0.01
0.00
0.01

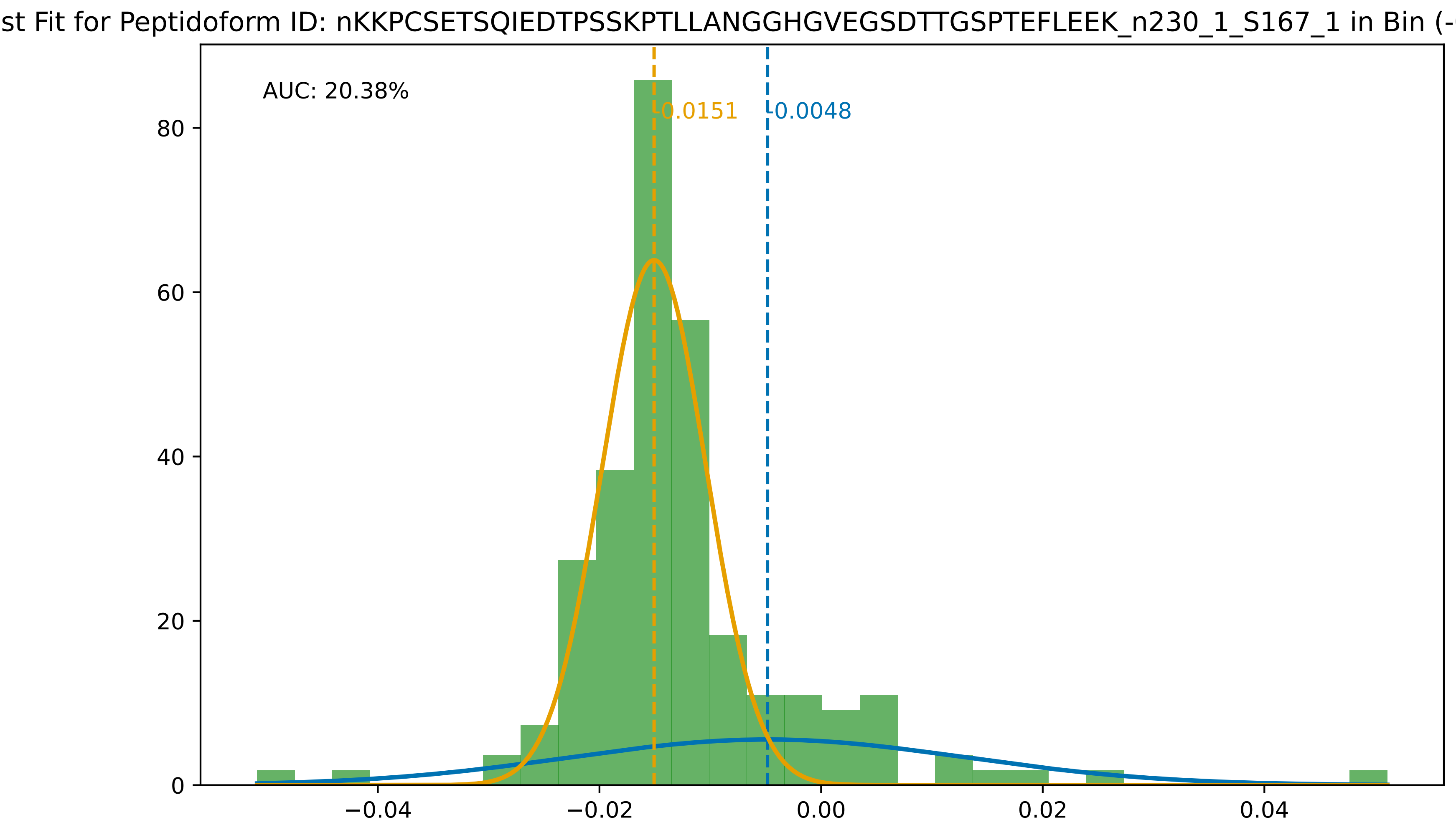

st Fit for Peptidoform ID: nKKPCSETSQIEDTPSSKPTLLANGGHGVEGSDTTGSPTEFLEEK_n230_1_S167_1 in Bin (-0.0225, -0.0
AUC: 20.38%
-0.0151
-0.0048
80
60
40
20
0
−0.04
−0.02
0.00
0.02
0.04

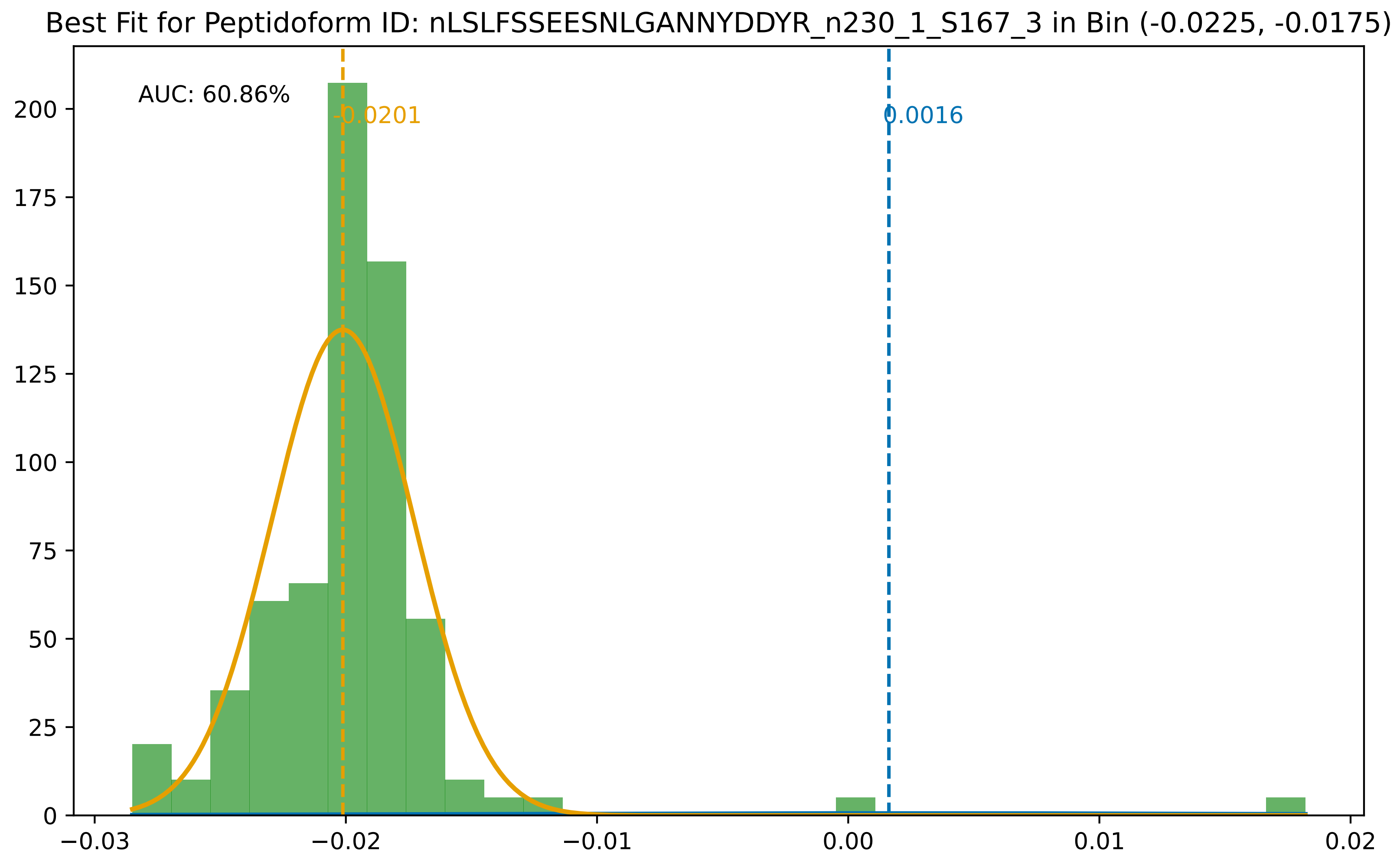

Best Fit for Peptidoform ID: nLSLFSSEESNLGANNYDDYR_n230_1_S167_3 in Bin (-0.0225, -0.0175)
AUC: 60.86%
-0.0201
0.0016
0
25
50
75
100
125
150
175
200
−0.03
−0.02
−0.01
0.00
0.01
0.02

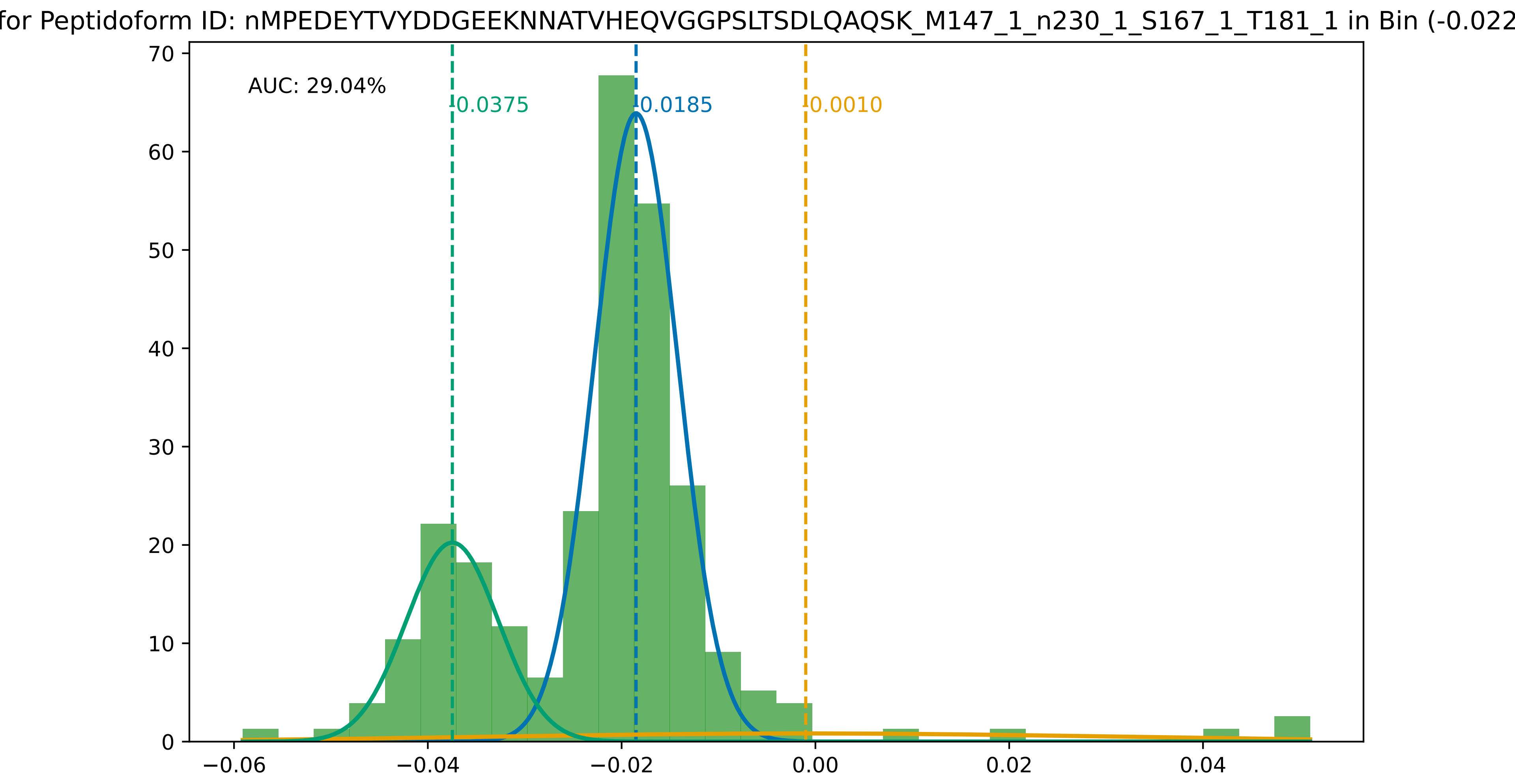
for Peptidoform ID: nMPEDEYTVYDDGEEKNNATVHEQVGGPSLTSDLQAQSK_M147_1_n230_1_S167_1_T181_1 in Bin (-0.022
AUC: 29.04%
0.0375
0.0185
0.0010
70
60
50
40
30
20
10
0
−0.06
−0.04
−0.02
0.00
0.02
0.04

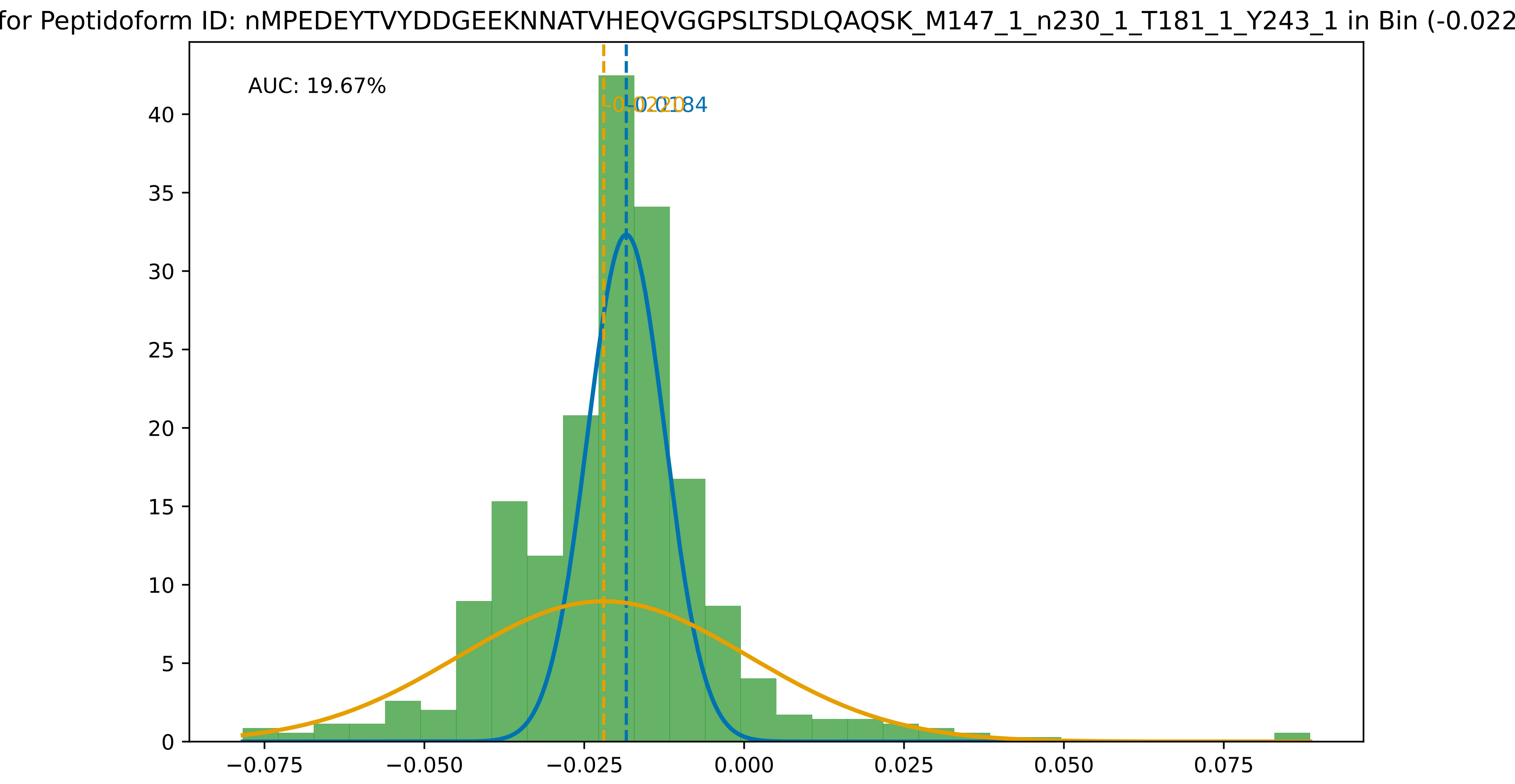
for Peptidoform ID: nMPEDEYTVYDDGEEKNNATVHEQVGGPSLTSDLQAQSK_M147_1_n230_1_T181_1_Y243_1 in Bin (-0.022
AUC: 19.67%
-0.0220
-0.0184
0
5
10
15
20
25
30
35
40
−0.075
−0.050
−0.025
0.000
0.025
0.050
0.075

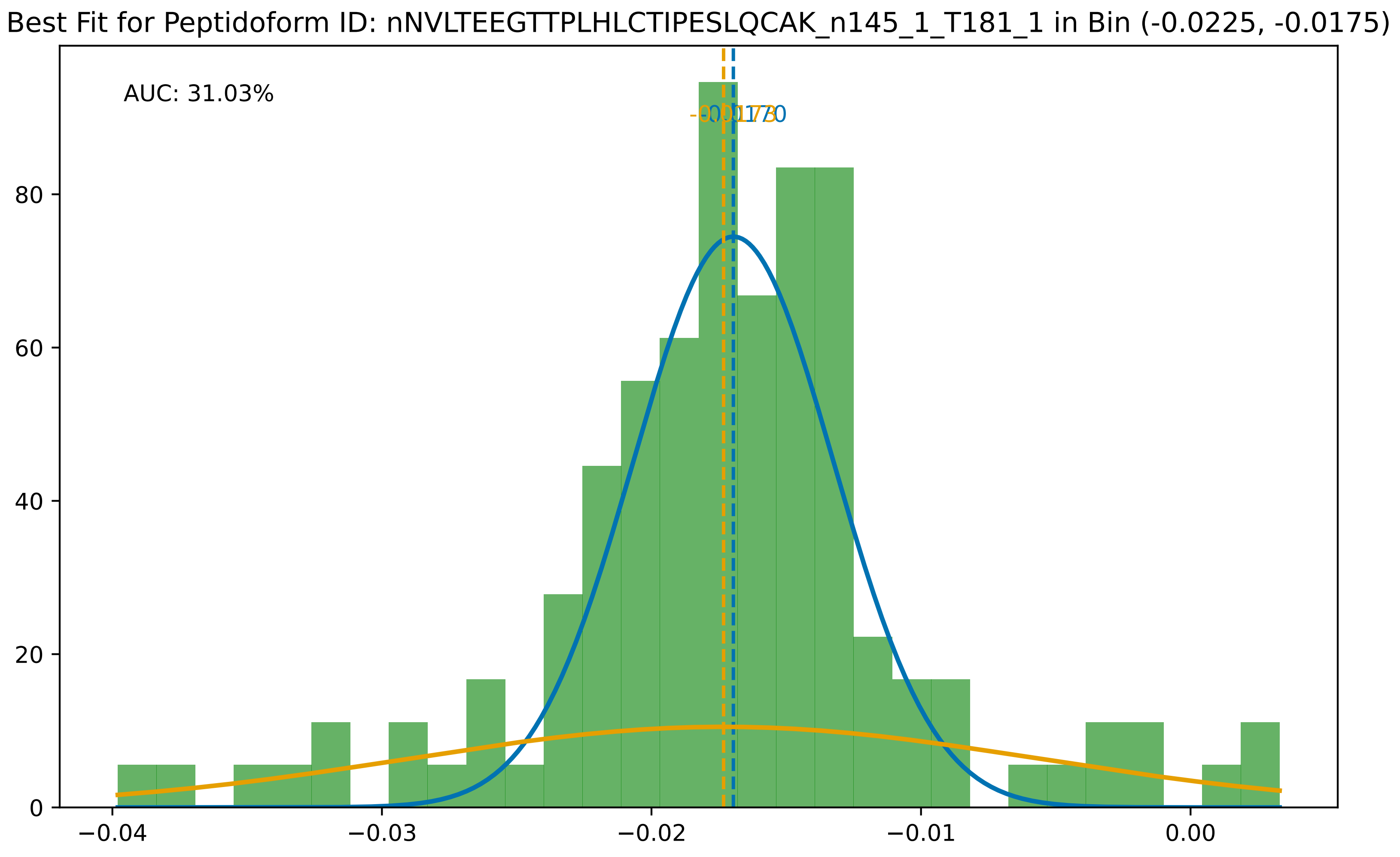

Best Fit for Peptidoform ID: nNVLTEEGTTPLHLCTIPESLQCAK_n145_1_T181_1 in Bin (-0.0225, -0.0175)
AUC: 31.03%
0
20
40
60
80
−0.04
−0.03
−0.02
−0.01
0.00

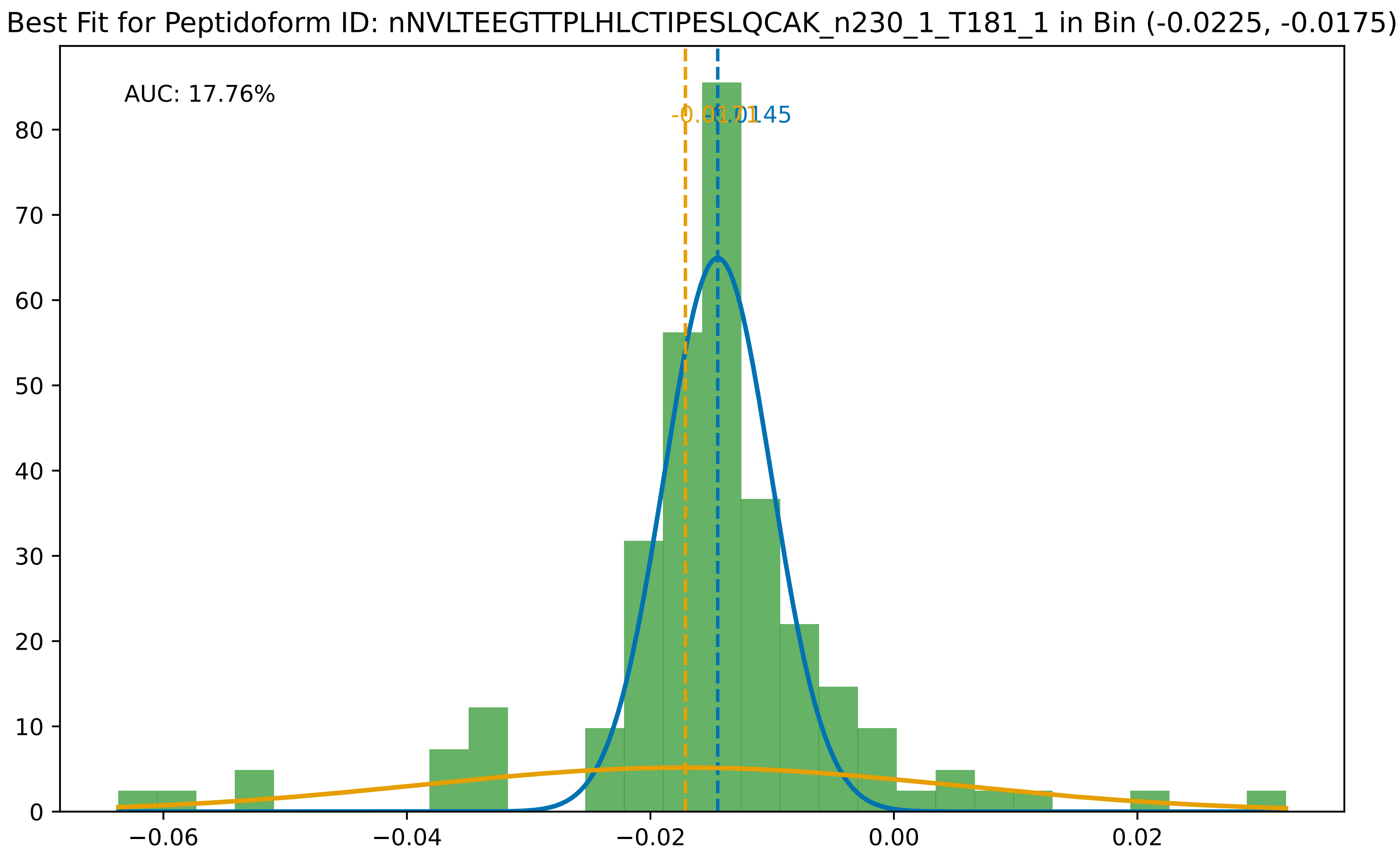

Best Fit for Peptidoform ID: nNVLTEEGTTPLHLCTIPESLQCAK_n230_1_T181_1 in Bin (-0.0225, -0.0175)
AUC: 17.76%
-0.0170
-0.0145
0
10
20
30
40
50
60
70
80
−0.06
−0.04
−0.02
0.00
0.02

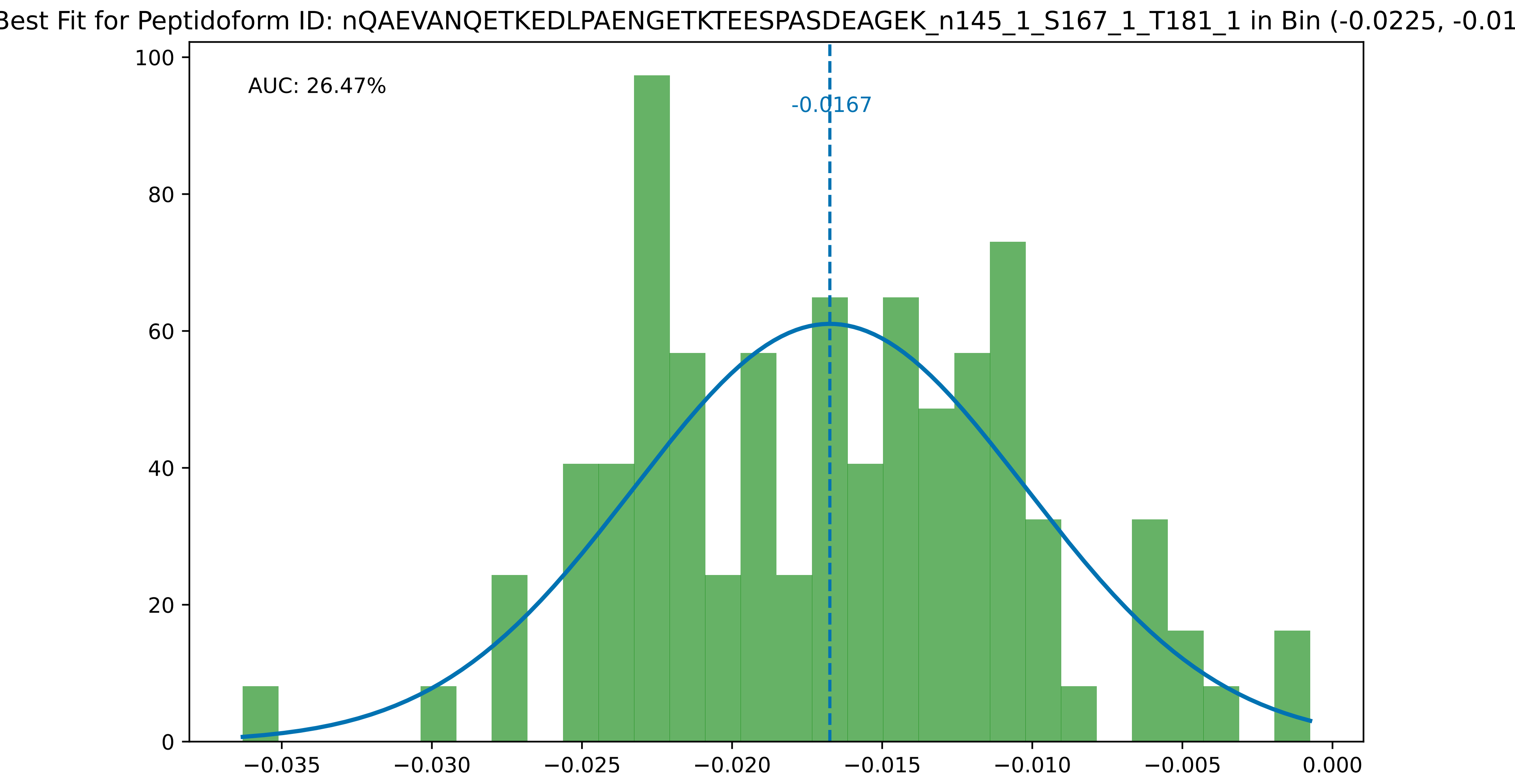

Best Fit for Peptidoform ID: nQAEVANQETKEDLPAENGETKTEESPASDEAGEK_n145_1_S167_1_T181_1 in Bin (-0.0225, -0.01
AUC: 26.47%
-0.0167
0
20
40
60
80
100
−0.035
−0.030
−0.025
−0.020
−0.015
−0.010
−0.005
0.000

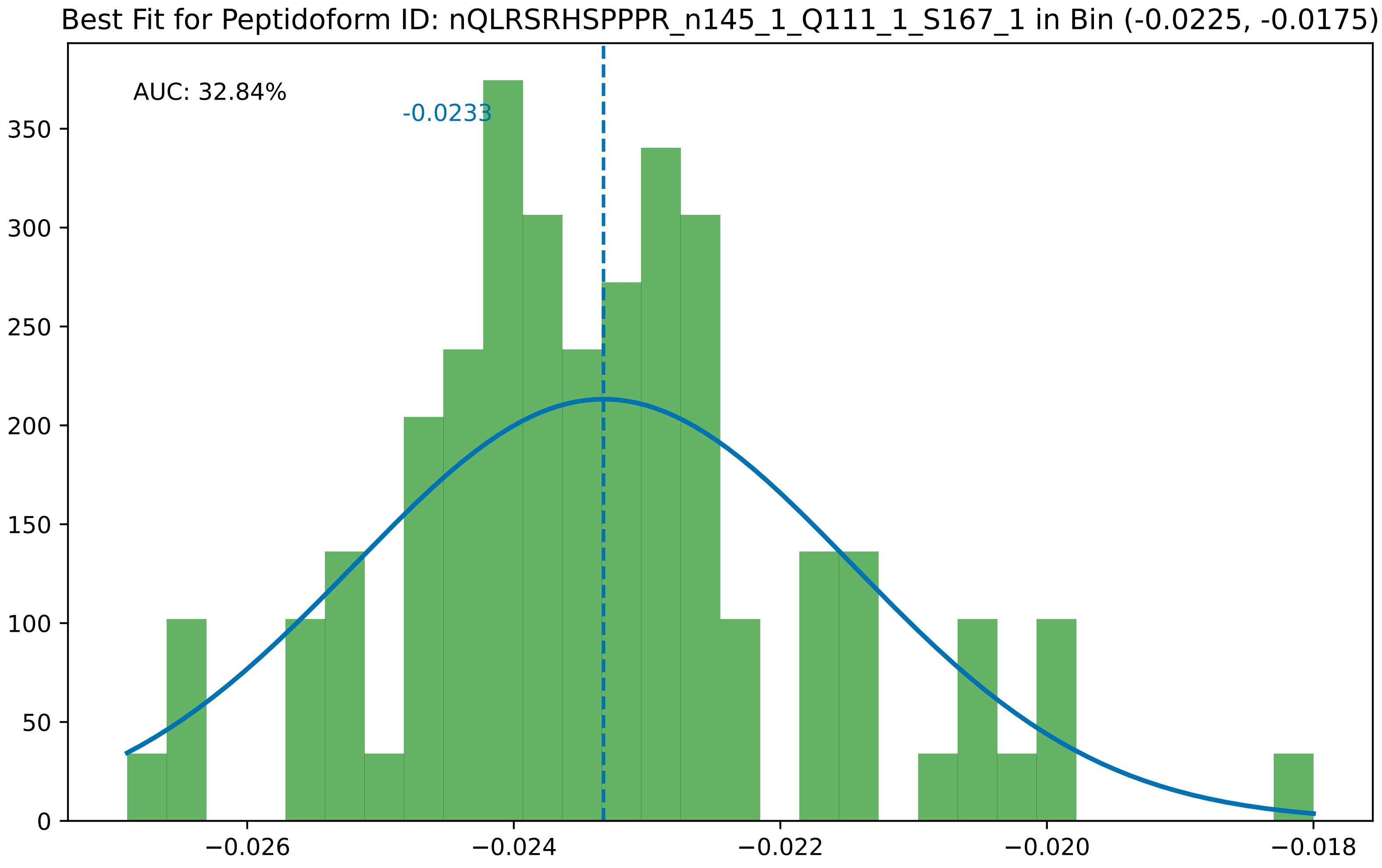

Best Fit for Peptidoform ID: nQLRSRHSPPPR_n145_1_Q111_1_S167_1 in Bin (-0.0225, -0.0175)
AUC: 32.84%
-0.0233
0
50
100
150
200
250
300
350
−0.026
−0.024
−0.022
−0.020
−0.018

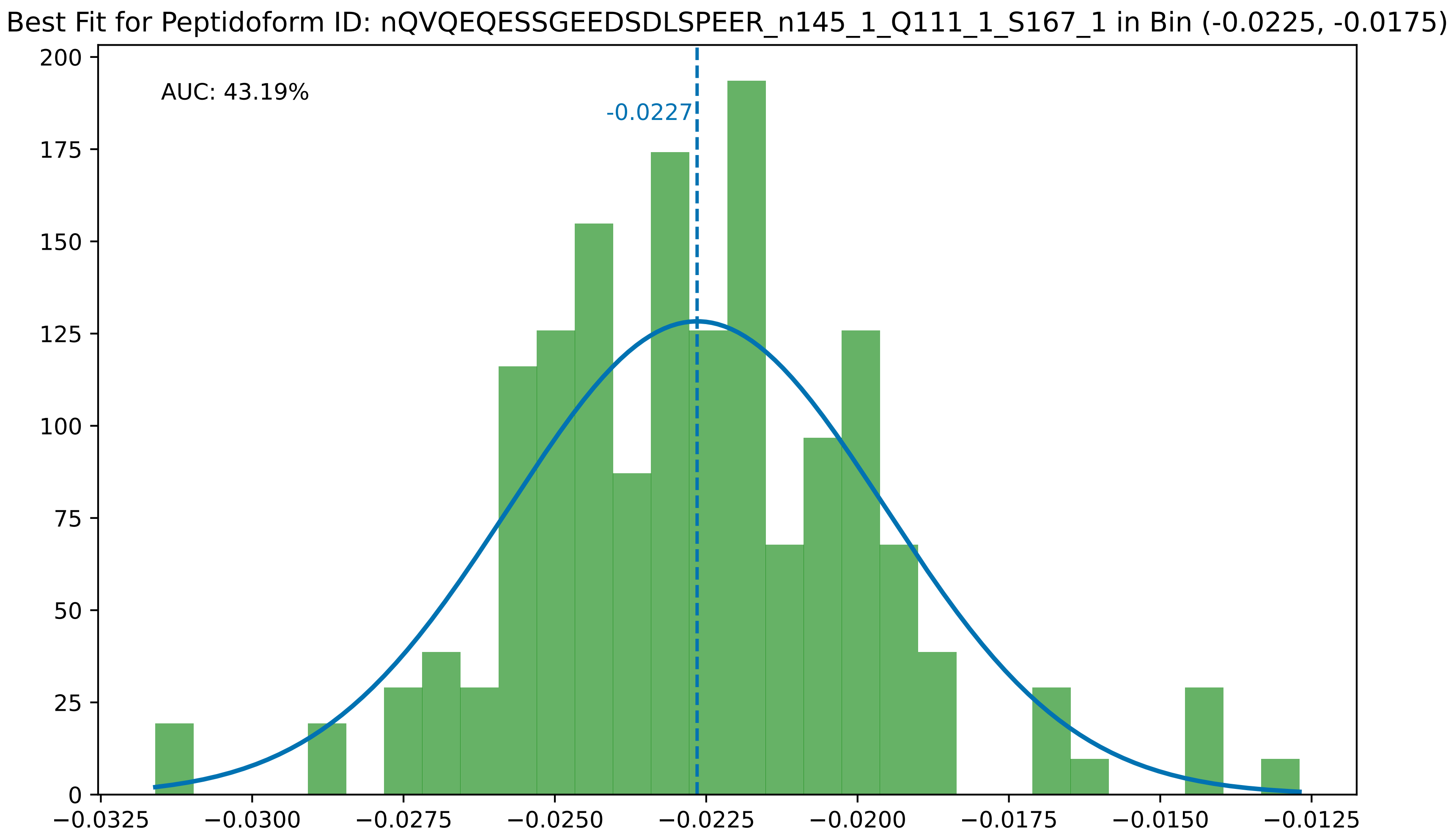

Best Fit for Peptidoform ID: nQVQEQESSGEEDSDLSPEER_n145_1_Q111_1_S167_1 in Bin (-0.0225, -0.0175)
AUC: 43.19%
-0.0227
0
25
50
75
100
125
150
175
200
−0.0325
−0.0300
−0.0275
−0.0250
−0.0225
−0.0200
−0.0175
−0.0150
−0.0125

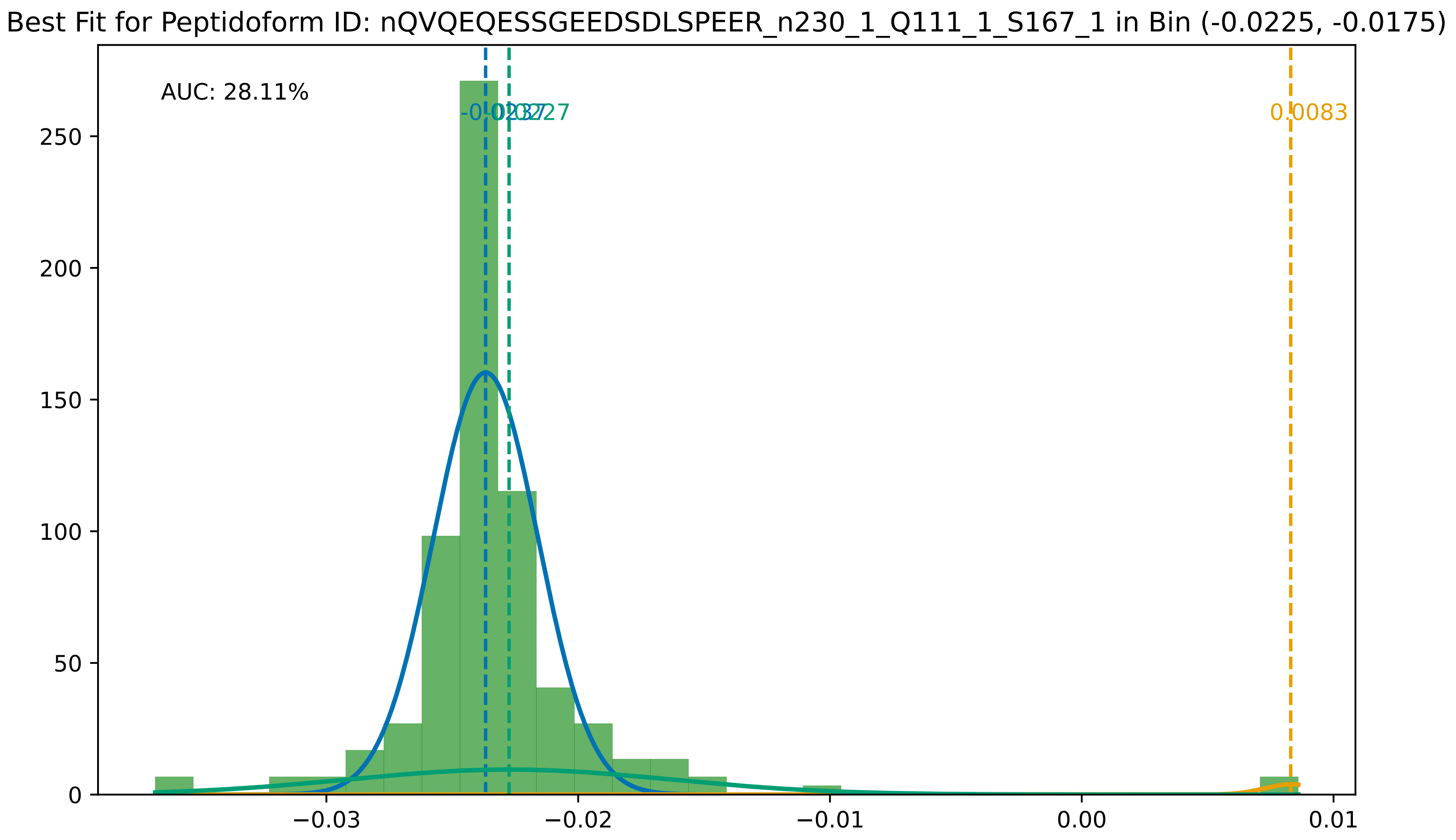

Best Fit for Peptidoform ID: nQVQEQESSGEEDSDLSPEER_n230_1_Q111_1_S167_1 in Bin (-0.0225, -0.0175)
AUC: 28.11%
-0.0237
-0.0227
0.0083
0
50
100
150
200
250
−0.03
−0.02
−0.01
0.00
0.01

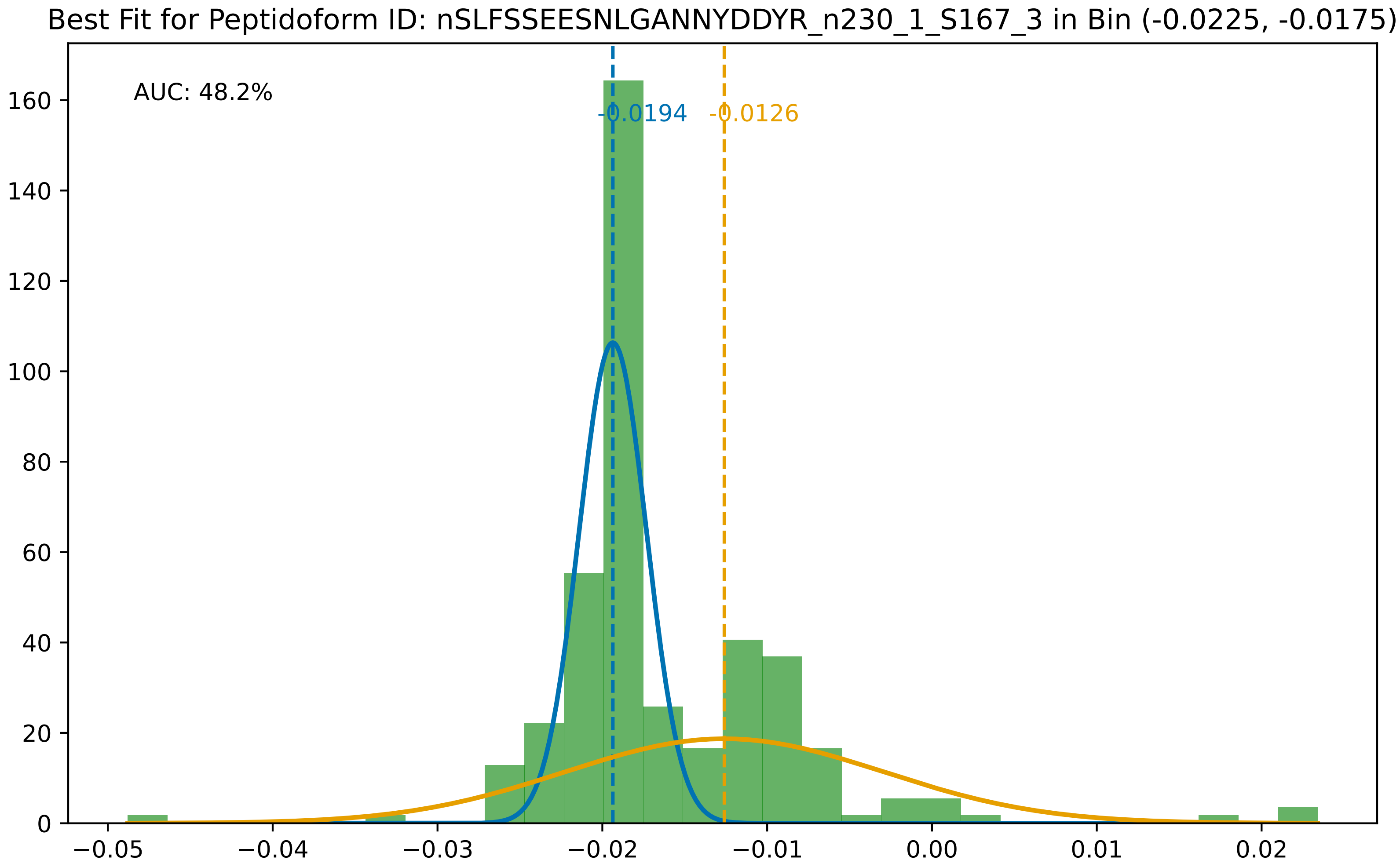

Best Fit for Peptidoform ID: nSLFSSEESNLGANNYDDYR_n230_1_S167_3 in Bin (-0.0225, -0.0175)
AUC: 48.2%
-0.0194
-0.0126
160
140
120
100
80
60
40
20
0
−0.05
−0.04
−0.03
−0.02
−0.01
0.00
0.01
0.02

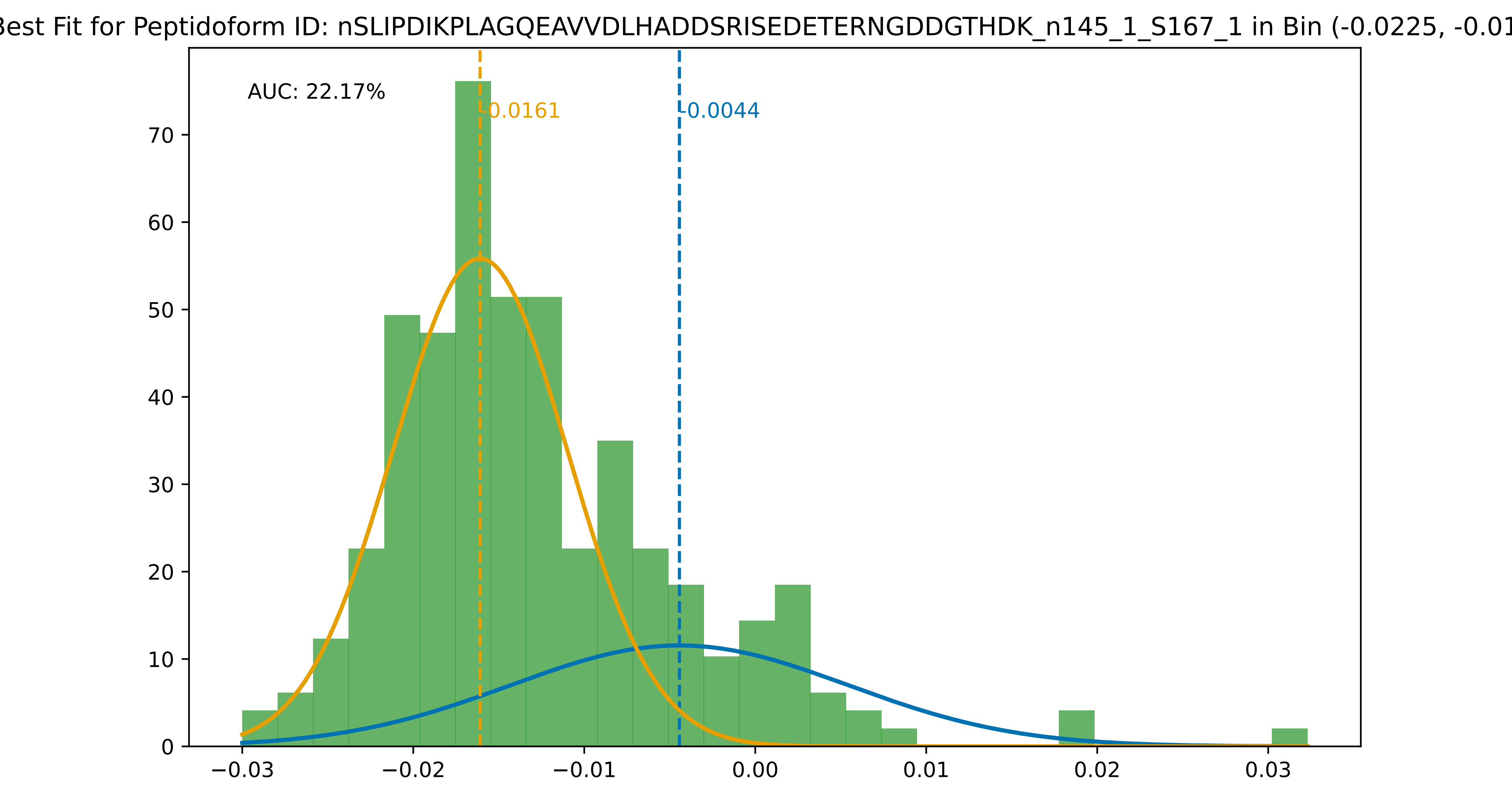

Best Fit for Peptidoform ID: nSLIPDIKPLAGQEAVVDLHADDSRISEDETERNGDDGTHDK_n145_1_S167_1 in Bin (-0.0225, -0.01
AUC: 22.17%
-0.0161
-0.0044
0
10
20
30
40
50
60
70
−0.03
−0.02
−0.01
0.00
0.01
0.02
0.03
