## Supplementary File 4 for "Searching for Sulfotyrosines (sY) in a HA(pY)STACK": README.txt

Each plot is a histogram of the calibrated mass errors for all pooled PSMs 
identified as the specified non-strict peptidoform. The colour-coded lines
represent the normal distribution density function for each component of 
the fitted GMM model (up to 3 components, component 1 in blue, component 2 
in orange, component 3 in green), and the dashed lines indicate the median
mass error for that component, with the value printed in the same colour.
The area under the curve value specifies the proportion of the density 
function that falls over the specified bin of interest. 
